# Rising minimum temperatures contribute to 50 years of shifting Arctic and boreal butterfly communities in North America

**DOI:** 10.1101/2023.04.24.538168

**Authors:** Vaughn Shirey, Naresh Neupane, Robert Guralnick, Leslie Ries

## Abstract

Global climate change has been identified as a major driver of observed insect declines, yet in many regions there are critical knowledge gaps for how communities are responding to climate. Poleward regions are of particular interest because warming is most rapid while biodiversity data are most sparse. Building on recent advances in occupancy modeling of presence-only data, we reconstructed 50 years (1970-2019) of butterfly population trends in response to rising minimum temperatures in one of the most under sampled regions of the continent. Among 90 modeled species, we found that cold-adapted species are far more often in decline compared to their warm-adapted, more southerly distributed counterparts. Further, in a post-hoc analysis using species’ traits, we find that species’ range-wide average annual temperature and wingspan are a consistent predictor of occupancy changes. Species with warmer ranges and larger wingspans were most likely to be increasing in occupancy. Our results provide the first look at macroscale butterfly biodiversity shifts in a critically under sampled region of North America. Further, these results highlight the potential of leveraging the wealth of presence only data, the most abundant source of historical insect biodiversity. New approaches to the modeling of presence only data will match recent increases in community science participation with sparse historical records to reconstruct trends even in poorly sampled regions.

## 1. INTRODUCTION

Multiple lines of scientific evidence have pointed to the potential for an alarming recent decline in insect biodiversity across the planet (Cardoso *et al*. 2020; Wagner *et al*. 2021a, b). Major changes in the abundance and composition of insect populations increase the risk of losing vital insect-mediated ecosystem services, including acting as prey items, and in pollination, decomposition, and pest control (Kremen *et al*. 2007, Zhou *et al*. 2023), and even the potential for collapse of some ecological networks (Dunne *et al*. 2002; Memmott *et al*. 2004; Grames *et al*. 2023). Although the list of potential agents driving insect decline is multifaceted, climate change has been put forth as a major direct contributor but also as a force that interacts with other global drivers (Wagner *et al*. 2021b). As ectotherms, insects are especially sensitive to local scale and macroscale climatic conditions in ways that may integrate to signals of change at macro-scales (Deutsch *et al*. 2008). Among insects, butterflies have by far the densest sources of distribution and natural history data for insects due to their high public profile and apparency in nature. Numerous foundational studies in global change biology have used butterflies to better understand how life history, ecological traits, and species interactions may drive responses to novel climatic conditions (Parmesan *et al*. 1999; Pöyry *et al*. 2009; Breed *et al*. 2013) .

Surface-level temperatures have risen more drastically in high-latitude regions than anywhere else (Holland & Bitz 2003), a trend that is expected to continue under nearly all climate change forecasts (Ono *et al*. 2022). Under current, “business-as-usual” carbon-dioxide emission scenarios, surface temperatures are projected to increase in the Arctic by an average of 10° Celsius through 2100, roughly four times greater than other global regions (You *et al*. 2021). Further, this will likely result in substantive shifts with profound impacts for insect development, such as rainier rather than snowy winters across polar region (Bintanja & Andry 2017;

McCrystall *et al*. 2021). For butterflies in high-latitude regions to persist under such drastically shifting climatic conditions, they must be able to either move to where conditions are favorable, adapt to novel climates *in situ* or face significant risk of population declines (Corlett & Westcott 2013; Kellermann & van Heerwaarden 2019). The rapid climatic changes in this region and the potential for significant barriers to movement (e.g., closed-canopy habitats) may contribute to a high degree of “climate debt” for butterflies (Lewthwaite *et al*. 2018), in other words, a lag between the pace of climate change and observed species responses.

Despite the magnitude of change happening in high-latitude regions, North American boreal and Arctic regions remain critically under sampled for all taxa, and are even more biased that in many other regions due to high concentration of human populations in the south (Shirey *et al*. 2021). These sampling gaps are major impediments to conservation, especially for invertebrates (Cardoso *et al*. 2011) . Historically, data that do exist in this region come largely from natural history museum collections while more contemporary data largely arise from community science platforms, such as records from online platforms like iNaturalist and are almost all opportunistic or “presence-only” data (hereafter, presence-only). Presence-only data rarely capture all species in the broader community (Kelling *et al*. 2019). For example, collectors/observers may more often record species of particular interest or those that are more detectable. Cryptic, small-bodied species may go under-reported (Meyer *et al*. 2016; Adamo *et al*. 2021). Further, there is no expectation of resampling at the same site over time. Finally, accounting for data biases is particularly challenging because absences (or non-detections) go unrecorded, and effort is unknown. Thus, these data have long been unsuitable for standard statistical models, which are largely rooted in generalized linear modeling paradigms that require abundance or presence-absence data. To confront the conundrum that the most highly available distribution data could not be used in typical statistical models, a new class of species distribution models, including popular software like MaxENT, which generate “pseudoabsence” data based on algorithms (Elith* *et al*. 2006), have been widely used and have helped expand our knowledge about large-scale distributions (Elith & Leathwick 2009); however, some studies have found these methods to be unreliable (Yackulic *et al*. 2013).

The issue of using presence-only data in large scale ecological analysis has resulted in many conversations about the most appropriate models and also best practices for the use of these data. These debates span multiple metrics of biodiversity including when or how presence-only data is sufficient to estimate abundances (Ries *et al*. 2019; Wepprich 2019), phenological patterns (Larsen & Shirey 2021), and range dynamics (Yackulic *et al*. 2013; Ascher *et al*. 2020; Guzman *et al*. 2021). For example, Ries *et al*. 2019 and Wepprich 2019 found that approaches used to estimate abundance declines in the Monarch butterfly, *Danaus plexippus* (Lepidoptera: Nymphalidae), were severely biased in an effort to track abundances over 100 years (Ries *et al*. 2019; Wepprich 2019). In addition, Larsen & Shirey (2021) found that lack of curation and inappropriate modeling of presence-only data to in a study to infer phenometrics resulted in inferences that clearly did not make ecological sense; specifically the study found that butterfly populations in high latitudes having similar emergence timing as populations at lower latitudes (Larsen & Shirey 2021). Finally, the strategy of imputing non-detections, not censoring inference to regions where species could plausibly occur, and not accounting for heterogeneous detection probability within an occupancy modeling framework can produce biased estimates of occupancy trends with a particular tendency to find declines (Ascher *et al*. 2020; Guzman *et al*. 2021). Underscoring all of these examples is, given the current biodiversity crisis, the pressing need for approaches to produce reliable trend estimates from presence-only data, by far the most abundant data source, both in the past and likely into the future.

New advances in statistical modeling to more robustly address these biases have the potential to unlock the wealth of presence-only for inferring population trends, even at continental extents. For example, occupancy-detection models, (MacKenzie *et al*. 2002; Kéry & Royle 2015) have long been used to reconstruct ecological signals over space and time from presence-absence, more accurately (and hereafter) detection/non-detection, where sites are repeatedly visited and the patterns in a series of detections and non-detections across all visits are used to model occupancy patterns by disentangling the observation process from underlying ecological patterns. Yet occupancy models demand non-detection data (specifically, zeros for all species in the community when not observed) and so their use with presence-only data has traditionally been considered inappropriate. Recently, proposals for the use of occupancy models with presence-only data center on explicit additions of zeros in the data set by leveraging records of other species has been used as a proxy for effort (Guzman *et al*. 2021; Jackson *et al*. 2022). A recent simulation study has confirmed that, when multiple species records in the same location are used as a proxy for community sampling, the inference of zeros in an occupancy modeling framework provides robust ecological signal from presence-only data (Shirey *et al*. 2022). This approach is increasing in popularity and has already been used to reconstruct sensible ecological trends from presence-only data using these and other non-standardized datasets (van Strien *et al*. 2013; Jönsson *et al*. 2021; Engelhardt *et al*. 2022; Jackson *et al*. 2022).

Here, we used an occupancy-detection approach to account for a complex and undocumented detection process to reconstruct patterns of change in North America above 45°N latitude over a 50-year period (1970-2019). Specifically, we focused on the influence of changes in minimum temperature on species-specific occupancy patterns through space and time. A rich body of research from Europe (Parmesan *et al*. 1999; Hill *et al*. 2002; Pöyry *et al*. 2009) based on structured surveys, where effort is known and all individuals observed are recorded, is available and serves as a guidepost for our *a priori* hypotheses of general expectations in these polar and sub-polar climates. Following observed trends, we hypothesized that warm-associated species are faring, on average, better than their cold-associated counterparts as warmer winters support proliferation of species at their northern range-edges but are less likely to show declines in the south because the intensity of warming is much lower (You *et al*. 2021). In contrast, cold-associated species may be pushed to the northernmost limits of their range and exhibit declines in occupancy probability, either through maladaptation to warmer climates or being pushed out by an influx of better adapted species (Parmesan *et al*. 1999; Hill *et al*. 2002; Pöyry *et al*. 2009; Heikkinen *et al*. 2010; Hällfors *et al*. 2021, 2023). In addition to the importance of understanding trends in poleward latitudes, as the first empirical implementation of this approach, there is the benefit that we are estimating spatial patterns which have been not only been demonstrated in other regions, but also make ecological sense. This is useful because in regions with presence-only data, there is rarely an opportunity to validate results with independent data sets. In this work, we specifically aimed to use sparse, presence-only data on butterflies in North America above 45°N to:

a. Reconstruct historical occupancy trends for butterflies over 50 years including species-specific responses to climate change.
b. Split these trends into three zones: southern, mid-latitude, and northern components of each species range above 45°N latitude to assess subrange trends.
c. Relate overall trends in species occupancy to species-specific ecological traits in order to make identify species most likely to experience declines/increases in occupancy using a model selection paradigm (Candidate models are shown in Table 1).

**Table 1.** Descriptions of the trait-based models of occupancy trend used in our post-hoc analysis. Model codes are provided as well as mathematical definitions of the model. Supporting literature for each model is also provided.

| Model ID | Model Specification | Rationale |
| --- | --- | --- |
| A<br>“Null” | $Trend \sim \alpha$ | Ecological “null” model, *i.e.*, all occupancy trends are predicted by an average intercept. |
| B<br>“Null + Phylo” | $Trend \sim \alpha + \alpha_{phylo}$ | Model A with a phylogenetic random intercept term, *i.e.*, species occupancy trends are predicted by an average intercept and species-specific, phylogenetic intercept. |
| C<br>“Temp” | $Trend \sim \alpha + \beta_1 \times RangeTemp$ | Occupancy trends are best predicted by species’ range-wide average annual temperature. Warmer, southern species will exhibit increases in occupancy probability. |
| D<br>“Temp + Phylo” | $Trend \sim \alpha + \alpha_{phylo} + \beta_1 \times RangeTemp$ | Occupancy trends are best predicted by species’ range-wide average annual temperature and a species-specific, phylogenetic intercept. Warmer, southern species will exhibit increases in occupancy probability. |
| E<br>“Range” | $Trend \sim \alpha + \beta_1 \times RangeSize$ | Occupancy trends are best predicted by species’ range-size. Widely distributed species will exhibit increases in occupancy probability due to wider environmental tolerance. |
| F<br>“Range + Phylo” | $Trend \sim \alpha + \alpha_{phylo} + \beta_1 \times RangeSize$ | Occupancy trends are best predicted by species’ range-size and a species-specific phylogenetic intercept. Widely distributed species will exhibit increases in occupancy probability due to wider environmental tolerance. |
| G<br>“Size” | $Trend \sim \alpha + \beta_1 \times Wingspan$ | Occupancy trends are best predicted by species’ average wingspan. Larger species will exhibit increases in occupancy probability due to greater mobility (Sekar 2012). |
| H<br>“Size + Phylo” | $Trend \sim \alpha + \alpha_{phylo} + \beta_1 \times Wingspan$ | Occupancy trends are best predicted by species’ average wingspan and a species-specific phylogenetic intercept. Larger species will exhibit increases in occupancy probability due to greater mobility (Sekar 2012). |
| I<br>“Resource” | $Trend \sim \alpha + \beta_1 \times HostPlantBreadth$ | Occupancy trends are best predicted by species’ family-level host plant breadth. Species with greater host plant breadth will exhibit increases in occupancy probability due to broader resource availability. |
| J<br>“Resource + Phylo” | $Trend \sim \alpha + \alpha_{phylo} + \beta_1 \times HostPlantBreadth$ | Occupancy trends are best predicted by species’ family-level host plant breadth and a species-specific phylogenetic intercept. Species with greater host plant breadth will exhibit increases in occupancy probability due to broader resource availability. |
| K<br>“Overwinter” | $Trend \sim \alpha + \beta_1 \times RangeTemp + \beta_2 \times OverwinterStage + \beta_3 \times (RangeTemp: OverwinterStage)$ | Occupancy trends are best predicted by range-wide temperature, overwintering life stage, and the interaction between those two factors. We included an interaction term here since physiological or behavioral overwintering strategies could vary by a species’ thermal adaptability. For example, some species of Lepidoptera possess the ability to synthesize antifreeze compounds while others may burrow underground or find other sheltering mechanisms to survive winter conditions (Downes 1965; Layne Jr & Kuharsky 2000; Brackley 2021). Further, responses to environmental cues can vary ontogenetically (Brackley *et al.* 2021). |
| L<br>“Overwinter + Phylo” | $Trend \sim \alpha + \alpha_{phylo} + \beta_1 \times RangeTemp + \beta_2 \times OverwinterStage + \beta_3 \times (RangeTemp: OverwinterStage)$ | Occupancy trends are best predicted by range-wide temperature, overwintering life stage, the interaction between those two factors and a species-specific, phylogenetic intercept. Following the same logic as Model K. |
| M<br>“Complex” | $Trend \sim \alpha + \beta_1 \times Trait_1 \dots \beta_n \times Trait_n$ | Occupancy trends are best predicted by a model that includes all available trait information. Many traits predict occupancy trend. |
| N<br>“Complex +<br>Phylo” | $Trend \sim \alpha + \alpha_{phylo} + \beta_1 \times Trait_1 \dots \beta_n \times Trait_n$ | Occupancy trends are best predicted by a model that includes all available trait information and a species-specific, phylogenetic intercept. Many traits predict occupancy trend. |

## 2. METHODOLOGY

Occupancy-detection models are designed to disentangle the process of observation from the underlying ecology (MacKenzie *et al*. 2002; van Strien *et al*. 2013; Kéry & Royle 2015). Further, multi-species occupancy models can be used to leverage community patterns to strengthen the inferences for multiple species at the same time (Dorazio & Royle 2005; Zipkin *et al*. 2010). In a recent simulation study, these models have demonstrated the ability to accurately reconstruct ecological trends from presence-only datasets that meet certain pre-qualifying conditions indicative of sampling multiple species at one location (Shirey *et al*. 2022). A graphical representation of our workflow with details about this implementation of our model, including methods for inferring zeros is illustrated in Supplemental Figure 1.

To implement occupancy-detection models for this study, we established grids of 100×100 and 200×200-kilometer square cells across our study region to determine if the spatial grain of our study impacted our results. We set our analysis to the 50 years between 1970 and 2019 and divided this period into ten, five-year-long intervals. Previous simulation results found that five temporal bins were found to provide sufficient granularity to detect trends, but ten were best able to reconstruct simulated trends and also provide more nuanced patterns through time. In contrast, using only two time periods, a common method in these studies, often produced poor results (Shirey et al., 2022). In each of these periods we estimated the probability that a given species is an occupant of a specific grid cell. These “occupancy intervals” were further broken down into five one-year-long “visit intervals” to provide a basis for separate estimation of detection and occupancy, which requires tracking multiple visits as repeated trials within the larger occupancy interval (MacKenzie *et al*. 2002).

### 2.2 Climate Data

We used minimum temperatures and precipitation data to inform the ecological process of our occupancy-detection model. Although land use and pesticides are also contributors to insect population shifts (Wagner 2021) we do not use those here since, to our knowledge, no comprehensive datasets of land cover or pesticide use exists for high-latitude regions of North America extending back into the 1970s. Monthly minimum temperatures from 1970 to 2015 were extracted from the National Oceanic and Atmospheric Administration’s (NOAA) 20th- century reanalysis, 20CRv3 (Slivinski *et al*. 2019). These data are available at 0.1-degree spatial and monthly temporal resolution from 1836 to 2015; reanalysis products (where observation data are synthesized with mechanistic climate models are not yet available for 2016 onward, variables were obtained from the Climate Prediction Center (CPC, which are available at 0.5-degree spatial and monthly temporal resolutions) (Climate Prediction Center 2022). Minimum average temperatures increased over the vast majority of our study range, up to 4° Celsius over the 50- year study period (Figure 1a). For precipitation, monthly precipitation data were extracted from the Climate Research Unit (CRU) TS4.04 global precipitation dataset which are also available at 0.5-degree spatial resolution, but only back to 1900 (Harris *et al*. 2020a, b). Trends in precipitation were generally more heterogenous but reflect, on average, increasing rainfall across the region.

**Figure 1.**
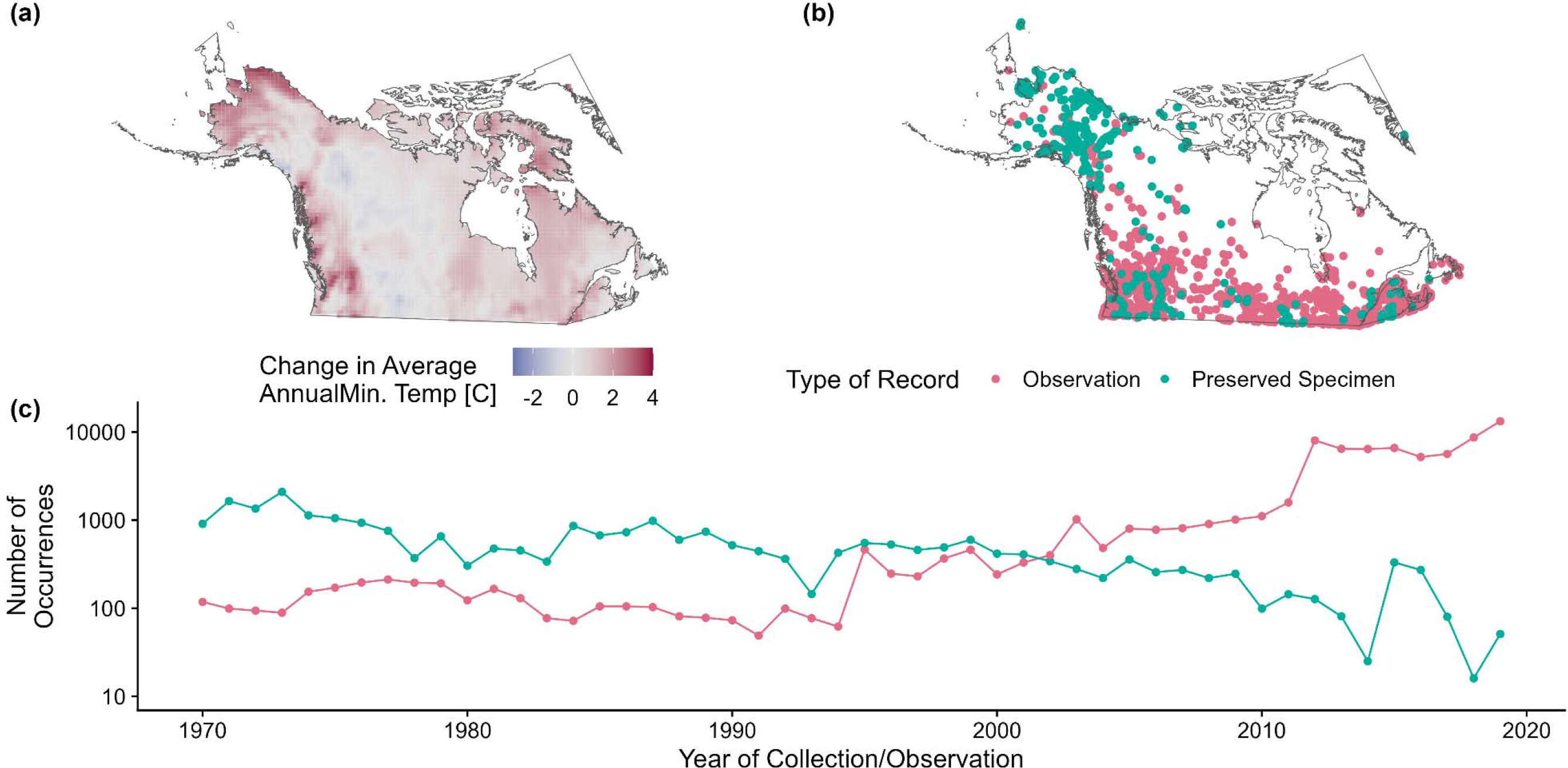
An overview of the study region encompassed by our analysis including (a) the change in average annual minimum temperature from the 1970s to 2010s, (b) the spatial distribution of occurrence records from natural history collections and community science platforms, and (c) the number of occurrence records from natural history museum collections and community science platforms by year over the same 50-year timeframe. A random sample of 5,000 of the occurrence records (c) are shown to indicate spatial bias in available records while avoiding overplotting.

Climate raster data reflecting monthly average minimum temperature and monthly precipitation for the years 1970 through 2019 were read into R using the package “raster” (Hijmans *et al*. 2015). All raster data were reprojected using this same package to the project coordinate reference system (North America Albers Equal Area Conic) and this was also used as the base projection for all distribution data in this analysis. The raster data were summarized within each 100×100-kilometer (or 200×200-kilometer) grid cell using the mean value across for each five-year occupancy interval (e.g., 1970-1974) weighted by their coverage of the cell. Finally, exploration of our climate data revealed that average minimum temperature and average precipitation were highly correlated (Pearson’s r = 0.67 for the 100×100-kilometer scale and Pearson’s r = 0.73 for the 200×200-kilometer scale), thus we opted to run our models separately for both climate covariates (results for temperature are shown in the main text and precipitation in the Supplementary material). We opted to focus on temperature over precipitation as prior research has demonstrated temperature to be more important for range dynamics in butterflies (Keret *et al*. 2020).

### 2.1 Species Range, Trait, Phylogeny, and Occurrence Data

We used species range boundaries to constrain model inference to historical regions applying a conservative 100-kilometer buffer to allow for range shifts. If any portion of the buffer intersected with a grid cell, that cell was assumed to be potentially occupied by that species. Prior work has shown that failure to censor locations where a species could not occur can produce biased and/or misleading estimates of occupancy (Guzman *et al*. 2021; Shirey *et al*. 2022).

We used the same range maps to calculate range size and range-wide climatic metrics which served as a basis for species range-related traits, including total range size, average, range-wide temperature and precipitation across all of North America. Species range map data were derived from published field guides on North American butterfly species, including The Kaufman Field Guide to Butterflies of North America (Brock & Kaufman 2006) and A Swift Guide to Butterflies of Mexico and Central America (Glassberg 2018). The range maps were digitized as part of work by Earl et al. (2021) and were re-used here. Range-wide average annual temperature and precipitation traits were extracted from each species’ range averaging conditions over each polygon using the WorldClim2/BioClim dataset (representing average climatic conditions from 1970-2000) (Fick & Hijmans, 2017). Along with this derived climatic trait data, we aggregated other ecological traits from the global LepTraits database (Shirey et al., 2022) to test how these traits are associated with observed population trajectories. Thus, the final dataset of butterfly traits consisted of average annual range-wide temperatures and precipitation, geographic range size, disturbance tolerance, host plant family breadth, average wingspan, and overwintering stage. Finally, we obtained a recently produced phylogeny (Earl *et al*. 2021) of North American butterfly species to be used in our *post hoc* analysis of occupancy trends. We used this phylogeny to test for phylogenetic signal in our modeled occupancy results.

Species occurrence data for all butterflies above 45°N from 1970 to 2019 were obtained from the Global Biodiversity Information Facility (GBIF)(GBIF.org 2022), Integrated Digitized Biocollections (iDigBio)(see Supplemental File S1 for a list of accessed collections), and the Symbiota Collections of Arthropods Network (SCAN) (Heinrich *et al*. 2015)(see Figure 1b, c for a summary of these data). Duplicate records for the same specimen were filtered out. We removed records with coordinate precision greater than 25 kilometers of a reported uncertainty radius. We reconciled the taxonomic names using the R package “taxize” (Scott Chamberlain & Eduard Szocs 2013) and, for non-matches, we resolved the taxonomy by hand where tractable (e.g. by matching unresolved synonymies to currently accepted names). We used the Lamas taxonomy as a backbone (Lamas 2015); because of the continued disputes within the *Celestrina* complex (Wright & Pavulaan 1999; Pavulaan & Wright 2000, 2005), we excluded this group from our analysis since we could not make redetermination for all occurrence records. We then limited our analysis to include only species with at least 500 reported occurrences across the spatiotemporal scope of our study. Finally, we removed all species from the analysis that are known migrants (*i.e.,* are not historically known to overwinter anywhere north of 45° in North America). In the iterative process of modeling, we noted several species for which the MCMC chains experienced notable convergence issues and removed these species from the analysis as well. This resulted in us modeling 90 species out of the total 291 species in the region or roughly 31% of the species pool.

### 2.3 Inferring Detection from Presence-only Observations

Occupancy-detection models use detection/non-detection data to reconstruct occupancy trends (MacKenzie *et al*. 2002; Kéry & Royle 2015). Our simulation study showed that if at least 50% of all spatiotemporal sampling bins contain community data, then the data set is sufficient to go forward with model implementation. If not, our simulation results suggest data are too sparse or not collected/observed under an appropriate methodology for this approach (Shirey et al., 2022) To proxy this probability that field observations in our dataset were community-focused in our dataset, we spatiotemporally aggregated all of our occurrence data at the point level. The occurrence records must have been collected during the same year and must also have been collected/observed within 5-kilometers of one another to be included in an aggregate cluster. We then used the number of species in these spatiotemporal aggregations as a means to classify clusters. “Community clusters” were point aggregate clusters where more than one distinct species was reported; “singleton clusters” were point clusters or single points where only one species was reported. We took the percentage of points that fell within community clusters compared to the filtered occurrence dataset as the proxy of community-focused visitation probability. This percentage was approximately 70% and which was above the 50% found to be a sufficient cutoff in simulation studies (Shirey et al., 2021).

Our approach for imputing non-detection is detailed in Supplemental Figure S1 from Shirey et al. (2022b) and we also provide a brief description here. For the purposes of this study, we assume that the detection of at least two species in a single, one-year visit interval within a 5- year temporal bin within each pixel (e.g., two different species recorded in the 1970 interval within the 1970-1974 occupancy bin) is sufficient to impute non-detection data for all other species that could potentially occur at that grid cell (*i.e.,* their range plus a 100-kilometer buffer that intersects the cell). Although a seemingly low bar for imputing zeros into a 5-year occupancy interval cell, simulations have shown that such an approach generates results that match simulated trends and so are a reasonable minimum requirement, useful when considering regions with particularly sparse data (Shirey *et al*. 2022).

### 2.4 Occupancy-detection Modeling

For the occupancy subcomponent of our model, we included one environmental predictor of occupancy, average minimum temperature (or, in a separate analysis provided in the supplement, average precipitation) across a 5-year-long period (e.g., 1970-1974), as a species-specific slope, and the terrestrial surface area of the grid cell (to account for cells along coastlines or with other large bodies of water). We modeled the butterflies as a single community, and we included species-specific intercepts in our model to account for difference in baseline occupancy among species. Thus, the occupancy component of our model is:

where is the probability that a given species, is an occupant of cell in a 5-year occupancy interval t; is the mean occupancy probability for butterflies (on a linear scale); is a species-specific intercept; is the effect of terrestrial surface area on occupancy probability; and is the species-specific effect (slope) of minimum temperature or precipitation on occupancy. The parameter is only estimated in the temperature model to estimate the potential quadratic effect of minimum temperature on occupancy probability (results for the precipitation analysis are presented in the Supplemental Material). We used uninformative priors for all parameters, thus assuming values were drawn from normal distributions with a mean of zero and a variance that was estimated at the parameter level from a uniform distribution.

The detection component of our model was informed by a random-effects structure which included an average intercept value estimated across all grid cell, occupancy interval, and species combinations, a fixed-effect of occupancy interval on detection (to account for potentially increasing detectability due to modern survey methods and digital platforms such as iNaturalist), and two random intercepts that varies by (a) species and (b) cell by occupancy interval.

Mathematically, our detection component is defined as:

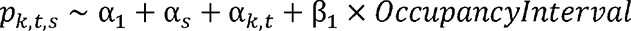

where *p_k,t,s_*, is the detection probability of a given species, c, during a given occupancy interval, at a particular cell, i; the parameter α_1_is the mean detection probability for all butterflies (on a linear scale); α is a random species-specific intercept; α_*k,t*_ is a cell by occupancy interval random intercept; and β_1_ is the effect of occupancy interval on detection probability. For all parameters, we assumed the values were to be drawn from normal distributions with a mean of zero and a variance that was estimated at the parameter level from a uniform distribution. Finally, the likelihood in our model was defined by the product of the latent occupancy state (either present or absent per a Bernoulli draw of the calculated occupancy probability ψ_k,t,s,_) and the probability of detection, *p*_*k,t,s*_.

We ran our occupancy models using JAGS (Plummer 2003) on each detection/non-detection dataset for 150,000 iterations, 50,000 of which were discarded as “burn-in,” retaining the samples of every 100 iterations across four chains for a total of 4,000 samples from the posterior distribution. We assessed convergence across these chains by examining both Gelman- Rubin diagnostic values using 1.1 as an upper threshold (Gelman & Rubin 1992) and by visually inspecting the trace plots for all parameters (provided in the Supplementary Material). We used the R packages “jagsUI” (Kellner 2021) and “MCMCvis” (Youngflesh 2018) to complete the majority of this work. Visualizations of model performance metrics are included in the Supplemental Material.

### 2.5 Post-hoc Trait Analysis

Species’ traits have been demonstrated to be associated with changes in species’ range over time and in the relative risk of decline (Keinath *et al*. 2017) or extinction (Fagan *et al*. 2001; Chichorro *et al*. 2019, 2022). We used our chosen traits (range size, range-wide mean temperature, wingspan, host plant family breadth, overwintering life-stage, and disturbance tolerance) to examine if any were associated with expected mean occupancy shifts from the 1970s to 2010s. Using a model-selection approach, we specified 14 models based on *a priori* hypotheses about which traits might be most important when predicting occupancy trends acros our study. Our first model, Model A, or our ecological “null” model, is an intercept-only model that assumes occupancy trend is best predicted by a singular mean value across all species. In other words, this model assumes that no trait is better able to account for differences in species-specific occupancy trends. To this model, we added species-specific intercept terms that are correlated by a variance-covariance matrix from our phylogeny (Model B, the “null + phylogenetic intercept” model). The additional models (Models C-N) follow a similar structure but in these cases, we selected trait(s) for predicting occupancy trend and implemented each combination with and without phylogenetic intercept terms. Table 1 describes each of these models and provides relevant background for the selection of trait information.

After running our models, we then used a predictive post-check using leave-one-out (LOO) cross-validation, where each pixel-year combination was dropped, and each dropped value was estimated from the model and compared to observations, in order to select the top candidate model (Vehtari *et al*. 2017). From our LOO scores, we calculated the expected log pointwise predictive density (ELPD-LOO) or a measure of predictive capacity (Vehtari *et al*. 2017). Higher values of this metric indicate better predictive capacity of the model. Phylogenetic signal among occupancy estimates was measured using Pagel’s where values close to zeroindicate no phylogenetic signal in the response and values close to one indicate strong correlation with phylogeny (Pagel 1999; Freckleton *et al*. 2002). This work was performed using the R package “brms” which interfaces with Stan (Bürkner 2017). We ran all of our post-hoc models for 200,000 iterations, discarding 100,000 as “burn-in” and thinning by 50 across four chains for a total of 4,000 samples from the posterior. We again assessed convergence by using the Gelman-Rubin diagnostic values and by inspecting trace plots. We provide all model code and complete data files used in our entire analyses via GitHub (https://github.com/vmshirey/HighLatitudeNorthAmericanButterflyOccupancy) and DataDryad

## 3. RESULTS

We modeled the response of 90 species of butterfly to changes to temperature from 1970- 2019. On average, occupancy probability increased with increasing minimum temperatures for the majority of warm-adapted species (Figure 2, 3). As expected, warm-adapted species showed the greatest positive response to increases in minimum temperature and cold-associated species showed acute declines in their response (Figure 2); however, modeled results also showed an unexpected reversal in response for cold-associated species at the highest range of observed temperature increases, although confidence was much lower for that response (Figure 2). When examined on a per-species basis (Figure 3), several cold-adapted species exhibit average occupancy declines across their modeled ranges, including *Boloria freija* (BOLFRE, -7.5% in mid-latitude sites), *B. chariclea* (BOLCHA, -6.8% in mid-latitude sites), *Boloria eunomia* (BOLEUN, -5.9% in mid-latitude sites), and *Agriades glandon* (AGRGLA, -3.1% in mid-latitude sites). In contrast, warm-adapted/southern species exhibited relative occupancy stability or average increases alongside rising minimum temperatures (Figure 3). For example, the species, *Pieris rapae* (PIERAP), *Pterourus rutulus* (PTERUT) and *Cercyonis pegala* (CERPEG) have all exhibited average occupancy increases of roughly 4% or greater across all geographic components of their modeled range. Several species for which occupancy trends appear relatively stable compared to 1970 occupancy levels include *Coenonympha tullia* (COETUL) and *Glaucopsyche lygdamus* (GLALYG).

**Figure 2.**
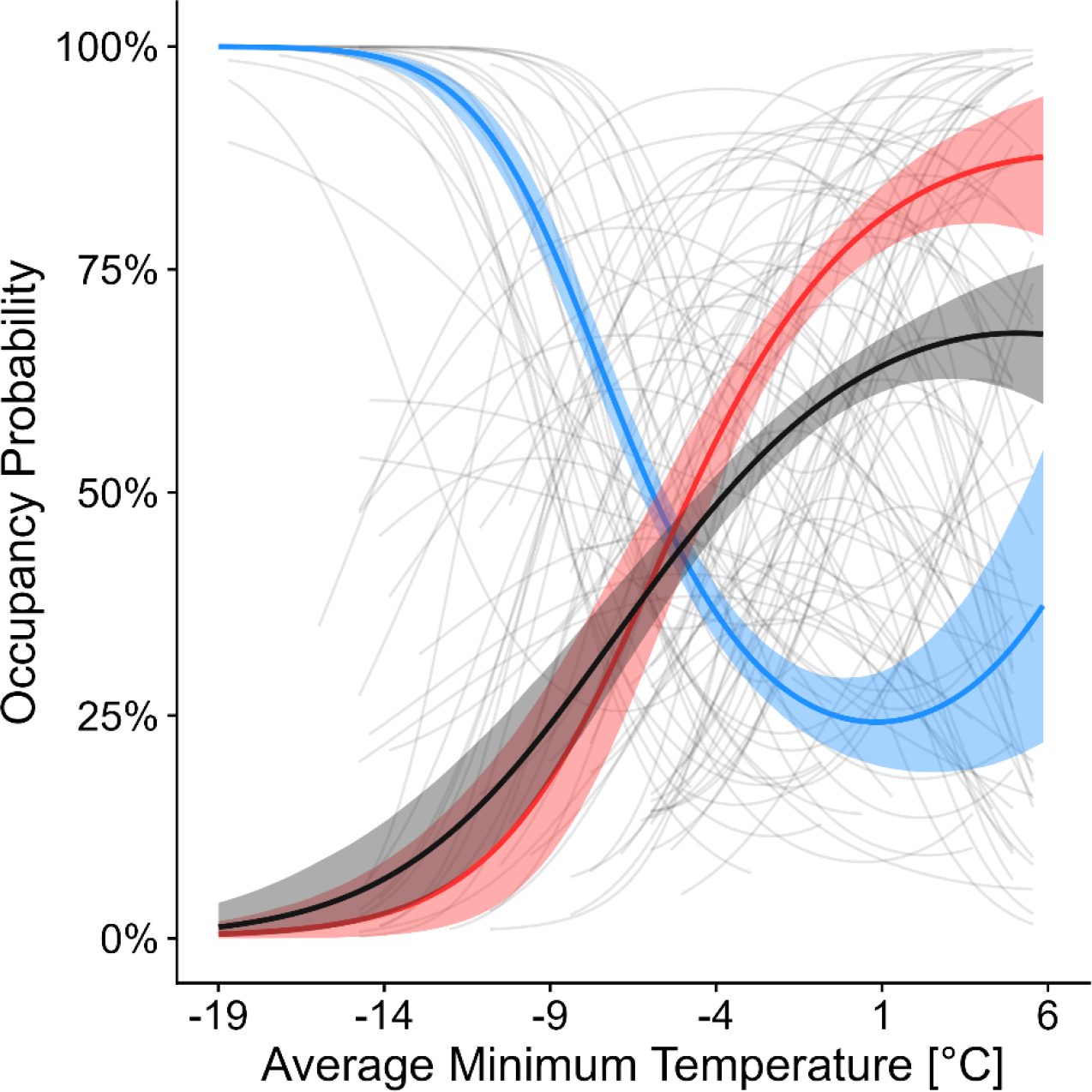
Species-specific responses to average minimum temperature (faded, grey lines) as well as the response of the butterflies with the coldest quartile of North American ranges when all species’ entire ranges are compared (blue, *n = 23*), warmest quartile of North American ranges (red, *n = 23*), and middle half of species or those species with average range-wide temperatures (grey, *n = 44*). Species-specific responses are shown in grey and truncated to show modeled results only for the range of temperatures that each species experienced within its range above 45°N between 1970-2019. All lines are derived from model estimated parameters (linear and quadratic) of species to the average minimum temperature covariate and species-specific responses are shown in Figure 3.

**Figure 3.**
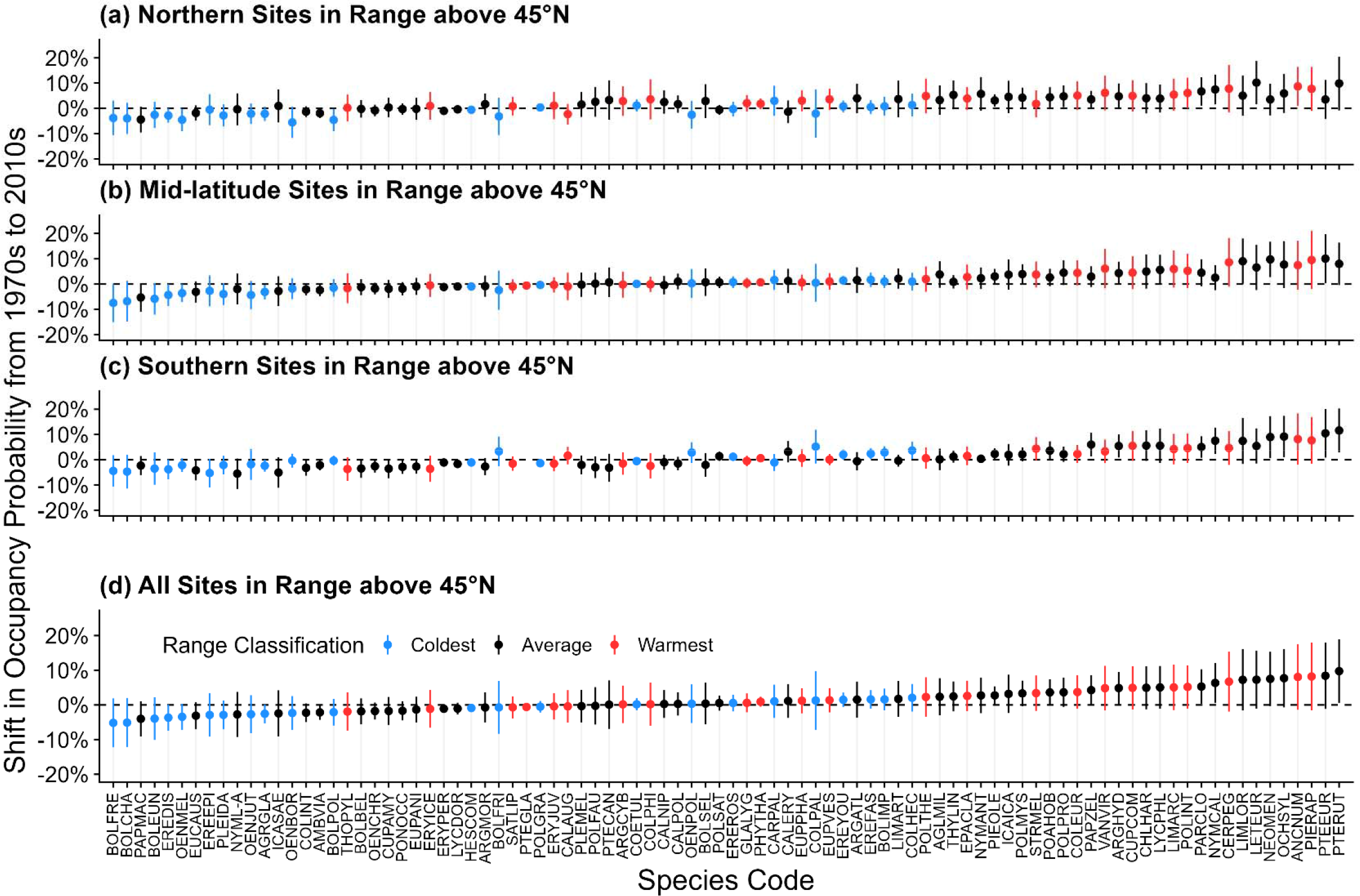
Species-specific occupancy shifts from the 1970s to 2010s where points indicate the mean occupancy shift and lines indicate one standard deviation of the variation in occupancy shift among (a) the northernmost third of cells within that species’ range in the study region, (b) the mid-latitude third of sites within that species’ range, (c) the southernmost third of sites within that species’ range, and (d) all of the sites within that species’ range (all ranges truncated at 45°N). Species are colored by quartiles of the average annual temperature in their North American ranges, so it is directly comparable to Figure 2.

In sum, we find trends at the northern third of each species’ model range are generally more positive than trends at the southern third (Figure 3) but those for cold-associated species are typically negative across all geographic subcomponents of their ranges (Figure 3a, b, c) while those for warm-associate species, the trends in each geographic subcomponent are stable or increasing. Due to the very small number of sites modeled for *P. glaucus*, a comparison could not be made between northern and southern sites. A full account of average occupancy shifts as well as maps reproducing the shifts from the 1970s for each species are included in the supplementary material for all models.

The top model from our post-hoc analysis revealed that a model including range-wide temperature (Models C/D) and a model including average wingspan (Model G/H) were the best models in terms of predicting occupancy shift at the 100-kilometer scale (Supplemental Table S2). Generally, species with warmer North American ranges and species that have larger wingspans exhibited larger gains/smaller losses in occupancy probability over the period of this research (Figure 4a, b). Across other models not presented in the main text, the Model C/D group were consistently the top predictive models of occupancy trend for temperature-based occupancy trends (Supplemental Table S2-S3). An analysis of the phylogenetic signal in occupancy trend revealed weak evidence for a strong phylogenetic signal with Pagel’s λ being estimated at 0.0 (0.0 – 0.01 95% Bayesian credible interval) for Model D and Model H across all models. This indicates that phylogenetic signal among occupancy trends is low.

**Figure 4.**
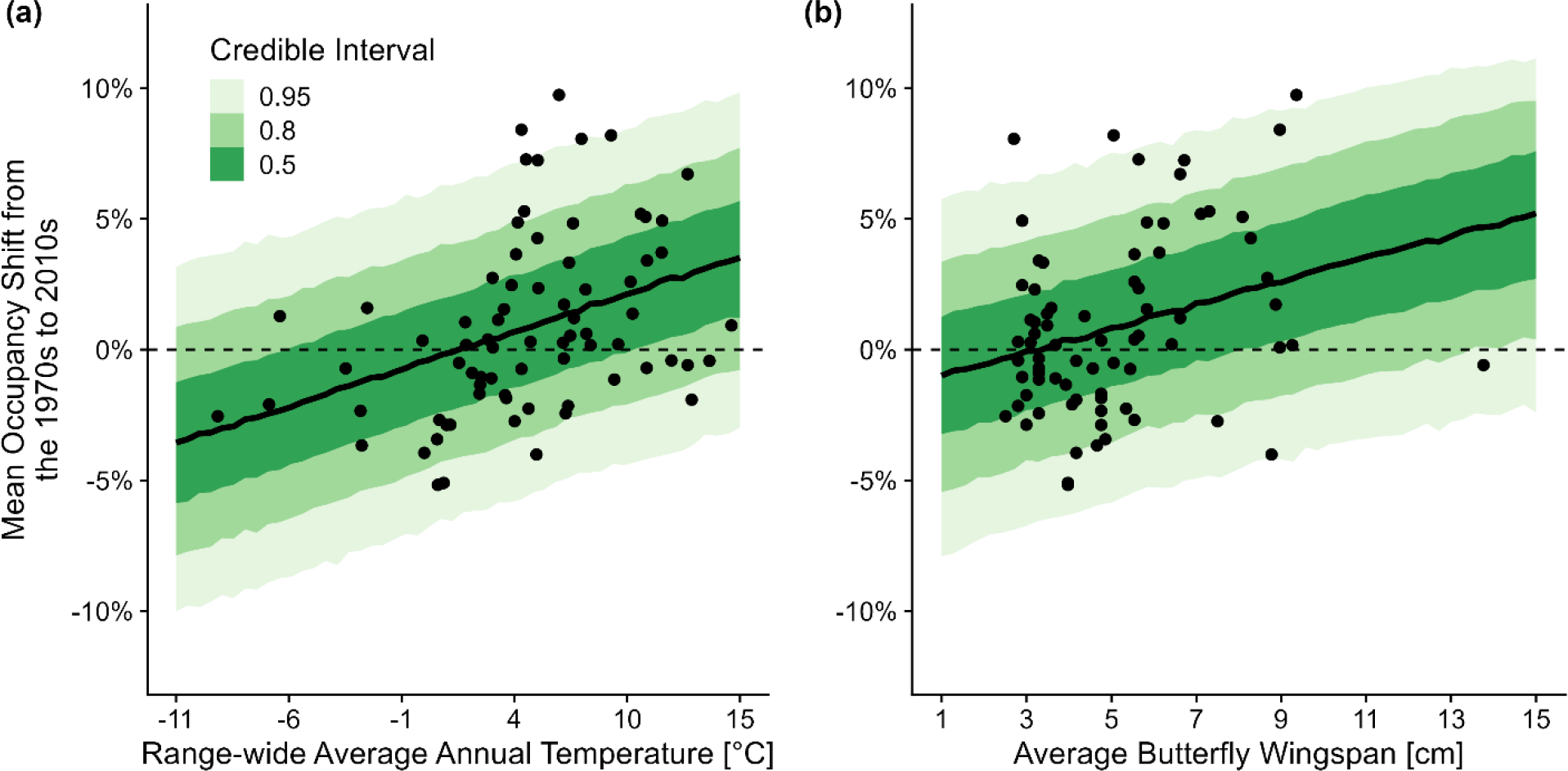
The predictive relationship between species’ (a) range-wide average annual temperature (Model C) and (b) average wingspan (Model G) for the mean occupancy probability shift from the 1970s to 2010s. Shaded regions indicate Bayesian credible intervals while points represent the individual species in our post-hoc analysis.

## 4. DISCUSSION

Our models demonstrate that minimum temperature (and in corollary, precipitation, see supplement) predicts the overall 50-year occupancy trajectory of butterfly species in our study region. On average, minimum temperatures have increased by an average of 0.86 degrees Celsius between the 1970s and 2010s (Figure 1). Increases in winter temperatures may influence butterfly species differently depending on the climates they are best suited to. First, elevated minimum temperatures may be detrimental to cold-associated species in several different ways. Lack of snow cover and/or false phenological cues of winter termination may increase the risk of exposure of diapausing butterflies to harsh winter temperatures and early spring frosts. Increased minimum temperatures may also be indicative of increased winter heatwaves (Beniston 2005; Vikhamar-Schuler *et al*. 2016), and the impact of winter heatwaves has been quantified in single species experimental studies and may be a core driver of population decline at the trailing southern edge of species’ ranges as has been found in the Baltimore Checkerspot, *Euphydryas phaeton* (Lepidoptera: Nymphalidae) (Abarca *et al*. 2019). For warm-adapted species, elevated minimum temperatures are likely beneficial as previously harsh winters become milder for these groups, allowing for northward range expansion as has been found in some species (Crozier 2003, 2004). Indeed, poleward shifts are a commonly noted phenomenon among species elsewhere as they aim to track their thermal optima within a changing climate (Parmesan *et al*. 1999; Pöyry *et al*. 2009; Breed *et al*. 2013). Spatially, our results confirm that stronger positive shifts in species occupancy are occurring for a majority of species (n = 60 or roughly 66%) in the northern geographic context of their modeled range (Figure 3). The estimated effect of minimum temperature on species-specific occupancy from our model supports these aforementioned hypotheses (Figure 2), and as evidenced by cold-associated species generally declining in occupancy probability as minimum temperatures rise (Figure 3). In contrast, many of the warm-associated species in our study benefit from the same increasing minimum temperatures (Figure 2). Further, species with intermediate range-wide temperatures exhibit a positive response to rising minimum temperatures (Figure 2). Finally, while cold-adapted species appear to be faring poorly across all components of their subranges (Figure 3); warm-adapted species tend to exhibit the largest increases in occupancy probability towards the northern periphery of their ranges (Figure 3, Supplemental File S1). While these southern species can still track their thermal niches poleward, cold-adapted species may already be approaching the final course of the “escalator to extinction” as they run out of potentially suitable habitat in which to track their thermal niche (Marris 2007; Freeman *et al*. 2018; Urban 2018).

Thermal niche is indeed a critically important predictor of occupancy dynamics in our study system. Our post-hoc analysis of species traits confirmed that a species’ range-wide temperature was the top predictor of 50-year occupancy shift across our study region (Supplemental Table S2; Model C/D). Indeed, species’ thermal niches are nearly ubiquitous predictors of range shifts across the tree of life (McMahon & Hays 2006; Scridel *et al*. 2017; Braschler *et al*. 2020). Further, given that we informed our estimates of occupancy probability by climatic variables, it is a positive confirmation that range-wide average annual temperature was a strong predictor of trends and that our methodology was reliable for reconstructing ecologically sensible trends from sparse, presence-only data. Taken together, these results suggest a more nuanced perspective of insect biodiversity decline in more northern regions and suggest that a scenario of climate “winners” and “losers” (Jackson *et al*. 2022) is the dominant pattern likely occurring as opposed to ubiquitous declines.

Since prediction is an important part of the scientific enterprise, we aimed to also test which traits, aside from range-wide temperature (if any), might also predict occupancy declines in our study region. Among our models, Model G/H, using average wingspan as a predictor, emerged as a near equivalents to Model C/D (range-wide temperatures) candidate for predicting occupancy trend (Supplemental Table S2). In these models, butterflies with larger wingspans are more likely to be increasing in occupancy probability with smaller butterflies more likely to be in decline (Figure 4b). Wingspan is often used as a proxy for mobility where species with larger wingspans are typically considered to have greater flying ability and, consequently, improved odds of navigating to new, suitable habitats (Sekar 2012). Further, we note that wingspan as a proxy for mobility appears to operate at both intra- and interspecific scales of variation. For example, in the Monarch butterfly (*Danaus plexippus*, not modeled here due to its exceptional migratory status), long-distance migrants exhibit larger wingspans than generations that do not make the long flight to the overwintering grounds in Mexico (Dockx 2007; Freedman & Dingle 2018; Freedman *et al*. 2020). Emerging work on the impacts of climate on morphology have shown that butterflies fluctuate in size (Bowden *et al*. 2015; Daly 2018) and thus future work should consider how intraspecific variation in morphology, driven by global change, may predict, exacerbate, or mitigate occupancy trends at various spatial scales. For example, if butterflies are becoming smaller due to climate change, this may, in turn, impact their mobility, and consequently, the ability for them to track their thermal niche. The degree to which these effects are not captured by models that don’t account for intraspecific variation in these factors across a species range remains to be tested. Our model results point to a scenario where larger butterflies typically fare better with respect to occupancy declines, which, in the absence of connectivity data, suggest by proxy that species’ mobility is an important factor for persistence under rapidly changing climates in the region. Limited ability to colonize newly available habitat in this region is the leading hypothesis surrounding why so much climate debt has accrued for butterflies here (Lewthwaite *et al*. 2017, 2018).

Limitations to species’ mobility may be intrinsic to the species (e.g., mobility) or to the landscape. While we examined the impacts of climate on species-specific occupancy trends, the importance of changing land-cover/use on boreal and Arctic butterflies cannot be ignored. Butterflies are especially sensitive to habitat type and disturbance as well as the availability of corridors to navigate to new, suitable habitats. In boreal Canada, butterfly abundance and species richness increases along human-made cutlines (Riva *et al*. 2018a). Additionally, increasing frequency and intensity of forest fire in the region may also contribute to both opportunities and challenges for high-latitude butterflies (Girardin & Mudelsee 2008; Hanes *et al*. 2019). Forest fire (of varying severity) may contribute to initial negative butterfly abundance patterns, but over time, butterflies may benefit from the presence of early successional and open canopy habitats (Johansson *et al*. 2020; Mason Jr *et al*. 2021; Ulyshen *et al*. 2022). Further, open canopy areas (including roadsides, smaller cutlines, and trails) may act as corridors for butterfly movement (Haddad 1999; Haddad & Tewksbury 2005; Riva *et al*. 2018b). Low connectivity of habitat/mobility of species can make it more challenging for more southern distributed butterflies to navigate and colonize new habitats as they track their thermal tolerances (Hodgson *et al*. 2012).

Recent work in the western United States has revealed parallel sensitivity of butterfly species to global climate change. In particular, warmer autumn months has been identified as a potential driver for fewer butterflies being seen by community scientists in the western United States (below our study region) (Forister *et al*. 2021). In comparison to work from the Forister team, we find that similar species in our study also exhibit declines including, the Large Marble, *Euchloe ausonides* (Lepidoptera: Pieridae) and the Anicia Checkerspot, *Euphydryas anicia* (Lepidoptera: Nymphalidae) are in declines. Our trends are an important addition to this work and provide context for some of the same species in the northernmost reaches of their ranges.

Finally, to contextualize our results back into the world of modeling practice, we note several key findings from researchers on the forefront of statistical modeling with presence-only data. First, several statistical advances have emerged from research teams to process community sourced survey data to estimate various biodiversity metrics (Dennis *et al*. 2021; Belitz *et al*. 2022). These advancements are exciting and with the integration of emerging technologies, including those from machine-learning (Joseph 2020) we may be approaching a technological boom in biodiversity science, especially as it pertains to reconstructing historical distributions. Our ecologically sensible trends found in this paper may mean that the occupancy-detection framework is readily extensible to other data poor regions of the planet. Finally, we note that there are the robust relationships between occupancy and abundance (Gaston *et al*. 2000; Zuckerberg *et al*. 2009; but see Dennis *et al*. 2019) which may support the use of occupancy dynamics as an appropriate proxy for abundance trajectories . Estimating abundance has been the statistical “Holy Grail” for ecologists working to reconstruct historical trends, especially since we can never go back in time to conduct structured surveys of historical populations. We are optimistic about the use of hierarchical and integrated modeling approaches in this space (Davis *et al*. 2023) and, to the extent that occupancy can be considered a proxy for underlying abundance patterns, suggest that well-crafted occupancy-detection approaches are a substantial leap forward in the use of presence-only data to assess insect declines.

## 5. CONCLUSION

Global climate change is impacting high-latitude butterfly communities across North America; however, it is important to recognize that not all species are being impacted equally. Our research, in tandem with recent occupancy analyses of other groups demonstrates the importance of understanding how climatic shifts will impact insects on a species-specific level. Further, we show that occupancy-detection models can be used with sparse, presence-only data to extract clear ecological signals over large spatiotemporal scales. As such, occupancy-detection models will be an important tool for ecologists and conservation biologists, especially for hyper- diverse groups like insects, where structured monitoring data are often unavailable. While an overall perspective shows that rising minimum temperatures may benefit the majority of butterflies; cold-adapted species typically do not benefit. Species’ range-wide temperature and wingspan are the best predictors of overall occupancy trend, pointing to both the ability of the occupancy-detection approach for reconstructing historical trends from presence-only data but also the importance of mobility for butterflies in the region as they aim to track their thermal tolerances. Further research on the interplay of traits including the role of intraspecific variation in thermal tolerance, phenological and morphological adaptation, and genetic variation/effective population size are needed to contextualize these broad scale patterns further and support predictive frameworks for insect biodiversity decline.

## Data Availability Statement

The code utilized in this analysis are freely available via GitHub at https://github.com/vmshirey/HighLatitudeNorthAmericanButterflyOccupancy and via DataDryad at the following DOI: xxx. Data related to model inferences can be found on GitHub and also at DataDryad. Range maps used in this analysis can be made available upon request.

## Authorship Contributions

VS came up with the study idea and design. NN obtained and processed climate data. VS conducted the analysis and original interpretation. RG, NN, and LR provided feedback on the analysis. VS wrote the original manuscript, and all authors edited the manuscript together.

## Acknowledgments

Firstly, we want to thank all of the museum staff, volunteers, and community scientists who aided in the mobilization of biodiversity data used in this study. We would also like to thank Laura Melissa Guzman, Rassim Khelifa, Leithen M’Gonigle, Sarah Johnson, Hanna Jackson, and Elijah Reyes for insightful conversations and collaboration on the use of occupancy models with presence-only data. Martha Weiss, Gina Wimp, Greg Breed and other members of the Ries Lab provided feedback on the original manuscript. Finally, model development, testing, and implementation were performed on the Georgetown High-performance Computing Cluster (aided by Woonki Chung), Georgetown Massive Data Institute (Lisa Singh), and the University of Florida HiperGator High-performance Computing System.

## SUPPLEMENTAL MATERIAL

### Table of Contents

**Supplemental Table S1.** Average occupancy trend estimates across all models/species

**Supplemental Table S2.** Model selection for post-hoc 100-km temperature analysis

**Supplemental Table S3.** Model selection for post-hoc 200-km temperature analysis

**Supplemental Table S4.** Model selection for post-hoc 100-km precipitation analysis

### ADDITIONAL SUPPLEMENTARY MATERIAL FILES

**Supplemental File S1:** Collections accessed through iDigBio.

**Supplemental File S2:** Trace plots and other model assessment metrics for our 100-kilometer temperature occupancy-detection model.

**Supplemental File S3:** Trace plots and other model assessment metrics for our 200-kilometer temperature occupancy-detection model.

**Supplemental File S4:** Trace plots and other model assessment metrics for our 100-kilometer precipitation occupancy-detection model.

**Supplemental File S5:** Trace plots and other model assessment metrics for our 200-kilometer precipitation occupancy-detection model.

**Supplementary File S6.** Mean occupancy shift maps for species modeled using the 100-kilometer temperature model.

**Supplementary File S7.** Mean occupancy shift maps for species modeled using the 100-kilometer precipitation model

**Supplementary File S8.** Mean occupancy shift maps for species modeled using the 200-kilometer temperature model

**Supplementary File S9.** Mean occupancy shift maps for species modeled using the 200-kilometer precipitation model

**Supplemental Table S5.** Model selection for post-hoc 200-km precipitation analysis

Supplemental Figure S1. Graphical illustration of the approach used in this study

Supplemental Figure S2. Occupancy trends from the 200km temperature model

Supplemental Figure S3. Occupancy trends from the 100km precipitation model

Supplemental Figure S4. Occupancy trends from the 200km precipitation model

Supplemental Figure S5. Parameter estimates for the 200km temperature post-hoc Model C.

Supplemental Figure S7. Parameter estimates for the 100km precipitation post-hoc Model C.

Supplemental Figure S8. Parameter estimates for the 200km precipitation post-hoc Model C.

**Supplemental Figure S1.**
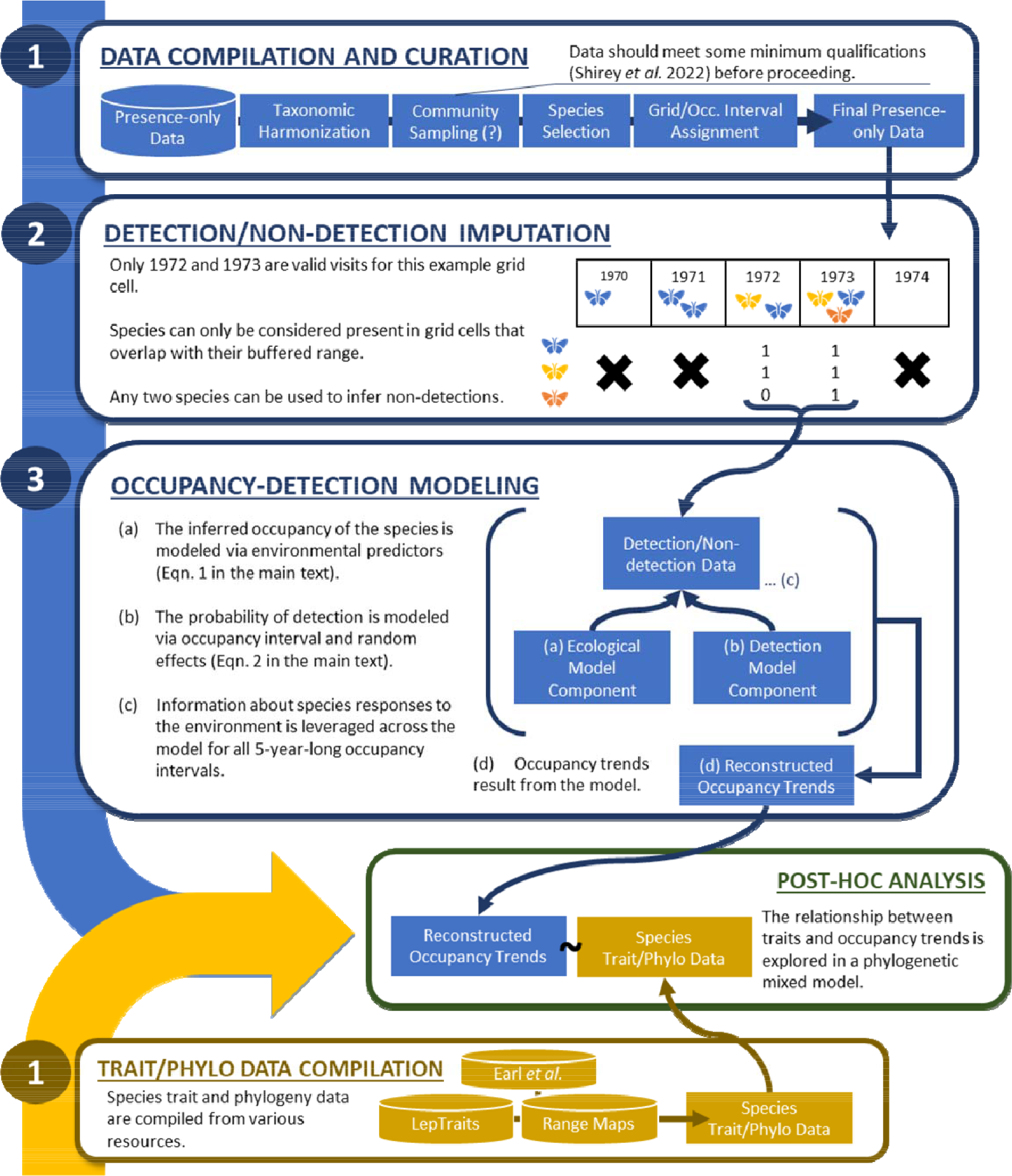
A summary of the methodological workflow used in this study from data compilation to non-detection imputation and occupancy-detection modeling. The post-hoc analysis using species traits and phylogeny is also visualized.

**Supplemental Figure S2.**
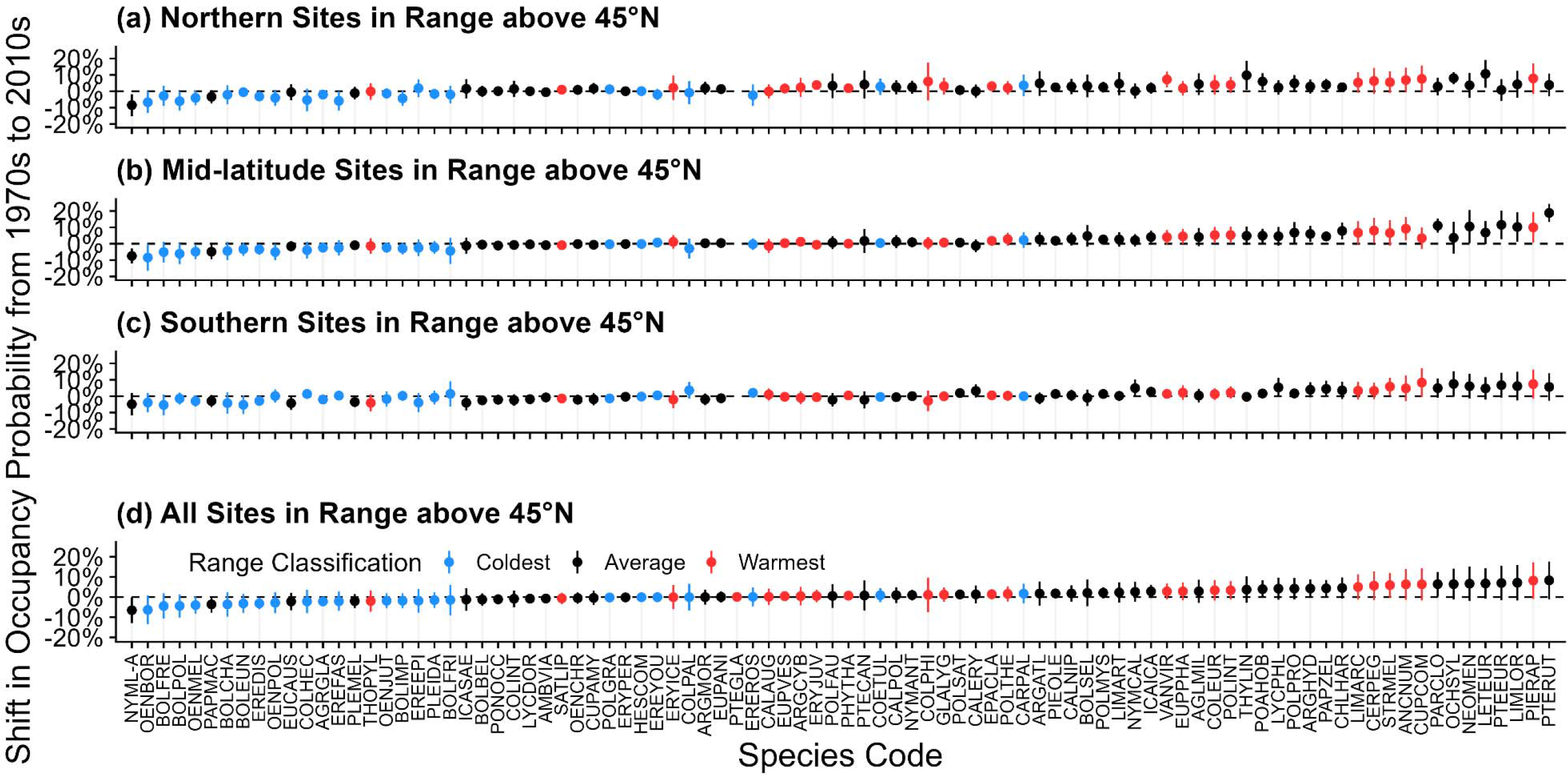
Species-specific occupancy shifts from the 1970s to 2010s where points indicate the mean occupancy shift and lines indicate one standard deviation of the variation in occupancy shift among (a) the northernmost third of cells within that species’ range in the study region, (b) the mid-latitude third of sites within that species’ range, (c) the southernmost third of sites within that species’ range, and (d) all of the sites within that species’ range (all ranges truncated at 45°N). Species are colored by quartiles of the average annual temperature in their North American ranges, so it is directly comparable to Figure 2.Results shown here are from the 200-kilometer temperature model. Note that there were too few sites to model *Pterourus glaucus* at this scale.

**Supplemental Figure S3.**
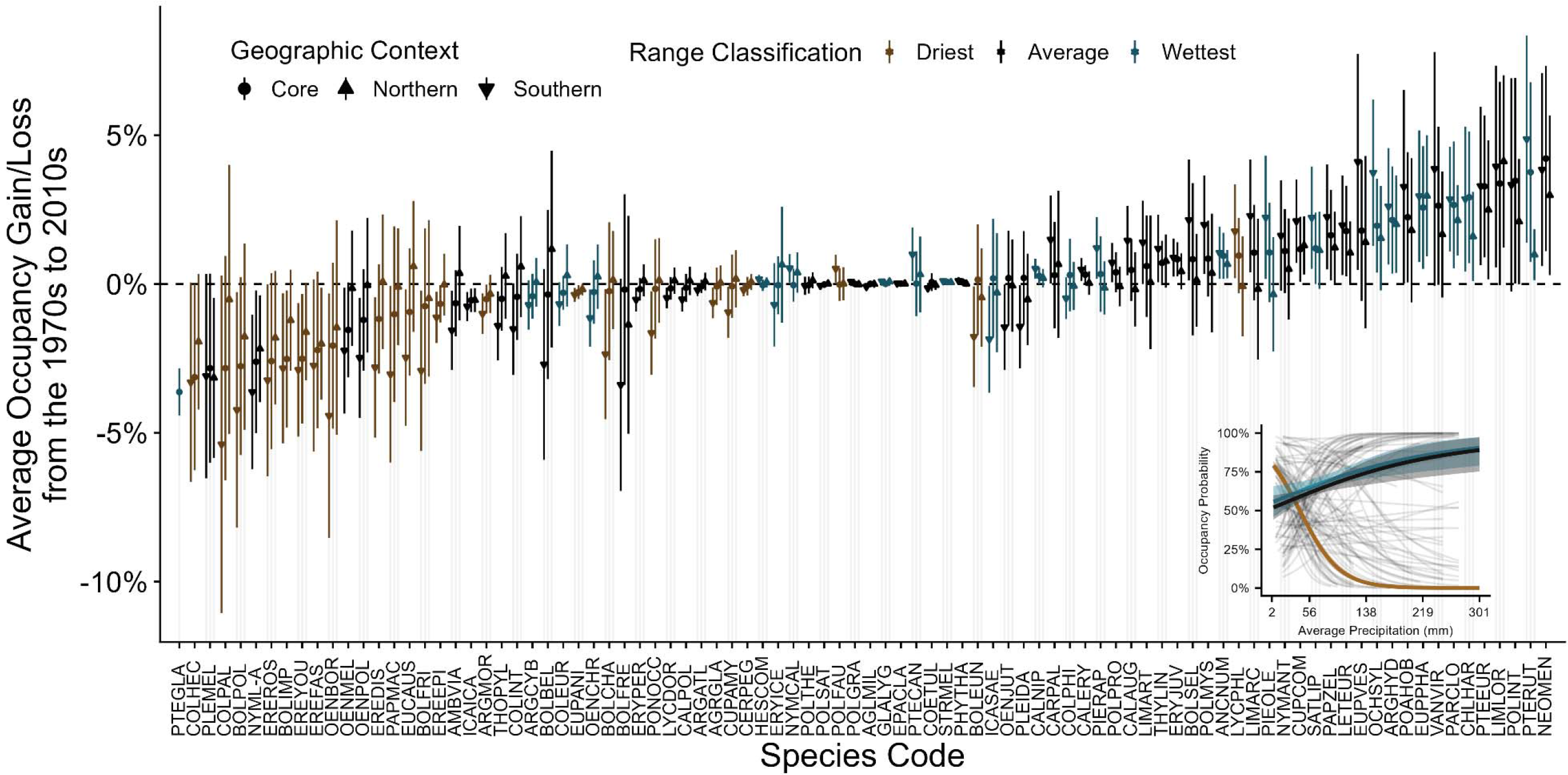
Species-specific occupancy shifts from the 1970s where points indicate the average occupancy shift for a given geographic context (core/mid-latitude, southern, and northern grid cells) and lines indicate one standard deviation of variation among relevant sites. The inset panel illustrates the relationship between precipitation and occupancy probability for each species (thin, grey lines), the middle 50% of butterflies (black line), the species with the wettest quarter of ranges (blue line), and the species with the driest quarter of ranges (brown line). Only precipitation that each species has experienced in its range are shown by the species-specific lines. Results shown here are from the 100-kilometer precipitation model.

**Supplemental Figure S4.**
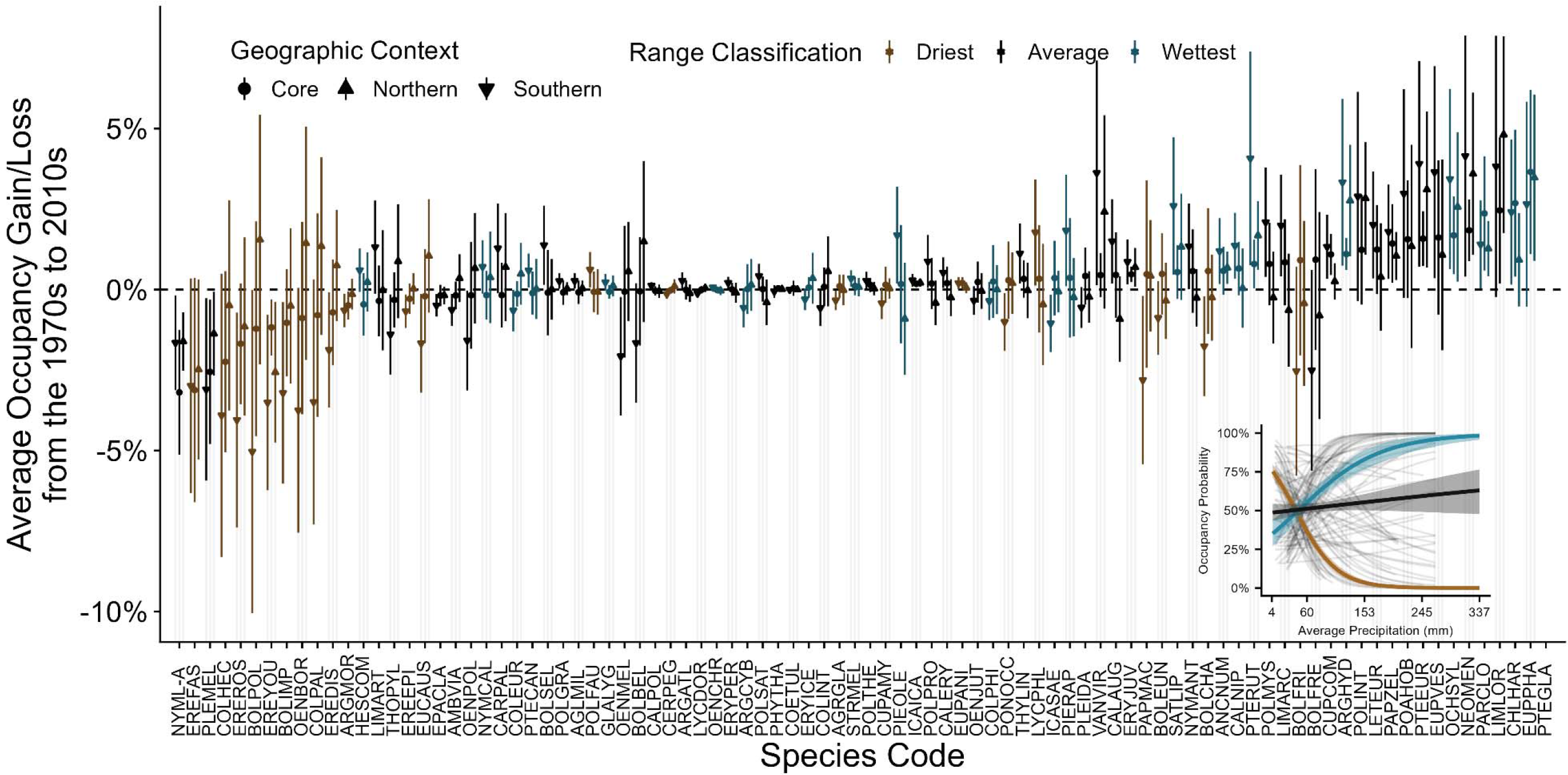
Species-specific occupancy shifts from the 1970s where points indicate the average occupancy shift for a given geographic context (core/mid-latitude, southern, and northern grid cells) and lines indicate one standard deviation of variation among relevant sites. The inset panel illustrates the relationship between precipitation and occupancy probability for each species (thin, grey lines), the middle 50% of butterflies (black line), the species with the wettest quarter of ranges (blue line), and the species with the driest quarter of ranges (red line). Only precipitation that each species has experienced in its range are shown by the species- specific lines. Results shown here are from the 200-kilometer precipitation model

**Supplemental Figure S5.**
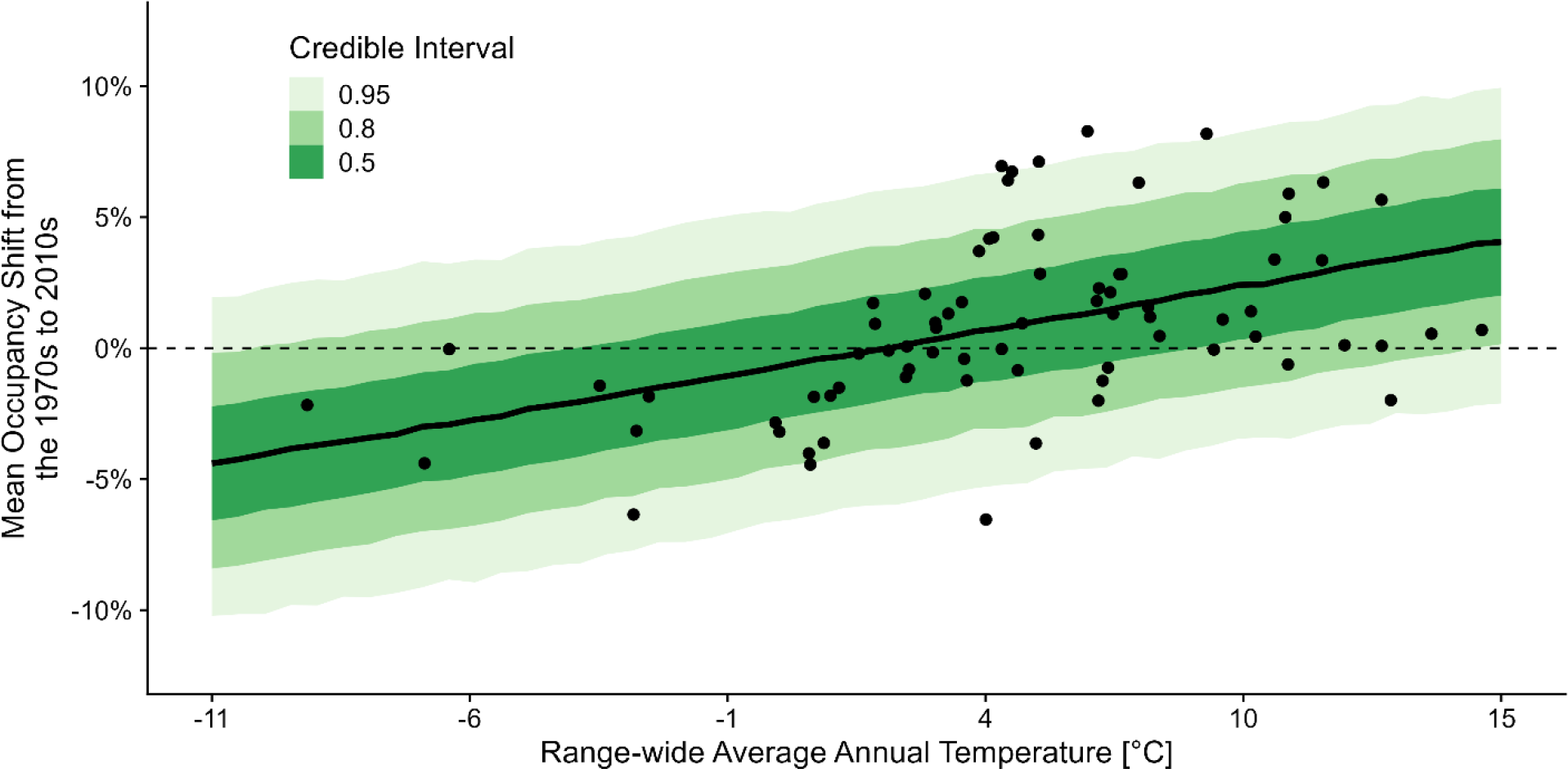
Model predicted relationship between species range-wide average annual precipitation and occupancy trend (Model C). Inference is based on trends from the 200- kilometer temperature occupancy analysis.

**Supplemental Figure S6.**
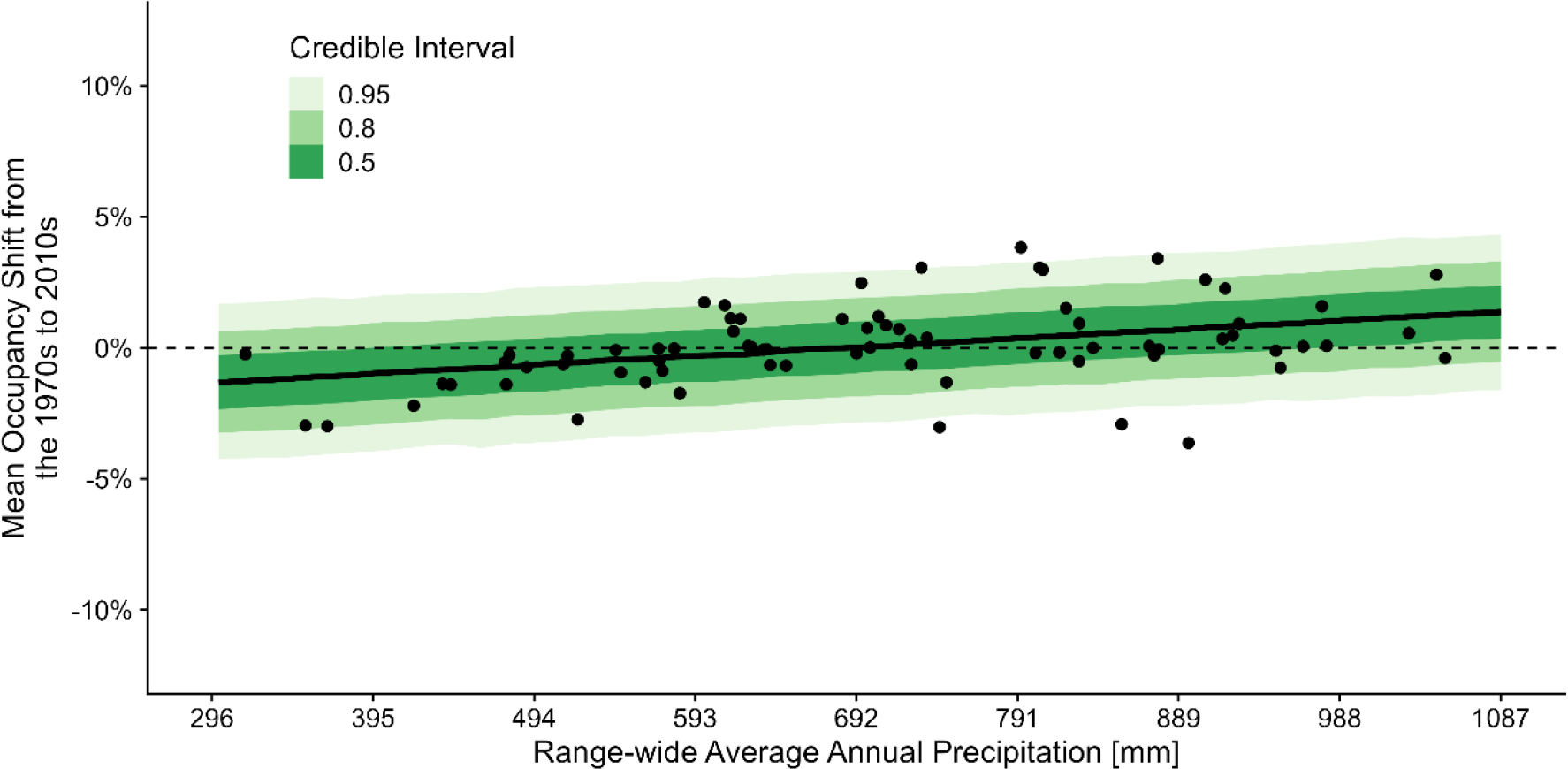
Model predicted relationship between species range-wide average annual precipitation and occupancy trend (Model C). Inference is based on trends from the 100- kilometer precipitation occupancy analysis.

**Supplemental Figure S7.**
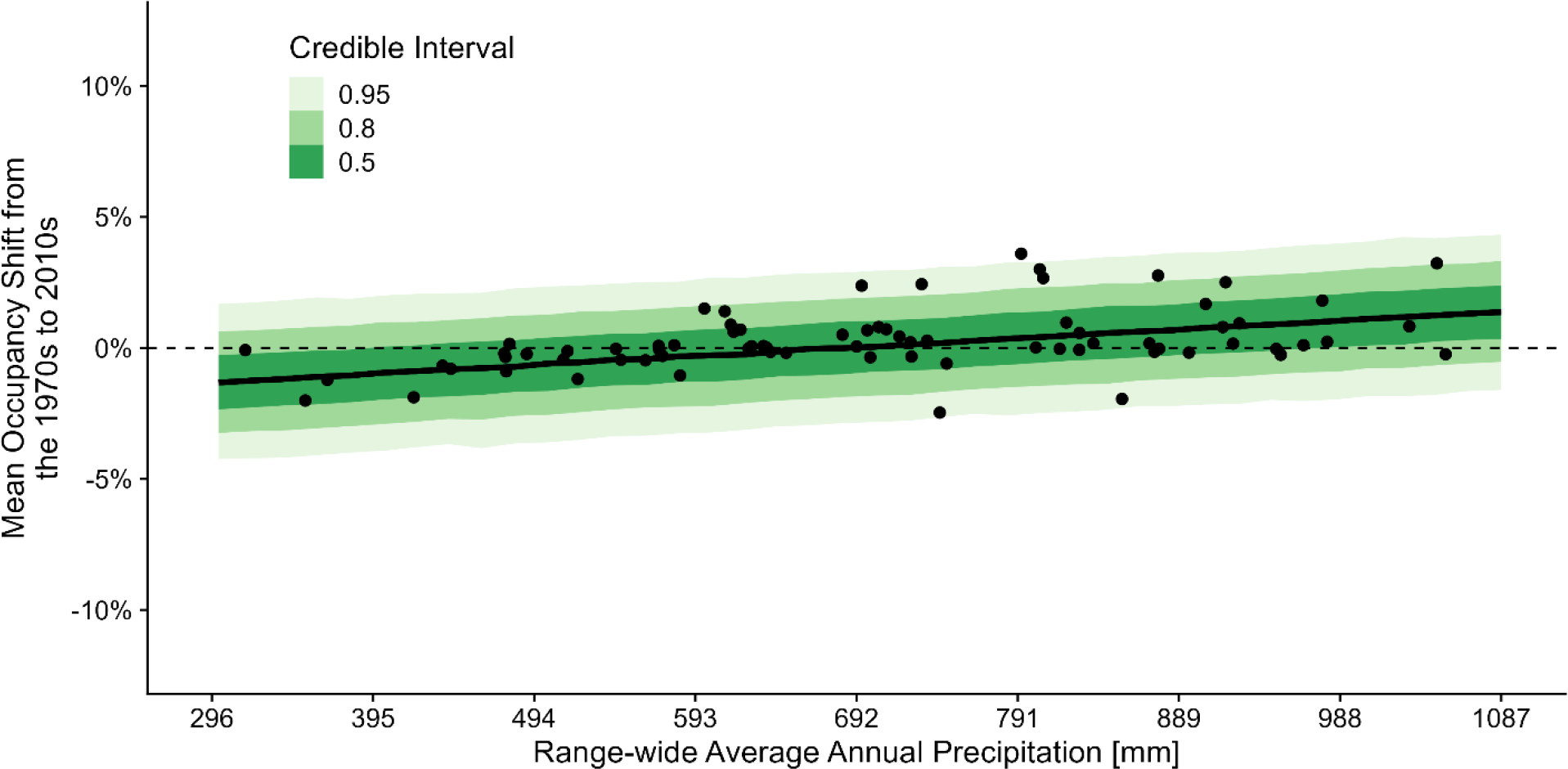
Model predicted relationship between species range-wide average annual precipitation and occupancy trend (Model C). Inference is based on trends from the 200- kilometer precipitation occupancy analysis.

**Supplemental Table S1.**
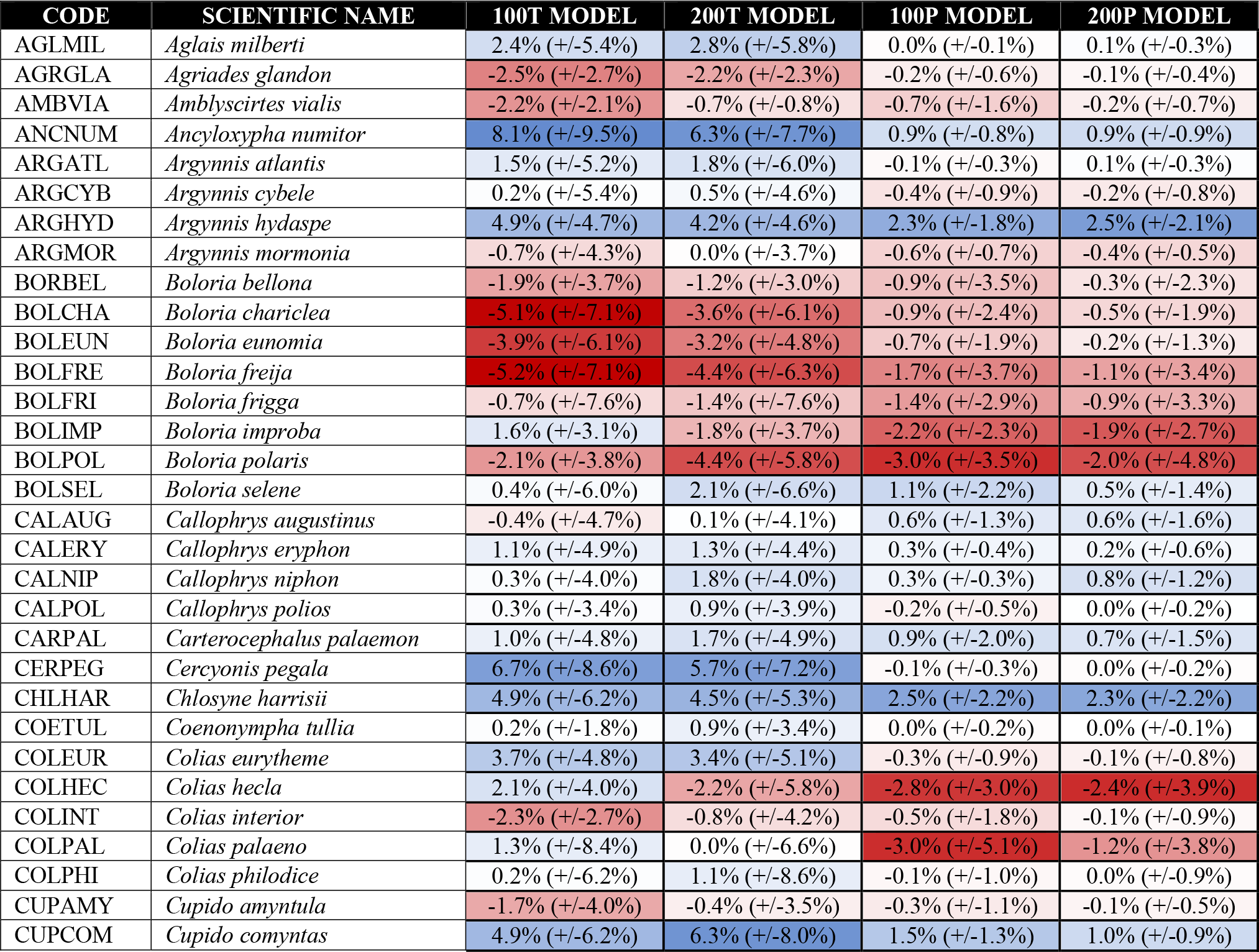
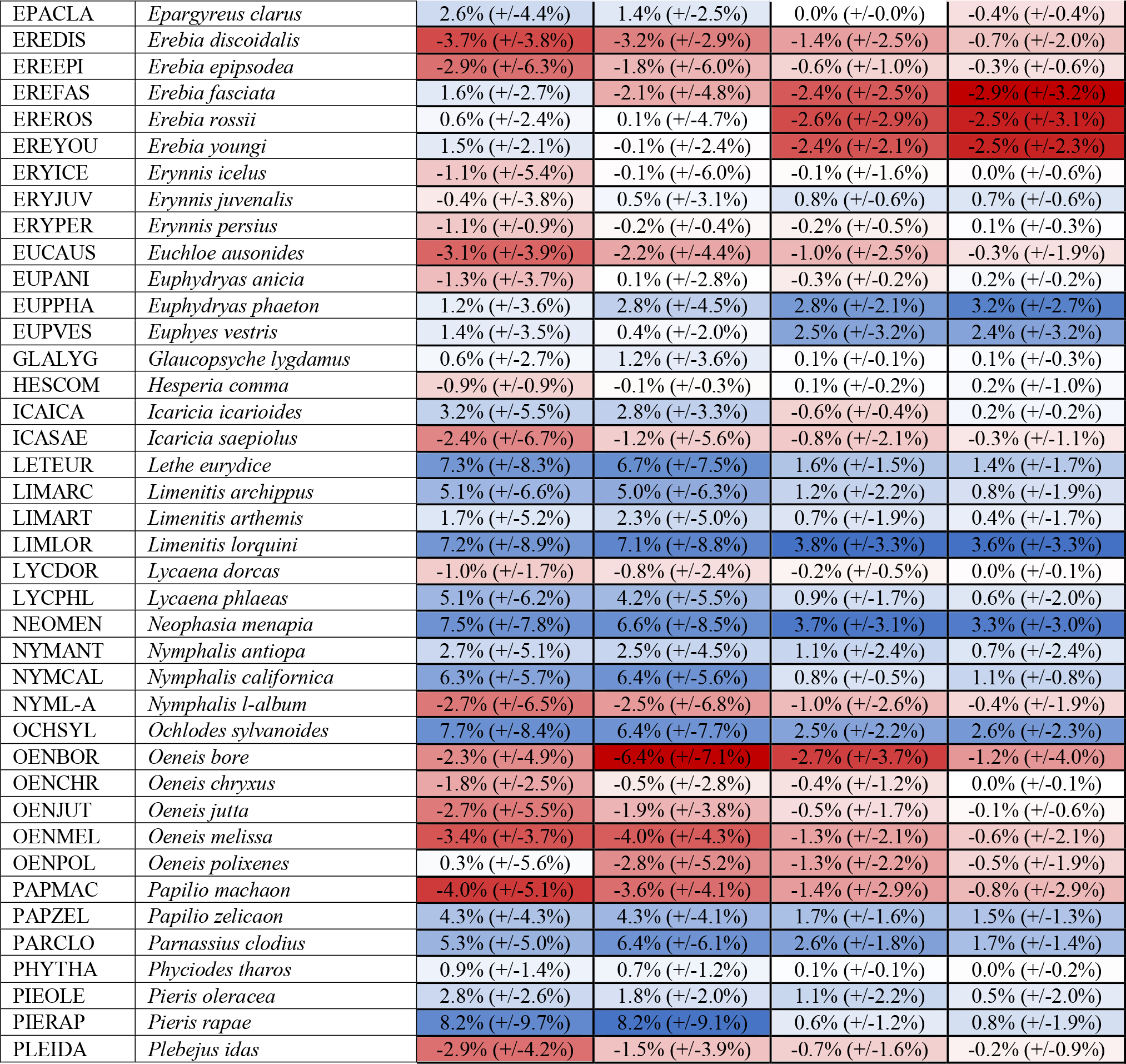
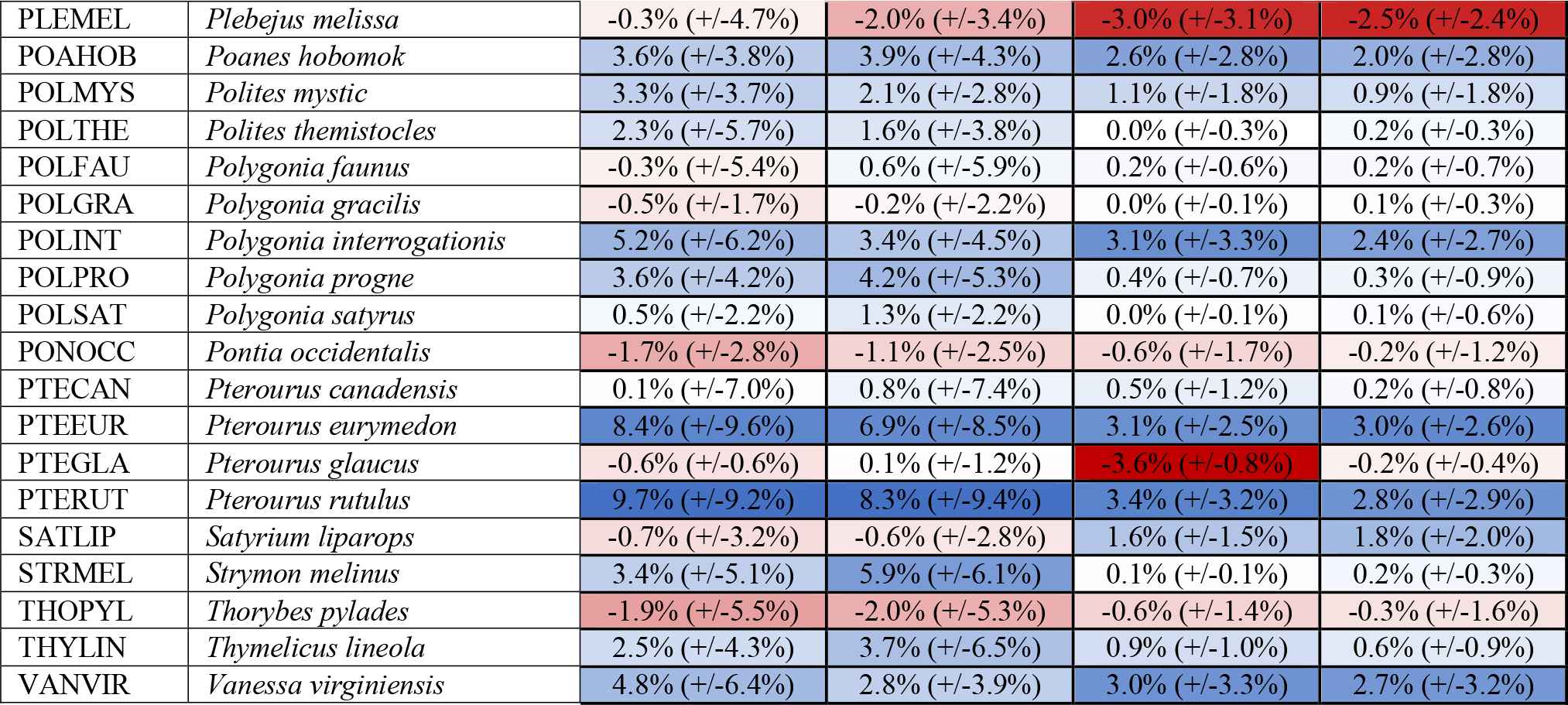
Mean occupancy trend for modeled butterfly species across all models in our analysis. Occupancy trends reflect the overall average trend and variance (+/- one standard deviation) across all cells modeled for each species. Inferences from each model type are denoted by column headers (100T = 100km temperature, 200T = 200km temperature, 100P = 100km precipitation, and 200P = 200km precipitation. Cells are conditionally formatted by the direction and magnitude of decline/increase where dark red indicates strong declines in occupancy probability and dark blue indicates strong increases in occupancy probability. White shading indicates marginal decline/increase.

| CODE | SCIENTIFIC NAME | 100T MODEL | 200T MODEL | 100P MODEL | 200P MODEL |
| --- | --- | --- | --- | --- | --- |
| AGLMIL | *Aglaia milberti* | 2.4% (+/-5.4%) | 2.8% (+/-5.8%) | 0.0% (+/-0.1%) | 0.1% (+/-0.3%) |
| AGRGLA | *Agriades glandon* | -2.5% (+/-2.7%) | -2.2% (+/-2.3%) | -0.2% (+/-0.6%) | -0.1% (+/-0.4%) |
| AMBVIA | *Amblyscirtes vialis* | -2.2% (+/-2.1%) | -0.7% (+/-0.8%) | -0.7% (+/-1.6%) | -0.2% (+/-0.7%) |
| ANCNUM | *Ancyloxypha numitor* | 8.1% (+/-9.5%) | 6.3% (+/-7.7%) | 0.9% (+/-0.8%) | 0.9% (+/-0.9%) |
| ARGATL | *Argynnis atlantis* | 1.5% (+/-5.2%) | 1.8% (+/-6.0%) | -0.1% (+/-0.3%) | 0.1% (+/-0.3%) |
| ARGCYB | *Argynnis cybele* | 0.2% (+/-5.4%) | 0.5% (+/-4.6%) | -0.4% (+/-0.9%) | -0.2% (+/-0.8%) |
| ARGHYD | *Argynnis hydaspe* | 4.9% (+/-4.7%) | 4.2% (+/-4.6%) | 2.3% (+/-1.8%) | 2.5% (+/-2.1%) |
| ARGMOR | *Argynnis mormonia* | -0.7% (+/-4.3%) | 0.0% (+/-3.7%) | -0.6% (+/-0.7%) | -0.4% (+/-0.5%) |
| BORBEL | *Boloria bellona* | -1.9% (+/-3.7%) | -1.2% (+/-3.0%) | -0.9% (+/-3.5%) | -0.3% (+/-2.3%) |
| BOLCHA | *Boloria chariclea* | -5.1% (+/-7.1%) | -3.6% (+/-6.1%) | -0.9% (+/-2.4%) | -0.5% (+/-1.9%) |
| BOLEUN | *Boloria eunomia* | -3.9% (+/-6.1%) | -3.2% (+/-4.8%) | -0.7% (+/-1.9%) | -0.2% (+/-1.3%) |
| BOLFRE | *Boloria freija* | -5.2% (+/-7.1%) | -4.4% (+/-6.3%) | -1.7% (+/-3.7%) | -1.1% (+/-3.4%) |
| BOLFRI | *Boloria frigga* | -0.7% (+/-7.6%) | -1.4% (+/-7.6%) | -1.4% (+/-2.9%) | -0.9% (+/-3.3%) |
| BOLIMP | *Boloria improba* | 1.6% (+/-3.1%) | -1.8% (+/-3.7%) | -2.2% (+/-2.3%) | -1.9% (+/-2.7%) |
| BOLPOL | *Boloria polaris* | -2.1% (+/-3.8%) | -4.4% (+/-5.8%) | -3.0% (+/-3.5%) | -2.0% (+/-4.8%) |
| BOLSEL | *Boloria selene* | 0.4% (+/-6.0%) | 2.1% (+/-6.6%) | 1.1% (+/-2.2%) | 0.5% (+/-1.4%) |
| CALAUG | *Callophrys augustinus* | -0.4% (+/-4.7%) | 0.1% (+/-4.1%) | 0.6% (+/-1.3%) | 0.6% (+/-1.6%) |
| CALERY | *Callophrys eryphon* | 1.1% (+/-4.9%) | 1.3% (+/-4.4%) | 0.3% (+/-0.4%) | 0.2% (+/-0.6%) |
| CALNIP | *Callophrys niphon* | 0.3% (+/-4.0%) | 1.8% (+/-4.0%) | 0.3% (+/-0.3%) | 0.8% (+/-1.2%) |
| CALPOL | *Callophrys polios* | 0.3% (+/-3.4%) | 0.9% (+/-3.9%) | -0.2% (+/-0.5%) | 0.0% (+/-0.2%) |
| CARPAL | *Carterocephalus palaemon* | 1.0% (+/-4.8%) | 1.7% (+/-4.9%) | 0.9% (+/-2.0%) | 0.7% (+/-1.5%) |
| CERPEG | *Cercyonis pegala* | 6.7% (+/-8.6%) | 5.7% (+/-7.2%) | -0.1% (+/-0.3%) | 0.0% (+/-0.2%) |
| CHLHAR | *Chlosyne harrisii* | 4.9% (+/-6.2%) | 4.5% (+/-5.3%) | 2.5% (+/-2.2%) | 2.3% (+/-2.2%) |
| COETUL | *Coenonympha tullia* | 0.2% (+/-1.8%) | 0.9% (+/-3.4%) | 0.0% (+/-0.2%) | 0.0% (+/-0.1%) |
| COLEUR | *Colias eurytheme* | 3.7% (+/-4.8%) | 3.4% (+/-5.1%) | -0.3% (+/-0.9%) | -0.1% (+/-0.8%) |
| COLHEC | *Colias hecla* | 2.1% (+/-4.0%) | -2.2% (+/-5.8%) | -2.8% (+/-3.0%) | -2.4% (+/-3.9%) |
| COLINT | *Colias interior* | -2.3% (+/-2.7%) | -0.8% (+/-4.2%) | -0.5% (+/-1.8%) | -0.1% (+/-0.9%) |
| COLPAL | *Colias palaeno* | 1.3% (+/-8.4%) | 0.0% (+/-6.6%) | -3.0% (+/-5.1%) | -1.2% (+/-3.8%) |
| COLPHI | *Colias philodice* | 0.2% (+/-6.2%) | 1.1% (+/-8.6%) | -0.1% (+/-1.0%) | 0.0% (+/-0.9%) |
| CUPAMY | *Cupido amyntula* | -1.7% (+/-4.0%) | -0.4% (+/-3.5%) | -0.3% (+/-1.1%) | -0.1% (+/-0.5%) |
| CUPCOM | *Cupido comyntas* | 4.9% (+/-6.2%) | 6.3% (+/-8.0%) | 1.5% (+/-1.3%) | 1.0% (+/-0.9%) |
| EPACLA | *Epargyreus clarus* | 2.6% (+/-4.4%) | 1.4% (+/-2.5%) | 0.0% (+/-0.0%) | -0.4% (+/-0.4%) |
| EREDIS | *Erebia discoidalis* | -3.7% (+/-3.8%) | -3.2% (+/-2.9%) | -1.4% (+/-2.5%) | -0.7% (+/-2.0%) |
| EREPI | *Erebia epipsodea* | -2.9% (+/-6.3%) | -1.8% (+/-6.0%) | -0.6% (+/-1.0%) | -0.3% (+/-0.6%) |
| EREFAS | *Erebia fasciata* | 1.6% (+/-2.7%) | -2.1% (+/-4.8%) | -2.4% (+/-2.5%) | -2.9% (+/-3.2%) |
| EREROS | *Erebia rossii* | 0.6% (+/-2.4%) | 0.1% (+/-4.7%) | -2.6% (+/-2.9%) | -2.5% (+/-3.1%) |
| EREYOU | *Erebia youngi* | 1.5% (+/-2.1%) | -0.1% (+/-2.4%) | -2.4% (+/-2.1%) | -2.5% (+/-2.3%) |
| ERYICE | *Erynnis icelus* | -1.1% (+/-5.4%) | -0.1% (+/-6.0%) | -0.1% (+/-1.6%) | 0.0% (+/-0.6%) |
| ERYJUV | *Erynnis juvenalis* | -0.4% (+/-3.8%) | 0.5% (+/-3.1%) | 0.8% (+/-0.6%) | 0.7% (+/-0.6%) |
| ERYPER | *Erynnis persius* | -1.1% (+/-0.9%) | -0.2% (+/-0.4%) | -0.2% (+/-0.5%) | 0.1% (+/-0.3%) |
| EUCAUS | *Euchloe ausonides* | -3.1% (+/-3.9%) | -2.2% (+/-4.4%) | -1.0% (+/-2.5%) | -0.3% (+/-1.9%) |
| EUPANI | *Euphydryas anicia* | -1.3% (+/-3.7%) | 0.1% (+/-2.8%) | -0.3% (+/-0.2%) | 0.2% (+/-0.2%) |
| EUPPHA | *Euphydryas phaeton* | 1.2% (+/-3.6%) | 2.8% (+/-4.5%) | 2.8% (+/-2.1%) | 3.2% (+/-2.7%) |
| EUPVES | *Euphyes vestris* | 1.4% (+/-3.5%) | 0.4% (+/-2.0%) | 2.5% (+/-3.2%) | 2.4% (+/-3.2%) |
| GLALYG | *Glaucopsyche lygdamus* | 0.6% (+/-2.7%) | 1.2% (+/-3.6%) | 0.1% (+/-0.1%) | 0.1% (+/-0.3%) |
| HESCOM | *Hesperia comma* | -0.9% (+/-0.9%) | -0.1% (+/-0.3%) | 0.1% (+/-0.2%) | 0.2% (+/-1.0%) |
| ICAICA | *Icaricia icarioides* | 3.2% (+/-5.5%) | 2.8% (+/-3.3%) | -0.6% (+/-0.4%) | 0.2% (+/-0.2%) |
| ICASAE | *Icaricia saepiolus* | -2.4% (+/-6.7%) | -1.2% (+/-5.6%) | -0.8% (+/-2.1%) | -0.3% (+/-1.1%) |
| LETEUR | *Lethe eurydice* | 7.3% (+/-8.3%) | 6.7% (+/-7.5%) | 1.6% (+/-1.5%) | 1.4% (+/-1.7%) |
| LIMARC | *Limenitis archippus* | 5.1% (+/-6.6%) | 5.0% (+/-6.3%) | 1.2% (+/-2.2%) | 0.8% (+/-1.9%) |
| LIMART | *Limenitis arthemis* | 1.7% (+/-5.2%) | 2.3% (+/-5.0%) | 0.7% (+/-1.9%) | 0.4% (+/-1.7%) |
| LIMLOR | *Limenitis lorquini* | 7.2% (+/-8.9%) | 7.1% (+/-8.8%) | 3.8% (+/-3.3%) | 3.6% (+/-3.3%) |
| LYCDOR | *Lycaena dorcas* | -1.0% (+/-1.7%) | -0.8% (+/-2.4%) | -0.2% (+/-0.5%) | 0.0% (+/-0.1%) |
| LYCPHL | *Lycaena phlaeas* | 5.1% (+/-6.2%) | 4.2% (+/-5.5%) | 0.9% (+/-1.7%) | 0.6% (+/-2.0%) |
| NEOMEN | *Neophasia menapia* | 7.5% (+/-7.8%) | 6.6% (+/-8.5%) | 3.7% (+/-3.1%) | 3.3% (+/-3.0%) |
| NYMANT | *Nymphalis antiopa* | 2.7% (+/-5.1%) | 2.5% (+/-4.5%) | 1.1% (+/-2.4%) | 0.7% (+/-2.4%) |
| NYMCAL | *Nymphalis californica* | 6.3% (+/-5.7%) | 6.4% (+/-5.6%) | 0.8% (+/-0.5%) | 1.1% (+/-0.8%) |
| NYML-A | *Nymphalis l-album* | -2.7% (+/-6.5%) | -2.5% (+/-6.8%) | -1.0% (+/-2.6%) | -0.4% (+/-1.9%) |
| OCHSYL | *Ochlodes sylvanoides* | 7.7% (+/-8.4%) | 6.4% (+/-7.7%) | 2.5% (+/-2.2%) | 2.6% (+/-2.3%) |
| OENBOR | *Oeneis bore* | -2.3% (+/-4.9%) | -6.4% (+/-7.1%) | -2.7% (+/-3.7%) | -1.2% (+/-4.0%) |
| OENCHR | *Oeneis chryxus* | -1.8% (+/-2.5%) | -0.5% (+/-2.8%) | -0.4% (+/-1.2%) | 0.0% (+/-0.1%) |
| OENJUT | *Oeneis jutta* | -2.7% (+/-5.5%) | -1.9% (+/-3.8%) | -0.5% (+/-1.7%) | -0.1% (+/-0.6%) |
| OENMEL | *Oeneis melissa* | -3.4% (+/-3.7%) | -4.0% (+/-4.3%) | -1.3% (+/-2.1%) | -0.6% (+/-2.1%) |
| OENPOL | *Oeneis polixenes* | 0.3% (+/-5.6%) | -2.8% (+/-5.2%) | -1.3% (+/-2.2%) | -0.5% (+/-1.9%) |
| PAPMAC | *Papilio machaon* | -4.0% (+/-5.1%) | -3.6% (+/-4.1%) | -1.4% (+/-2.9%) | -0.8% (+/-2.9%) |
| PAPZEL | *Papilio zelicaon* | 4.3% (+/-4.3%) | 4.3% (+/-4.1%) | 1.7% (+/-1.6%) | 1.5% (+/-1.3%) |
| PARCLO | *Parnassius clodius* | 5.3% (+/-5.0%) | 6.4% (+/-6.1%) | 2.6% (+/-1.8%) | 1.7% (+/-1.4%) |
| PHYTHA | *Phyciodes tharos* | 0.9% (+/-1.4%) | 0.7% (+/-1.2%) | 0.1% (+/-0.1%) | 0.0% (+/-0.2%) |
| PIEOLE | *Pieris oleracea* | 2.8% (+/-2.6%) | 1.8% (+/-2.0%) | 1.1% (+/-2.2%) | 0.5% (+/-2.0%) |
| PIERAP | *Pieris rapae* | 8.2% (+/-9.7%) | 8.2% (+/-9.1%) | 0.6% (+/-1.2%) | 0.8% (+/-1.9%) |
| PLEIDA | *Plebejus idas* | -2.9% (+/-4.2%) | -1.5% (+/-3.9%) | -0.7% (+/-1.6%) | -0.2% (+/-0.9%) |
| PLEMEL | *Plebejus melissa* | -0.3% (+/-4.7%) | -2.0% (+/-3.4%) | -3.0% (+/-3.1%) | -2.5% (+/-2.4%) |
| POAHOB | *Poanes hobomok* | 3.6% (+/-3.8%) | 3.9% (+/-4.3%) | 2.6% (+/-2.8%) | 2.0% (+/-2.8%) |
| POLMYS | *Polites mystic* | 3.3% (+/-3.7%) | 2.1% (+/-2.8%) | 1.1% (+/-1.8%) | 0.9% (+/-1.8%) |
| POLTHE | *Polites themistocles* | 2.3% (+/-5.7%) | 1.6% (+/-3.8%) | 0.0% (+/-0.3%) | 0.2% (+/-0.3%) |
| POLFAU | *Polygonia faunus* | -0.3% (+/-5.4%) | 0.6% (+/-5.9%) | 0.2% (+/-0.6%) | 0.2% (+/-0.7%) |
| POLGRA | *Polygonia gracilis* | -0.5% (+/-1.7%) | -0.2% (+/-2.2%) | 0.0% (+/-0.1%) | 0.1% (+/-0.3%) |
| POLINT | *Polygonia interrogationis* | 5.2% (+/-6.2%) | 3.4% (+/-4.5%) | 3.1% (+/-3.3%) | 2.4% (+/-2.7%) |
| POLPRO | *Polygonia progne* | 3.6% (+/-4.2%) | 4.2% (+/-5.3%) | 0.4% (+/-0.7%) | 0.3% (+/-0.9%) |
| POLSAT | *Polygonia satyrus* | 0.5% (+/-2.2%) | 1.3% (+/-2.2%) | 0.0% (+/-0.1%) | 0.1% (+/-0.6%) |
| PONOCC | *Pontia occidentalis* | -1.7% (+/-2.8%) | -1.1% (+/-2.5%) | -0.6% (+/-1.7%) | -0.2% (+/-1.2%) |
| PTECAN | *Pterourus canadensis* | 0.1% (+/-7.0%) | 0.8% (+/-7.4%) | 0.5% (+/-1.2%) | 0.2% (+/-0.8%) |
| PTEEUR | *Pterourus eurymedon* | 8.4% (+/-9.6%) | 6.9% (+/-8.5%) | 3.1% (+/-2.5%) | 3.0% (+/-2.6%) |
| PTGLA | *Pterourus glaucus* | -0.6% (+/-0.6%) | 0.1% (+/-1.2%) | -3.6% (+/-0.8%) | -0.2% (+/-0.4%) |
| PTERUT | *Pterourus rutulus* | 9.7% (+/-9.2%) | 8.3% (+/-9.4%) | 3.4% (+/-3.2%) | 2.8% (+/-2.9%) |
| SATLIP | *Satyrrium liparops* | -0.7% (+/-3.2%) | -0.6% (+/-2.8%) | 1.6% (+/-1.5%) | 1.8% (+/-2.0%) |
| STRMEL | *Strymon melinus* | 3.4% (+/-5.1%) | 5.9% (+/-6.1%) | 0.1% (+/-0.1%) | 0.2% (+/-0.3%) |
| THOPYL | *Thorybes pylades* | -1.9% (+/-5.5%) | -2.0% (+/-5.3%) | -0.6% (+/-1.4%) | -0.3% (+/-1.6%) |
| THYLIN | *Thymelicus lineola* | 2.5% (+/-4.3%) | 3.7% (+/-6.5%) | 0.9% (+/-1.0%) | 0.6% (+/-0.9%) |
| VANVIR | *Vanessa virginiensis* | 4.8% (+/-6.4%) | 2.8% (+/-3.9%) | 3.0% (+/-3.3%) | 2.7% (+/-3.2%) |

**Supplemental Table S2.** Model comparison metrics for models predicting average occupancy shift in our study region since the 1970s. Results are shown for the 100-kilometer scale, temperature analysis. Top candidate model is underlined and in bold. We considered models equivalent if the difference in ELPD +/- the standard error of the ELPD estimate overlapped zero and favored models which used fewer predictors if this was the case.

| MODEL | ELDP-LOO | DELTA ELPD | SE DELTA ELPD |
| --- | --- | --- | --- |
| A | 145.6 | -5.1 | 2.5 |
| B | 147.0 | -3.7 | 3.2 |
| <u>**C**</u> | <u>**150.7**</u> | <u>**0.0**</u> | <u>**0.0**</u> |
| <u>**D**</u> | <u>**150.1**</u> | <u>**-0.6**</u> | <u>**1.5**</u> |
| E | 144.7 | -6.1 | 2.9 |
| F | 145.9 | -4.9 | 3.6 |
| <u>**G**</u> | <u>**147.5**</u> | <u>**-3.3**</u> | <u>**3.8**</u> |
| <u>**H**</u> | <u>**147.1**</u> | <u>**-3.7**</u> | <u>**4.0**</u> |
| I | 144.1 | -6.6 | 2.6 |
| J | 145.6 | -5.1 | 3.3 |
| K | 144.3 | -6.4 | 2.9 |
| L | 145.0 | -5.8 | 2.6 |
| M | 146.0 | -4.7 | 4.3 |
| N | 145.1 | -5.7 | 4.4 |

**Supplemental Table S3.** Model comparison metrics for models predicting average occupancy shift since the 1970s. Results are shown for the 200-kilometer scale, temperature analysis. Top candidate model is underlinedand bold. We considered models equivalent if the difference in ELPD +/- the standard error of the ELPD estimate overlapped zero and favored models which used fewer predictors if this was the case.

| MODEL | ELDP-LOO | DELTA ELPD | SE DELTA ELPD |
| --- | --- | --- | --- |
| A | 147.5 | -8.3 | 3.3 |
| B | 150.9 | -5.0 | 4.1 |
| <u>C</u> | <u>155.8</u> | <u>0.0</u> | <u>0.0</u> |
| <u>D</u> | <u>155.0</u> | <u>-0.8</u> | <u>1.6</u> |
| E | 146.5 | -9.3 | 3.8 |
| F | 149.4 | -6.5 | 4.2 |
| G | 148.3 | -7.5 | 3.9 |
| H | 149.9 | -5.9 | 4.3 |
| I | 146.1 | -9.7 | 3.4 |
| J | 149.0 | -6.8 | 4.2 |
| K | 148.9 | -6.9 | 3.2 |
| L | 150.4 | -5.4 | 2.7 |
| M | 149.9 | -6.0 | 4.2 |
| N | 149.3 | -6.5 | 4.3 |

**Supplemental Table S4.** Model comparison metrics for models predicting average occupancy shift since the 1970s. Results are shown for the 100-kilometer scale, precipitation analysis. Top candidate model is underlined and bold. We considered models equivalent if the difference in ELPD +/- the standard error of the ELPD estimate overlapped zero and favored models which used fewer predictors if this was the case.

| MODEL | ELDP-LOO | DELTA ELPD | SE DELTA ELPD |
| --- | --- | --- | --- |
| A | -408.3 | -5.4 | 3.6 |
| B | -408.0 | -5.6 | 2.7 |
| **<u>C</u>** | **<u>-419.1</u>** | **<u>0.0</u>** | **<u>0.0</u>** |
| **<u>D</u>** | **<u>-416.9</u>** | **<u>-1.1</u>** | **<u>1.7</u>** |
| E | -406.5 | -6.3 | 3.5 |
| F | -405.9 | -6.6 | 2.7 |
| G | -404.8 | -7.1 | 3.0 |
| H | -403.9 | -7.6 | 3.7 |
| I | -406.5 | -6.3 | 3.4 |
| J | -405.8 | -6.7 | 3.0 |
| K | -408.5 | -5.3 | 3.2 |
| L | -406.5 | -6.3 | 3.4 |
| M | -406.9 | -6.1 | 4.2 |
| N | -404.7 | -7.2 | 4.4 |

**Supplemental Table S5.** Model comparison metrics for models predicting average occupancy shift since the 1970s. Results are shown for the 200-kilometer scale, precipitation analysis. Top candidate model is underlined and bold. We considered models equivalent if the difference in ELPD +/- the standard error of the ELPD estimate overlapped zero and favored models which used fewer predictors if this was the case.

| MODEL | ELDP-LOO | DELTA ELPD | SE DELTA ELPD |
| --- | --- | --- | --- |
| A | 225.0 | -6.9 | 3.5 |
| B | 226.1 | -5.8 | 3.6 |
| <u>C</u> | <u>231.9</u> | <u>0.0</u> | <u>0.0</u> |
| <u>D</u> | <u>231.6</u> | <u>-0.3</u> | <u>1.3</u> |
| E | 225.0 | -6.9 | 3.7 |
| F | 225.4 | -6.5 | 3.5 |
| G | 226.1 | -5.8 | 3.5 |
| H | 226.1 | -5.8 | 3.7 |
| I | 224.4 | -7.5 | 3.7 |
| J | 225.3 | -6.6 | 4.0 |
| K | 225.1 | -6.9 | 3.1 |
| L | 224.8 | -7.1 | 3.7 |
| <u>M</u> | <u>228.2</u> | <u>-3.7</u> | <u>4.7</u> |
| <u>N</u> | <u>227.2</u> | <u>-4.7</u> | <u>4.7</u> |

