## Supplemental File S2 for "Rising minimum temperatures contribute to 50 years of shifting Arctic and boreal butterfly communities in North America"

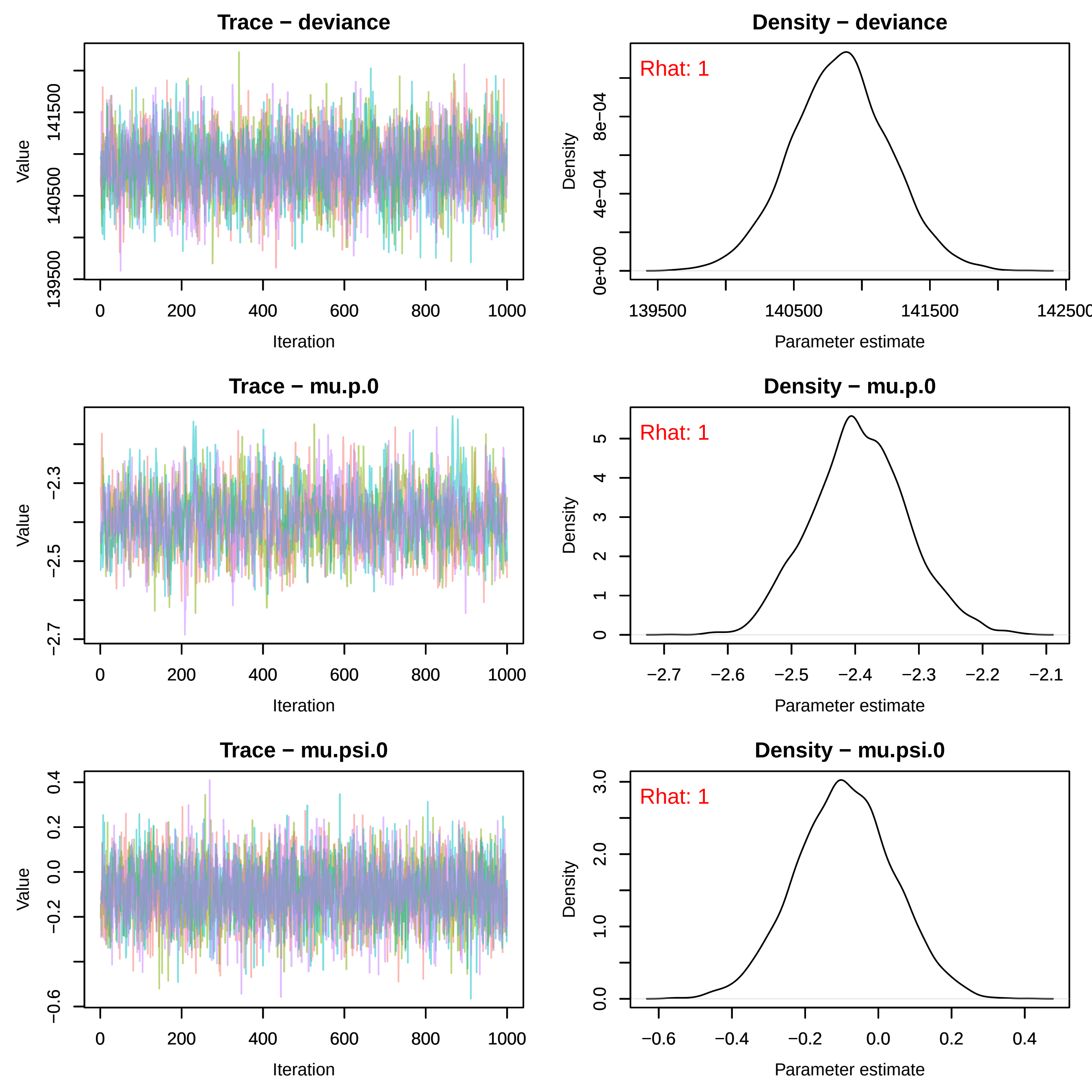
Trace − deviance
Density − deviance
Rhat: 1
Trace − mu.p.0
Density − mu.p.0
Rhat: 1
Trace − mu.psi.0
Density − mu.psi.0
Rhat: 1
Value
Iteration
Density
Parameter estimate

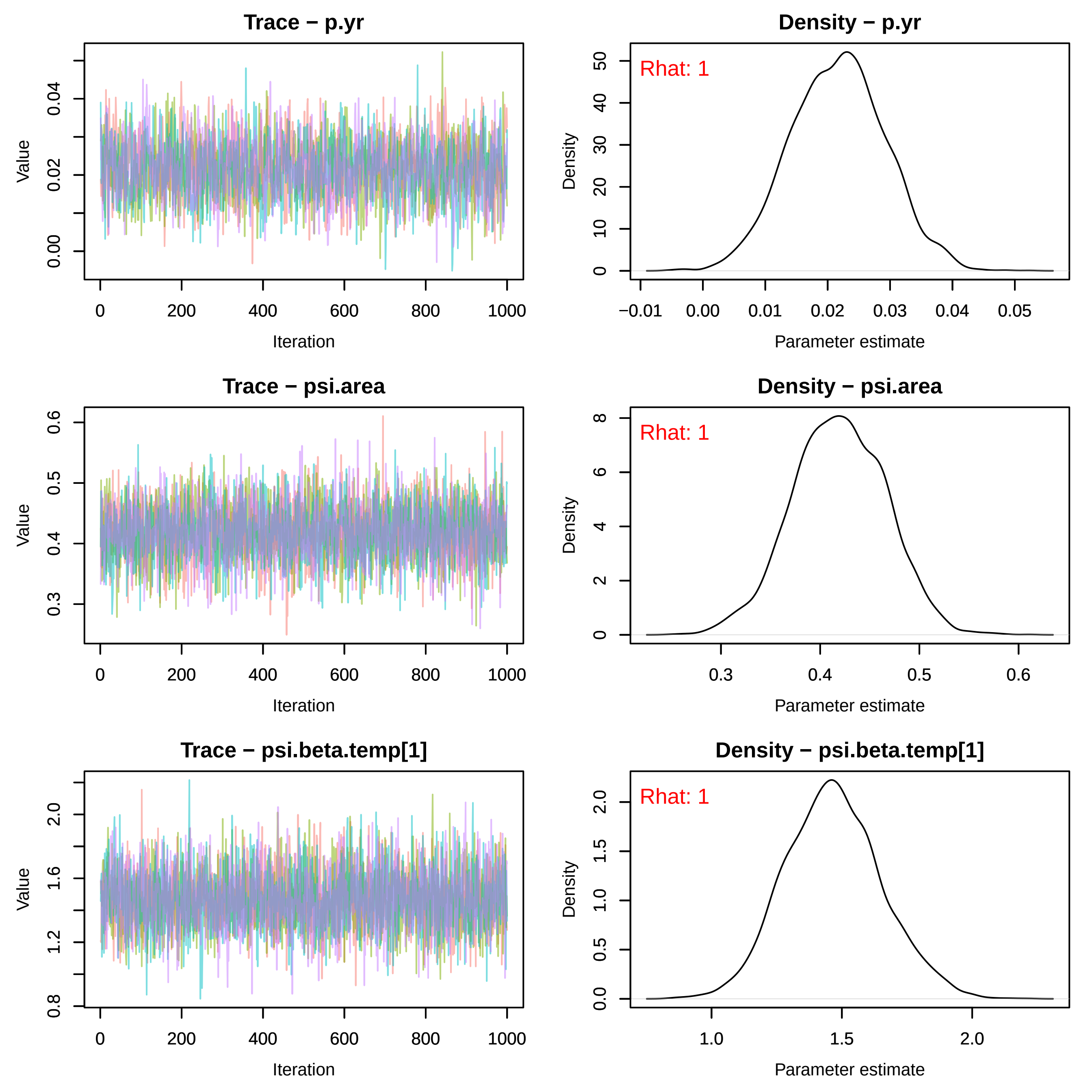
Trace − p.yr
Density − p.yr
Rhat: 1
Trace − psi.area
Density − psi.area
Rhat: 1
Trace − psi.beta.temp[1]
Density − psi.beta.temp[1]
Rhat: 1
Value
Density
Iteration
Parameter estimate

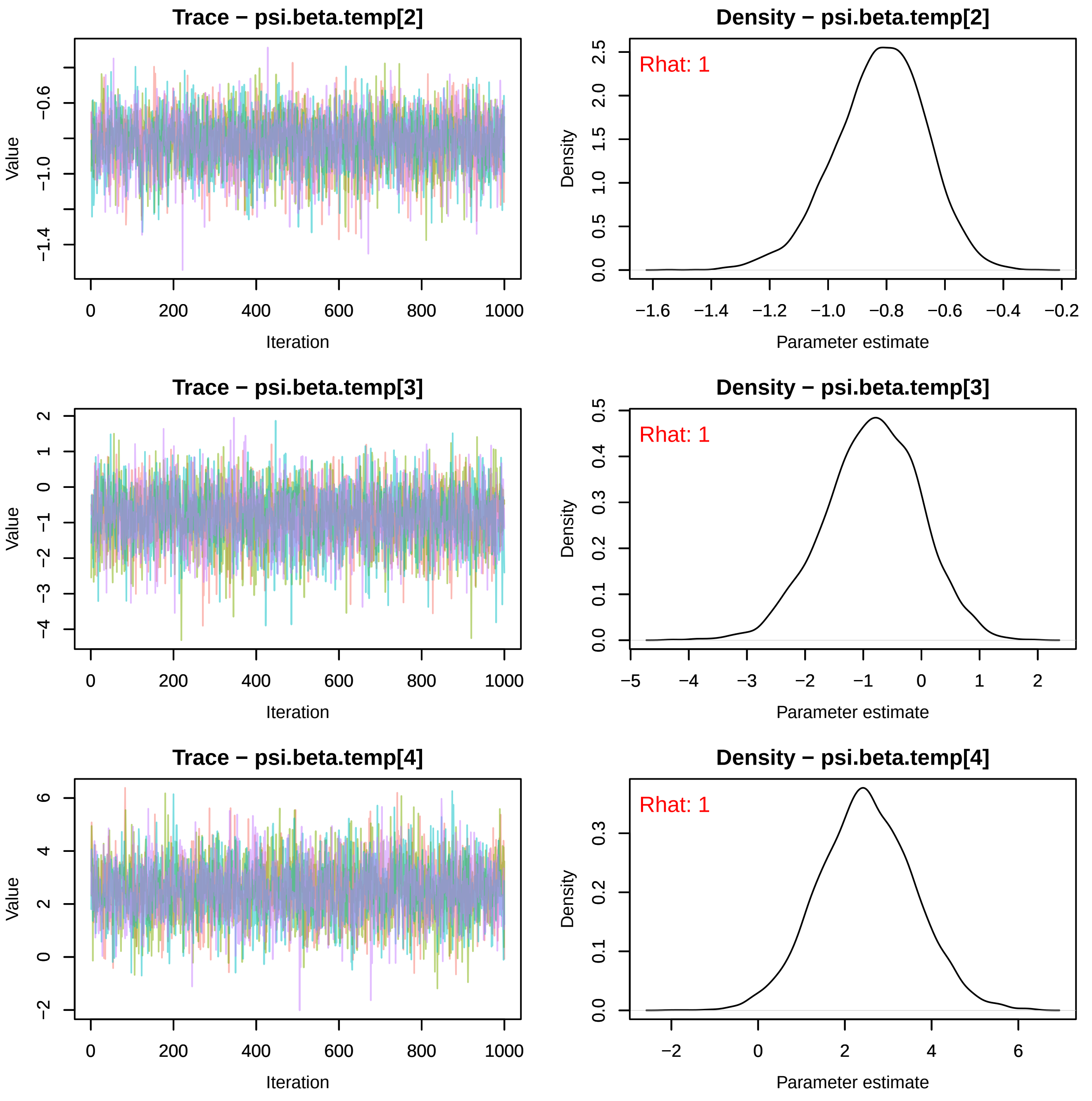
Trace − psi.beta.temp[2]
Density − psi.beta.temp[2]
Trace − psi.beta.temp[3]
Density − psi.beta.temp[3]
Trace − psi.beta.temp[4]
Density − psi.beta.temp[4]
Value
Iteration
Density
Parameter estimate
Rhat: 1
Rhat: 1
Rhat: 1
0
200
400
600
800
1000
−0.6
−1.0
−1.4
0.0
0.5
1.0
1.5
2.0
2.5
−1.6
−1.4
−1.2
−1.0
−0.8
−0.6
−0.4
−0.2
2
1
0
−1
−2
−3
−4
0.1
0.2
0.3
0.4
−5
6
4

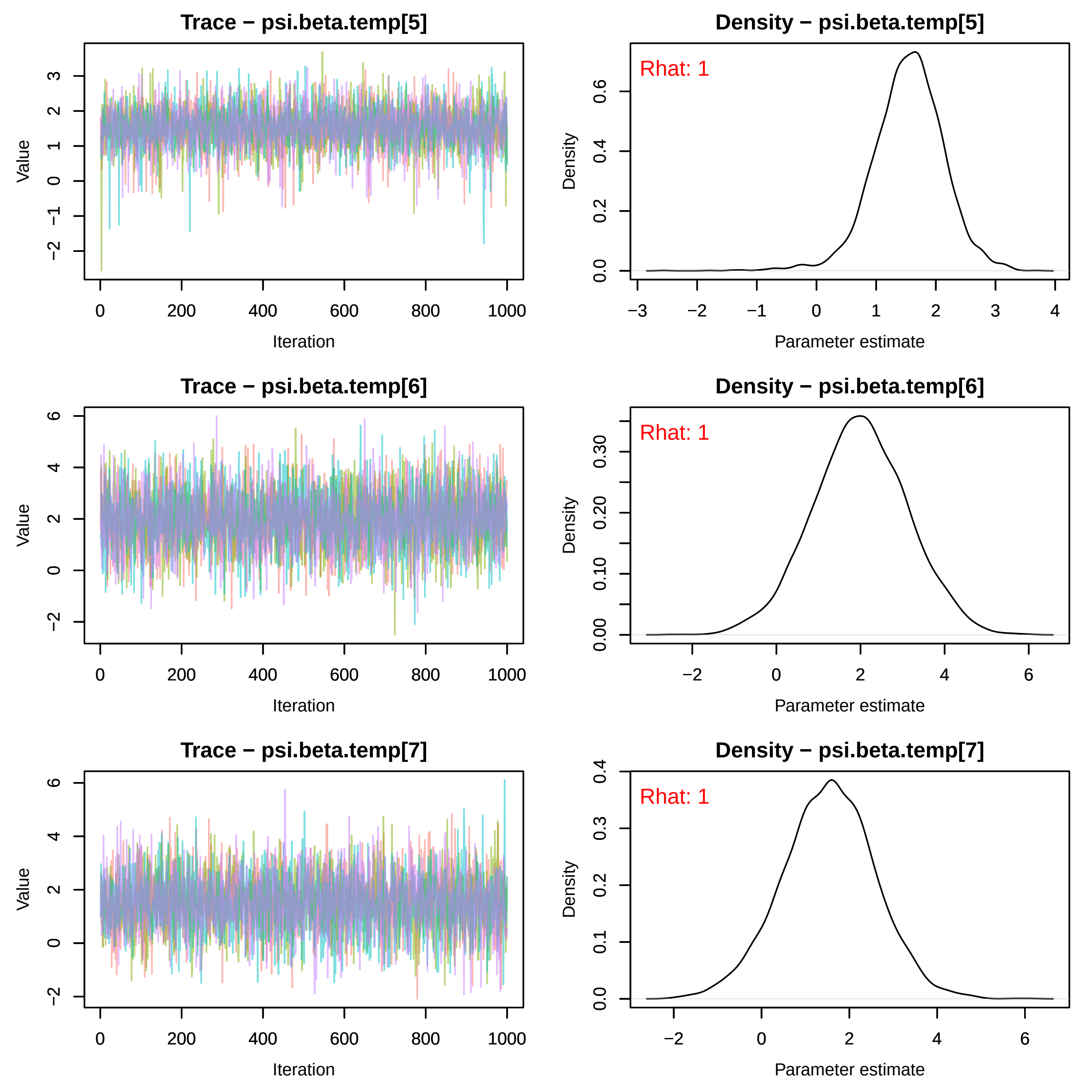
Trace − psi.beta.temp[5]
Value
Iteration
0 200 400 600 800 1000
−2 −1 0 1 2 3
Density − psi.beta.temp[5]
Rhat: 1
Density
Parameter estimate
−3 −2 −1 0 1 2 3 4
0.0 0.2 0.4 0.6
Trace − psi.beta.temp[6]
Value
Iteration
0 200 400 600 800 1000
−2 0 2 4 6
Density − psi.beta.temp[6]
Rhat: 1
Density
Parameter estimate
−2 0 2 4 6
0.00 0.10 0.20 0.30
Trace − psi.beta.temp[7]
Value
Iteration
0 200 400 600 800 1000
−2 0 2 4 6
Density − psi.beta.temp[7]
Rhat: 1
Density
Parameter estimate
−2 0 2 4 6
0.0 0.1 0.2 0.3 0.4

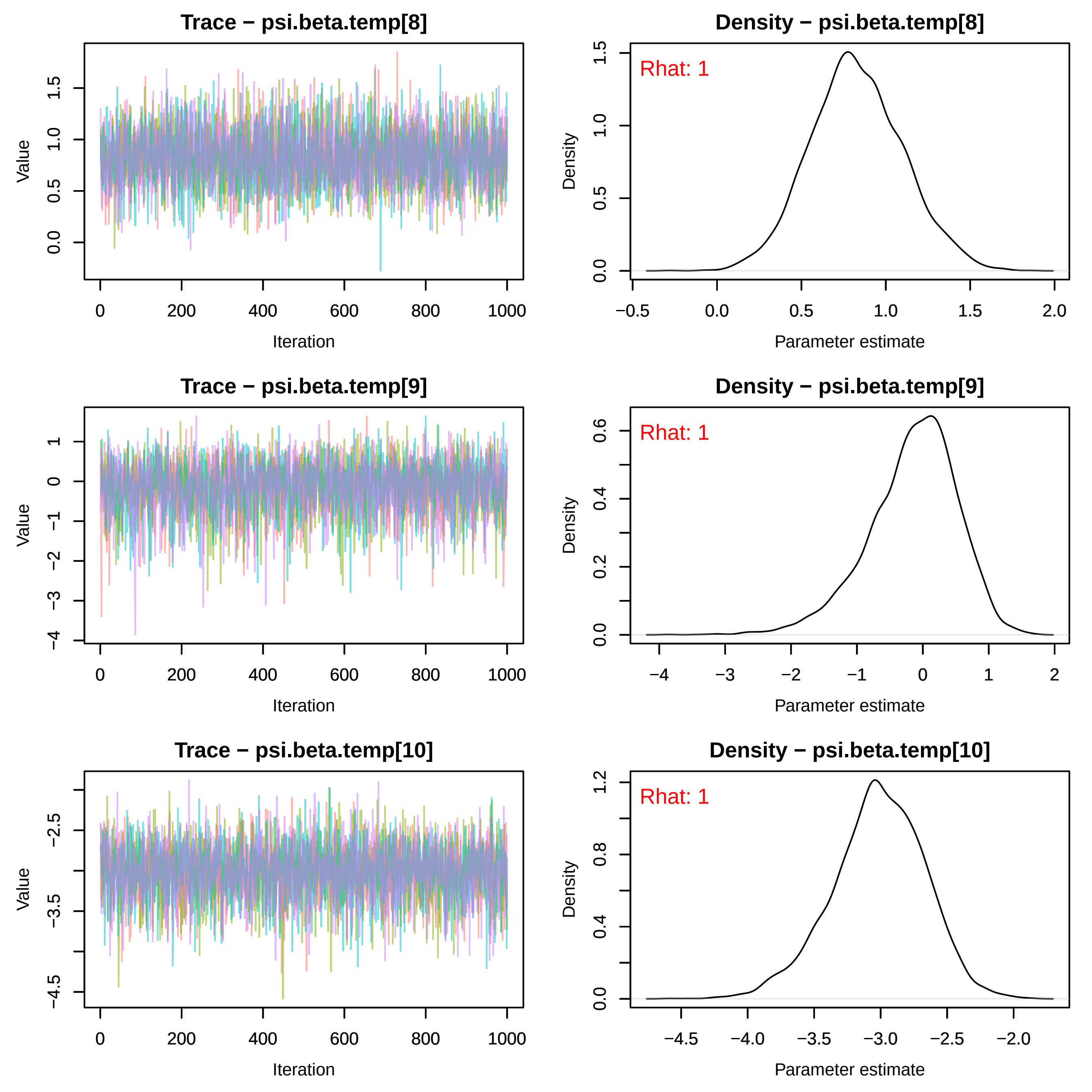
Trace − psi.beta.temp[8]
Value
Iteration
Density − psi.beta.temp[8]
Rhat: 1
Density
Parameter estimate
Trace − psi.beta.temp[9]
Value
Iteration
Density − psi.beta.temp[9]
Rhat: 1
Density
Parameter estimate
Trace − psi.beta.temp[10]
Value
Iteration
Density − psi.beta.temp[10]
Rhat: 1
Density
Parameter estimate

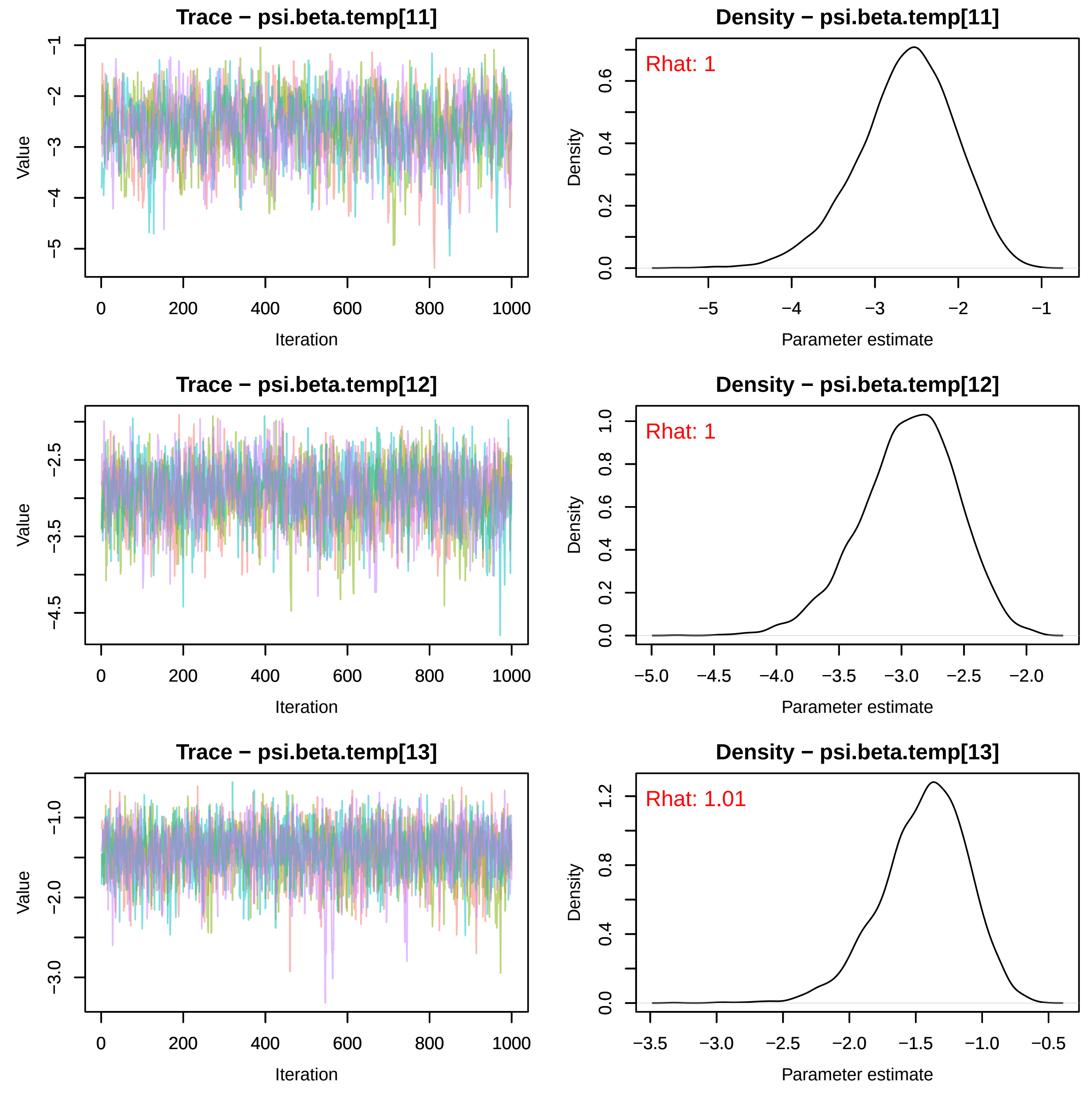
Trace − psi.beta.temp[11]
Value
Iteration
0
200
400
600
800
1000
−1
−2
−3
−4
−5
Density − psi.beta.temp[11]
Rhat: 1
Density
Parameter estimate
0.0
0.2
0.4
0.6
−5
−4
−3
−2
−1
Trace − psi.beta.temp[12]
Value
Iteration
−2.5
−3.5
−4.5
Density − psi.beta.temp[12]
Rhat: 1
Density
Parameter estimate
0.0
0.2
0.4
0.6
0.8
1.0
−5.0
−4.5
−4.0
−3.5
−3.0
−2.5
−2.0
Trace − psi.beta.temp[13]
Value
Iteration
−1.0
−2.0
−3.0
Density − psi.beta.temp[13]
Rhat: 1.01
Density
Parameter estimate
0.0
0.4
0.8
1.2
−3.5
−3.0
−2.5
−2.0
−1.5
−1.0
−0.5

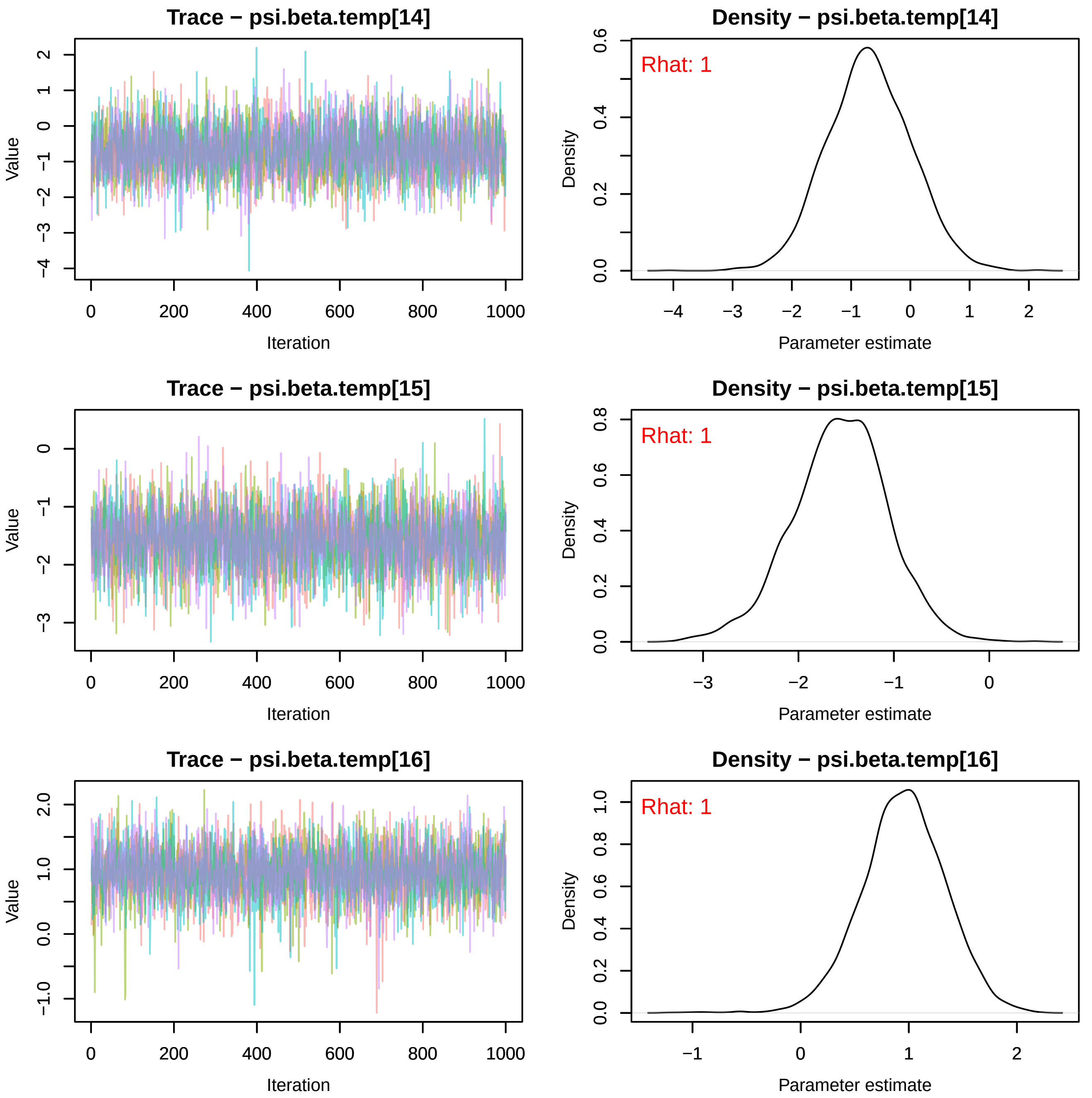
Trace − psi.beta.temp[14]
Density − psi.beta.temp[14]
Rhat: 1
Trace − psi.beta.temp[15]
Density − psi.beta.temp[15]
Rhat: 1
Trace − psi.beta.temp[16]
Density − psi.beta.temp[16]
Rhat: 1
Value
Iteration
Density
Parameter estimate

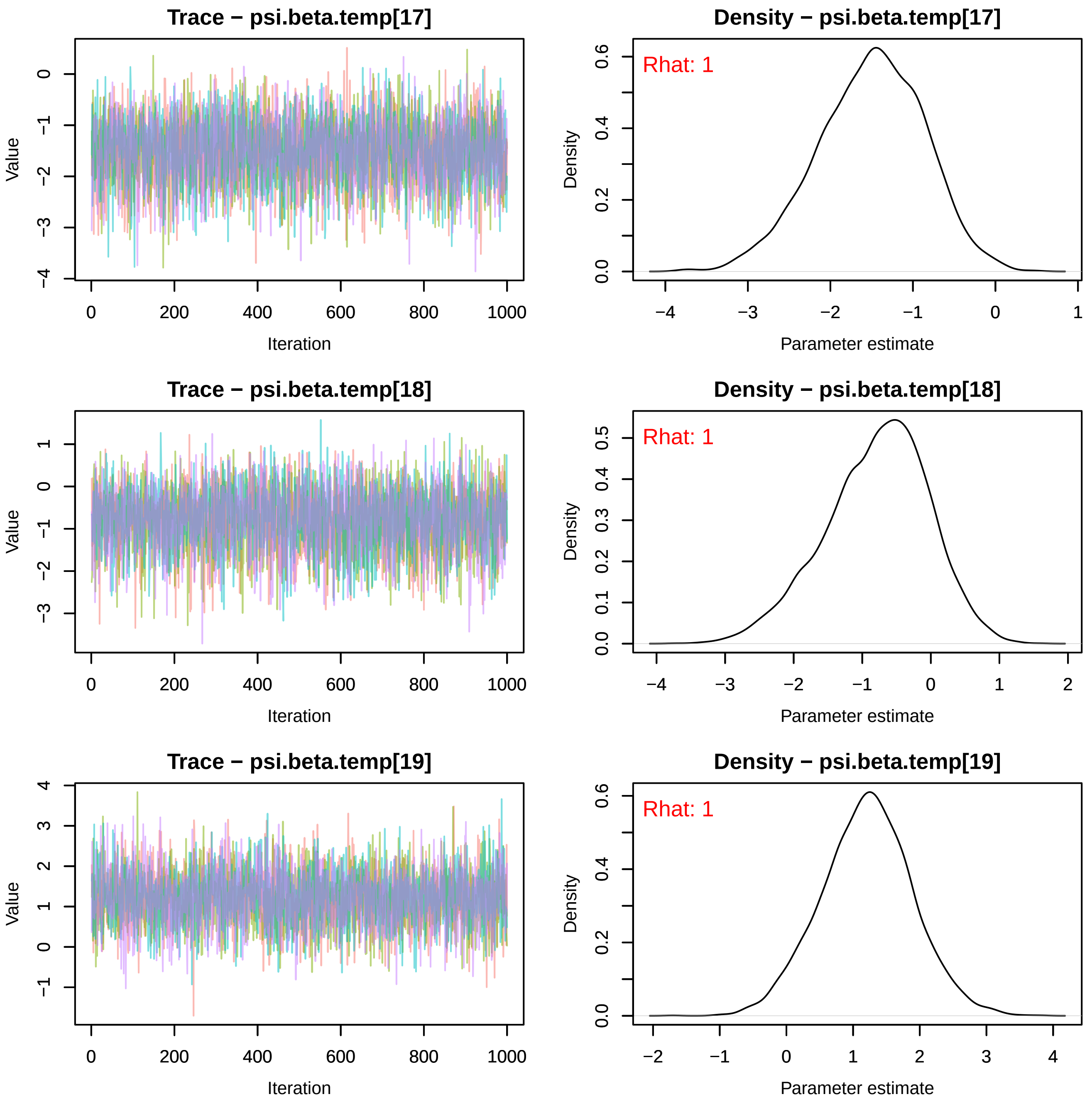
Trace − psi.beta.temp[17]
Value
Iteration
Density − psi.beta.temp[17]
Rhat: 1
Density
Parameter estimate
Trace − psi.beta.temp[18]
Value
Iteration
Density − psi.beta.temp[18]
Rhat: 1
Density
Parameter estimate
Trace − psi.beta.temp[19]
Value
Iteration
Density − psi.beta.temp[19]
Rhat: 1
Density
Parameter estimate

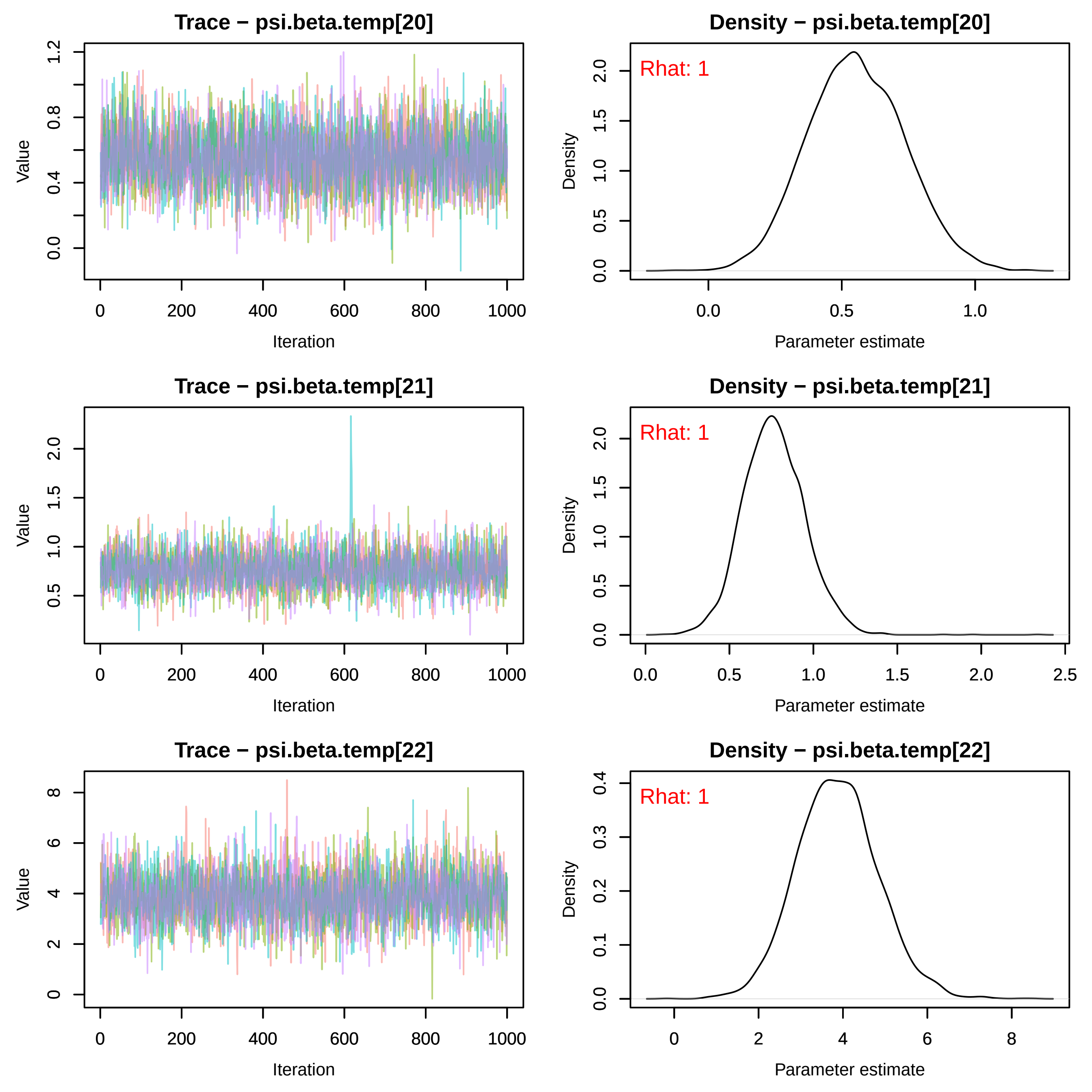
Trace − psi.beta.temp[20]
Density − psi.beta.temp[20]
Rhat: 1
Trace − psi.beta.temp[21]
Density − psi.beta.temp[21]
Rhat: 1
Trace − psi.beta.temp[22]
Density − psi.beta.temp[22]
Rhat: 1
Value
Iteration
Density
Parameter estimate

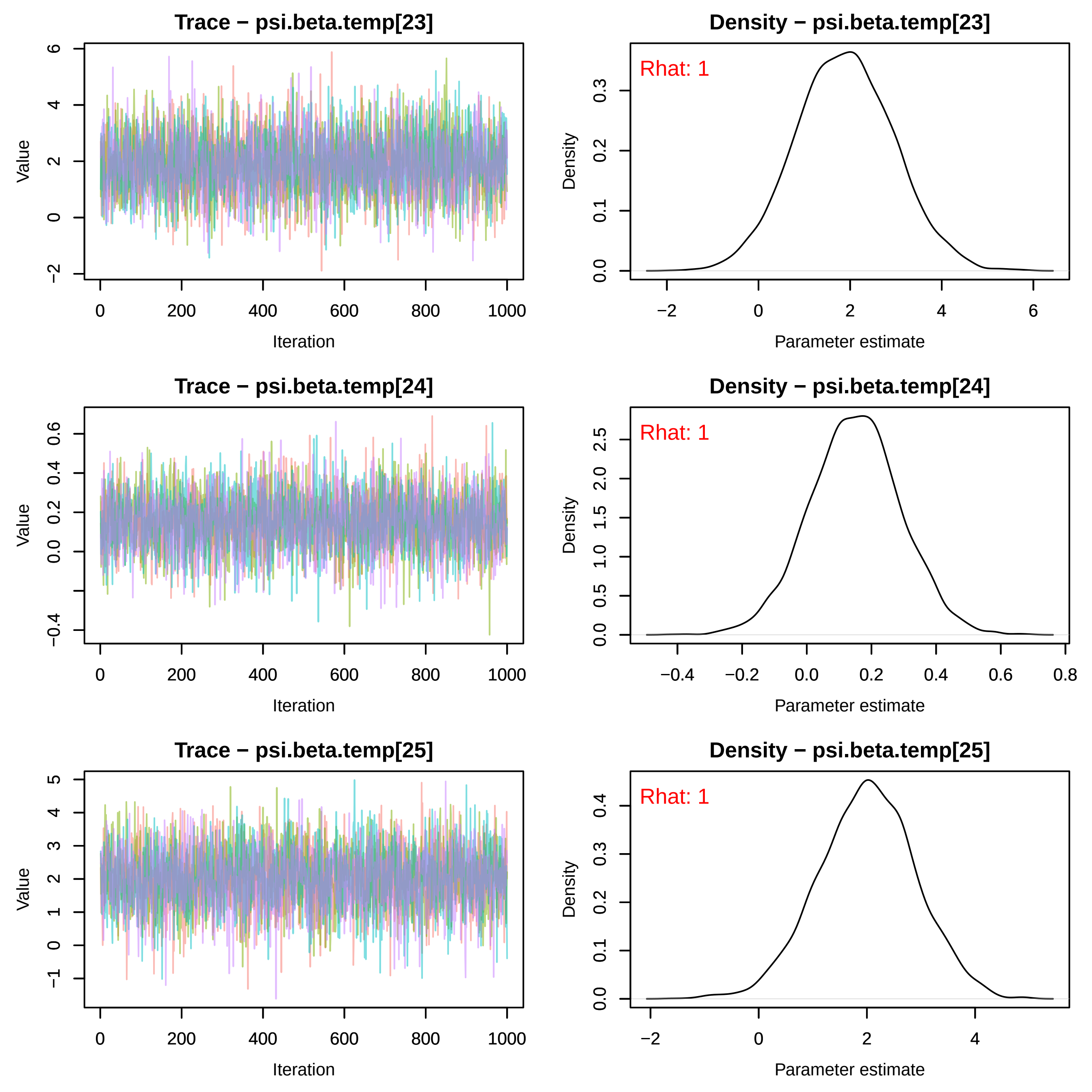
Trace − psi.beta.temp[23]
Value
Iteration
Density − psi.beta.temp[23]
Rhat: 1
Density
Parameter estimate
Trace − psi.beta.temp[24]
Value
Iteration
Density − psi.beta.temp[24]
Rhat: 1
Density
Parameter estimate
Trace − psi.beta.temp[25]
Value
Iteration
Density − psi.beta.temp[25]
Rhat: 1
Density
Parameter estimate
0
200
400
600
800
1000
−2
0
2
4
6
0.0
0.1
0.2
0.3
−0.4
0.0
0.2
0.4
0.6
−0.2
0.8
0.5
1.0
1.5
2.0
2.5
−1
1
3
5

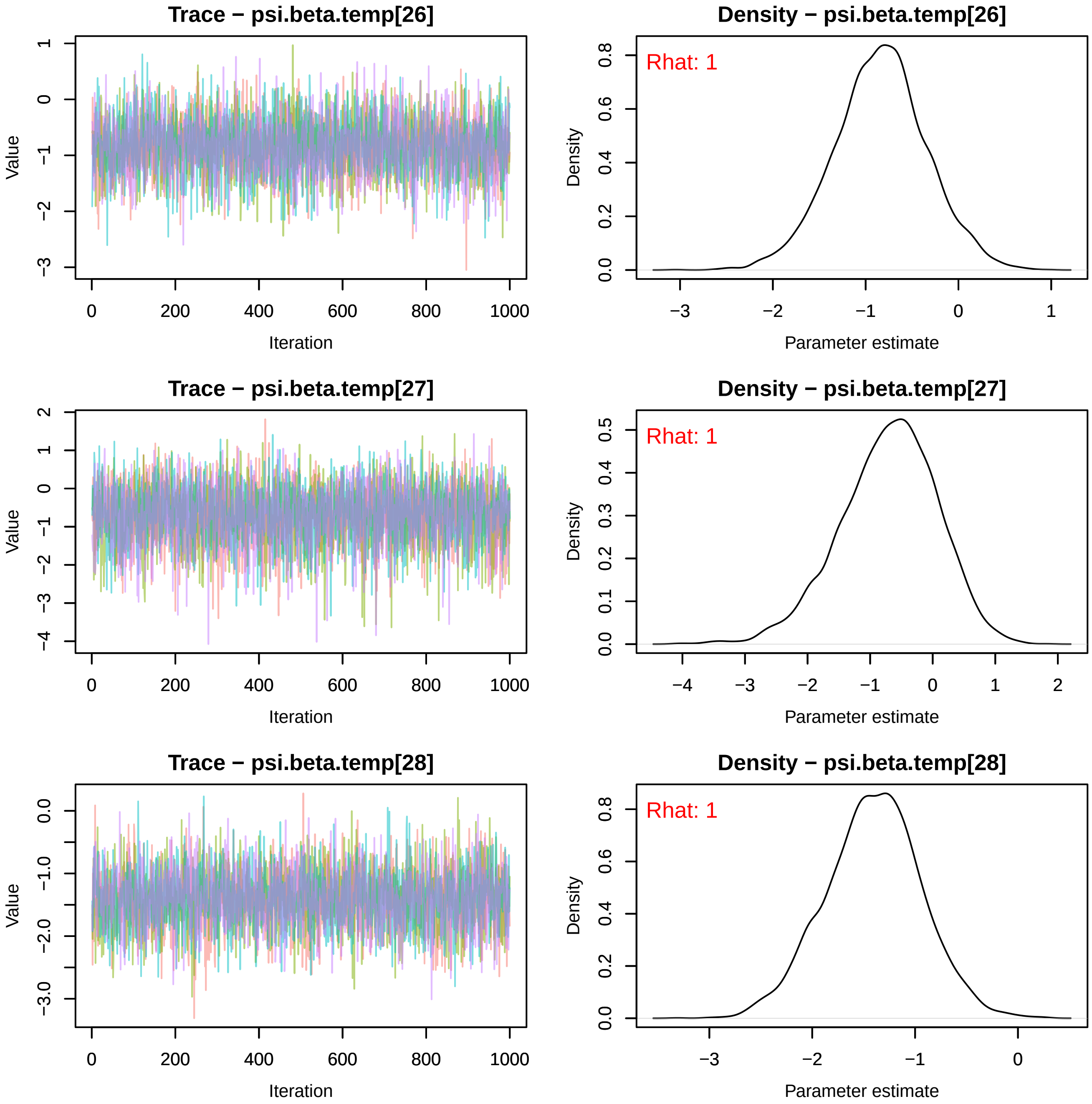
Trace − psi.beta.temp[26]
Value
Iteration
Density − psi.beta.temp[26]
Rhat: 1
Density
Parameter estimate
Trace − psi.beta.temp[27]
Value
Iteration
Density − psi.beta.temp[27]
Rhat: 1
Density
Parameter estimate
Trace − psi.beta.temp[28]
Value
Iteration
Density − psi.beta.temp[28]
Rhat: 1
Density
Parameter estimate

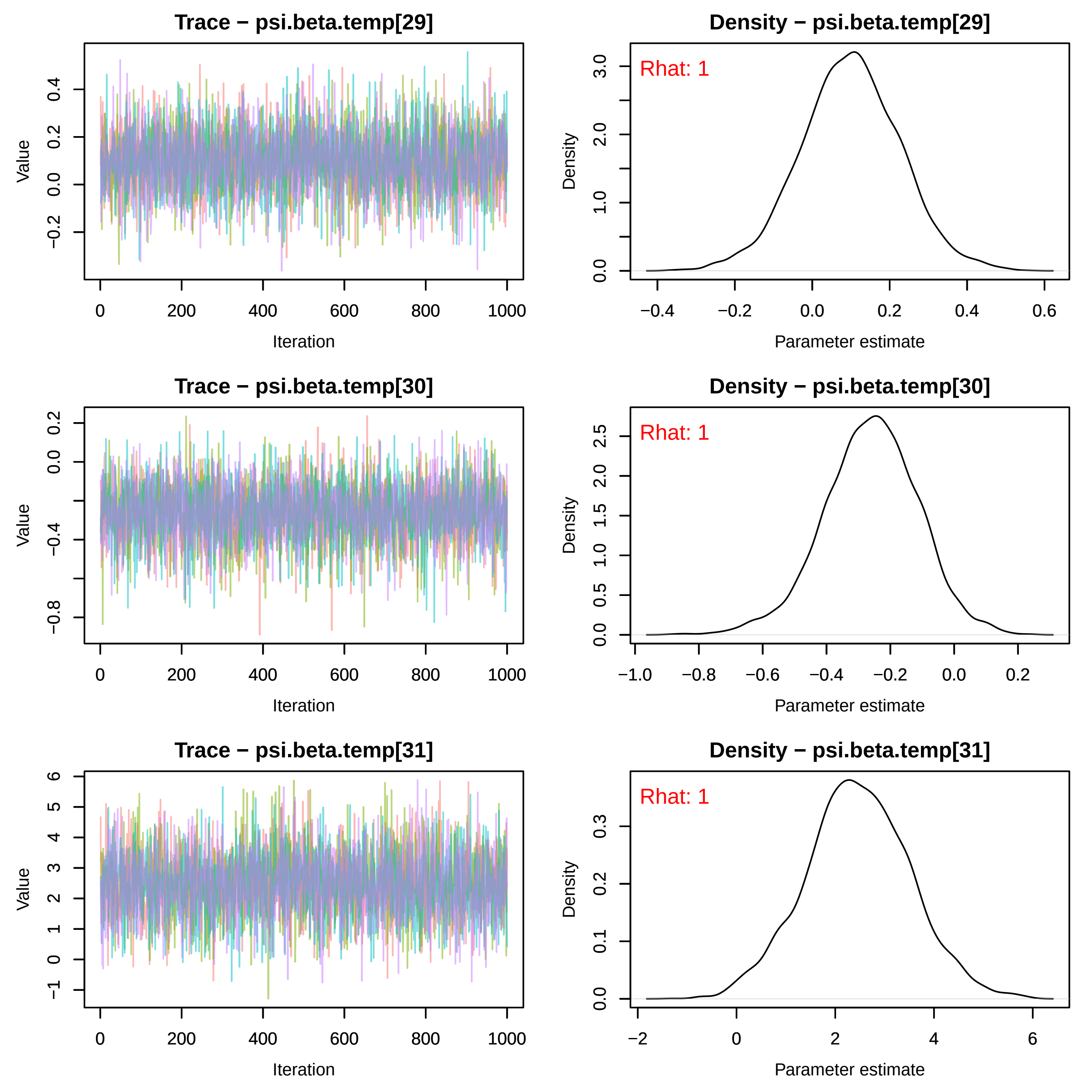
Trace − psi.beta.temp[29]
Value
Iteration
Density − psi.beta.temp[29]
Rhat: 1
Density
Parameter estimate
Trace − psi.beta.temp[30]
Value
Iteration
Density − psi.beta.temp[30]
Rhat: 1
Density
Parameter estimate
Trace − psi.beta.temp[31]
Value
Iteration
Density − psi.beta.temp[31]
Rhat: 1
Density
Parameter estimate

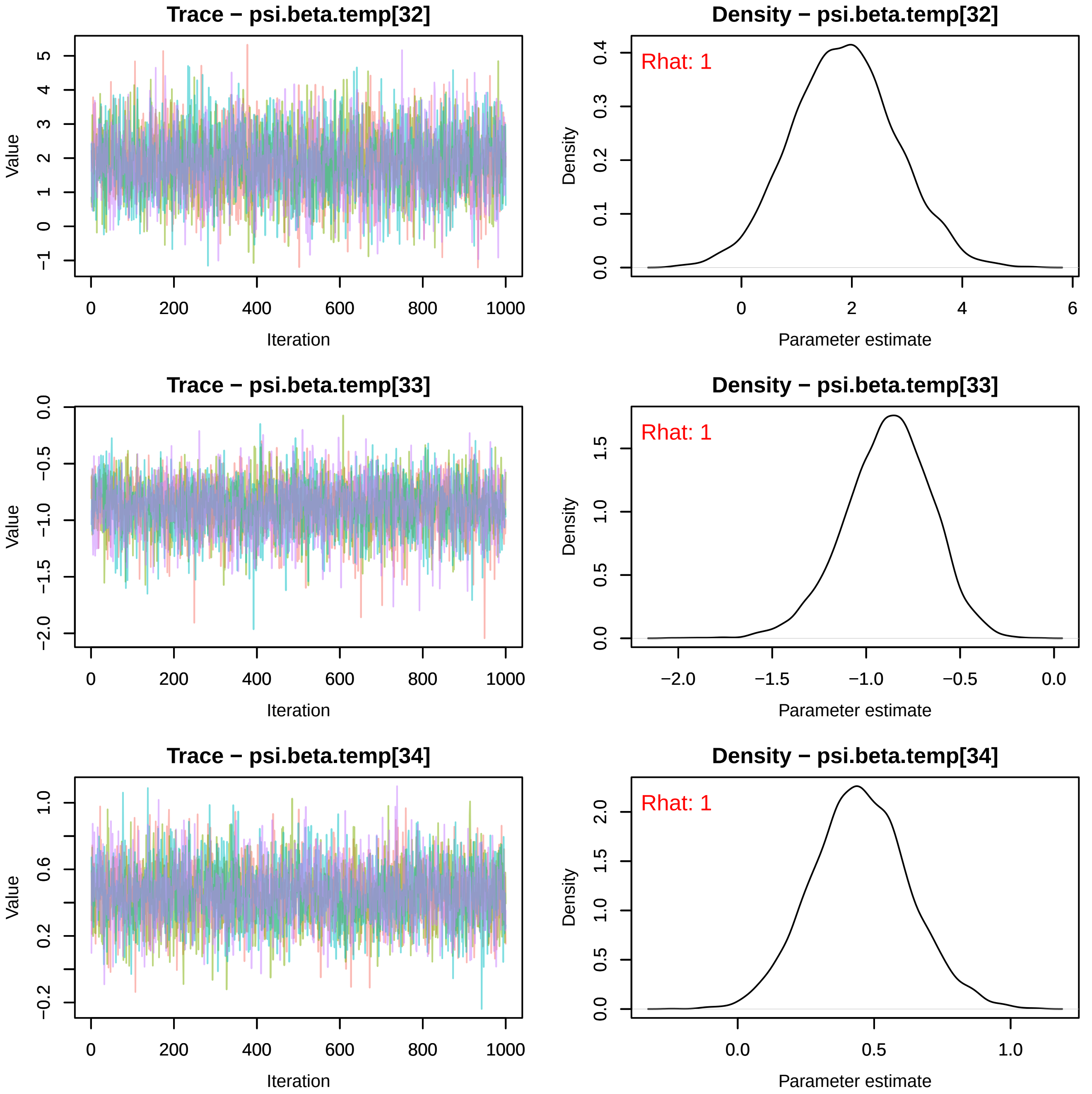
Trace − psi.beta.temp[32]
Value
−1
0
1
2
3
4
5
0
200
400
600
800
1000
Iteration
Density − psi.beta.temp[32]
Rhat: 1
Density
0.0
0.1
0.2
0.3
0.4
0
2
4
6
Parameter estimate
Trace − psi.beta.temp[33]
Value
−2.0
−1.5
−1.0
−0.5
0.0
0
200
400
600
800
1000
Iteration
Density − psi.beta.temp[33]
Rhat: 1
Density
0.0
0.5
1.0
1.5
−2.0
−1.5
−1.0
−0.5
0.0
Parameter estimate
Trace − psi.beta.temp[34]
Value
−0.2
0.2
0.6
1.0
0
200
400
600
800
1000
Iteration
Density − psi.beta.temp[34]
Rhat: 1
Density
0.0
0.5
1.0
1.5
2.0
0.0
0.5
1.0
Parameter estimate

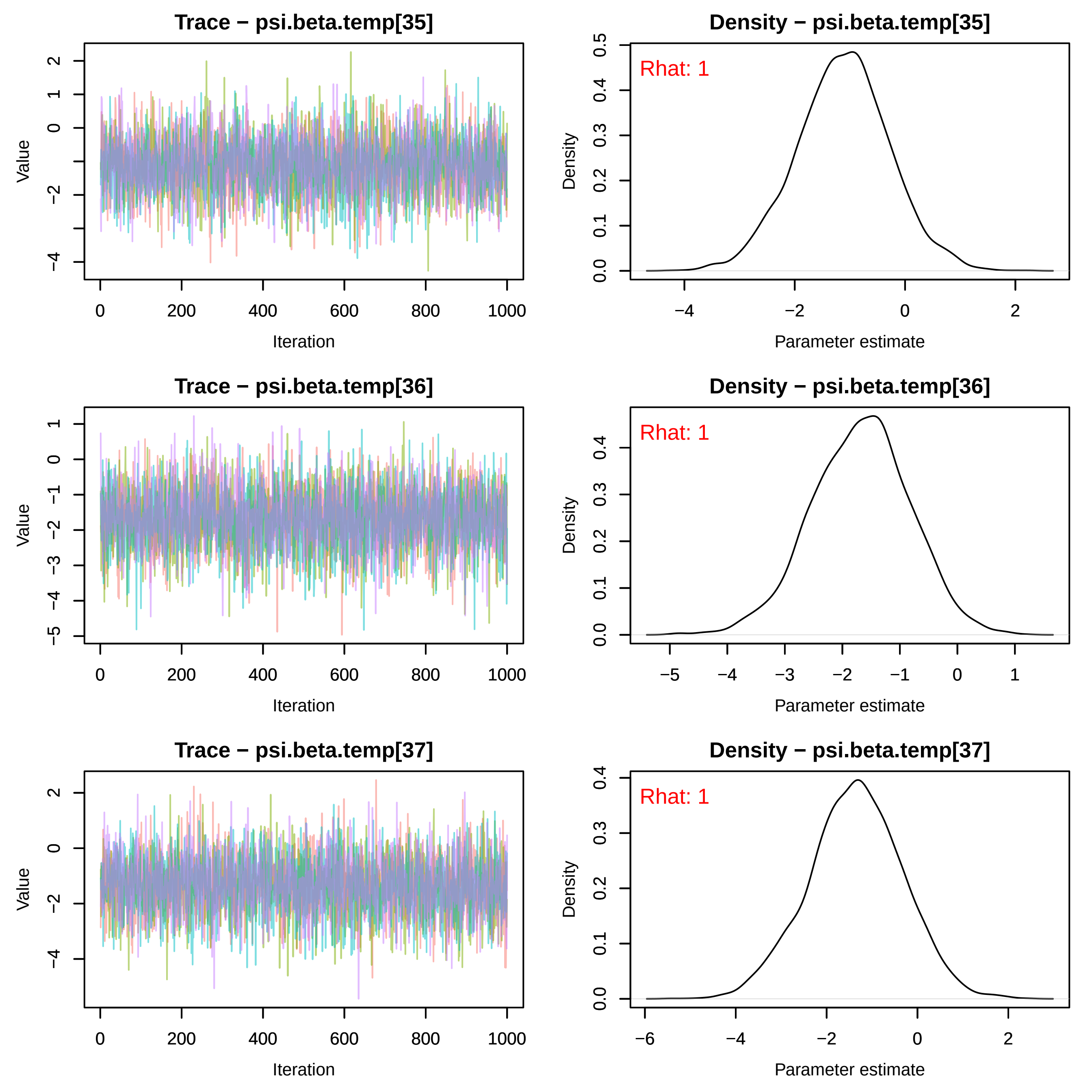

Trace − psi.beta.temp[35]
Density − psi.beta.temp[35]
Rhat: 1
Trace − psi.beta.temp[36]
Density − psi.beta.temp[36]
Rhat: 1
Trace − psi.beta.temp[37]
Density − psi.beta.temp[37]
Rhat: 1
Value
Iteration
Density
Parameter estimate

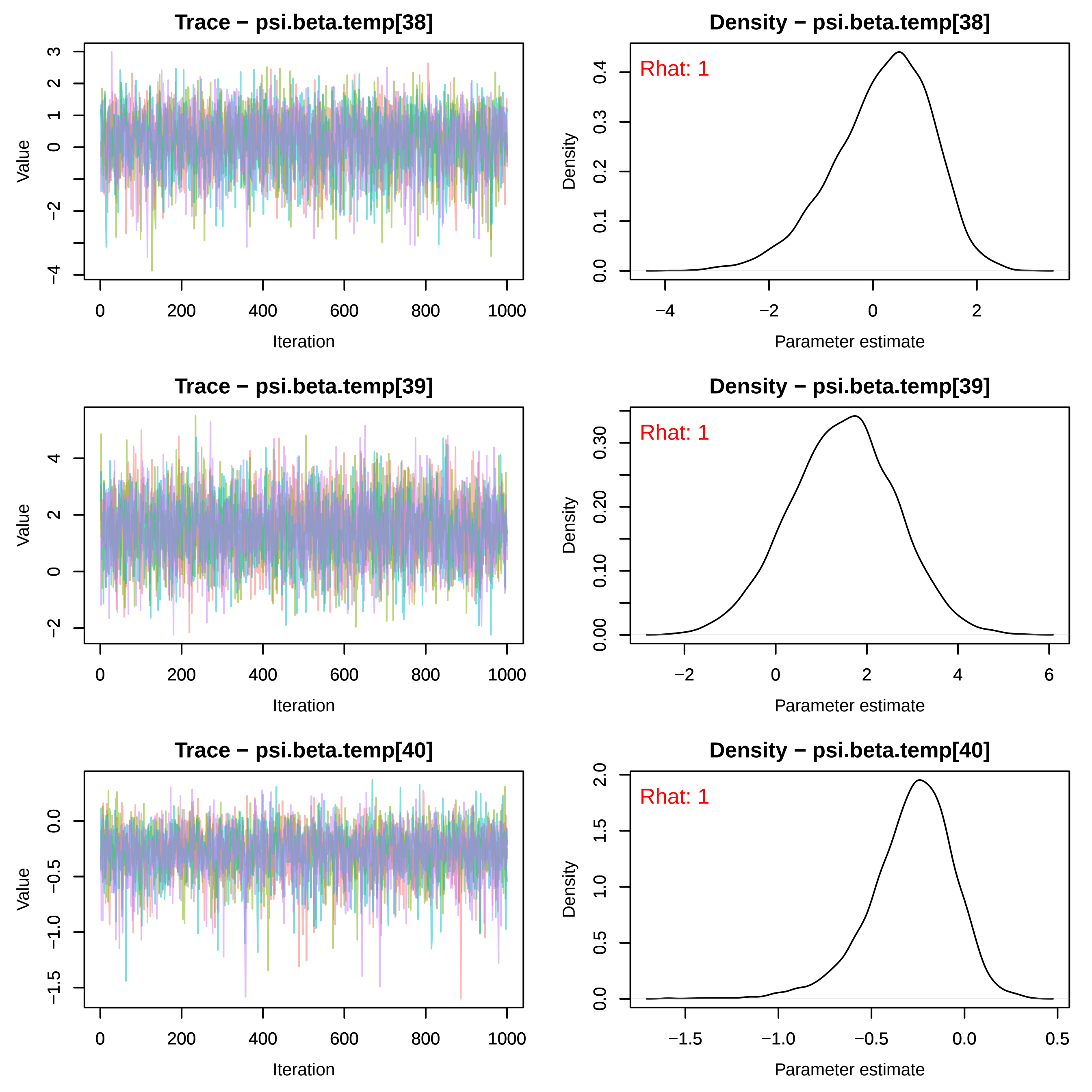
Trace − psi.beta.temp[38]
Value
Iteration
Density − psi.beta.temp[38]
Rhat: 1
Density
Parameter estimate
Trace − psi.beta.temp[39]
Value
Iteration
Density − psi.beta.temp[39]
Rhat: 1
Density
Parameter estimate
Trace − psi.beta.temp[40]
Value
Iteration
Density − psi.beta.temp[40]
Rhat: 1
Density
Parameter estimate

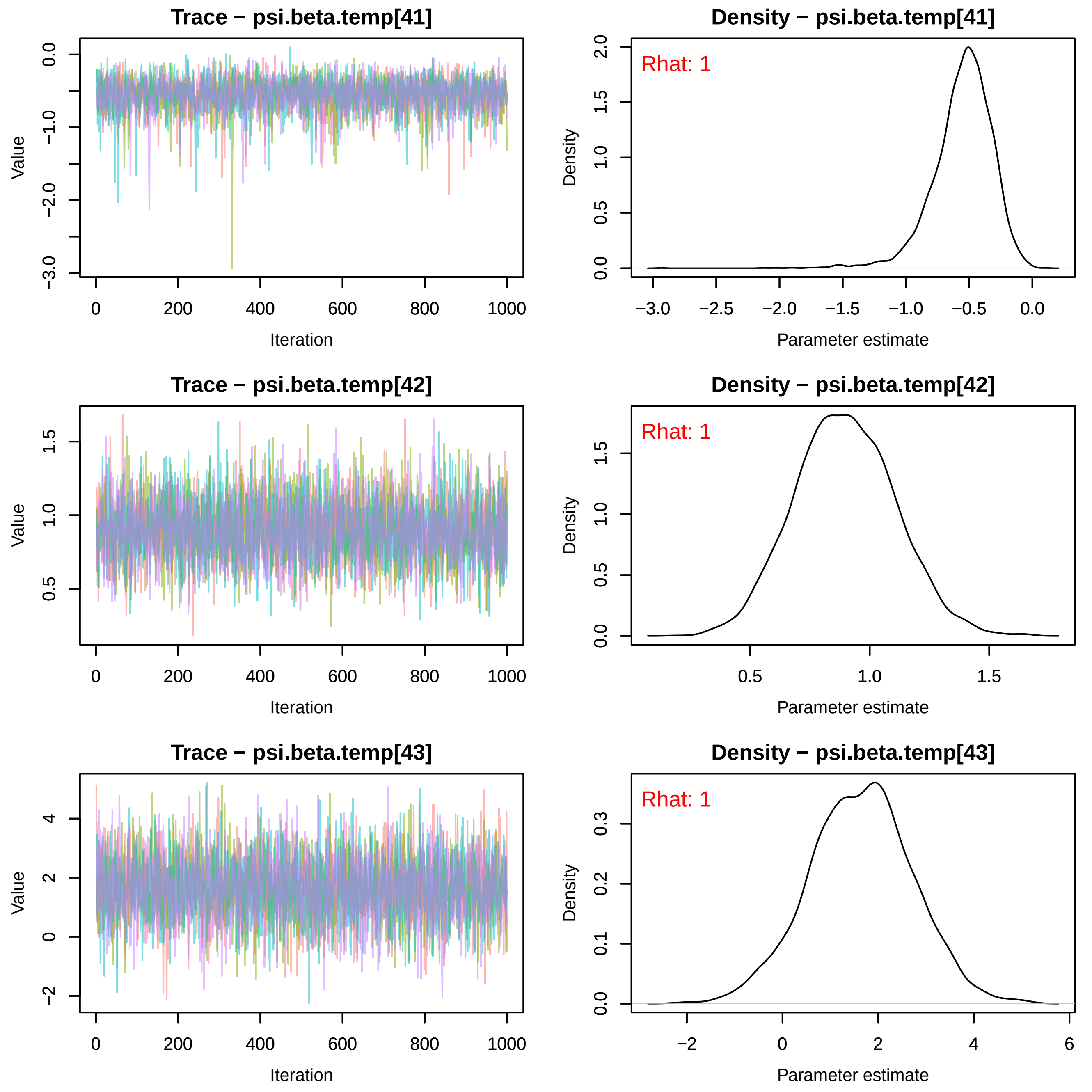
Trace − psi.beta.temp[41]
Value
Iteration
0.0
−1.0
−2.0
−3.0
0
200
400
600
800
1000
Density − psi.beta.temp[41]
Rhat: 1
Density
Parameter estimate
2.0
1.5
1.0
0.5
0.0
−3.0
−2.5
−2.0
−1.5
−1.0
−0.5
0.0
Trace − psi.beta.temp[42]
Value
Iteration
1.5
1.0
0.5
0
200
400
600
800
1000
Density − psi.beta.temp[42]
Rhat: 1
Density
Parameter estimate
1.5
1.0
0.5
0.0
0.5
1.0
1.5
Trace − psi.beta.temp[43]
Value
Iteration
4
2
0
−2
0
200
400
600
800
1000
Density − psi.beta.temp[43]
Rhat: 1
Density
Parameter estimate
0.3
0.2
0.1
0.0
−2
0
2
4
6

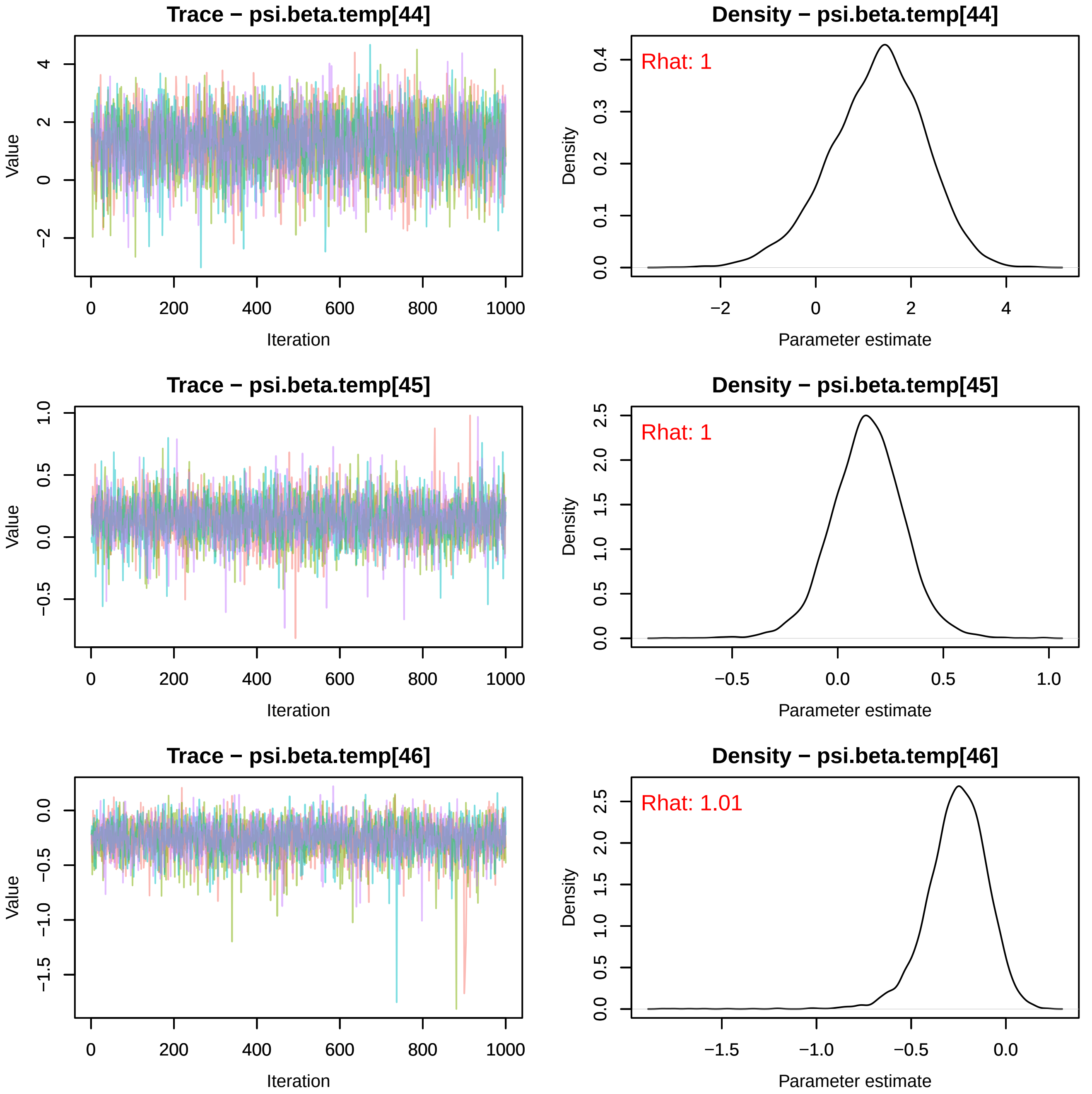
Trace − psi.beta.temp[44]
Value
Iteration
Density − psi.beta.temp[44]
Rhat: 1
Density
Parameter estimate
Trace − psi.beta.temp[45]
Value
Iteration
Density − psi.beta.temp[45]
Rhat: 1
Density
Parameter estimate
Trace − psi.beta.temp[46]
Value
Iteration
Density − psi.beta.temp[46]
Rhat: 1.01
Density
Parameter estimate

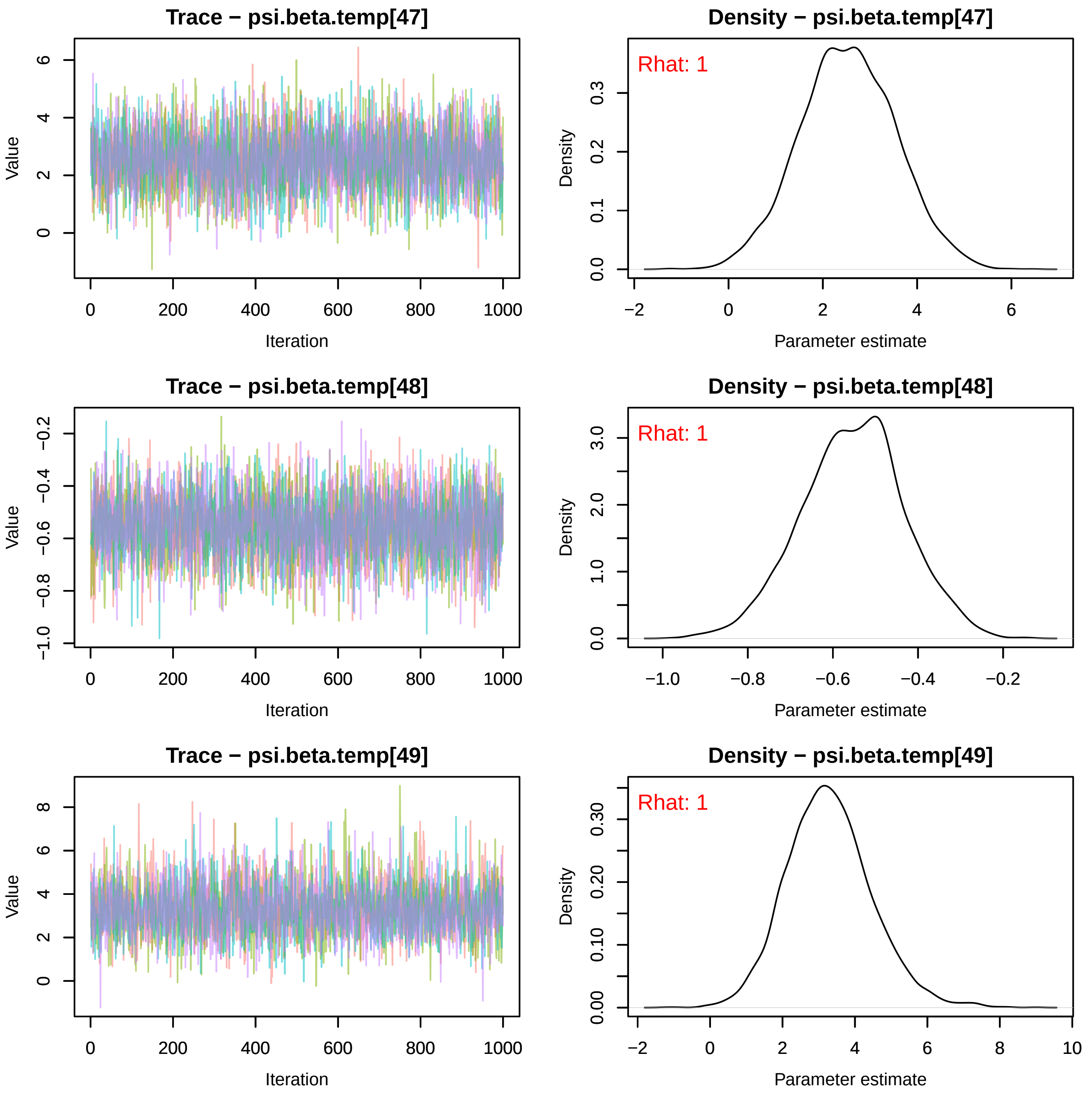
Trace − psi.beta.temp[47]
Density − psi.beta.temp[47]
Rhat: 1
Trace − psi.beta.temp[48]
Density − psi.beta.temp[48]
Rhat: 1
Trace − psi.beta.temp[49]
Density − psi.beta.temp[49]
Rhat: 1
Value
Iteration
Density
Parameter estimate

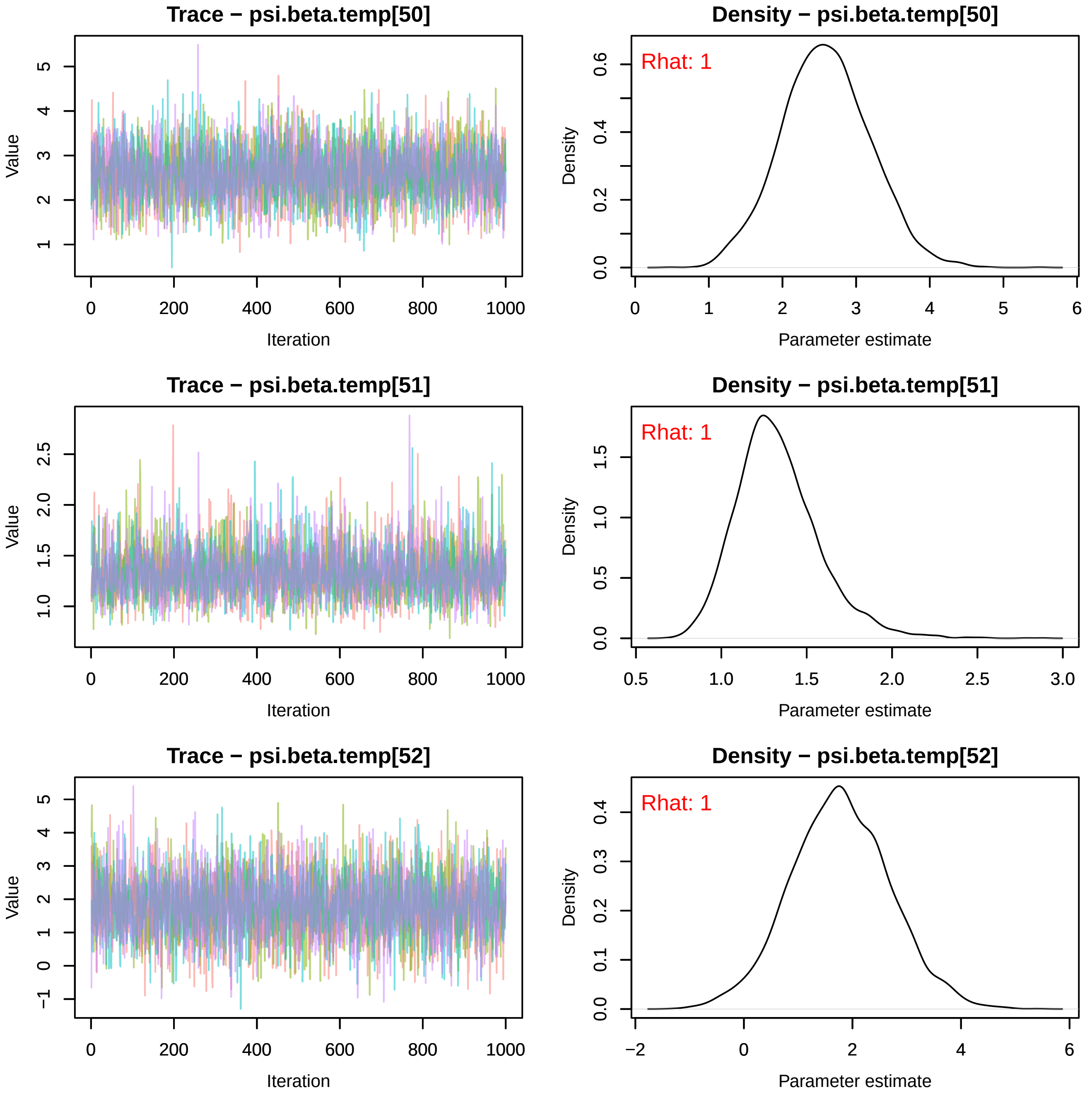

Trace − psi.beta.temp[50]
Density − psi.beta.temp[50]
Rhat: 1
Trace − psi.beta.temp[51]
Density − psi.beta.temp[51]
Rhat: 1
Trace − psi.beta.temp[52]
Density − psi.beta.temp[52]
Rhat: 1
Value
Iteration
Density
Parameter estimate

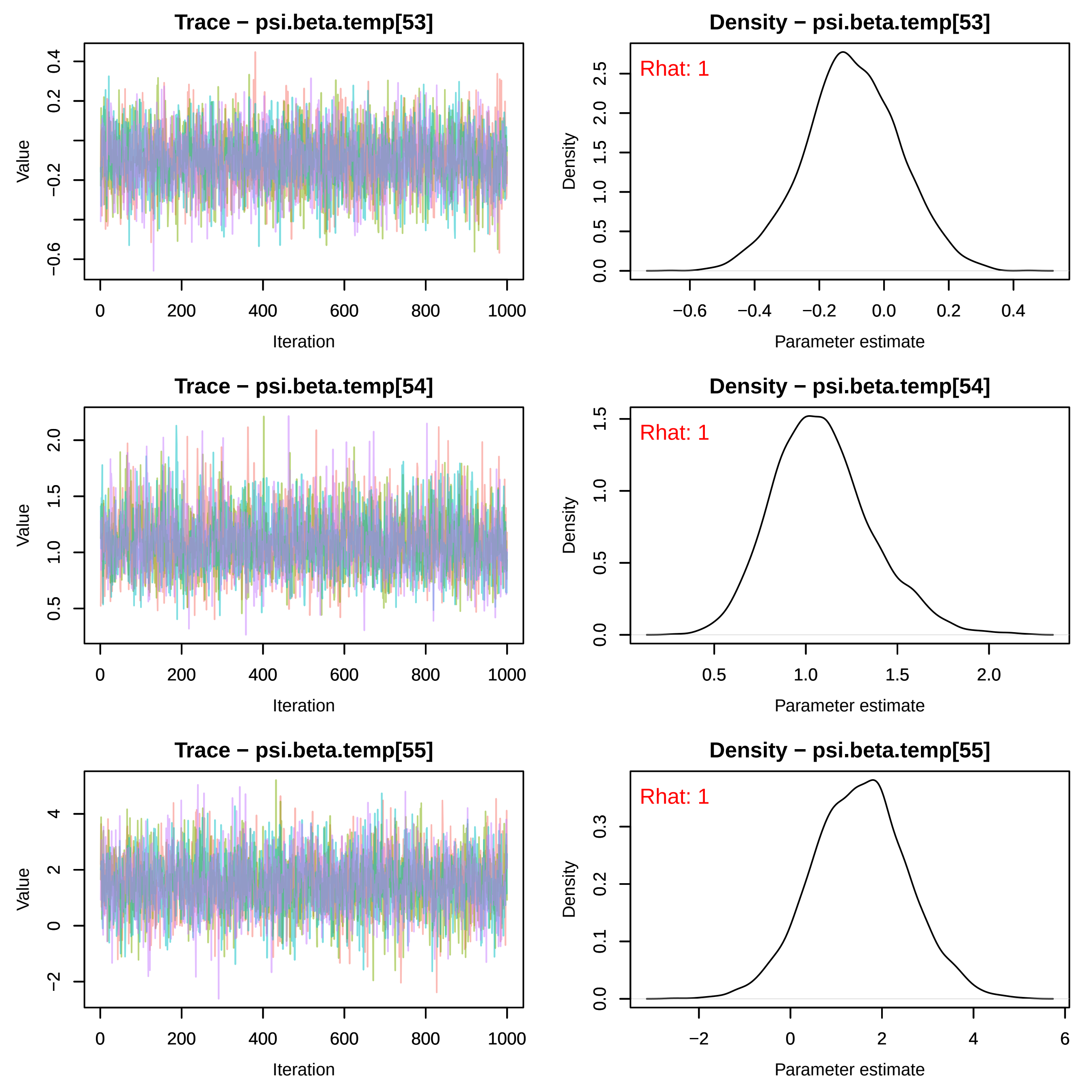
Trace − psi.beta.temp[53]
Value
Iteration
Density − psi.beta.temp[53]
Rhat: 1
Density
Parameter estimate
Trace − psi.beta.temp[54]
Value
Iteration
Density − psi.beta.temp[54]
Rhat: 1
Density
Parameter estimate
Trace − psi.beta.temp[55]
Value
Iteration
Density − psi.beta.temp[55]
Rhat: 1
Density
Parameter estimate

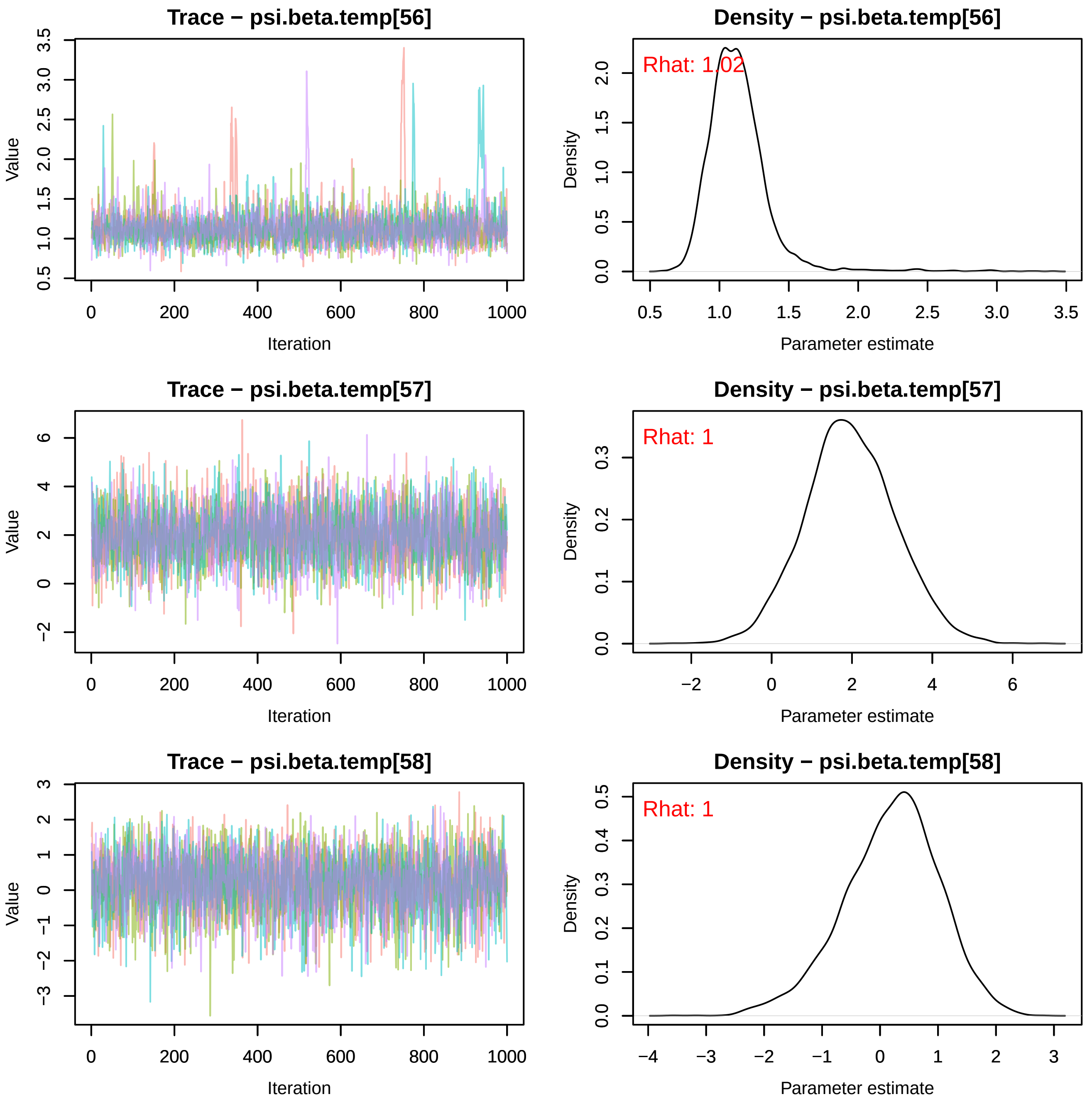
Trace − psi.beta.temp[56]
Density − psi.beta.temp[56]
Rhat: 1.02
Trace − psi.beta.temp[57]
Density − psi.beta.temp[57]
Rhat: 1
Trace − psi.beta.temp[58]
Density − psi.beta.temp[58]
Rhat: 1
Value
Iteration
Density
Parameter estimate

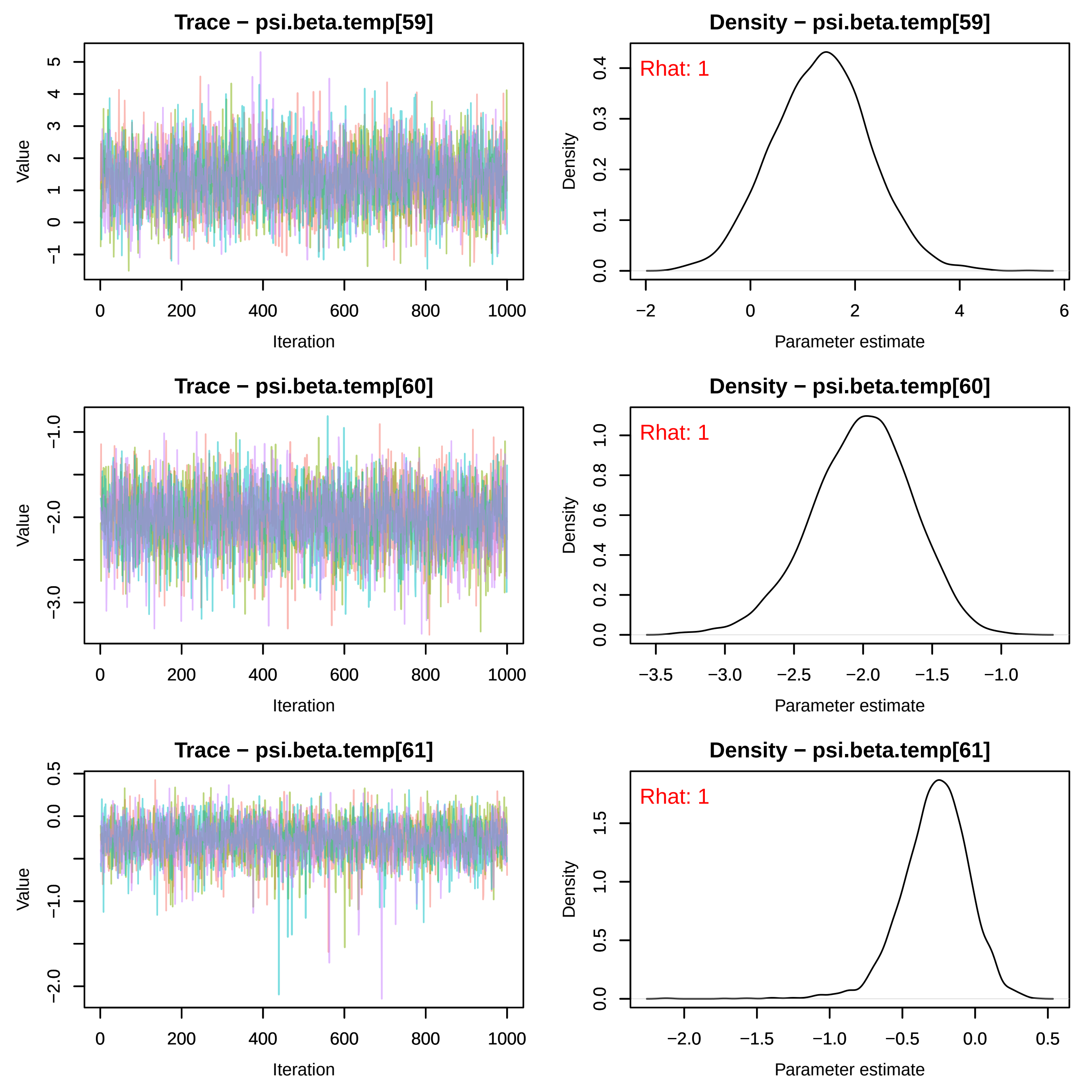
Trace − psi.beta.temp[59]
Density − psi.beta.temp[59]
Rhat: 1
Trace − psi.beta.temp[60]
Density − psi.beta.temp[60]
Rhat: 1
Trace − psi.beta.temp[61]
Density − psi.beta.temp[61]
Rhat: 1
Value
Density
Iteration
Parameter estimate

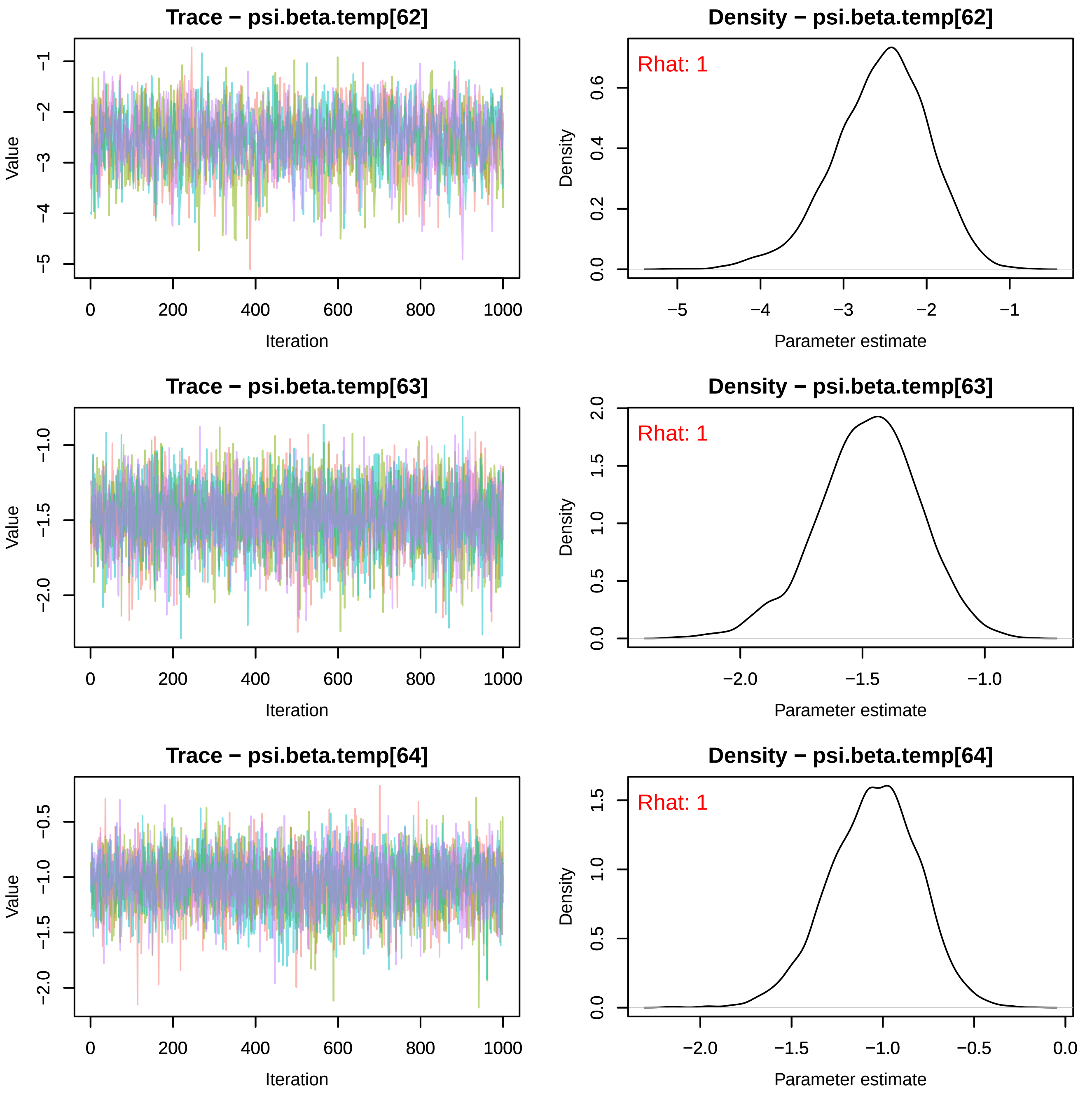
Trace − psi.beta.temp[62]
Value
Iteration
Density − psi.beta.temp[62]
Rhat: 1
Density
Parameter estimate
Trace − psi.beta.temp[63]
Value
Iteration
Density − psi.beta.temp[63]
Rhat: 1
Density
Parameter estimate
Trace − psi.beta.temp[64]
Value
Iteration
Density − psi.beta.temp[64]
Rhat: 1
Density
Parameter estimate

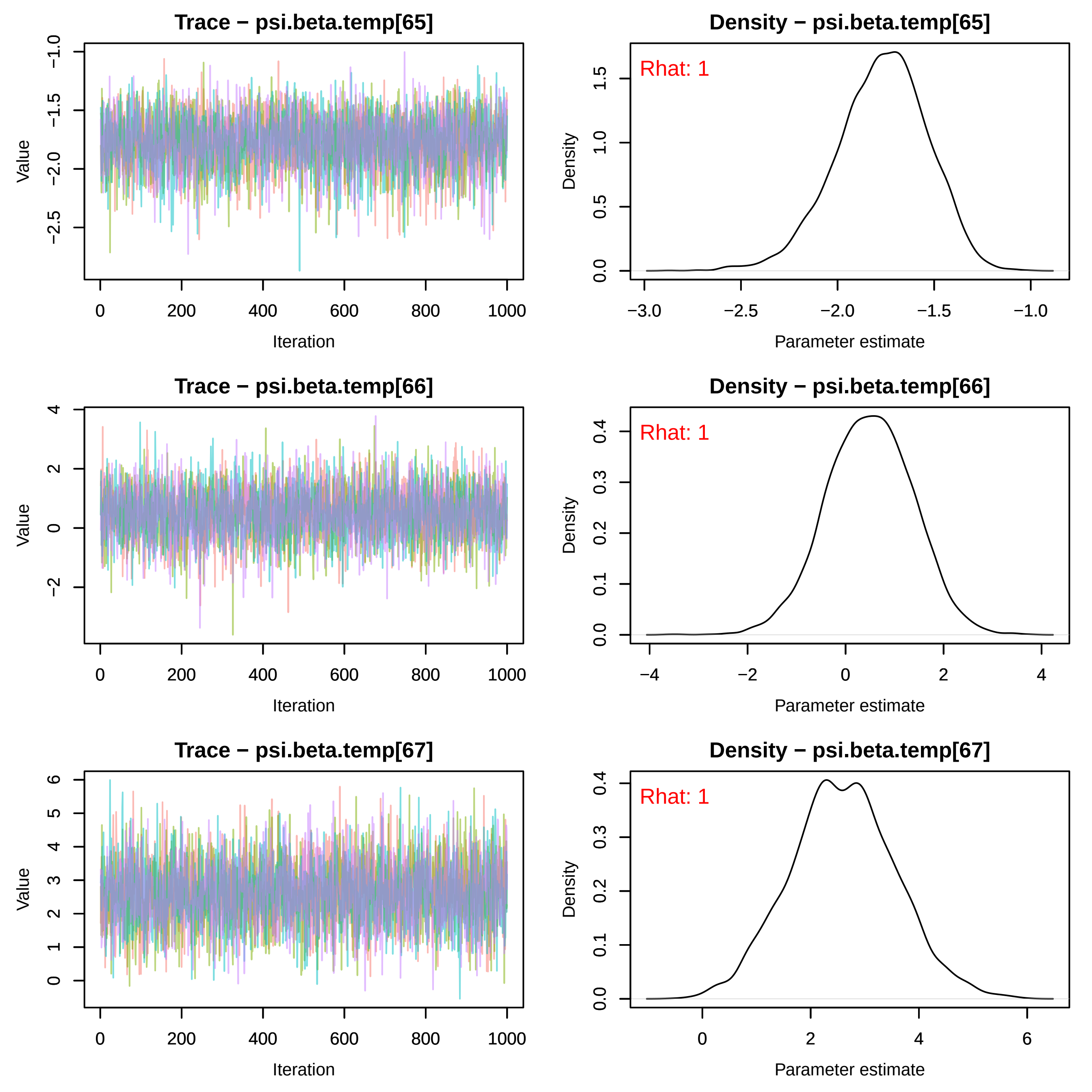
Trace − psi.beta.temp[65]
Value
Iteration
Density − psi.beta.temp[65]
Rhat: 1
Density
Parameter estimate
Trace − psi.beta.temp[66]
Value
Iteration
Density − psi.beta.temp[66]
Rhat: 1
Density
Parameter estimate
Trace − psi.beta.temp[67]
Value
Iteration
Density − psi.beta.temp[67]
Rhat: 1
Density
Parameter estimate

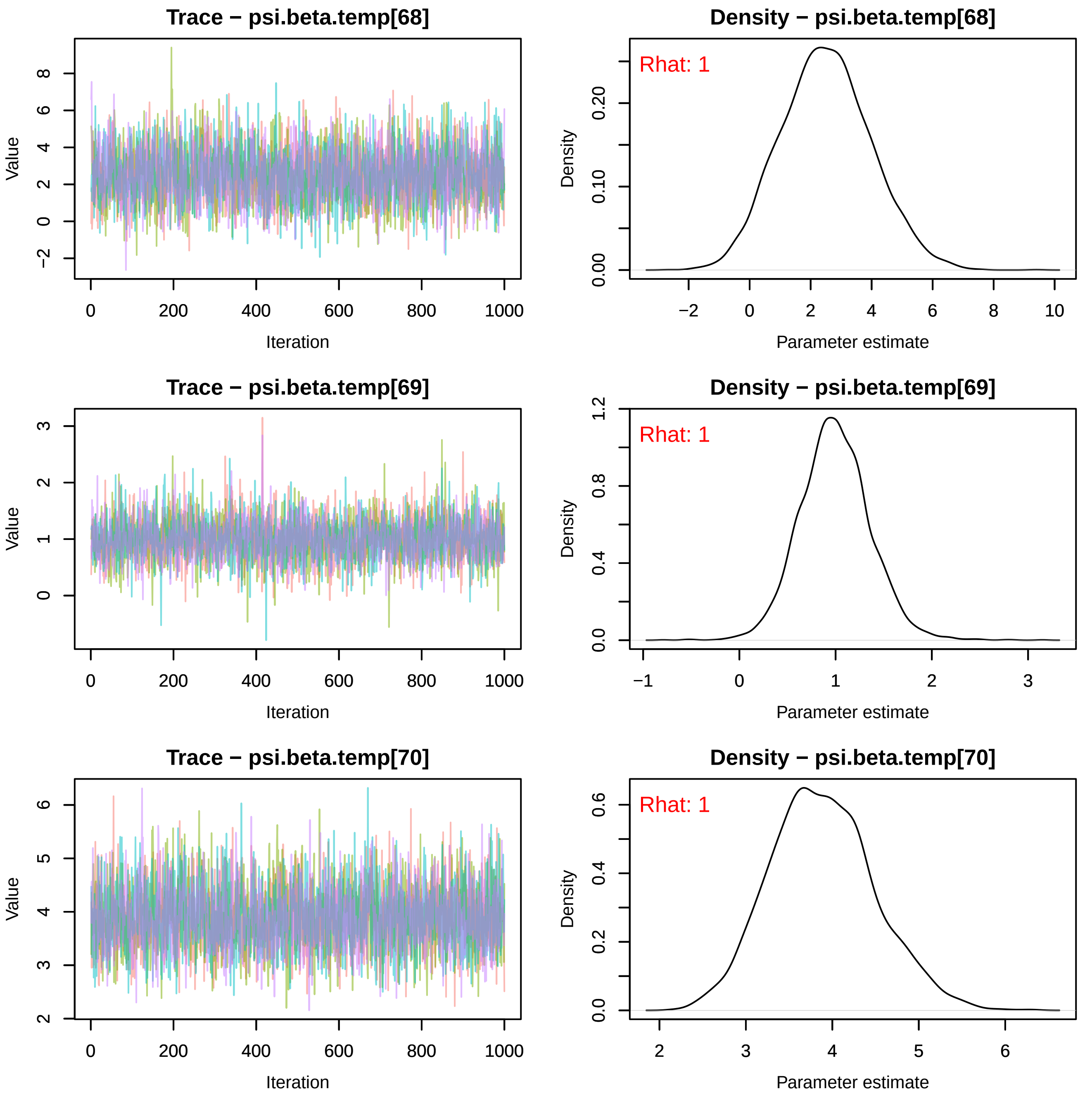
Trace − psi.beta.temp[68]
Value
Iteration
Density − psi.beta.temp[68]
Rhat: 1
Density
Parameter estimate
Trace − psi.beta.temp[69]
Value
Iteration
Density − psi.beta.temp[69]
Rhat: 1
Density
Parameter estimate
Trace − psi.beta.temp[70]
Value
Iteration
Density − psi.beta.temp[70]
Rhat: 1
Density
Parameter estimate

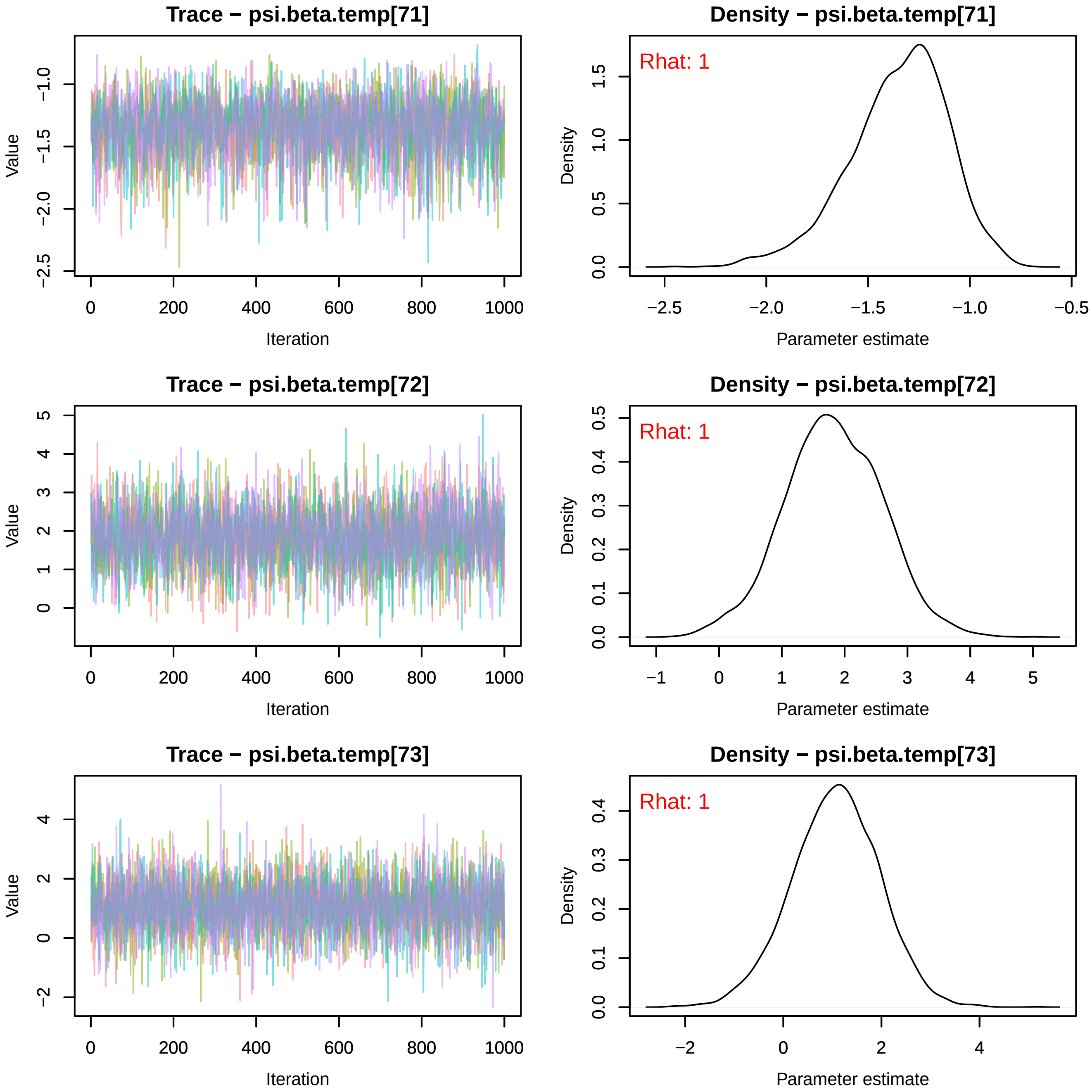

Trace − psi.beta.temp[71]
Value
Iteration
Density − psi.beta.temp[71]
Rhat: 1
Density
Parameter estimate
Trace − psi.beta.temp[72]
Value
Iteration
Density − psi.beta.temp[72]
Rhat: 1
Density
Parameter estimate
Trace − psi.beta.temp[73]
Value
Iteration
Density − psi.beta.temp[73]
Rhat: 1
Density
Parameter estimate

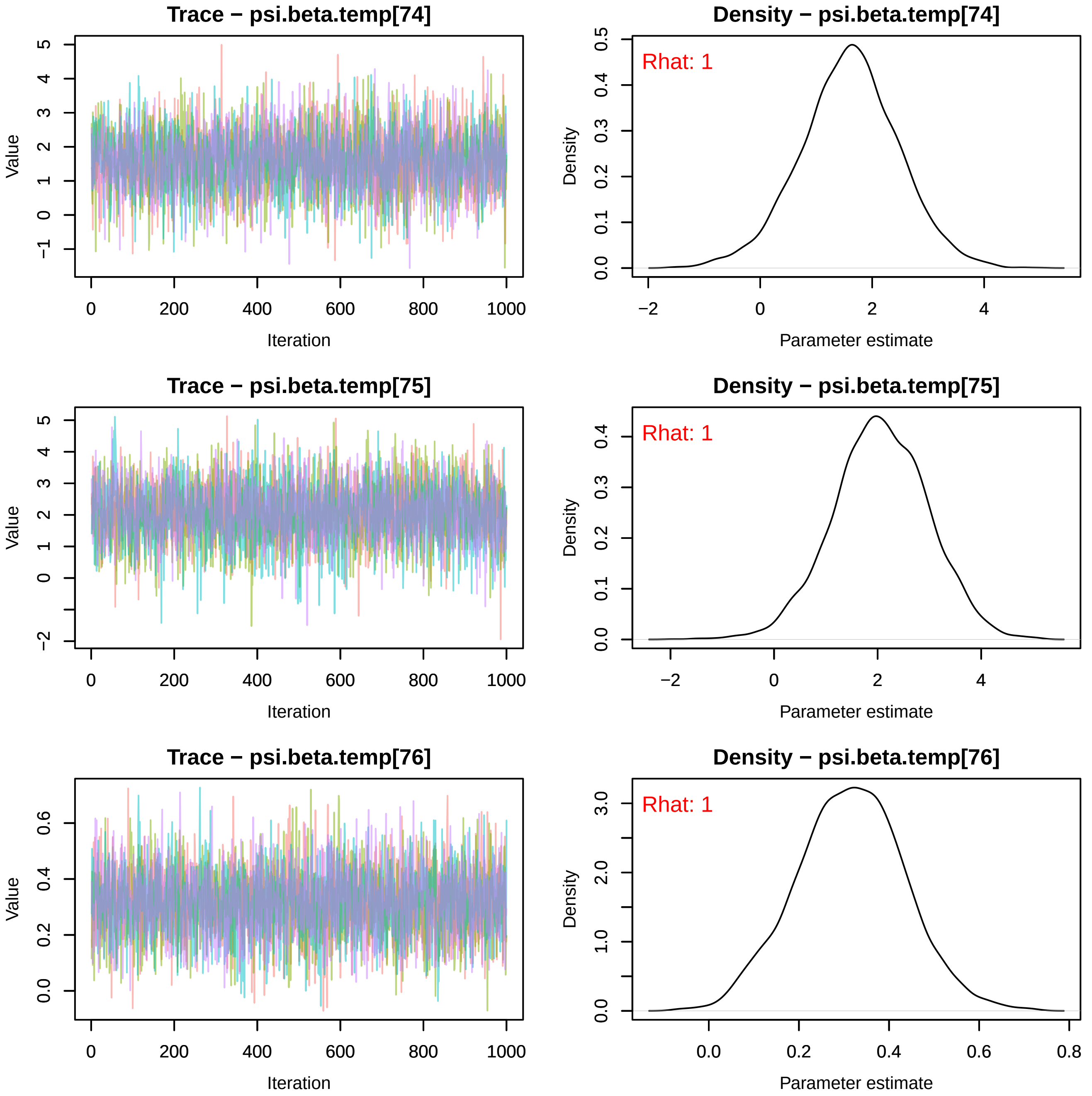
Trace − psi.beta.temp[74]
Value
Iteration
Density − psi.beta.temp[74]
Rhat: 1
Density
Parameter estimate
Trace − psi.beta.temp[75]
Value
Iteration
Density − psi.beta.temp[75]
Rhat: 1
Density
Parameter estimate
Trace − psi.beta.temp[76]
Value
Iteration
Density − psi.beta.temp[76]
Rhat: 1
Density
Parameter estimate

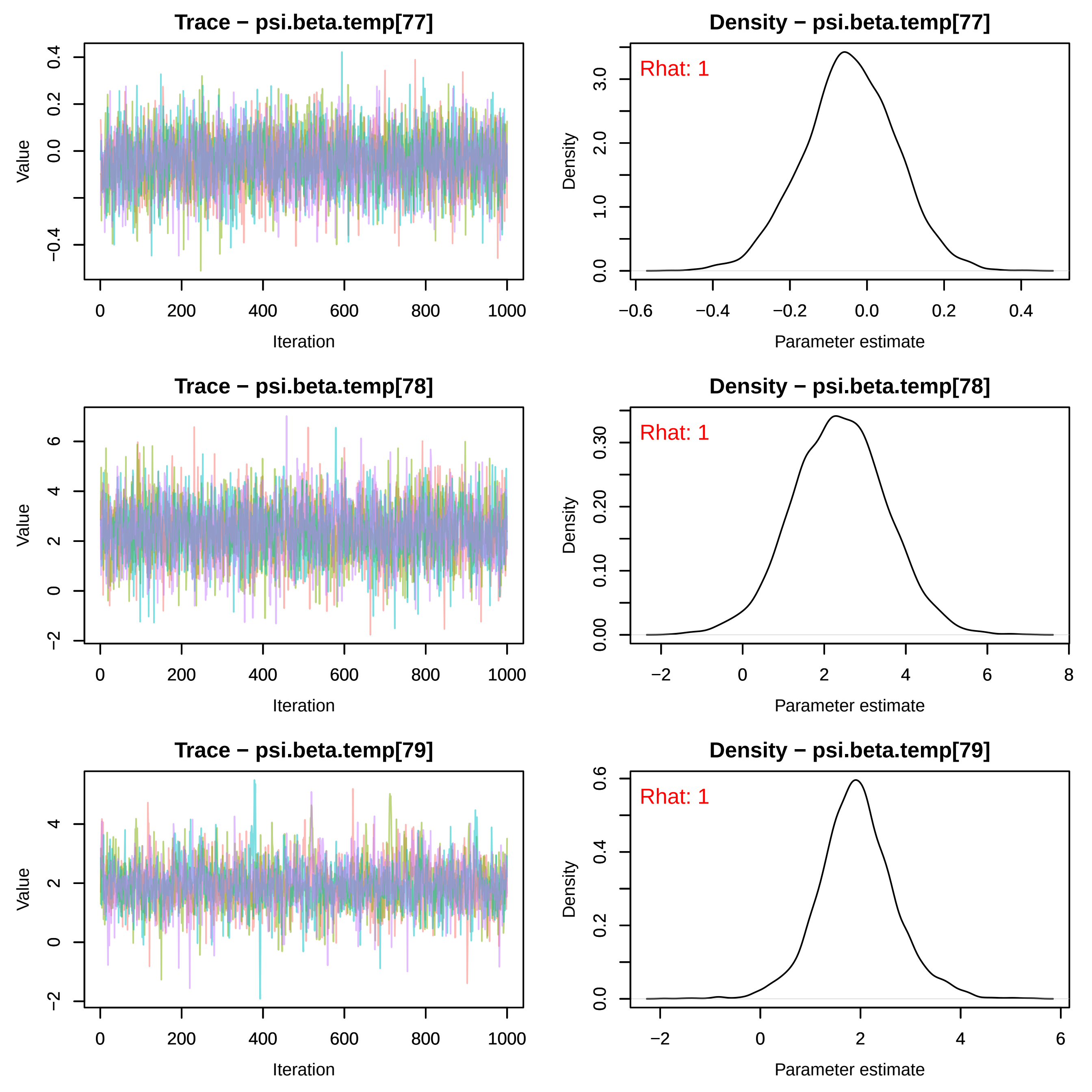
Trace − psi.beta.temp[77]
Value
Iteration
Density − psi.beta.temp[77]
Rhat: 1
Density
Parameter estimate
Trace − psi.beta.temp[78]
Value
Iteration
Density − psi.beta.temp[78]
Rhat: 1
Density
Parameter estimate
Trace − psi.beta.temp[79]
Value
Iteration
Density − psi.beta.temp[79]
Rhat: 1
Density
Parameter estimate

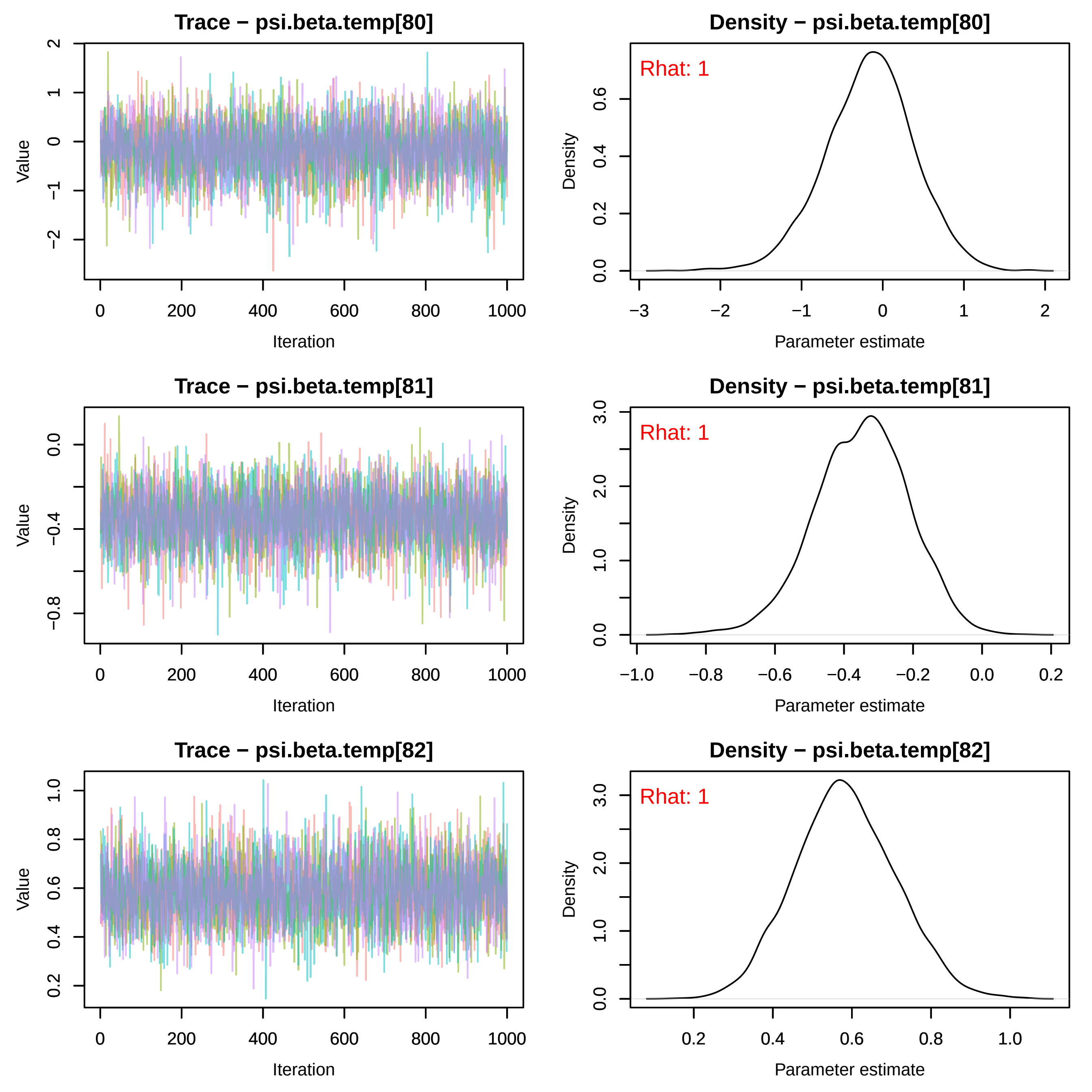
Trace − psi.beta.temp[80]
Value
Iteration
0
200
400
600
800
1000
2
1
0
−1
−2
Density − psi.beta.temp[80]
Rhat: 1
Density
Parameter estimate
0.0
0.2
0.4
0.6
−3
−2
−1
0
1
2
Trace − psi.beta.temp[81]
Value
Iteration
0.0
−0.4
−0.8
0
200
400
600
800
1000
Density − psi.beta.temp[81]
Rhat: 1
Density
Parameter estimate
0.0
1.0
2.0
3.0
−1.0
−0.8
−0.6
−0.4
−0.2
0.0
0.2
Trace − psi.beta.temp[82]
Value
Iteration
0.2
0.4
0.6
0.8
1.0
0
200
400
600
800
1000
Density − psi.beta.temp[82]
Rhat: 1
Density
Parameter estimate
0.0
1.0
2.0
3.0
0.2
0.4
0.6
0.8
1.0

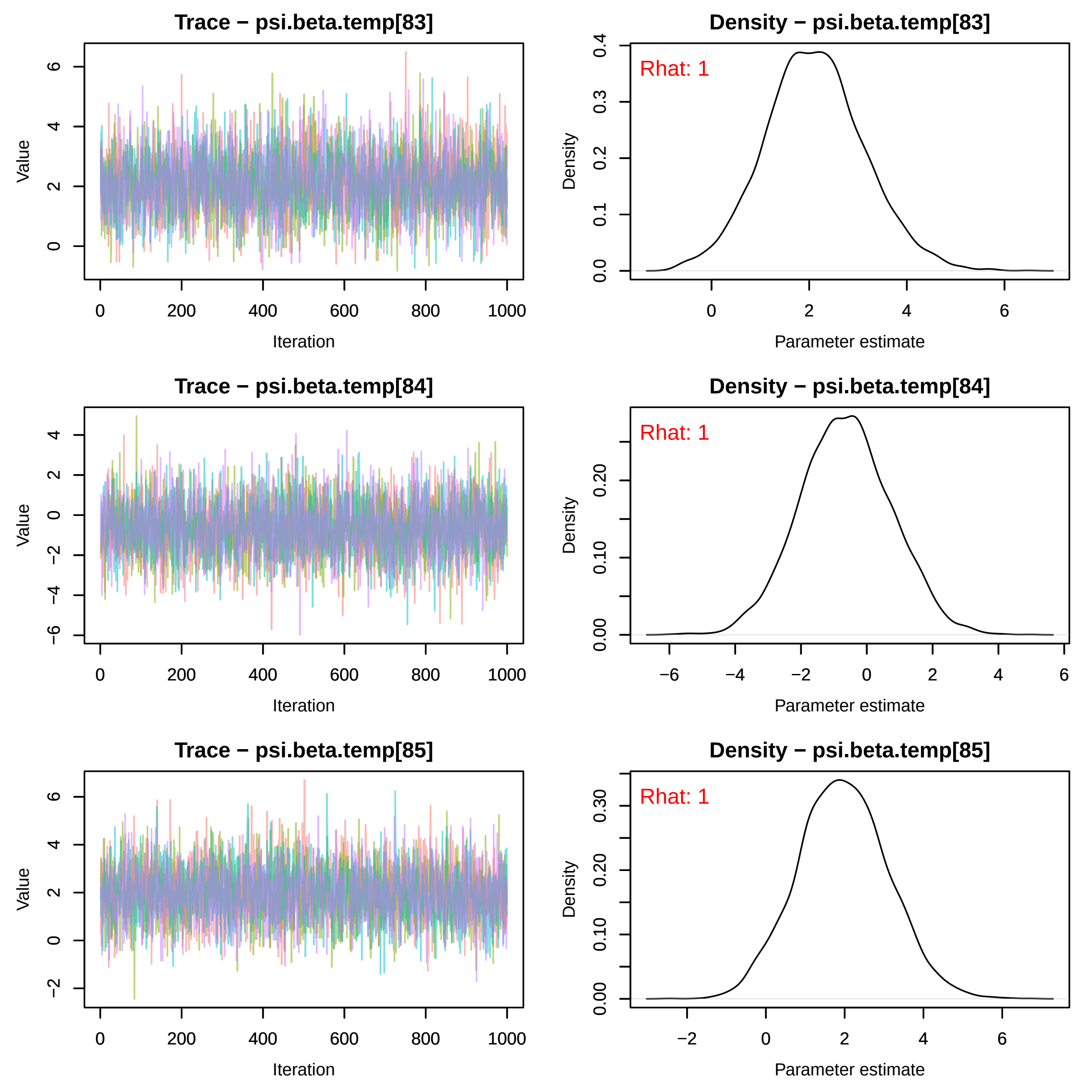

Trace − psi.beta.temp[83]
Value
Iteration
Density − psi.beta.temp[83]
Rhat: 1
Density
Parameter estimate
Trace − psi.beta.temp[84]
Value
Iteration
Density − psi.beta.temp[84]
Rhat: 1
Density
Parameter estimate
Trace − psi.beta.temp[85]
Value
Iteration
Density − psi.beta.temp[85]
Rhat: 1
Density
Parameter estimate

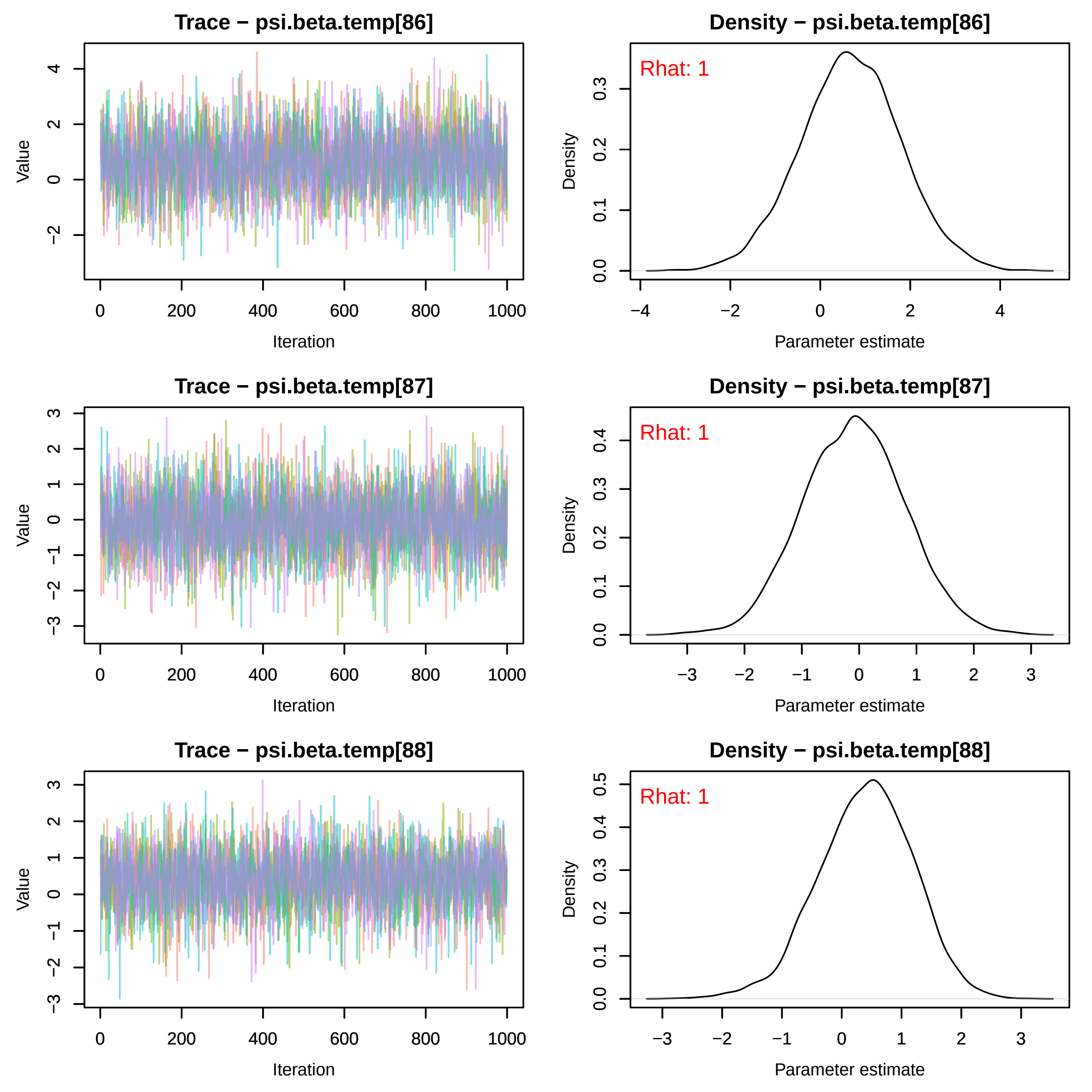
Trace − psi.beta.temp[86]
Density − psi.beta.temp[86]
Rhat: 1
Value
Iteration
Density
Parameter estimate
Trace − psi.beta.temp[87]
Density − psi.beta.temp[87]
Rhat: 1
Value
Iteration
Density
Parameter estimate
Trace − psi.beta.temp[88]
Density − psi.beta.temp[88]
Rhat: 1
Value
Iteration
Density
Parameter estimate

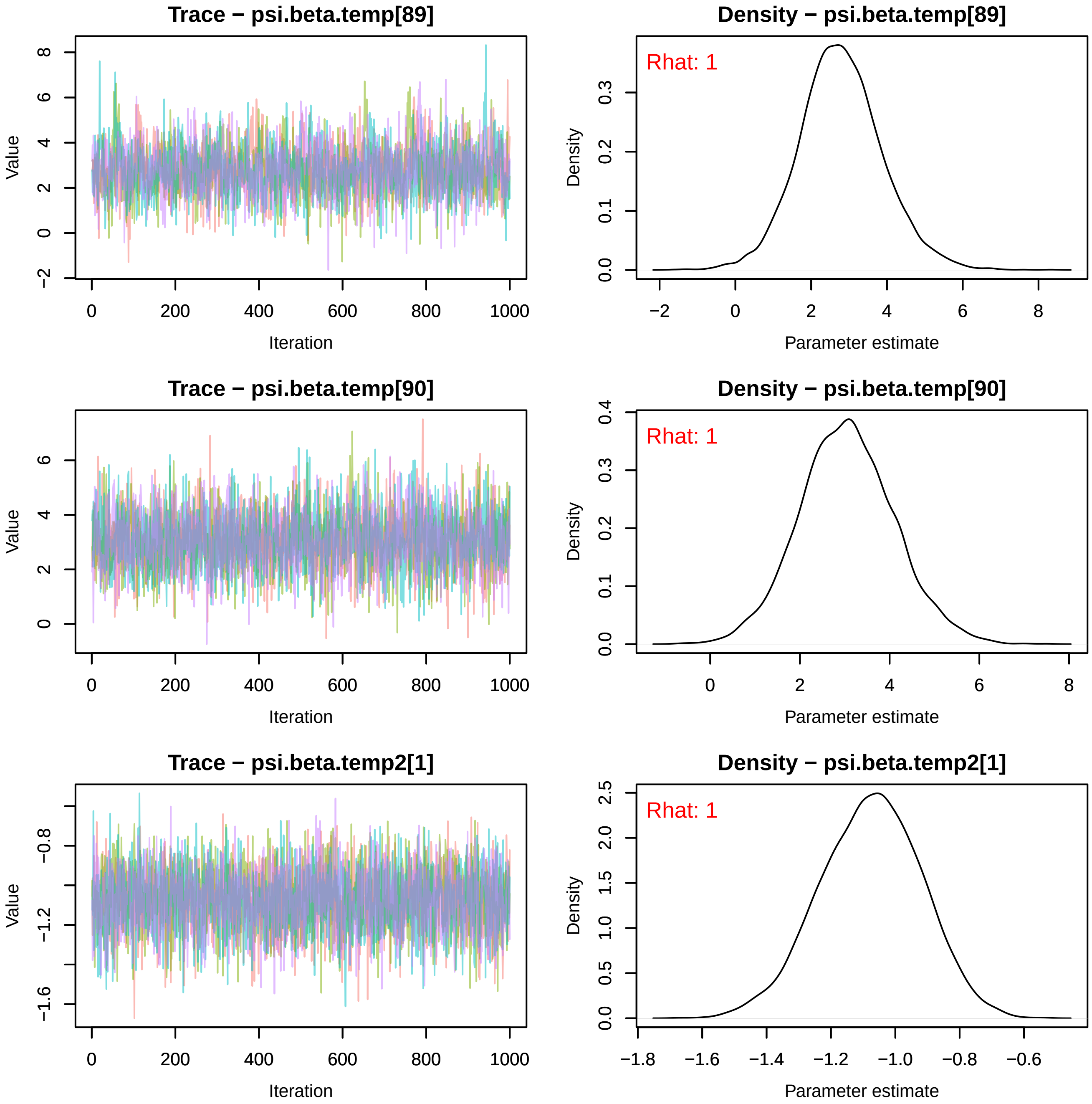
Trace − psi.beta.temp[89]
Value
Iteration
−2
0
2
4
6
8
0
200
400
600
800
1000
Density − psi.beta.temp[89]
Rhat: 1
Density
Parameter estimate
0.0
0.1
0.2
0.3
−2
0
2
4
6
8
Trace − psi.beta.temp[90]
Value
Iteration
0
2
4
6
0
200
400
600
800
1000
Density − psi.beta.temp[90]
Rhat: 1
Density
Parameter estimate
0.0
0.1
0.2
0.3
0.4
0
2
4
6
8
Trace − psi.beta.temp2[1]
Value
Iteration
−1.6
−1.2
−0.8
0
200
400
600
800
1000
Density − psi.beta.temp2[1]
Rhat: 1
Density
Parameter estimate
0.0
0.5
1.0
1.5
2.0
2.5
−1.8
−1.6
−1.4
−1.2
−1.0
−0.8
−0.6

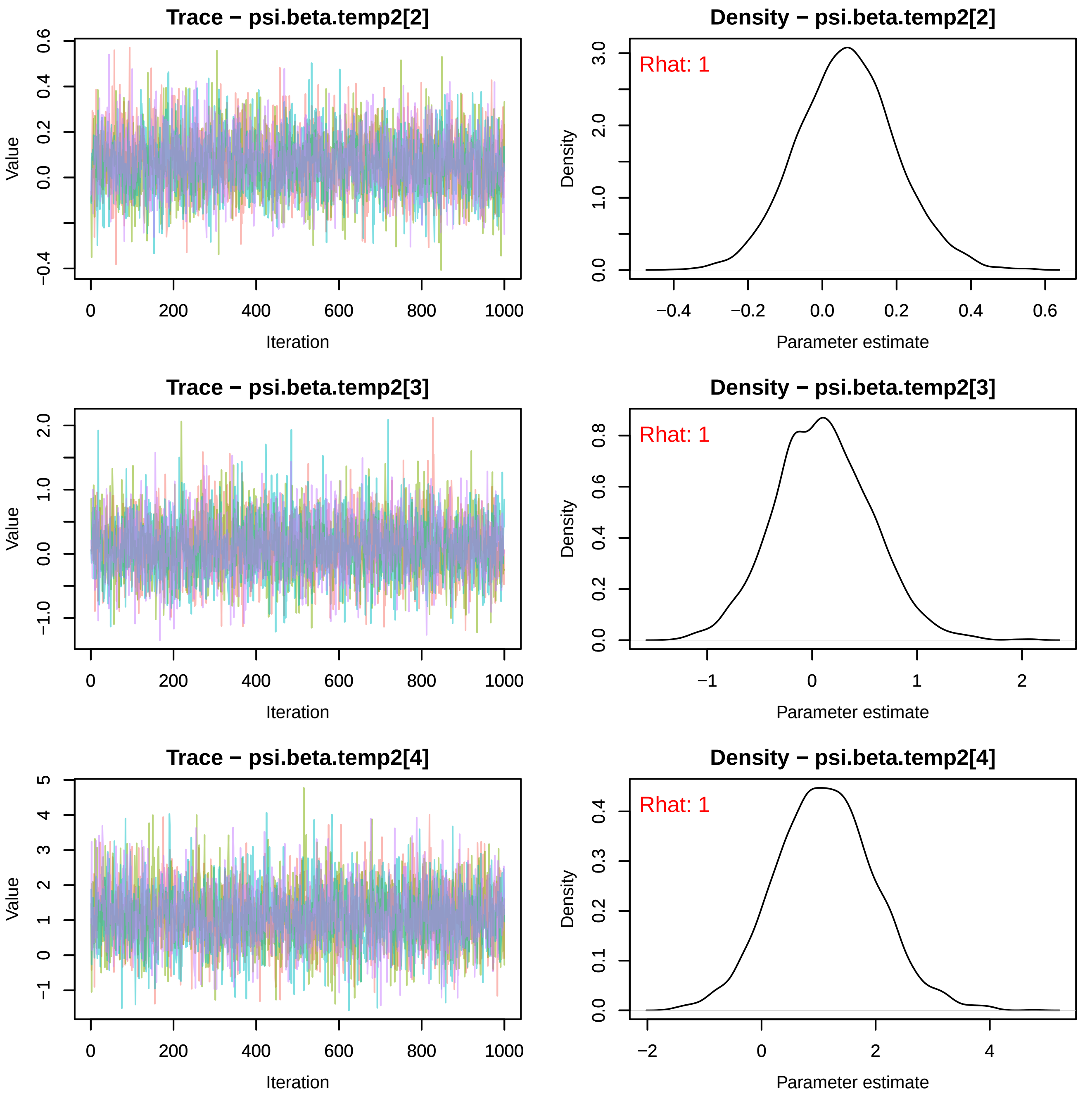
Trace − psi.beta.temp2[2]
Density − psi.beta.temp2[2]
Rhat: 1
Value
Iteration
Density
Parameter estimate
Trace − psi.beta.temp2[3]
Density − psi.beta.temp2[3]
Rhat: 1
Value
Iteration
Density
Parameter estimate
Trace − psi.beta.temp2[4]
Density − psi.beta.temp2[4]
Rhat: 1
Value
Iteration
Density
Parameter estimate

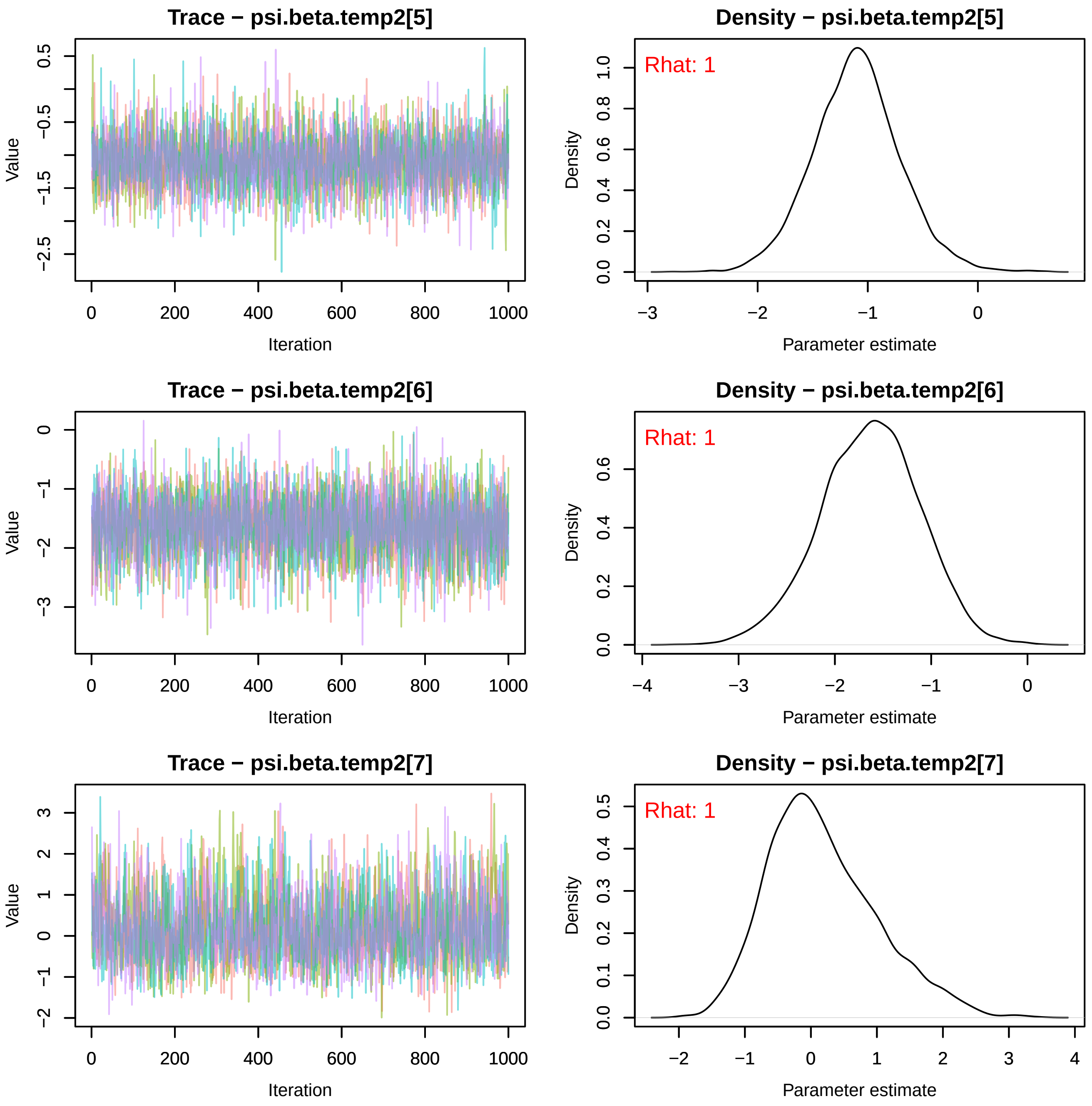
Trace − psi.beta.temp2[5]
Value
Iteration
0.5
−0.5
−1.5
−2.5
0
200
400
600
800
1000
Density − psi.beta.temp2[5]
Rhat: 1
Density
Parameter estimate
0.0
0.2
0.4
0.6
0.8
1.0
−3
−2
−1
0
Trace − psi.beta.temp2[6]
Value
Iteration
0
−1
−2
−3
0
200
400
600
800
1000
Density − psi.beta.temp2[6]
Rhat: 1
Density
Parameter estimate
0.0
0.2
0.4
0.6
−4
−3
−2
−1
0
Trace − psi.beta.temp2[7]
Value
Iteration
3
2
1
0
−1
−2
0
200
400
600
800
1000
Density − psi.beta.temp2[7]
Rhat: 1
Density
Parameter estimate
0.0
0.1
0.2
0.3
0.4
0.5
−2
−1
0
1
2
3
4

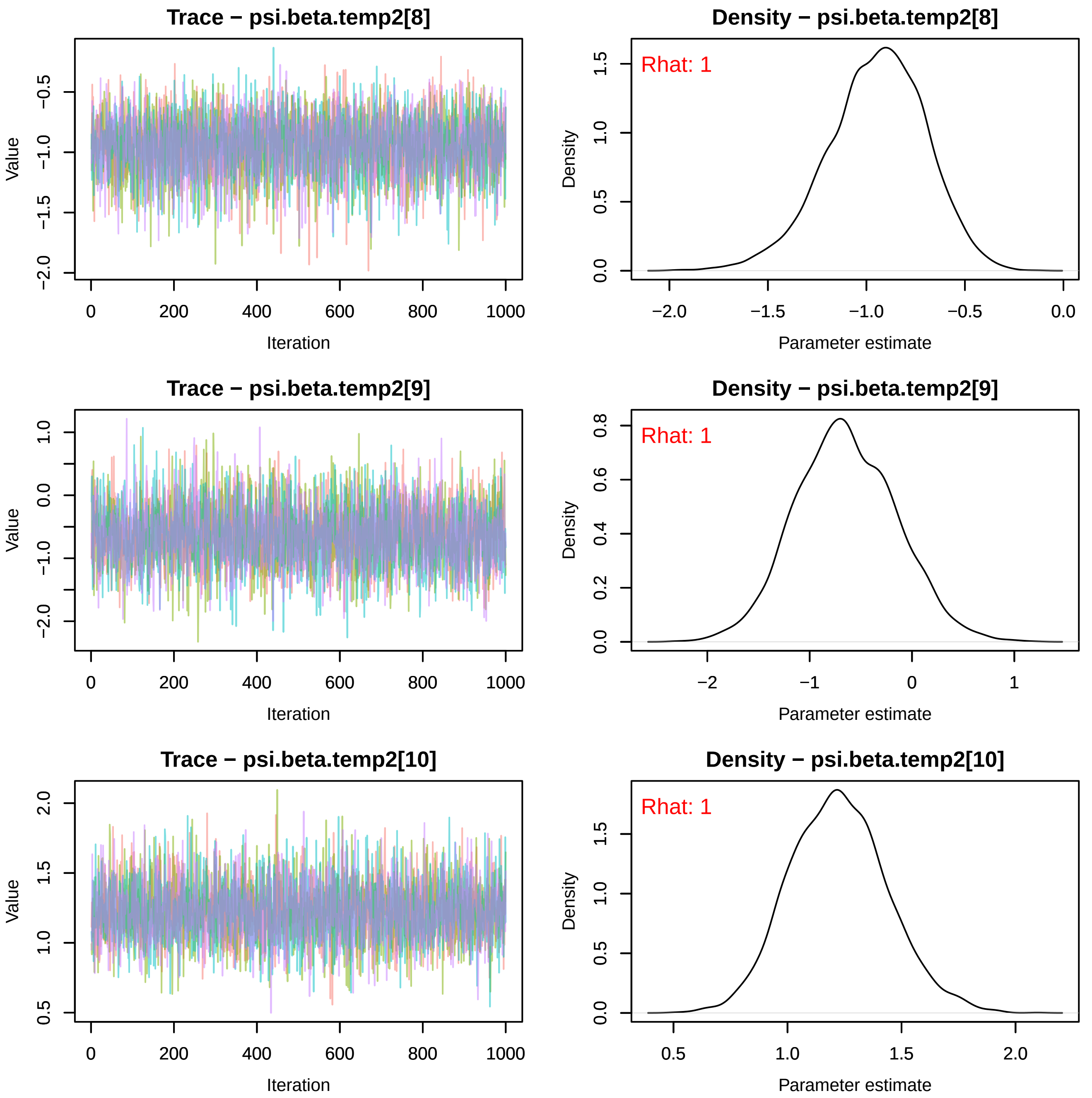
Trace − psi.beta.temp2[8]
Value
Iteration
Density − psi.beta.temp2[8]
Rhat: 1
Density
Parameter estimate
Trace − psi.beta.temp2[9]
Value
Iteration
Density − psi.beta.temp2[9]
Rhat: 1
Density
Parameter estimate
Trace − psi.beta.temp2[10]
Value
Iteration
Density − psi.beta.temp2[10]
Rhat: 1
Density
Parameter estimate

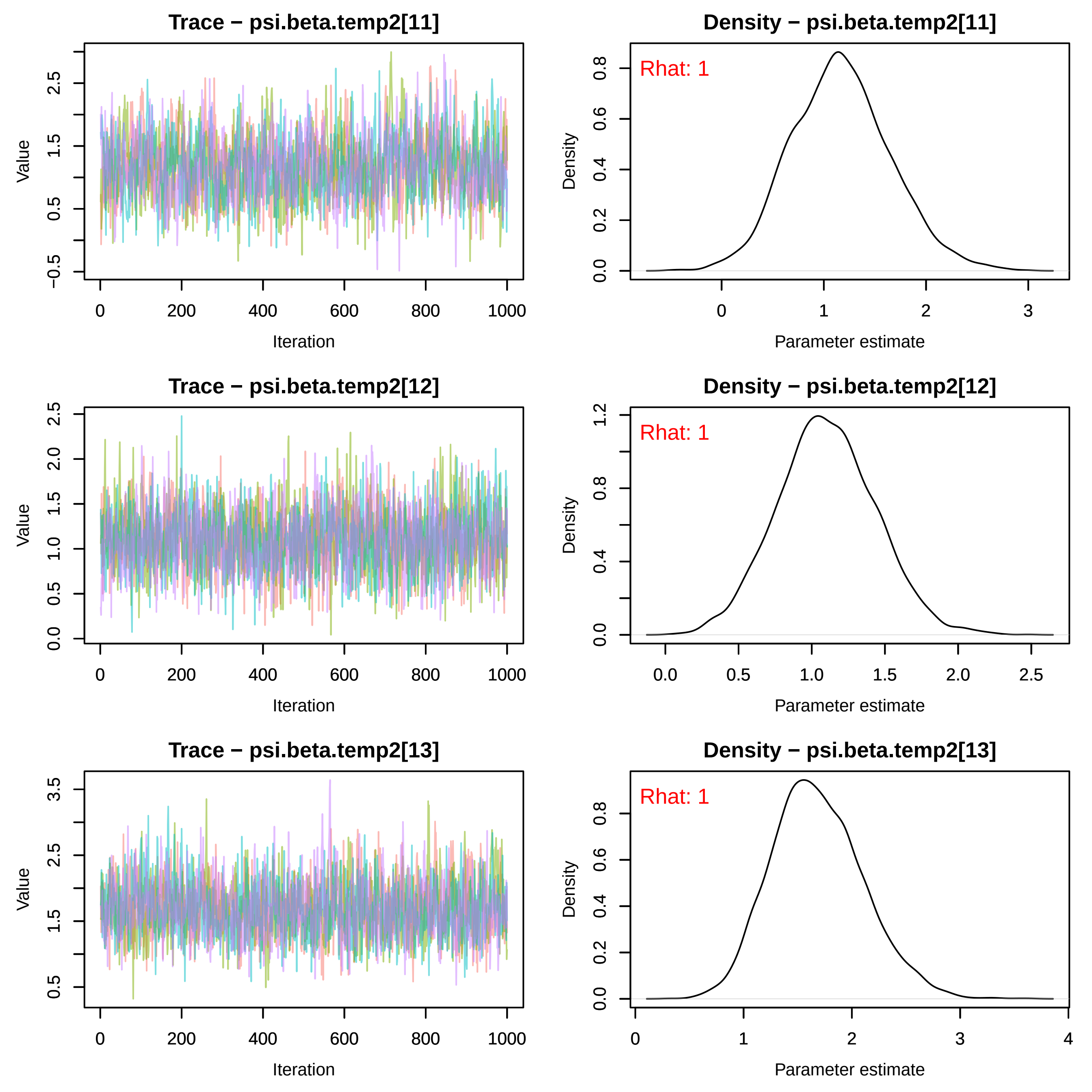
Trace − psi.beta.temp2[11]
Density − psi.beta.temp2[11]
Rhat: 1
Trace − psi.beta.temp2[12]
Density − psi.beta.temp2[12]
Rhat: 1
Trace − psi.beta.temp2[13]
Density − psi.beta.temp2[13]
Rhat: 1
Value
Iteration
Density
Parameter estimate

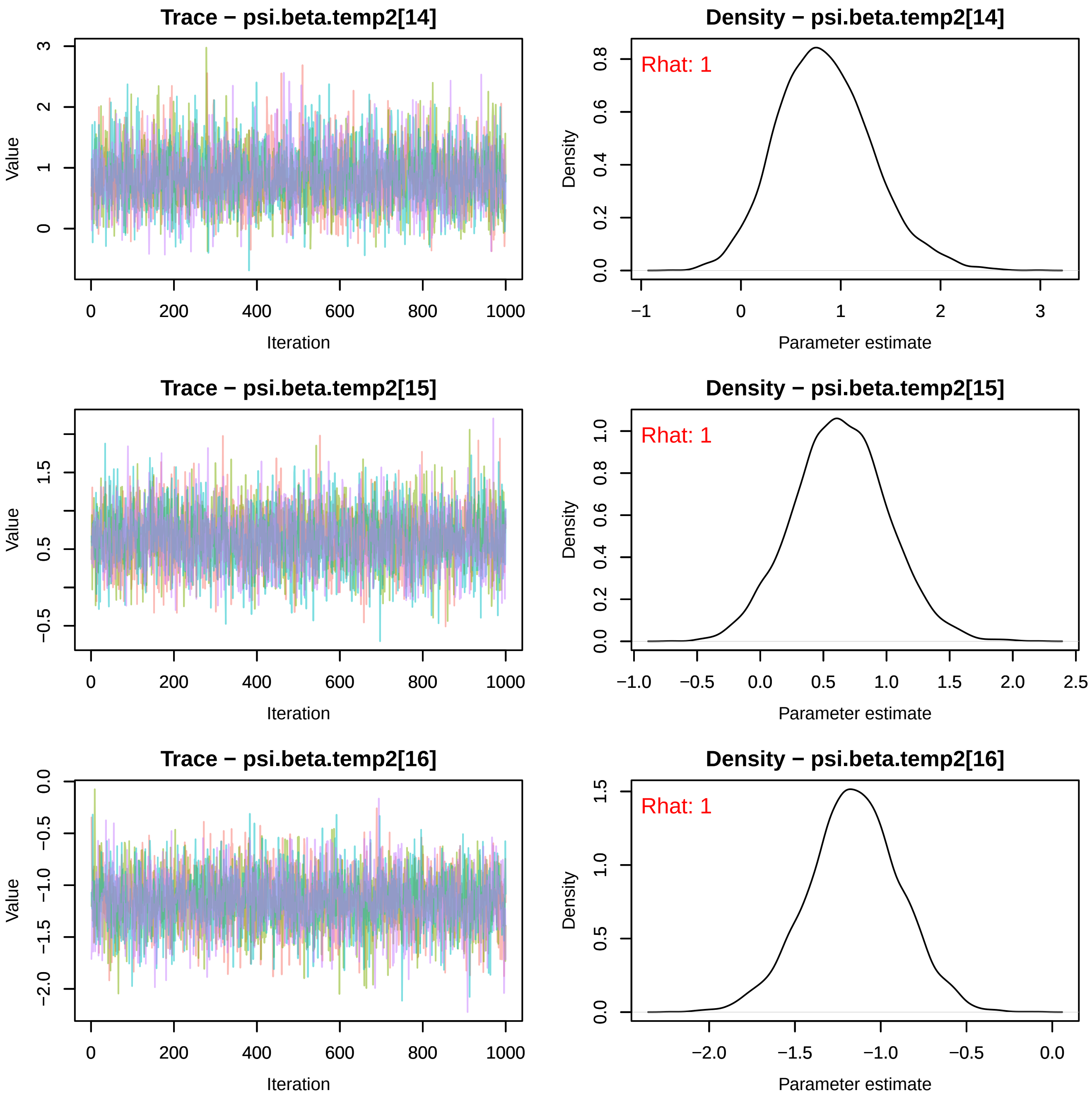
Trace − psi.beta.temp2[14]
Value
Iteration
Density − psi.beta.temp2[14]
Rhat: 1
Density
Parameter estimate
Trace − psi.beta.temp2[15]
Value
Iteration
Density − psi.beta.temp2[15]
Rhat: 1
Density
Parameter estimate
Trace − psi.beta.temp2[16]
Value
Iteration
Density − psi.beta.temp2[16]
Rhat: 1
Density
Parameter estimate

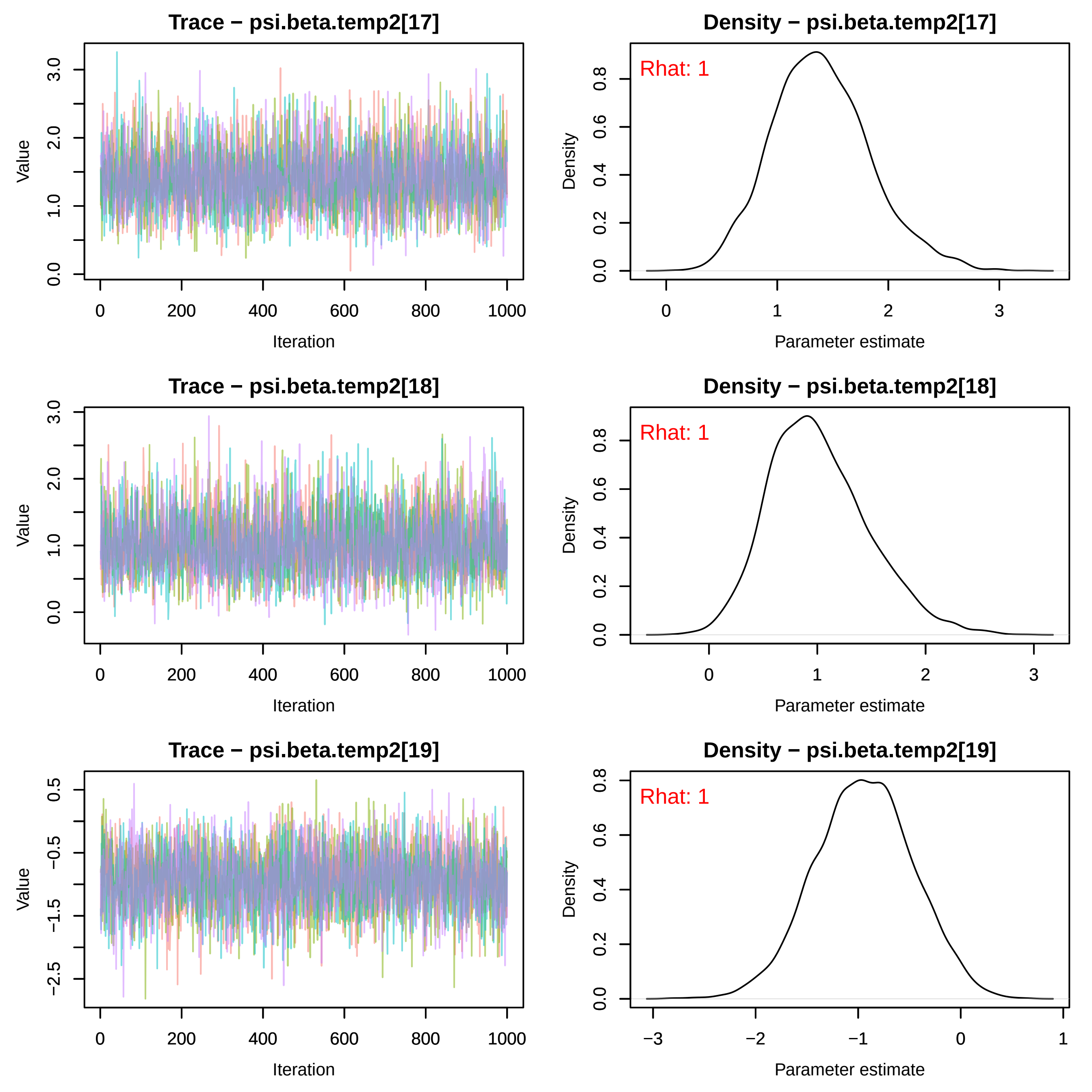

Trace − psi.beta.temp2[17]
Value
Iteration
0.0
1.0
2.0
3.0
0
200
400
600
800
1000
Density − psi.beta.temp2[17]
Rhat: 1
Density
Parameter estimate
0.0
0.2
0.4
0.6
0.8
0
1
2
3
Trace − psi.beta.temp2[18]
Value
Iteration
0.0
1.0
2.0
3.0
0
200
400
600
800
1000
Density − psi.beta.temp2[18]
Rhat: 1
Density
Parameter estimate
0.0
0.2
0.4
0.6
0.8
0
1
2
3
Trace − psi.beta.temp2[19]
Value
Iteration
−2.5
−1.5
−0.5
0.5
0
200
400
600
800
1000
Density − psi.beta.temp2[19]
Rhat: 1
Density
Parameter estimate
0.0
0.2
0.4
0.6
0.8
−3
−2
−1
0
1

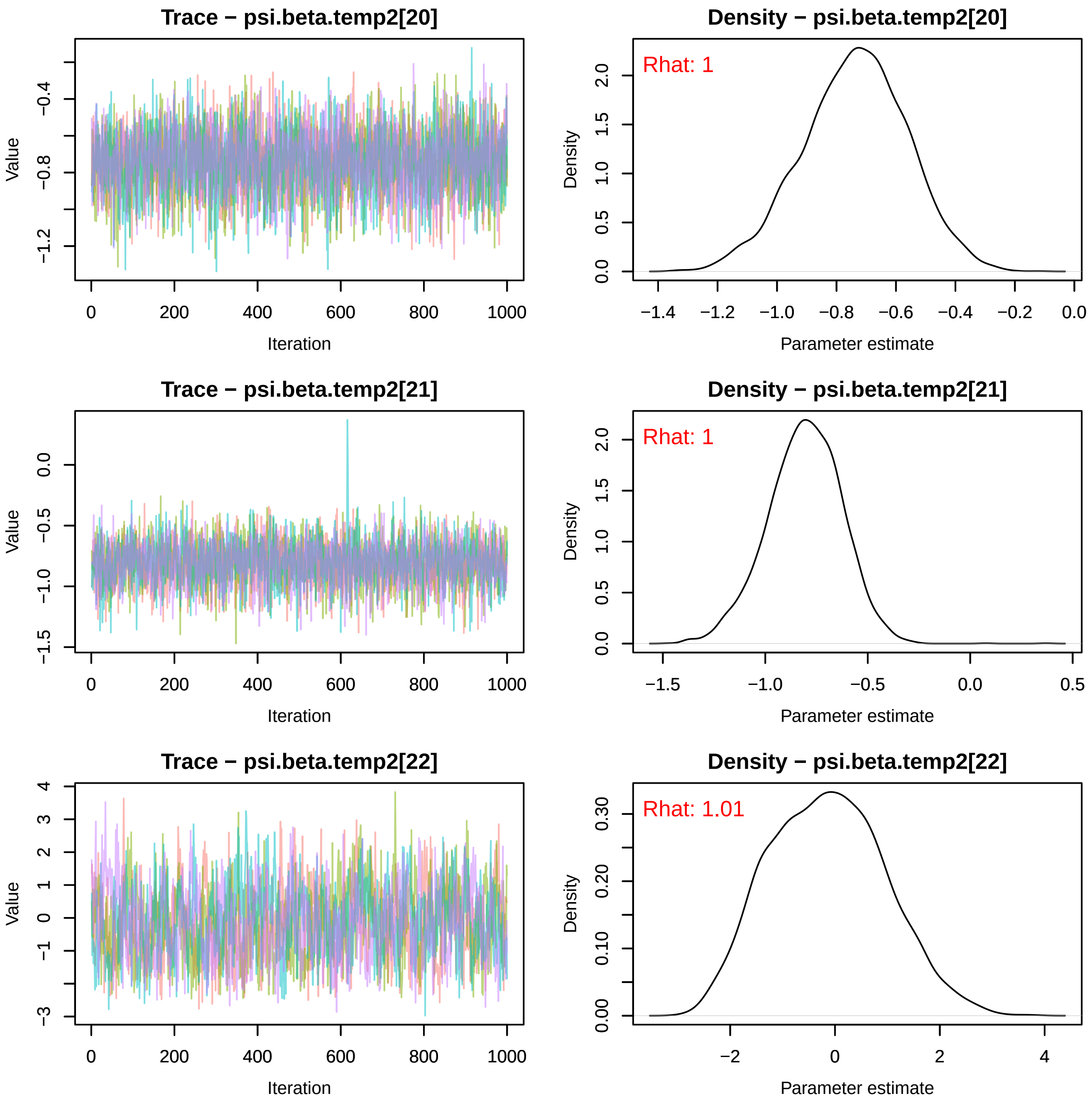
Trace − psi.beta.temp2[20]
Value
Iteration
Density − psi.beta.temp2[20]
Rhat: 1
Density
Parameter estimate
Trace − psi.beta.temp2[21]
Value
Iteration
Density − psi.beta.temp2[21]
Rhat: 1
Density
Parameter estimate
Trace − psi.beta.temp2[22]
Value
Iteration
Density − psi.beta.temp2[22]
Rhat: 1.01
Density
Parameter estimate

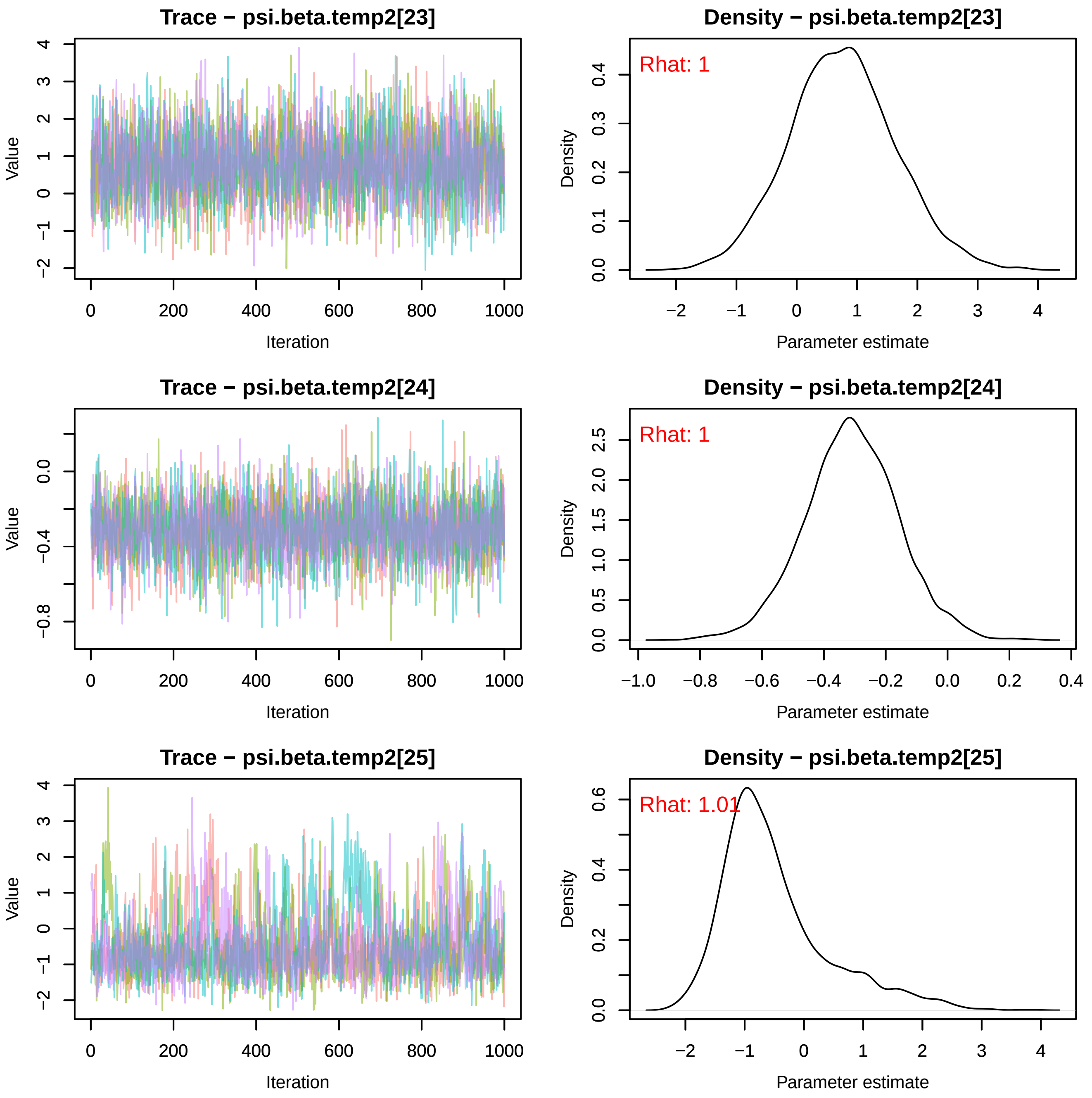
Trace − psi.beta.temp2[23]
Value
Iteration
Density − psi.beta.temp2[23]
Rhat: 1
Density
Parameter estimate
Trace − psi.beta.temp2[24]
Value
Iteration
Density − psi.beta.temp2[24]
Rhat: 1
Density
Parameter estimate
Trace − psi.beta.temp2[25]
Value
Iteration
Density − psi.beta.temp2[25]
Rhat: 1.01
Density
Parameter estimate

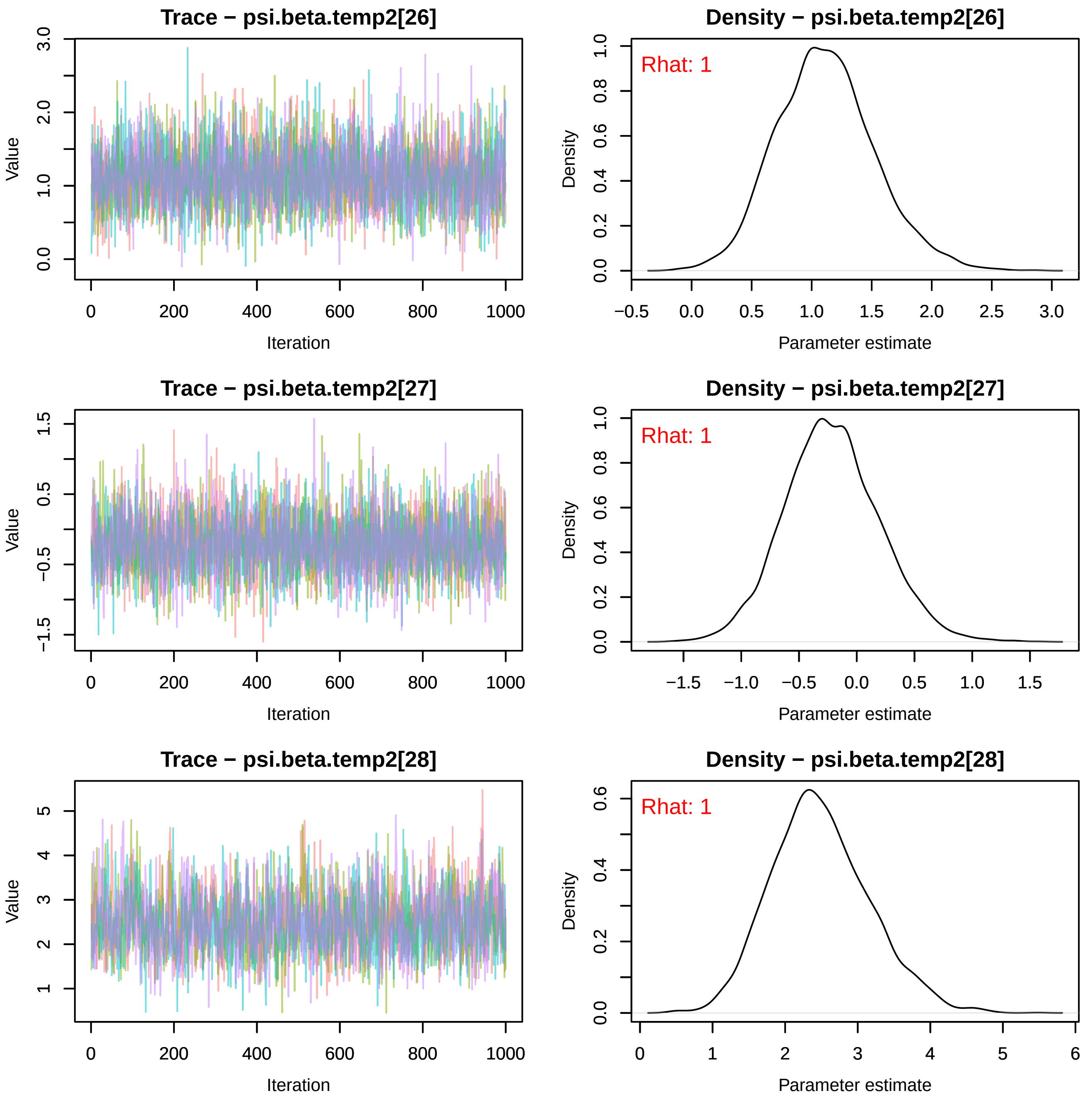
Trace − psi.beta.temp2[26]
Value
Iteration
Density − psi.beta.temp2[26]
Rhat: 1
Density
Parameter estimate
Trace − psi.beta.temp2[27]
Value
Iteration
Density − psi.beta.temp2[27]
Rhat: 1
Density
Parameter estimate
Trace − psi.beta.temp2[28]
Value
Iteration
Density − psi.beta.temp2[28]
Rhat: 1
Density
Parameter estimate

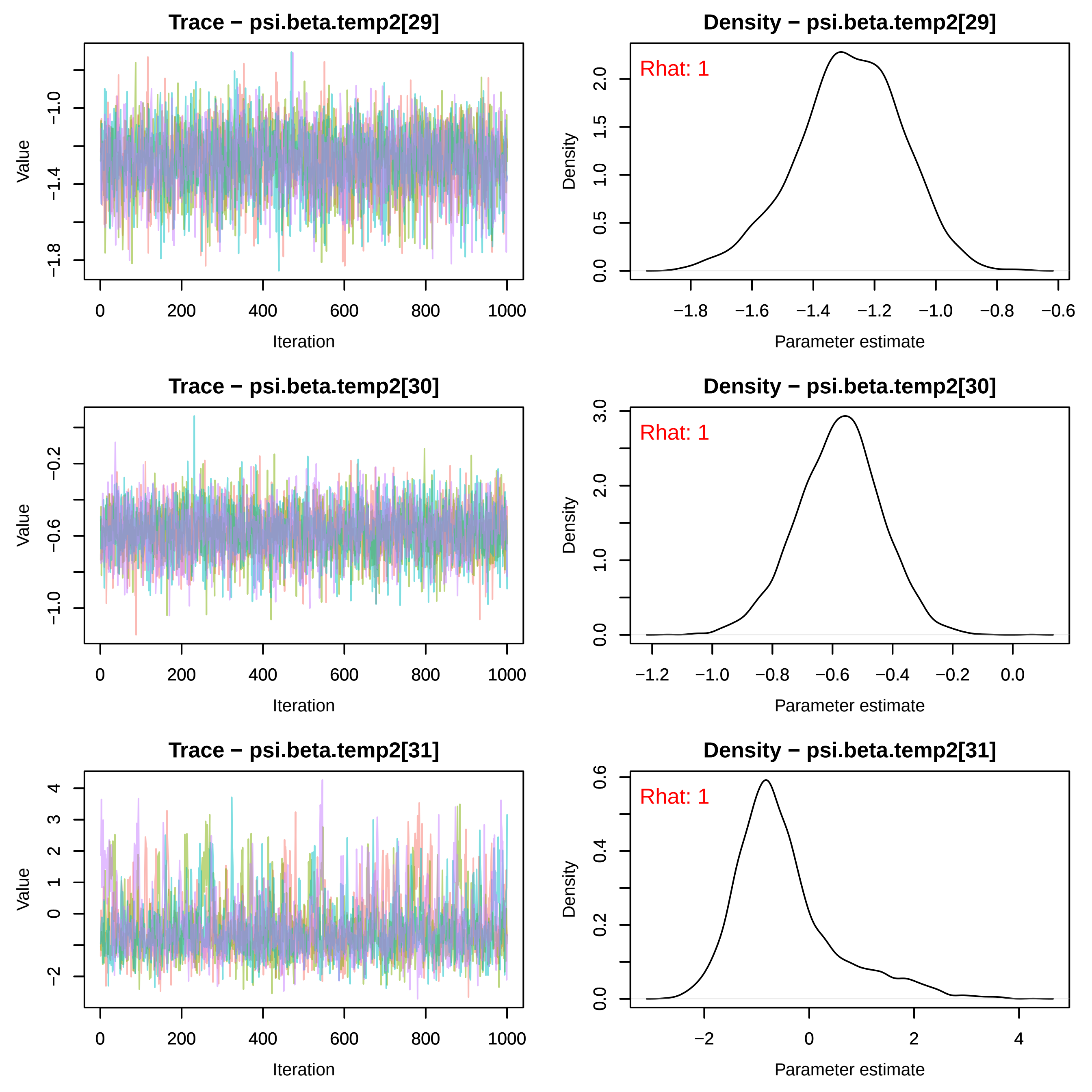
Trace − psi.beta.temp2[29]
Density − psi.beta.temp2[29]
Rhat: 1
Trace − psi.beta.temp2[30]
Density − psi.beta.temp2[30]
Rhat: 1
Trace − psi.beta.temp2[31]
Density − psi.beta.temp2[31]
Rhat: 1
Value
Iteration
Density
Parameter estimate

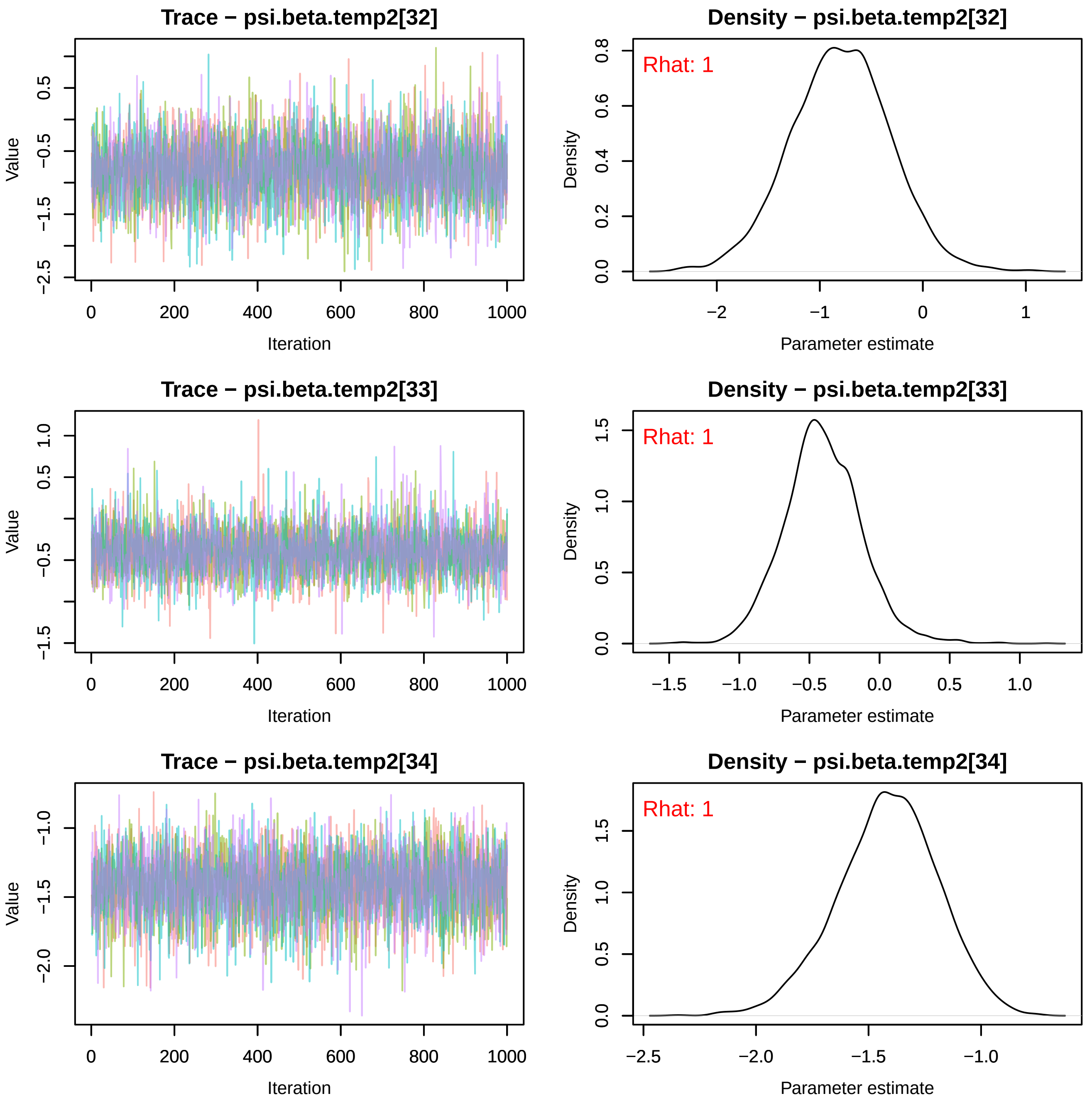

Trace − psi.beta.temp2[32]
Value
Iteration
Density − psi.beta.temp2[32]
Rhat: 1
Density
Parameter estimate
Trace − psi.beta.temp2[33]
Value
Iteration
Density − psi.beta.temp2[33]
Rhat: 1
Density
Parameter estimate
Trace − psi.beta.temp2[34]
Value
Iteration
Density − psi.beta.temp2[34]
Rhat: 1
Density
Parameter estimate

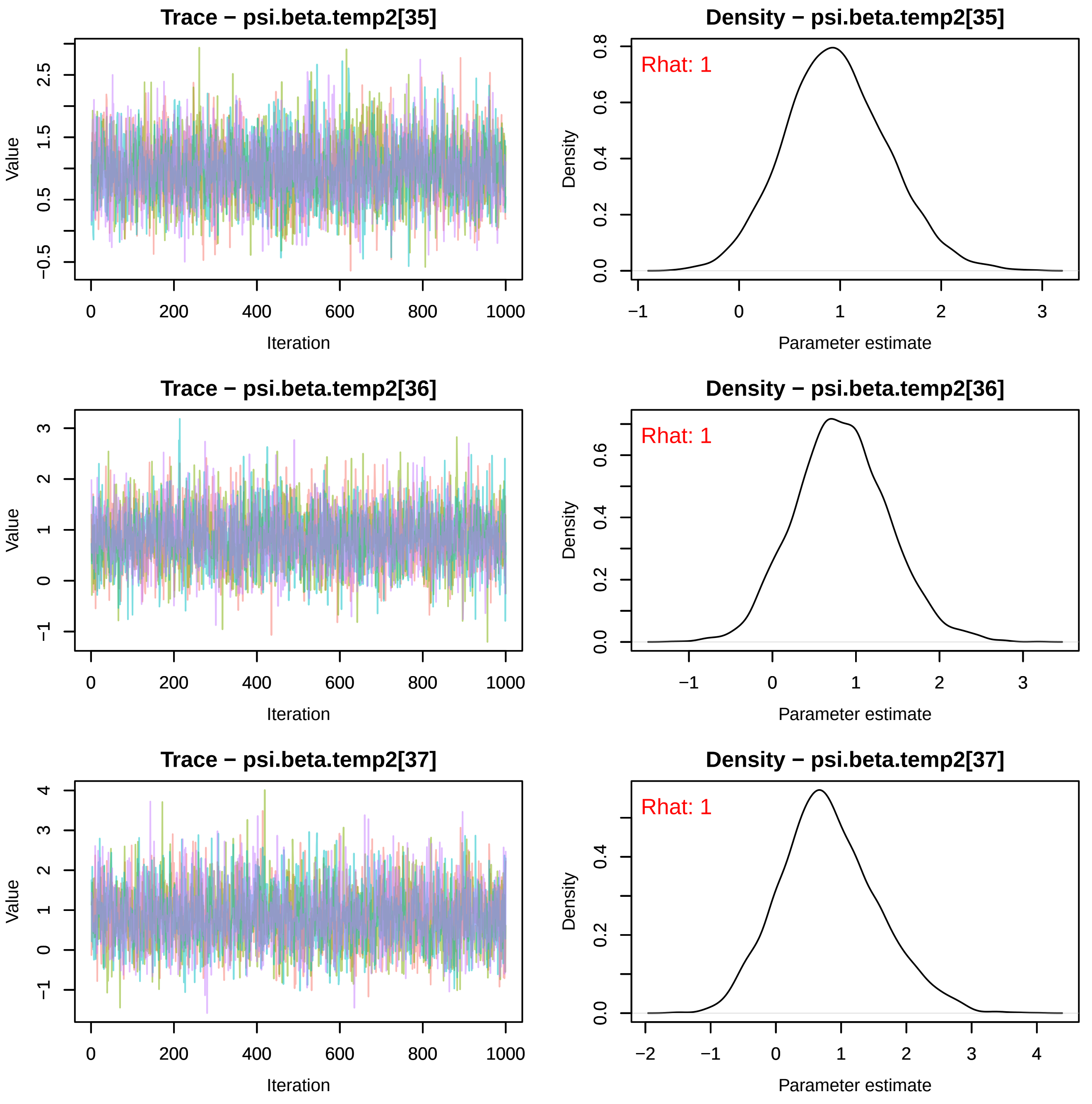
Trace − psi.beta.temp2[35]
Density − psi.beta.temp2[35]
Rhat: 1
Trace − psi.beta.temp2[36]
Density − psi.beta.temp2[36]
Rhat: 1
Trace − psi.beta.temp2[37]
Density − psi.beta.temp2[37]
Rhat: 1
Value
Iteration
Density
Parameter estimate

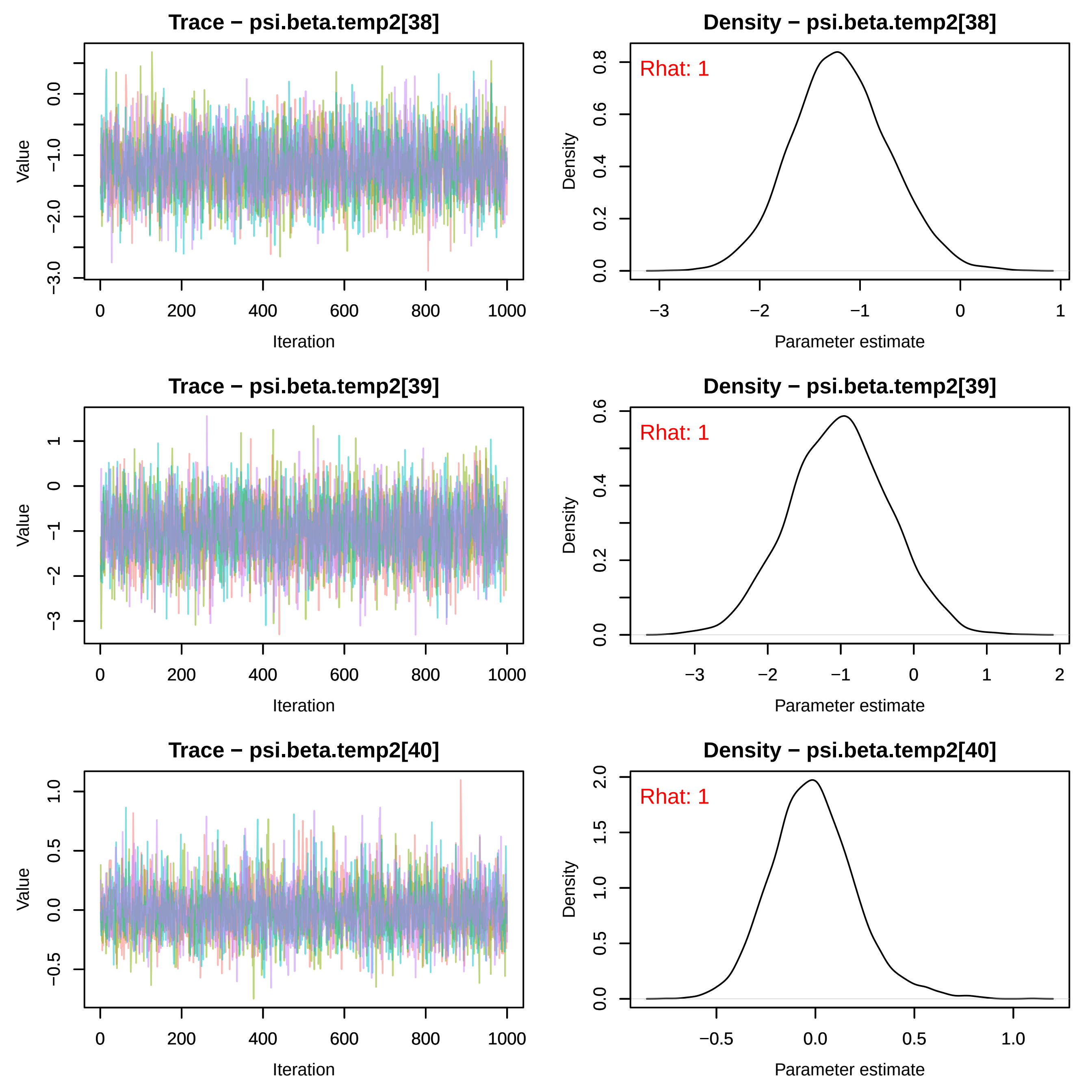

Trace − psi.beta.temp2[38]
Density − psi.beta.temp2[38]
Rhat: 1
Trace − psi.beta.temp2[39]
Density − psi.beta.temp2[39]
Rhat: 1
Trace − psi.beta.temp2[40]
Density − psi.beta.temp2[40]
Rhat: 1
Value
Iteration
Density
Parameter estimate

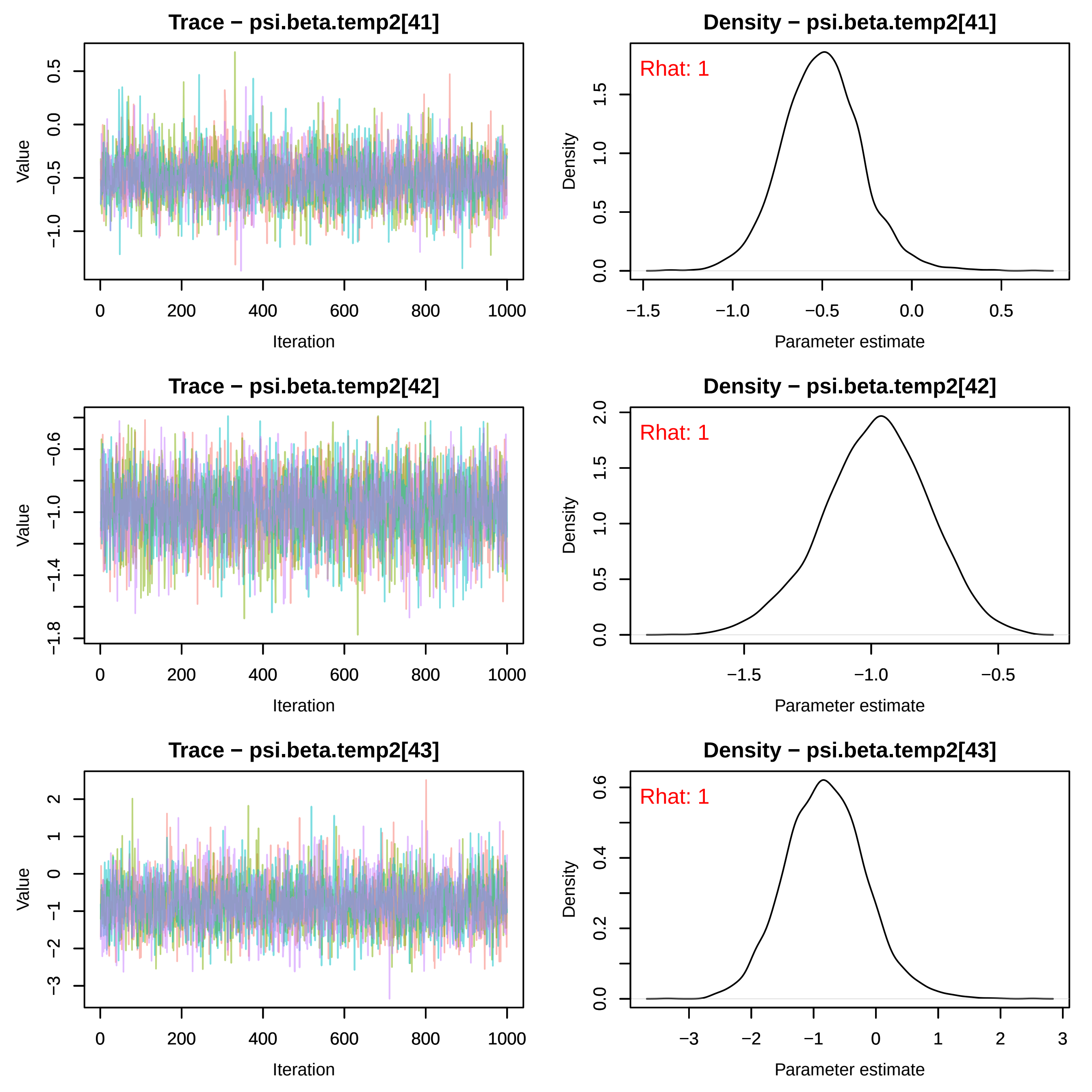
Trace − psi.beta.temp2[41]
Density − psi.beta.temp2[41]
Rhat: 1
Trace − psi.beta.temp2[42]
Density − psi.beta.temp2[42]
Rhat: 1
Trace − psi.beta.temp2[43]
Density − psi.beta.temp2[43]
Rhat: 1
Value
Iteration
Density
Parameter estimate

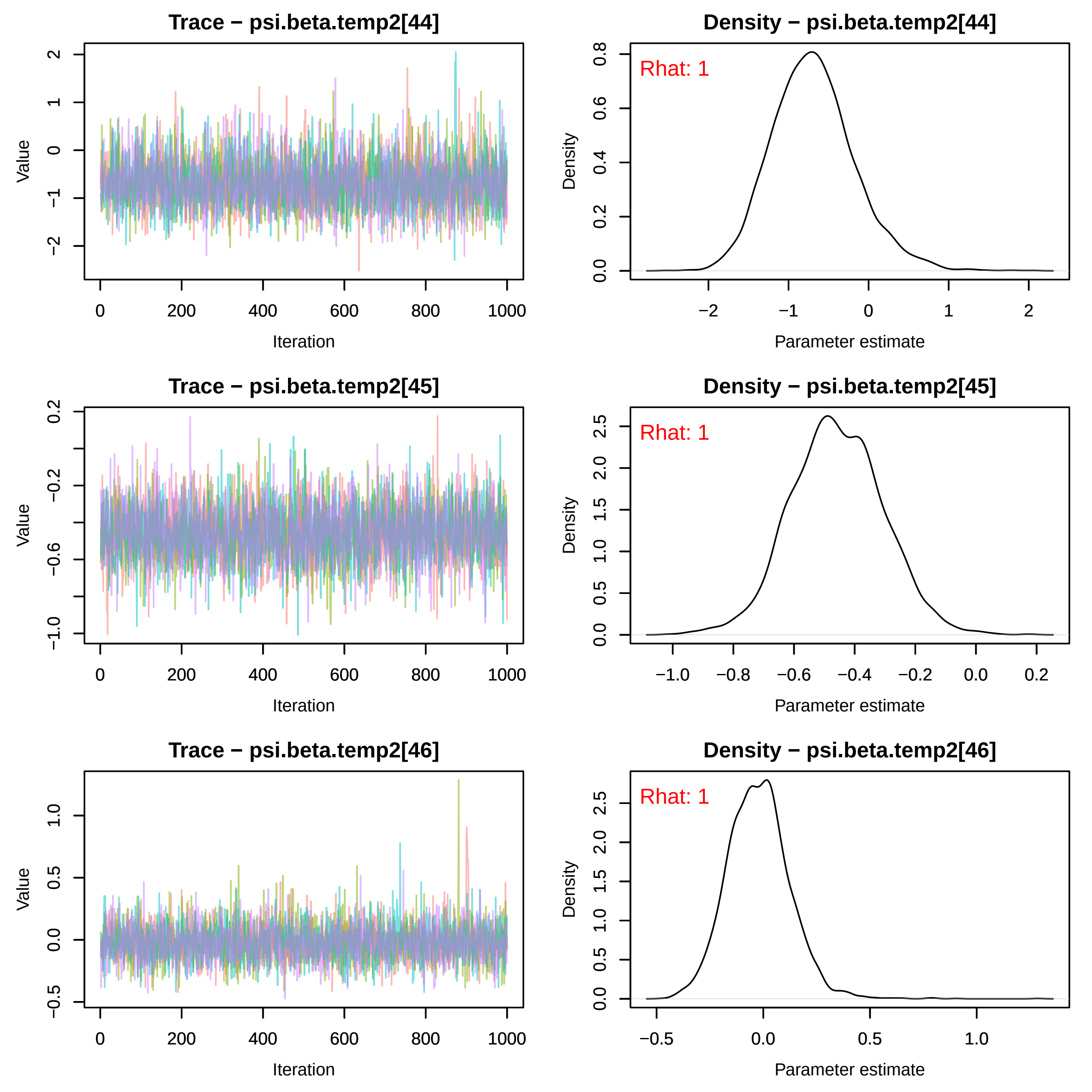
Trace − psi.beta.temp2[44]
Density − psi.beta.temp2[44]
Rhat: 1
Trace − psi.beta.temp2[45]
Density − psi.beta.temp2[45]
Rhat: 1
Trace − psi.beta.temp2[46]
Density − psi.beta.temp2[46]
Rhat: 1
Value
Iteration
Density
Parameter estimate

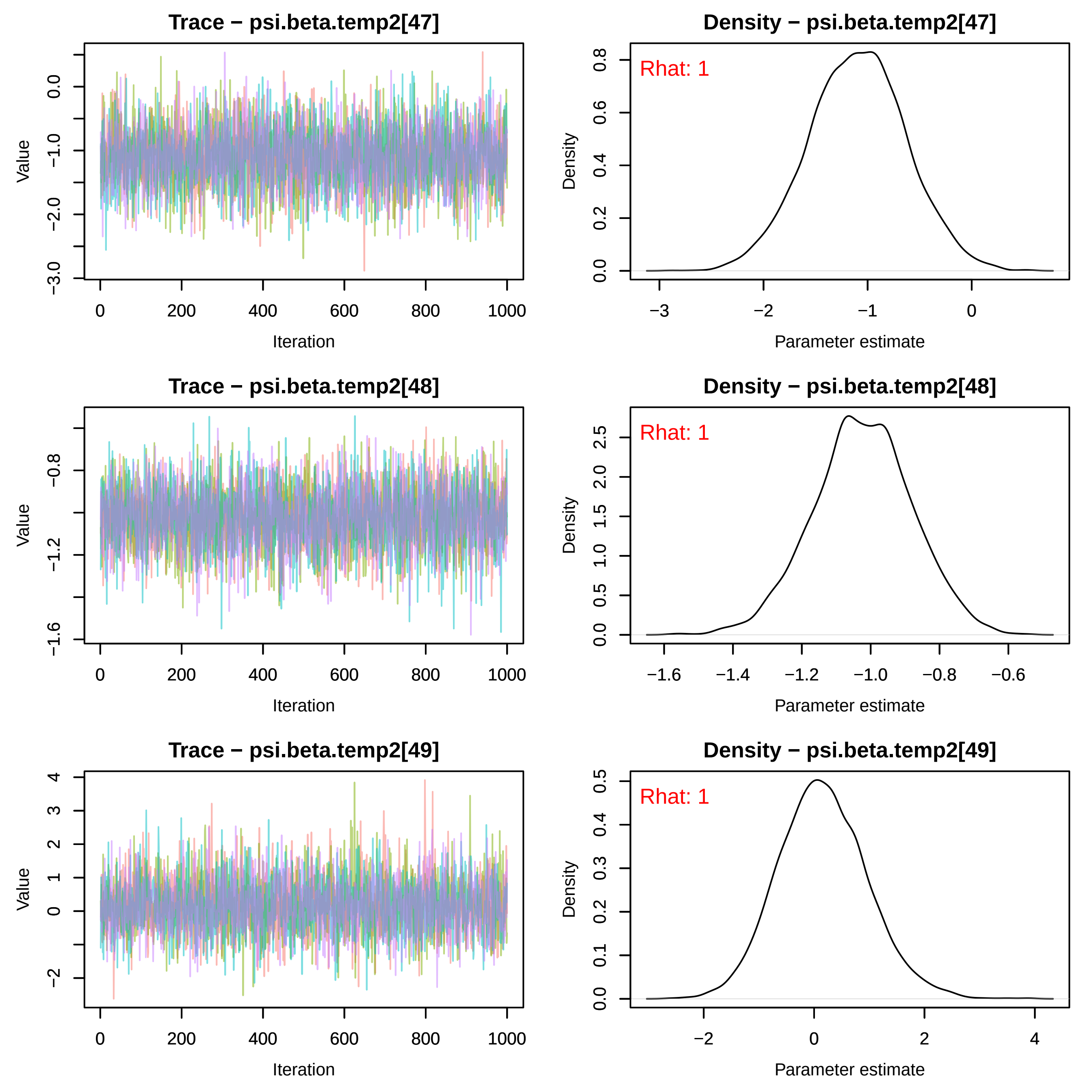
Trace − psi.beta.temp2[47]
Value
Iteration
Density − psi.beta.temp2[47]
Rhat: 1
Density
Parameter estimate
Trace − psi.beta.temp2[48]
Value
Iteration
Density − psi.beta.temp2[48]
Rhat: 1
Density
Parameter estimate
Trace − psi.beta.temp2[49]
Value
Iteration
Density − psi.beta.temp2[49]
Rhat: 1
Density
Parameter estimate

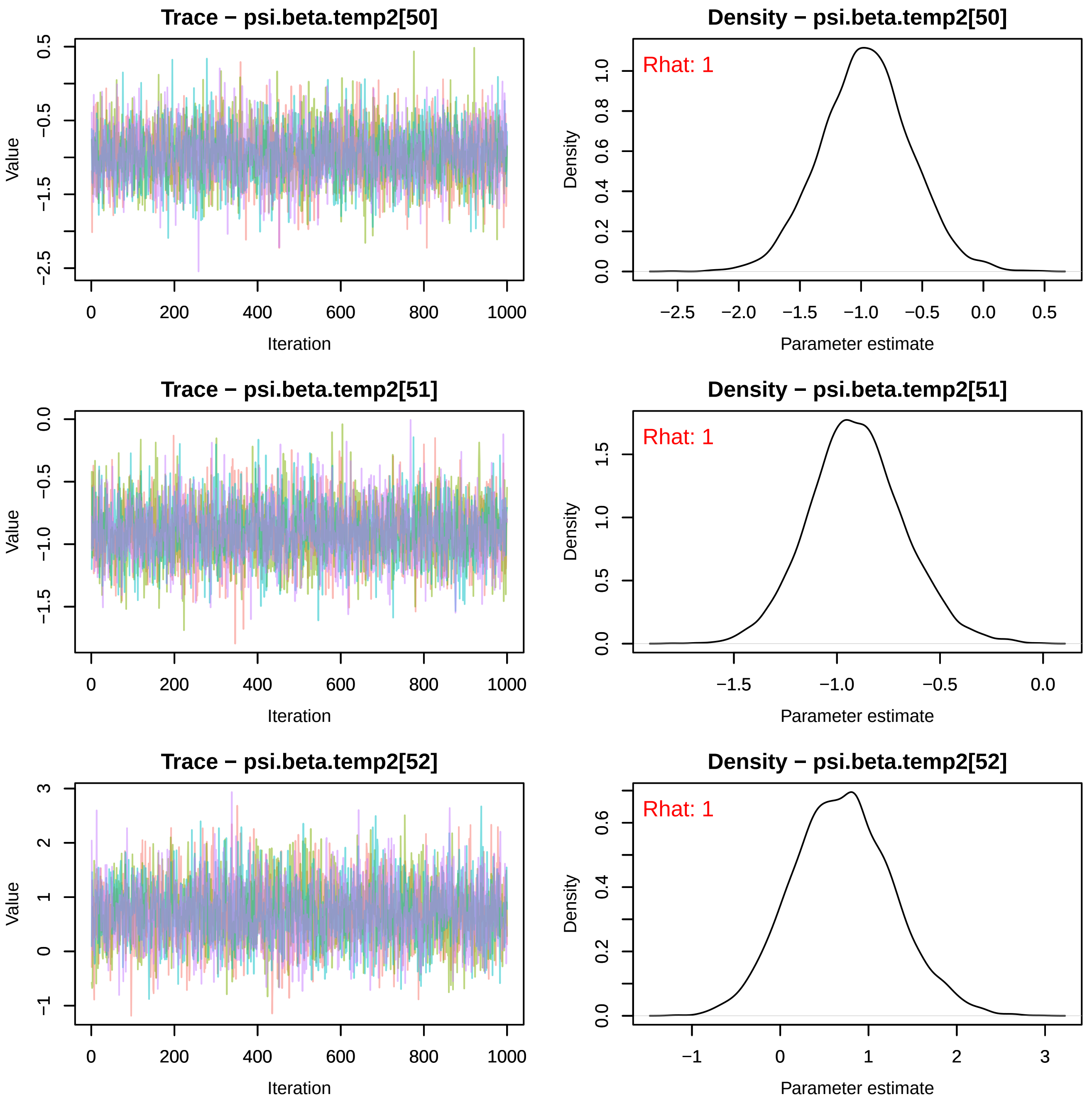

Trace − psi.beta.temp2[50]
Value
Iteration
Density − psi.beta.temp2[50]
Rhat: 1
Density
Parameter estimate
Trace − psi.beta.temp2[51]
Value
Iteration
Density − psi.beta.temp2[51]
Rhat: 1
Density
Parameter estimate
Trace − psi.beta.temp2[52]
Value
Iteration
Density − psi.beta.temp2[52]
Rhat: 1
Density
Parameter estimate

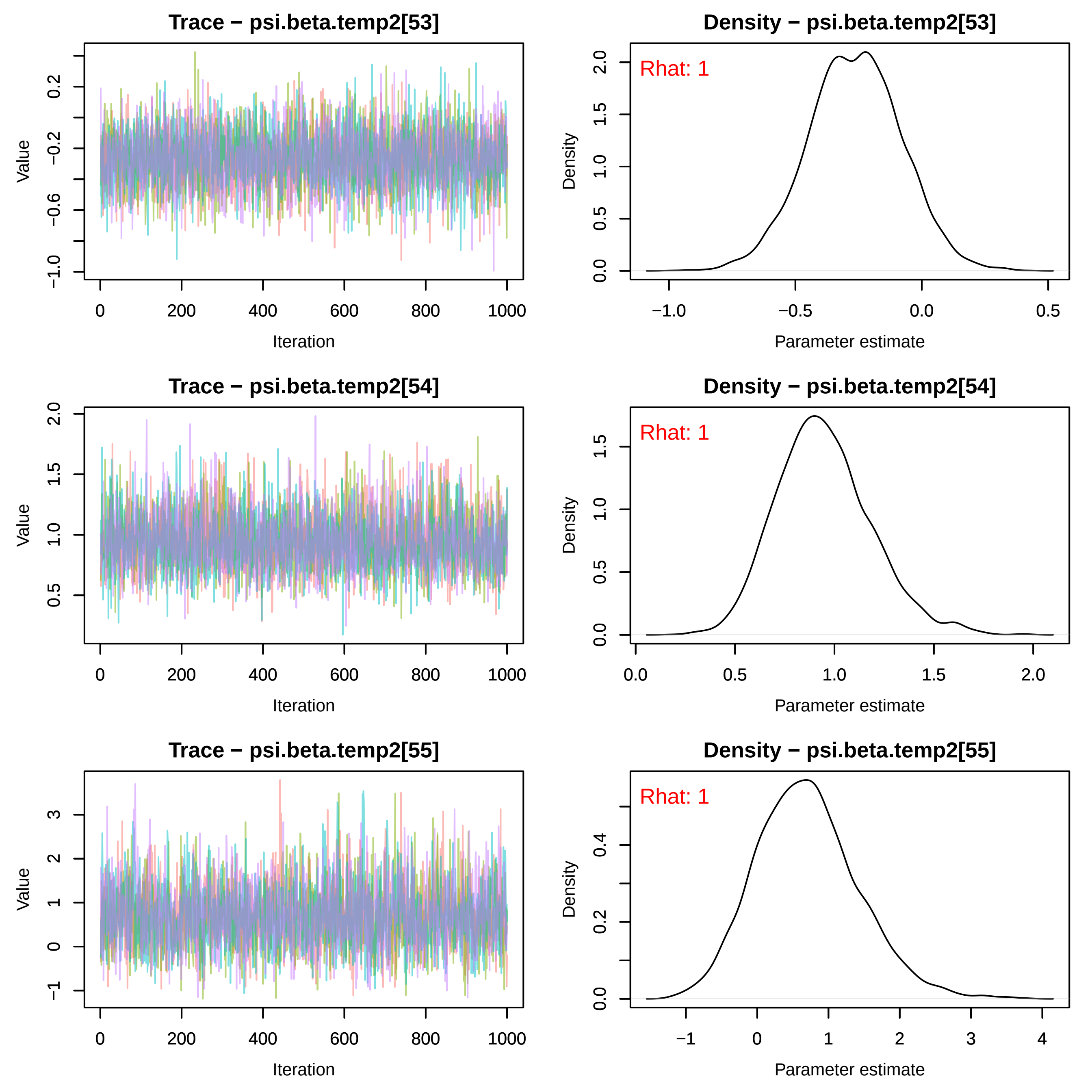
Trace − psi.beta.temp2[53]
Density − psi.beta.temp2[53]
Rhat: 1
Trace − psi.beta.temp2[54]
Density − psi.beta.temp2[54]
Rhat: 1
Trace − psi.beta.temp2[55]
Density − psi.beta.temp2[55]
Rhat: 1
Value
Iteration
Density
Parameter estimate

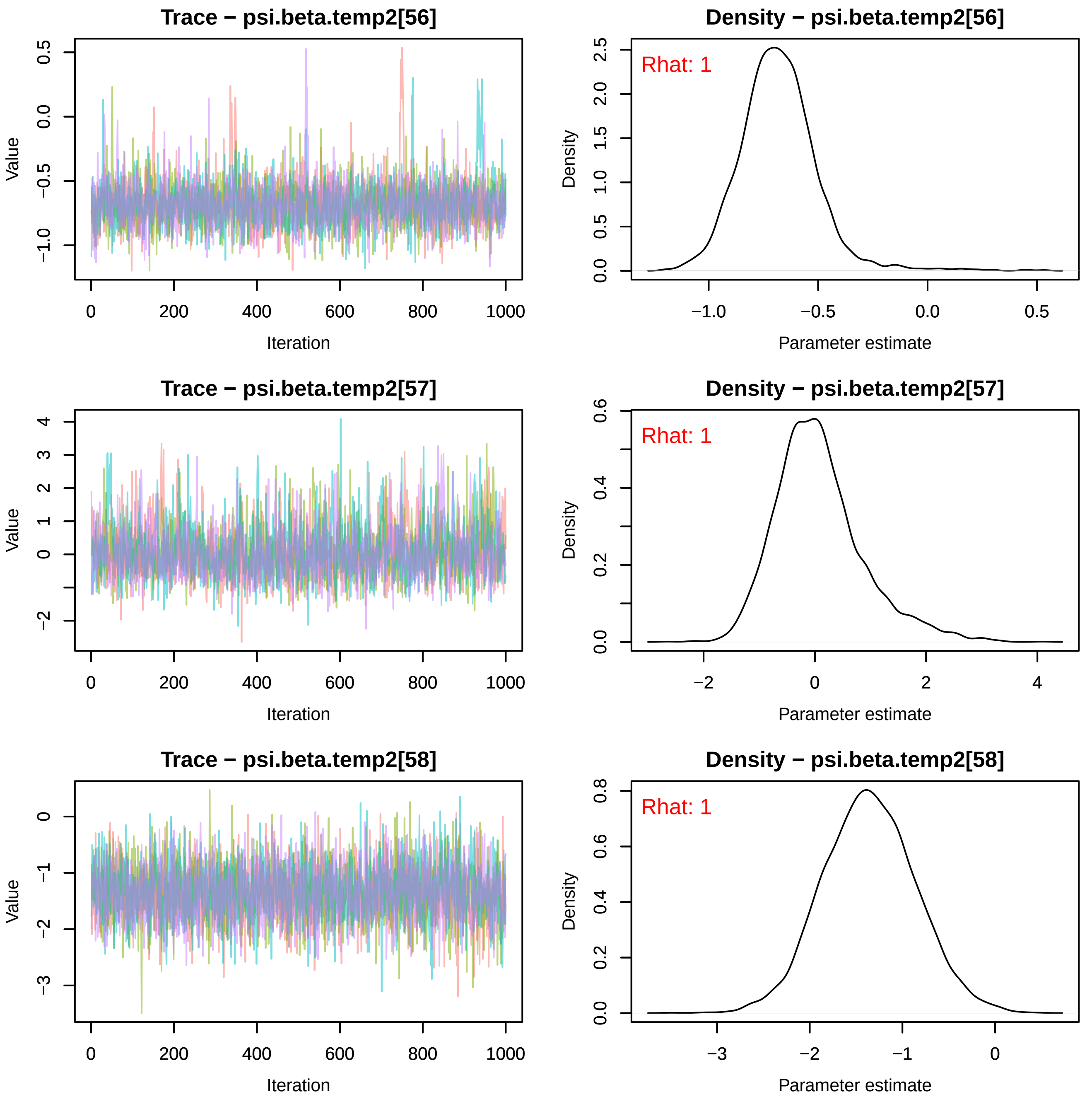
Trace − psi.beta.temp2[56]
Value
Iteration
Density − psi.beta.temp2[56]
Rhat: 1
Density
Parameter estimate
Trace − psi.beta.temp2[57]
Value
Iteration
Density − psi.beta.temp2[57]
Rhat: 1
Density
Parameter estimate
Trace − psi.beta.temp2[58]
Value
Iteration
Density − psi.beta.temp2[58]
Rhat: 1
Density
Parameter estimate

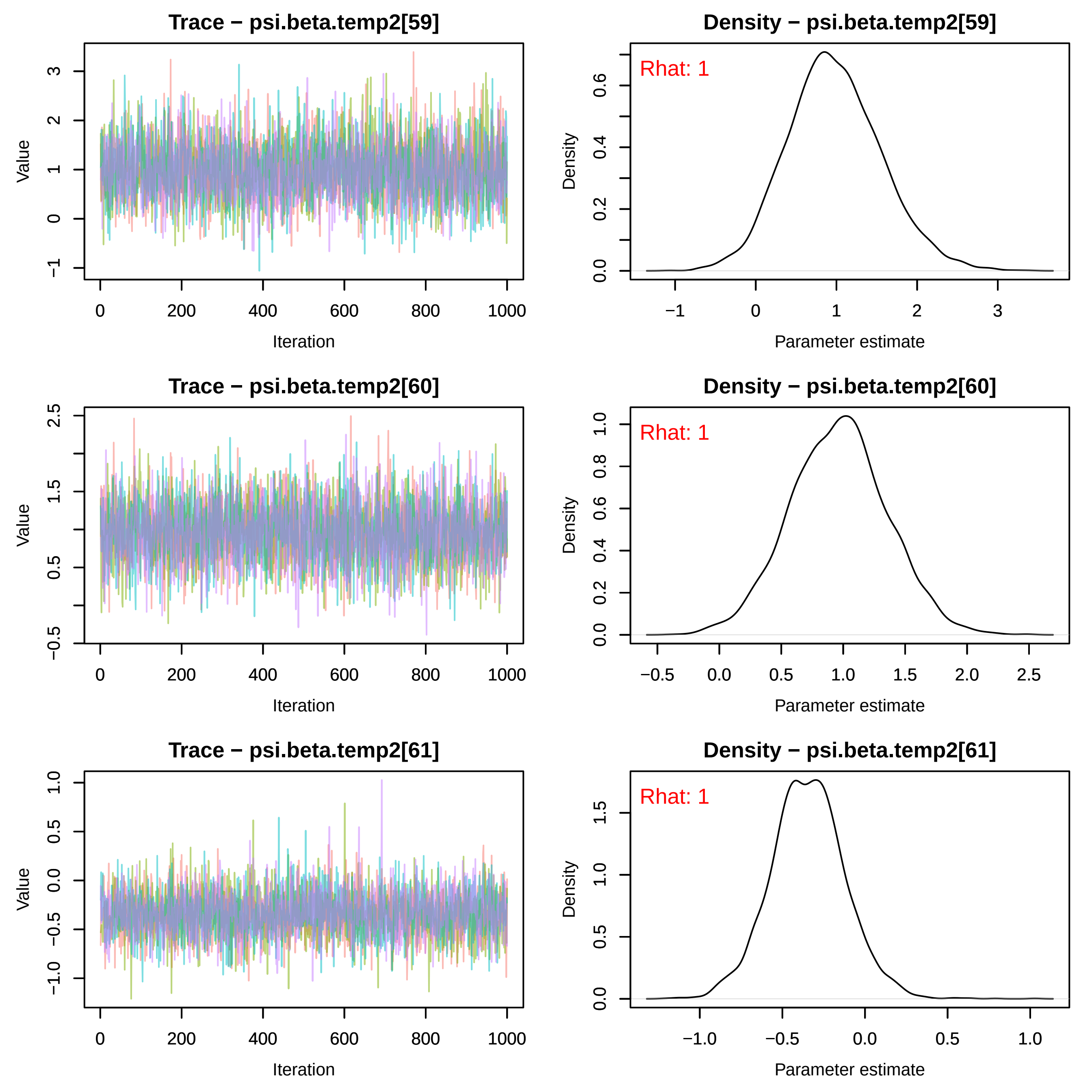

Trace − psi.beta.temp2[59]
Value
Iteration
Density − psi.beta.temp2[59]
Rhat: 1
Density
Parameter estimate
Trace − psi.beta.temp2[60]
Value
Iteration
Density − psi.beta.temp2[60]
Rhat: 1
Density
Parameter estimate
Trace − psi.beta.temp2[61]
Value
Iteration
Density − psi.beta.temp2[61]
Rhat: 1
Density
Parameter estimate

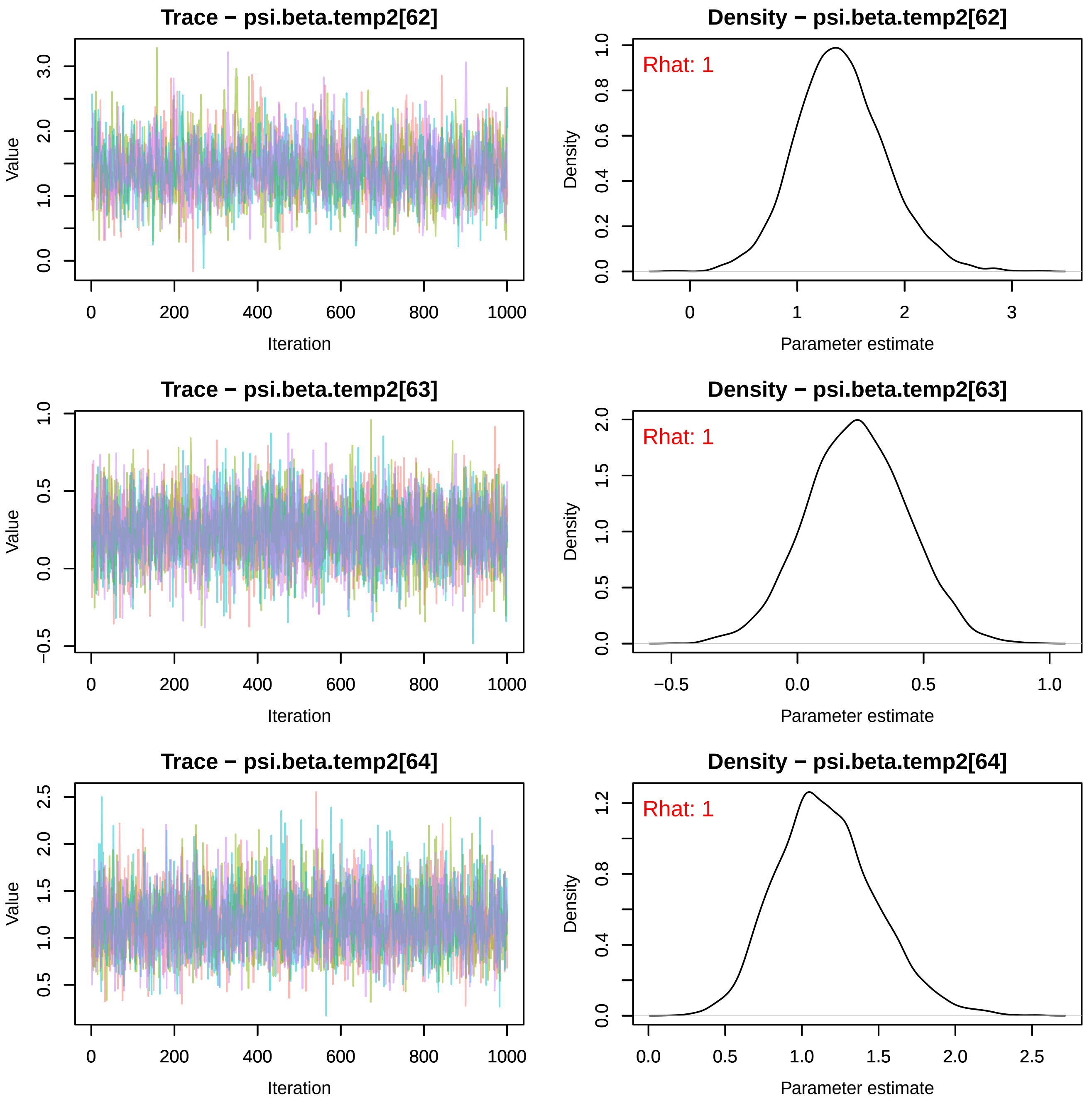
Trace − psi.beta.temp2[62]
Density − psi.beta.temp2[62]
Rhat: 1
Trace − psi.beta.temp2[63]
Density − psi.beta.temp2[63]
Rhat: 1
Trace − psi.beta.temp2[64]
Density − psi.beta.temp2[64]
Rhat: 1
Value
Iteration
Density
Parameter estimate

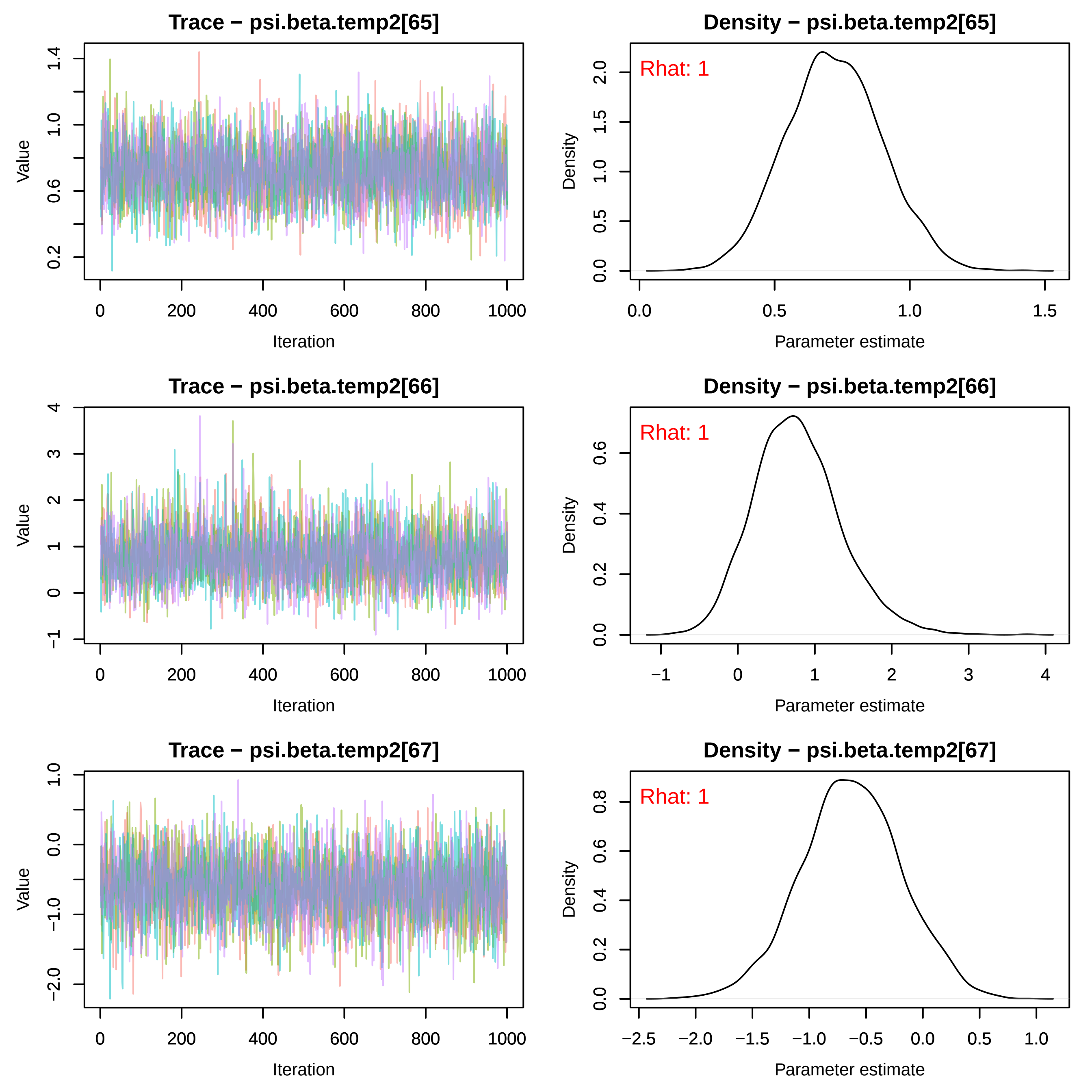

Trace − psi.beta.temp2[65]
Value
Iteration
Density − psi.beta.temp2[65]
Rhat: 1
Density
Parameter estimate
Trace − psi.beta.temp2[66]
Value
Iteration
Density − psi.beta.temp2[66]
Rhat: 1
Density
Parameter estimate
Trace − psi.beta.temp2[67]
Value
Iteration
Density − psi.beta.temp2[67]
Rhat: 1
Density
Parameter estimate

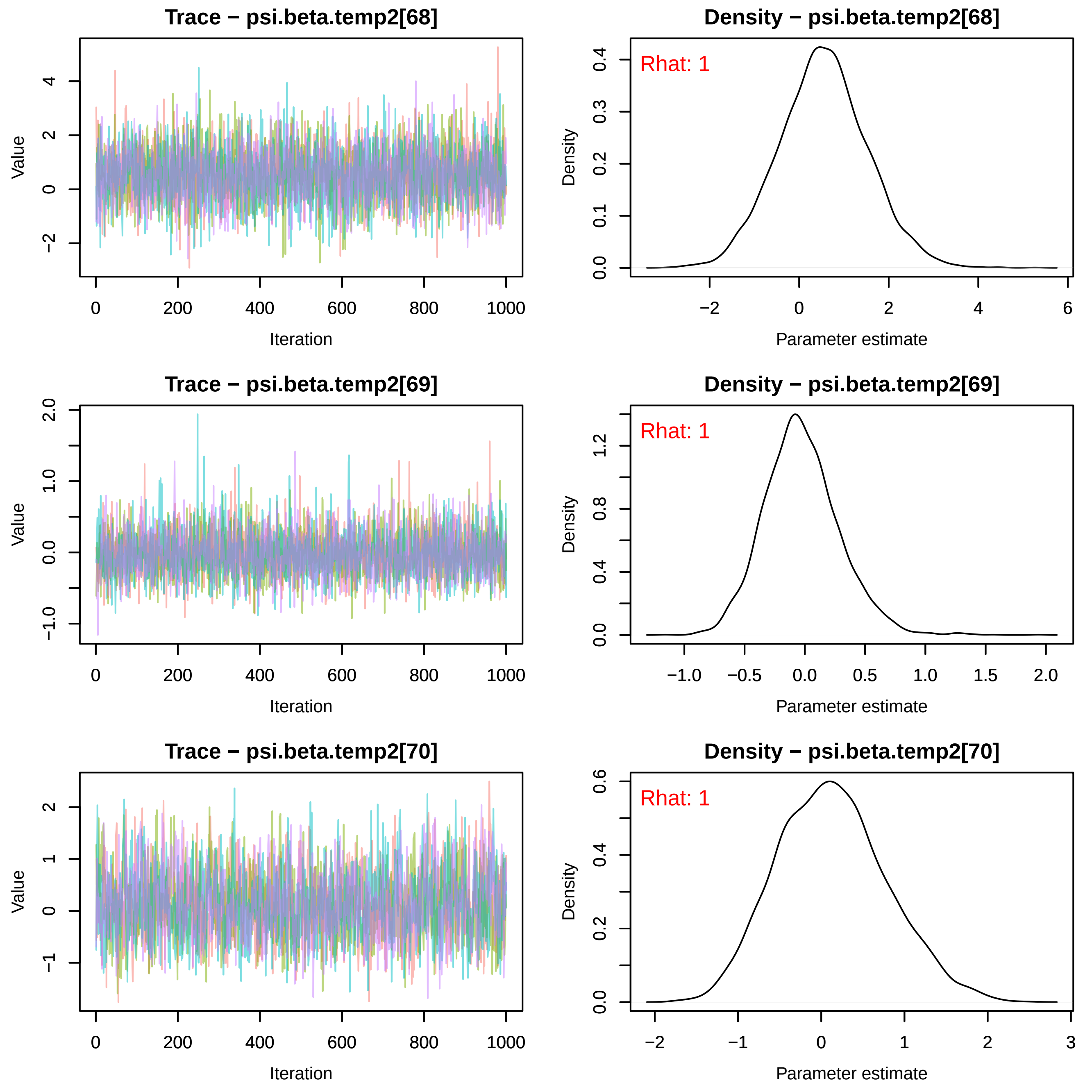
Trace − psi.beta.temp2[68]
Density − psi.beta.temp2[68]
Rhat: 1
Value
Iteration
Density
Parameter estimate
Trace − psi.beta.temp2[69]
Density − psi.beta.temp2[69]
Rhat: 1
Value
Iteration
Density
Parameter estimate
Trace − psi.beta.temp2[70]
Density − psi.beta.temp2[70]
Rhat: 1
Value
Iteration
Density
Parameter estimate

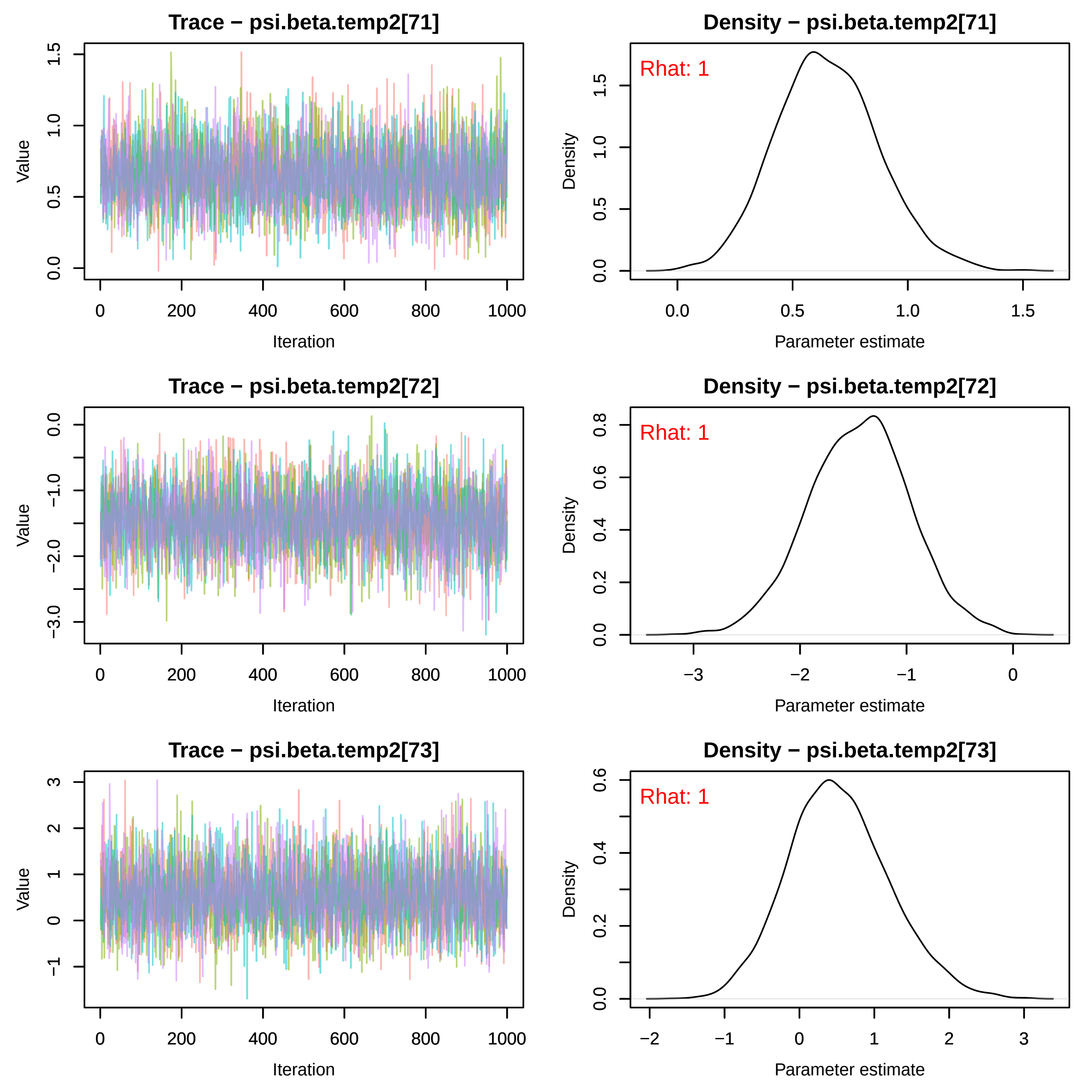

Trace − psi.beta.temp2[71]
Value
Iteration
0.0
0.5
1.0
1.5
0
200
400
600
800
1000
Density − psi.beta.temp2[71]
Rhat: 1
Density
Parameter estimate
0.0
0.5
1.0
1.5
Trace − psi.beta.temp2[72]
Value
Iteration
0.0
−1.0
−2.0
−3.0
0
200
400
600
800
1000
Density − psi.beta.temp2[72]
Rhat: 1
Density
Parameter estimate
0.0
0.2
0.4
0.6
0.8
−3
−2
−1
0
Trace − psi.beta.temp2[73]
Value
Iteration
−1
0
1
2
3
0
200
400
600
800
1000
Density − psi.beta.temp2[73]
Rhat: 1
Density
Parameter estimate
0.0
0.2
0.4
0.6
−2
−1
0
1
2
3

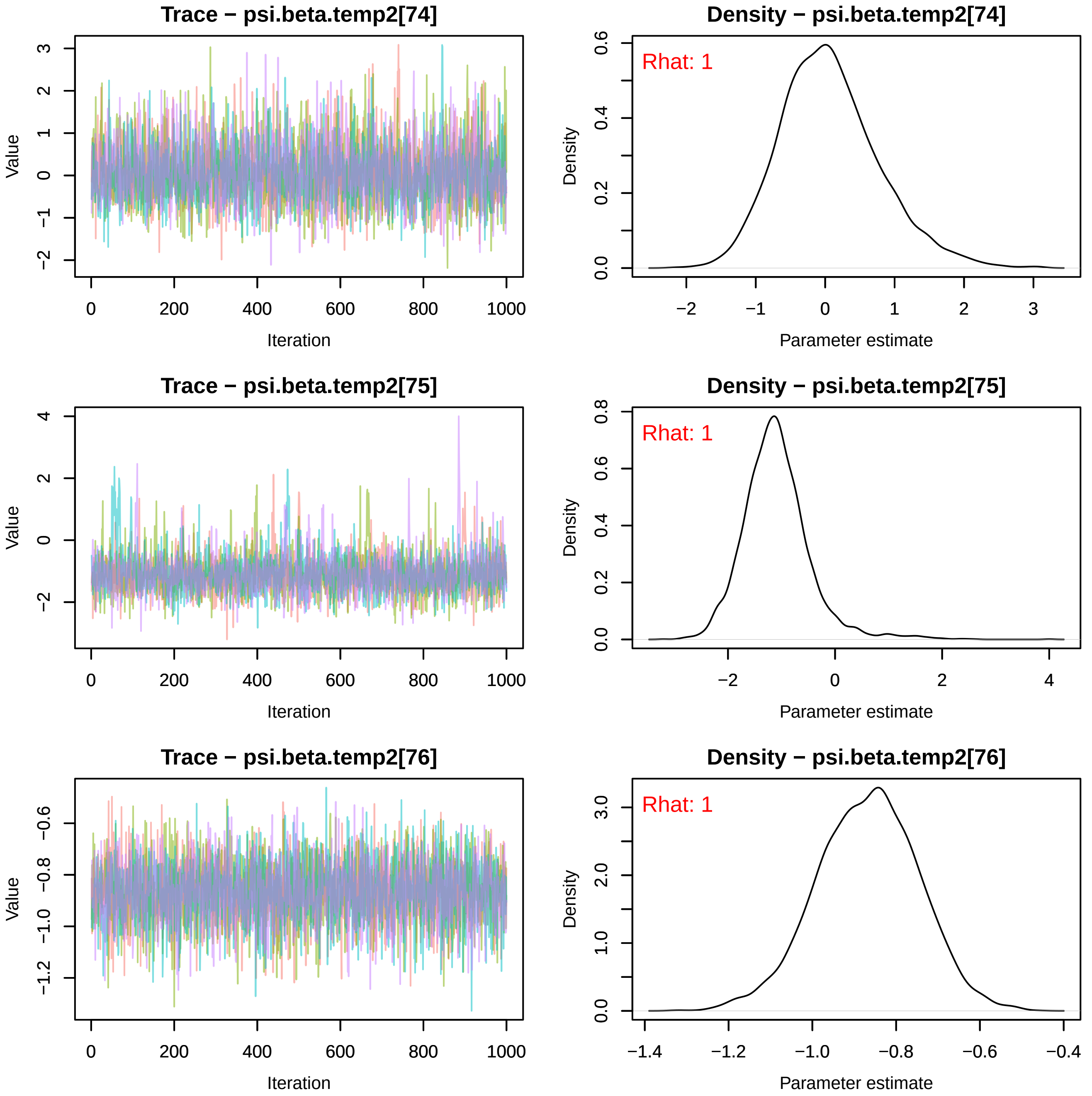
Trace − psi.beta.temp2[74]
Value
Iteration
0
200
400
600
800
1000
−2
−1
0
1
2
3
Density − psi.beta.temp2[74]
Rhat: 1
Density
Parameter estimate
−2
−1
0
1
2
3
0.0
0.2
0.4
0.6
Trace − psi.beta.temp2[75]
Value
Iteration
0
200
400
600
800
1000
−2
0
2
4
Density − psi.beta.temp2[75]
Rhat: 1
Density
Parameter estimate
−2
0
2
4
0.0
0.2
0.4
0.6
0.8
Trace − psi.beta.temp2[76]
Value
Iteration
0
200
400
600
800
1000
−1.2
−1.0
−0.8
−0.6
Density − psi.beta.temp2[76]
Rhat: 1
Density
Parameter estimate
−1.4
−1.2
−1.0
−0.8
−0.6
−0.4
0.0
1.0
2.0
3.0

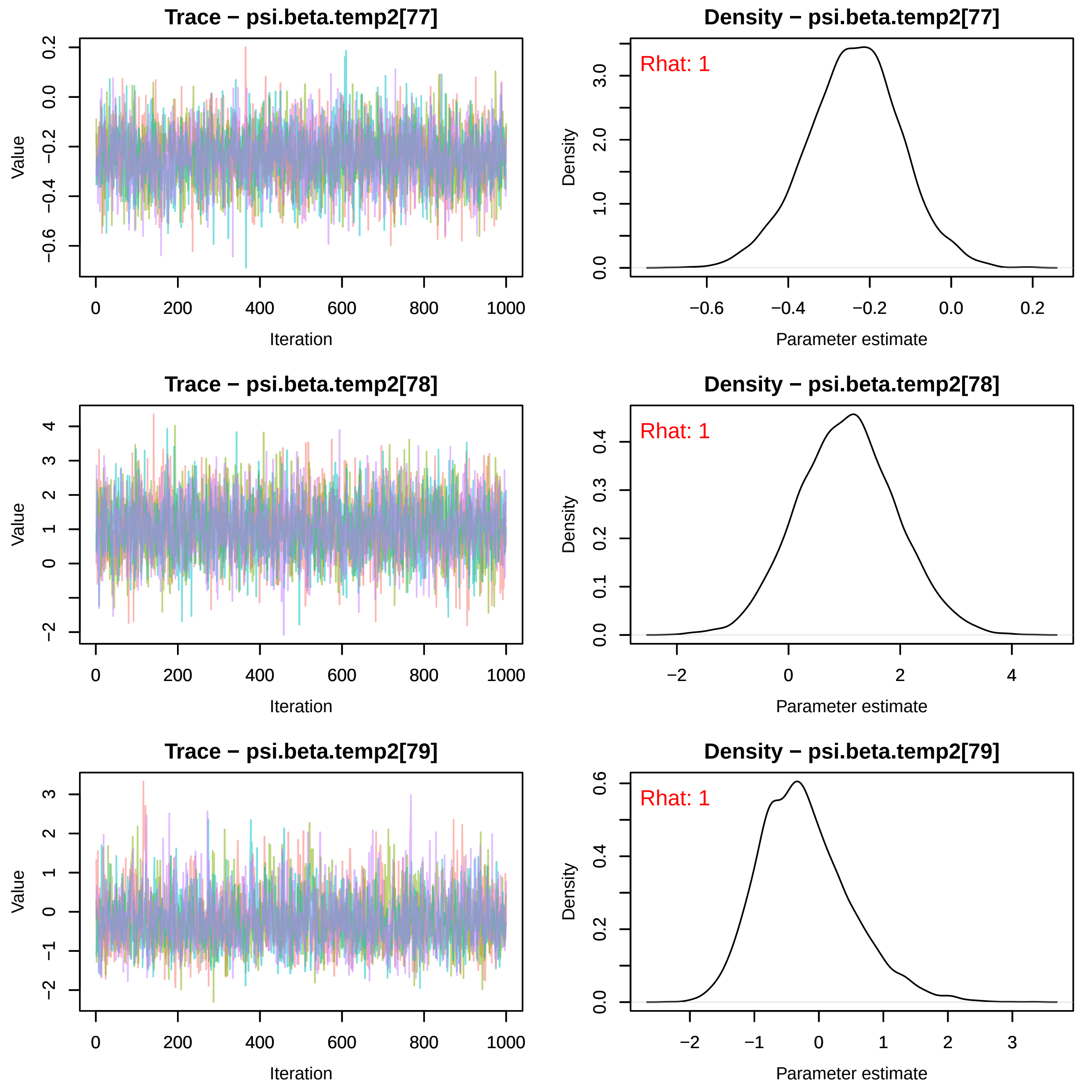
Trace − psi.beta.temp2[77]
Value
Iteration
0 200 400 600 800 1000
0.2 0.0 −0.2 −0.4 −0.6
Density − psi.beta.temp2[77]
Rhat: 1
Density
3.0 2.0 1.0 0.0
−0.6 −0.4 −0.2 0.0 0.2
Parameter estimate
Trace − psi.beta.temp2[78]
Value
4 3 2 1 0 −2
Iteration
0 200 400 600 800 1000
Density − psi.beta.temp2[78]
Rhat: 1
Density
0.4 0.3 0.2 0.1 0.0
−2 0 2 4
Parameter estimate
Trace − psi.beta.temp2[79]
Value
3 2 1 0 −1 −2
Iteration
0 200 400 600 800 1000
Density − psi.beta.temp2[79]
Rhat: 1
Density
0.6 0.4 0.2 0.0
−2 −1 0 1 2 3
Parameter estimate

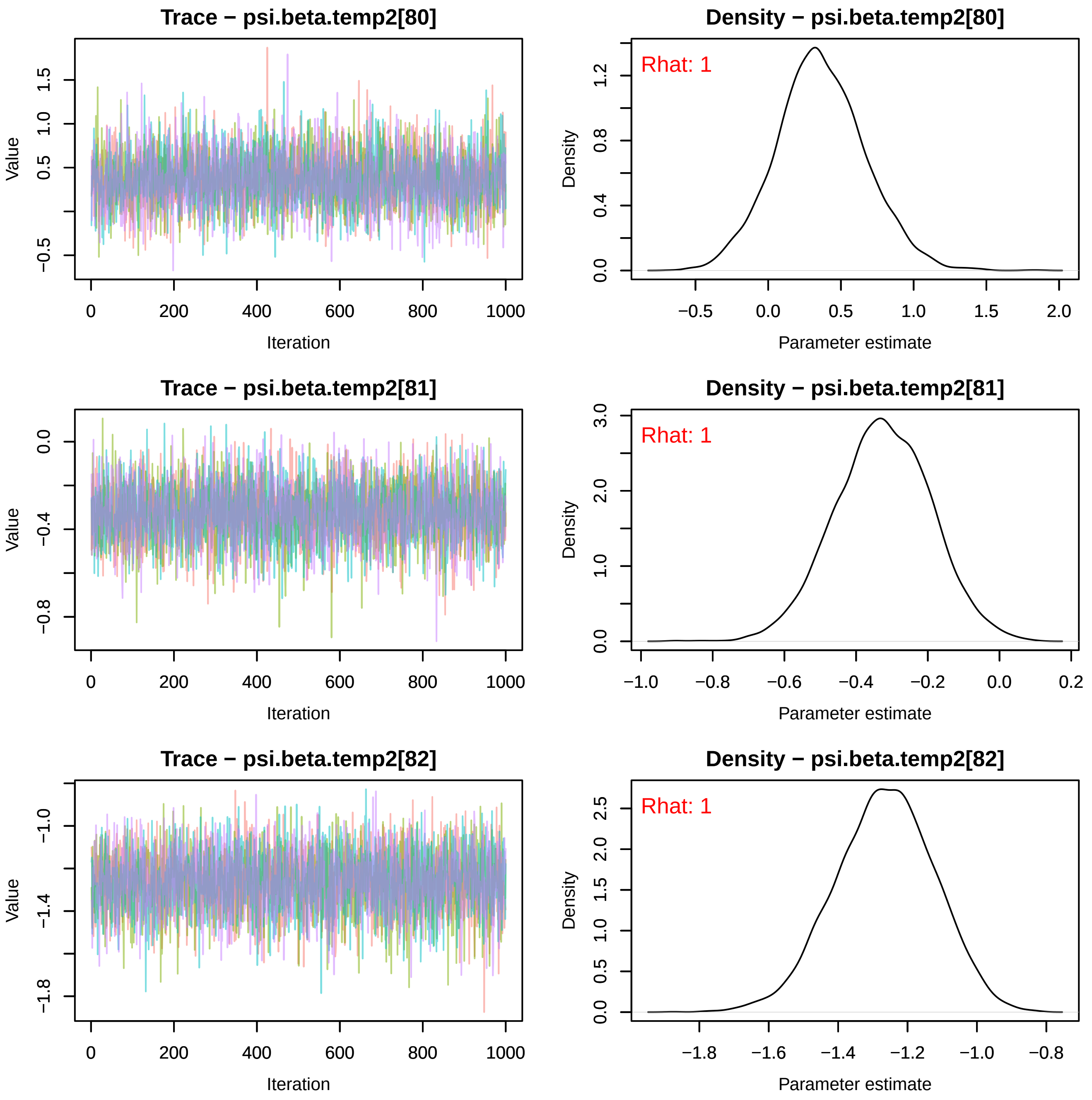
Trace − psi.beta.temp2[80]
Value
Iteration
Density − psi.beta.temp2[80]
Rhat: 1
Density
Parameter estimate
Trace − psi.beta.temp2[81]
Value
Iteration
Density − psi.beta.temp2[81]
Rhat: 1
Density
Parameter estimate
Trace − psi.beta.temp2[82]
Value
Iteration
Density − psi.beta.temp2[82]
Rhat: 1
Density
Parameter estimate

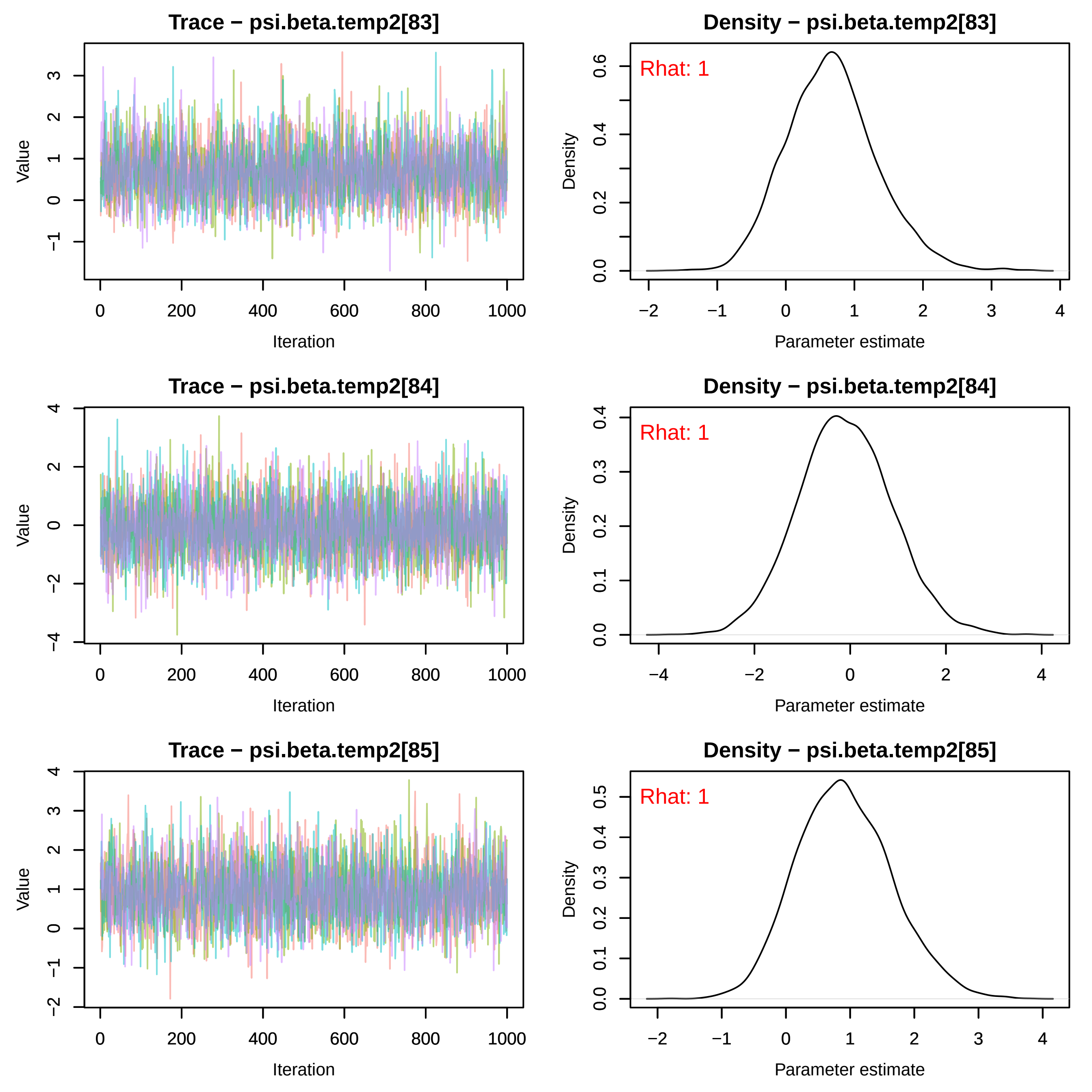

Trace − psi.beta.temp2[83]
Density − psi.beta.temp2[83]
Rhat: 1
Trace − psi.beta.temp2[84]
Density − psi.beta.temp2[84]
Rhat: 1
Trace − psi.beta.temp2[85]
Density − psi.beta.temp2[85]
Rhat: 1
Value
Iteration
Density
Parameter estimate

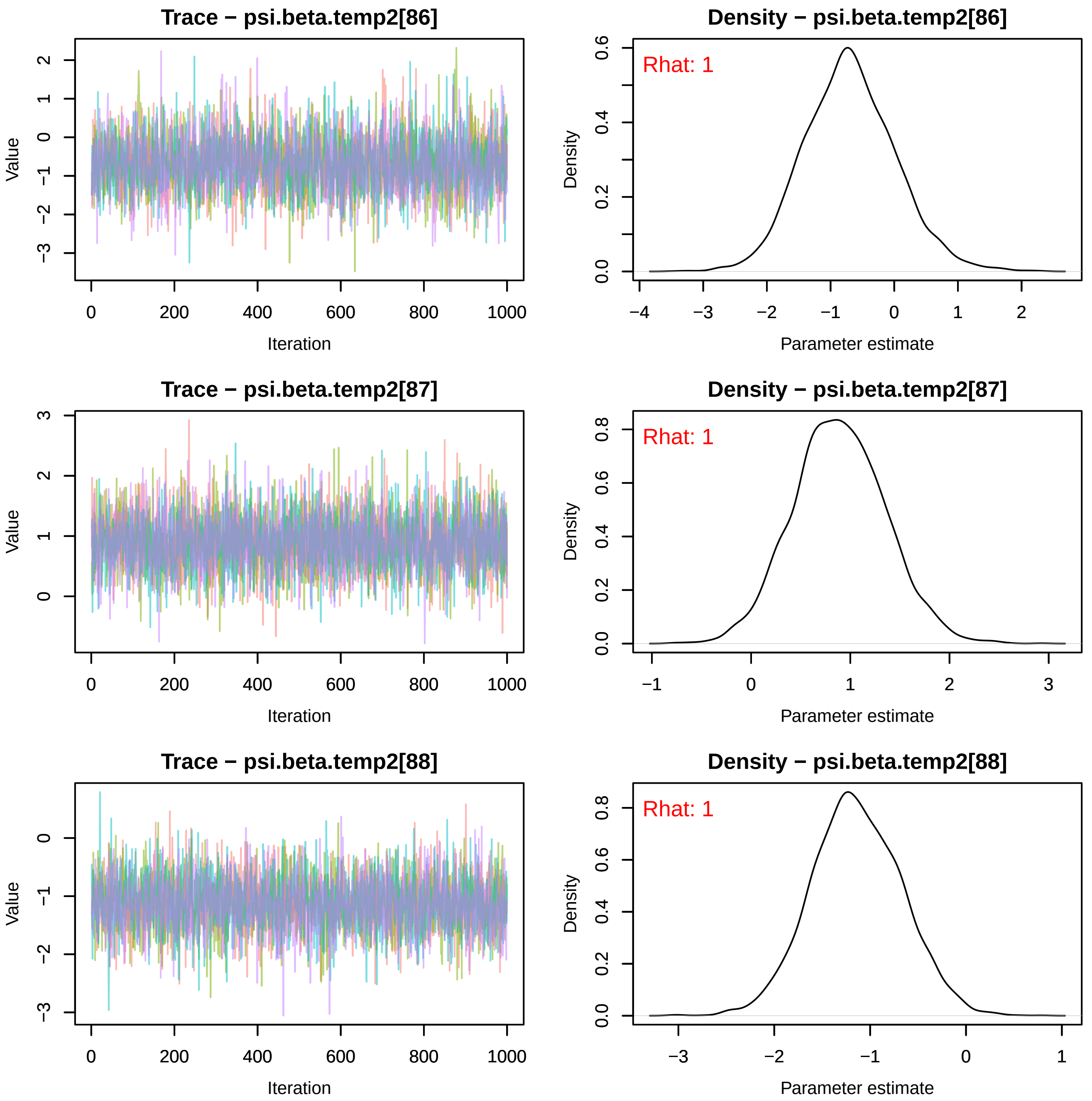
Trace − psi.beta.temp2[86]
Density − psi.beta.temp2[86]
Rhat: 1
Trace − psi.beta.temp2[87]
Density − psi.beta.temp2[87]
Rhat: 1
Trace − psi.beta.temp2[88]
Density − psi.beta.temp2[88]
Rhat: 1
Value
Iteration
Density
Parameter estimate

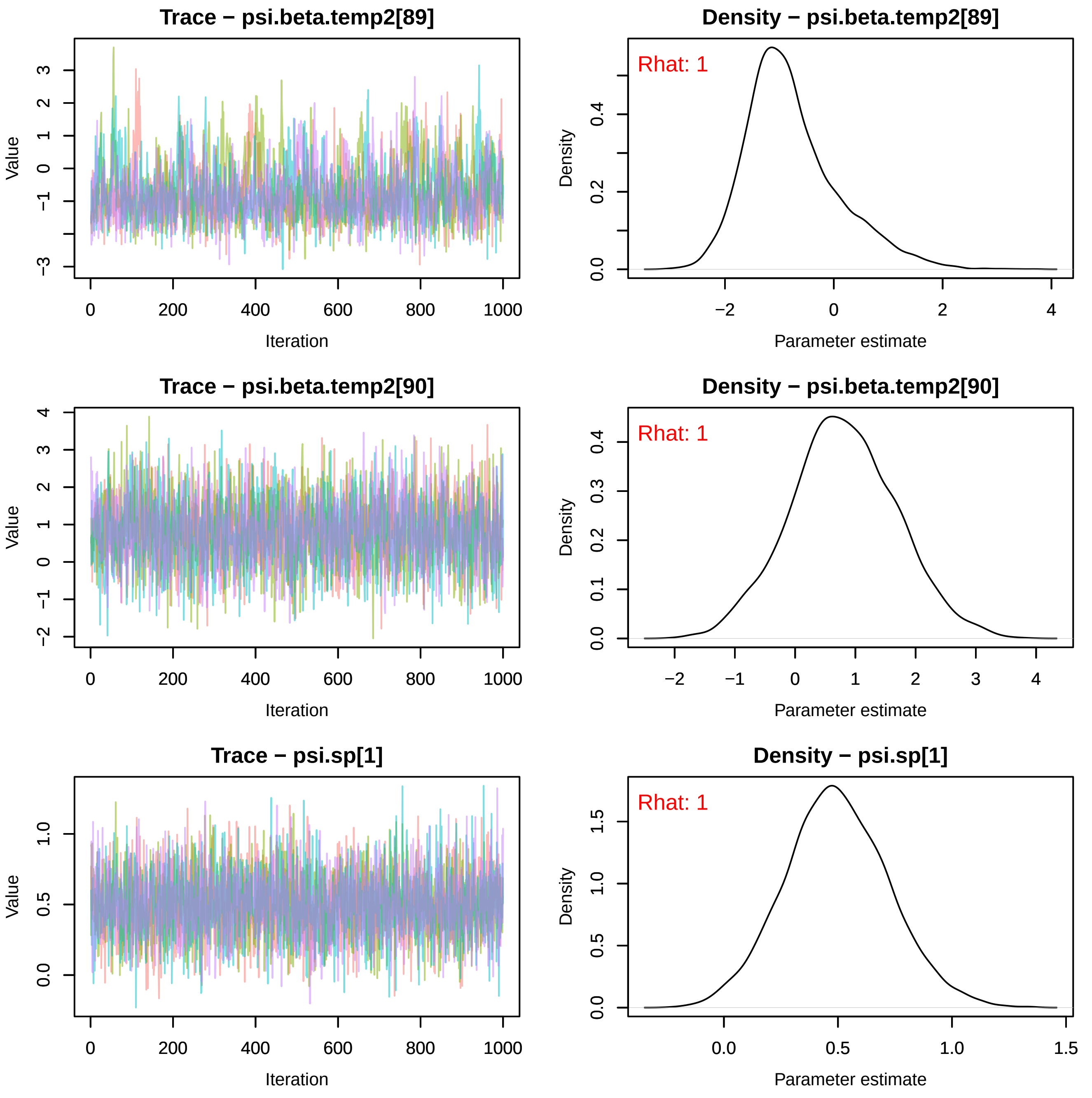

Trace − psi.beta.temp2[89]
Value
Iteration
0
200
400
600
800
1000
3
2
1
0
−1
−3
Density − psi.beta.temp2[89]
Rhat: 1
Density
Parameter estimate
0.0
0.2
0.4
−2
0
2
4
Trace − psi.beta.temp2[90]
Value
Iteration
4
3
2
1
0
−1
−2
0
200
400
600
800
1000
Density − psi.beta.temp2[90]
Rhat: 1
Density
Parameter estimate
0.0
0.1
0.2
0.3
0.4
−2
−1
0
1
2
3
4
Trace − psi.sp[1]
Value
Iteration
0.0
0.5
1.0
0
200
400
600
800
1000
Density − psi.sp[1]
Rhat: 1
Density
Parameter estimate
0.0
0.5
1.0
1.5
0.0
0.5
1.0
1.5

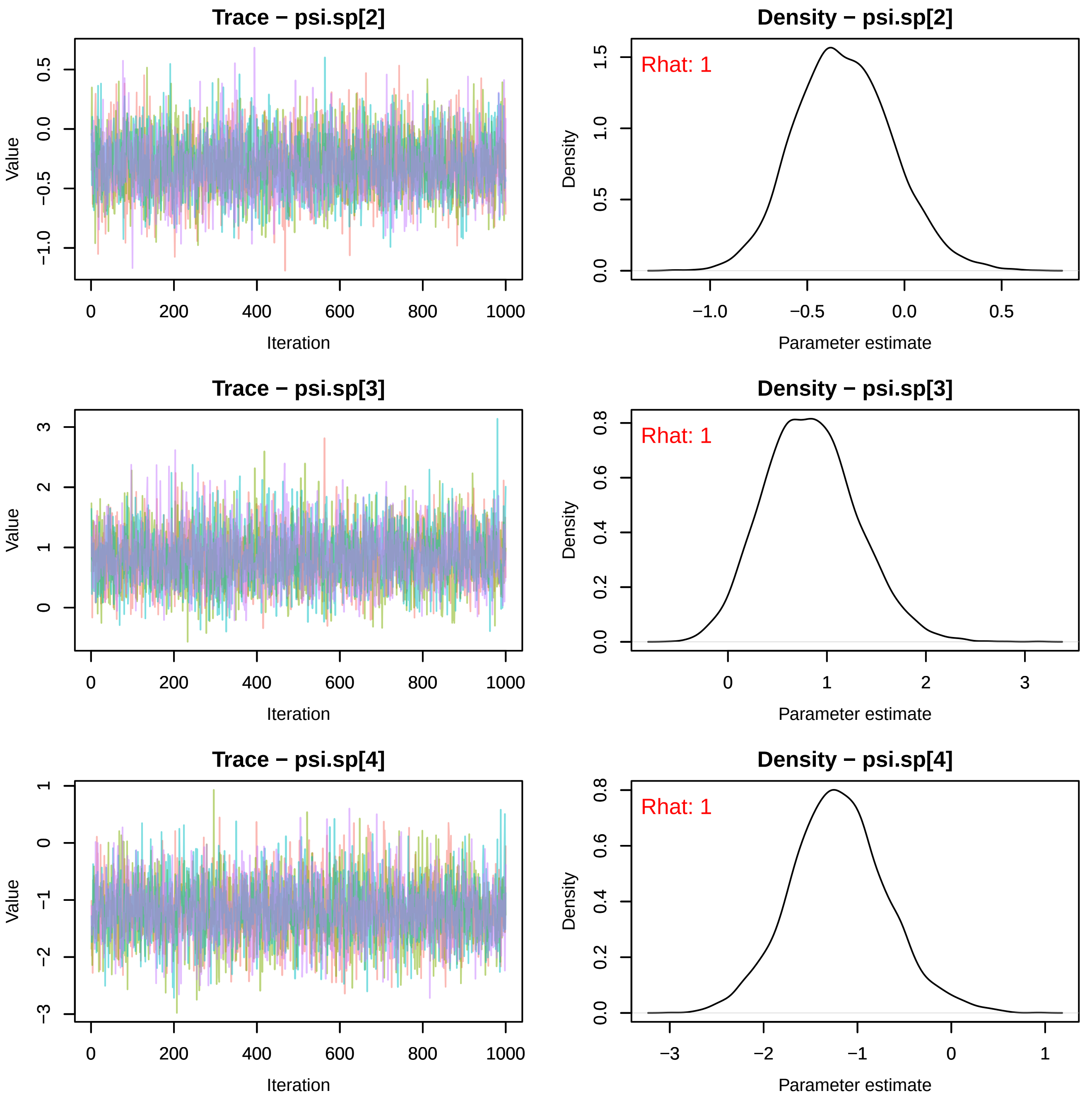
Trace − psi.sp[2]
Value
Iteration
Density − psi.sp[2]
Rhat: 1
Density
Parameter estimate
Trace − psi.sp[3]
Value
Iteration
Density − psi.sp[3]
Rhat: 1
Density
Parameter estimate
Trace − psi.sp[4]
Value
Iteration
Density − psi.sp[4]
Rhat: 1
Density
Parameter estimate

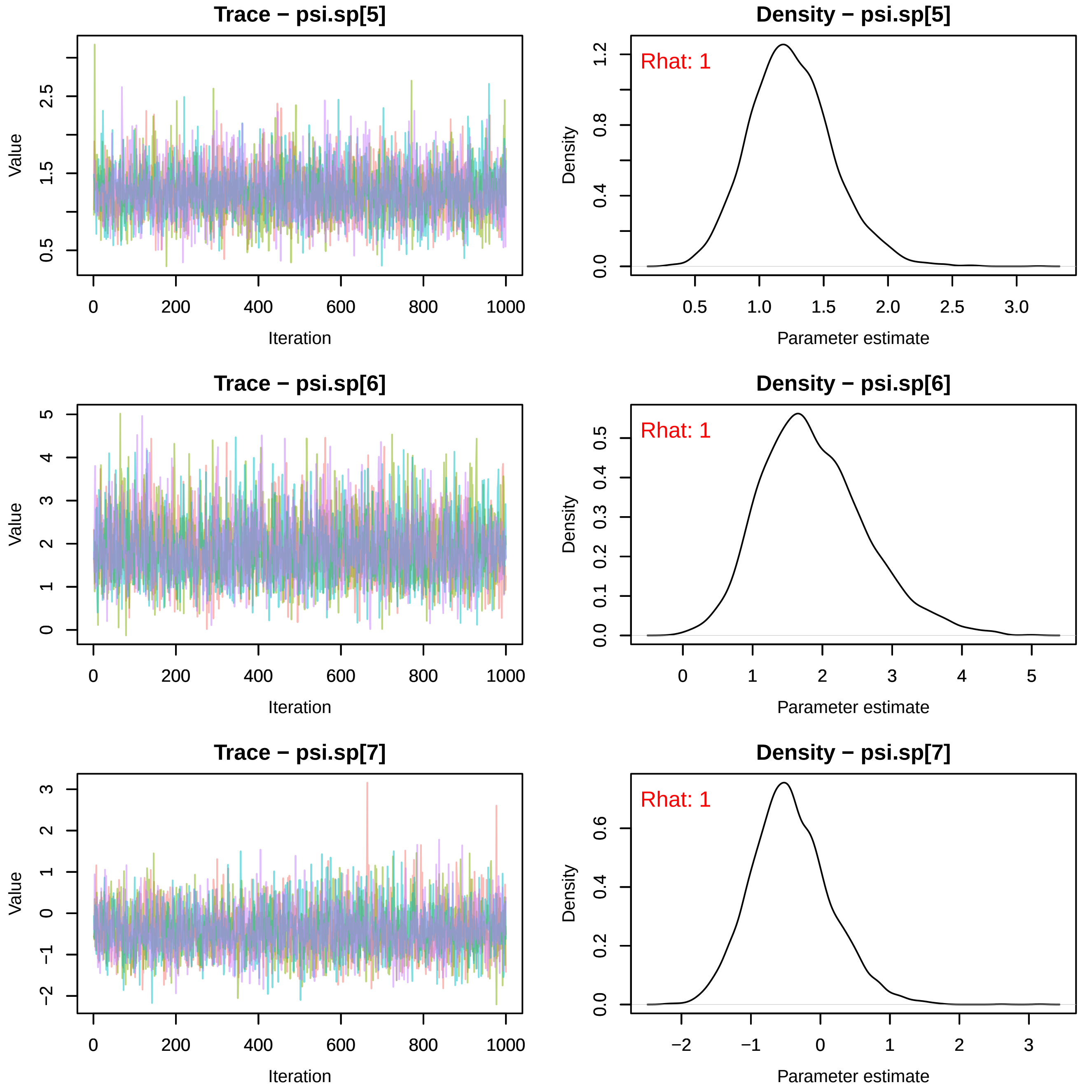
Trace − psi.sp[5]
Value
Iteration
0.5
1.5
2.5
0
200
400
600
800
1000
Density − psi.sp[5]
Rhat: 1
Density
Parameter estimate
0.0
0.4
0.8
1.2
0.5
1.0
1.5
2.0
2.5
3.0
Trace − psi.sp[6]
Value
Iteration
0
1
2
3
4
5
0
200
400
600
800
1000
Density − psi.sp[6]
Rhat: 1
Density
Parameter estimate
0.0
0.1
0.2
0.3
0.4
0.5
0
1
2
3
4
5
Trace − psi.sp[7]
Value
Iteration
−2
−1
0
1
2
3
0
200
400
600
800
1000
Density − psi.sp[7]
Rhat: 1
Density
Parameter estimate
0.0
0.2
0.4
0.6
−2
−1
0
1
2
3

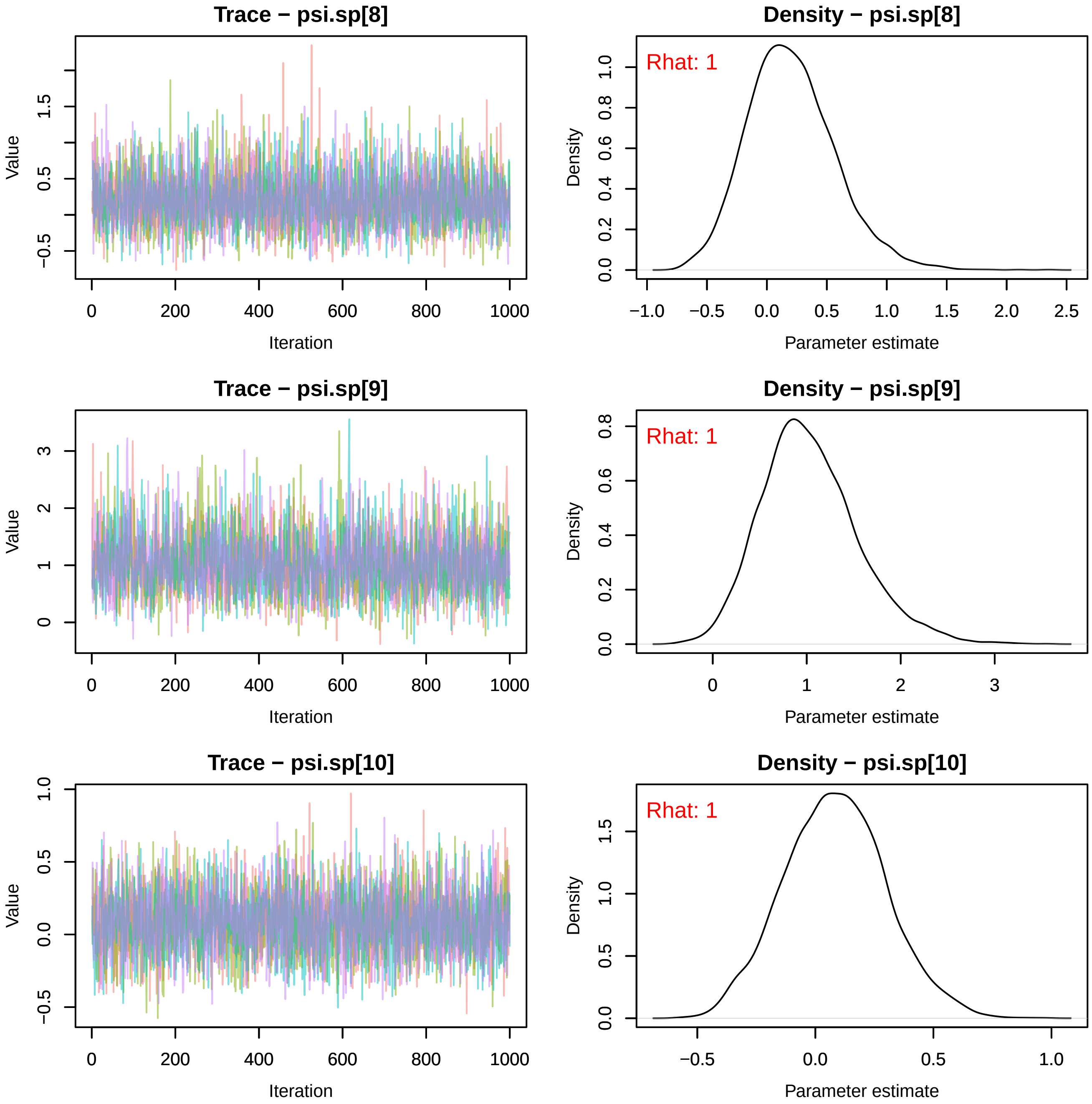
Trace − psi.sp[8]
Value
Iteration
0
200
400
600
800
1000
−0.5
0.5
1.5
Density − psi.sp[8]
Rhat: 1
Density
Parameter estimate
0.0
0.2
0.4
0.6
0.8
1.0
−1.0
−0.5
0.0
0.5
1.0
1.5
2.0
2.5
Trace − psi.sp[9]
Value
Iteration
0
1
2
3
Density − psi.sp[9]
Rhat: 1
Density
Parameter estimate
0.0
0.2
0.4
0.6
0.8
0
1
2
3
Trace − psi.sp[10]
Value
Iteration
−0.5
0.0
0.5
1.0
Density − psi.sp[10]
Rhat: 1
Density
Parameter estimate
0.0
0.5
1.0
1.5
−0.5
0.0
0.5
1.0

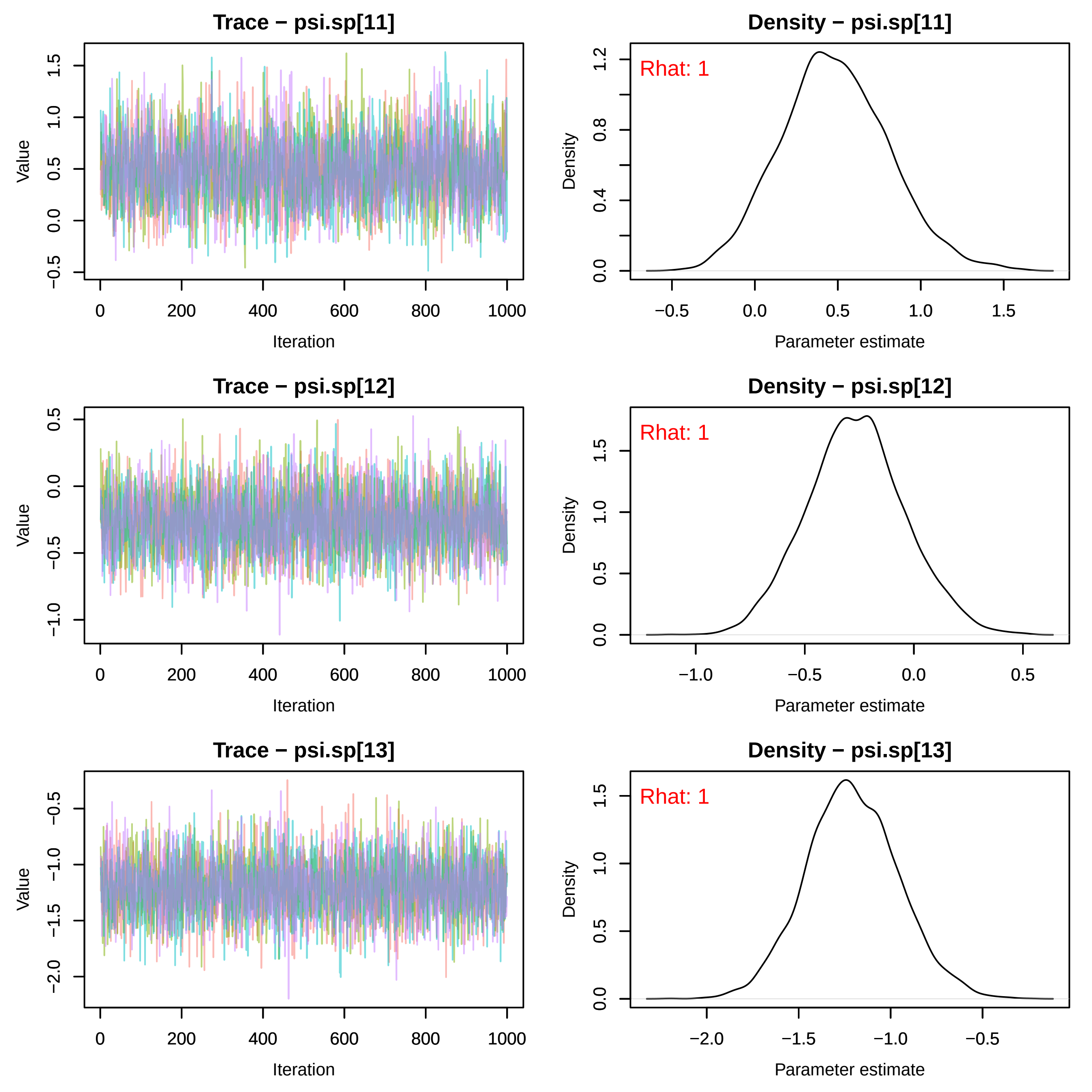
Trace − psi.sp[11]
Value
Iteration
Density − psi.sp[11]
Rhat: 1
Density
Parameter estimate
Trace − psi.sp[12]
Value
Iteration
Density − psi.sp[12]
Rhat: 1
Density
Parameter estimate
Trace − psi.sp[13]
Value
Iteration
Density − psi.sp[13]
Rhat: 1
Density
Parameter estimate

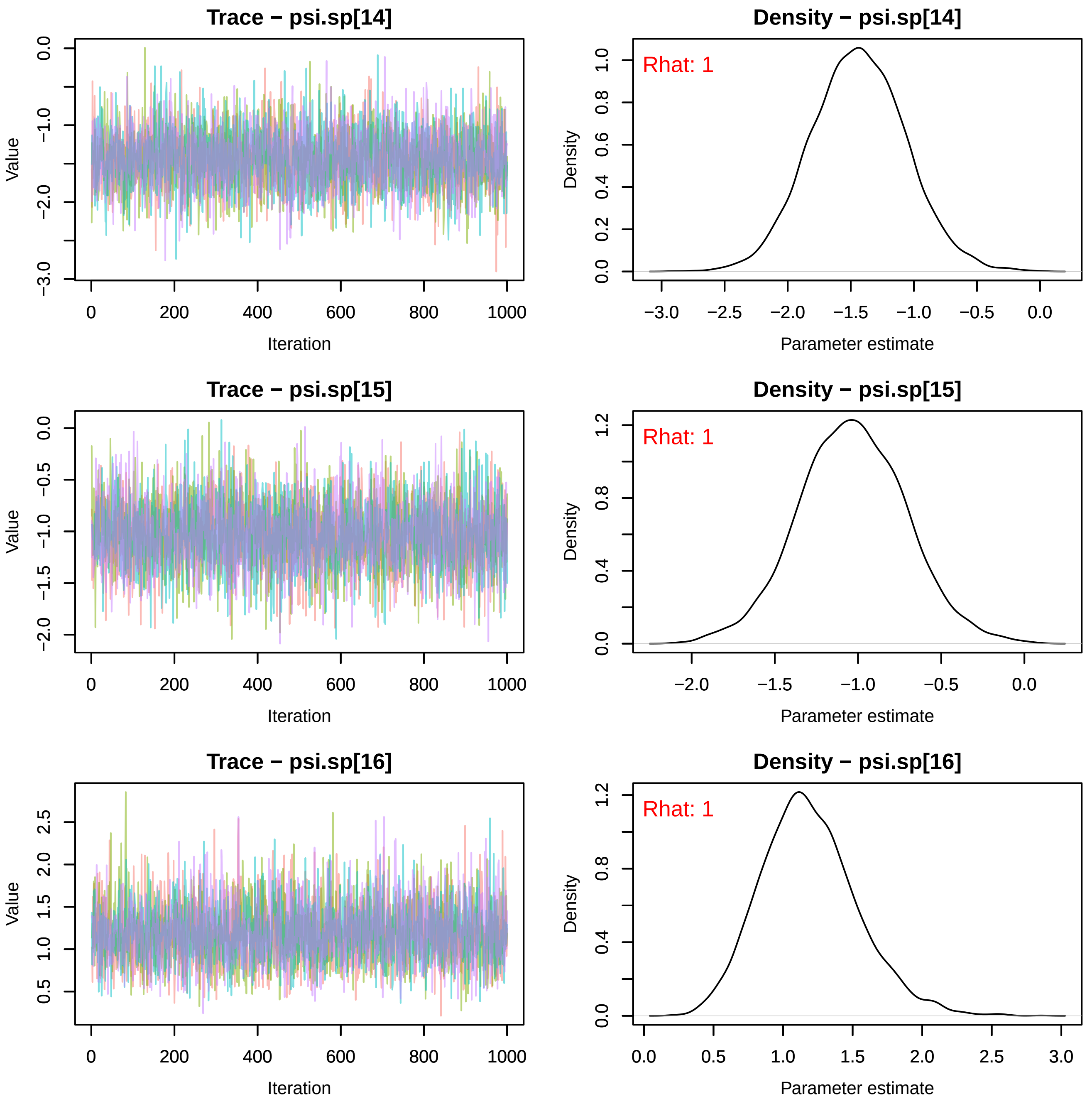
Trace − psi.sp[14]
Value
Iteration
Density − psi.sp[14]
Rhat: 1
Density
Parameter estimate
Trace − psi.sp[15]
Value
Iteration
Density − psi.sp[15]
Rhat: 1
Density
Parameter estimate
Trace − psi.sp[16]
Value
Iteration
Density − psi.sp[16]
Rhat: 1
Density
Parameter estimate

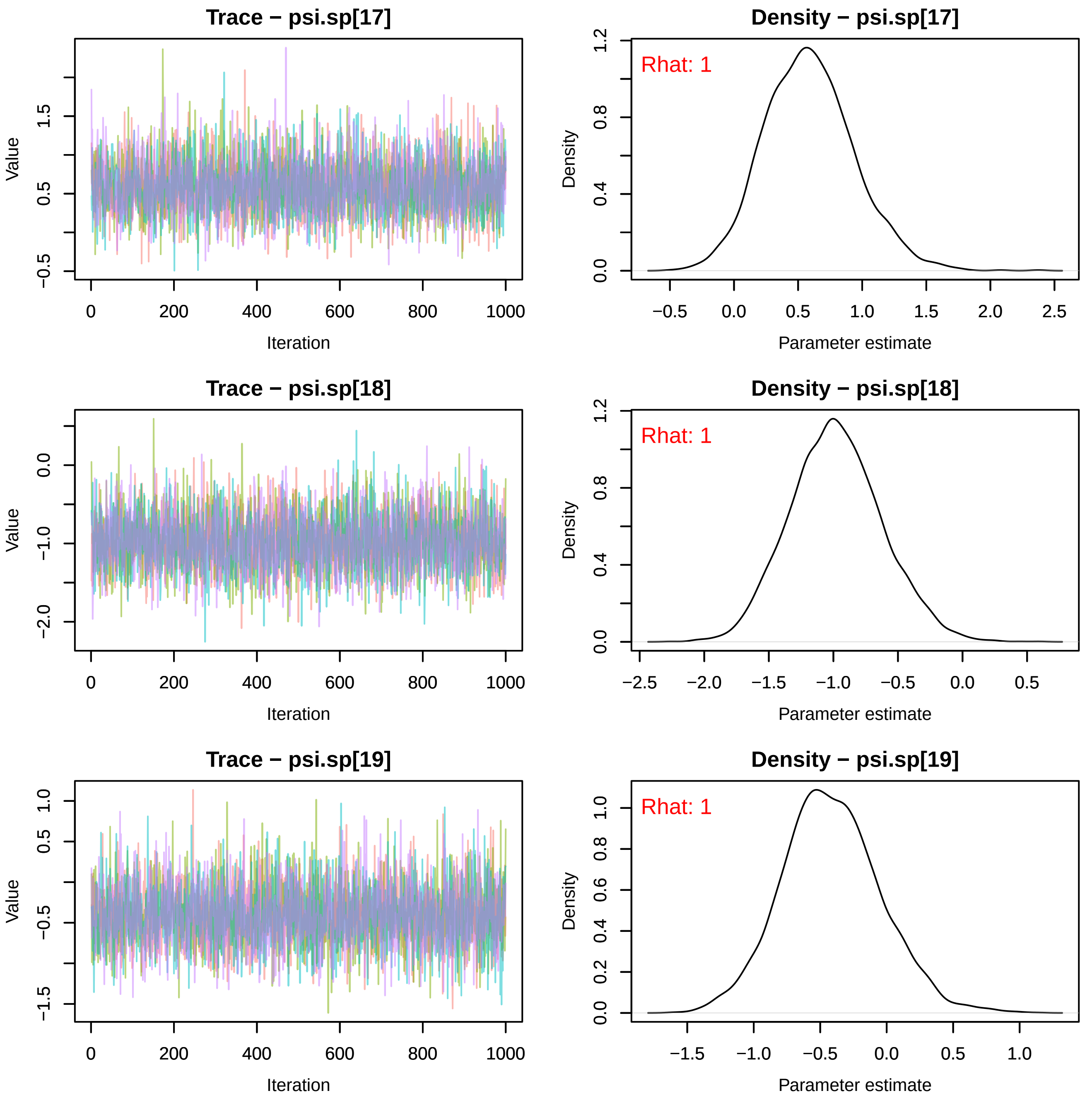

Trace − psi.sp[17]
Density − psi.sp[17]
Rhat: 1
Trace − psi.sp[18]
Density − psi.sp[18]
Rhat: 1
Trace − psi.sp[19]
Density − psi.sp[19]
Rhat: 1
Value
Iteration
Density
Parameter estimate

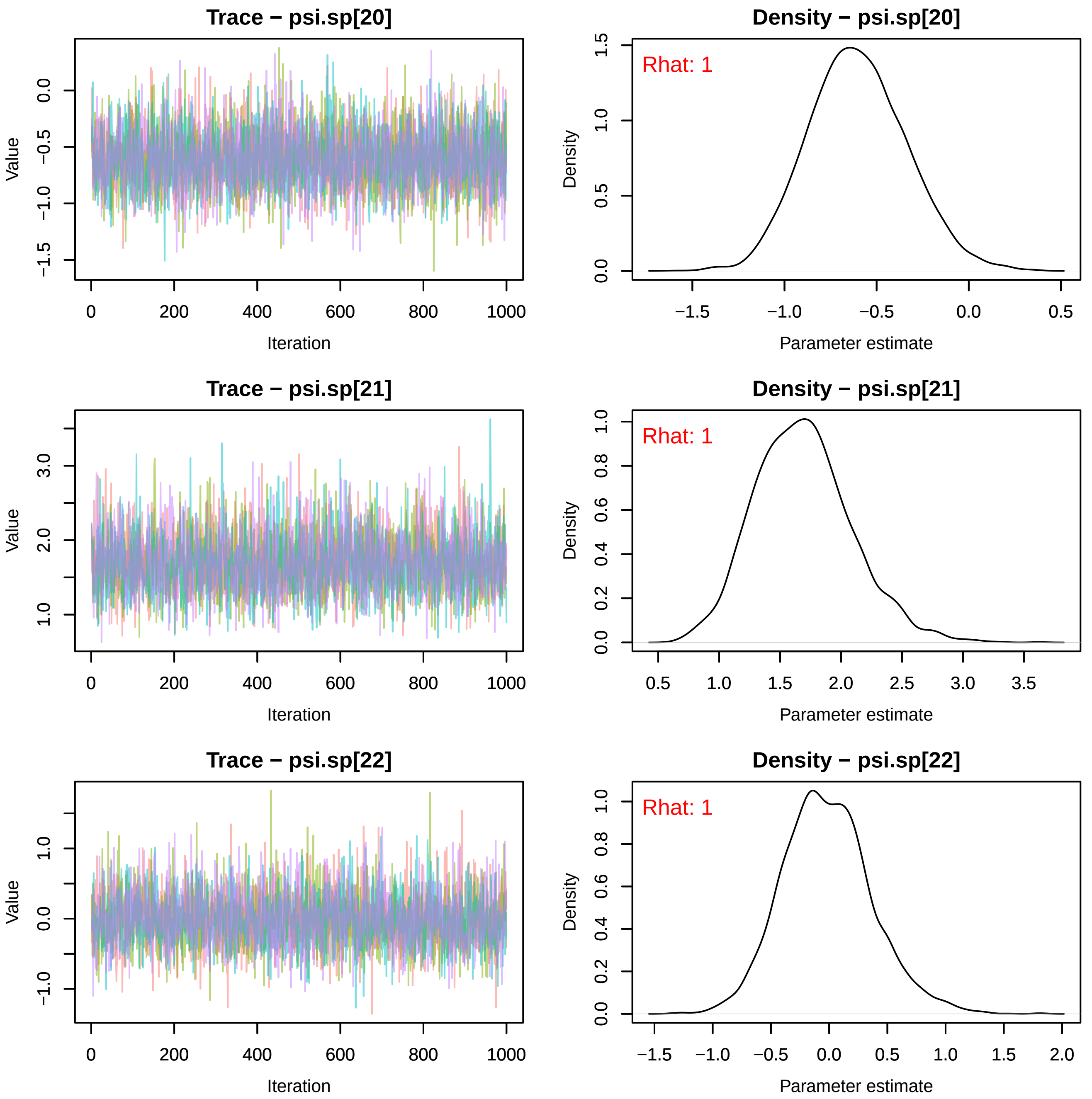
Trace − psi.sp[20]
Density − psi.sp[20]
Rhat: 1
Trace − psi.sp[21]
Density − psi.sp[21]
Rhat: 1
Trace − psi.sp[22]
Density − psi.sp[22]
Rhat: 1
Value
Iteration
Density
Parameter estimate
0
200
400
600
800
1000

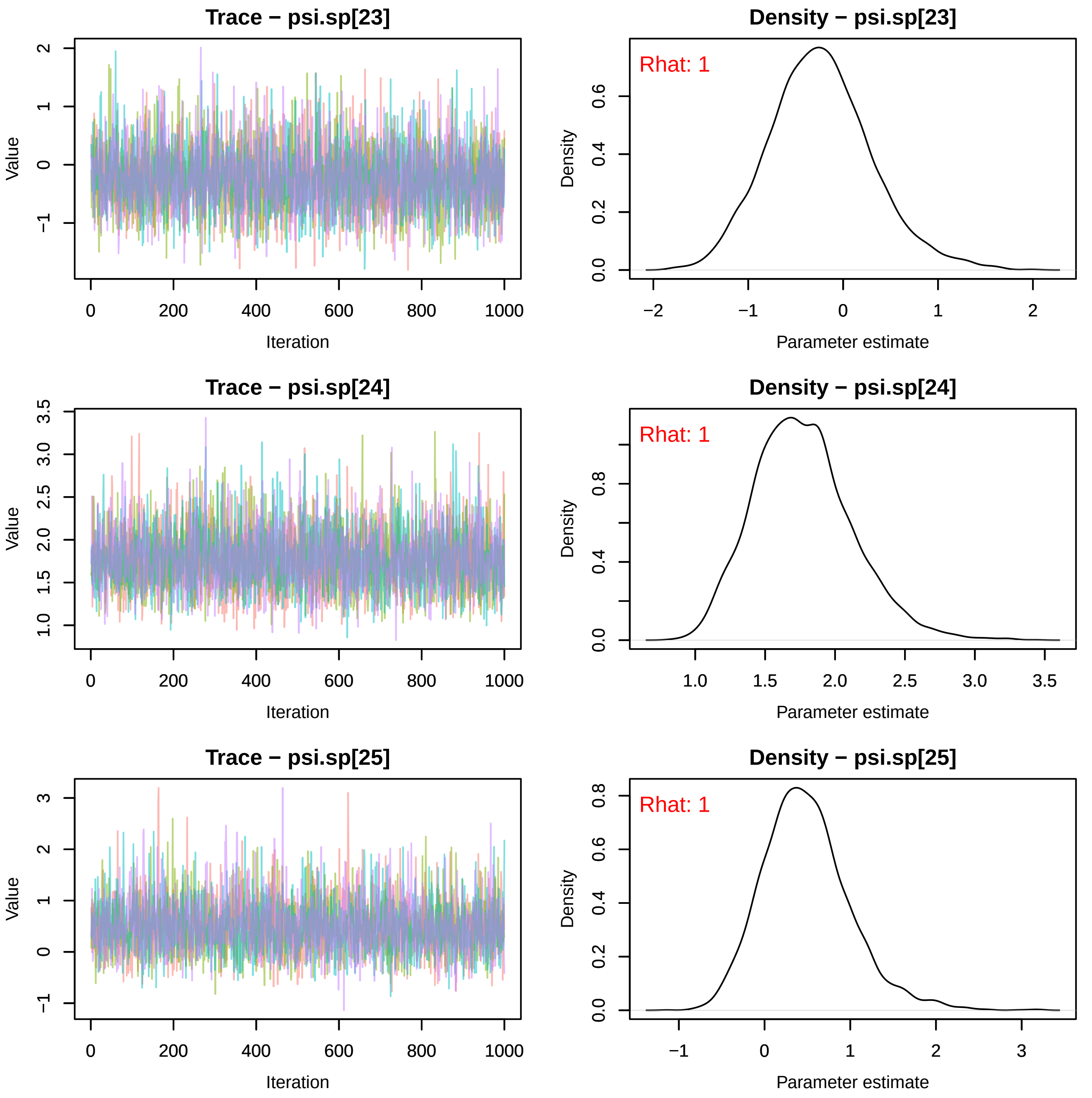

Trace − psi.sp[23]
Density − psi.sp[23]
Rhat: 1
Trace − psi.sp[24]
Density − psi.sp[24]
Rhat: 1
Trace − psi.sp[25]
Density − psi.sp[25]
Rhat: 1
Value
Iteration
Density
Parameter estimate

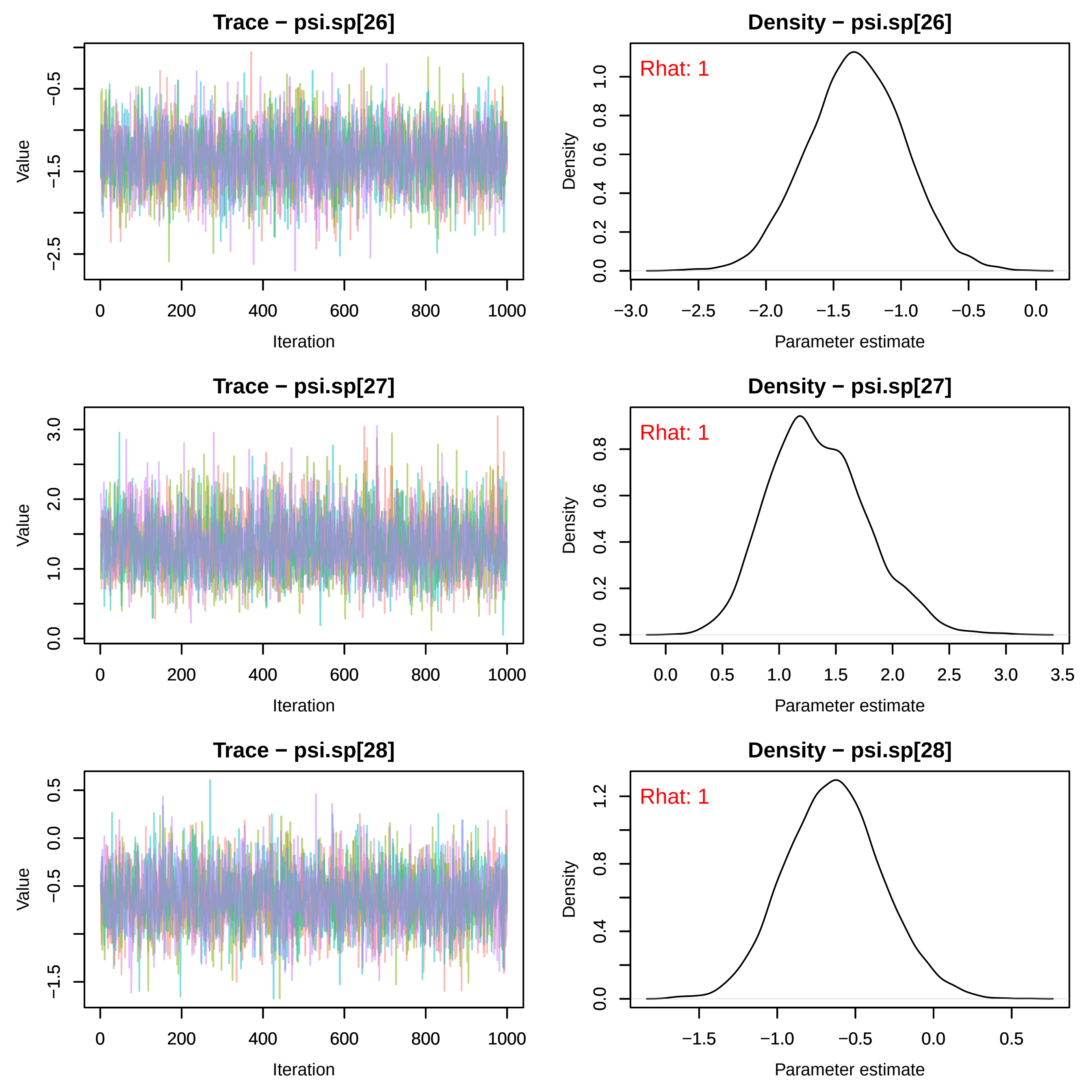
Trace − psi.sp[26]
Value
Iteration
Density − psi.sp[26]
Rhat: 1
Density
Parameter estimate
Trace − psi.sp[27]
Value
Iteration
Density − psi.sp[27]
Rhat: 1
Density
Parameter estimate
Trace − psi.sp[28]
Value
Iteration
Density − psi.sp[28]
Rhat: 1
Density
Parameter estimate

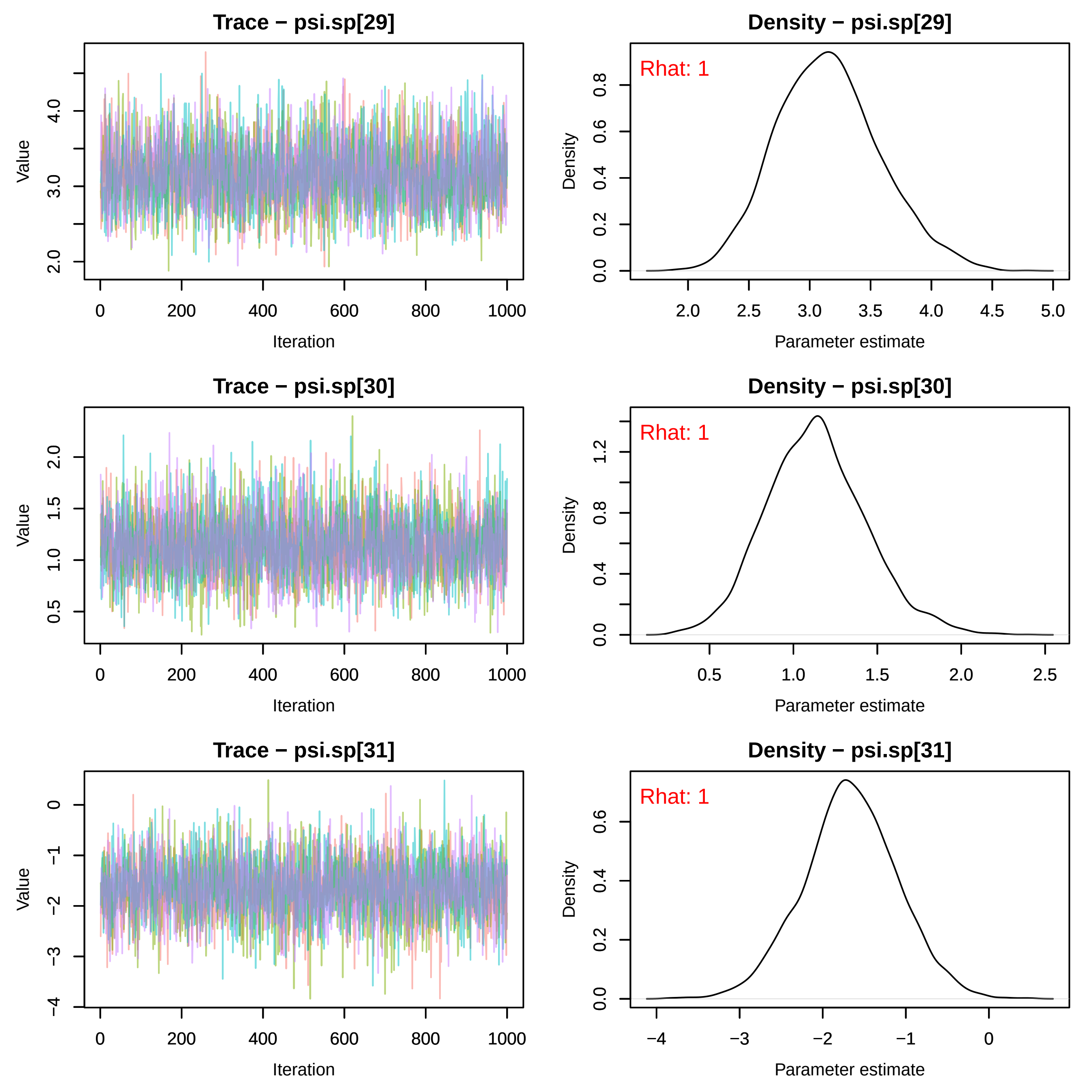
Trace − psi.sp[29]
Value
Iteration
Density − psi.sp[29]
Rhat: 1
Density
Parameter estimate
Trace − psi.sp[30]
Value
Iteration
Density − psi.sp[30]
Rhat: 1
Density
Parameter estimate
Trace − psi.sp[31]
Value
Iteration
Density − psi.sp[31]
Rhat: 1
Density
Parameter estimate

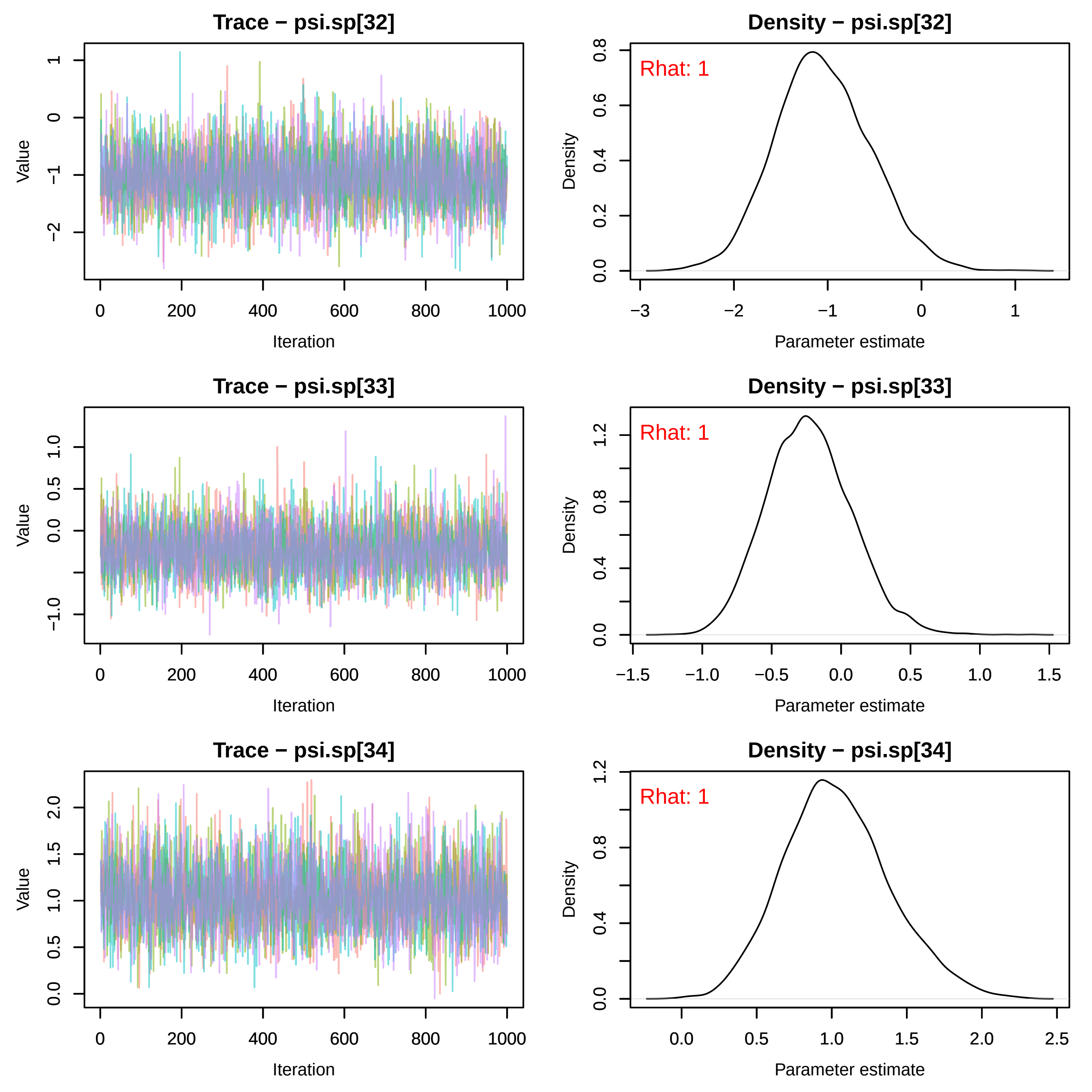

Trace − psi.sp[32]
Value
Iteration
Density − psi.sp[32]
Rhat: 1
Density
Parameter estimate
Trace − psi.sp[33]
Value
Iteration
Density − psi.sp[33]
Rhat: 1
Density
Parameter estimate
Trace − psi.sp[34]
Value
Iteration
Density − psi.sp[34]
Rhat: 1
Density
Parameter estimate

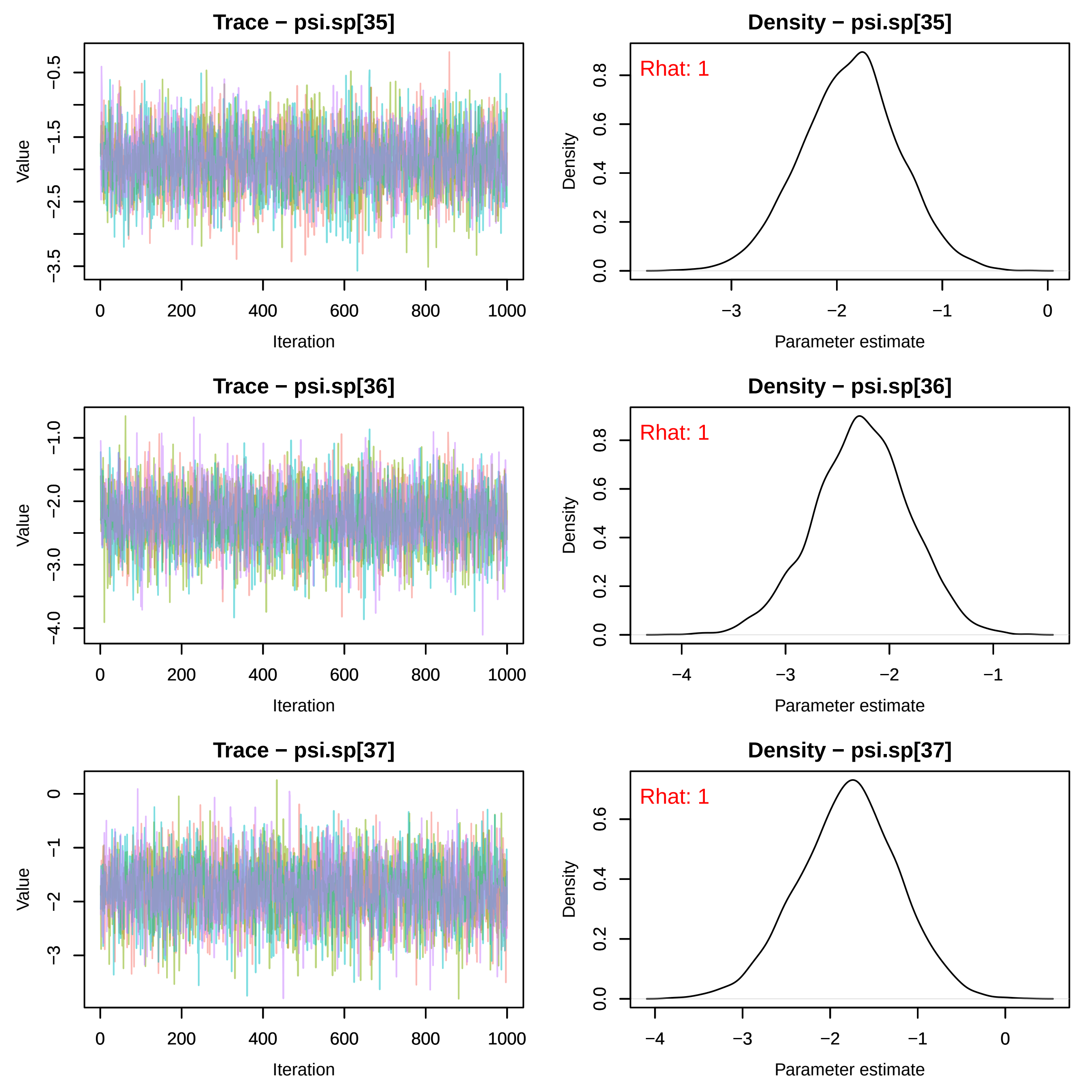
Trace − psi.sp[35]
Value
Iteration
Density − psi.sp[35]
Rhat: 1
Density
Parameter estimate
Trace − psi.sp[36]
Value
Iteration
Density − psi.sp[36]
Rhat: 1
Density
Parameter estimate
Trace − psi.sp[37]
Value
Iteration
Density − psi.sp[37]
Rhat: 1
Density
Parameter estimate

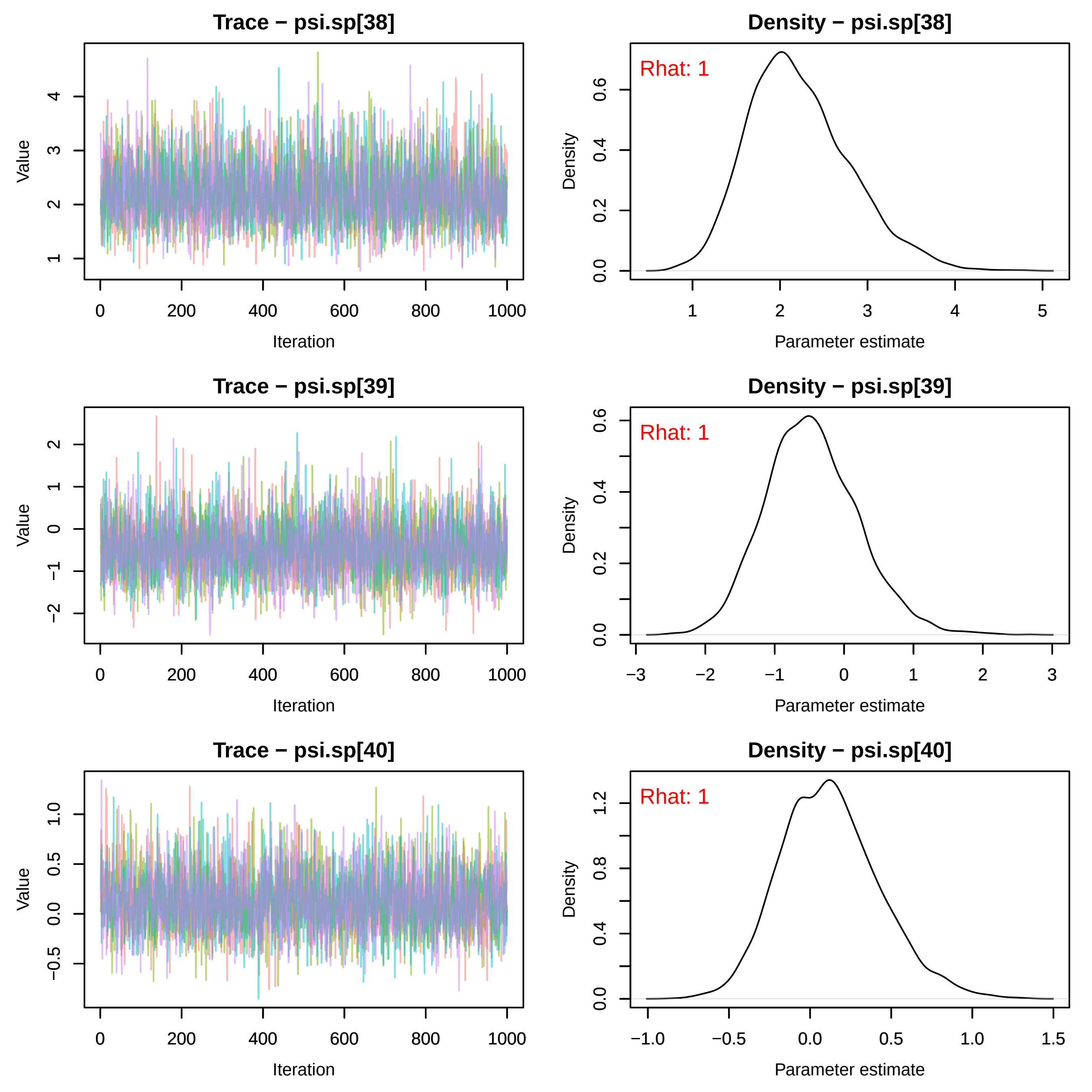

Trace − psi.sp[38]
Value
Iteration
Density − psi.sp[38]
Rhat: 1
Density
Parameter estimate
Trace − psi.sp[39]
Value
Iteration
Density − psi.sp[39]
Rhat: 1
Density
Parameter estimate
Trace − psi.sp[40]
Value
Iteration
Density − psi.sp[40]
Rhat: 1
Density
Parameter estimate

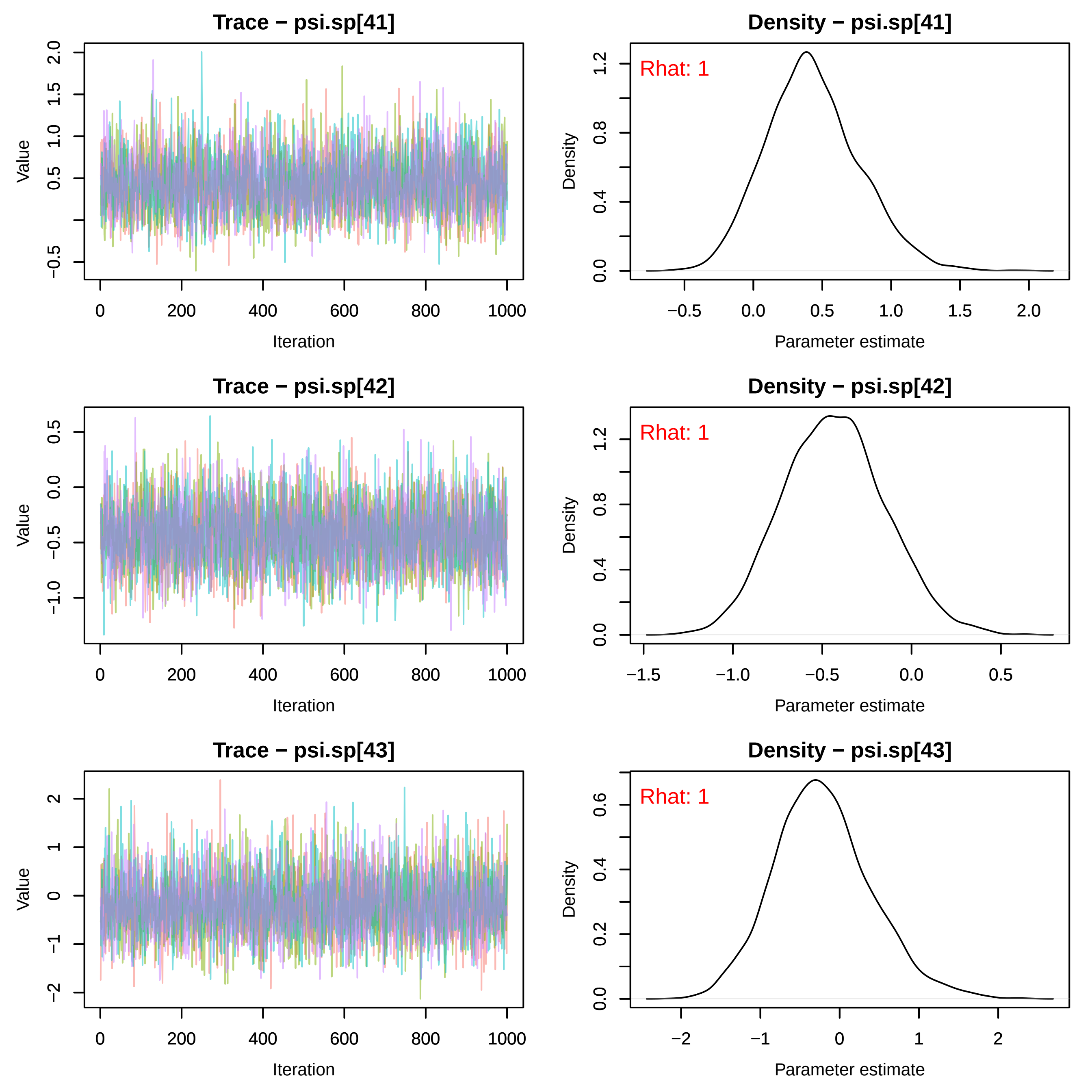
Trace − psi.sp[41]
Value
Iteration
Density − psi.sp[41]
Rhat: 1
Density
Parameter estimate
Trace − psi.sp[42]
Value
Iteration
Density − psi.sp[42]
Rhat: 1
Density
Parameter estimate
Trace − psi.sp[43]
Value
Iteration
Density − psi.sp[43]
Rhat: 1
Density
Parameter estimate

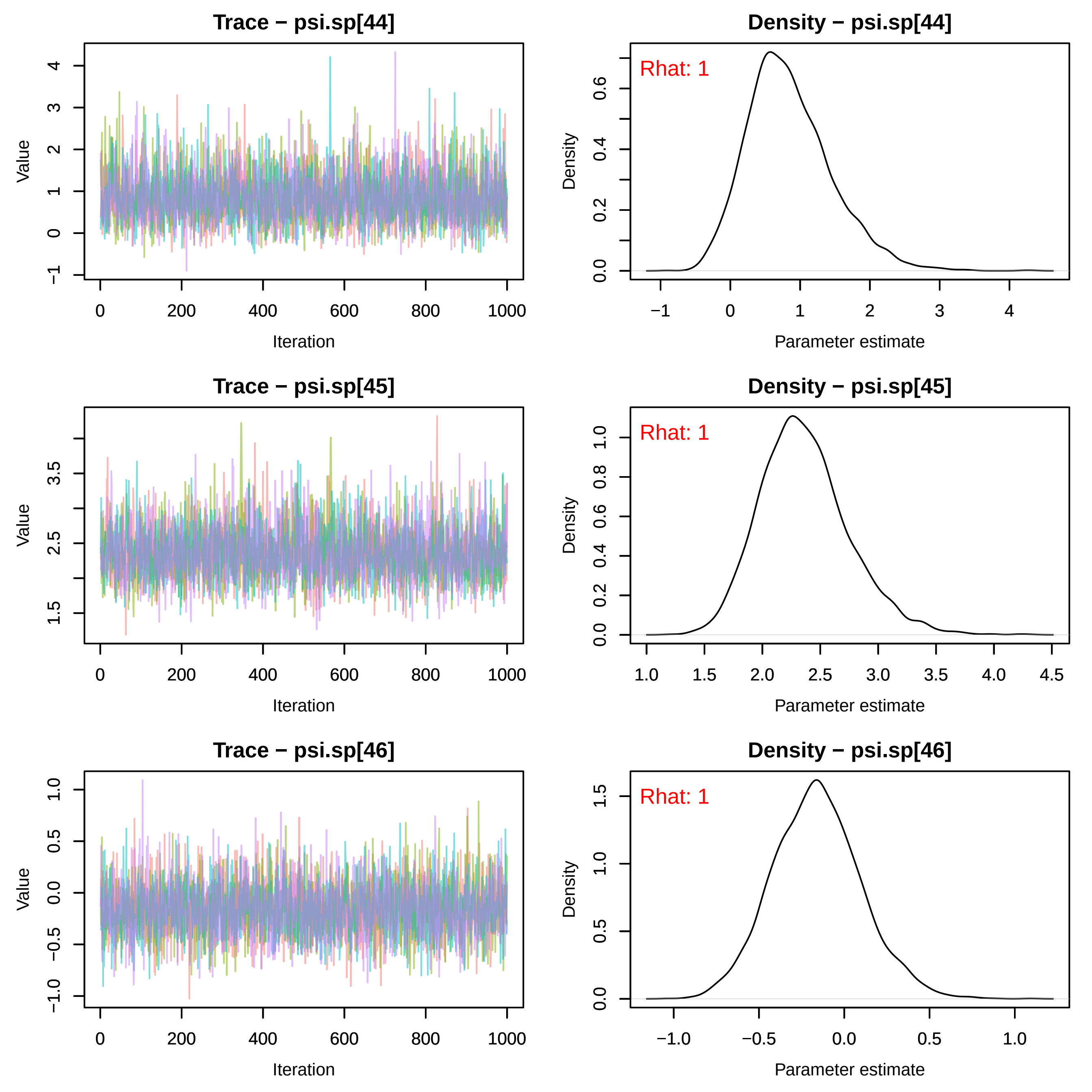
Trace − psi.sp[44]
Value
Iteration
Density − psi.sp[44]
Rhat: 1
Density
Parameter estimate
Trace − psi.sp[45]
Value
Iteration
Density − psi.sp[45]
Rhat: 1
Density
Parameter estimate
Trace − psi.sp[46]
Value
Iteration
Density − psi.sp[46]
Rhat: 1
Density
Parameter estimate

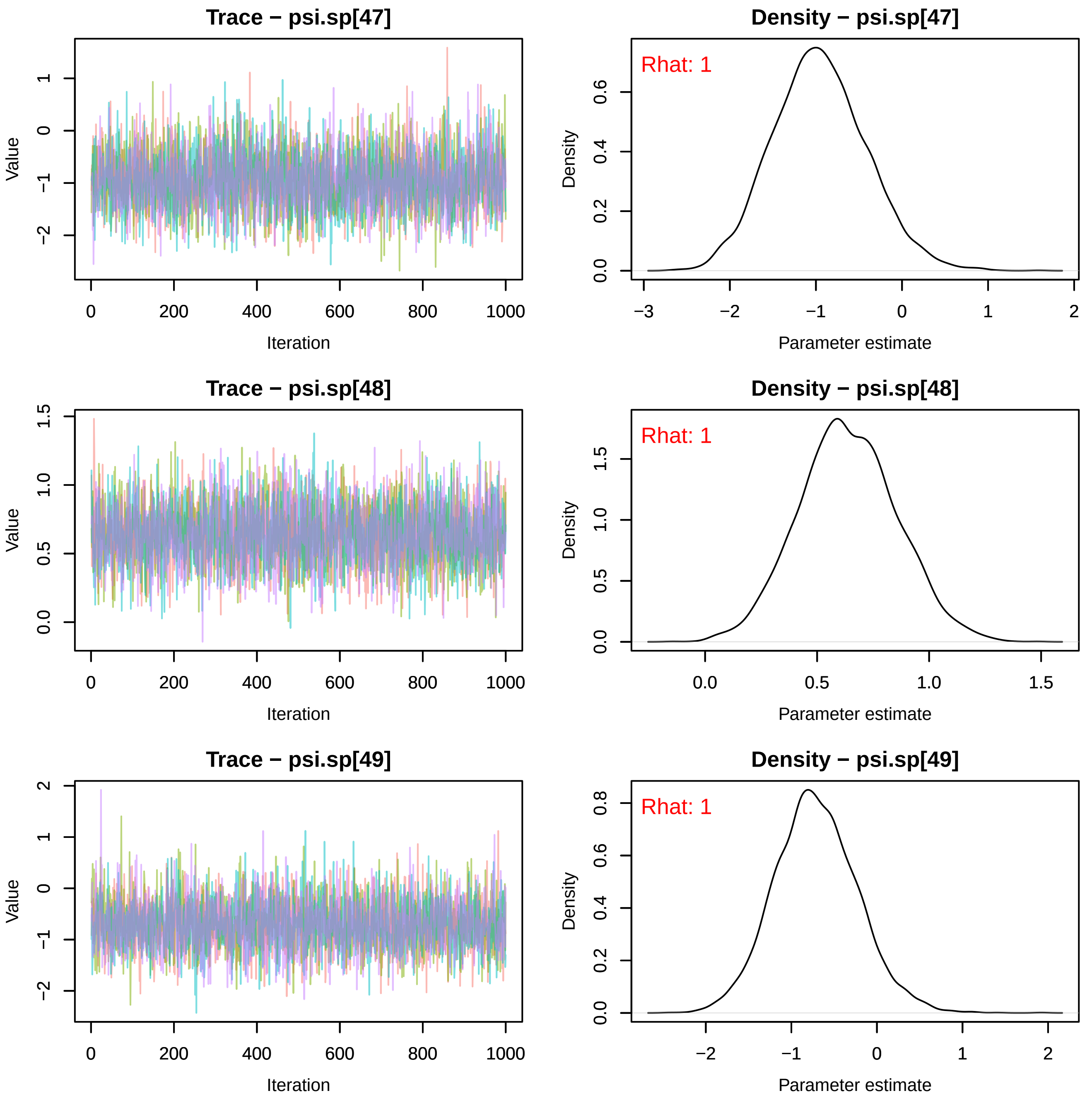
Trace − psi.sp[47]
Density − psi.sp[47]
Rhat: 1
Trace − psi.sp[48]
Density − psi.sp[48]
Rhat: 1
Trace − psi.sp[49]
Density − psi.sp[49]
Rhat: 1
Value
Iteration
Density
Parameter estimate

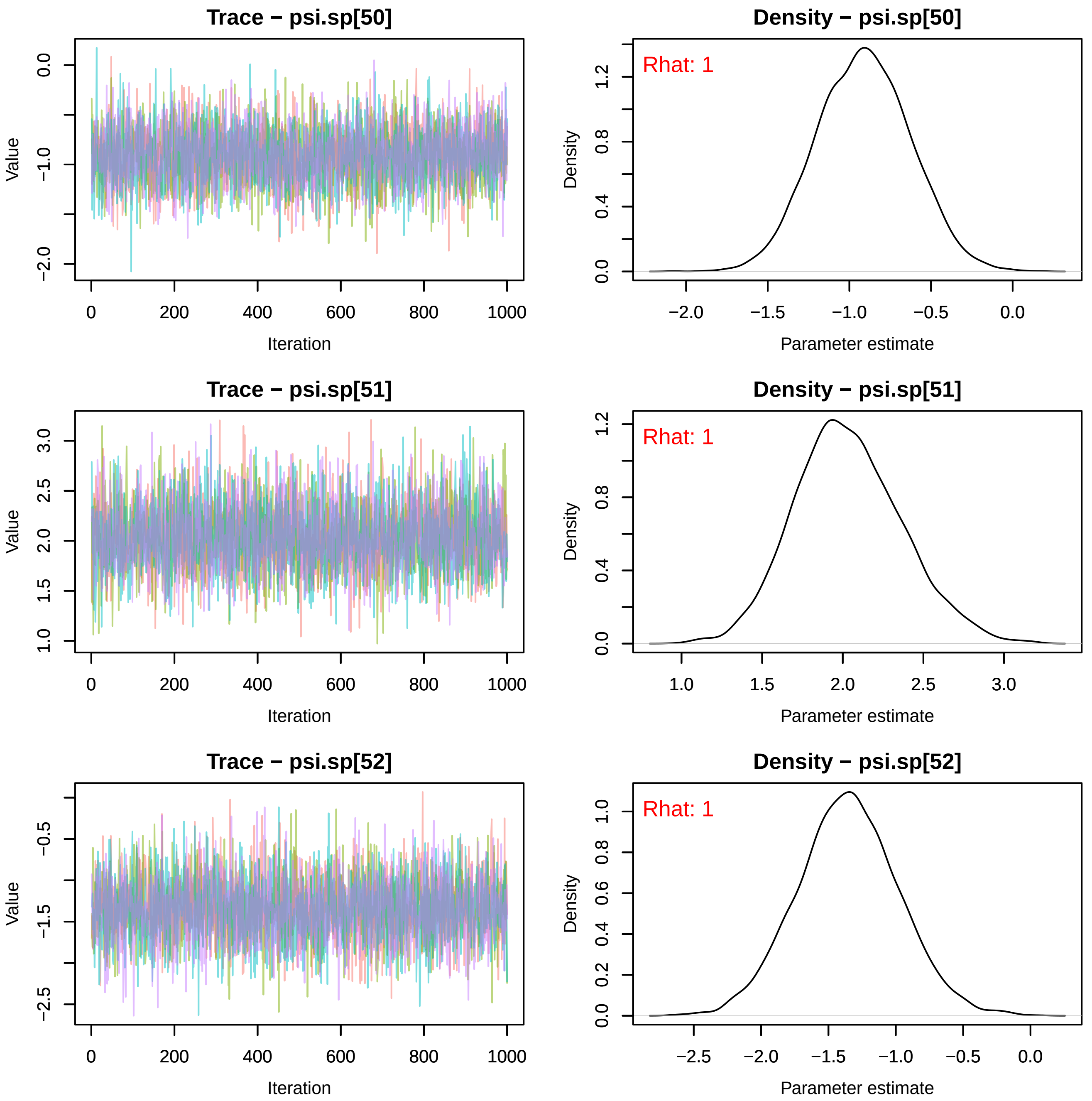

Trace − psi.sp[50]
Value
Iteration
Density − psi.sp[50]
Rhat: 1
Density
Parameter estimate
Trace − psi.sp[51]
Value
Iteration
Density − psi.sp[51]
Rhat: 1
Density
Parameter estimate
Trace − psi.sp[52]
Value
Iteration
Density − psi.sp[52]
Rhat: 1
Density
Parameter estimate

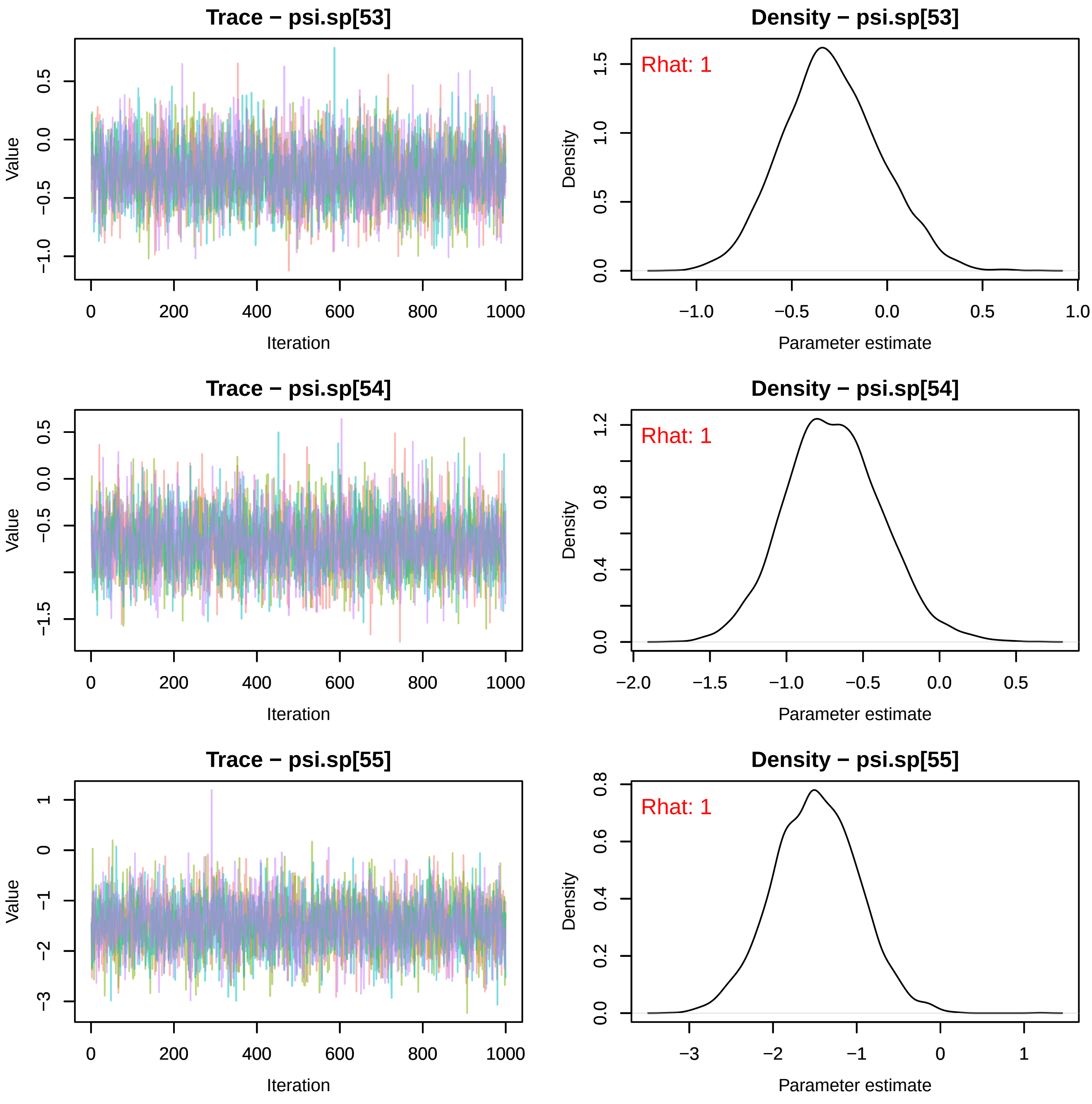
Trace − psi.sp[53]
Value
Iteration
Density − psi.sp[53]
Rhat: 1
Density
Parameter estimate
Trace − psi.sp[54]
Value
Iteration
Density − psi.sp[54]
Rhat: 1
Density
Parameter estimate
Trace − psi.sp[55]
Value
Iteration
Density − psi.sp[55]
Rhat: 1
Density
Parameter estimate

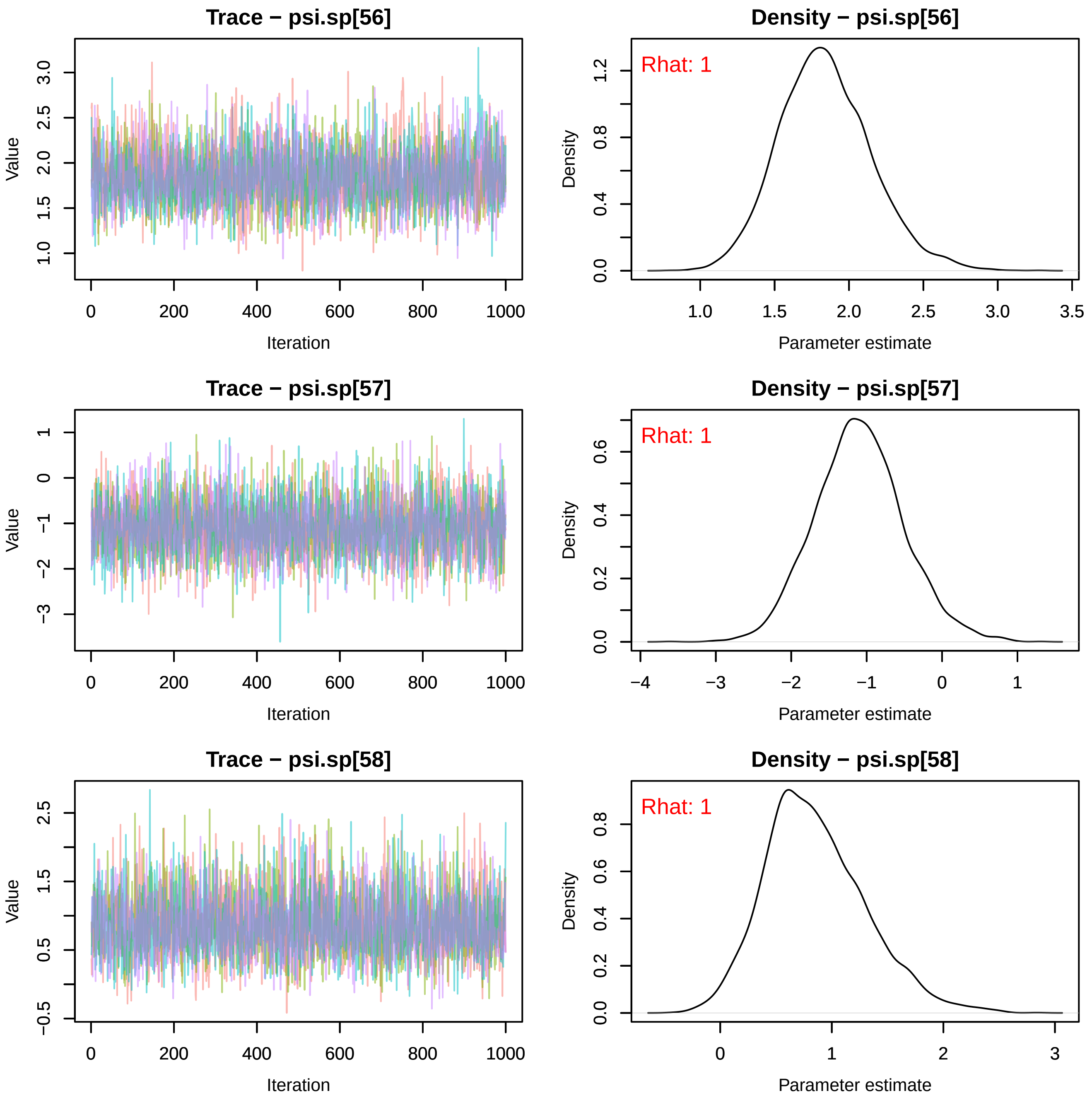
Trace − psi.sp[56]
Value
Iteration
Density − psi.sp[56]
Rhat: 1
Density
Parameter estimate
Trace − psi.sp[57]
Value
Iteration
Density − psi.sp[57]
Rhat: 1
Density
Parameter estimate
Trace − psi.sp[58]
Value
Iteration
Density − psi.sp[58]
Rhat: 1
Density
Parameter estimate

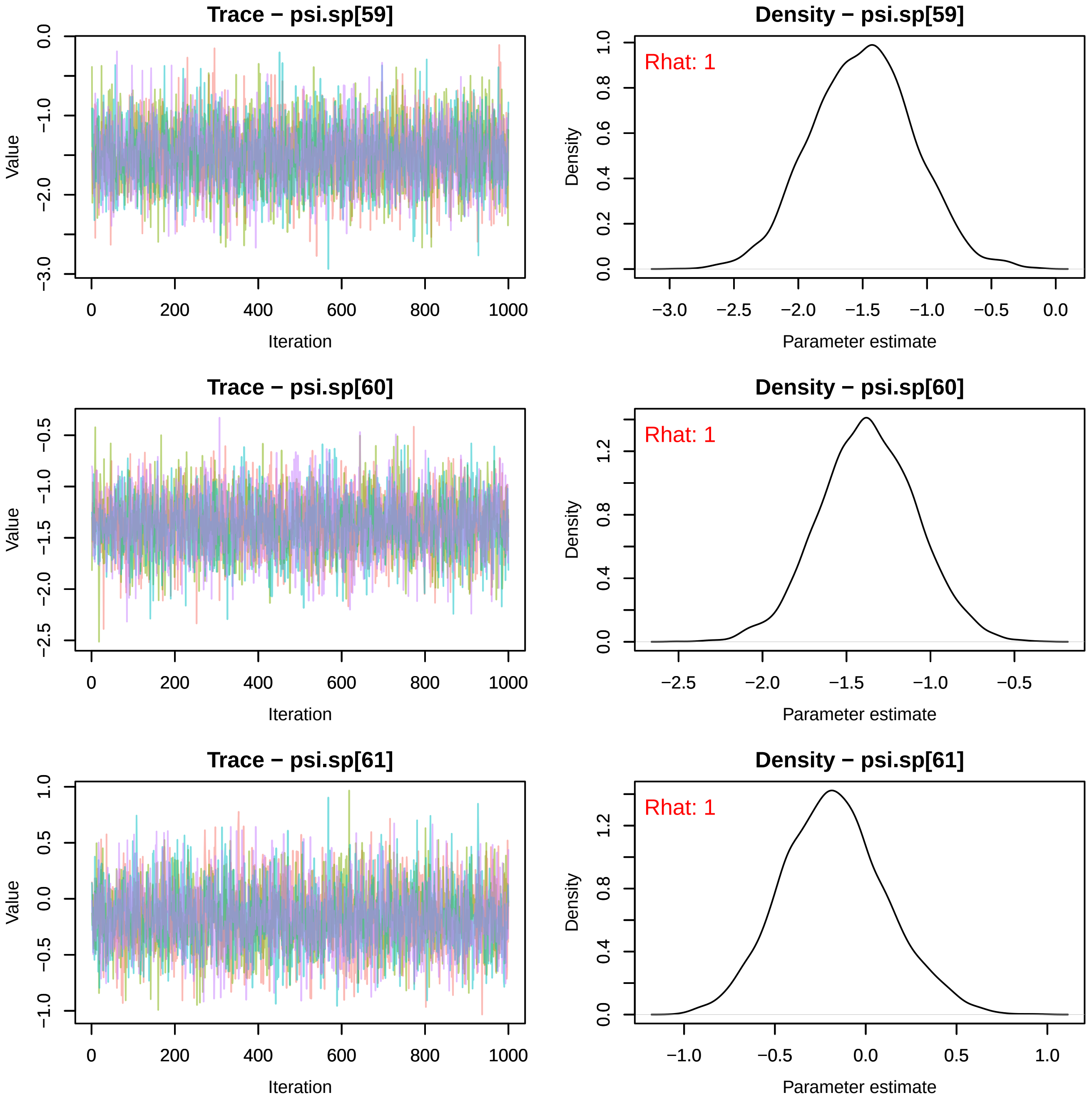
Trace − psi.sp[59]
Value
Iteration
Density − psi.sp[59]
Rhat: 1
Density
Parameter estimate
Trace − psi.sp[60]
Value
Iteration
Density − psi.sp[60]
Rhat: 1
Density
Parameter estimate
Trace − psi.sp[61]
Value
Iteration
Density − psi.sp[61]
Rhat: 1
Density
Parameter estimate

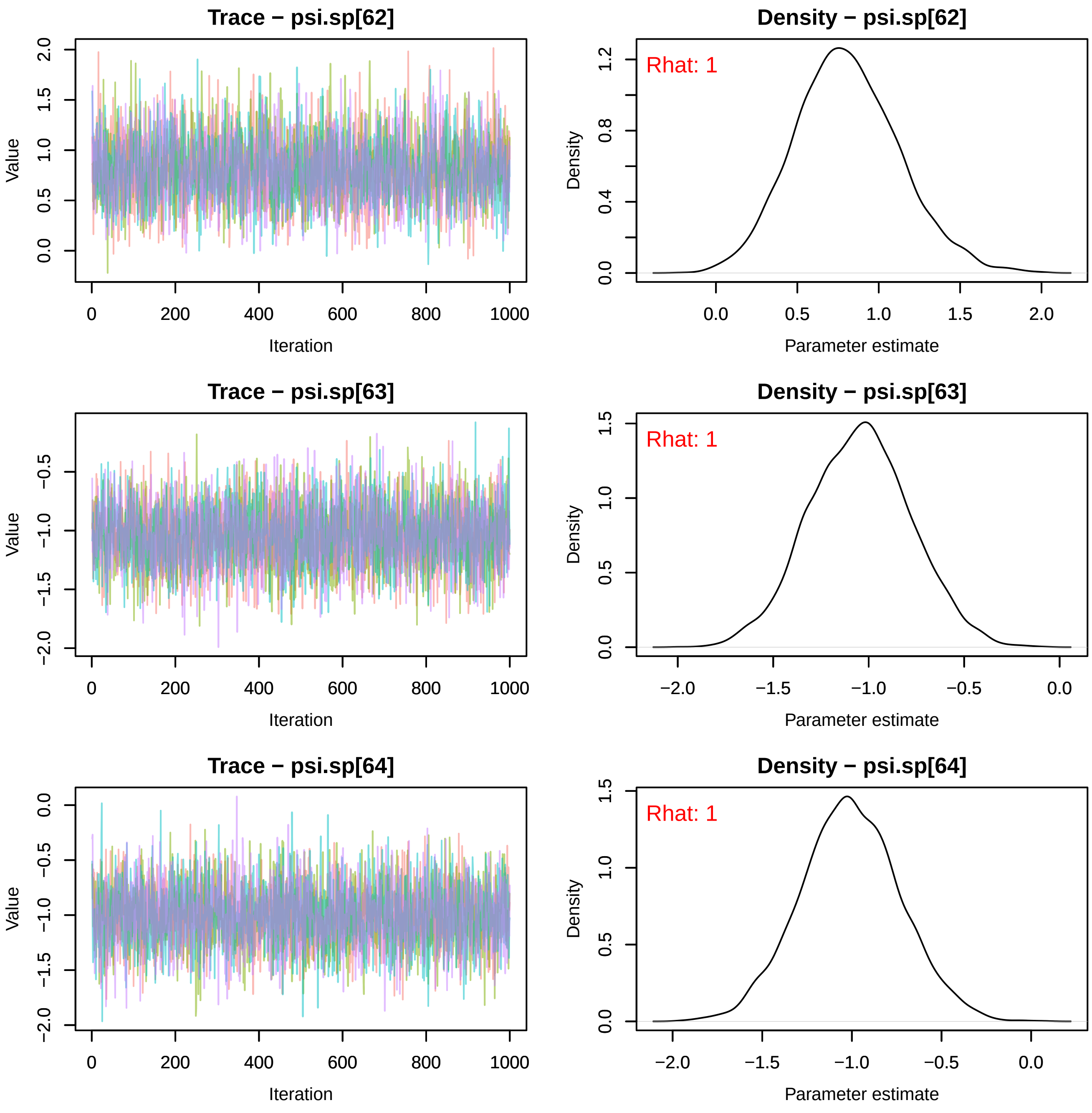
Trace − psi.sp[62]
Value
Iteration
0 200 400 600 800 1000
0.0 0.5 1.0 1.5 2.0
Density − psi.sp[62]
Rhat: 1
Density
Parameter estimate
0.0 0.5 1.0 1.5 2.0
0.0 0.4 0.8 1.2
Trace − psi.sp[63]
Value
Iteration
0 200 400 600 800 1000
−2.0 −1.5 −1.0 −0.5
Density − psi.sp[63]
Rhat: 1
Density
Parameter estimate
−2.0 −1.5 −1.0 −0.5 0.0
0.0 0.5 1.0 1.5
Trace − psi.sp[64]
Value
Iteration
0 200 400 600 800 1000
−2.0 −1.5 −1.0 −0.5 0.0
Density − psi.sp[64]
Rhat: 1
Density
Parameter estimate
−2.0 −1.5 −1.0 −0.5 0.0
0.0 0.5 1.0 1.5

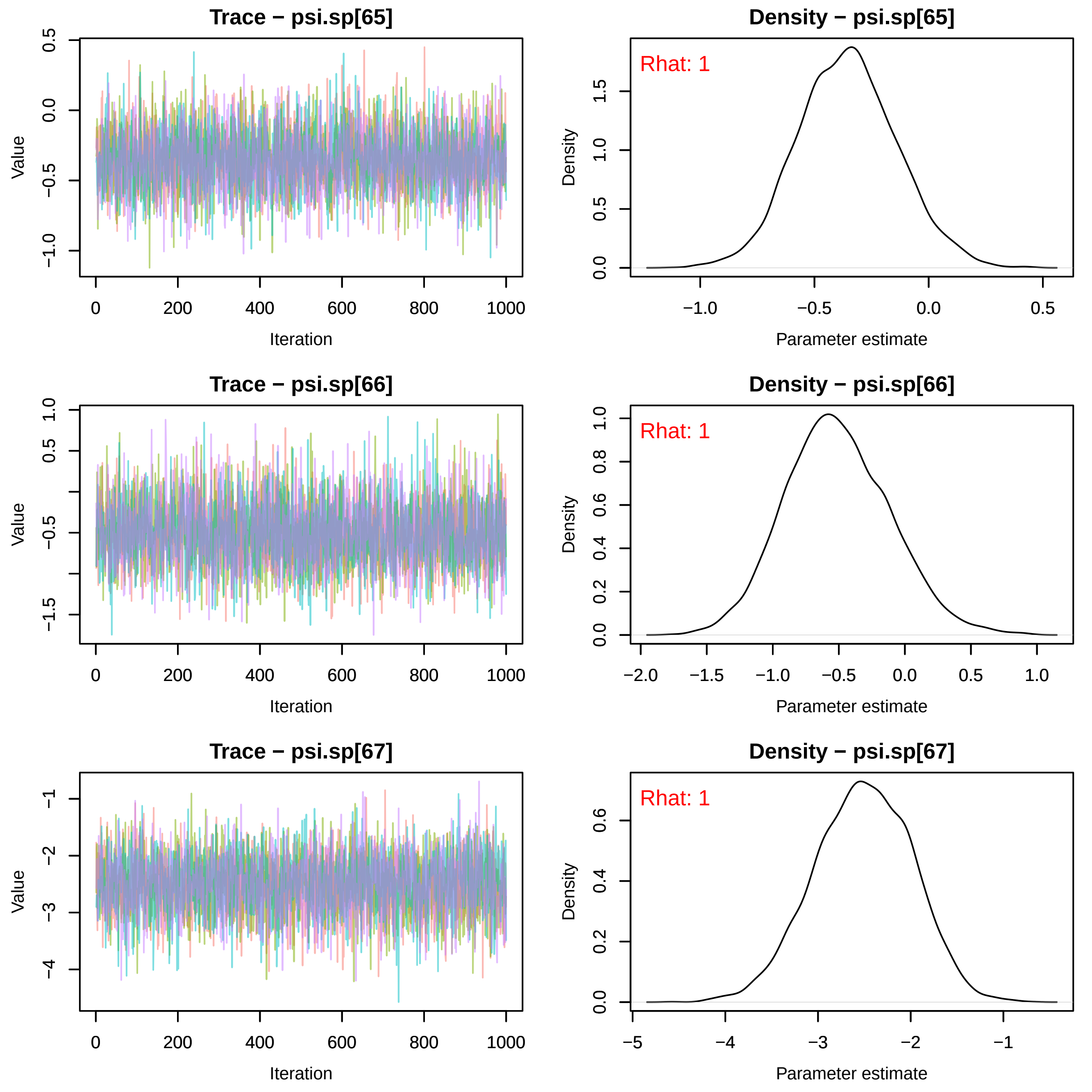

Trace − psi.sp[65]
Value
Iteration
Density − psi.sp[65]
Rhat: 1
Density
Parameter estimate
Trace − psi.sp[66]
Value
Iteration
Density − psi.sp[66]
Rhat: 1
Density
Parameter estimate
Trace − psi.sp[67]
Value
Iteration
Density − psi.sp[67]
Rhat: 1
Density
Parameter estimate

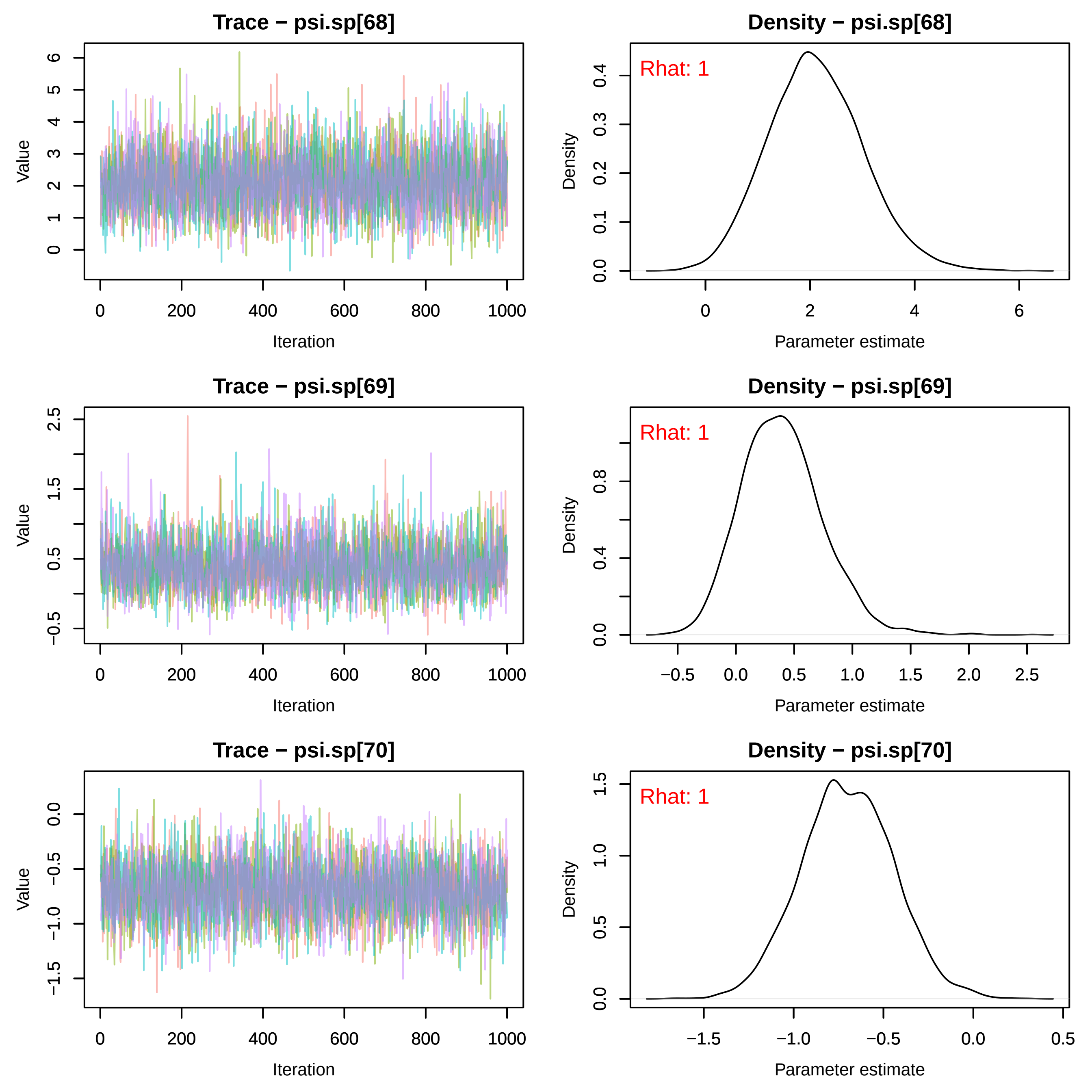
Trace − psi.sp[68]
Value
Iteration
Density − psi.sp[68]
Rhat: 1
Density
Parameter estimate
Trace − psi.sp[69]
Value
Iteration
Density − psi.sp[69]
Rhat: 1
Density
Parameter estimate
Trace − psi.sp[70]
Value
Iteration
Density − psi.sp[70]
Rhat: 1
Density
Parameter estimate

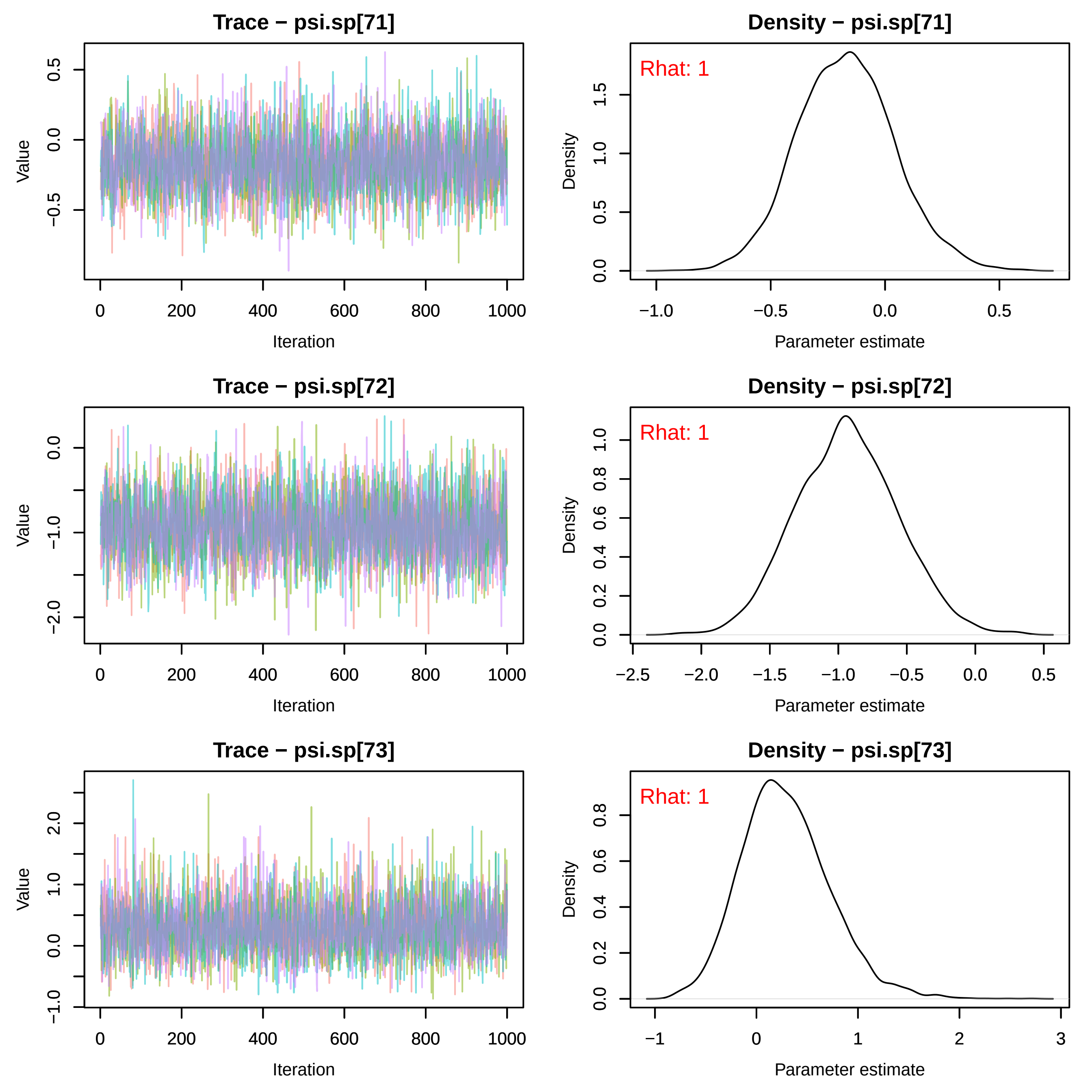
Trace − psi.sp[71]
Value
Iteration
Density − psi.sp[71]
Rhat: 1
Density
Parameter estimate
Trace − psi.sp[72]
Value
Iteration
Density − psi.sp[72]
Rhat: 1
Density
Parameter estimate
Trace − psi.sp[73]
Value
Iteration
Density − psi.sp[73]
Rhat: 1
Density
Parameter estimate

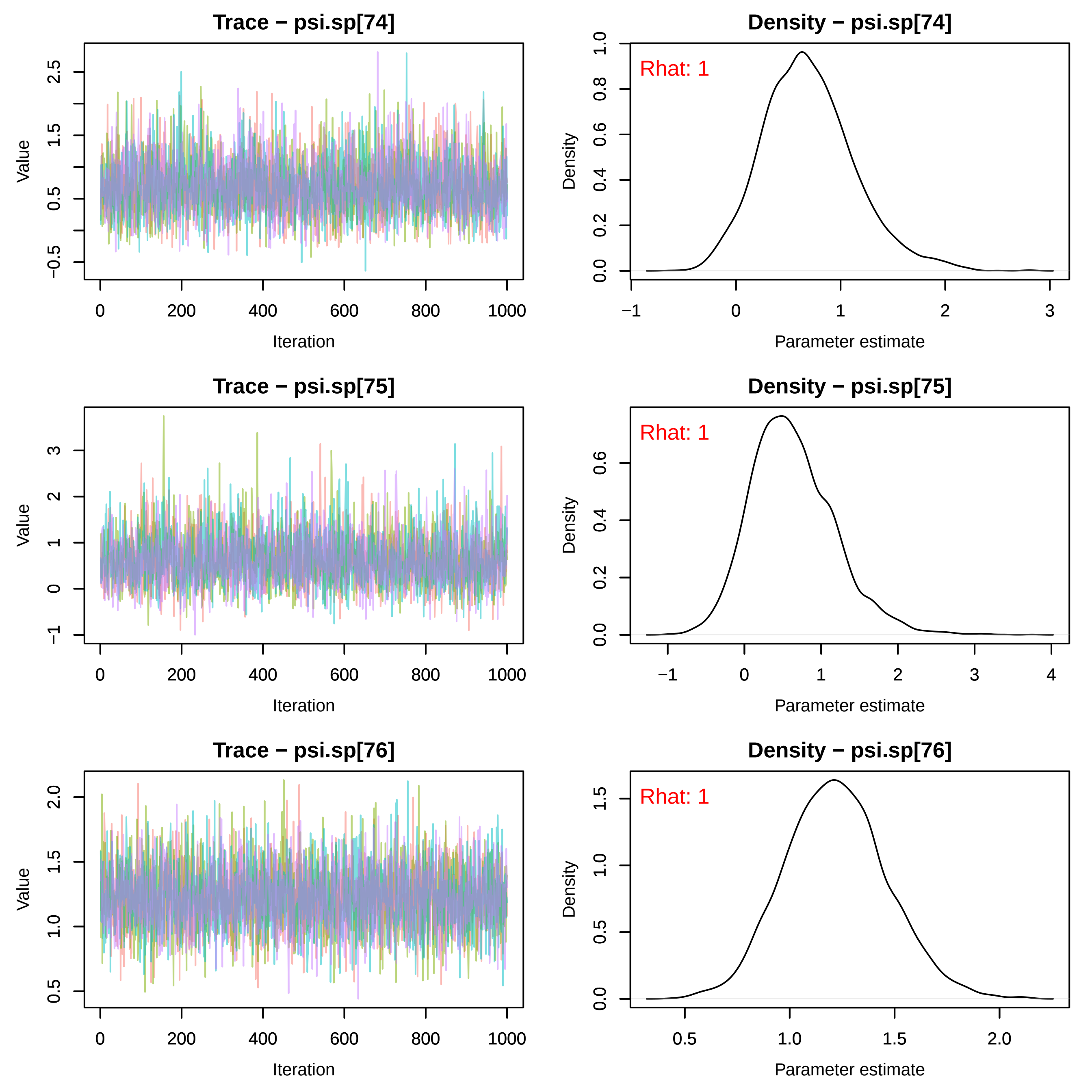
Trace − psi.sp[74]
Value
−0.5
0.5
1.5
2.5
0
200
400
600
800
1000
Iteration
Density − psi.sp[74]
Rhat: 1
Density
0.0
0.2
0.4
0.6
0.8
1.0
−1
0
1
2
3
Parameter estimate
Trace − psi.sp[75]
Value
−1
0
1
2
3
0
200
400
600
800
1000
Iteration
Density − psi.sp[75]
Rhat: 1
Density
0.0
0.2
0.4
0.6
−1
0
1
2
3
4
Parameter estimate
Trace − psi.sp[76]
Value
0.5
1.0
1.5
2.0
0
200
400
600
800
1000
Iteration
Density − psi.sp[76]
Rhat: 1
Density
0.0
0.5
1.0
1.5
0.5
1.0
1.5
2.0
Parameter estimate

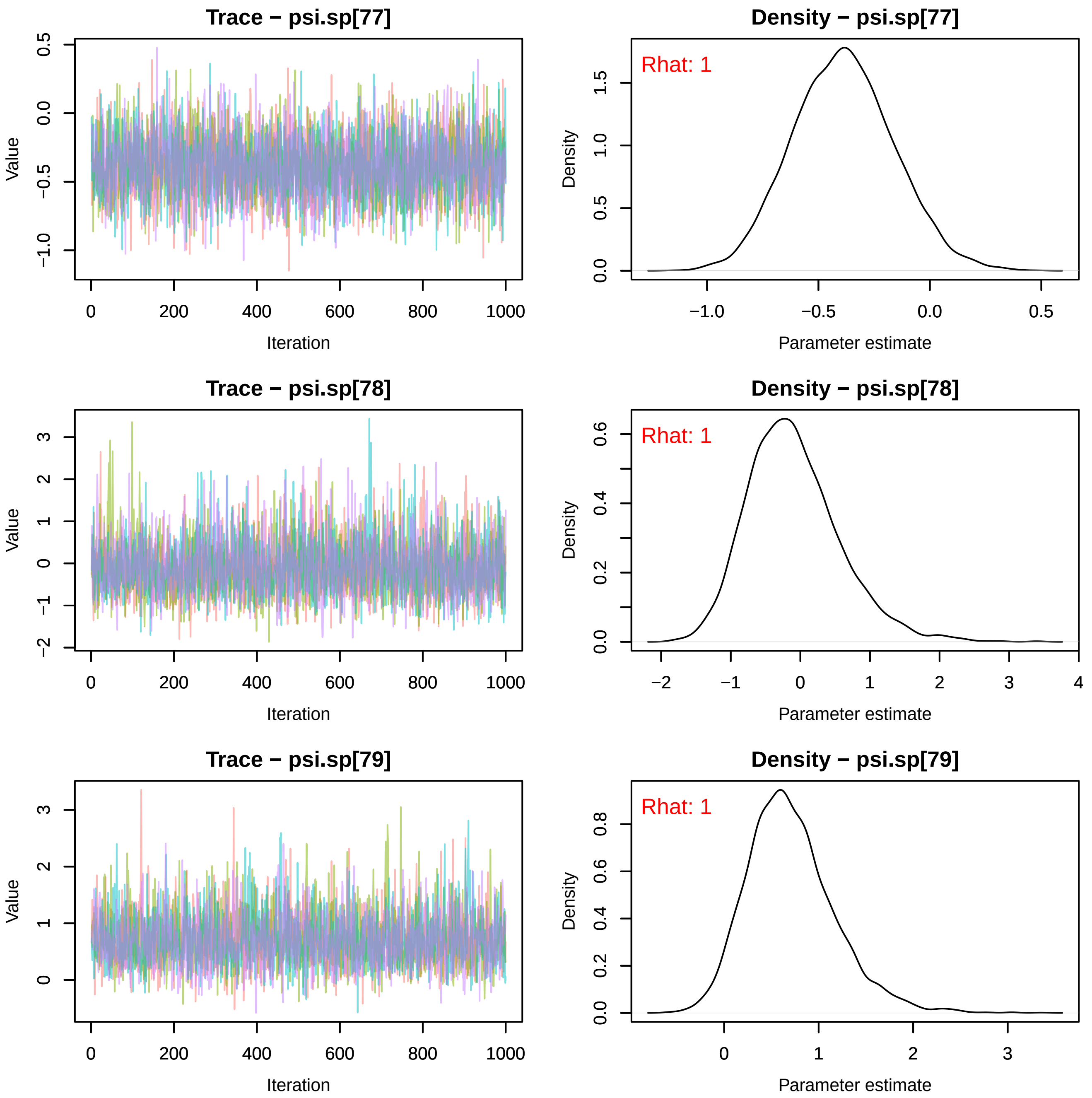
Trace − psi.sp[77]
Value
Iteration
Density − psi.sp[77]
Rhat: 1
Density
Parameter estimate
Trace − psi.sp[78]
Value
Iteration
Density − psi.sp[78]
Rhat: 1
Density
Parameter estimate
Trace − psi.sp[79]
Value
Iteration
Density − psi.sp[79]
Rhat: 1
Density
Parameter estimate

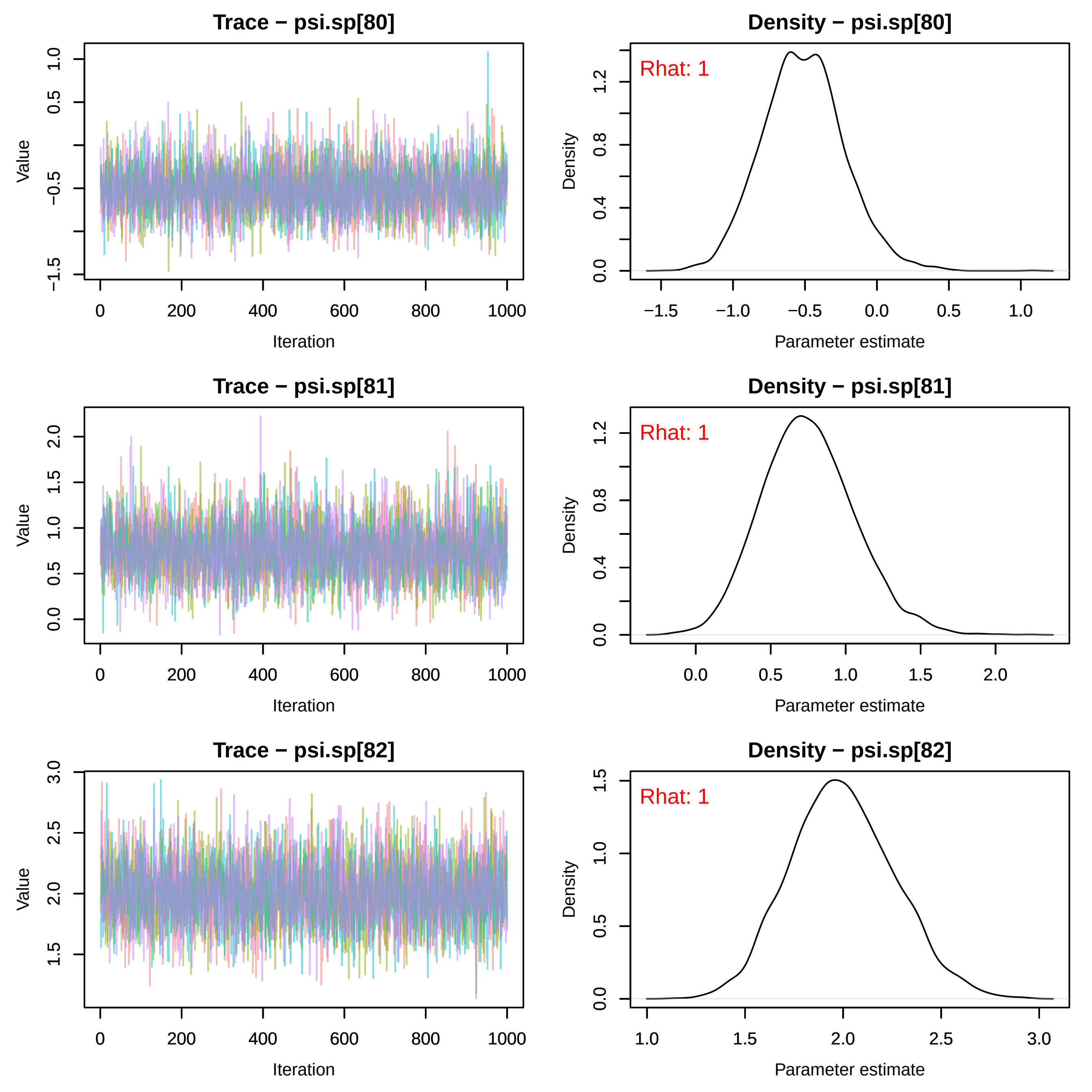
Trace − psi.sp[80]
Density − psi.sp[80]
Rhat: 1
Trace − psi.sp[81]
Density − psi.sp[81]
Rhat: 1
Trace − psi.sp[82]
Density − psi.sp[82]
Rhat: 1
Value
Iteration
Density
Parameter estimate

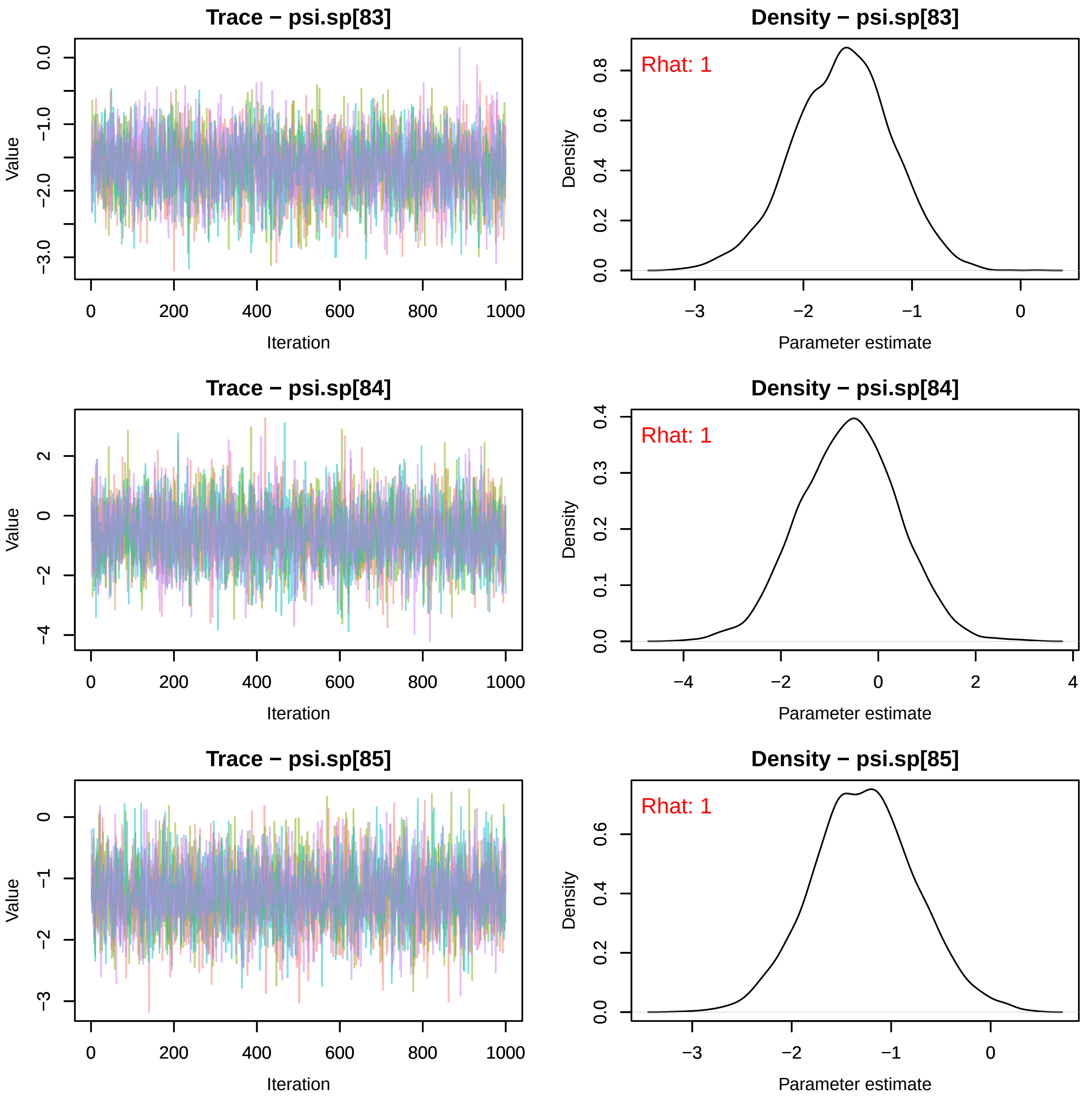
Trace − psi.sp[83]
Density − psi.sp[83]
Rhat: 1
Value
Iteration
Density
Parameter estimate
Trace − psi.sp[84]
Density − psi.sp[84]
Rhat: 1
Value
Iteration
Density
Parameter estimate
Trace − psi.sp[85]
Density − psi.sp[85]
Rhat: 1
Value
Iteration
Density
Parameter estimate

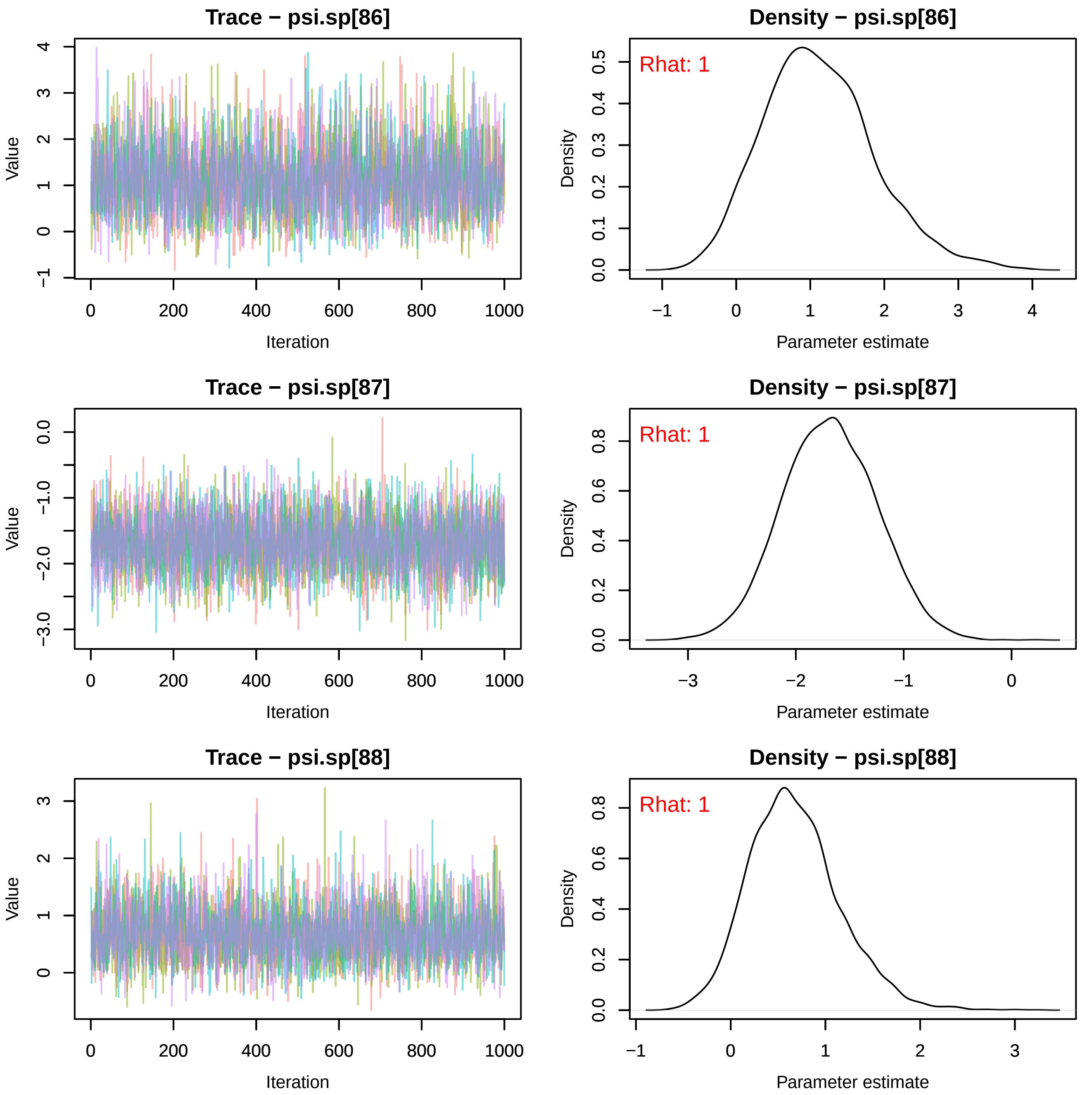
Trace − psi.sp[86]
Value
Iteration
Density − psi.sp[86]
Rhat: 1
Density
Parameter estimate
Trace − psi.sp[87]
Value
Iteration
Density − psi.sp[87]
Rhat: 1
Density
Parameter estimate
Trace − psi.sp[88]
Value
Iteration
Density − psi.sp[88]
Rhat: 1
Density
Parameter estimate

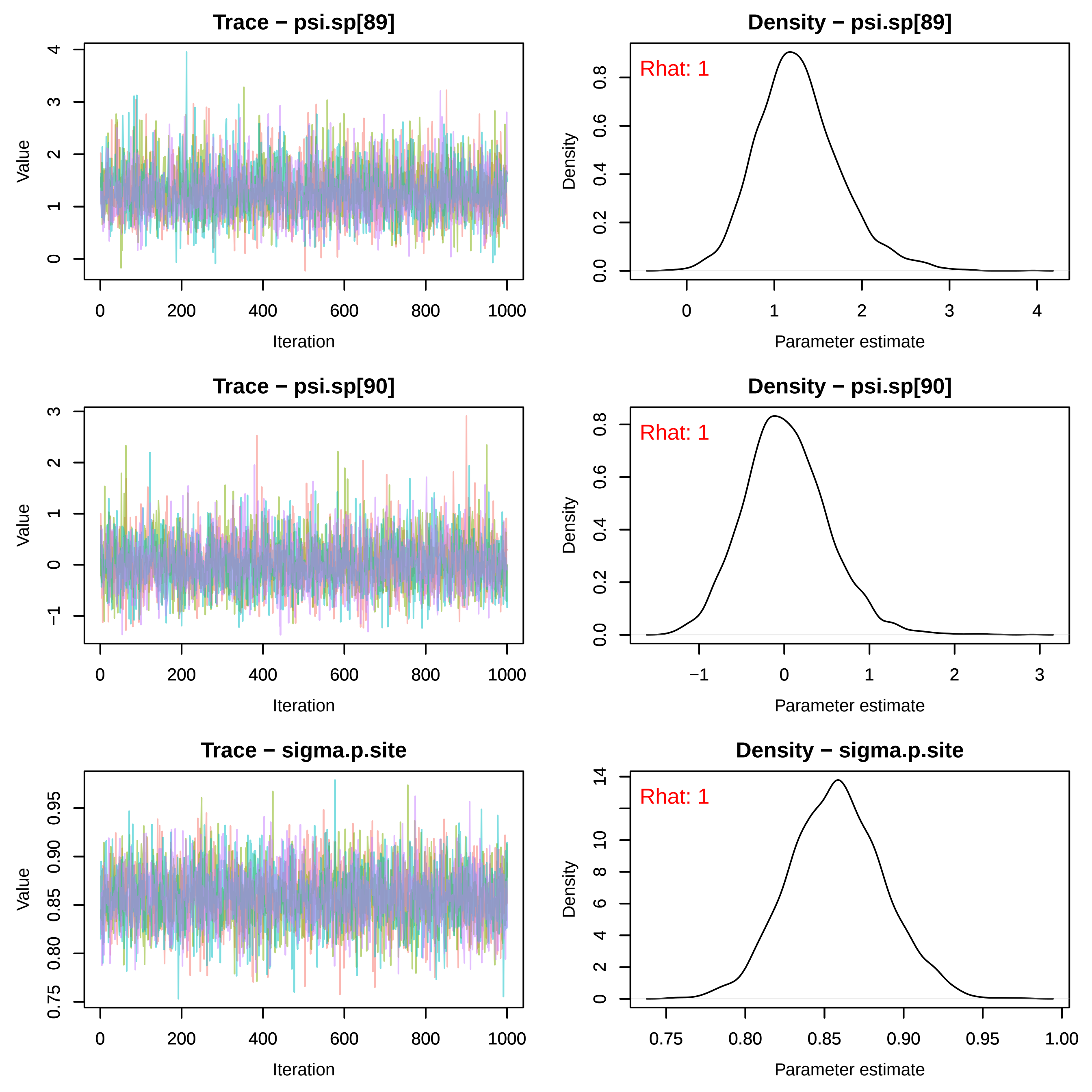

Trace − psi.sp[89]
Value
Iteration
Density − psi.sp[89]
Rhat: 1
Density
Parameter estimate
Trace − psi.sp[90]
Value
Iteration
Density − psi.sp[90]
Rhat: 1
Density
Parameter estimate
Trace − sigma.p.site
Value
Iteration
Density − sigma.p.site
Rhat: 1
Density
Parameter estimate

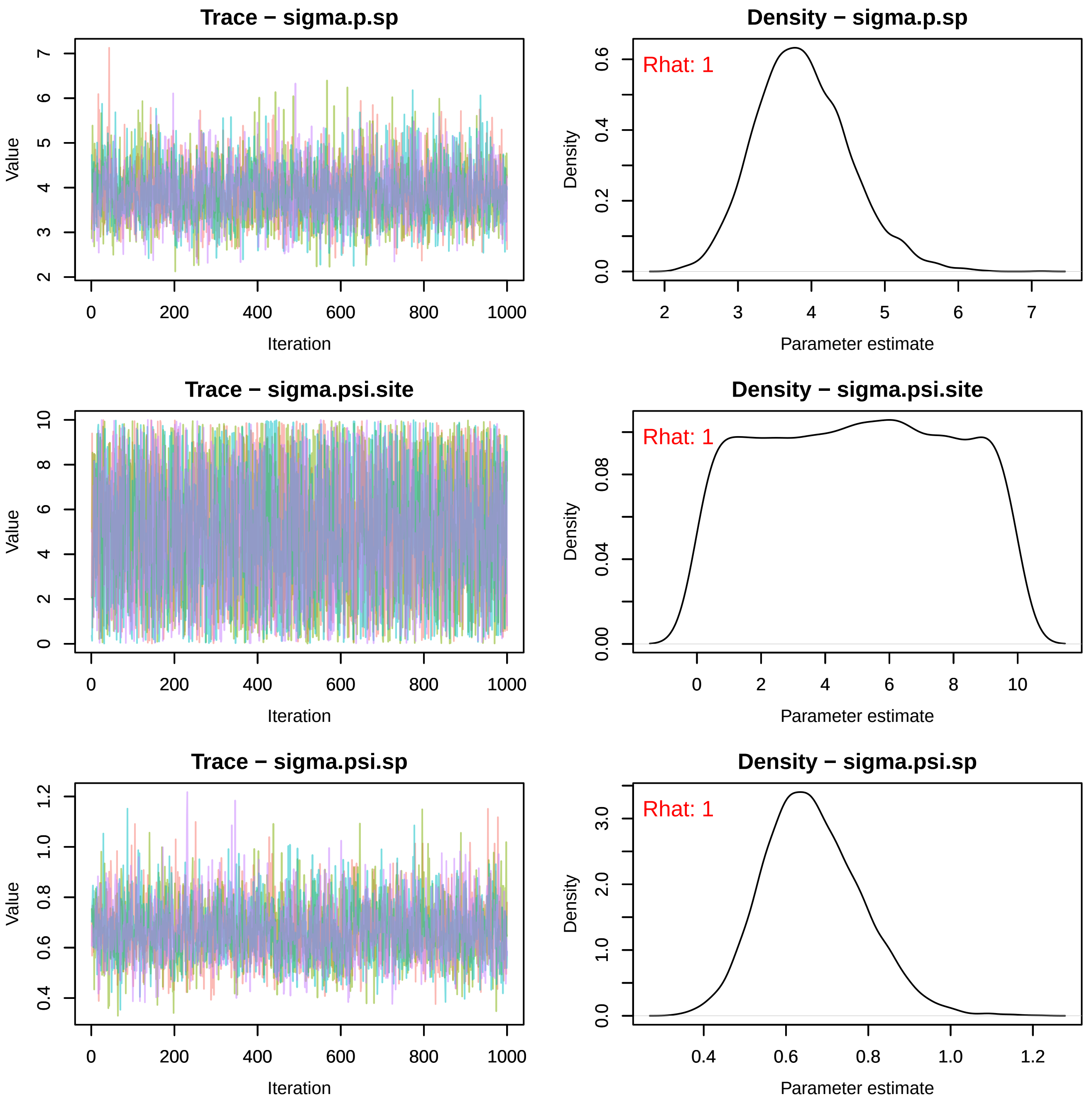
Trace − sigma.p.sp
Density − sigma.p.sp
Rhat: 1
Trace − sigma.psi.site
Density − sigma.psi.site
Rhat: 1
Trace − sigma.psi.sp
Density − sigma.psi.sp
Rhat: 1
Value
Iteration
Density
Parameter estimate

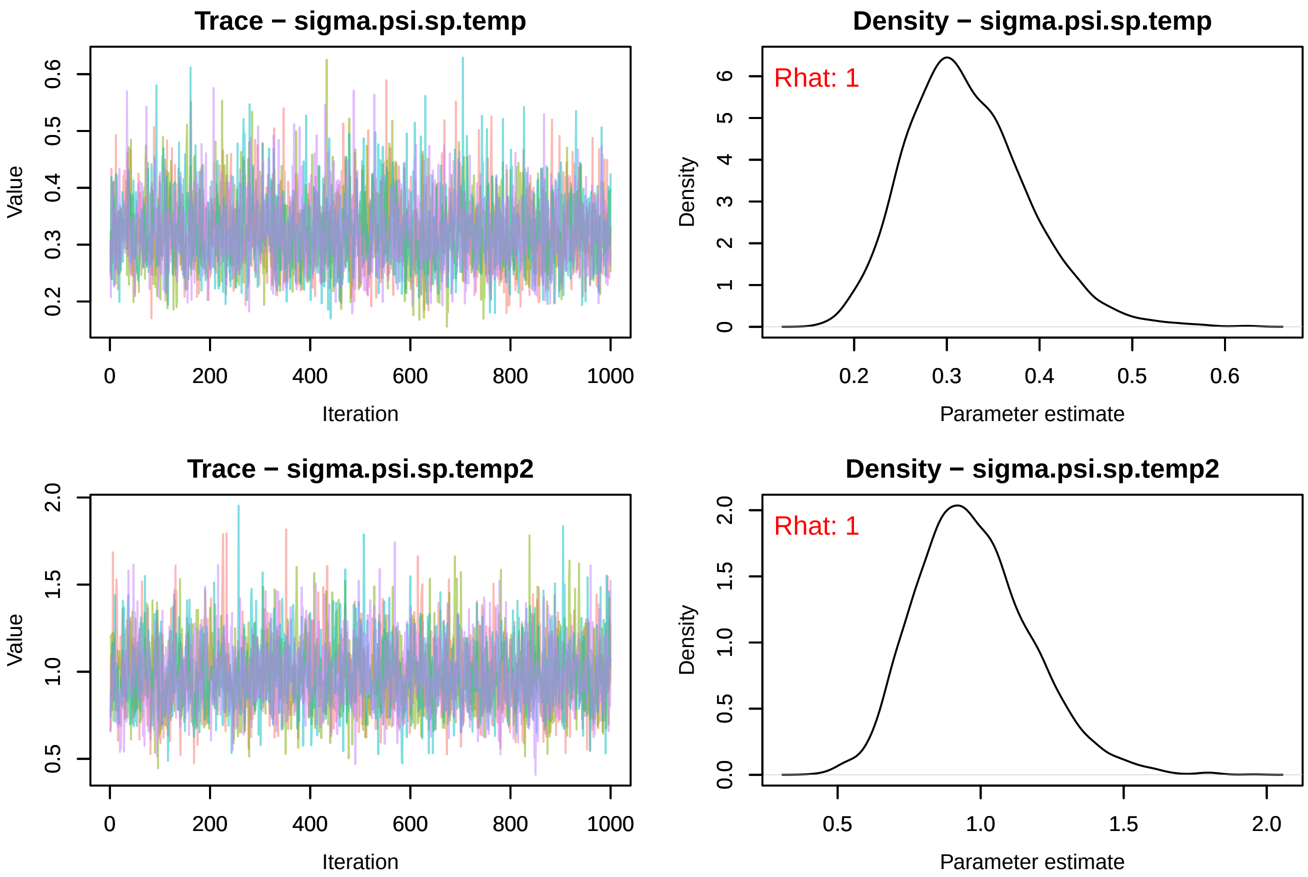

Trace − sigma.psi.sp.temp
Value
Iteration
Density − sigma.psi.sp.temp
Rhat: 1
Density
Parameter estimate
Trace − sigma.psi.sp.temp2
Value
Iteration
Density − sigma.psi.sp.temp2
Rhat: 1
Density
Parameter estimate
