## Supplemental File S4 for "Rising minimum temperatures contribute to 50 years of shifting Arctic and boreal butterfly communities in North America"

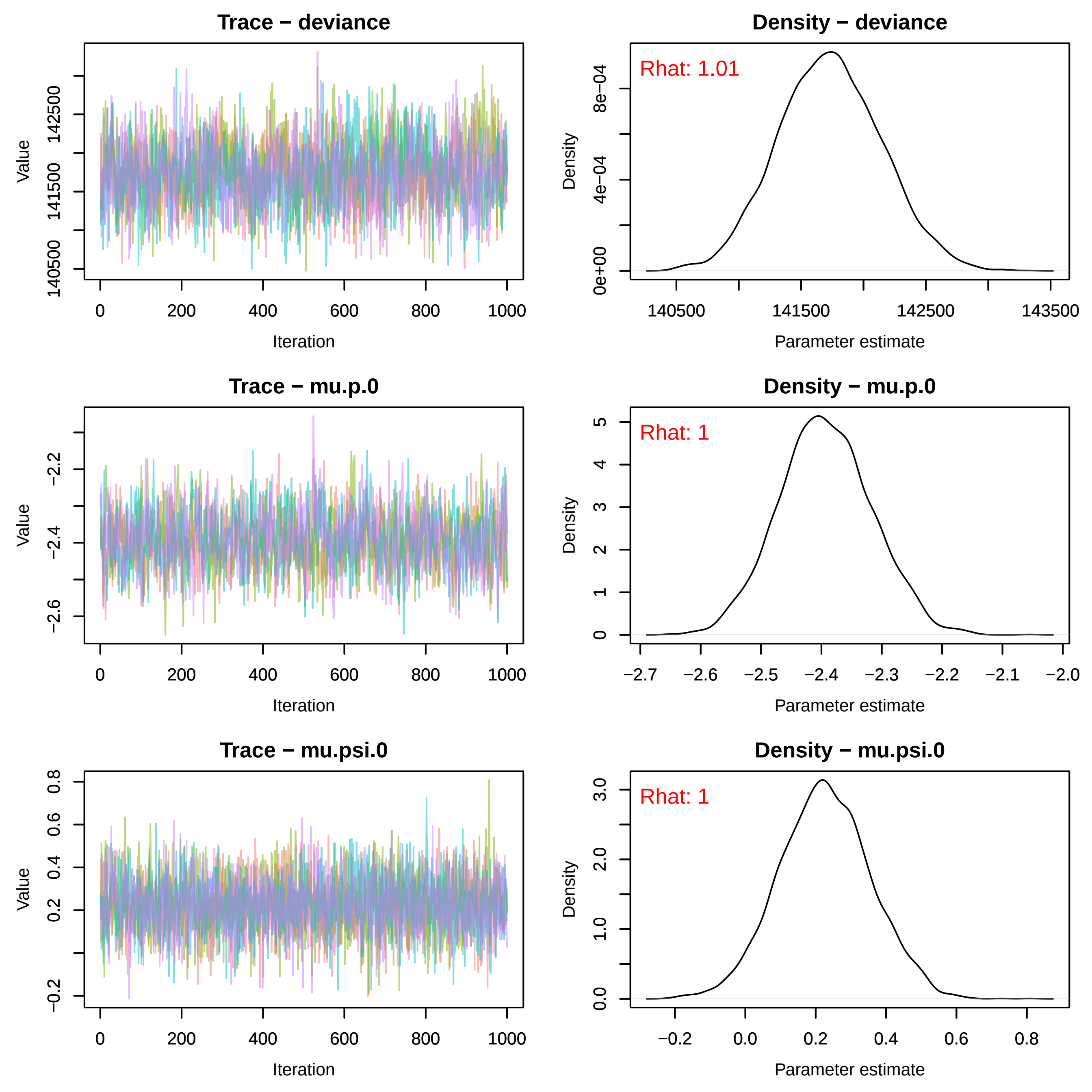
Trace − deviance
Value
Iteration
Density − deviance
Rhat: 1.01
Density
Parameter estimate
Trace − mu.p.0
Value
Iteration
Density − mu.p.0
Rhat: 1
Density
Parameter estimate
Trace − mu.psi.0
Value
Iteration
Density − mu.psi.0
Rhat: 1
Density
Parameter estimate

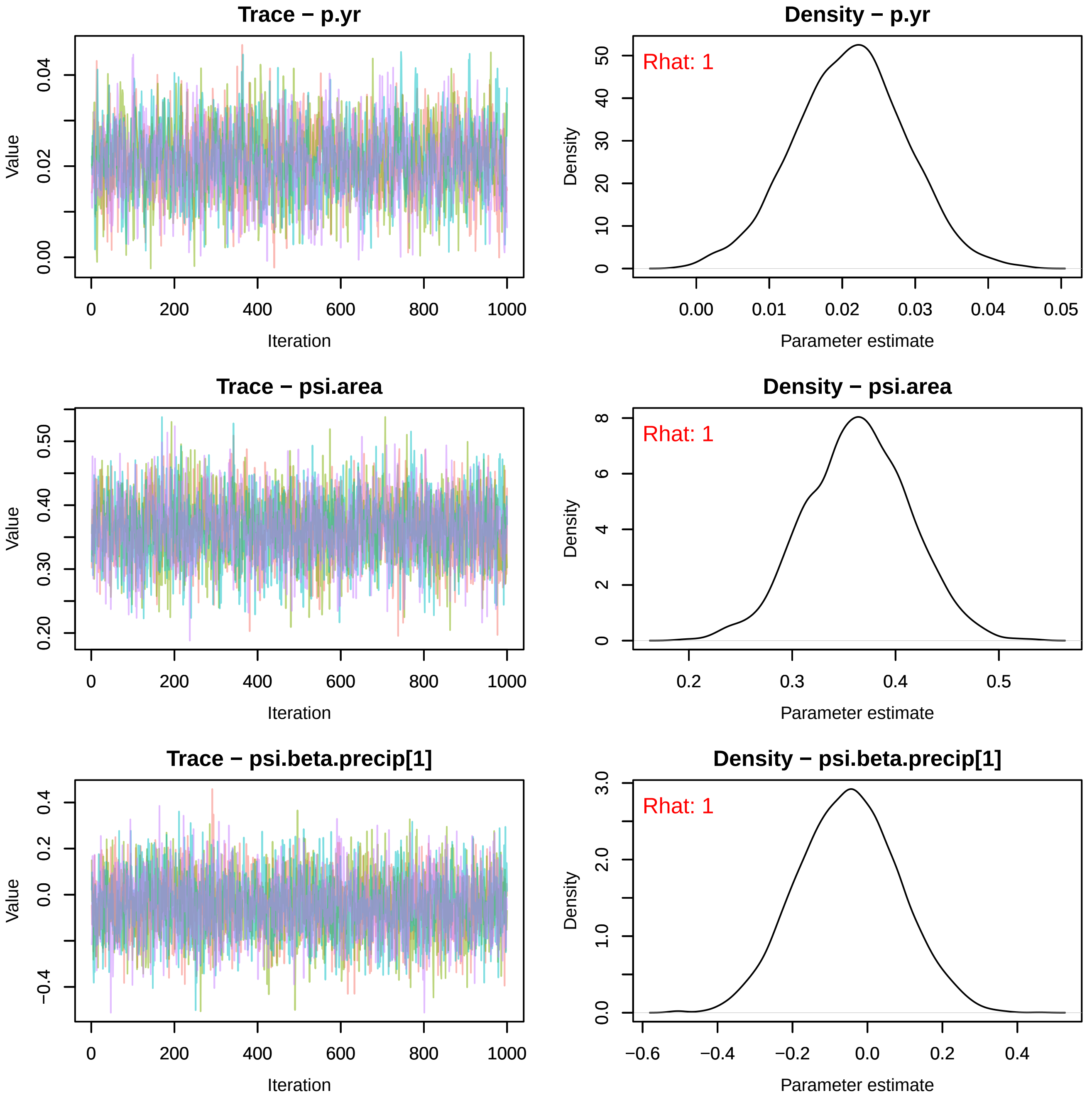
Trace − p.yr
Value
0.00
0.02
0.04
Iteration
0
200
400
600
800
1000
Density − p.yr
Rhat: 1
Density
0
10
20
30
40
50
Parameter estimate
0.00
0.01
0.02
0.03
0.04
0.05
Trace − psi.area
Value
0.20
0.30
0.40
0.50
Iteration
0
200
400
600
800
1000
Density − psi.area
Rhat: 1
Density
0
2
4
6
8
Parameter estimate
0.2
0.3
0.4
0.5
Trace − psi.beta.precip[1]
Value
−0.4
0.0
0.2
0.4
Iteration
0
200
400
600
800
1000
Density − psi.beta.precip[1]
Rhat: 1
Density
0.0
1.0
2.0
3.0
Parameter estimate
−0.6
−0.4
−0.2
0.0
0.2
0.4

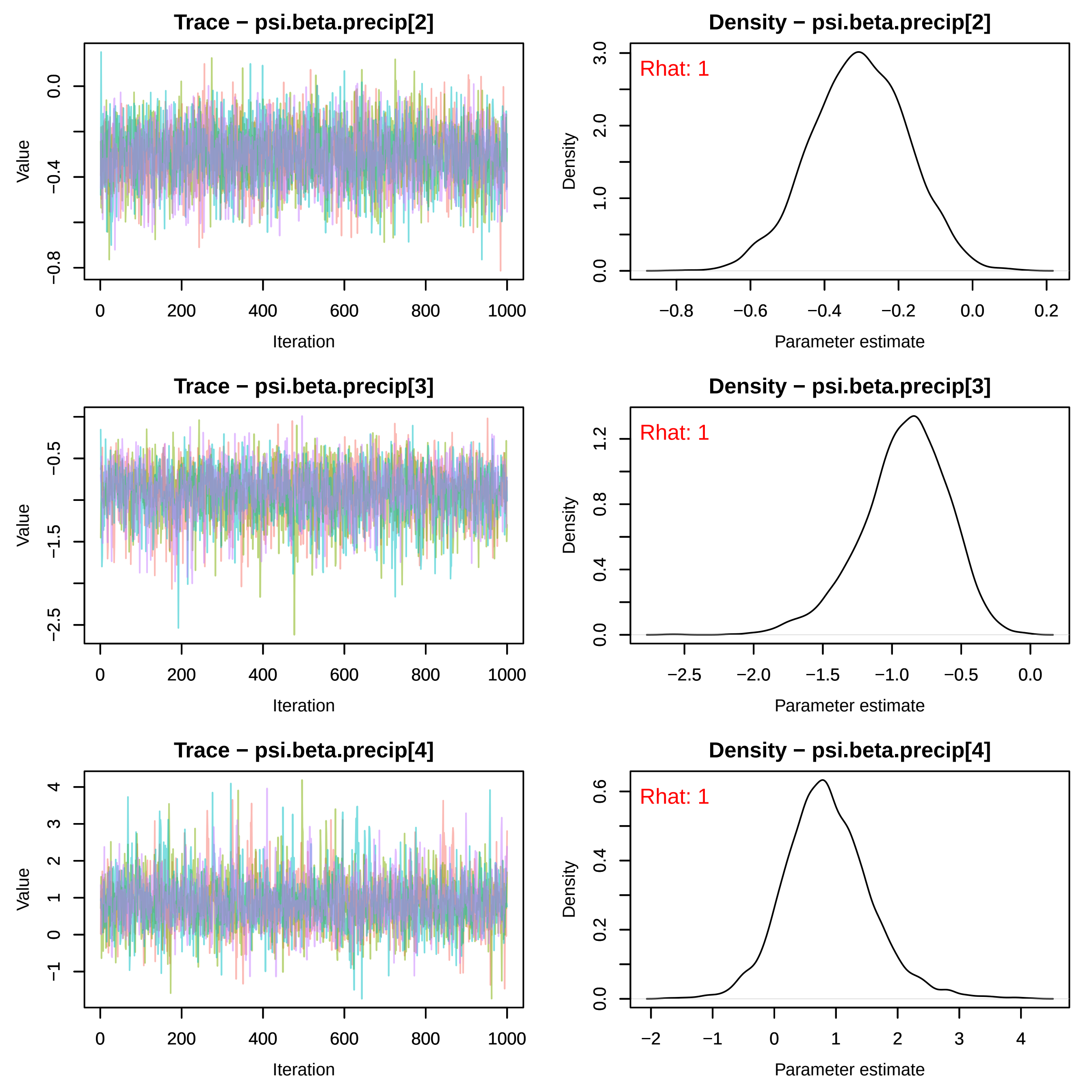
Trace − psi.beta.precip[2]
Density − psi.beta.precip[2]
Rhat: 1
Trace − psi.beta.precip[3]
Density − psi.beta.precip[3]
Rhat: 1
Trace − psi.beta.precip[4]
Density − psi.beta.precip[4]
Rhat: 1
Value
Iteration
Density
Parameter estimate

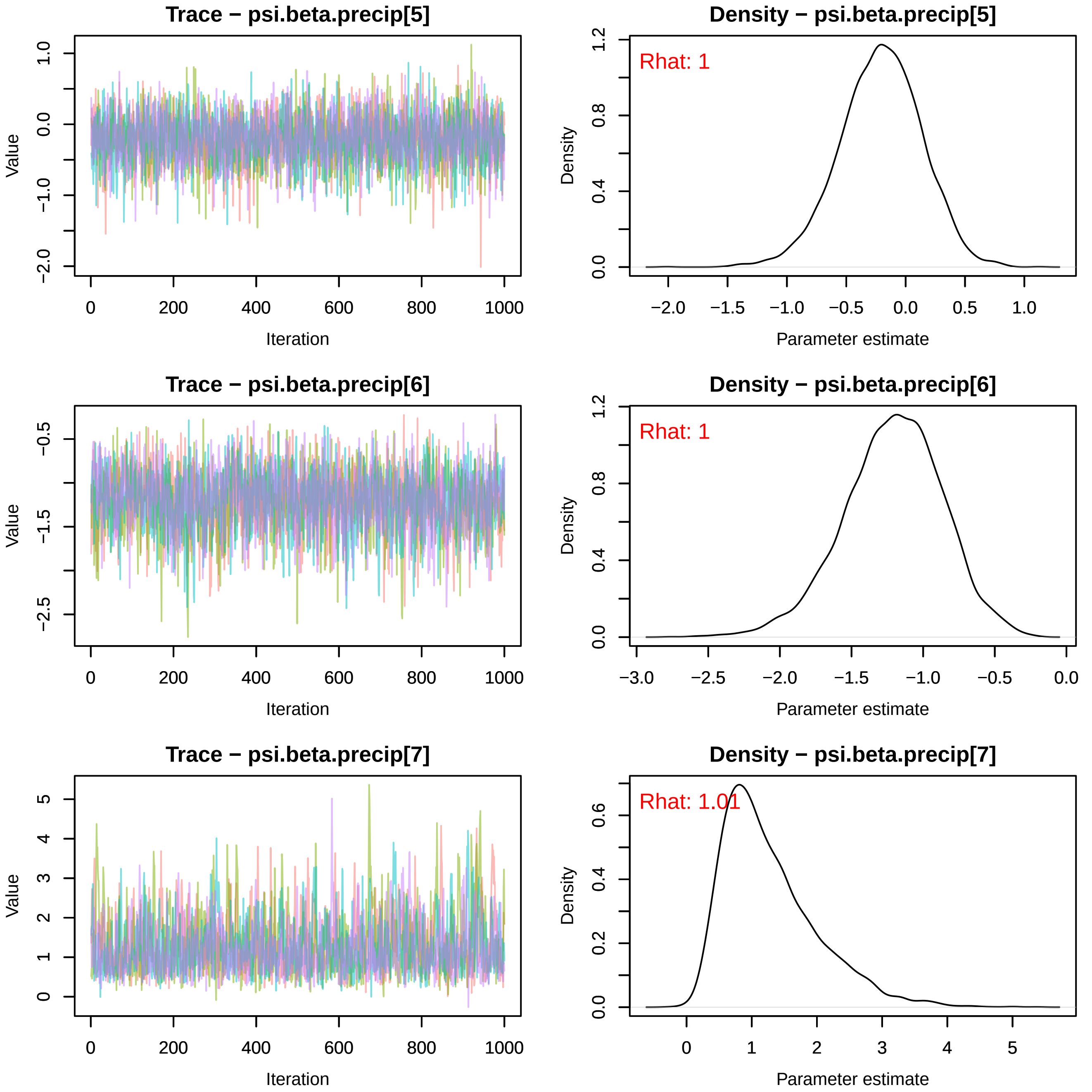
Trace − psi.beta.precip[5]
Value
Iteration
0
200
400
600
800
1000
−2.0
−1.0
0.0
1.0
Density − psi.beta.precip[5]
Rhat: 1
Density
Parameter estimate
−2.0
−1.5
−1.0
−0.5
0.0
0.5
1.0
0.0
0.4
0.8
1.2
Trace − psi.beta.precip[6]
Value
Iteration
−2.5
−1.5
−0.5
Density − psi.beta.precip[6]
Rhat: 1
Density
Parameter estimate
−3.0
−2.5
−2.0
−1.5
−1.0
−0.5
0.0
Trace − psi.beta.precip[7]
Value
Iteration
0
1
2
3
4
5
Density − psi.beta.precip[7]
Rhat: 1.01
Density
Parameter estimate
0.0
0.2
0.4
0.6

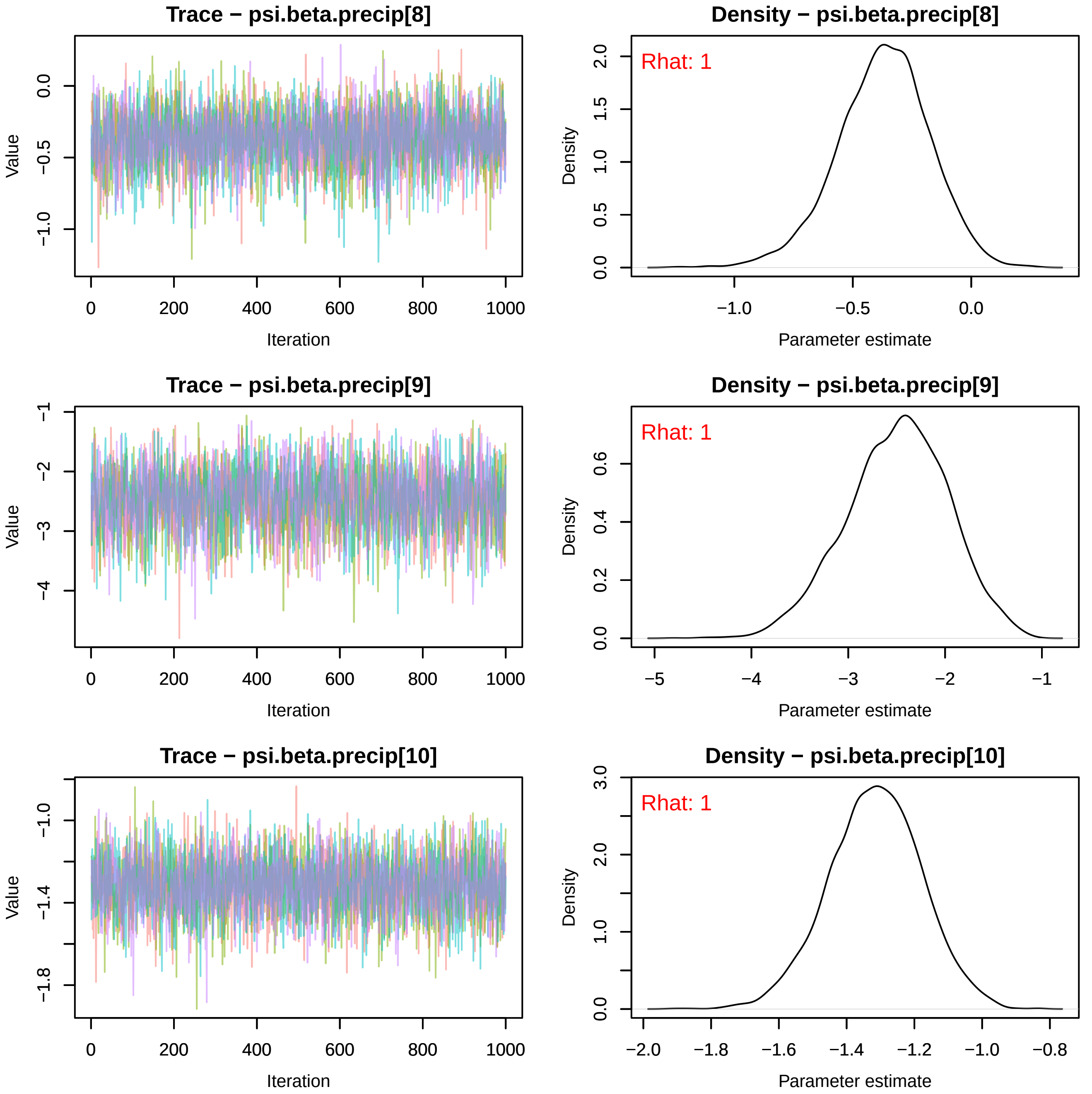

Trace − psi.beta.precip[8]
Value
Iteration
Density − psi.beta.precip[8]
Rhat: 1
Density
Parameter estimate
Trace − psi.beta.precip[9]
Value
Iteration
Density − psi.beta.precip[9]
Rhat: 1
Density
Parameter estimate
Trace − psi.beta.precip[10]
Value
Iteration
Density − psi.beta.precip[10]
Rhat: 1
Density
Parameter estimate

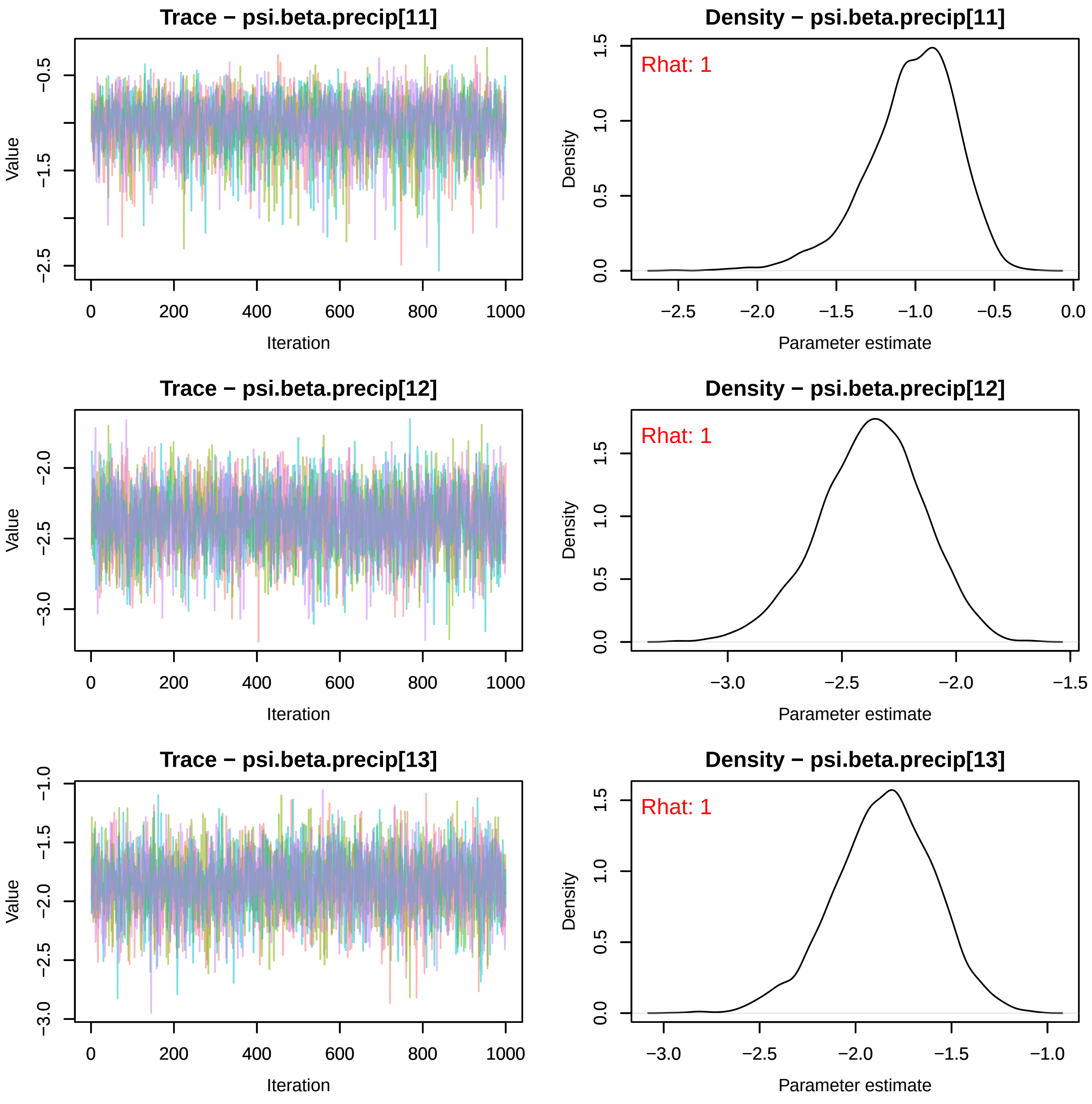

Trace − psi.beta.precip[11]
Density − psi.beta.precip[11]
Rhat: 1
Trace − psi.beta.precip[12]
Density − psi.beta.precip[12]
Rhat: 1
Trace − psi.beta.precip[13]
Density − psi.beta.precip[13]
Rhat: 1
Value
Density
Iteration
Parameter estimate

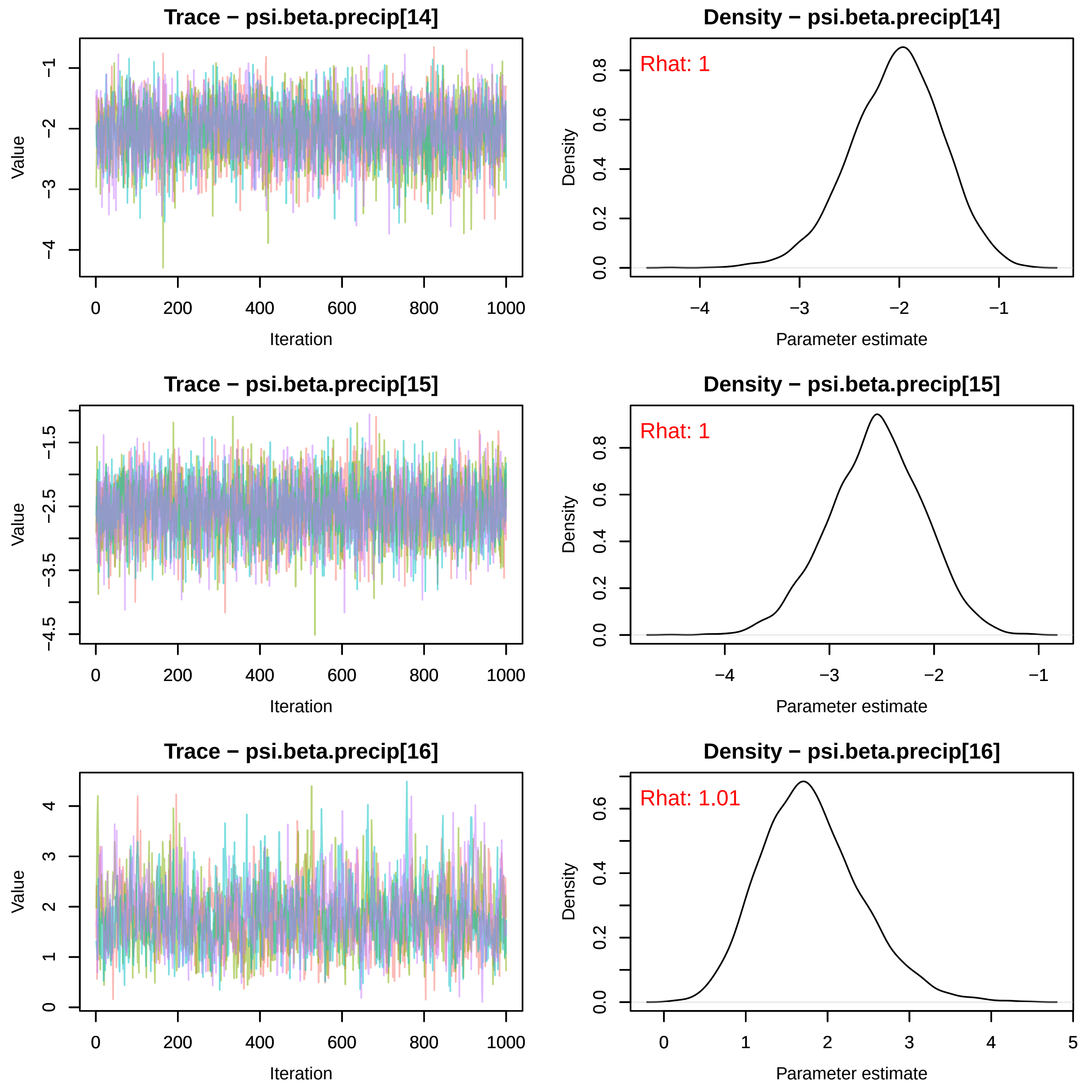
Trace − psi.beta.precip[14]
Value
Iteration
Density − psi.beta.precip[14]
Rhat: 1
Density
Parameter estimate
Trace − psi.beta.precip[15]
Value
Iteration
Density − psi.beta.precip[15]
Rhat: 1
Density
Parameter estimate
Trace − psi.beta.precip[16]
Value
Iteration
Density − psi.beta.precip[16]
Rhat: 1.01
Density
Parameter estimate

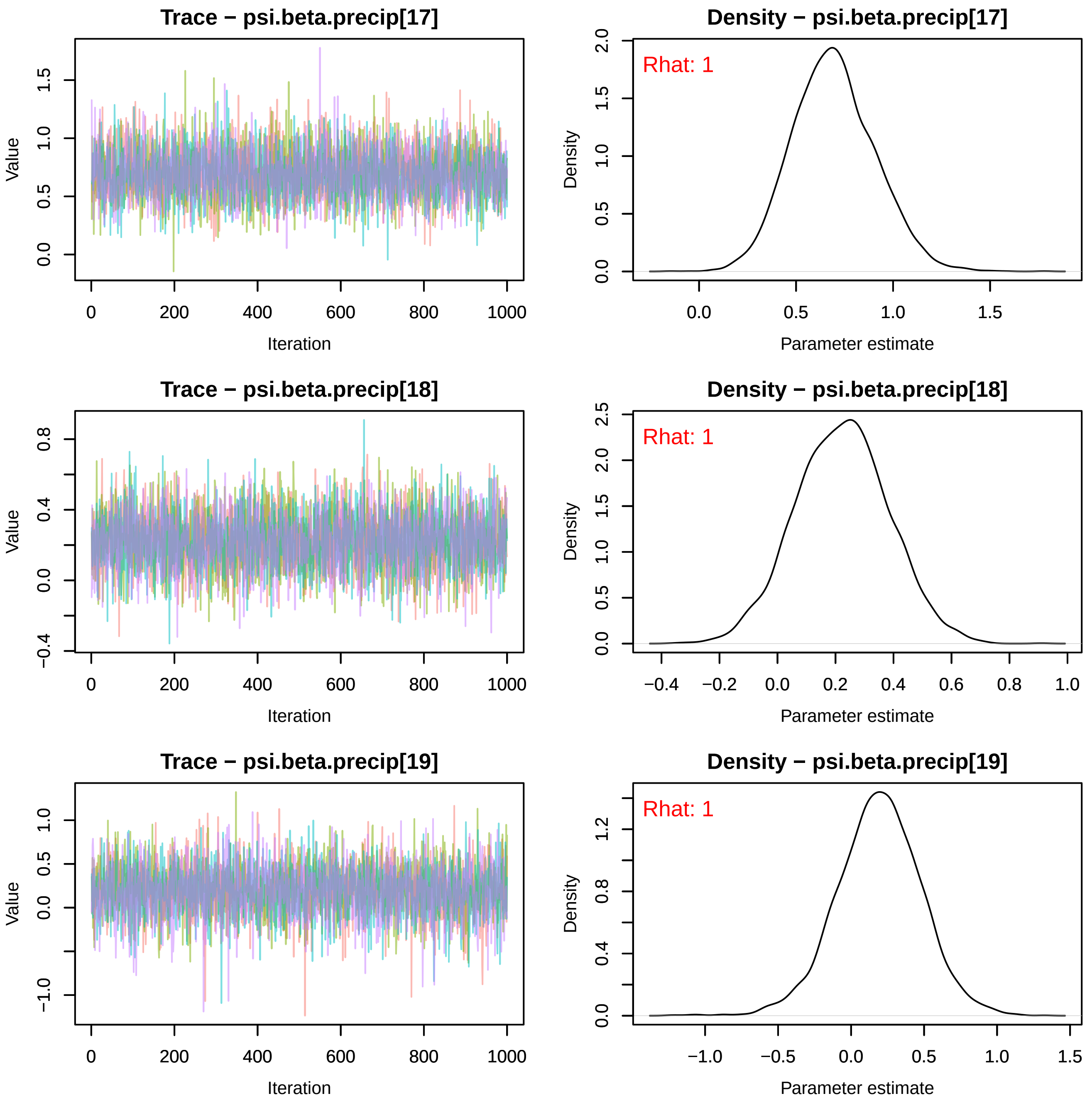
Trace − psi.beta.precip[17]
Value
Iteration
Density − psi.beta.precip[17]
Rhat: 1
Density
Parameter estimate
Trace − psi.beta.precip[18]
Value
Iteration
Density − psi.beta.precip[18]
Rhat: 1
Density
Parameter estimate
Trace − psi.beta.precip[19]
Value
Iteration
Density − psi.beta.precip[19]
Rhat: 1
Density
Parameter estimate

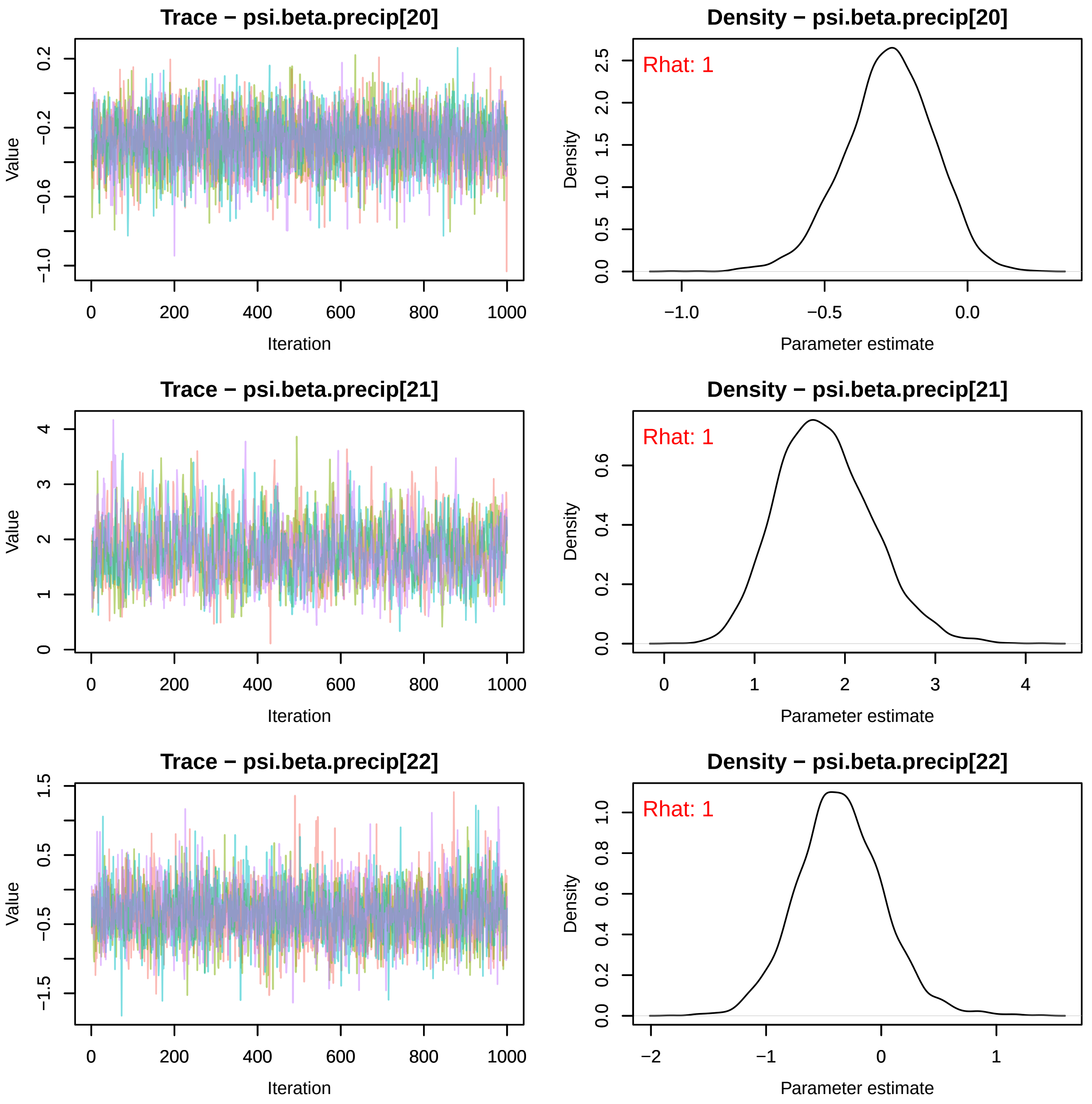
Trace − psi.beta.precip[20]
Density − psi.beta.precip[20]
Rhat: 1
Trace − psi.beta.precip[21]
Density − psi.beta.precip[21]
Rhat: 1
Trace − psi.beta.precip[22]
Density − psi.beta.precip[22]
Rhat: 1
Value
Iteration
Density
Parameter estimate
0
200
400
600
800
1000
−1.0
−0.5
0.0
0.5
1.0
1.5
2.0
2.5
0.2
−0.2
−0.6
1
2
3
4
0.2
0.4
0.6
0.8
−2
−1
−1.5

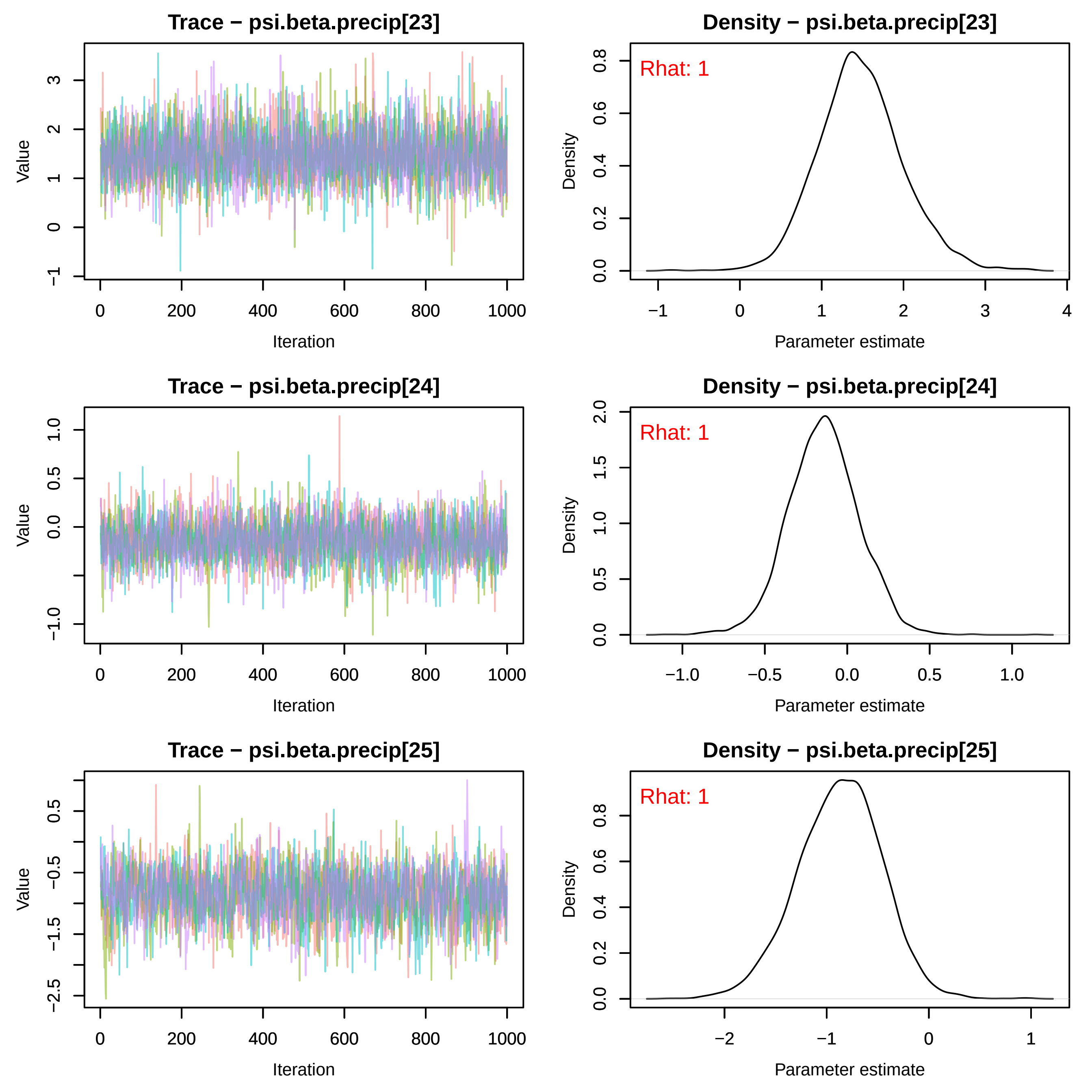
Trace − psi.beta.precip[23]
Value
Iteration
0
200
400
600
800
1000
−1
0
1
2
3
Density − psi.beta.precip[23]
Rhat: 1
Density
Parameter estimate
0.0
0.2
0.4
0.6
0.8
−1
0
1
2
3
4
Trace − psi.beta.precip[24]
Value
Iteration
−1.0
0.0
0.5
1.0
Density − psi.beta.precip[24]
Rhat: 1
Density
Parameter estimate
0.0
0.5
1.0
1.5
2.0
−1.0
−0.5
0.0
0.5
1.0
Trace − psi.beta.precip[25]
Value
Iteration
−2.5
−1.5
−0.5
0.5
Density − psi.beta.precip[25]
Rhat: 1
Density
Parameter estimate
0.0
0.2
0.4
0.6
0.8
−2
−1
0
1

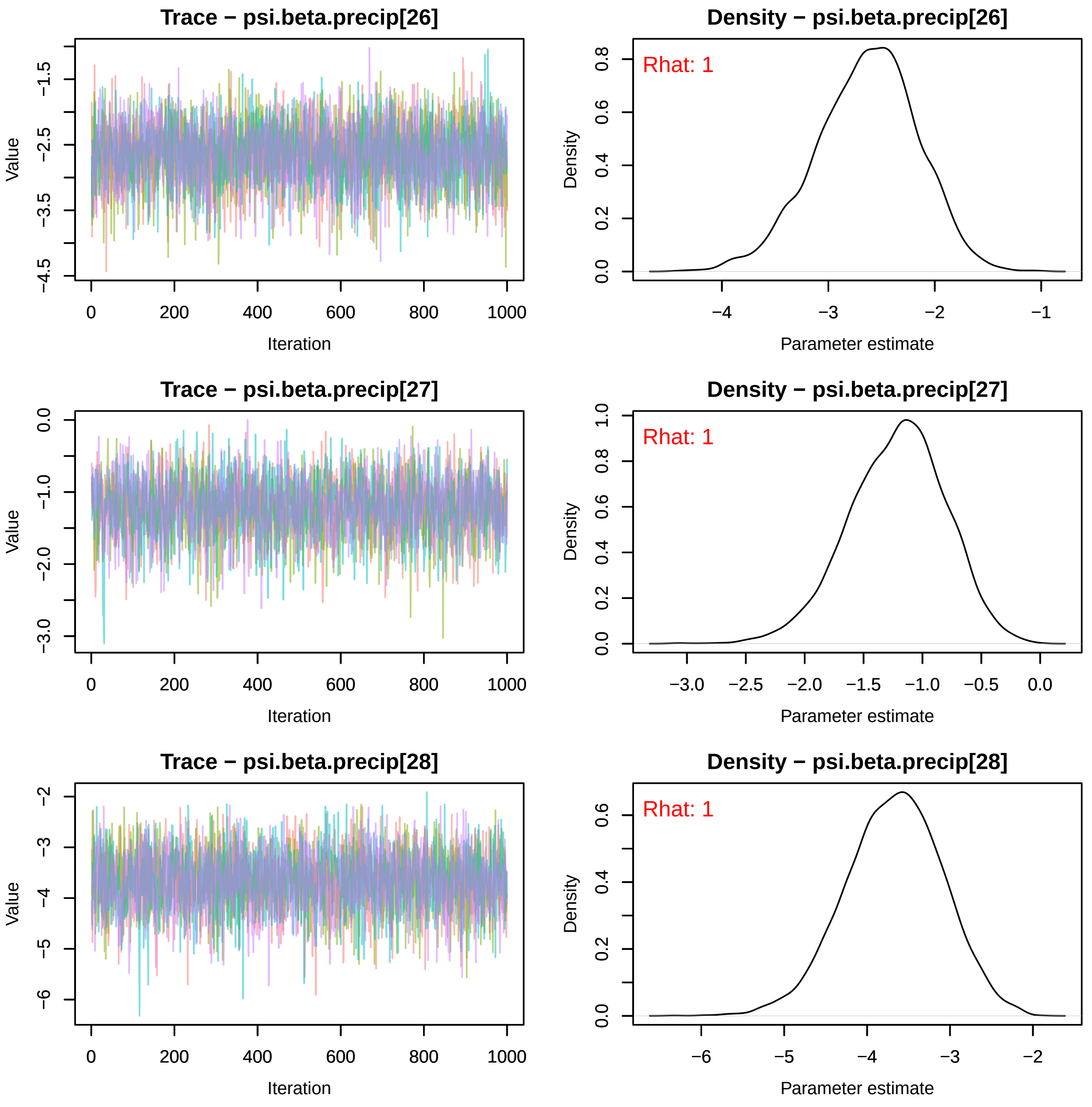
Trace − psi.beta.precip[26]
Value
Iteration
Density − psi.beta.precip[26]
Rhat: 1
Density
Parameter estimate
Trace − psi.beta.precip[27]
Value
Iteration
Density − psi.beta.precip[27]
Rhat: 1
Density
Parameter estimate
Trace − psi.beta.precip[28]
Value
Iteration
Density − psi.beta.precip[28]
Rhat: 1
Density
Parameter estimate

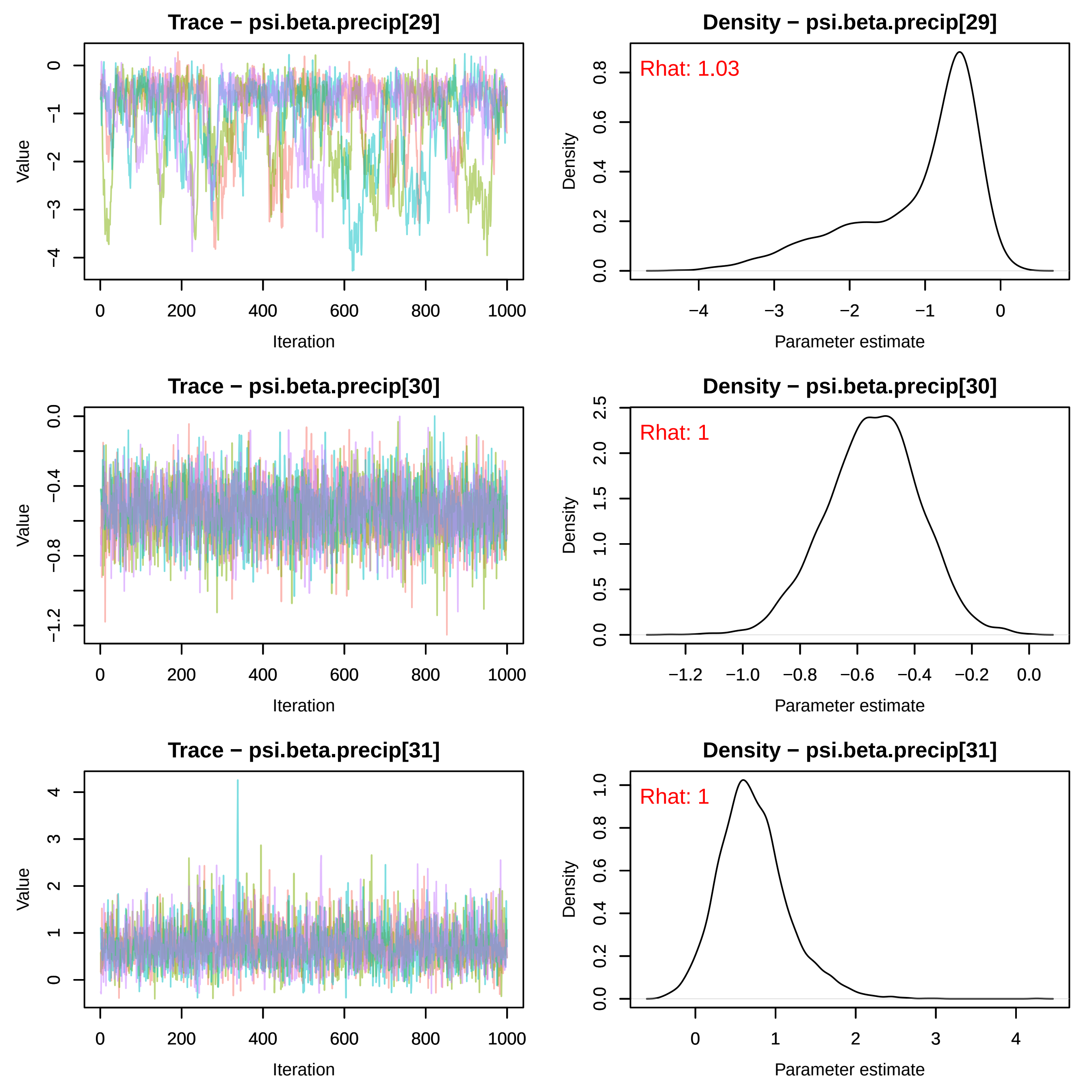

Trace − psi.beta.precip[29]
Value
Iteration
Density − psi.beta.precip[29]
Rhat: 1.03
Density
Parameter estimate
Trace − psi.beta.precip[30]
Value
Iteration
Density − psi.beta.precip[30]
Rhat: 1
Density
Parameter estimate
Trace − psi.beta.precip[31]
Value
Iteration
Density − psi.beta.precip[31]
Rhat: 1
Density
Parameter estimate

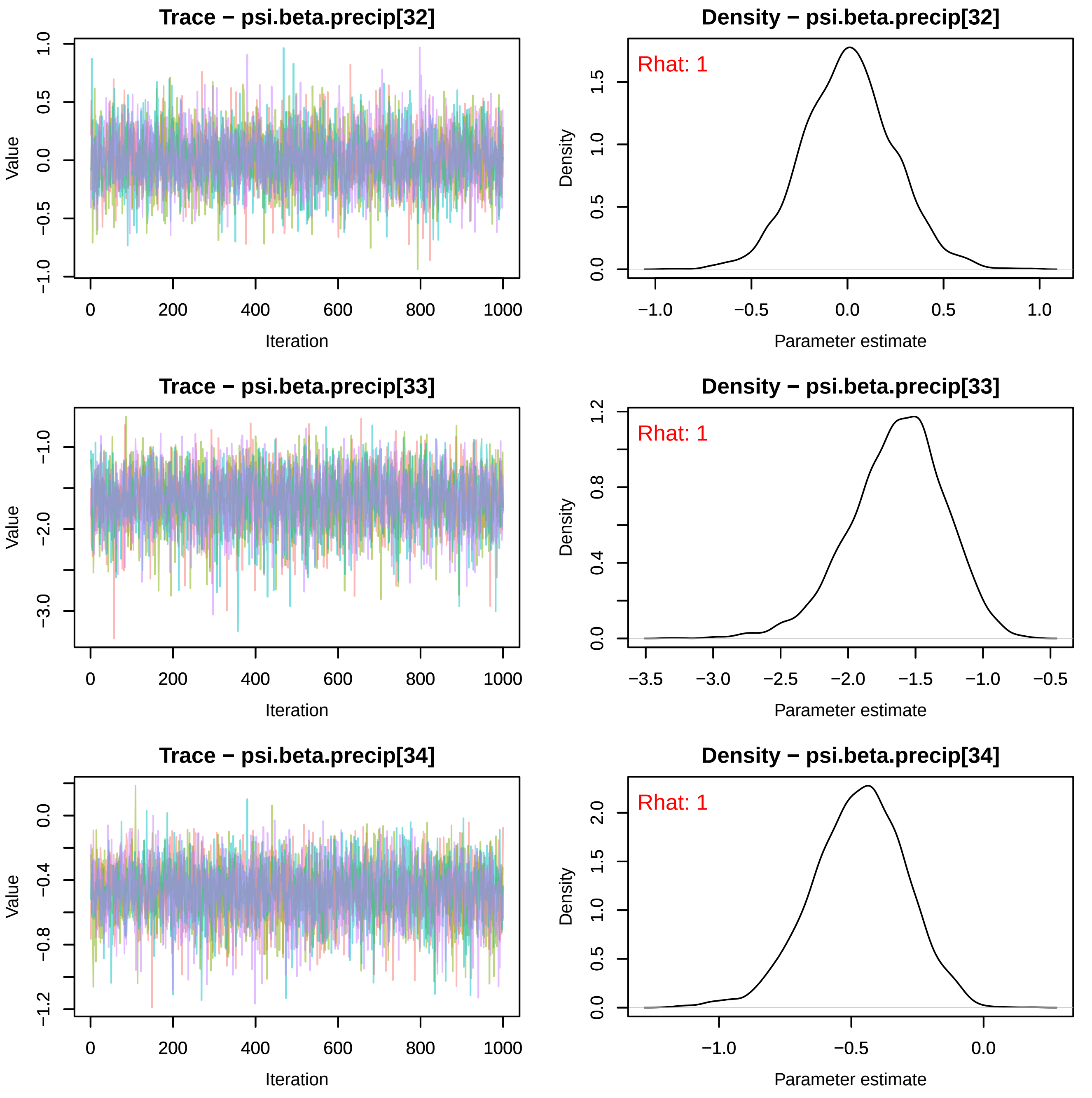
Trace − psi.beta.precip[32]
Density − psi.beta.precip[32]
Rhat: 1
Trace − psi.beta.precip[33]
Density − psi.beta.precip[33]
Rhat: 1
Trace − psi.beta.precip[34]
Density − psi.beta.precip[34]
Rhat: 1
Value
Iteration
Density
Parameter estimate

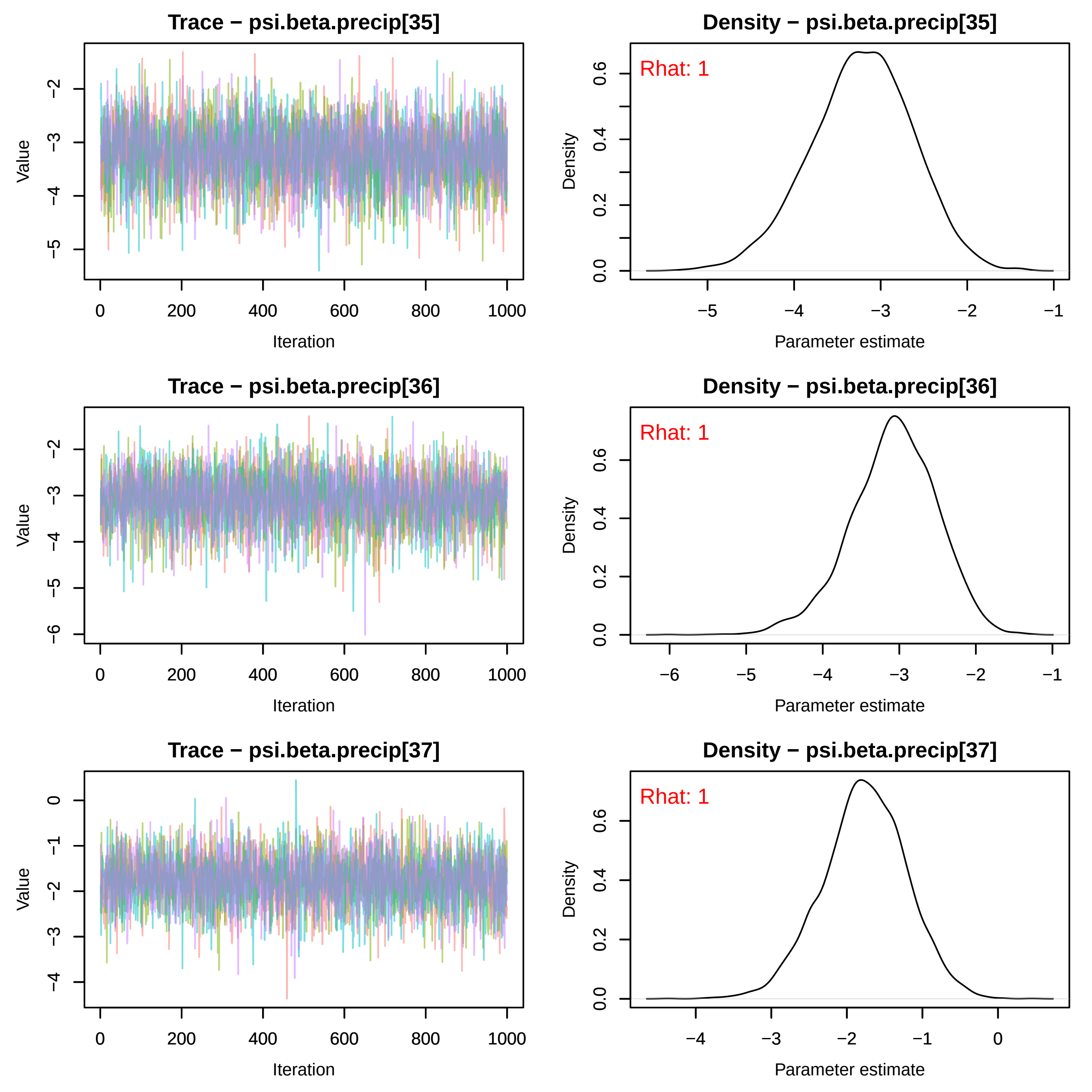
Trace − psi.beta.precip[35]
Value
Iteration
Density − psi.beta.precip[35]
Rhat: 1
Density
Parameter estimate
Trace − psi.beta.precip[36]
Value
Iteration
Density − psi.beta.precip[36]
Rhat: 1
Density
Parameter estimate
Trace − psi.beta.precip[37]
Value
Iteration
Density − psi.beta.precip[37]
Rhat: 1
Density
Parameter estimate

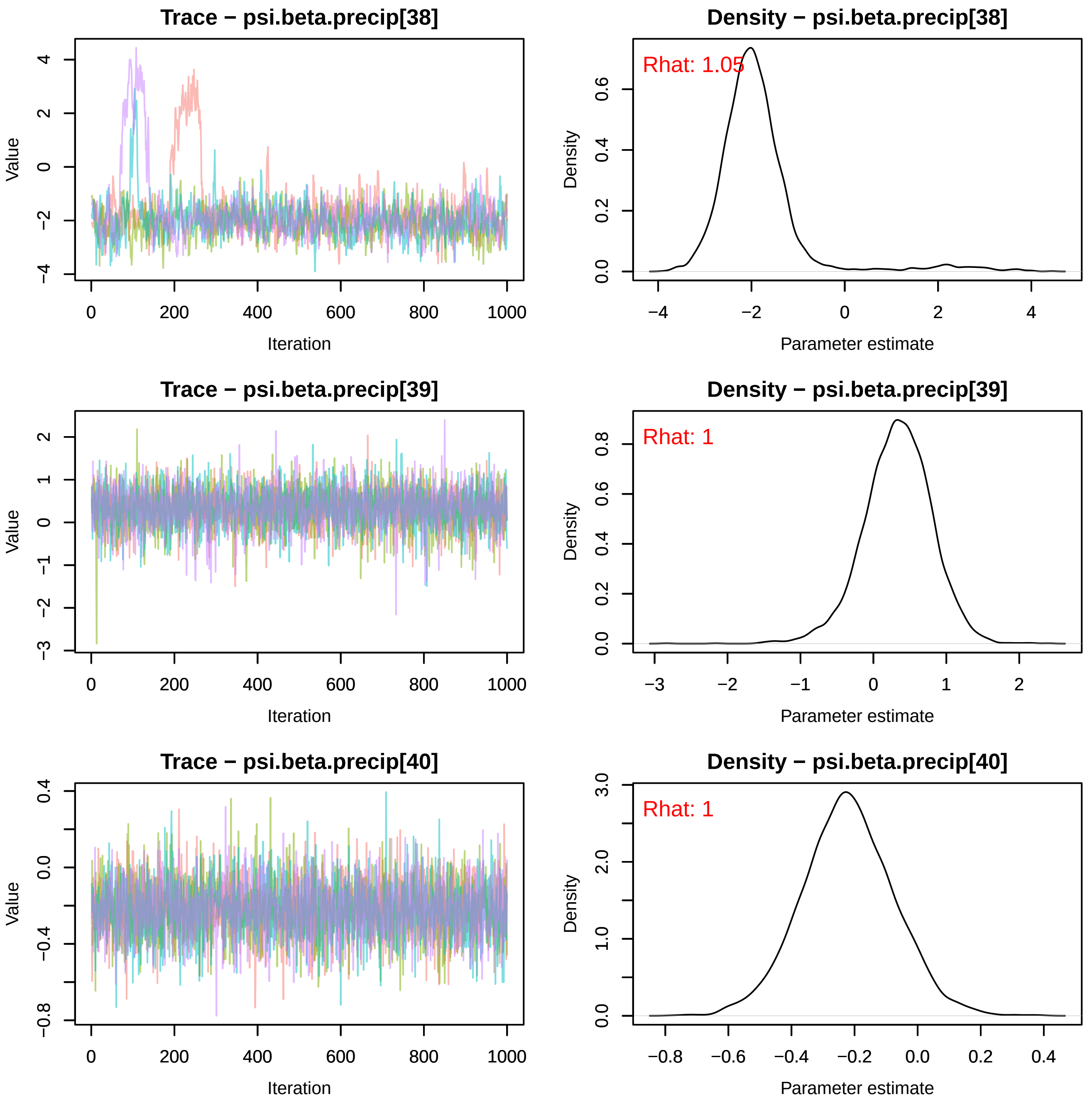

Trace − psi.beta.precip[38]
Density − psi.beta.precip[38]
Rhat: 1.05
Trace − psi.beta.precip[39]
Density − psi.beta.precip[39]
Rhat: 1
Trace − psi.beta.precip[40]
Density − psi.beta.precip[40]
Rhat: 1
Value
Iteration
Density
Parameter estimate

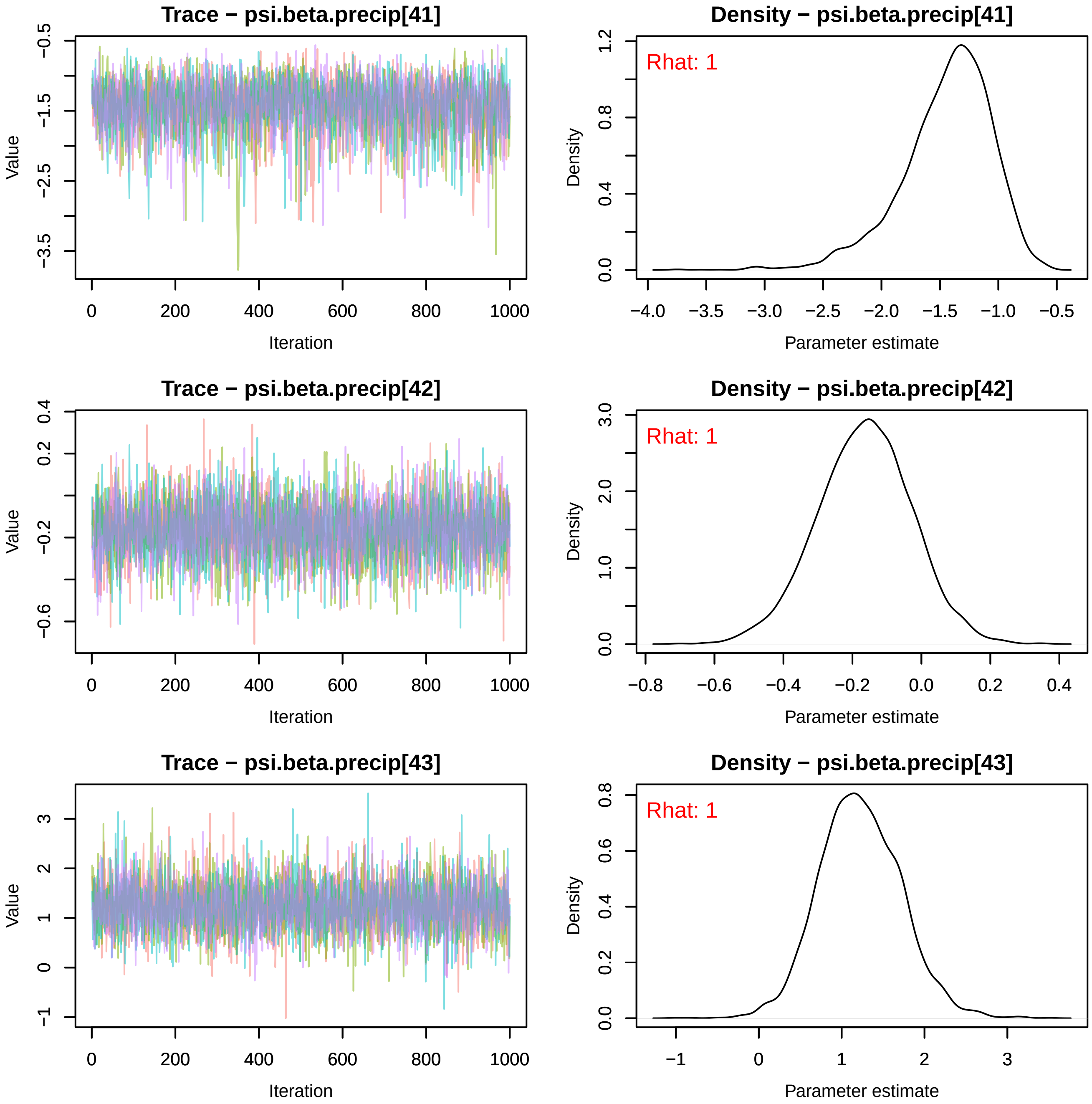
Trace − psi.beta.precip[41]
Value
Iteration
Density − psi.beta.precip[41]
Rhat: 1
Density
Parameter estimate
Trace − psi.beta.precip[42]
Value
Iteration
Density − psi.beta.precip[42]
Rhat: 1
Density
Parameter estimate
Trace − psi.beta.precip[43]
Value
Iteration
Density − psi.beta.precip[43]
Rhat: 1
Density
Parameter estimate

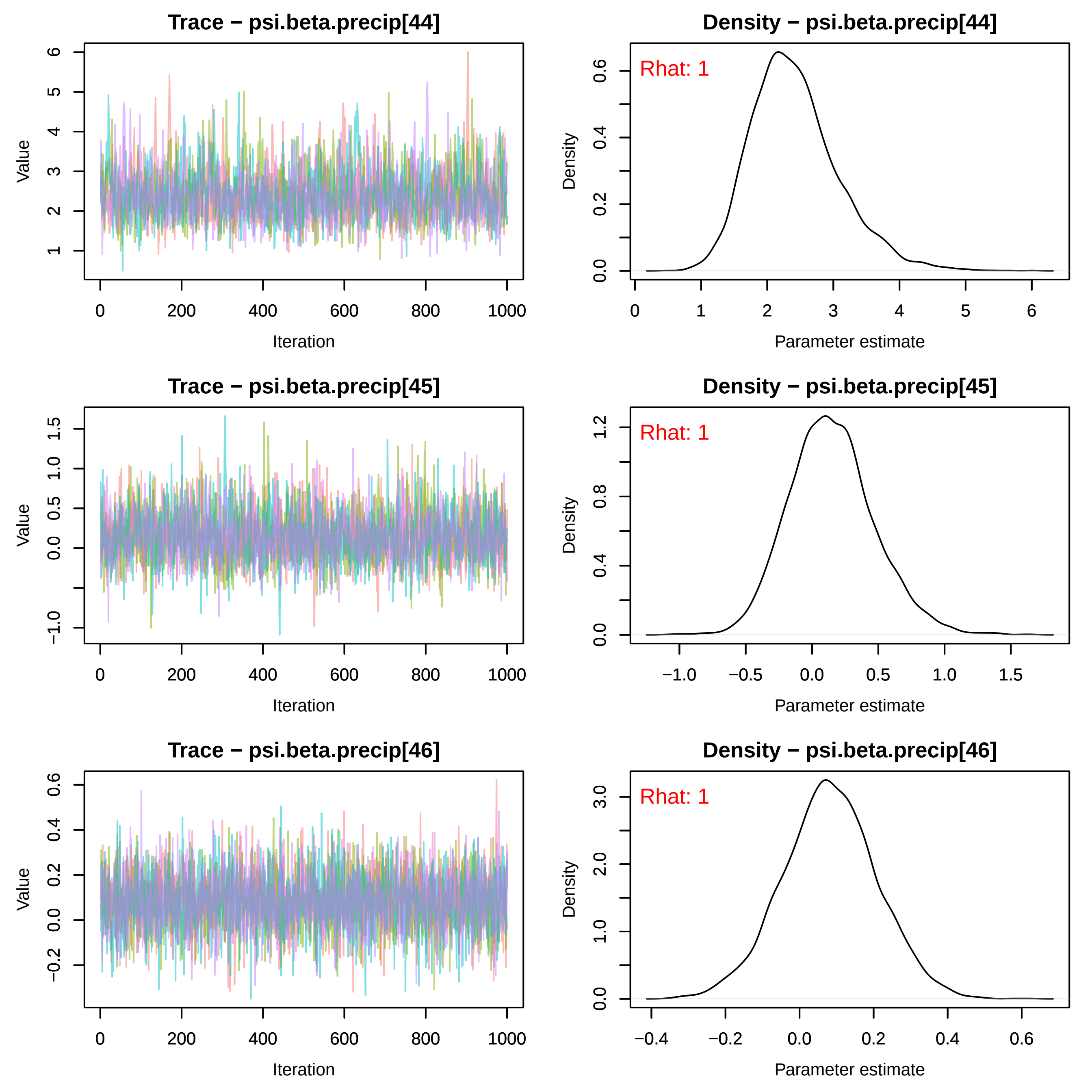
Trace − psi.beta.precip[44]
Density − psi.beta.precip[44]
Rhat: 1
Trace − psi.beta.precip[45]
Density − psi.beta.precip[45]
Rhat: 1
Trace − psi.beta.precip[46]
Density − psi.beta.precip[46]
Rhat: 1
Value
Iteration
Density
Parameter estimate

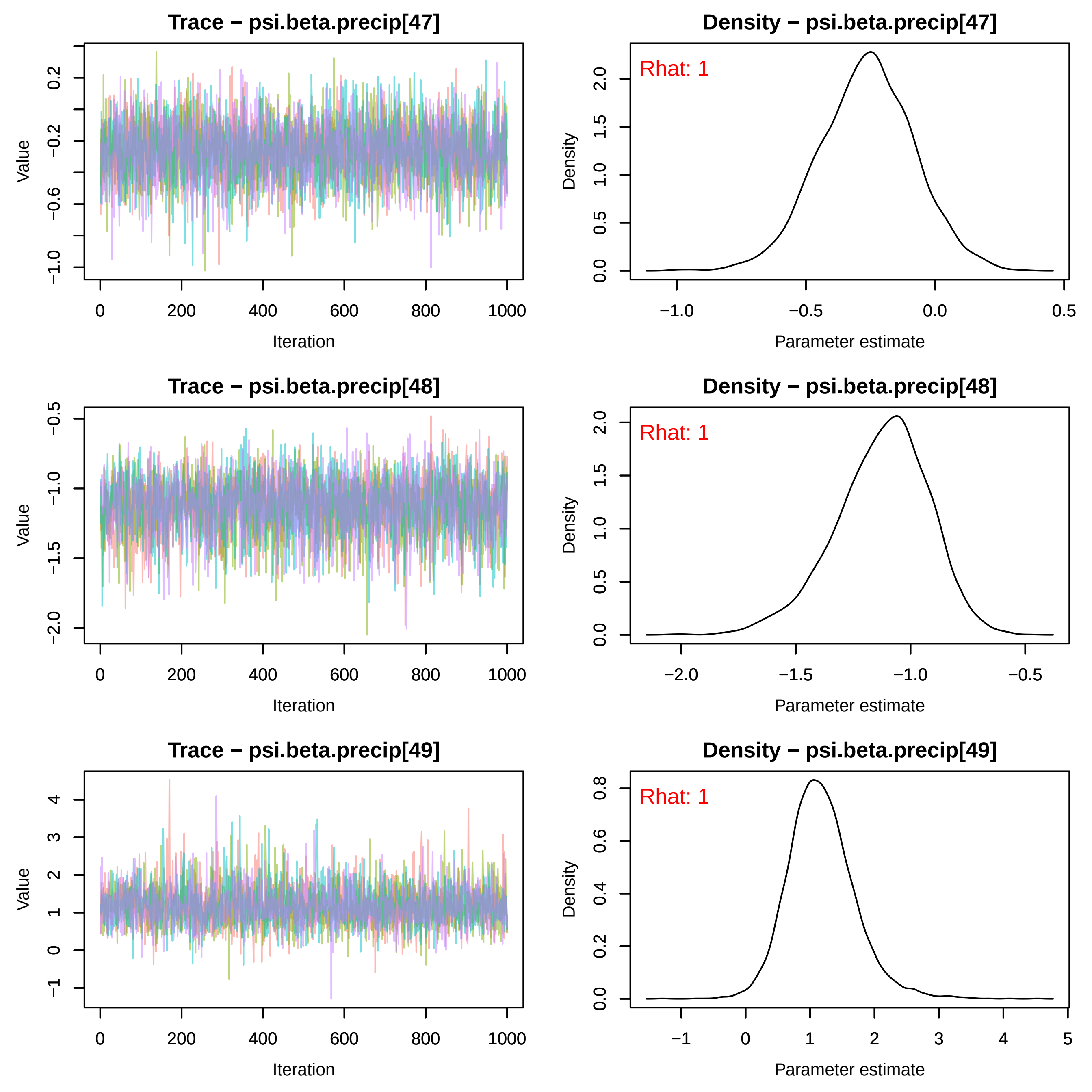
Trace − psi.beta.precip[47]
Value
Iteration
0.2
−0.2
−0.6
−1.0
0
200
400
600
800
1000
Density − psi.beta.precip[47]
Rhat: 1
Density
Parameter estimate
2.0
1.5
1.0
0.5
0.0
−1.0
−0.5
0.0
0.5
Trace − psi.beta.precip[48]
Value
Iteration
−0.5
−1.0
−1.5
−2.0
0
200
400
600
800
1000
Density − psi.beta.precip[48]
Rhat: 1
Density
Parameter estimate
2.0
1.5
1.0
0.5
0.0
−2.0
−1.5
−1.0
−0.5
Trace − psi.beta.precip[49]
Value
Iteration
4
3
2
1
0
−1
0
200
400
600
800
1000
Density − psi.beta.precip[49]
Rhat: 1
Density
Parameter estimate
0.8
0.6
0.4
0.2
0.0
−1
0
1
2
3
4
5

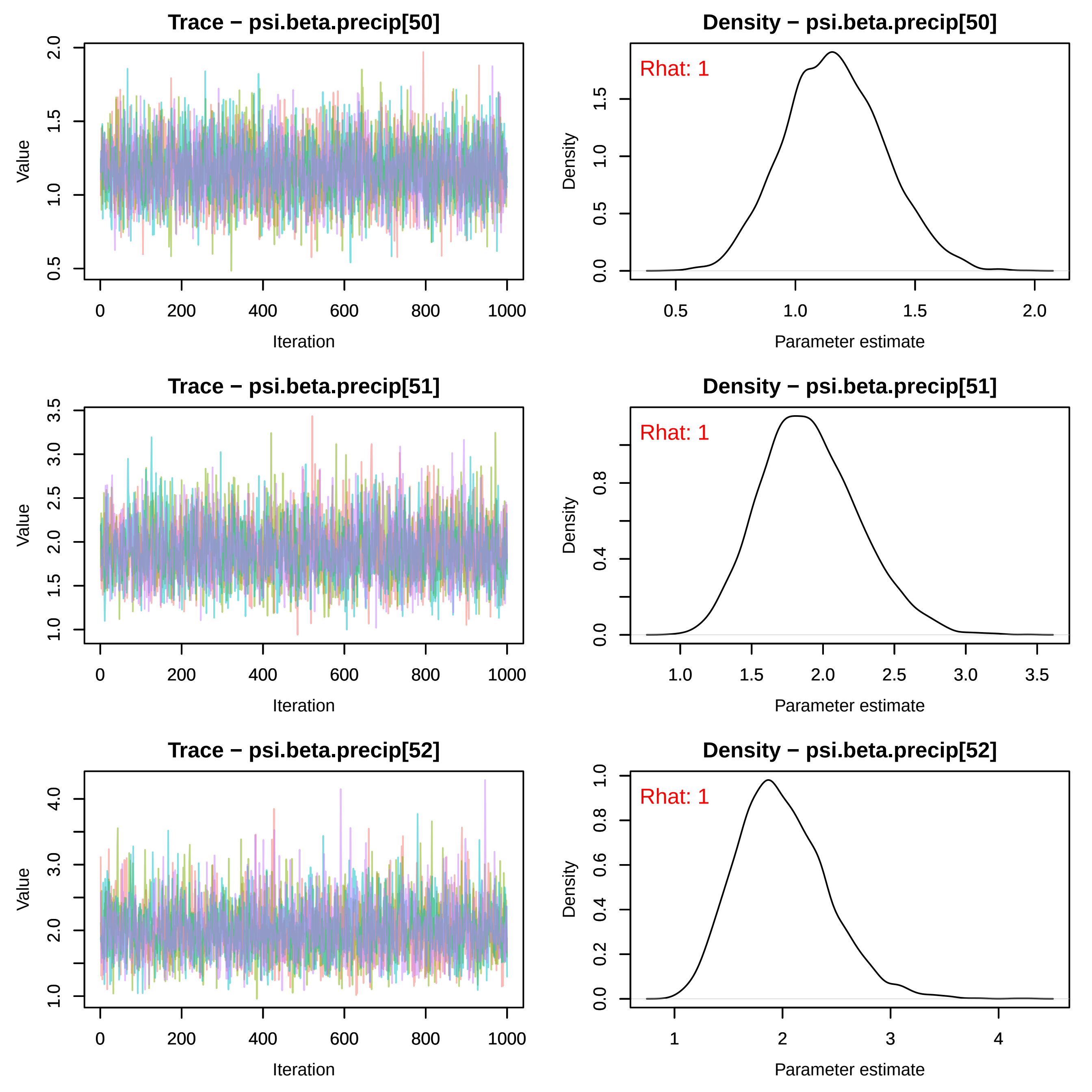
Trace − psi.beta.precip[50]
Value
Iteration
Density − psi.beta.precip[50]
Rhat: 1
Density
Parameter estimate
Trace − psi.beta.precip[51]
Value
Iteration
Density − psi.beta.precip[51]
Rhat: 1
Density
Parameter estimate
Trace − psi.beta.precip[52]
Value
Iteration
Density − psi.beta.precip[52]
Rhat: 1
Density
Parameter estimate

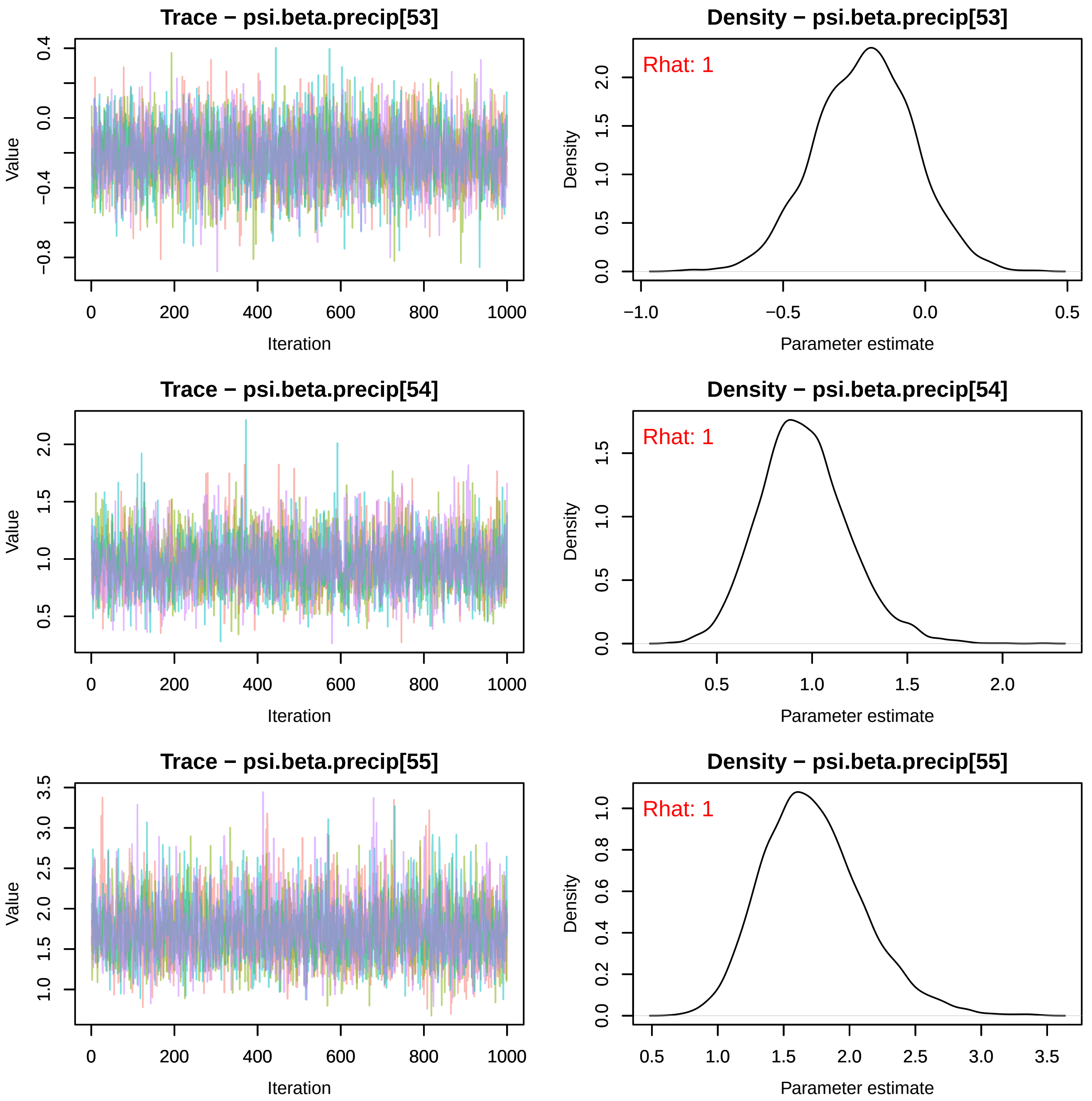
Trace − psi.beta.precip[53]
Density − psi.beta.precip[53]
Rhat: 1
Trace − psi.beta.precip[54]
Density − psi.beta.precip[54]
Rhat: 1
Trace − psi.beta.precip[55]
Density − psi.beta.precip[55]
Rhat: 1
Value
Iteration
Density
Parameter estimate

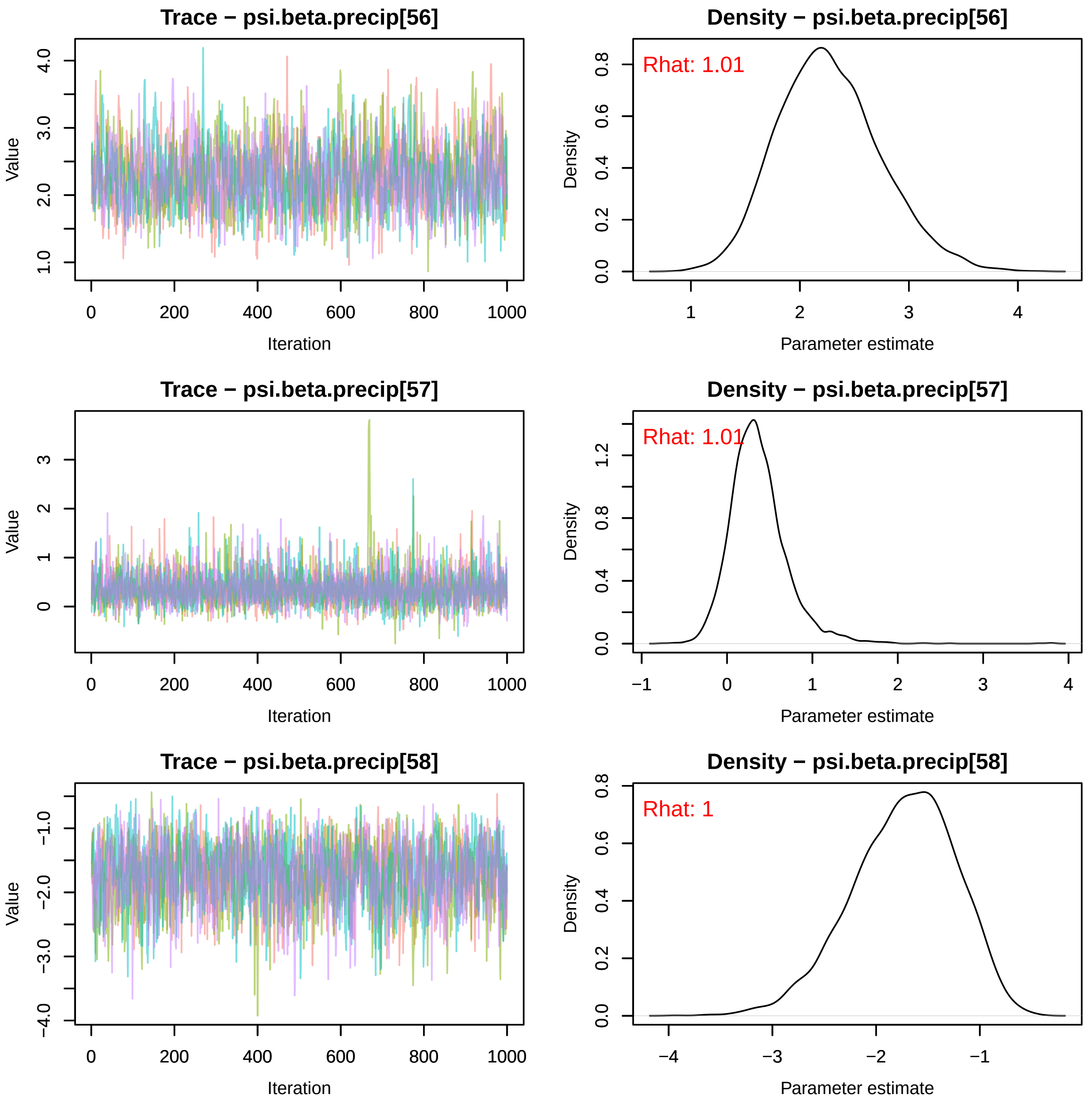
Trace − psi.beta.precip[56]
Density − psi.beta.precip[56]
Rhat: 1.01
Trace − psi.beta.precip[57]
Density − psi.beta.precip[57]
Rhat: 1.01
Trace − psi.beta.precip[58]
Density − psi.beta.precip[58]
Rhat: 1
Value
Iteration
Density
Parameter estimate

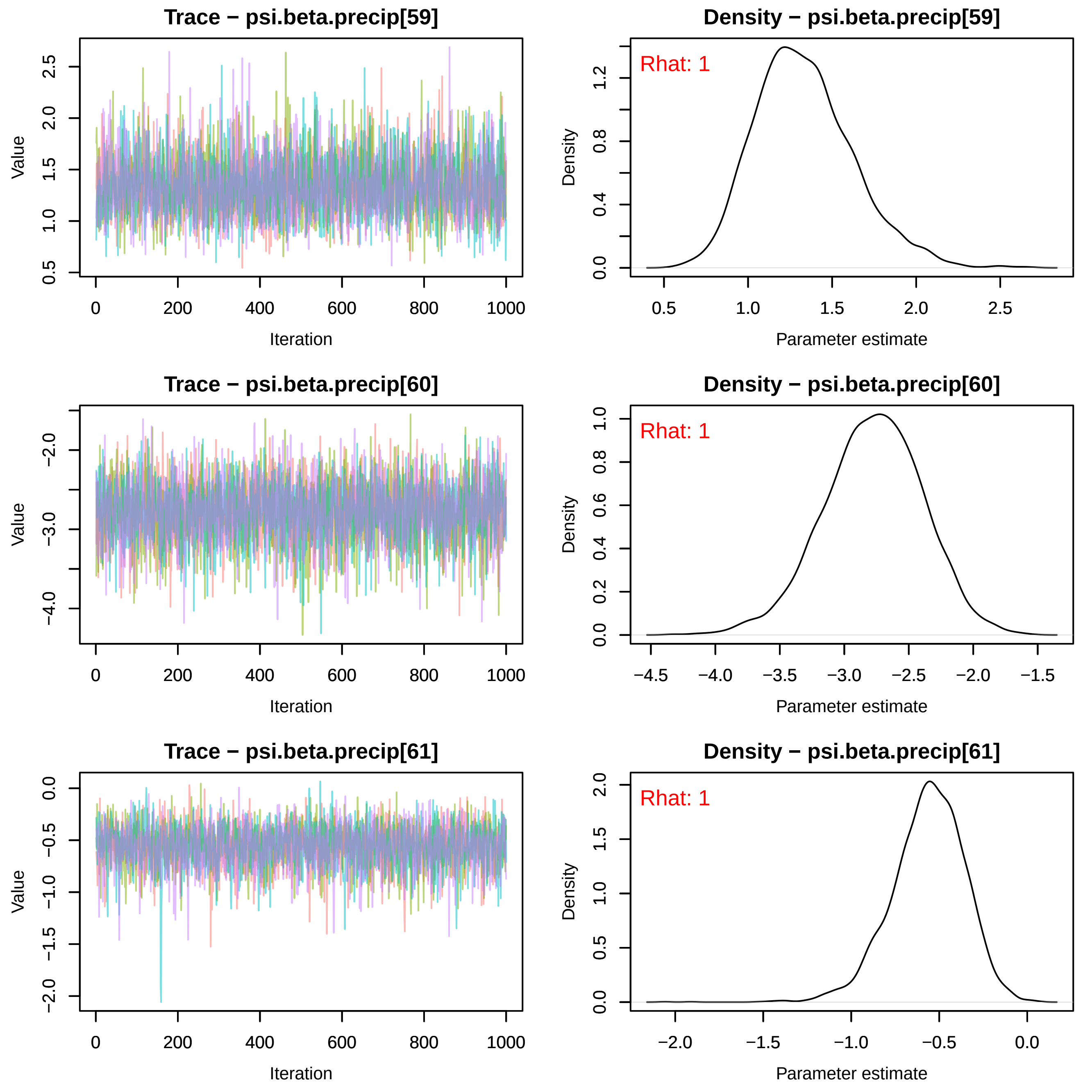
Trace − psi.beta.precip[59]
Density − psi.beta.precip[59]
Rhat: 1
Trace − psi.beta.precip[60]
Density − psi.beta.precip[60]
Rhat: 1
Trace − psi.beta.precip[61]
Density − psi.beta.precip[61]
Rhat: 1
Value
Iteration
Density
Parameter estimate

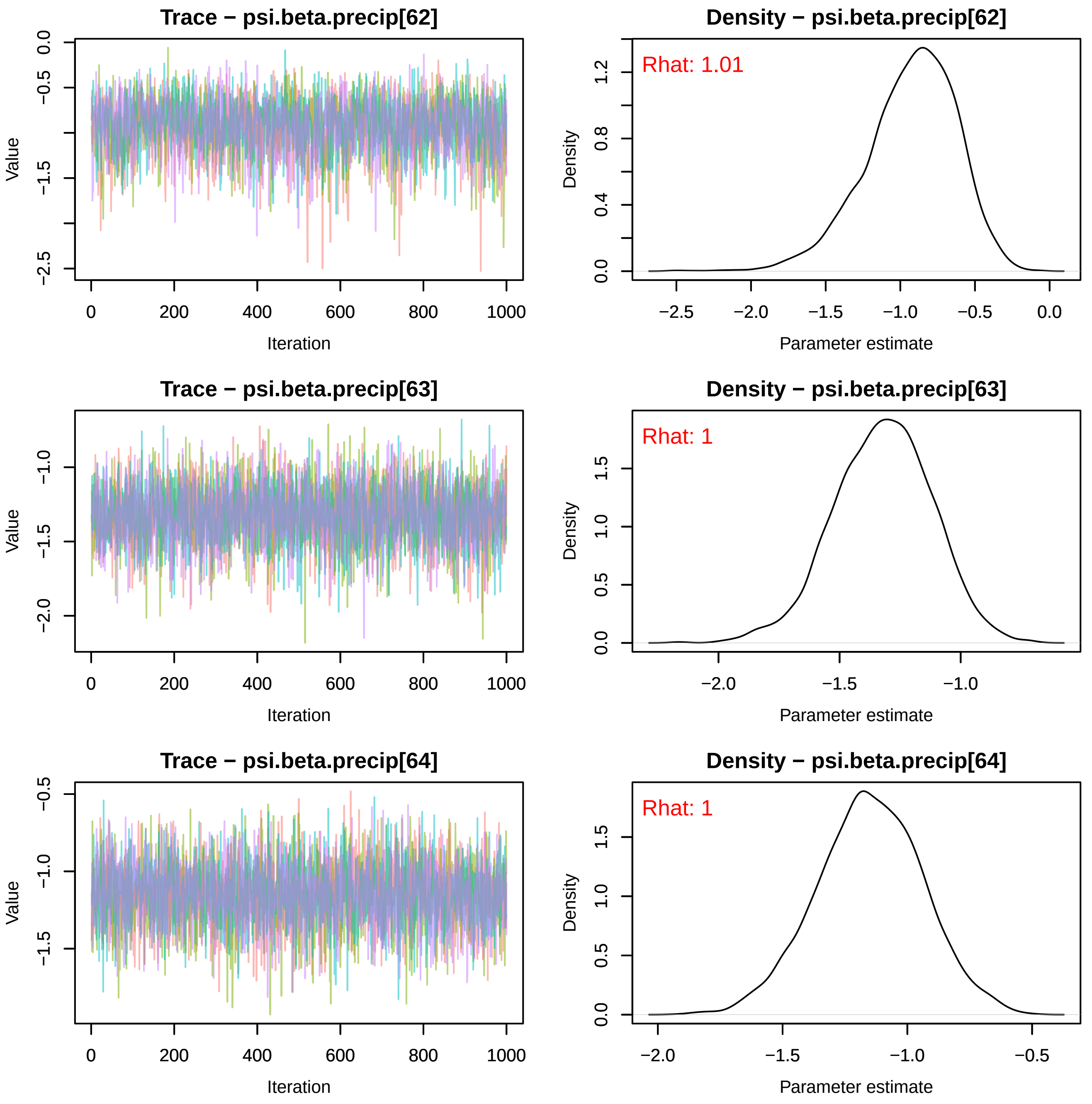
Trace − psi.beta.precip[62]
Value
Iteration
Density − psi.beta.precip[62]
Rhat: 1.01
Density
Parameter estimate
Trace − psi.beta.precip[63]
Value
Iteration
Density − psi.beta.precip[63]
Rhat: 1
Density
Parameter estimate
Trace − psi.beta.precip[64]
Value
Iteration
Density − psi.beta.precip[64]
Rhat: 1
Density
Parameter estimate

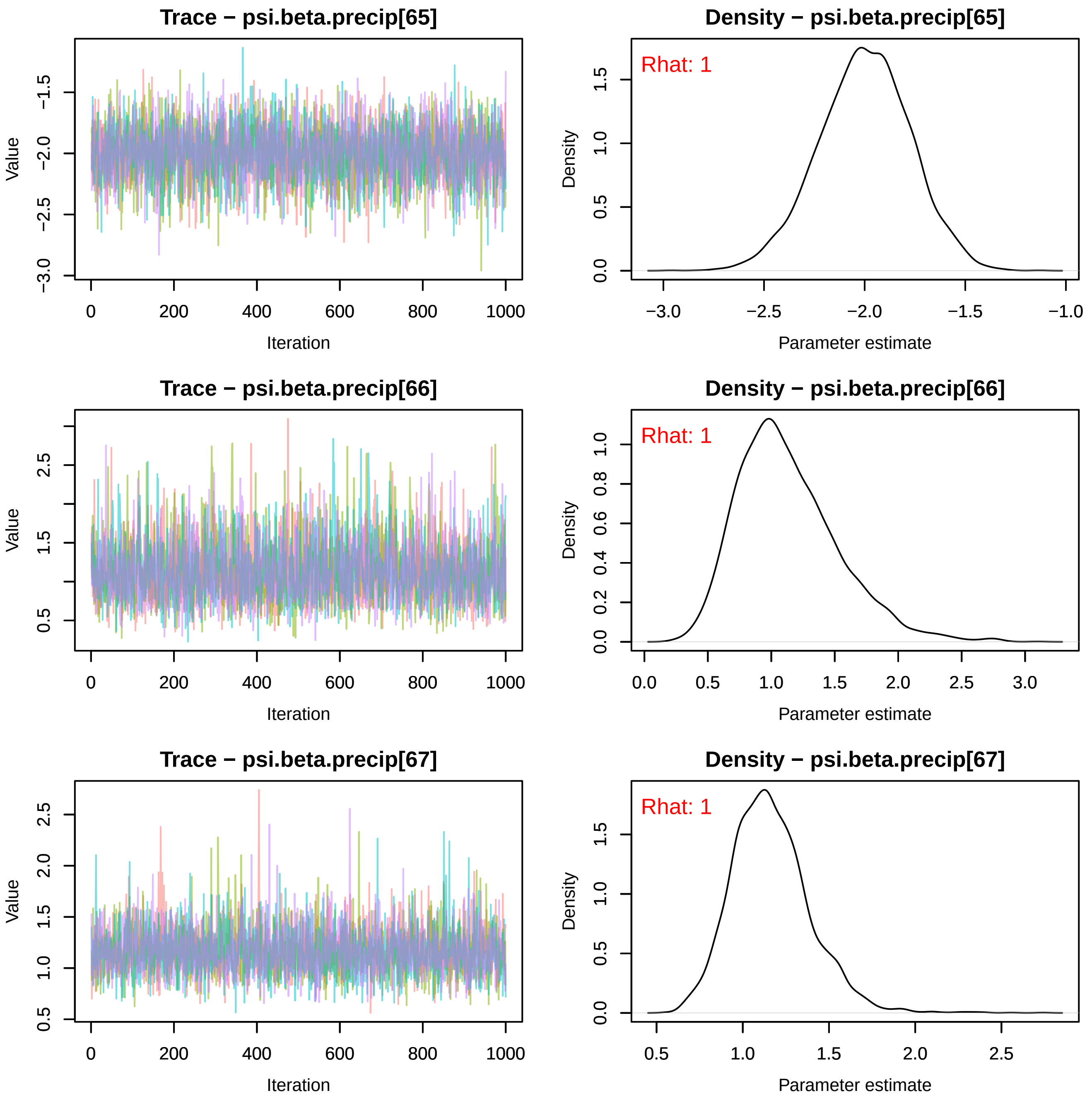
Trace − psi.beta.precip[65]
Density − psi.beta.precip[65]
Rhat: 1
Value
Iteration
Density
Parameter estimate
Trace − psi.beta.precip[66]
Density − psi.beta.precip[66]
Rhat: 1
Value
Iteration
Density
Parameter estimate
Trace − psi.beta.precip[67]
Density − psi.beta.precip[67]
Rhat: 1
Value
Iteration
Density
Parameter estimate

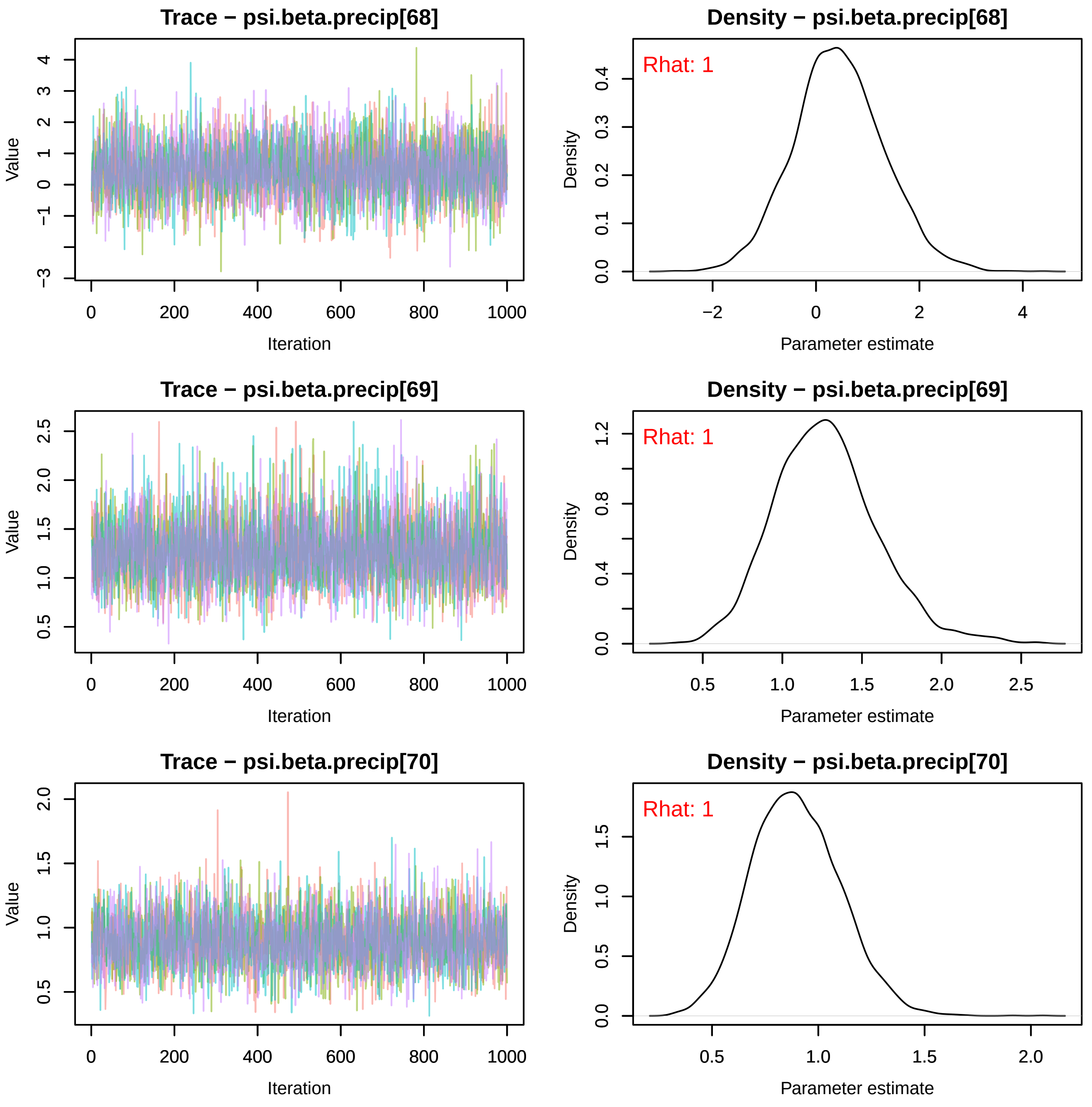
Trace − psi.beta.precip[68]
Density − psi.beta.precip[68]
Rhat: 1
Trace − psi.beta.precip[69]
Density − psi.beta.precip[69]
Rhat: 1
Trace − psi.beta.precip[70]
Density − psi.beta.precip[70]
Rhat: 1
Value
Iteration
Density
Parameter estimate

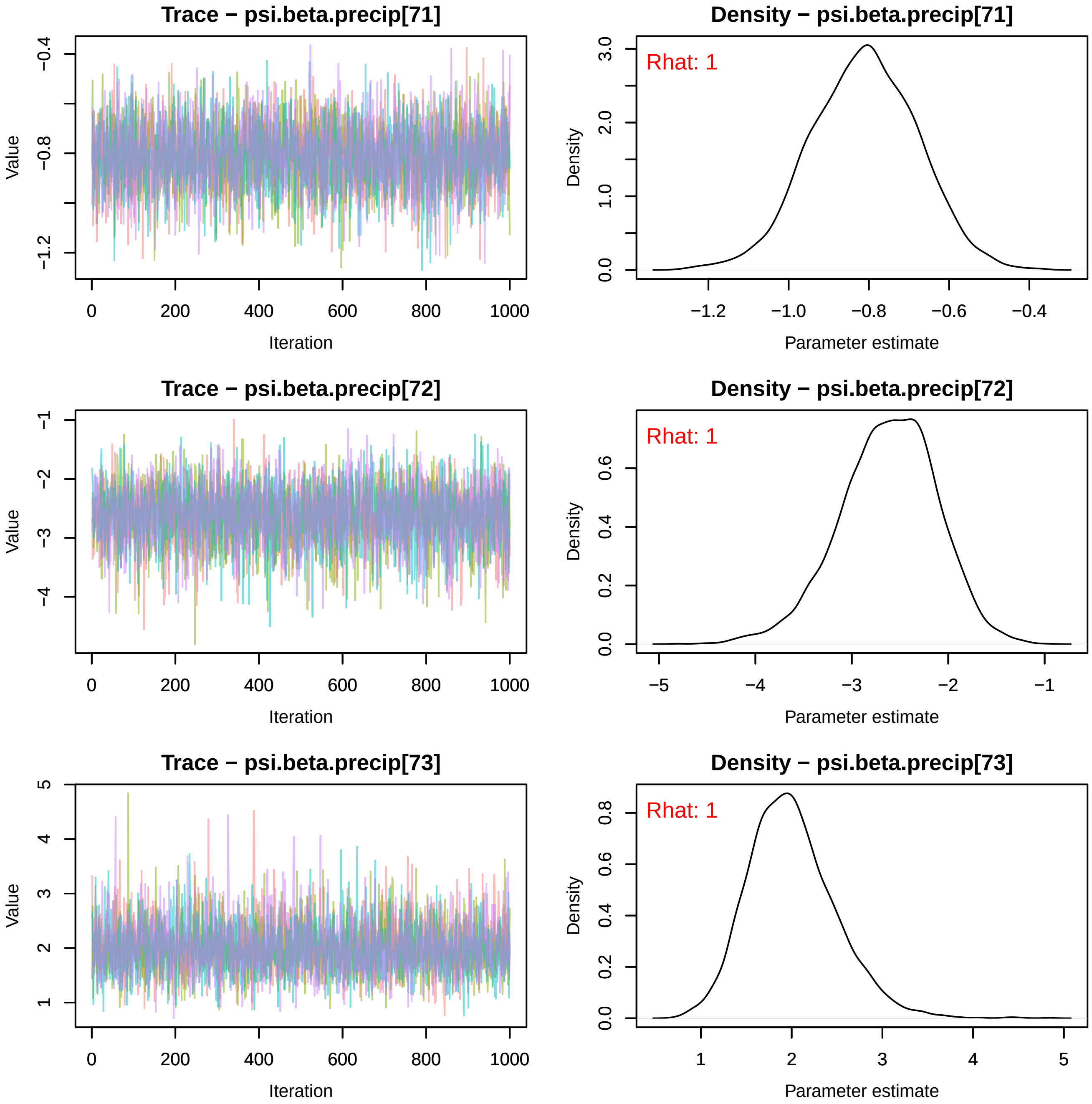
Trace − psi.beta.precip[71]
Value
Iteration
Density − psi.beta.precip[71]
Rhat: 1
Density
Parameter estimate
Trace − psi.beta.precip[72]
Value
Iteration
Density − psi.beta.precip[72]
Rhat: 1
Density
Parameter estimate
Trace − psi.beta.precip[73]
Value
Iteration
Density − psi.beta.precip[73]
Rhat: 1
Density
Parameter estimate

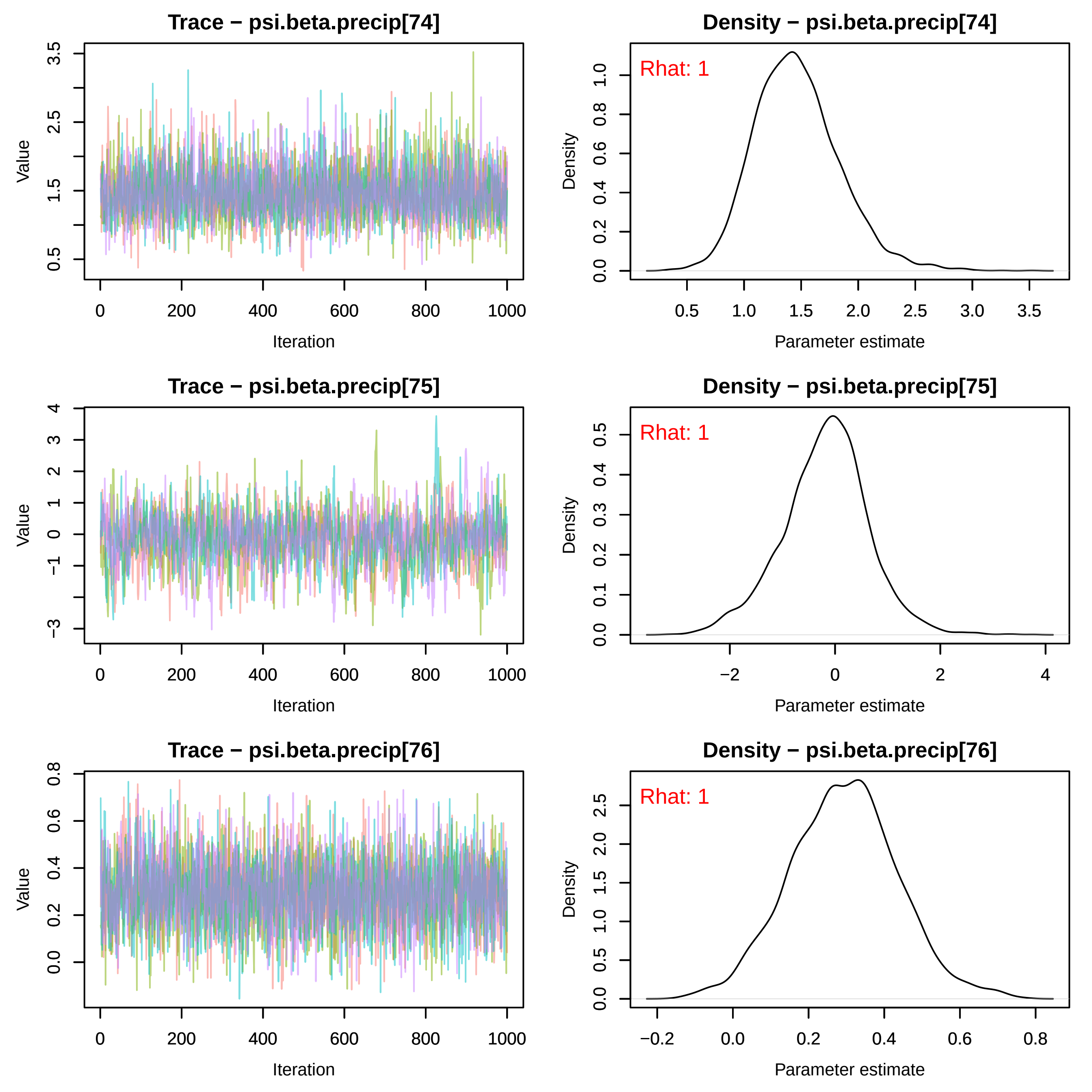
Trace − psi.beta.precip[74]
Value
Iteration
Density − psi.beta.precip[74]
Rhat: 1
Density
Parameter estimate
Trace − psi.beta.precip[75]
Value
Iteration
Density − psi.beta.precip[75]
Rhat: 1
Density
Parameter estimate
Trace − psi.beta.precip[76]
Value
Iteration
Density − psi.beta.precip[76]
Rhat: 1
Density
Parameter estimate

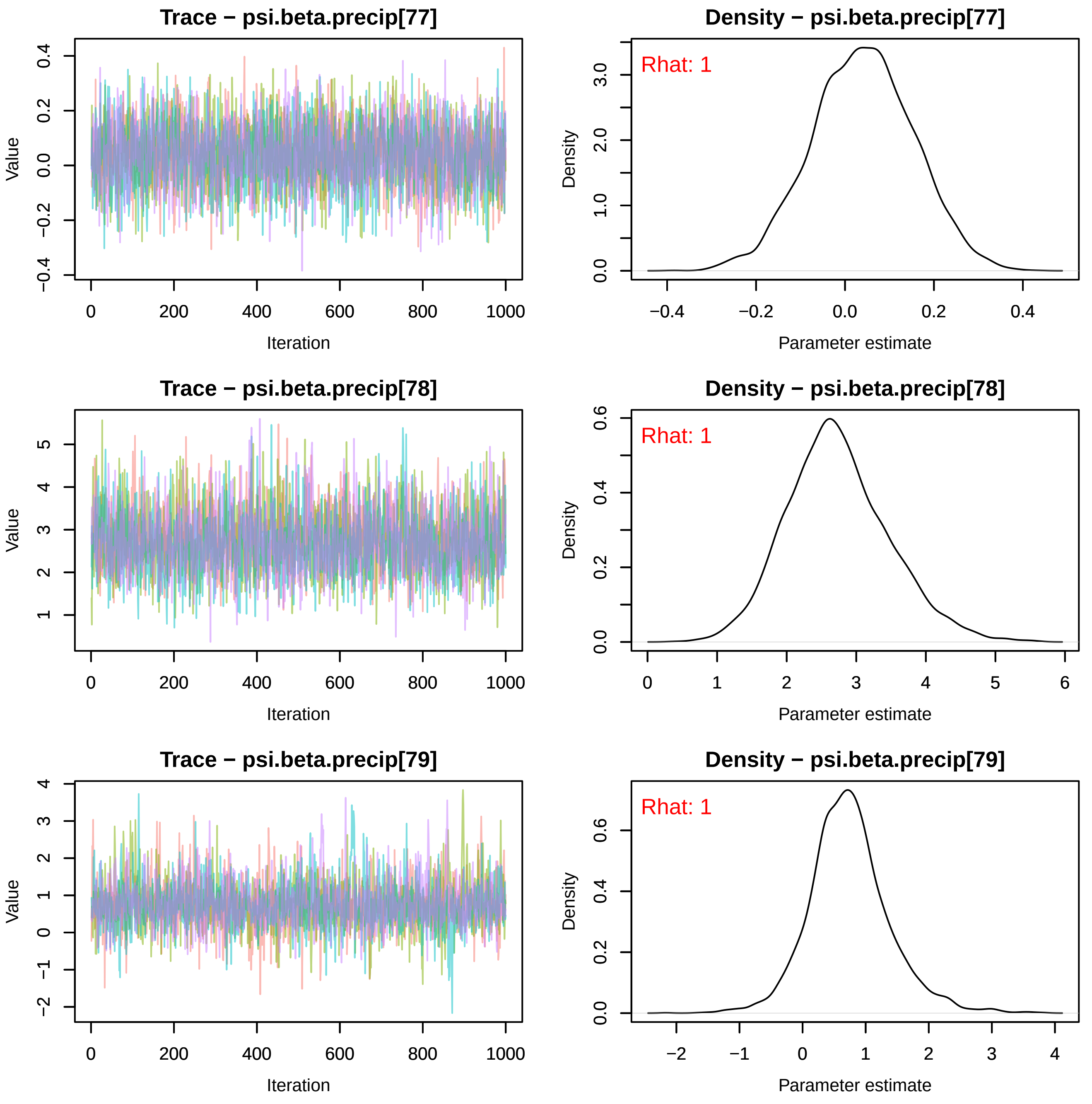

Trace − psi.beta.precip[77]
Density − psi.beta.precip[77]
Rhat: 1
Trace − psi.beta.precip[78]
Density − psi.beta.precip[78]
Rhat: 1
Trace − psi.beta.precip[79]
Density − psi.beta.precip[79]
Rhat: 1
Value
Iteration
Density
Parameter estimate

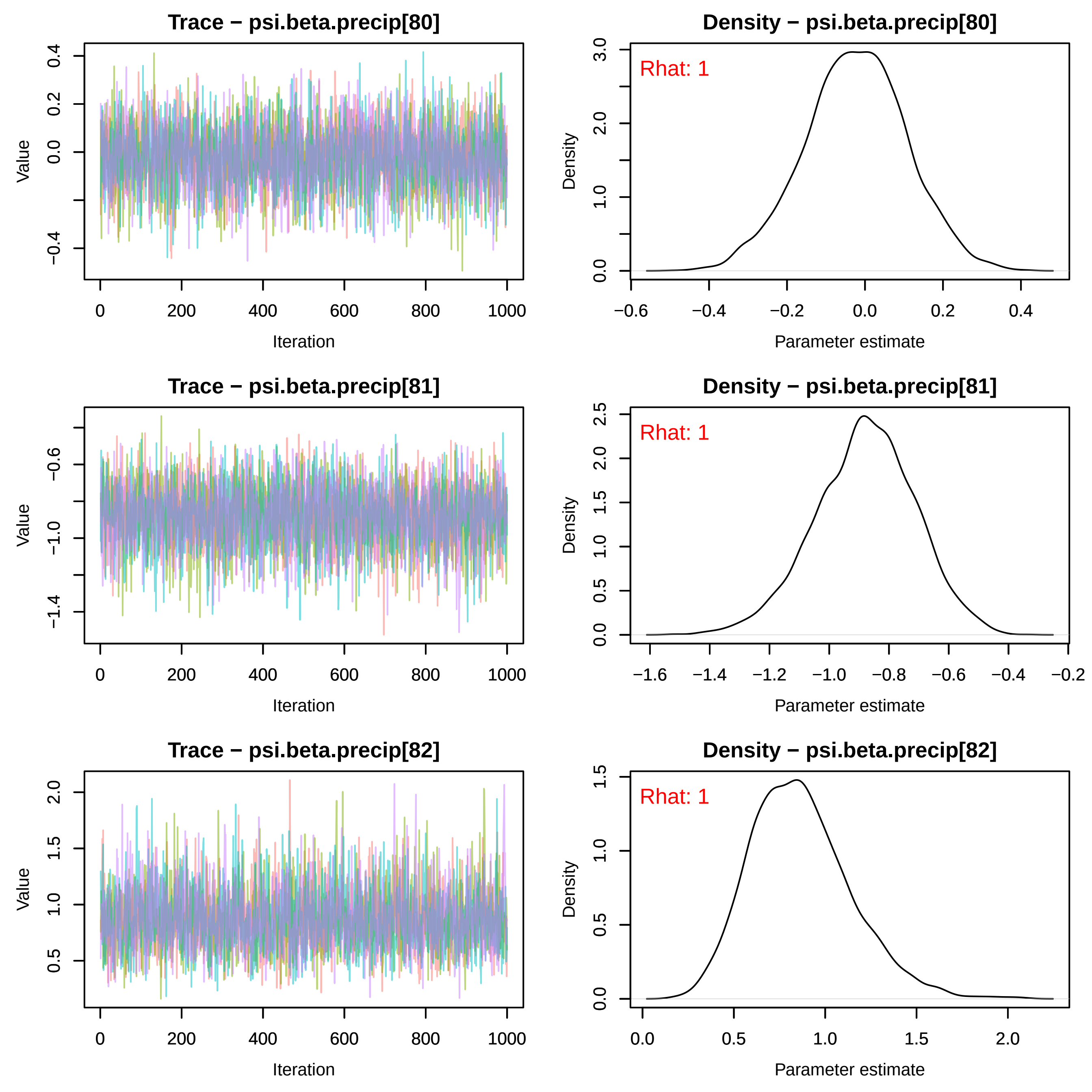

Trace − psi.beta.precip[80]
Value
Iteration
Density − psi.beta.precip[80]
Rhat: 1
Density
Parameter estimate
Trace − psi.beta.precip[81]
Value
Iteration
Density − psi.beta.precip[81]
Rhat: 1
Density
Parameter estimate
Trace − psi.beta.precip[82]
Value
Iteration
Density − psi.beta.precip[82]
Rhat: 1
Density
Parameter estimate

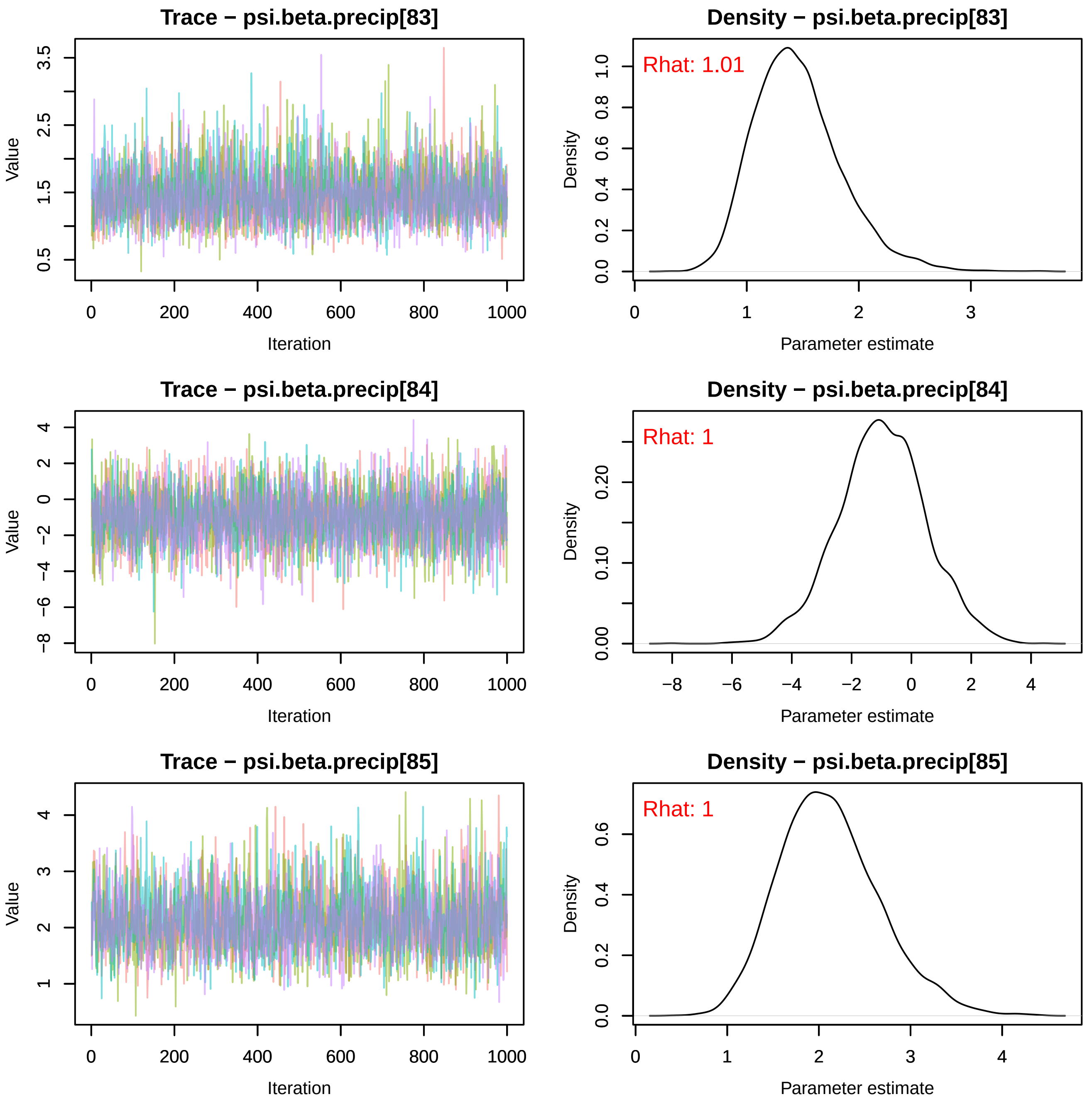
Trace − psi.beta.precip[83]
Density − psi.beta.precip[83]
Rhat: 1.01
Trace − psi.beta.precip[84]
Density − psi.beta.precip[84]
Rhat: 1
Trace − psi.beta.precip[85]
Density − psi.beta.precip[85]
Rhat: 1
Value
Iteration
Density
Parameter estimate

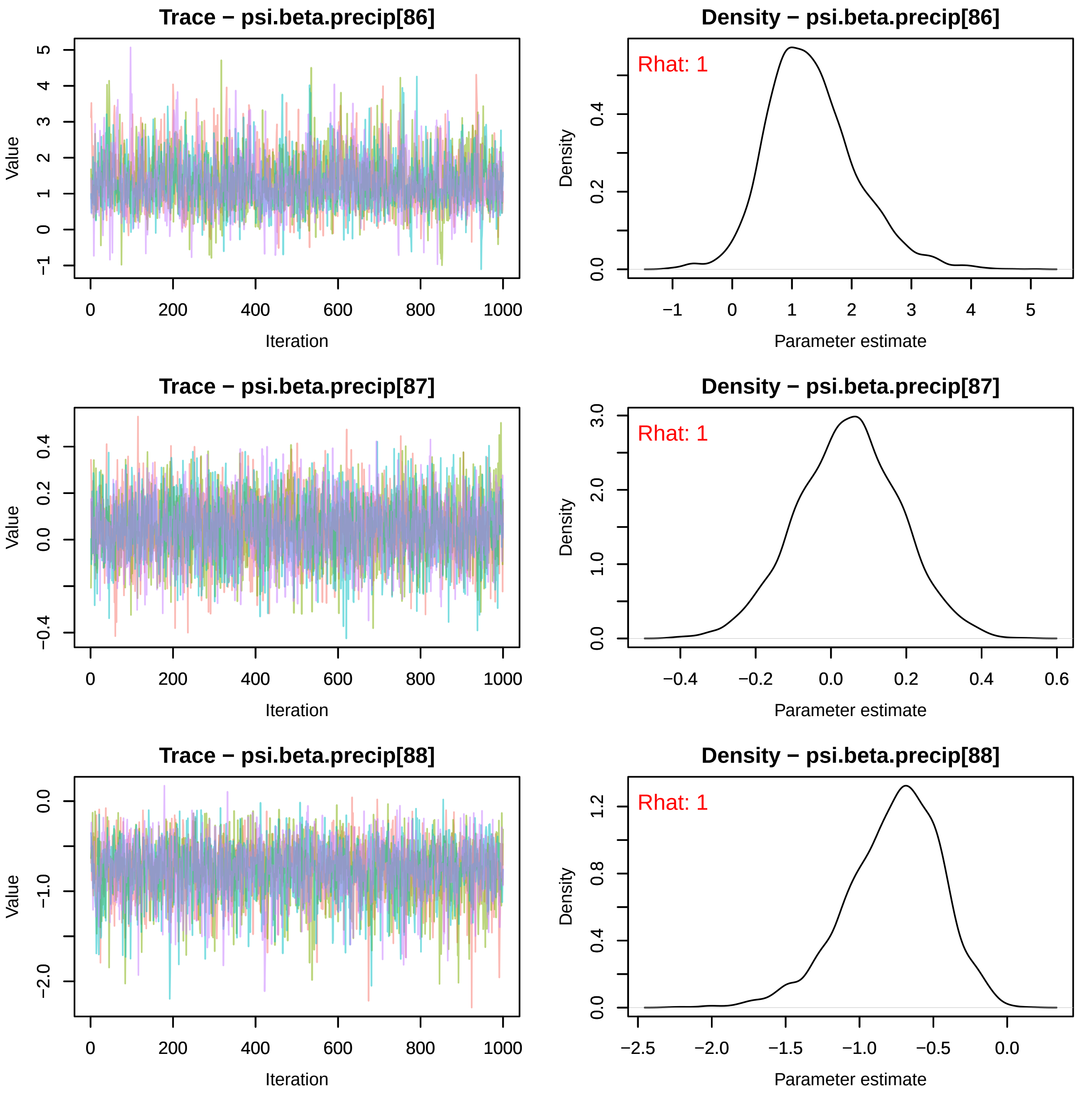
Trace − psi.beta.precip[86]
Value
Iteration
Density − psi.beta.precip[86]
Rhat: 1
Density
Parameter estimate
Trace − psi.beta.precip[87]
Value
Iteration
Density − psi.beta.precip[87]
Rhat: 1
Density
Parameter estimate
Trace − psi.beta.precip[88]
Value
Iteration
Density − psi.beta.precip[88]
Rhat: 1
Density
Parameter estimate

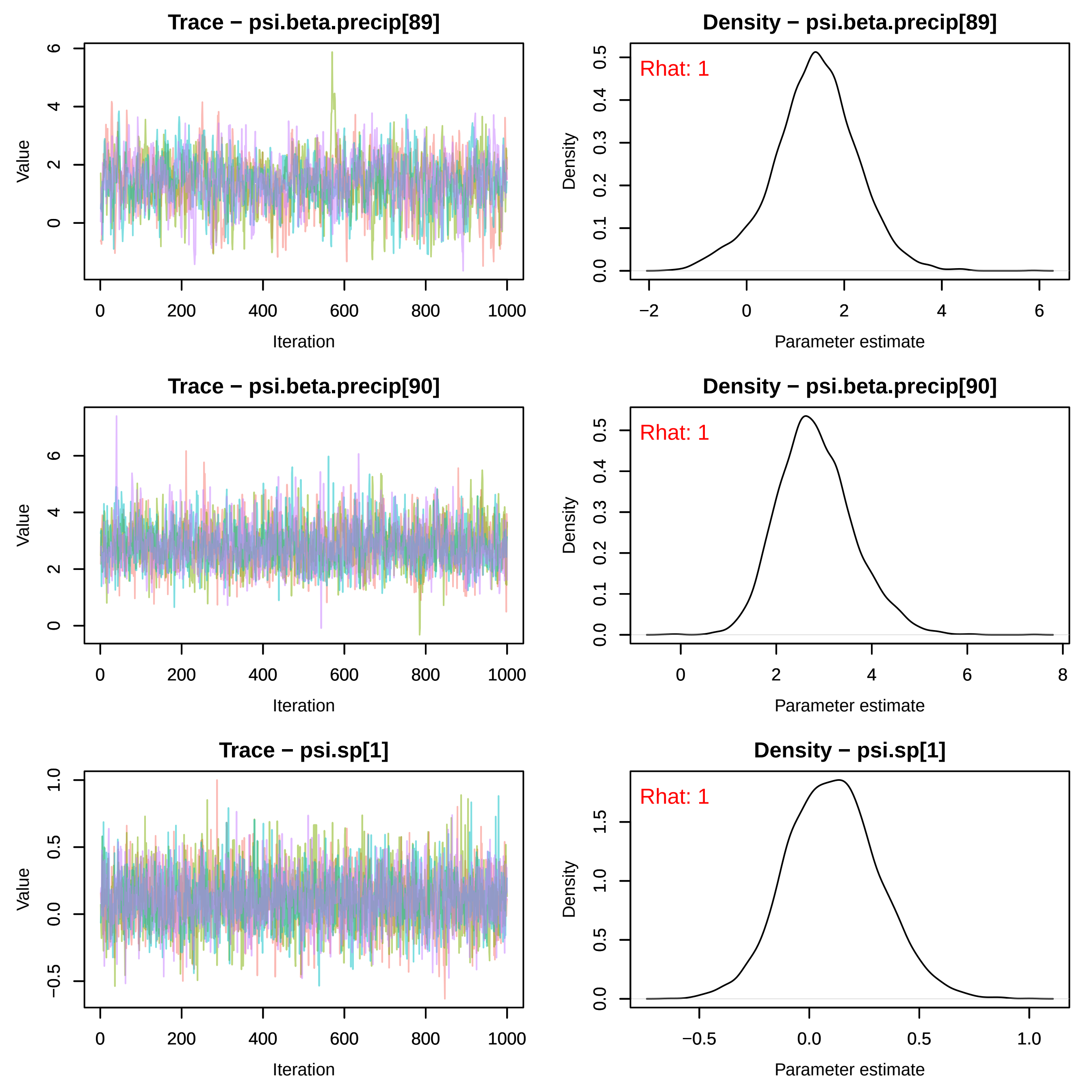
Trace − psi.beta.precip[89]
Value
Iteration
Density − psi.beta.precip[89]
Rhat: 1
Density
Parameter estimate
Trace − psi.beta.precip[90]
Value
Iteration
Density − psi.beta.precip[90]
Rhat: 1
Density
Parameter estimate
Trace − psi.sp[1]
Value
Iteration
Density − psi.sp[1]
Rhat: 1
Density
Parameter estimate

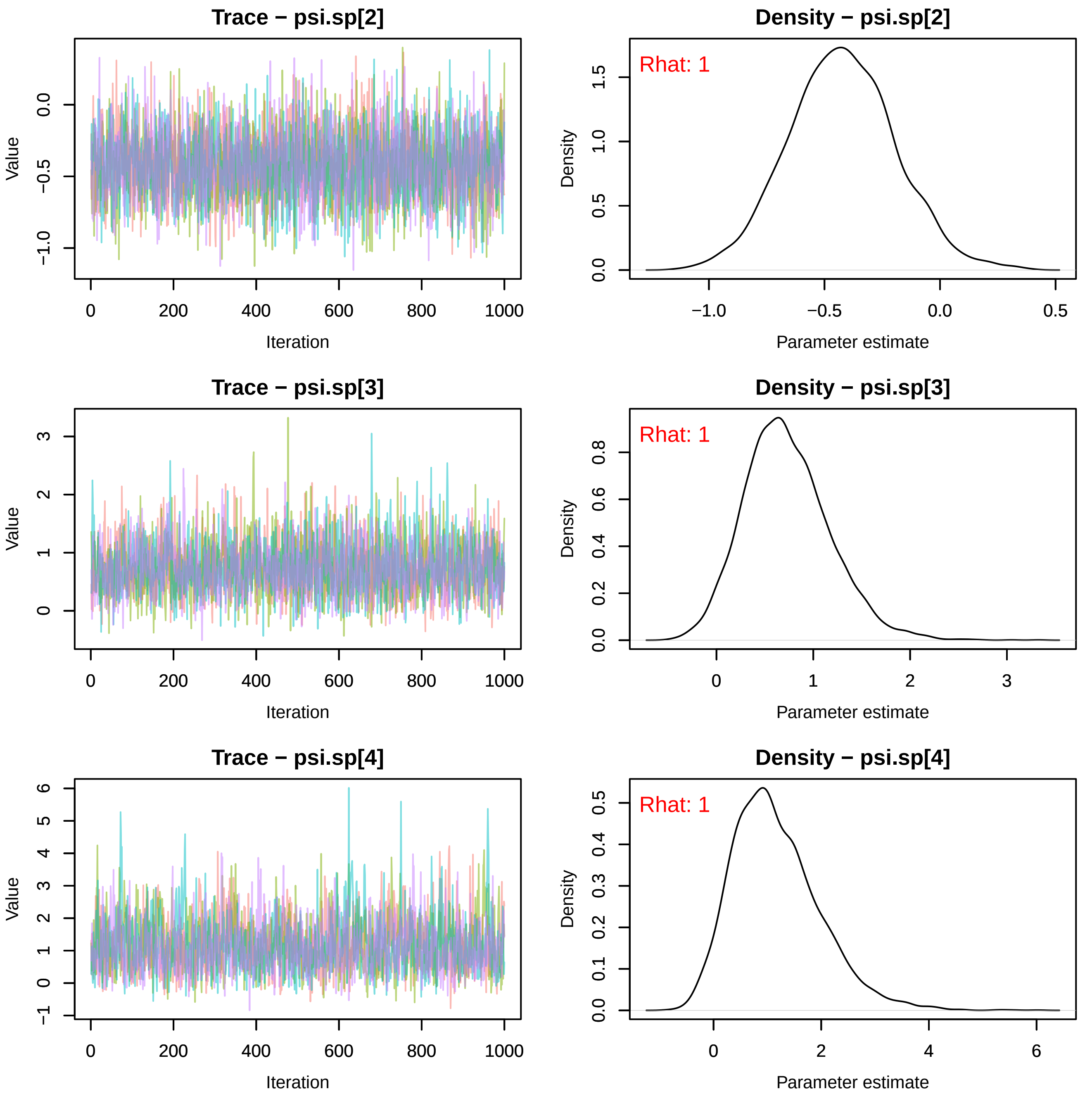
Trace − psi.sp[2]
Value
Iteration
Density − psi.sp[2]
Rhat: 1
Density
Parameter estimate
Trace − psi.sp[3]
Value
Iteration
Density − psi.sp[3]
Rhat: 1
Density
Parameter estimate
Trace − psi.sp[4]
Value
Iteration
Density − psi.sp[4]
Rhat: 1
Density
Parameter estimate

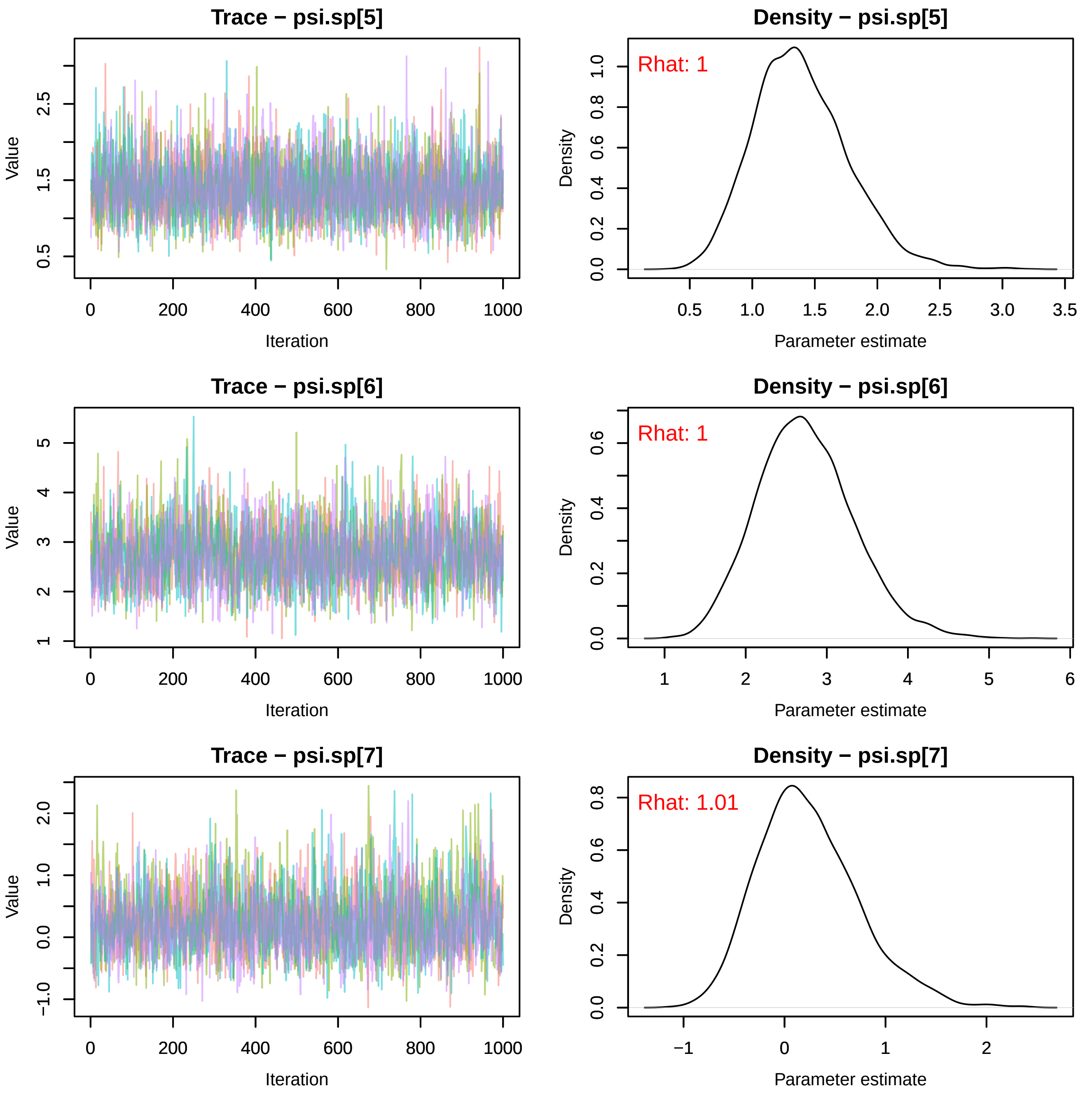
Trace − psi.sp[5]
Value
0.5
1.5
2.5
0
200
400
600
800
1000
Iteration
Density − psi.sp[5]
Rhat: 1
Density
0.0
0.2
0.4
0.6
0.8
1.0
0.5
1.0
1.5
2.0
2.5
3.0
3.5
Parameter estimate
Trace − psi.sp[6]
Value
1
2
3
4
5
0
200
400
600
800
1000
Iteration
Density − psi.sp[6]
Rhat: 1
Density
0.0
0.2
0.4
0.6
1
2
3
4
5
6
Parameter estimate
Trace − psi.sp[7]
Value
−1.0
0.0
1.0
2.0
0
200
400
600
800
1000
Iteration
Density − psi.sp[7]
Rhat: 1.01
Density
0.0
0.2
0.4
0.6
0.8
−1
0
1
2
Parameter estimate

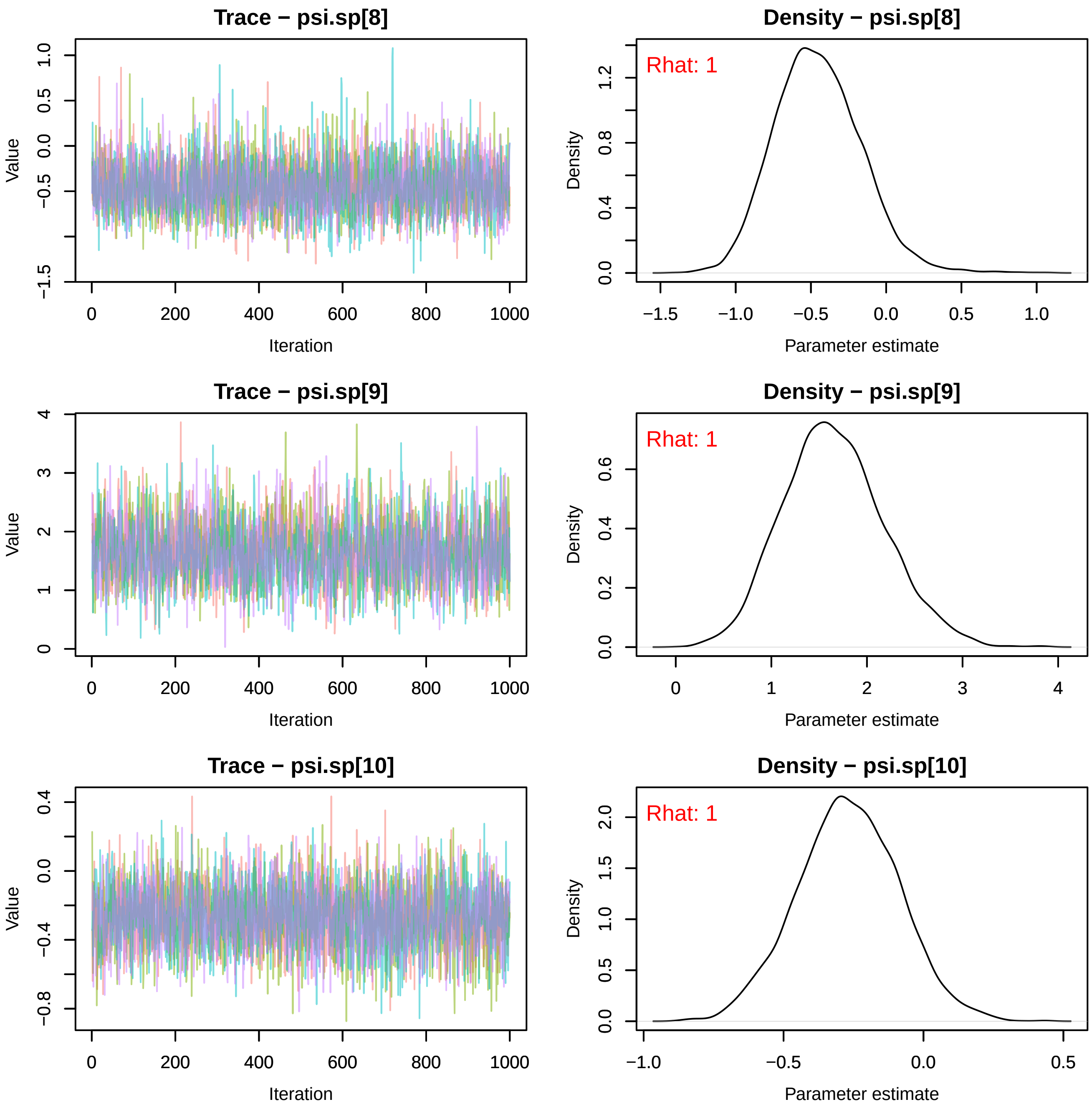
Trace − psi.sp[8]
Value
Iteration
Density − psi.sp[8]
Rhat: 1
Density
Parameter estimate
Trace − psi.sp[9]
Value
Iteration
Density − psi.sp[9]
Rhat: 1
Density
Parameter estimate
Trace − psi.sp[10]
Value
Iteration
Density − psi.sp[10]
Rhat: 1
Density
Parameter estimate

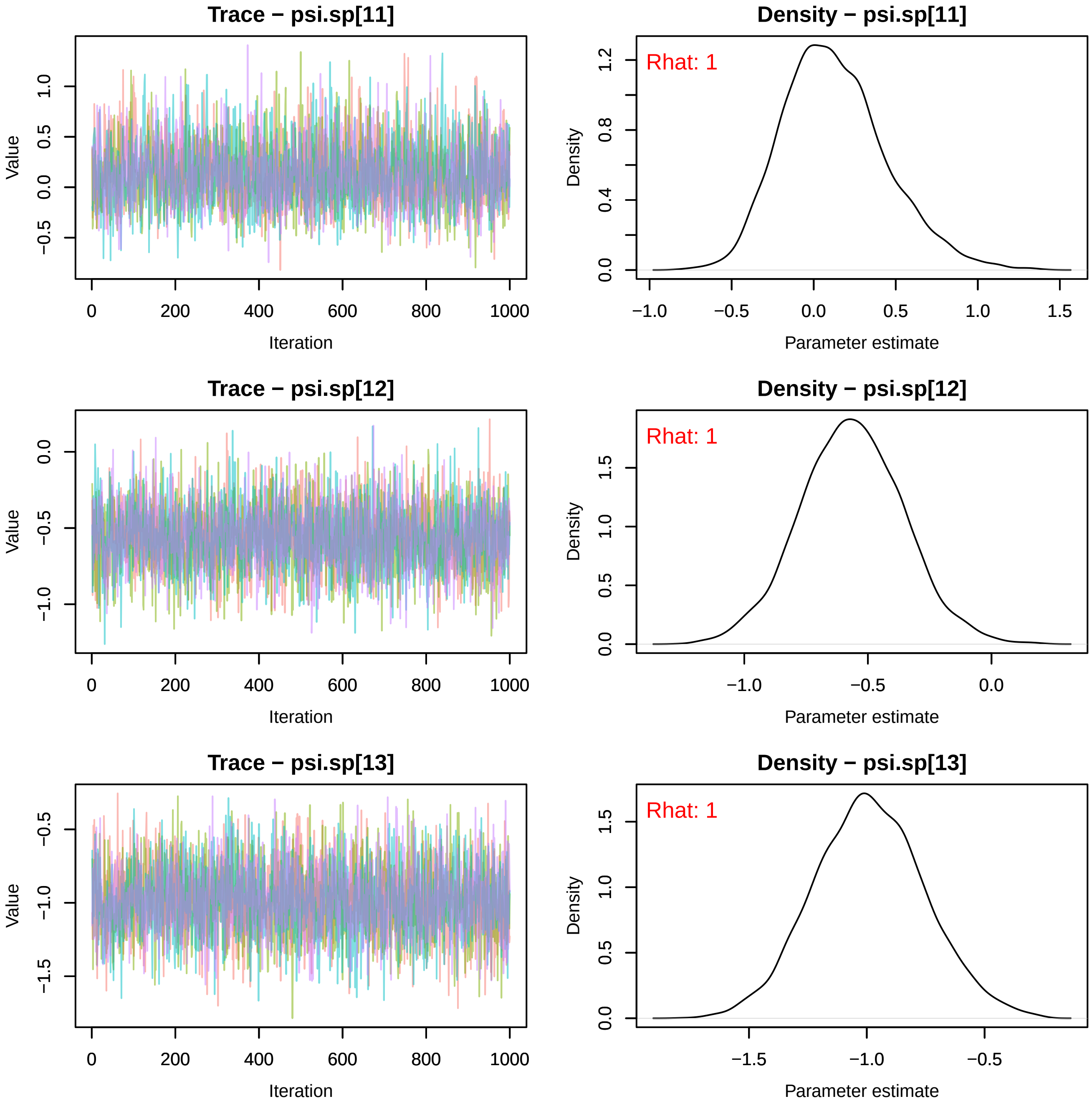
Trace − psi.sp[11]
Value
Iteration
Density − psi.sp[11]
Rhat: 1
Density
Parameter estimate
Trace − psi.sp[12]
Value
Iteration
Density − psi.sp[12]
Rhat: 1
Density
Parameter estimate
Trace − psi.sp[13]
Value
Iteration
Density − psi.sp[13]
Rhat: 1
Density
Parameter estimate

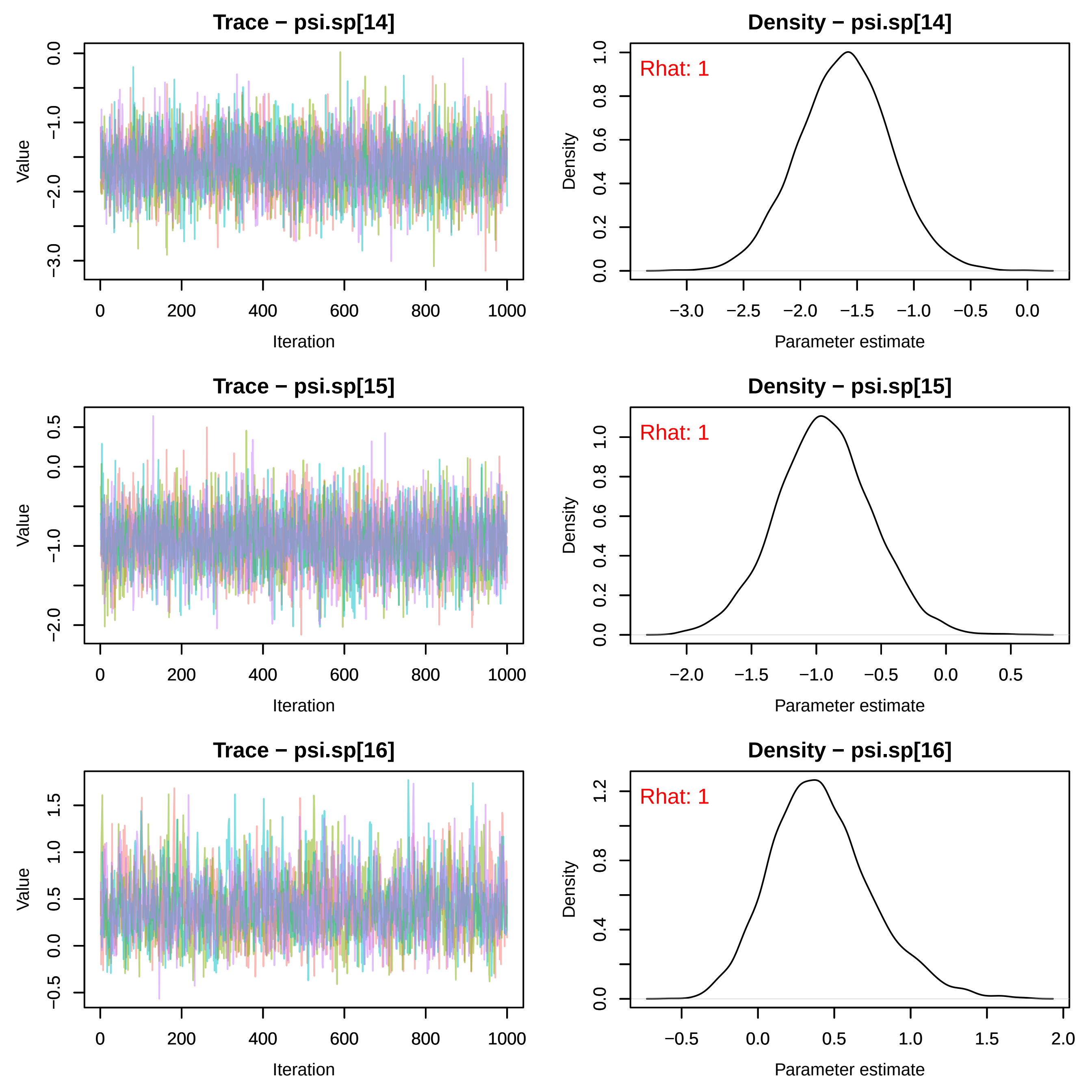
Trace − psi.sp[14]
Value
Iteration
Density − psi.sp[14]
Rhat: 1
Density
Parameter estimate
Trace − psi.sp[15]
Value
Iteration
Density − psi.sp[15]
Rhat: 1
Density
Parameter estimate
Trace − psi.sp[16]
Value
Iteration
Density − psi.sp[16]
Rhat: 1
Density
Parameter estimate

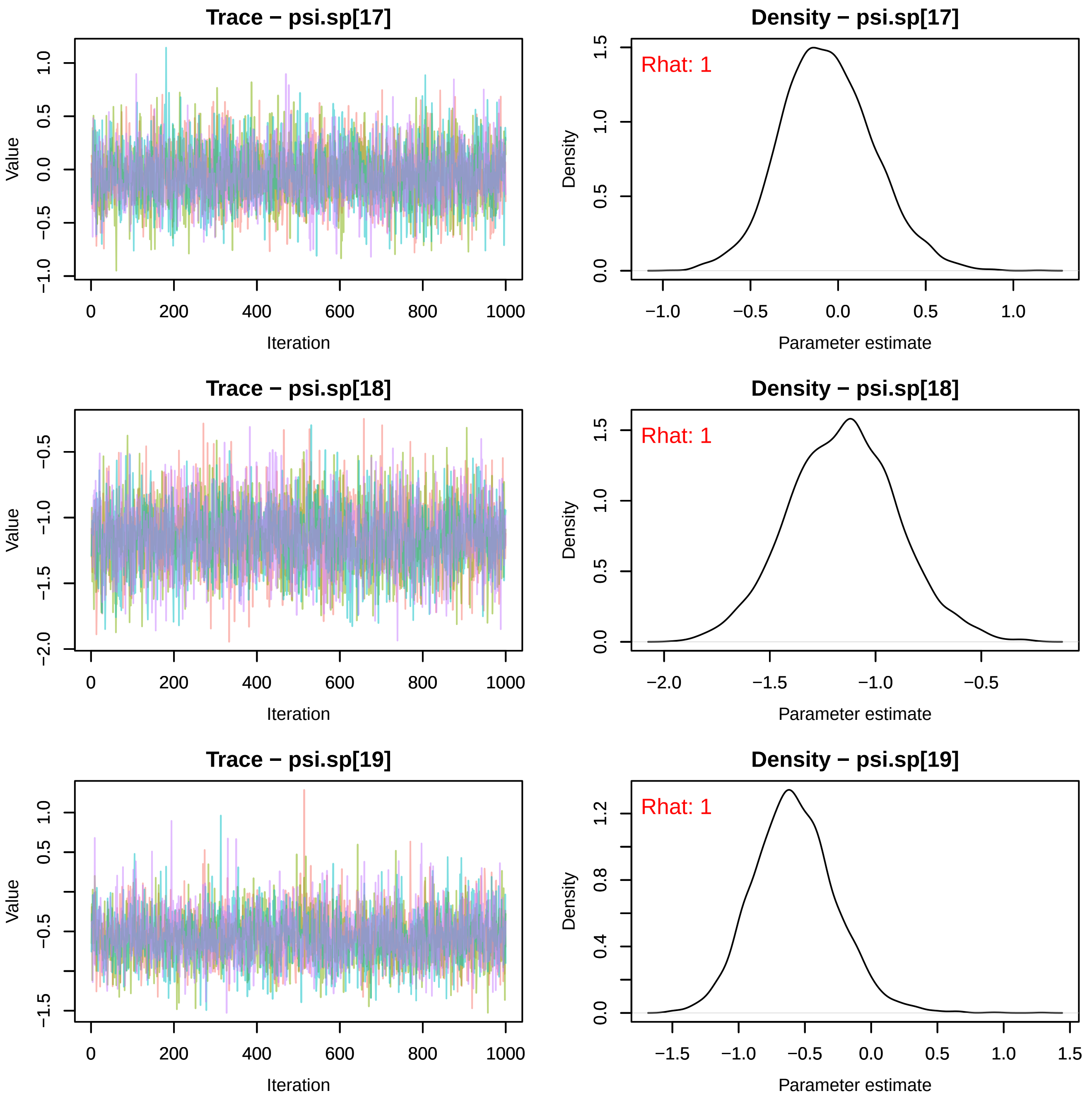

Trace − psi.sp[17]
Value
Iteration
Density − psi.sp[17]
Rhat: 1
Density
Parameter estimate
Trace − psi.sp[18]
Value
Iteration
Density − psi.sp[18]
Rhat: 1
Density
Parameter estimate
Trace − psi.sp[19]
Value
Iteration
Density − psi.sp[19]
Rhat: 1
Density
Parameter estimate

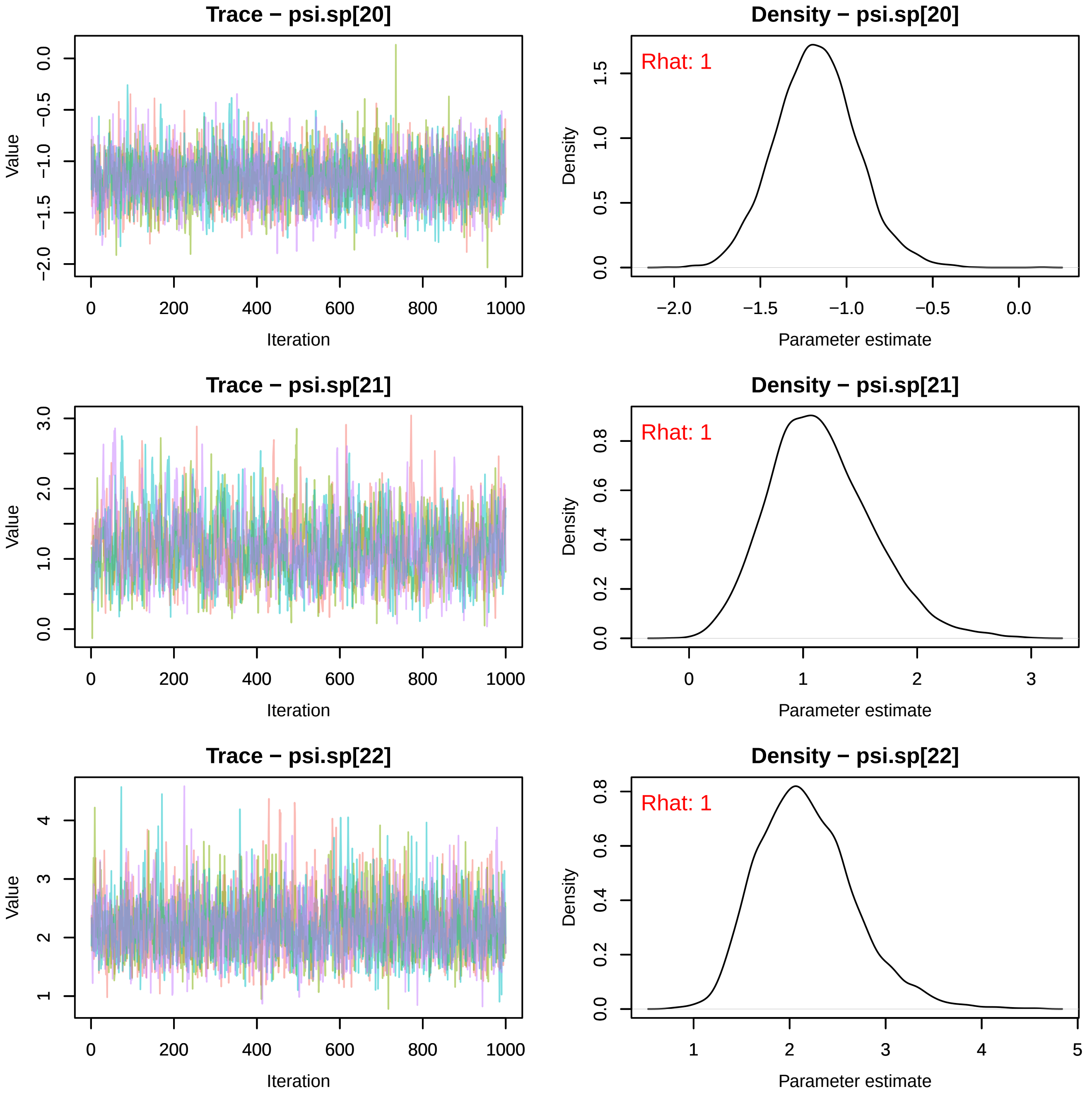
Trace − psi.sp[20]
Value
Iteration
Density − psi.sp[20]
Rhat: 1
Density
Parameter estimate
Trace − psi.sp[21]
Value
Iteration
Density − psi.sp[21]
Rhat: 1
Density
Parameter estimate
Trace − psi.sp[22]
Value
Iteration
Density − psi.sp[22]
Rhat: 1
Density
Parameter estimate

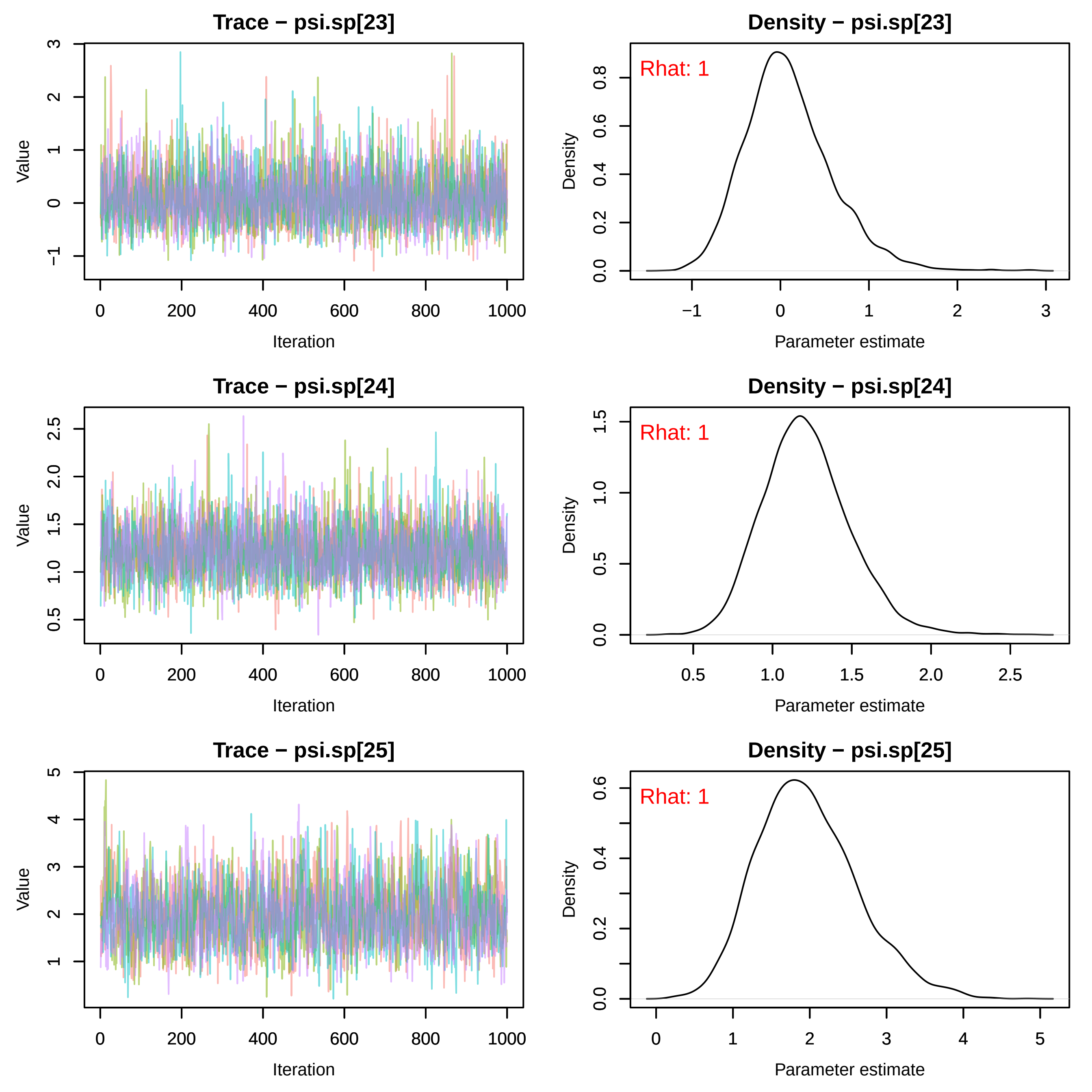
Trace − psi.sp[23]
Value
Iteration
Density − psi.sp[23]
Rhat: 1
Density
Parameter estimate
Trace − psi.sp[24]
Value
Iteration
Density − psi.sp[24]
Rhat: 1
Density
Parameter estimate
Trace − psi.sp[25]
Value
Iteration
Density − psi.sp[25]
Rhat: 1
Density
Parameter estimate

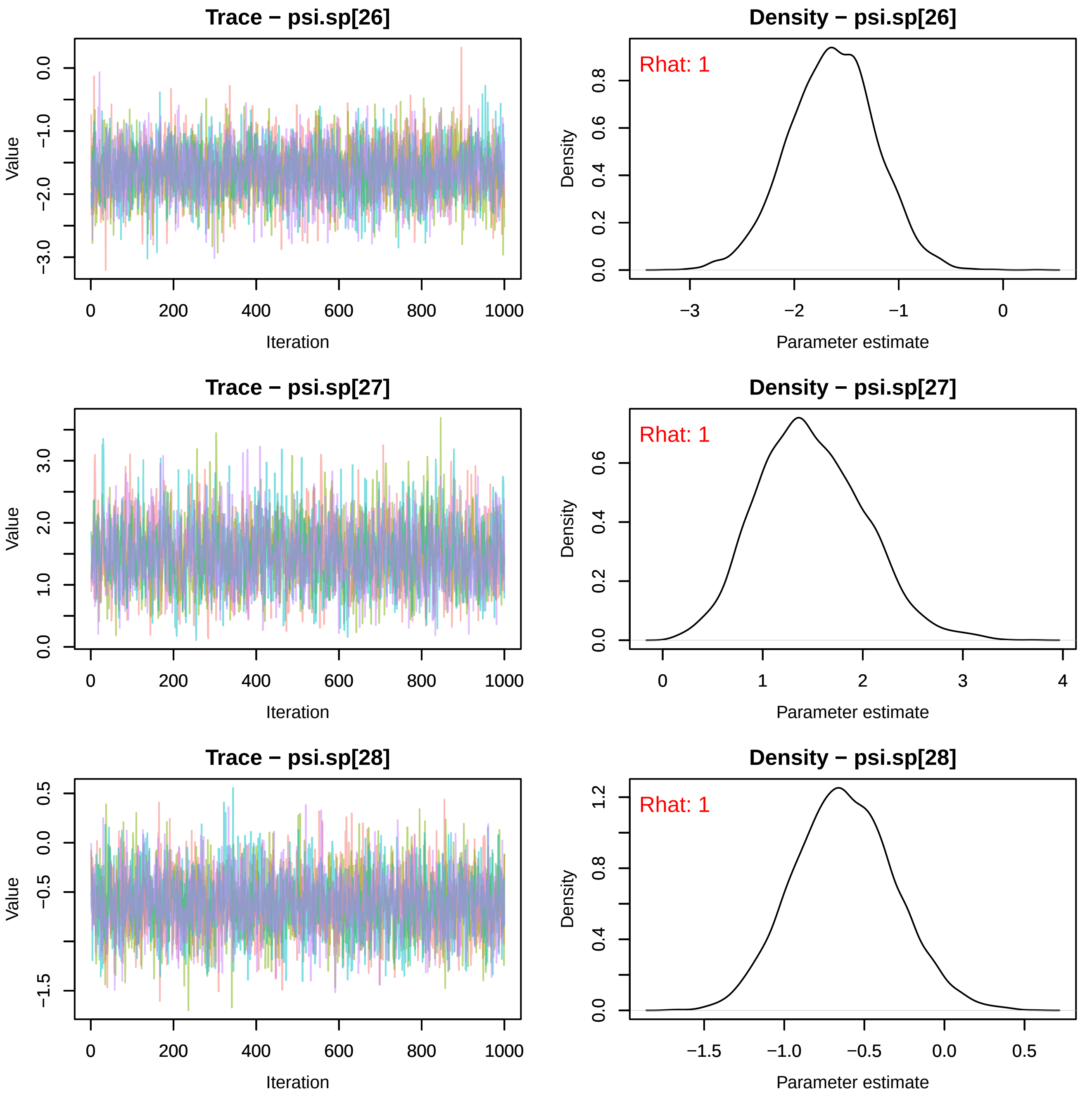

Trace − psi.sp[26]
Value
Iteration
Density − psi.sp[26]
Rhat: 1
Density
Parameter estimate
Trace − psi.sp[27]
Value
Iteration
Density − psi.sp[27]
Rhat: 1
Density
Parameter estimate
Trace − psi.sp[28]
Value
Iteration
Density − psi.sp[28]
Rhat: 1
Density
Parameter estimate

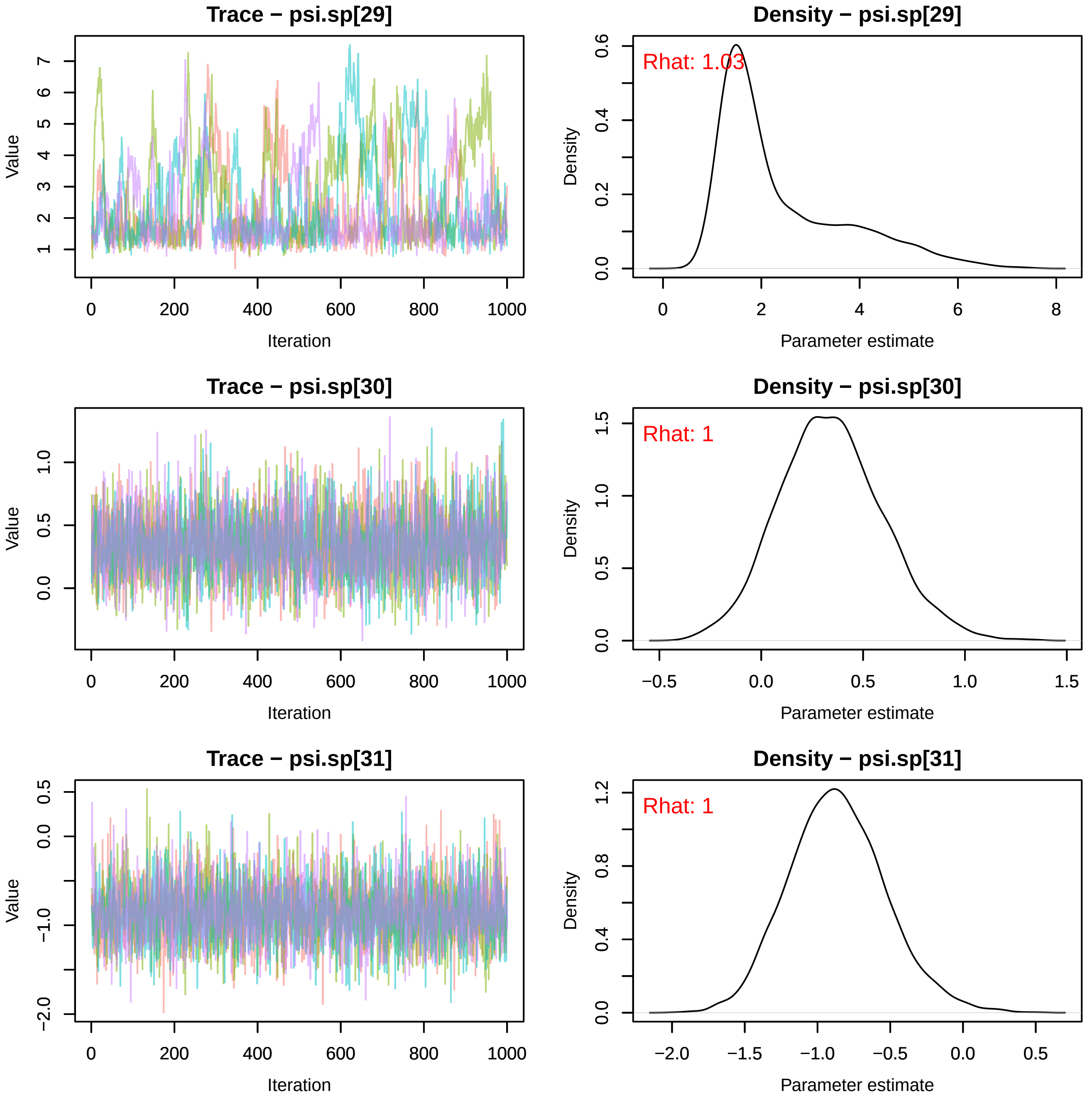
Trace − psi.sp[29]
Value
Iteration
Density − psi.sp[29]
Rhat: 1.03
Density
Parameter estimate
Trace − psi.sp[30]
Value
Iteration
Density − psi.sp[30]
Rhat: 1
Density
Parameter estimate
Trace − psi.sp[31]
Value
Iteration
Density − psi.sp[31]
Rhat: 1
Density
Parameter estimate

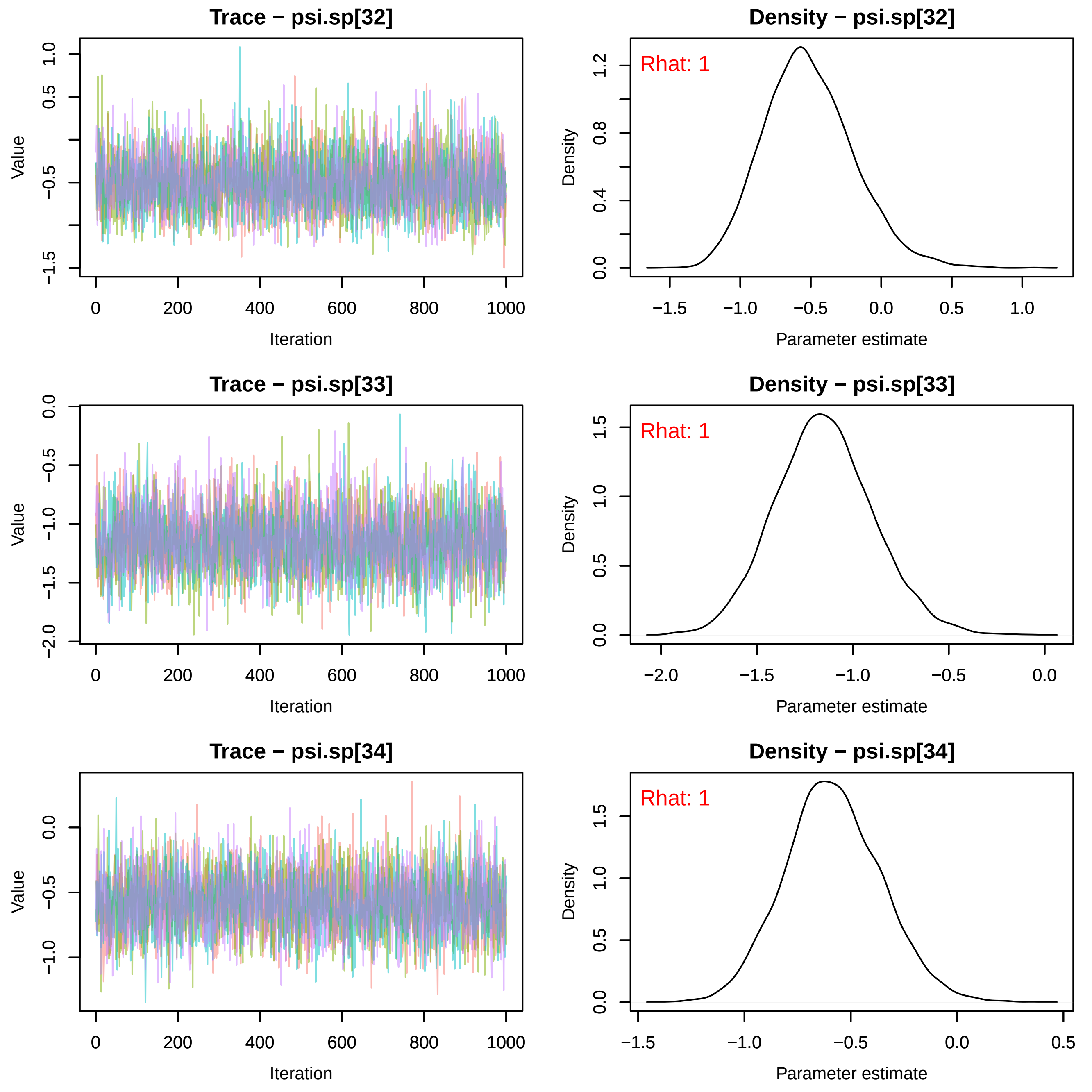
Trace − psi.sp[32]
Density − psi.sp[32]
Rhat: 1
Trace − psi.sp[33]
Density − psi.sp[33]
Rhat: 1
Trace − psi.sp[34]
Density − psi.sp[34]
Rhat: 1
Value
Iteration
Density
Parameter estimate

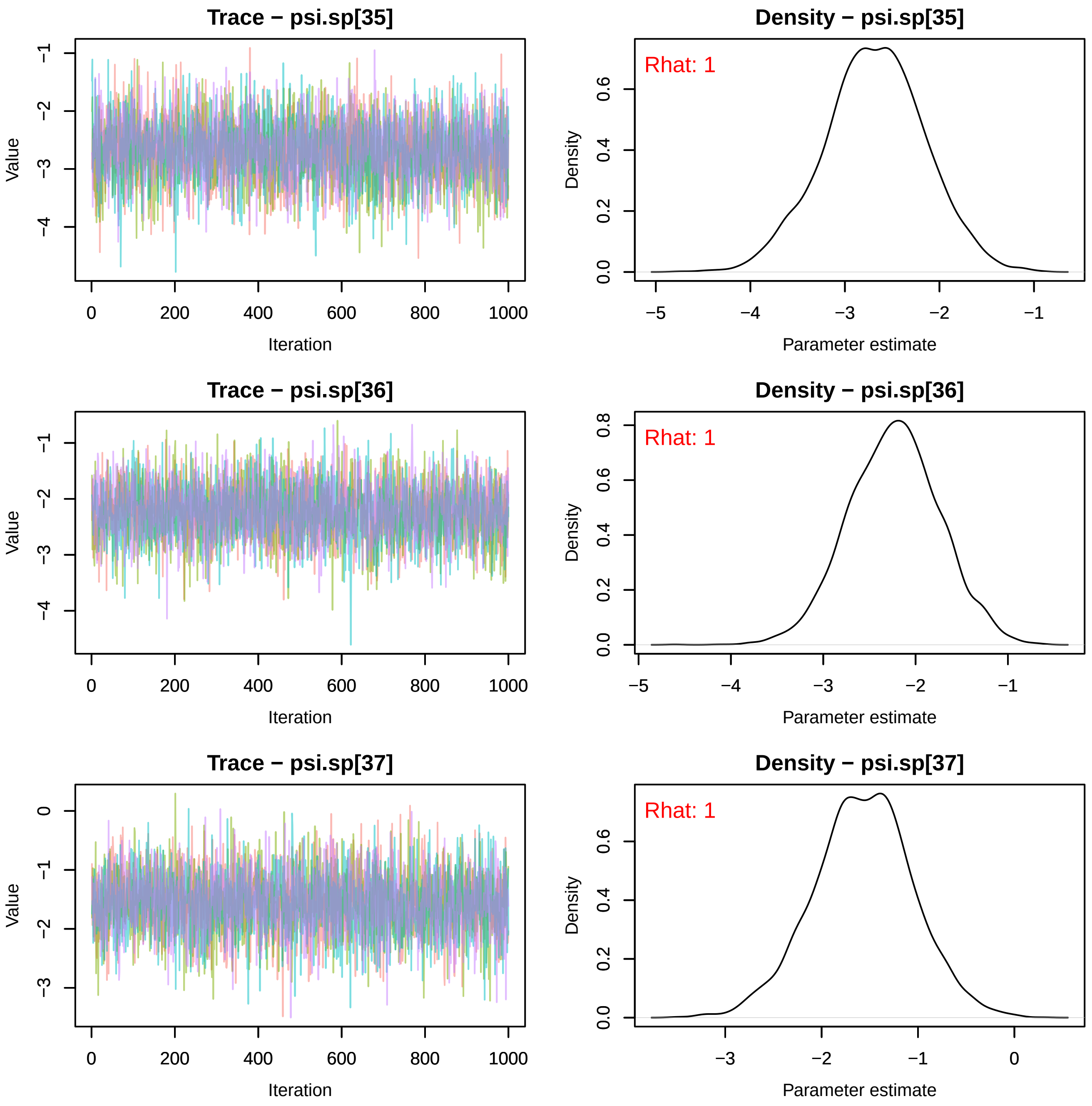
Trace − psi.sp[35]
Value
Iteration
Density − psi.sp[35]
Rhat: 1
Density
Parameter estimate
Trace − psi.sp[36]
Value
Iteration
Density − psi.sp[36]
Rhat: 1
Density
Parameter estimate
Trace − psi.sp[37]
Value
Iteration
Density − psi.sp[37]
Rhat: 1
Density
Parameter estimate

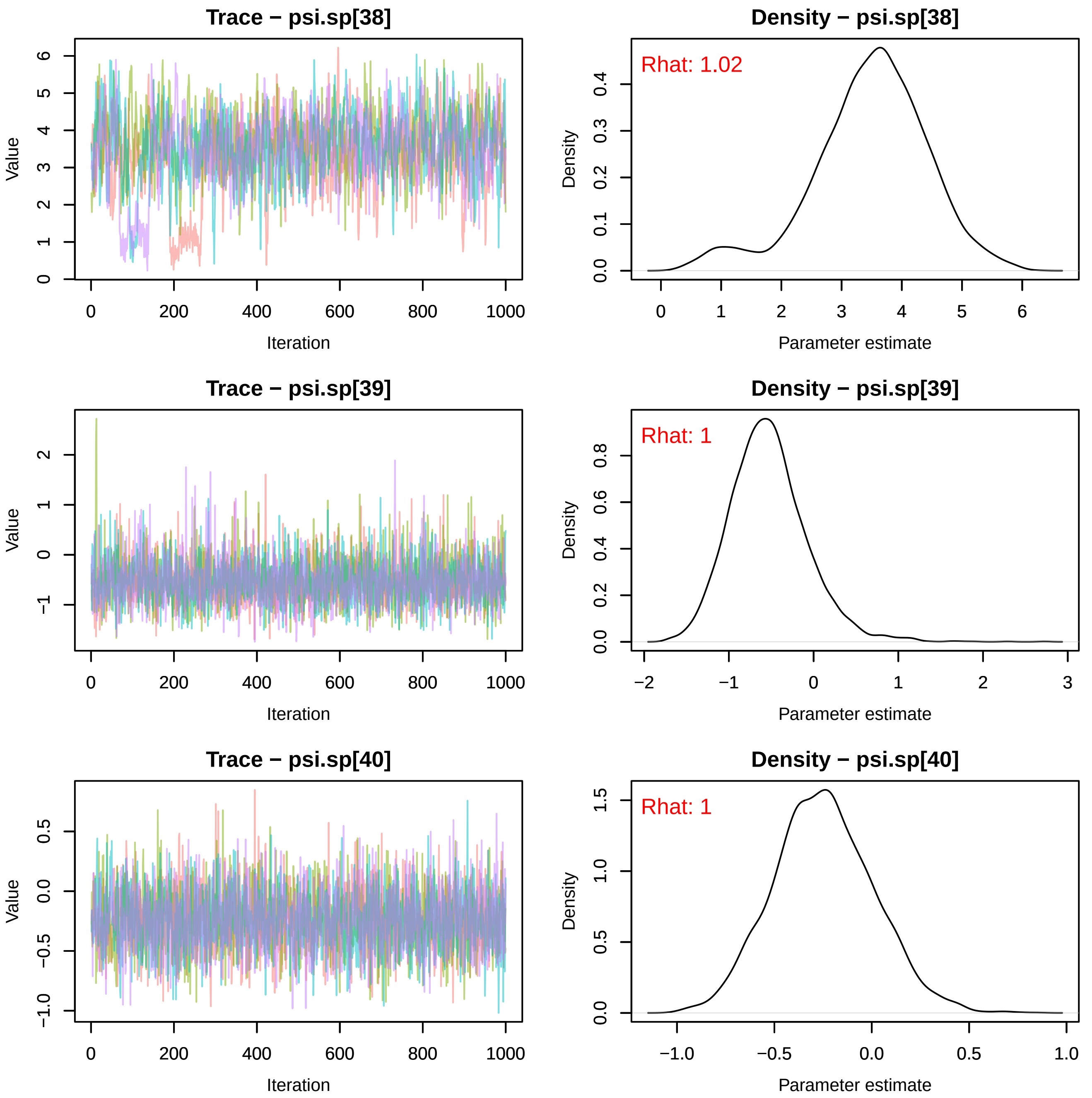
Trace − psi.sp[38]
Density − psi.sp[38]
Rhat: 1.02
Trace − psi.sp[39]
Density − psi.sp[39]
Rhat: 1
Trace − psi.sp[40]
Density − psi.sp[40]
Rhat: 1
Value
Iteration
Density
Parameter estimate

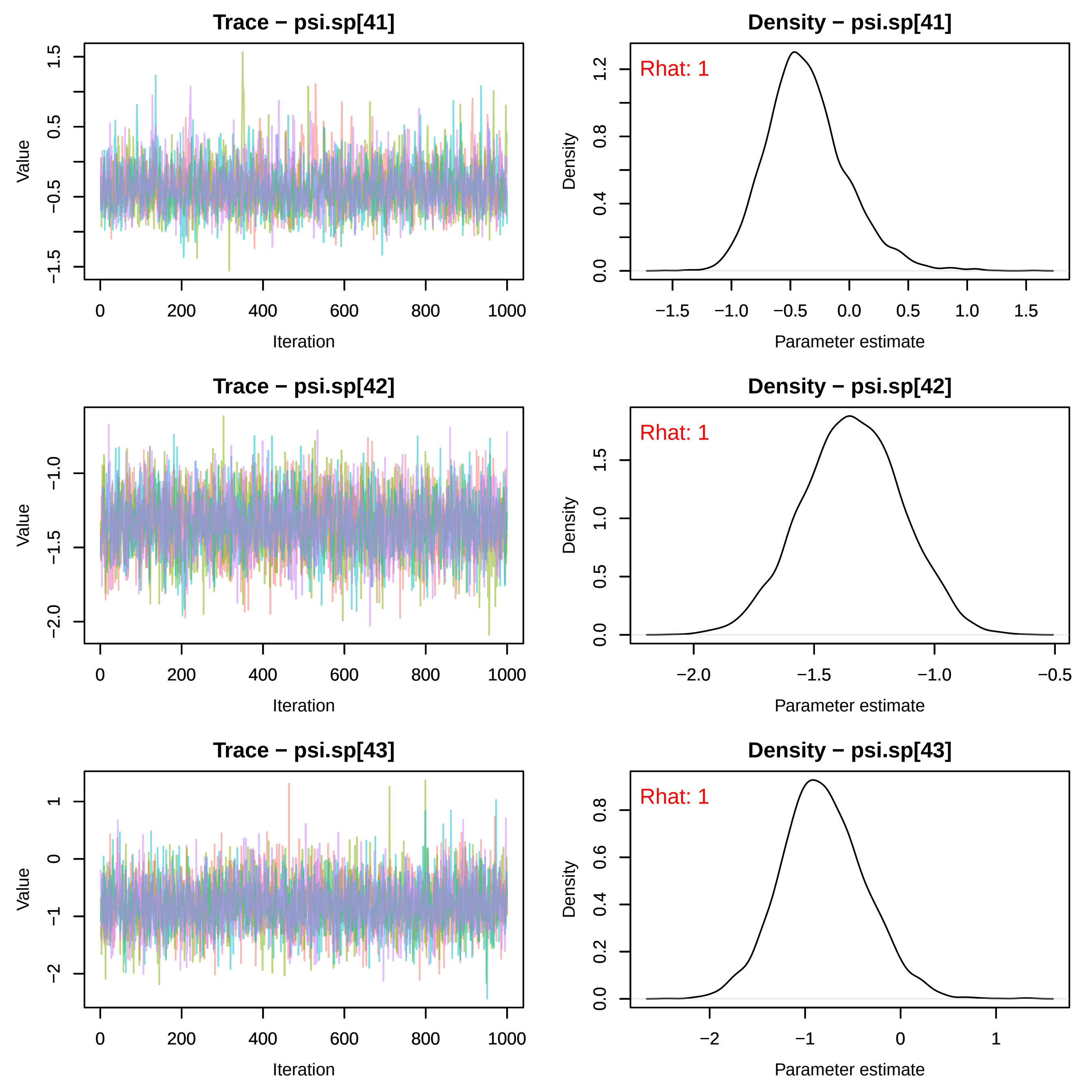
Trace − psi.sp[41]
Value
Iteration
Density − psi.sp[41]
Rhat: 1
Density
Parameter estimate
Trace − psi.sp[42]
Value
Iteration
Density − psi.sp[42]
Rhat: 1
Density
Parameter estimate
Trace − psi.sp[43]
Value
Iteration
Density − psi.sp[43]
Rhat: 1
Density
Parameter estimate

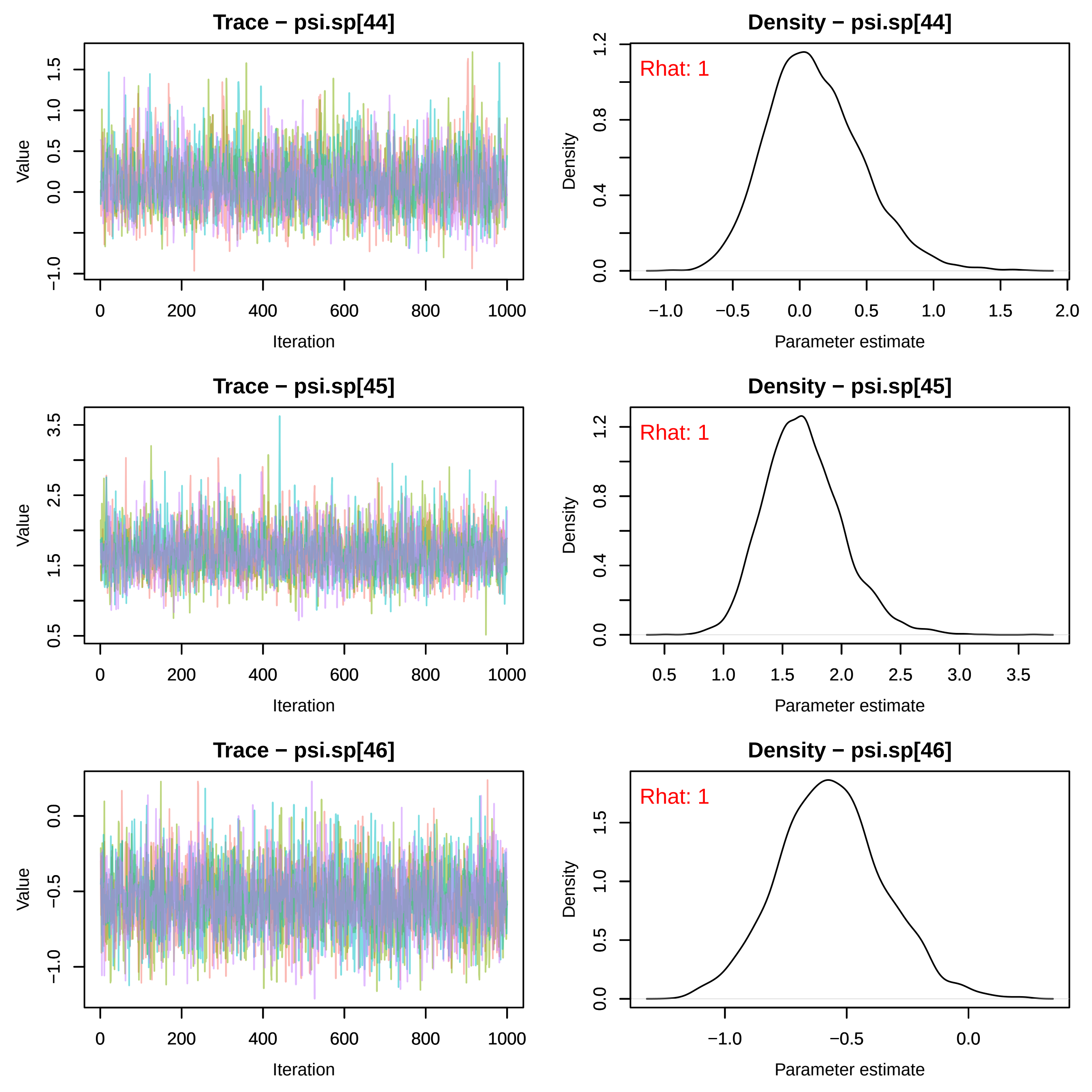
Trace − psi.sp[44]
Value
Iteration
Density − psi.sp[44]
Rhat: 1
Density
Parameter estimate
Trace − psi.sp[45]
Value
Iteration
Density − psi.sp[45]
Rhat: 1
Density
Parameter estimate
Trace − psi.sp[46]
Value
Iteration
Density − psi.sp[46]
Rhat: 1
Density
Parameter estimate

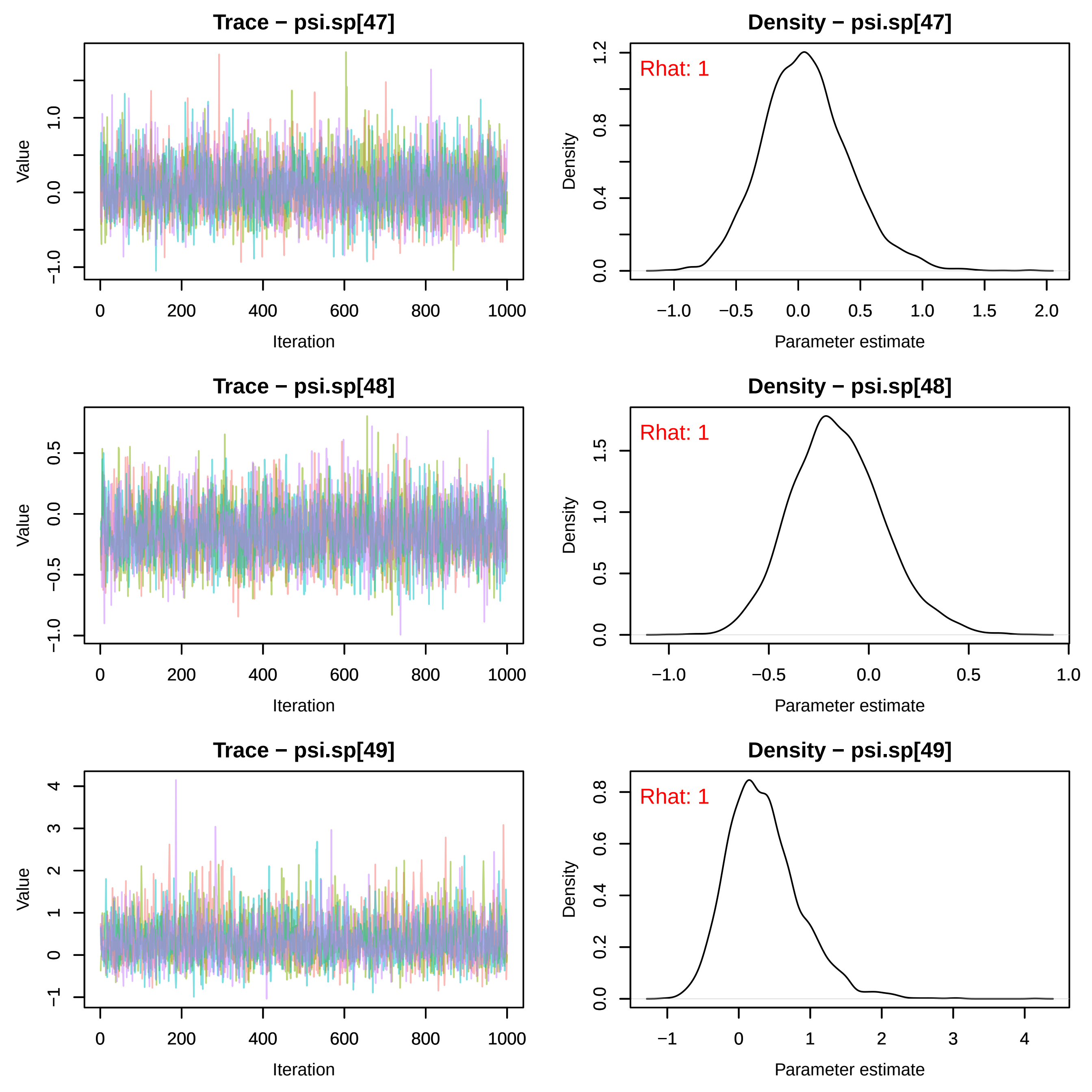
Trace − psi.sp[47]
Density − psi.sp[47]
Rhat: 1
Trace − psi.sp[48]
Density − psi.sp[48]
Rhat: 1
Trace − psi.sp[49]
Density − psi.sp[49]
Rhat: 1
Value
Iteration
Density
Parameter estimate

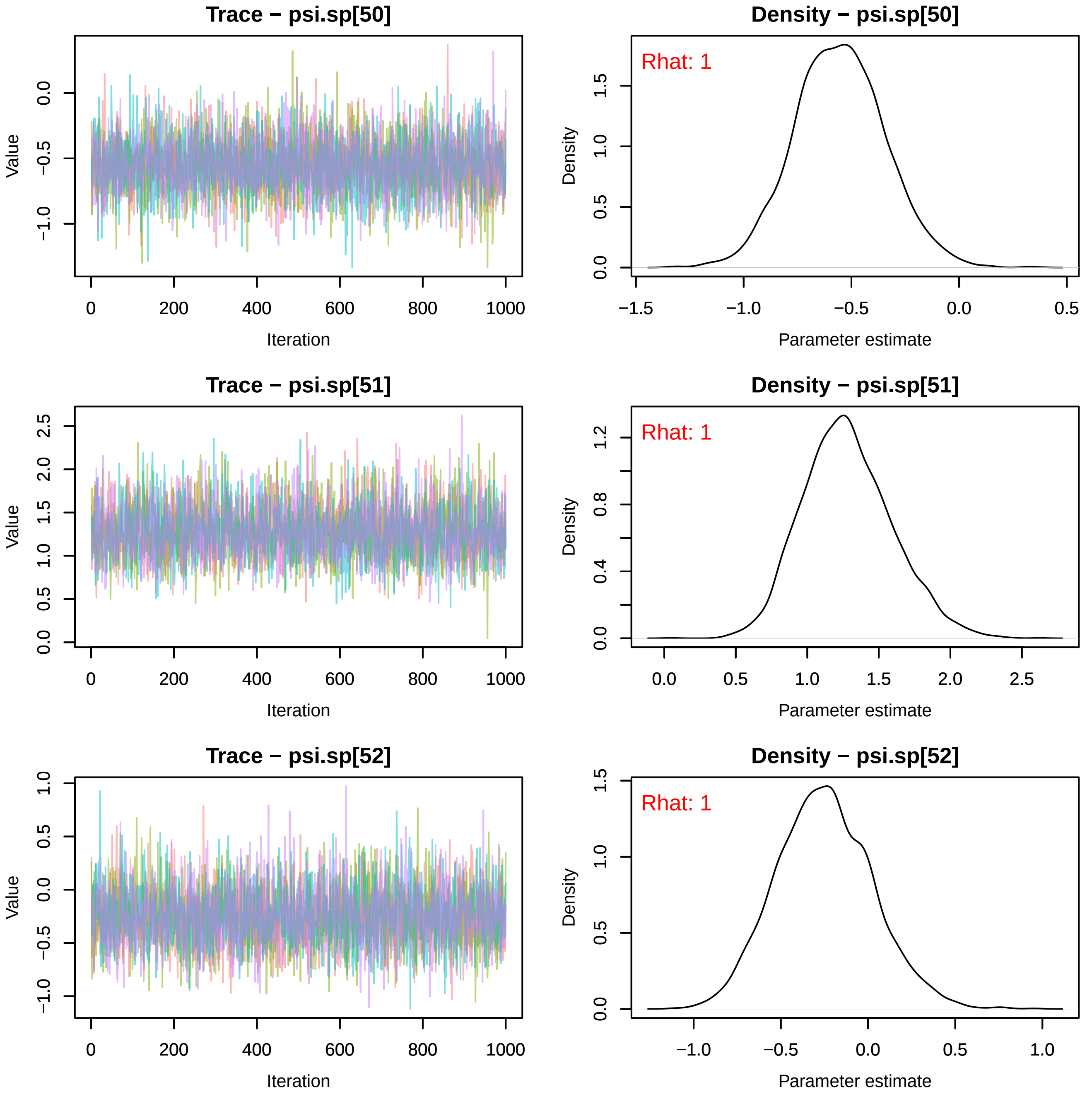
Trace − psi.sp[50]
Density − psi.sp[50]
Rhat: 1
Trace − psi.sp[51]
Density − psi.sp[51]
Rhat: 1
Trace − psi.sp[52]
Density − psi.sp[52]
Rhat: 1
Value
Iteration
Density
Parameter estimate

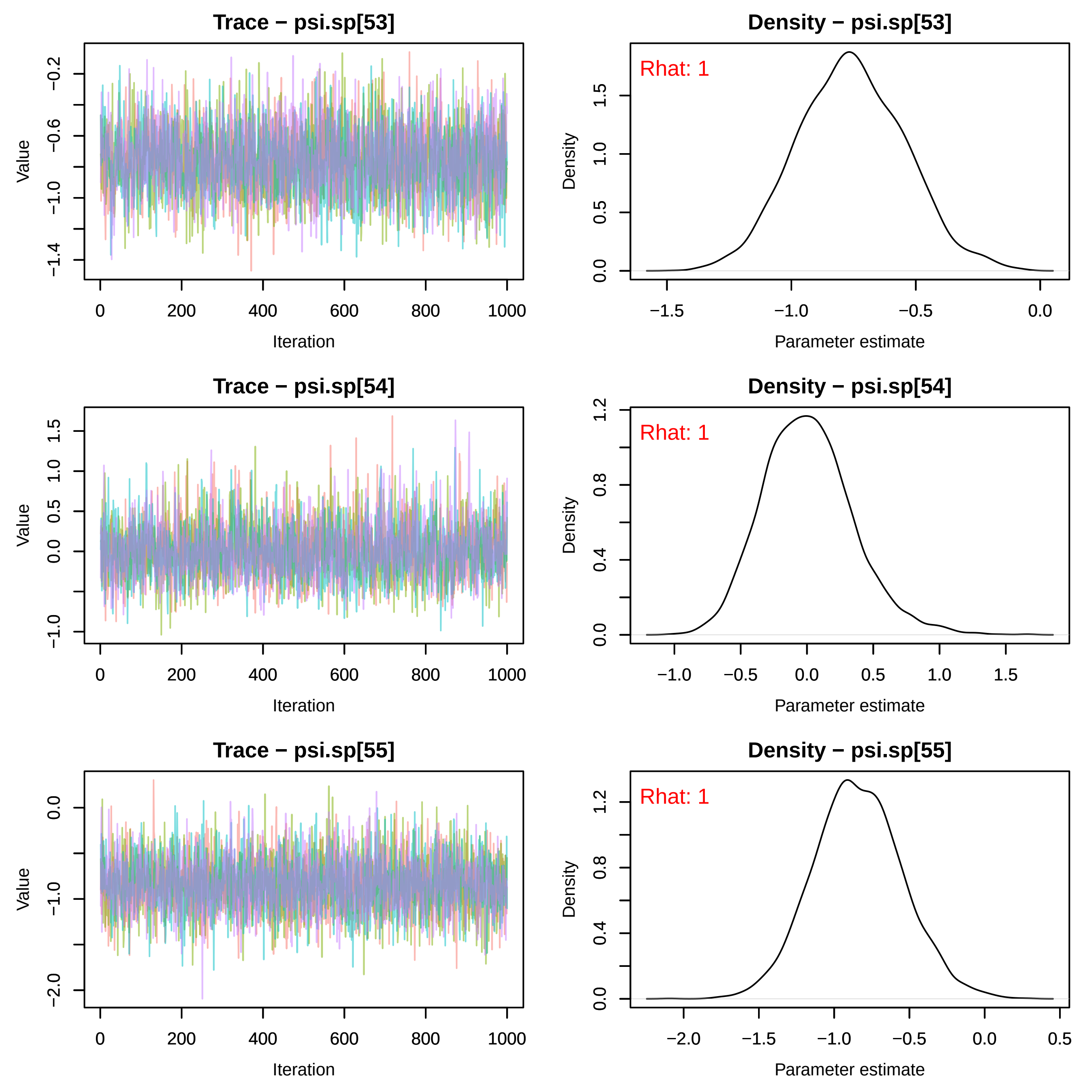
Trace − psi.sp[53]
Value
−0.2
−0.6
−1.0
−1.4
0
200
400
600
800
1000
Iteration
Density − psi.sp[53]
Rhat: 1
Density
0.0
0.5
1.0
1.5
−1.5
−1.0
−0.5
0.0
Parameter estimate
Trace − psi.sp[54]
Value
−1.0
0.0
0.5
1.0
1.5
0
200
400
600
800
1000
Iteration
Density − psi.sp[54]
Rhat: 1
Density
0.0
0.4
0.8
1.2
−1.0
−0.5
0.0
0.5
1.0
1.5
Parameter estimate
Trace − psi.sp[55]
Value
−2.0
−1.0
0.0
0
200
400
600
800
1000
Iteration
Density − psi.sp[55]
Rhat: 1
Density
0.0
0.4
0.8
1.2
−2.0
−1.5
−1.0
−0.5
0.0
0.5
Parameter estimate

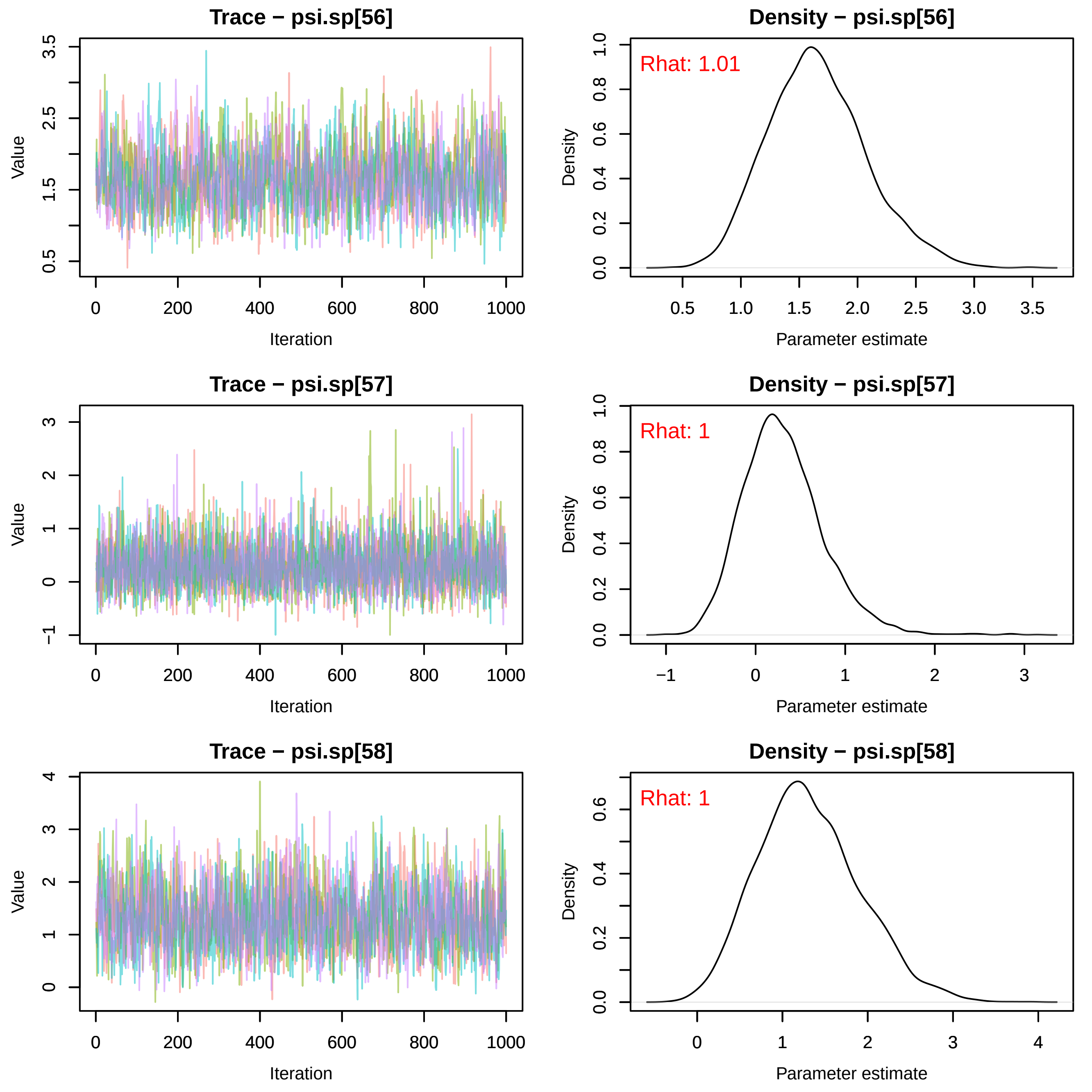
Trace − psi.sp[56]
Value
Iteration
Density − psi.sp[56]
Rhat: 1.01
Density
Parameter estimate
Trace − psi.sp[57]
Value
Iteration
Density − psi.sp[57]
Rhat: 1
Density
Parameter estimate
Trace − psi.sp[58]
Value
Iteration
Density − psi.sp[58]
Rhat: 1
Density
Parameter estimate

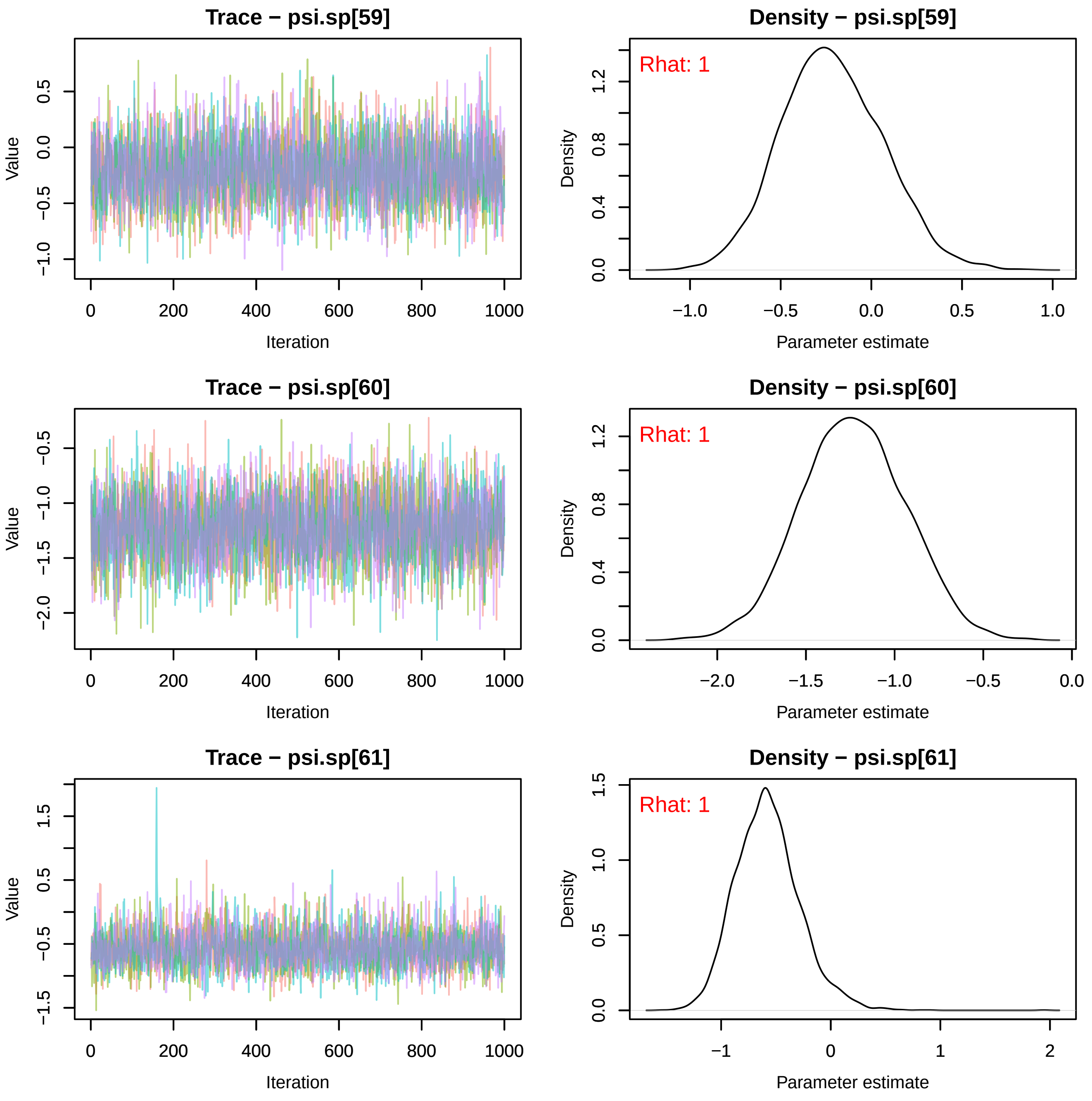
Trace − psi.sp[59]
Density − psi.sp[59]
Rhat: 1
Trace − psi.sp[60]
Density − psi.sp[60]
Rhat: 1
Trace − psi.sp[61]
Density − psi.sp[61]
Rhat: 1
Value
Iteration
Density
Parameter estimate

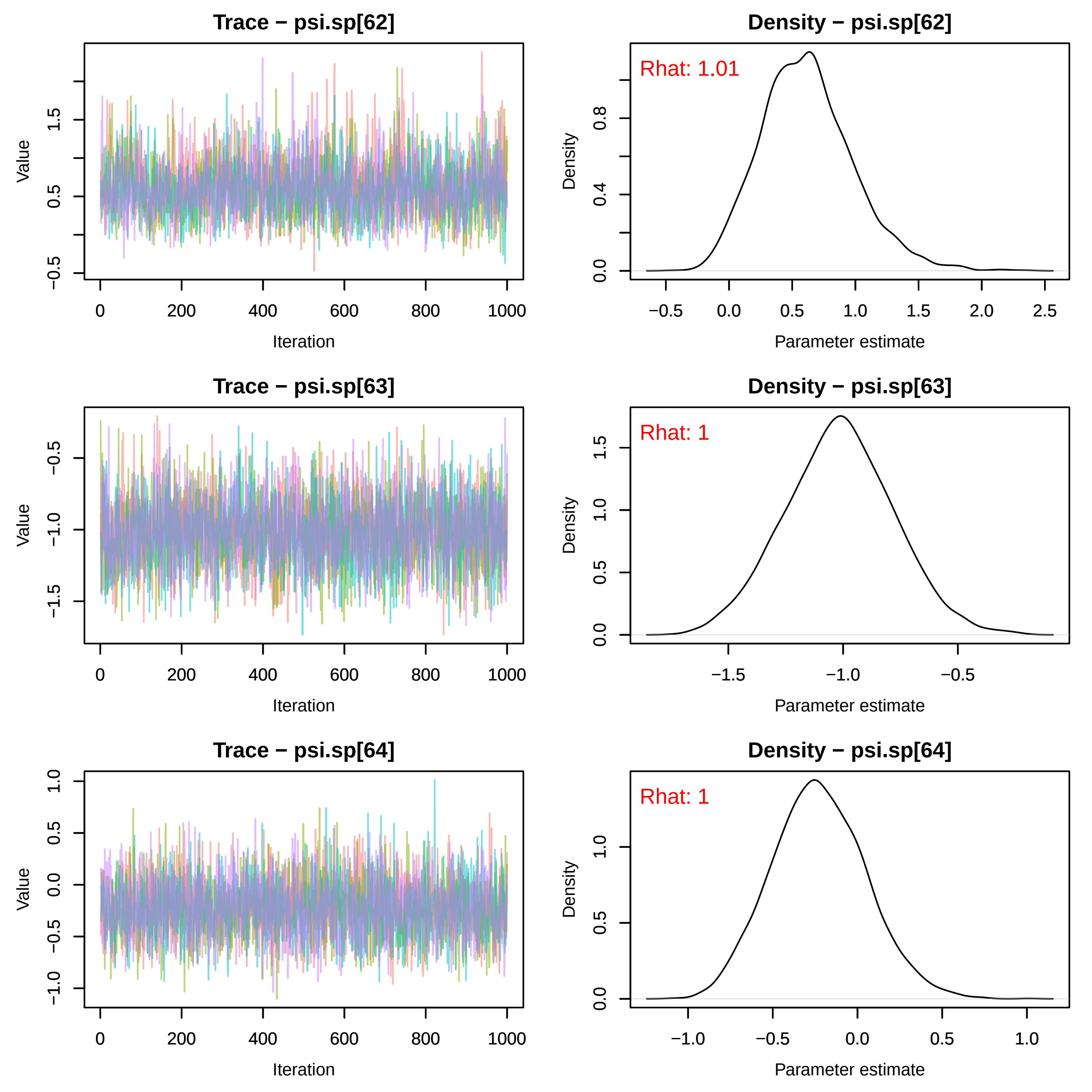
Trace − psi.sp[62]
Value
Iteration
Density − psi.sp[62]
Rhat: 1.01
Density
Parameter estimate
Trace − psi.sp[63]
Value
Iteration
Density − psi.sp[63]
Rhat: 1
Density
Parameter estimate
Trace − psi.sp[64]
Value
Iteration
Density − psi.sp[64]
Rhat: 1
Density
Parameter estimate

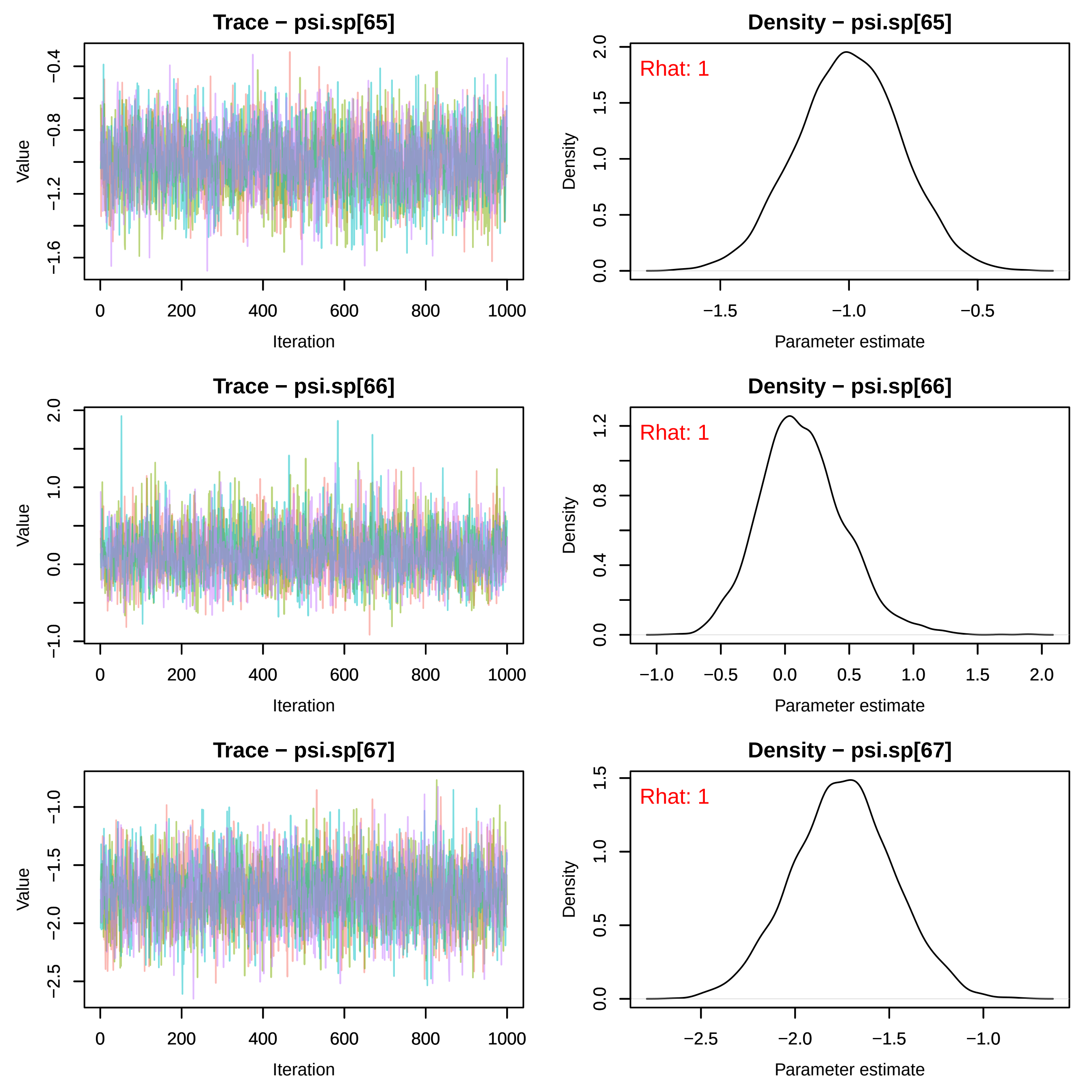
Trace − psi.sp[65]
Density − psi.sp[65]
Rhat: 1
Trace − psi.sp[66]
Density − psi.sp[66]
Rhat: 1
Trace − psi.sp[67]
Density − psi.sp[67]
Rhat: 1
Value
Iteration
Density
Parameter estimate

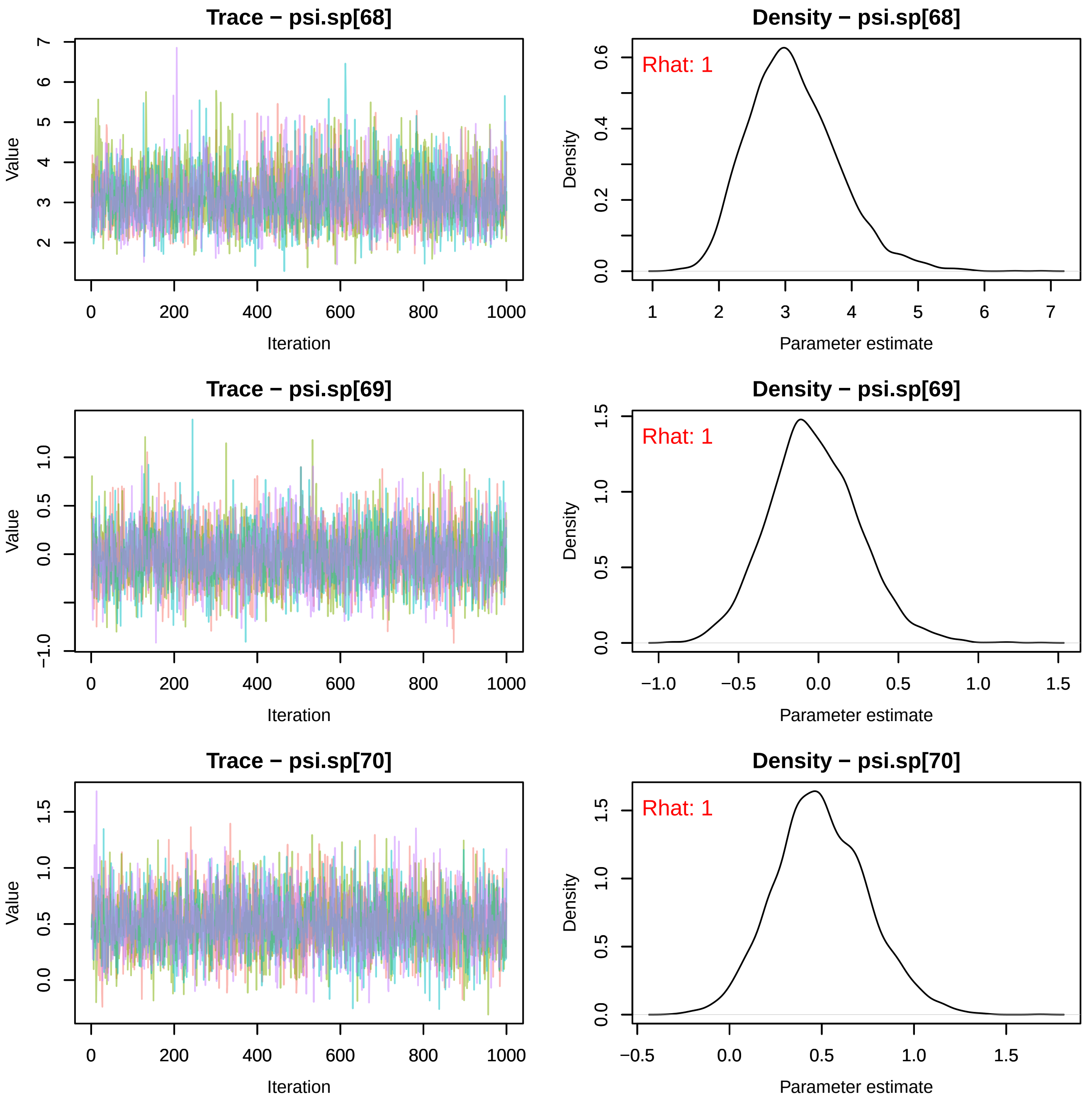

Trace − psi.sp[68]
Value
Iteration
Density − psi.sp[68]
Rhat: 1
Density
Parameter estimate
Trace − psi.sp[69]
Value
Iteration
Density − psi.sp[69]
Rhat: 1
Density
Parameter estimate
Trace − psi.sp[70]
Value
Iteration
Density − psi.sp[70]
Rhat: 1
Density
Parameter estimate

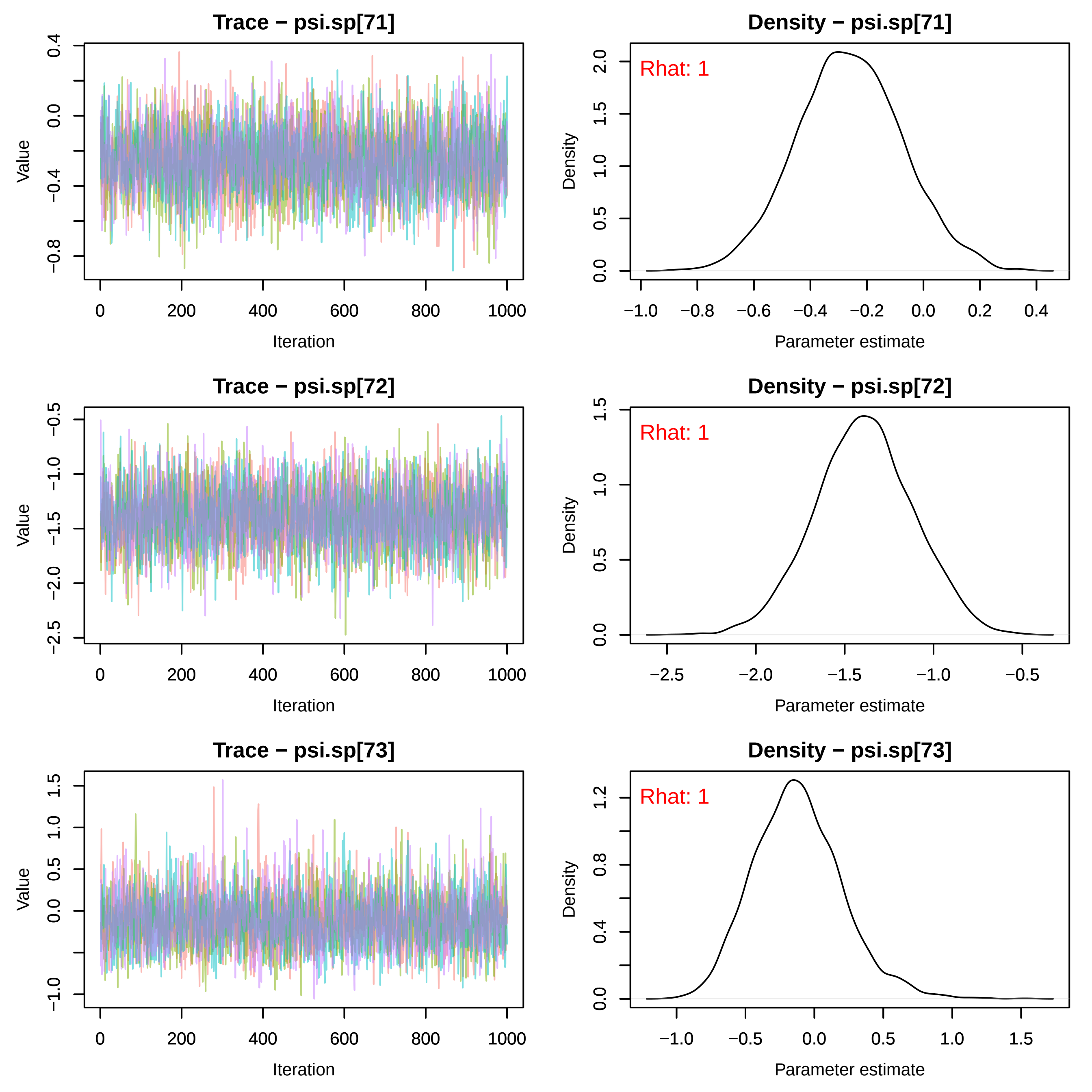
Trace − psi.sp[71]
Value
Iteration
Density − psi.sp[71]
Rhat: 1
Density
Parameter estimate
Trace − psi.sp[72]
Value
Iteration
Density − psi.sp[72]
Rhat: 1
Density
Parameter estimate
Trace − psi.sp[73]
Value
Iteration
Density − psi.sp[73]
Rhat: 1
Density
Parameter estimate

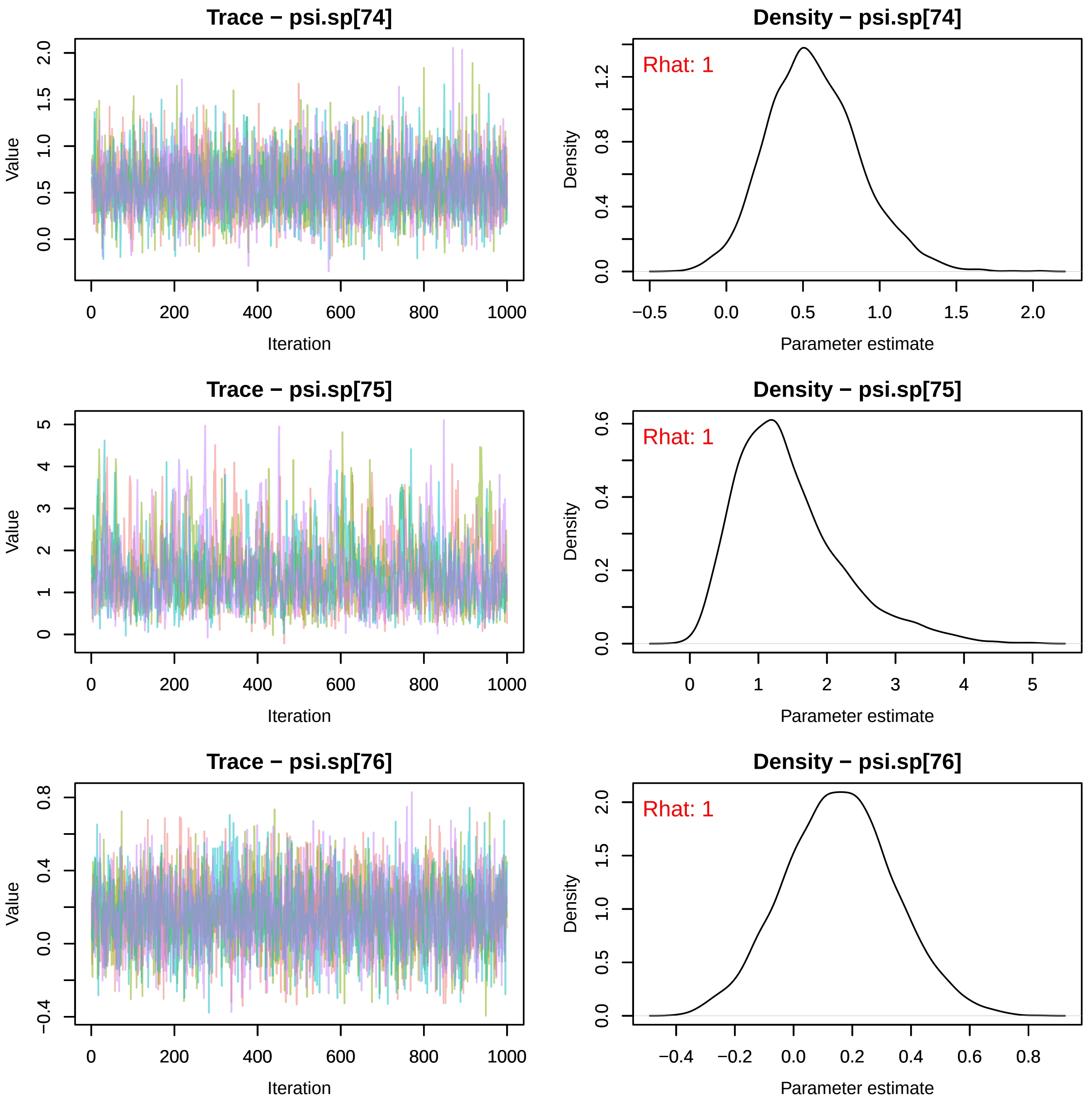
Trace − psi.sp[74]
Density − psi.sp[74]
Rhat: 1
Trace − psi.sp[75]
Density − psi.sp[75]
Rhat: 1
Trace − psi.sp[76]
Density − psi.sp[76]
Rhat: 1
Value
Iteration
Density
Parameter estimate

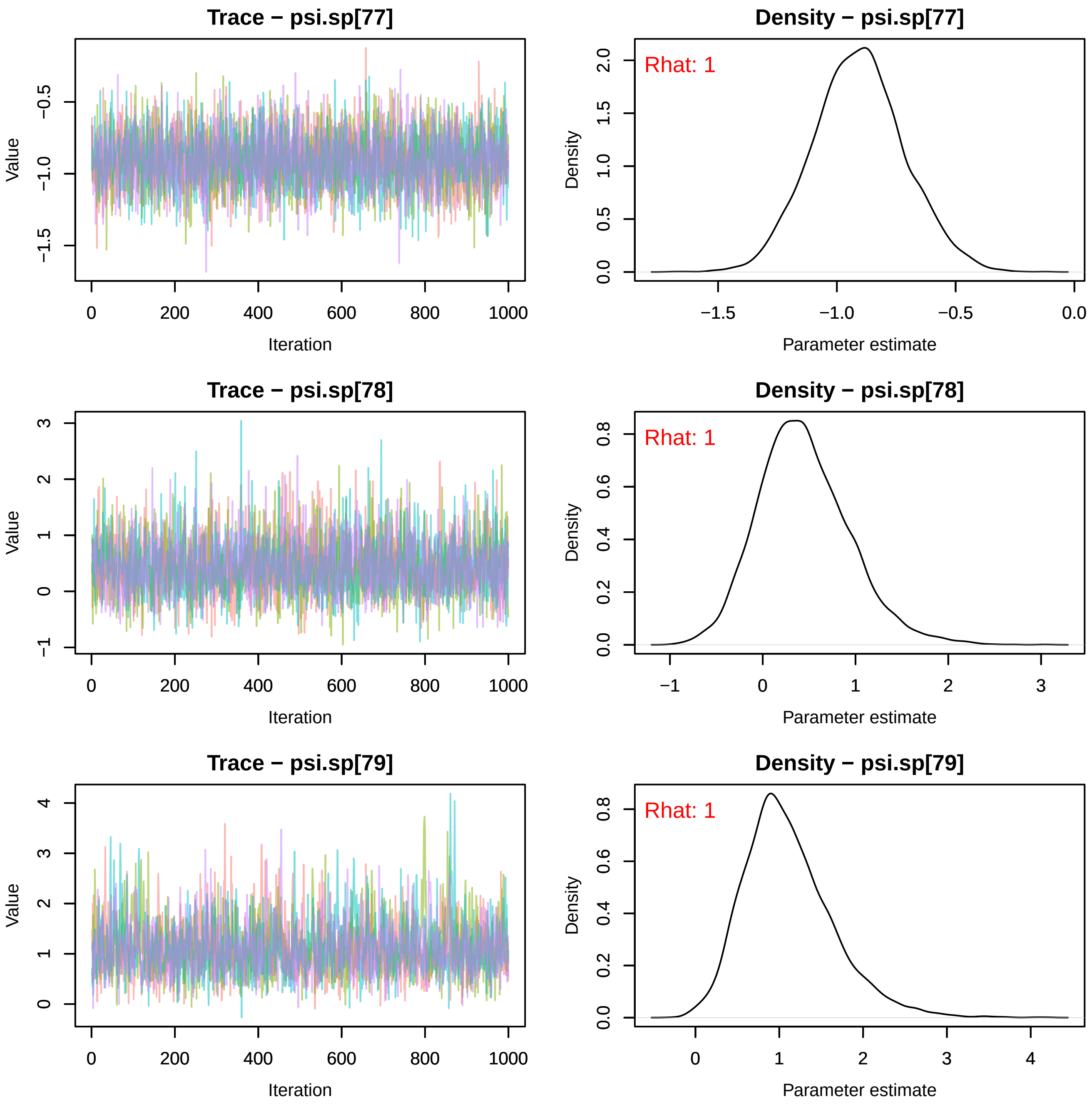
Trace − psi.sp[77]
Value
Iteration
Density − psi.sp[77]
Rhat: 1
Density
Parameter estimate
Trace − psi.sp[78]
Value
Iteration
Density − psi.sp[78]
Rhat: 1
Density
Parameter estimate
Trace − psi.sp[79]
Value
Iteration
Density − psi.sp[79]
Rhat: 1
Density
Parameter estimate

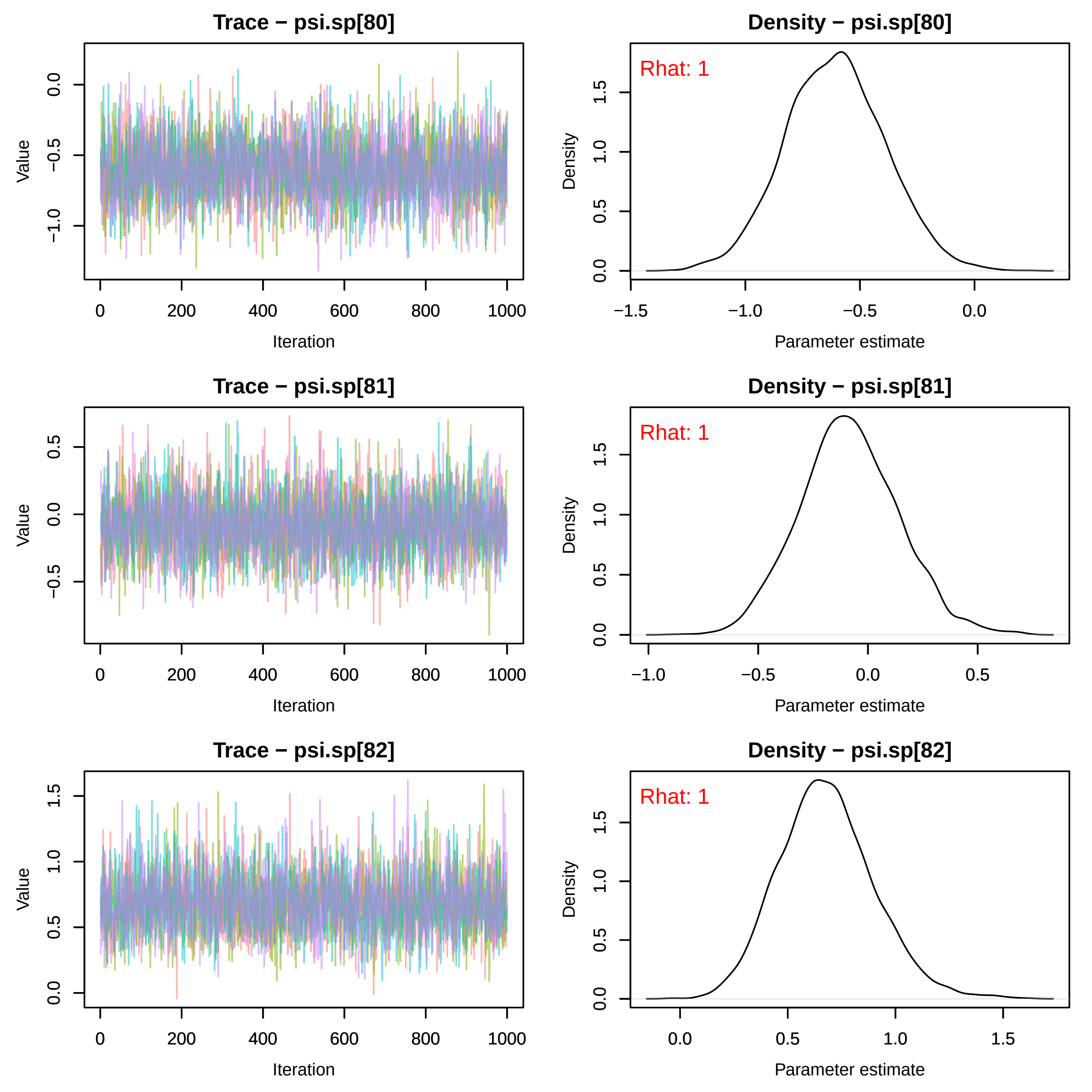
Trace − psi.sp[80]
Density − psi.sp[80]
Rhat: 1
Trace − psi.sp[81]
Density − psi.sp[81]
Rhat: 1
Trace − psi.sp[82]
Density − psi.sp[82]
Rhat: 1
Value
Iteration
Density
Parameter estimate

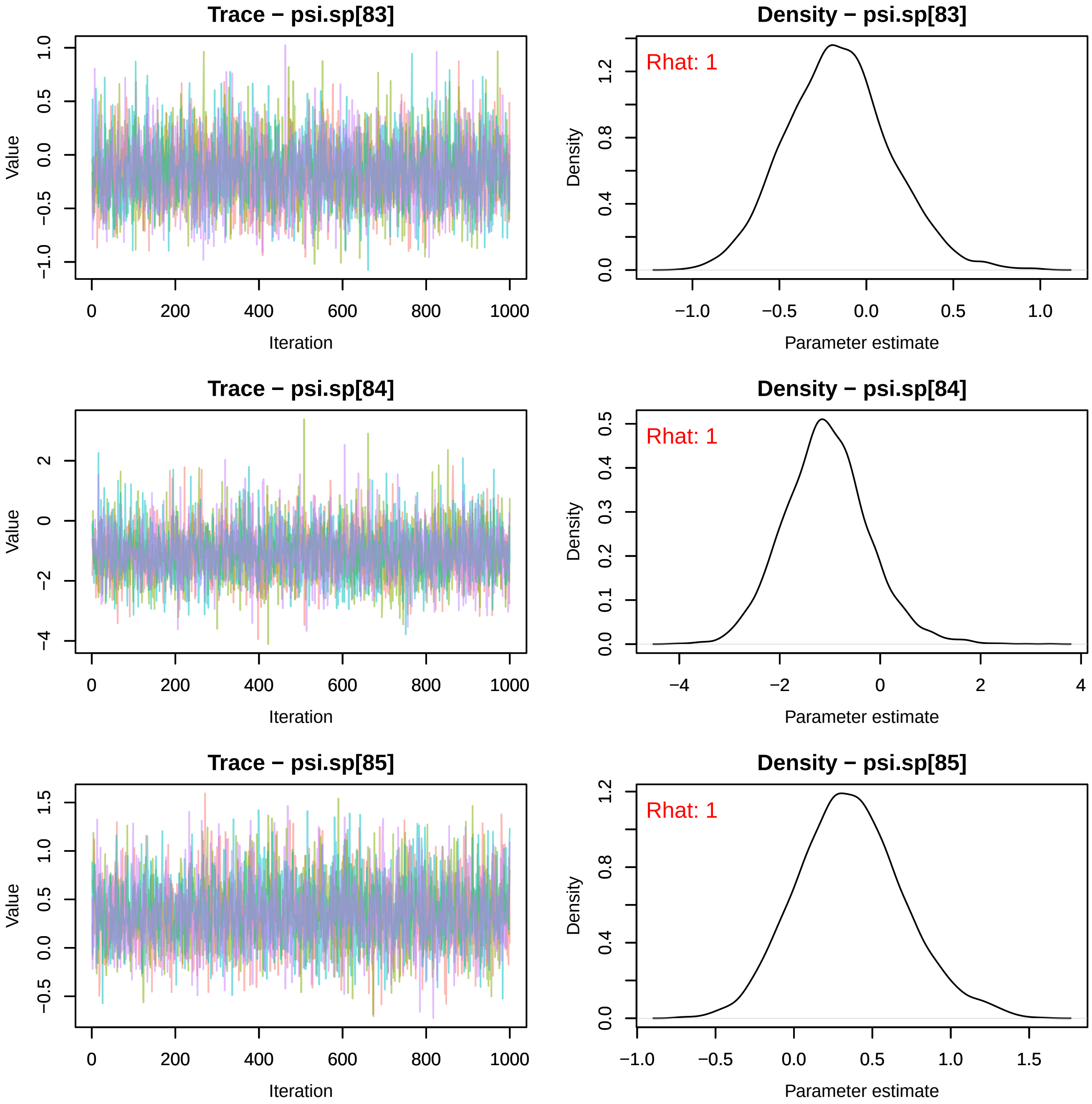
Trace − psi.sp[83]
Density − psi.sp[83]
Rhat: 1
Trace − psi.sp[84]
Density − psi.sp[84]
Rhat: 1
Trace − psi.sp[85]
Density − psi.sp[85]
Rhat: 1
Value
Iteration
Density
Parameter estimate

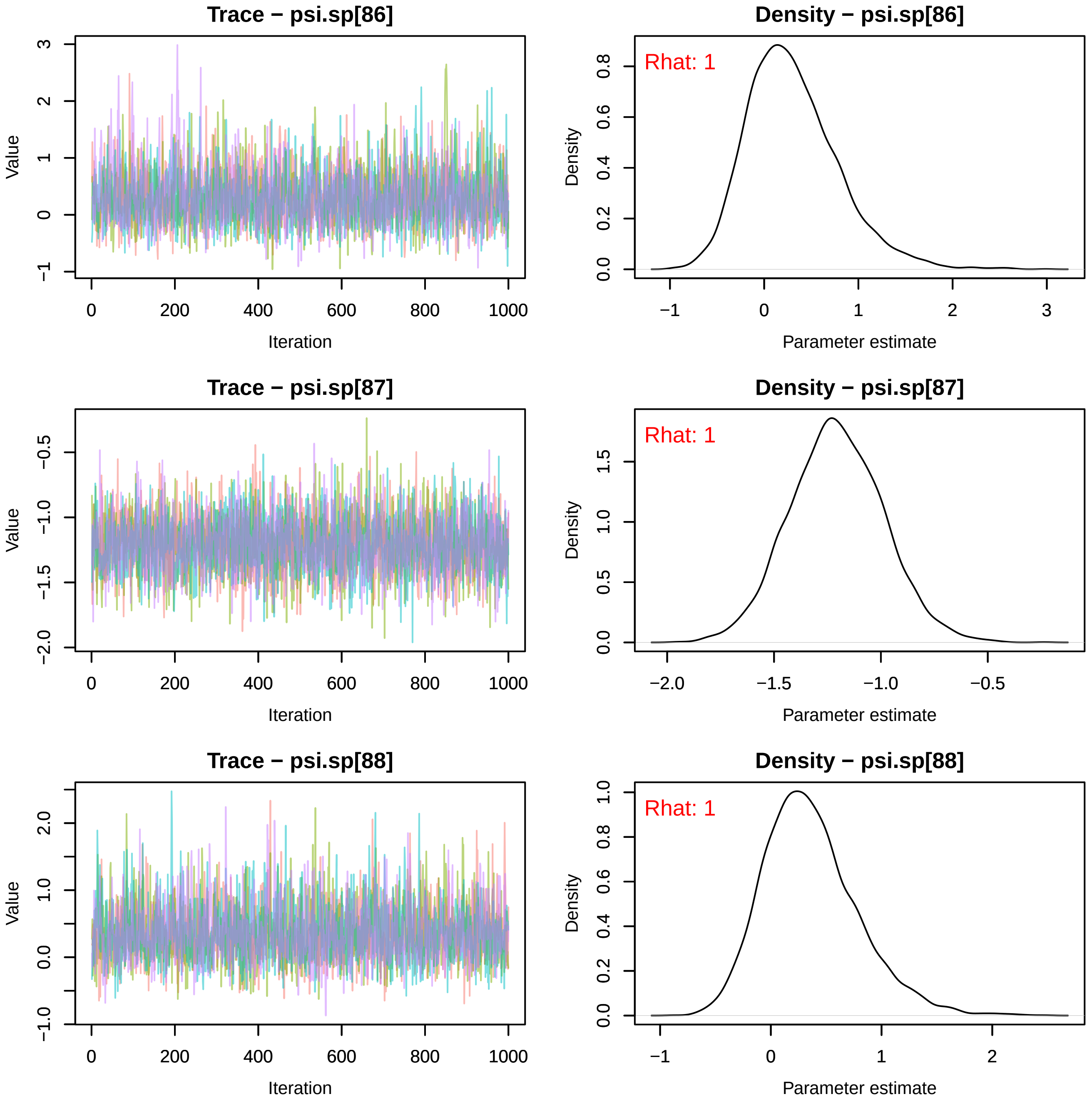

Trace − psi.sp[86]
Value
Iteration
Density − psi.sp[86]
Rhat: 1
Density
Parameter estimate
Trace − psi.sp[87]
Value
Iteration
Density − psi.sp[87]
Rhat: 1
Density
Parameter estimate
Trace − psi.sp[88]
Value
Iteration
Density − psi.sp[88]
Rhat: 1
Density
Parameter estimate

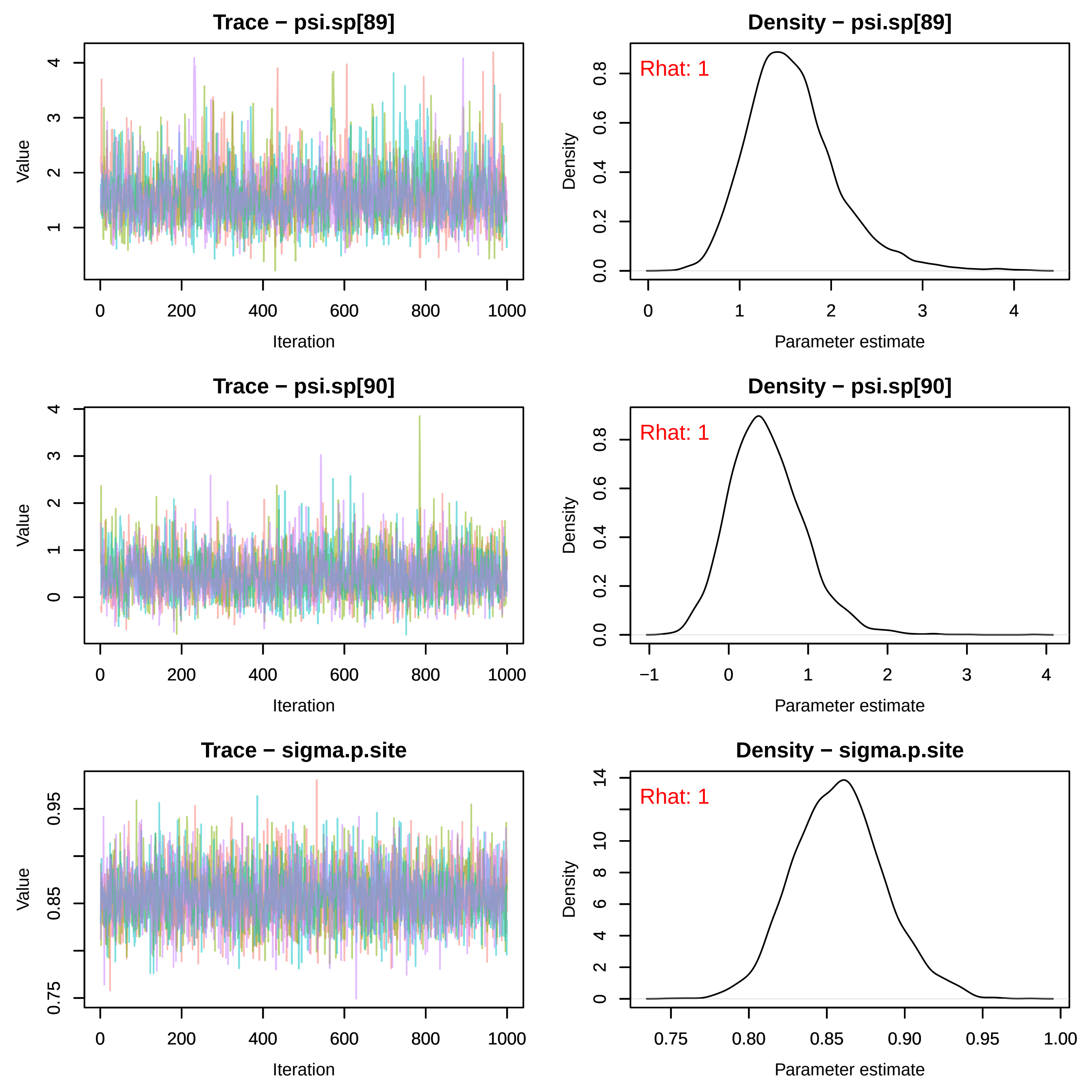
Trace − psi.sp[89]
Density − psi.sp[89]
Rhat: 1
Trace − psi.sp[90]
Density − psi.sp[90]
Rhat: 1
Trace − sigma.p.site
Density − sigma.p.site
Rhat: 1
Value
Iteration
Density
Parameter estimate

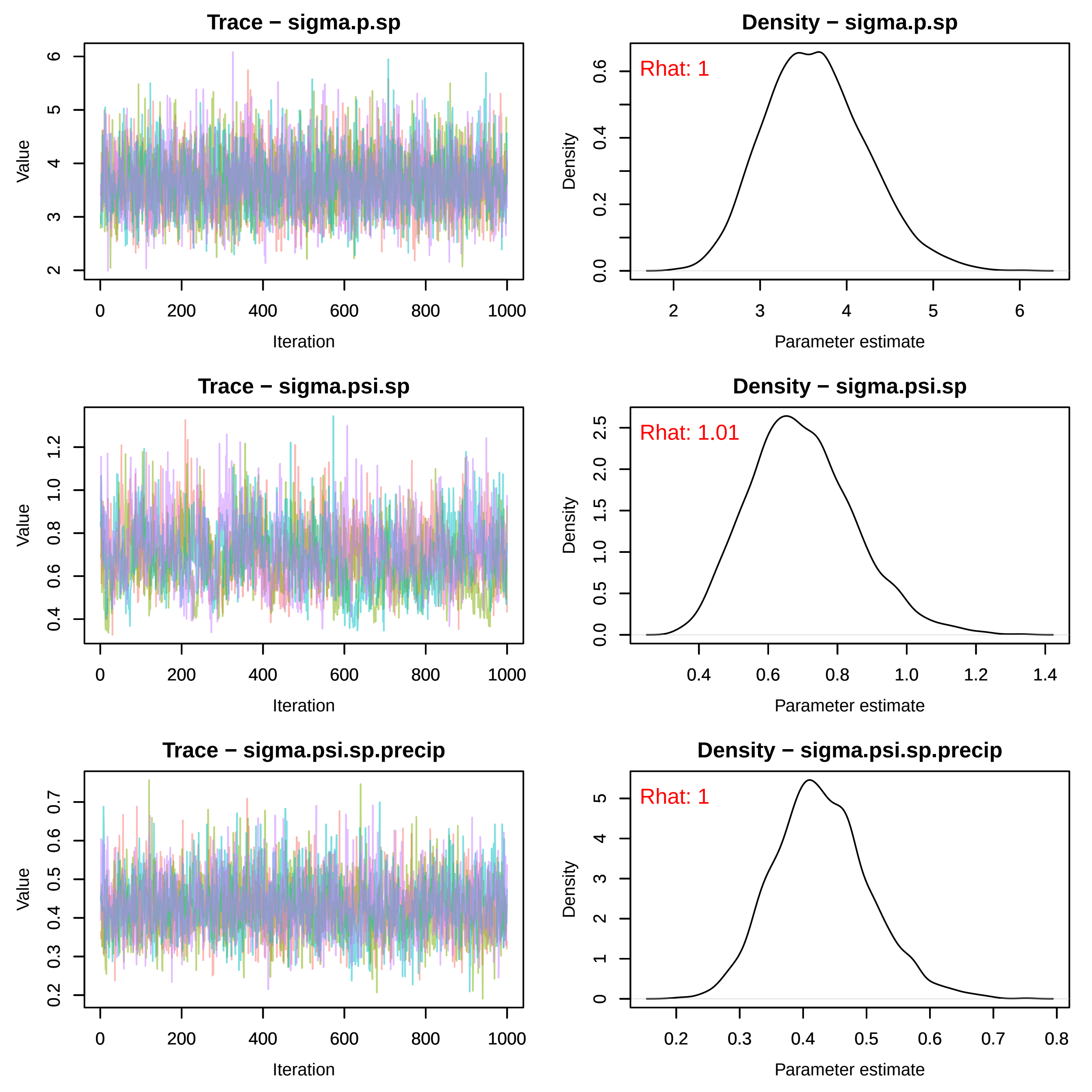
Trace − sigma.p.sp
Value
Iteration
Density − sigma.p.sp
Rhat: 1
Density
Parameter estimate
Trace − sigma.psi.sp
Value
Iteration
Density − sigma.psi.sp
Rhat: 1.01
Density
Parameter estimate
Trace − sigma.psi.sp.precip
Value
Iteration
Density − sigma.psi.sp.precip
Rhat: 1
Density
Parameter estimate
