## Supplemental File S6 for "Rising minimum temperatures contribute to 50 years of shifting Arctic and boreal butterfly communities in North America"

*Vanessa virginiensis*

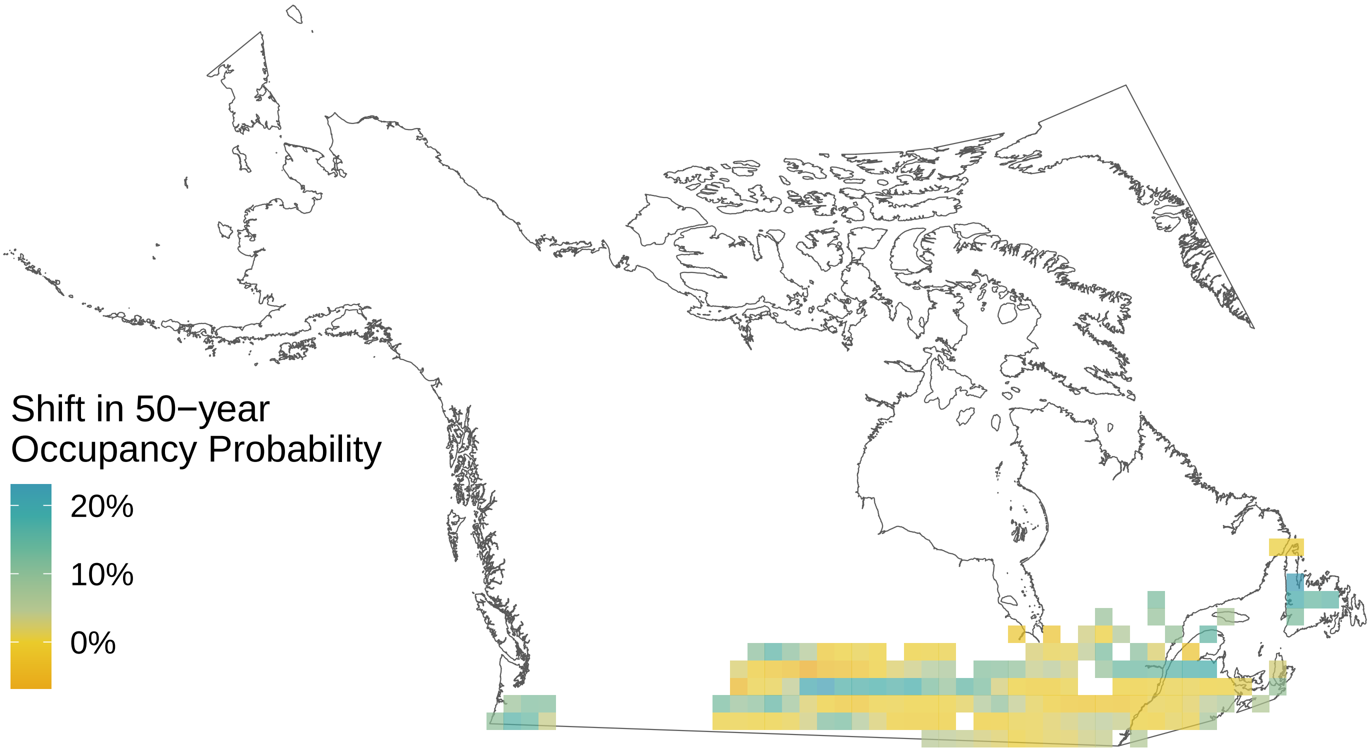

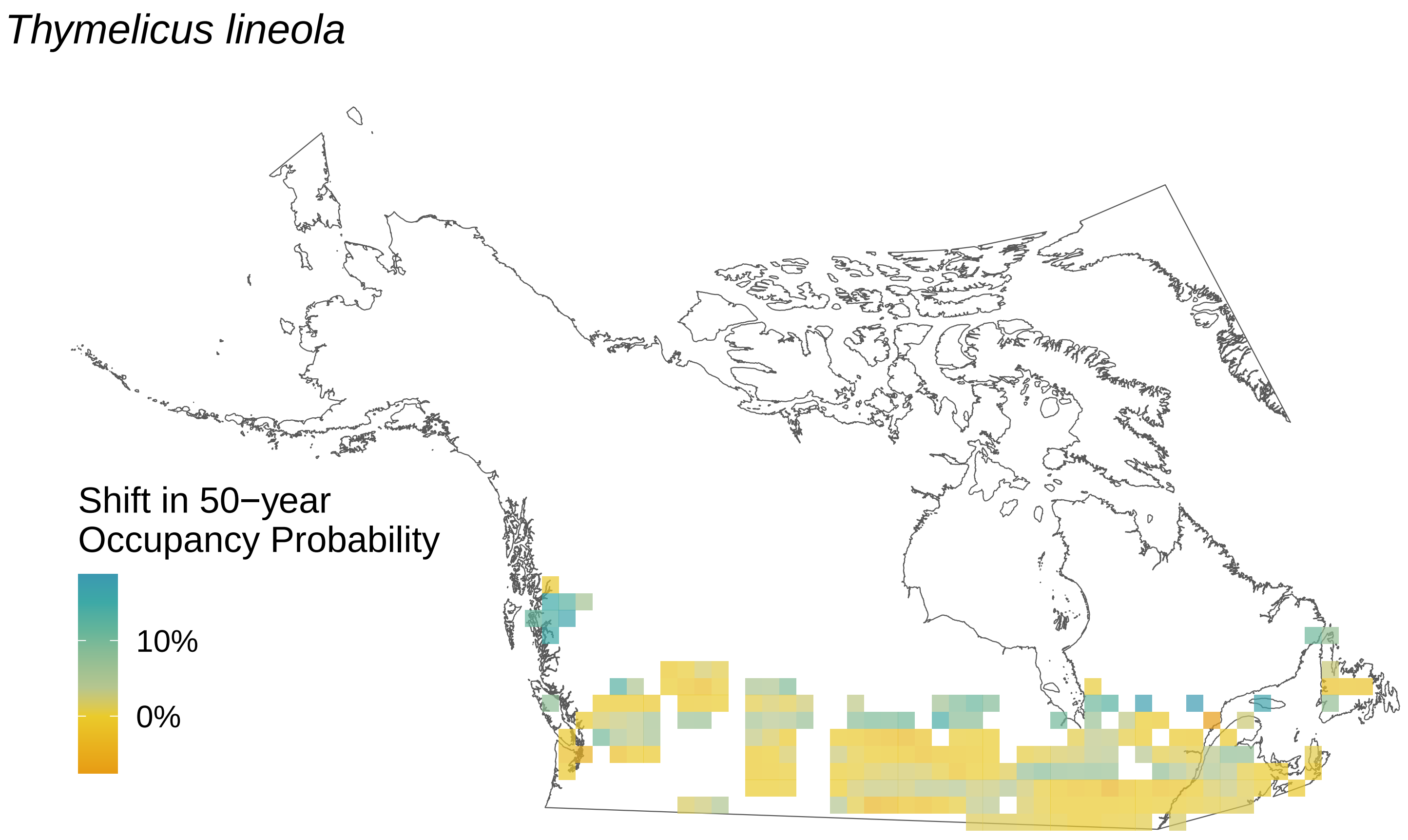
Thymelicus lineola
Shift in 50−year
Occupancy Probability
10%
0%

*Thorybes pylades*

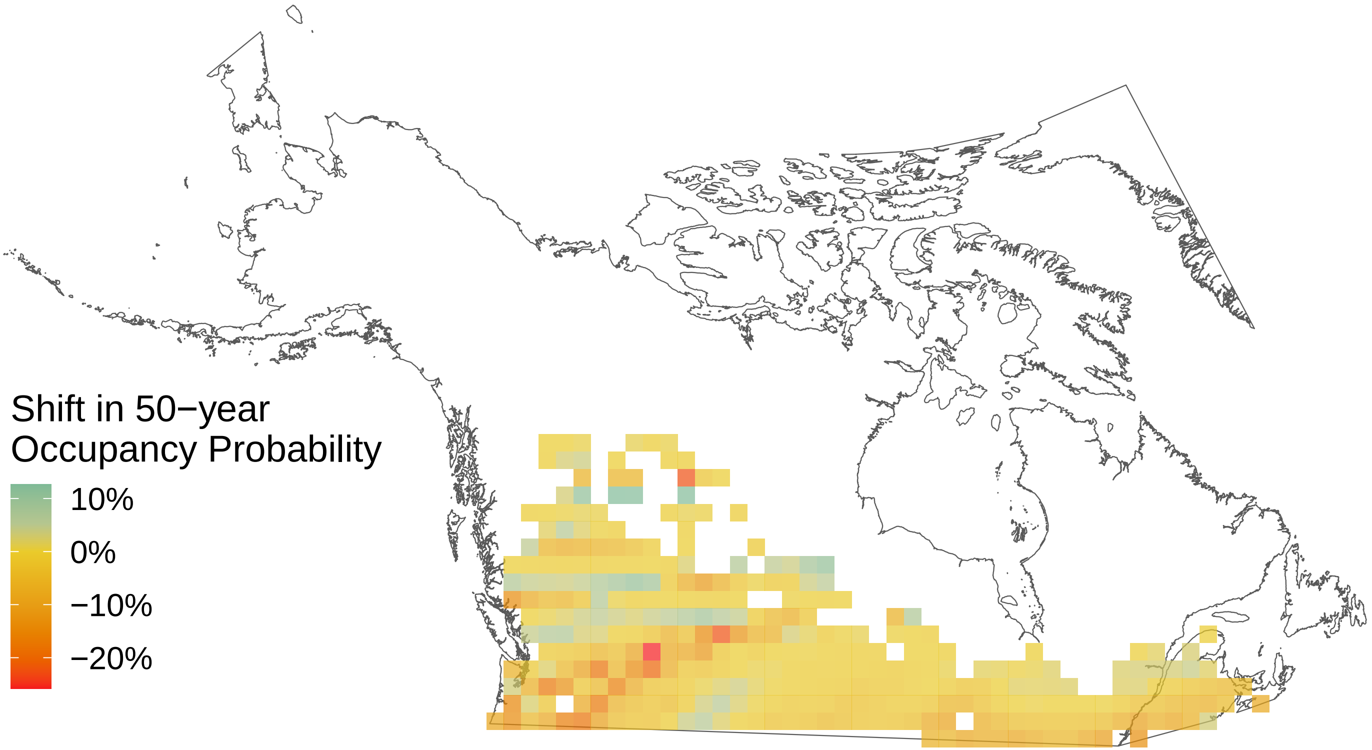

*Strymon melinus*

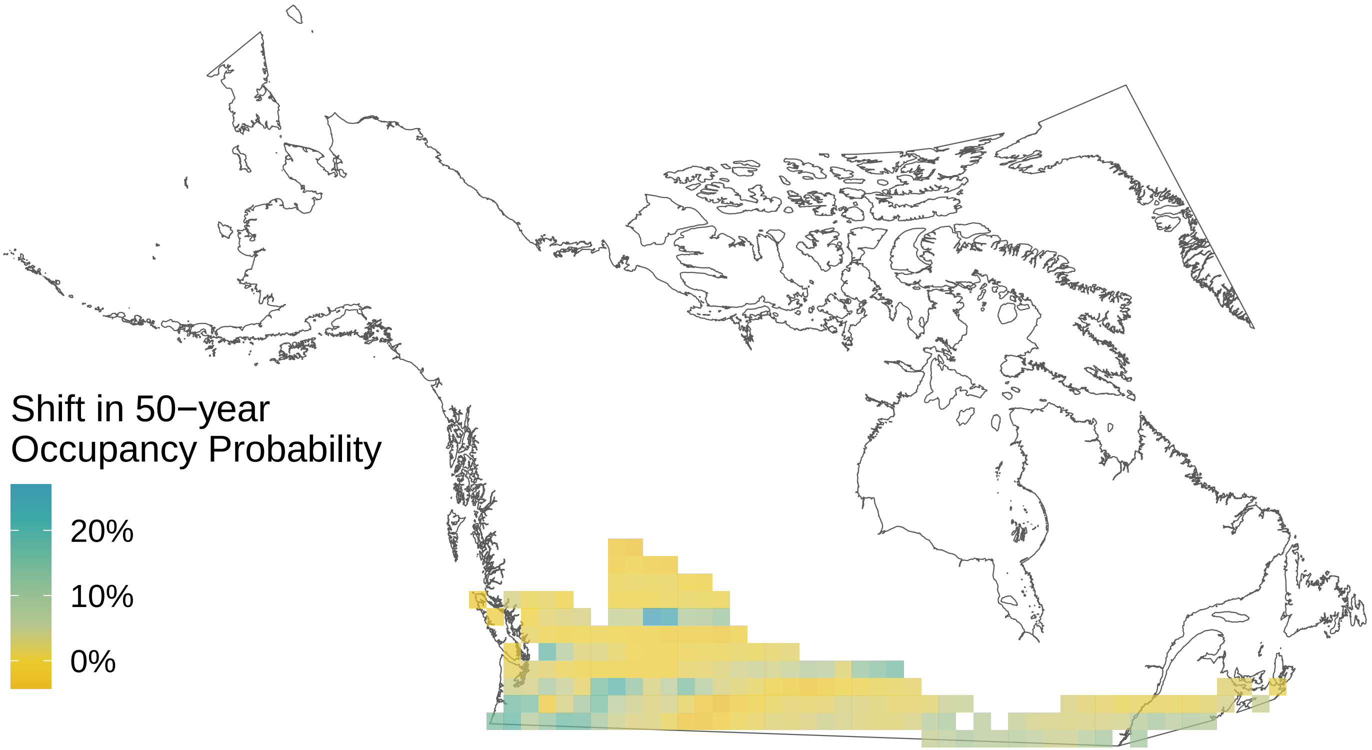

*Pterourus rutulus*

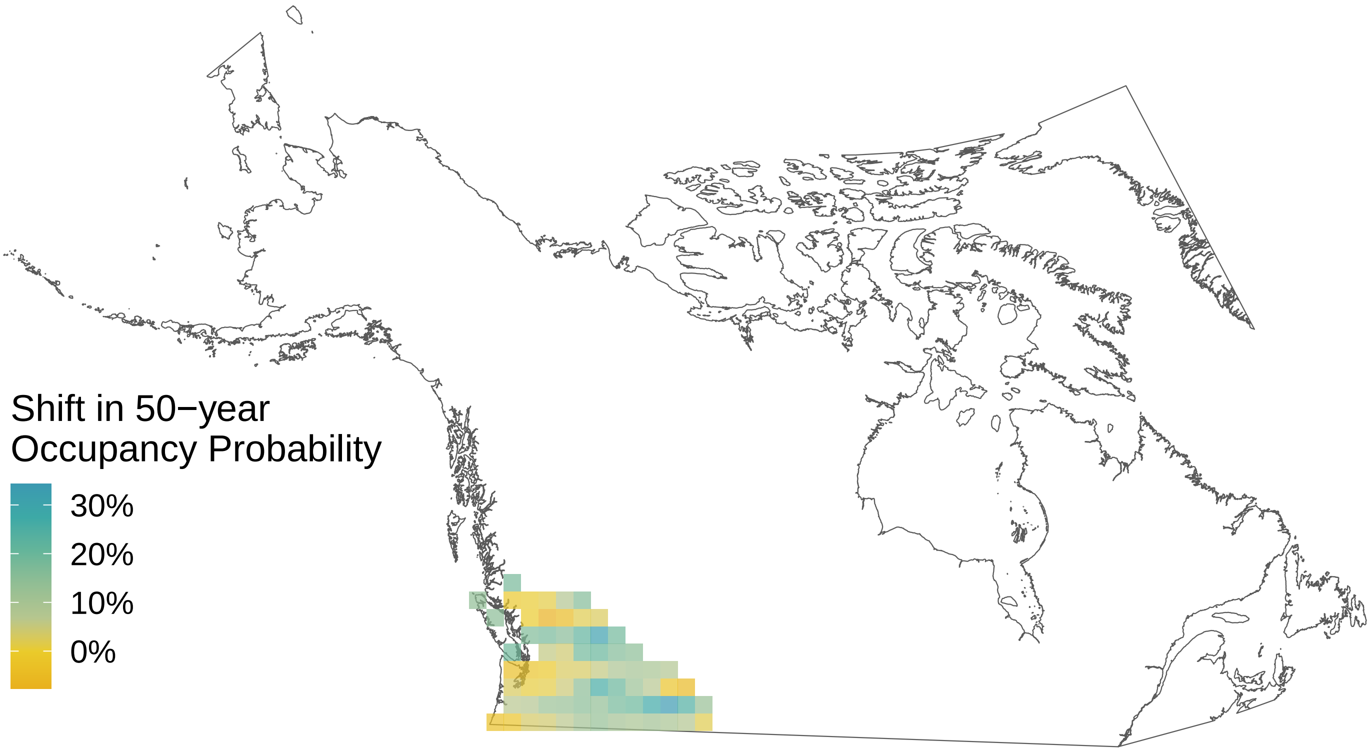

*Satyrium liparops*

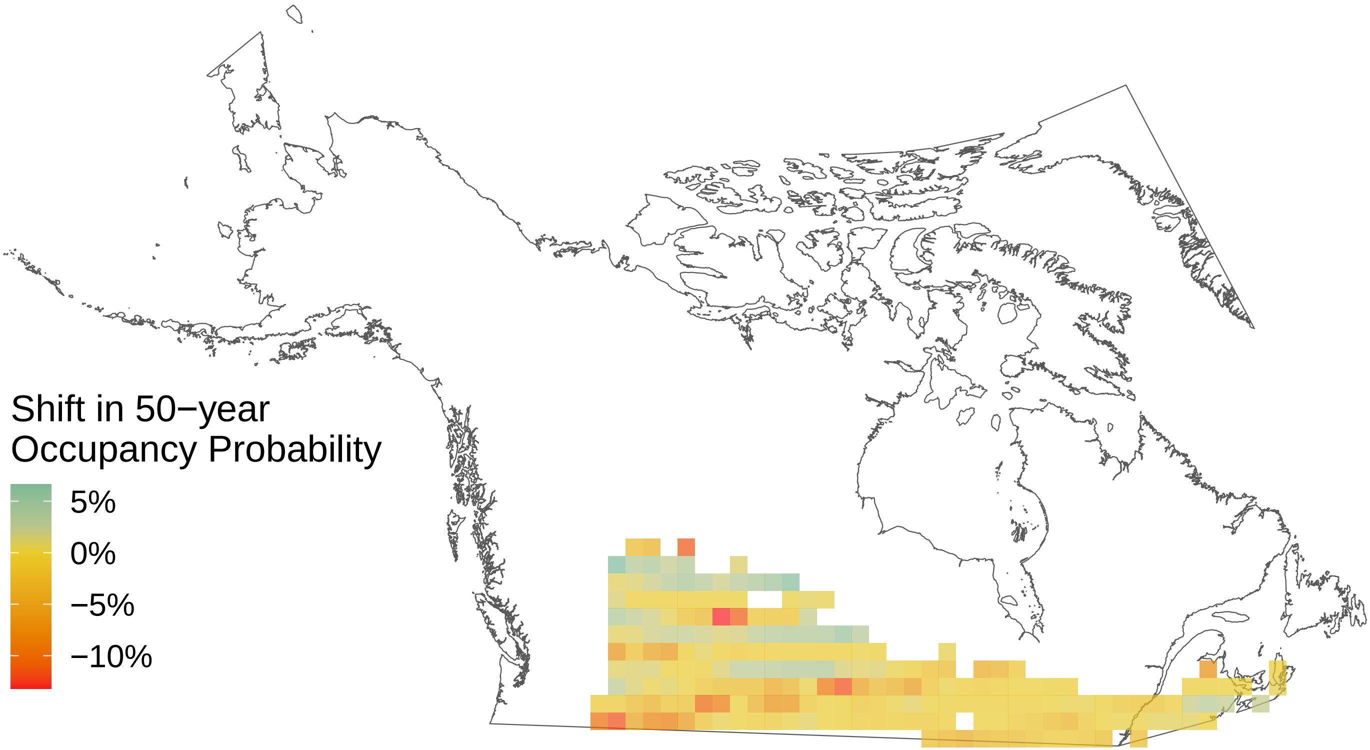

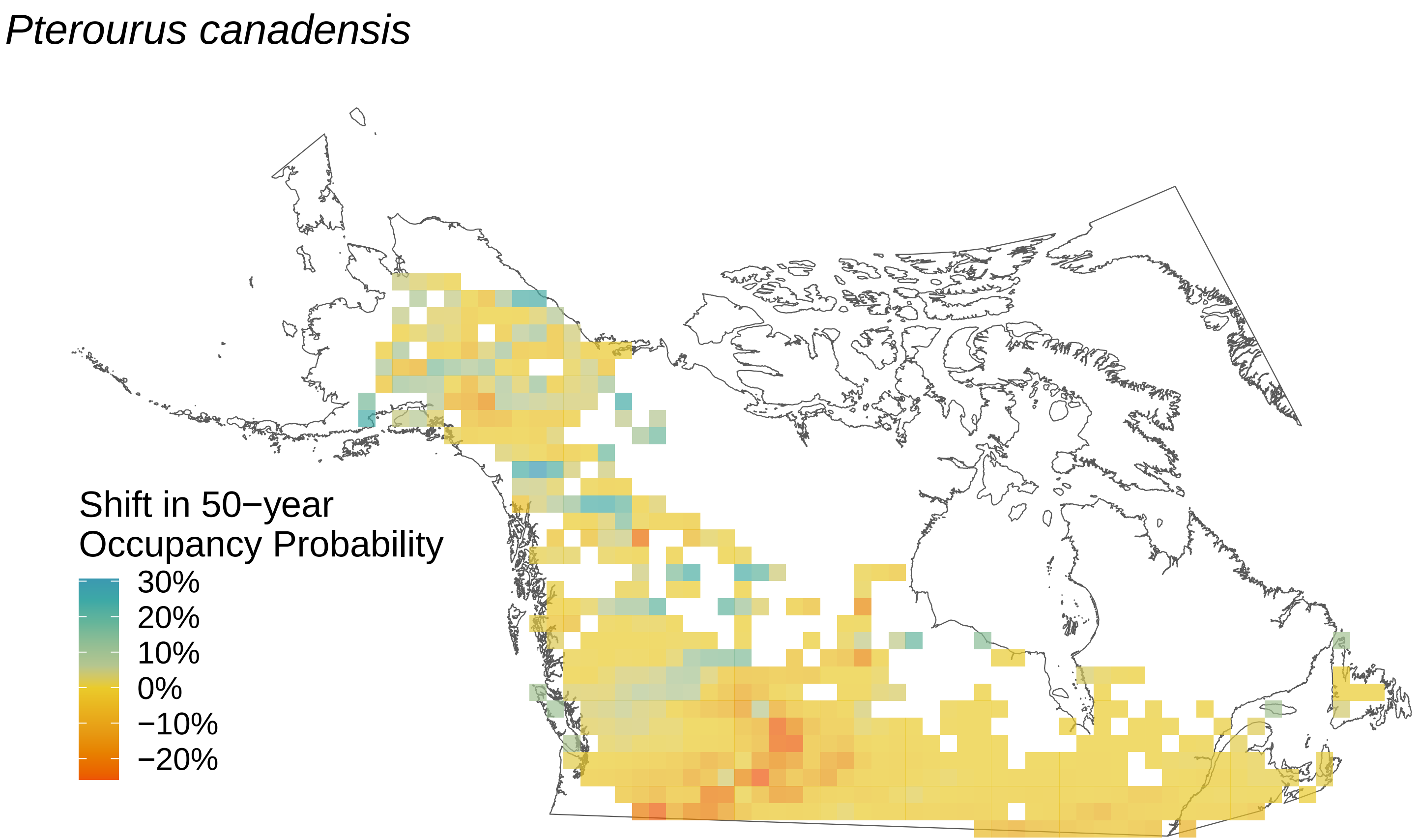
*Pterourus canadensis*
Shift in 50−year
Occupancy Probability
30%
20%
10%
0%
−10%
−20%

### *Pterourus eurymedon*

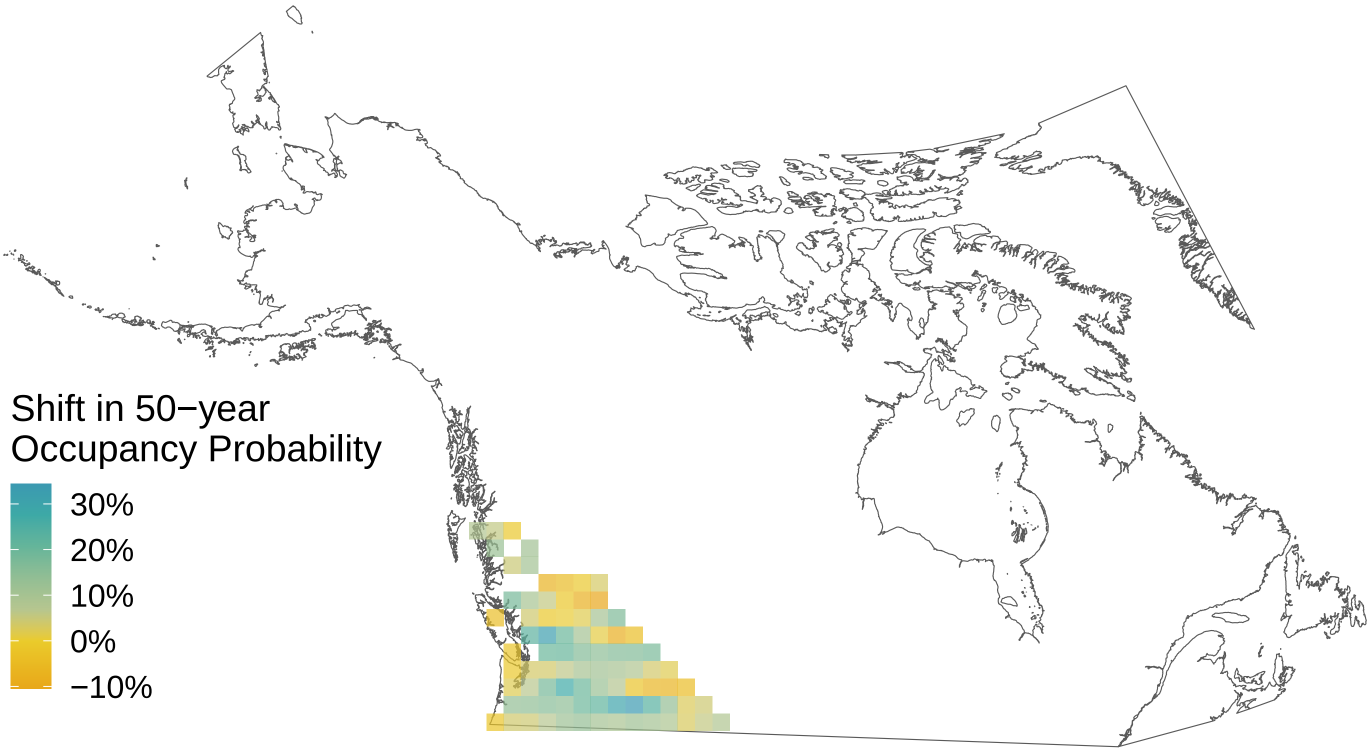

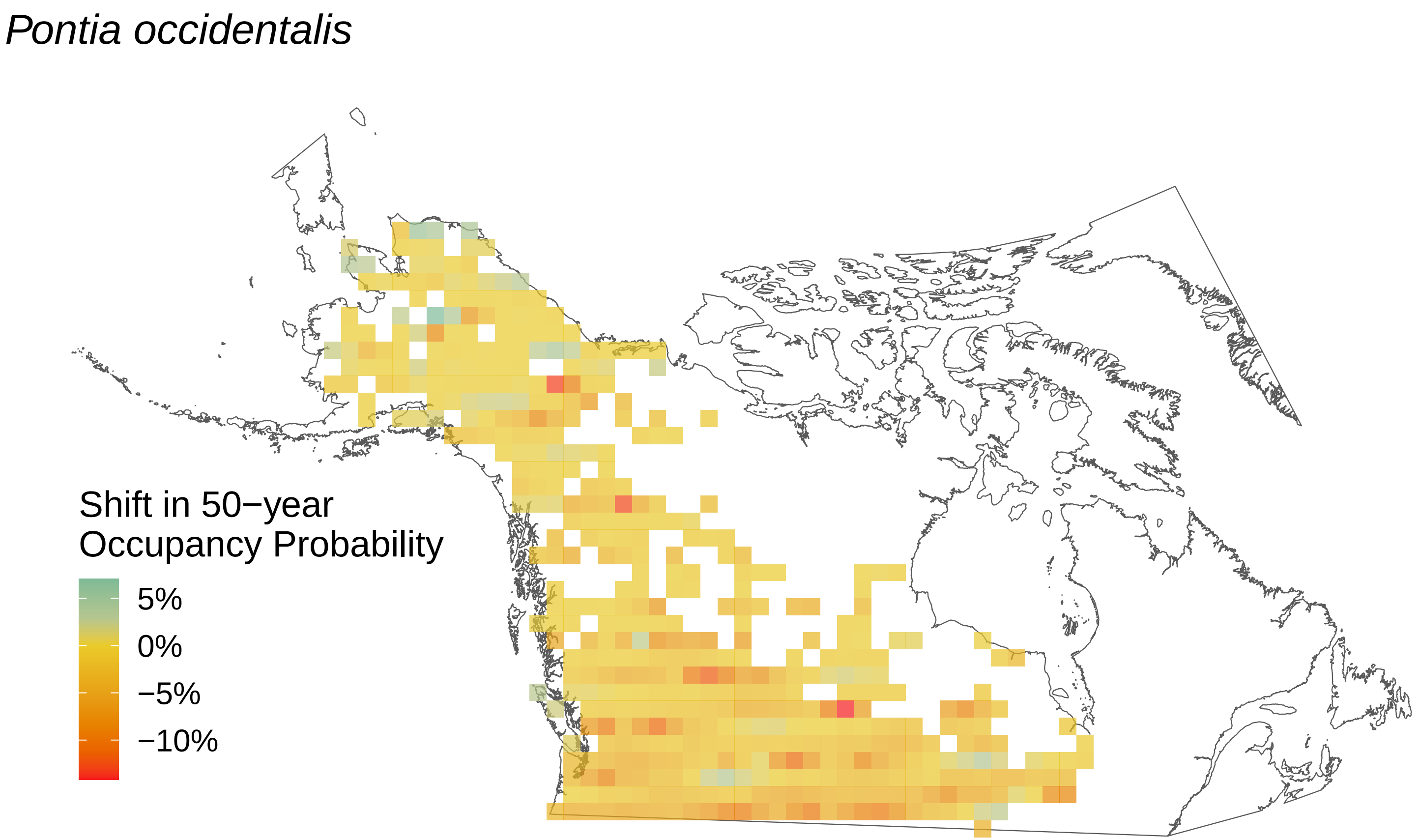
Pontia occidentalis
Shift in 50−year
Occupancy Probability
5%
0%
−5%
−10%

*Polygonia satyrus*

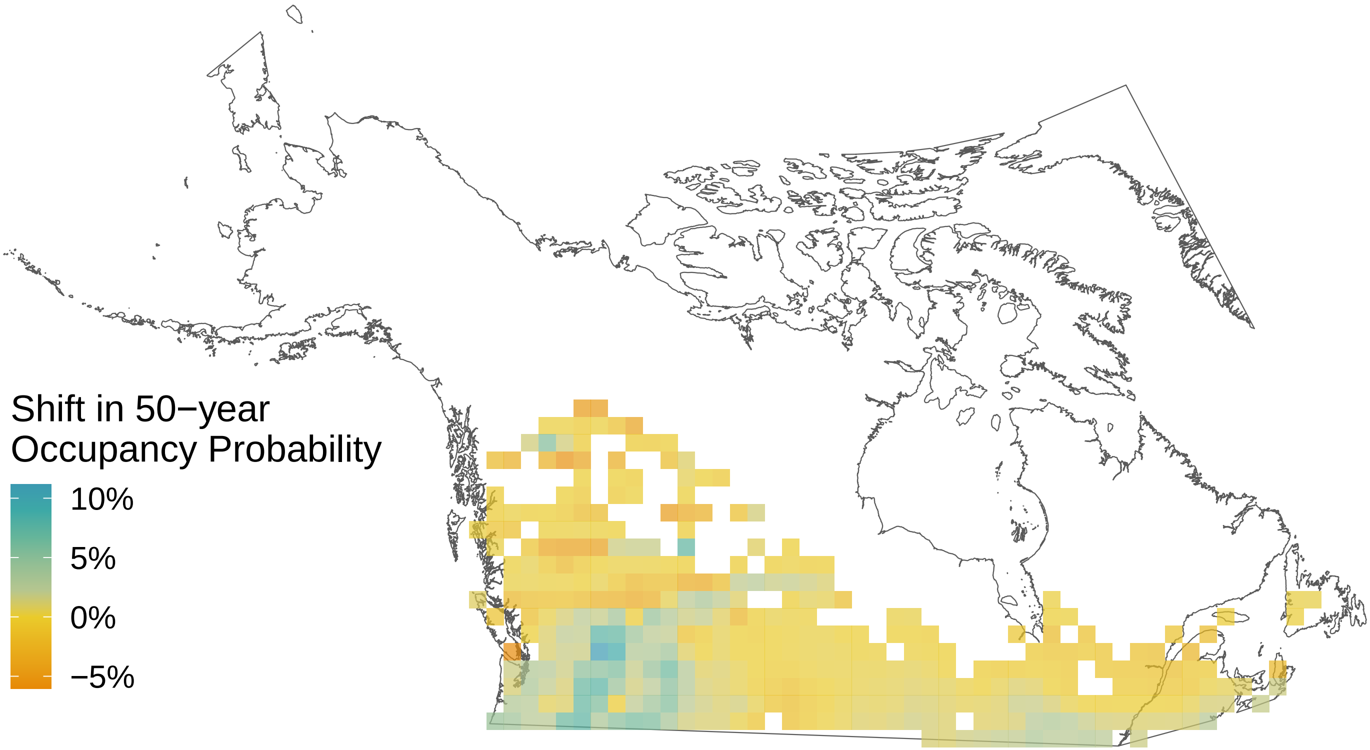

*Polygonia interrogationis*

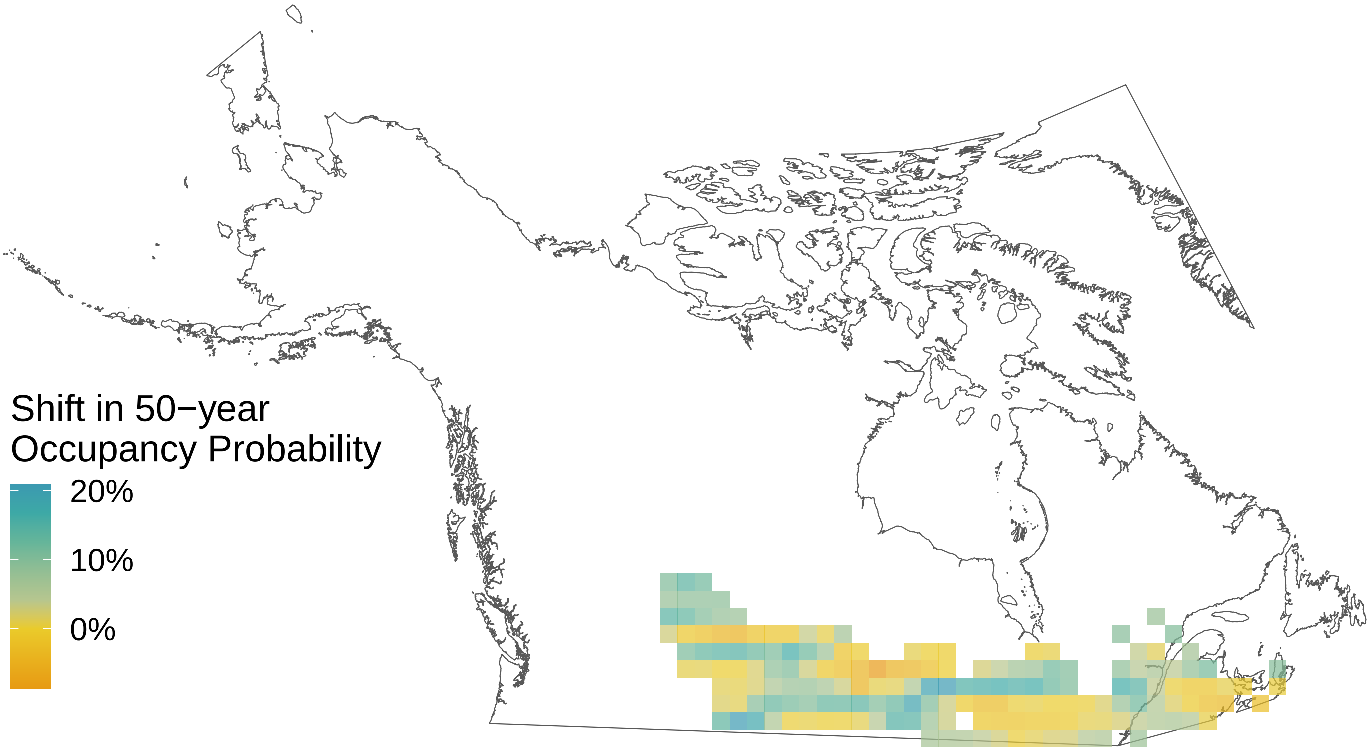

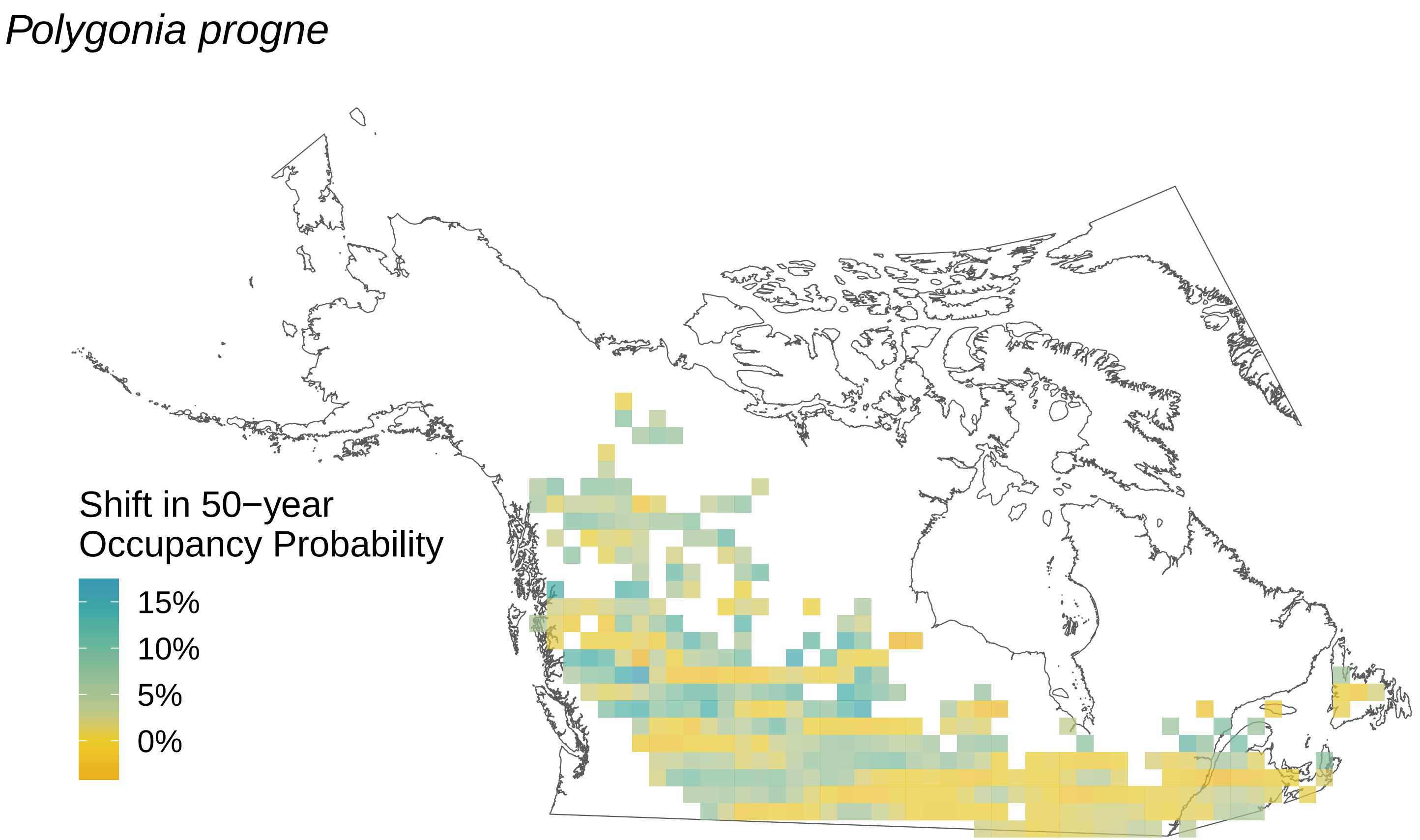
Polygonia progne
Shift in 50−year
Occupancy Probability
15%
10%
5%
0%

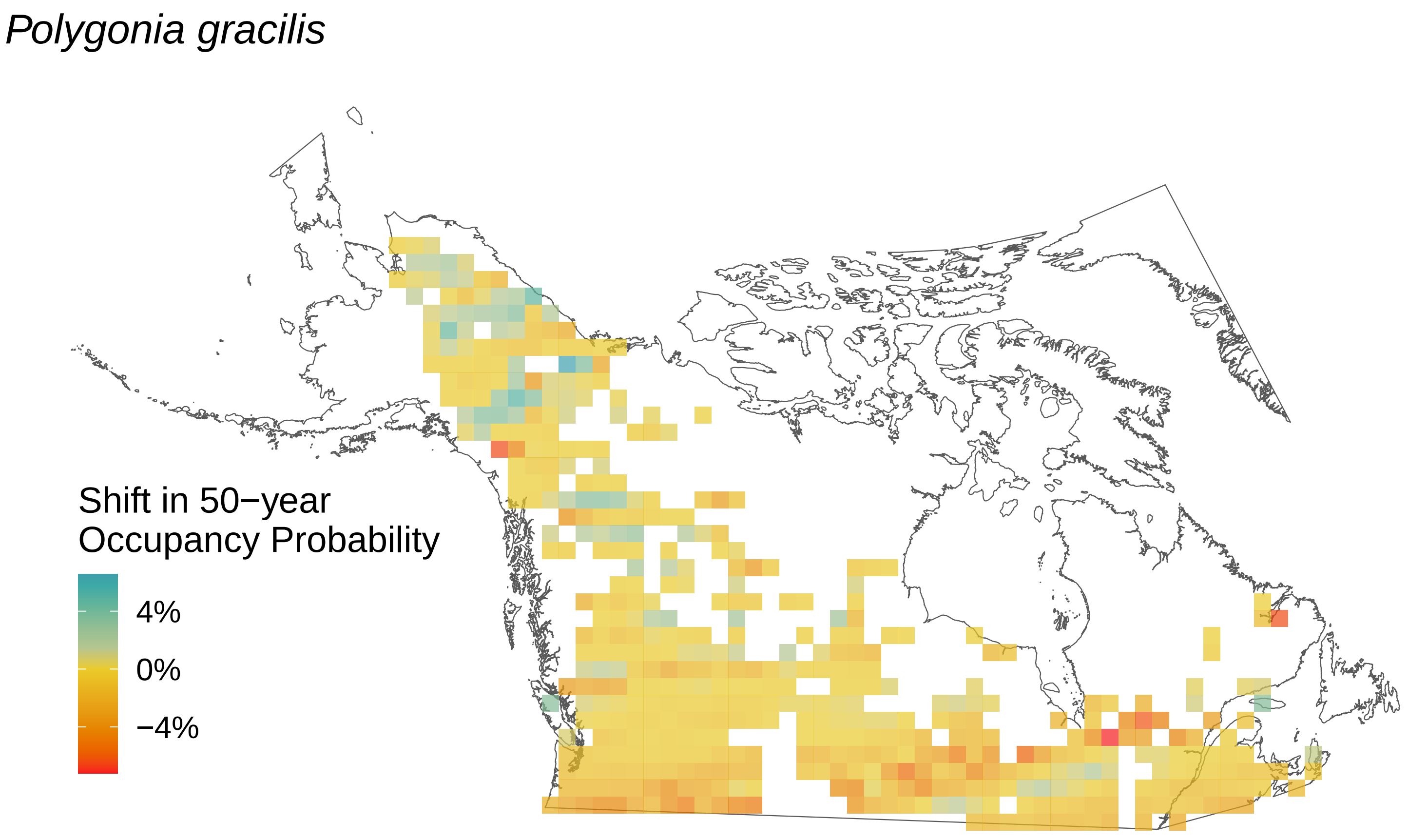

Polygonia gracilis
Shift in 50−year
Occupancy Probability
4%
0%
−4%

### *Polygonia faunus*

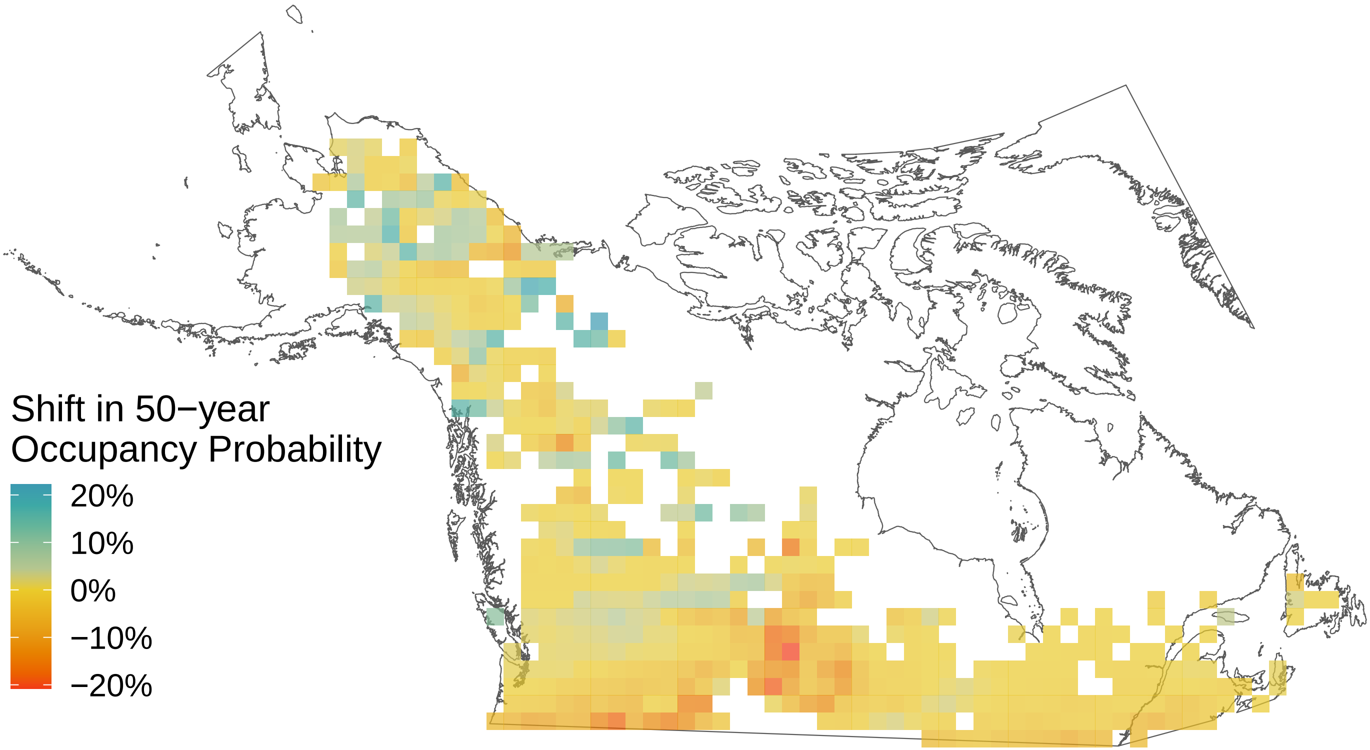

*Polites mystic*

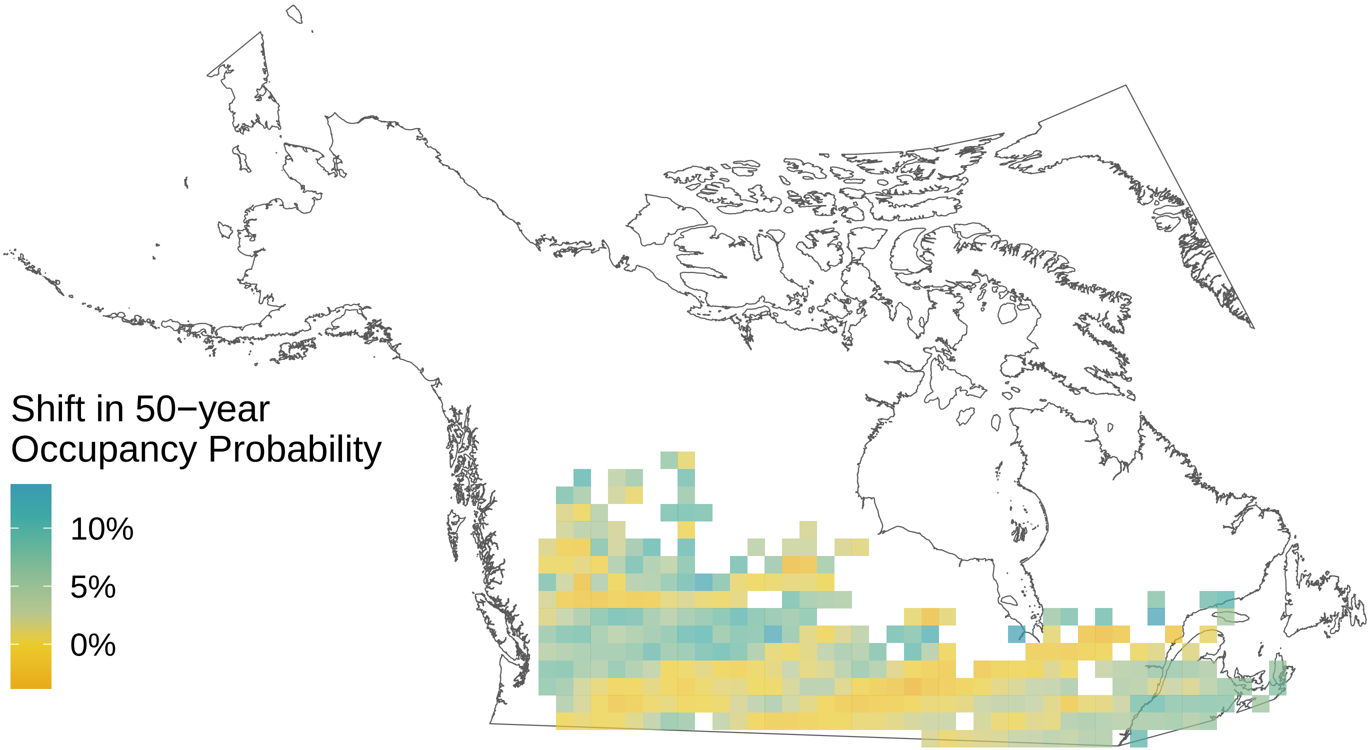

*Polites themistocles*

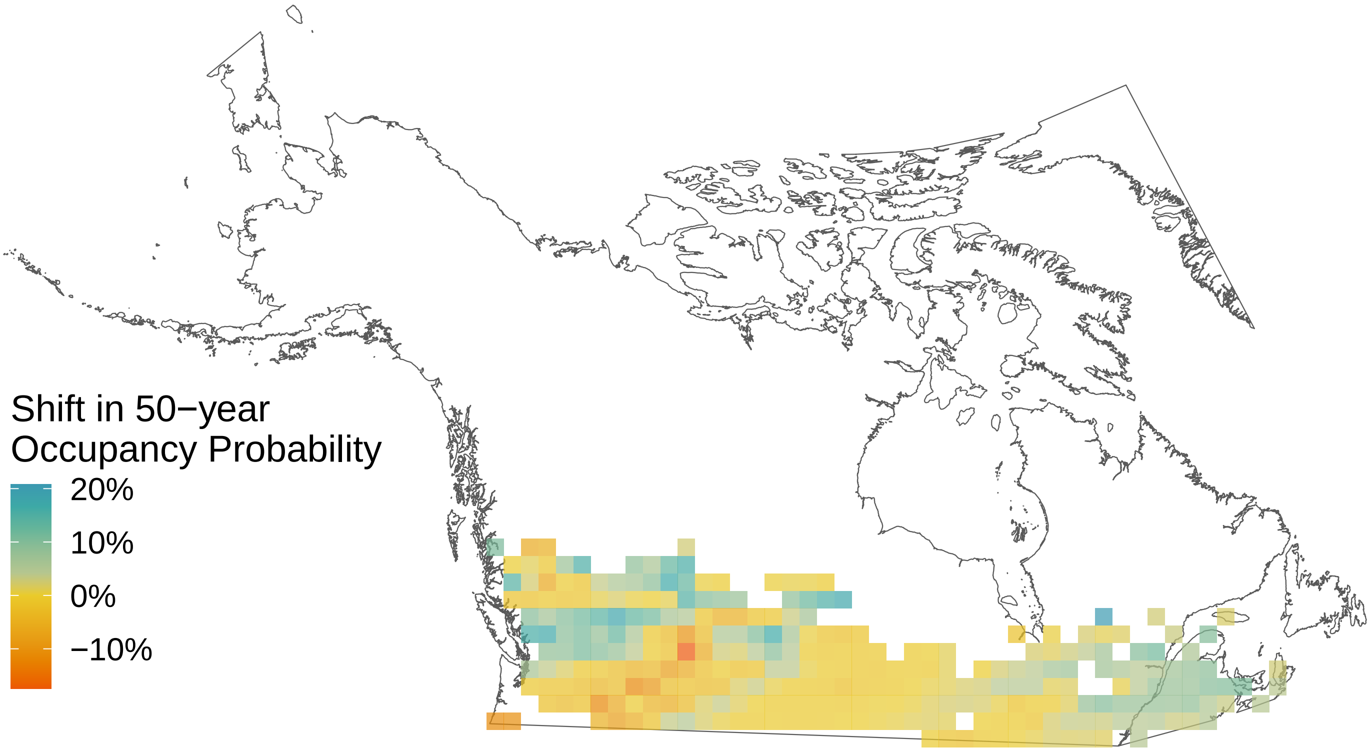

*Poanes hobomok*

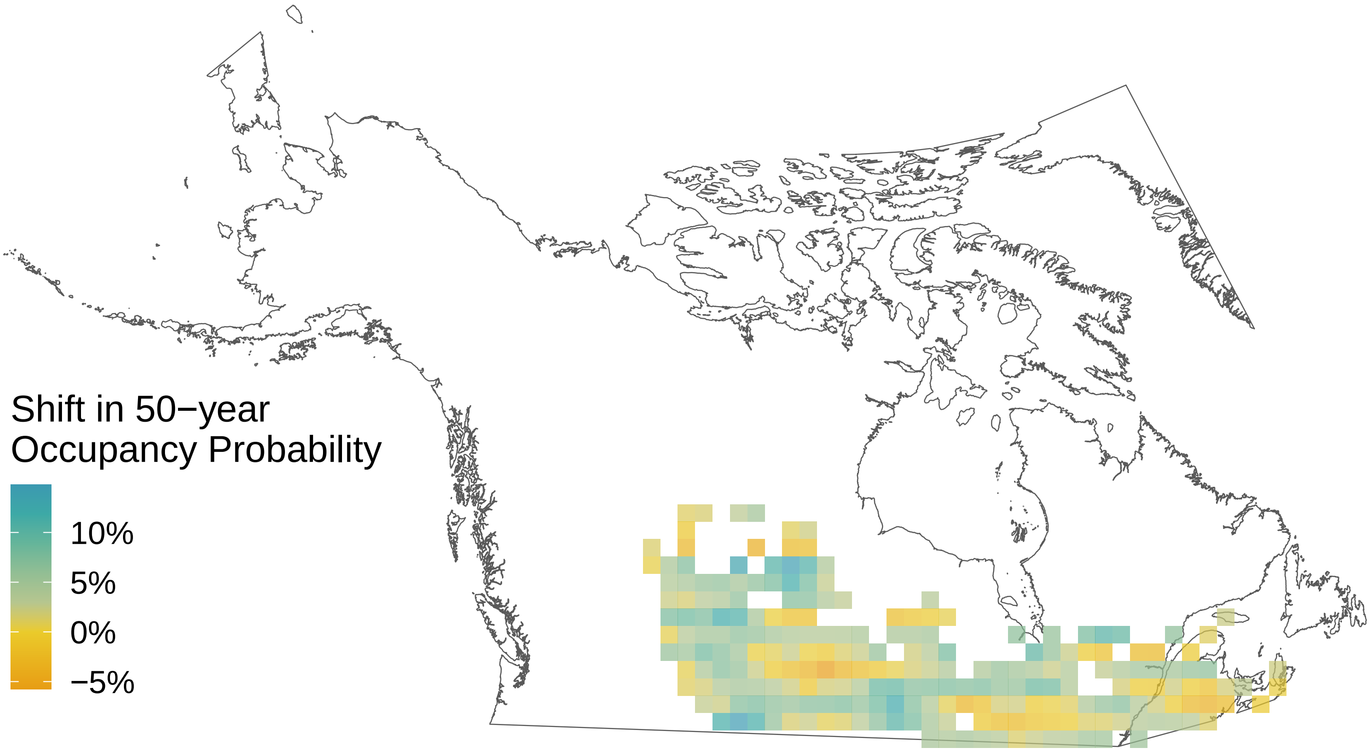

*Plebejus melissa*

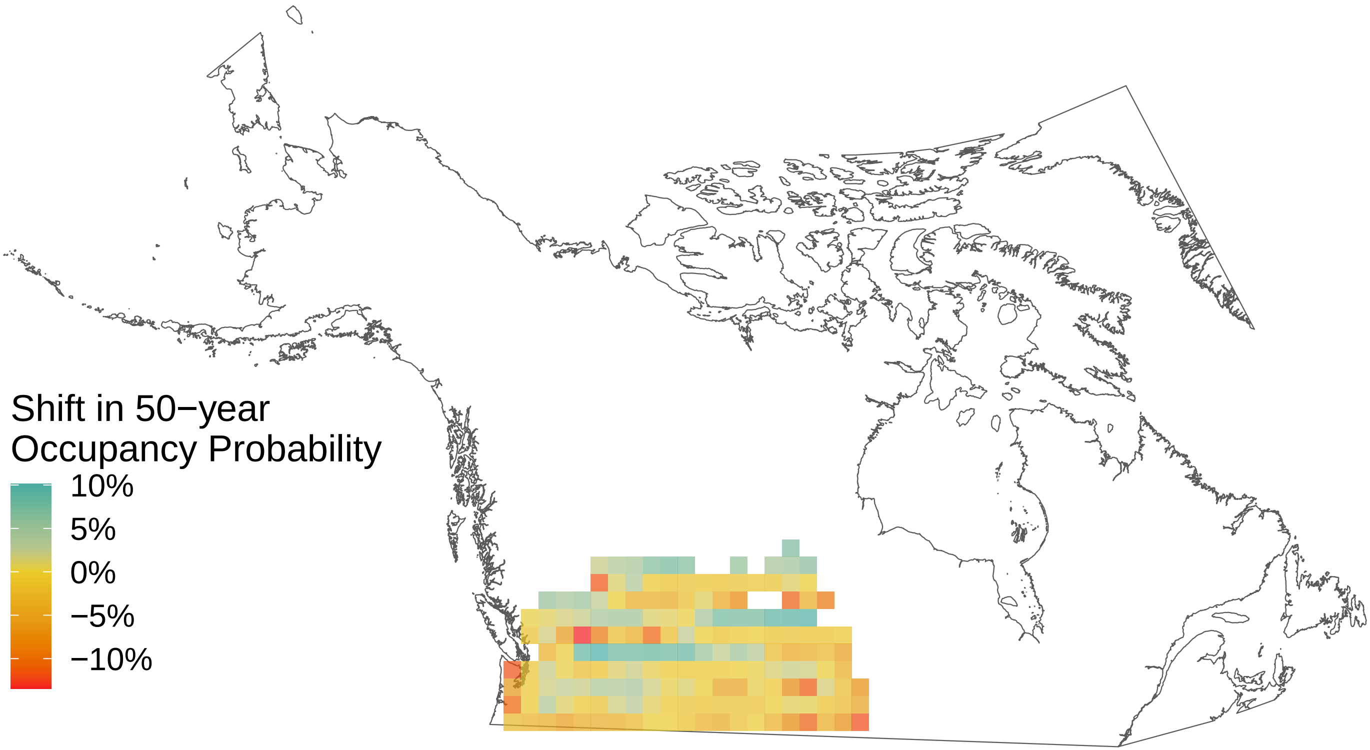

*Pieris rapae*

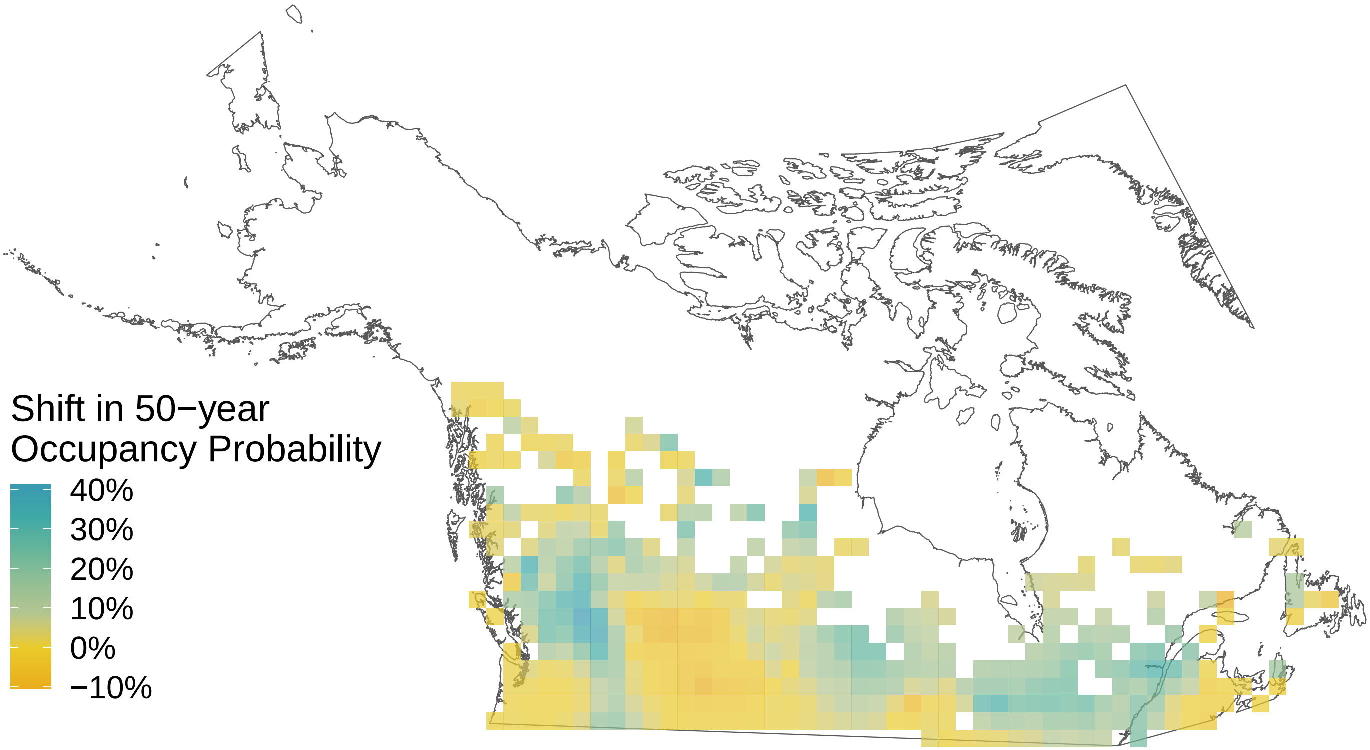

*Plebejus idas*

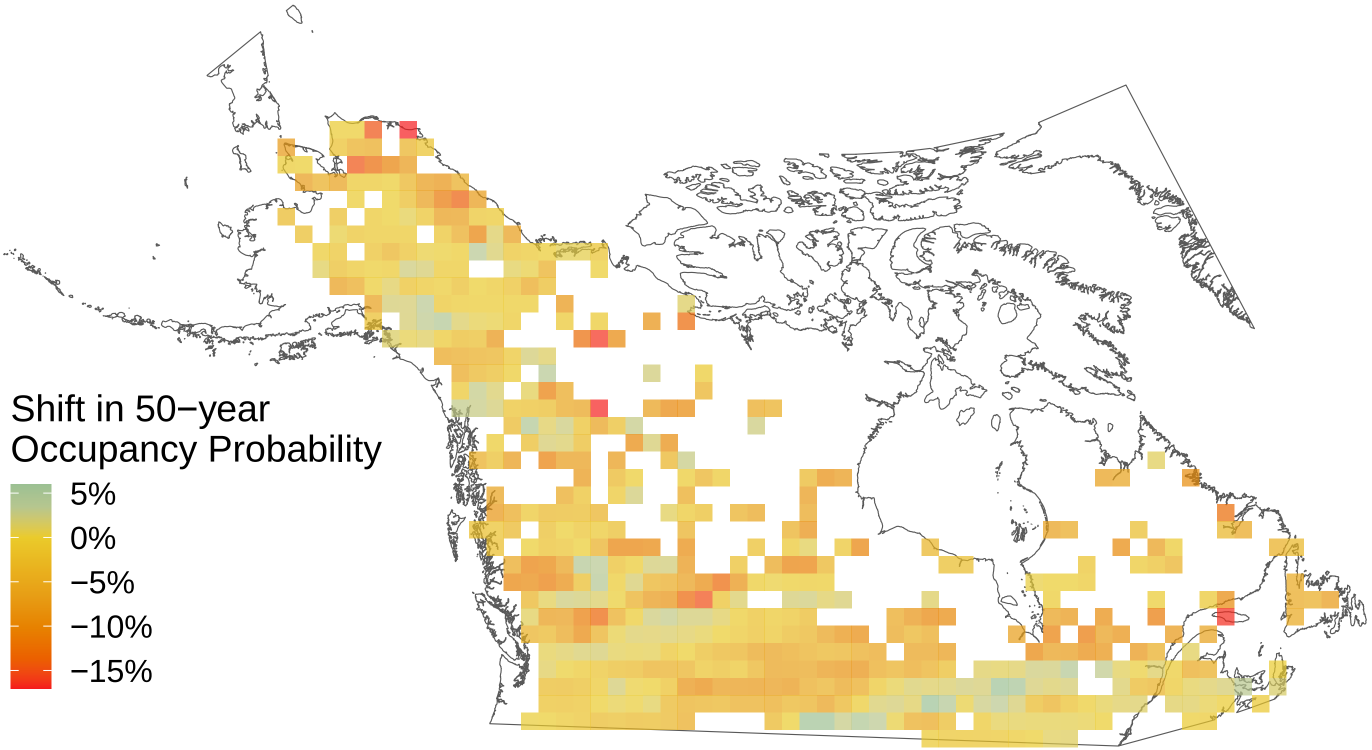

*Pieris oleracea*

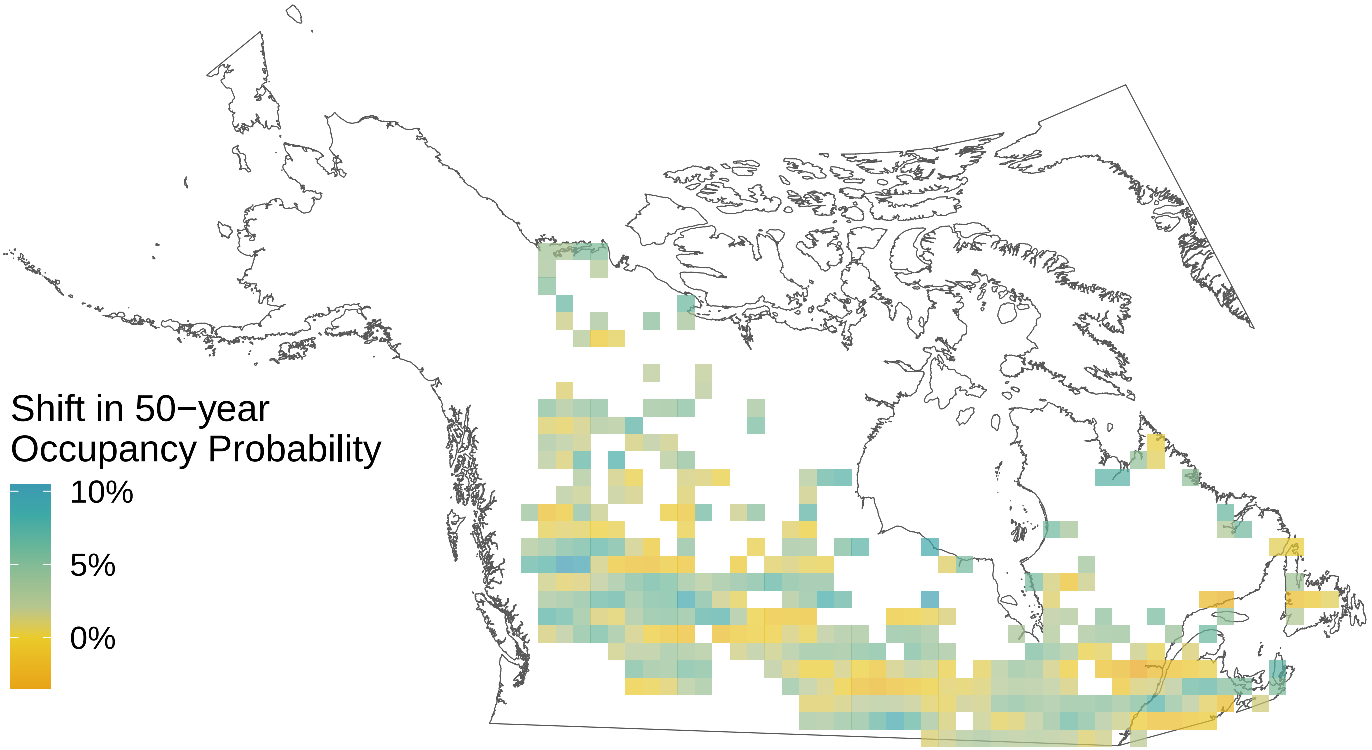

*Phyciodes tharos*

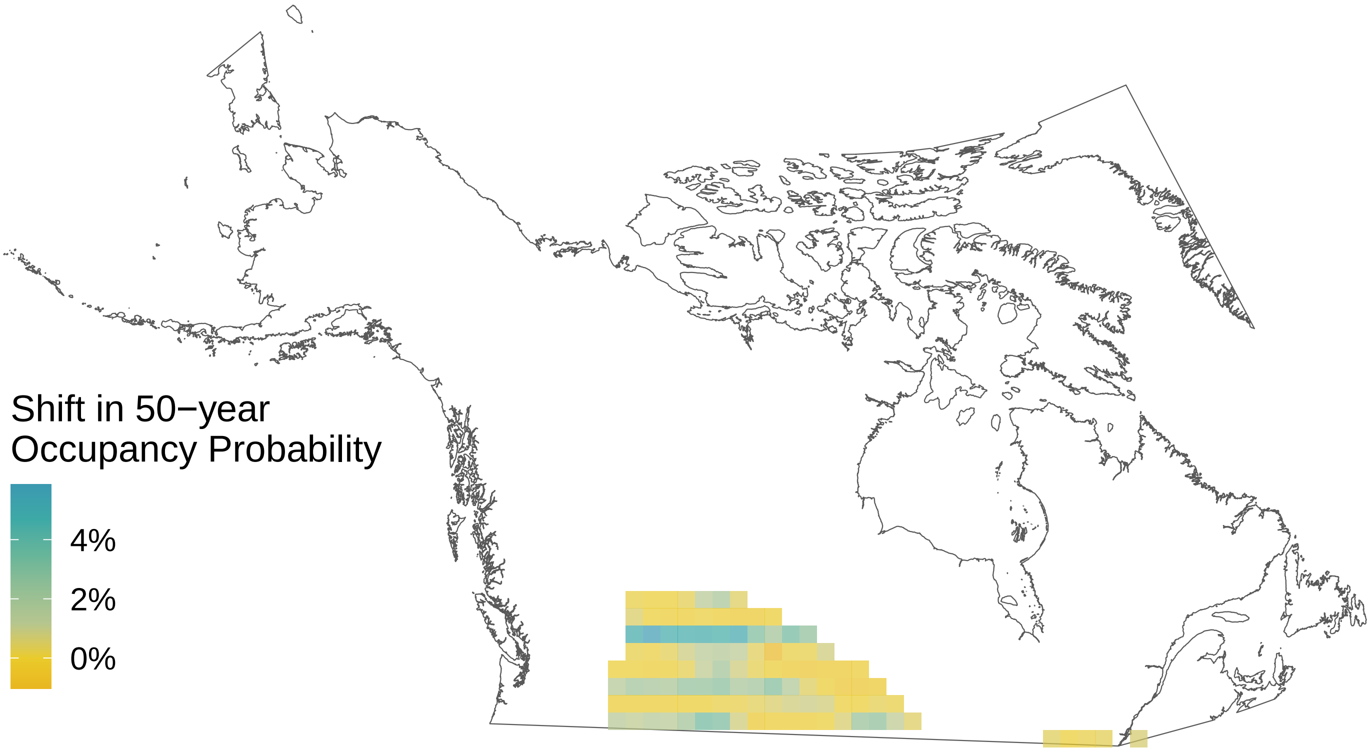

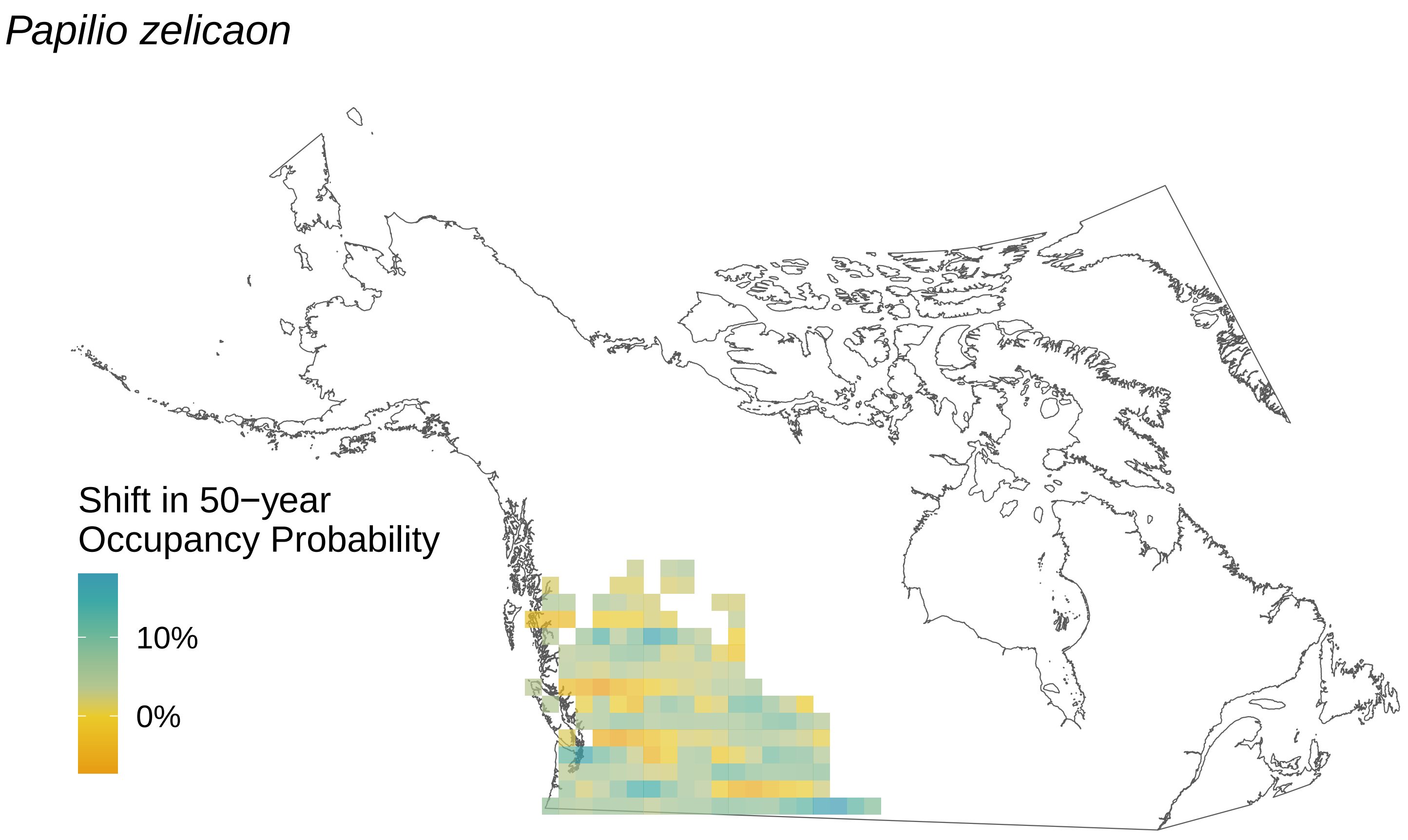
Papilio zelicaon
Shift in 50−year
Occupancy Probability
10%
0%

*Parnassius clodius*

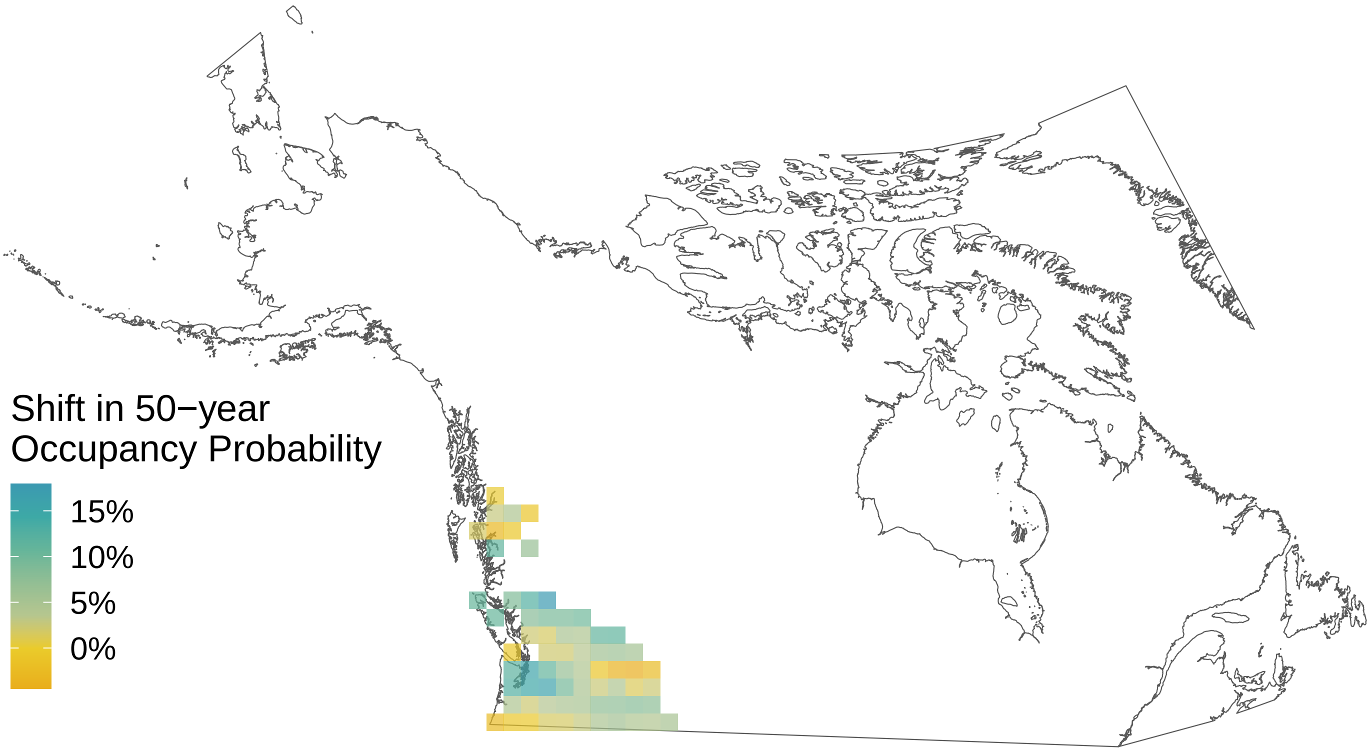

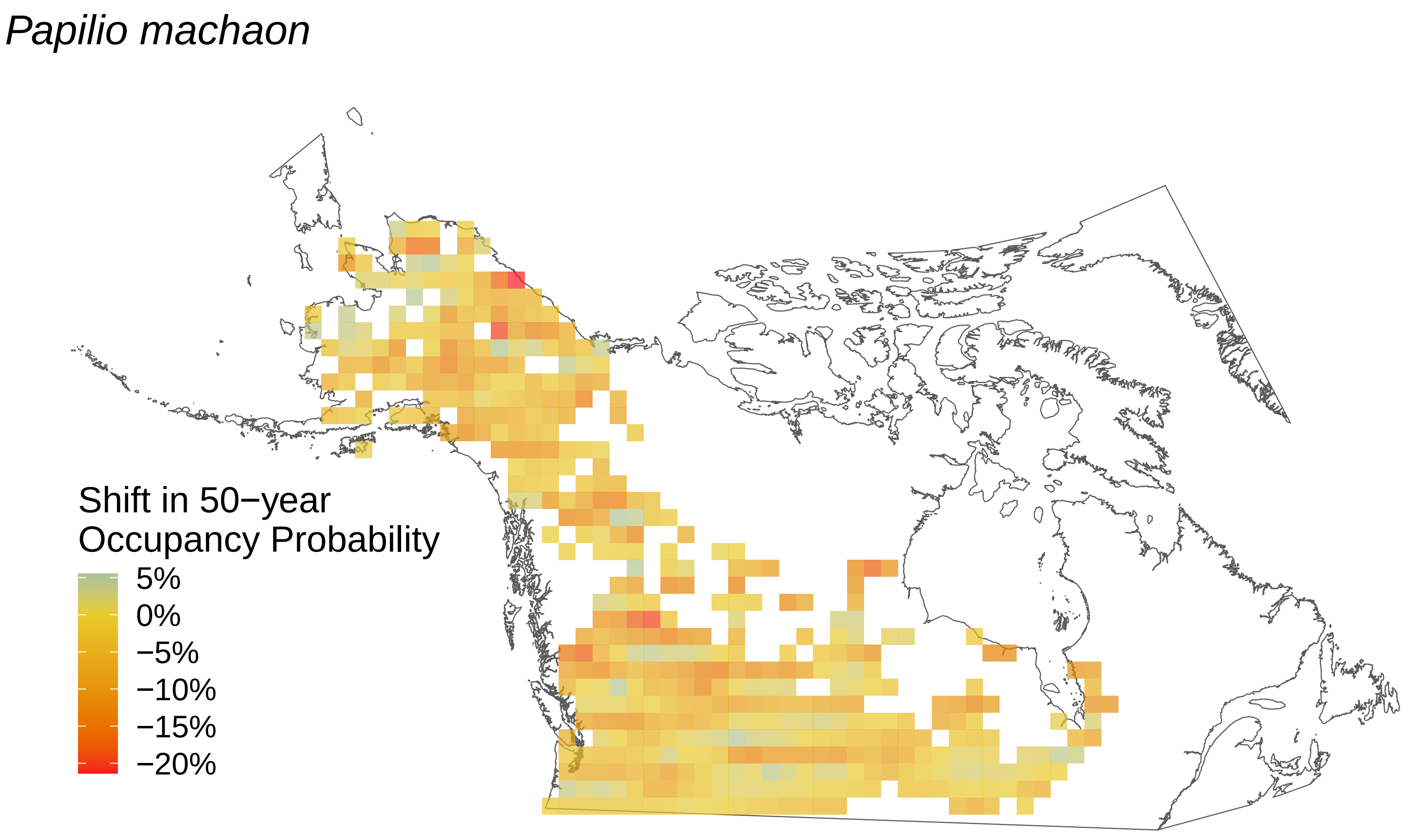
Papilio machaon
Shift in 50−year
Occupancy Probability
5%
0%
−5%
−10%
−15%
−20%

*Oeneis melissa*

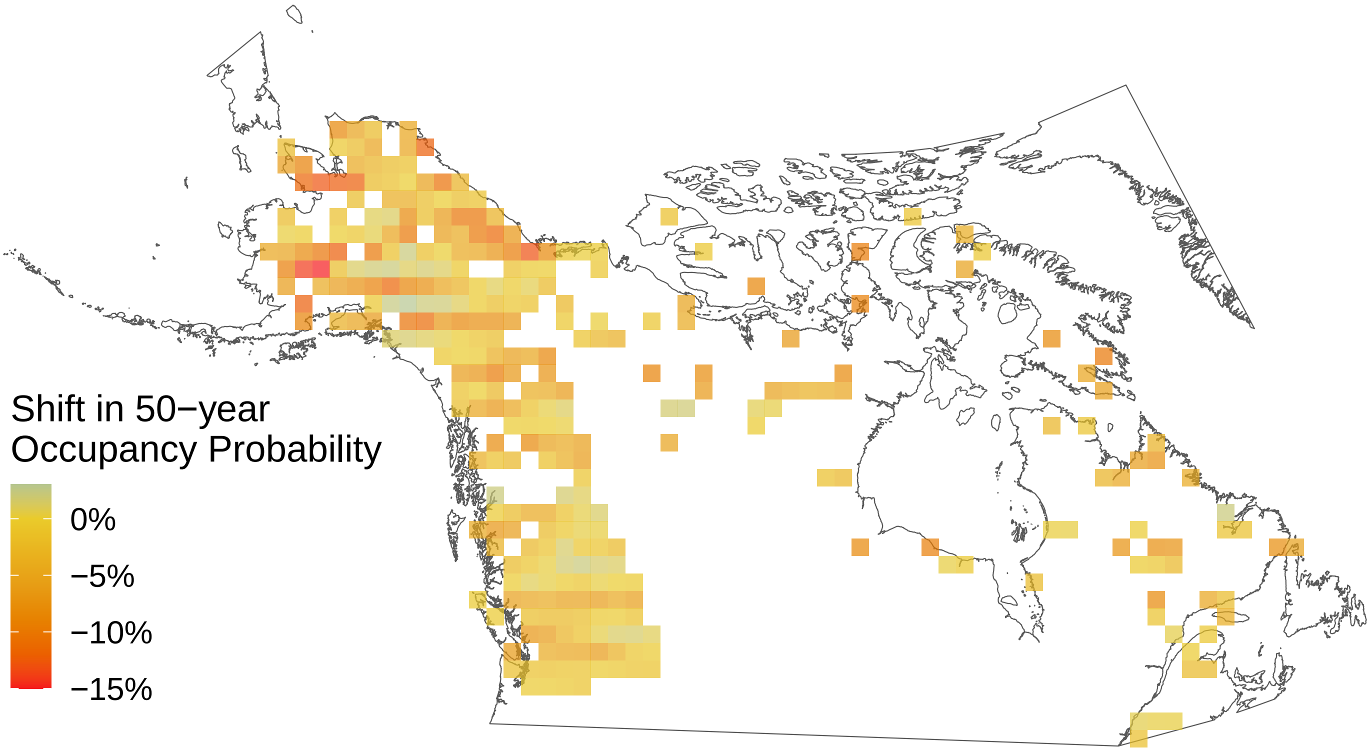

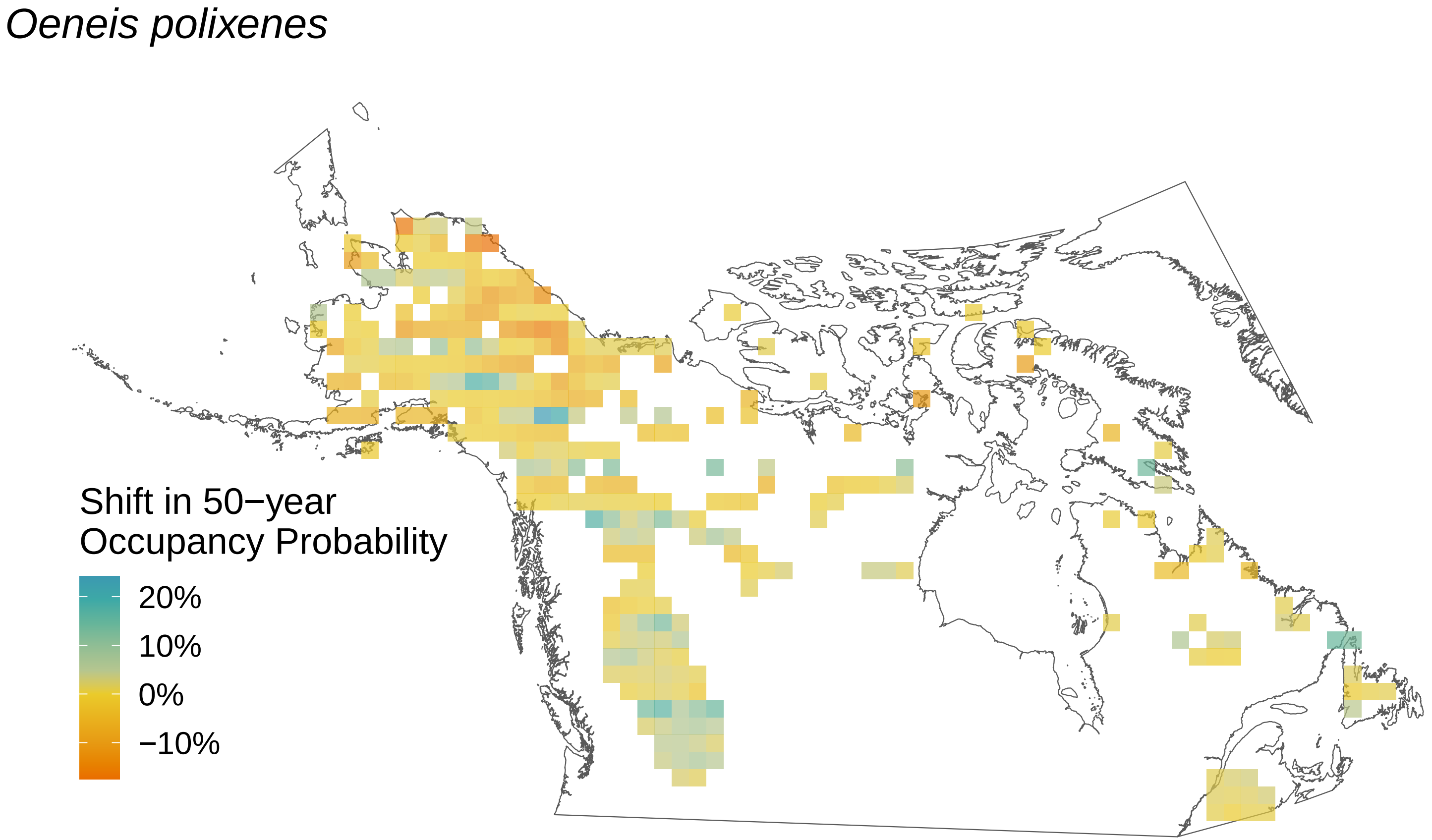

*Oeneis polixenes*
Shift in 50−year
Occupancy Probability
20%
10%
0%
−10%

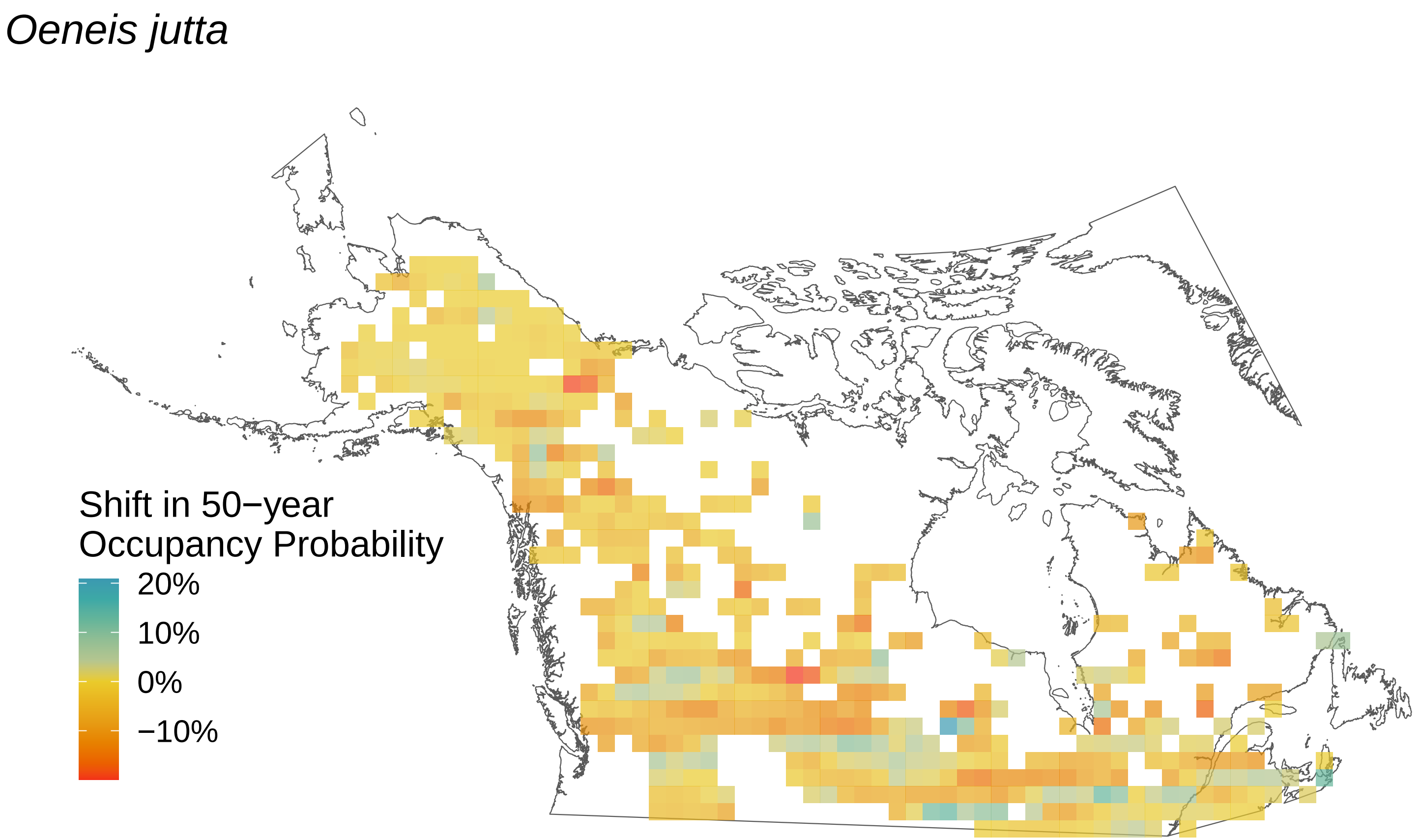

Oeneis jutta
Shift in 50−year
Occupancy Probability
20%
10%
0%
−10%

*Oeneis chryxus*

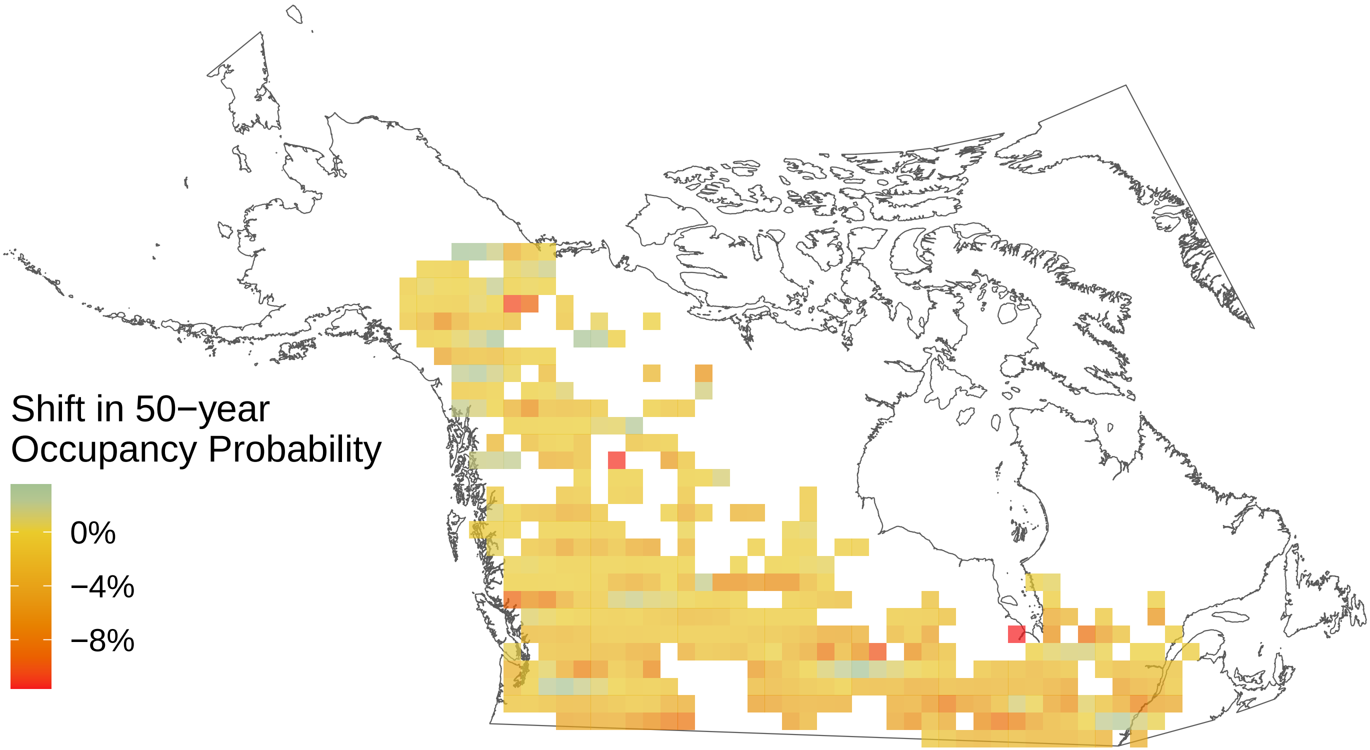

*Ochlodes sylvanoides*

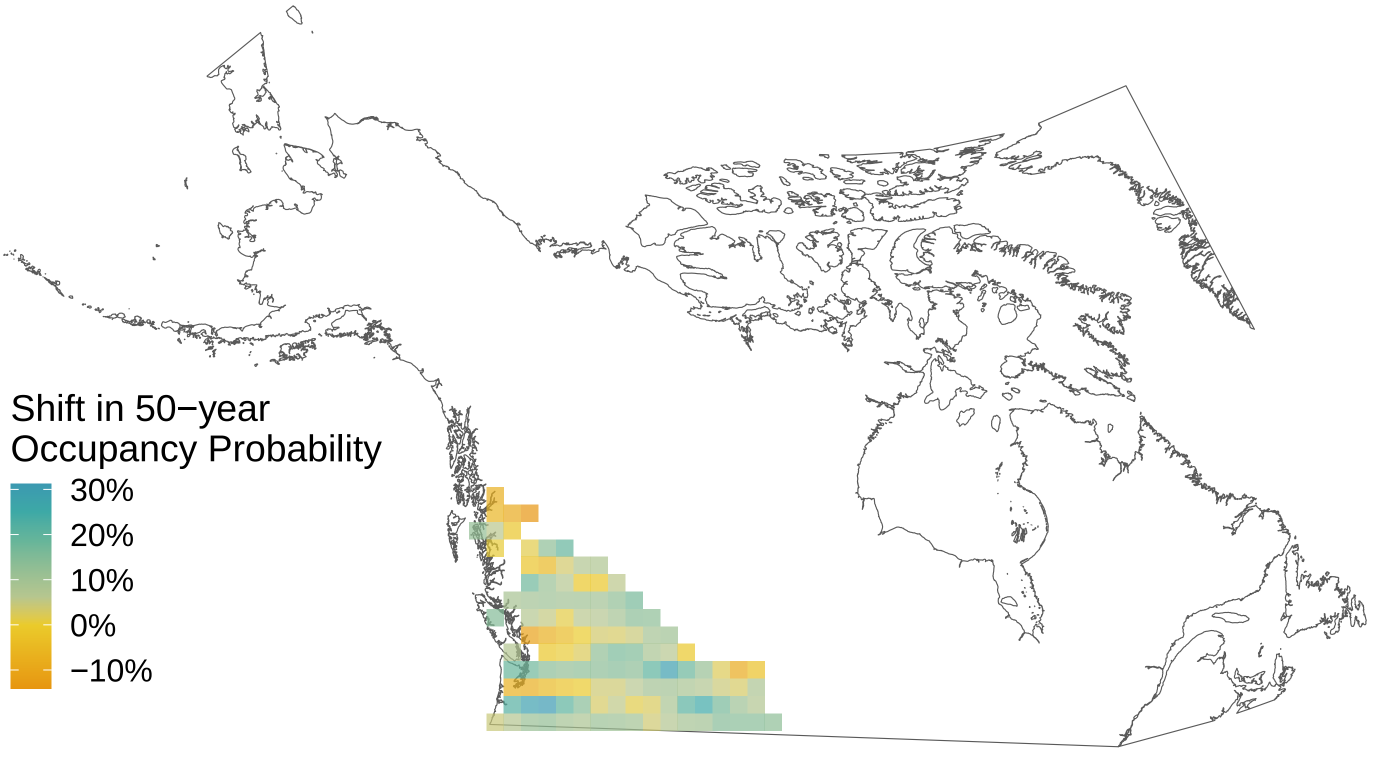

*Oeneis bore*

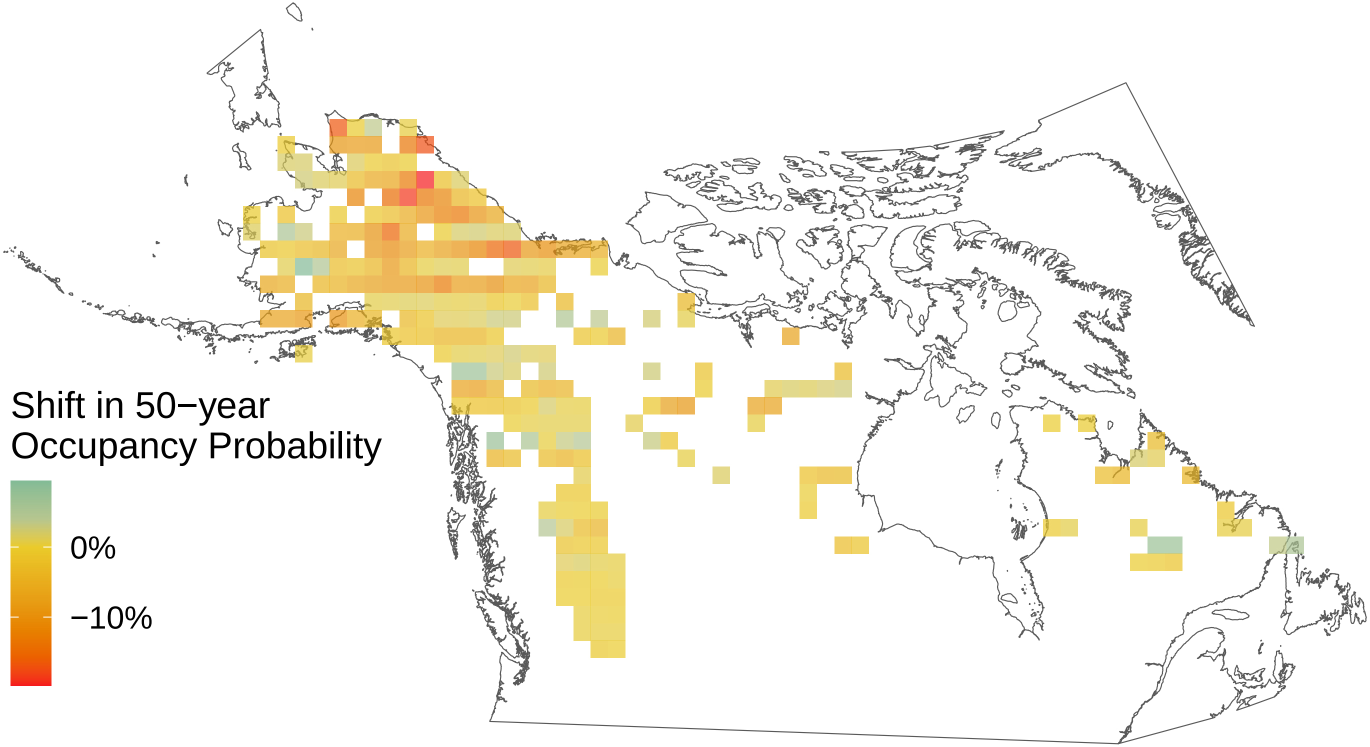

*Nymphalis l−album*

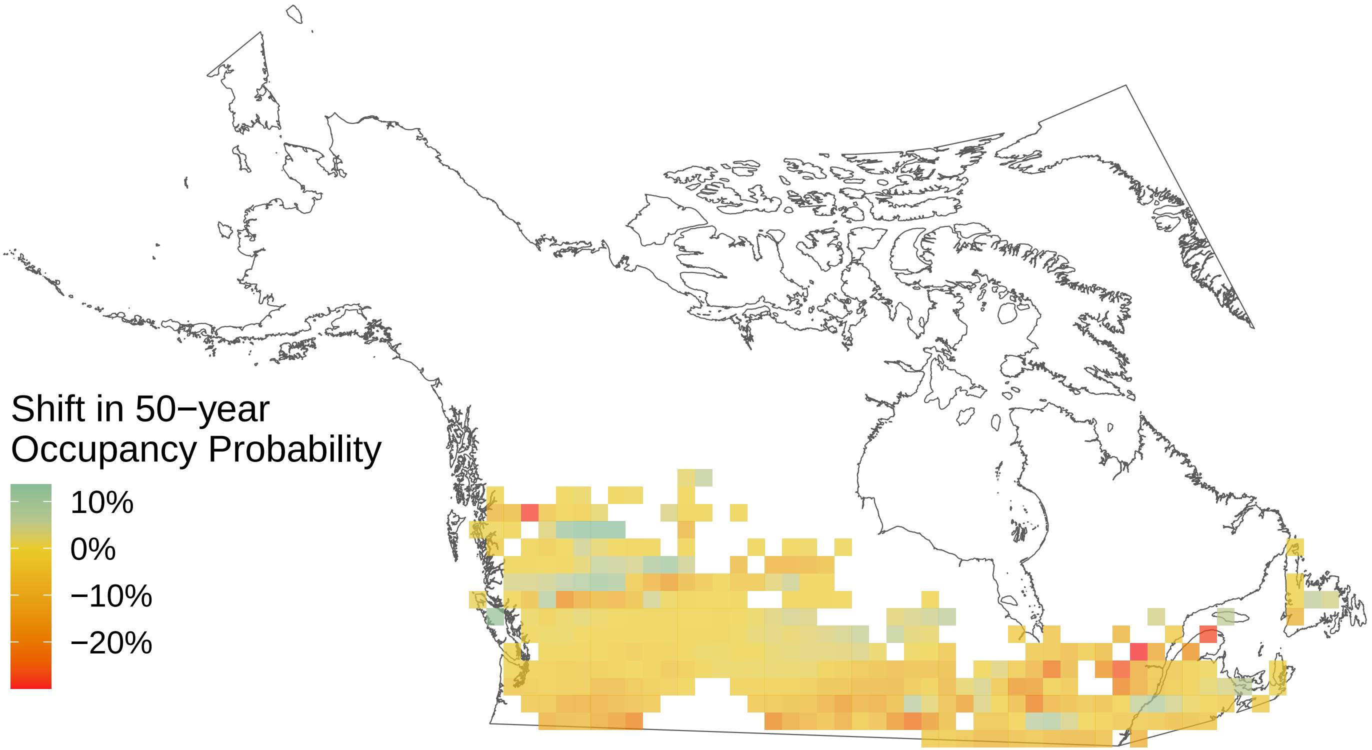

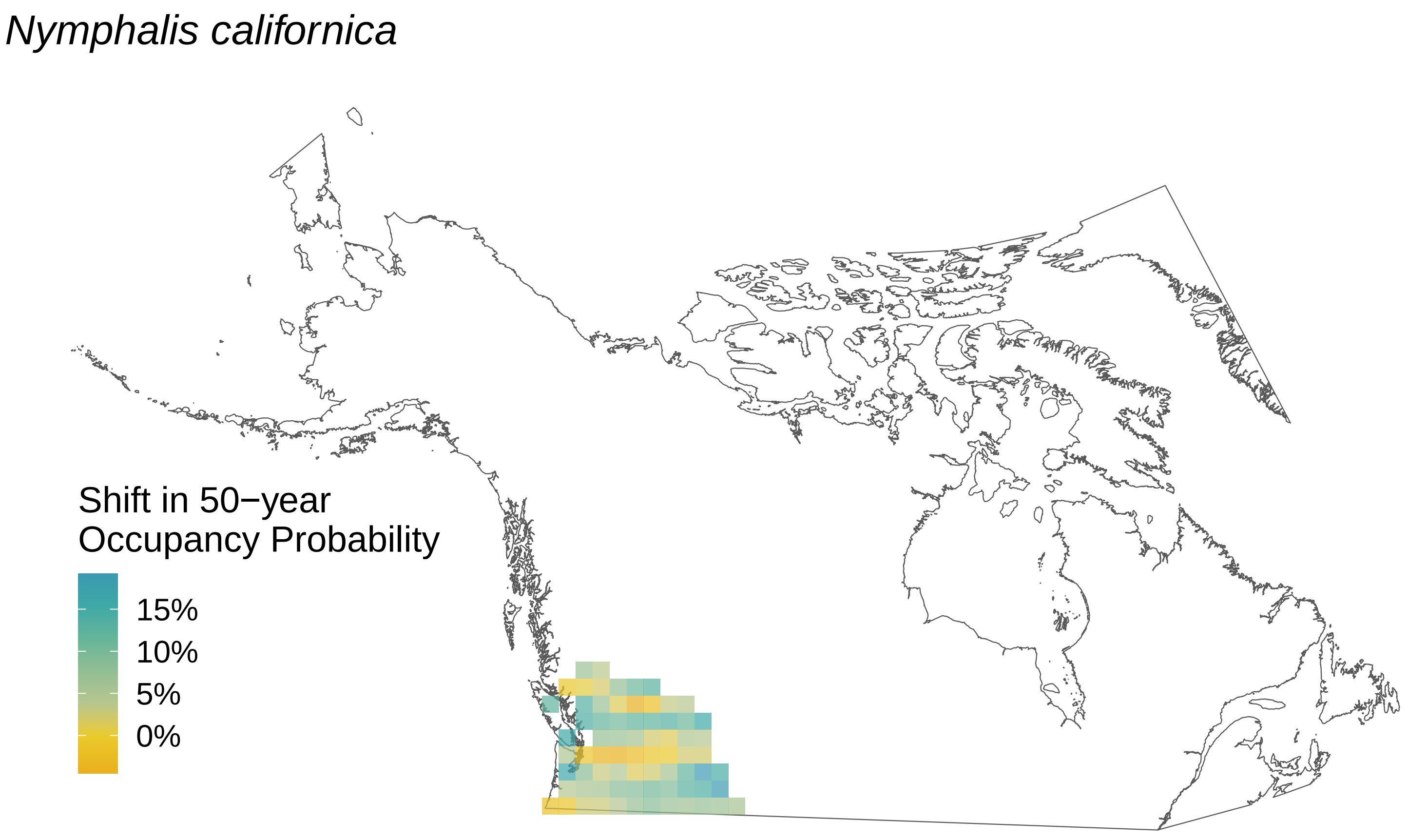
Nymphalis californica
Shift in 50−year
Occupancy Probability
15%
10%
5%
0%

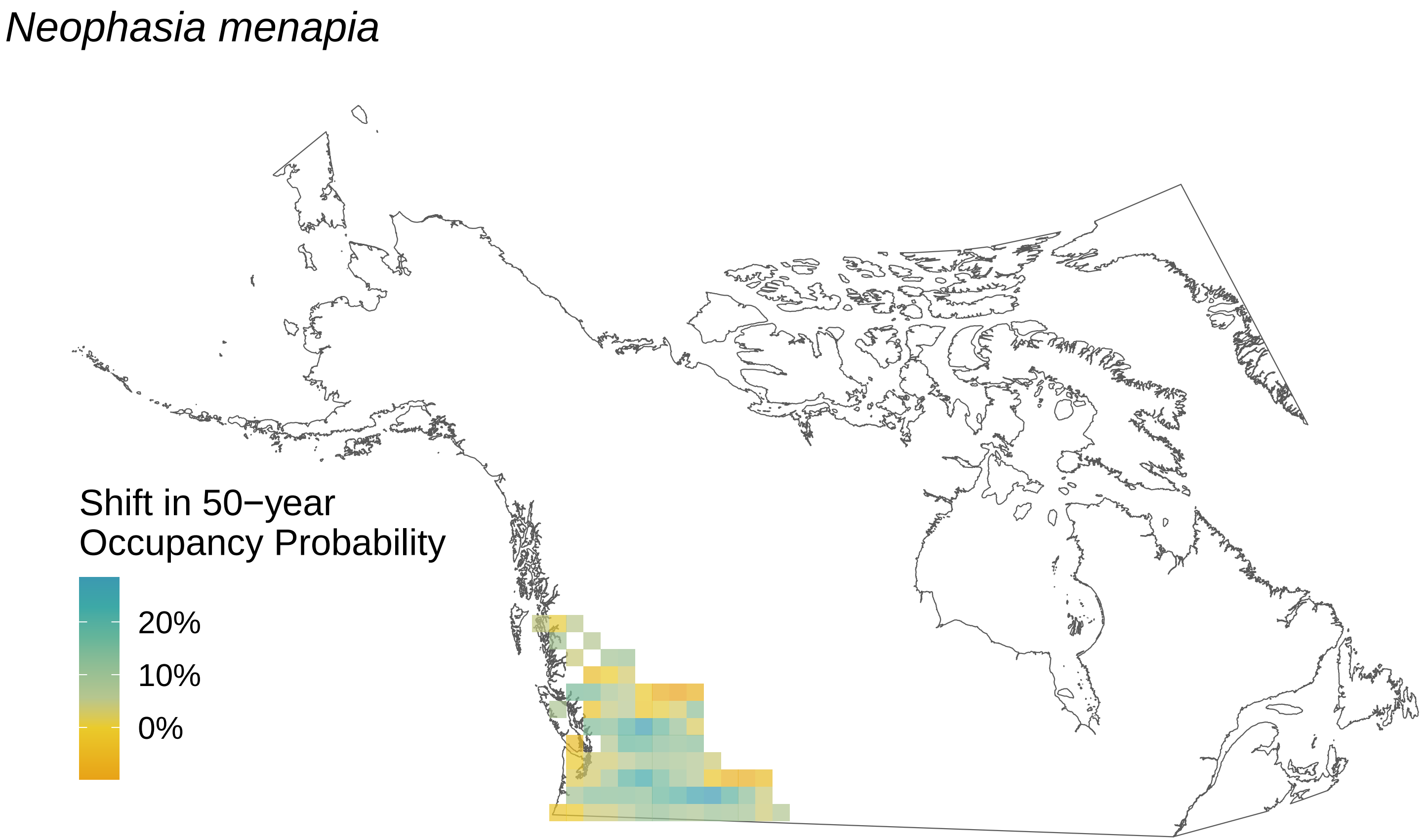

Neophasia menapia
Shift in 50−year
Occupancy Probability
20%
10%
0%

*Nymphalis antiopa*

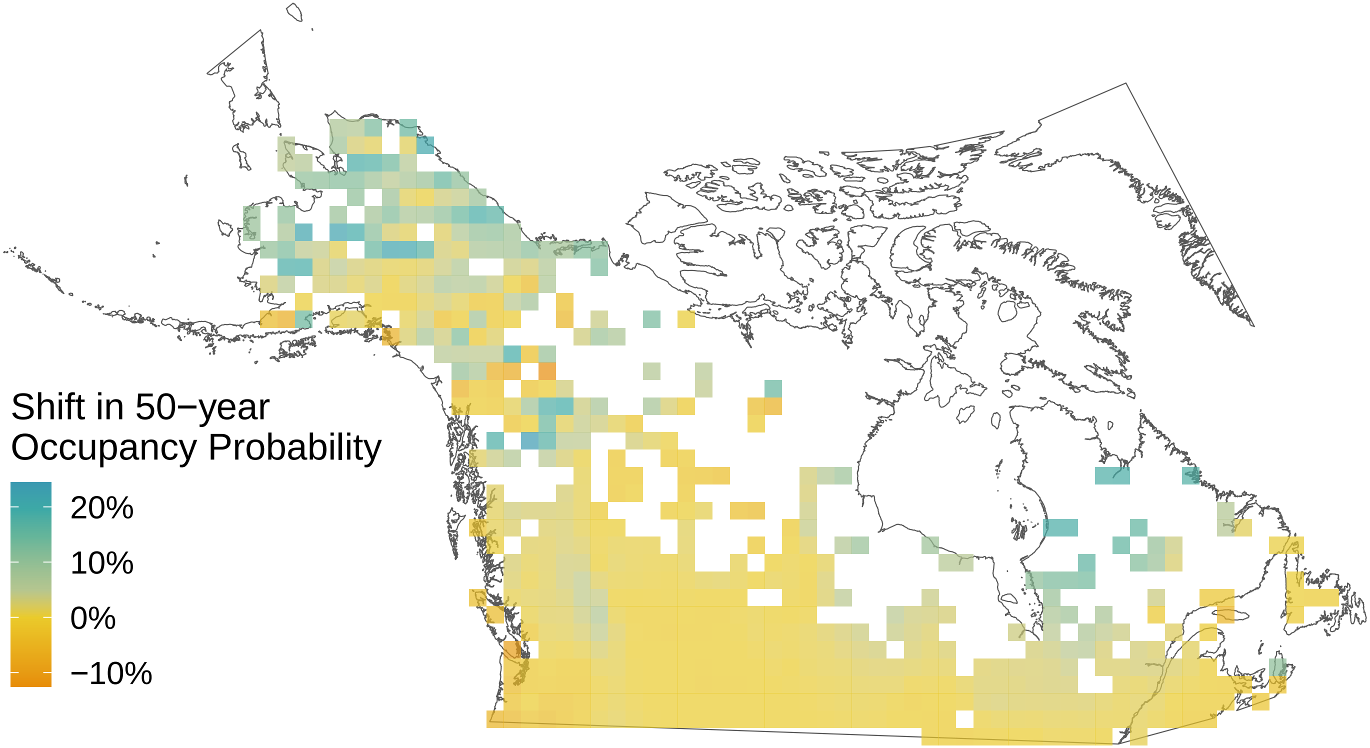

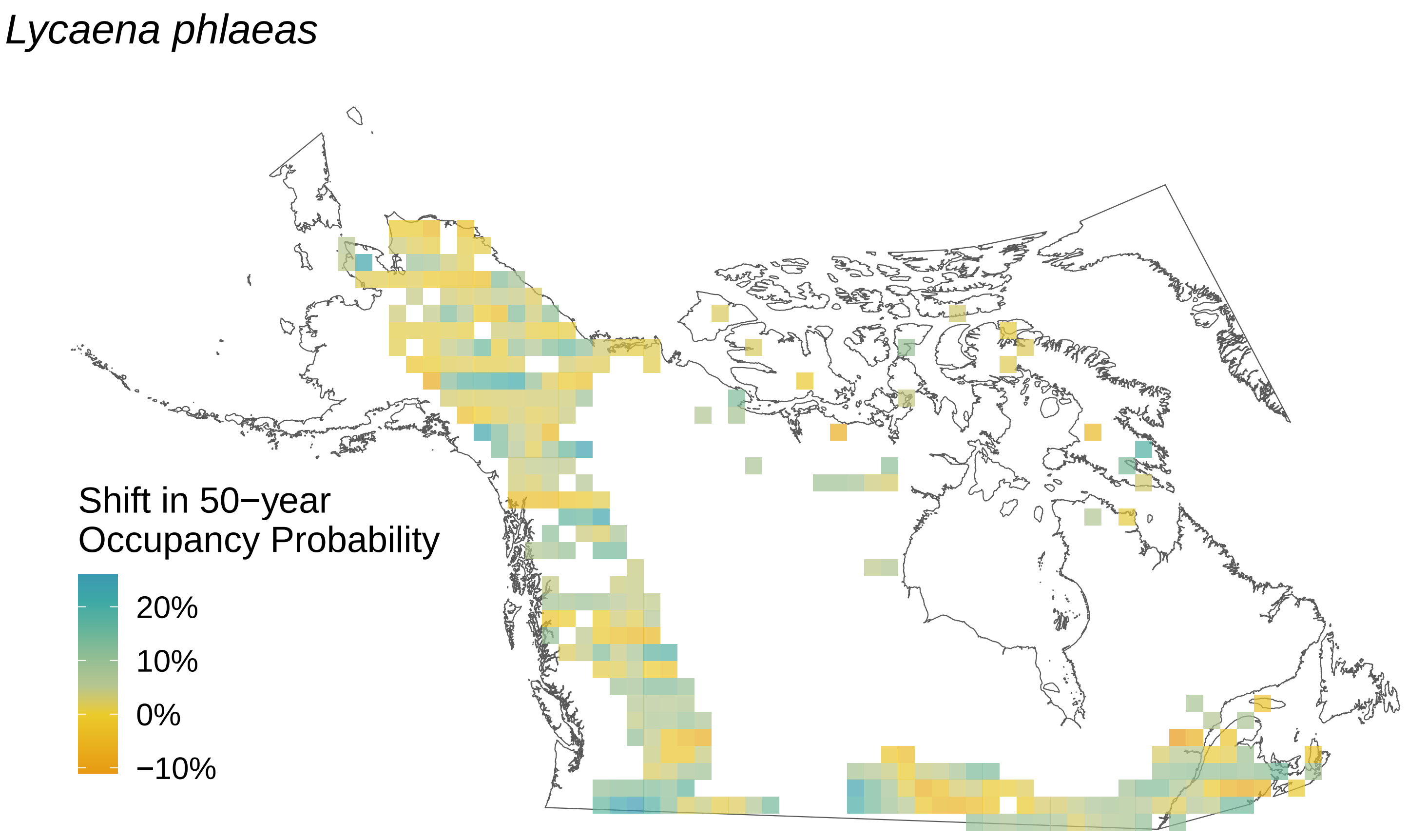
Lycaena phlaeas
Shift in 50−year
Occupancy Probability
20%
10%
0%
−10%

*Lycaena dorcas*

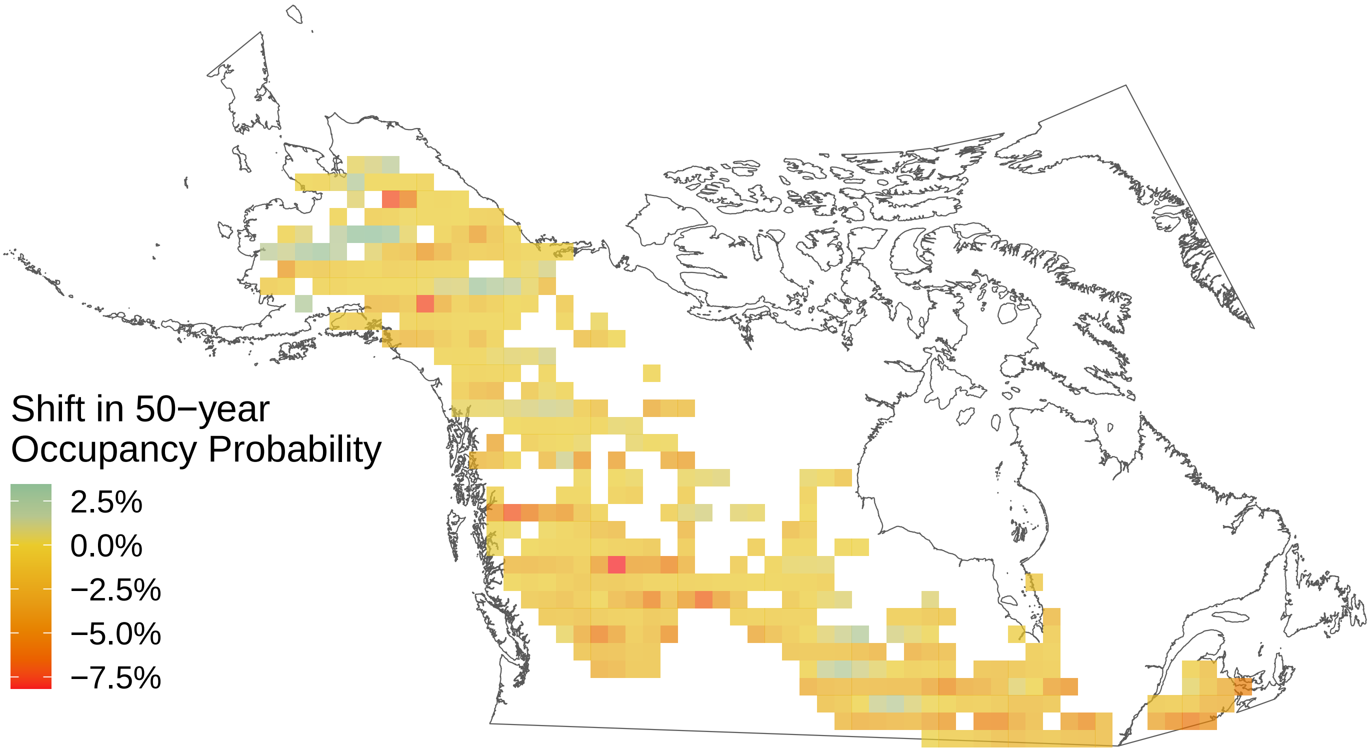

*Limenitis arthemis*

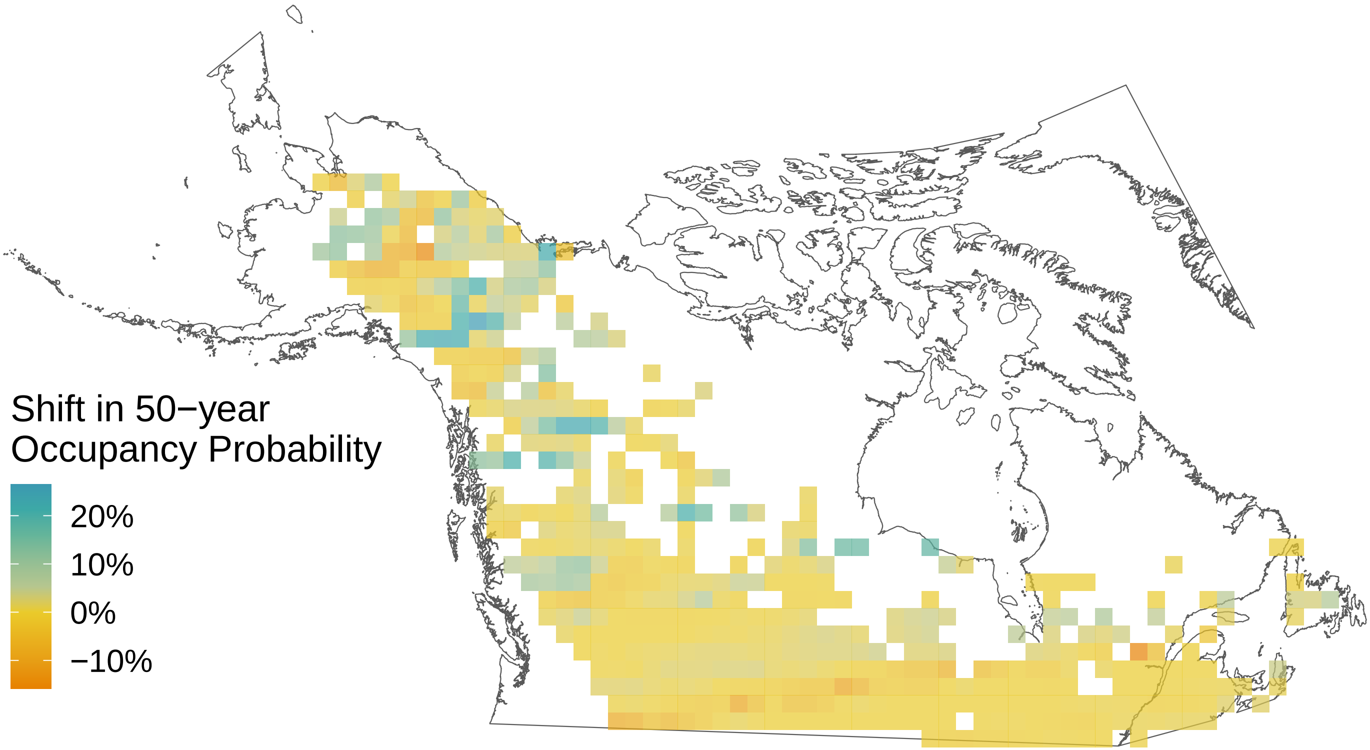

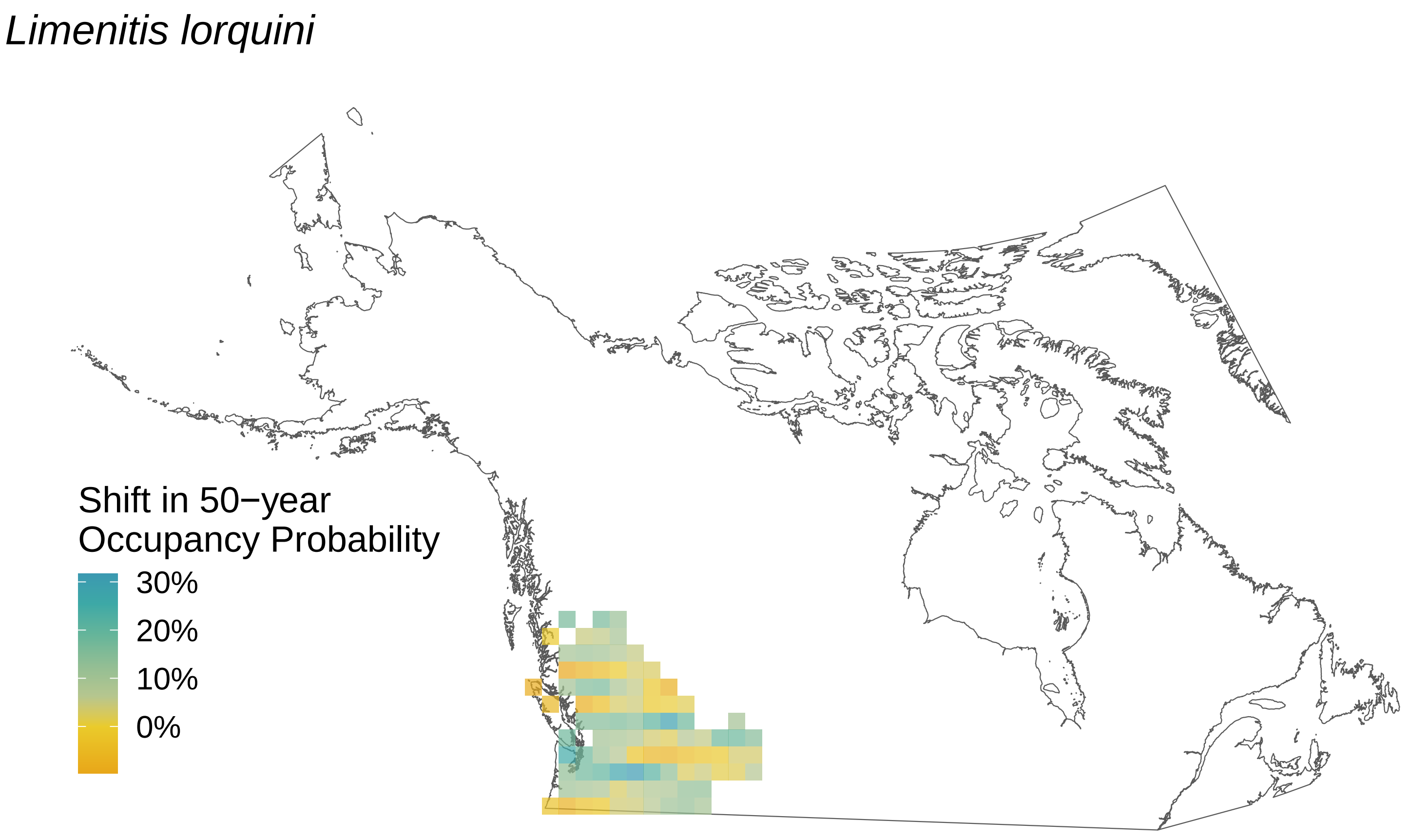
*Limenitis lorquini*
Shift in 50−year
Occupancy Probability
30%
20%
10%
0%

*Limenitis archippus*

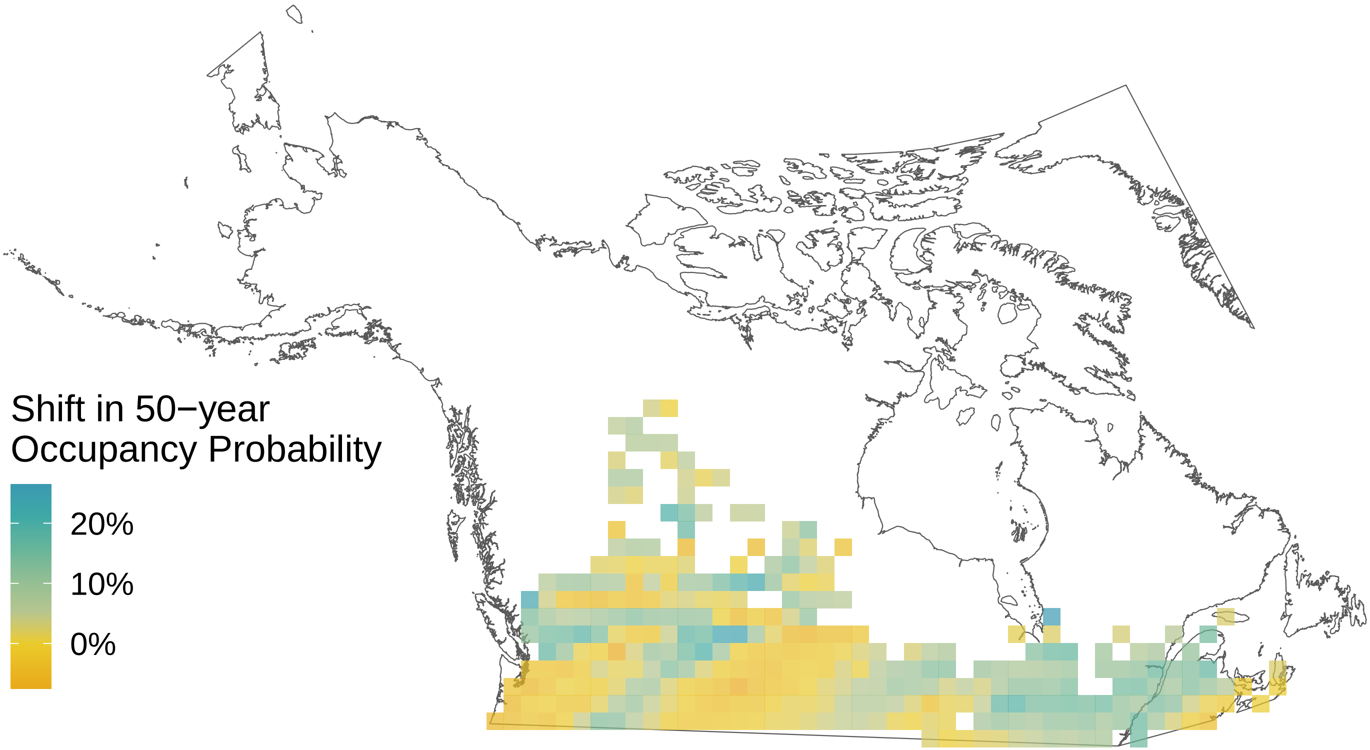

*Icaricia saepiolus*

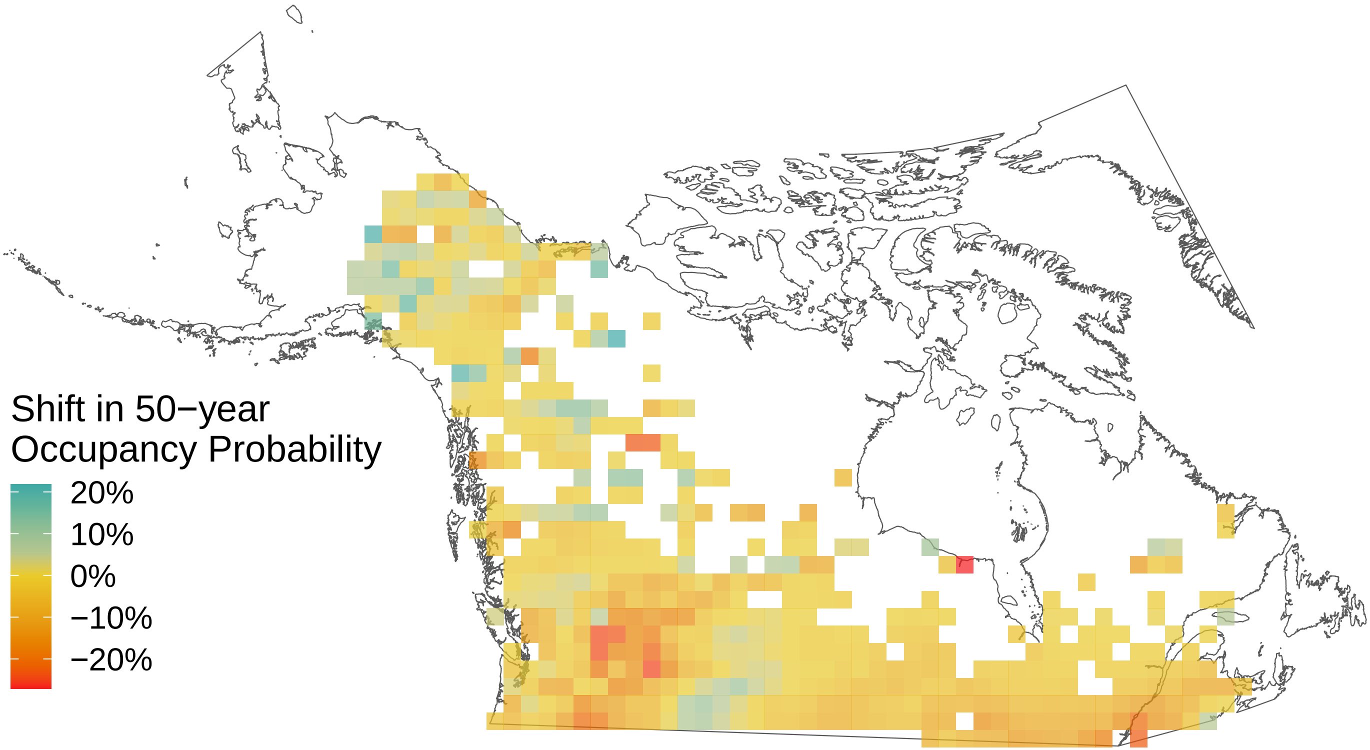

*Lethe eurydice*

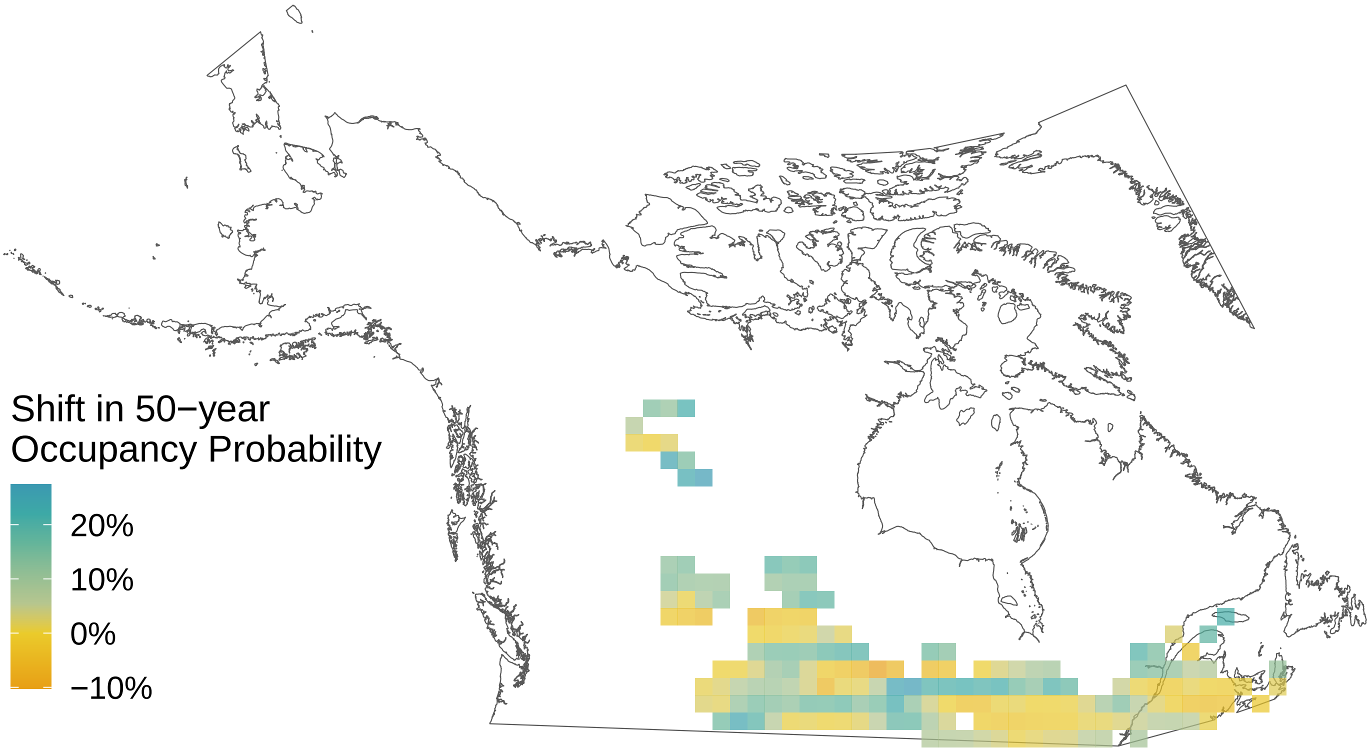

*Icaricia icarioides*

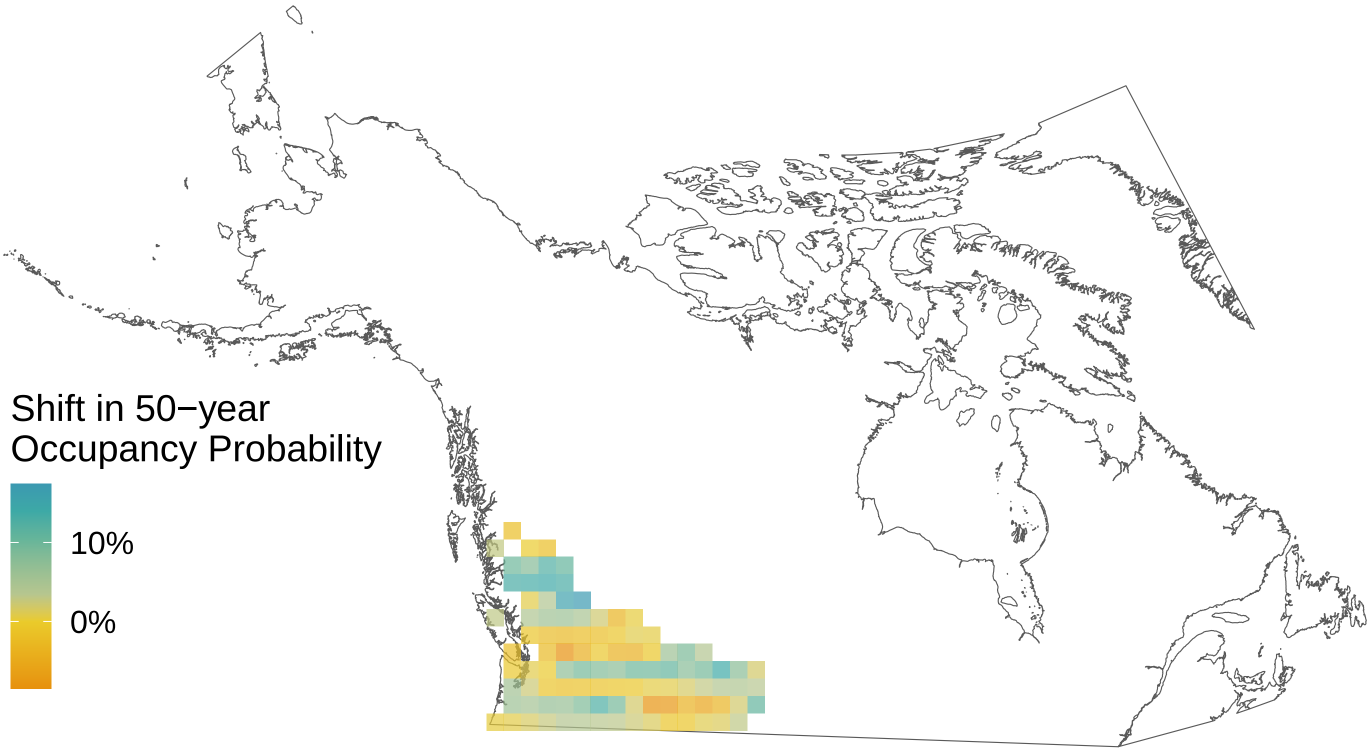

*Hesperia comma*

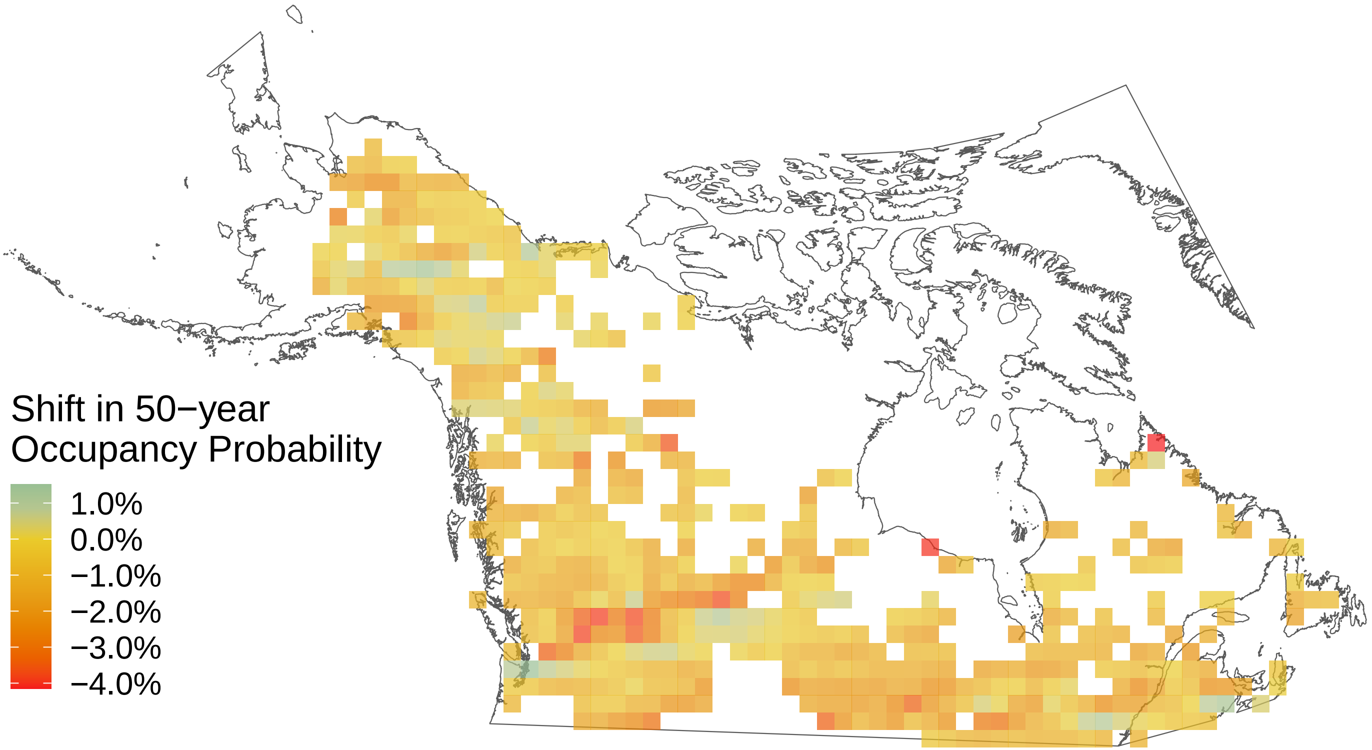

*Glaucopsyche lygdamus*

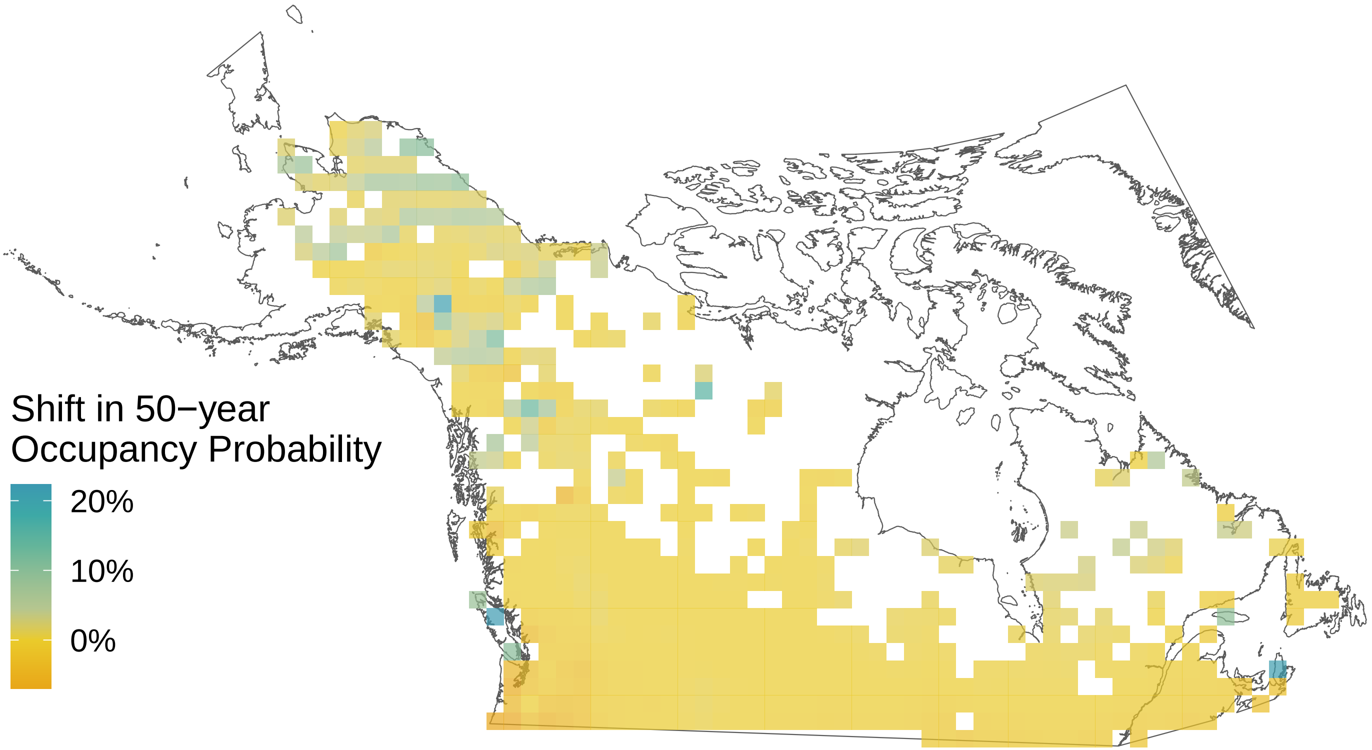

#### *Euphydryas phaeton*

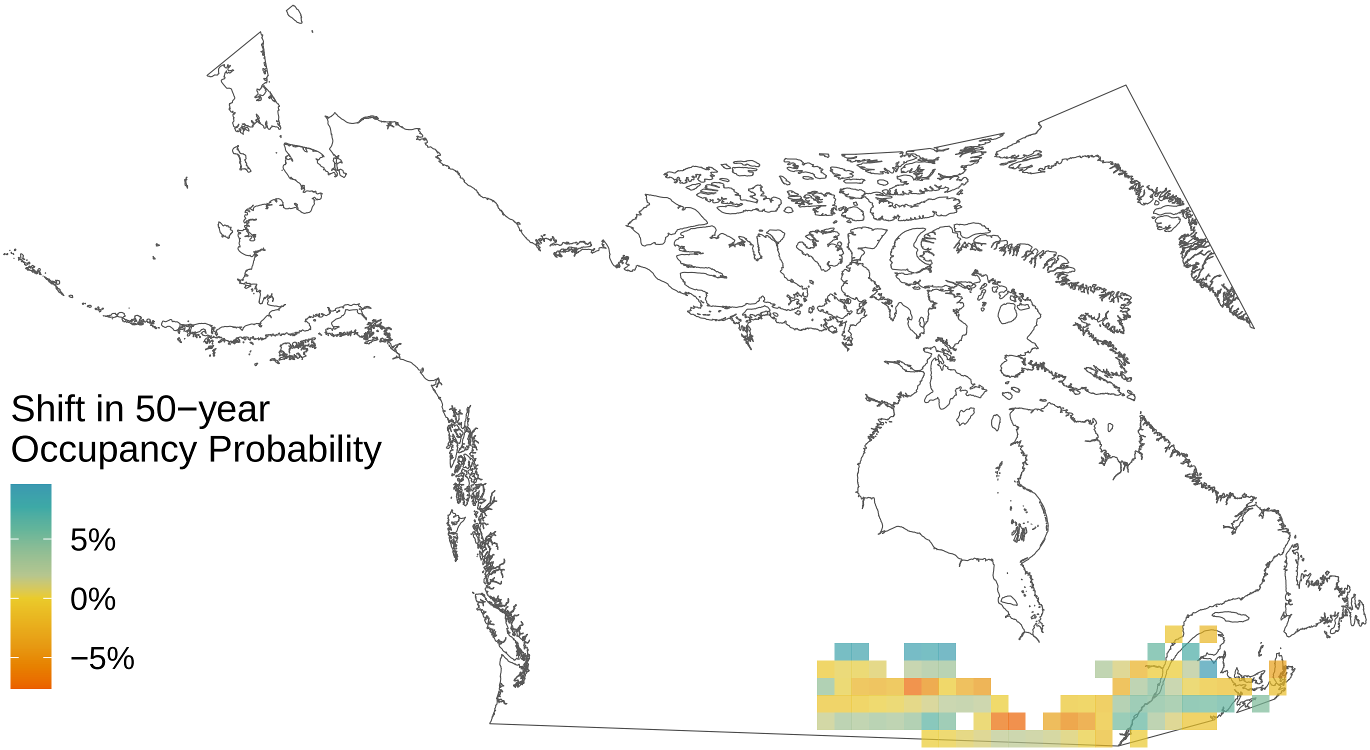

*Euphyes vestris*

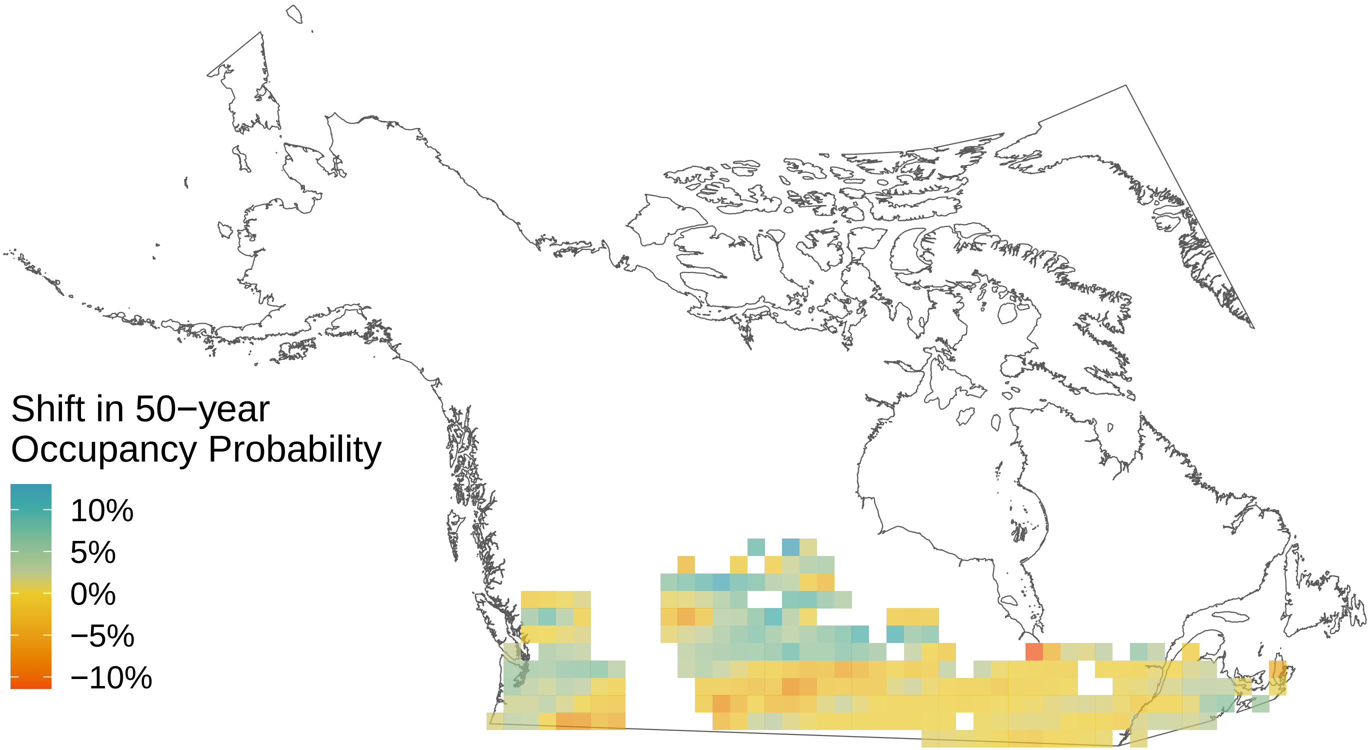

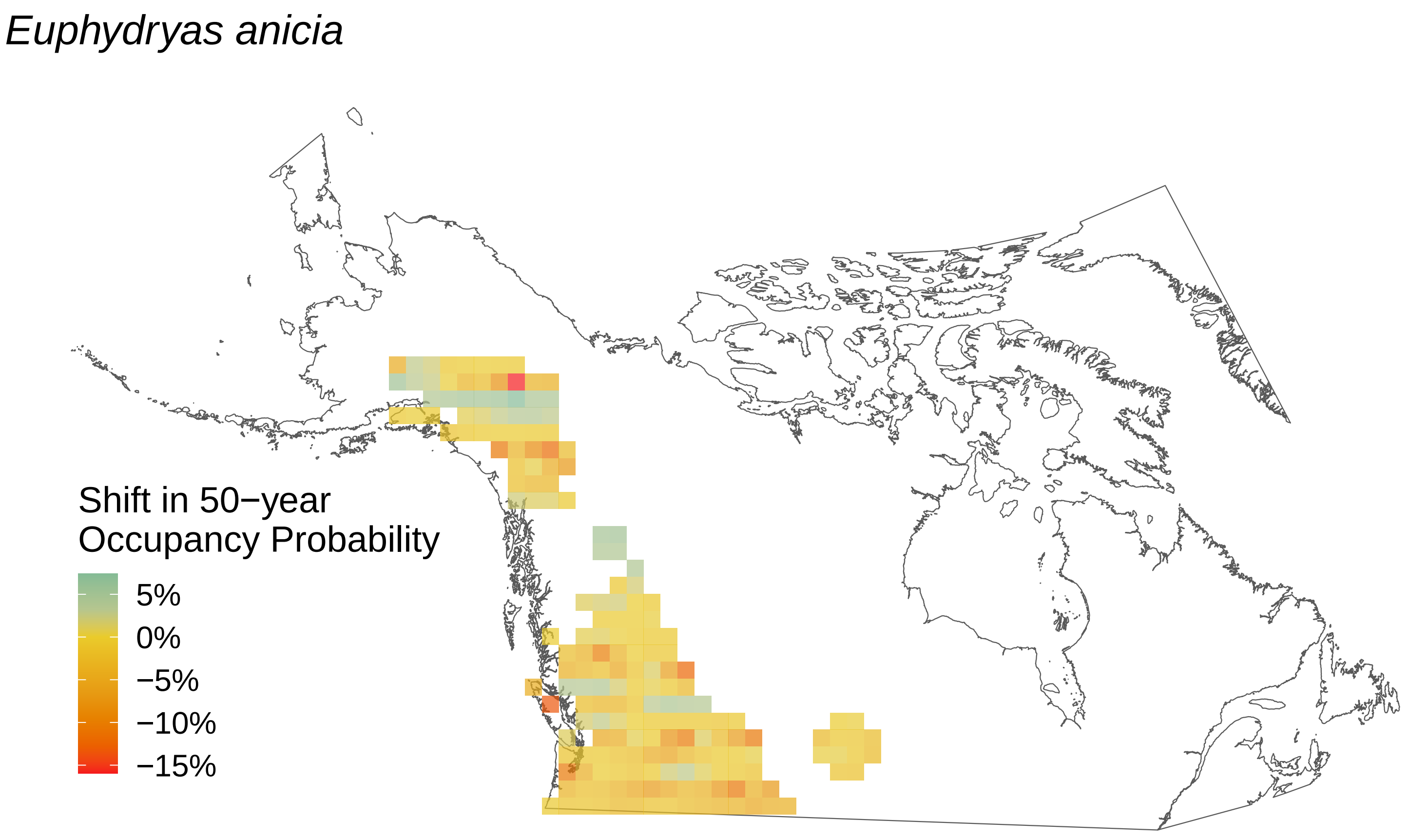
Euphydryas anicia
Shift in 50−year
Occupancy Probability
5%
0%
−5%
−10%
−15%

*Erynnis persius*

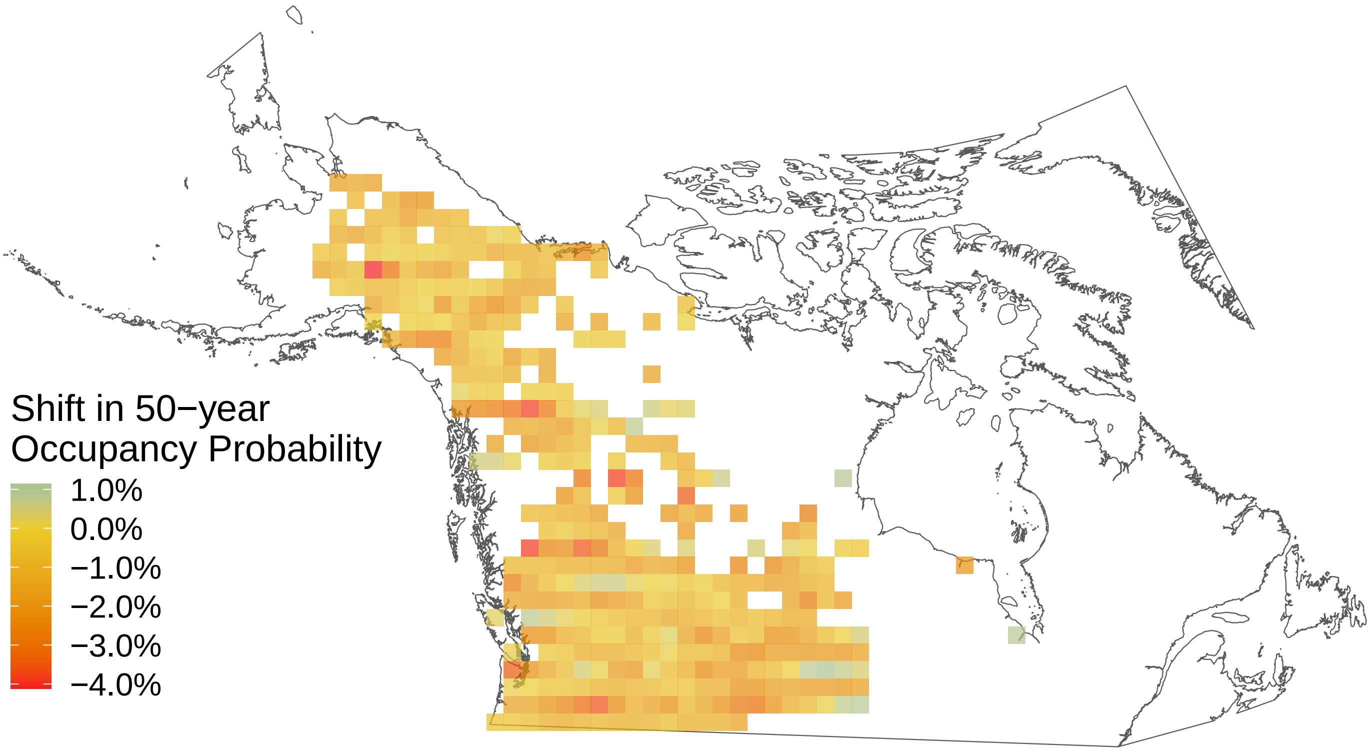

*Euchloe ausonides*

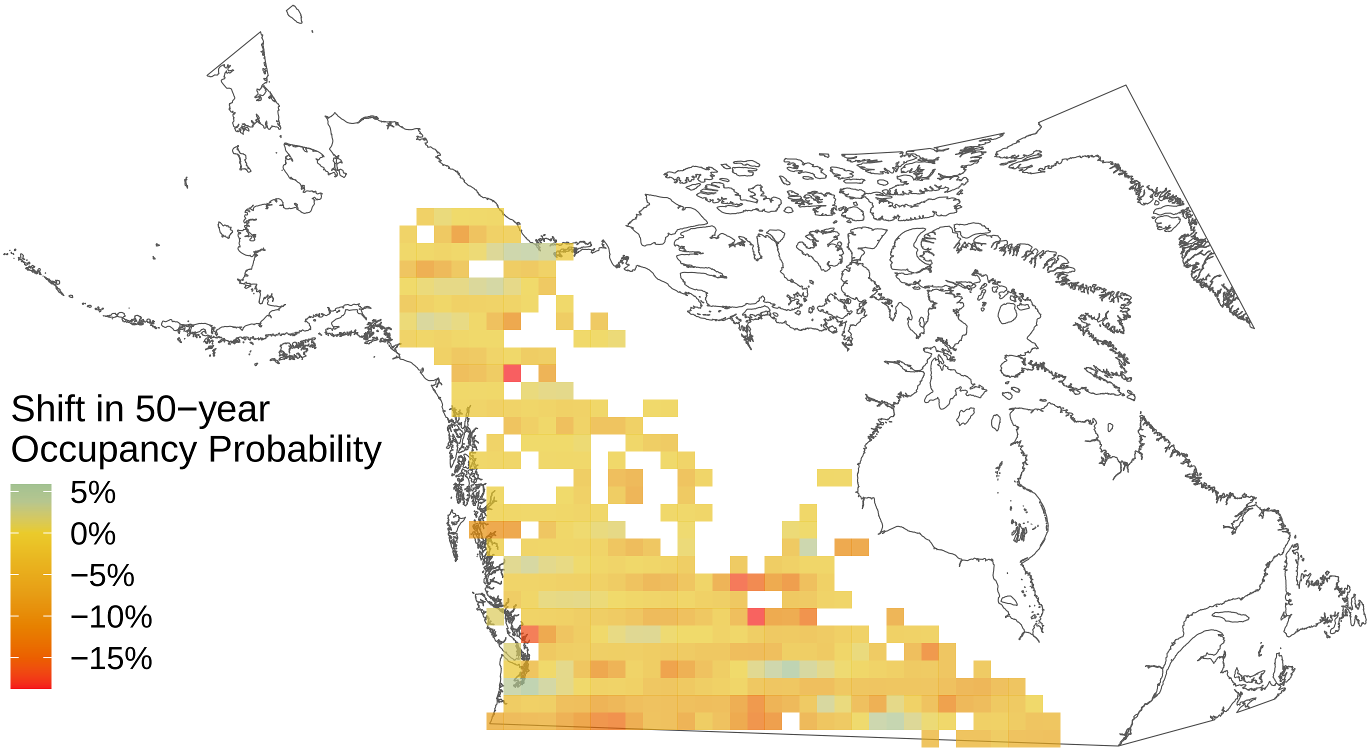

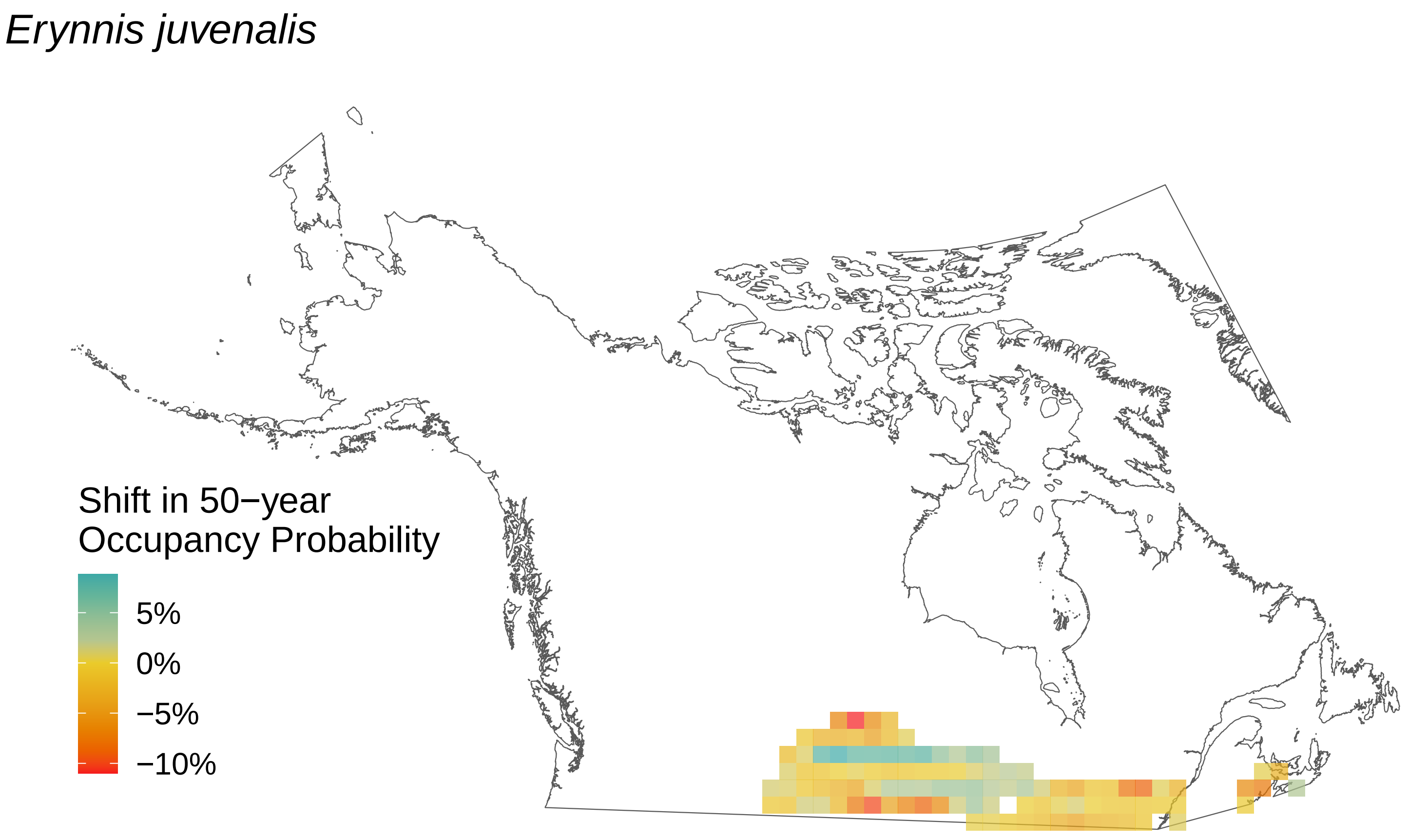

Erynnis juvenalis
Shift in 50−year Occupancy Probability
5%
0%
−5%
−10%

*Erynnis icelus*

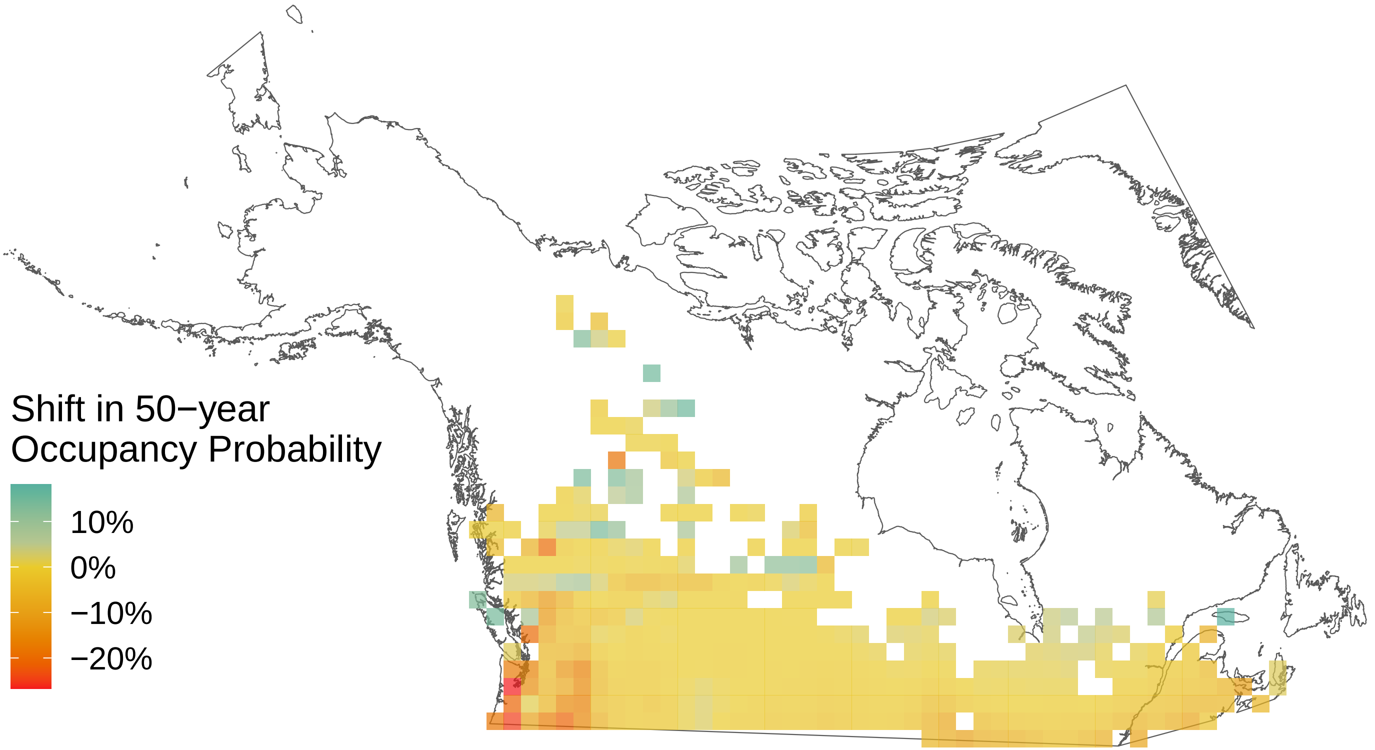

*Erebia rossii*

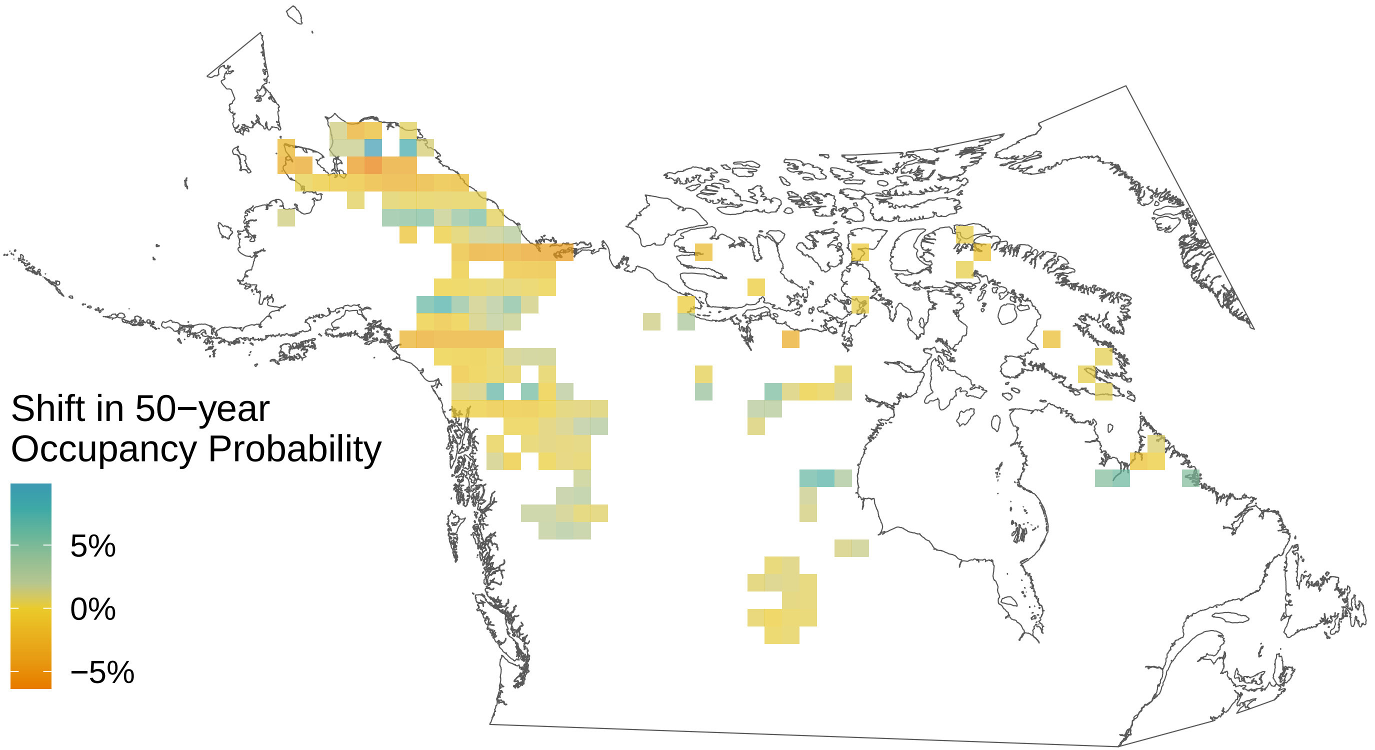

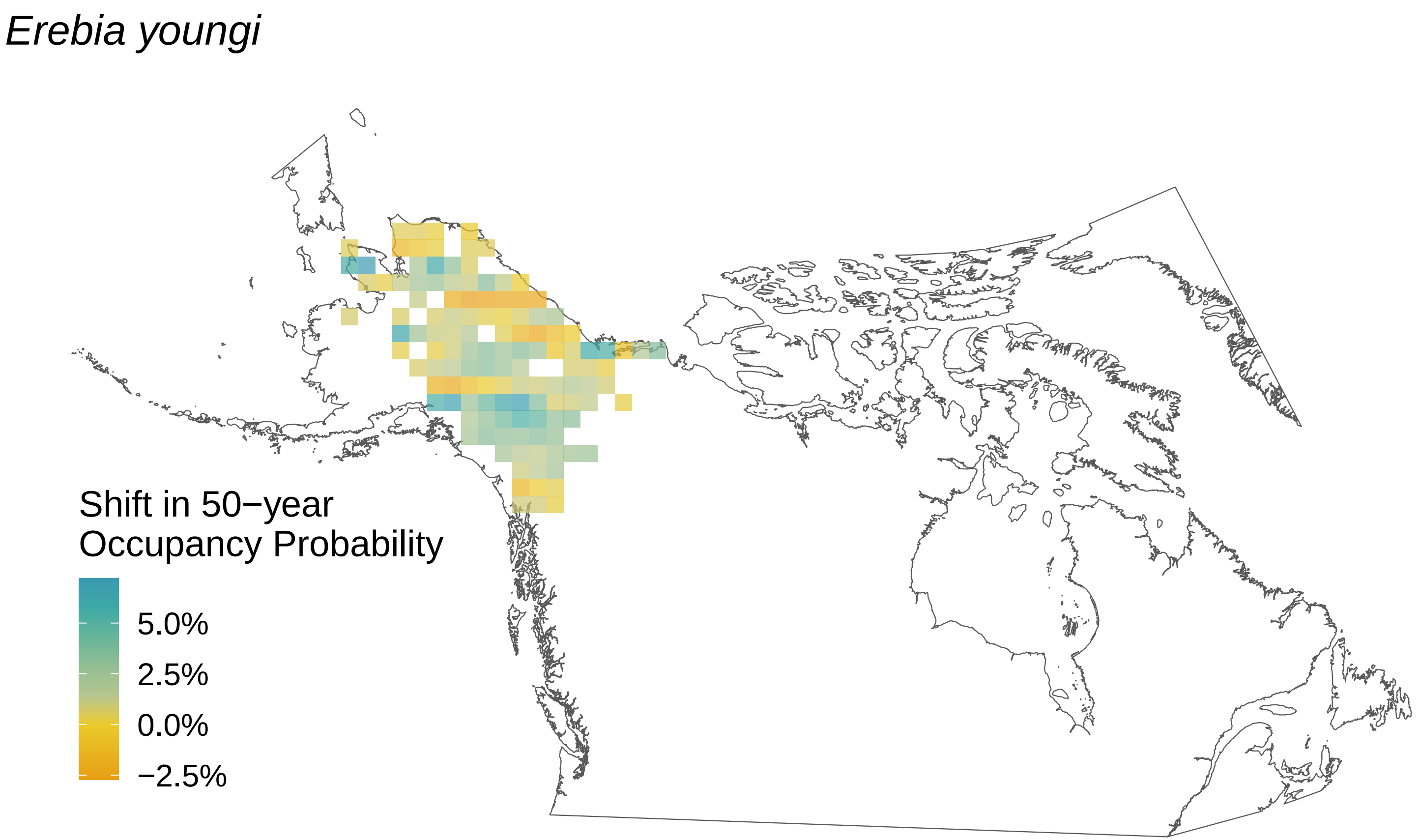

Erebia youngi
Shift in 50−year Occupancy Probability
5.0%
2.5%
0.0%
−2.5%

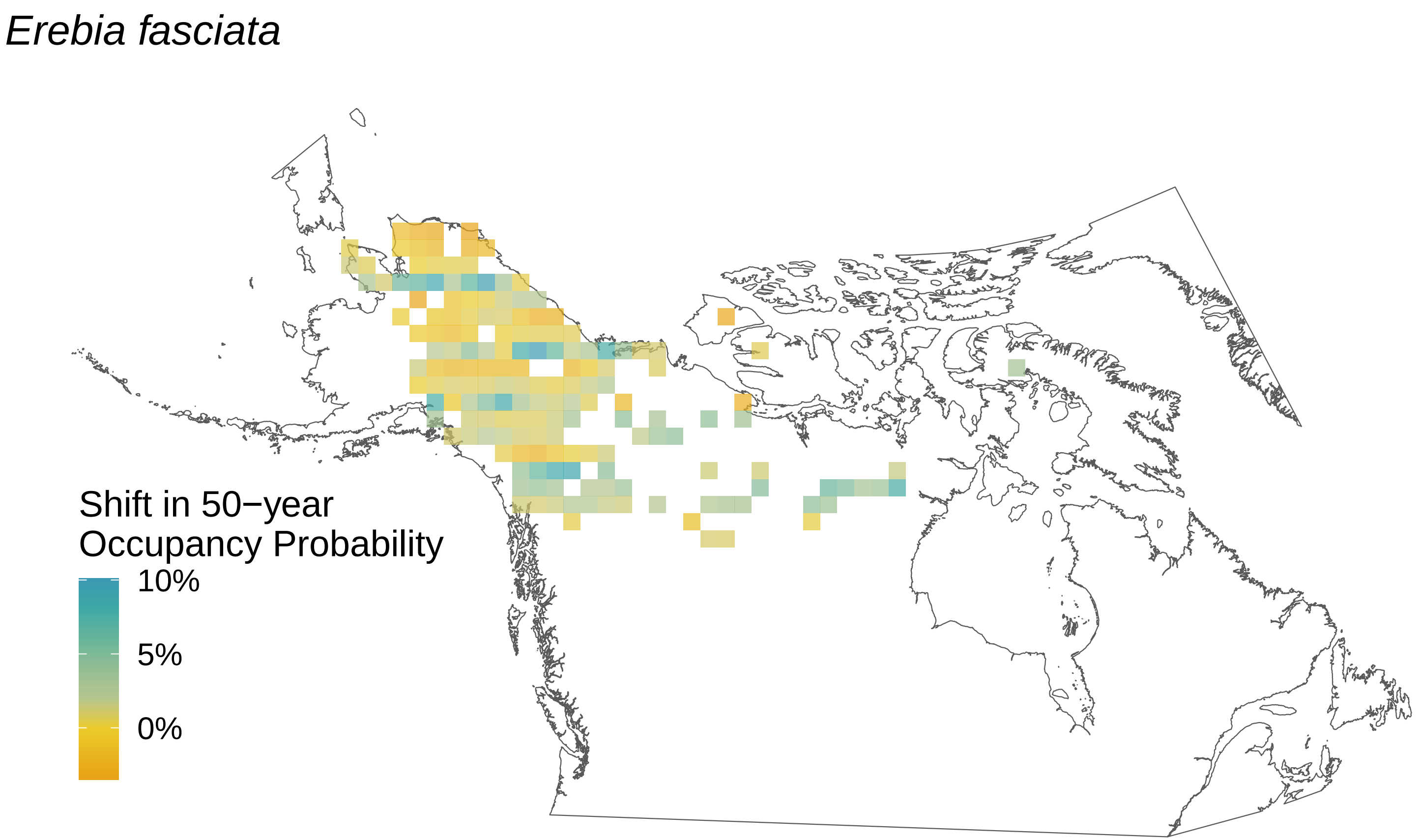
Erebia fasciata
Shift in 50−year Occupancy Probability
10%
5%
0%

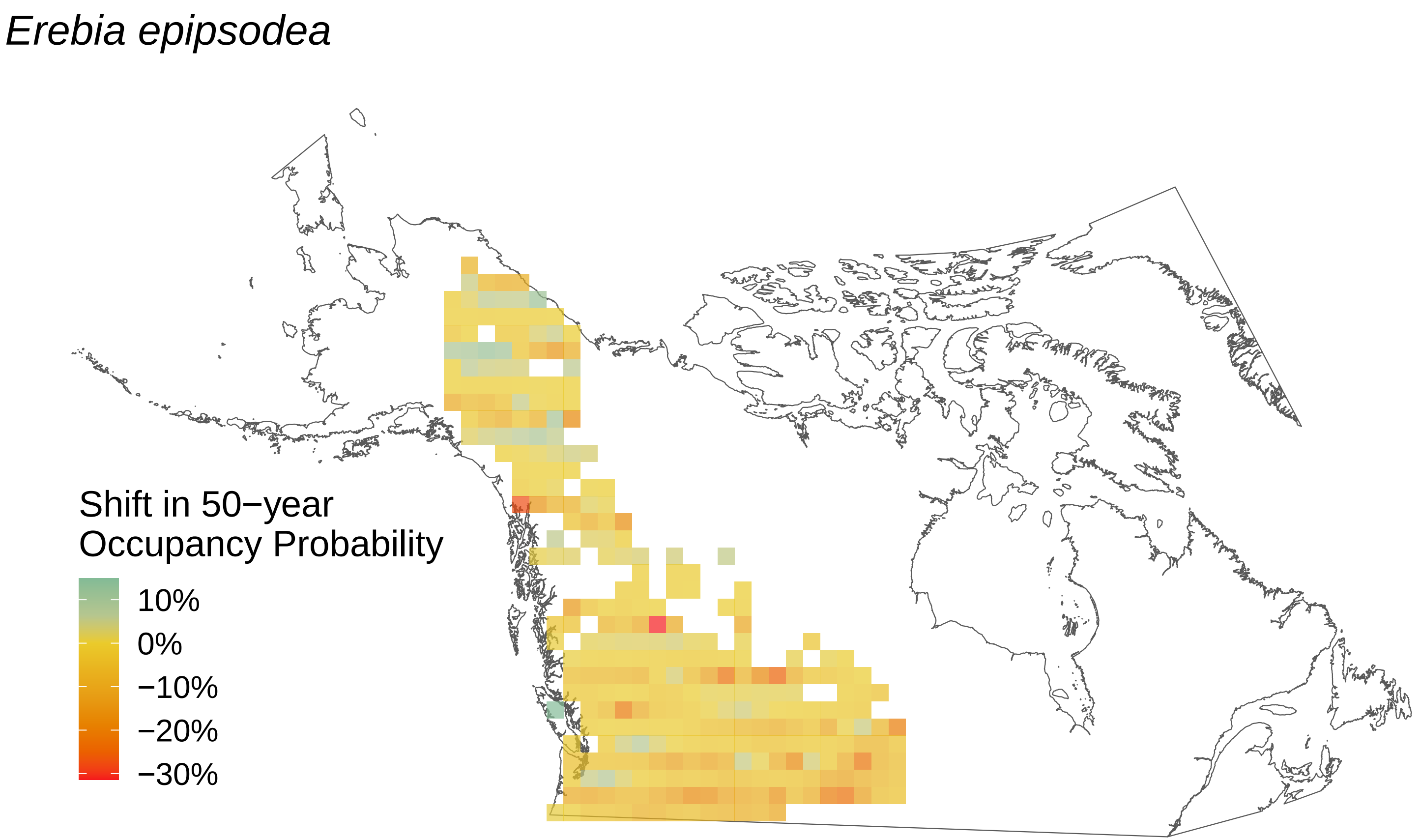

*Erebia epipsodea*
Shift in 50−year
Occupancy Probability
10%
0%
−10%
−20%
−30%

*Epargyreus clarus*

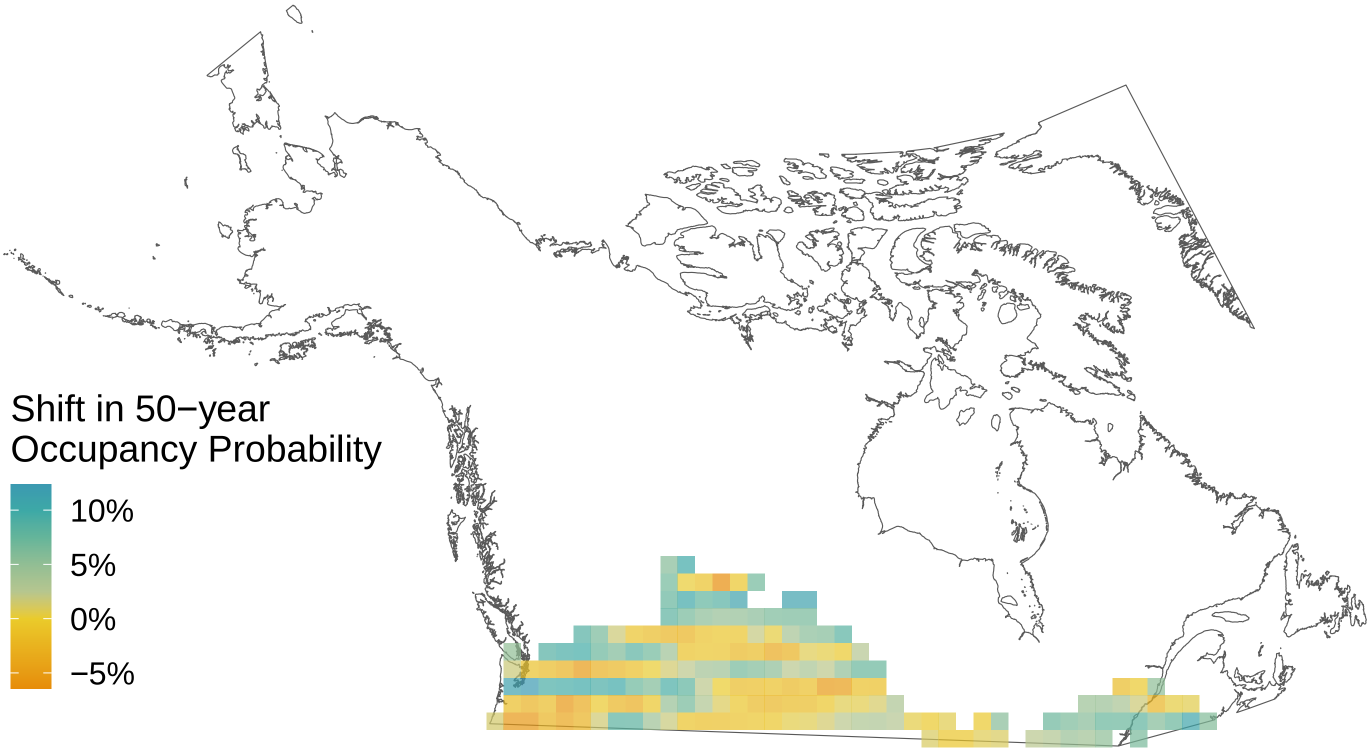

*Erebia discoidalis*

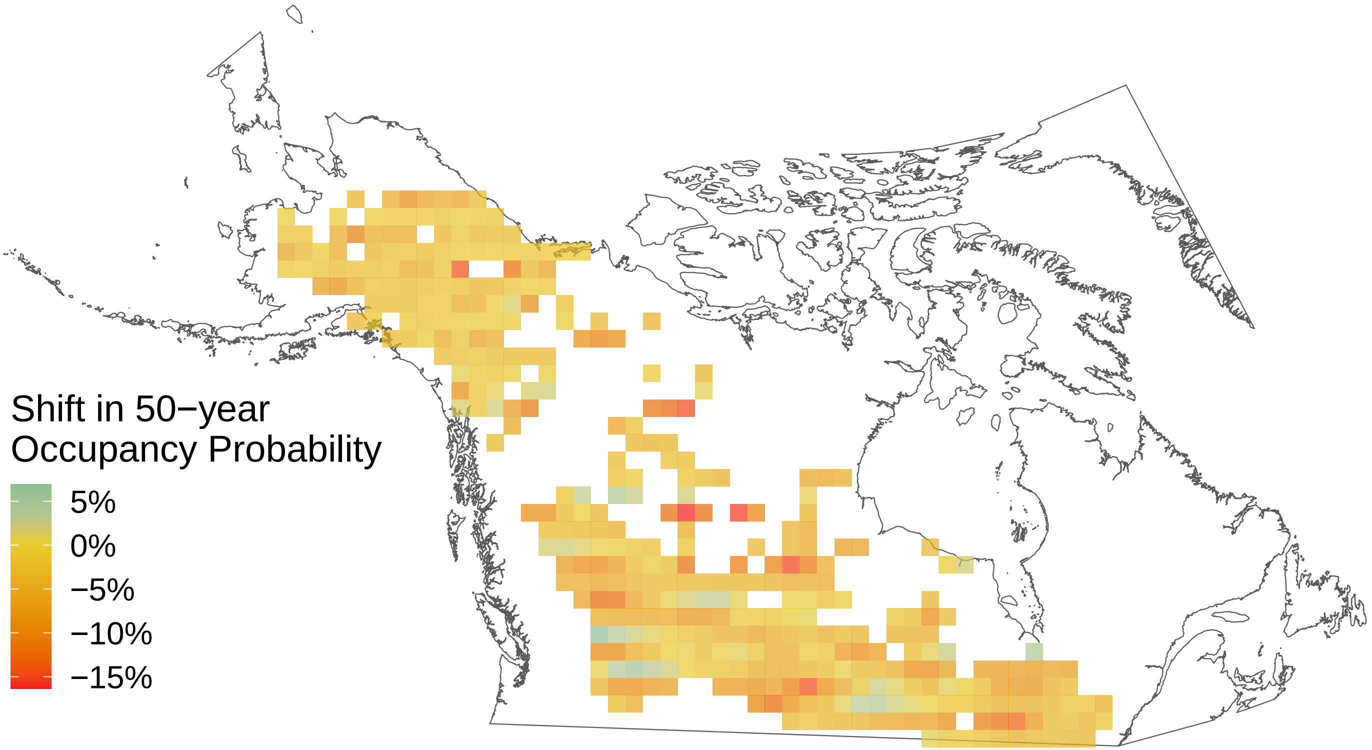

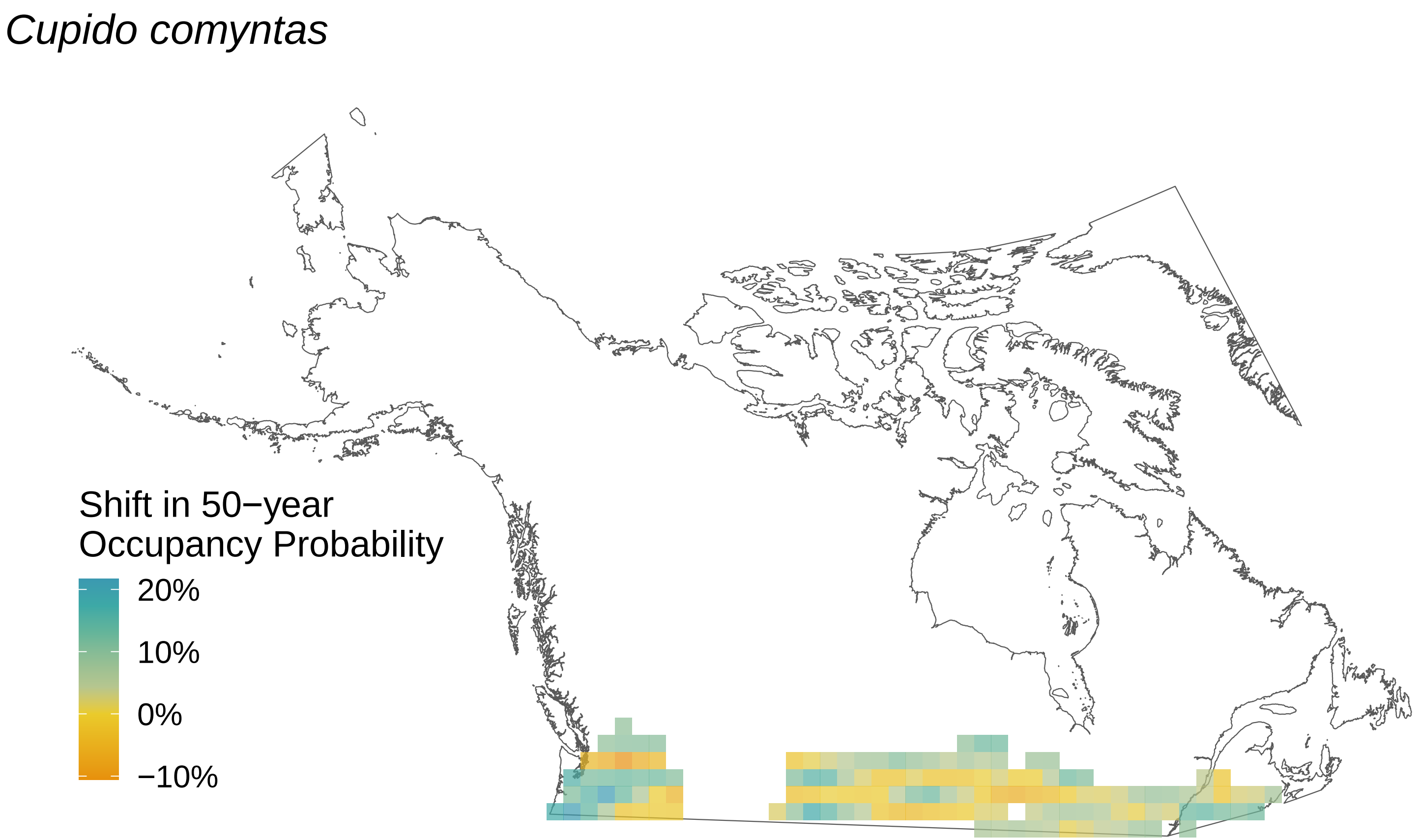
Cupido comyntas
Shift in 50−year
Occupancy Probability
20%
10%
0%
−10%

*Colias philodice*

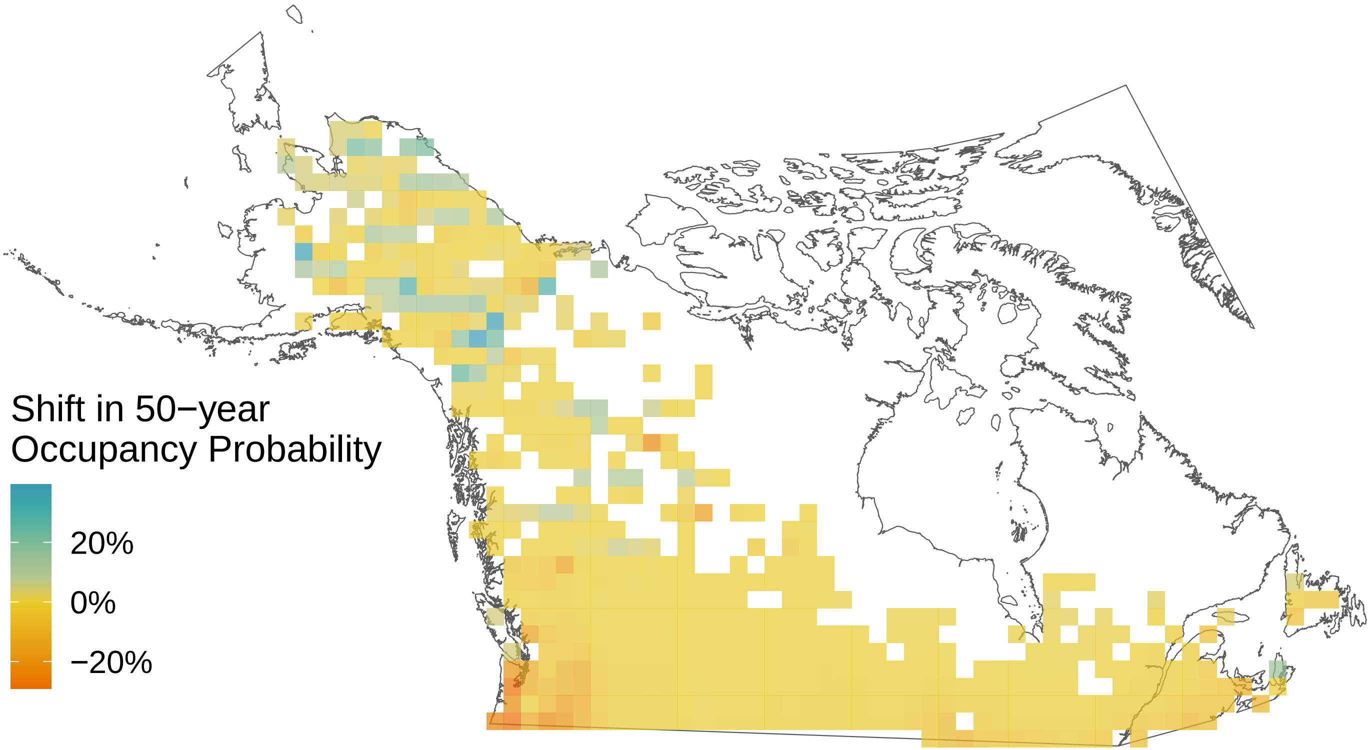

### *Cupido amyntula*

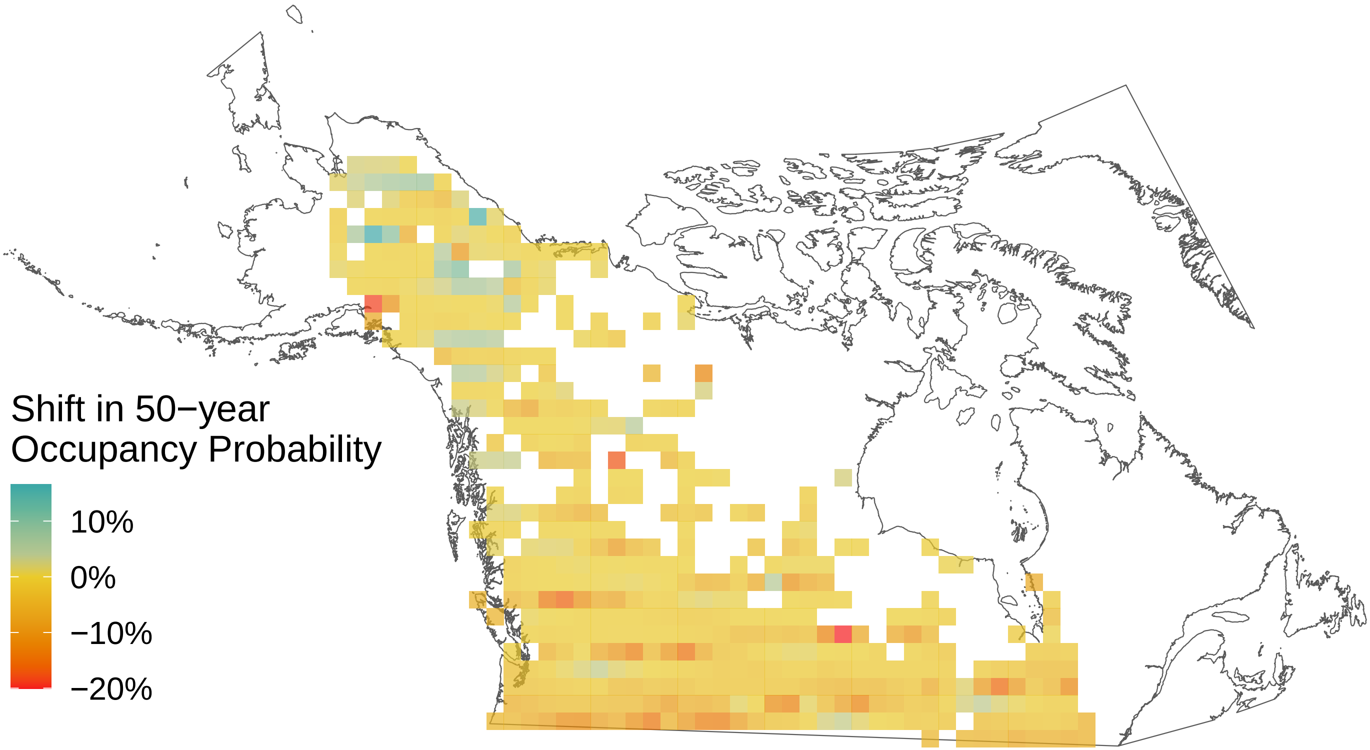

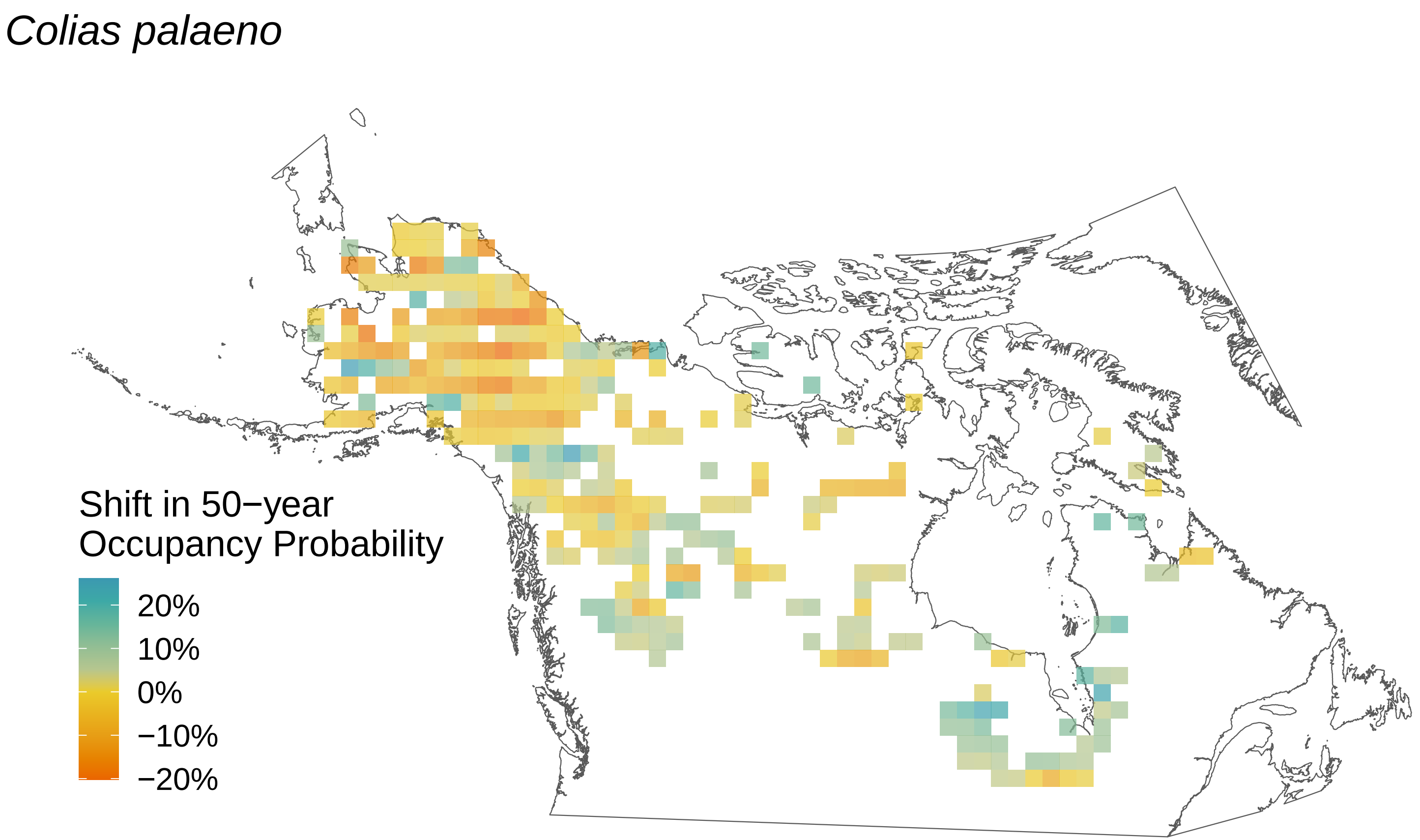

*Colias palaeno*
Shift in 50−year
Occupancy Probability
20%
10%
0%
−10%
−20%

*Colias interior*

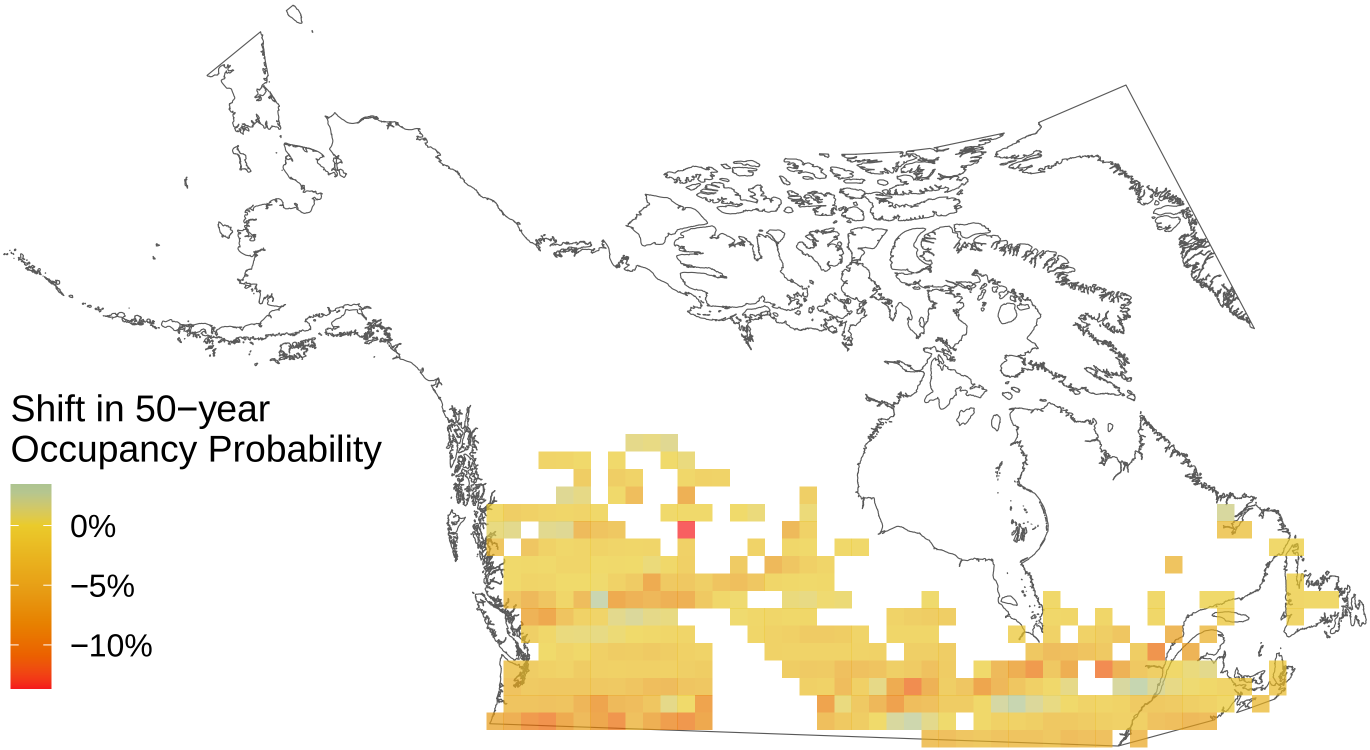

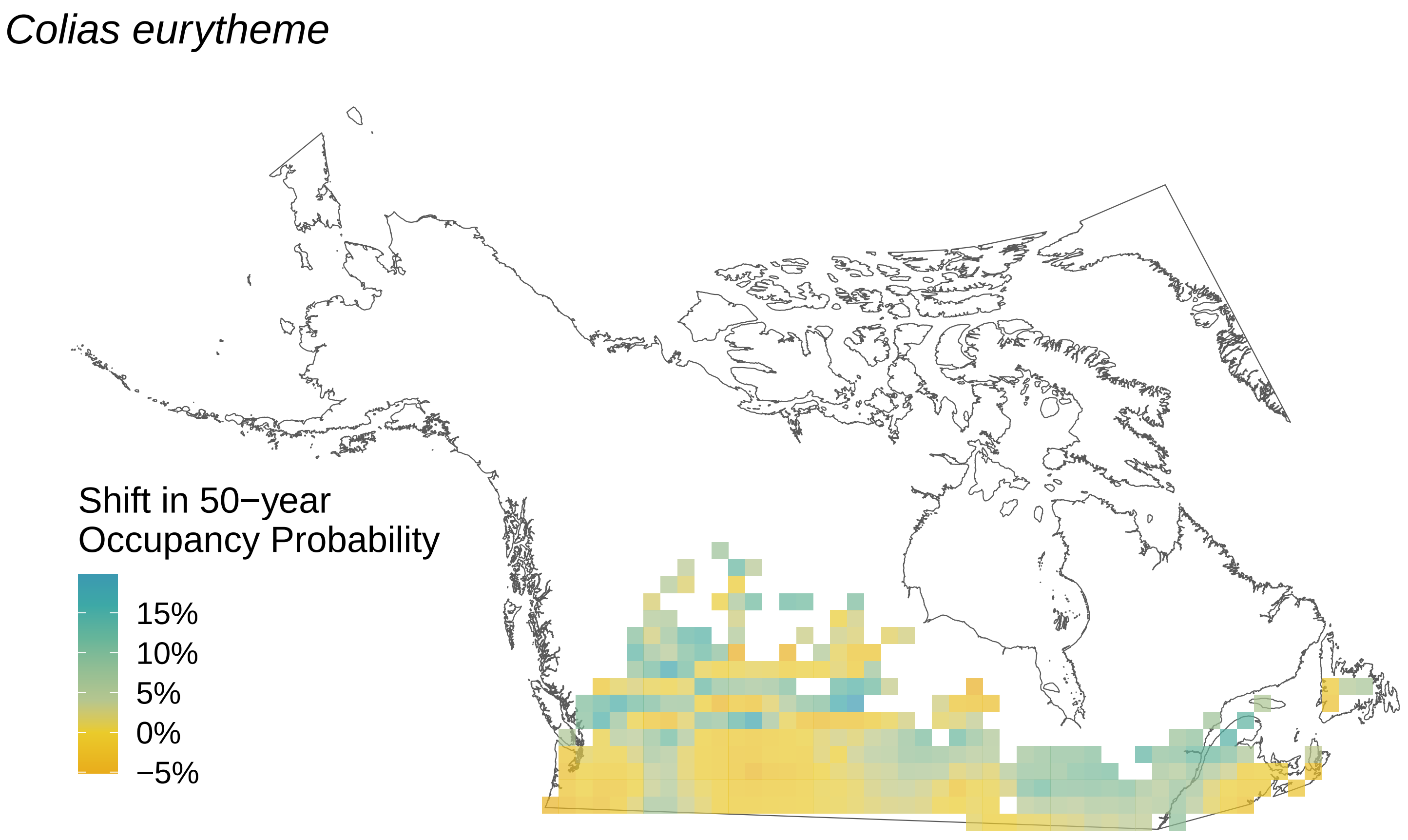
*Colias eurytheme*
Shift in 50−year
Occupancy Probability
15%
10%
5%
0%
−5%

### *Colias hecla*

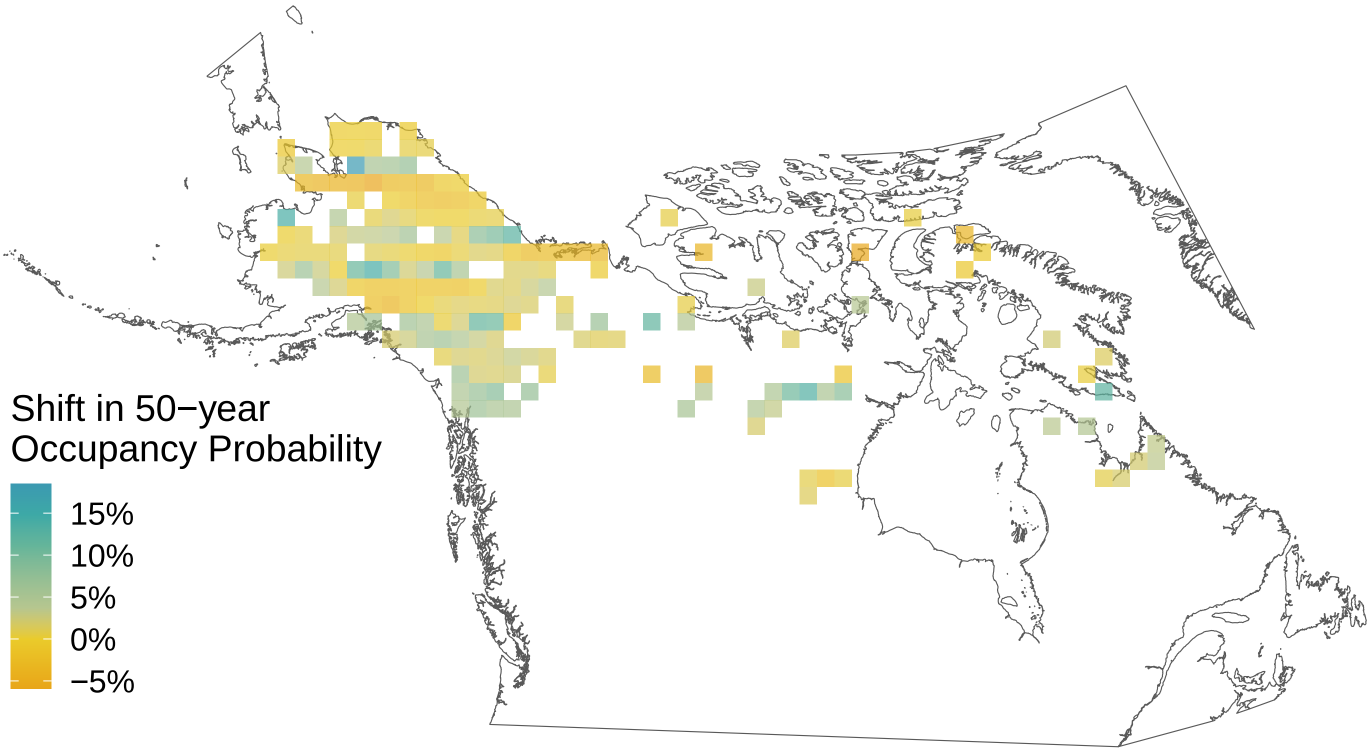

*Coenonympha tullia*

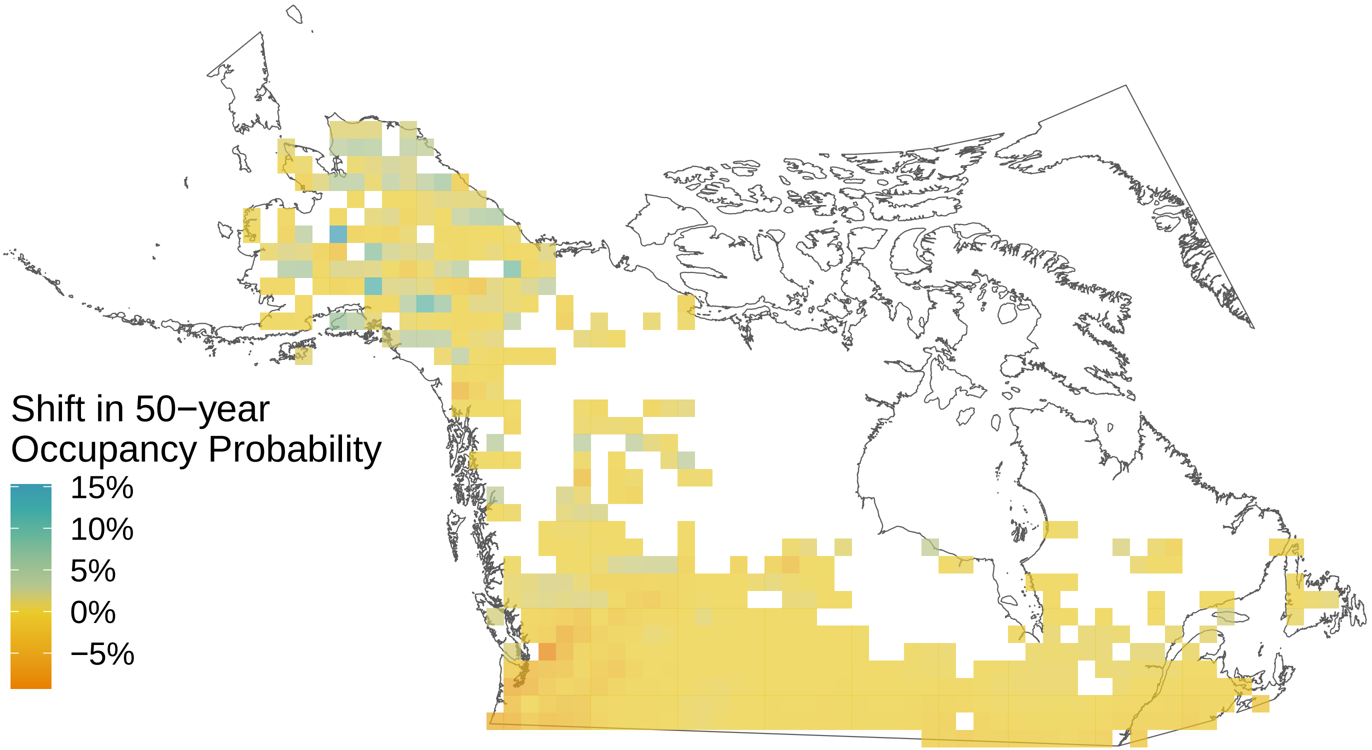

*Cercyonis pegala*

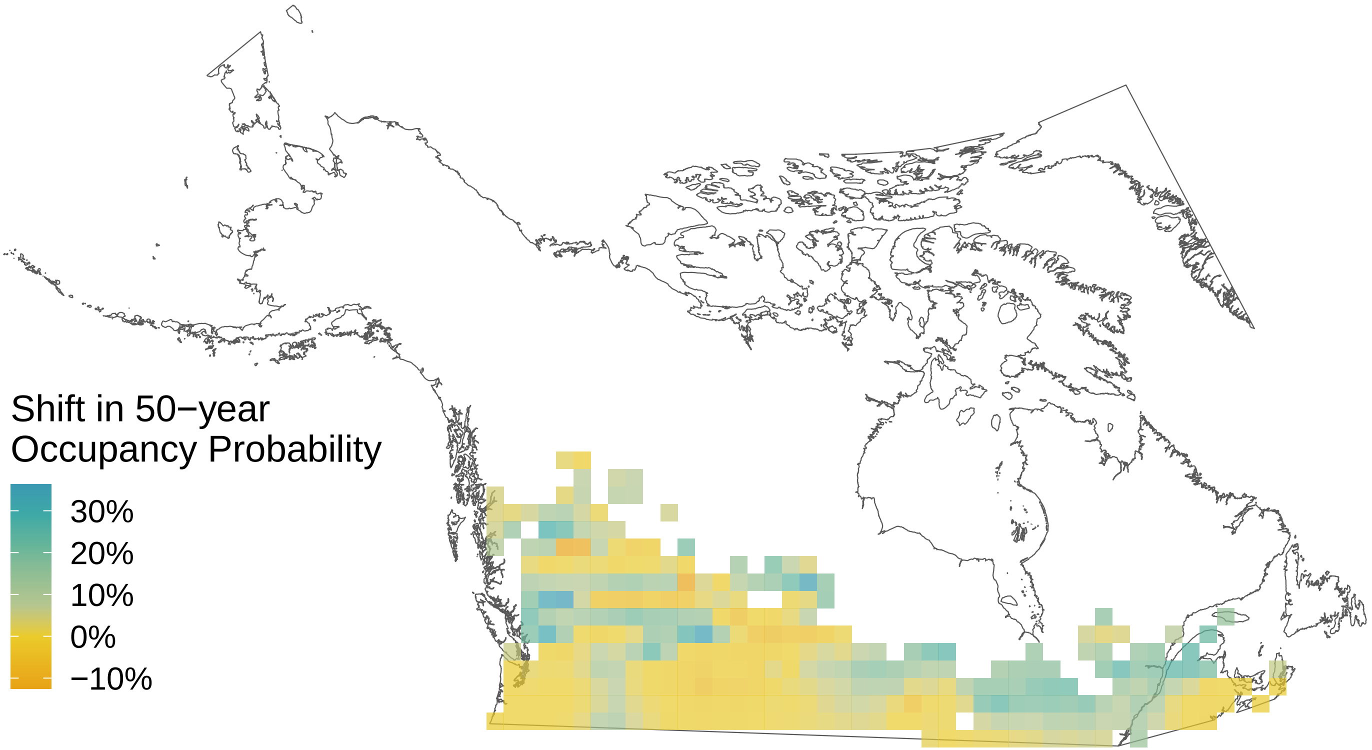

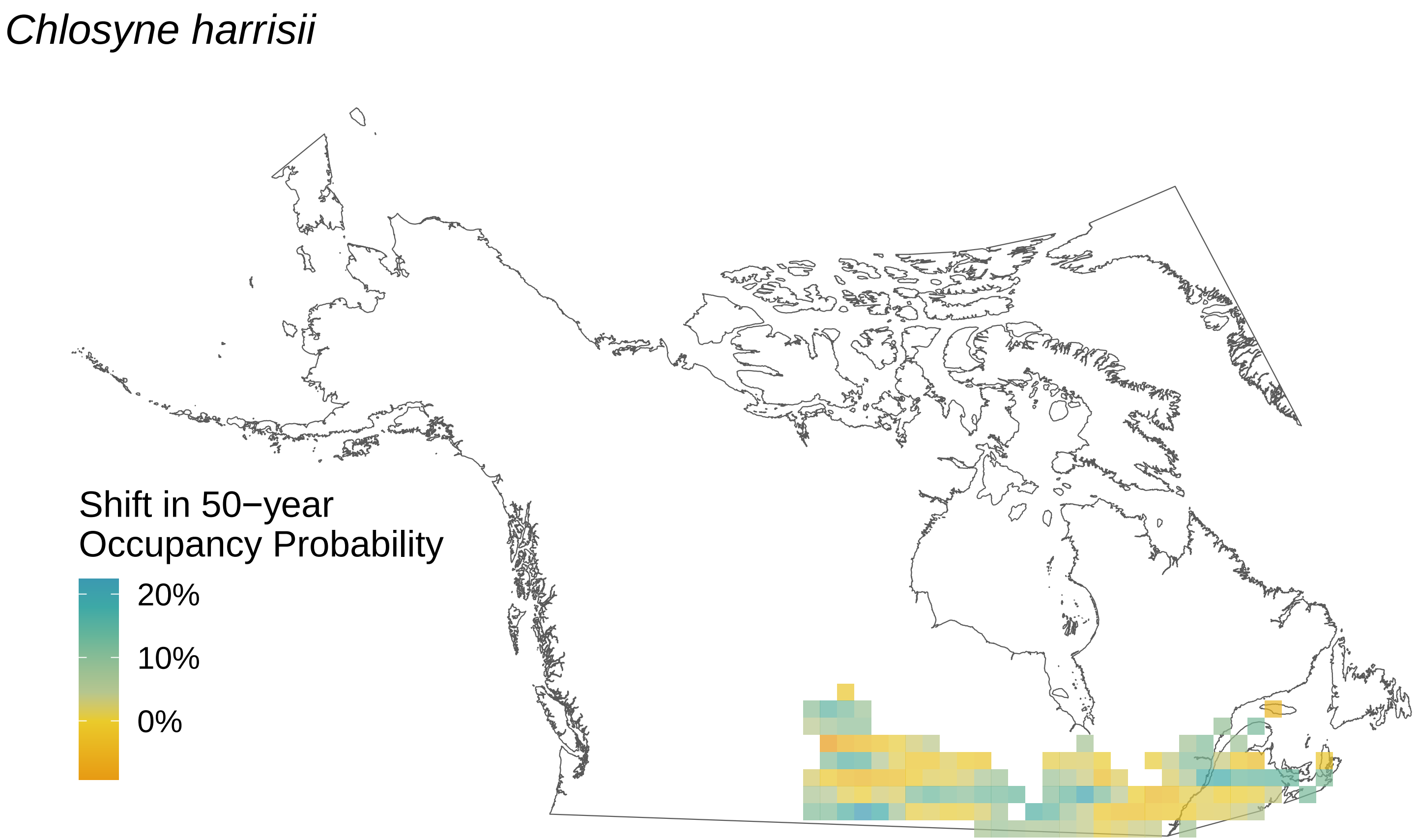
Chlosyne harrisii
Shift in 50−year
Occupancy Probability
20%
10%
0%

*Carterocephalus palaemon*

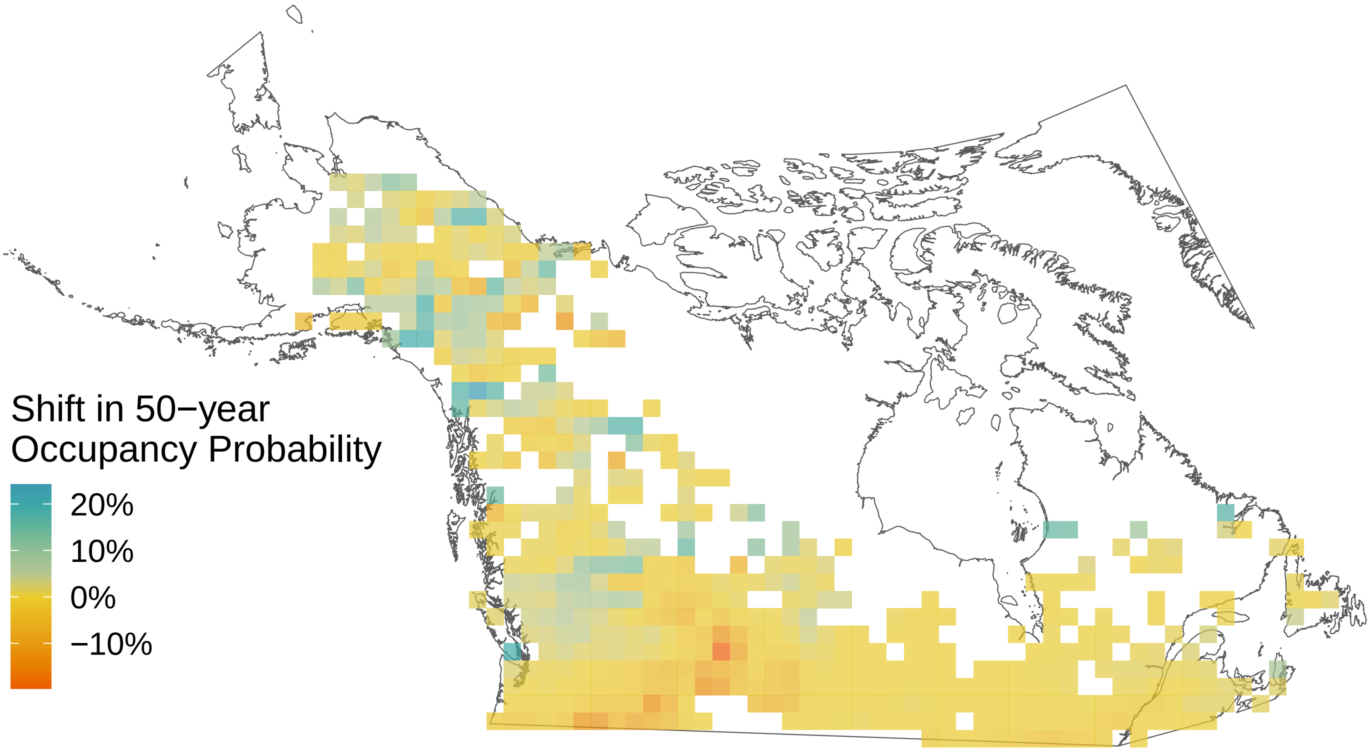

*Callophrys polios*

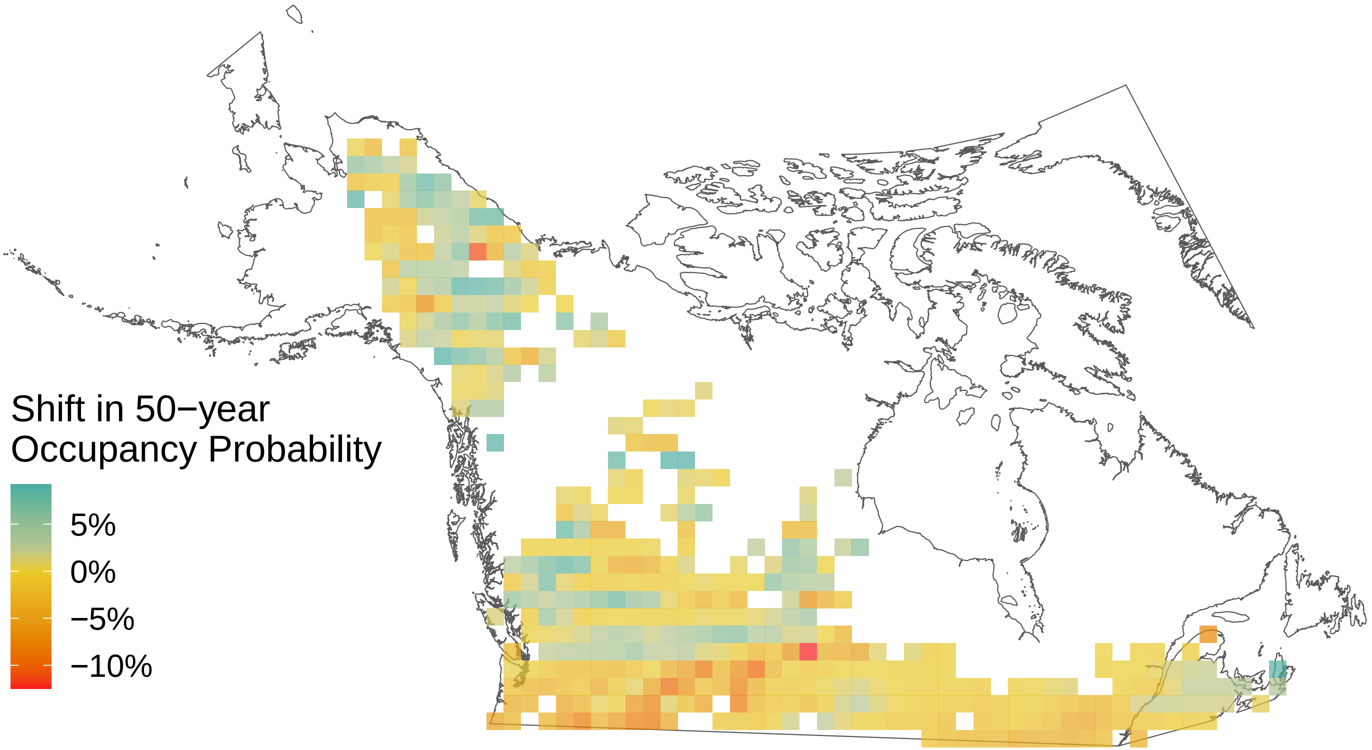

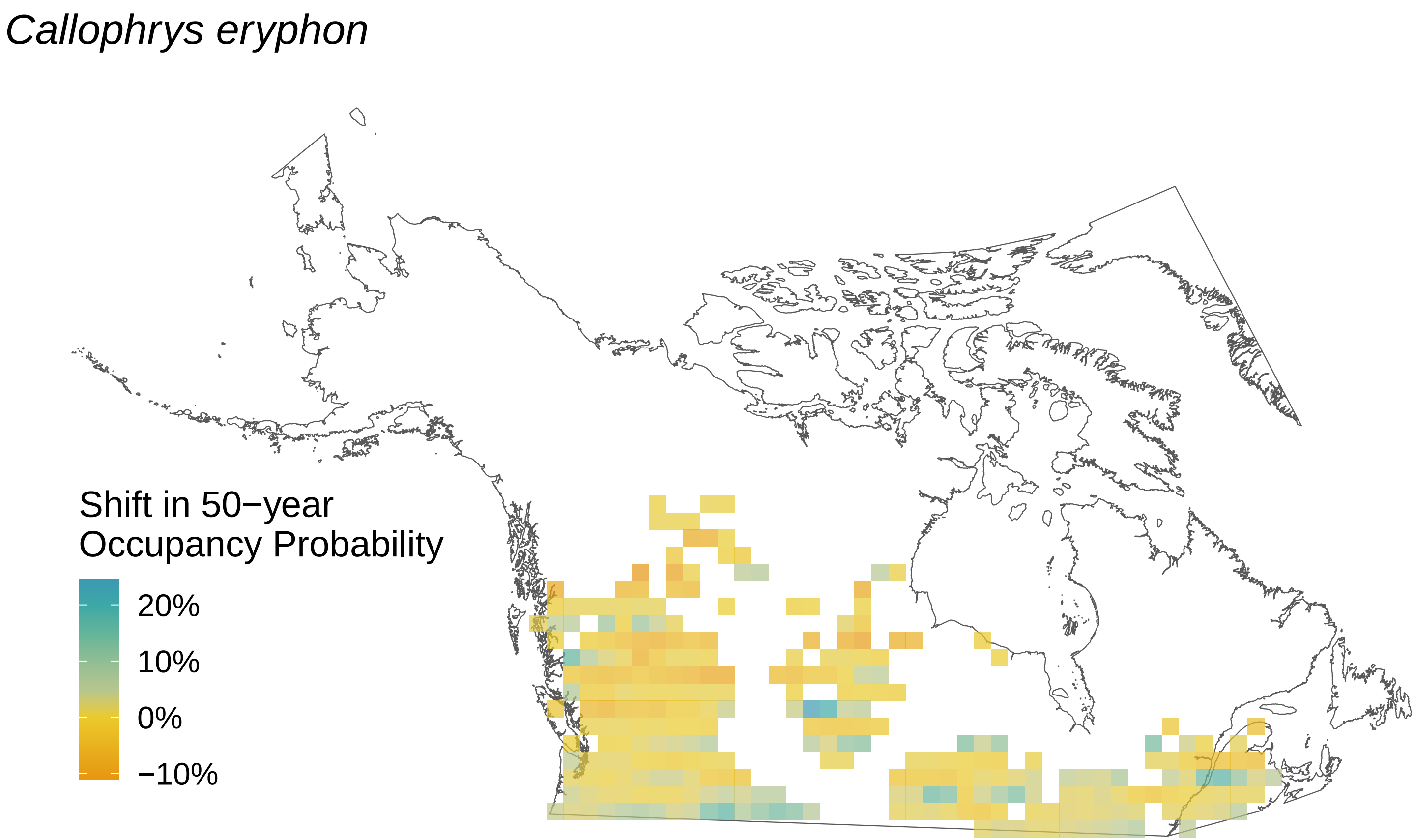
Callophrys eryphon
Shift in 50−year
Occupancy Probability
20%
10%
0%
−10%

*Callophrys niphon*

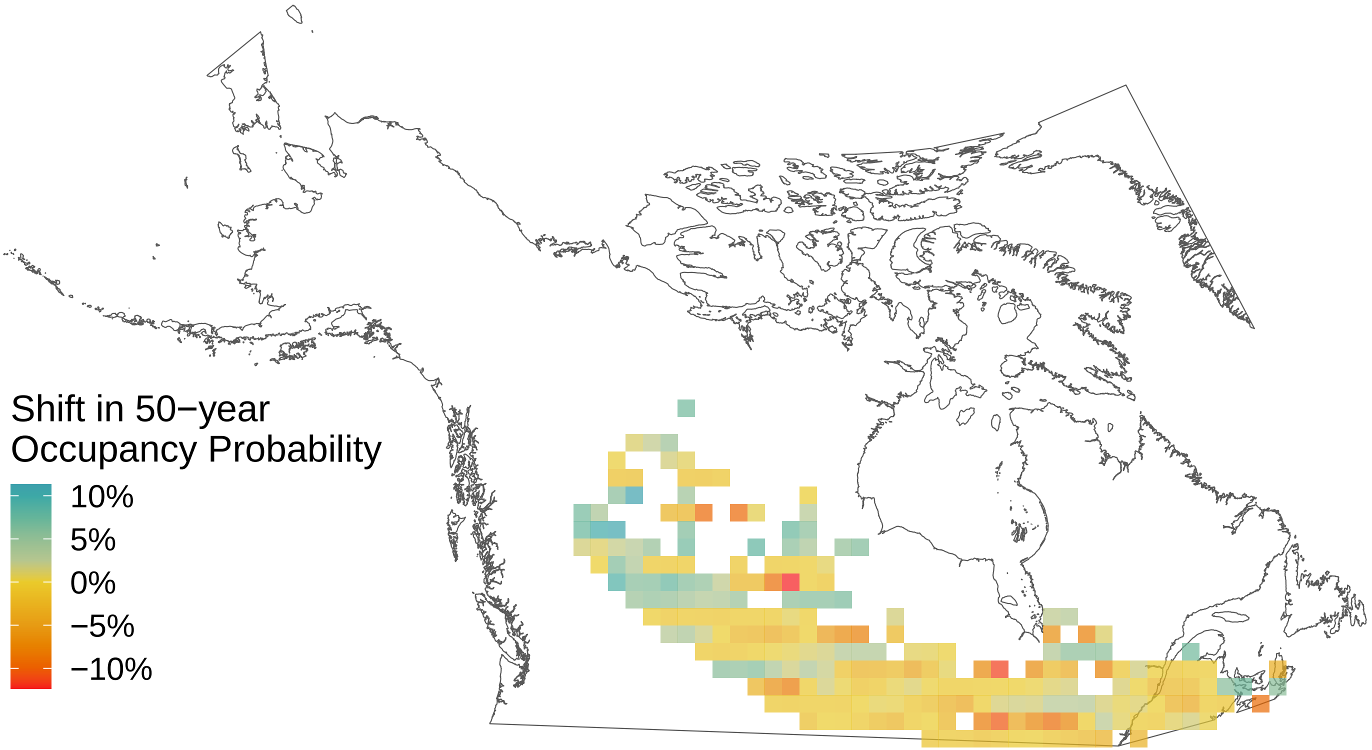

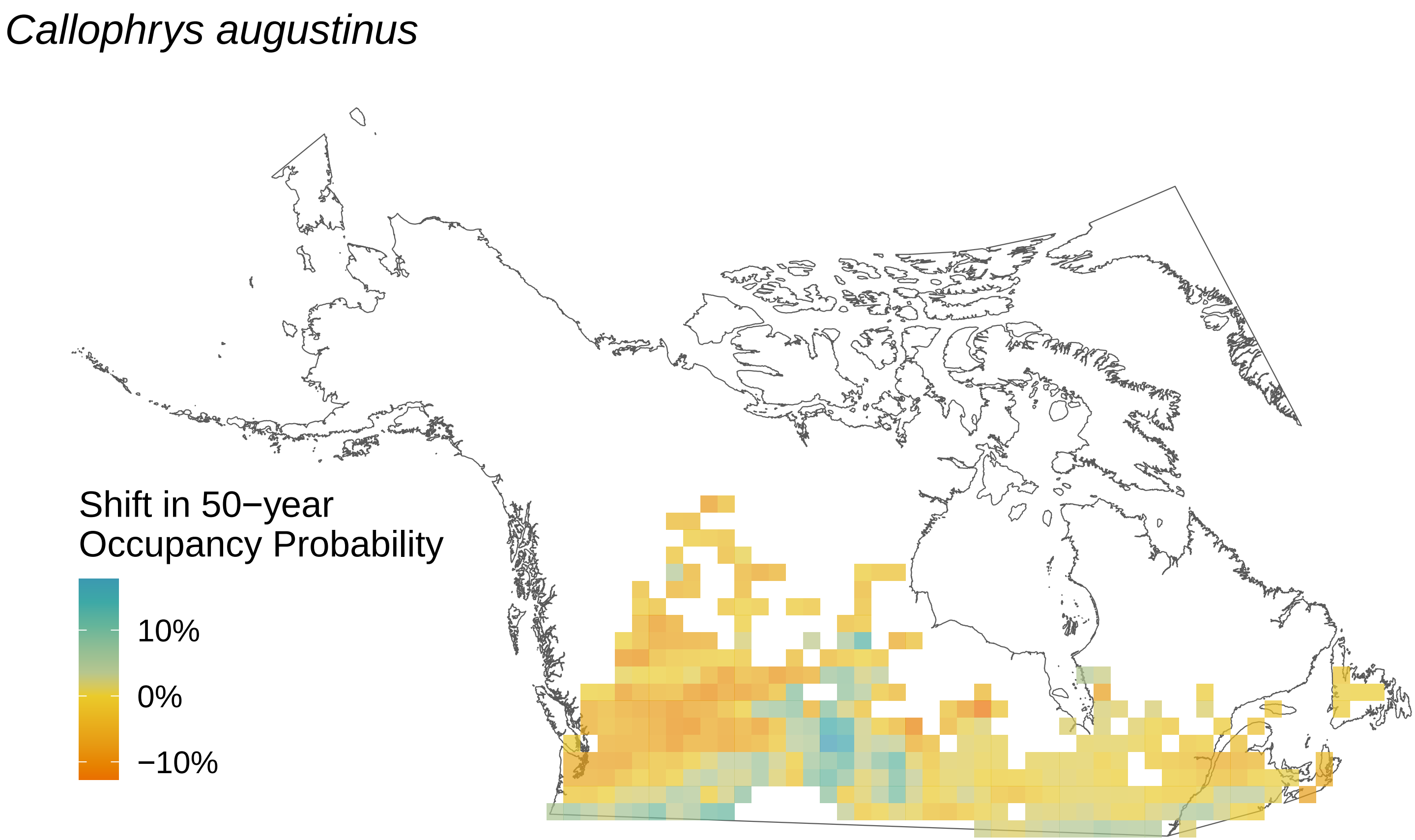

Callophrys augustinus
Shift in 50−year Occupancy Probability
10%
0%
−10%

*Boloria selene*

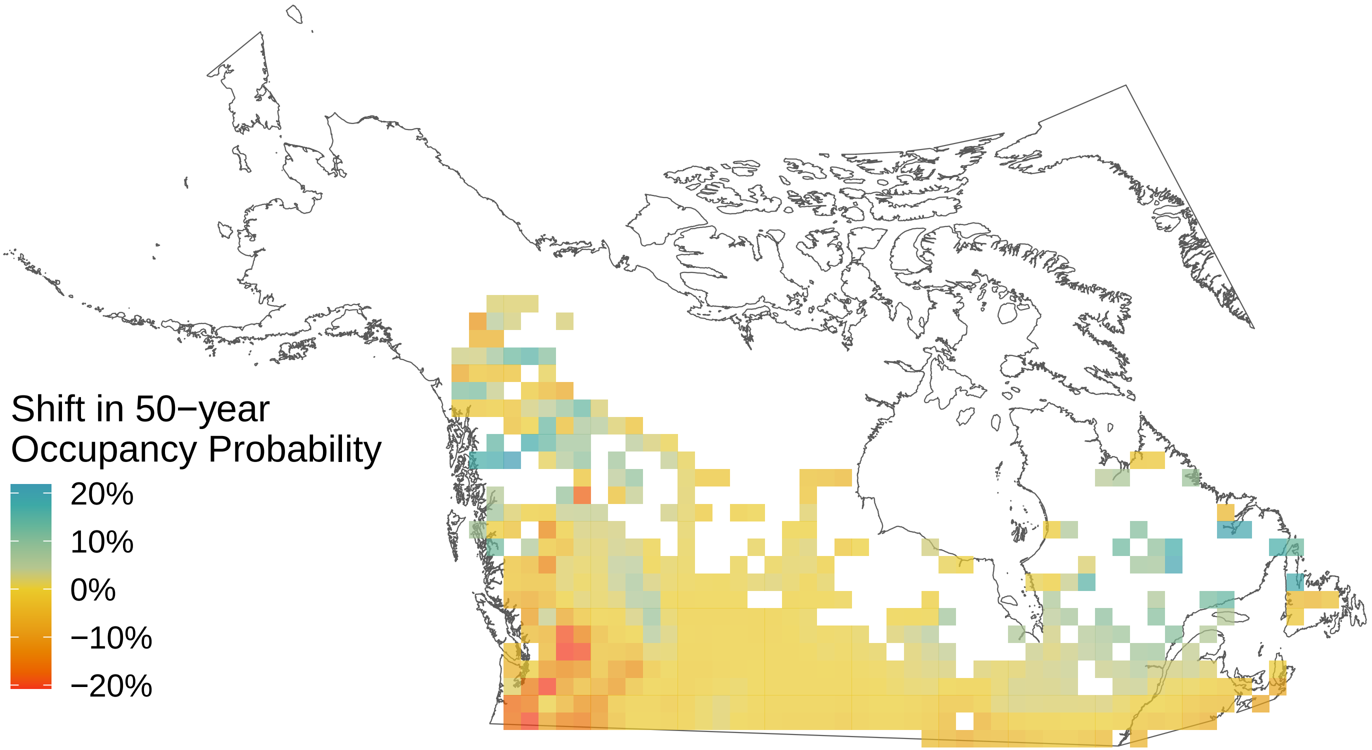

*Boloria improba*

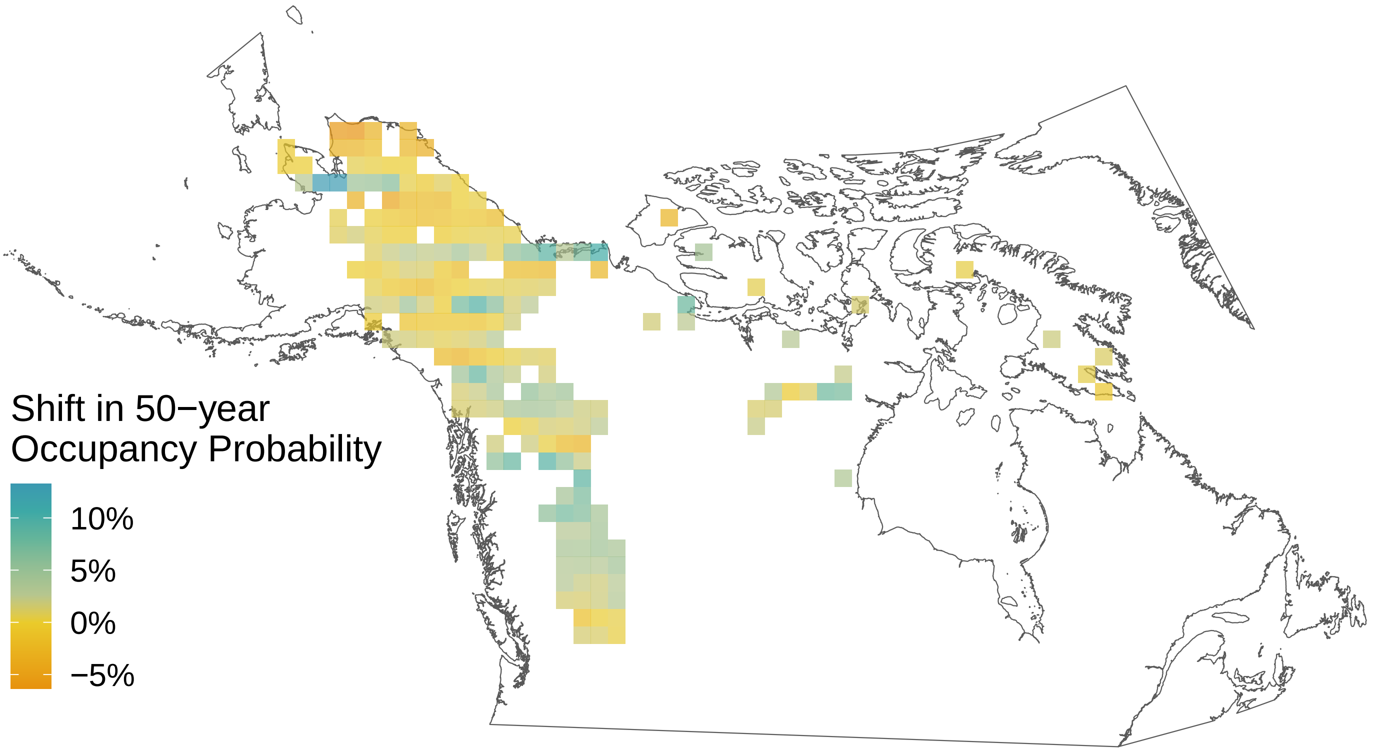

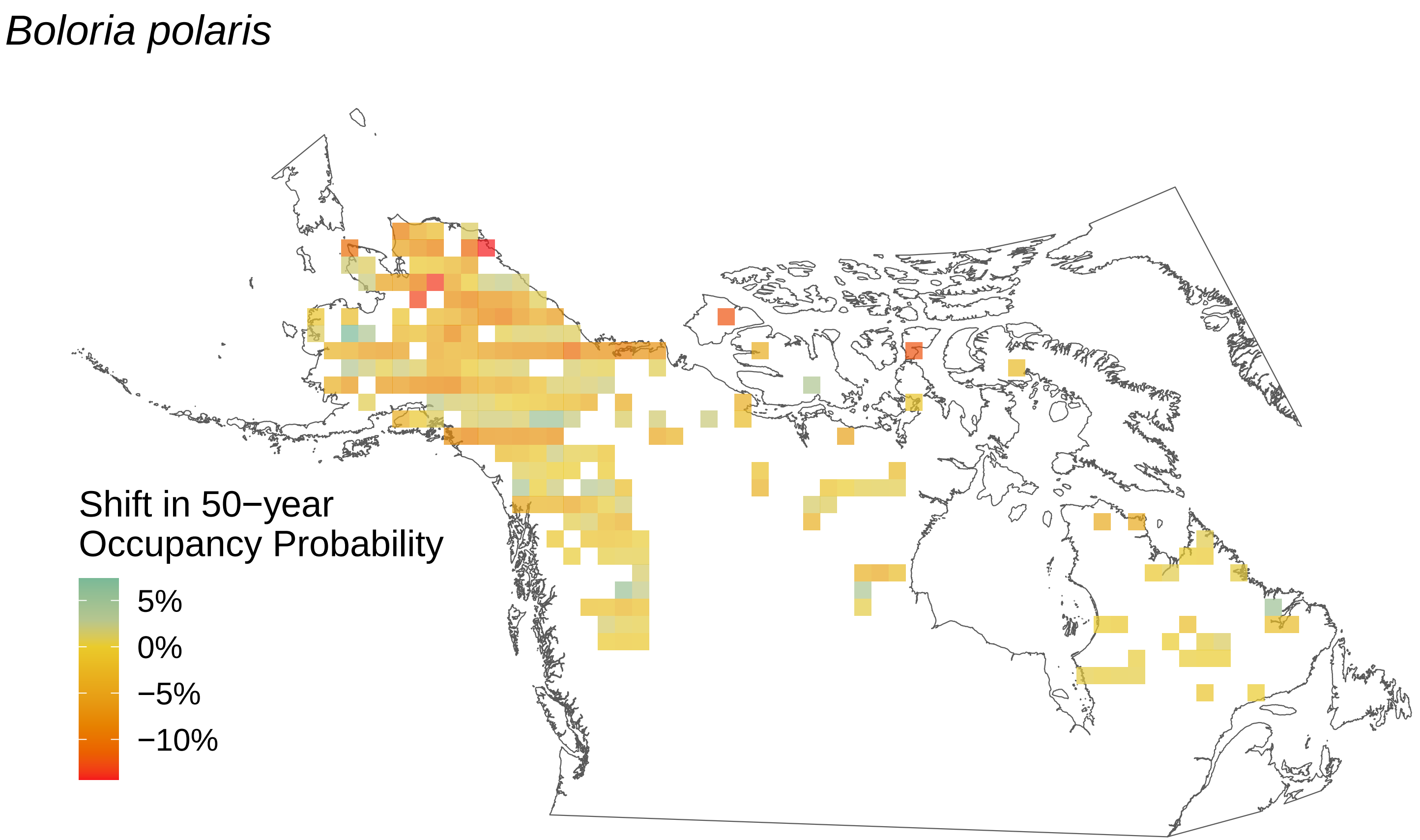

*Boloria polaris*
Shift in 50−year
Occupancy Probability
5%
0%
−5%
−10%

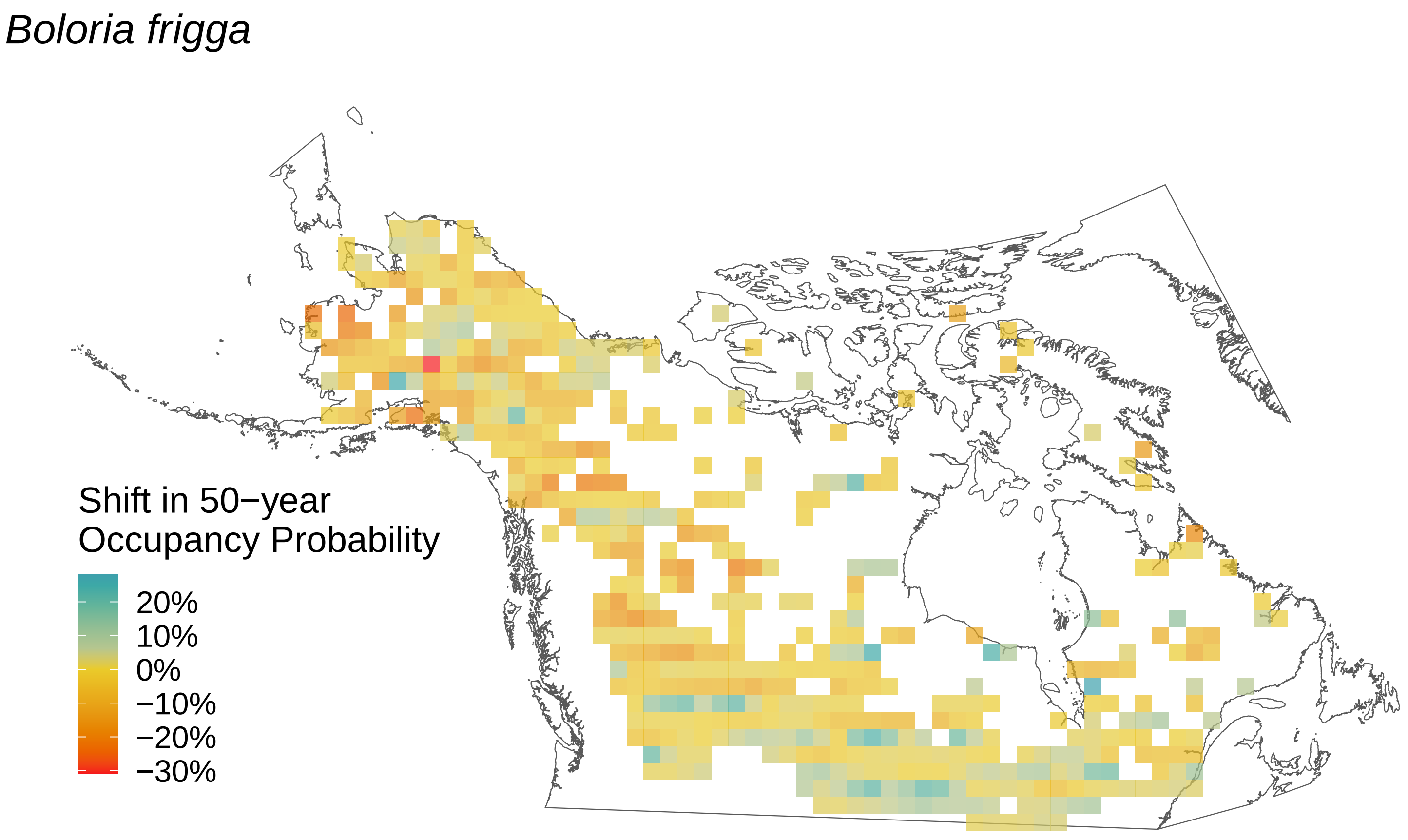

Boloria frigga
Shift in 50−year
Occupancy Probability
20%
10%
0%
−10%
−20%
−30%

*Boloria eunomia*

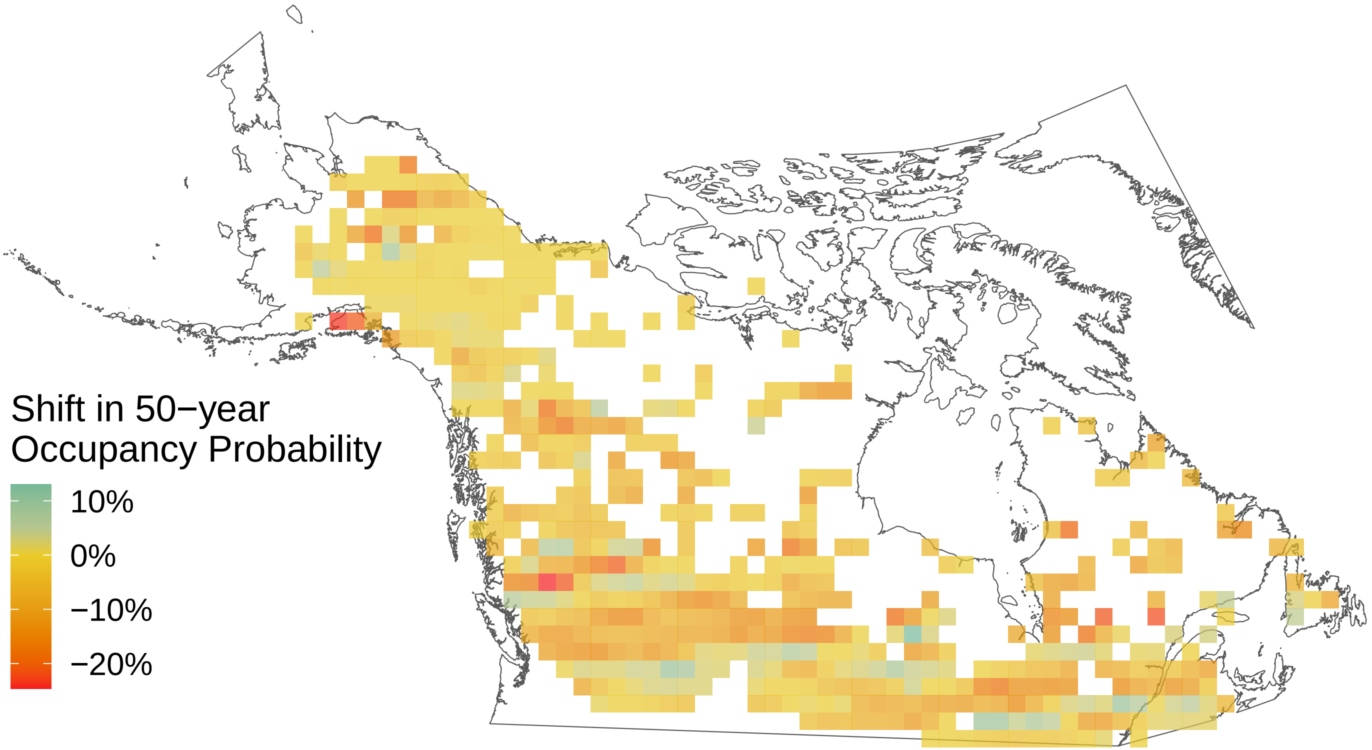

*Boloria freija*

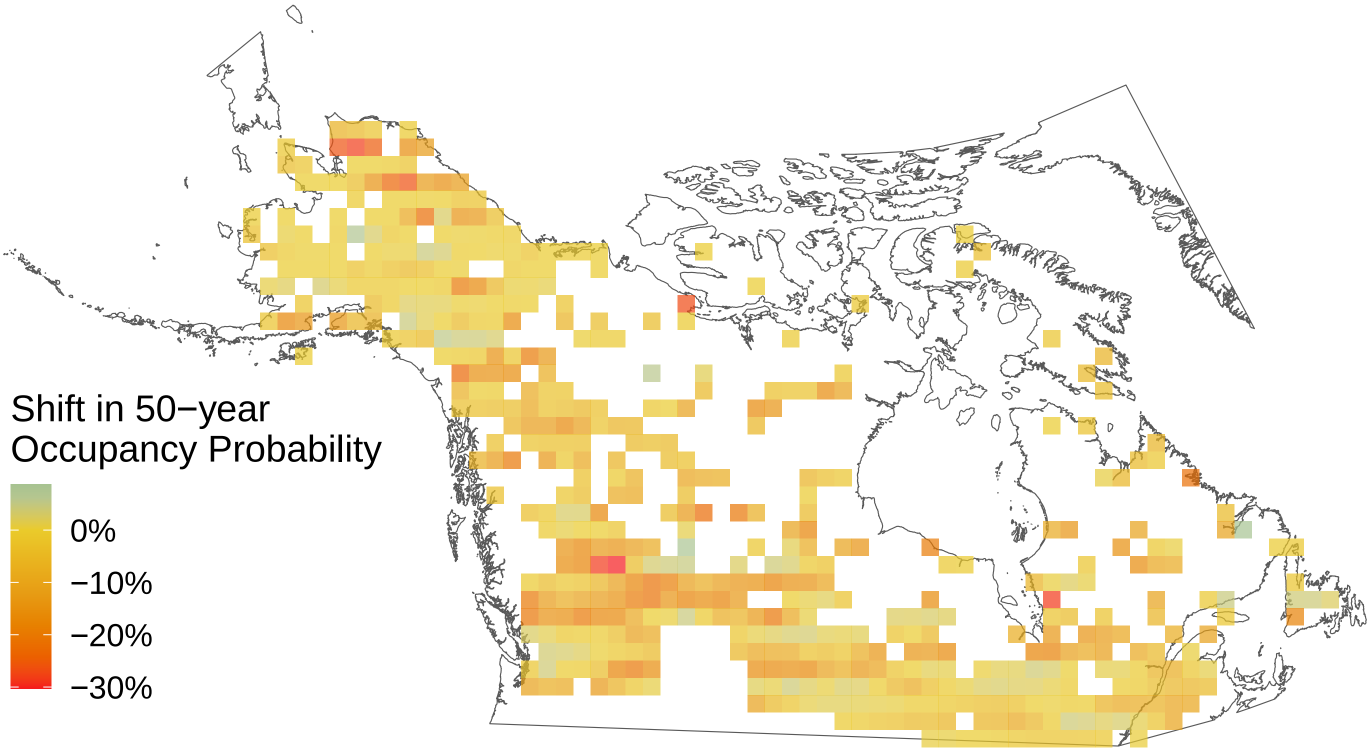

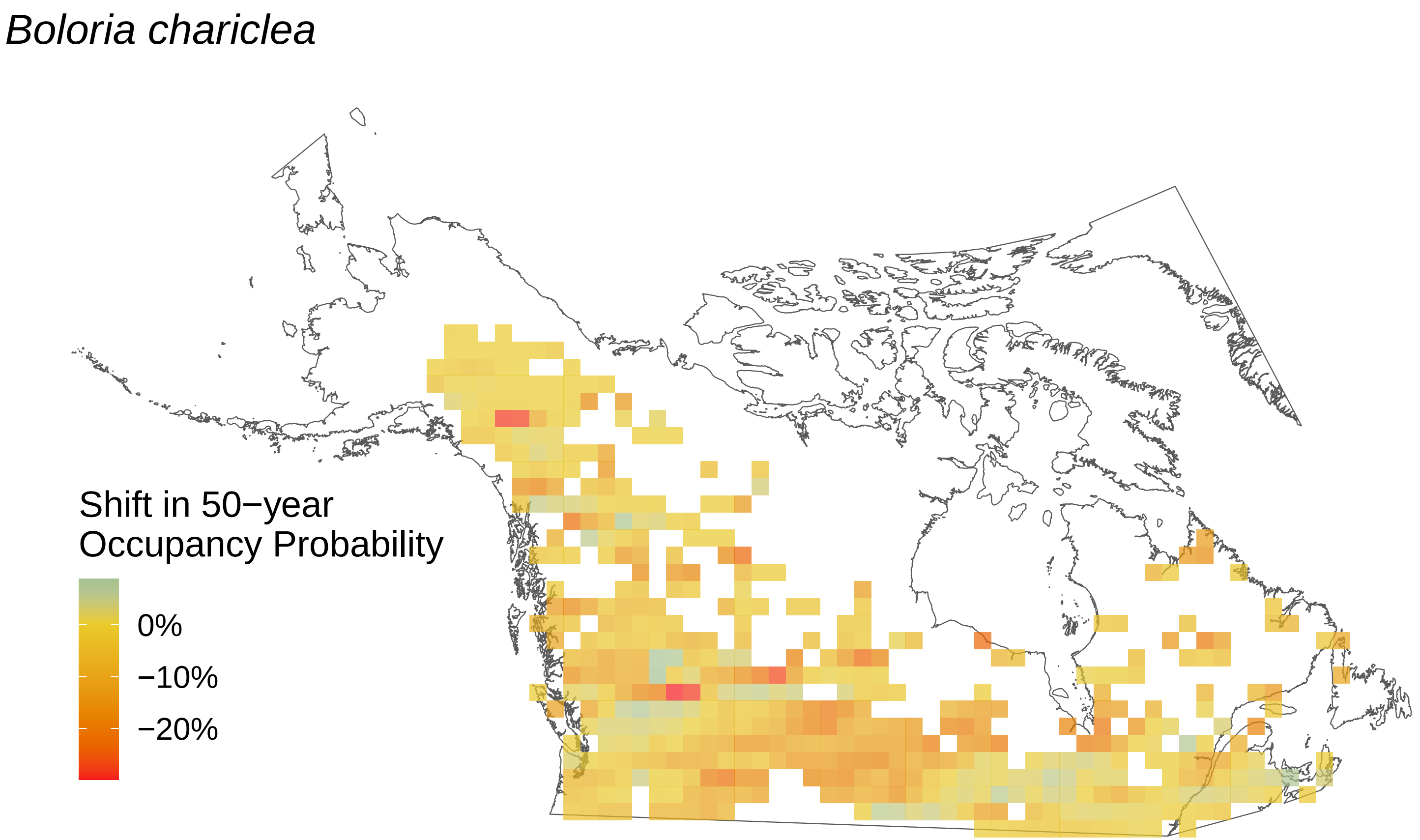

*Boloria chariclea*
Shift in 50−year
Occupancy Probability
0%
−10%
−20%

*Boloria bellona*

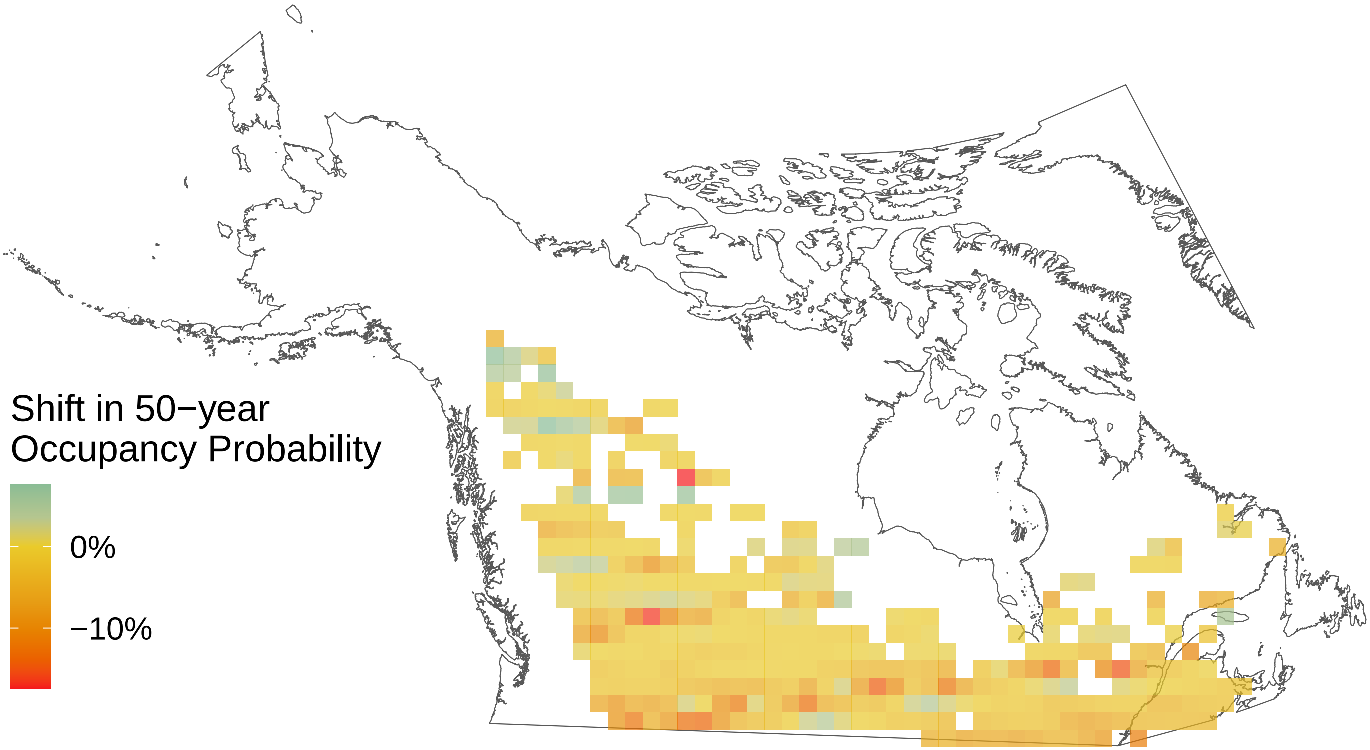

*Argynnis hydaspe*

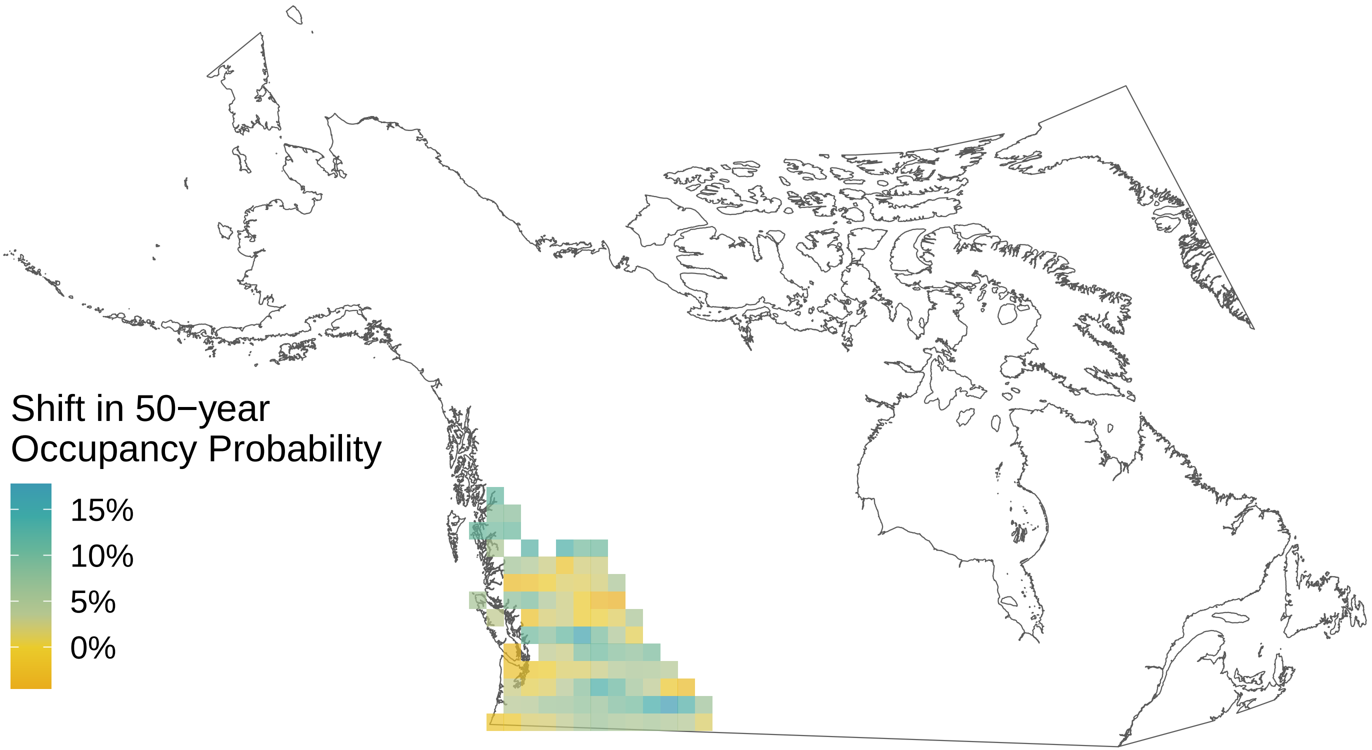

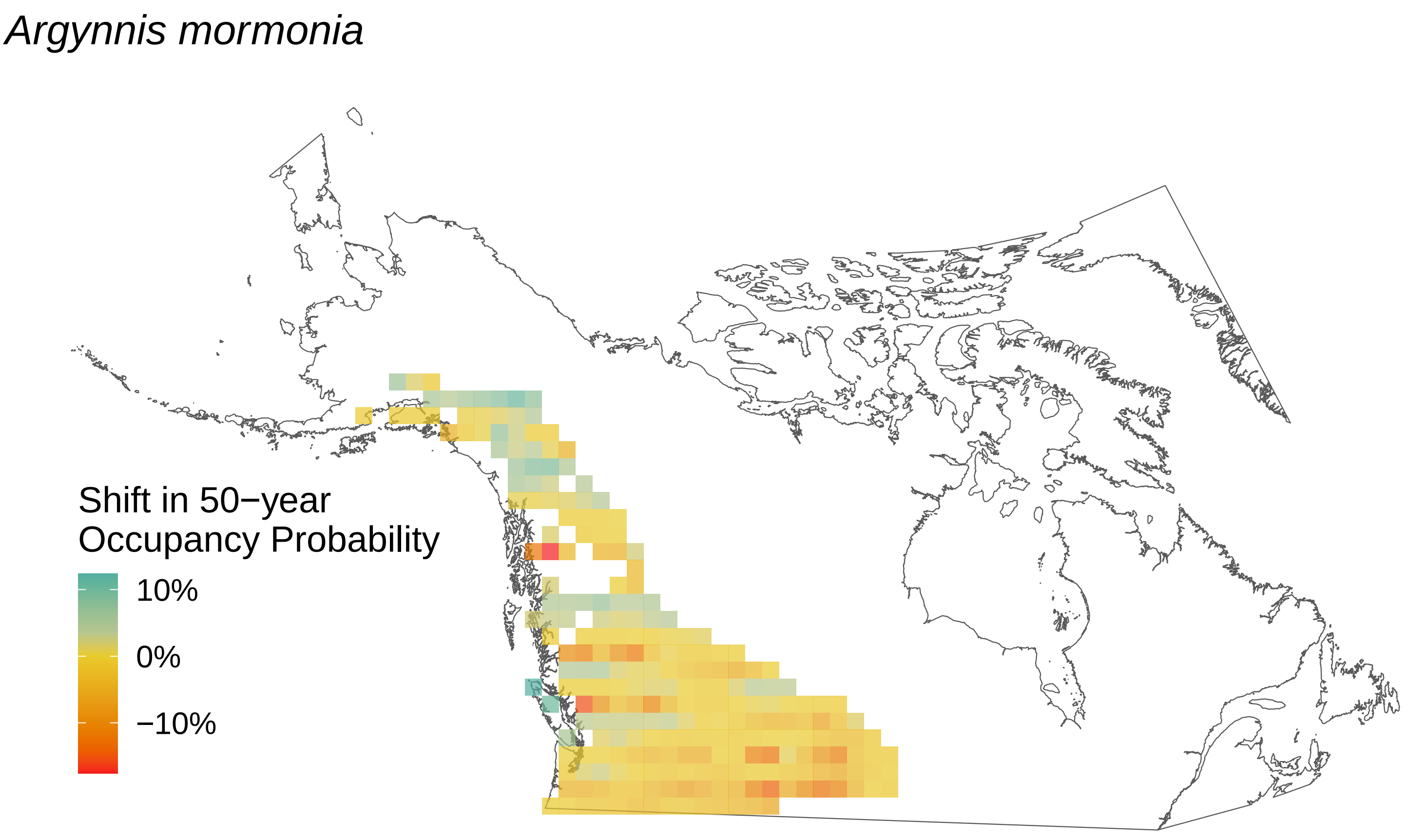

Argynnis mormonia
Shift in 50−year
Occupancy Probability
10%
0%
−10%

### *Argynnis cybele*

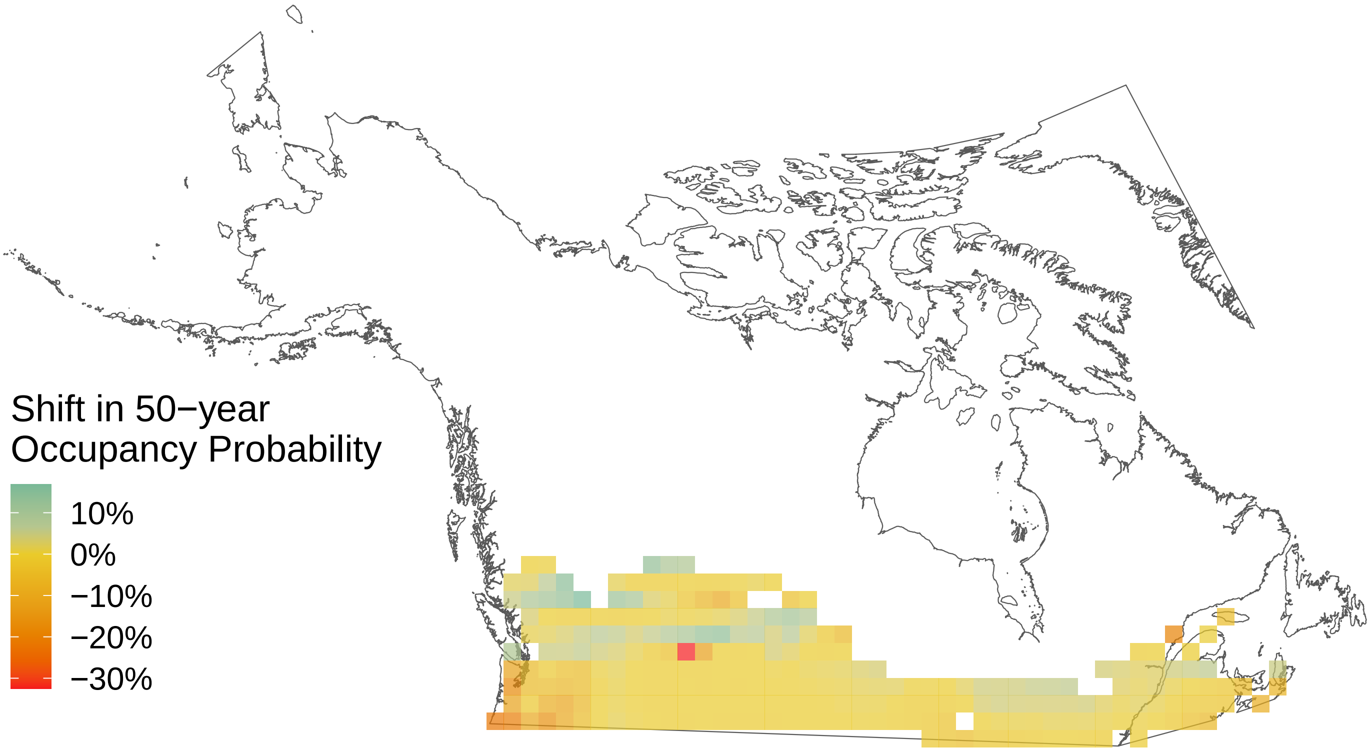

*Argynnis atlantis*

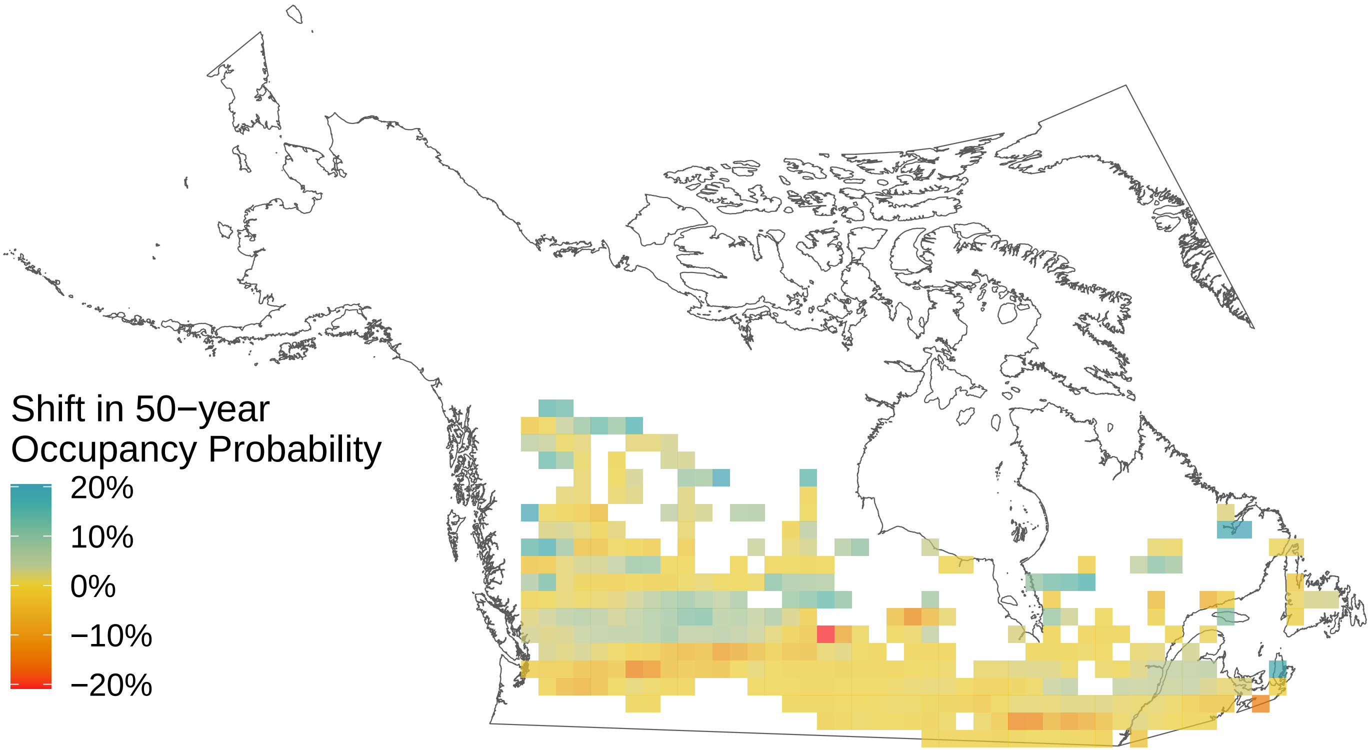

*Amblyscirtes vialis*

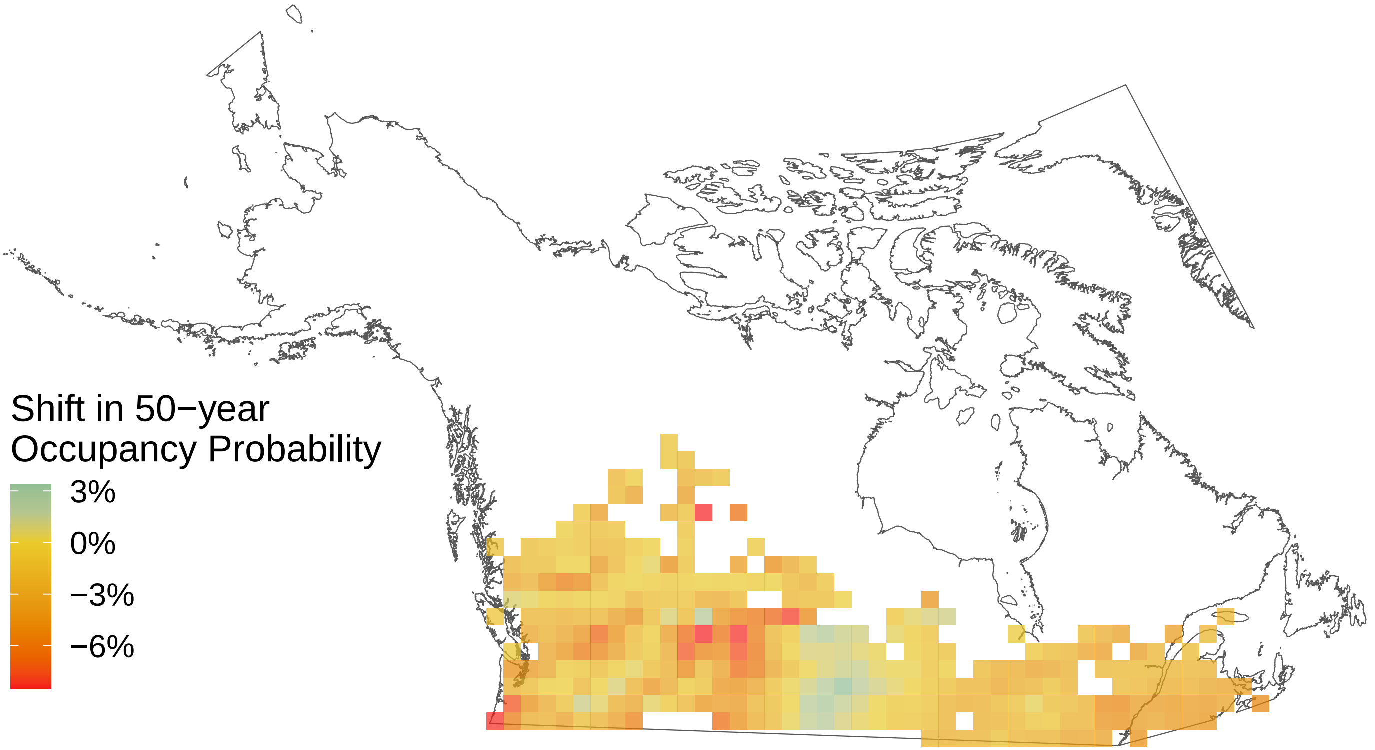

*Ancyloxypha numitor*

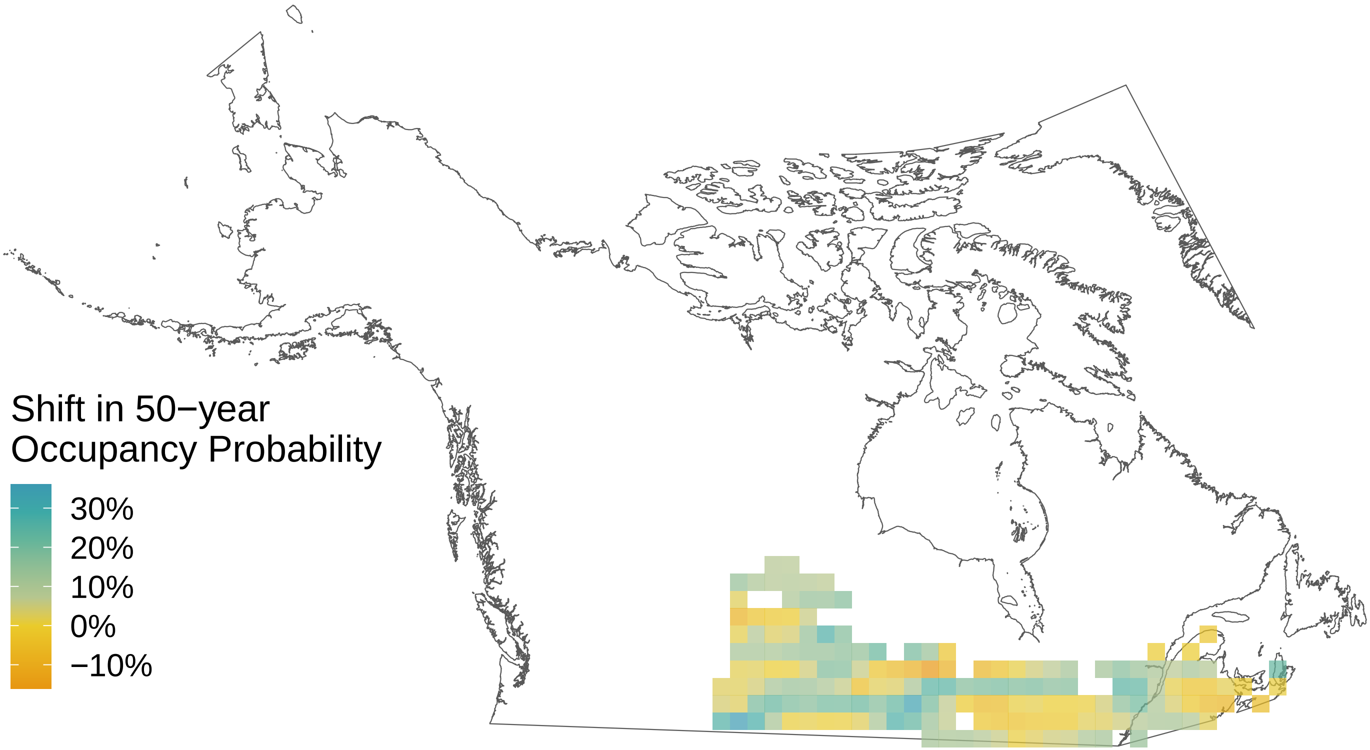

*Agriades glandon*

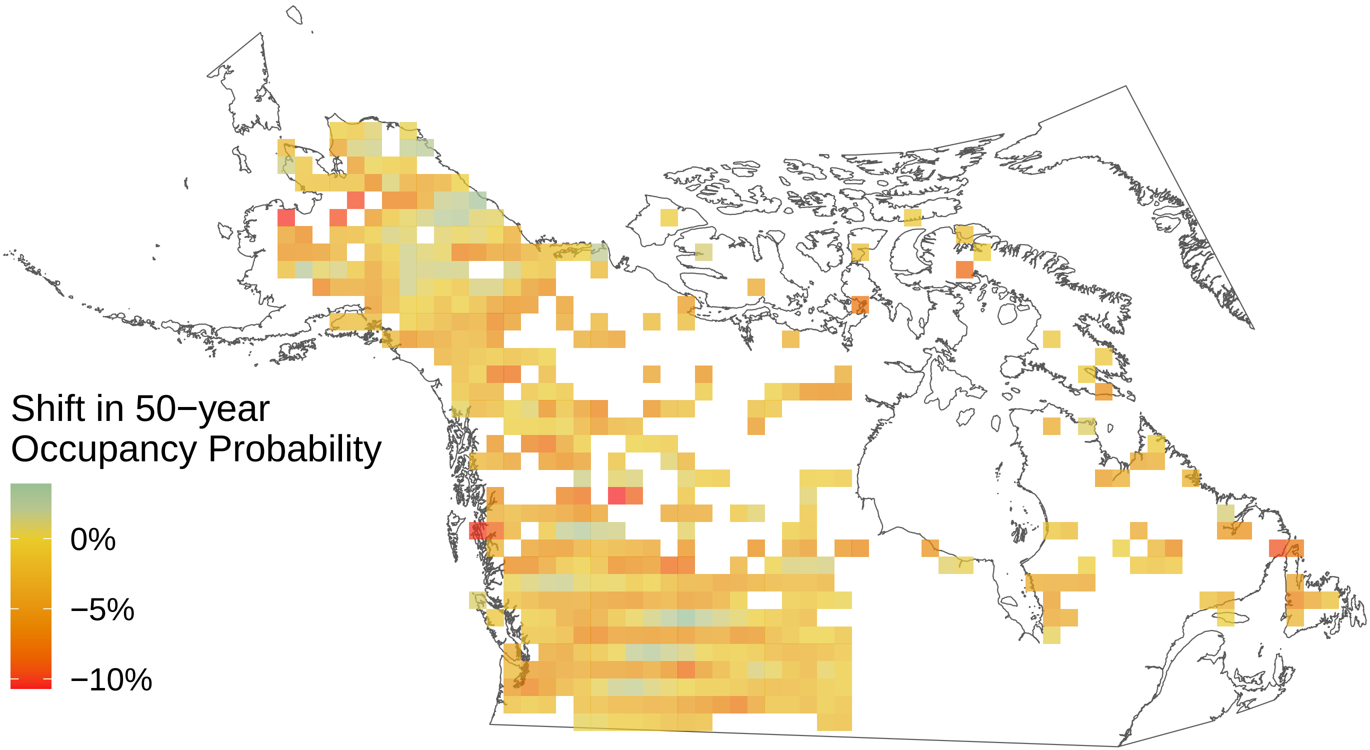

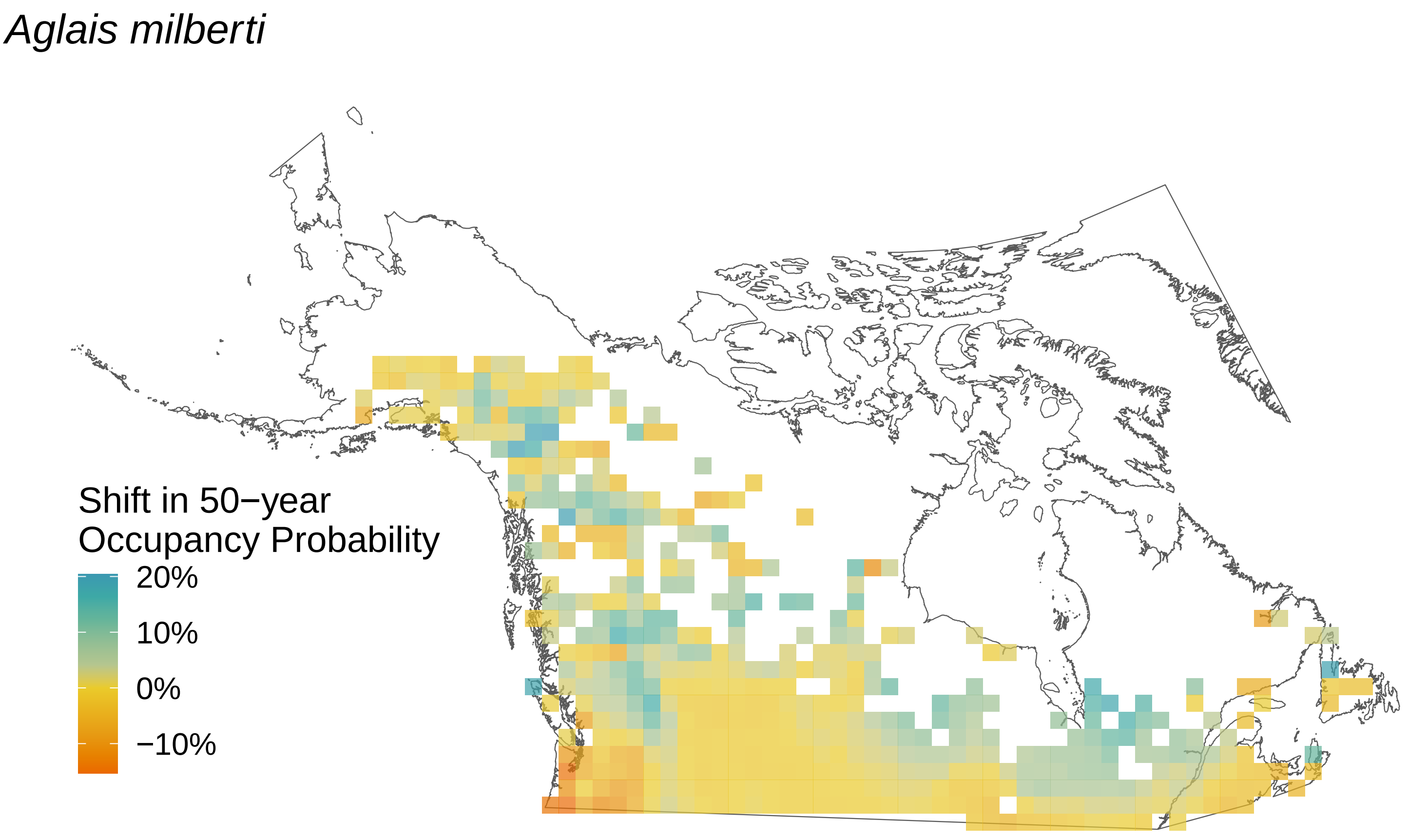

Aglais milberti
Shift in 50−year Occupancy Probability
20%
10%
0%
−10%
