## Supplemental File S7 for "Rising minimum temperatures contribute to 50 years of shifting Arctic and boreal butterfly communities in North America"

### *Vanessa virginiensis*

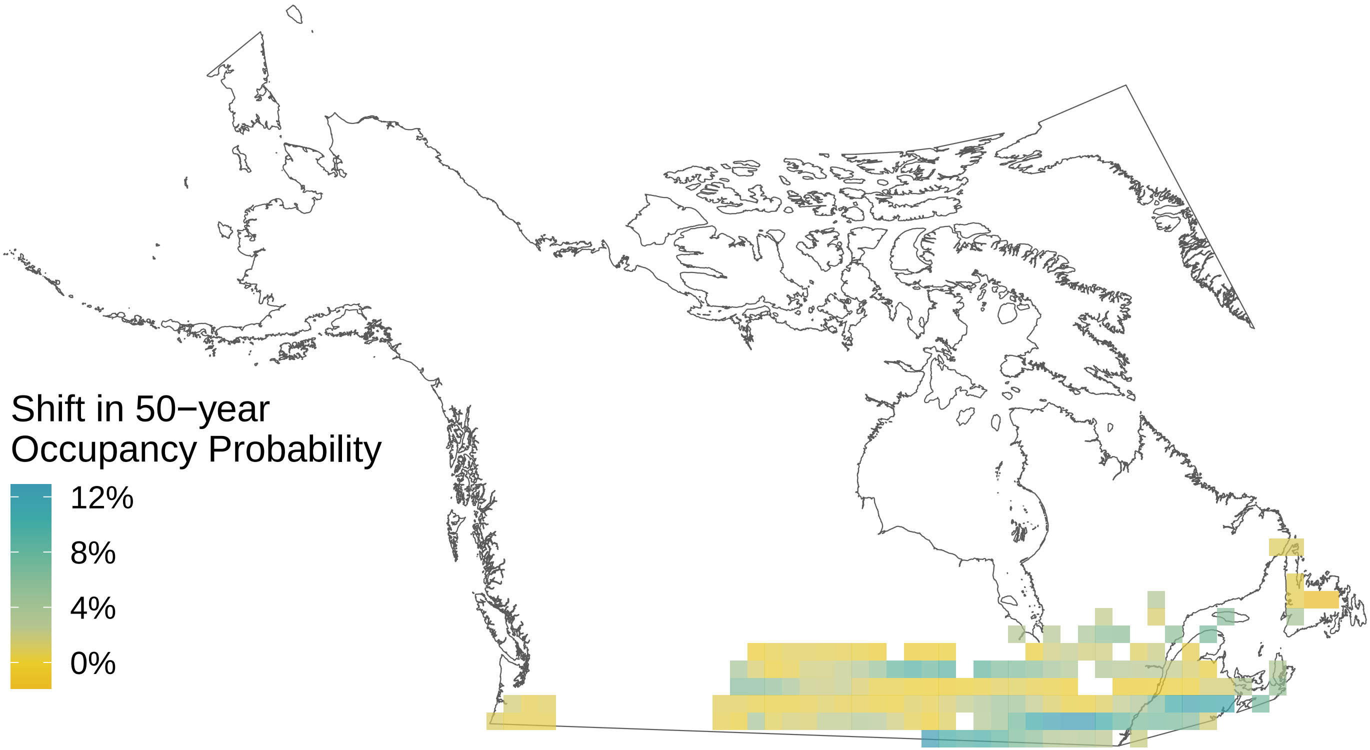

*Thymelicus lineola*

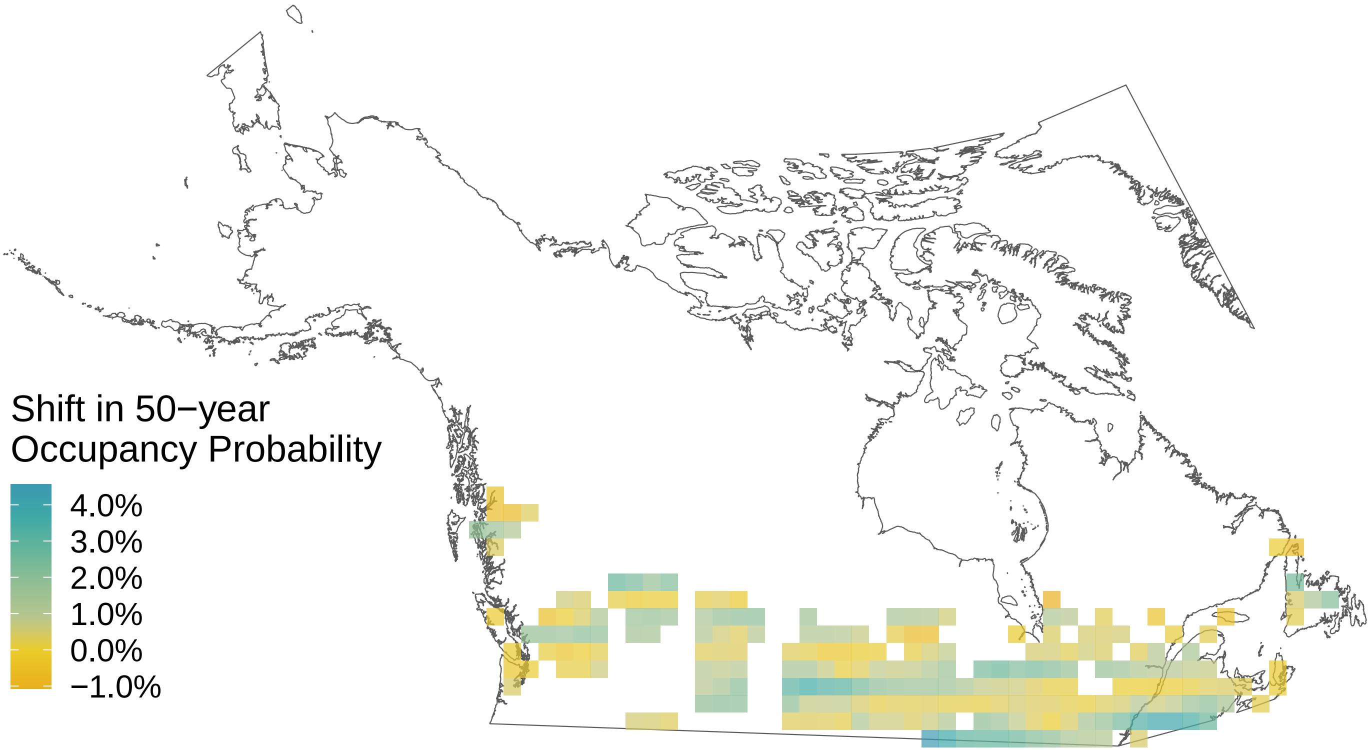

*Thorybes pylades*

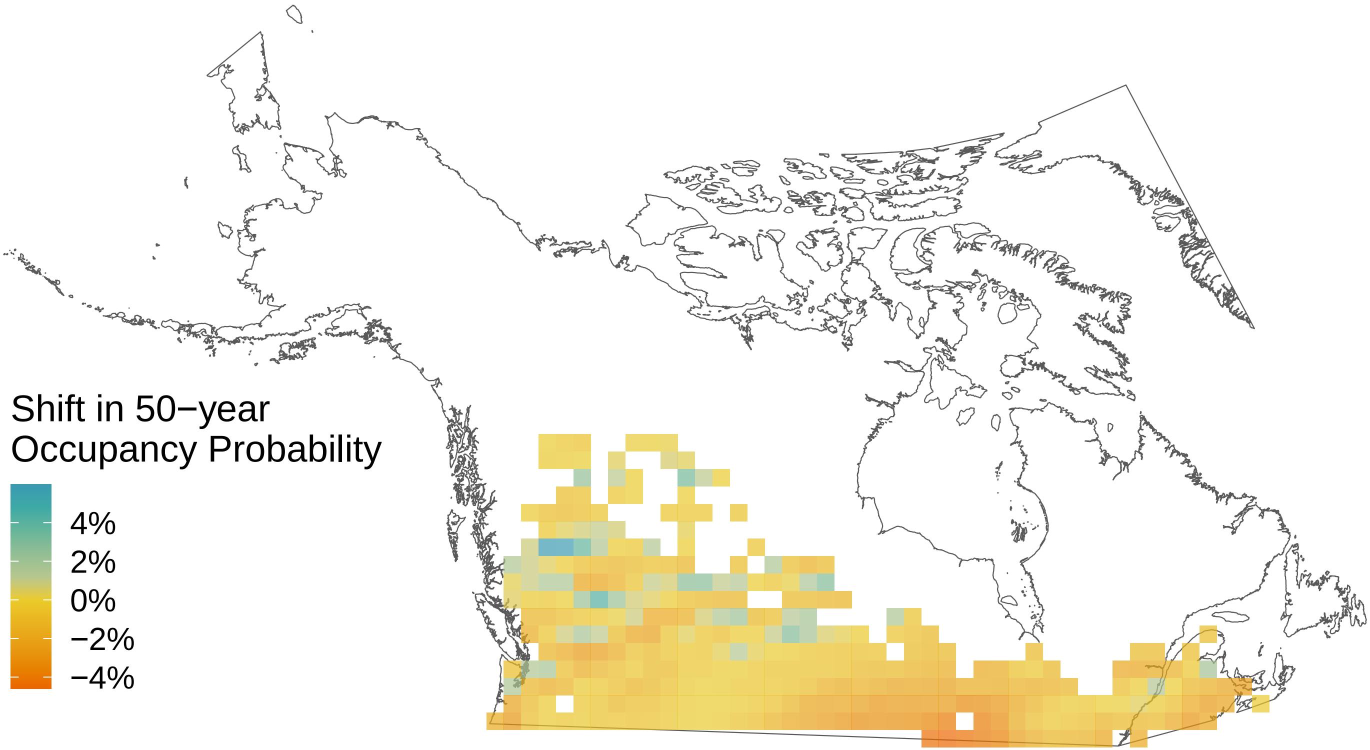

*Strymon melinus*

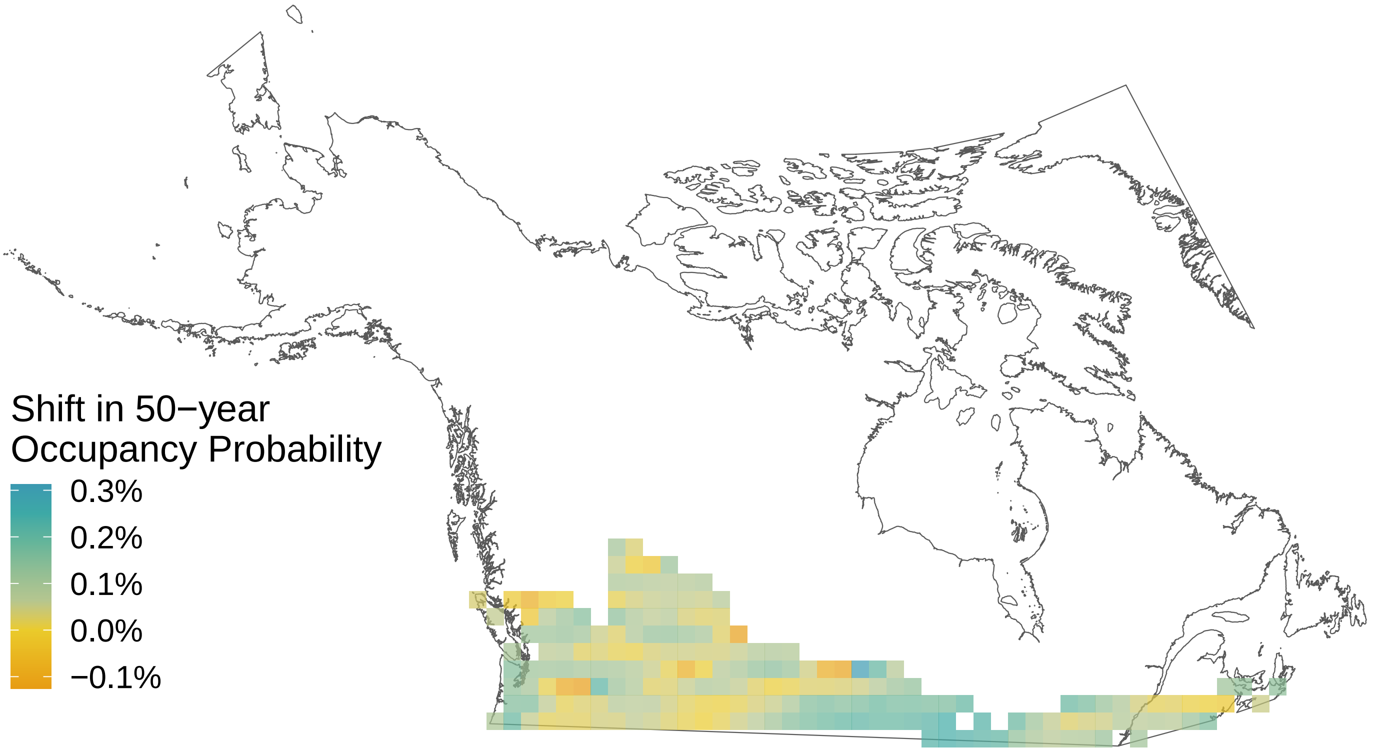

*Satyrium liparops*

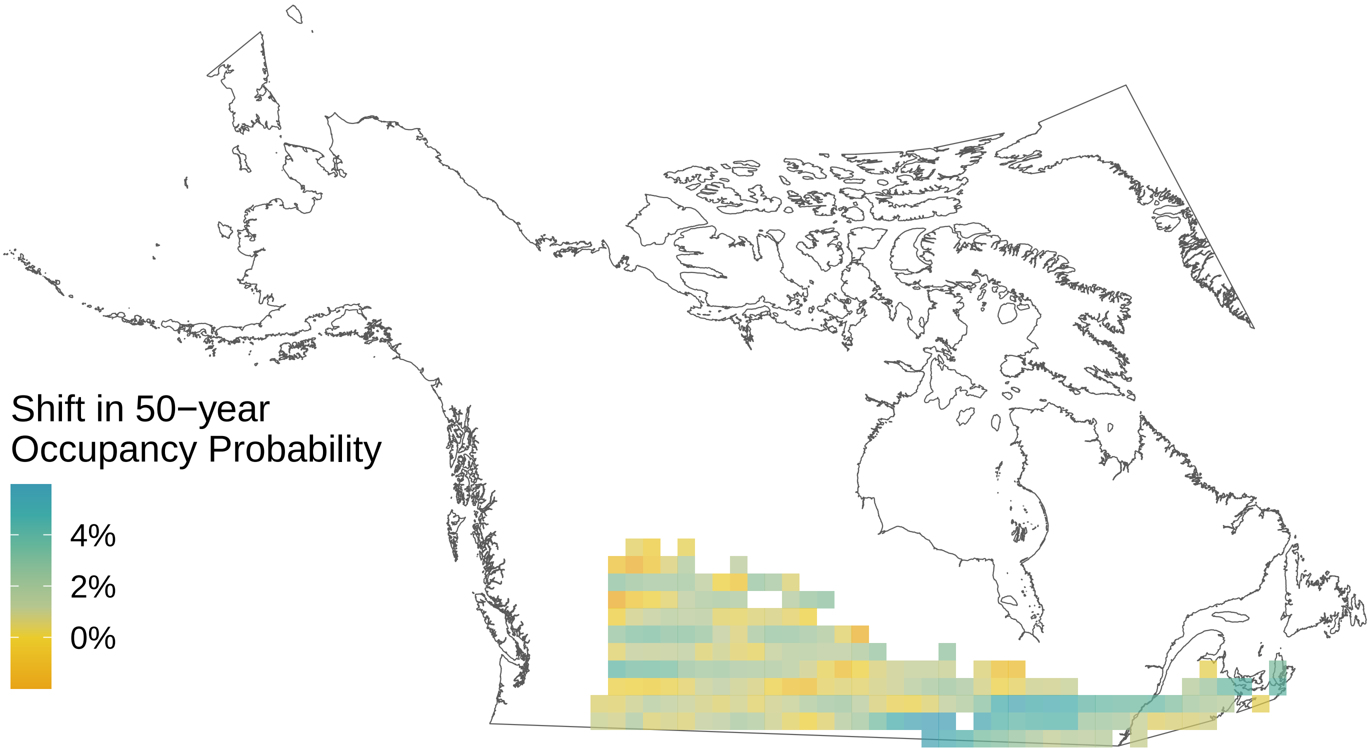

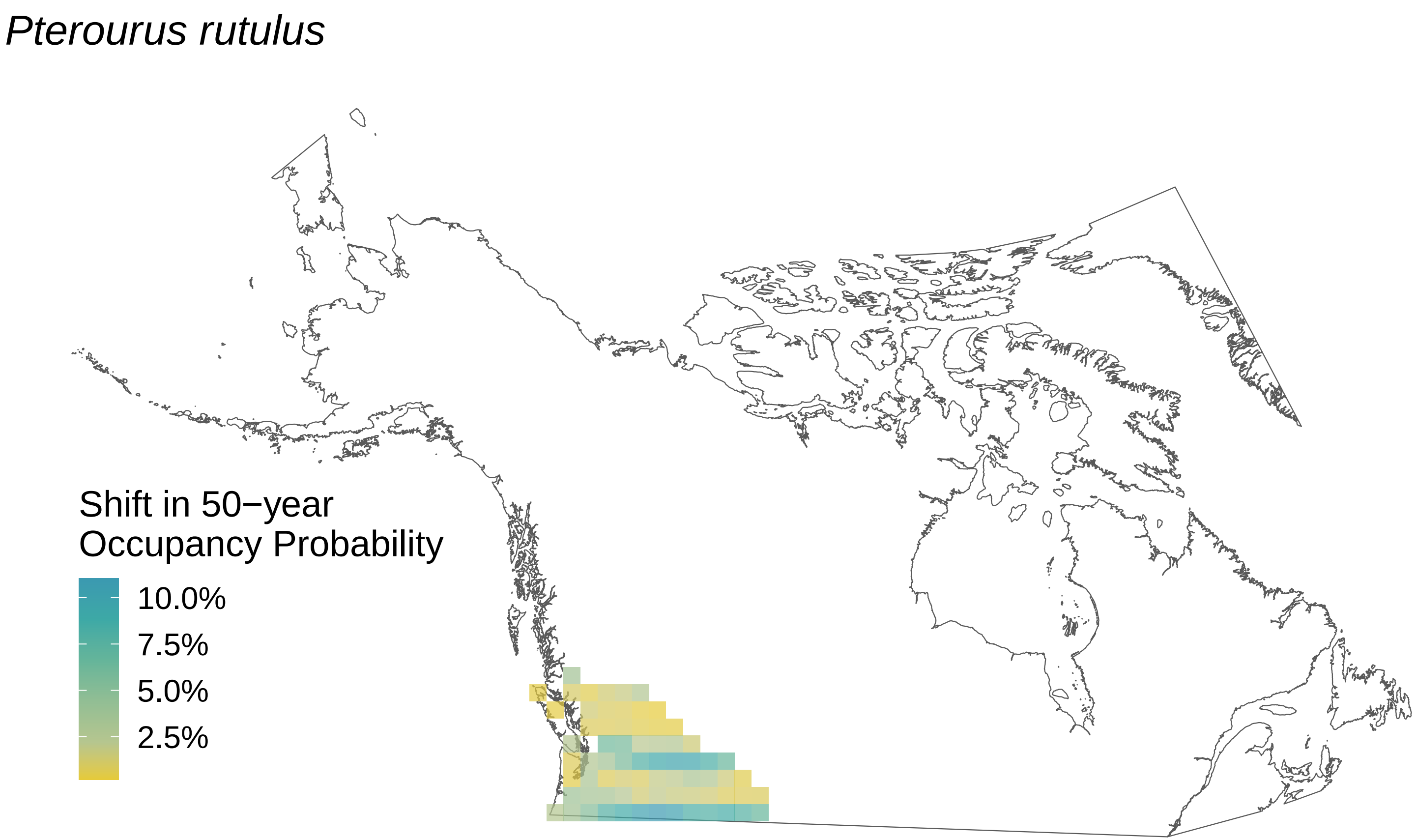
*Pterourus rutulus*
Shift in 50−year
Occupancy Probability
10.0%
7.5%
5.0%
2.5%

*Pterourus eurymedon*

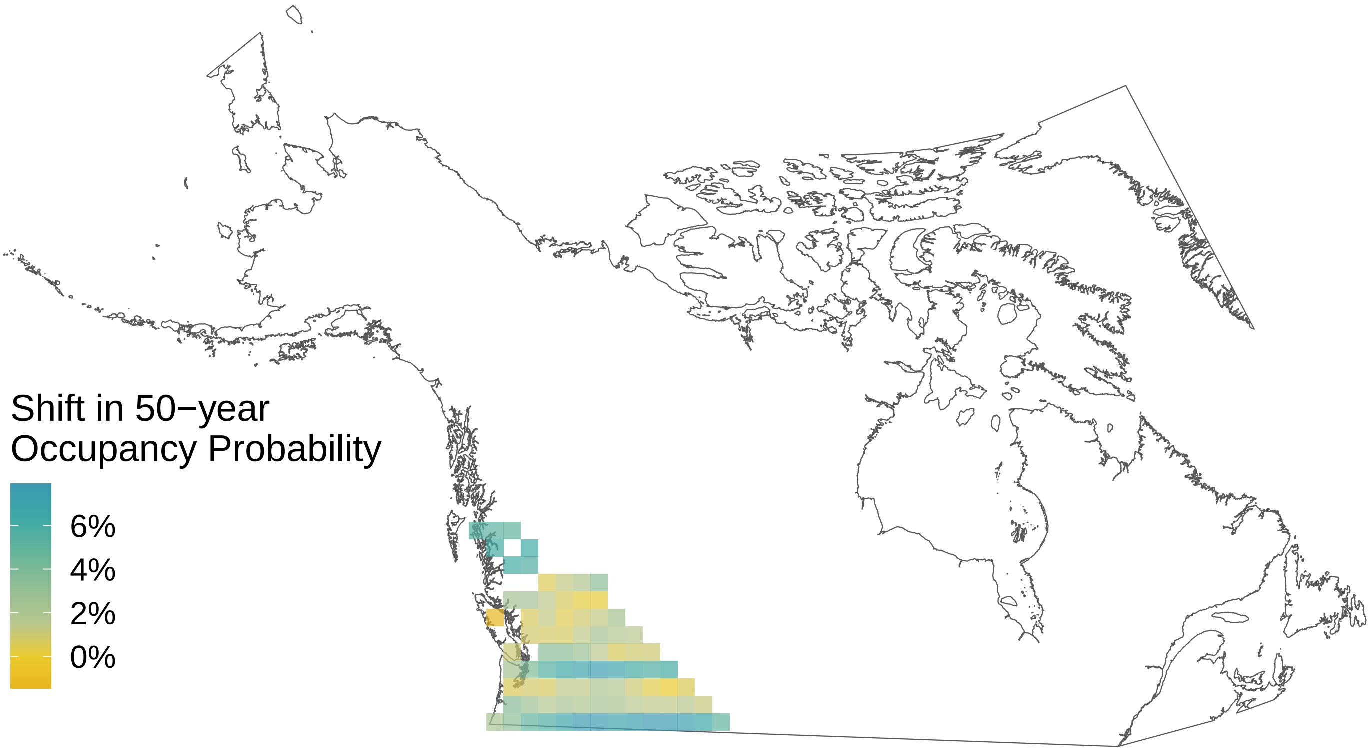

*Pterourus canadensis*

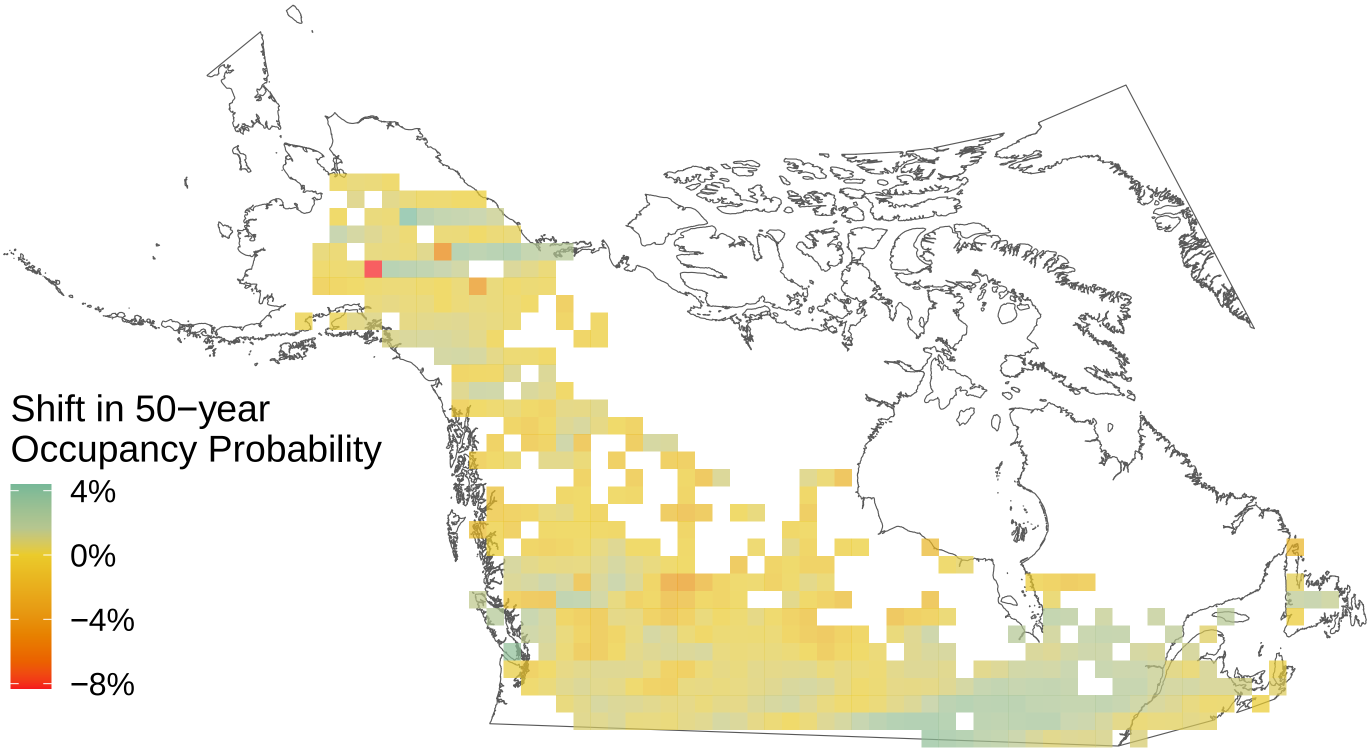

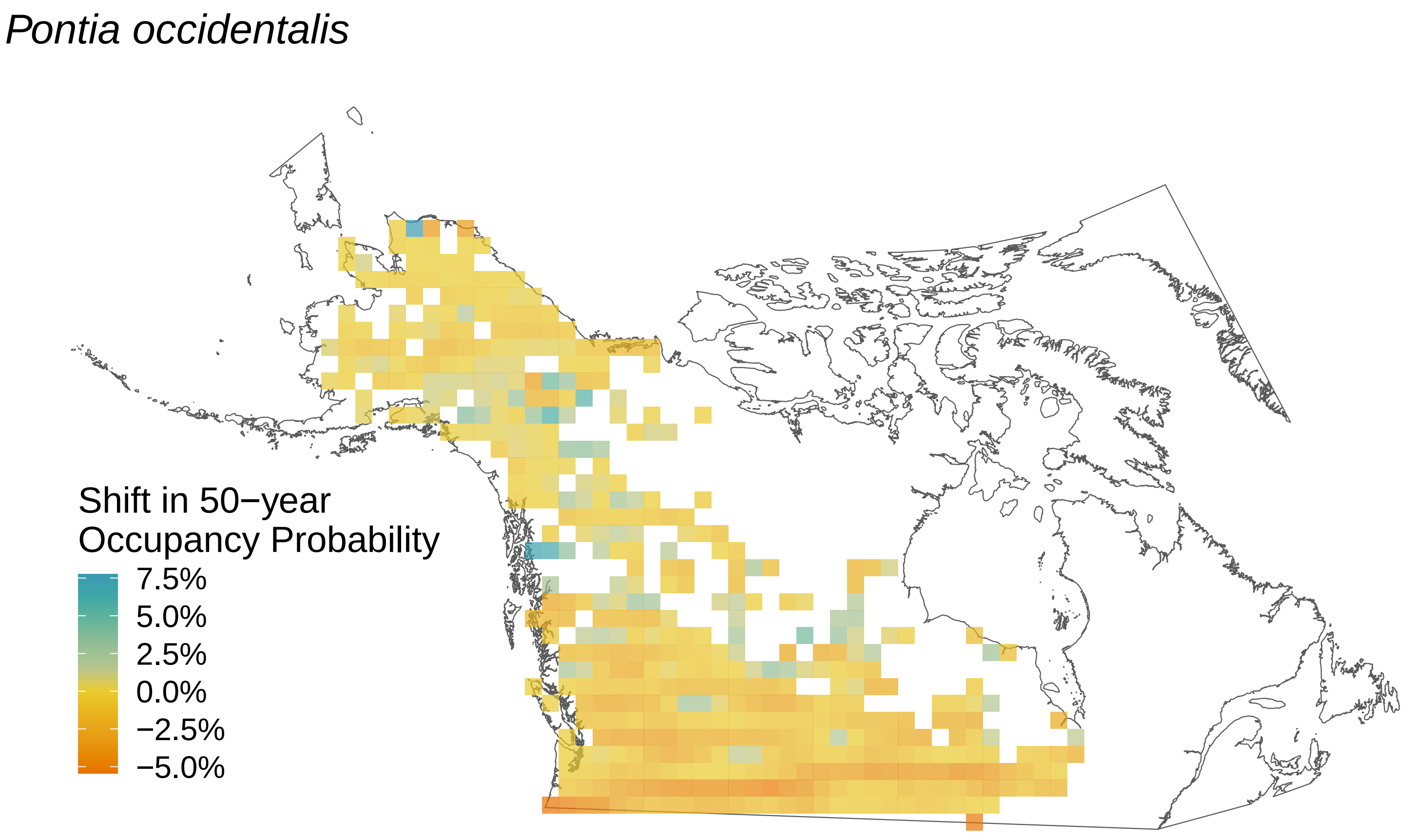

Pontia occidentalis
Shift in 50−year
Occupancy Probability
7.5%
5.0%
2.5%
0.0%
−2.5%
−5.0%

*Polygonia satyrus*

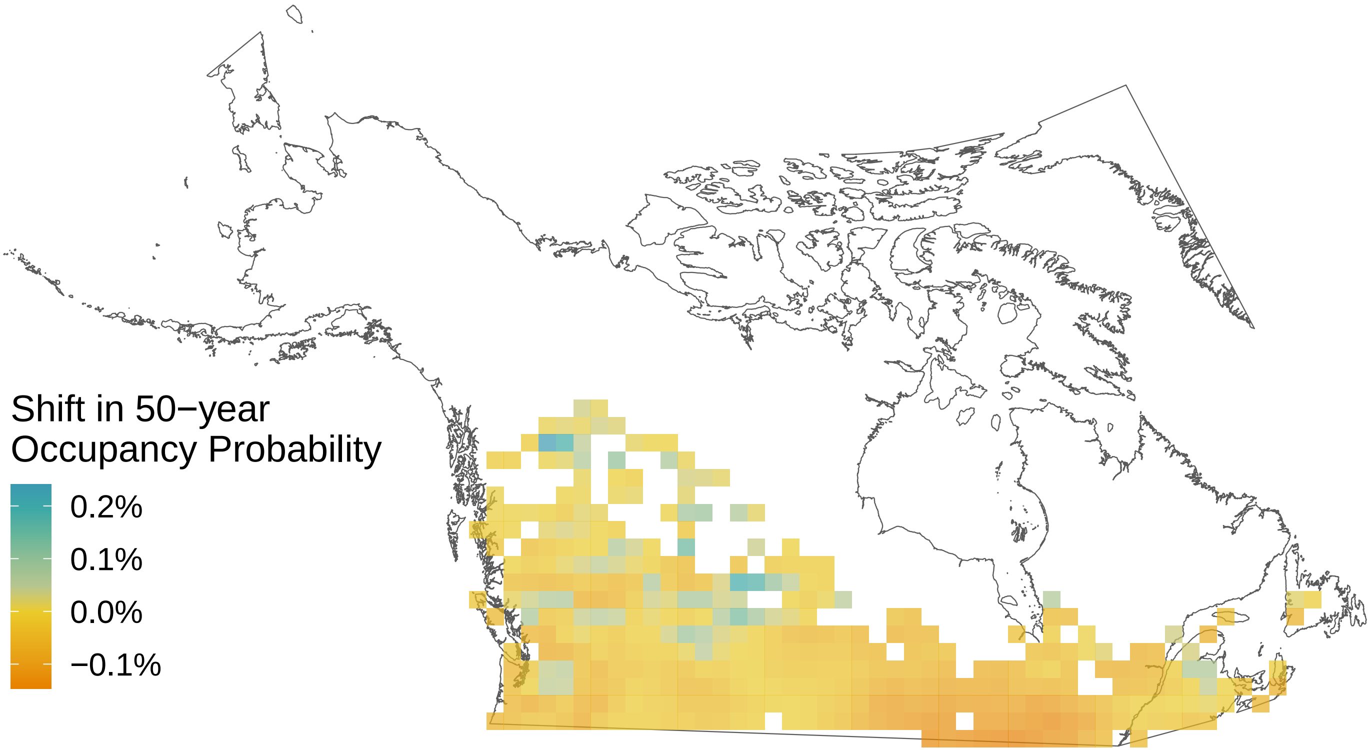

*Polygonia progne*

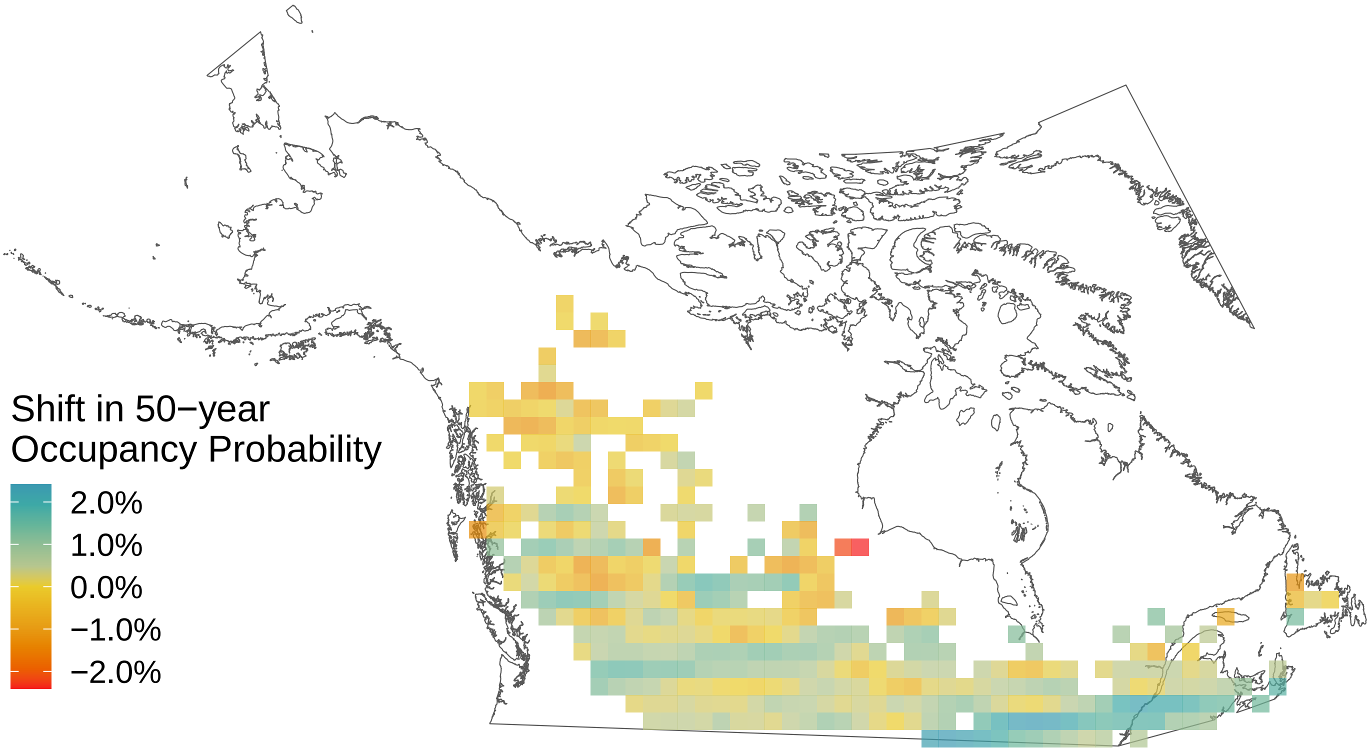

*Polygonia interrogationis*

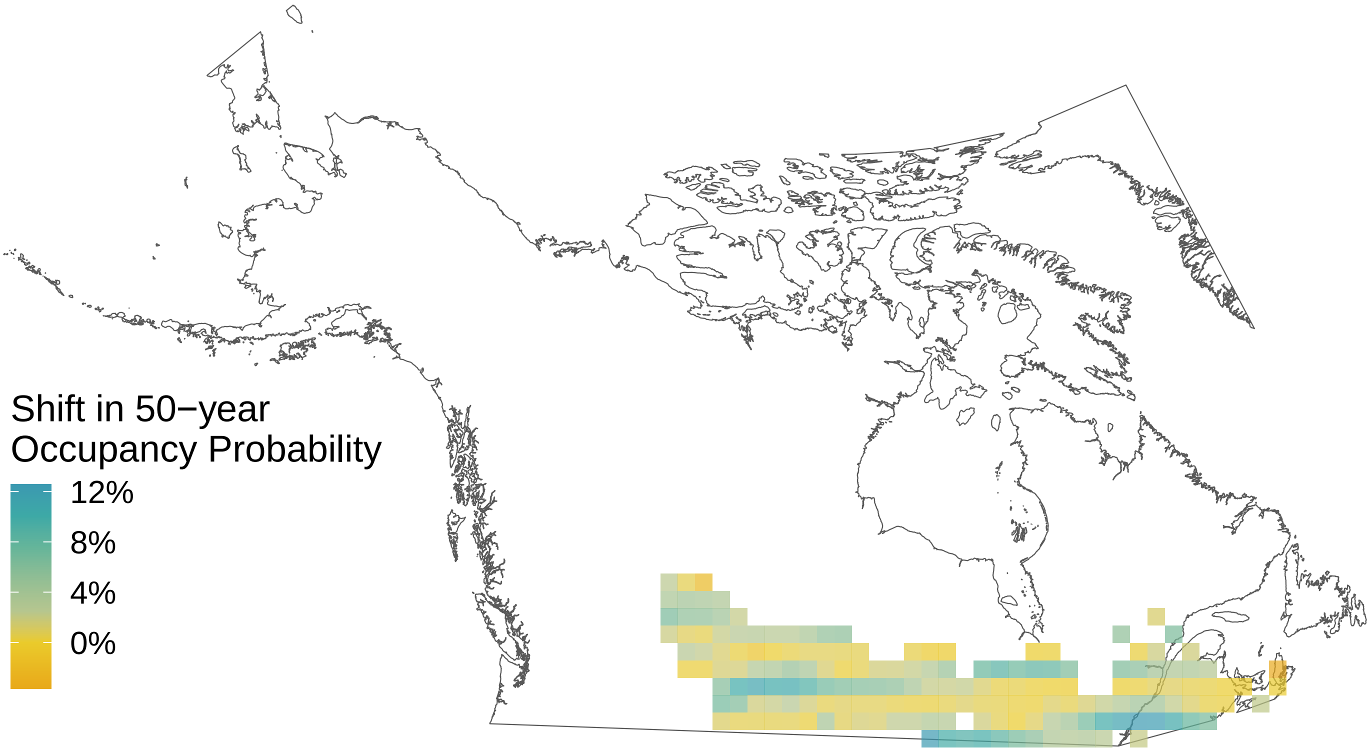

*Polygonia gracilis*

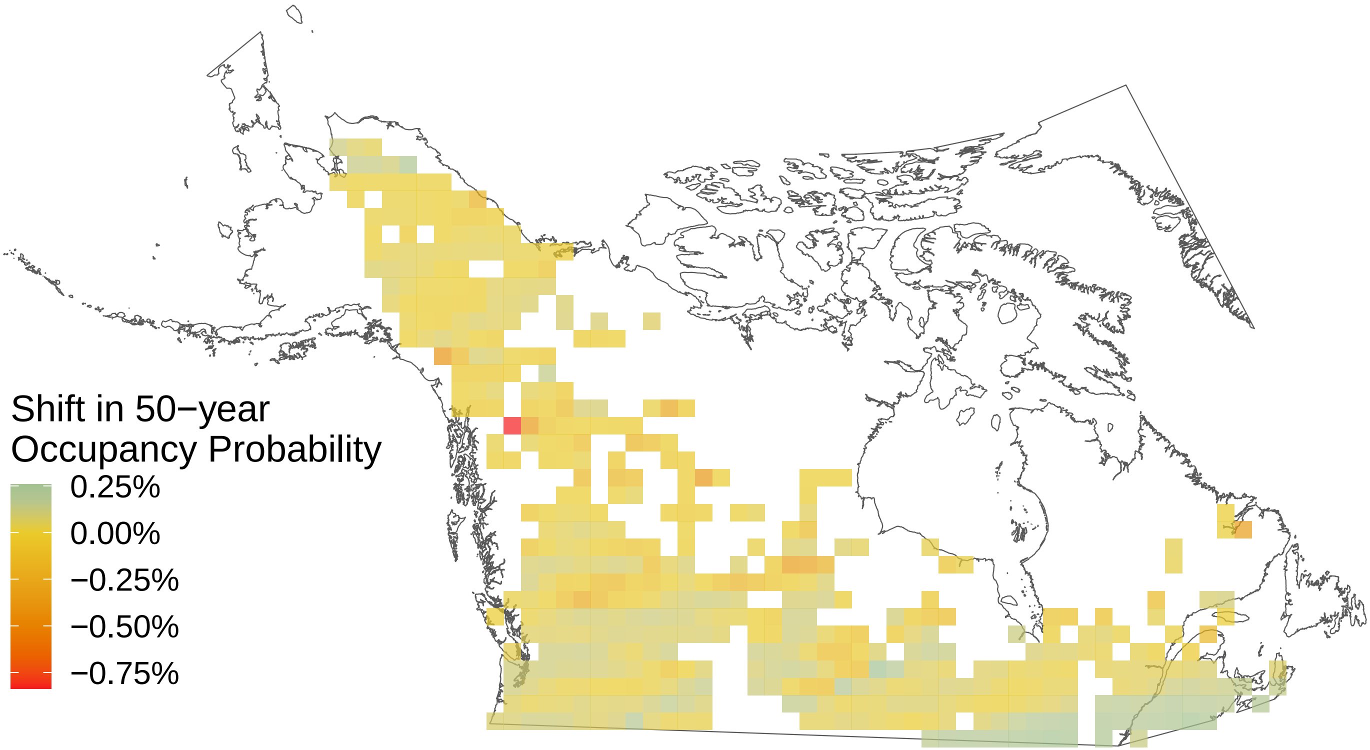

*Polygonia faunus*

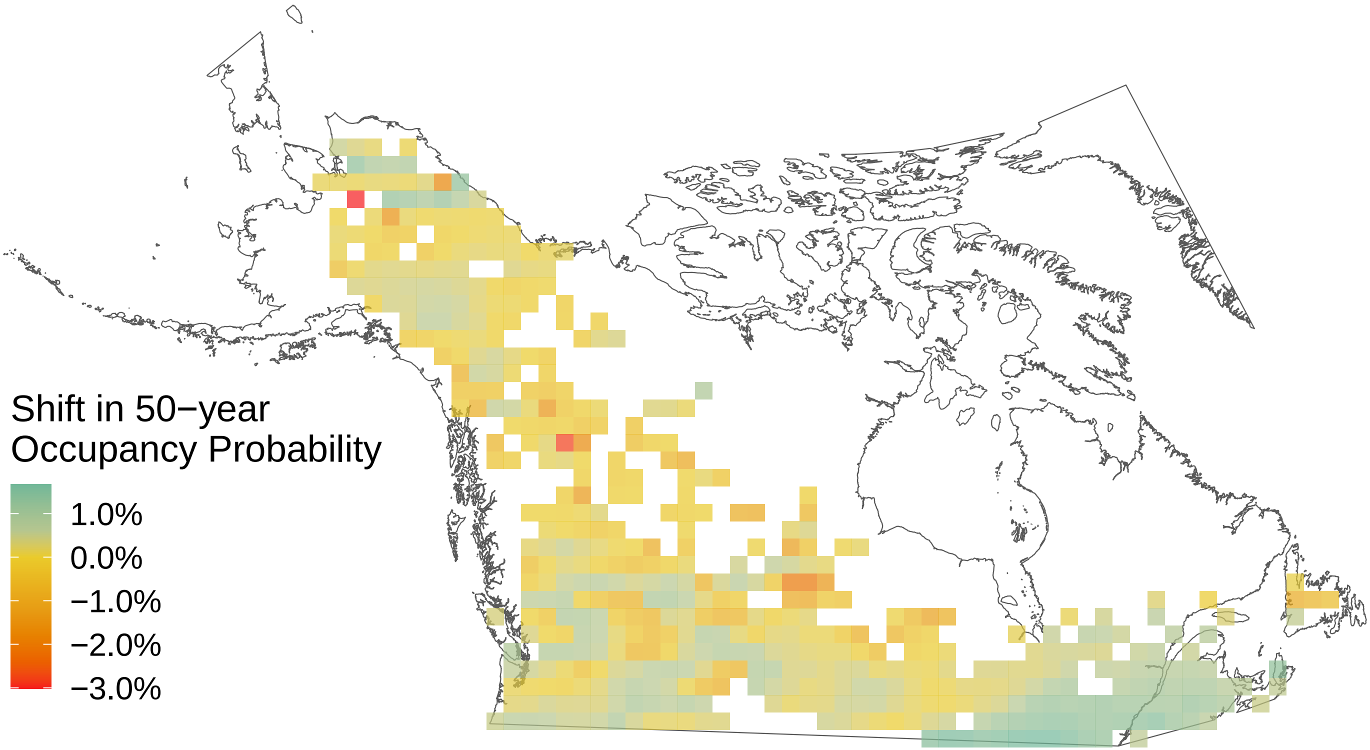

*Polites themistocles*

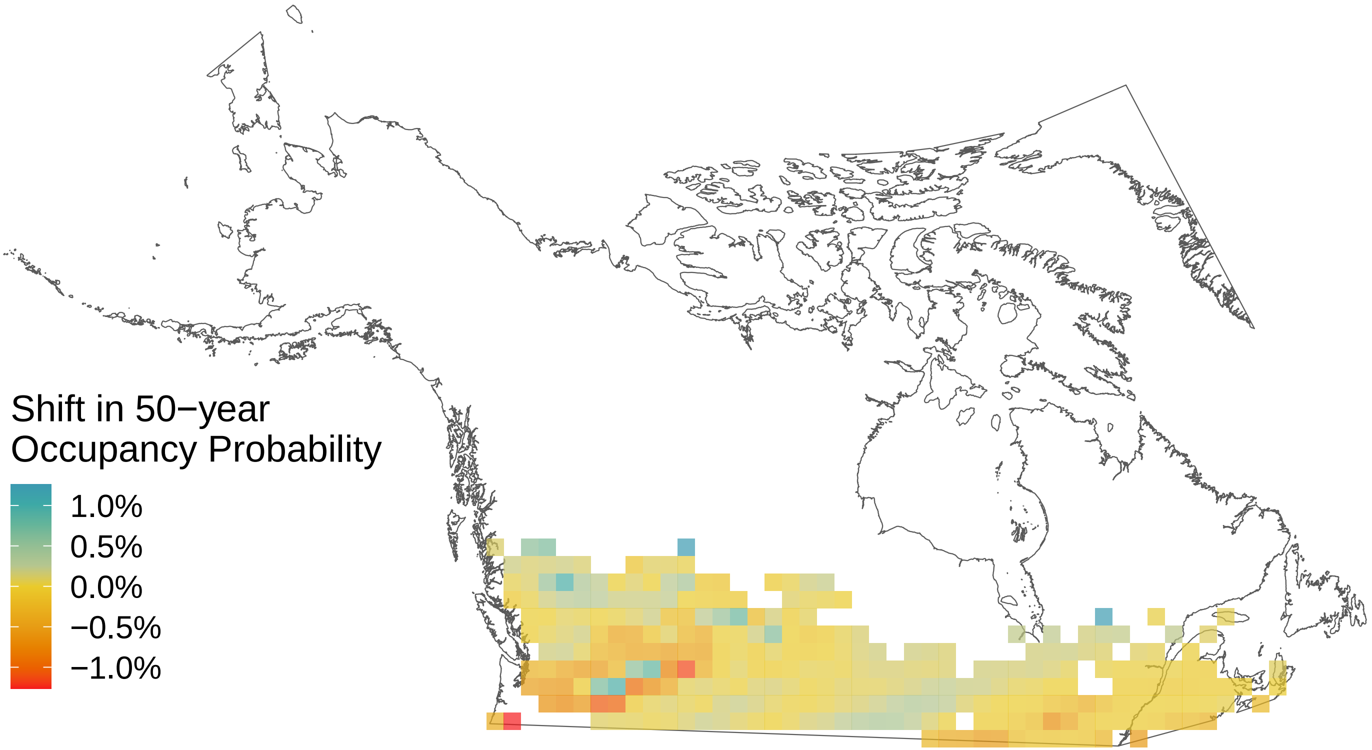

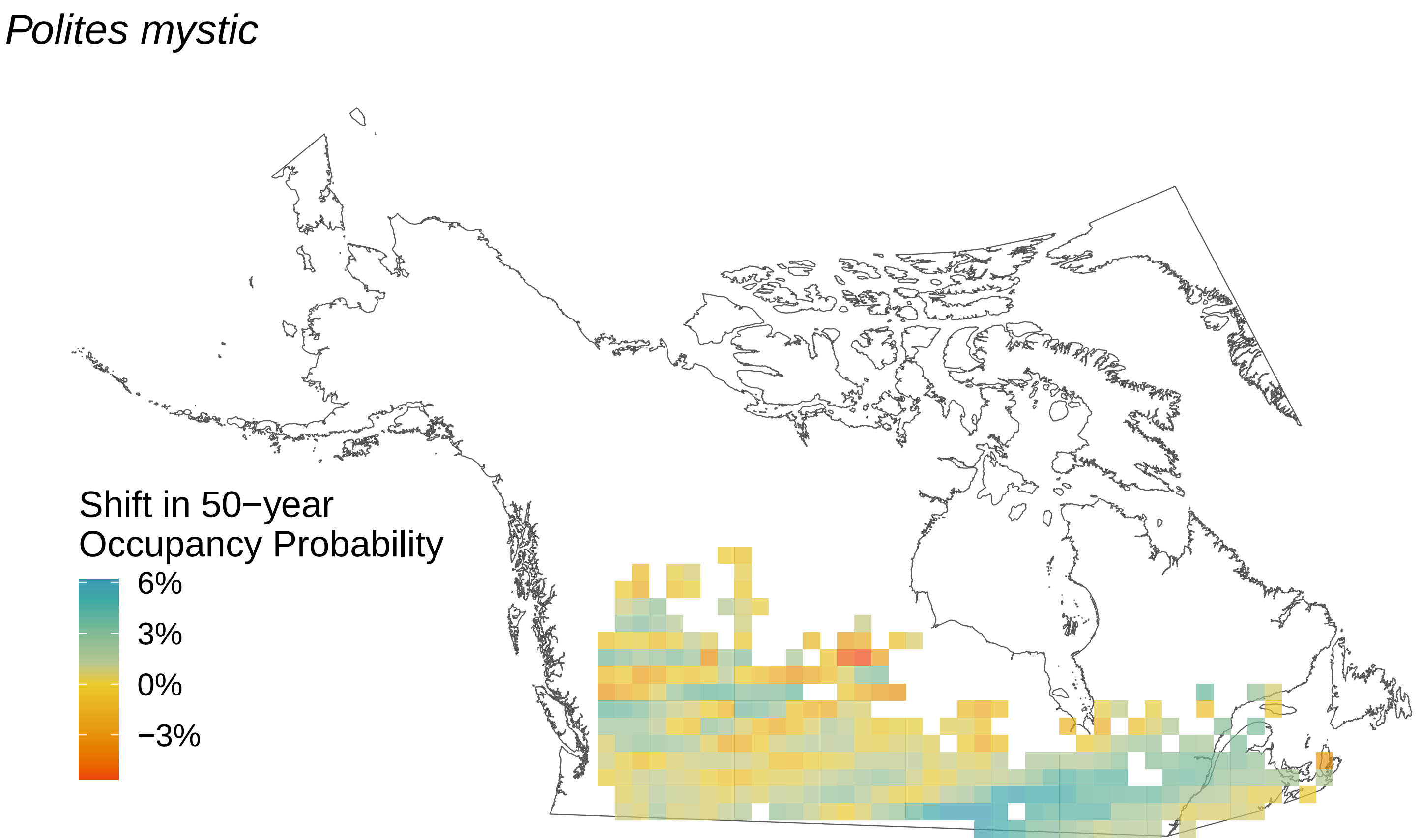
Polites mystic
Shift in 50−year
Occupancy Probability
6%
3%
0%
−3%

*Poanes hobomok*

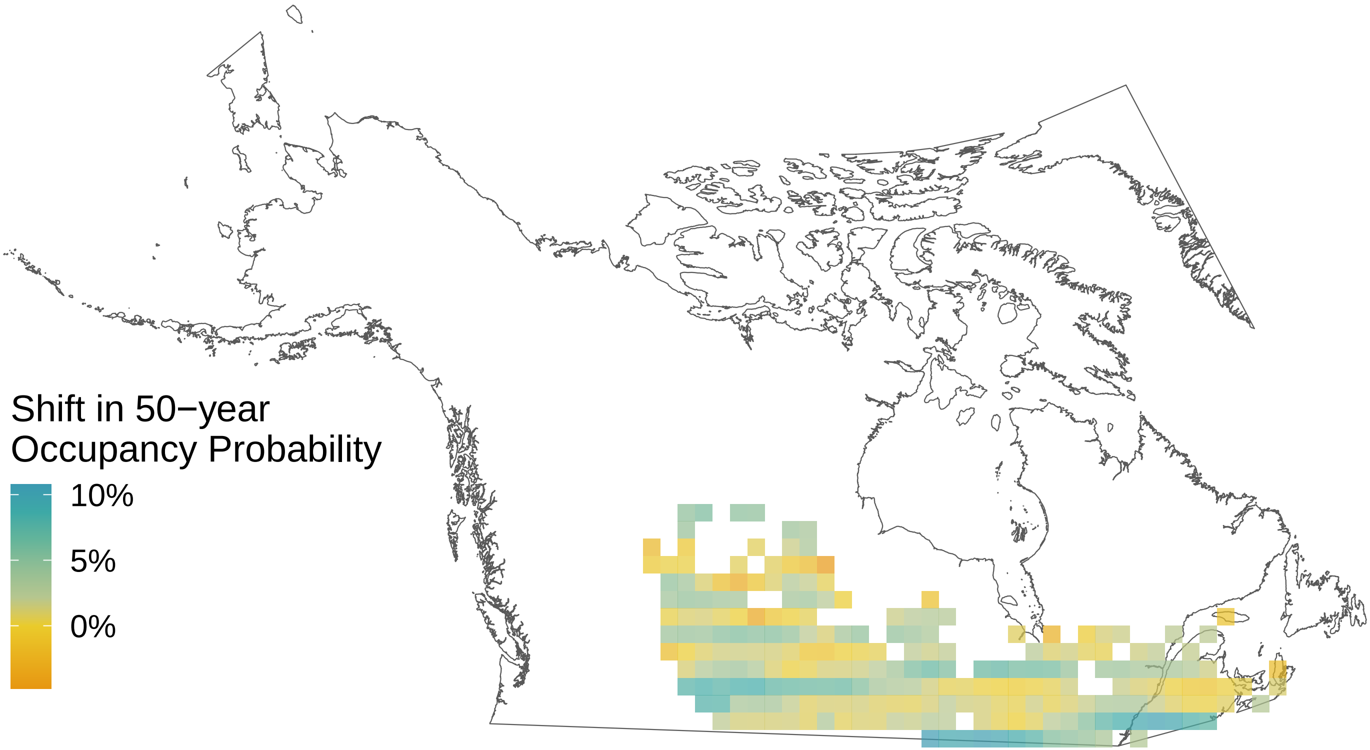

*Plebejus melissa*

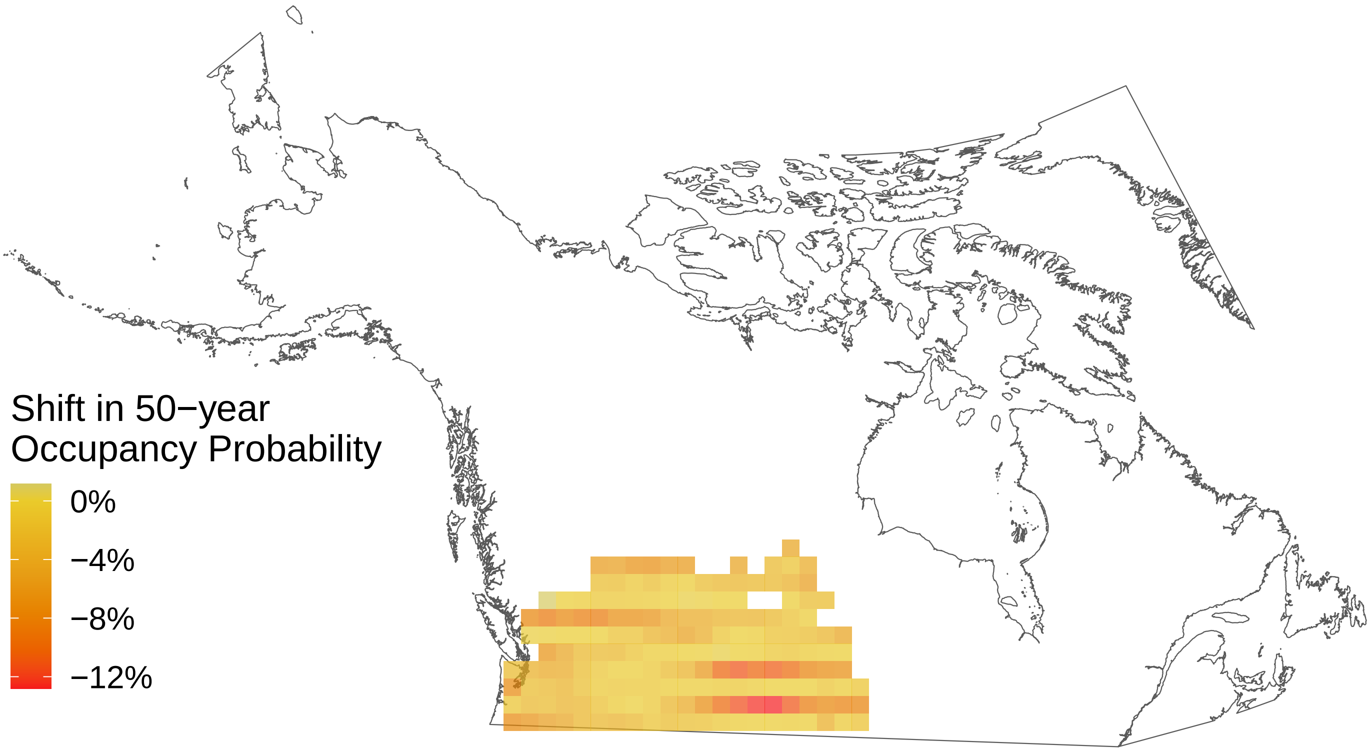

*Plebejus idas*

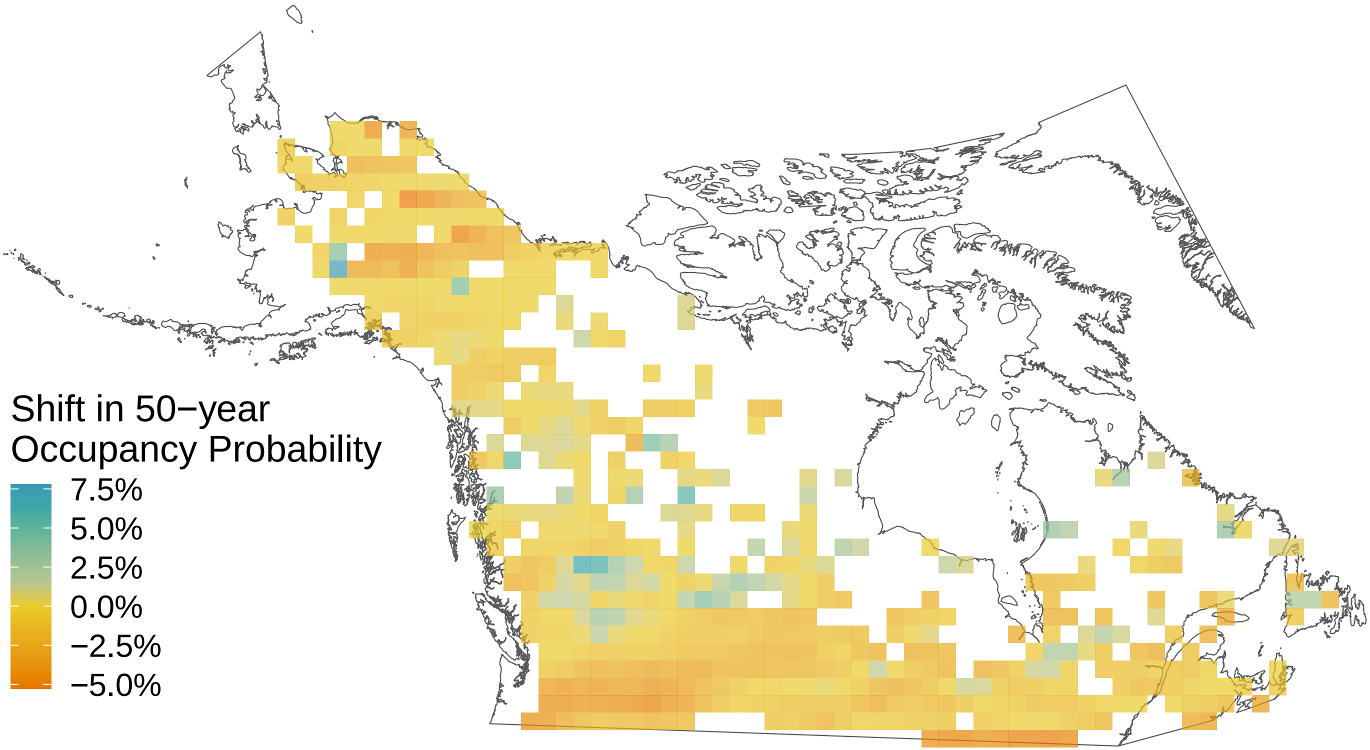

*Pieris rapae*

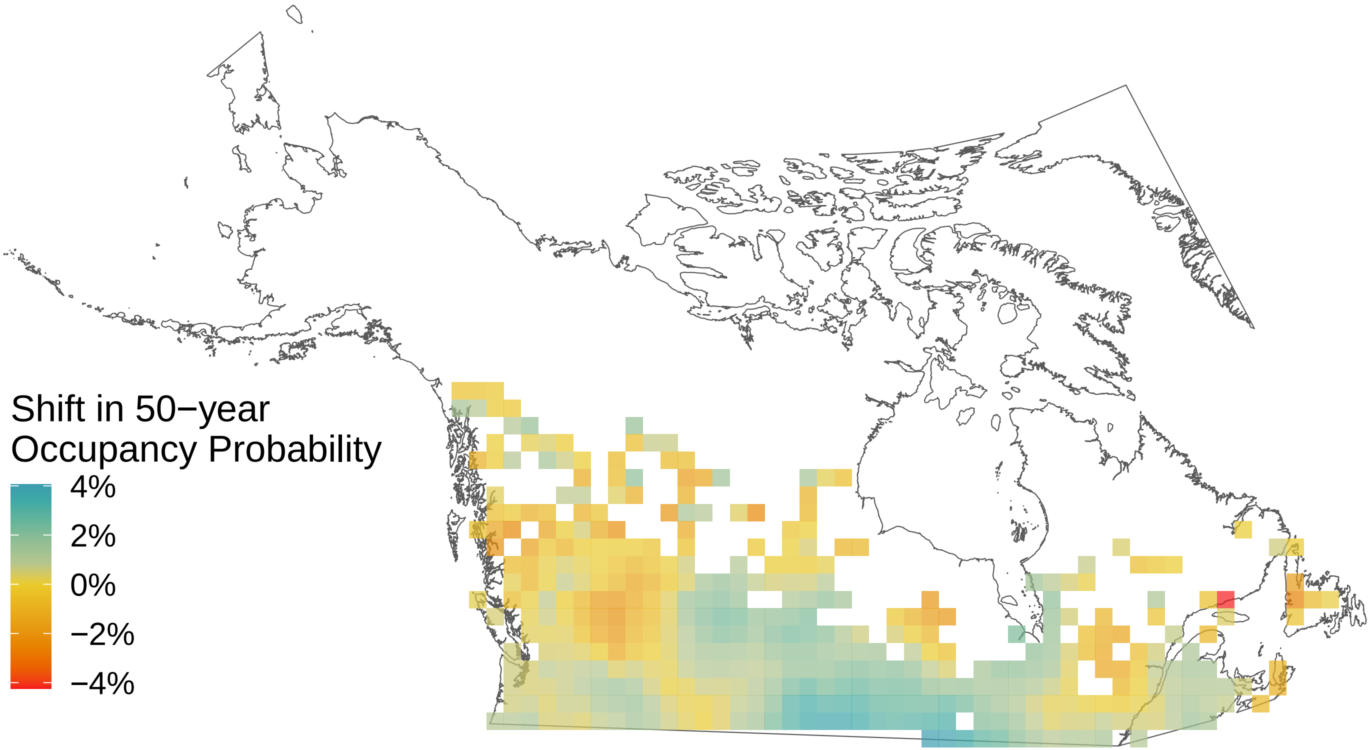

*Pieris oleracea*

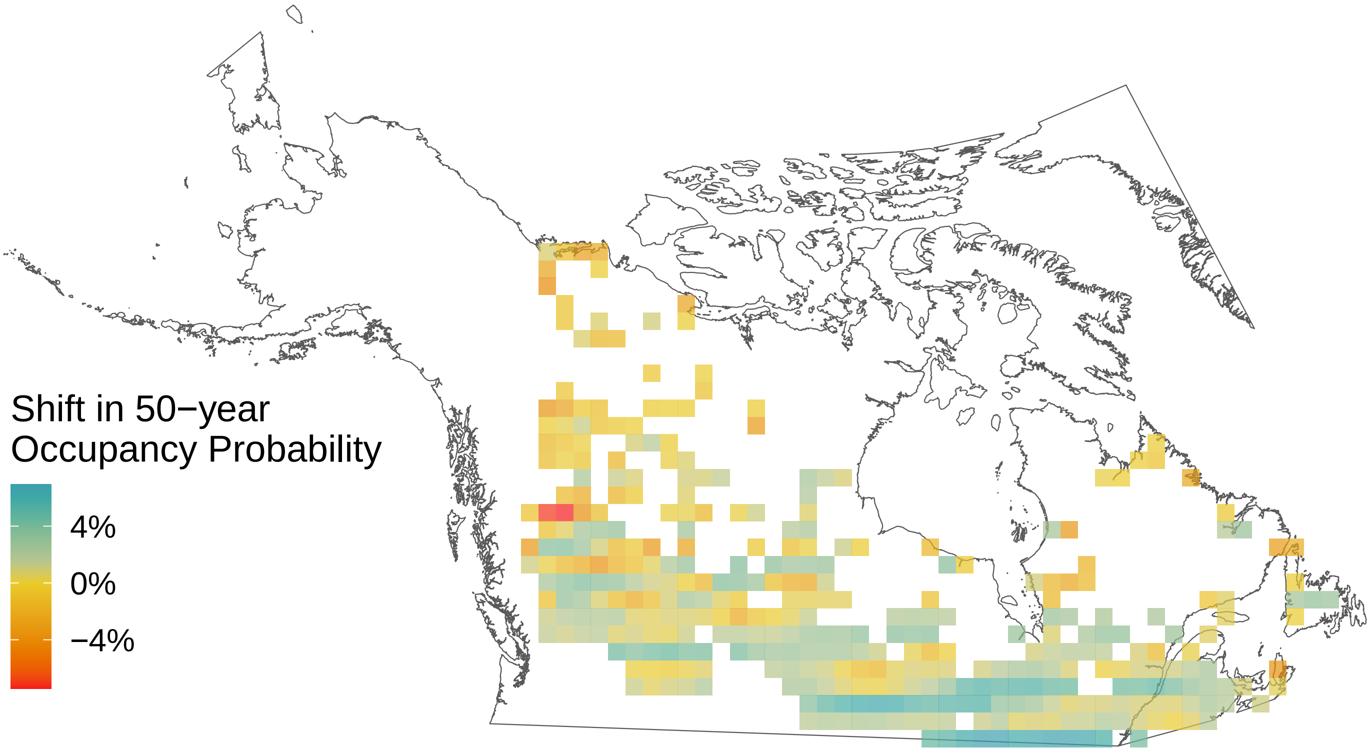

*Phyciodes tharos*

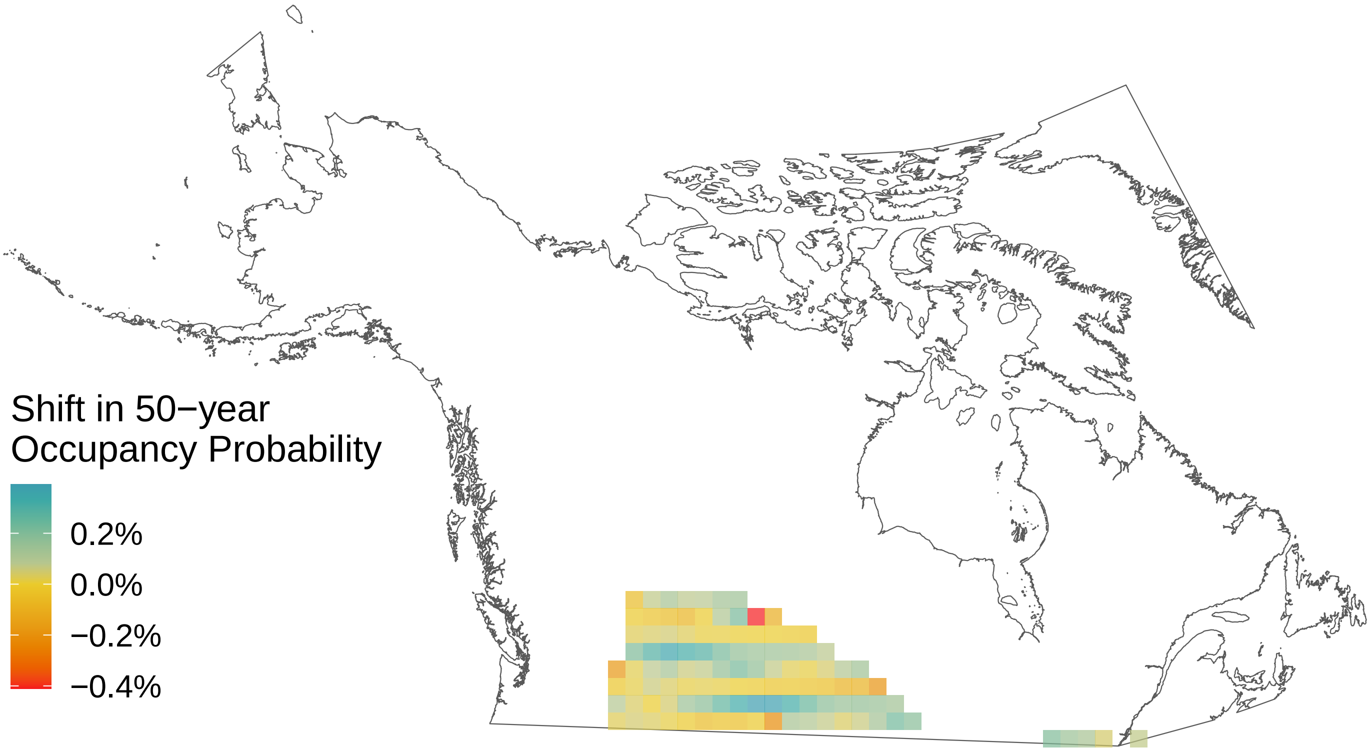

*Parnassius clodius*

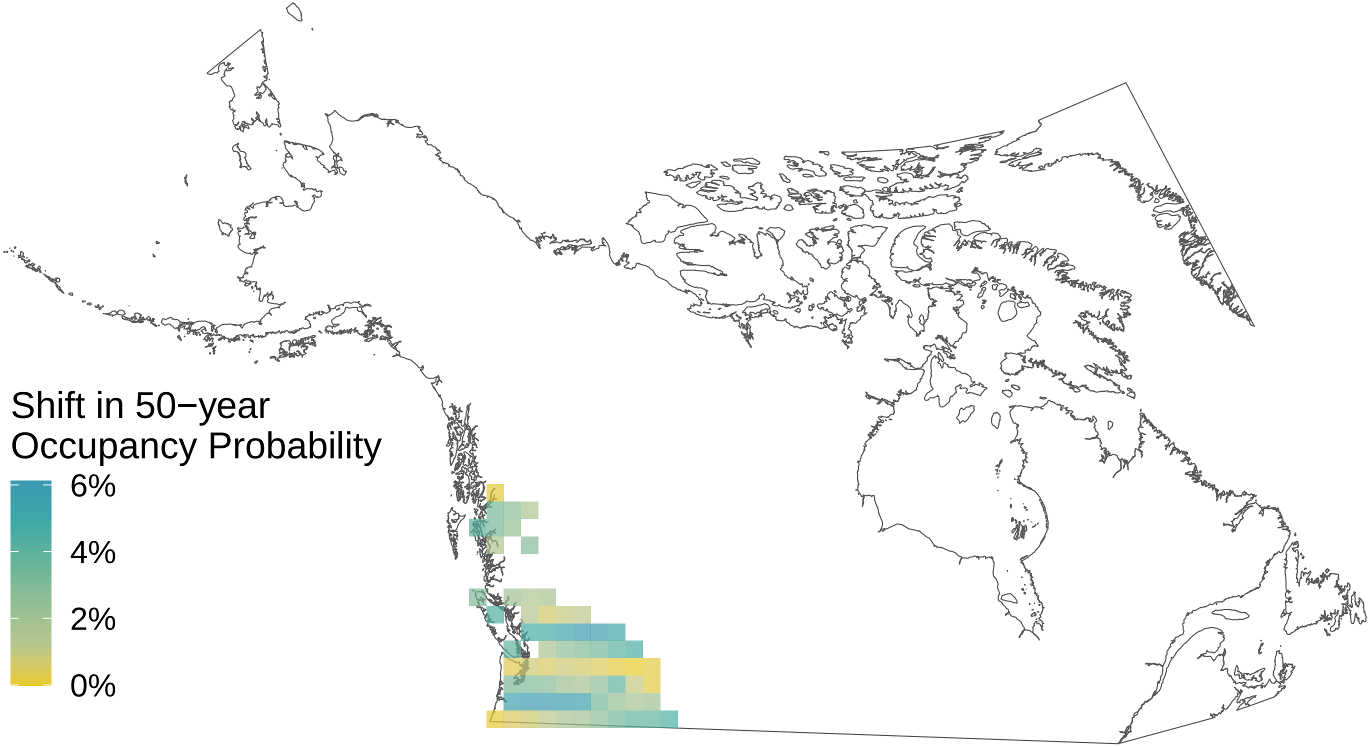

*Papilio zelicaon*

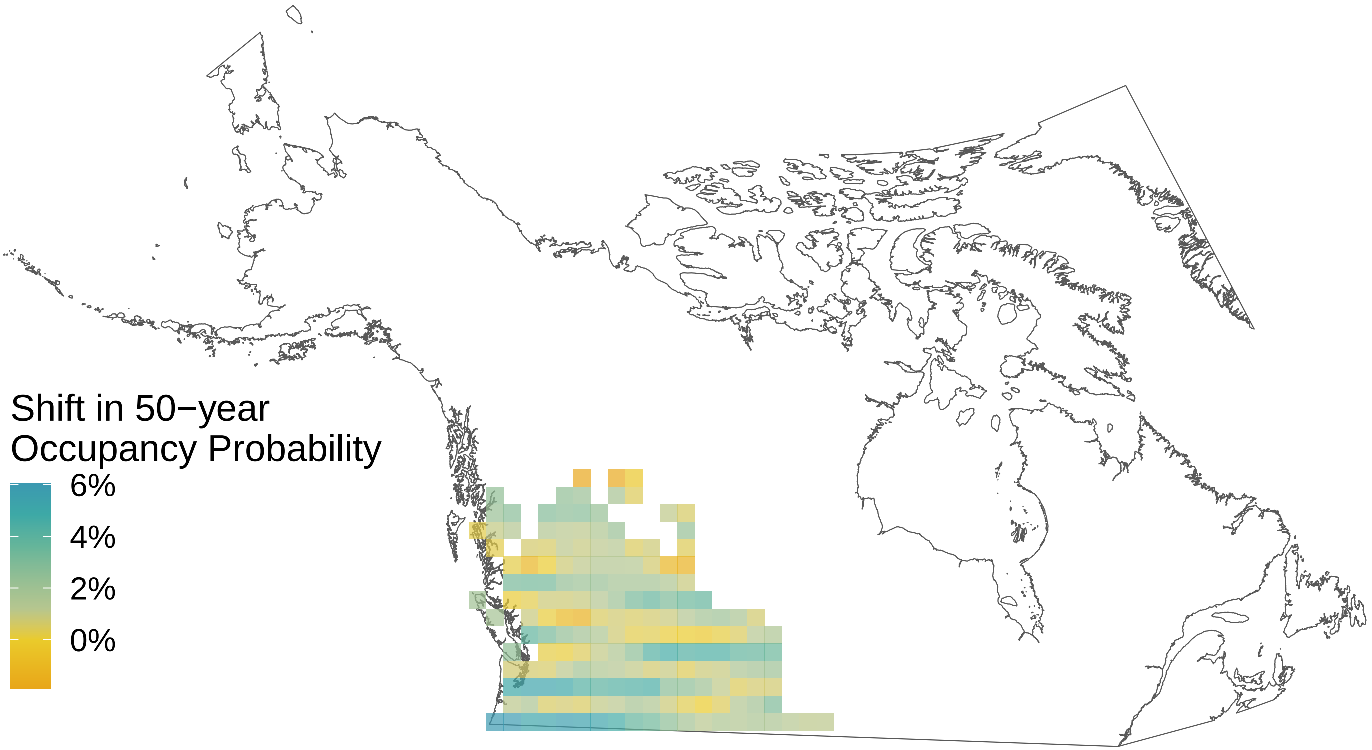

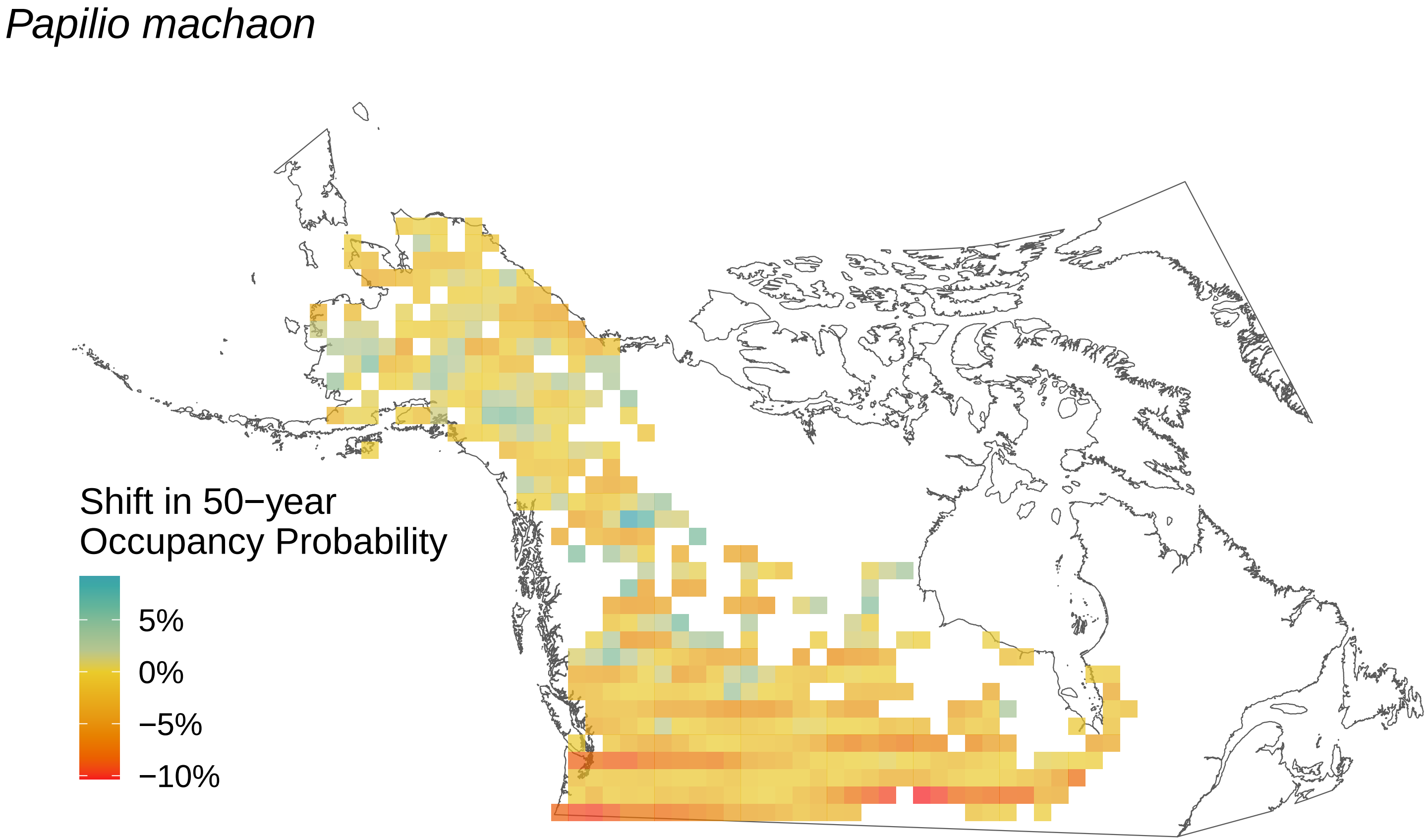

*Papilio machaon*
Shift in 50−year
Occupancy Probability
5%
0%
−5%
−10%

*Oeneis polixenes*

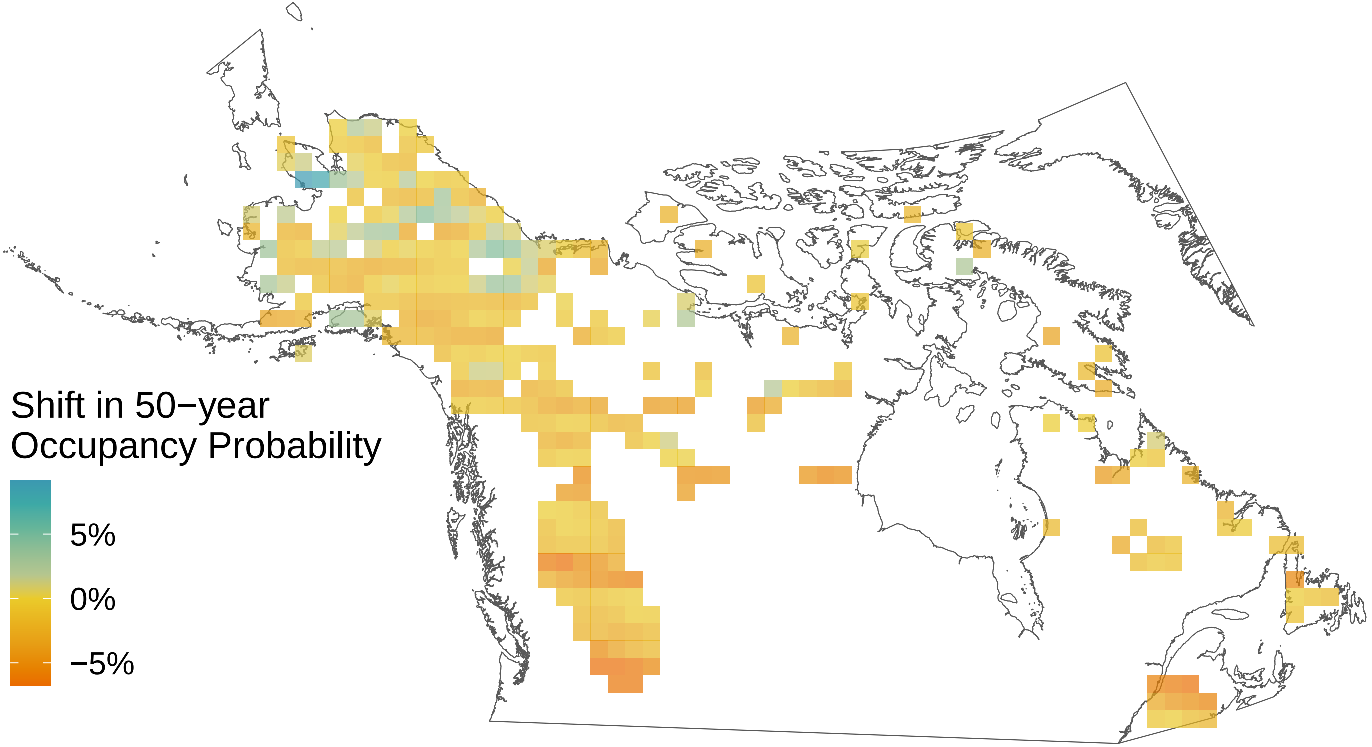

*Oeneis melissa*

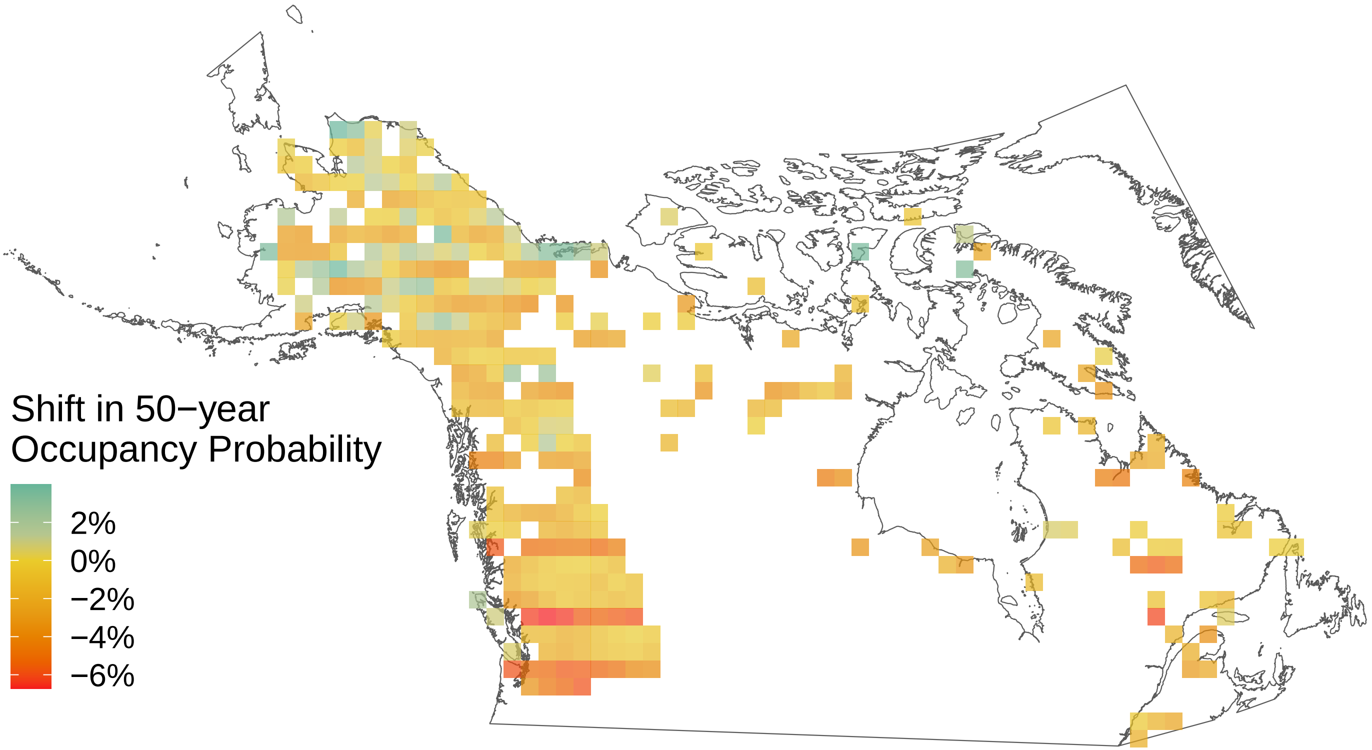

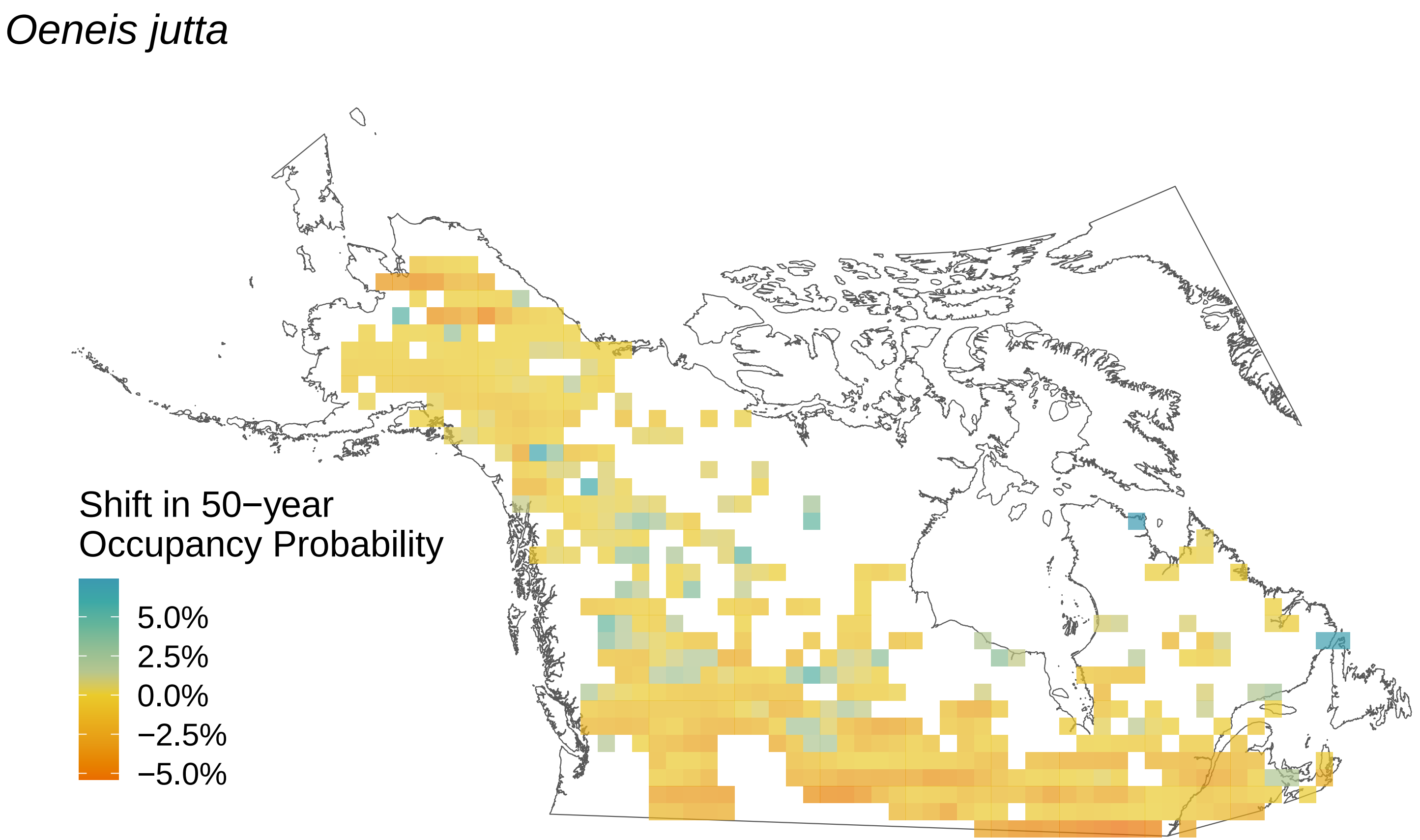
Oeneis jutta
Shift in 50−year Occupancy Probability
5.0%
2.5%
0.0%
−2.5%
−5.0%

*Oeneis chryxus*

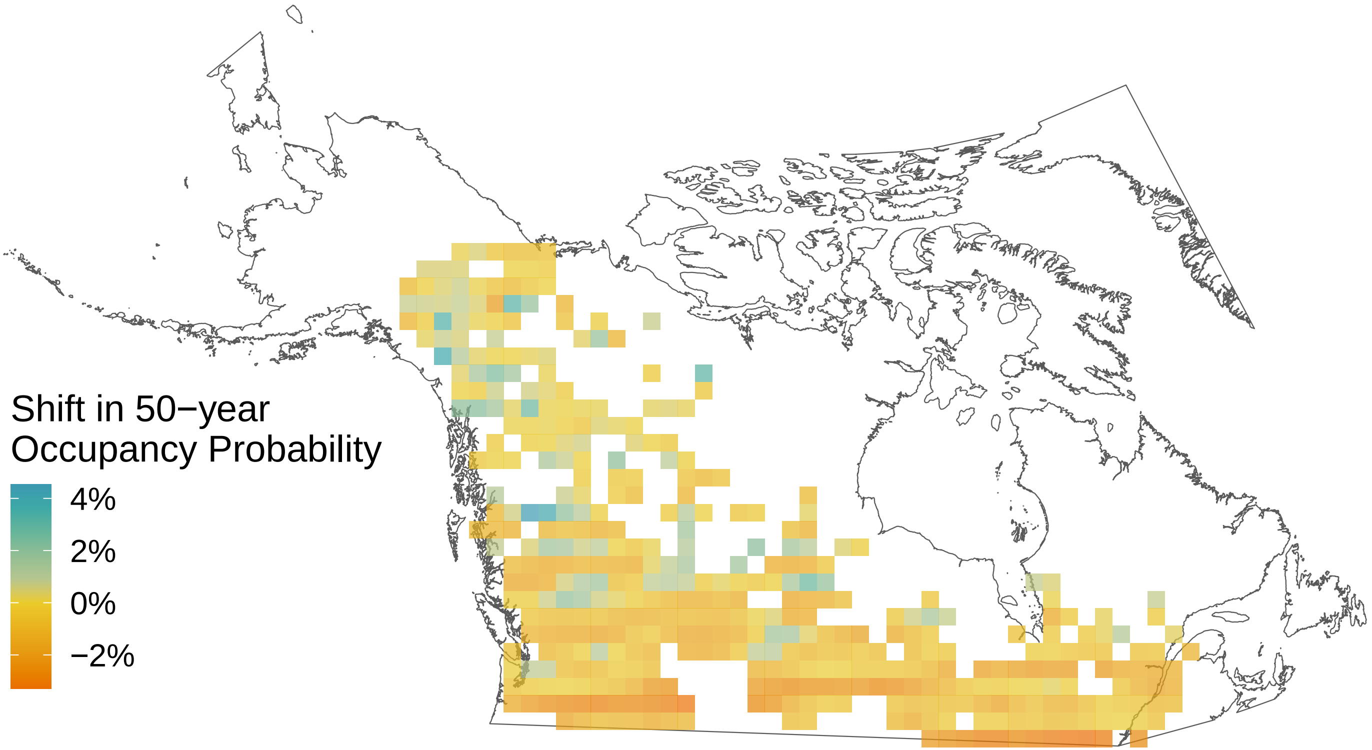

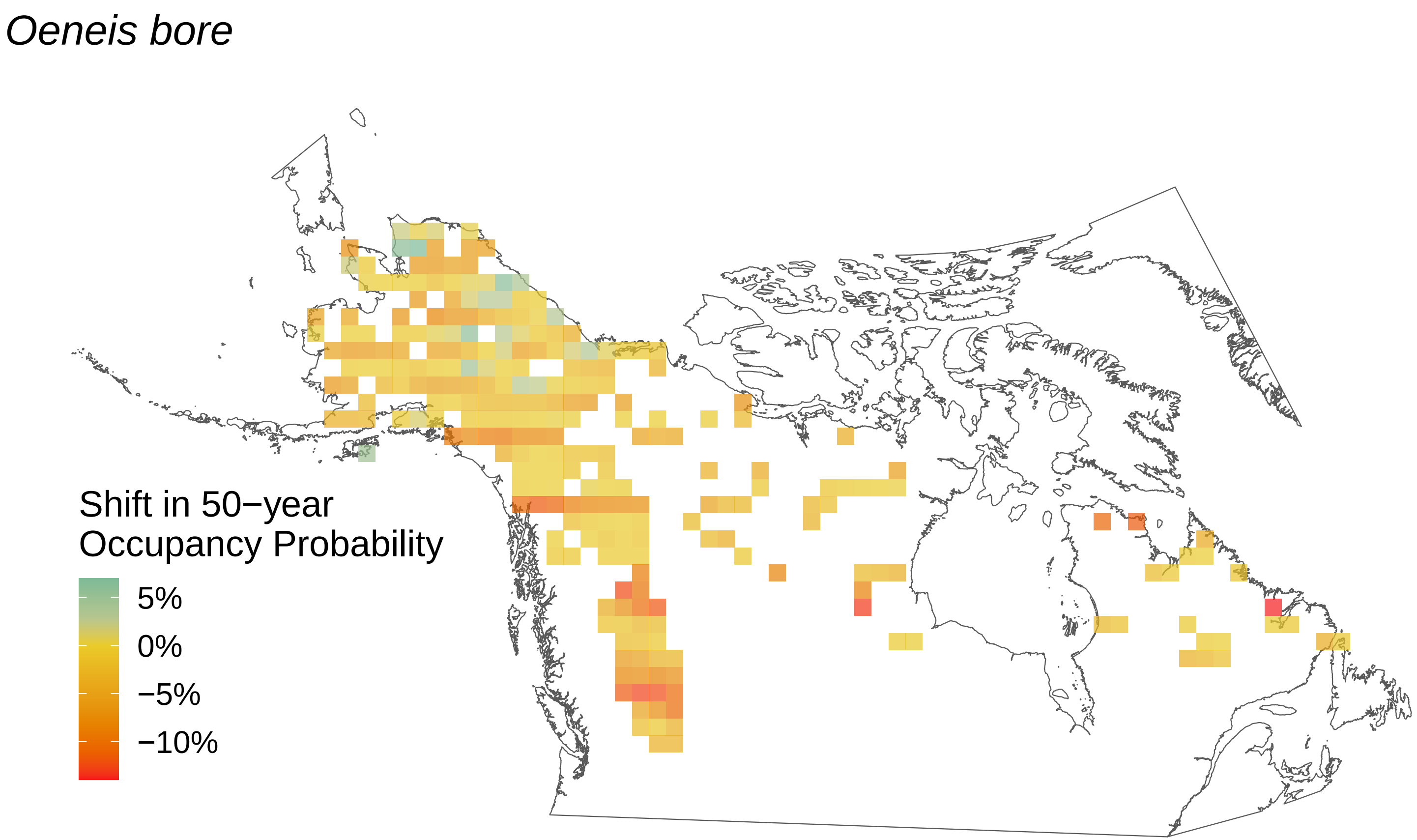
Oeneis bore
Shift in 50−year
Occupancy Probability
5%
0%
−5%
−10%

*Ochlodes sylvanoides*

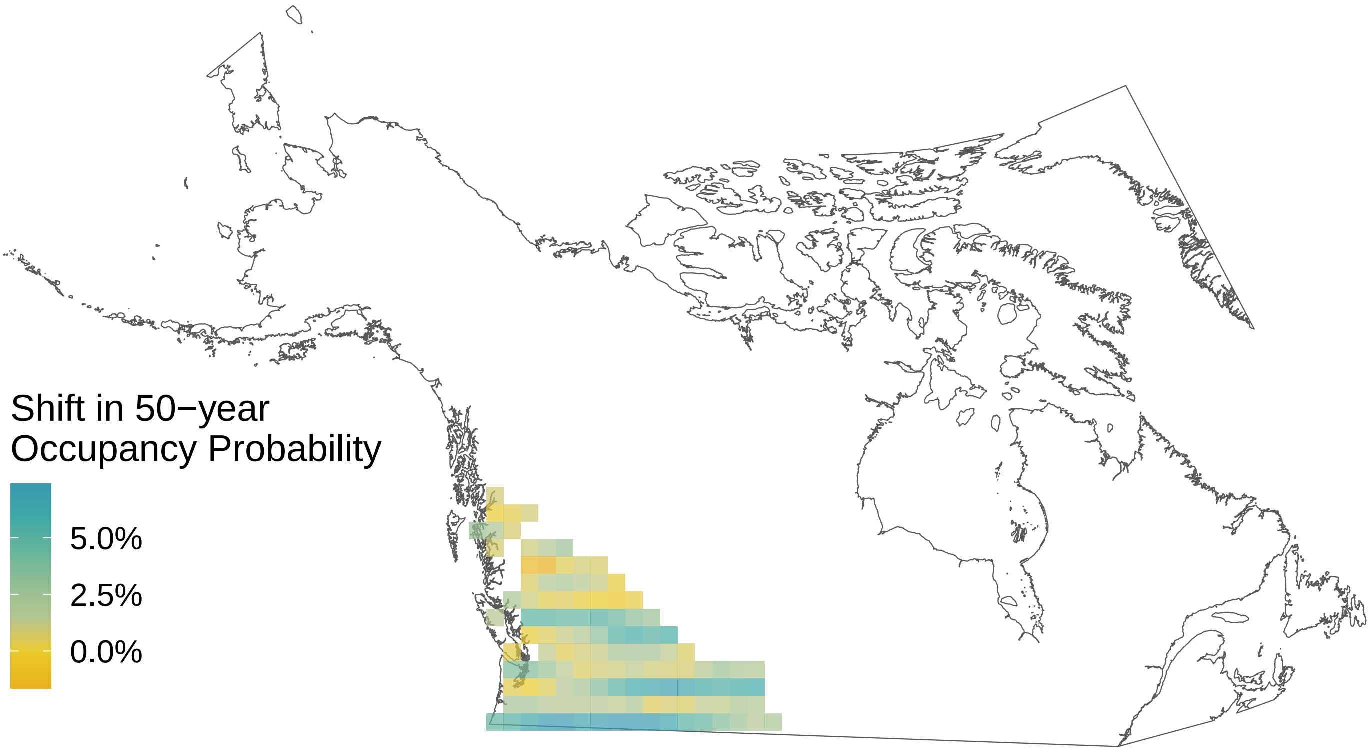

*Nymphalis l−album*

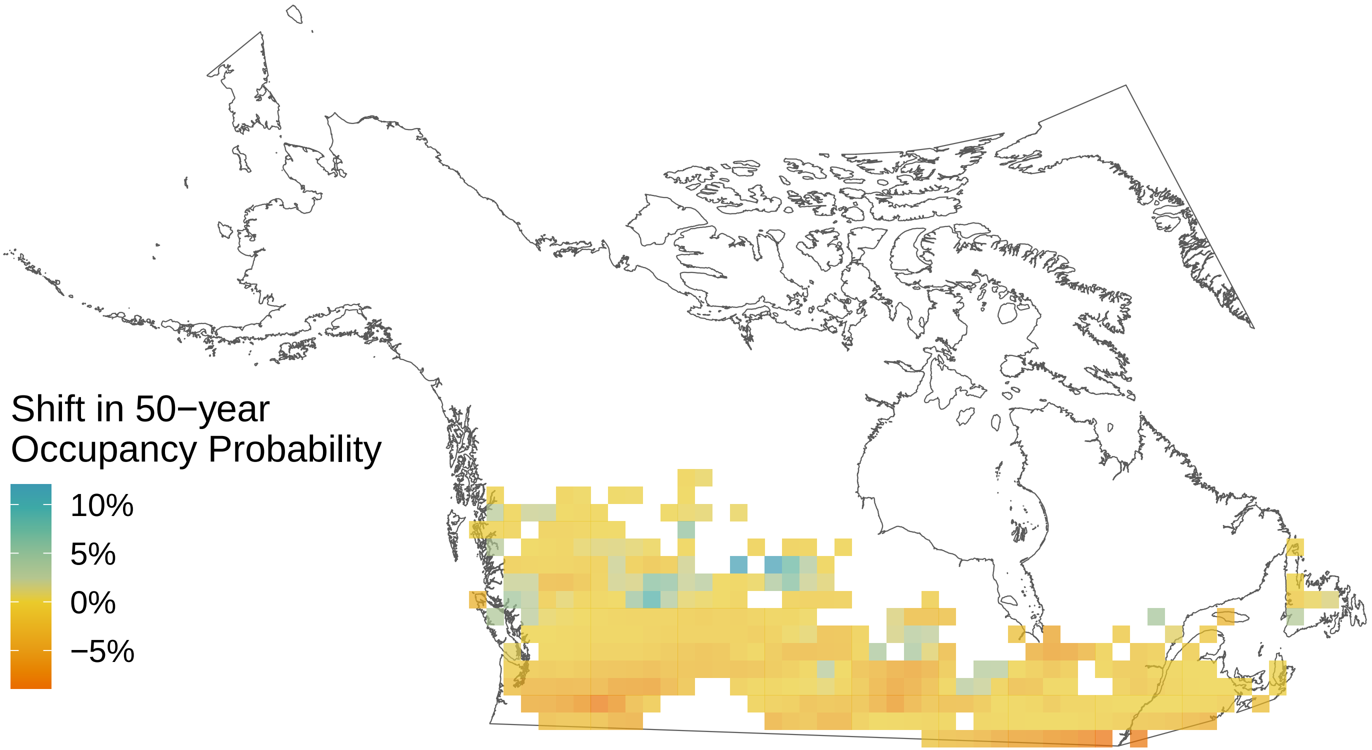

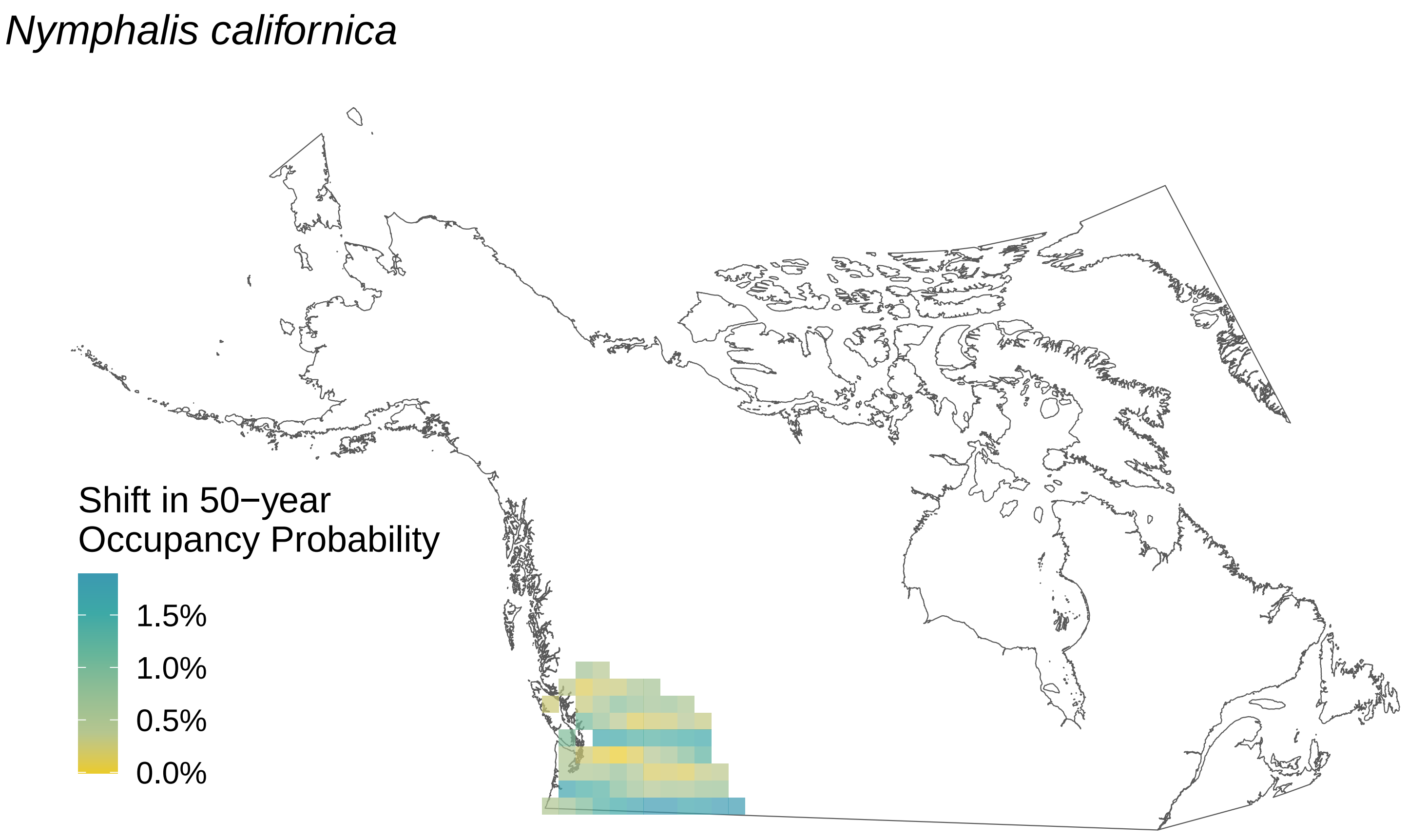
Nymphalis californica
Shift in 50−year
Occupancy Probability
1.5%
1.0%
0.5%
0.0%

*Nymphalis antiopa*

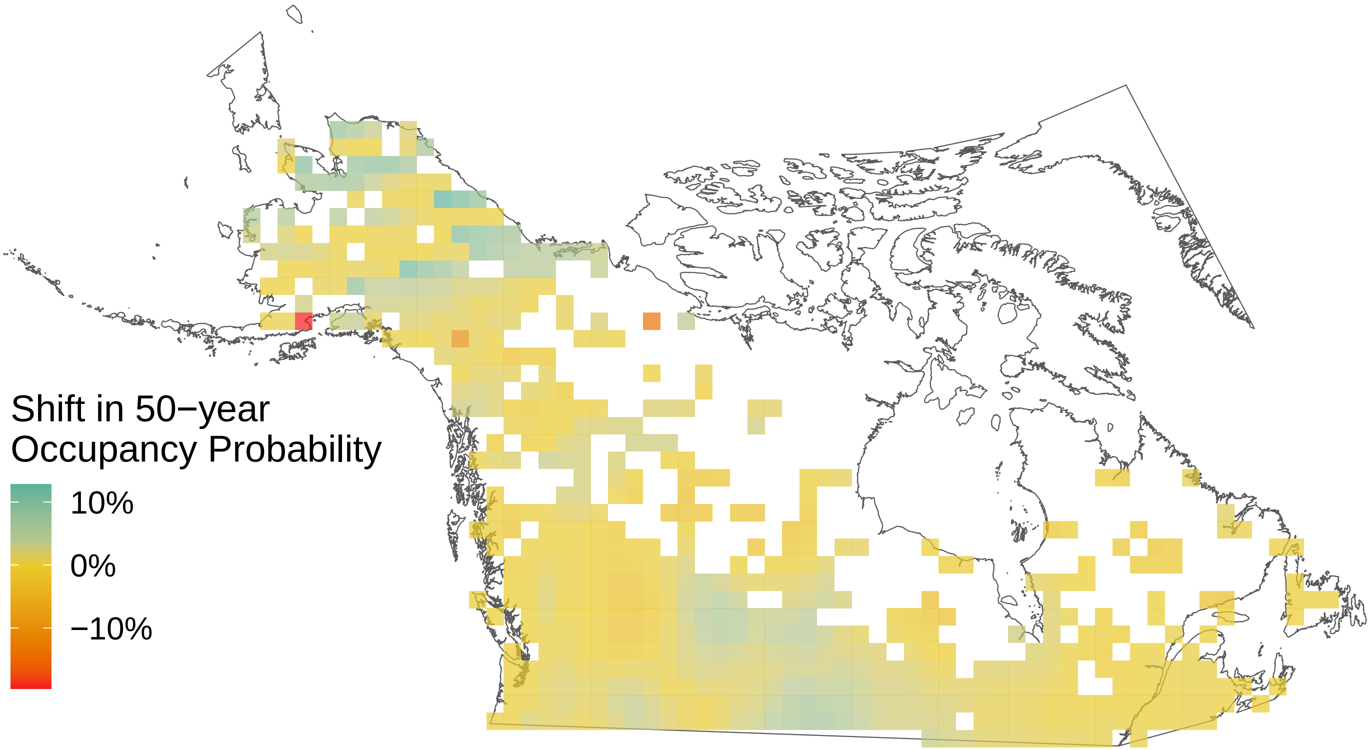

*Neophasia menapia*

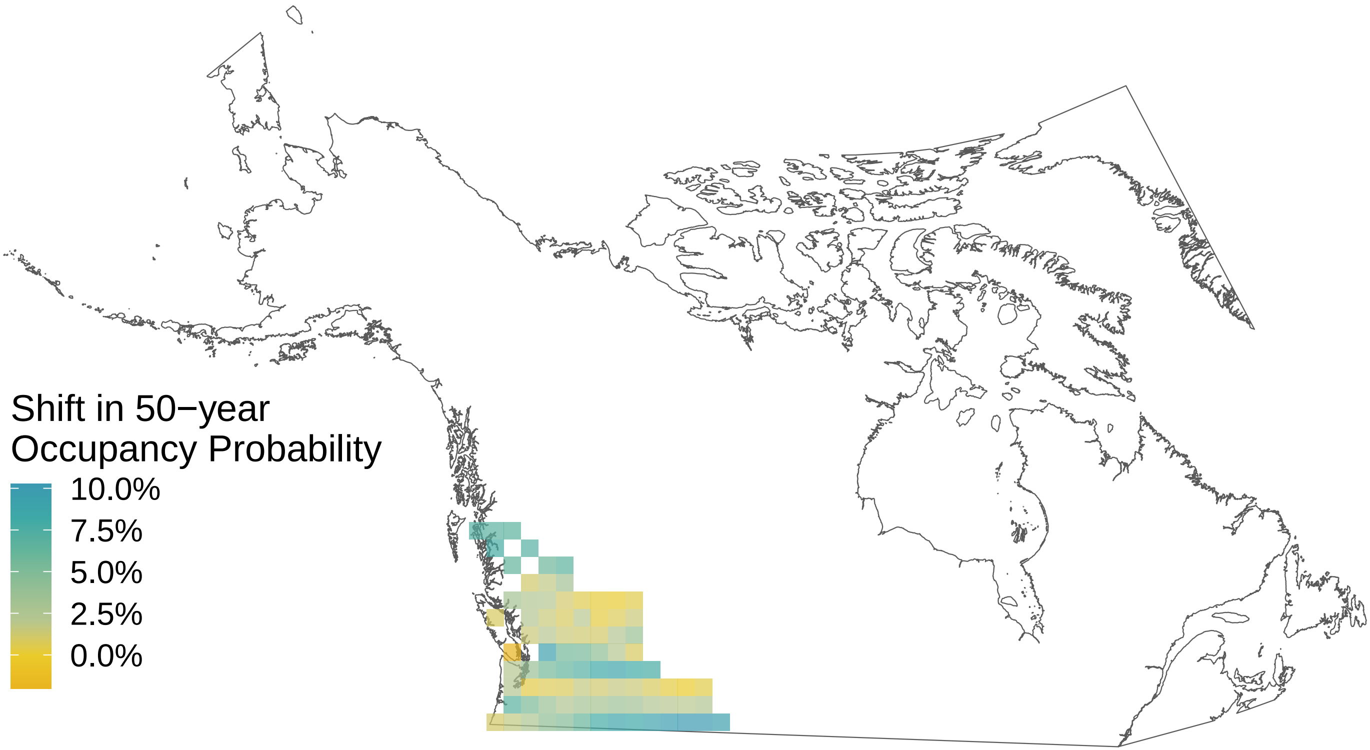

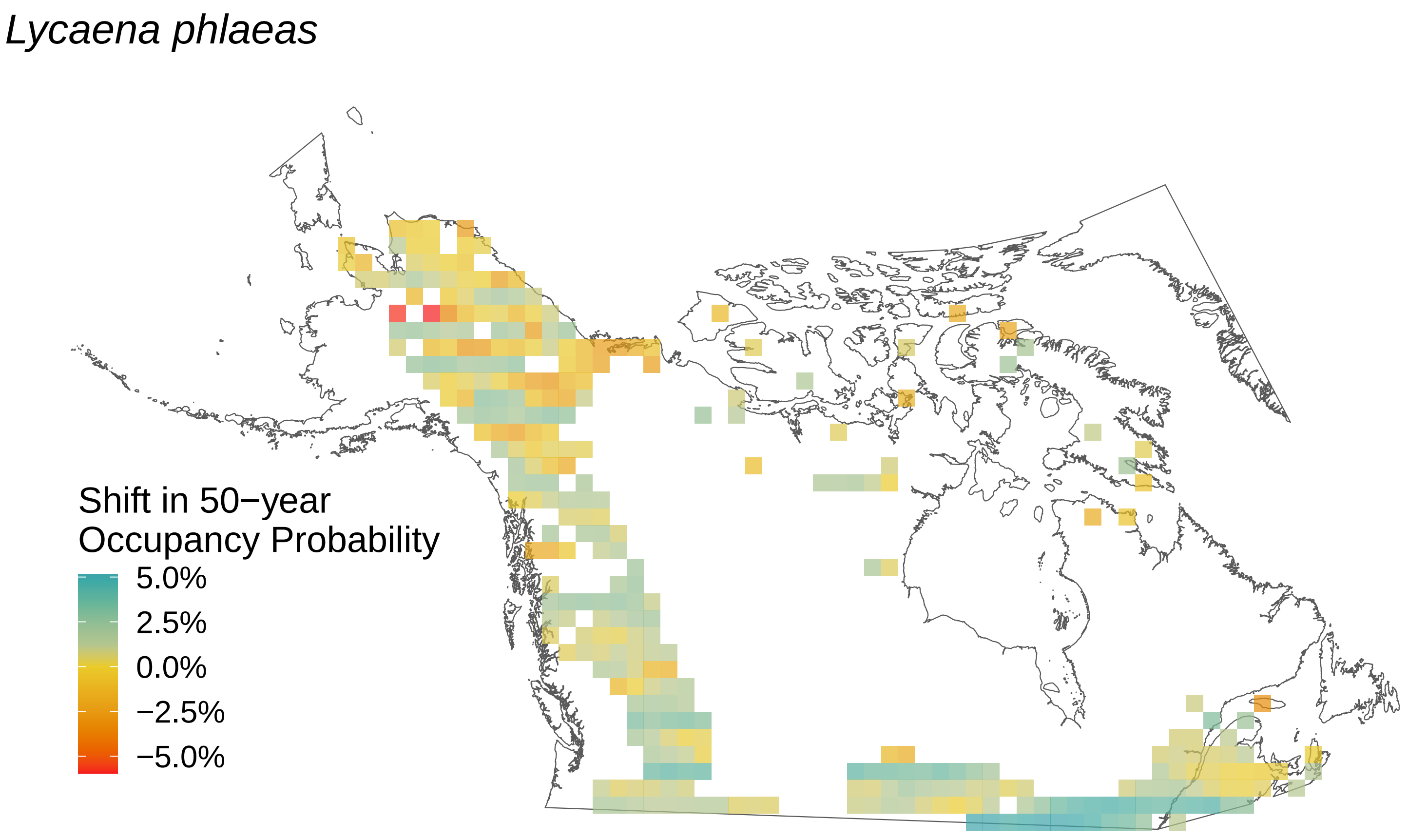
*Lycaena phlaeas*
Shift in 50−year
Occupancy Probability
5.0%
2.5%
0.0%
−2.5%
−5.0%

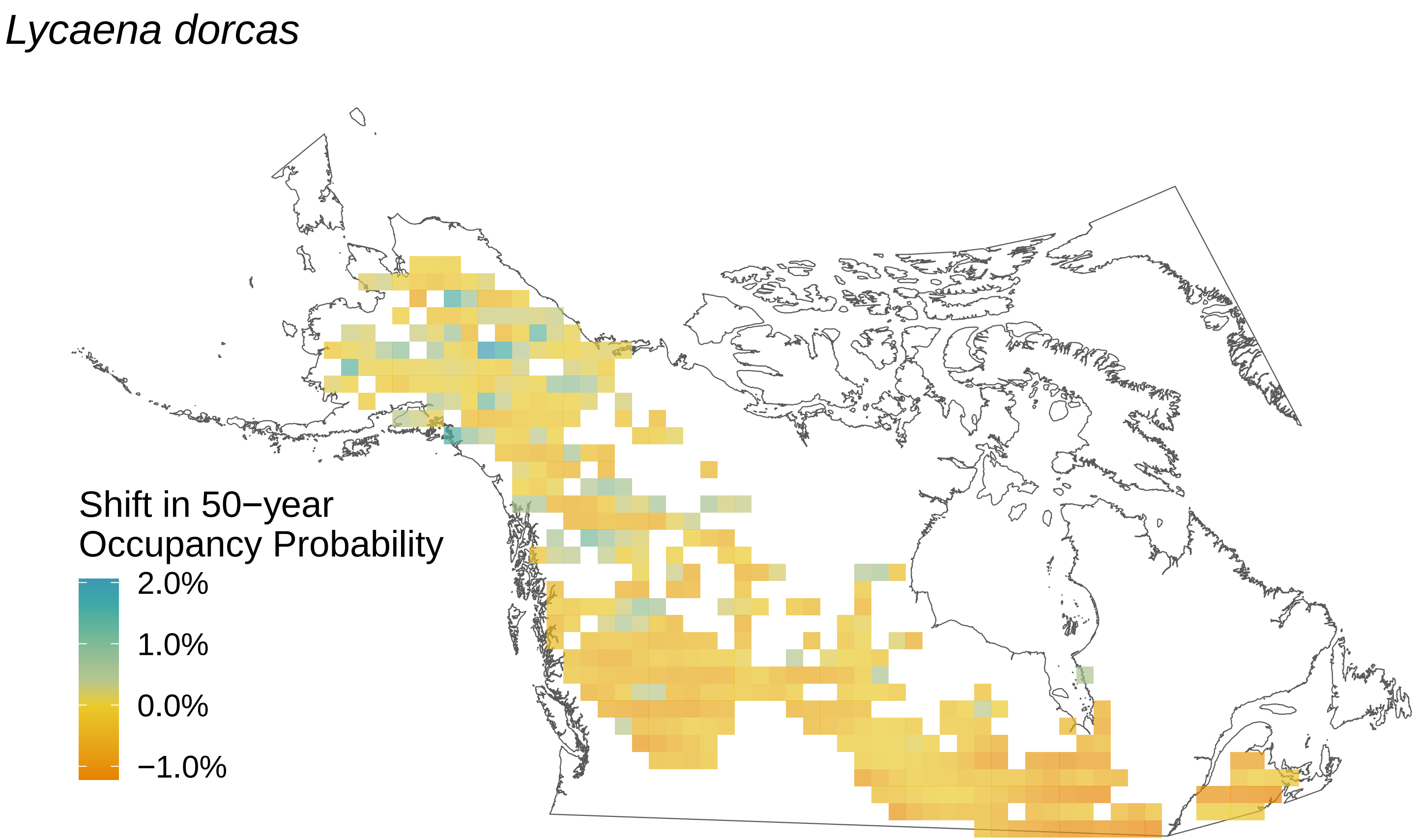
Lycaena dorcas
Shift in 50−year
Occupancy Probability
2.0%
1.0%
0.0%
−1.0%

*Limenitis lorquini*

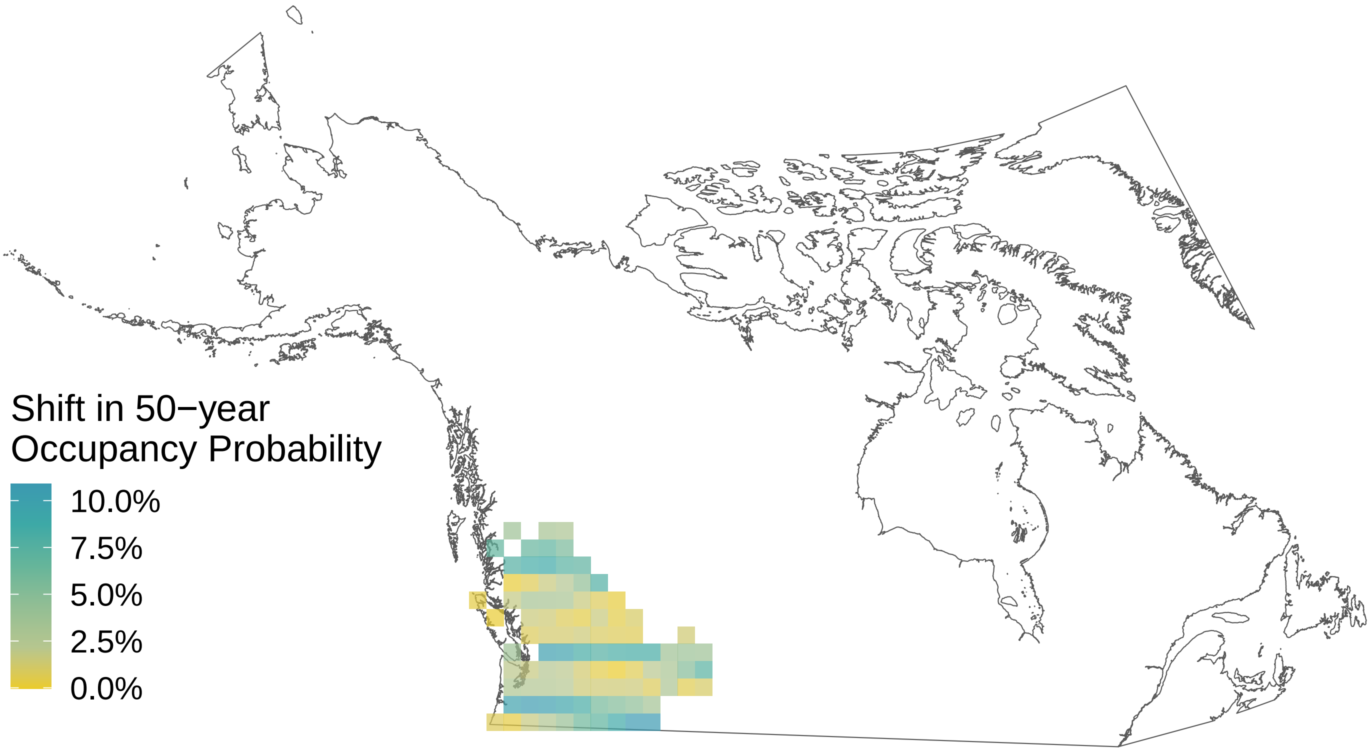

*Limenitis arthemis*

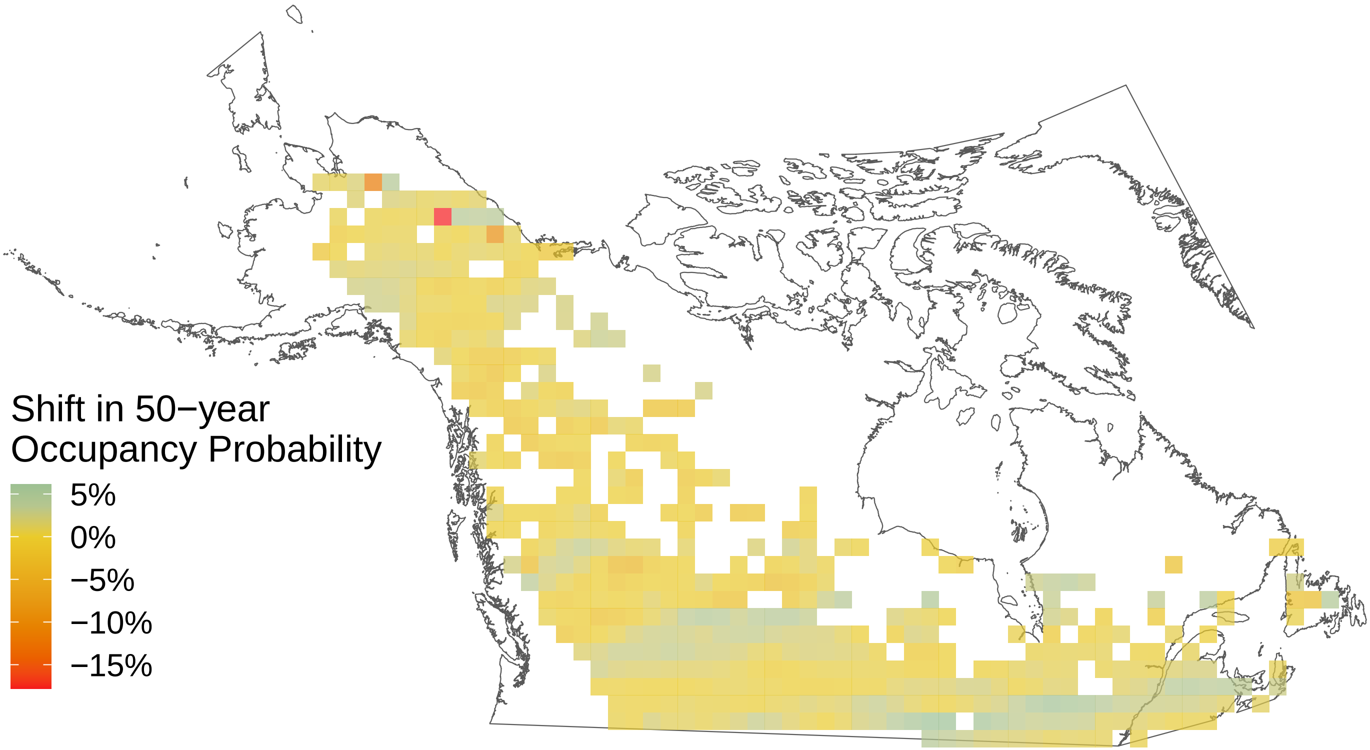

*Limenitis archippus*

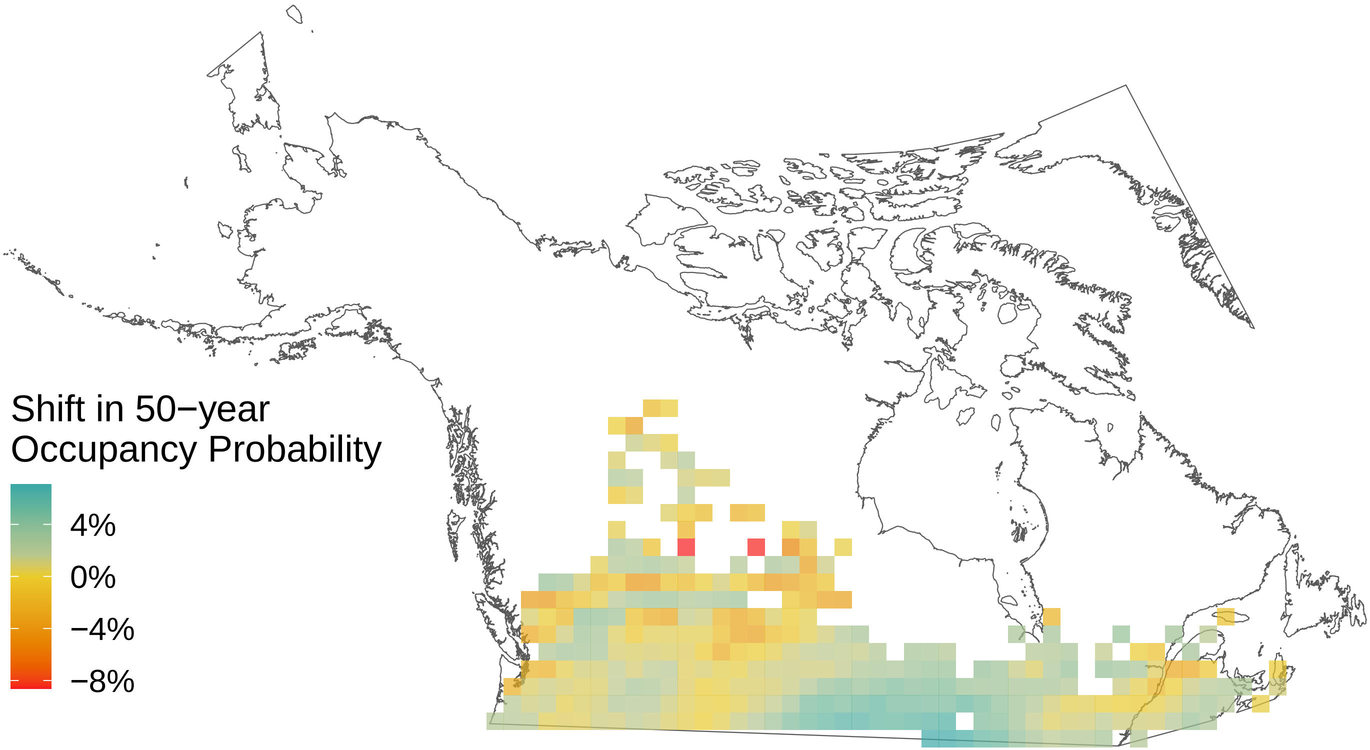

### *Lethe eurydice*

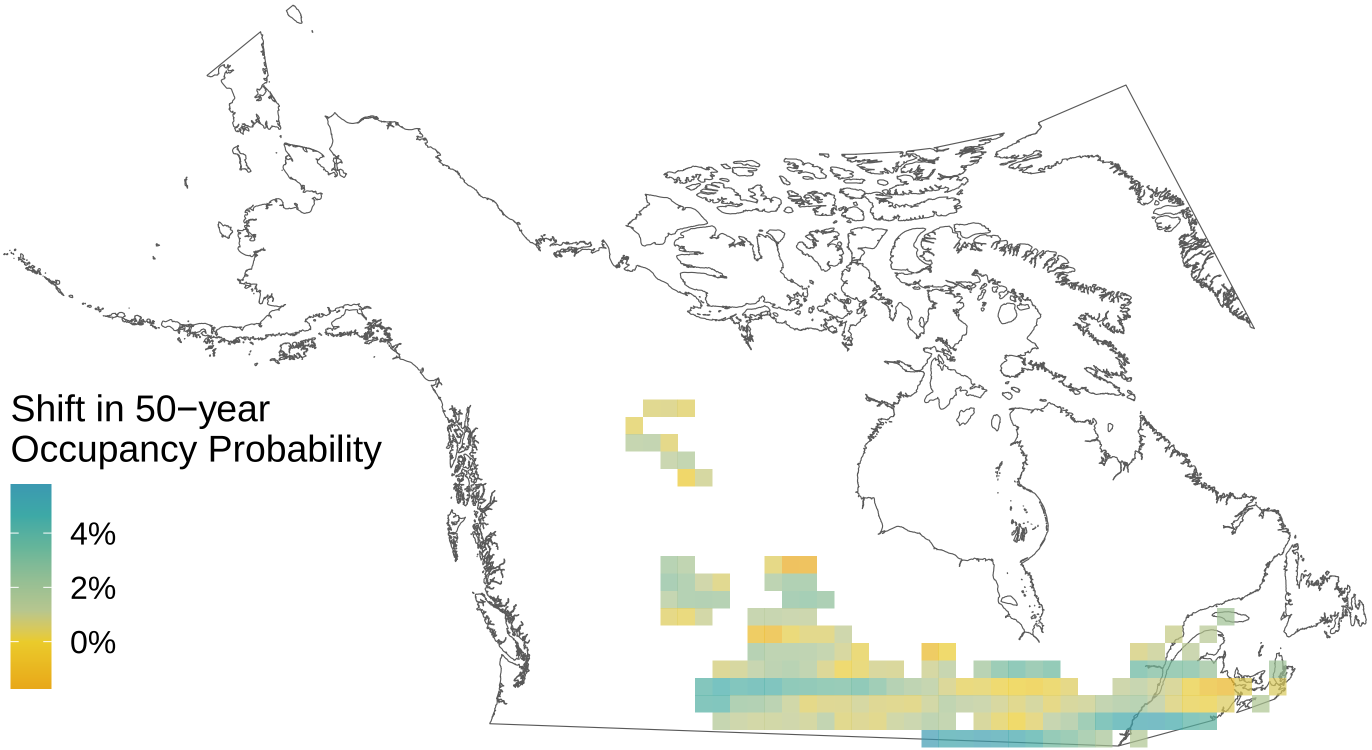

*Icaricia saepiolus*

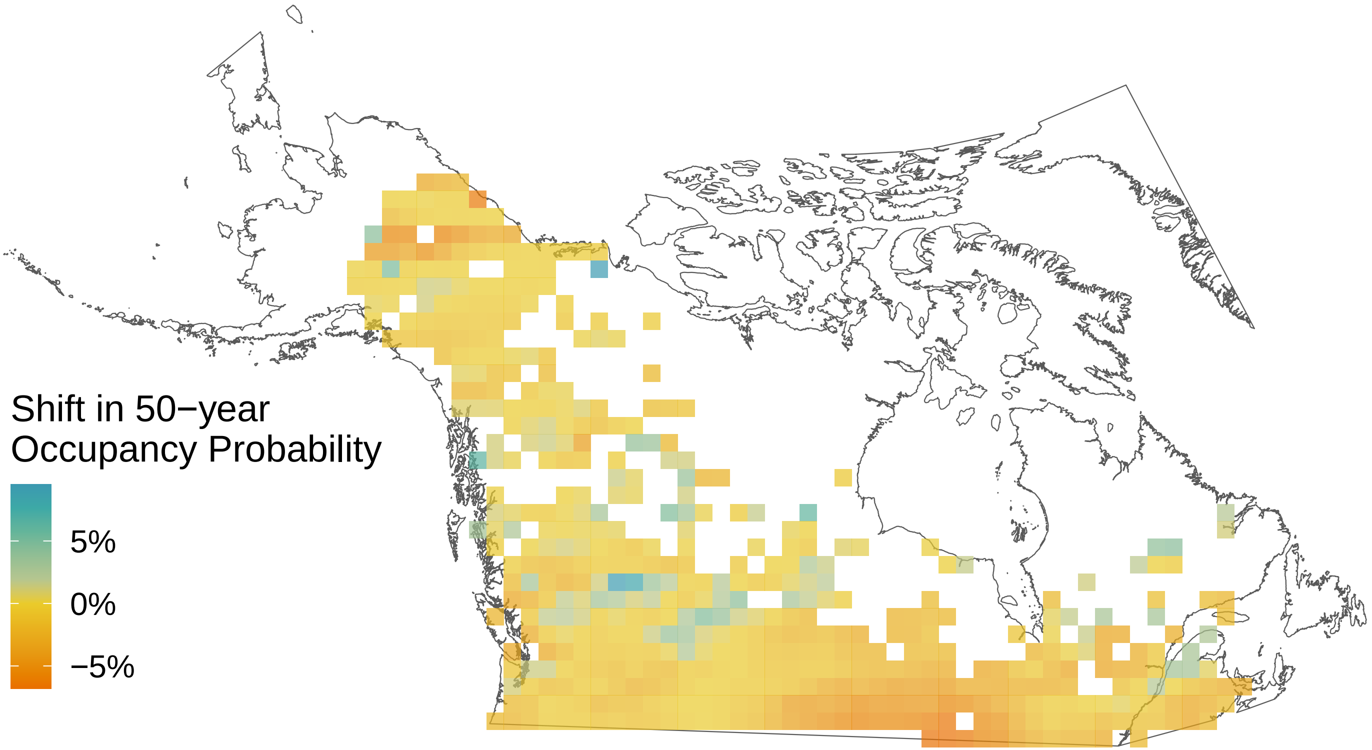

*Icaricia icarioides*

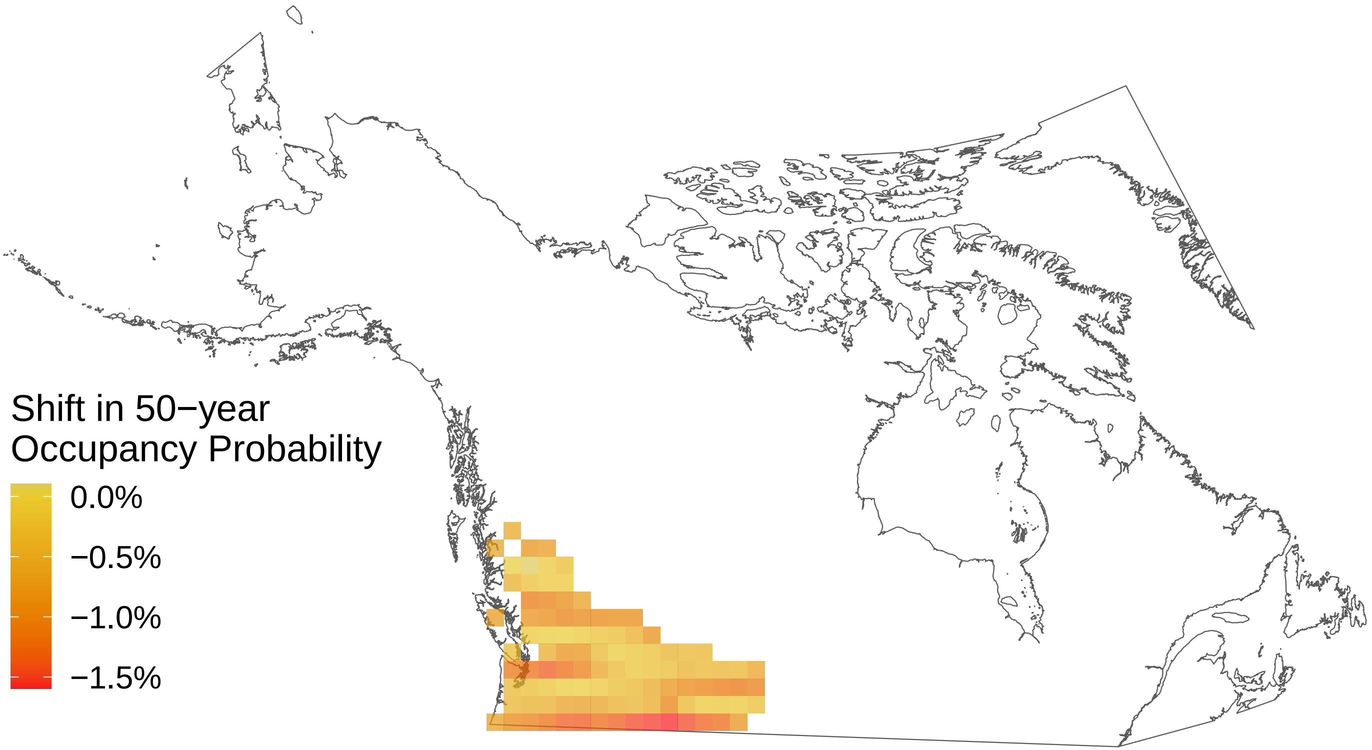

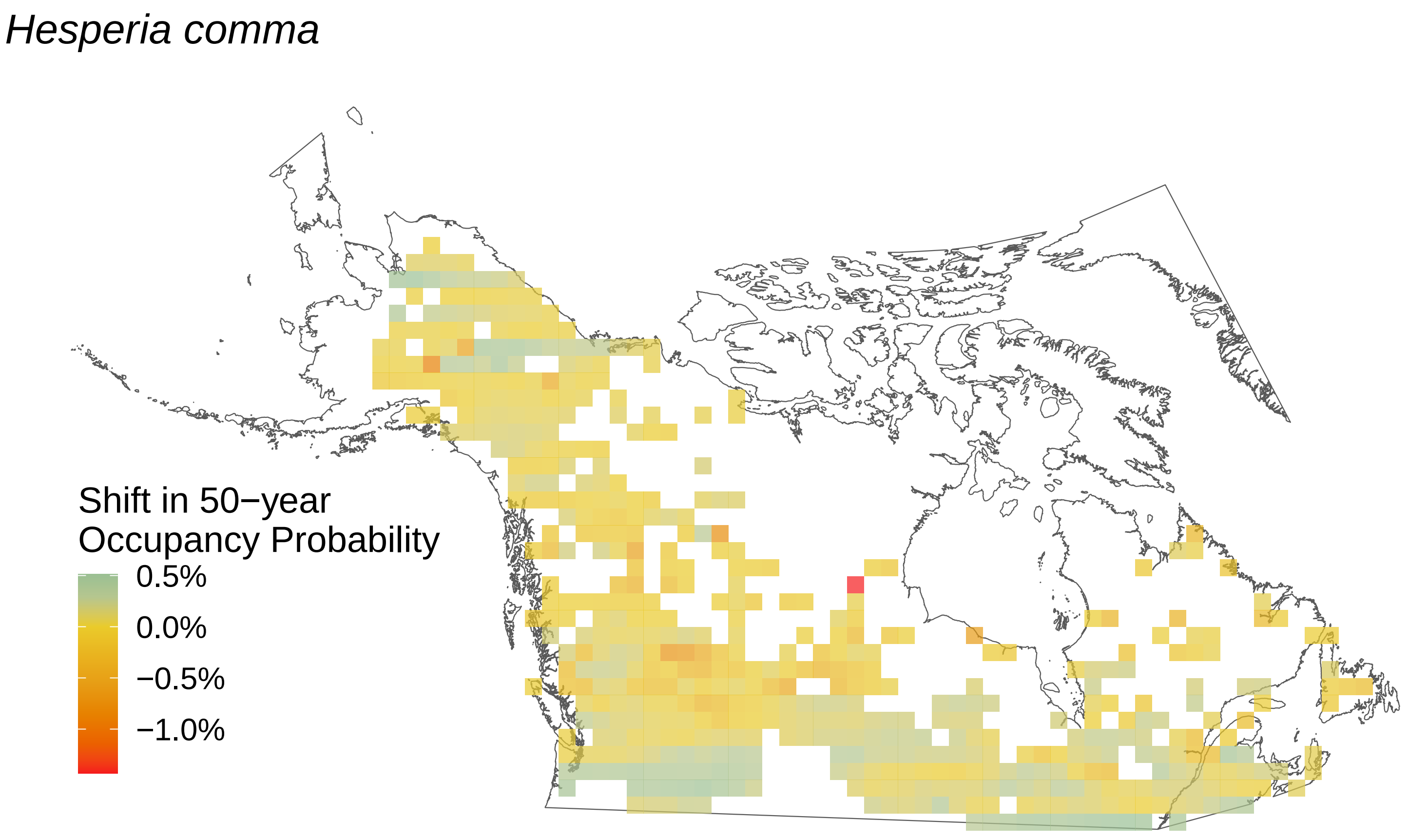
Hesperia comma
Shift in 50−year
Occupancy Probability
0.5%
0.0%
−0.5%
−1.0%

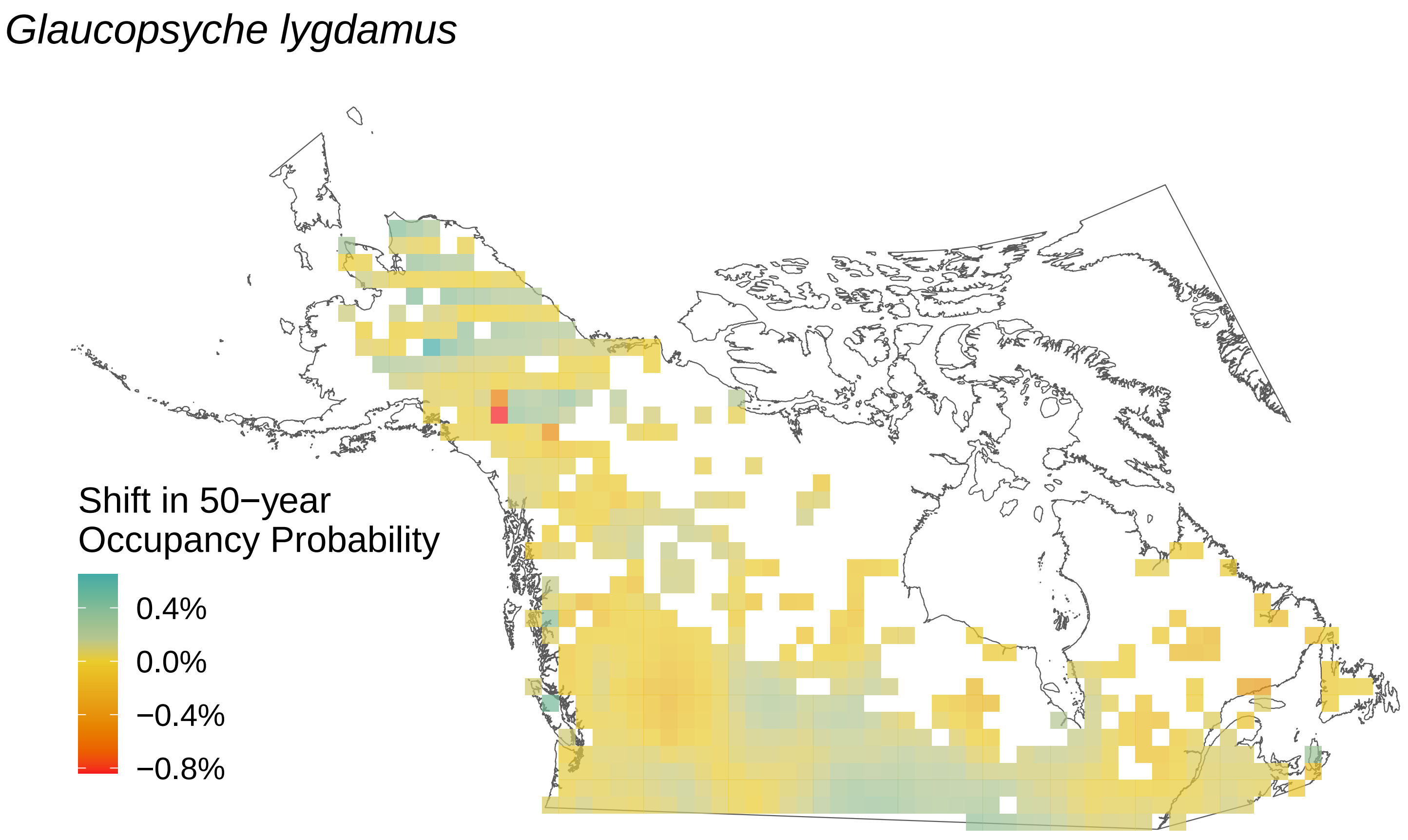

Glaucopsyche lygdamus
Shift in 50−year
Occupancy Probability
0.4%
0.0%
−0.4%
−0.8%

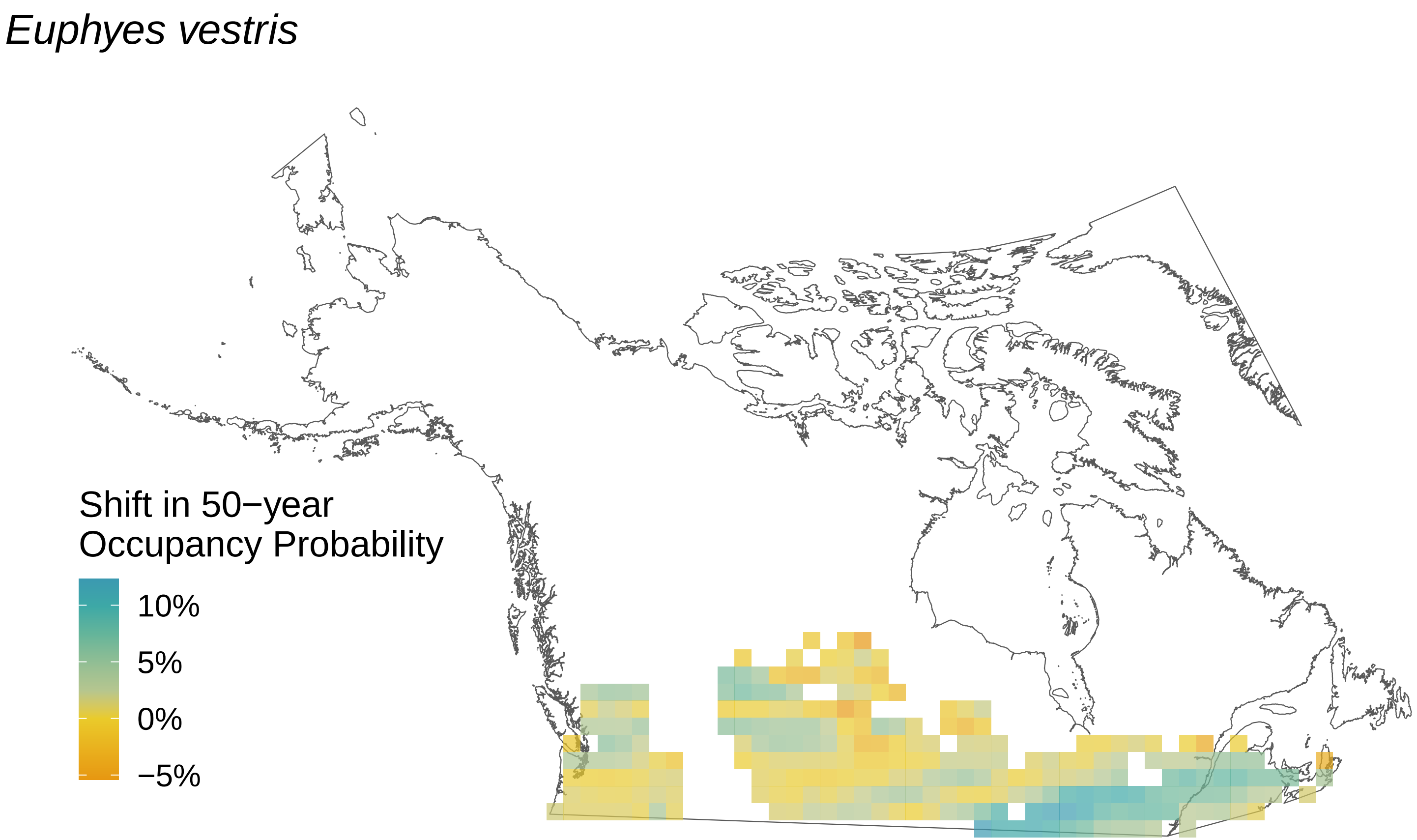

Euphyes vestris
Shift in 50−year
Occupancy Probability
10%
5%
0%
−5%

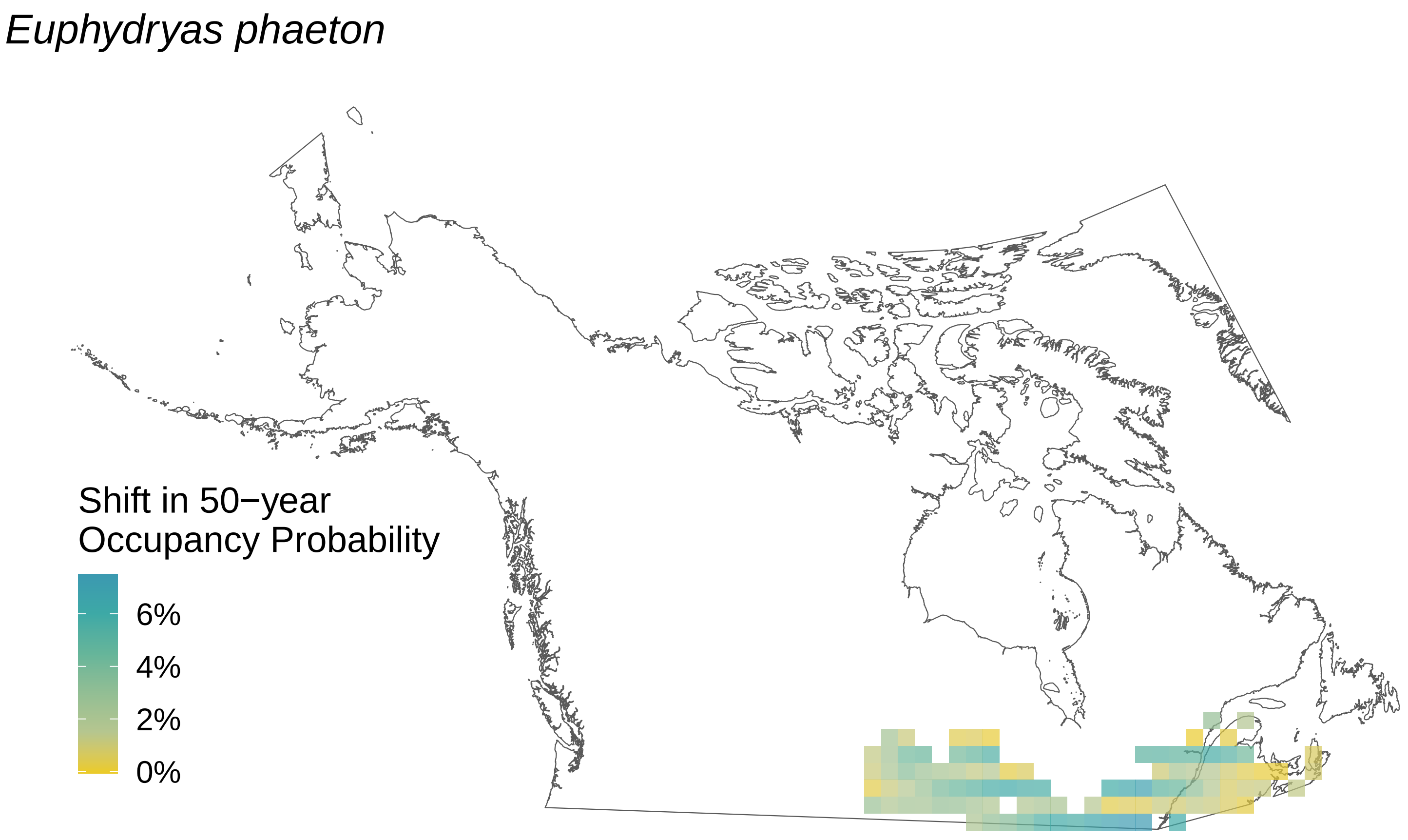

Euphydryas phaeton
Shift in 50−year
Occupancy Probability
6%
4%
2%
0%

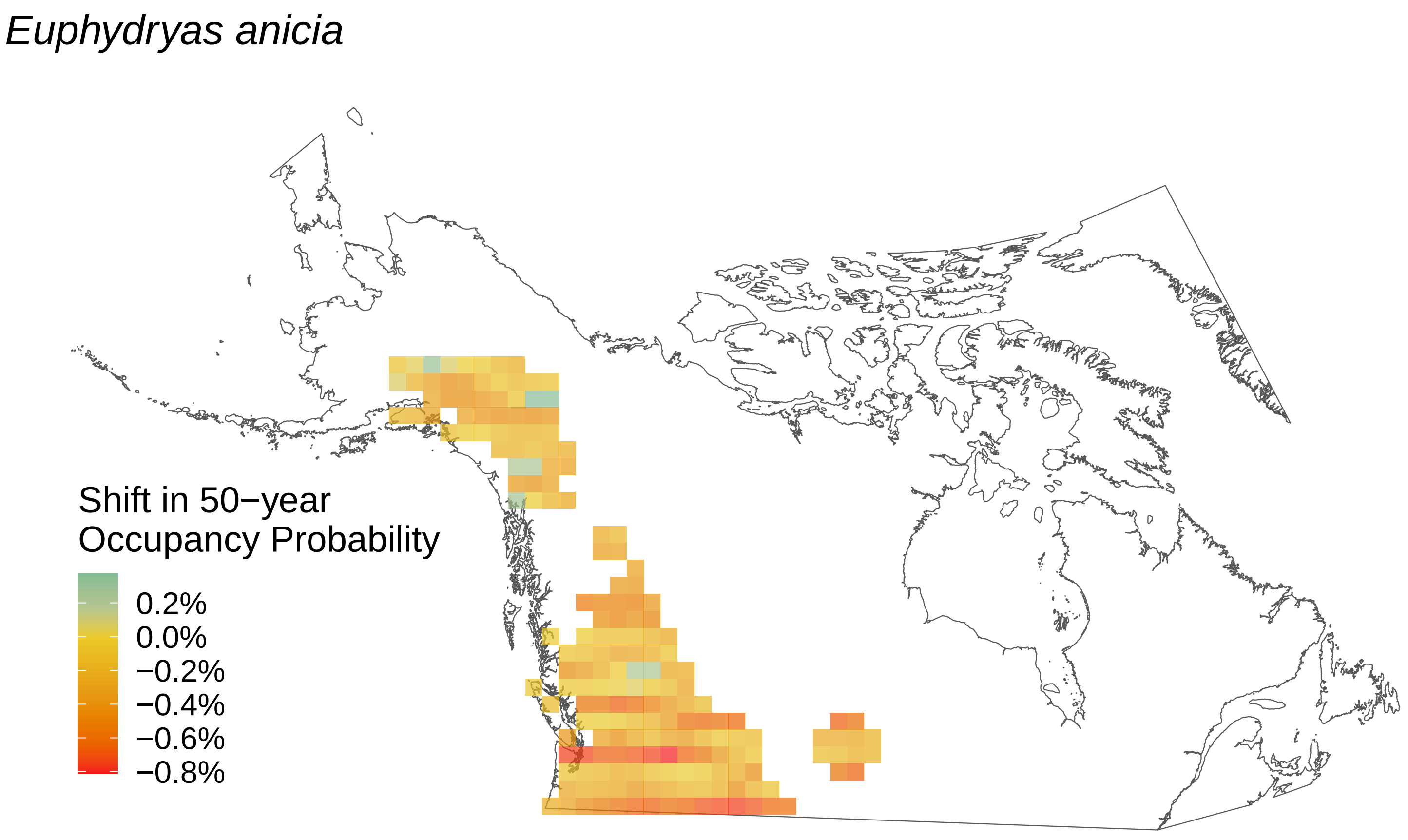
Euphydryas anicia
Shift in 50−year
Occupancy Probability
0.2%
0.0%
−0.2%
−0.4%
−0.6%
−0.8%

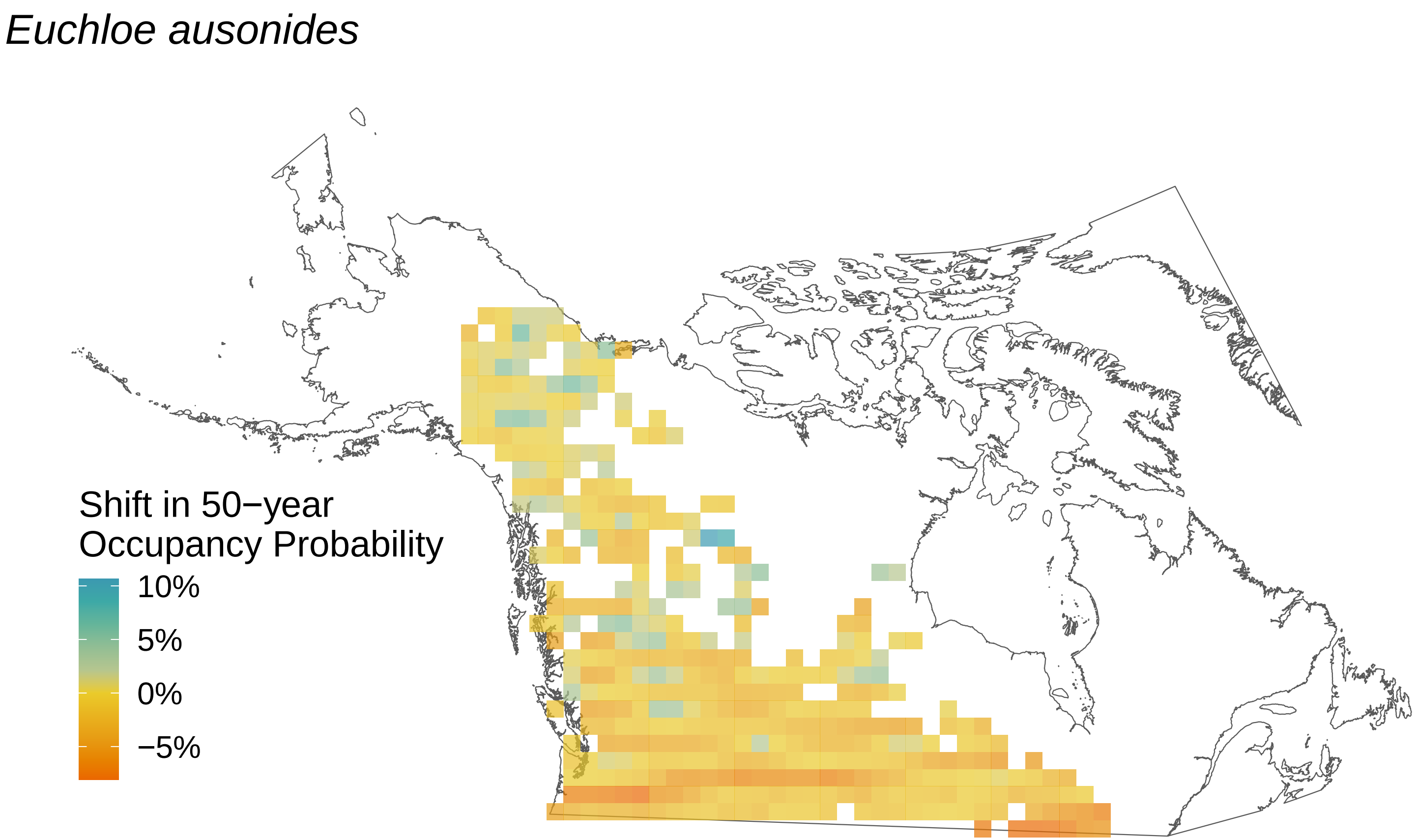
Euchloe ausonides
Shift in 50−year Occupancy Probability
10%
5%
0%
−5%

*Erynnis persius*

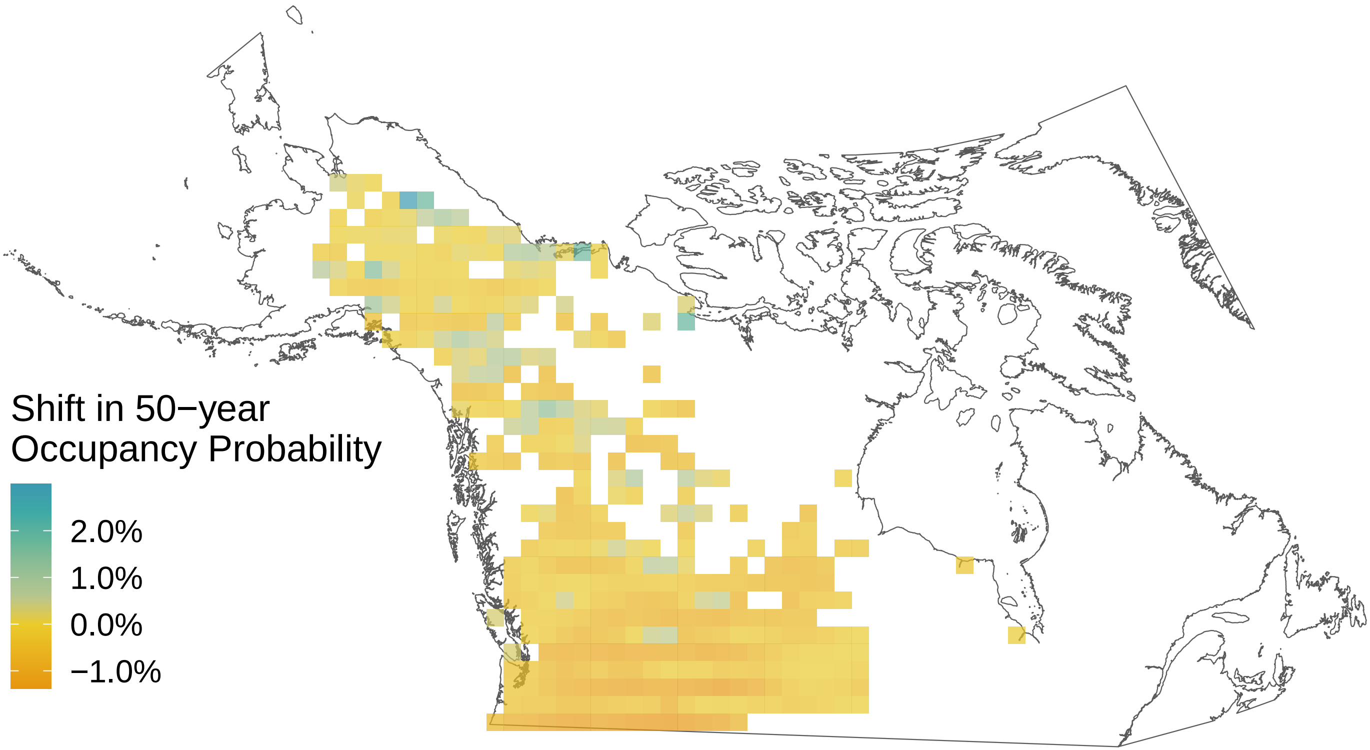

*Erynnis juvenalis*

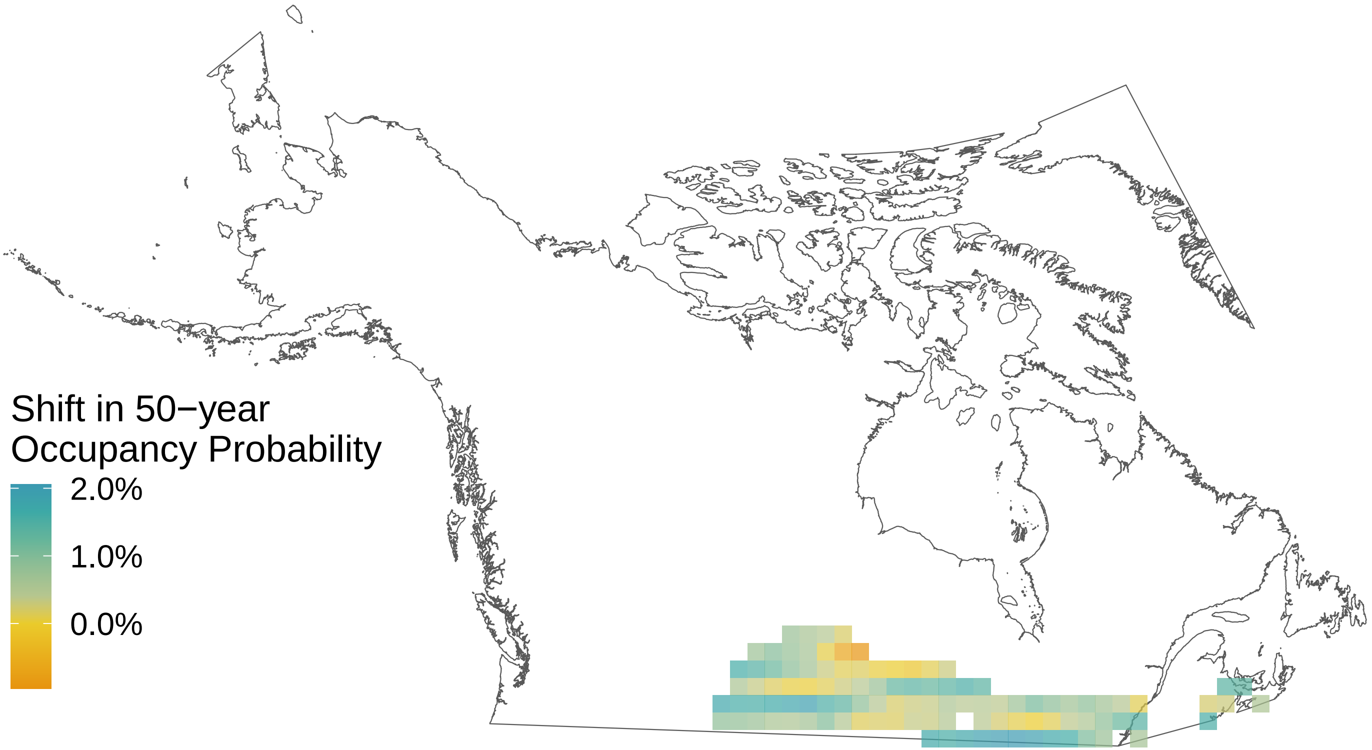

*Erynnis icelus*

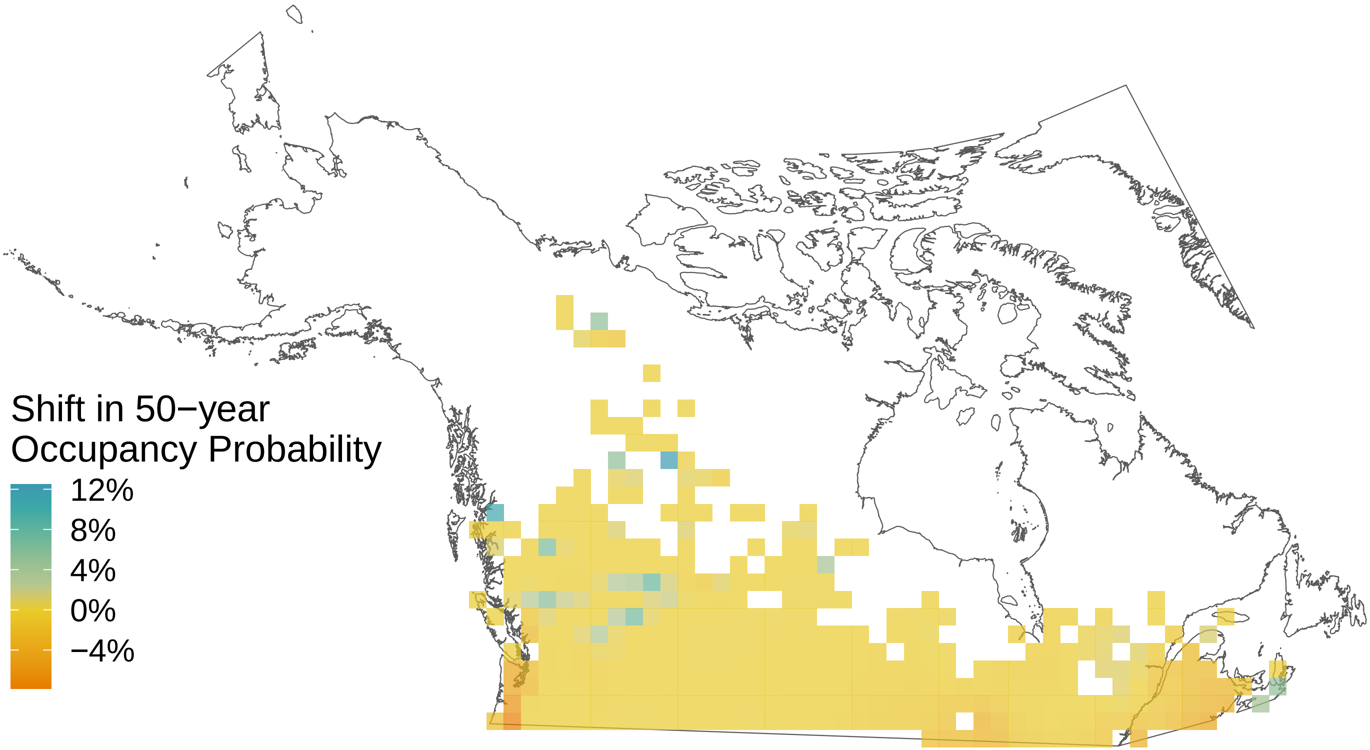

*Erebia youngi*

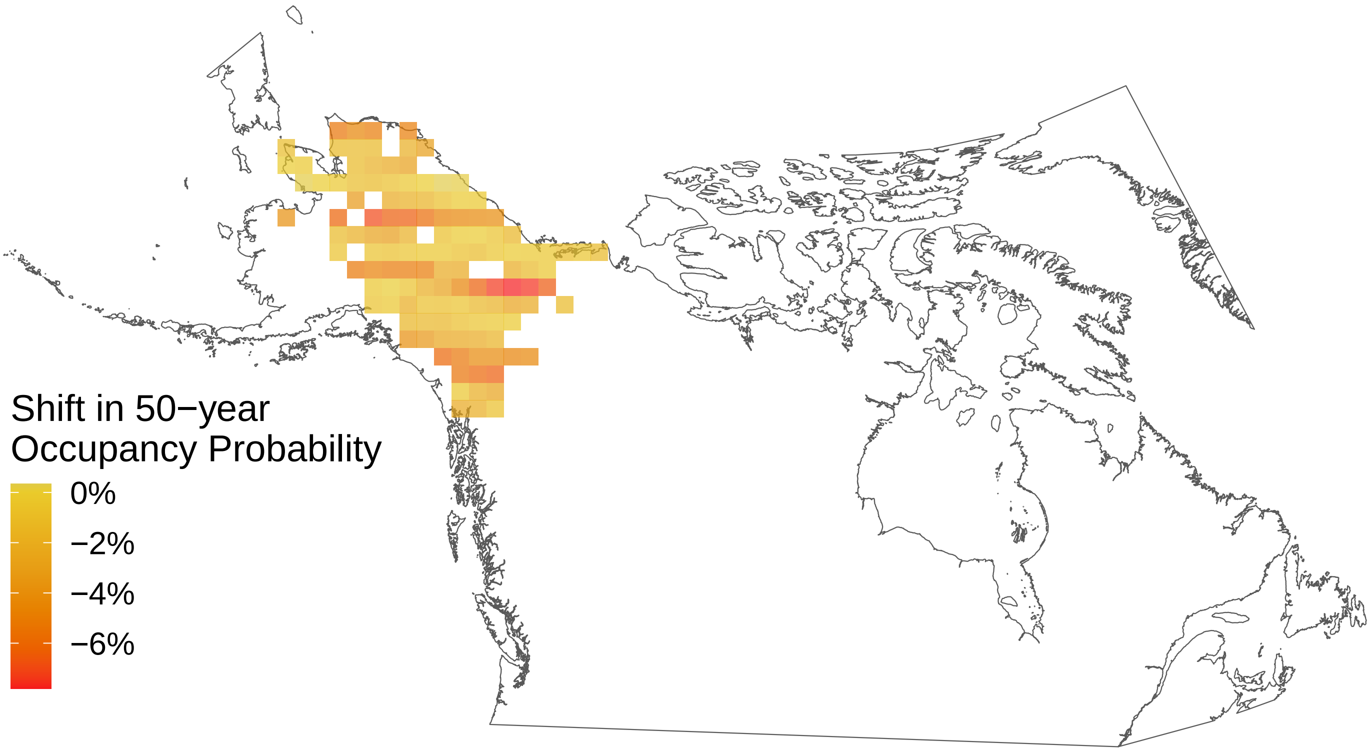

*Erebia rossii*

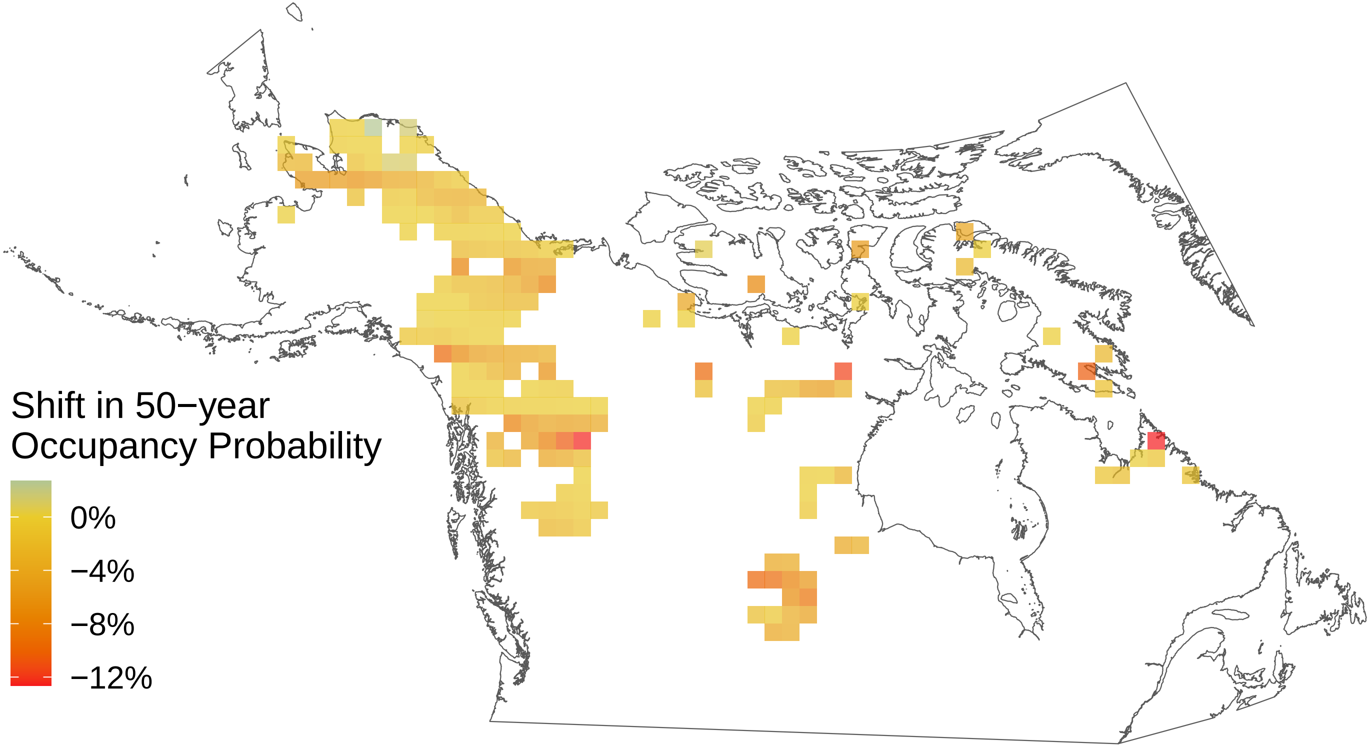

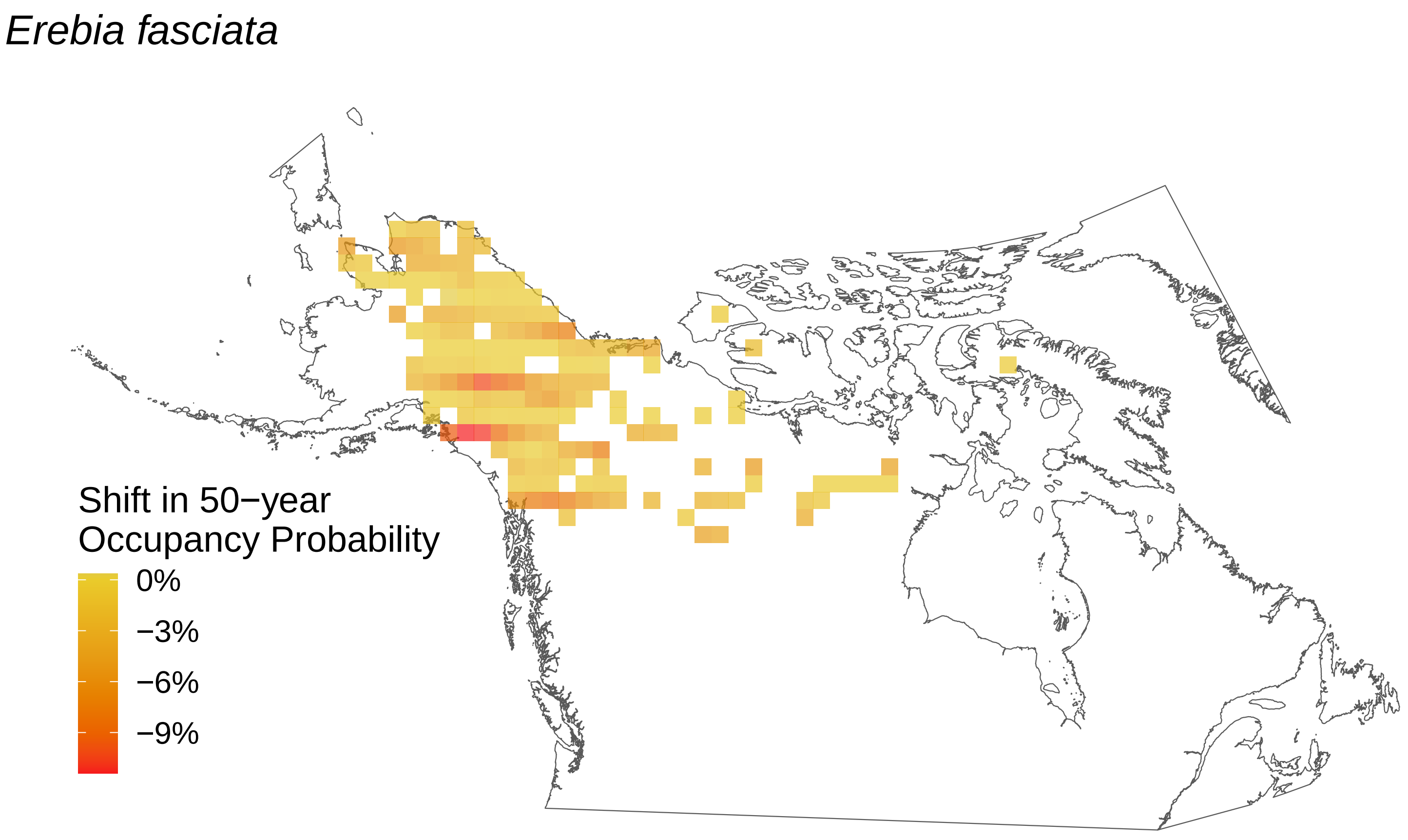

Erebia fasciata
Shift in 50−year Occupancy Probability
0%
−3%
−6%
−9%

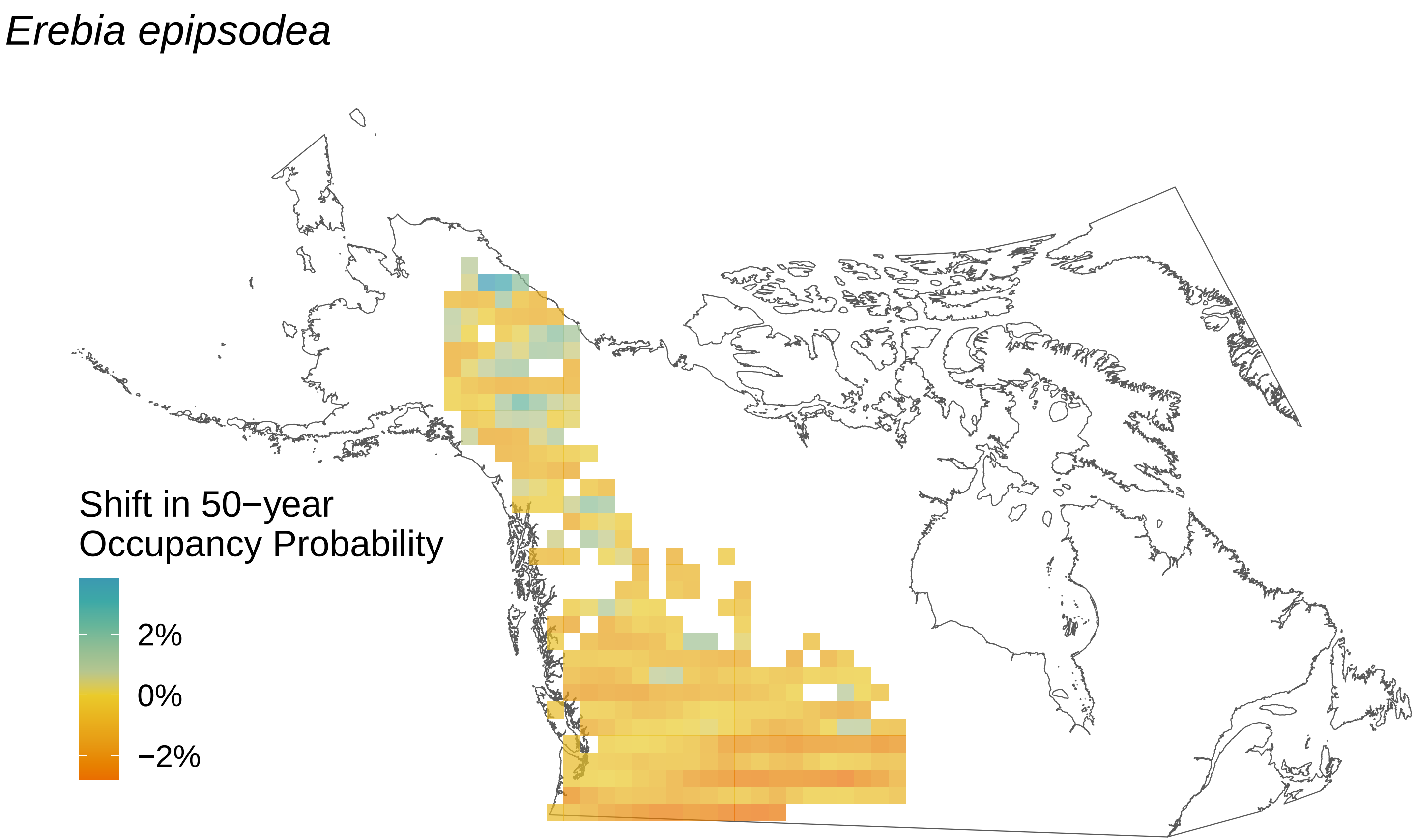
Erebia epipsodea
Shift in 50−year Occupancy Probability
2%
0%
−2%

*Erebia discoidalis*

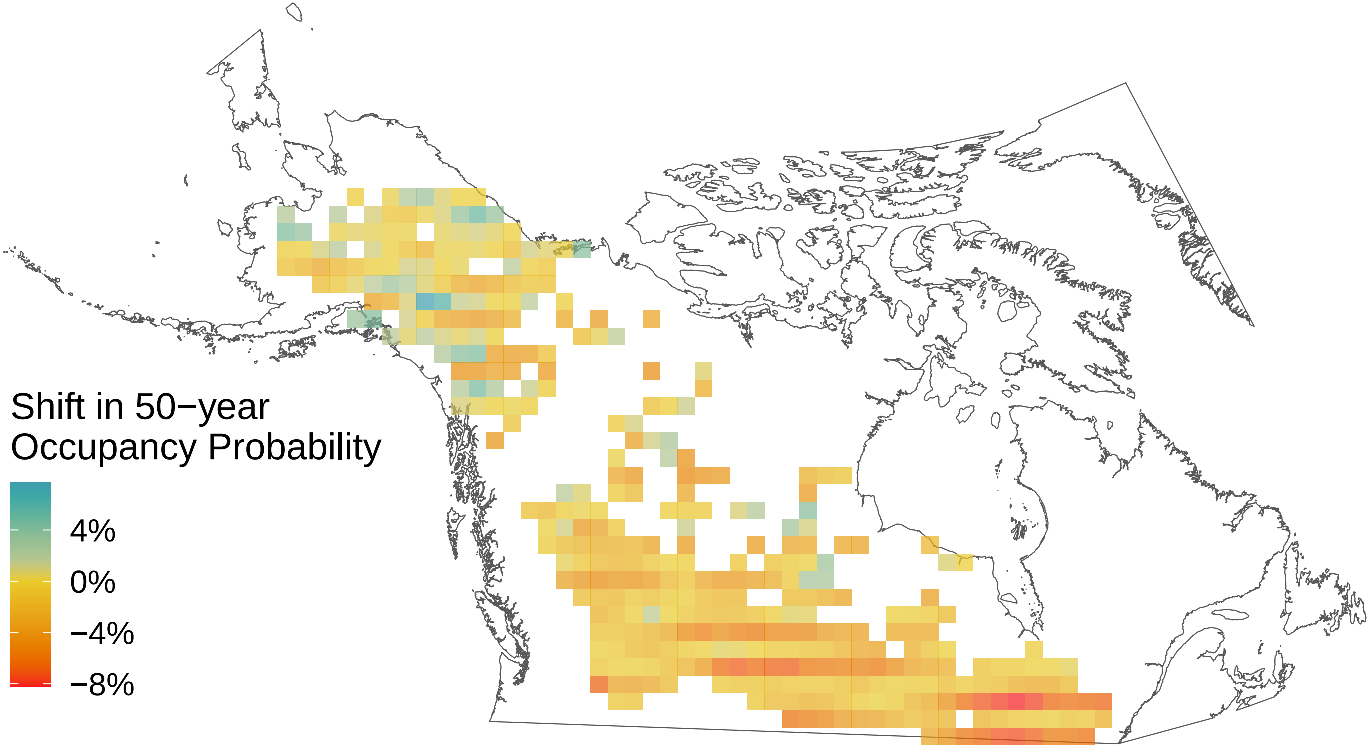

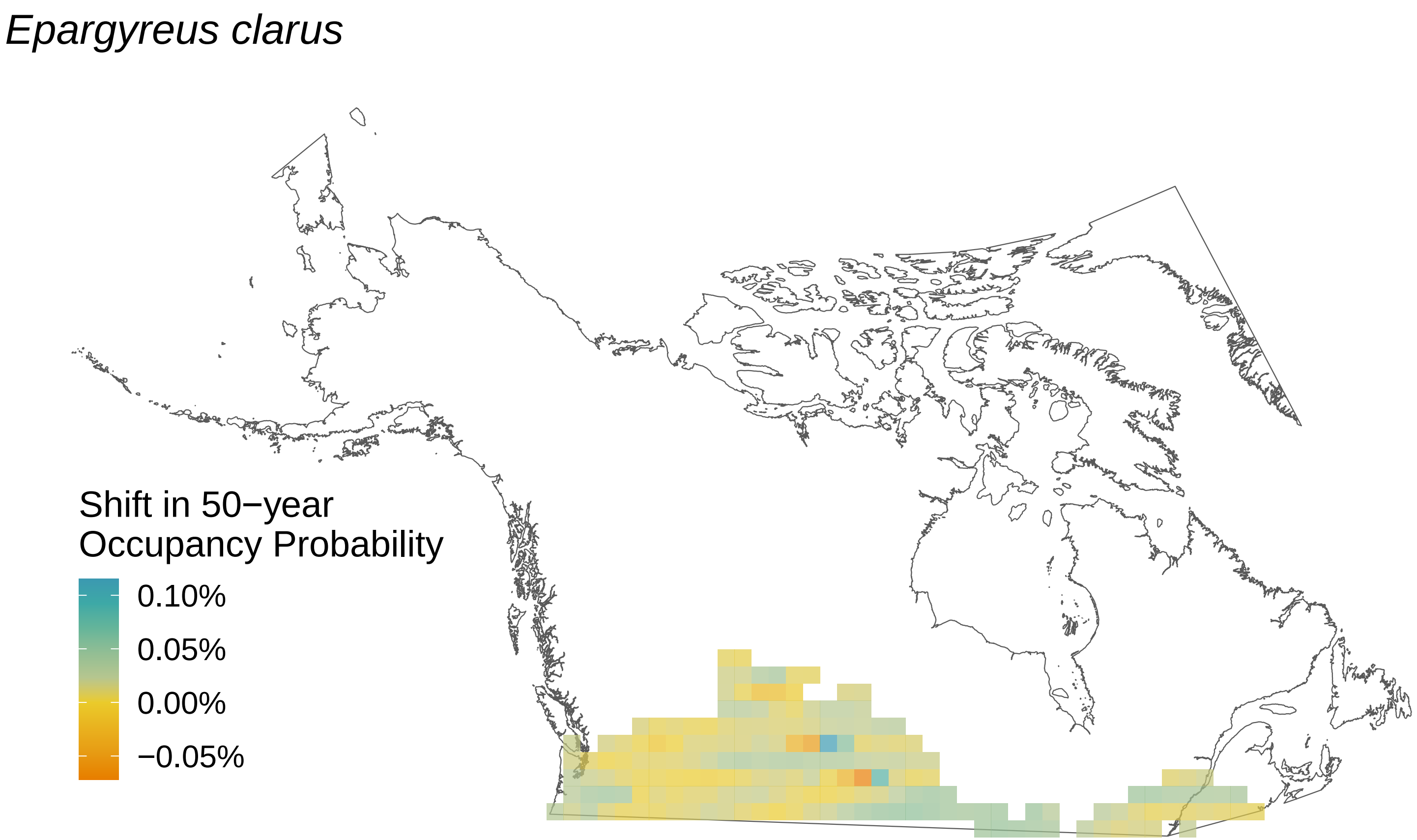

Epargyreus clarus
Shift in 50−year
Occupancy Probability
0.10%
0.05%
0.00%
−0.05%

*Cupido comyntas*

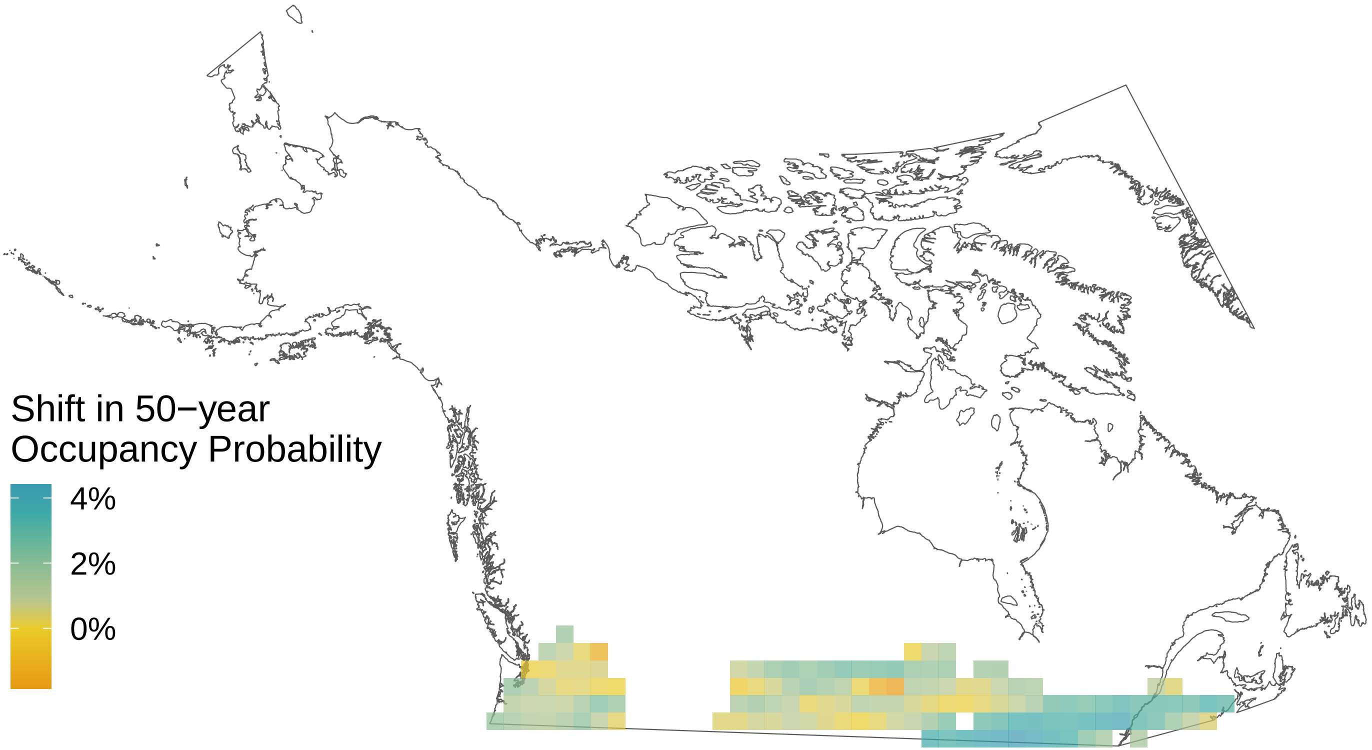

*Cupido amyntula*

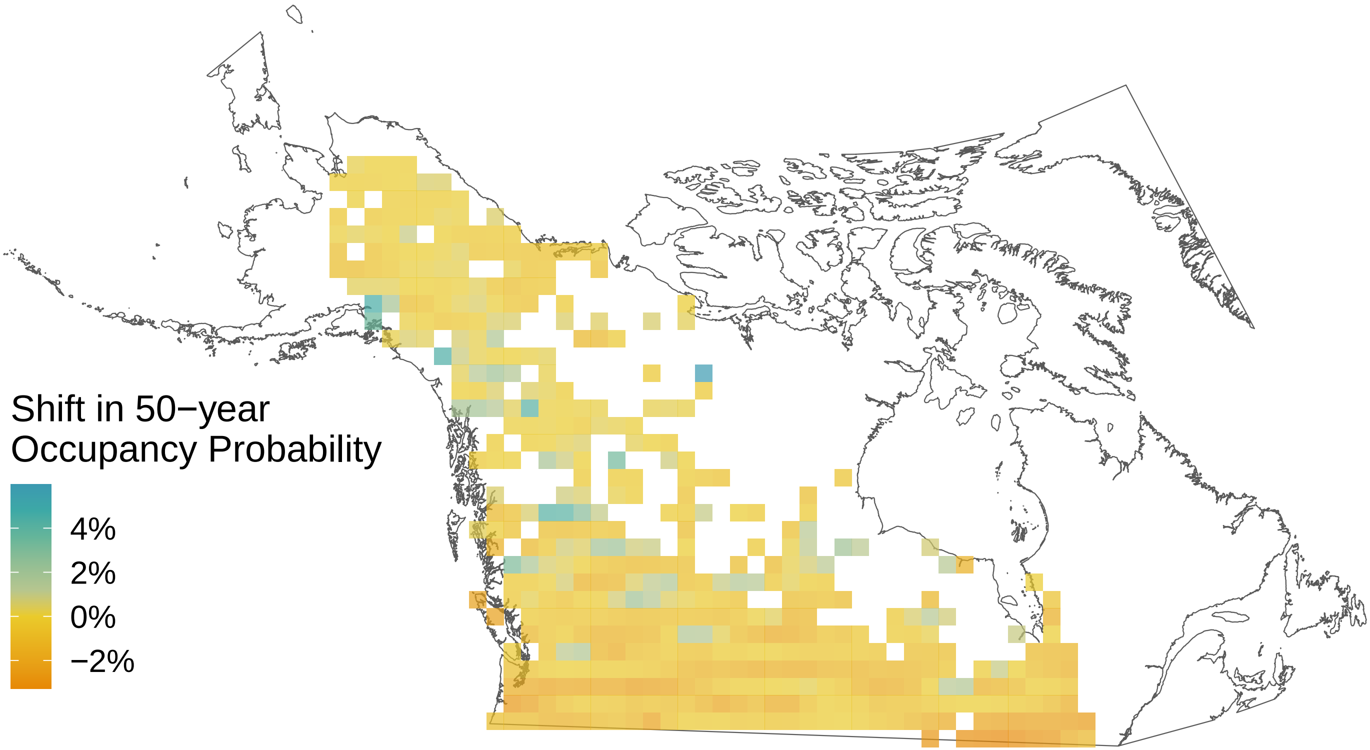

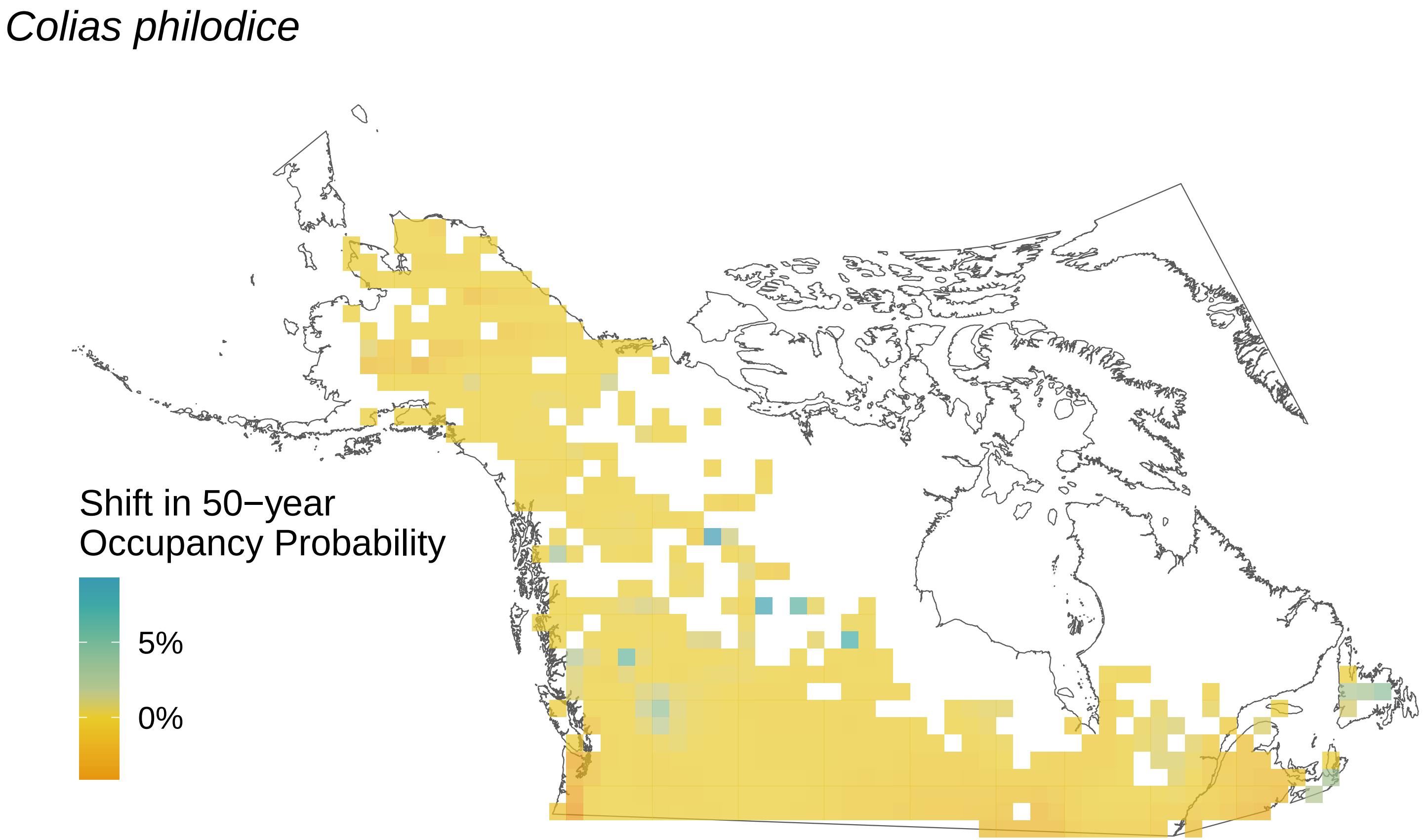
Colias philodice
Shift in 50−year
Occupancy Probability
5%
0%

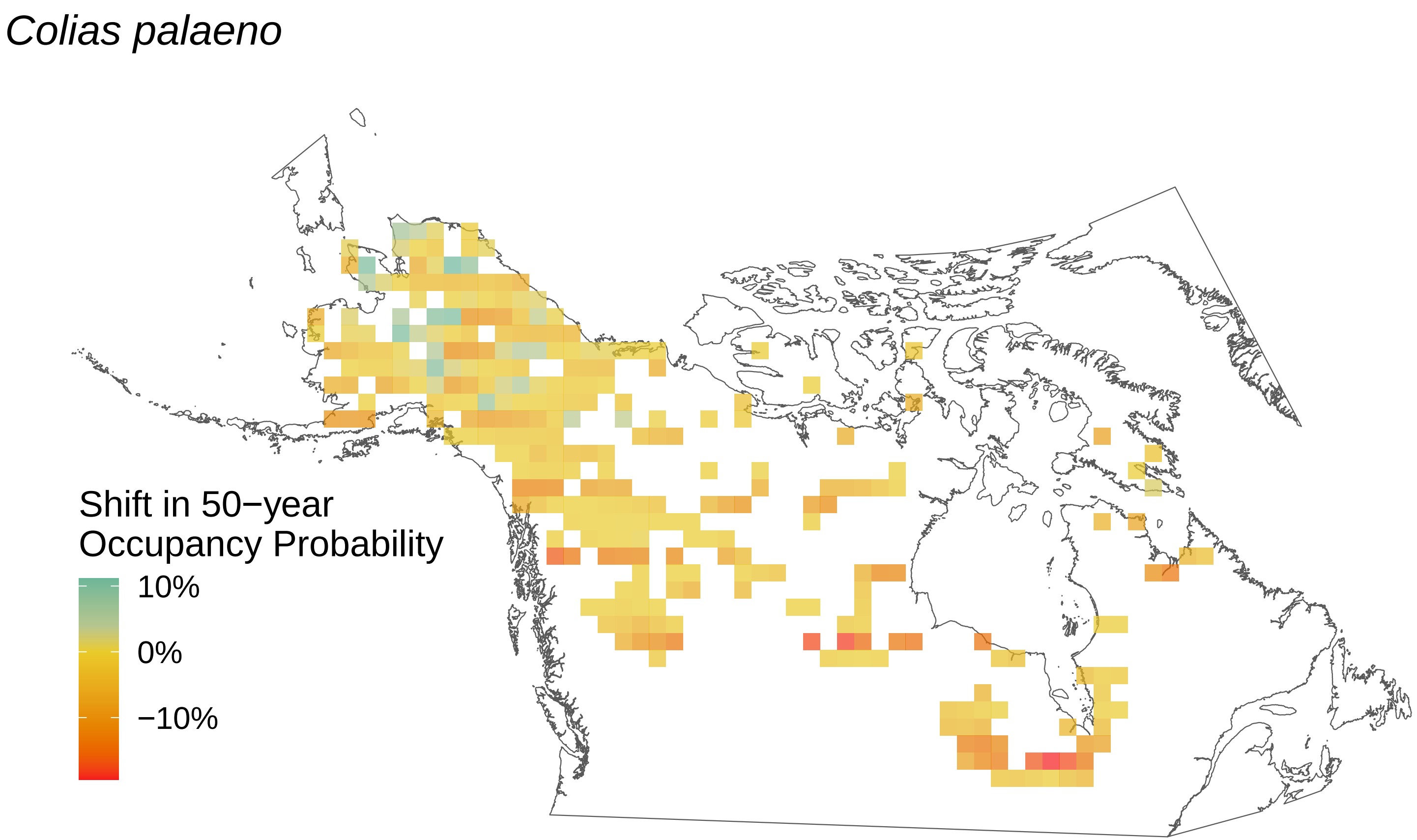
Colias palaeno
Shift in 50−year
Occupancy Probability
10%
0%
−10%

*Colias interior*

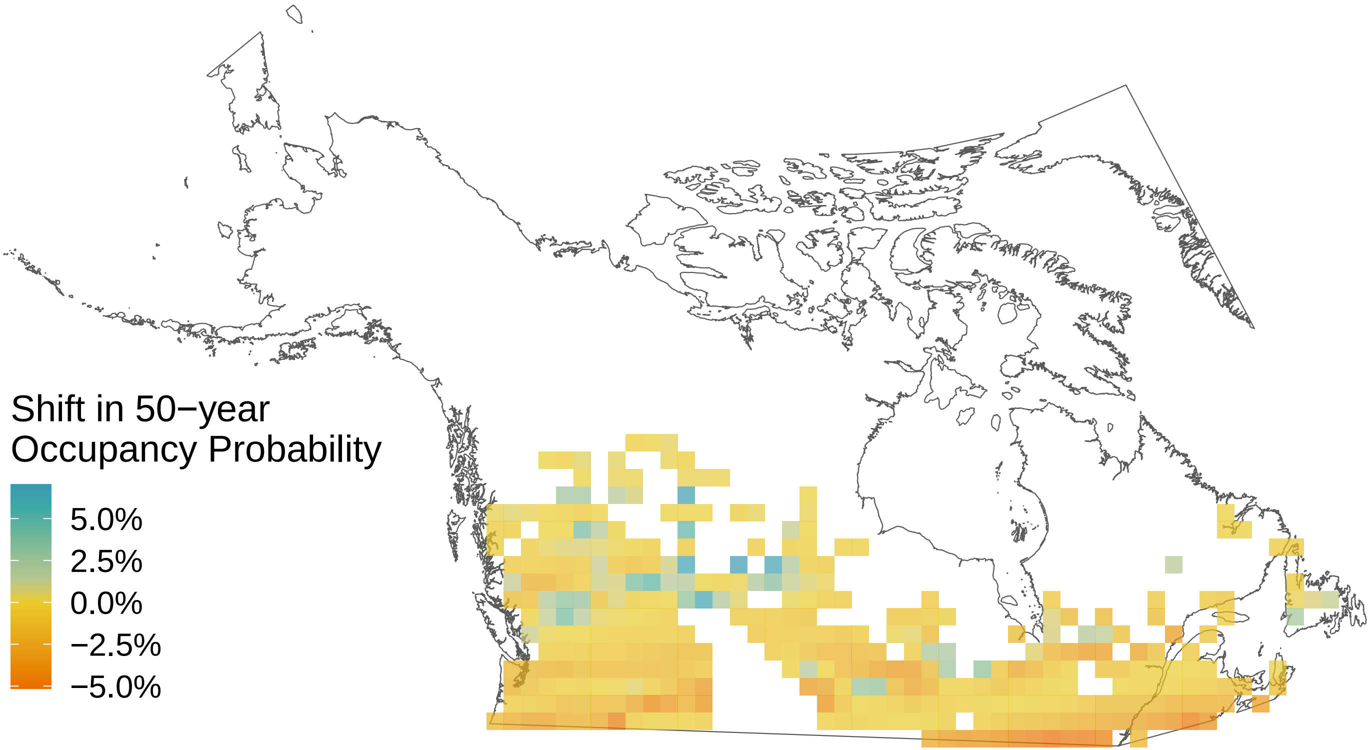

*Colias hecla*

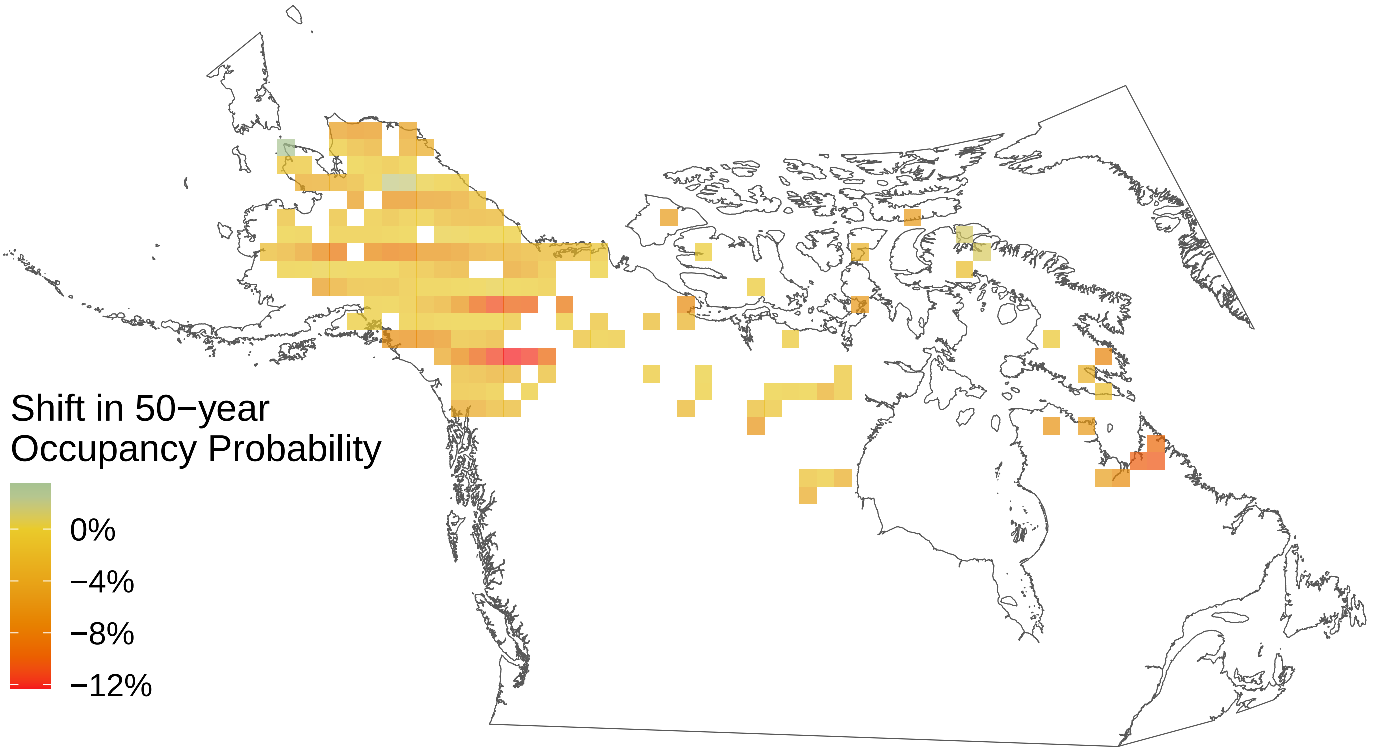

*Colias eurytheme*

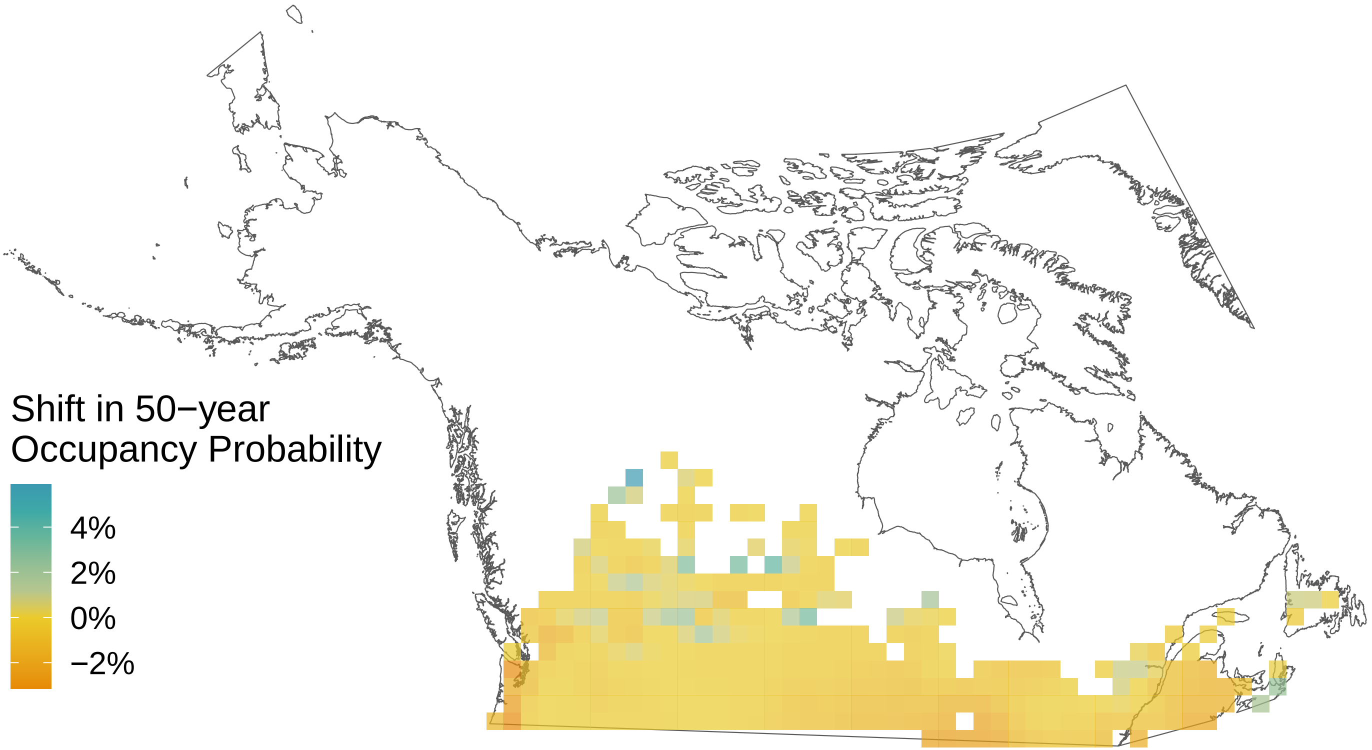

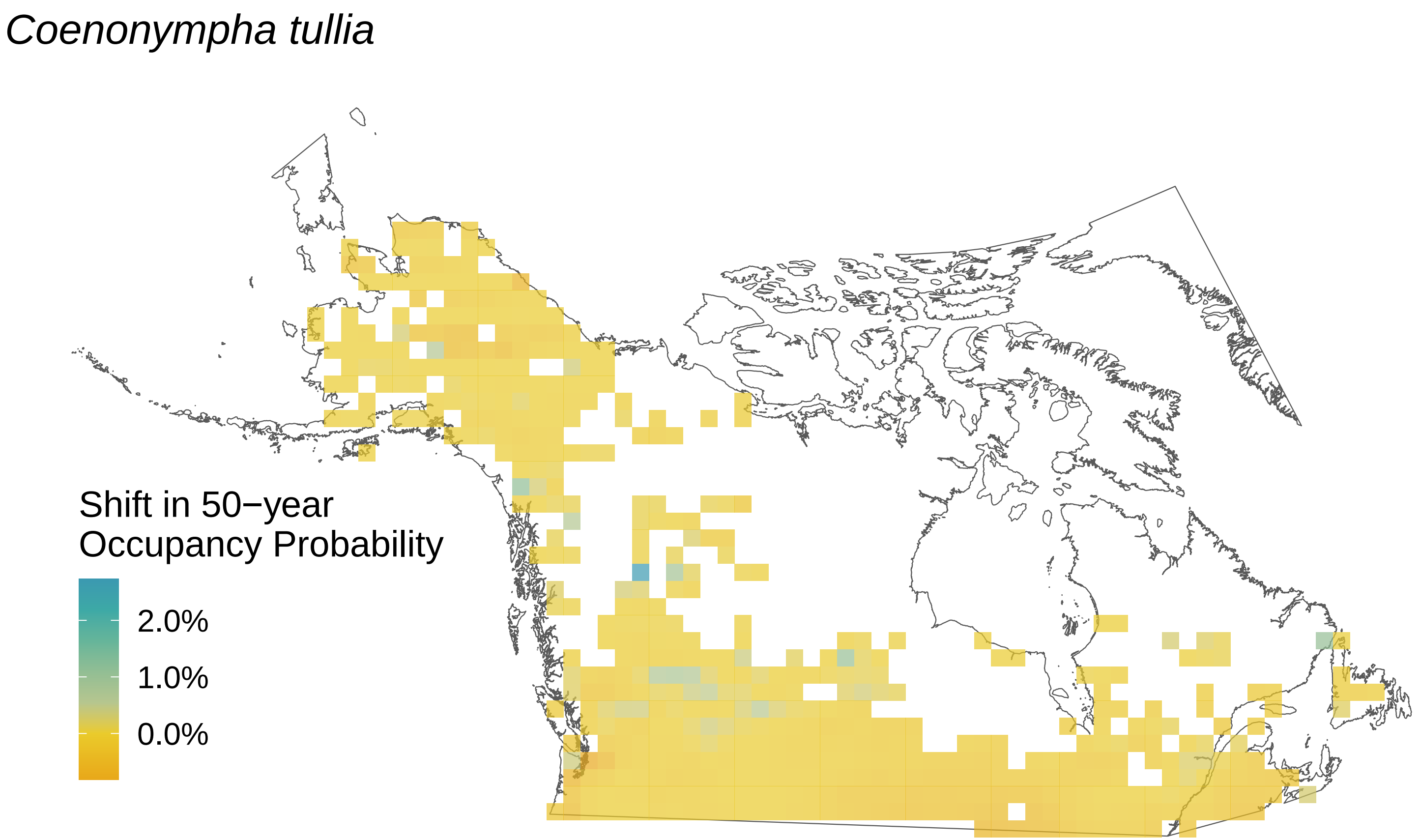
*Coenonympha tullia*
Shift in 50−year
Occupancy Probability
2.0%
1.0%
0.0%

*Chlosyne harrisii*

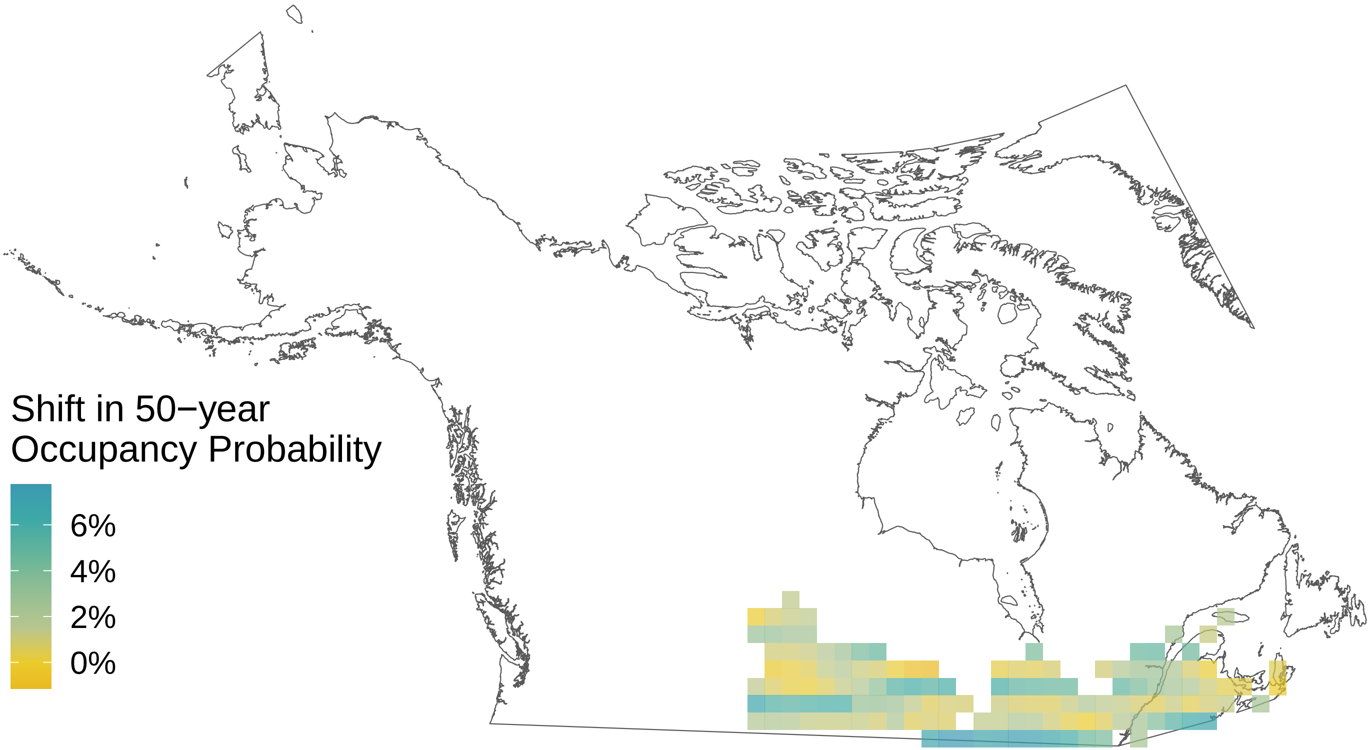

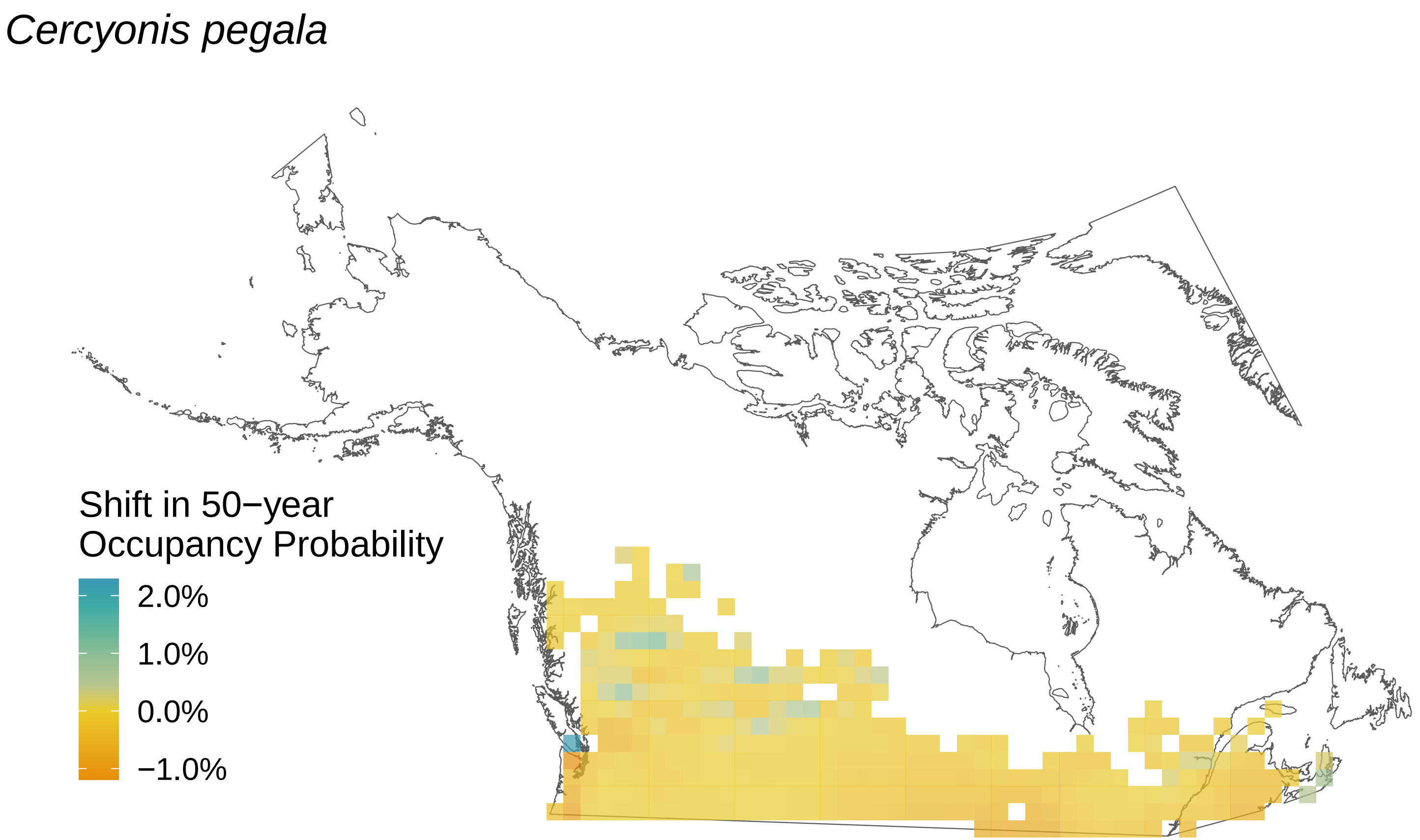

*Cercyonis pegala*
Shift in 50−year
Occupancy Probability
2.0%
1.0%
0.0%
−1.0%

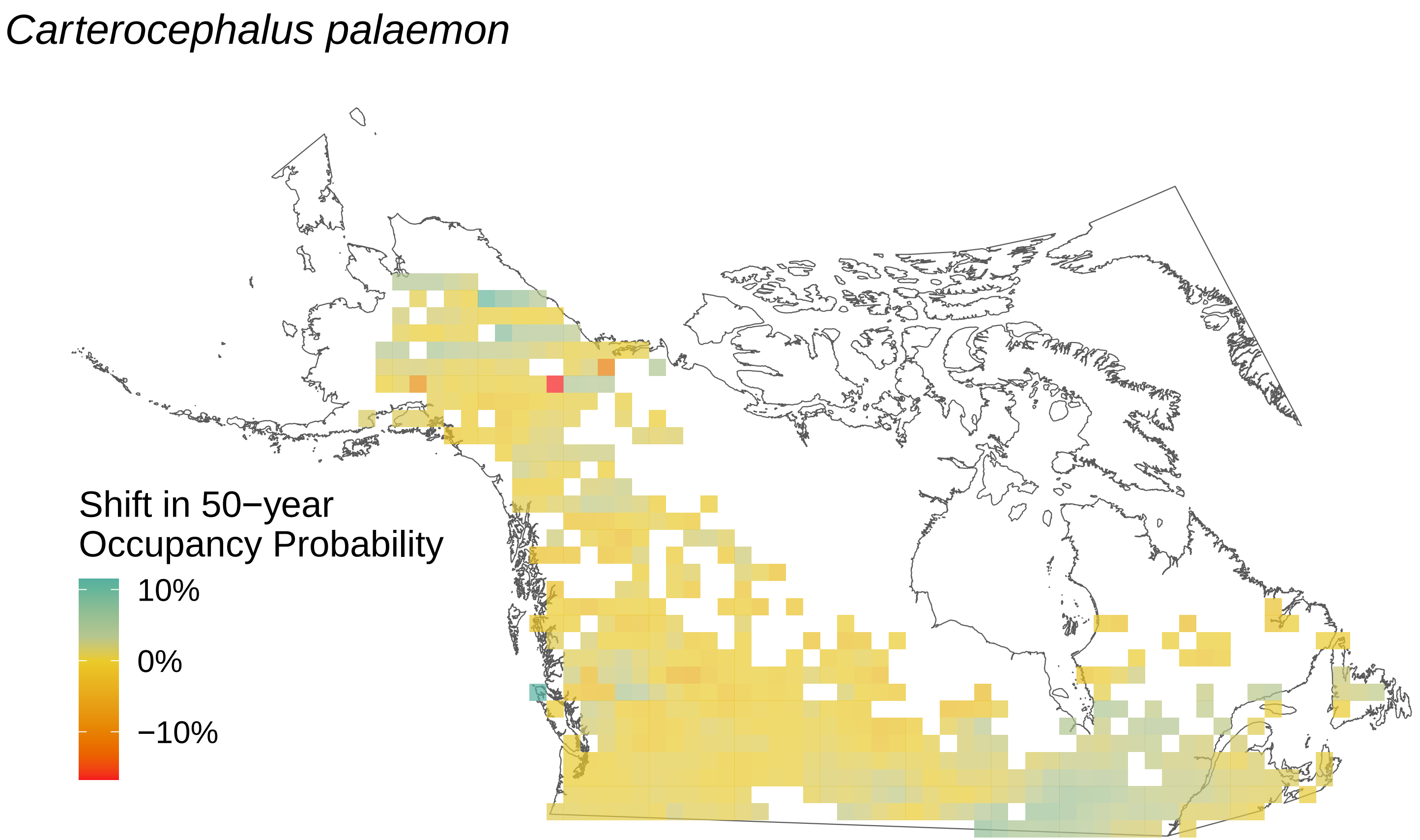
*Carterocephalus palaemon*
Shift in 50−year
Occupancy Probability
10%
0%
−10%

*Callophrys polios*

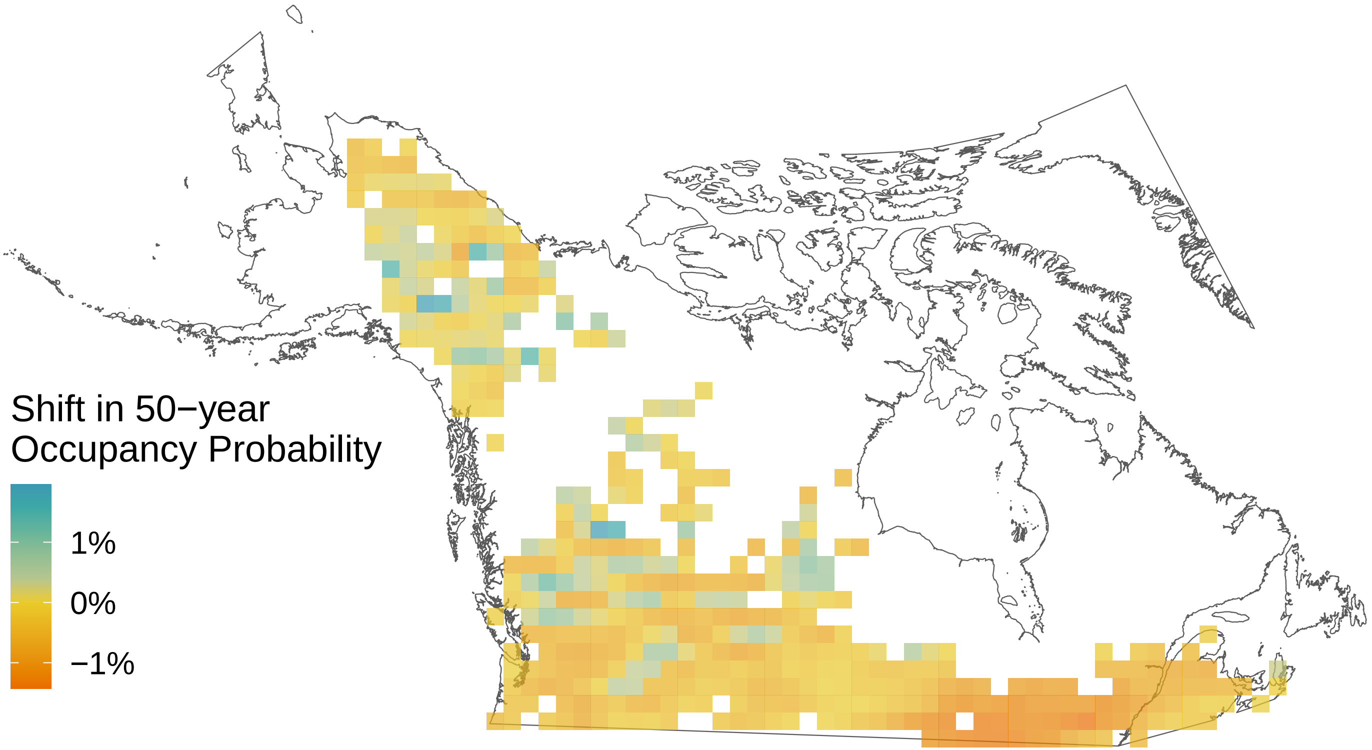

*Callophrys niphon*

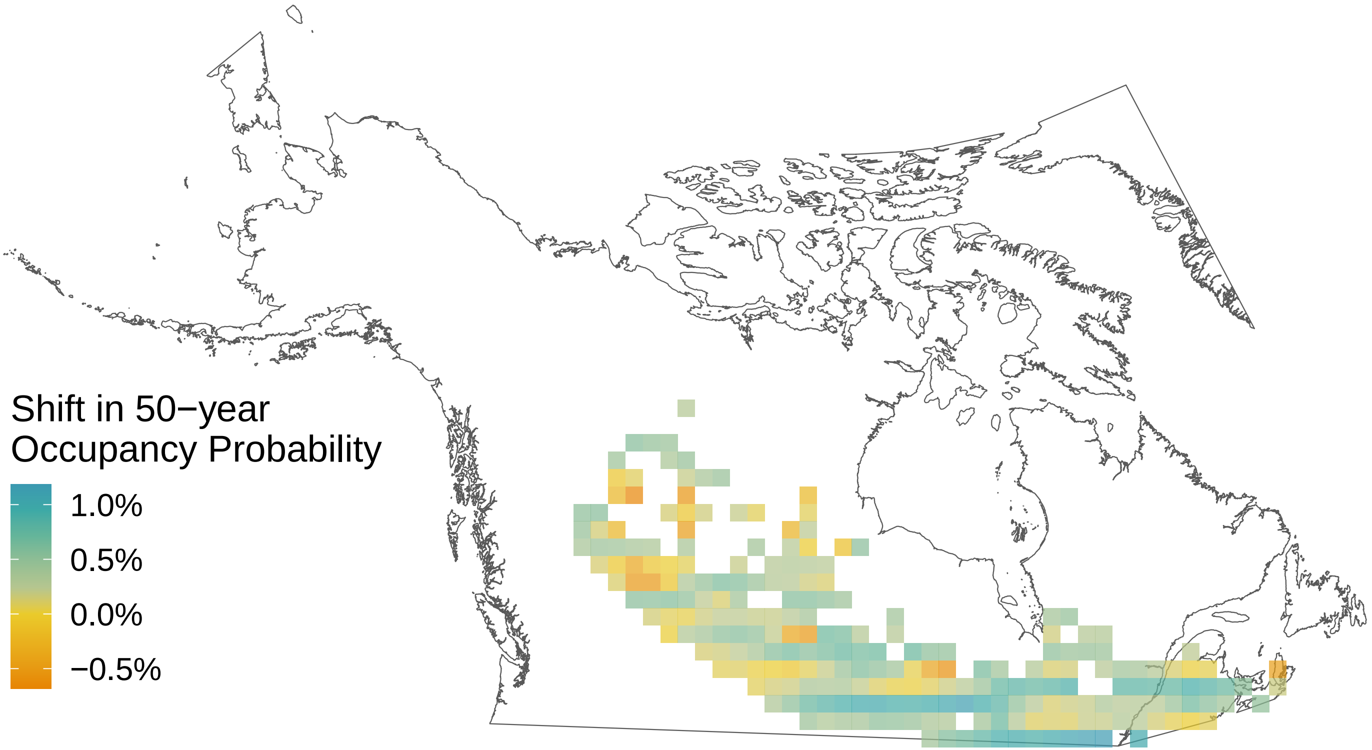

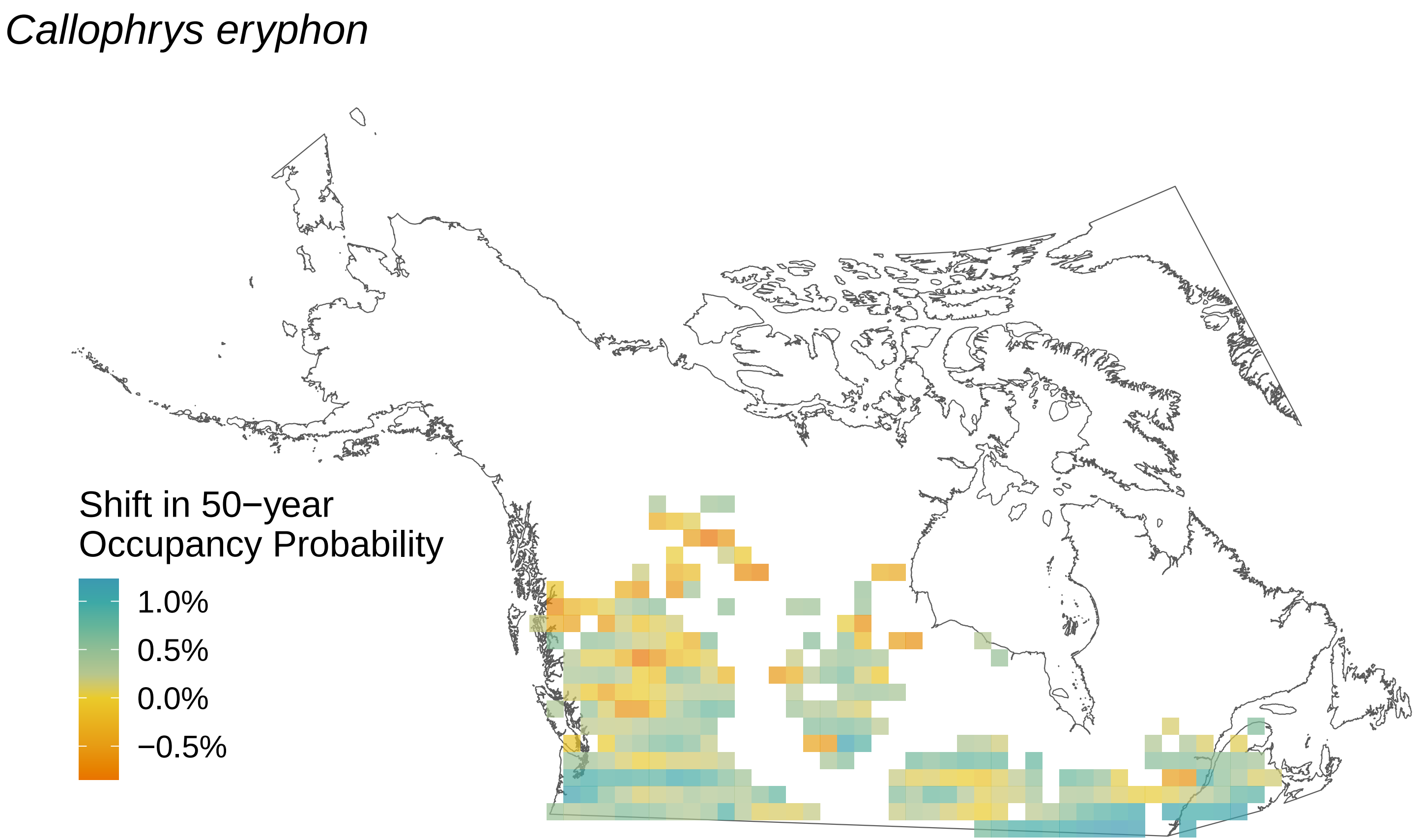
Callophrys eryphon
Shift in 50−year
Occupancy Probability
1.0%
0.5%
0.0%
−0.5%

*Callophrys augustinus*

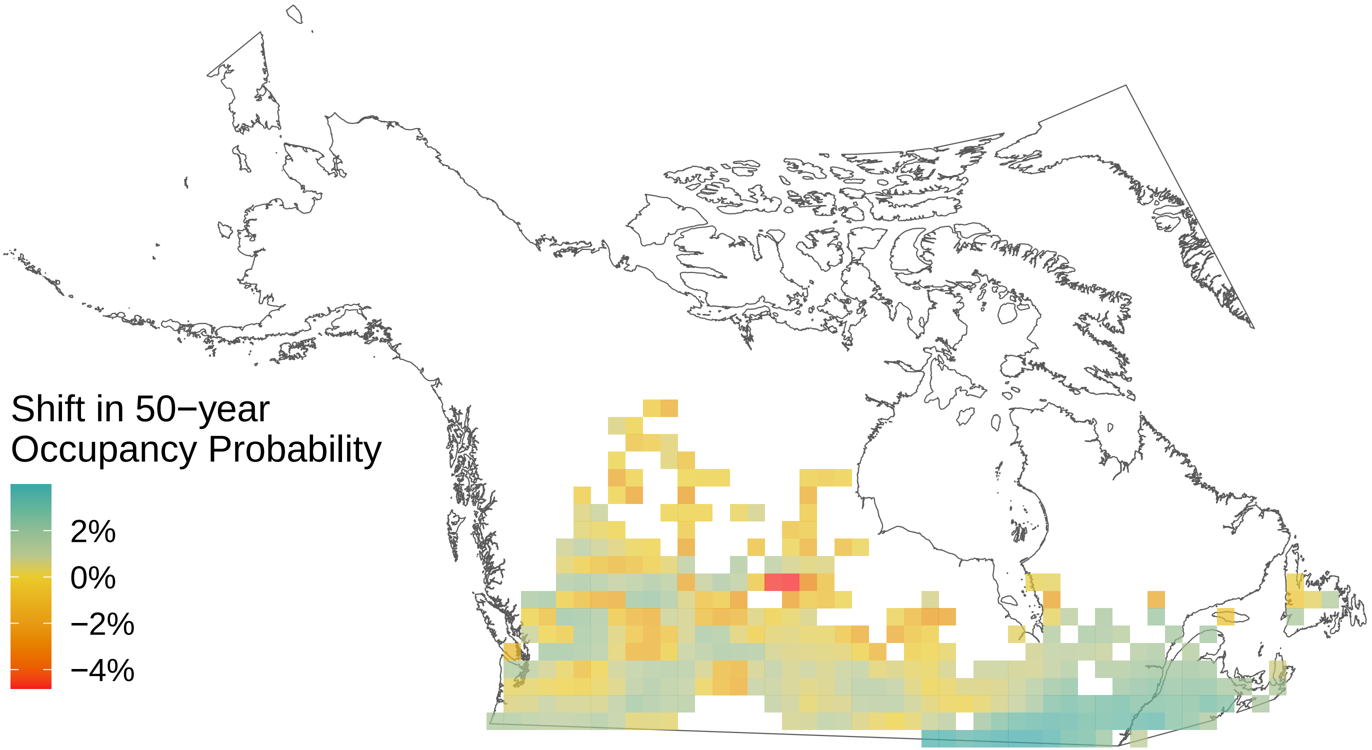

*Boloria selene*

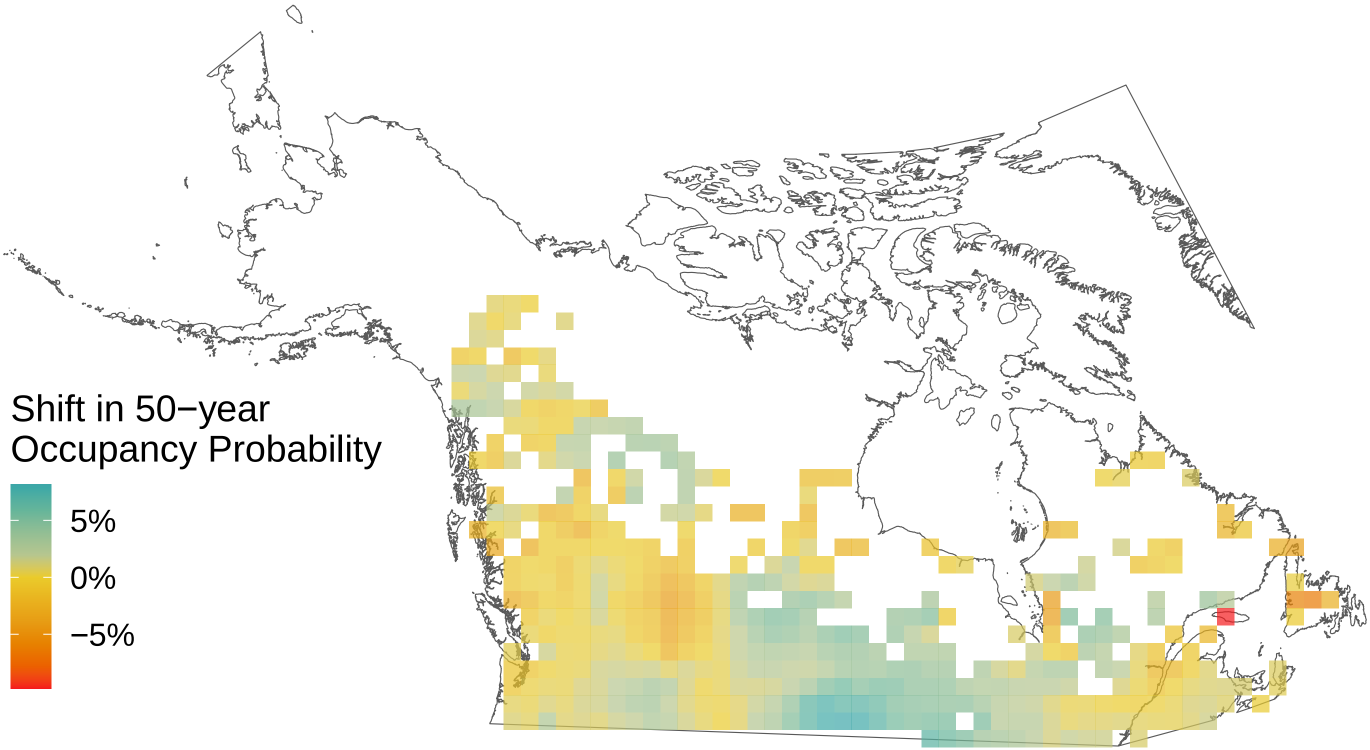

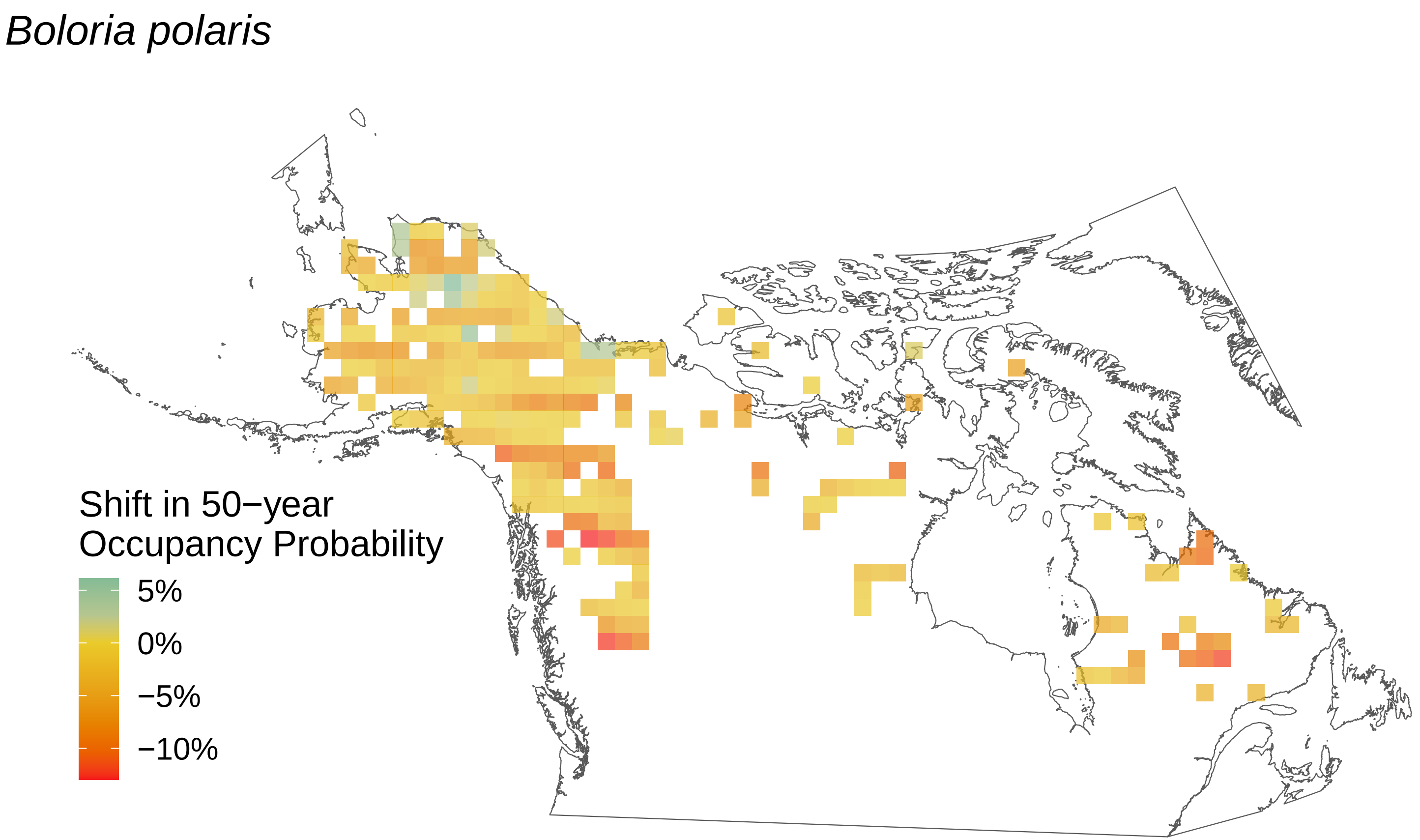
Boloria polaris
Shift in 50−year
Occupancy Probability
5%
0%
−5%
−10%

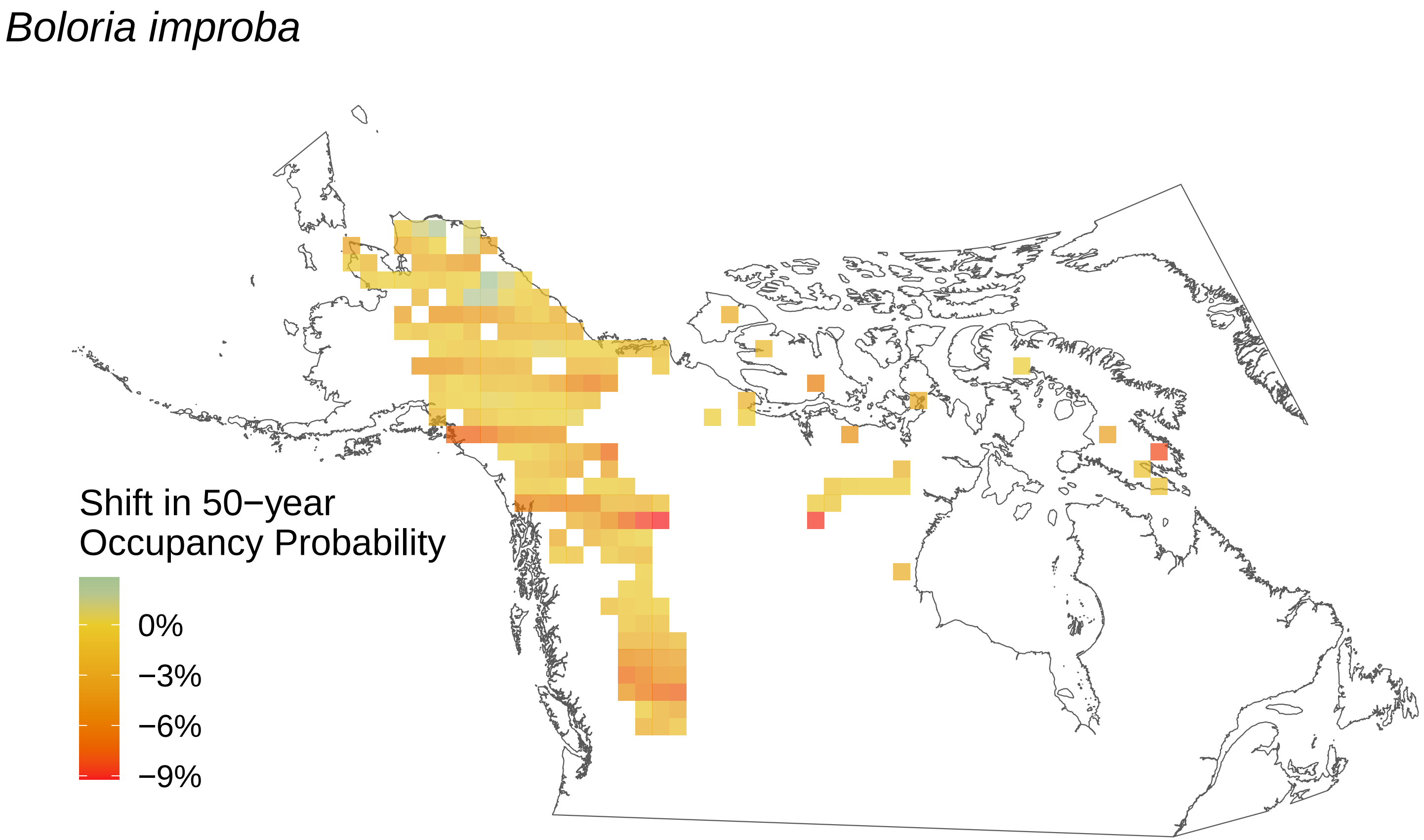
Boloria improba
Shift in 50−year Occupancy Probability
0%
−3%
−6%
−9%

### *Boloria frigga*

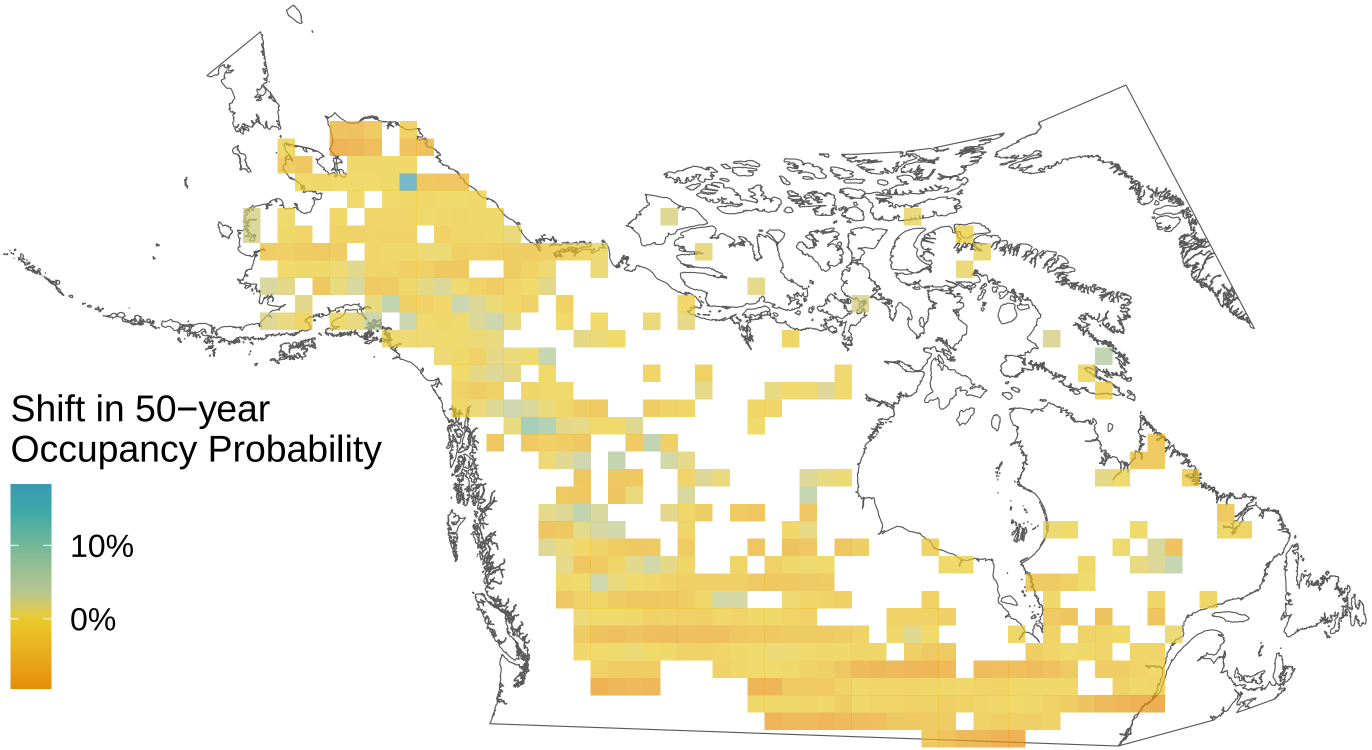

### *Boloria freija*

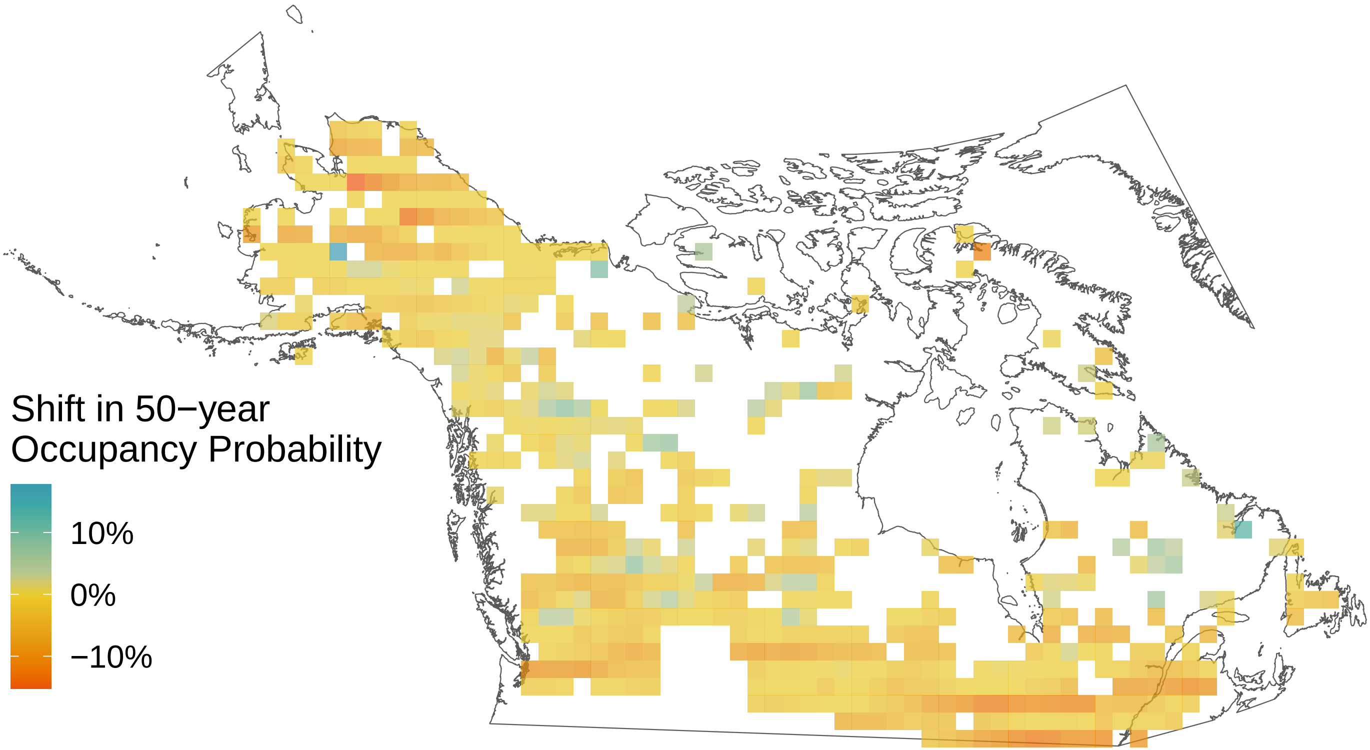

*Boloria eunomia*

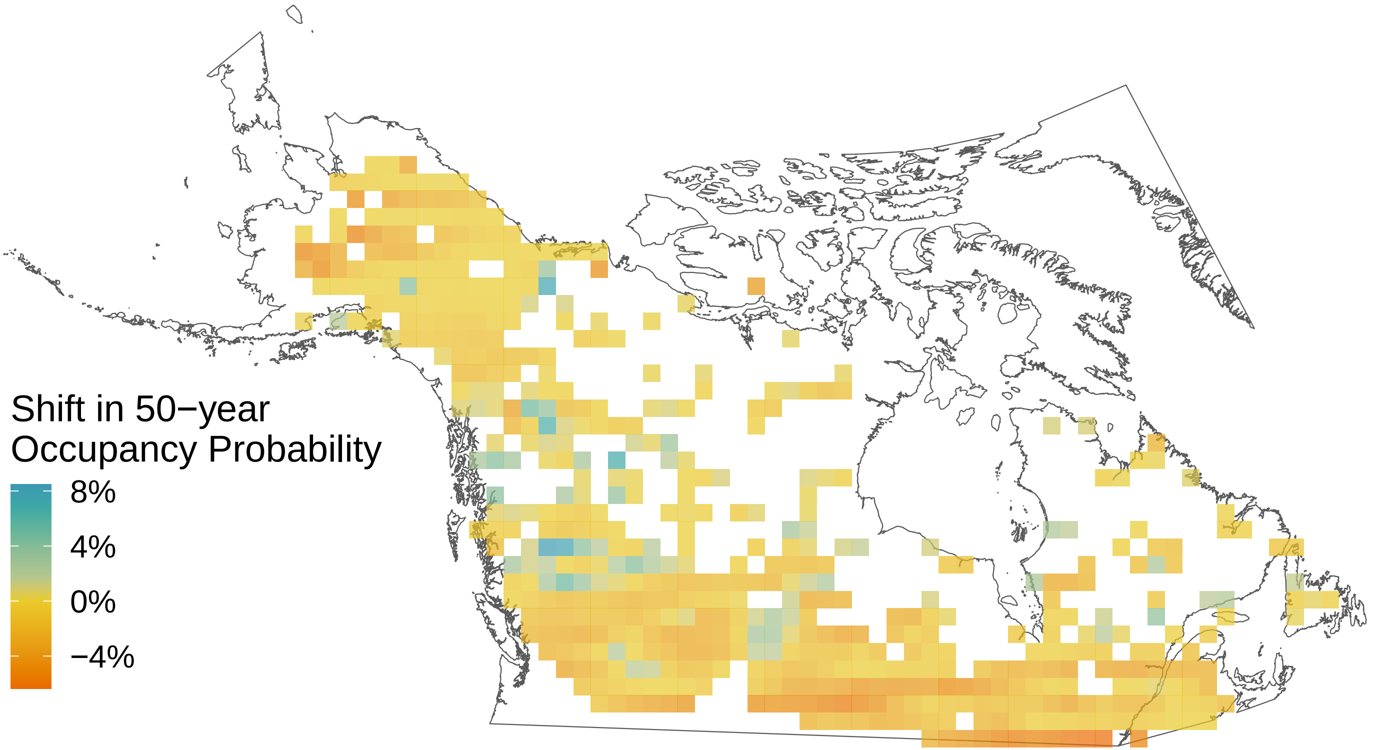

*Boloria chariclea*

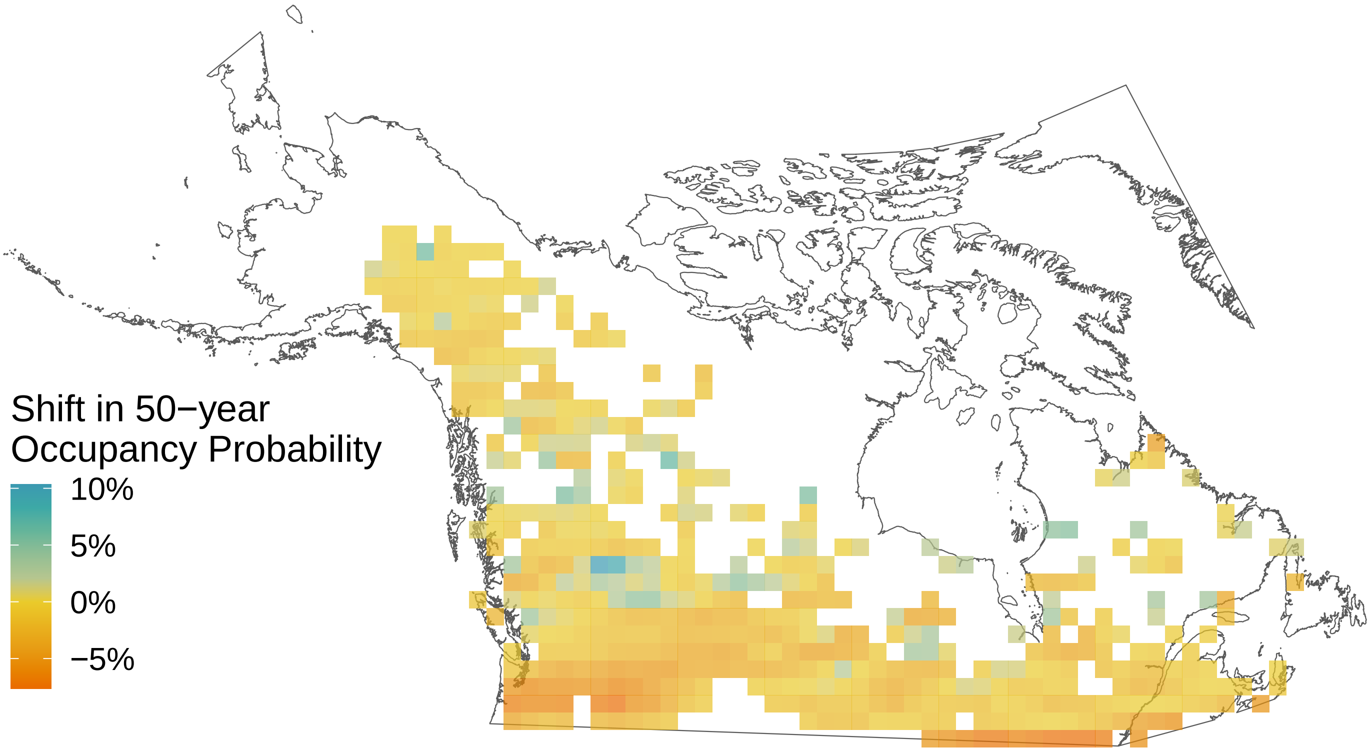

*Boloria bellona*

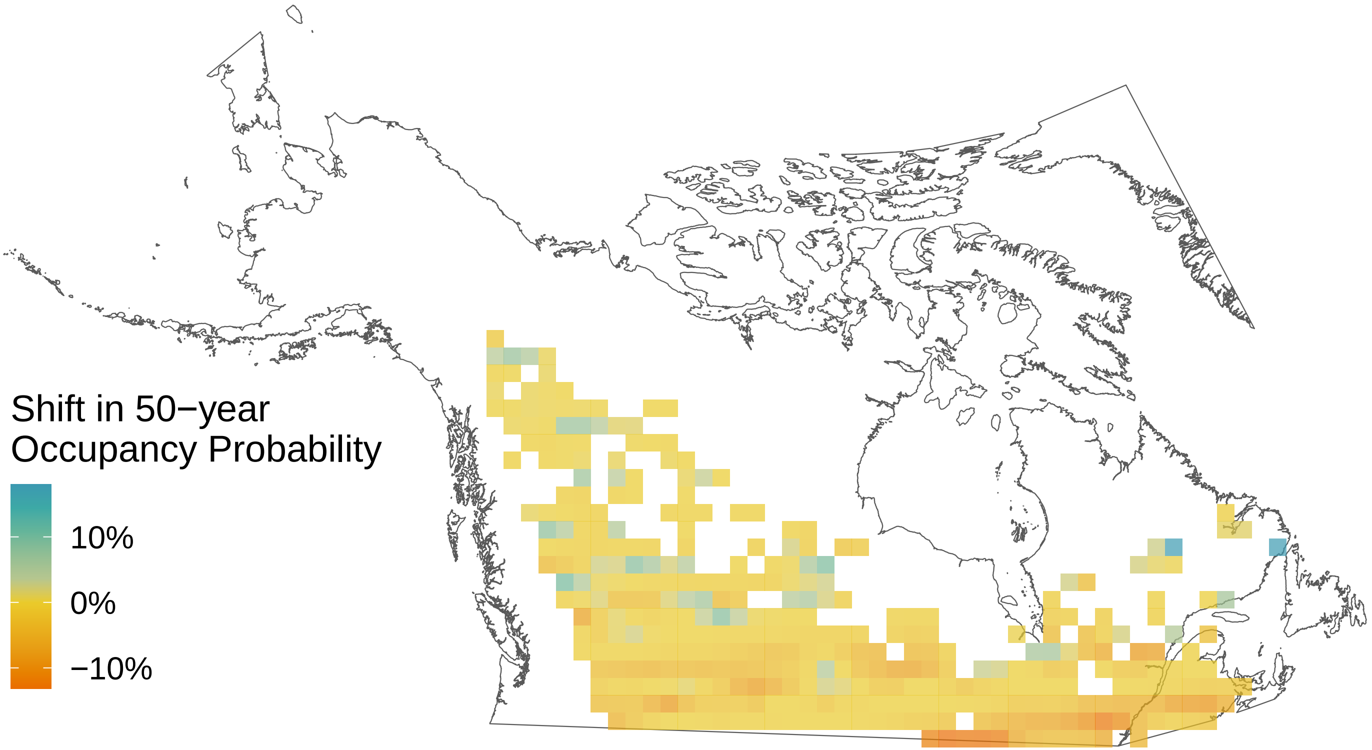

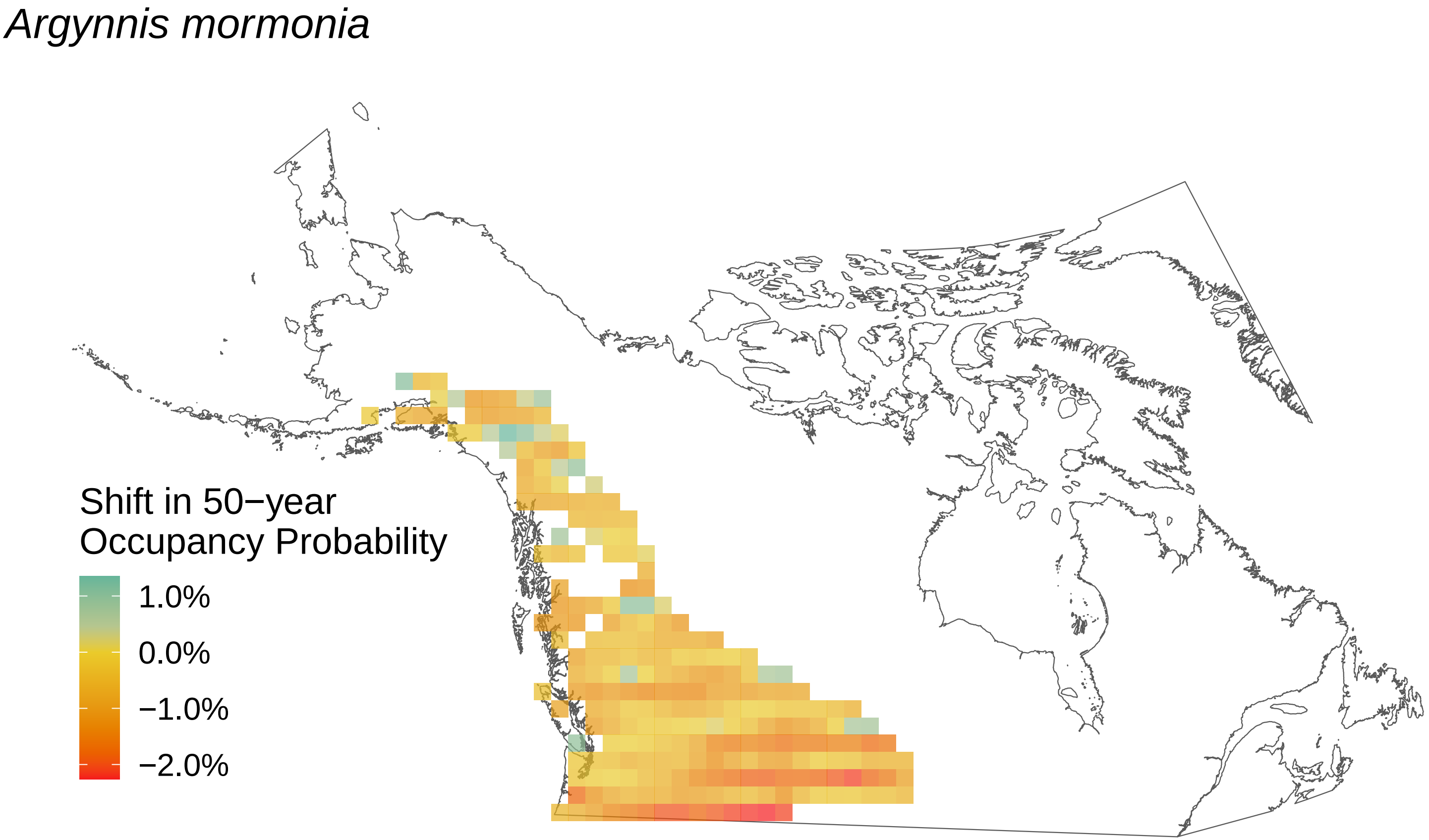
Argynnis mormonia
Shift in 50−year
Occupancy Probability
1.0%
0.0%
−1.0%
−2.0%

*Argynnis hydaspe*

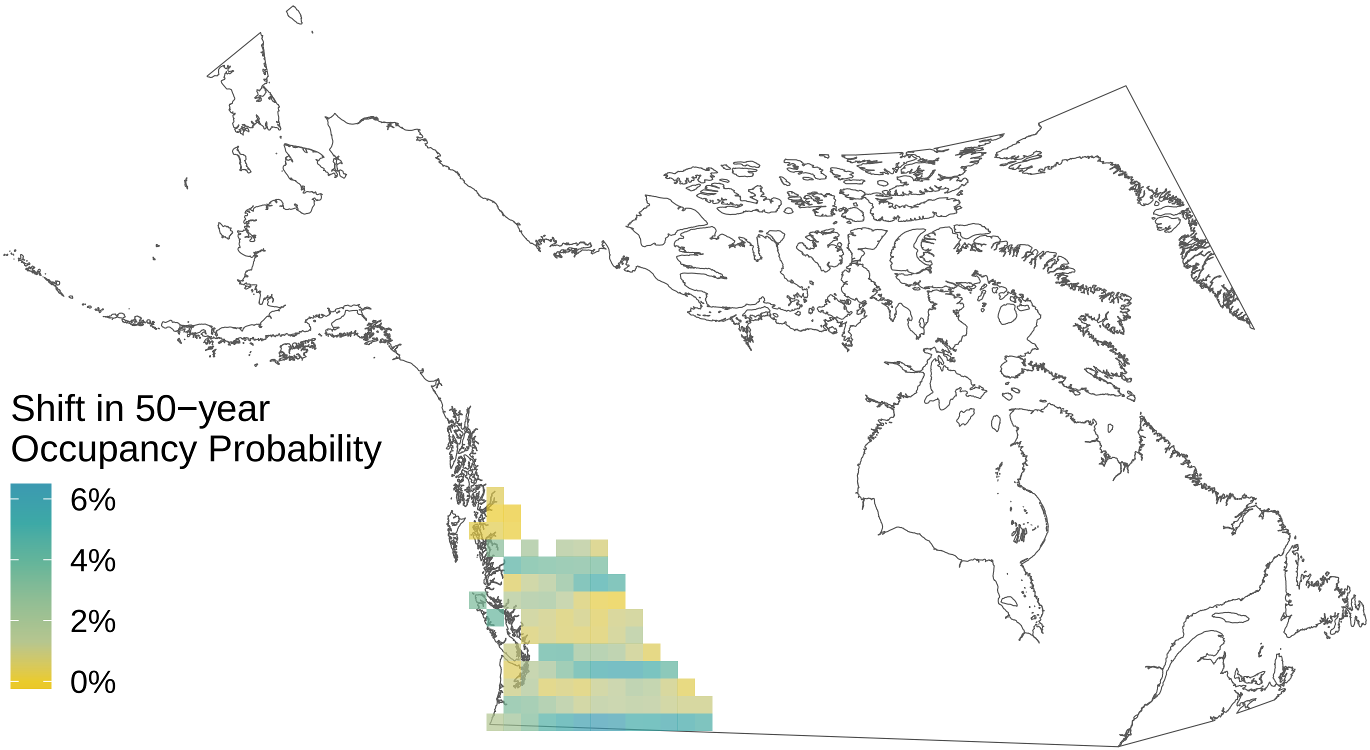

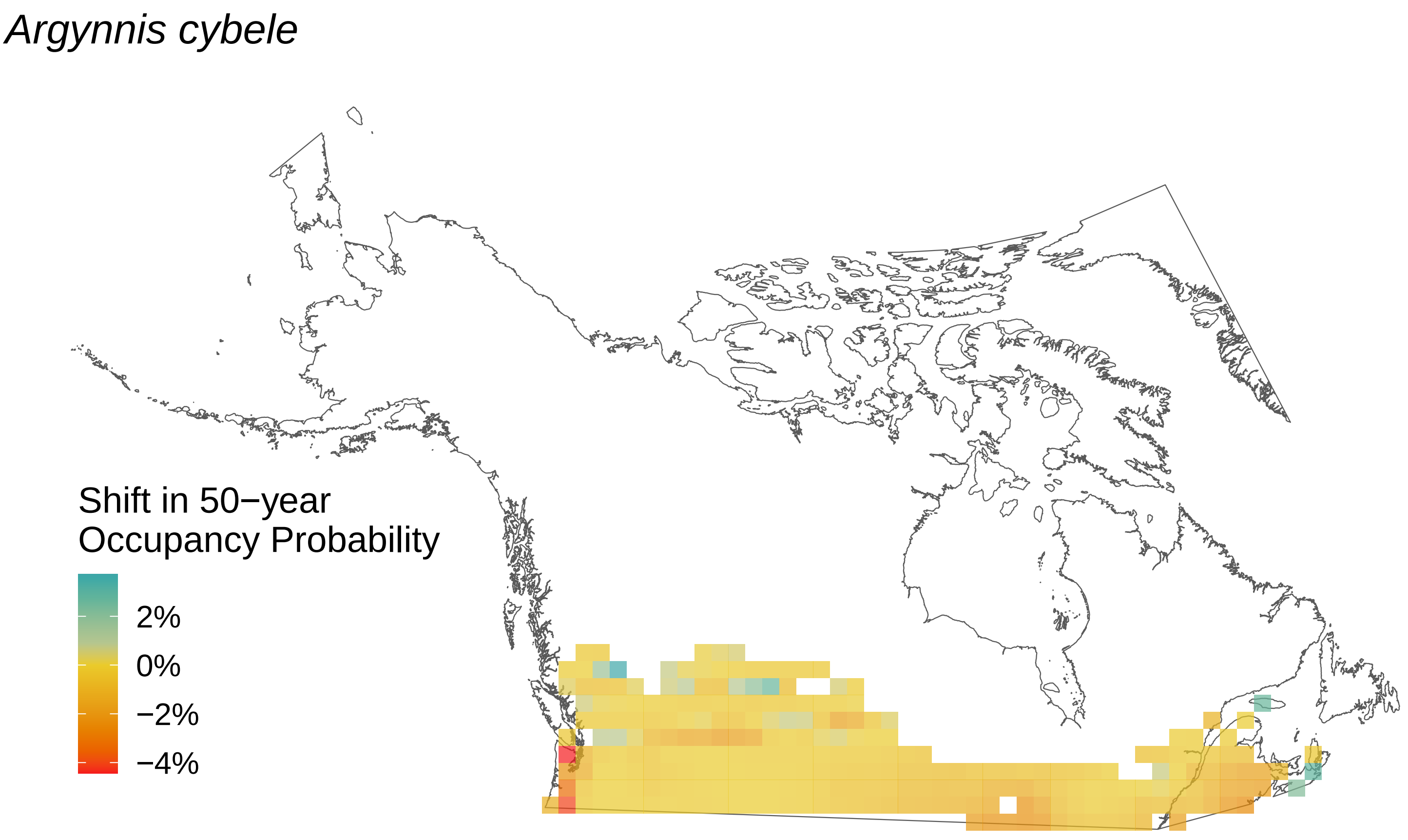
Argynnis cybele
Shift in 50−year
Occupancy Probability
2%
0%
−2%
−4%

*Argynnis atlantis*

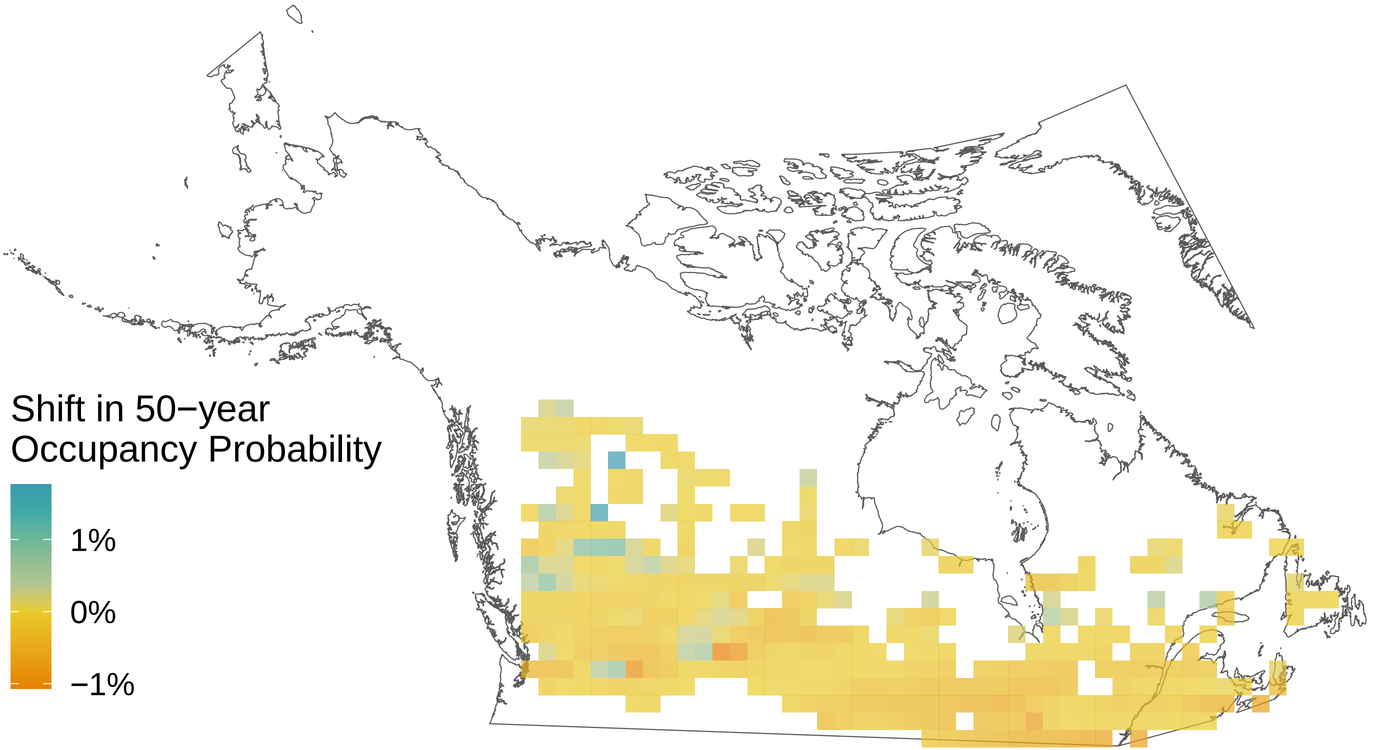

*Ancyloxypha numitor*

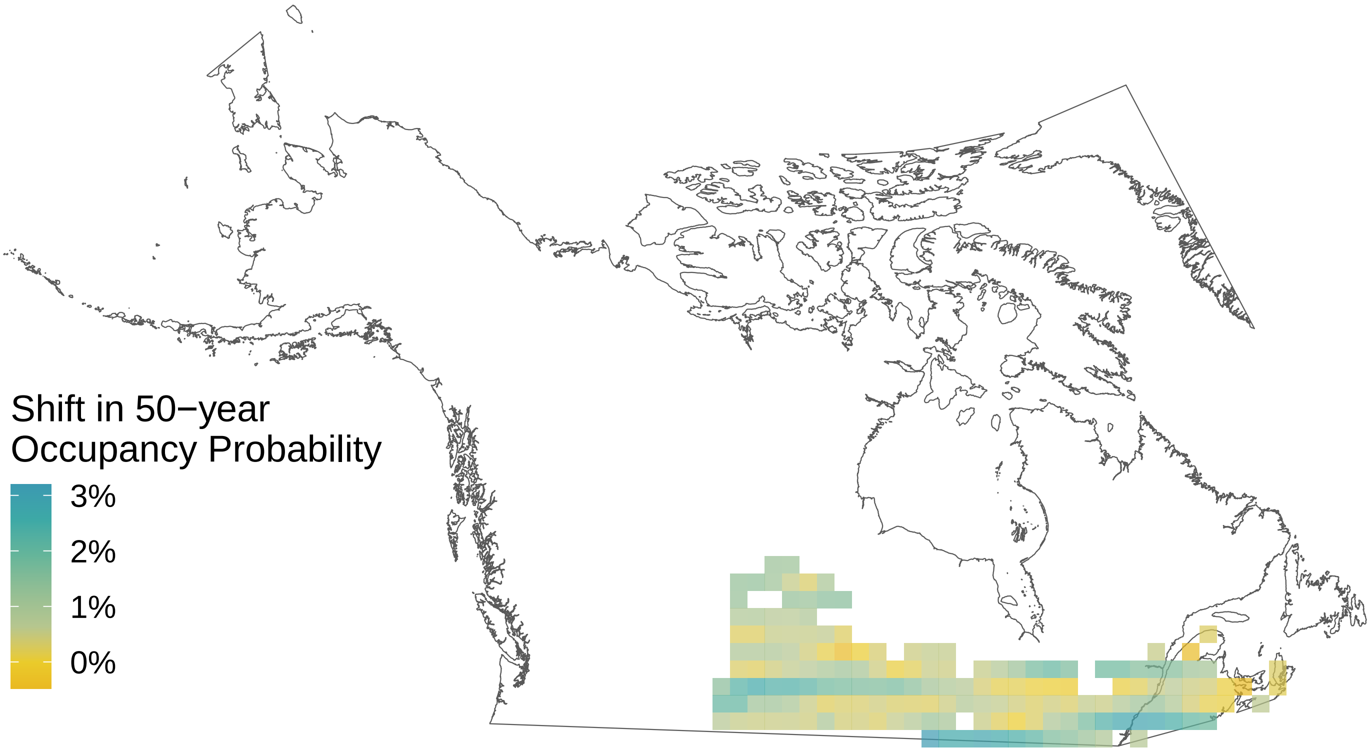

*Amblyscirtes vialis*

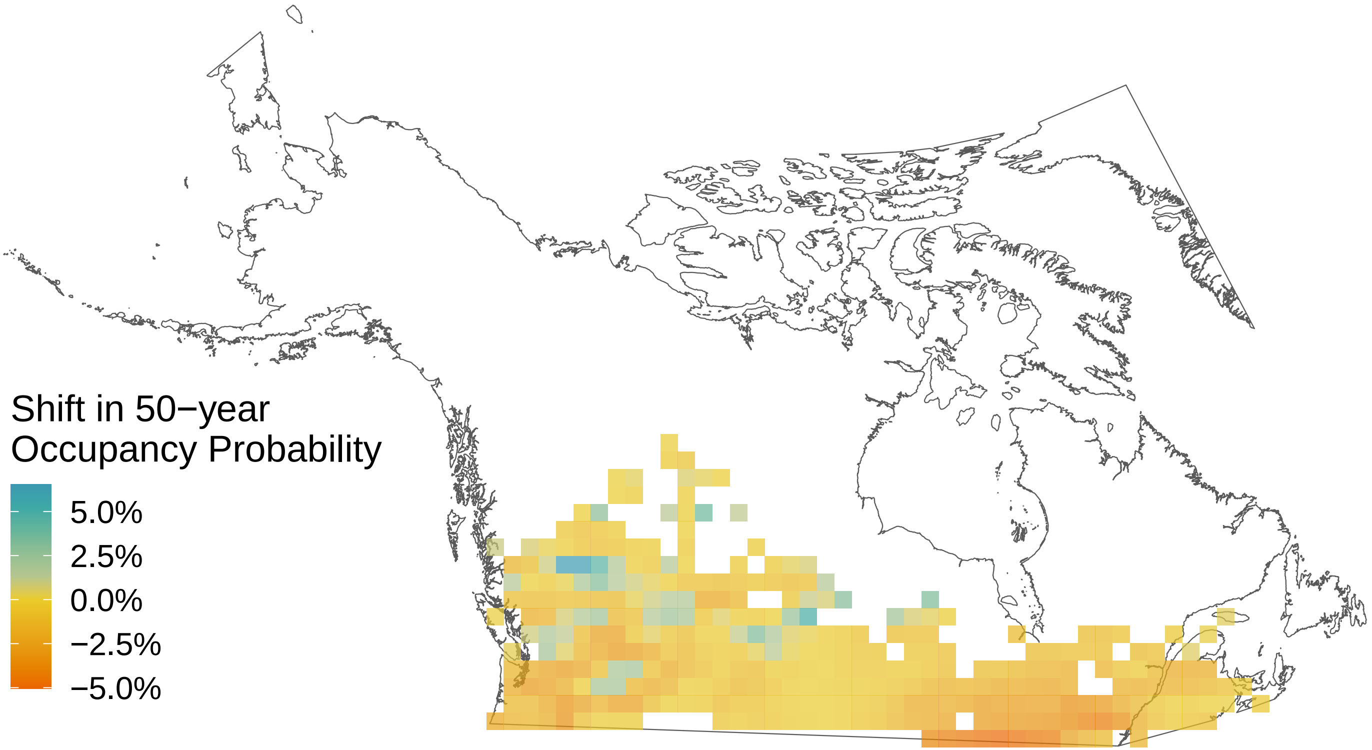

*Agriades glandon*

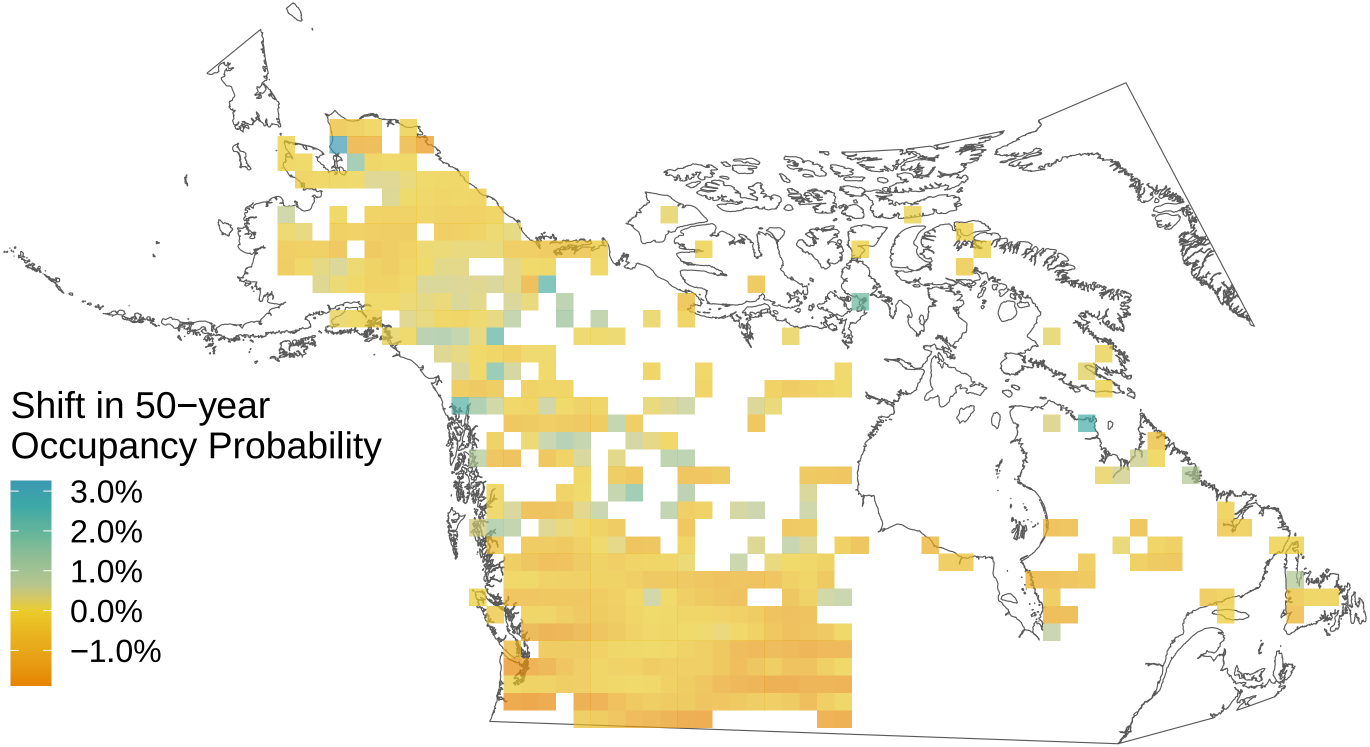

*Aglais milberti*

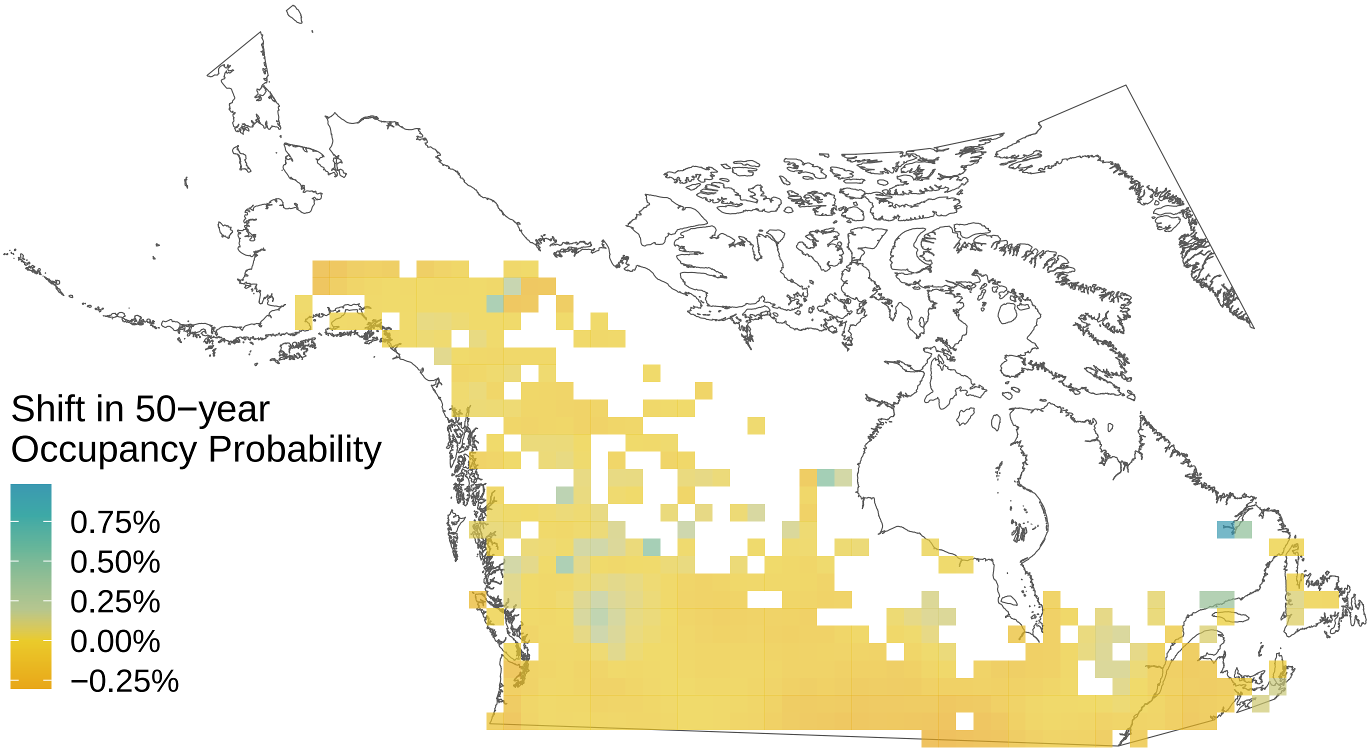
