## Supplemental File S8 for "Rising minimum temperatures contribute to 50 years of shifting Arctic and boreal butterfly communities in North America"

*Vanessa virginiensis*

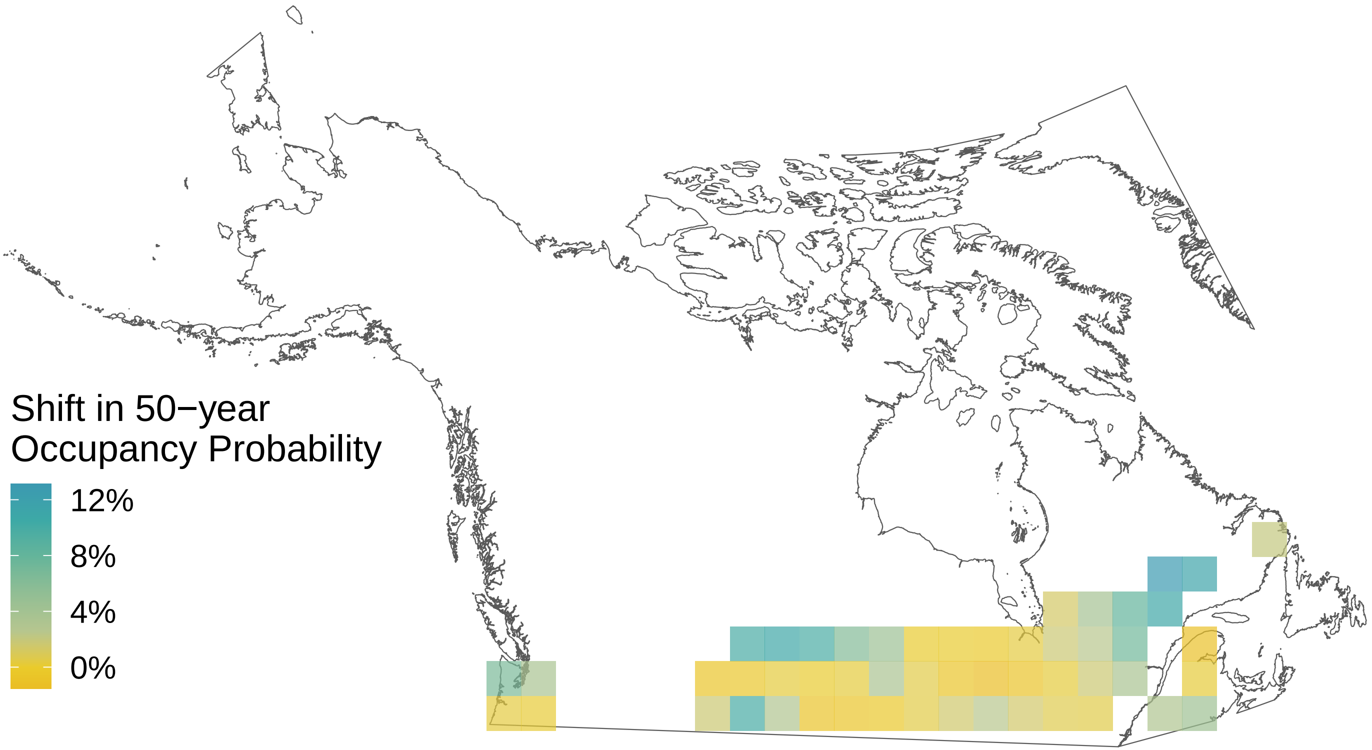

*Thymelicus lineola*

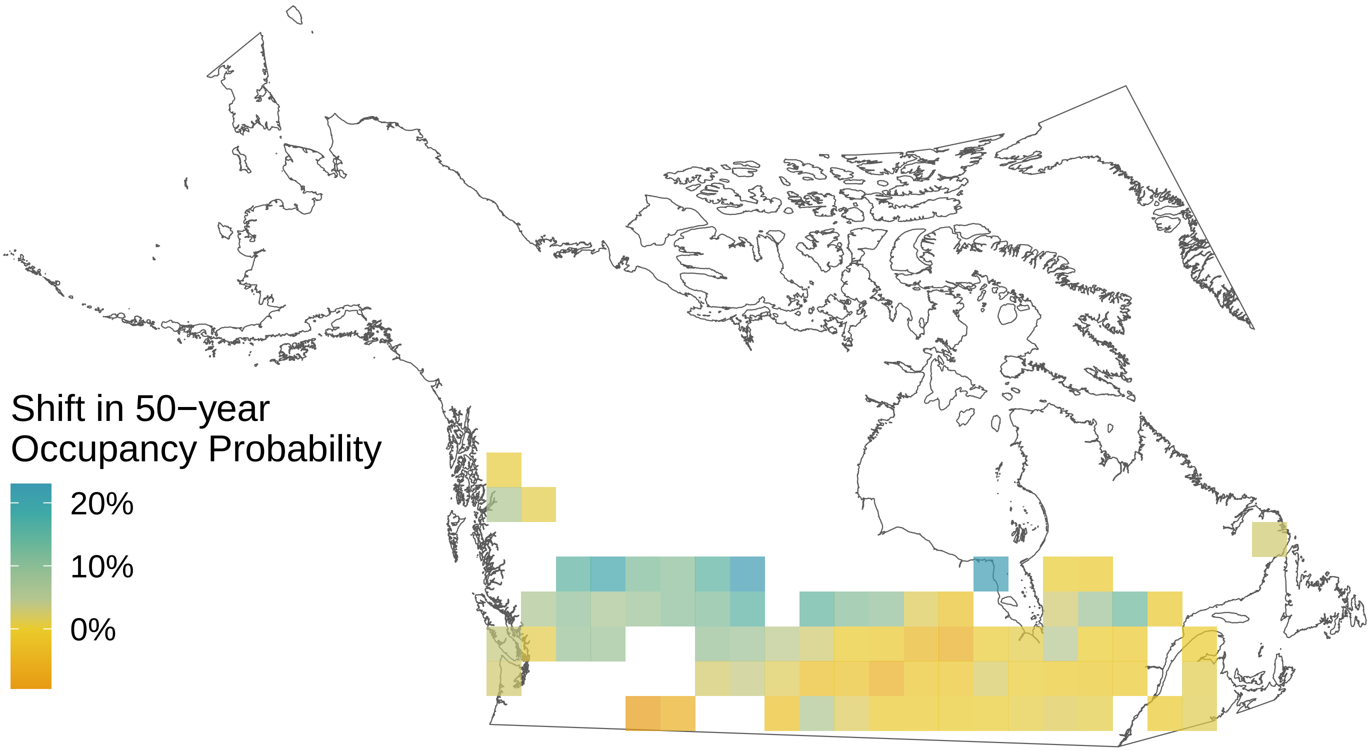

*Thorybes pylades*

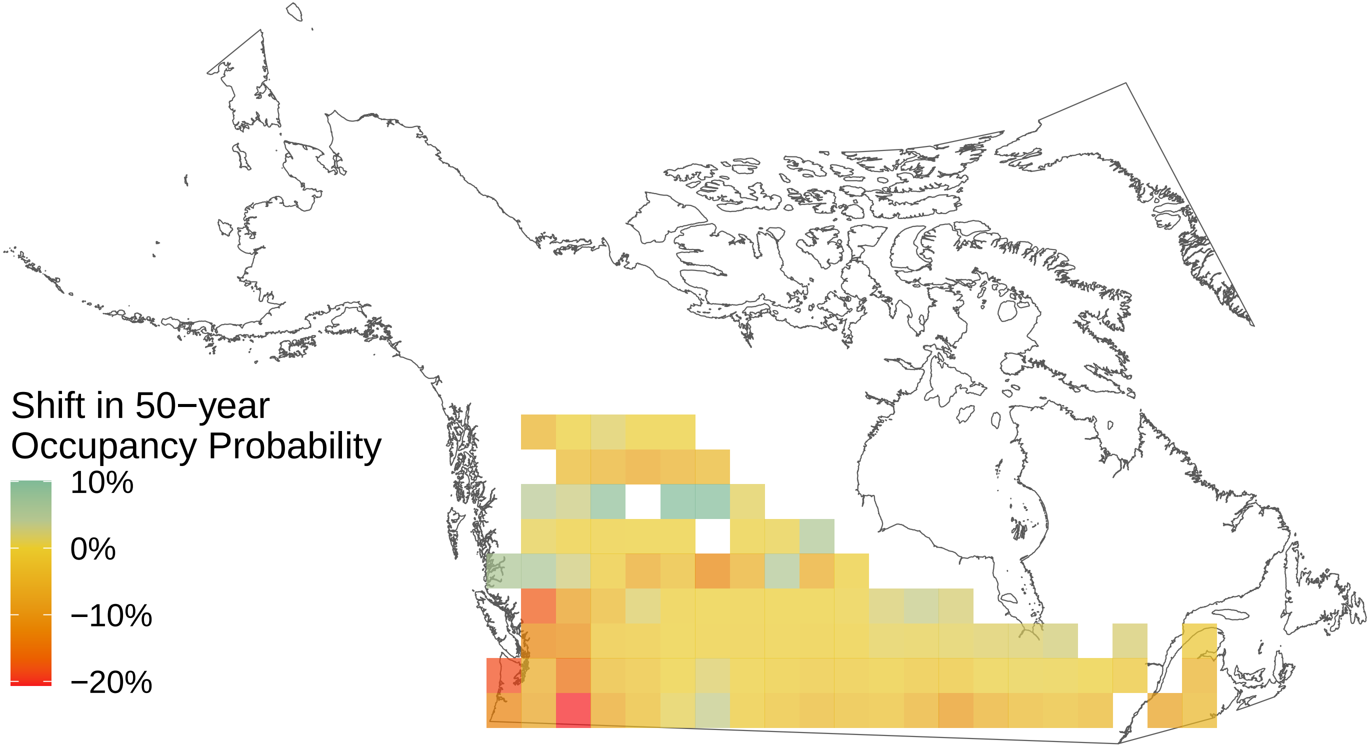

*Strymon melinus*

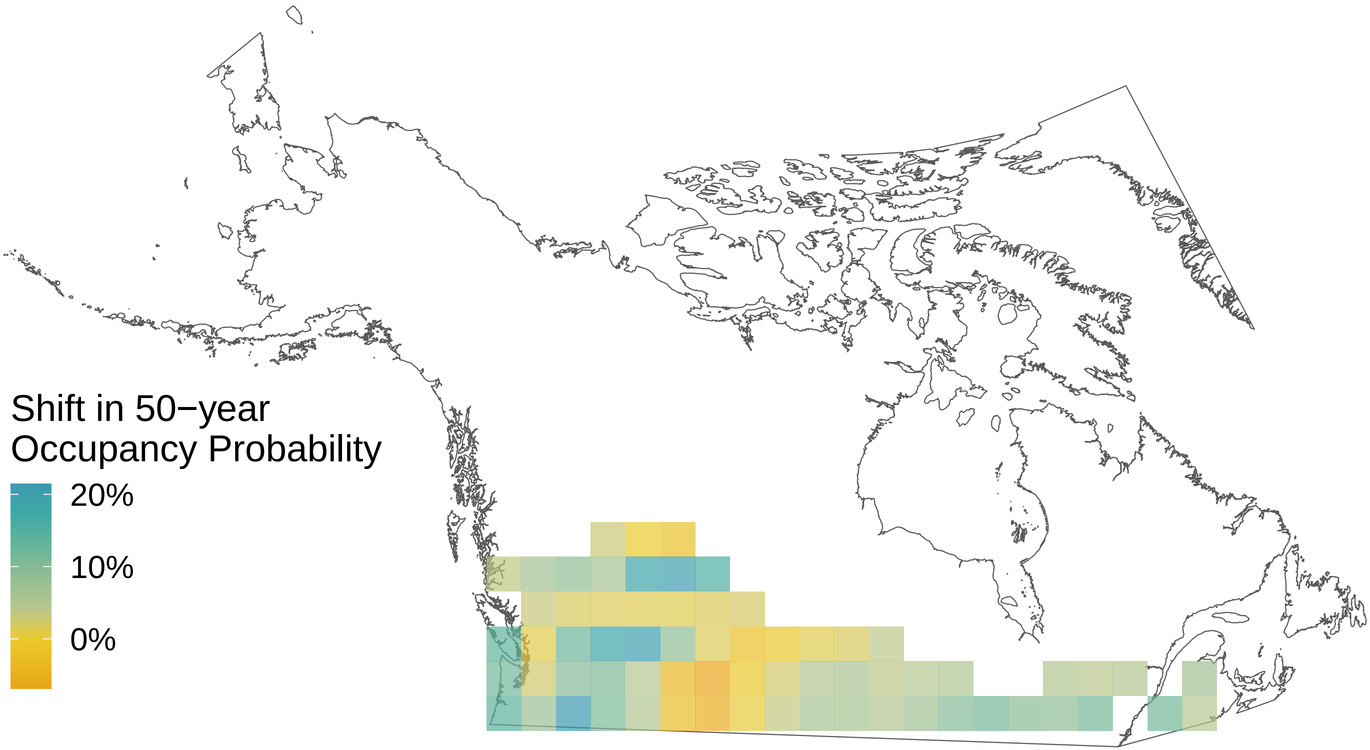

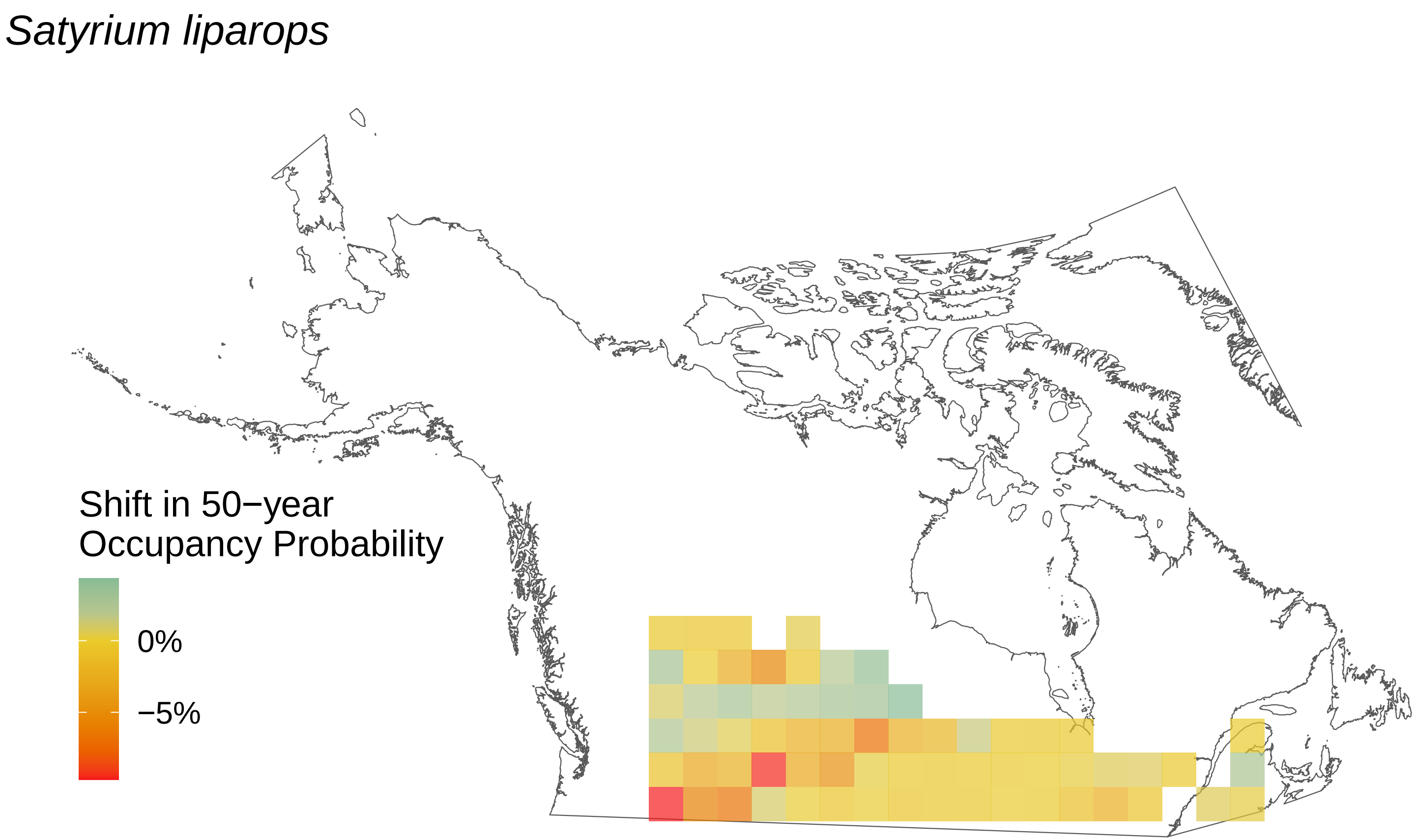

Satyrium liparops
Shift in 50−year Occupancy Probability
0%
−5%

*Pterourus rutulus*

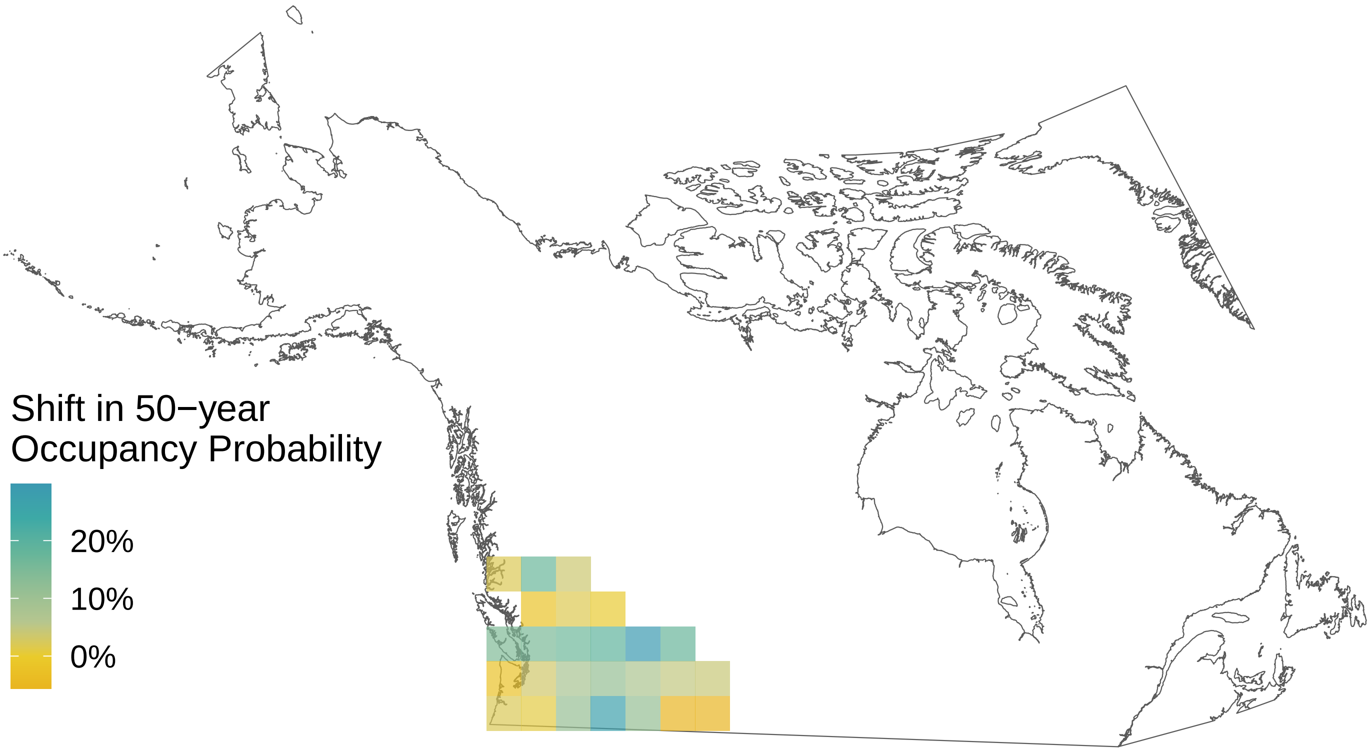

*Pterourus eurymedon*

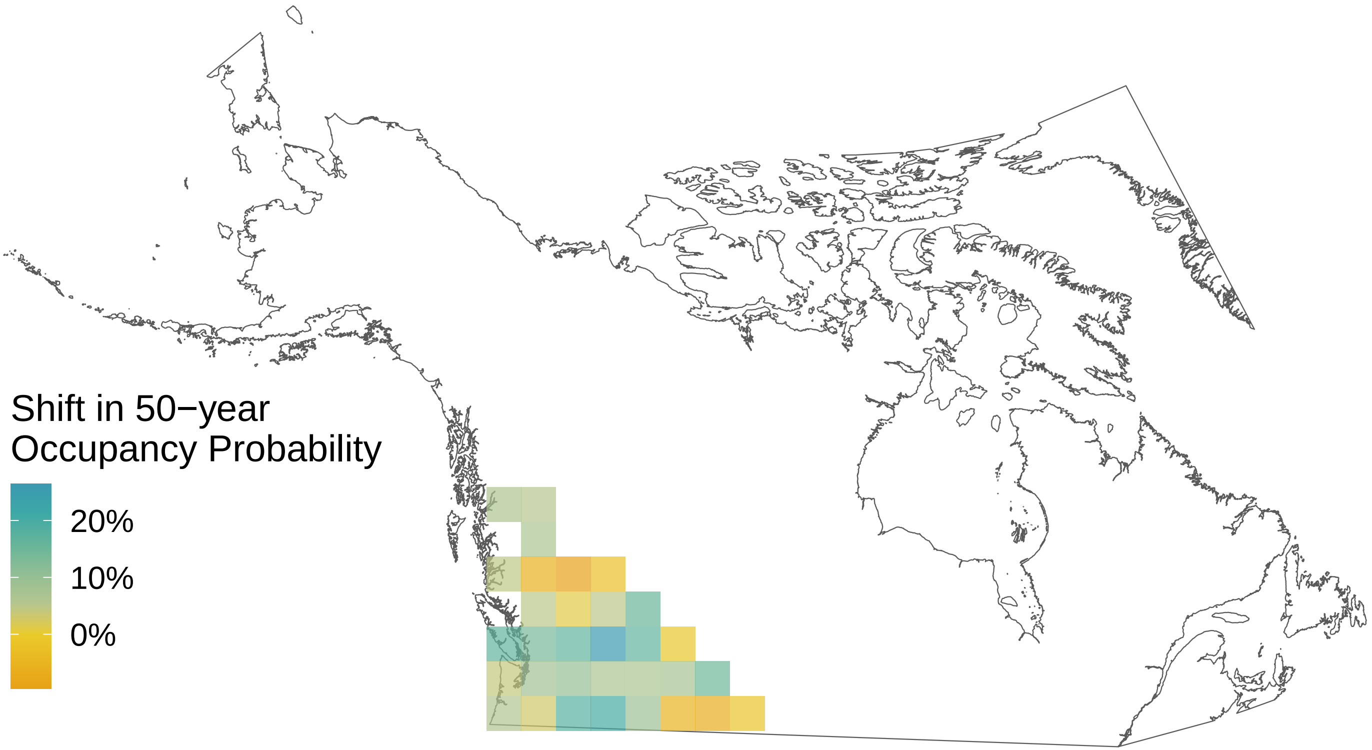

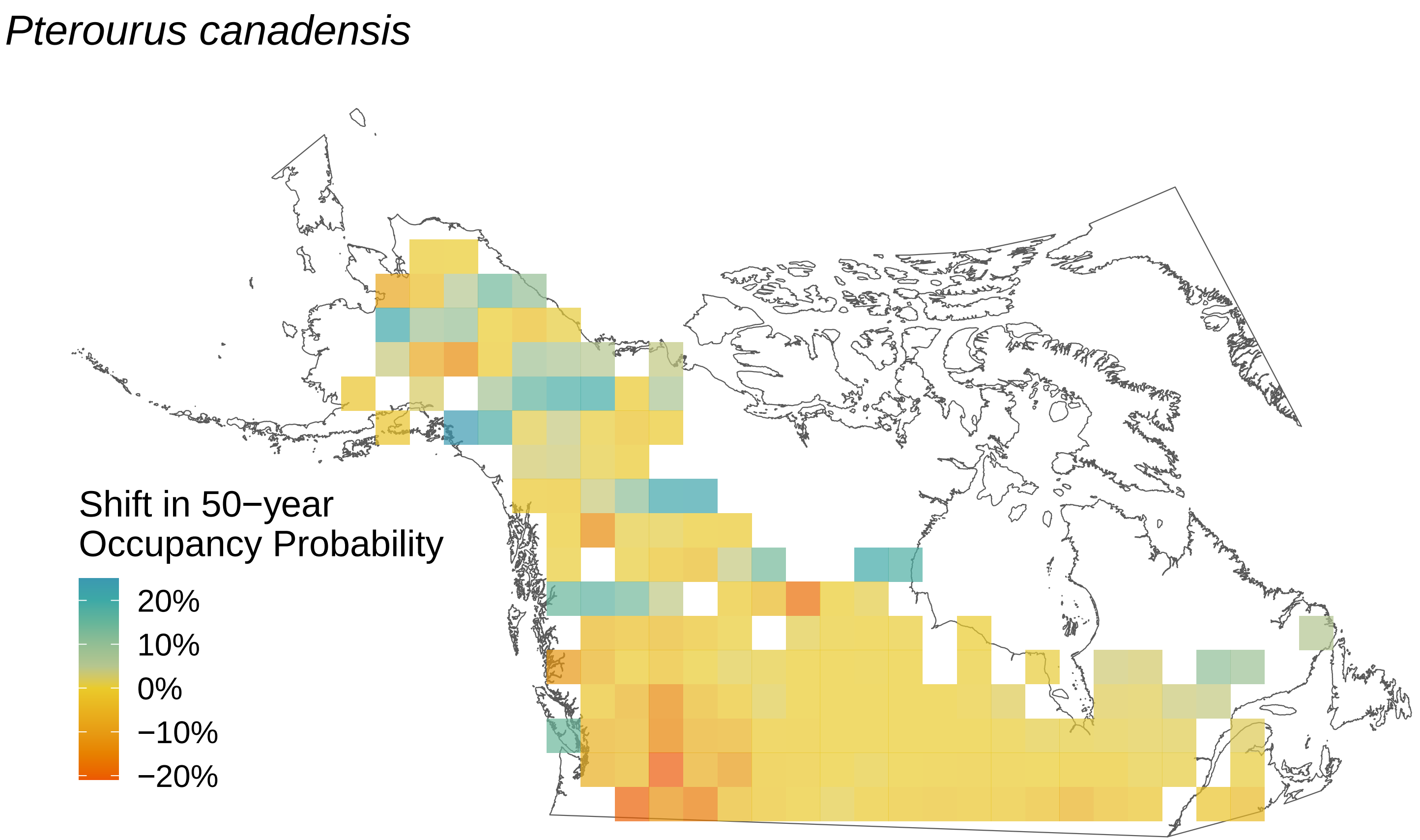
Pterourus canadensis
Shift in 50−year
Occupancy Probability
20%
10%
0%
−10%
−20%

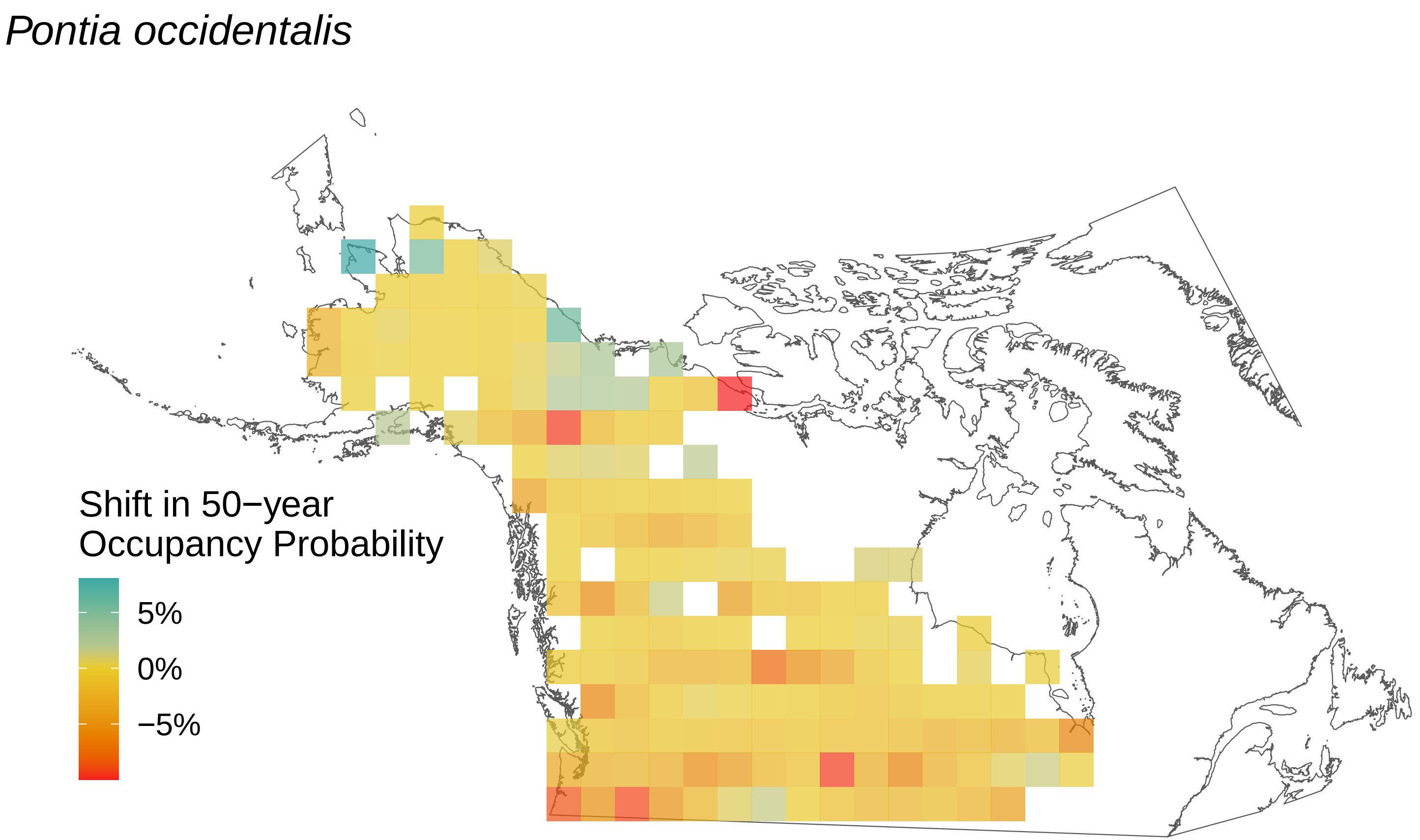

*Pontia occidentalis*
Shift in 50−year Occupancy Probability
5%
0%
−5%

*Polygonia satyrus*

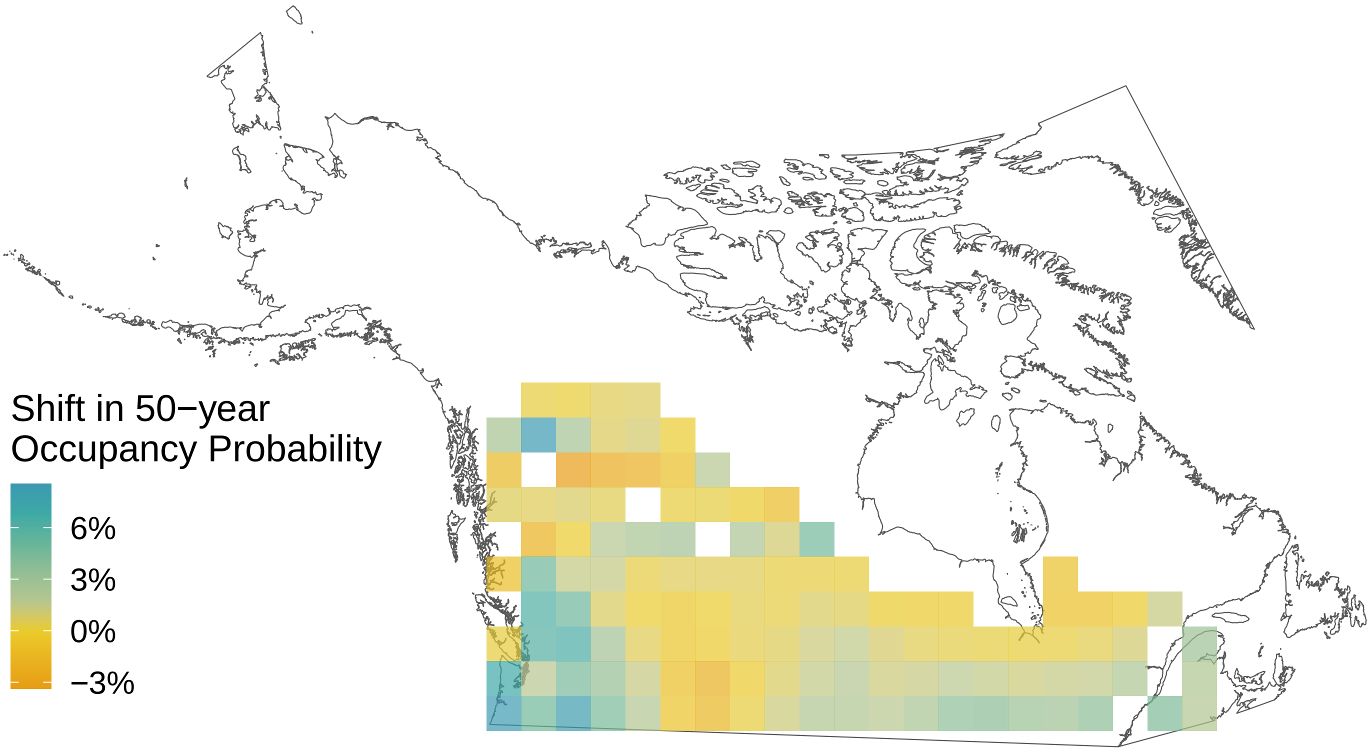

*Polygonia progne*

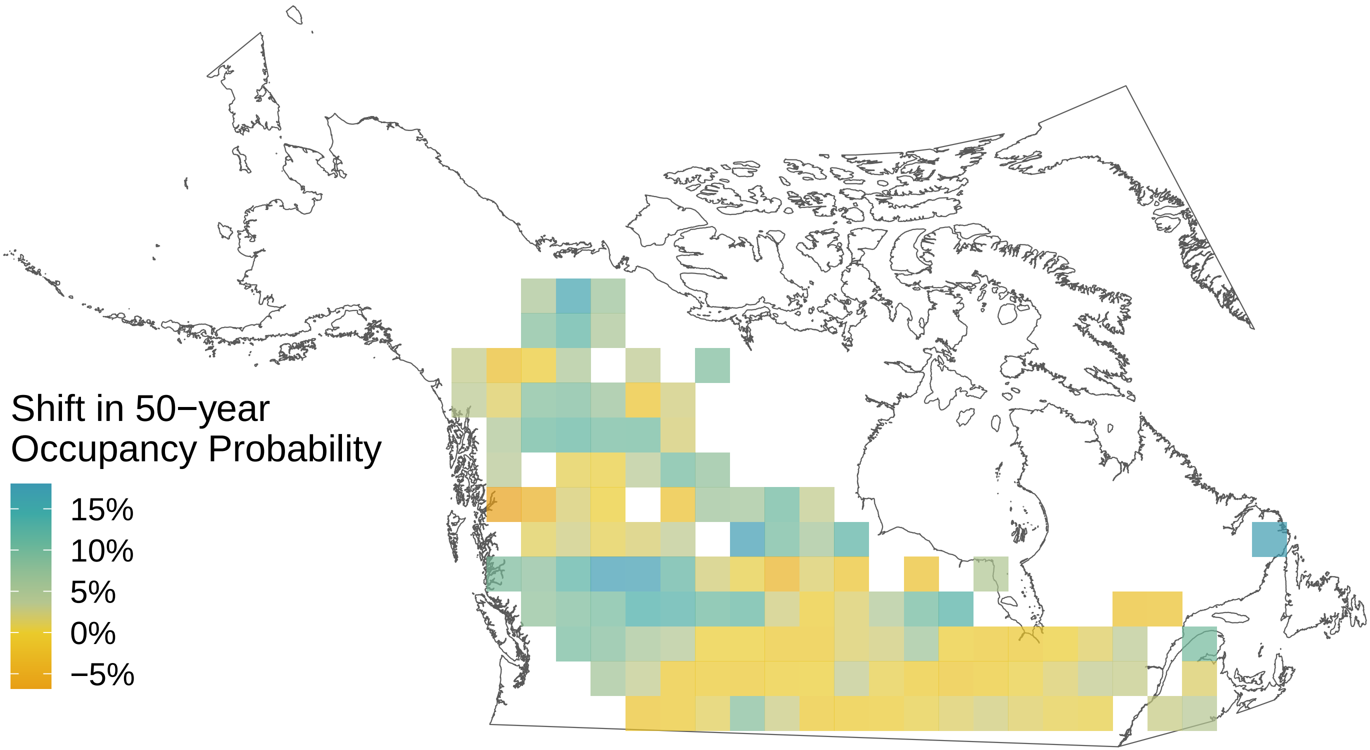

*Polygonia interrogationis*

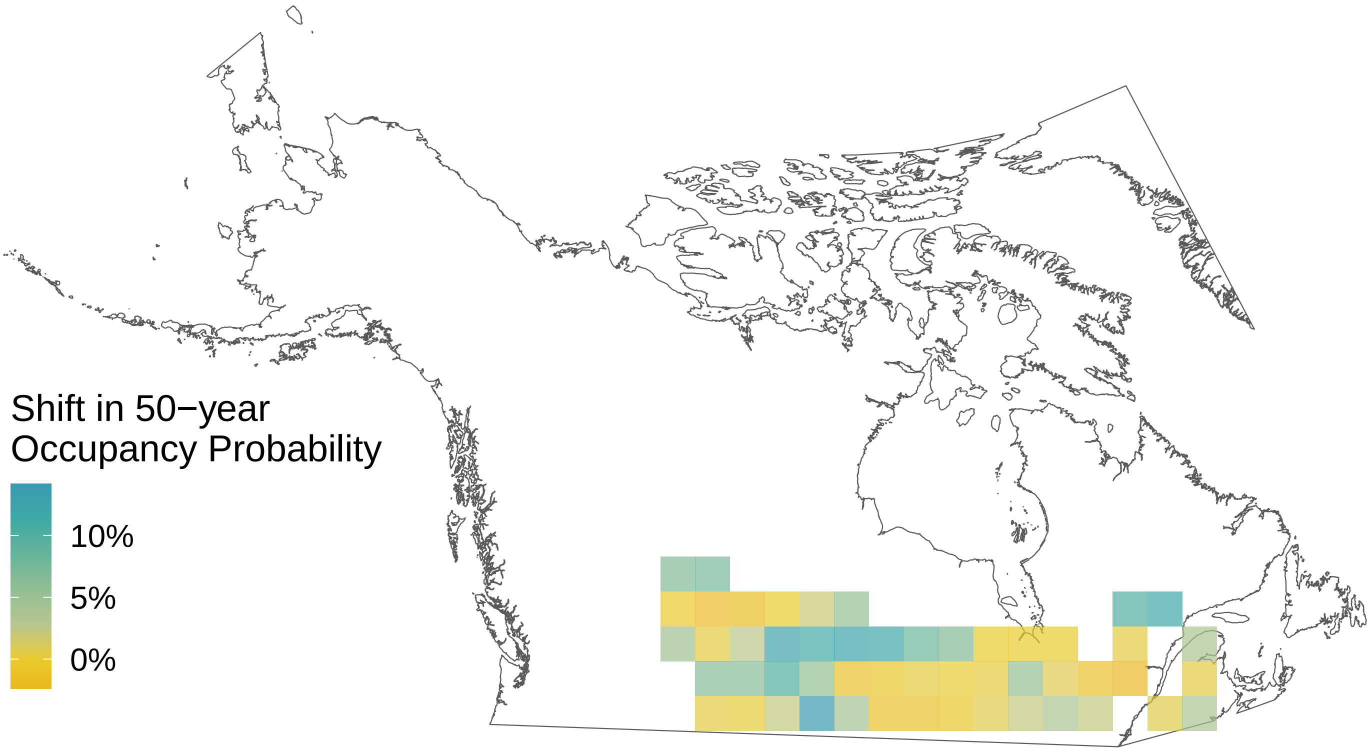

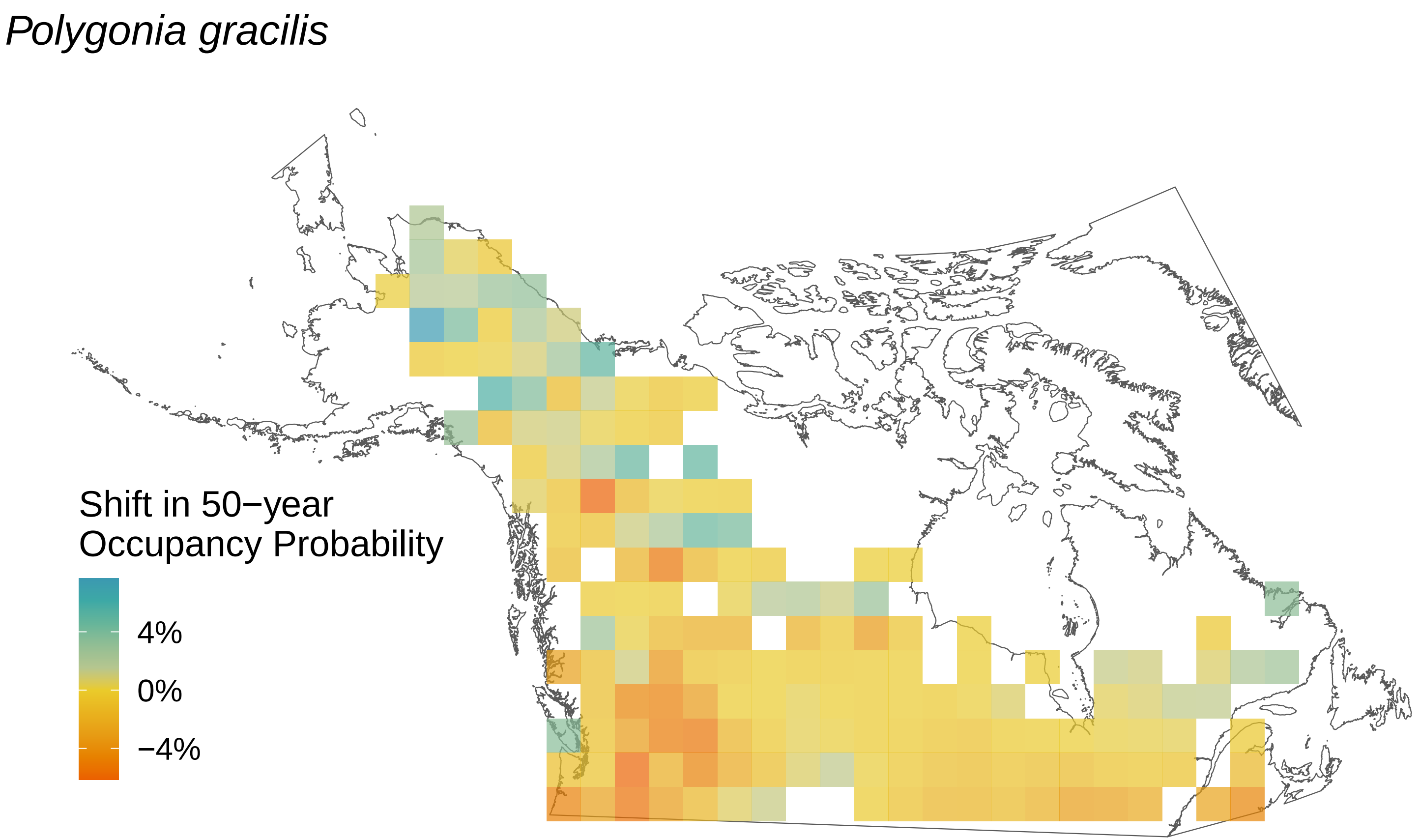
Polygonia gracilis
Shift in 50−year
Occupancy Probability
4%
0%
−4%

*Polygonia faunus*

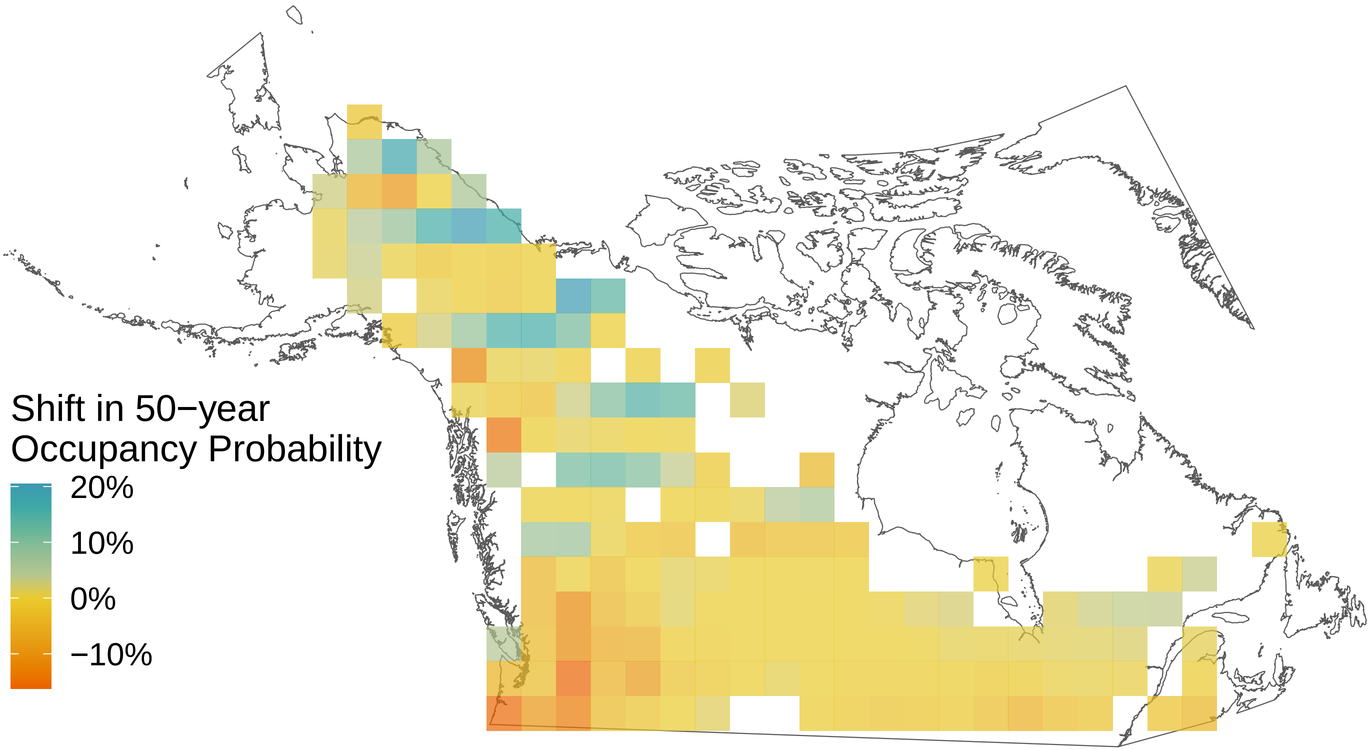

*Polites themistocles*

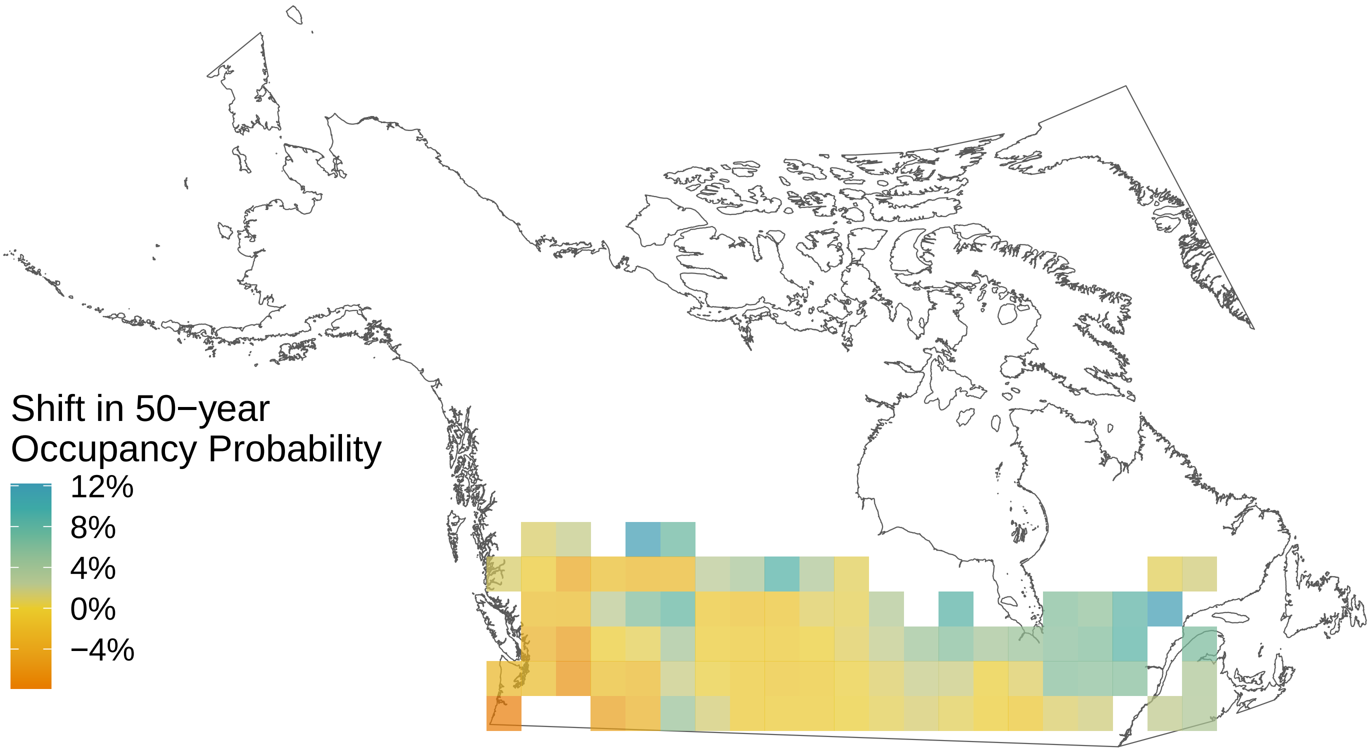

*Polites mystic*

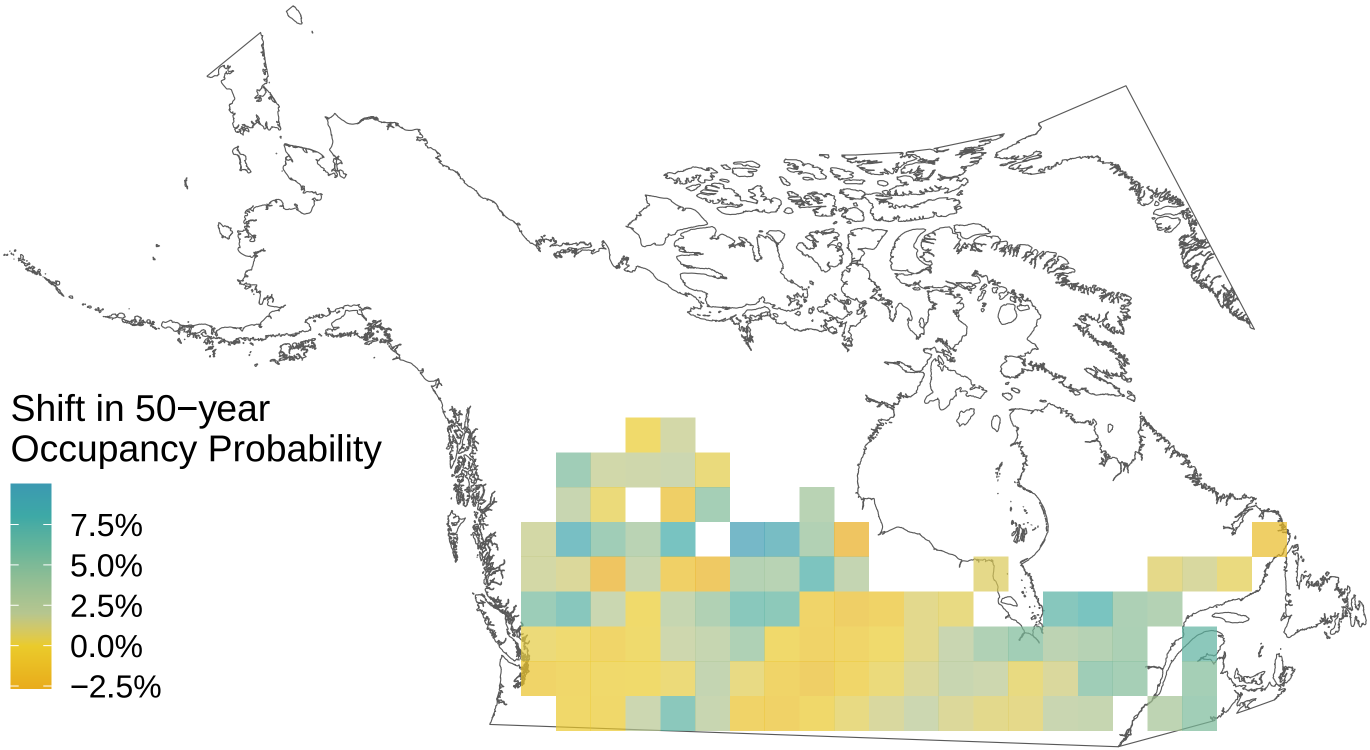

*Poanes hobomok*

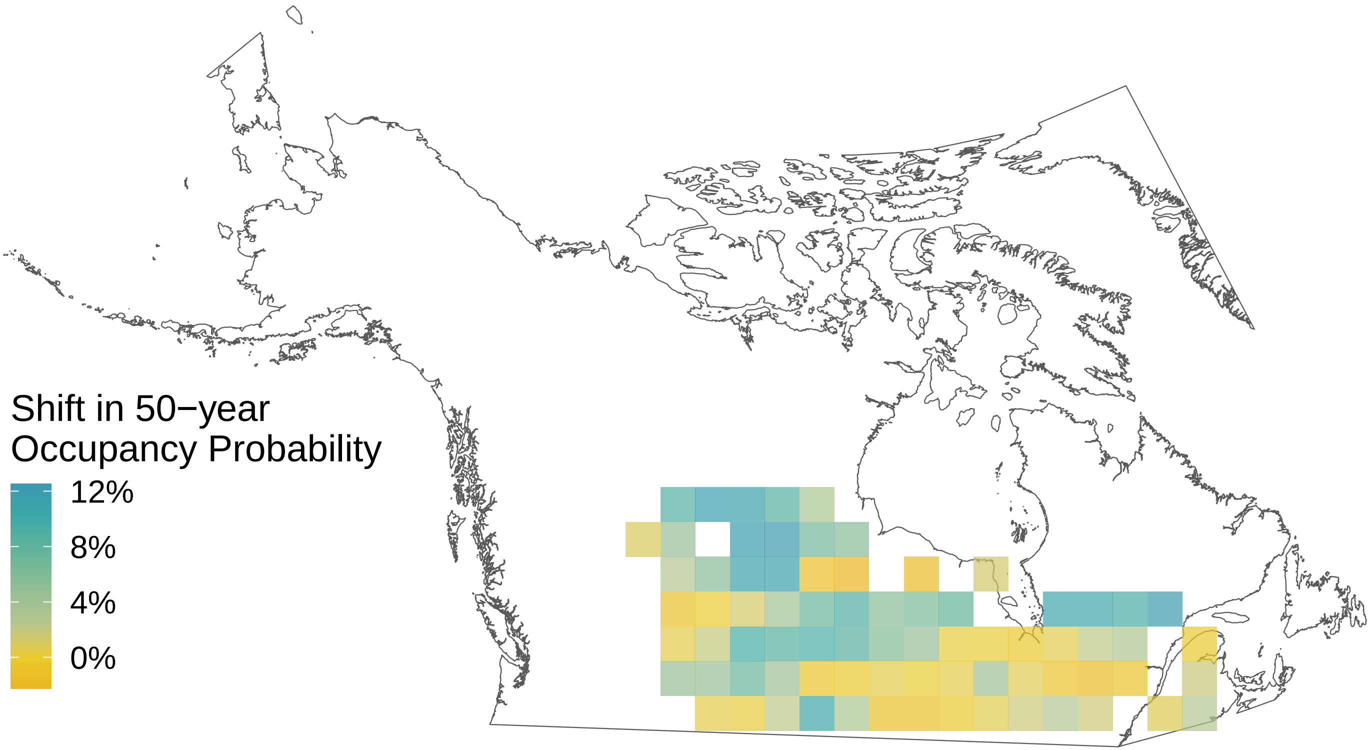

*Plebejus melissa*

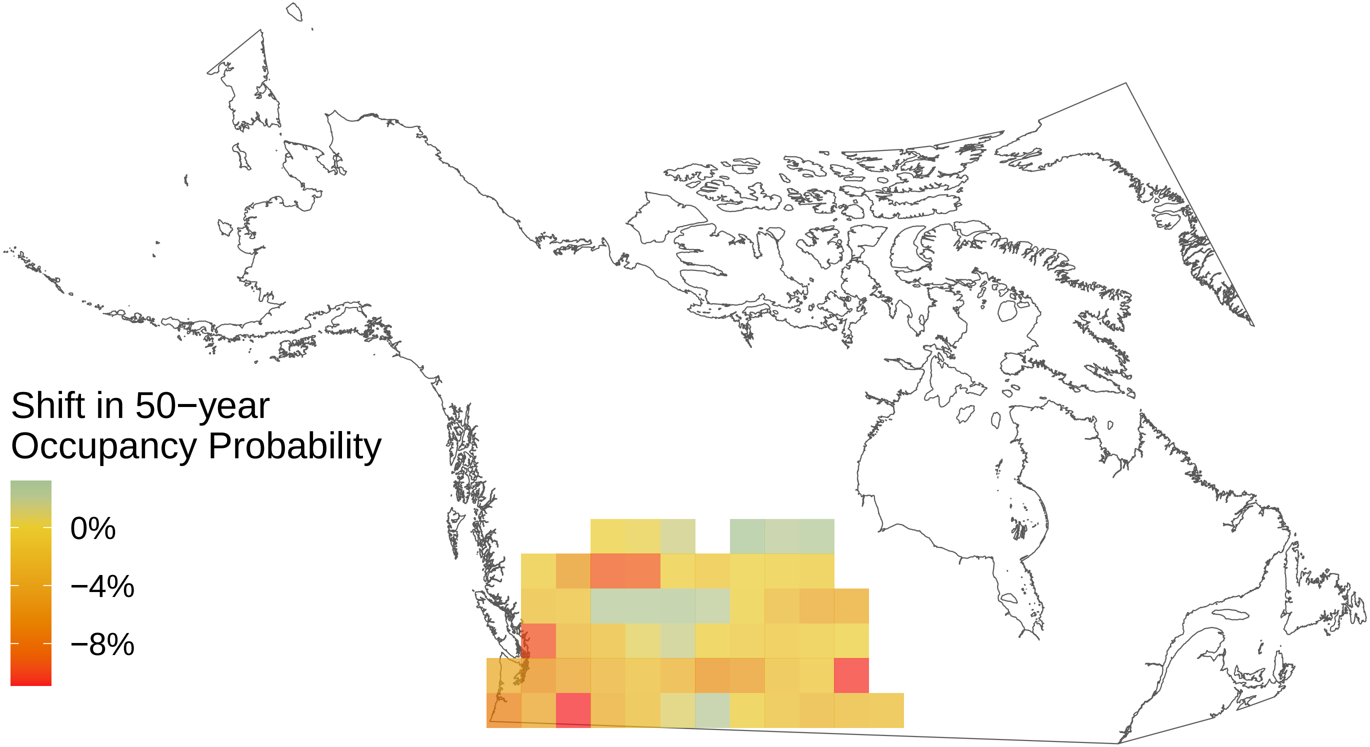

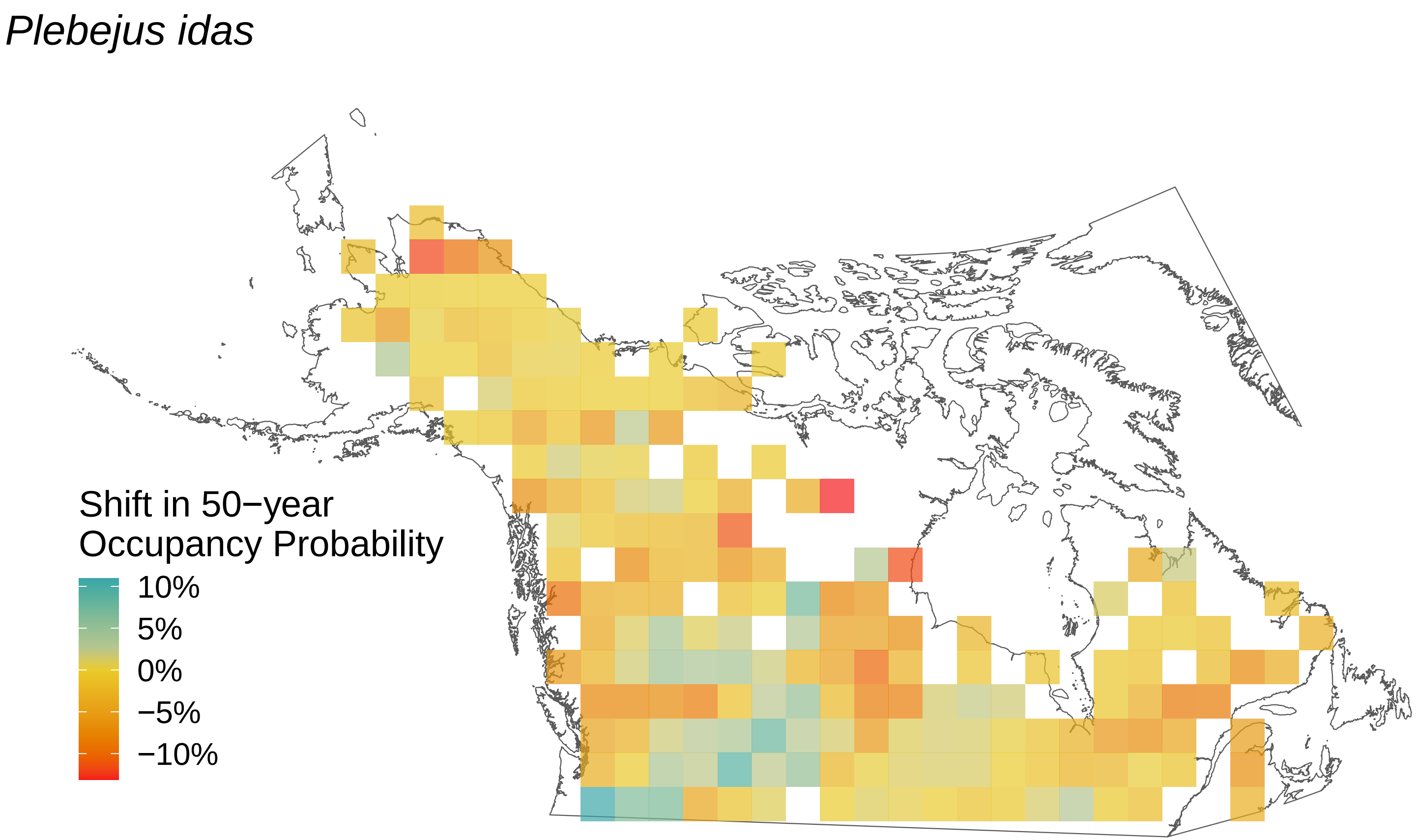

Plebejus idas
Shift in 50−year
Occupancy Probability
10%
5%
0%
−5%
−10%

*Pieris rapae*

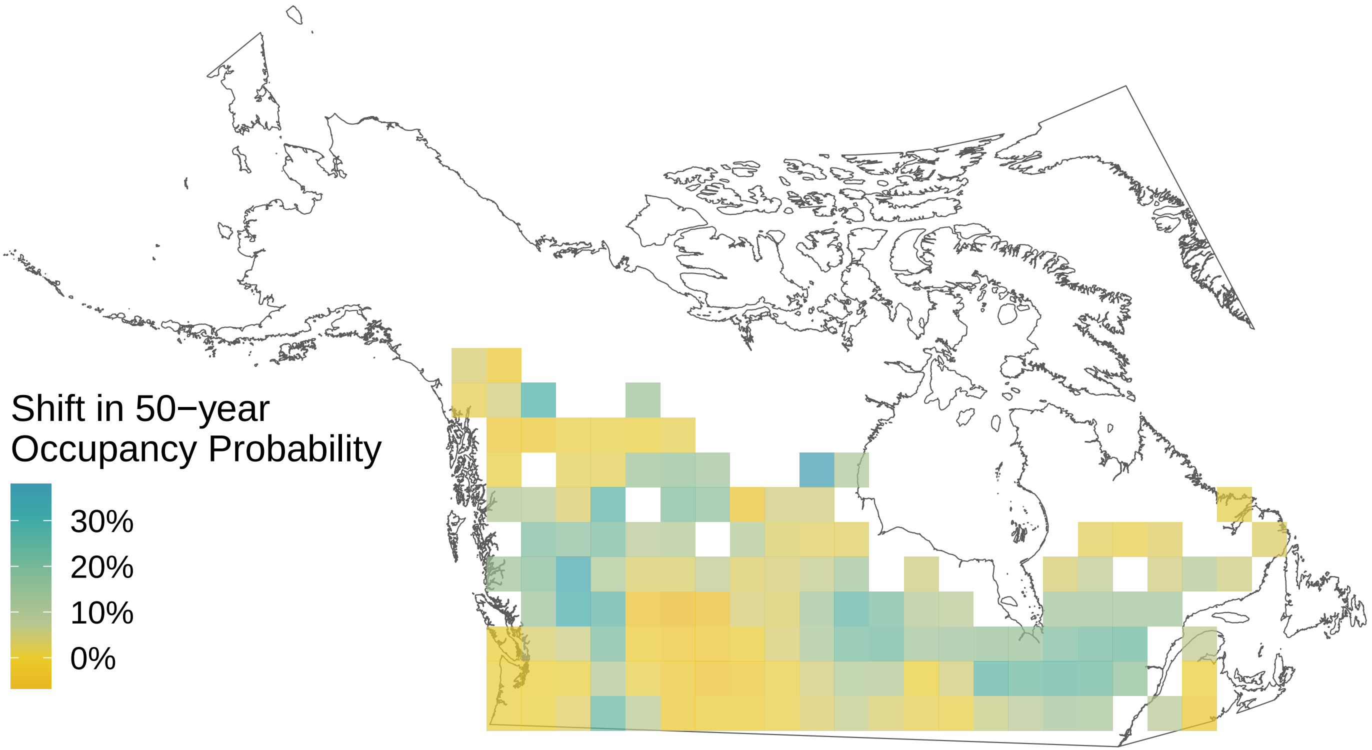

*Pieris oleracea*

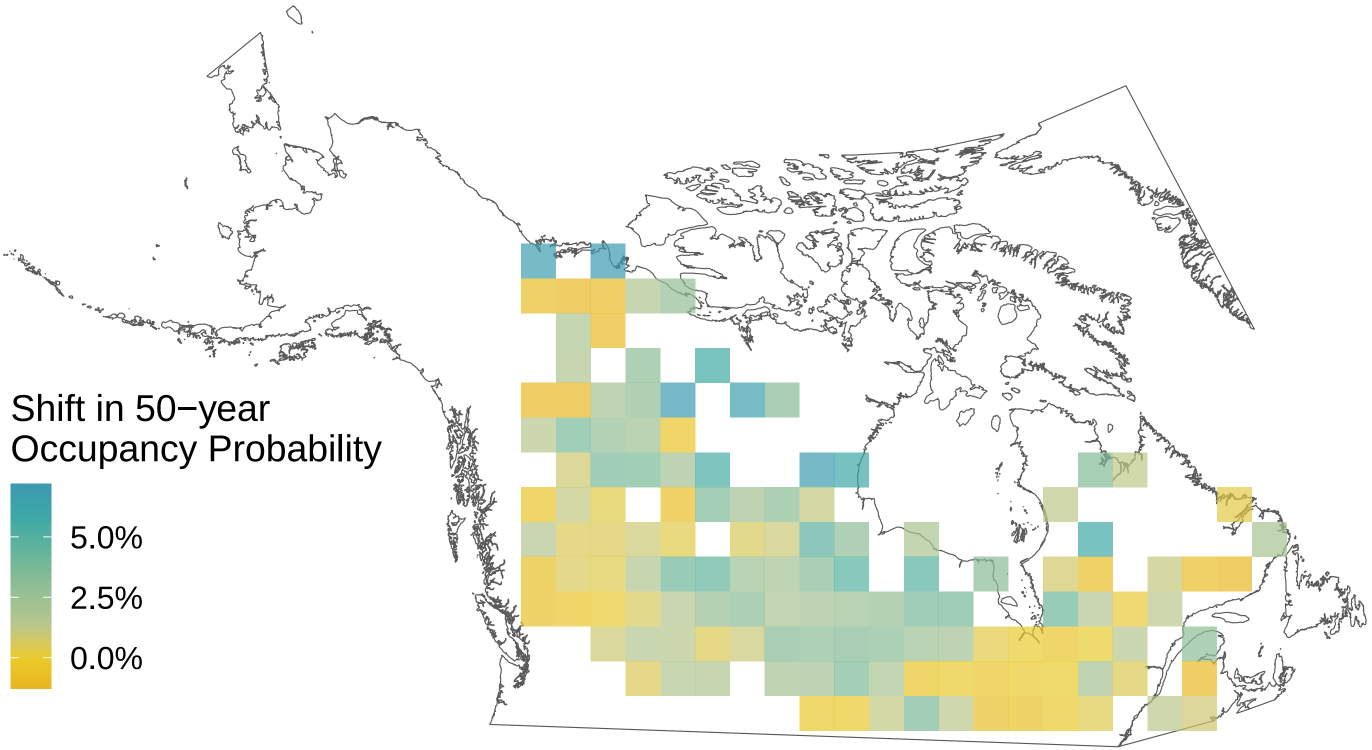

*Phyciodes tharos*

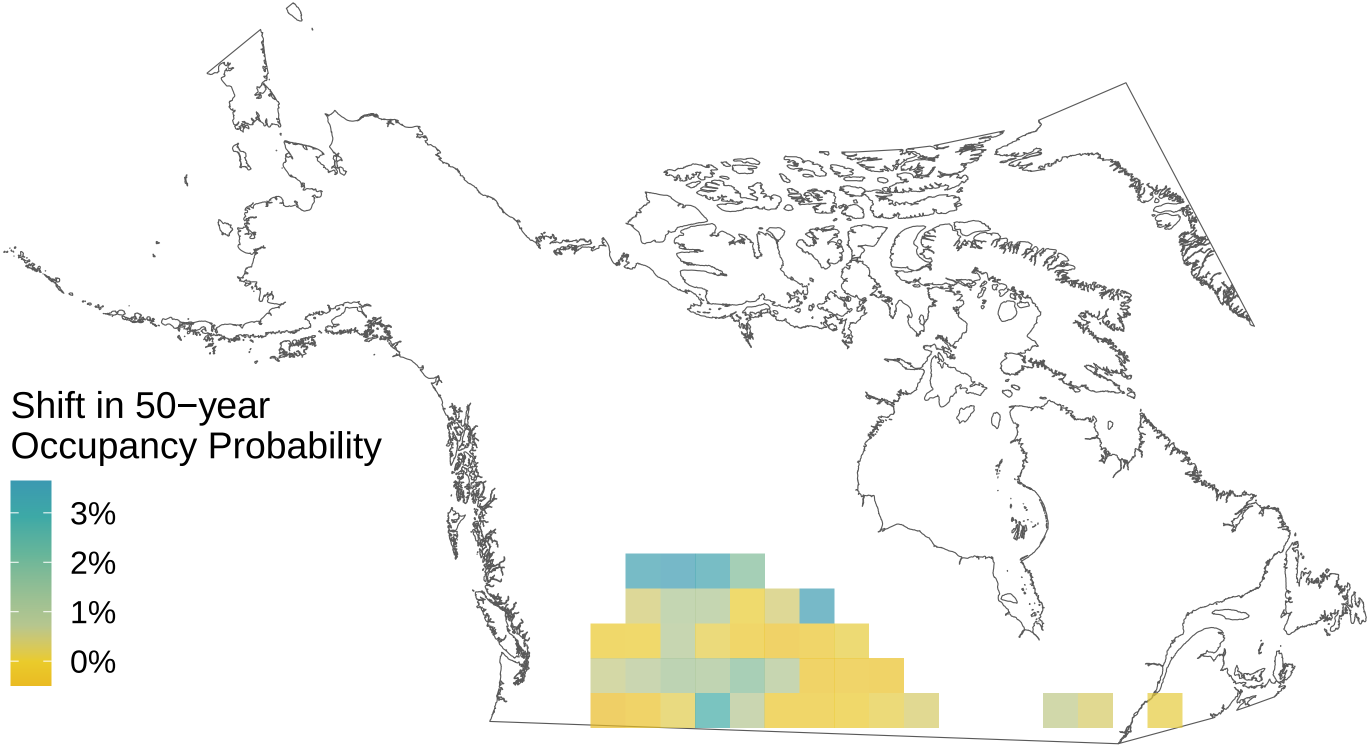

*Parnassius clodius*

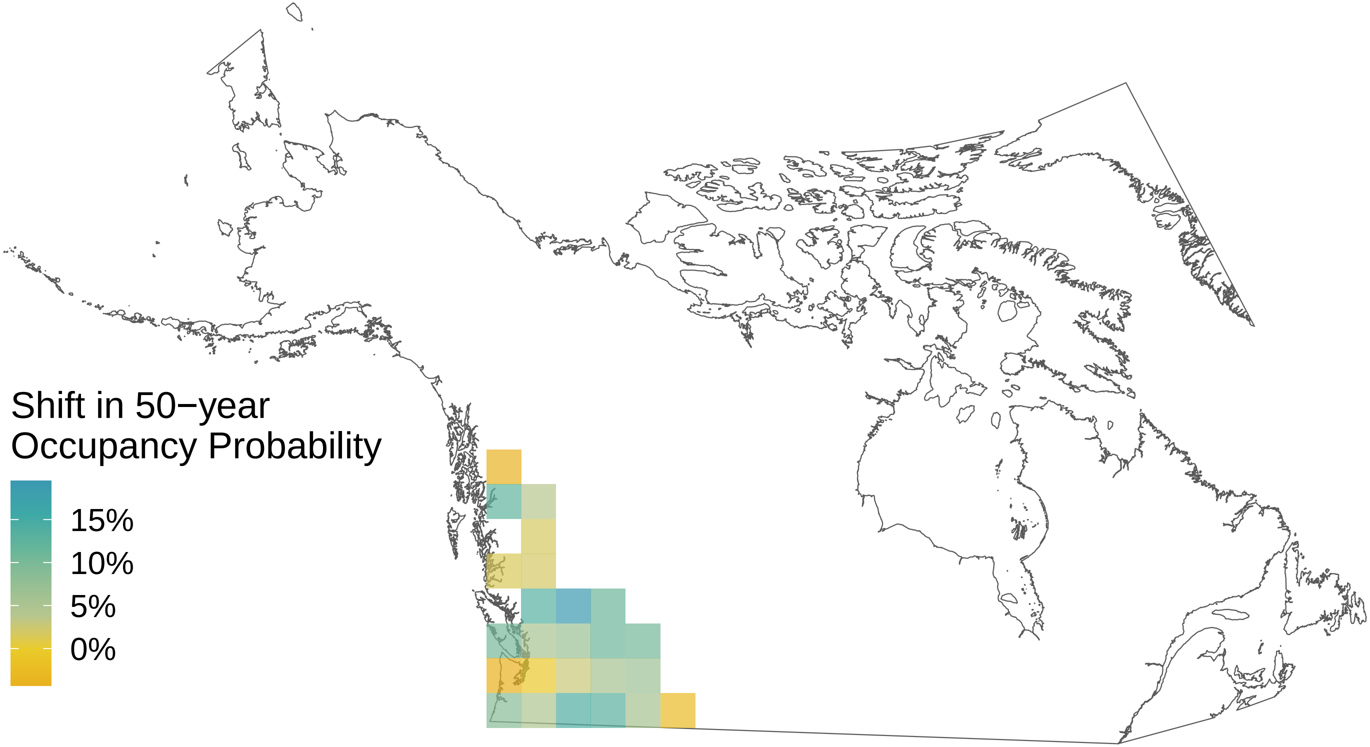

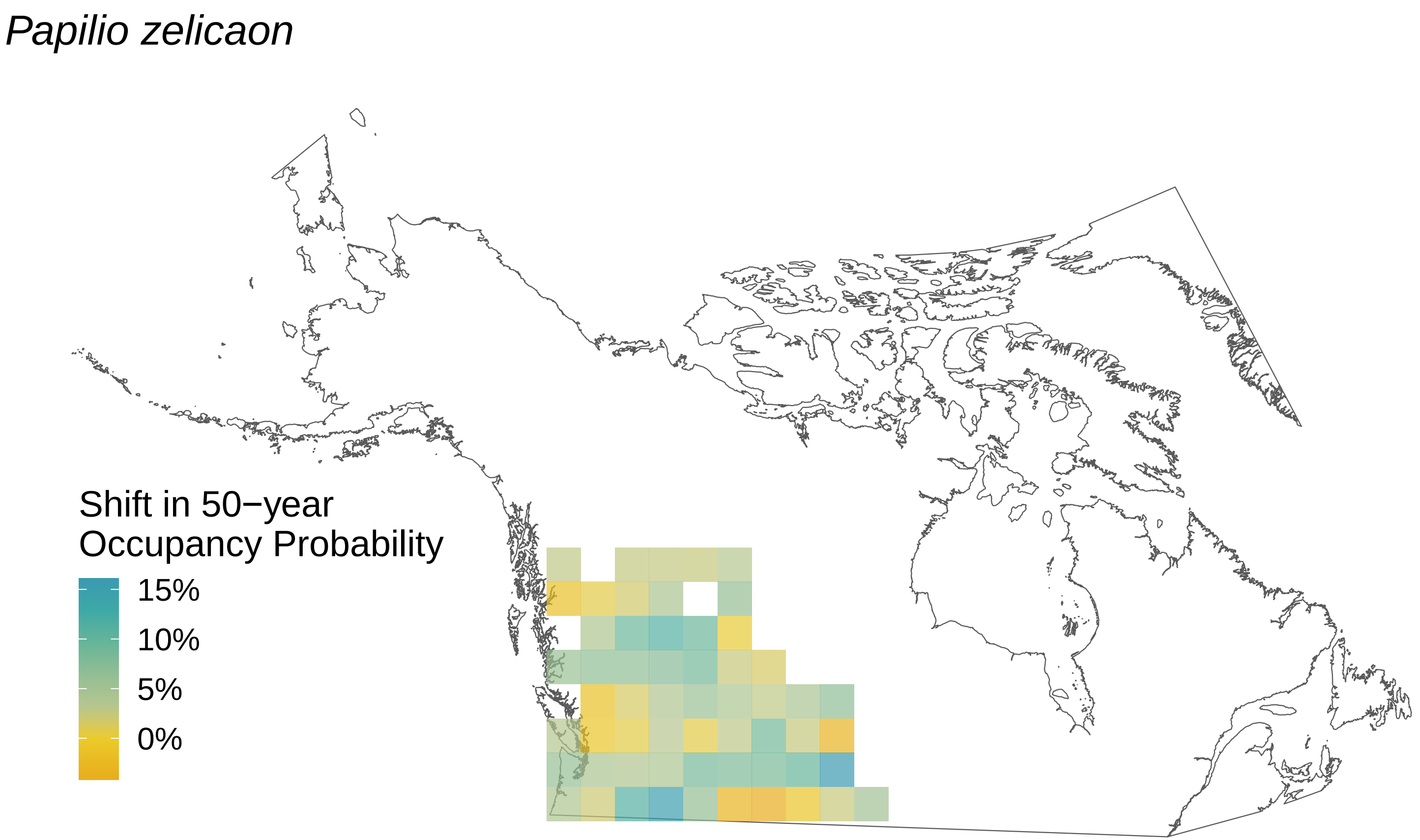
Papilio zelicaon
Shift in 50−year Occupancy Probability
15%
10%
5%
0%

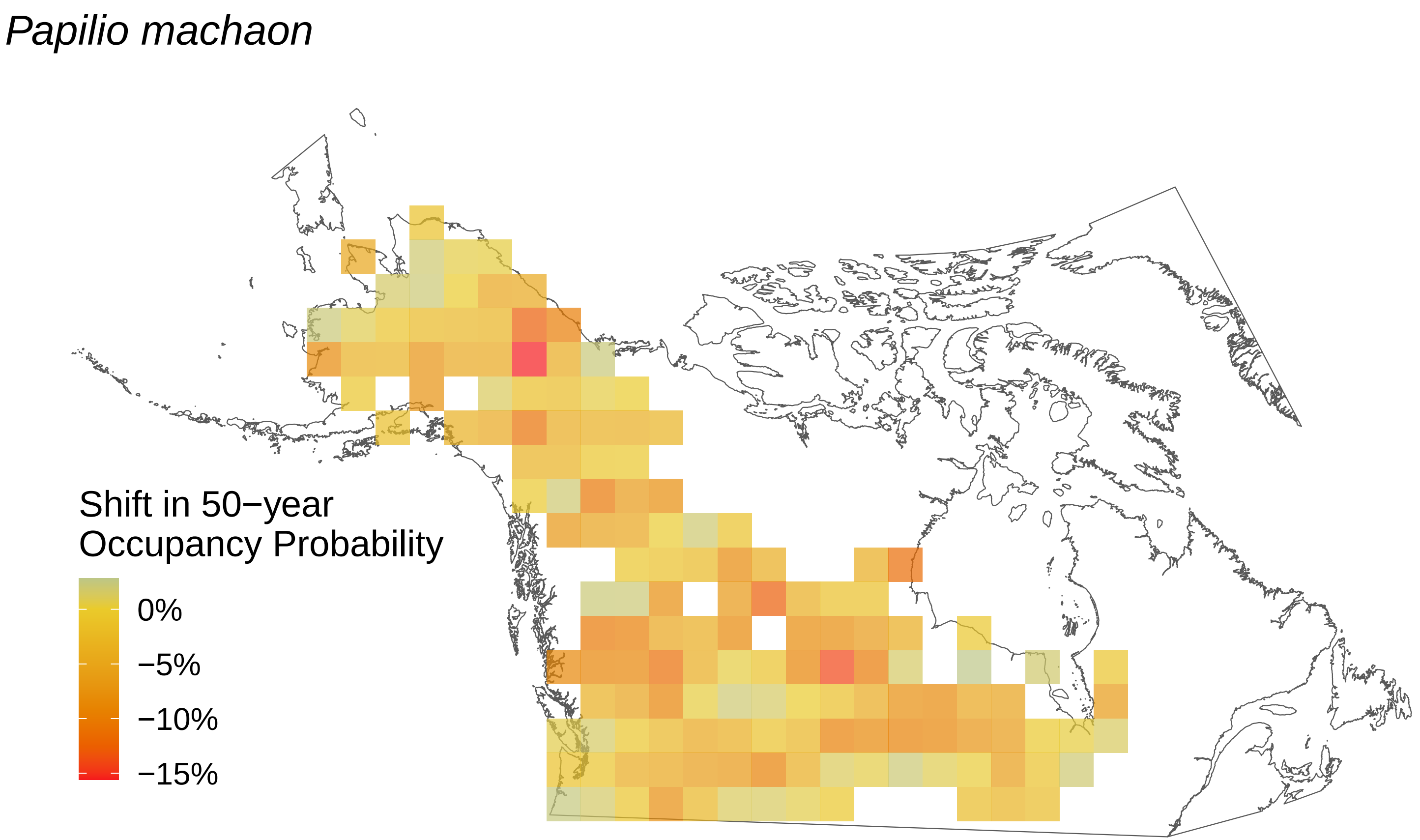
Papilio machaon
Shift in 50−year
Occupancy Probability
0%
−5%
−10%
−15%

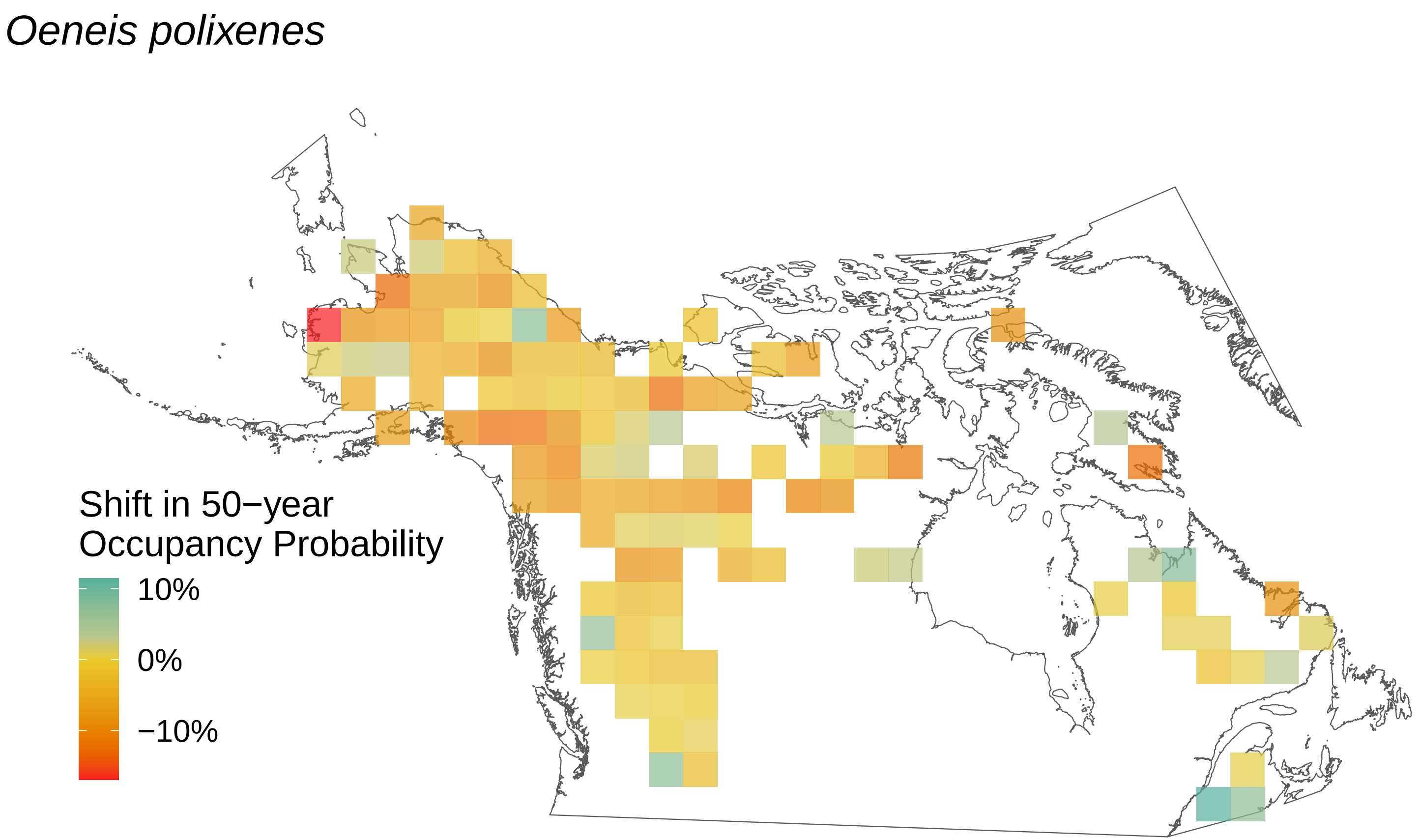

Oeneis polixenes
Shift in 50−year
Occupancy Probability
10%
0%
−10%

*Oeneis melissa*

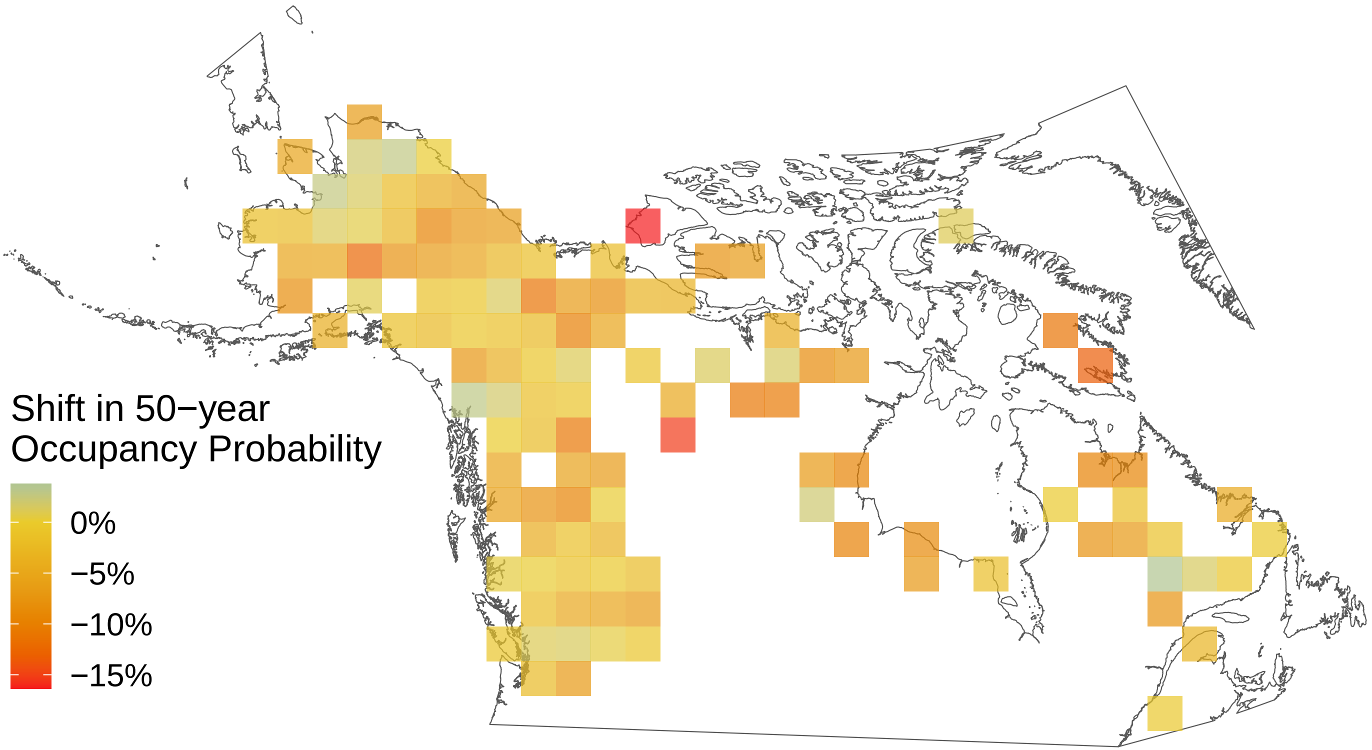

*Oeneis jutta*

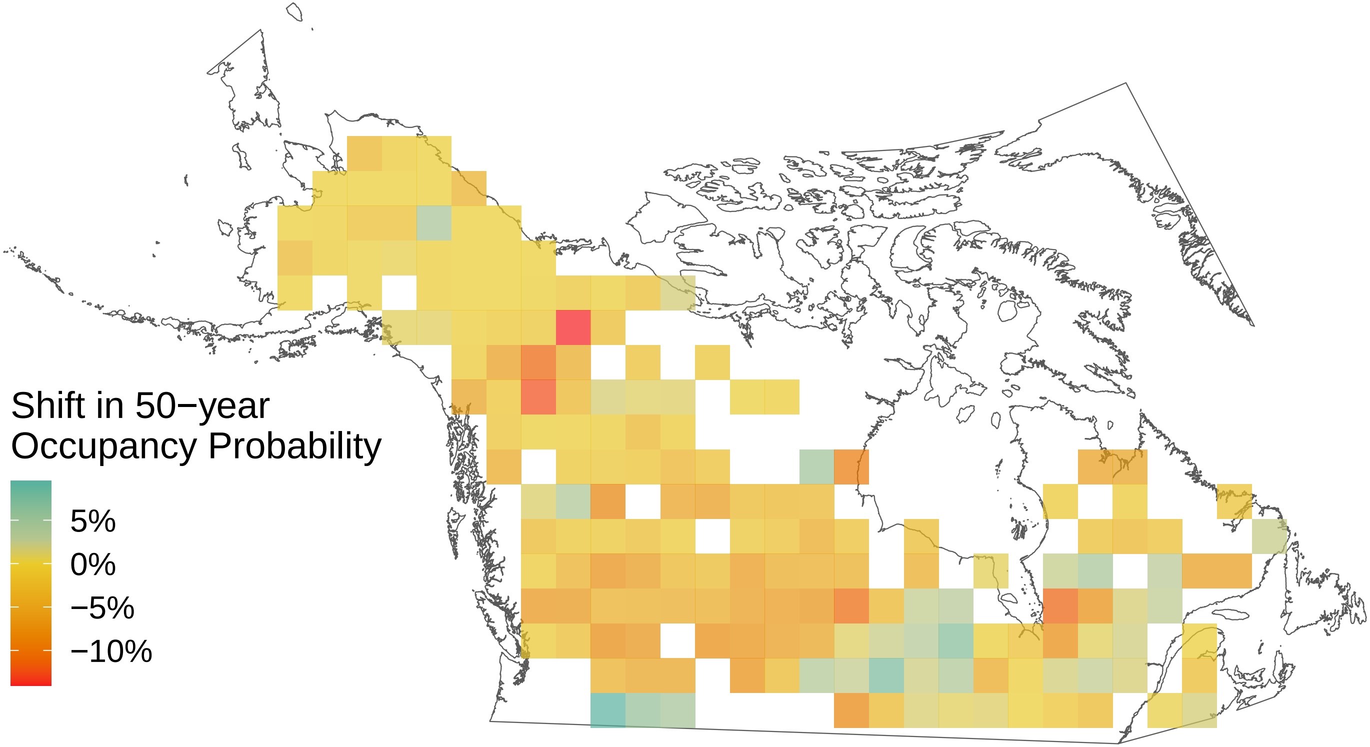

*Oeneis chryxus*

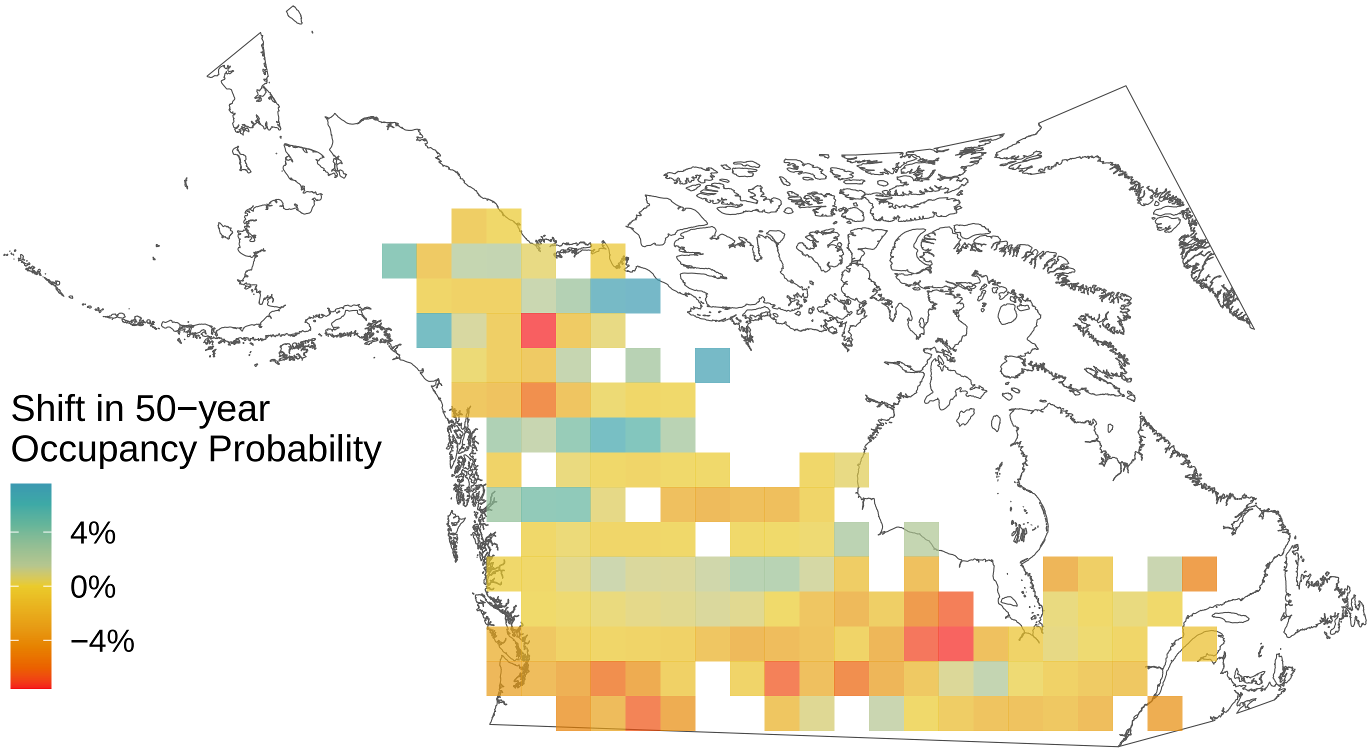

*Oeneis bore*

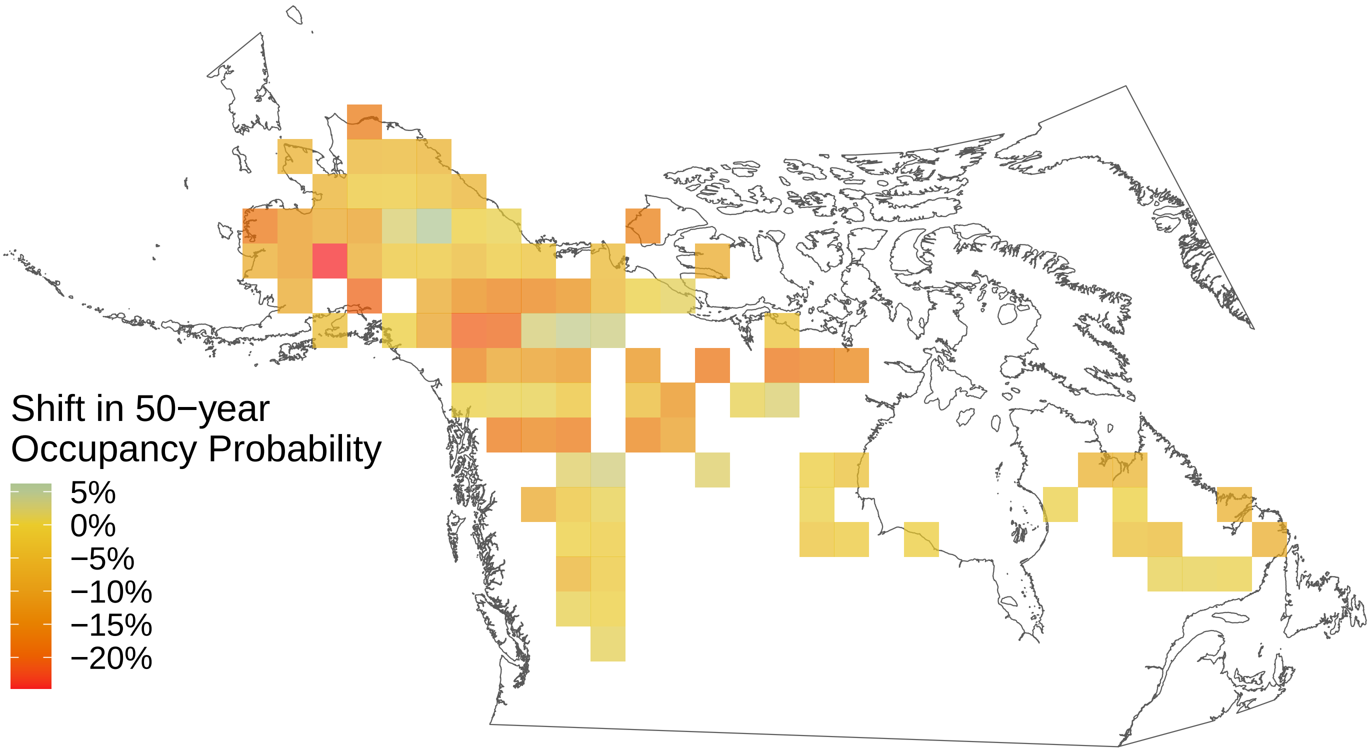

*Ochlodes sylvanoides*

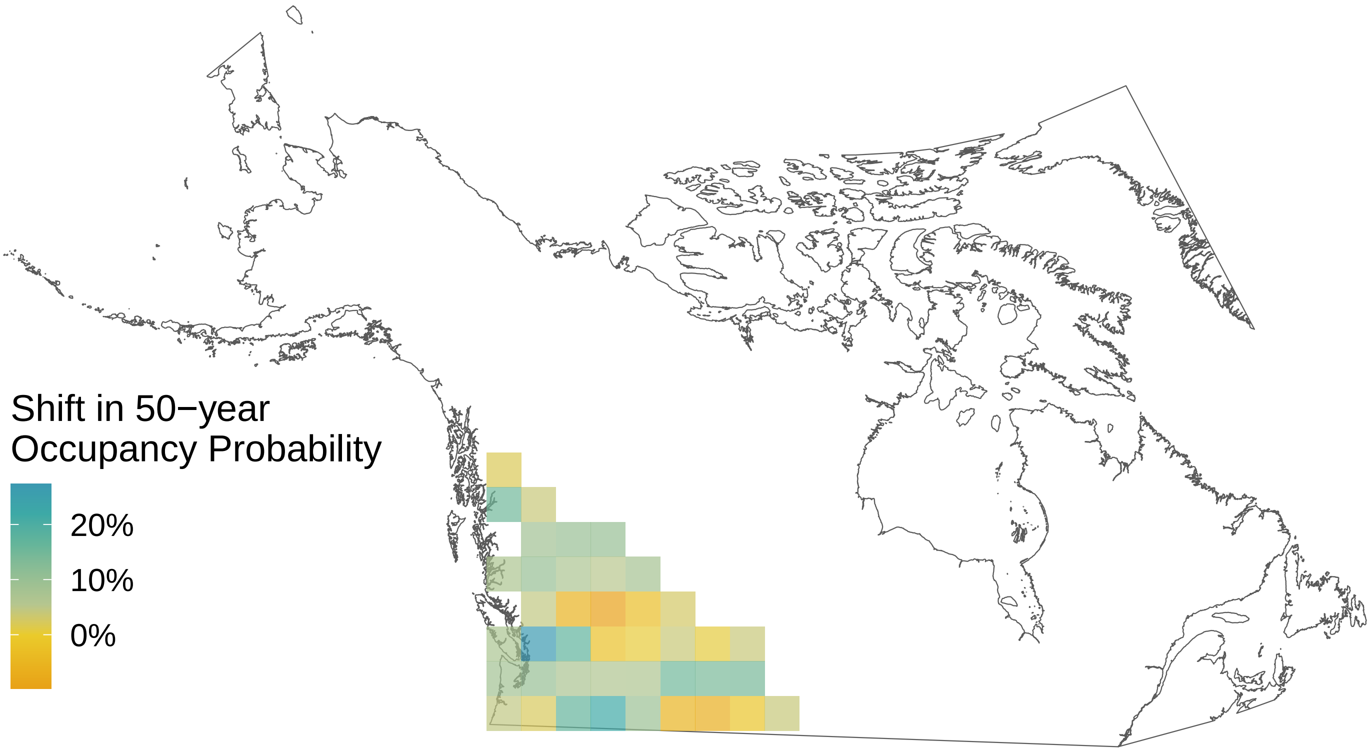

*Nymphalis l−album*

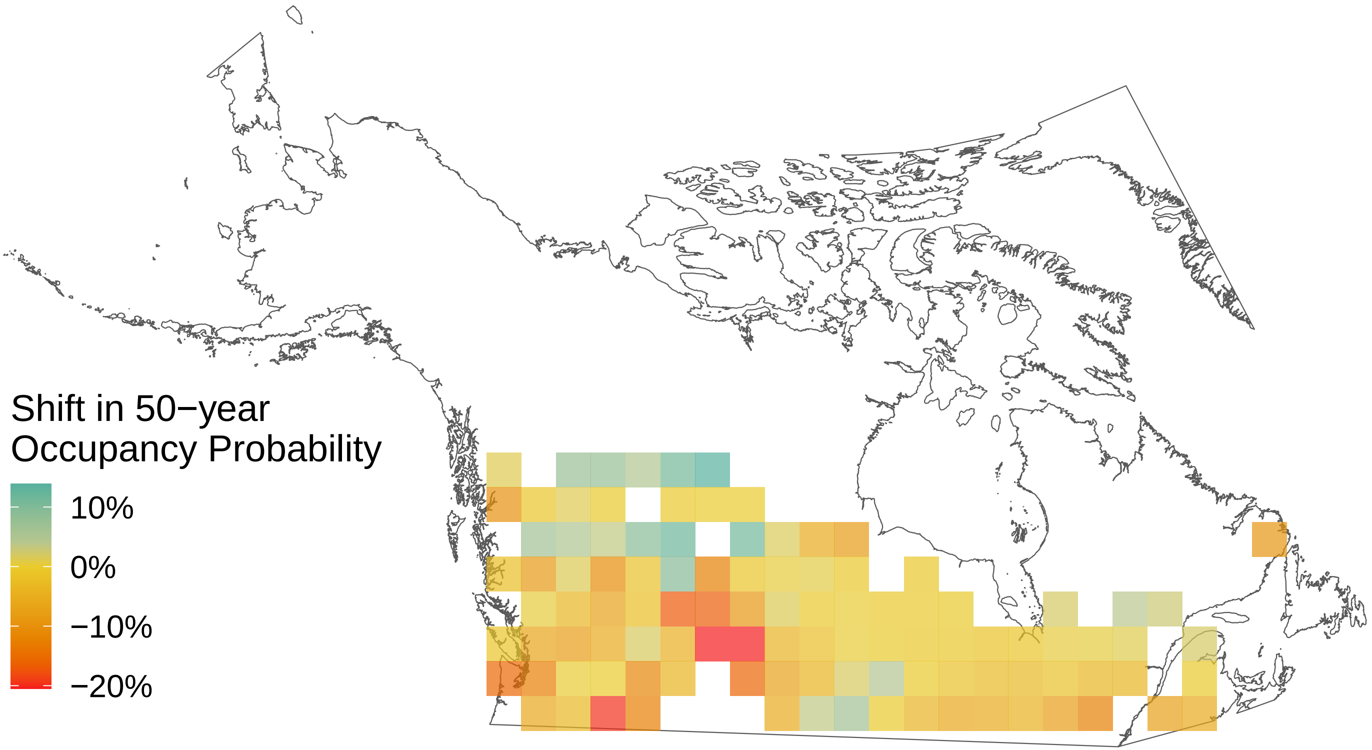

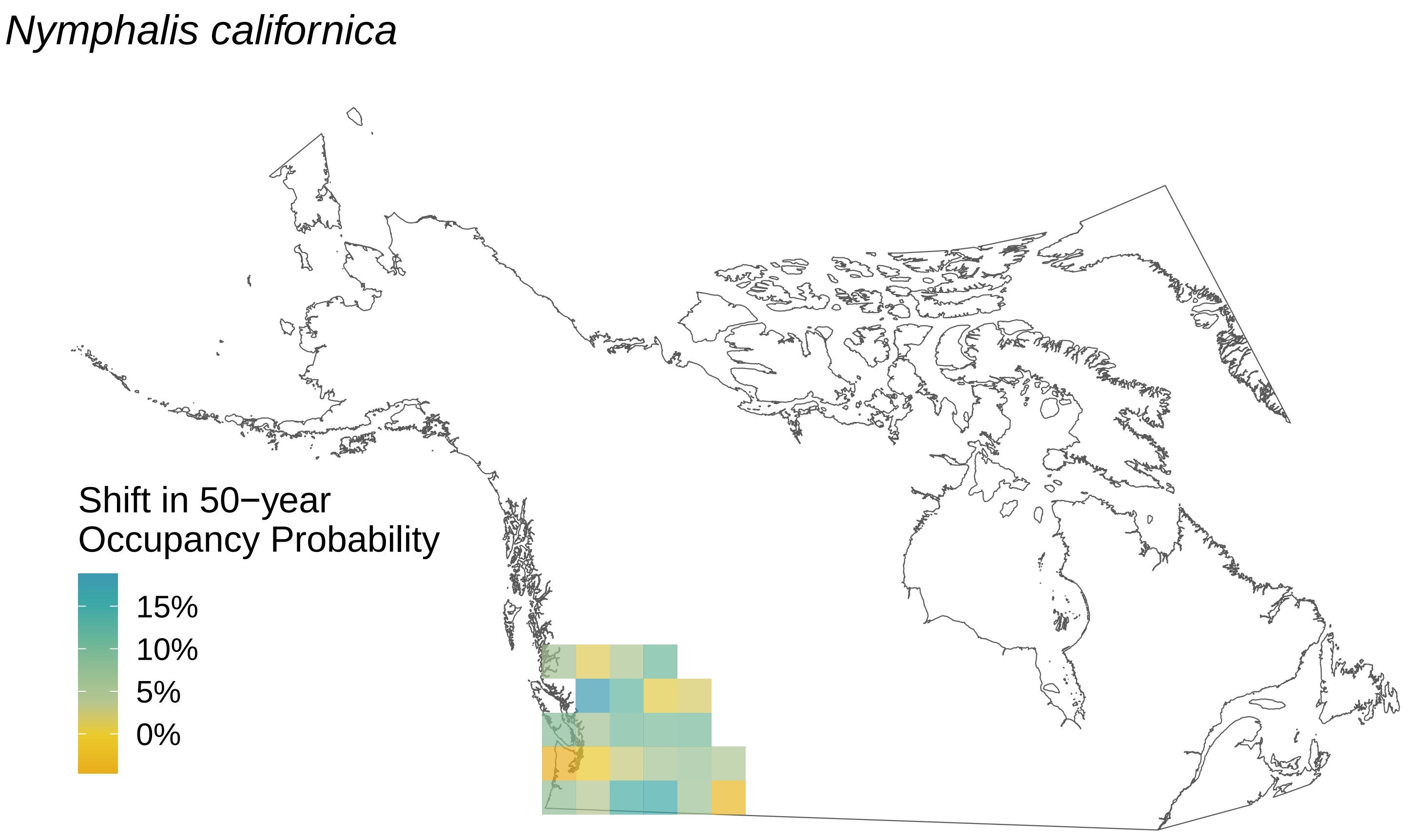
Nymphalis californica
Shift in 50−year
Occupancy Probability
15%
10%
5%
0%

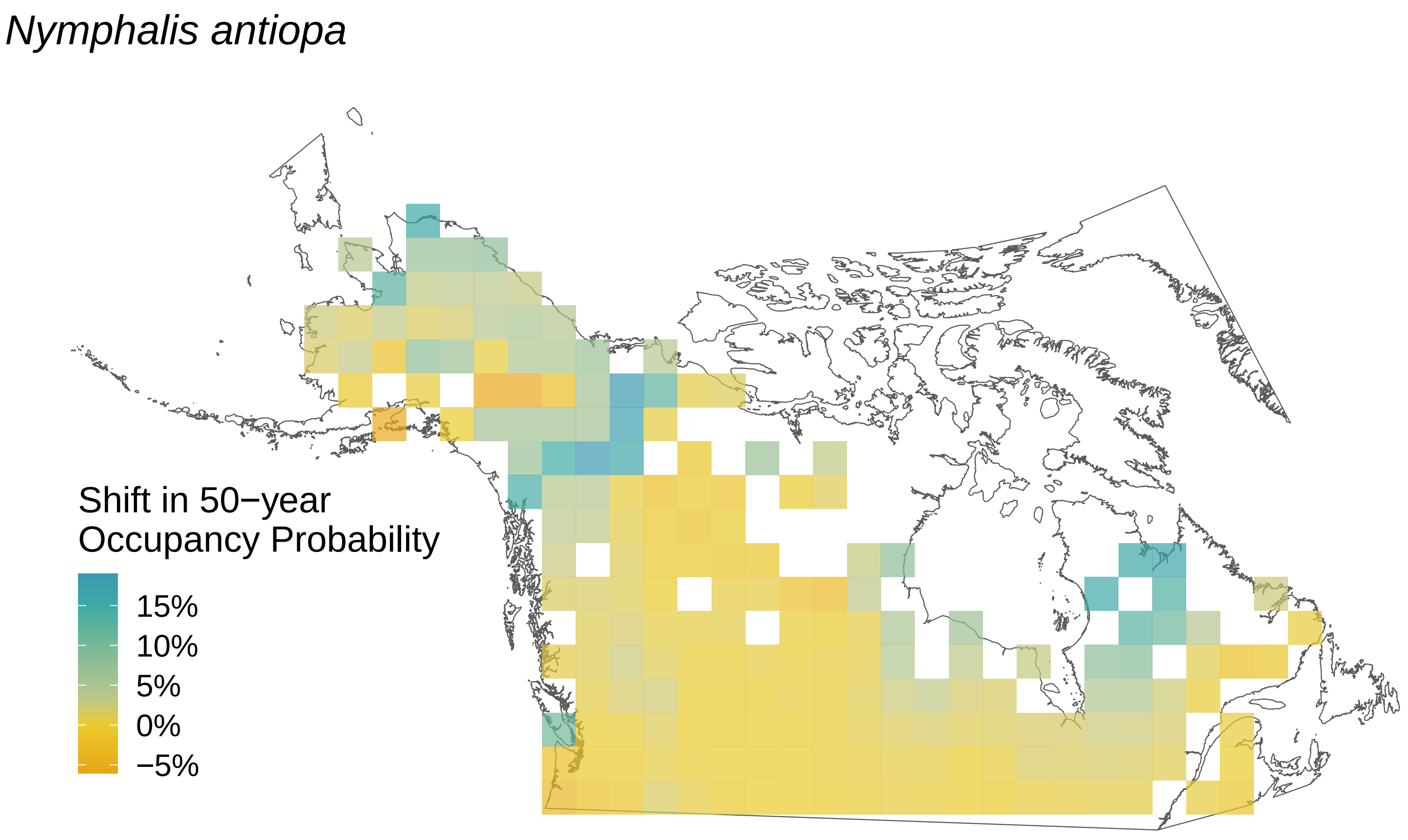

Nymphalis antiopa
Shift in 50−year
Occupancy Probability
15%
10%
5%
0%
−5%

*Neophasia menapia*

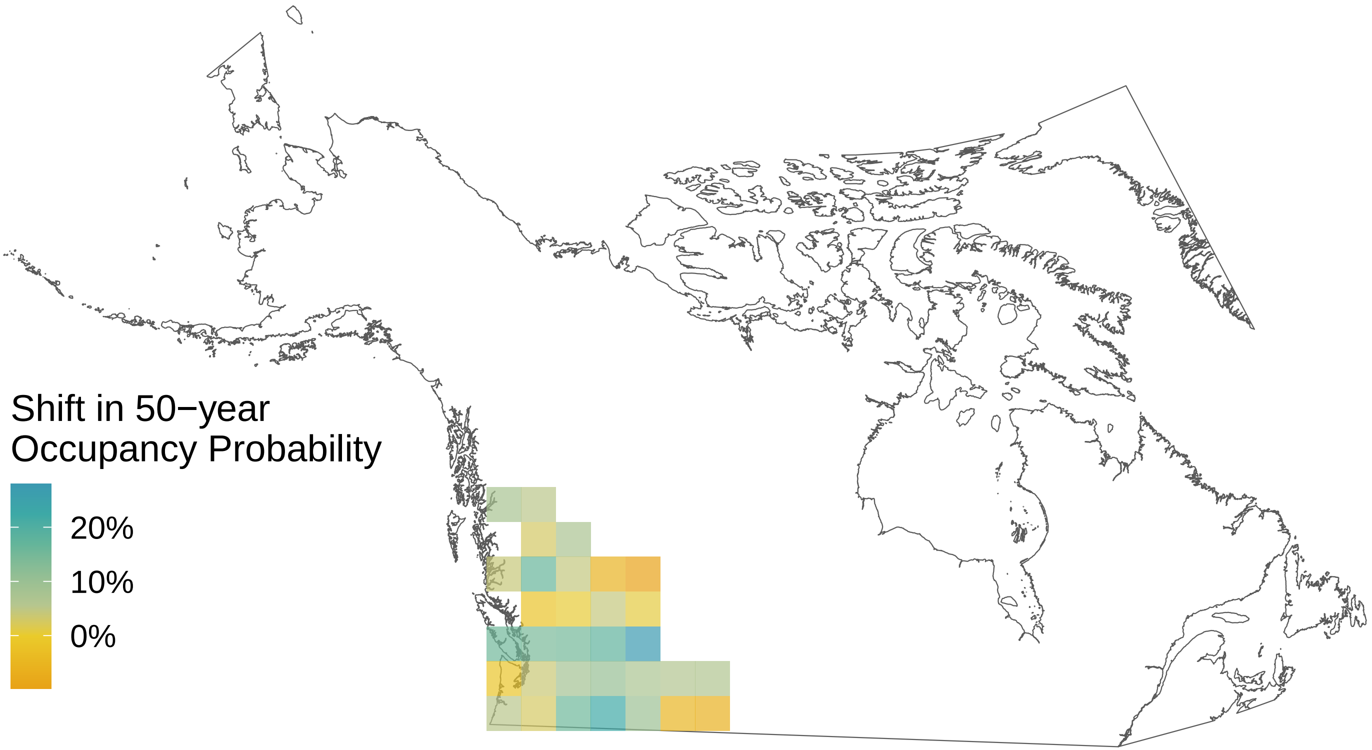

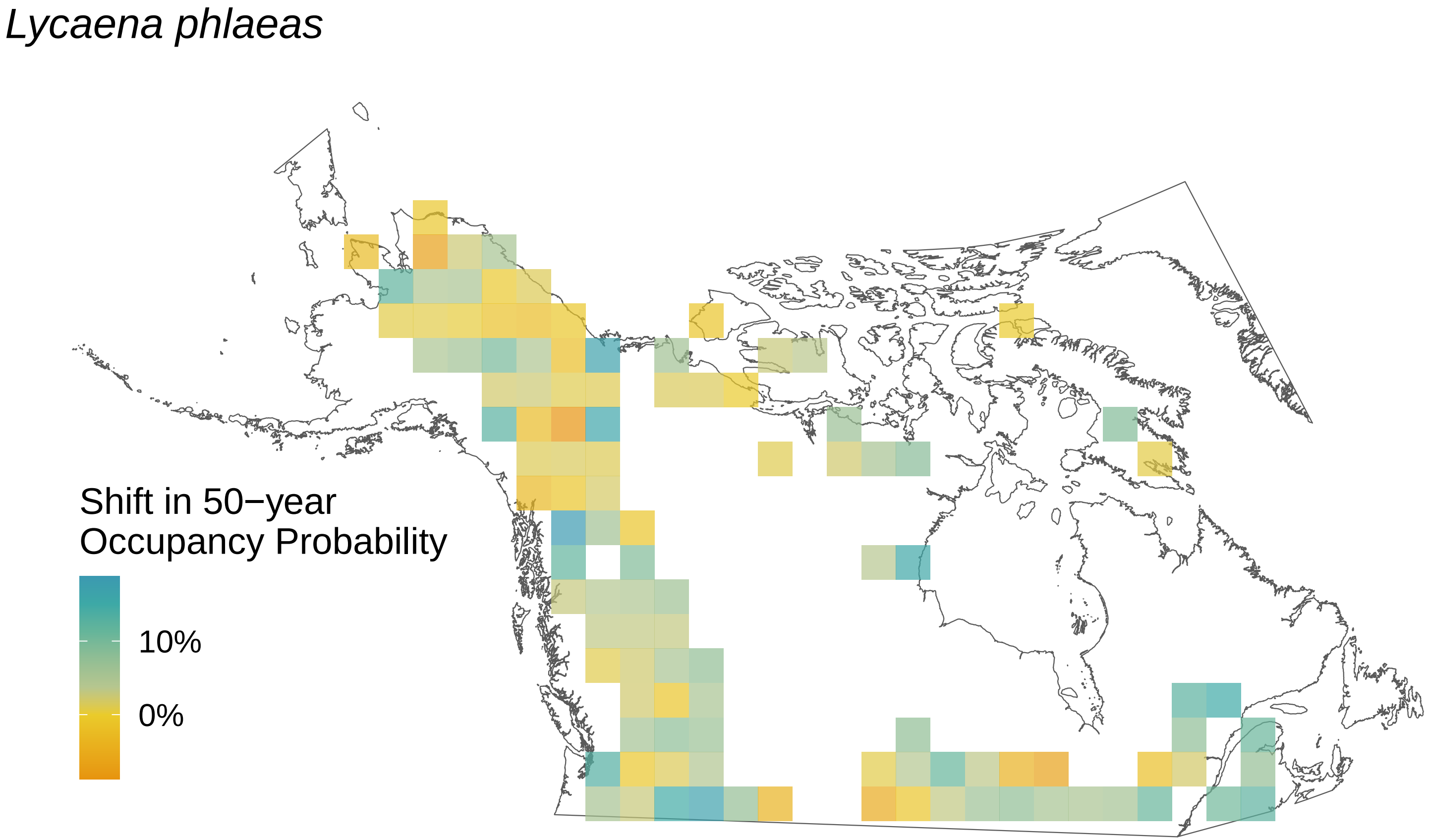
Lycaena phlaeas
Shift in 50−year
Occupancy Probability
10%
0%

*Lycaena dorcas*

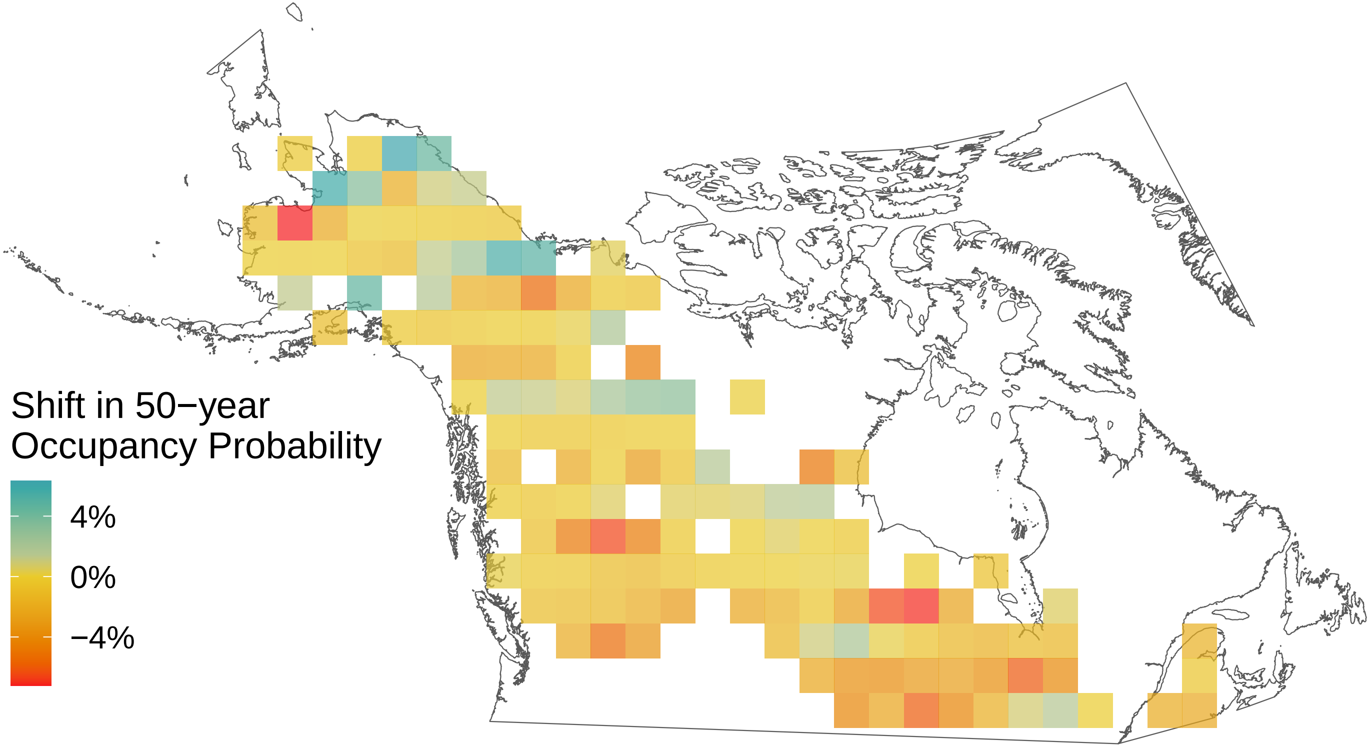

*Limenitis lorquini*

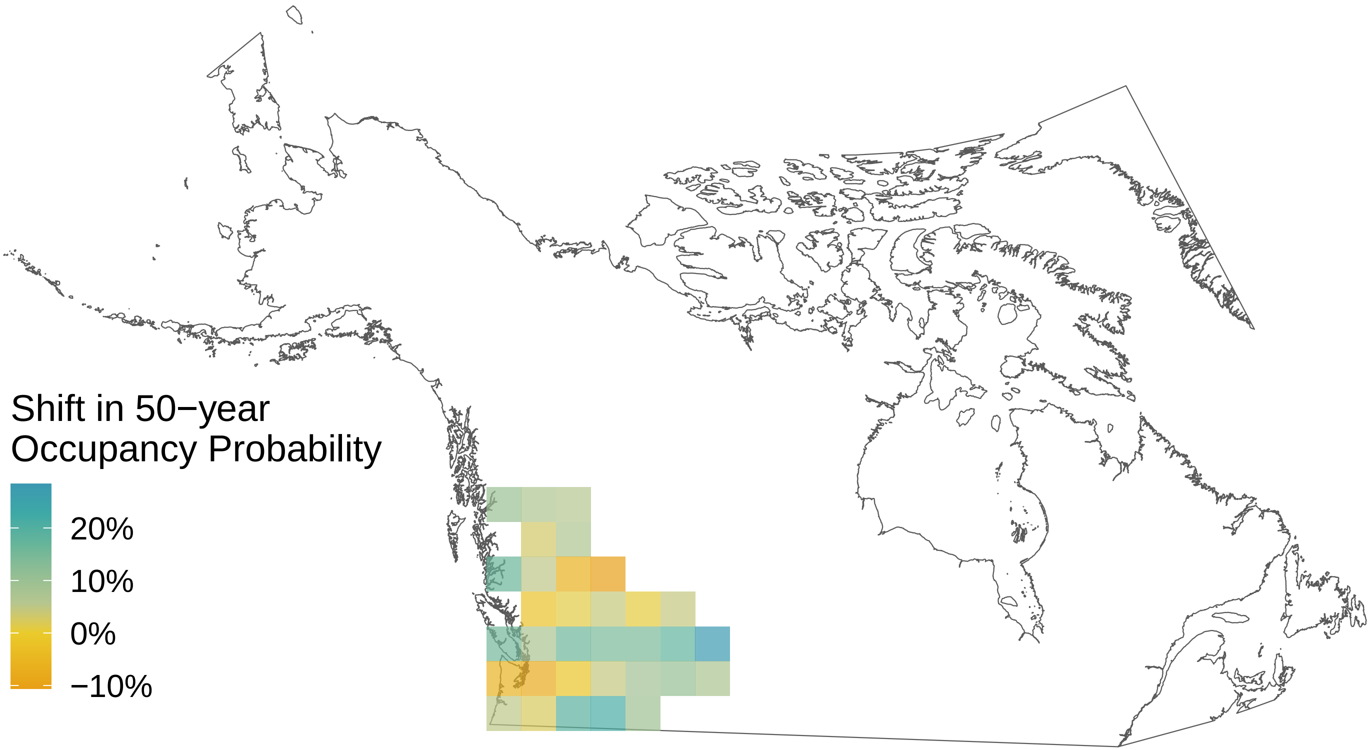

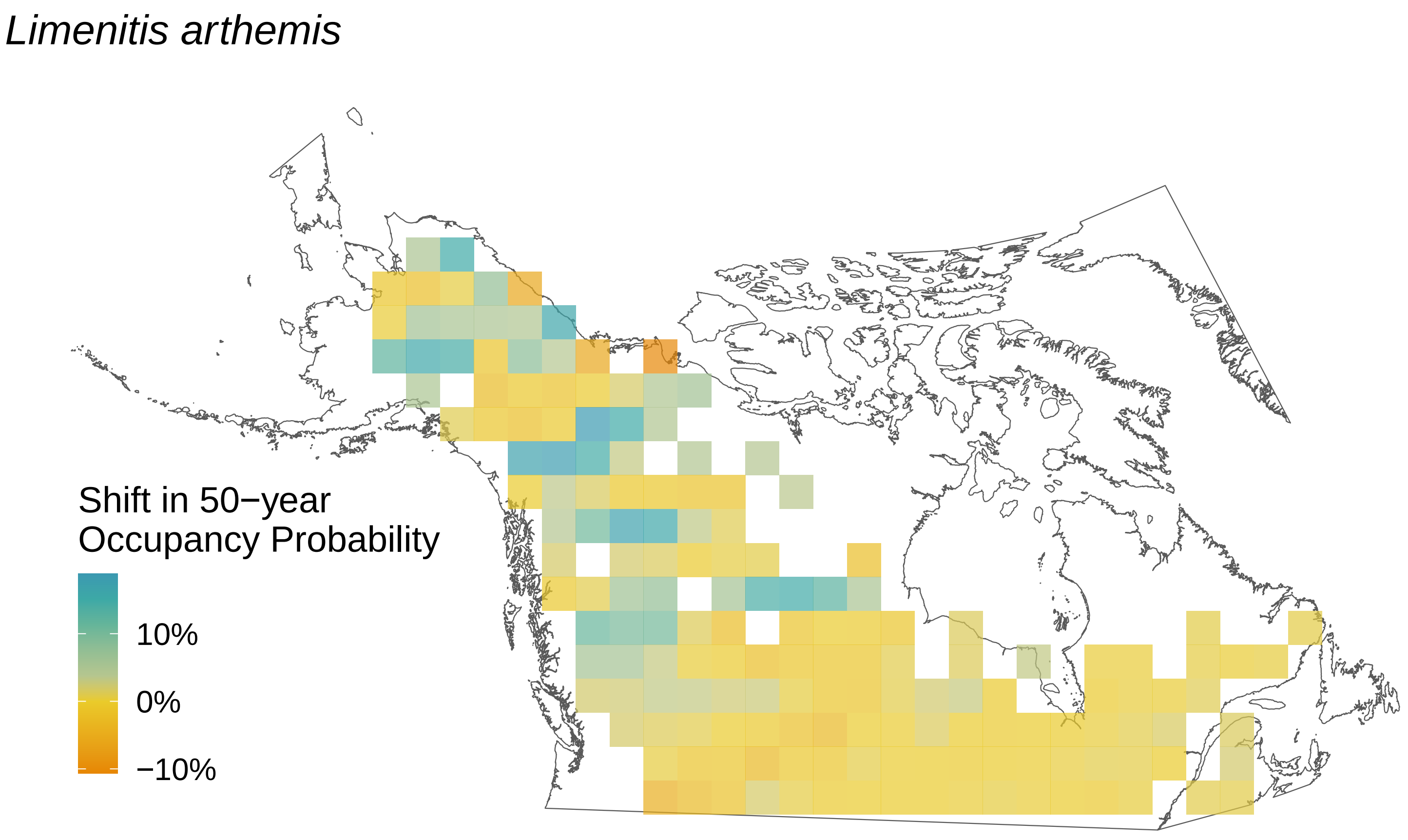
*Limenitis arthemis*
Shift in 50−year
Occupancy Probability
10%
0%
−10%

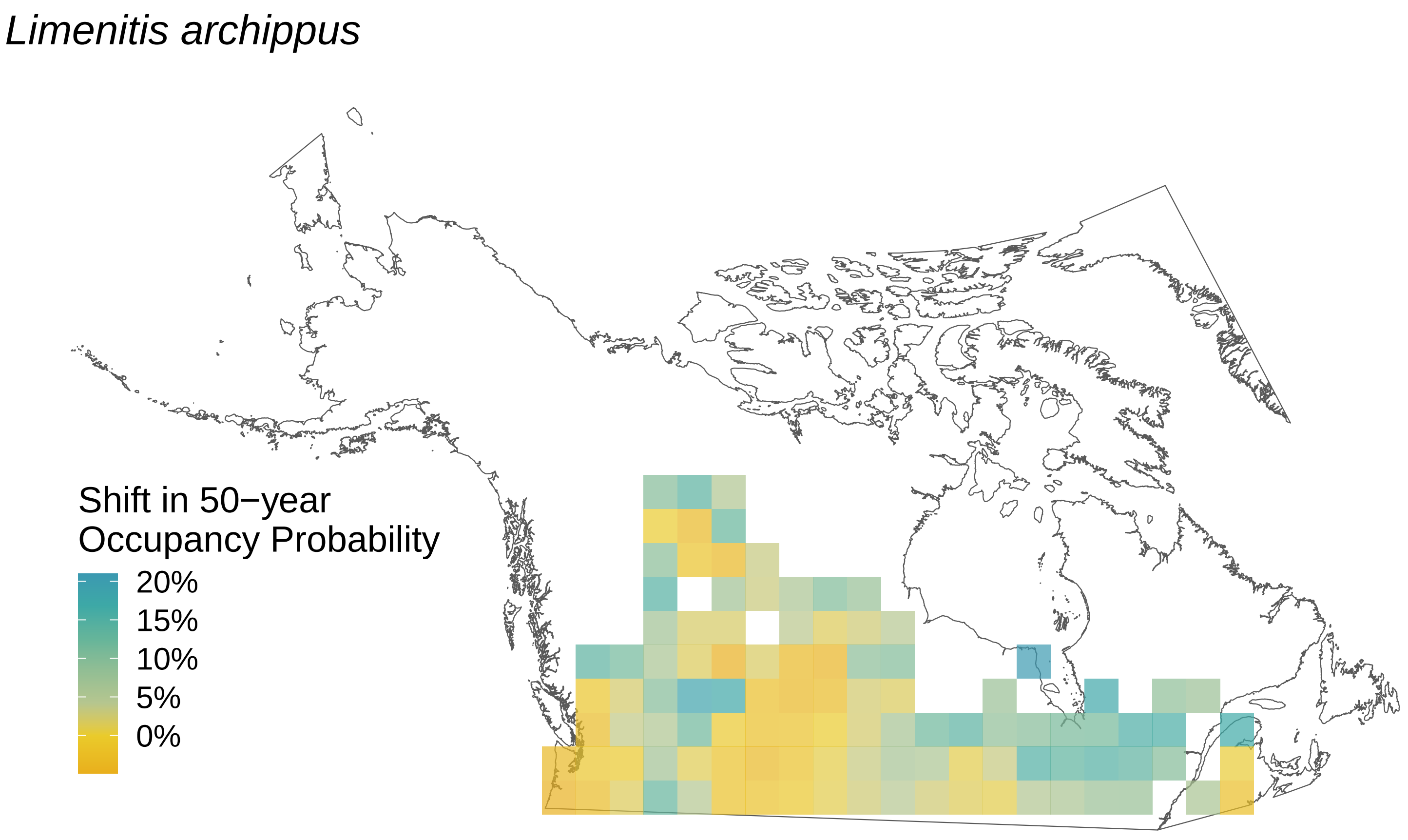

Limenitis archippus
Shift in 50−year
Occupancy Probability
20%
15%
10%
5%
0%

*Lethe eurydice*

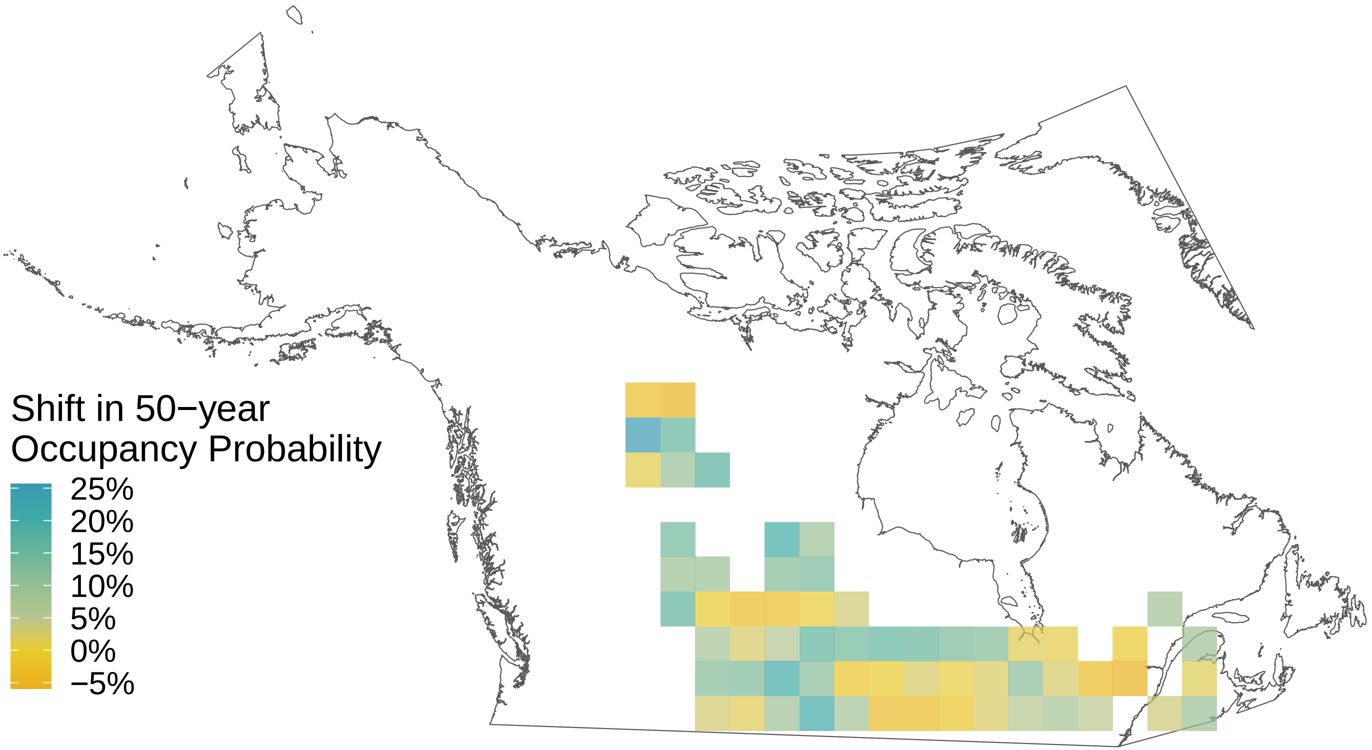

*Icaricia saepiolus*

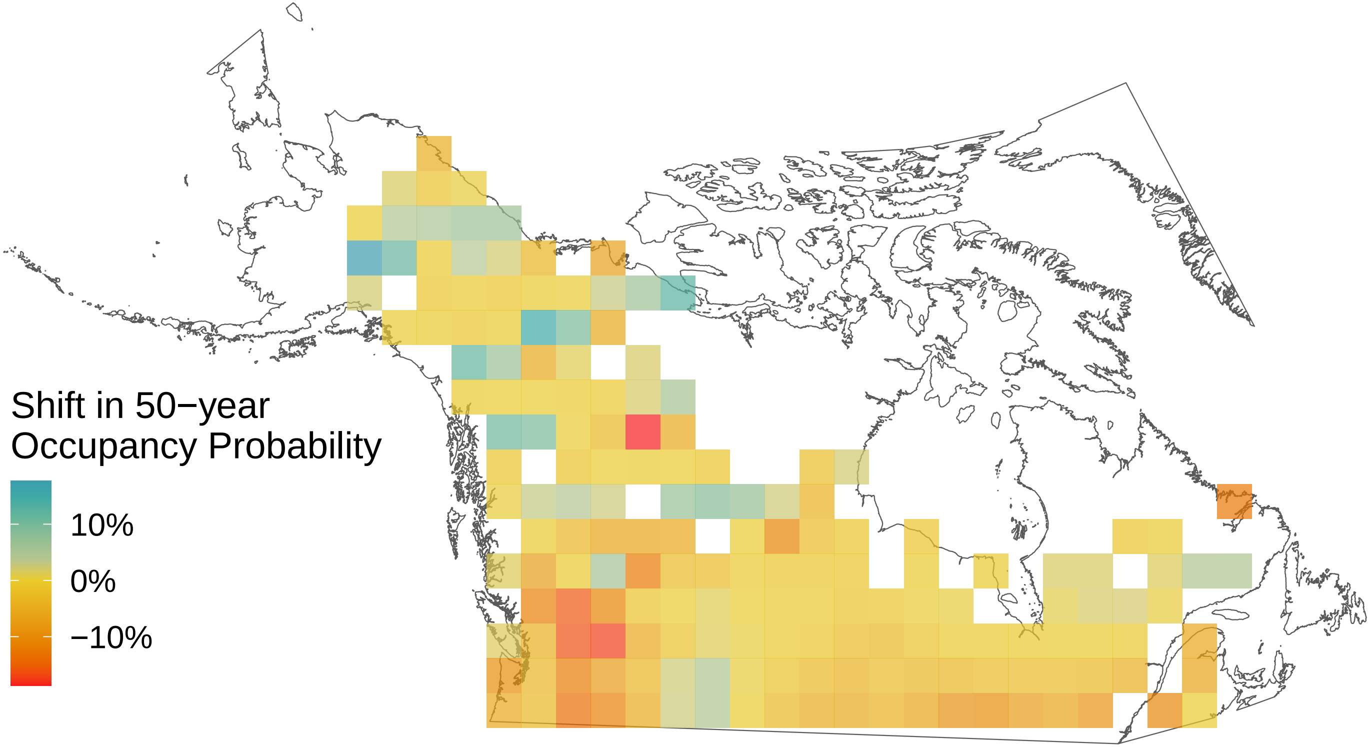

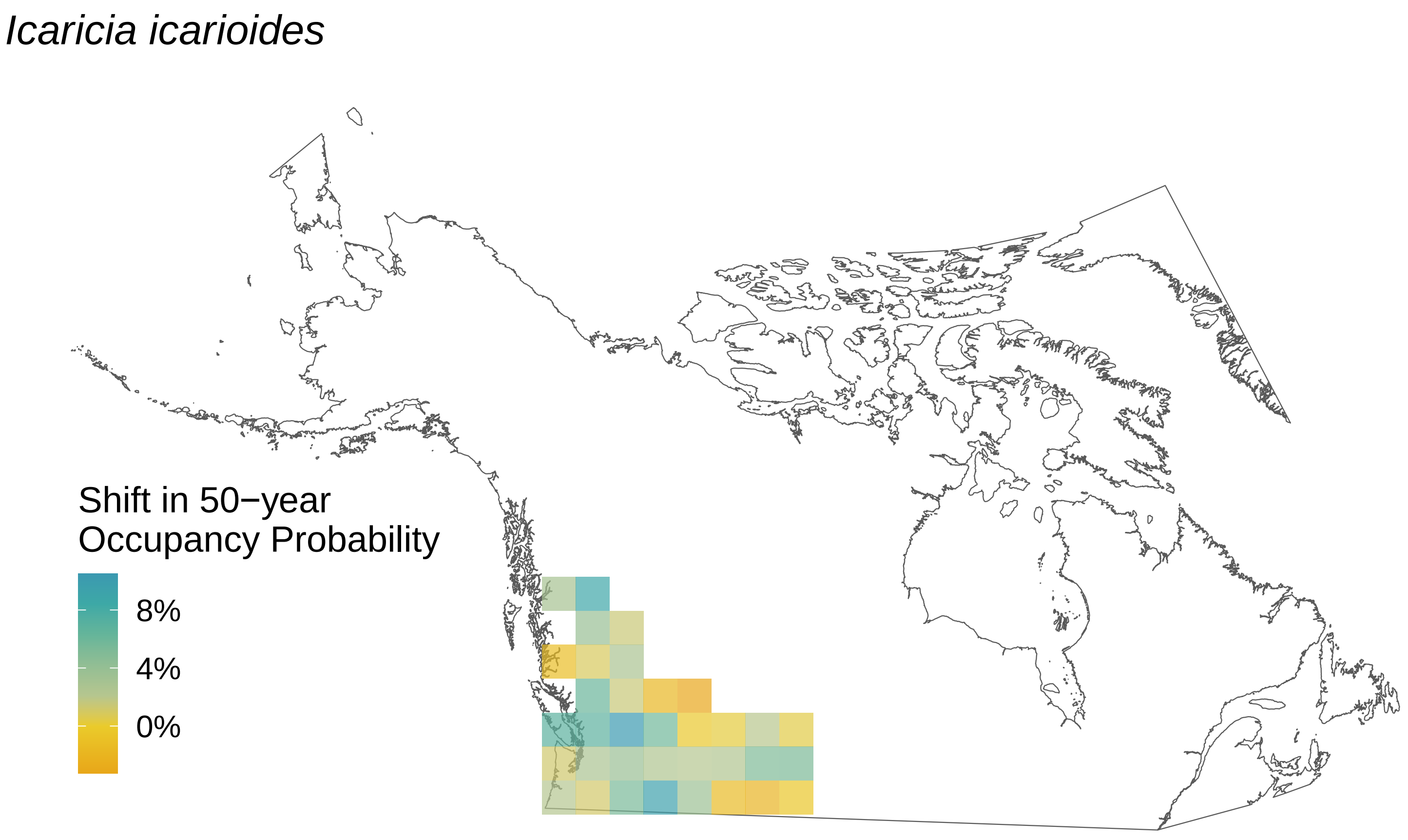

Icaricia icarioides
Shift in 50−year
Occupancy Probability
8%
4%
0%

*Hesperia comma*

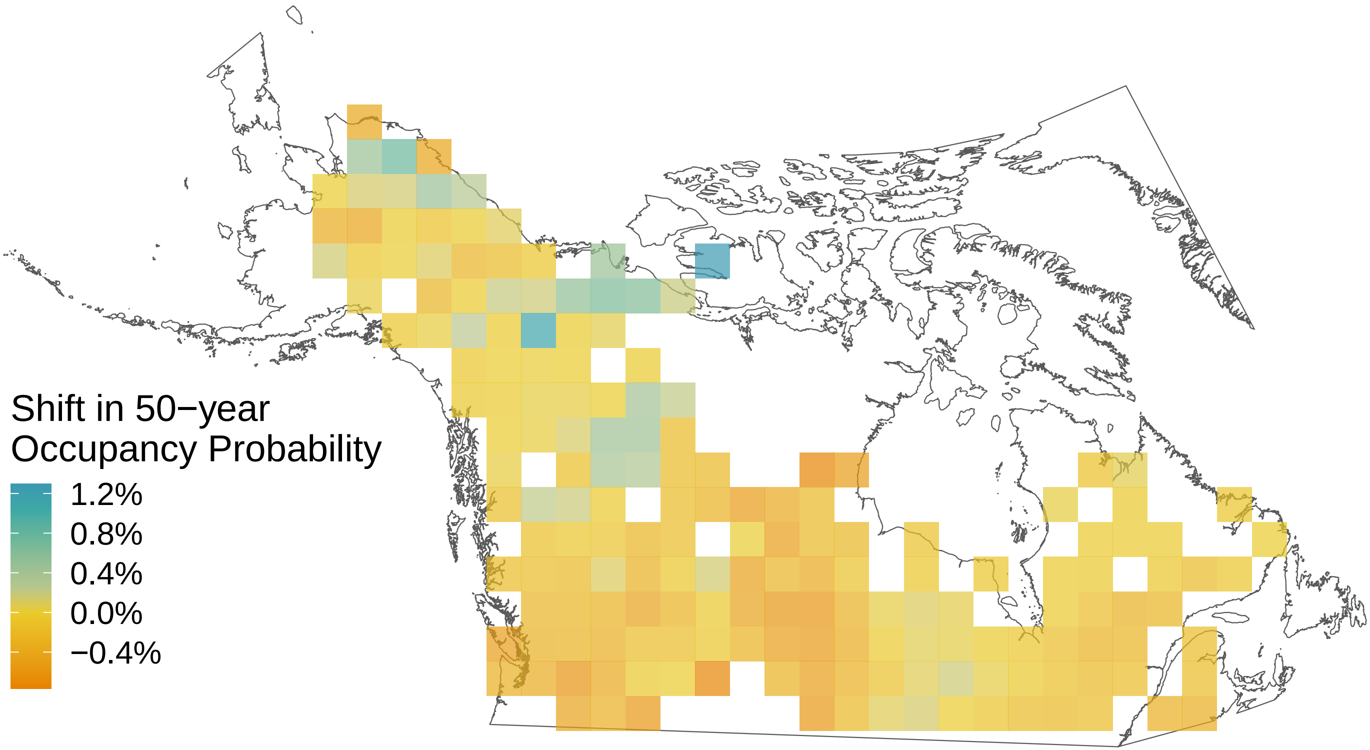

*Glaucopsyche lygdamus*

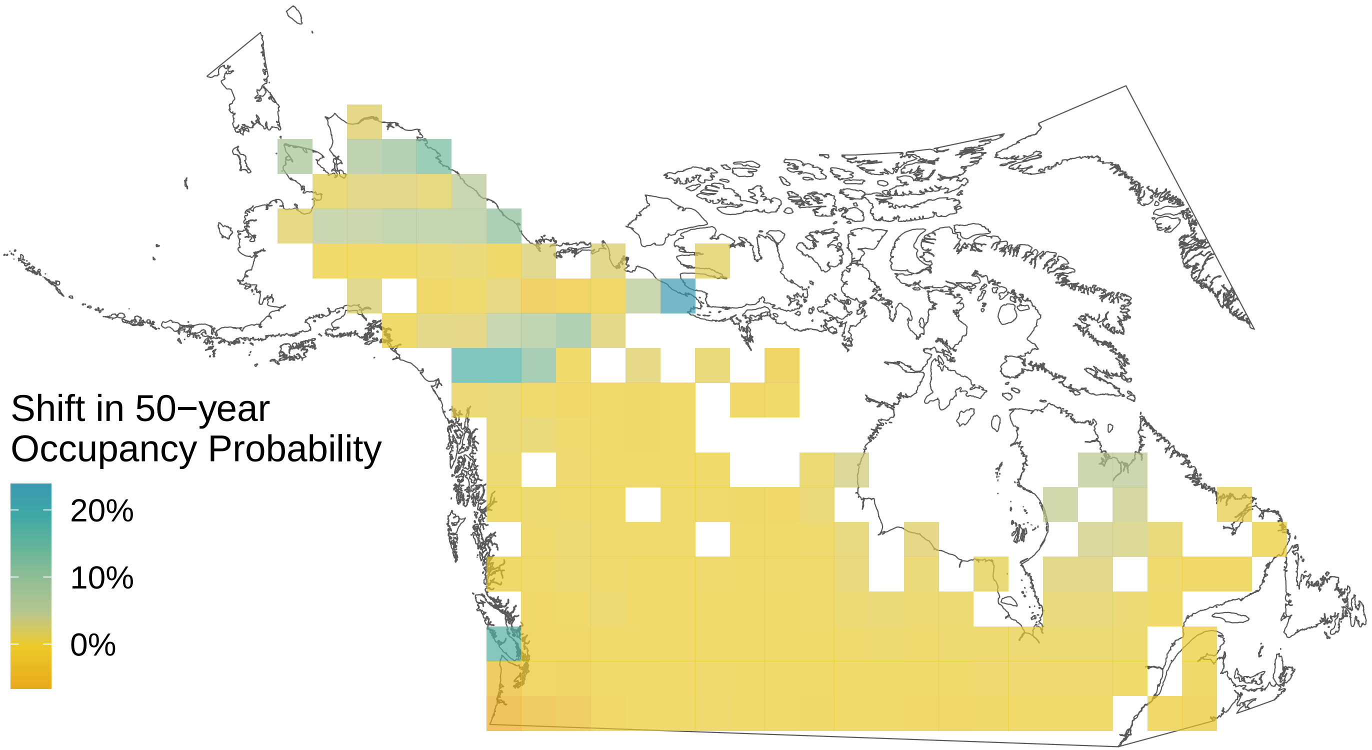

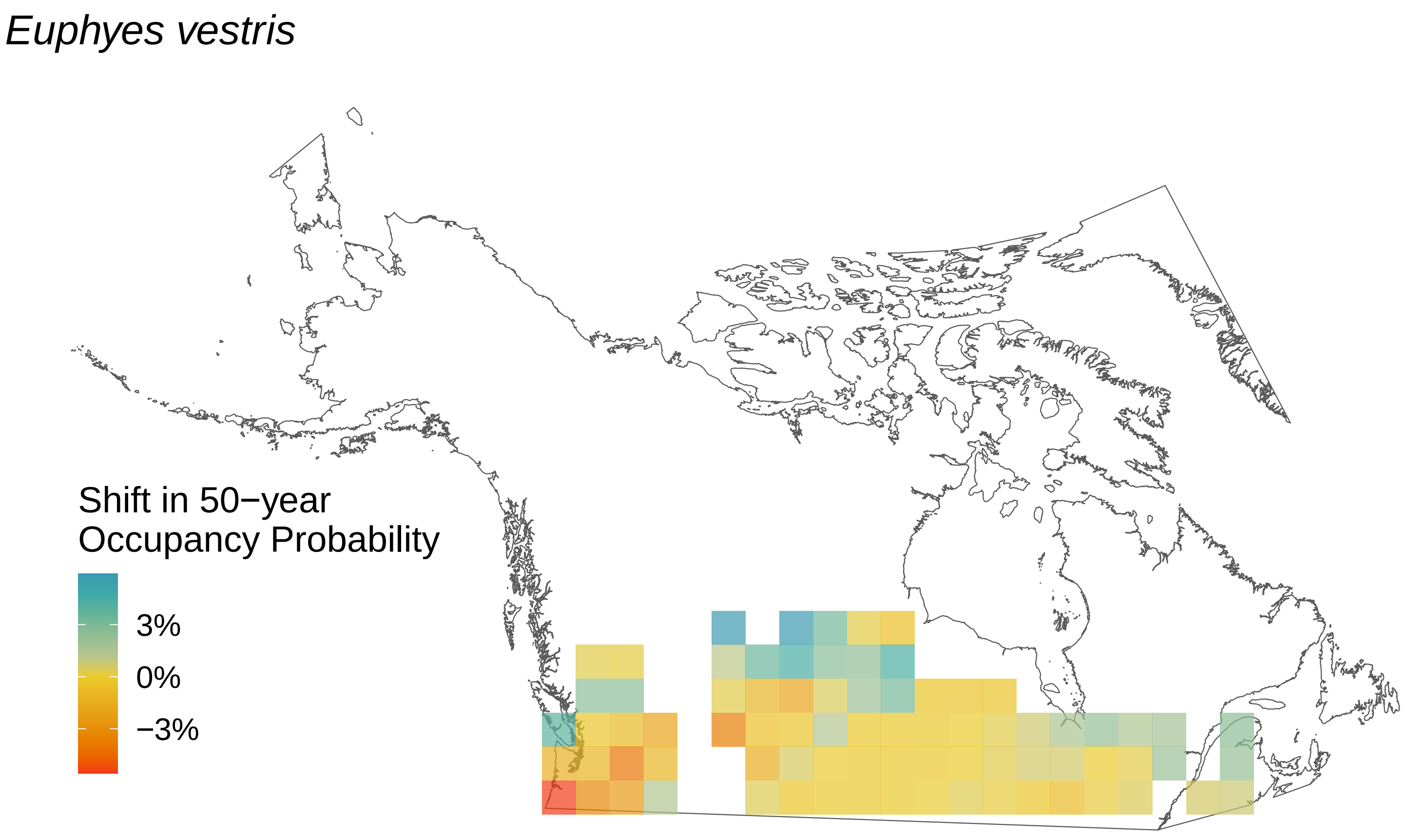

Euphyes vestris
Shift in 50−year
Occupancy Probability
3%
0%
−3%

*Euphydryas phaeton*

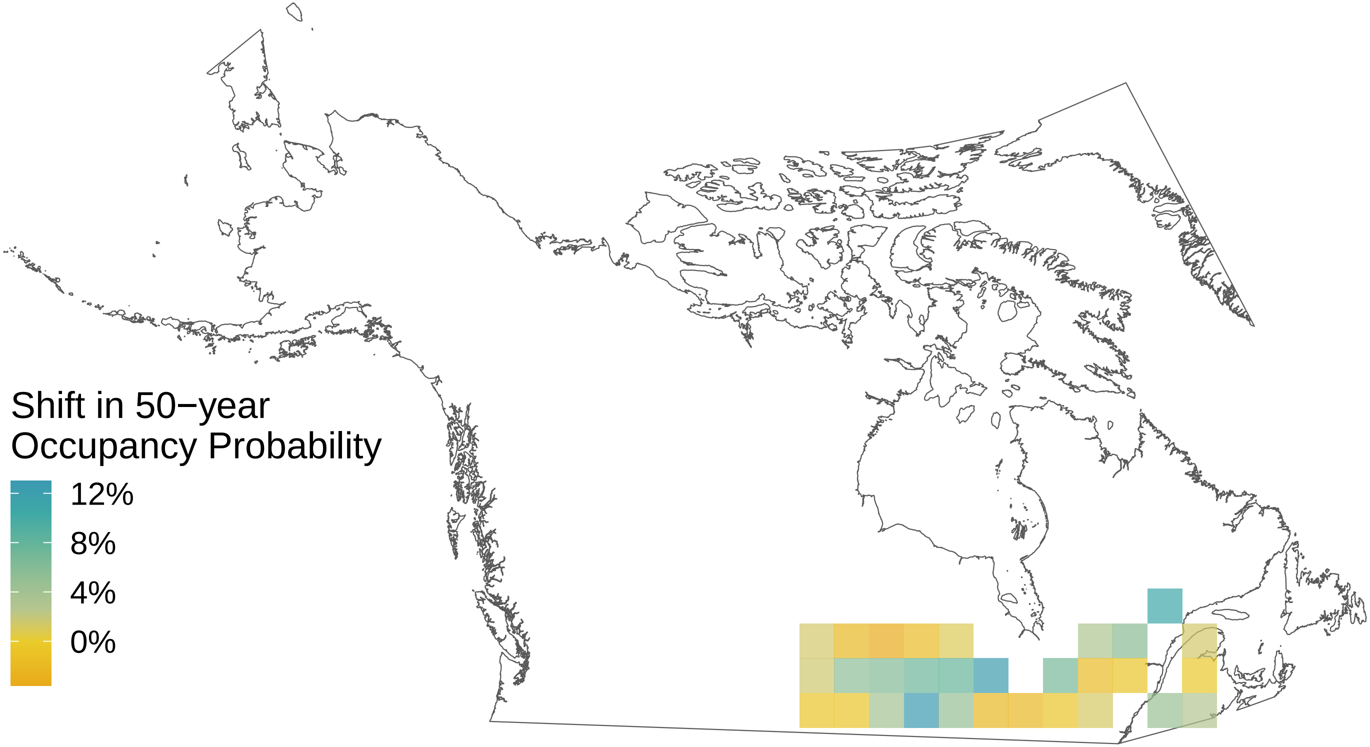

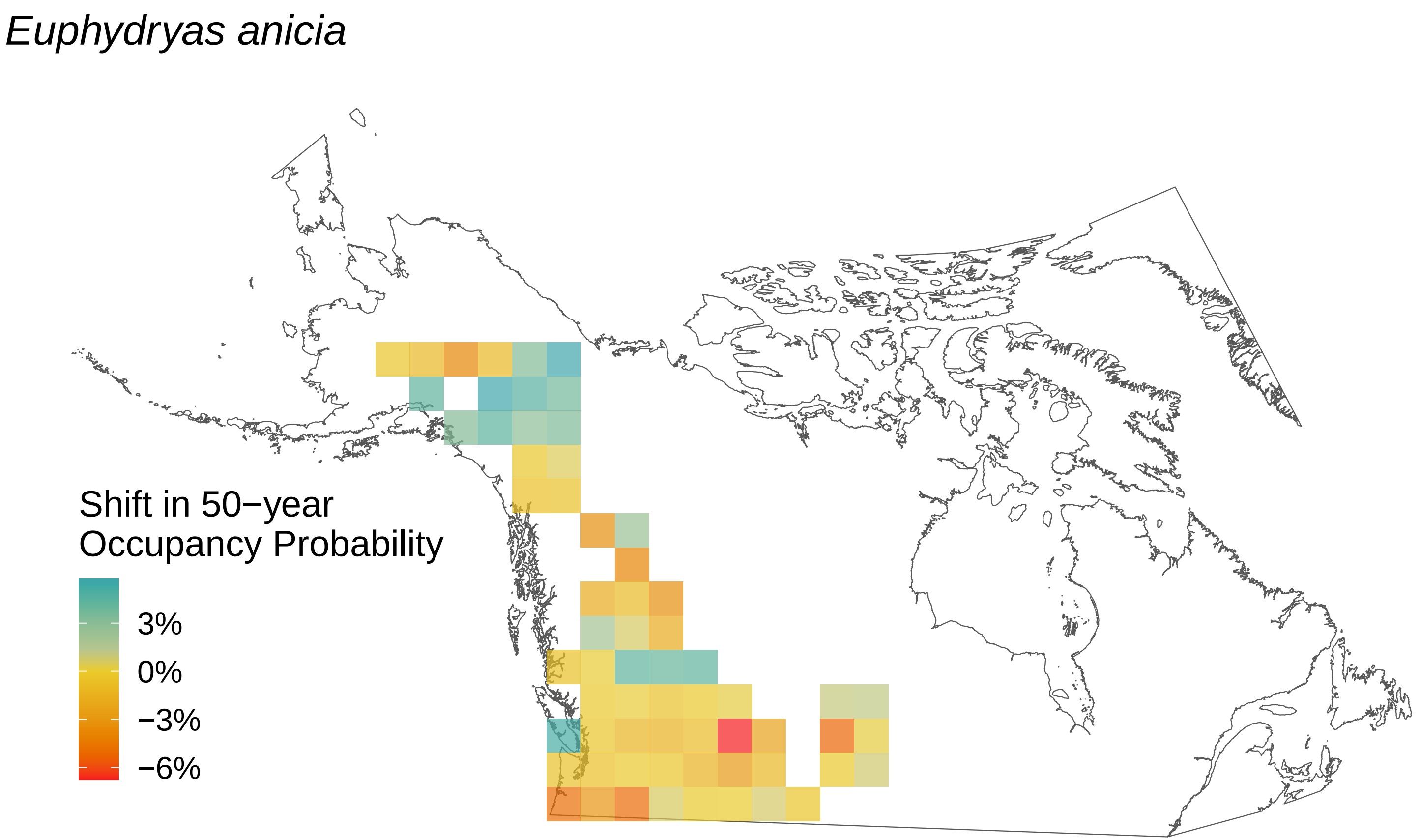
Euphydryas anicia
Shift in 50−year
Occupancy Probability
3%
0%
−3%
−6%

*Euchloe ausonides*

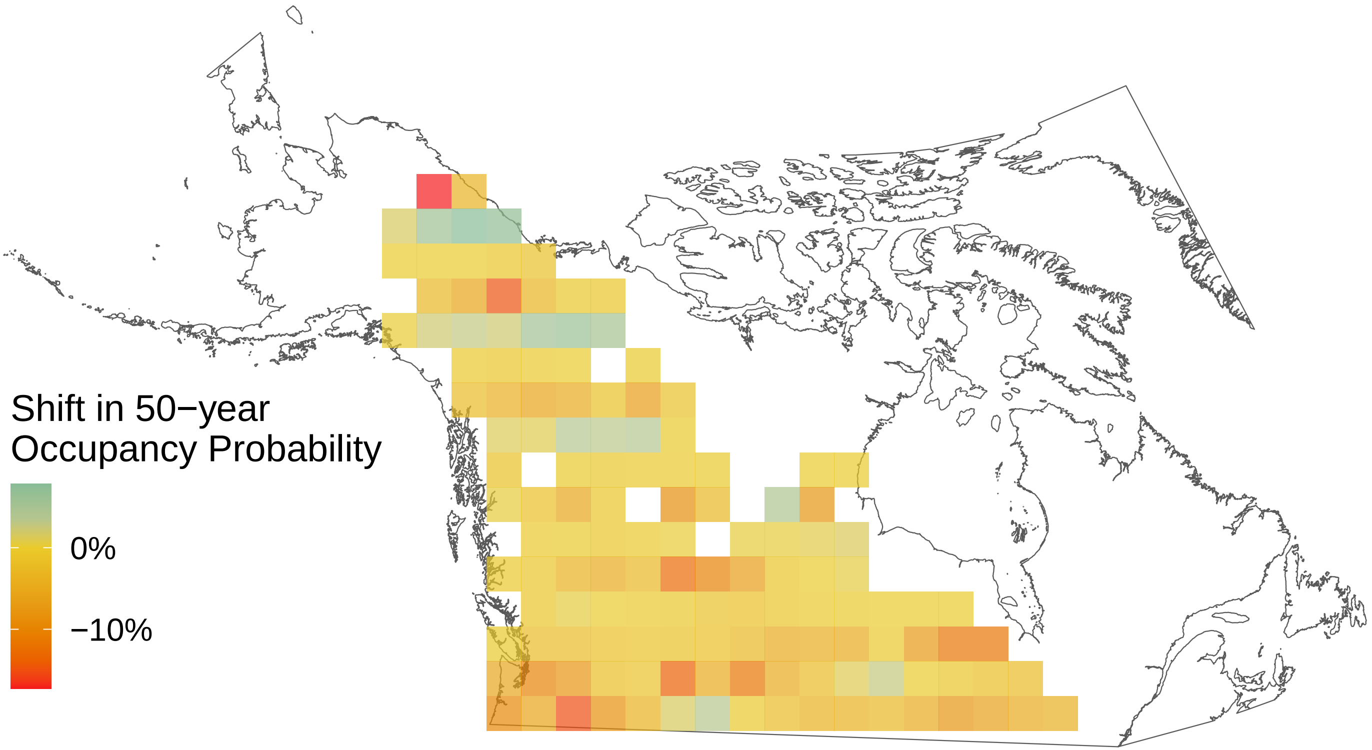

*Erynnis persius*

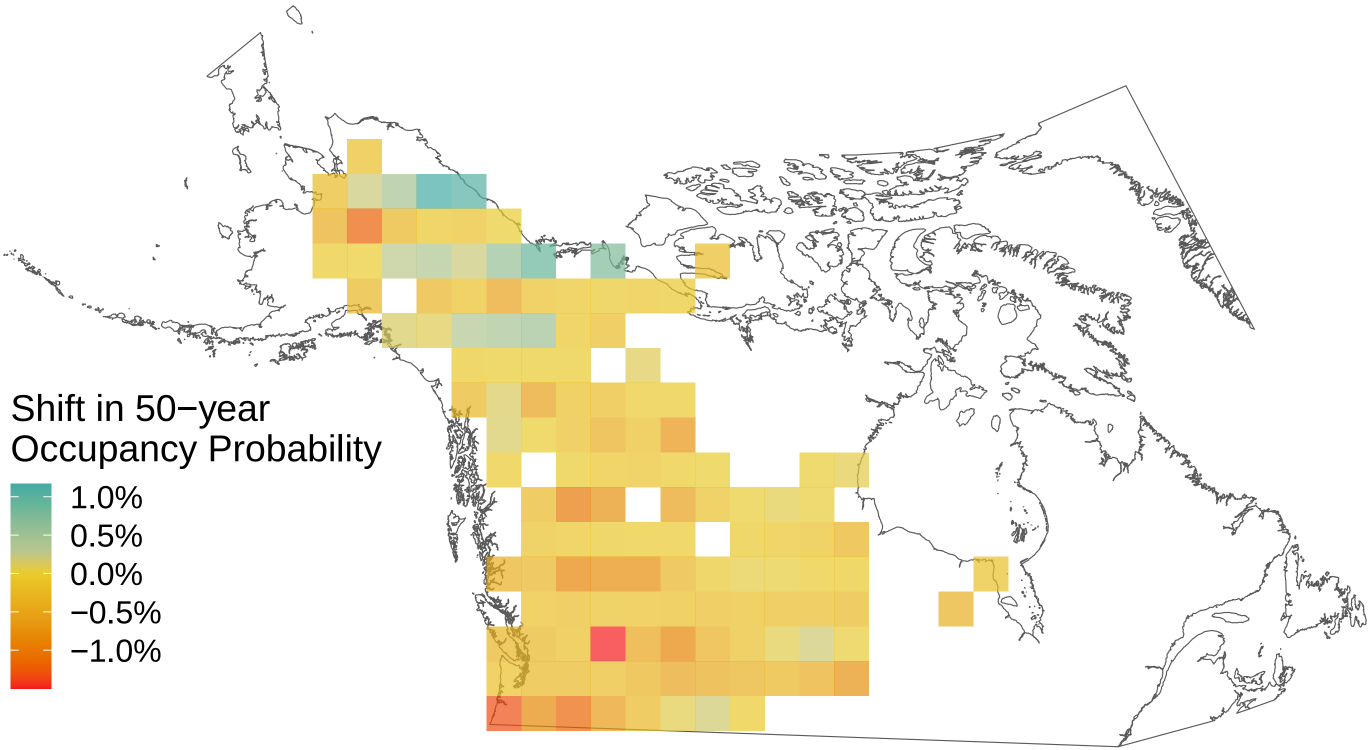

*Erynnis juvenalis*

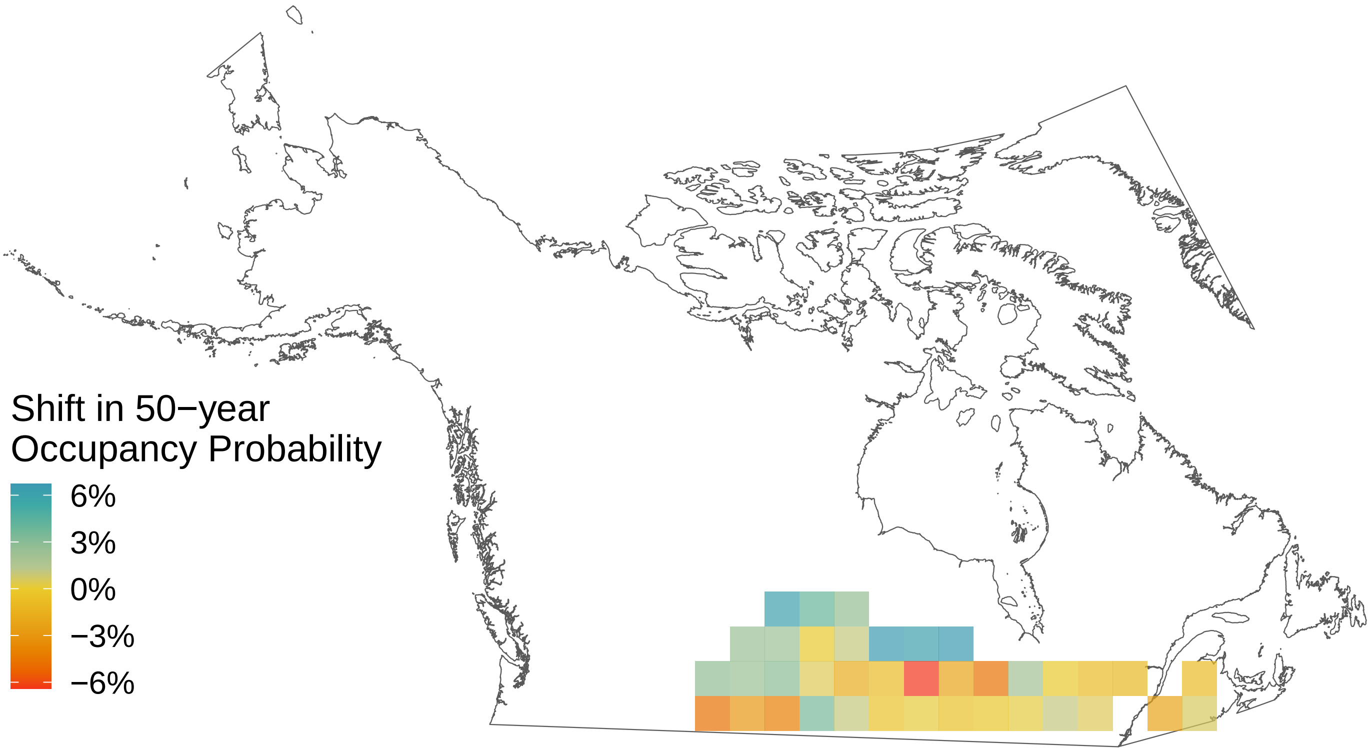

*Erynnis icelus*

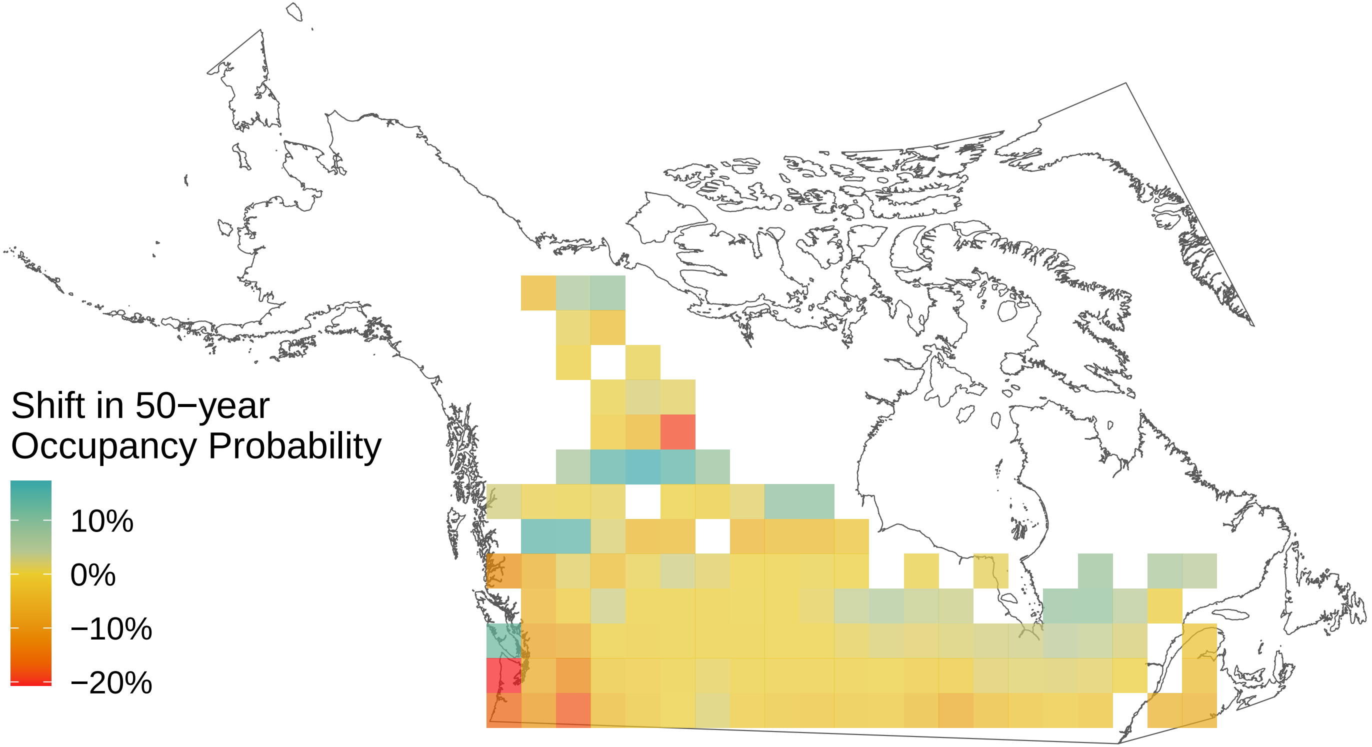

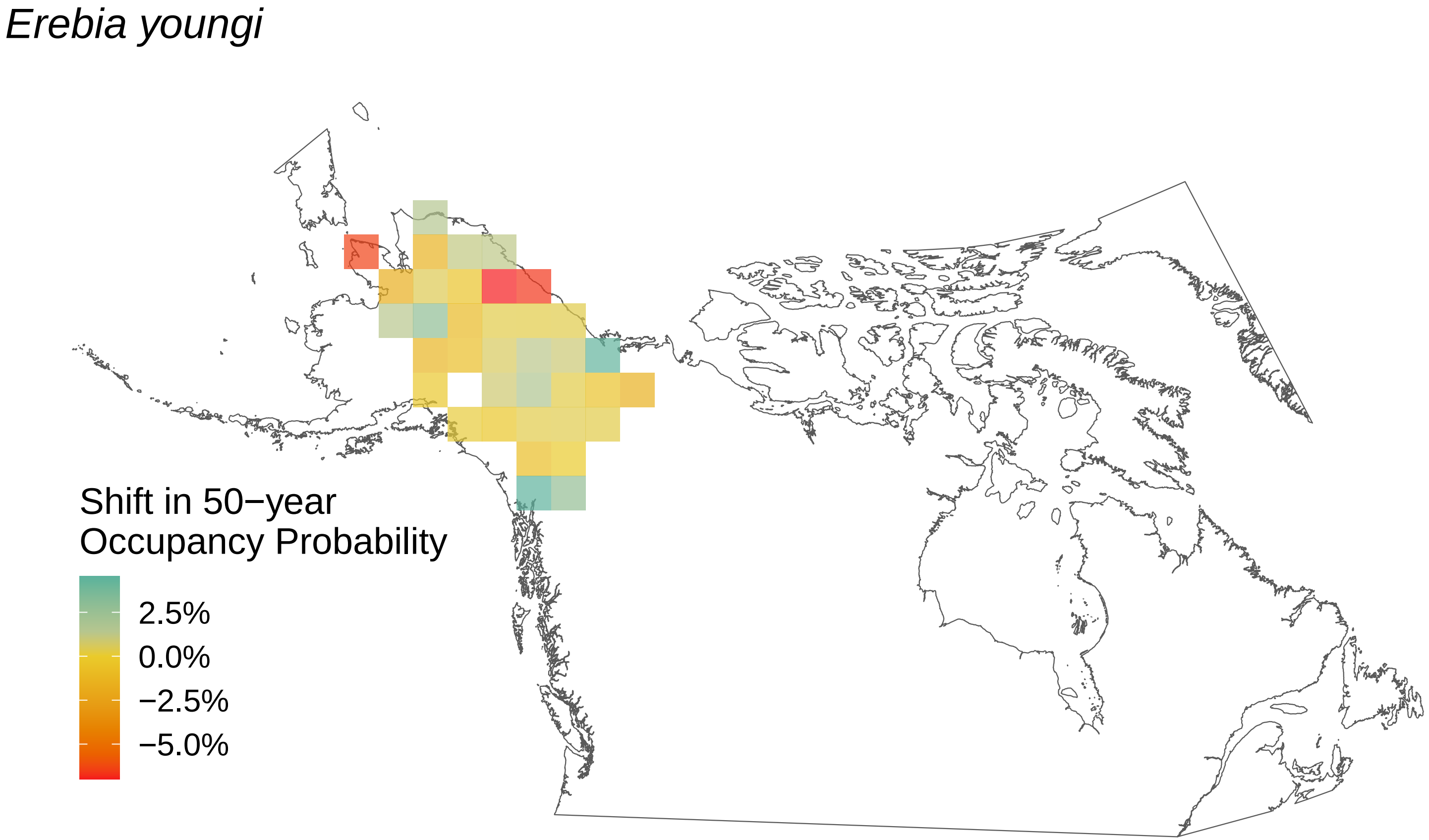
Erebia youngi
Shift in 50−year
Occupancy Probability
2.5%
0.0%
−2.5%
−5.0%

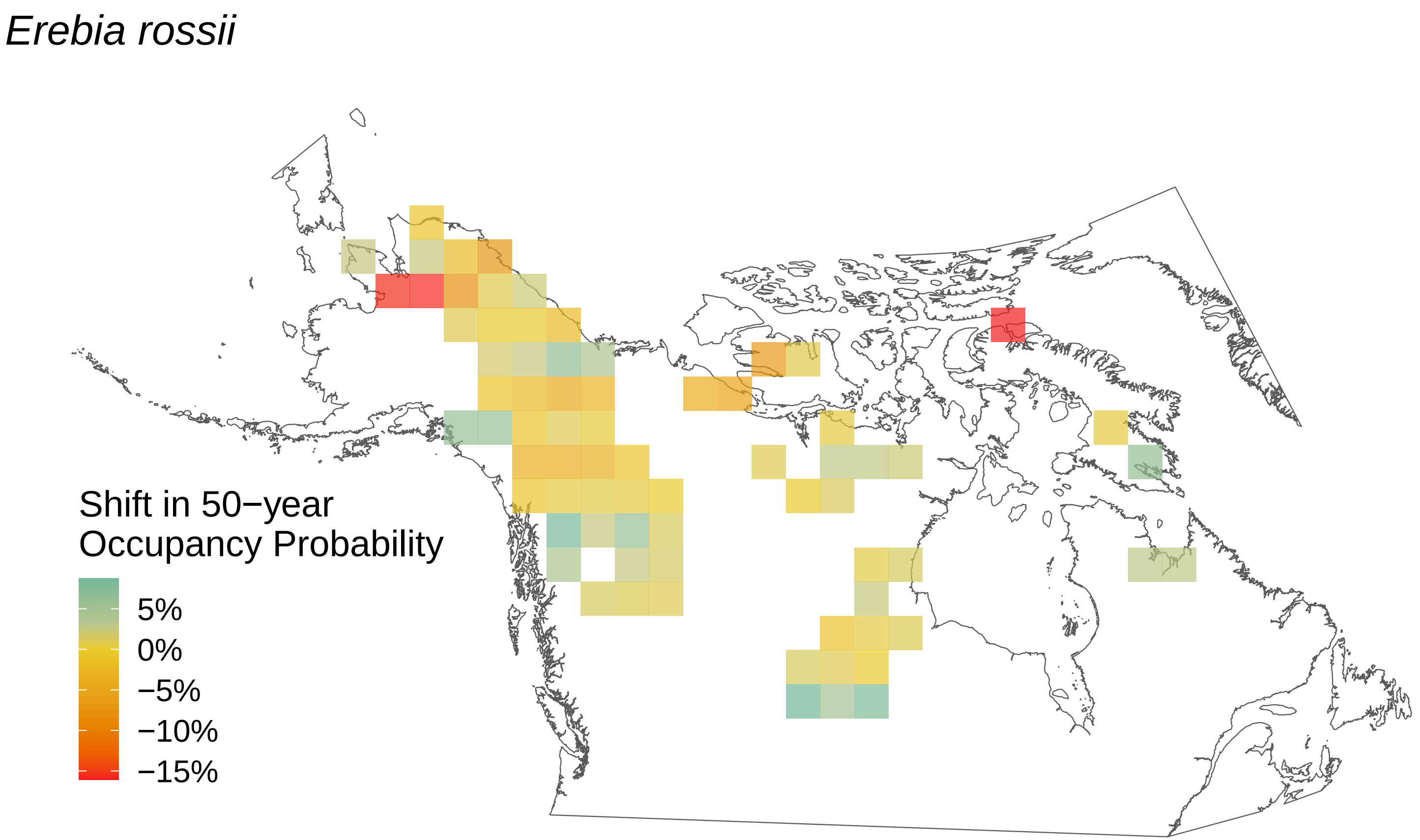
Erebia rossii
Shift in 50−year
Occupancy Probability
5%
0%
−5%
−10%
−15%

*Erebia fasciata*

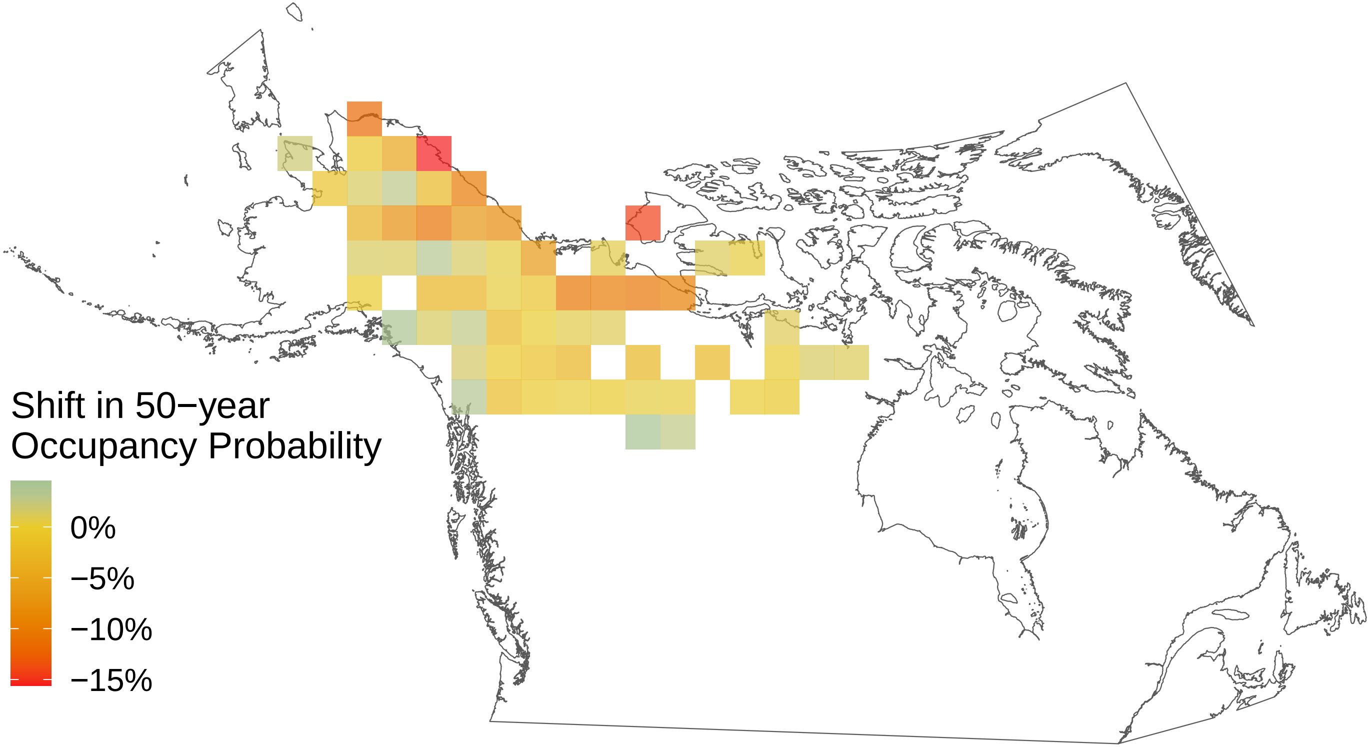

### *Erebia epipsodea*

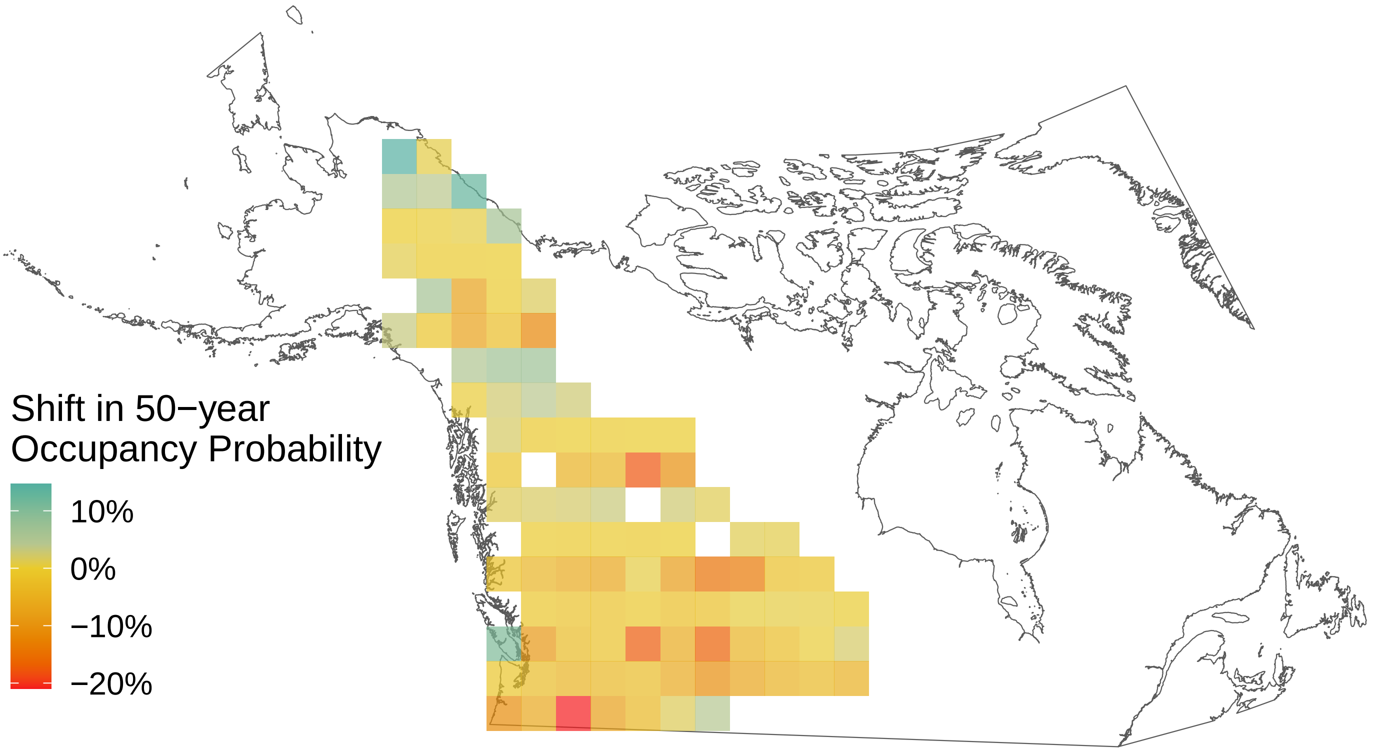

*Erebia discoidalis*

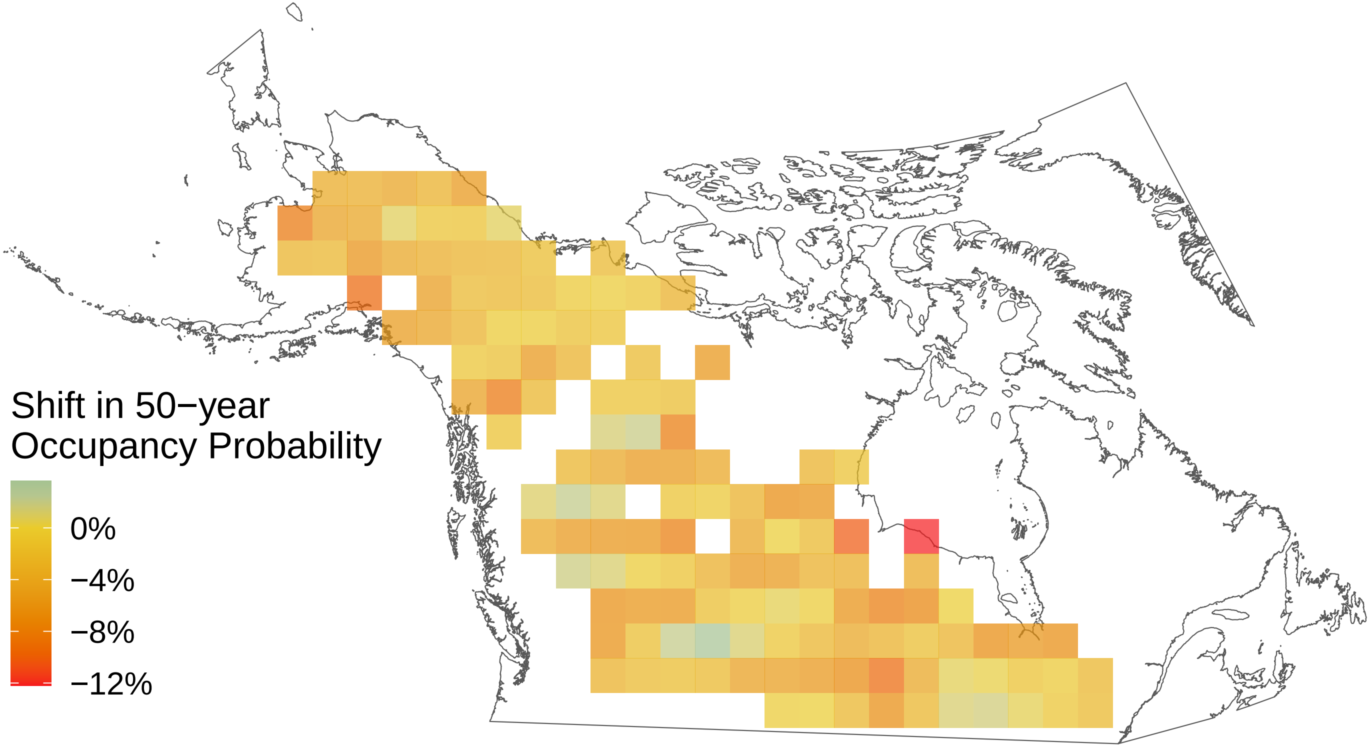

*Epargyreus clarus*

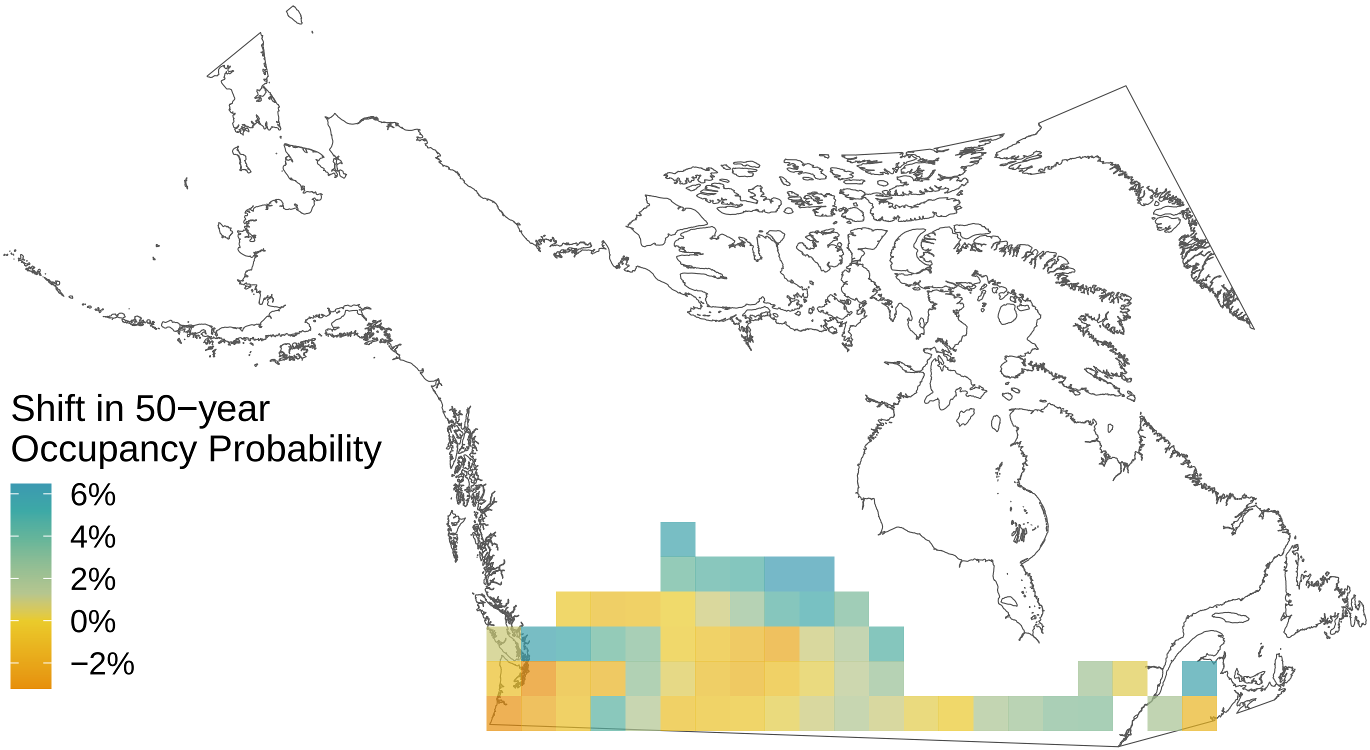

*Cupido comyntas*

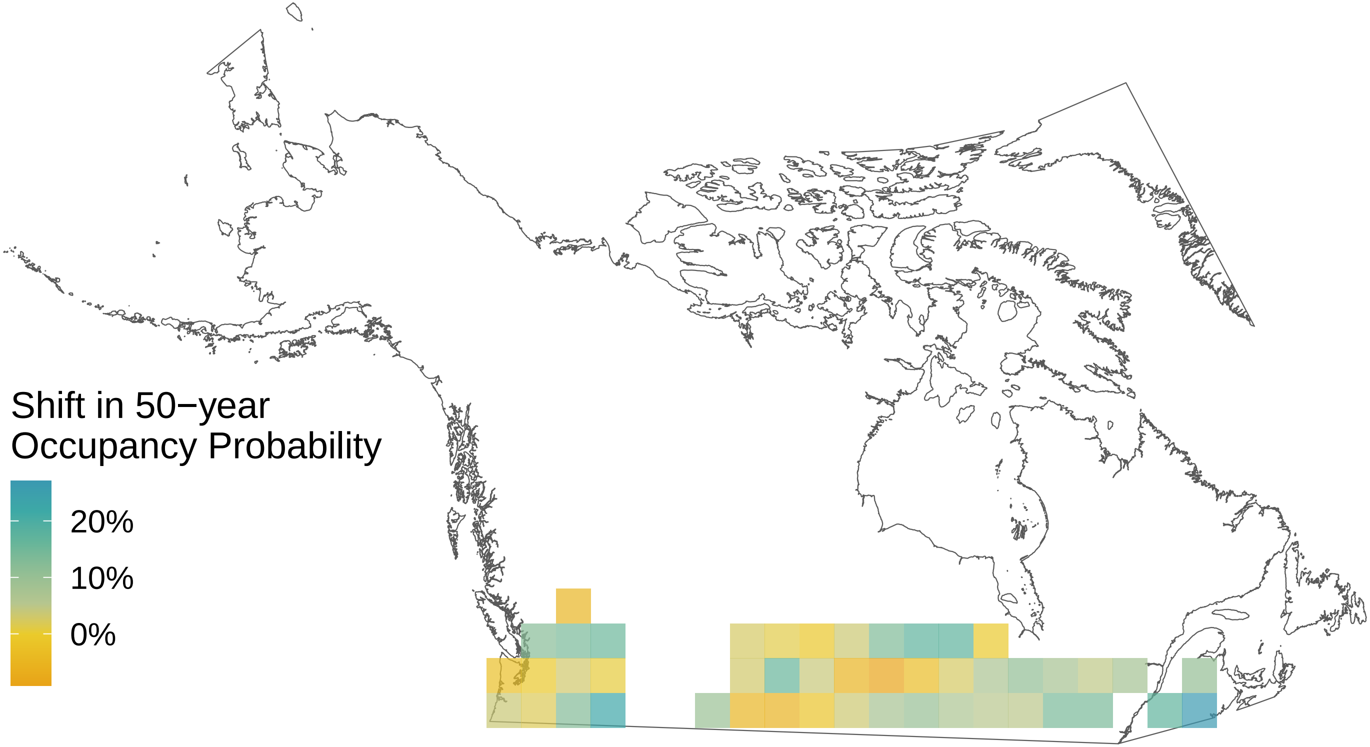

*Cupido amyntula*

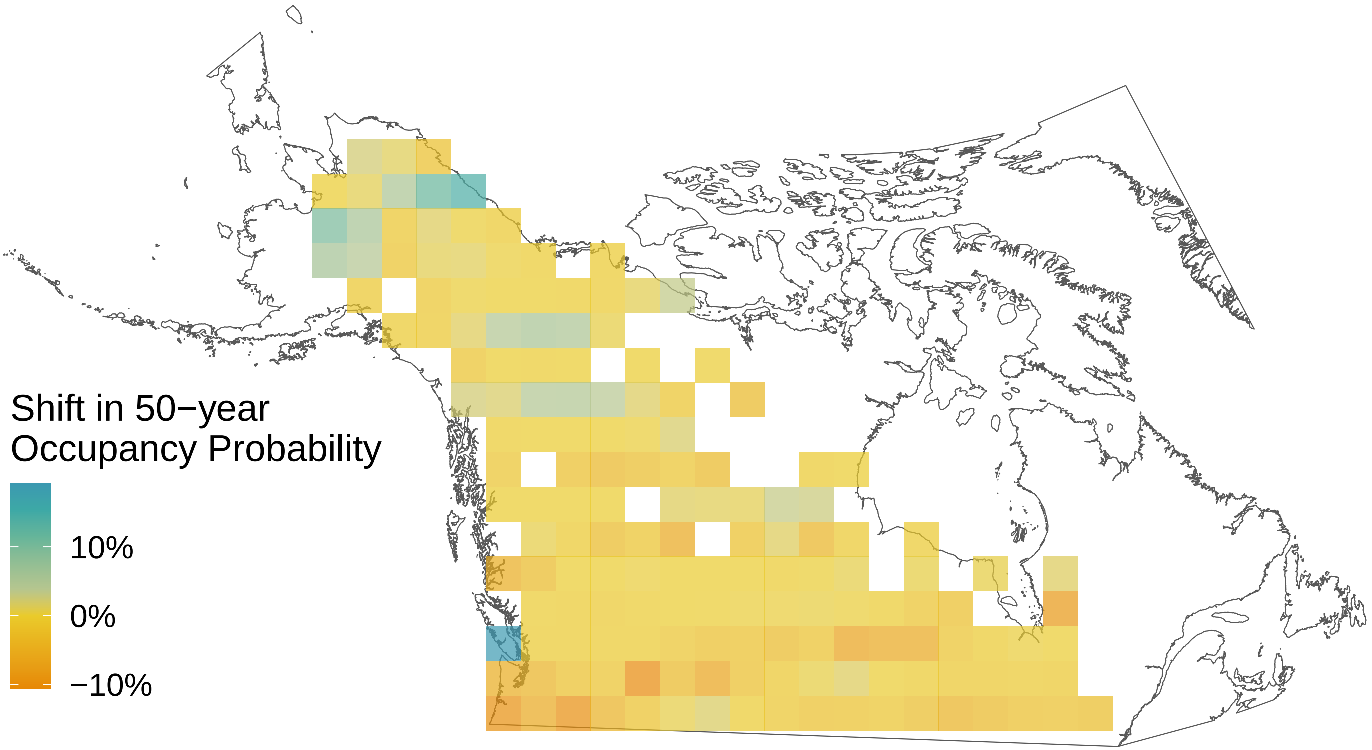

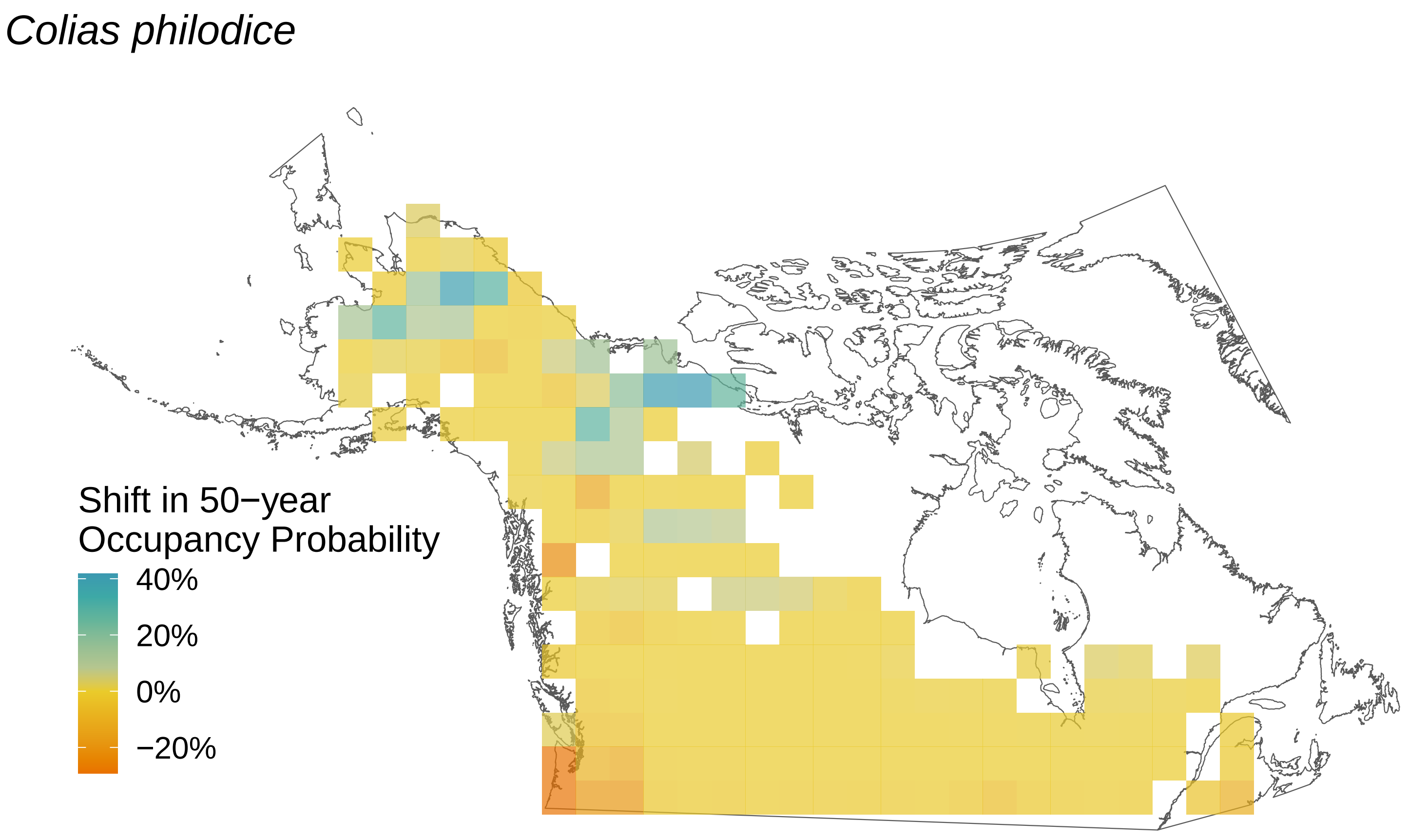

Colias philodice
Shift in 50−year
Occupancy Probability
40%
20%
0%
−20%

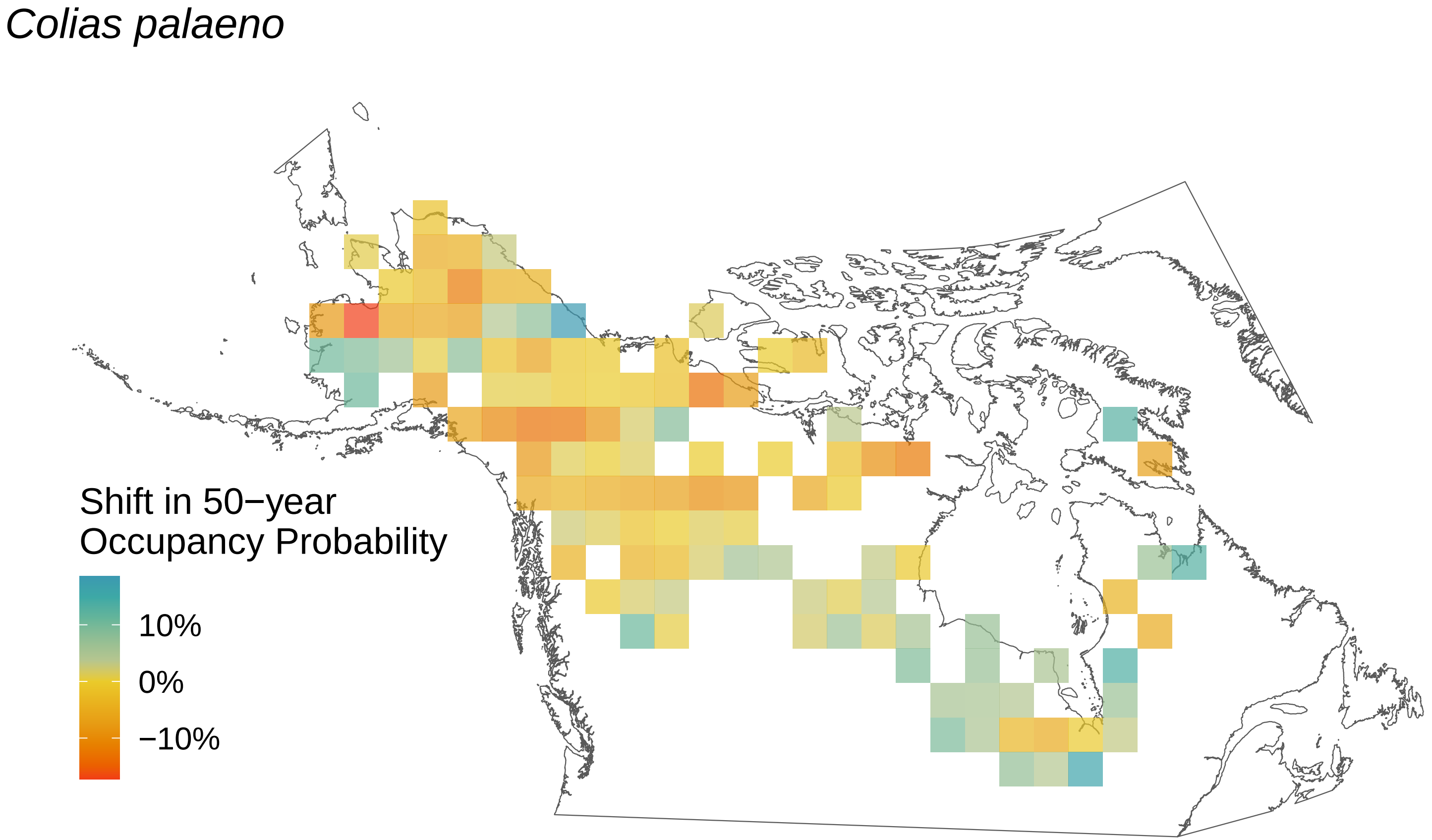
Colias palaeno
Shift in 50−year Occupancy Probability
10%
0%
−10%

*Colias interior*

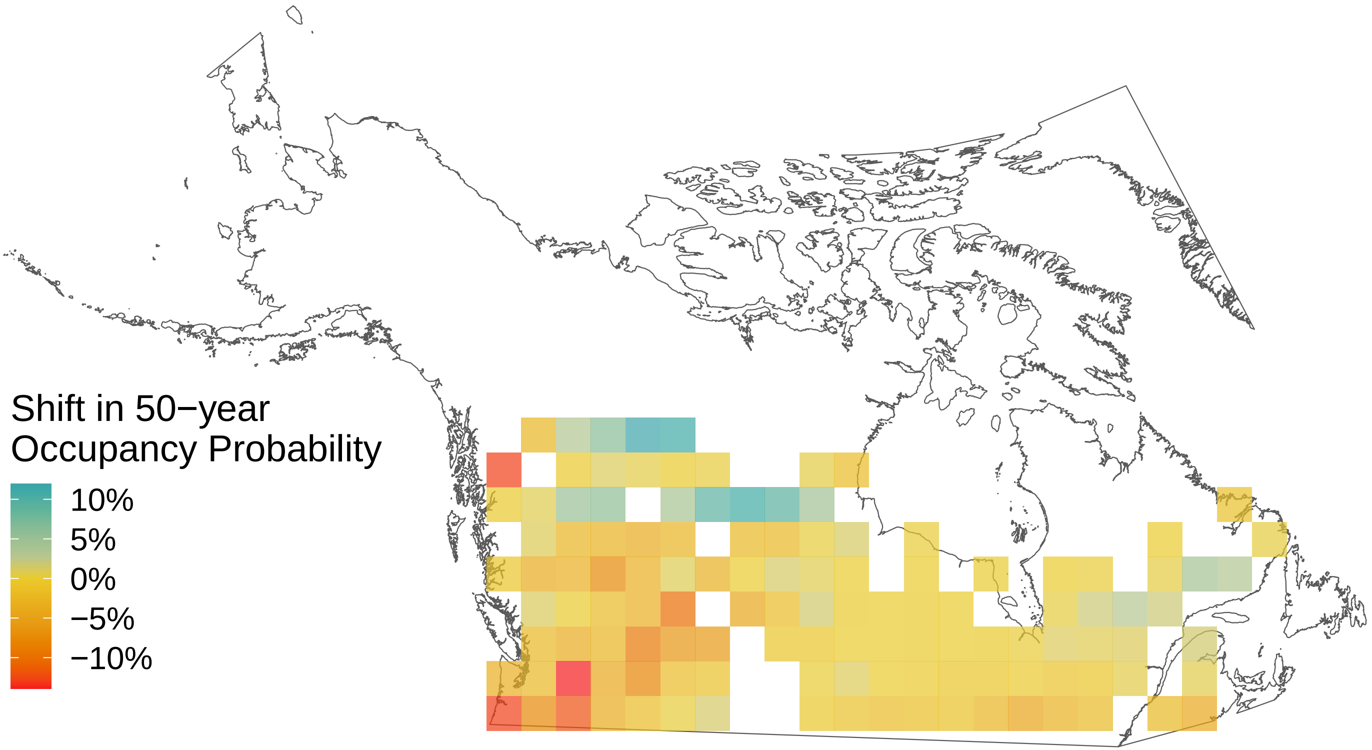

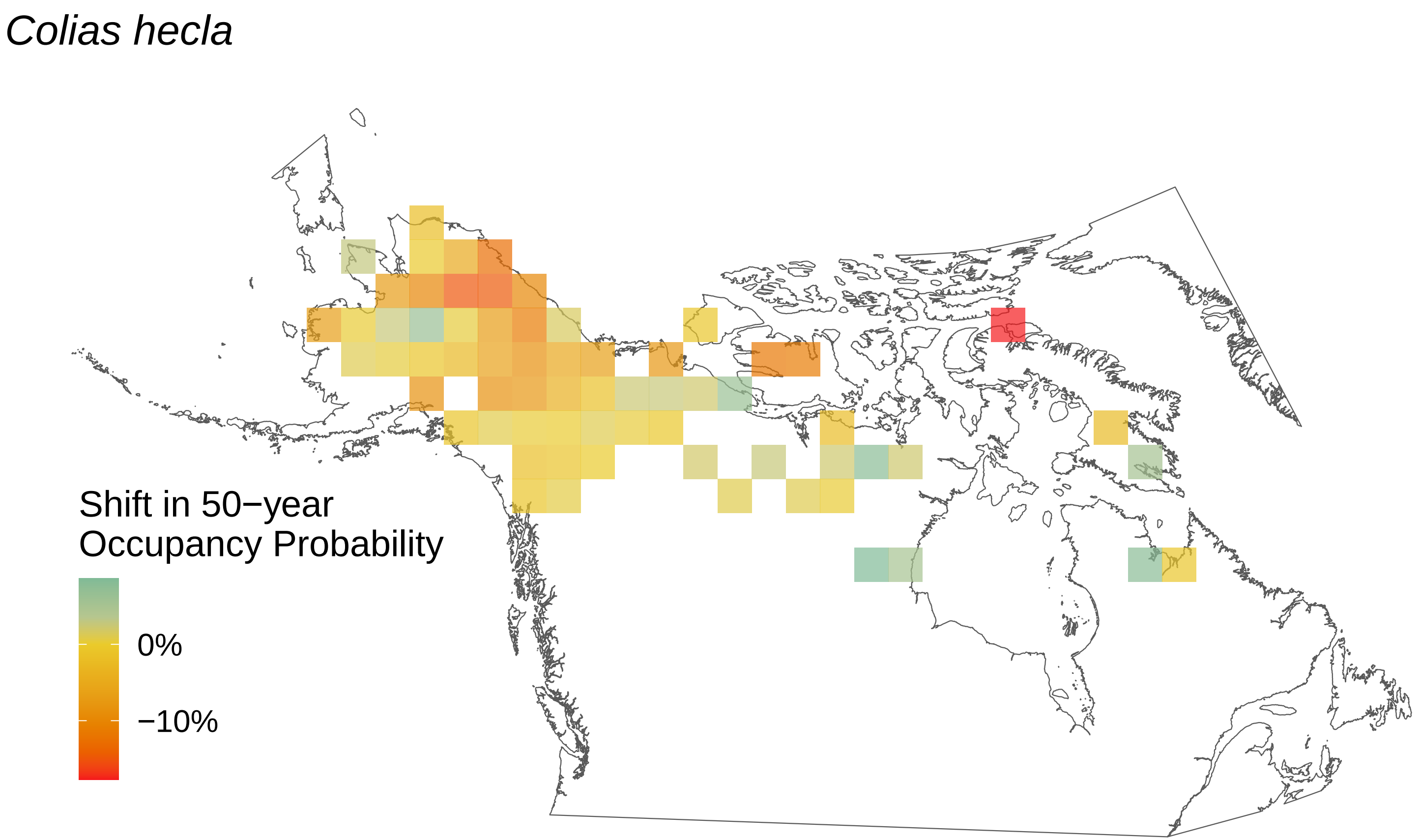

*Colias hecla*
Shift in 50−year Occupancy Probability
0%
−10%

*Colias eurytheme*

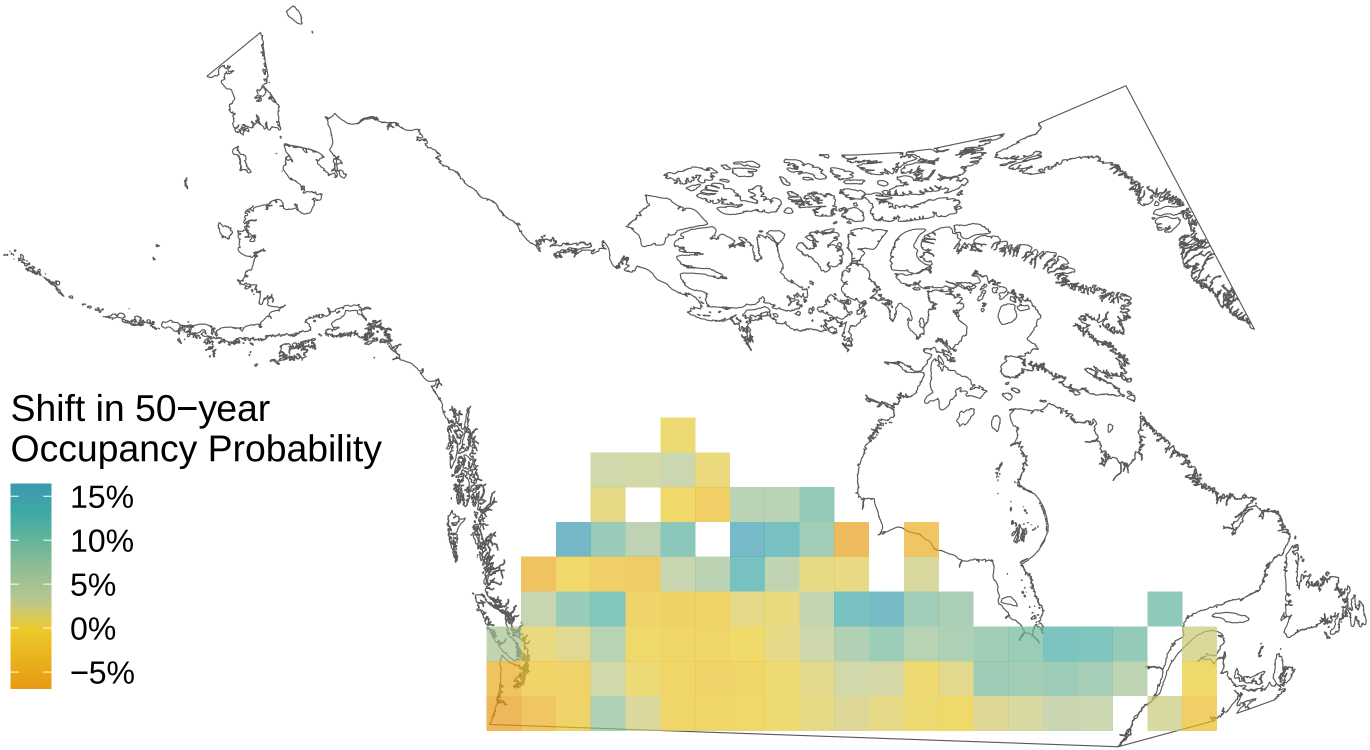

*Coenonympha tullia*

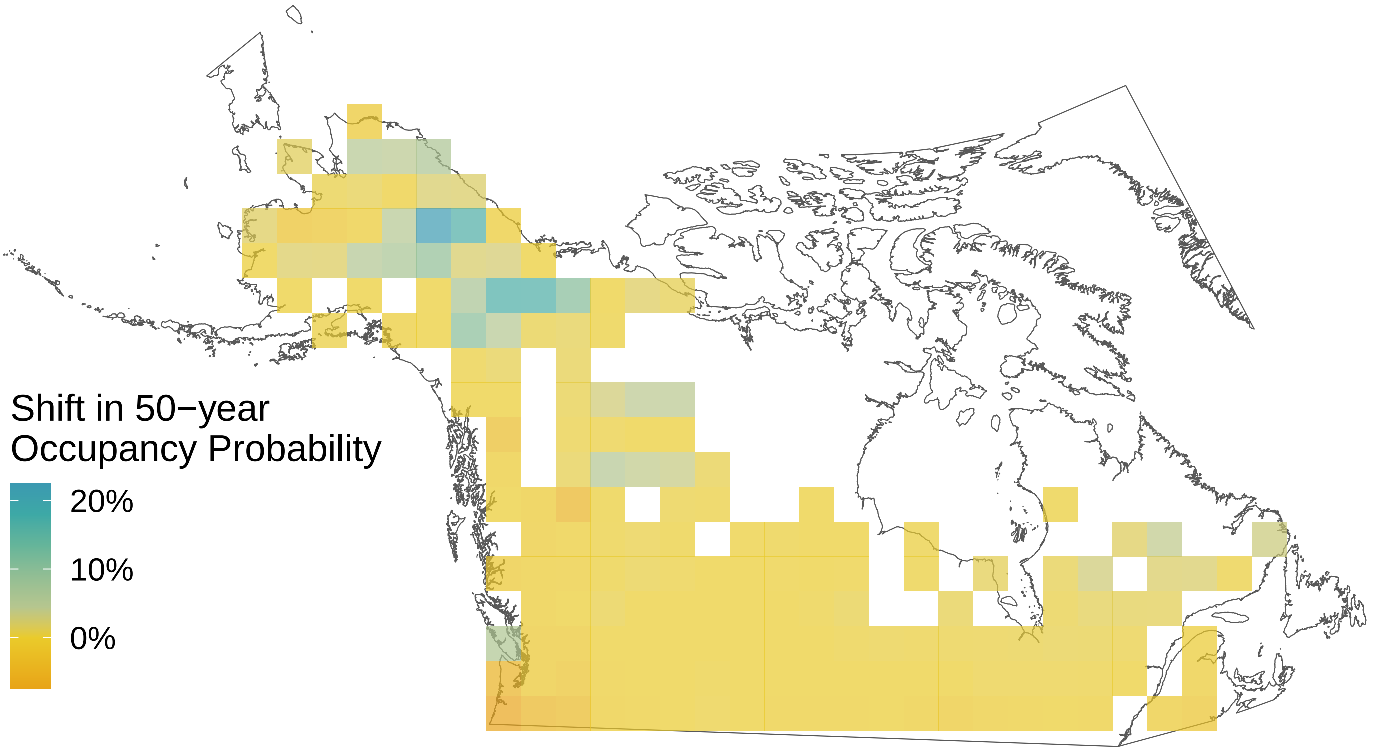

*Chlosyne harrisii*

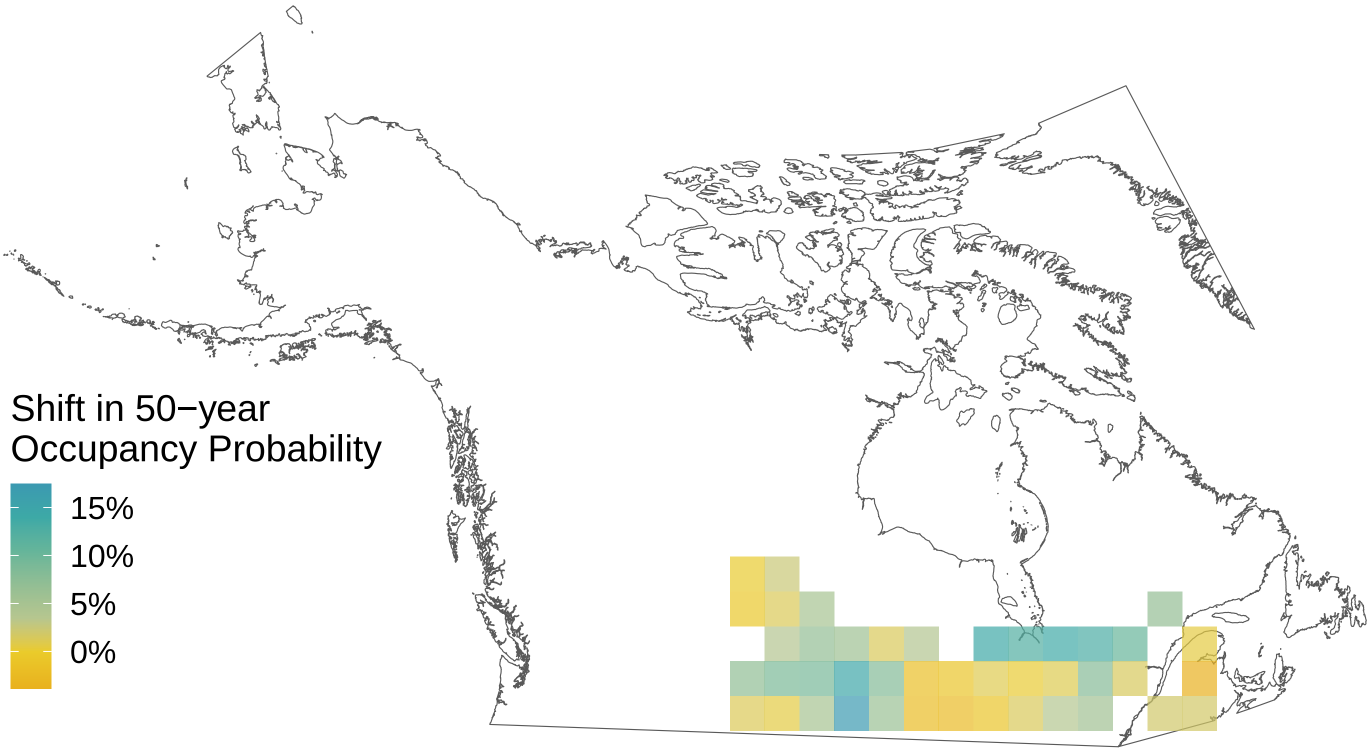

*Cercyonis pegala*

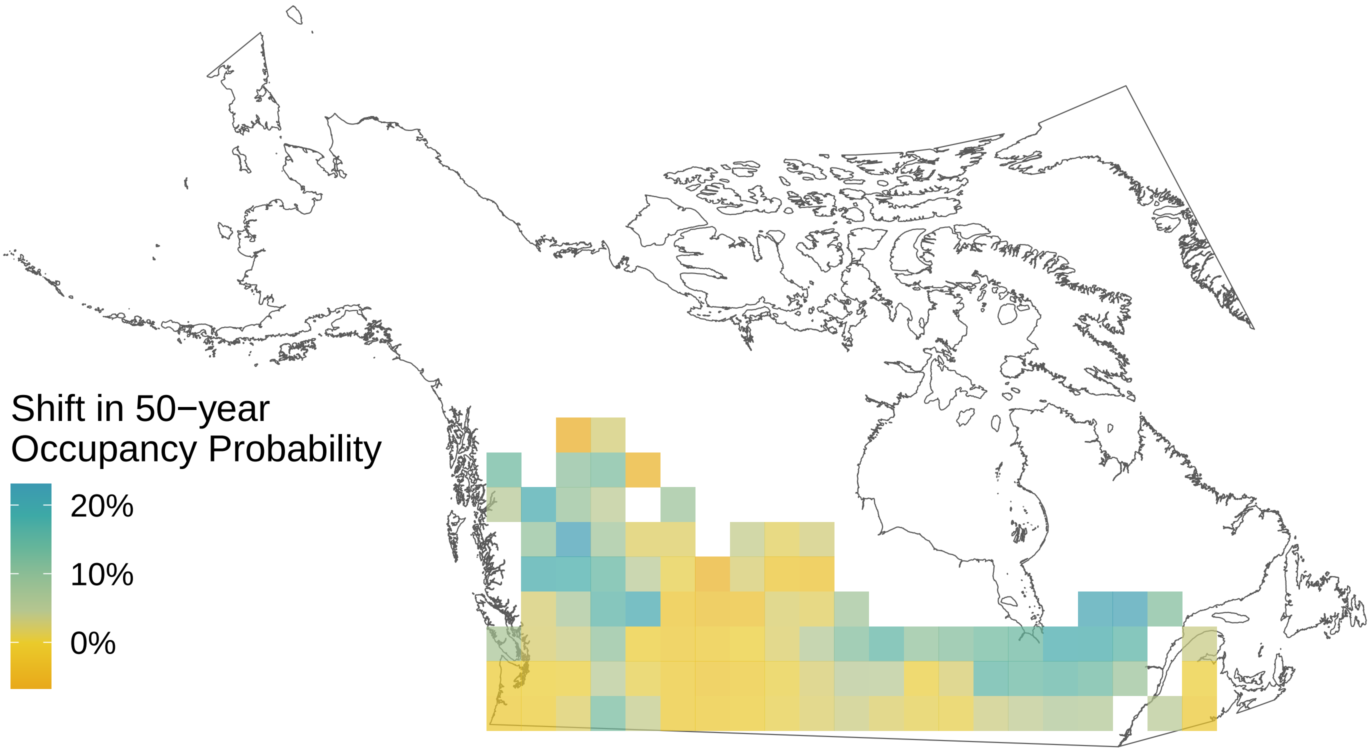

*Carterocephalus palaemon*

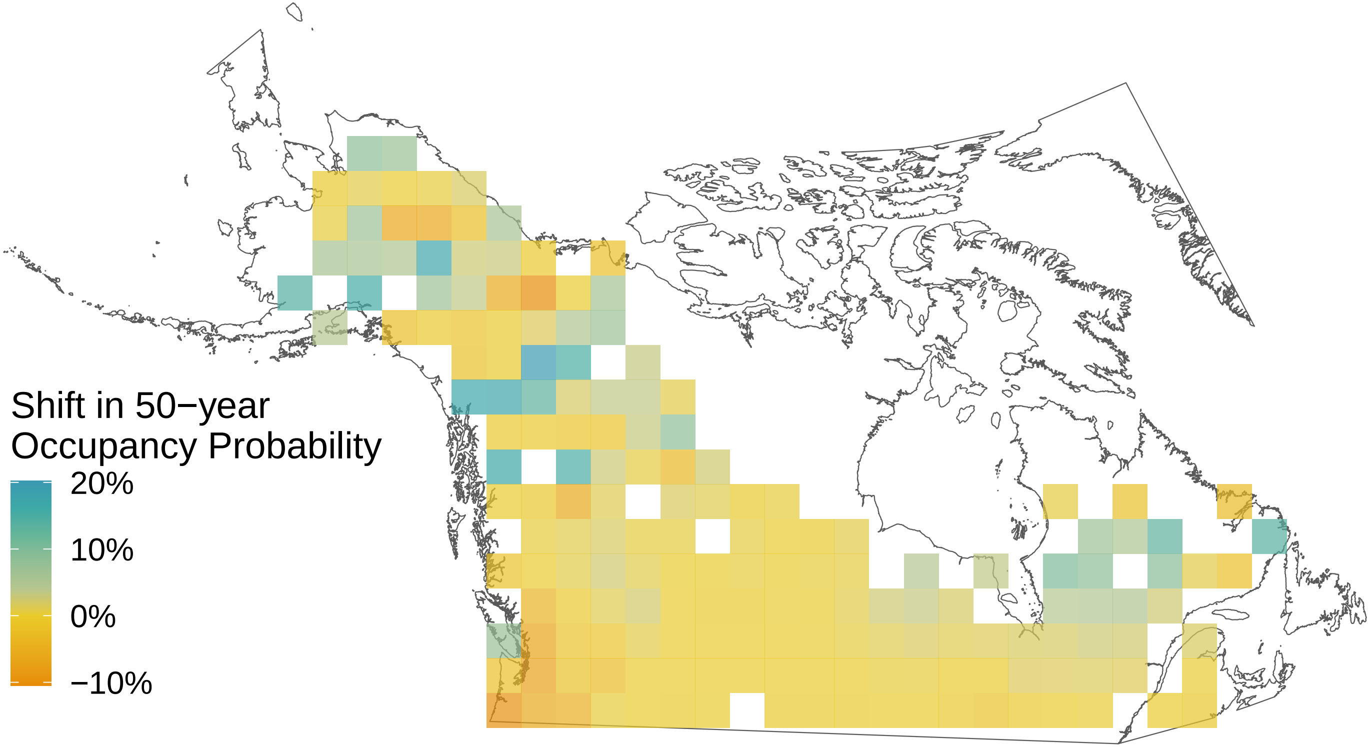

*Callophrys polios*

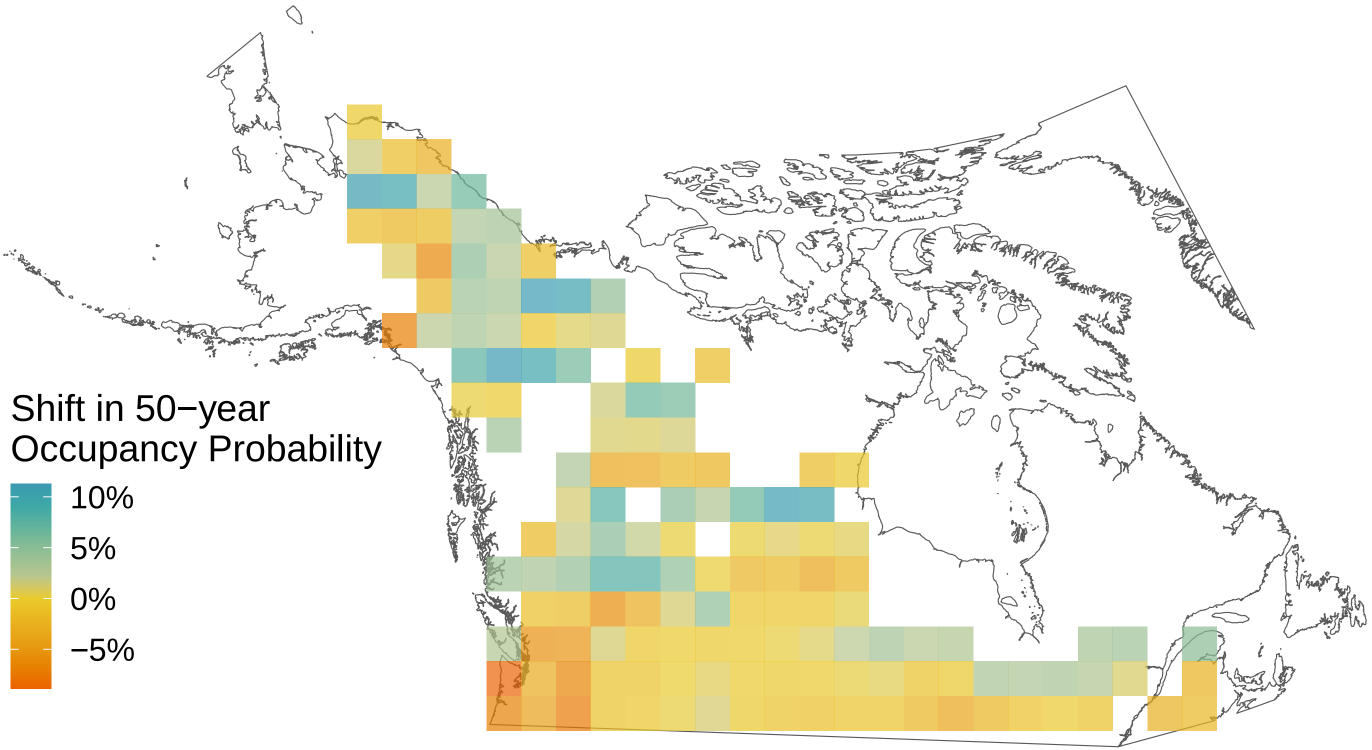

*Callophrys niphon*

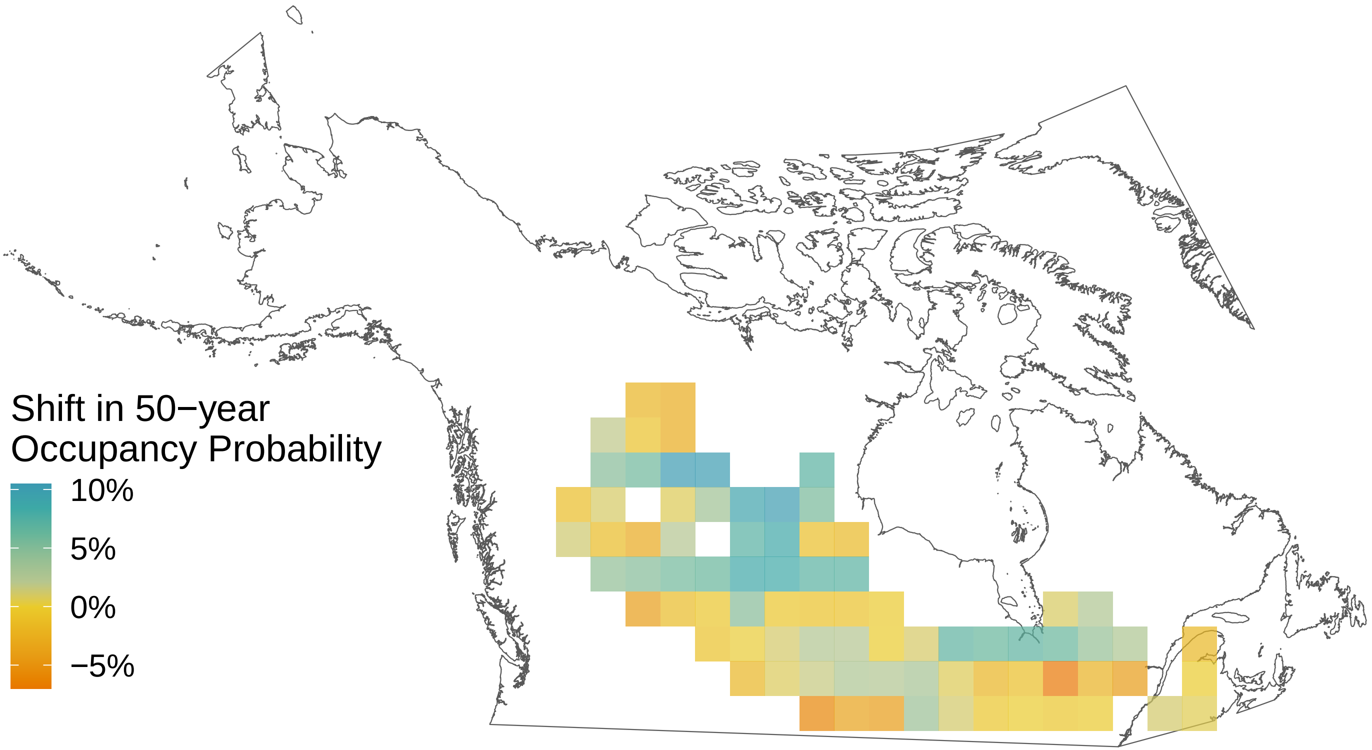

*Callophrys eryphon*

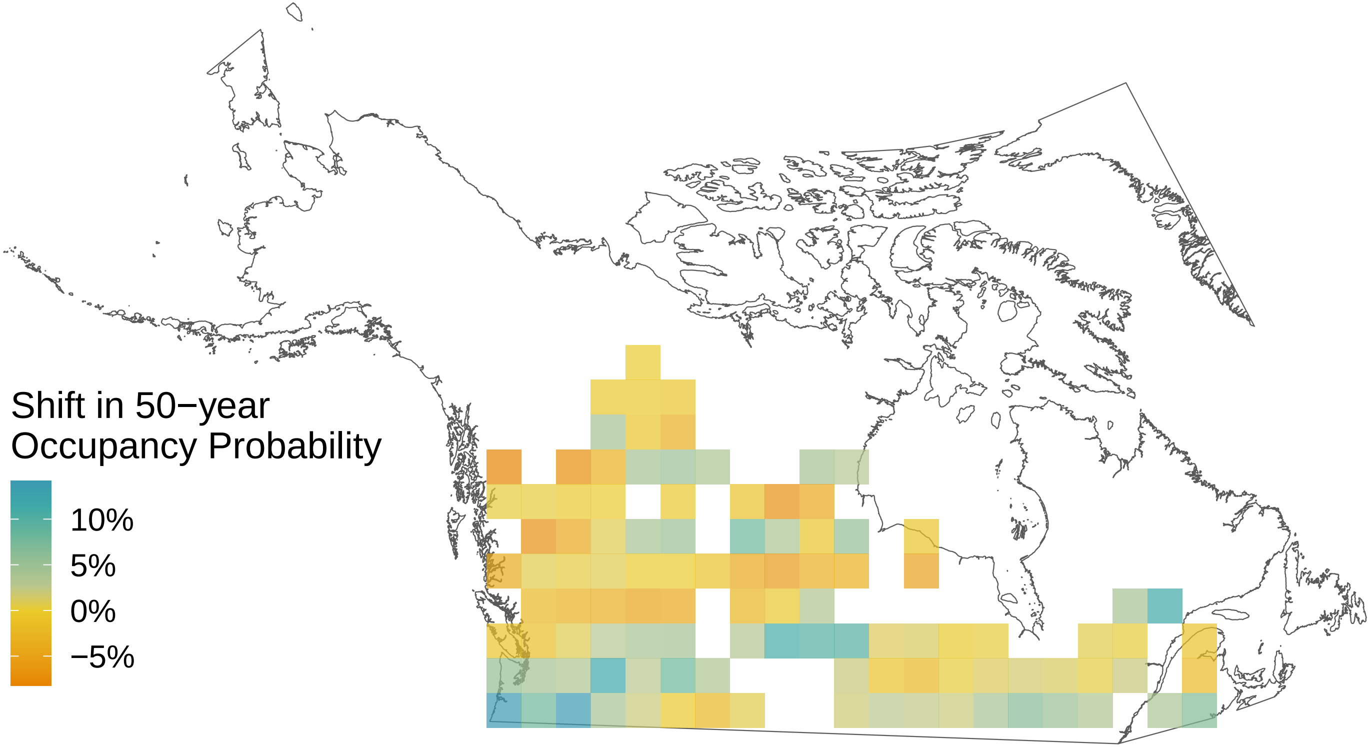

*Callophrys augustinus*

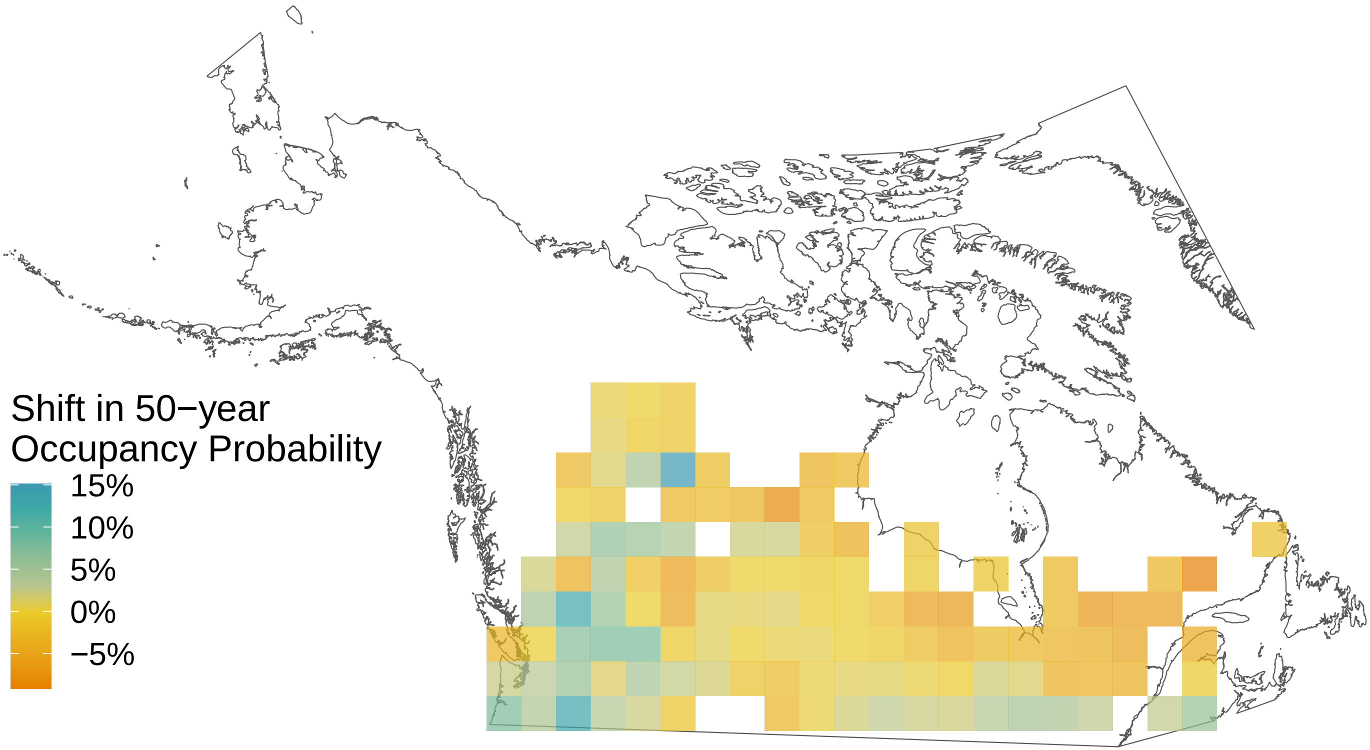

*Boloria selene*

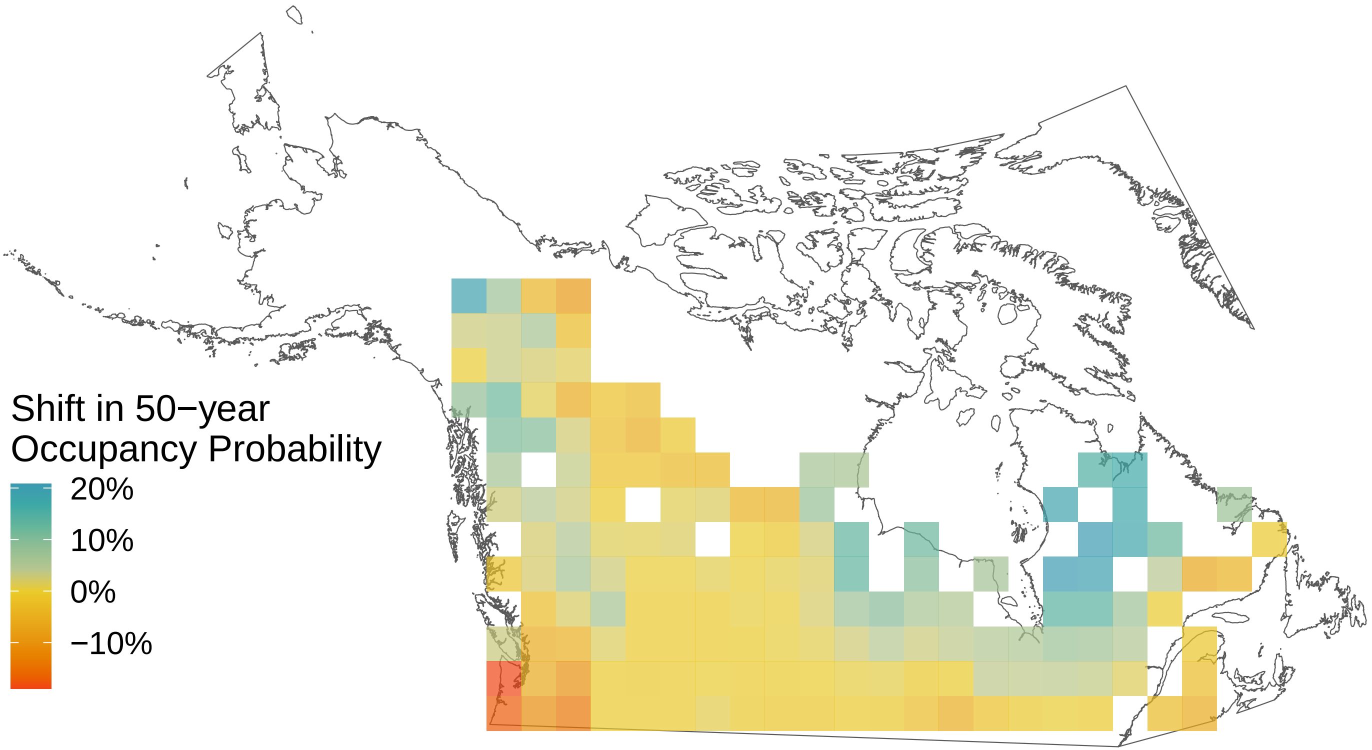

*Boloria polaris*

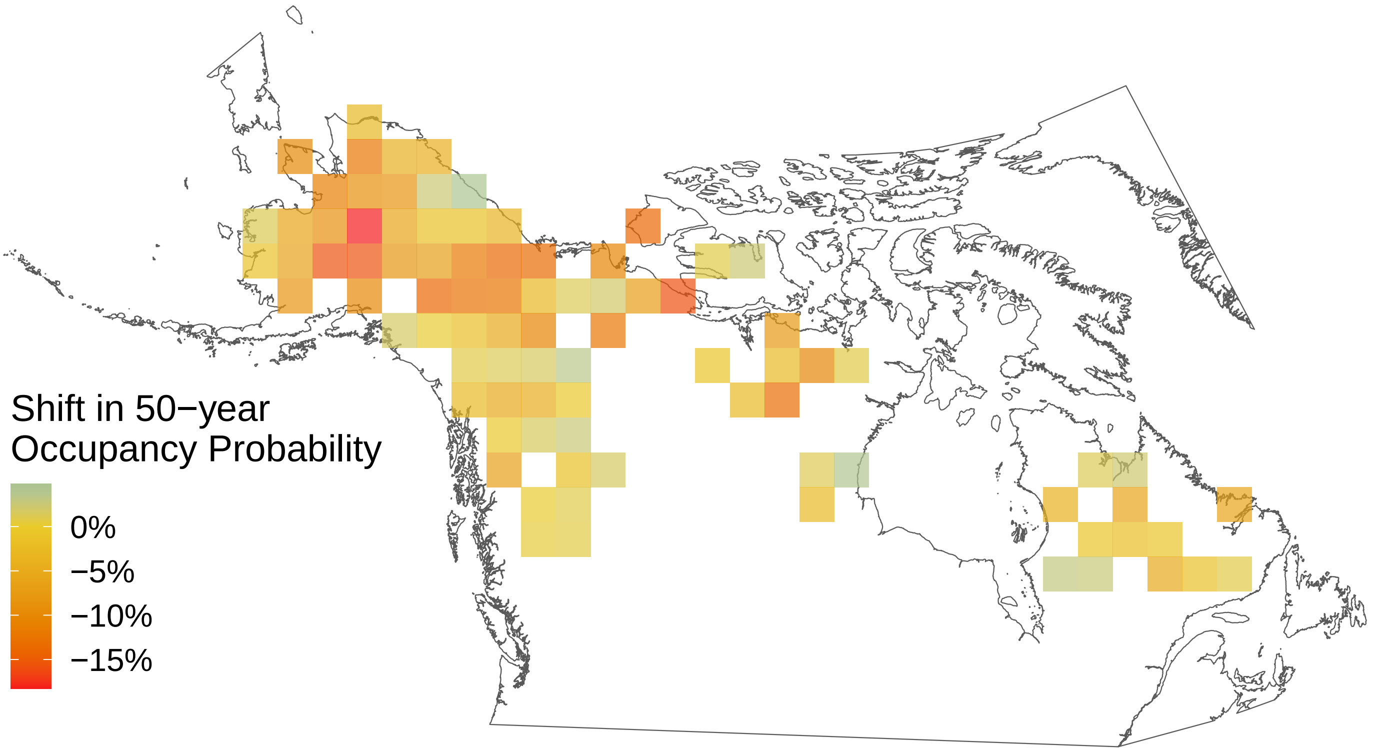

*Boloria improba*

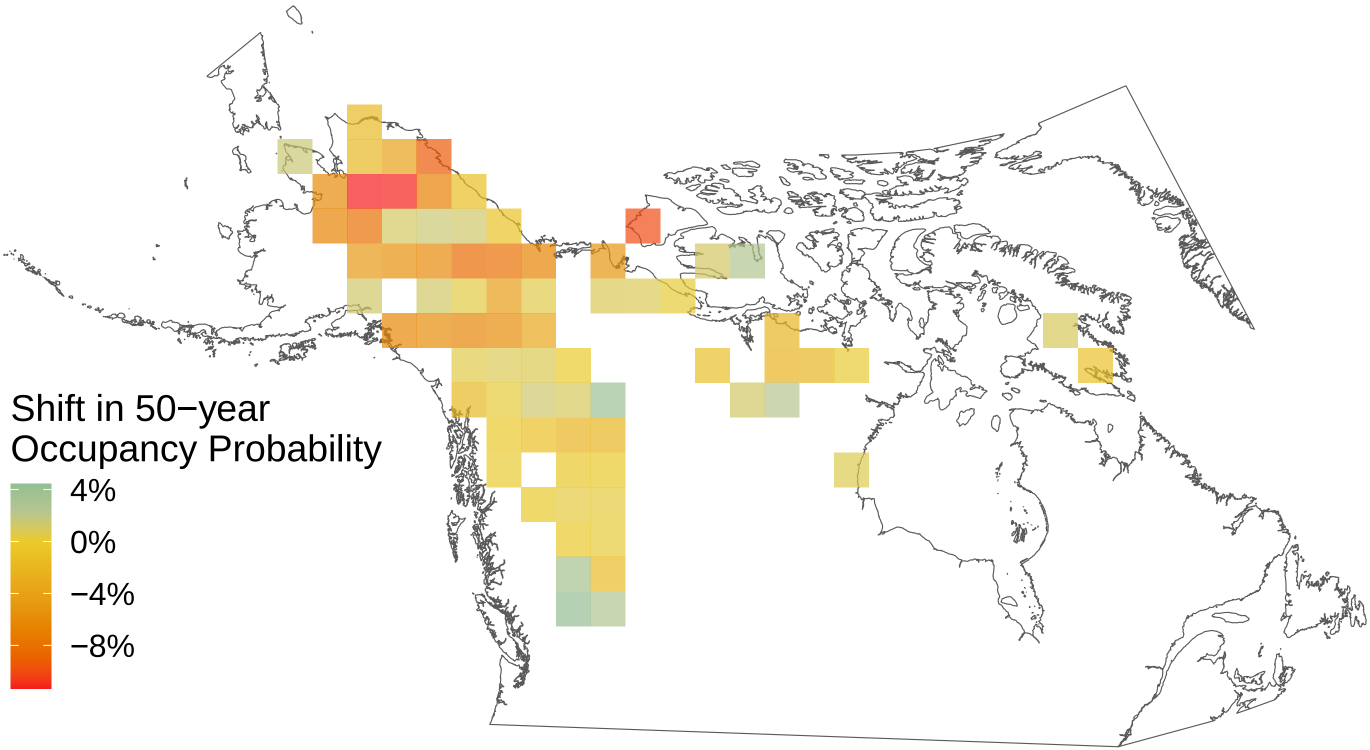

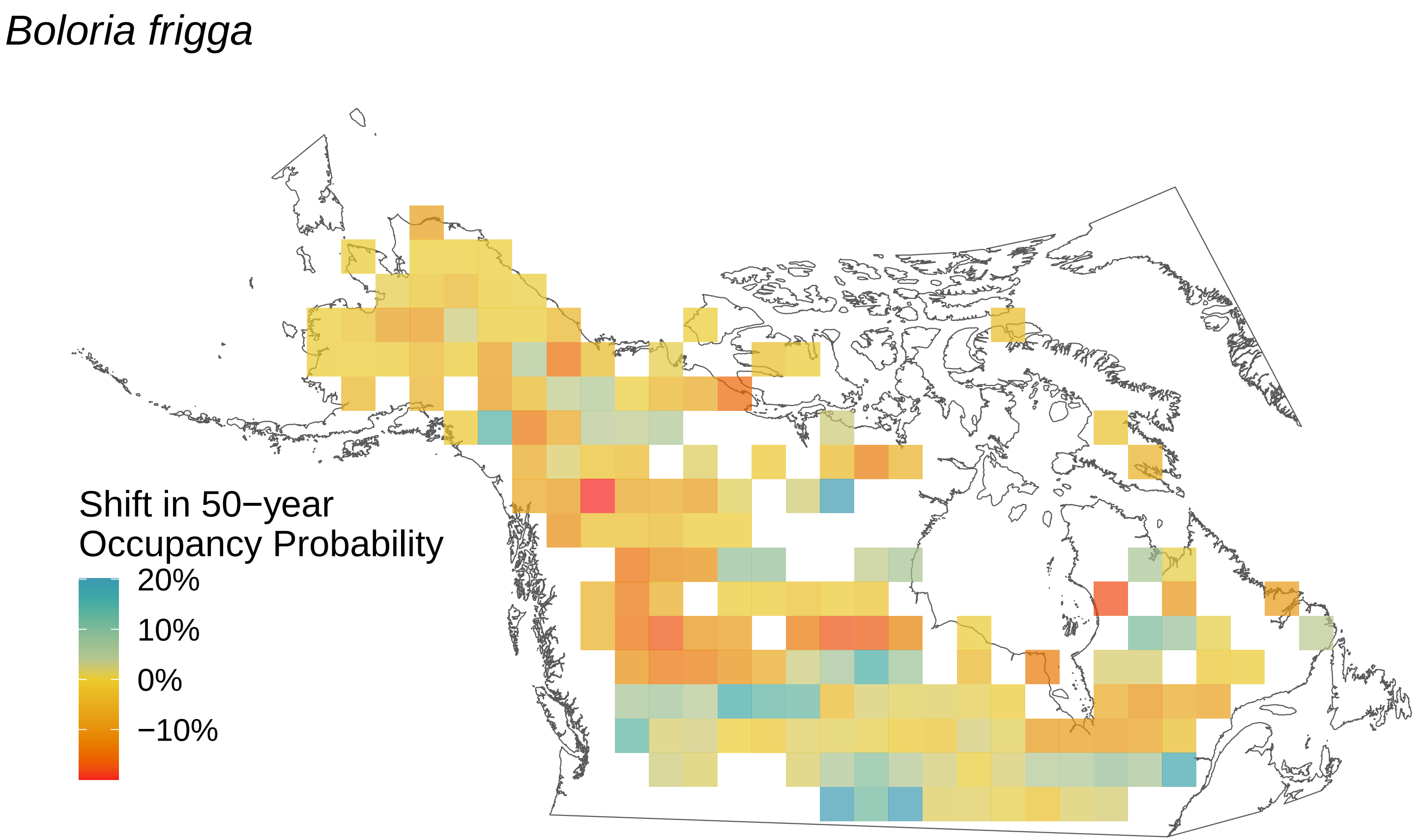
Boloria frigga
Shift in 50−year Occupancy Probability
20%
10%
0%
−10%

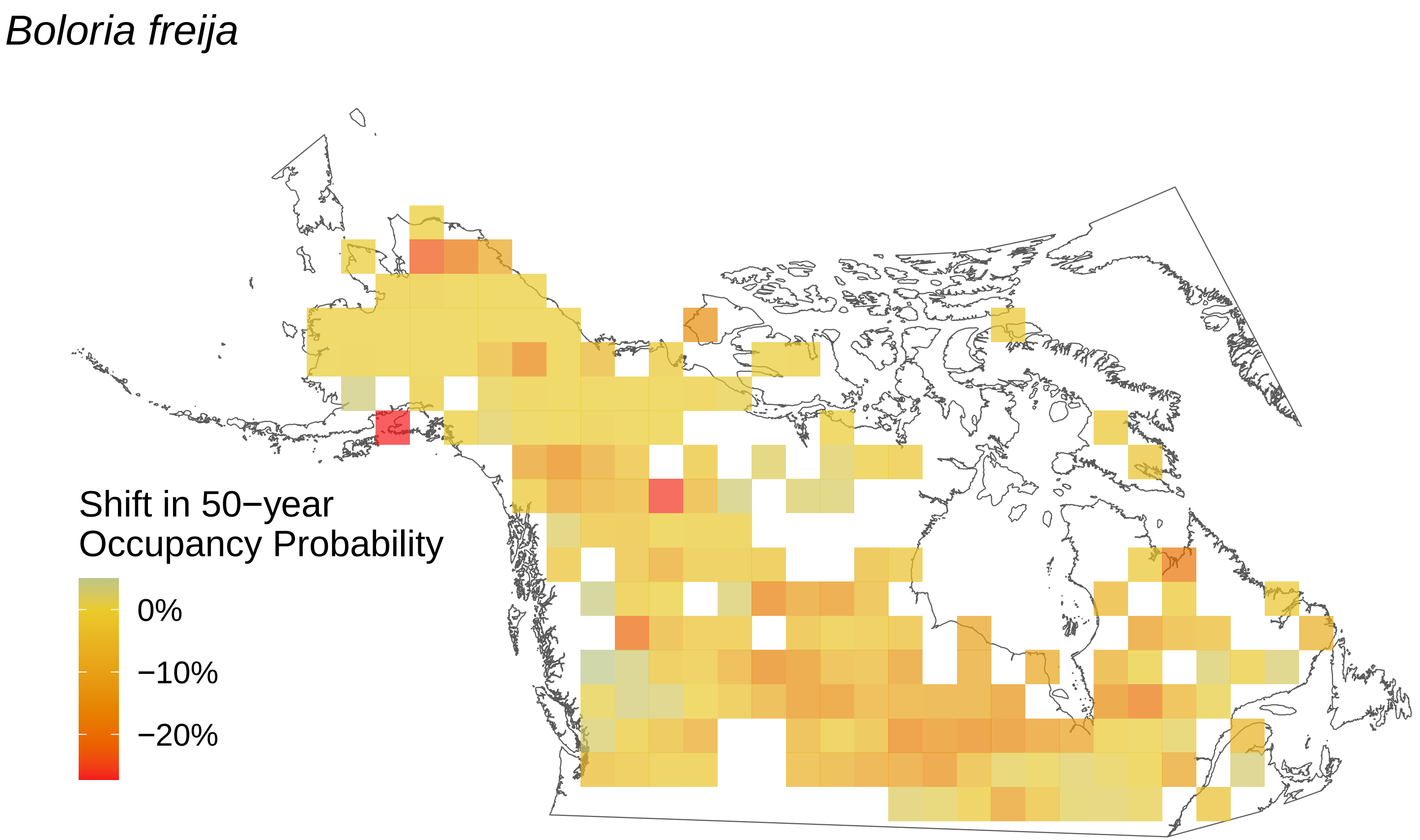

Boloria freija
Shift in 50−year
Occupancy Probability
0%
−10%
−20%

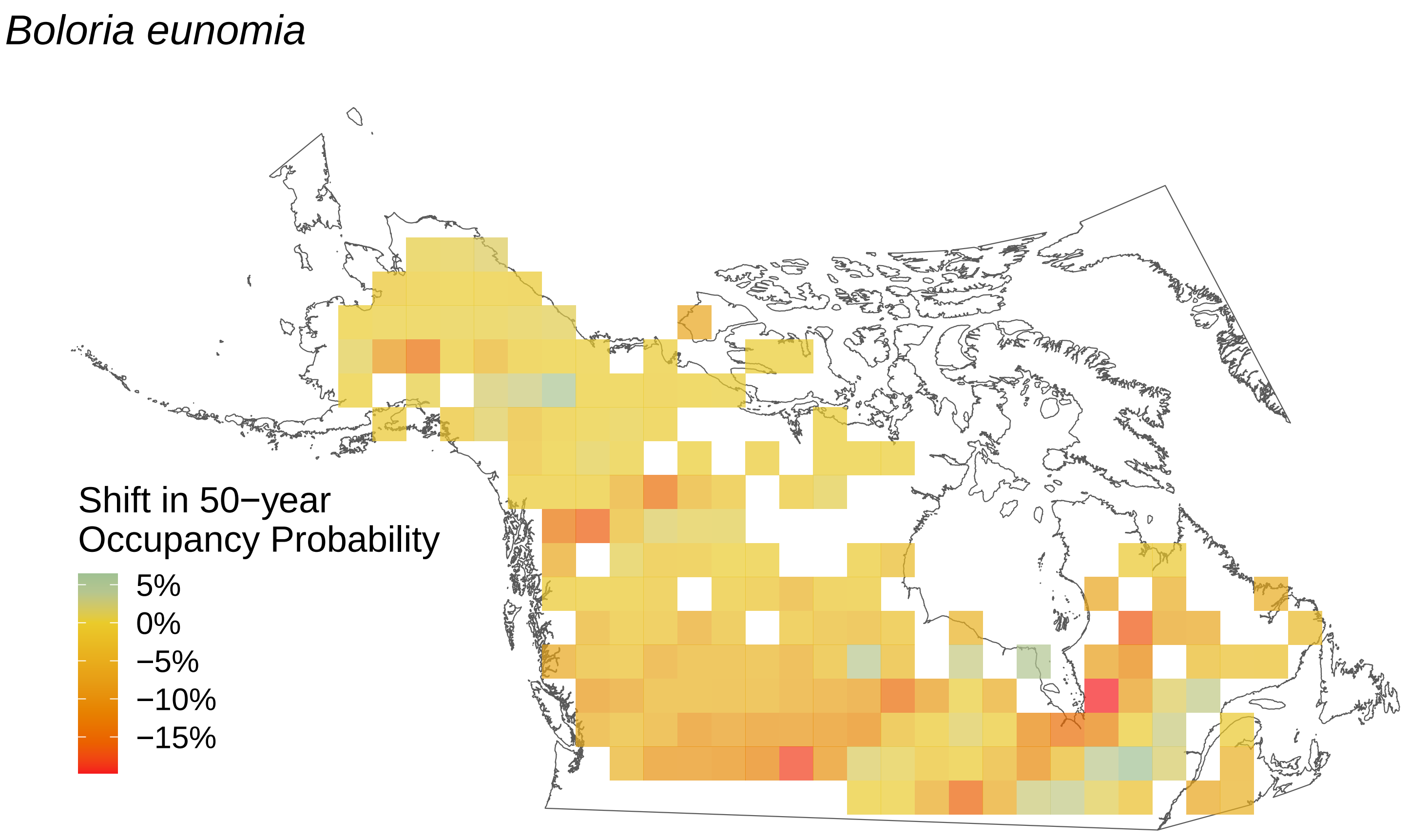
Boloria eunomia
Shift in 50−year
Occupancy Probability
5%
0%
−5%
−10%
−15%

*Boloria chariclea*

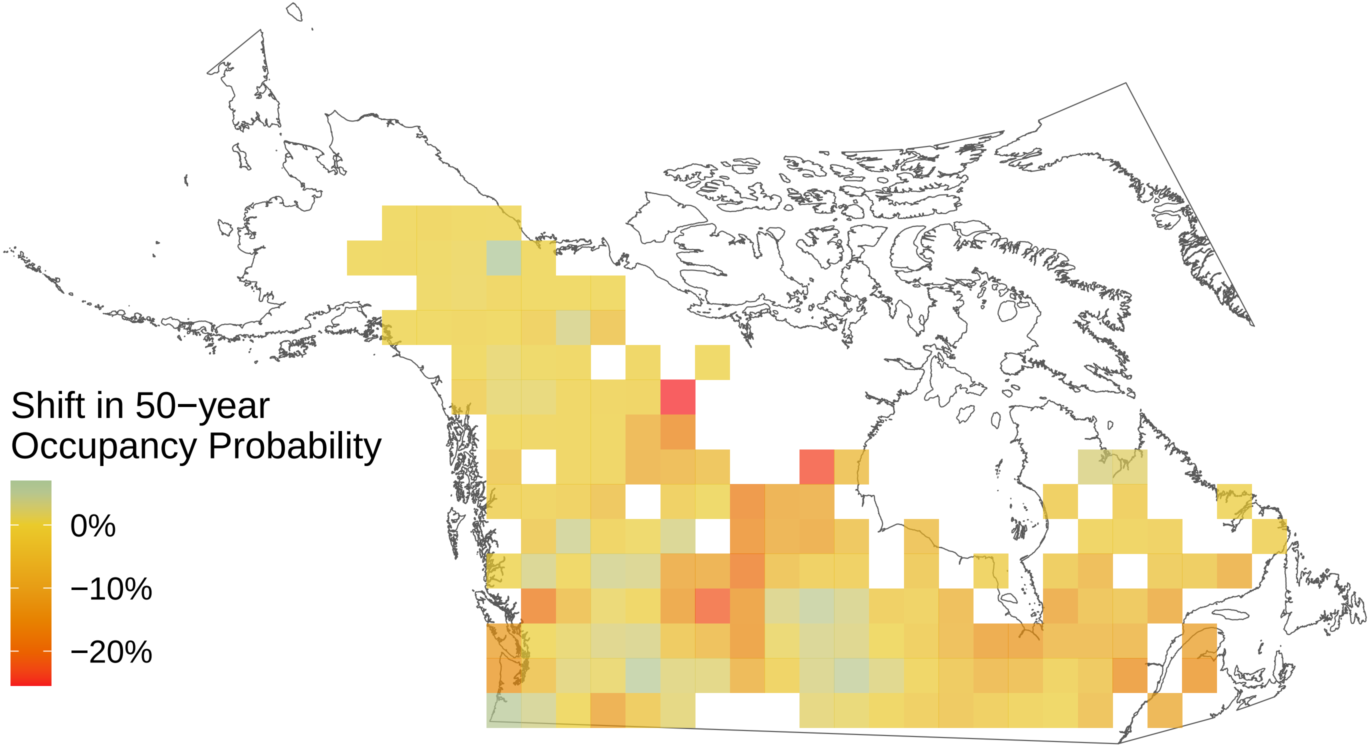

*Boloria bellona*

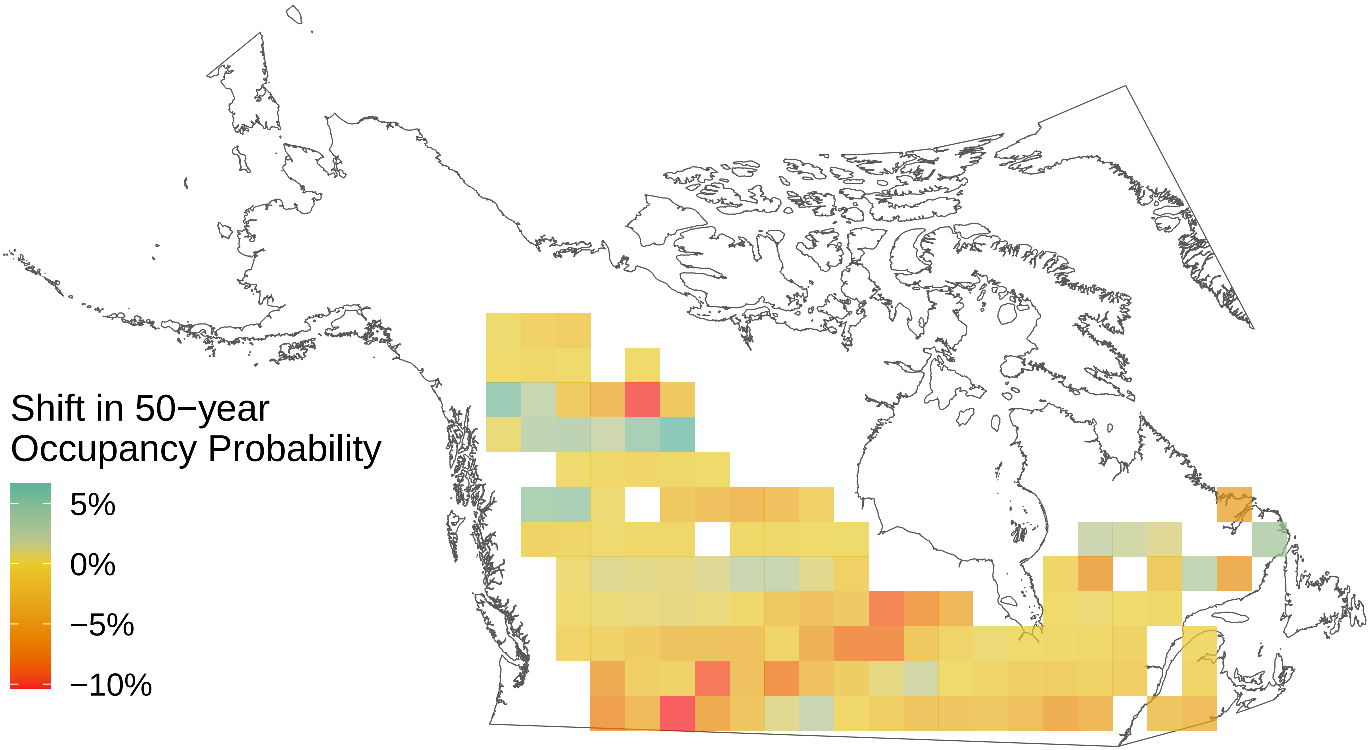

*Argynnis mormonia*

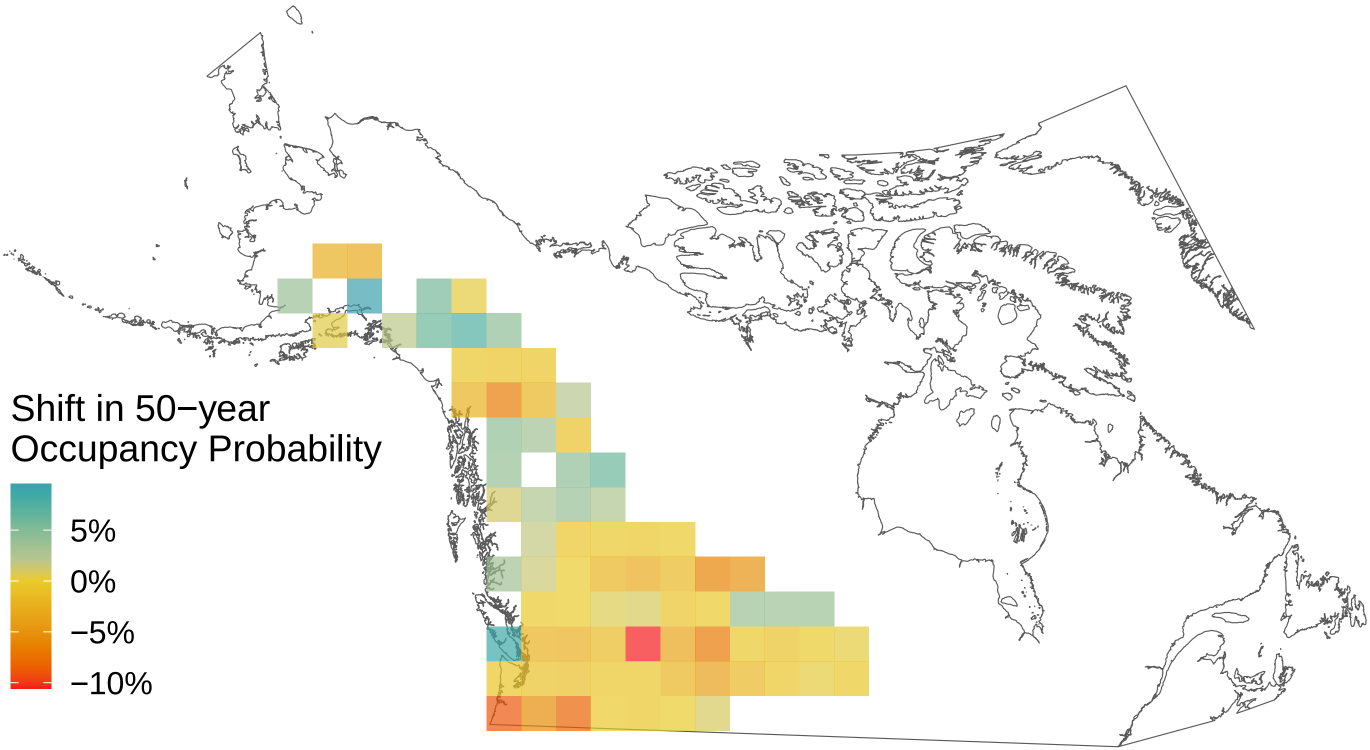

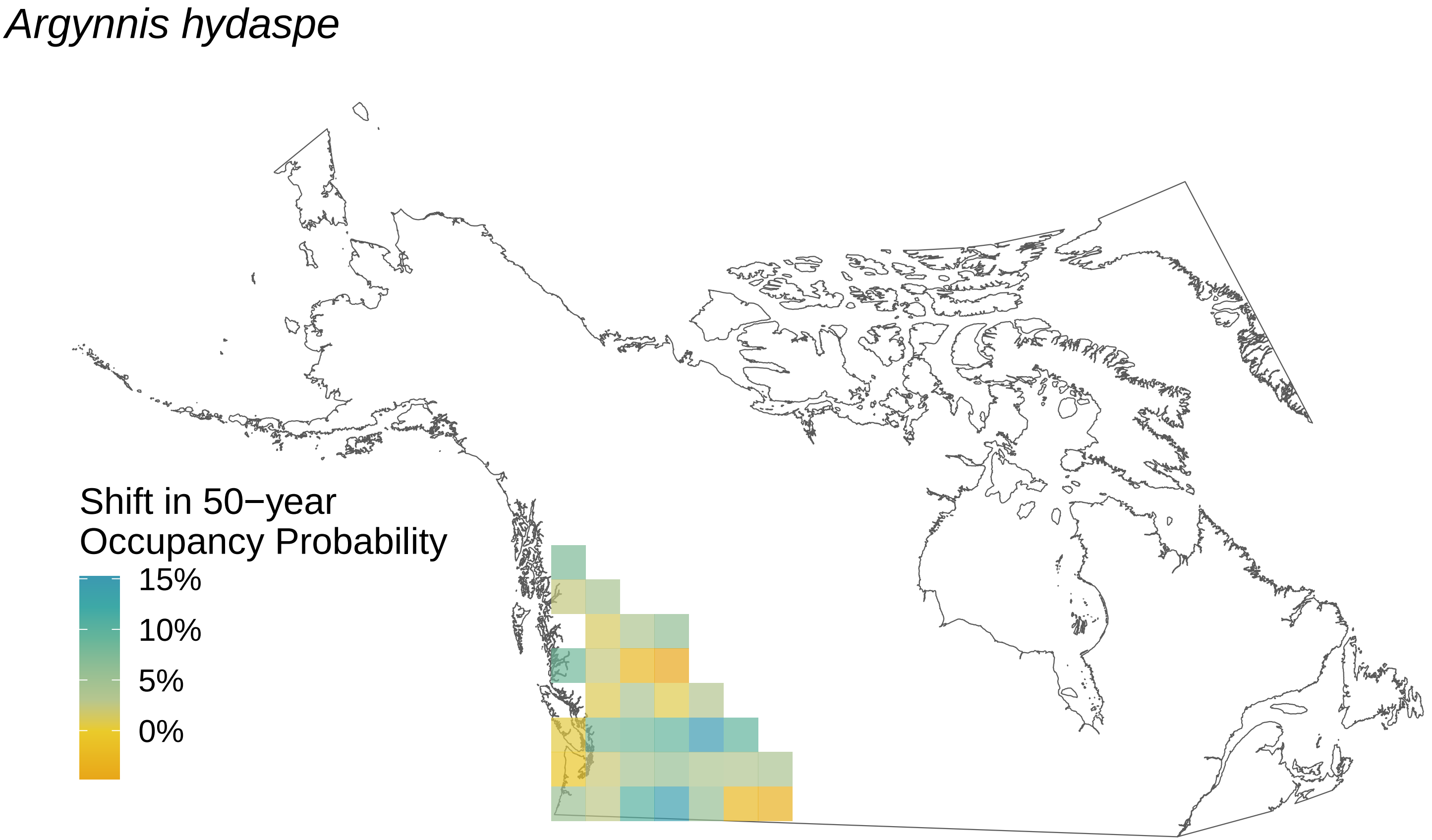
Argynnis hydaspe
Shift in 50−year
Occupancy Probability
15%
10%
5%
0%

*Argynnis cybele*

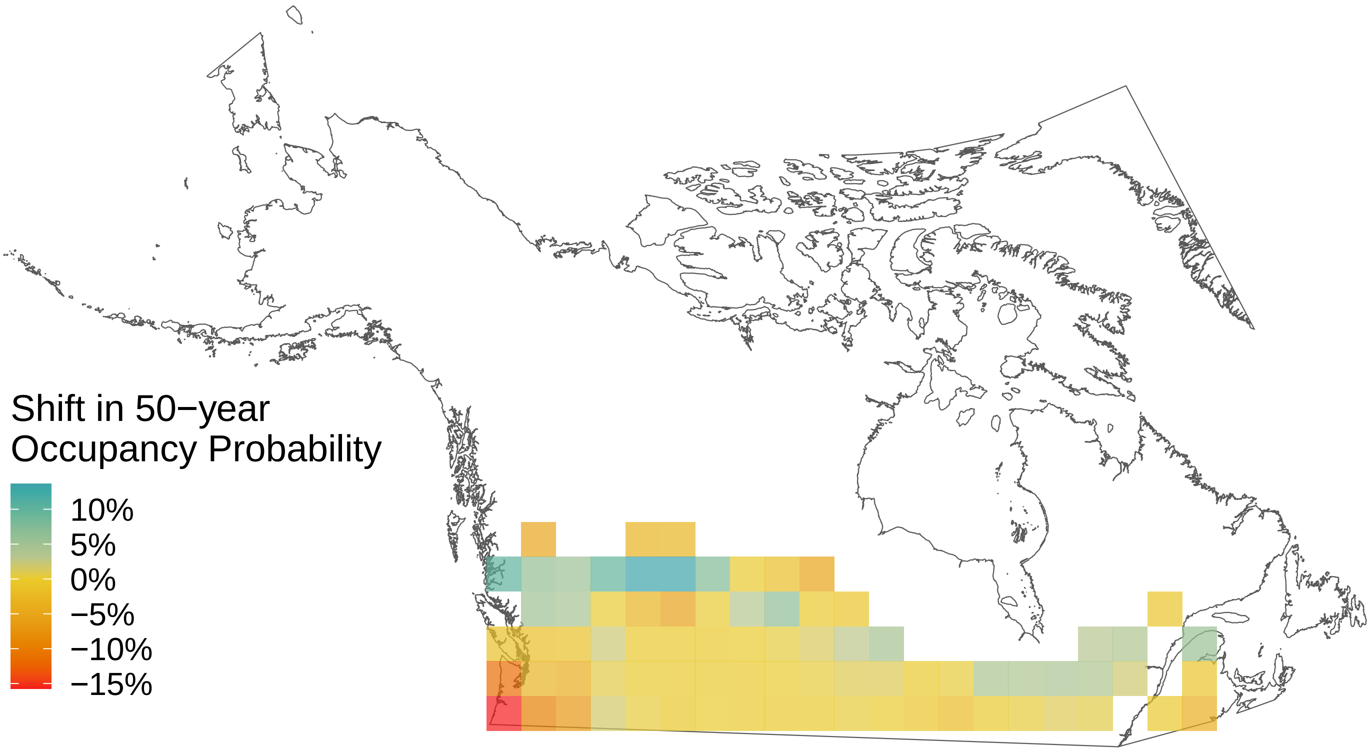

*Argynnis atlantis*

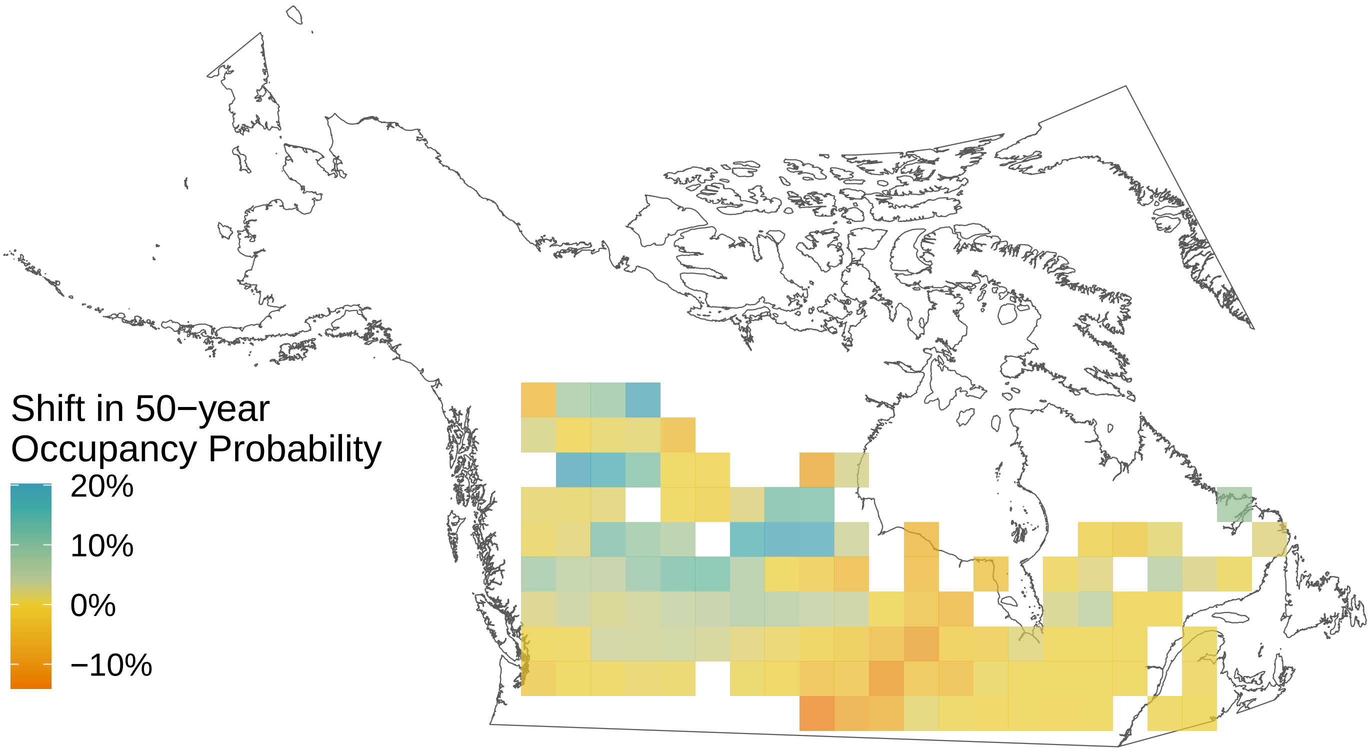

*Ancyloxypha numitor*

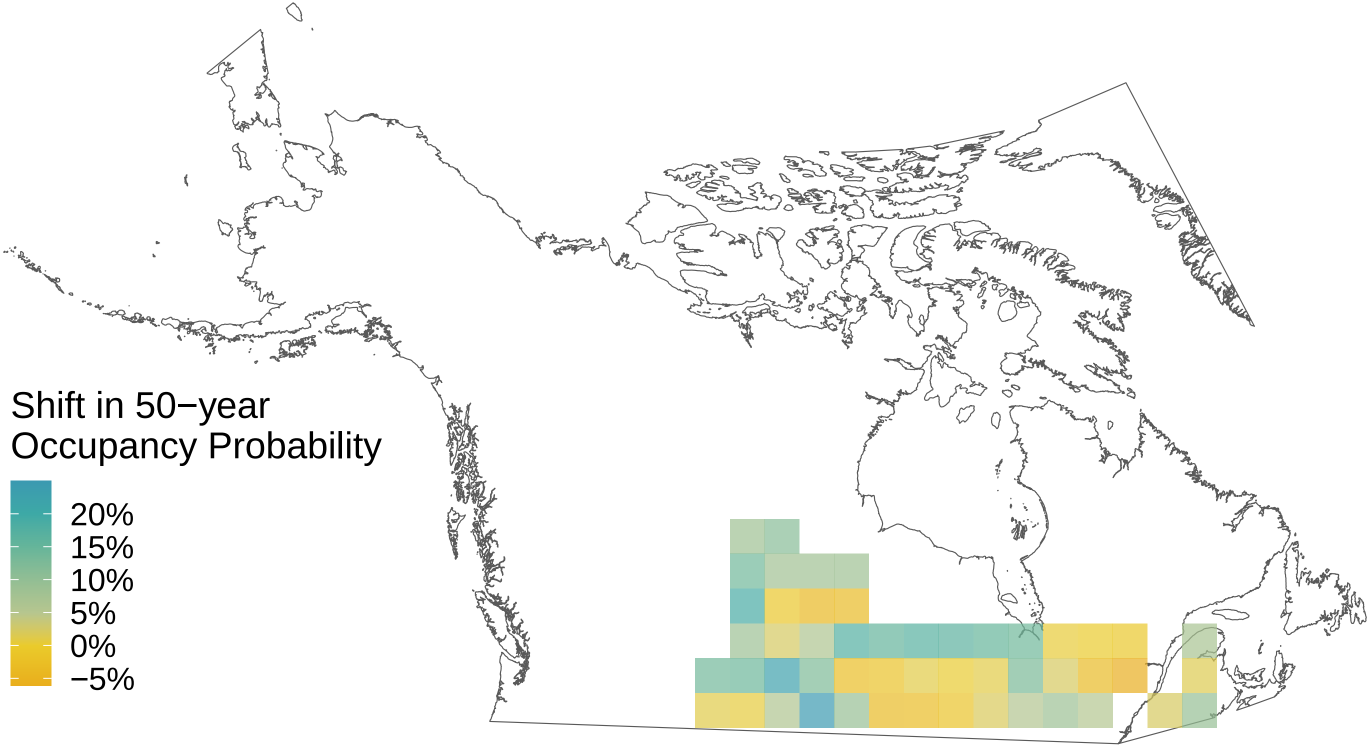

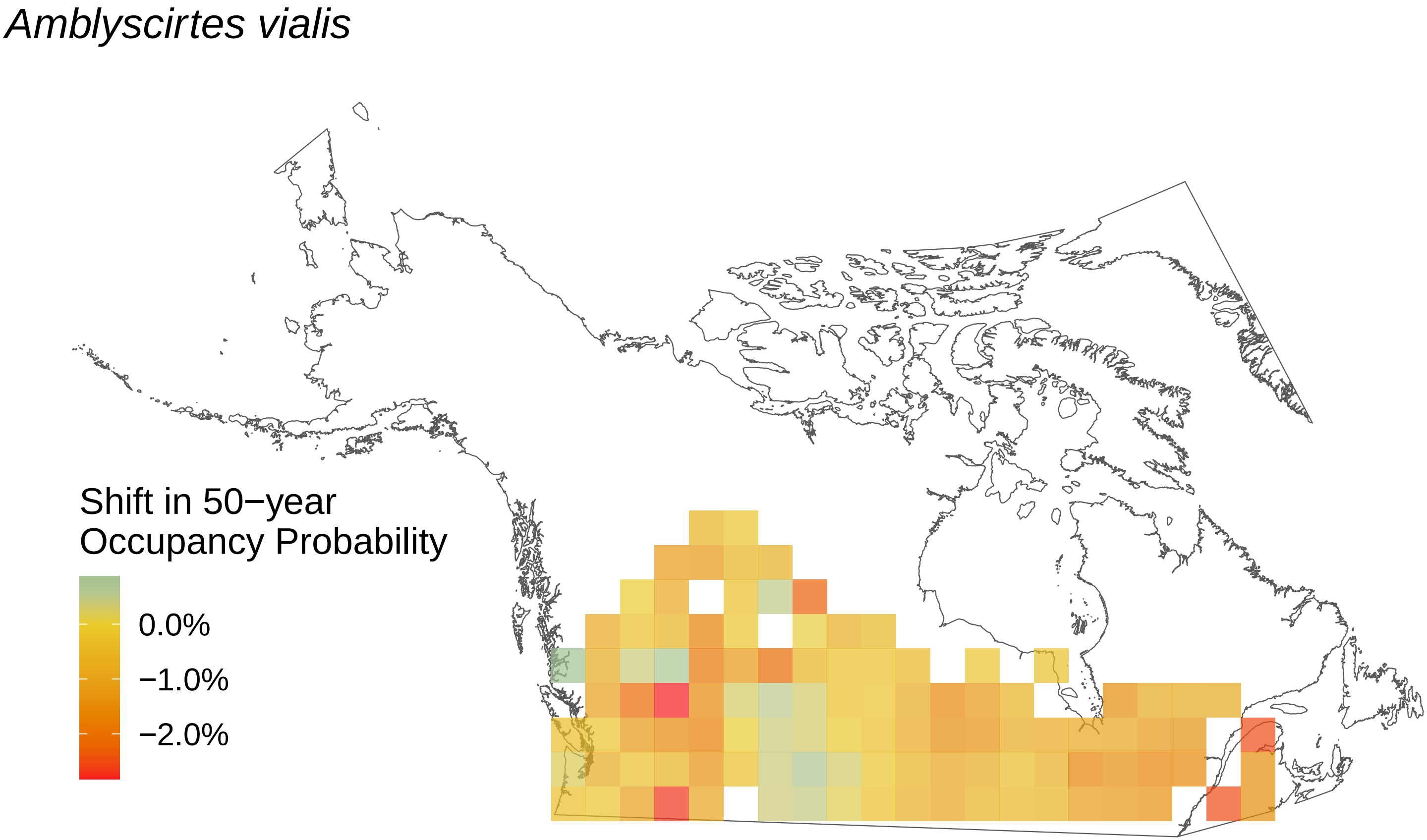
*Amblyscirtes vialis*
Shift in 50−year
Occupancy Probability
0.0%
−1.0%
−2.0%

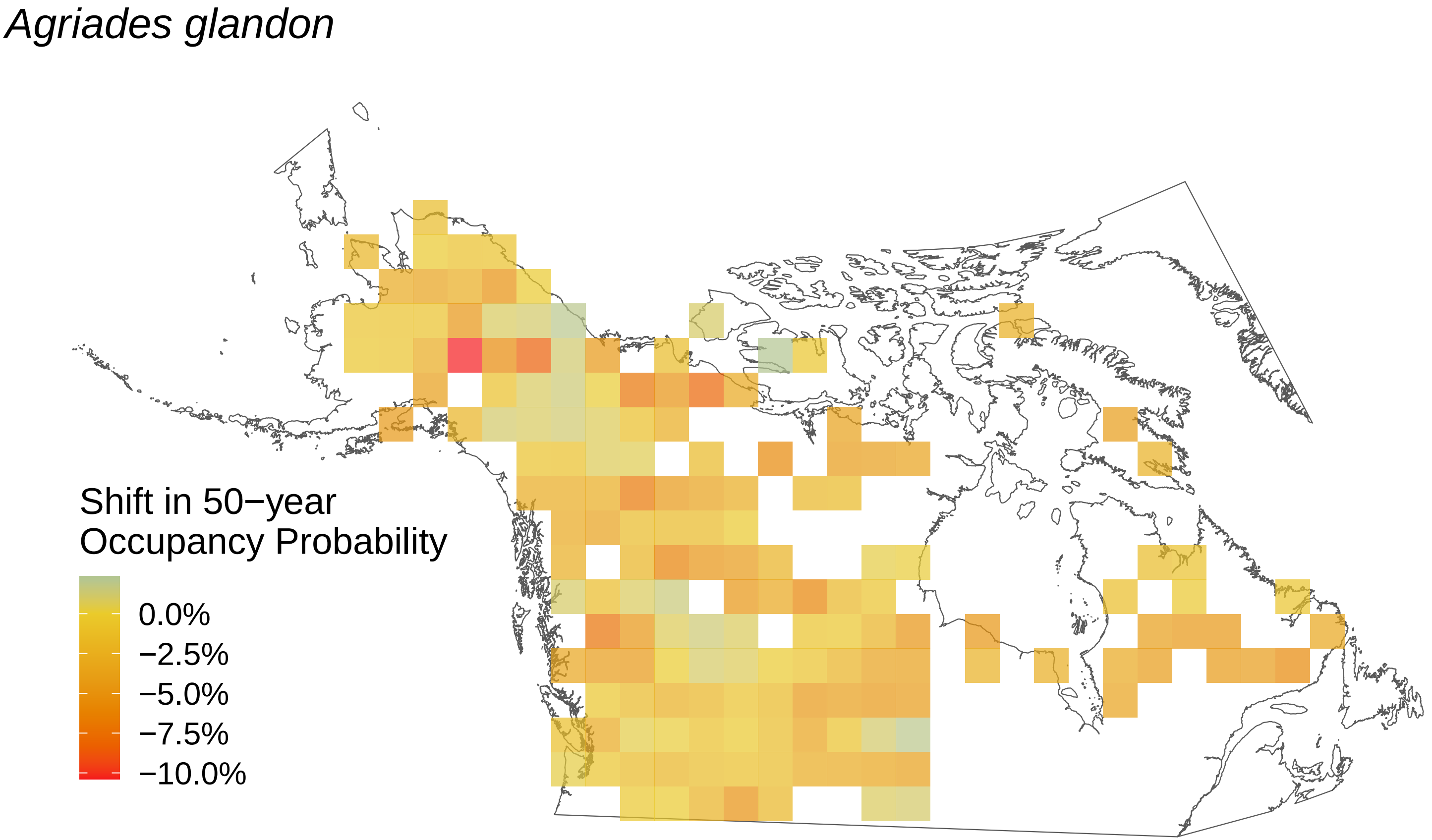
Agriades glandon
Shift in 50−year
Occupancy Probability
0.0%
−2.5%
−5.0%
−7.5%
−10.0%

*Aglais milberti*

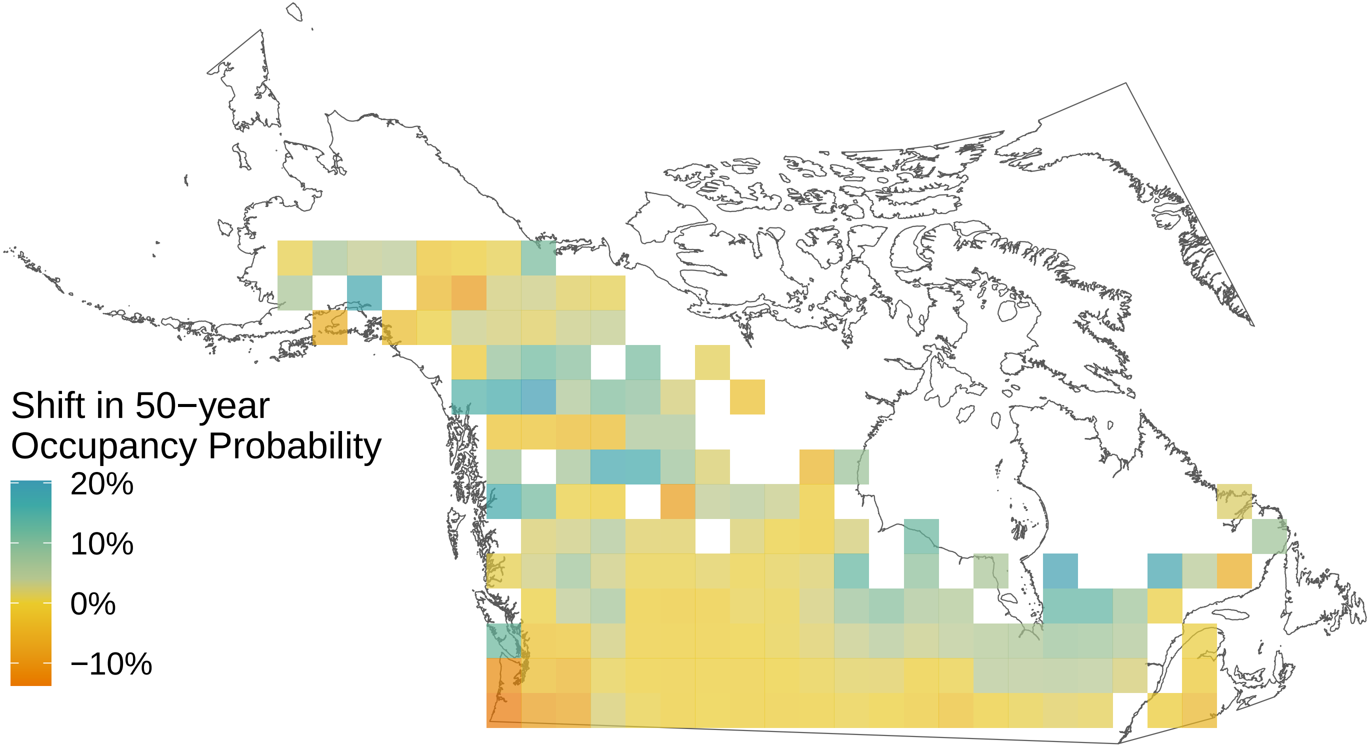
