## Supplemental File S9 for "Rising minimum temperatures contribute to 50 years of shifting Arctic and boreal butterfly communities in North America"

*Vanessa virginiensis*

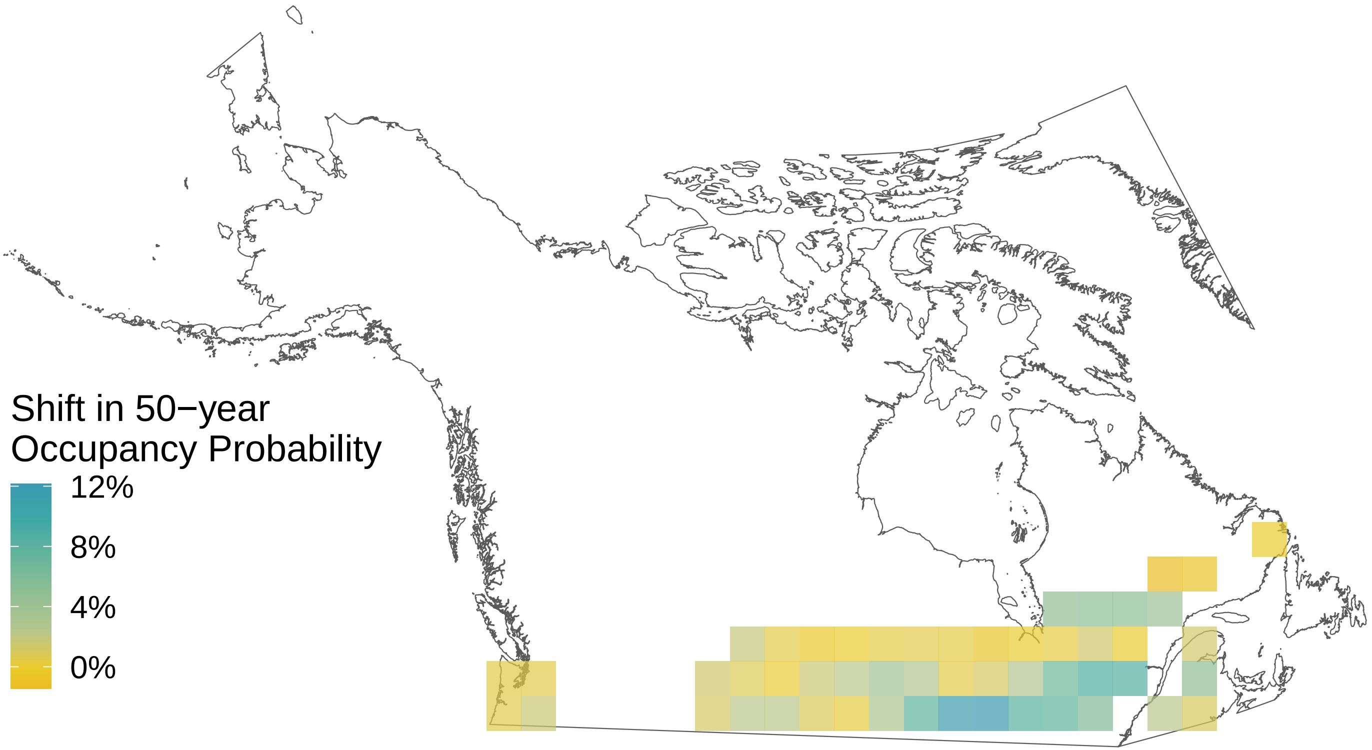

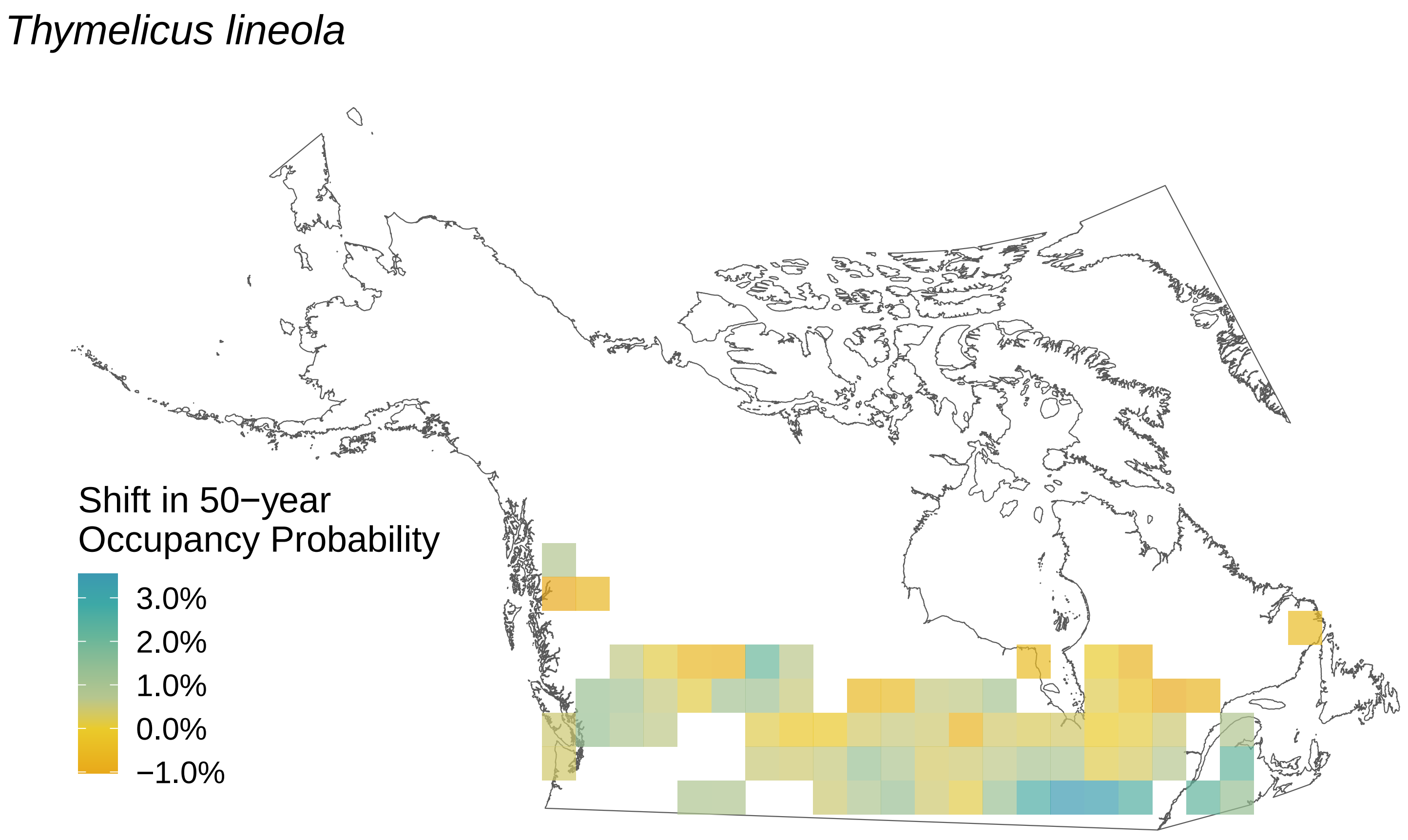
Thymelicus lineola
Shift in 50−year Occupancy Probability
3.0%
2.0%
1.0%
0.0%
−1.0%

*Thorybes pylades*

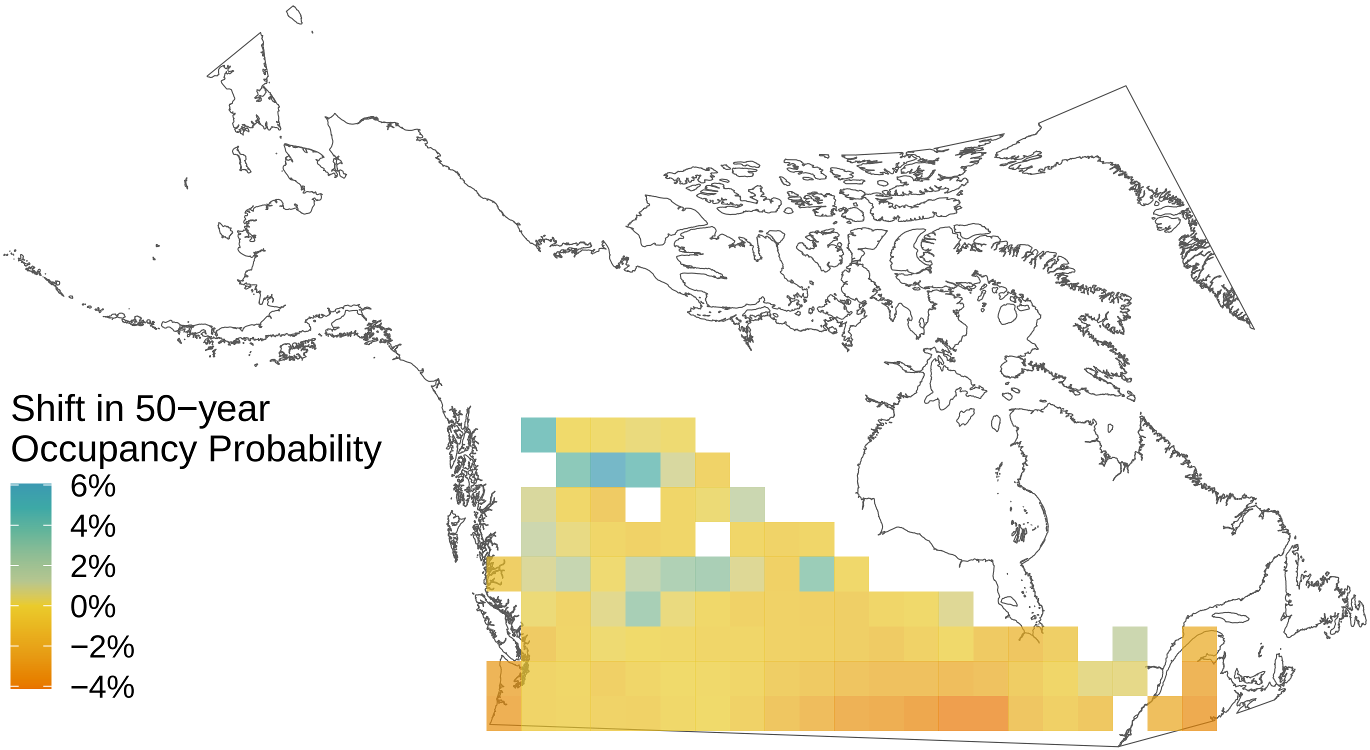

*Strymon melinus*

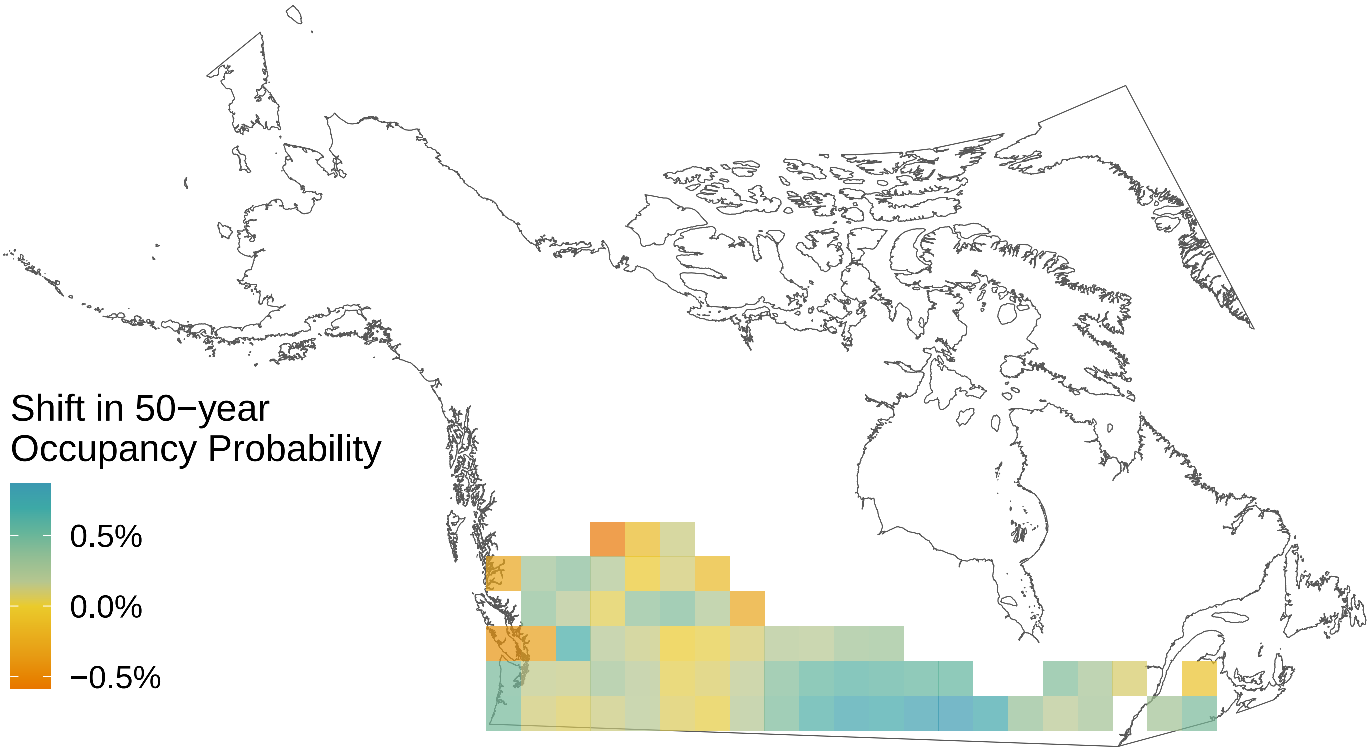

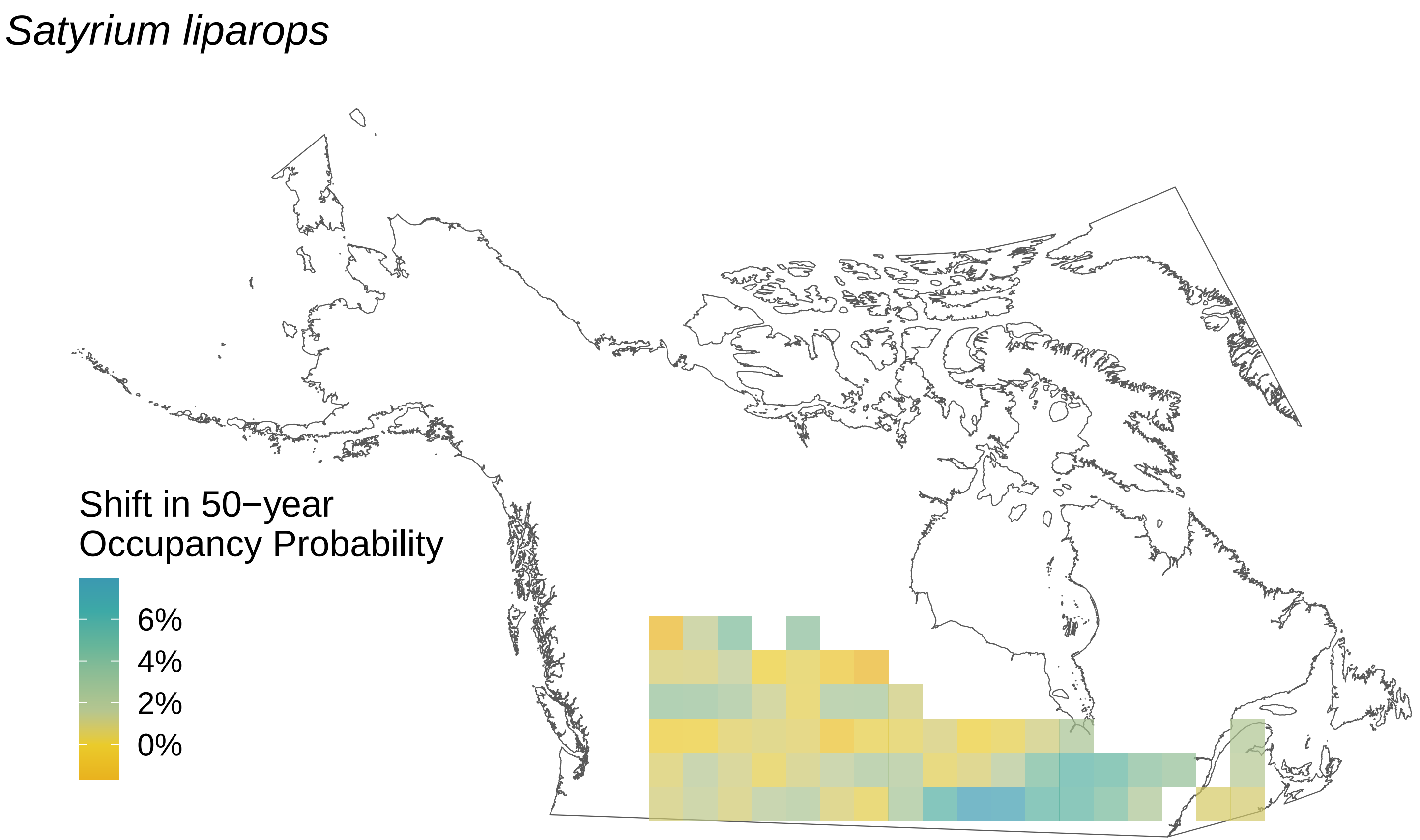

Satyrium liparops
Shift in 50−year
Occupancy Probability
6%
4%
2%
0%

*Pterourus rutulus*

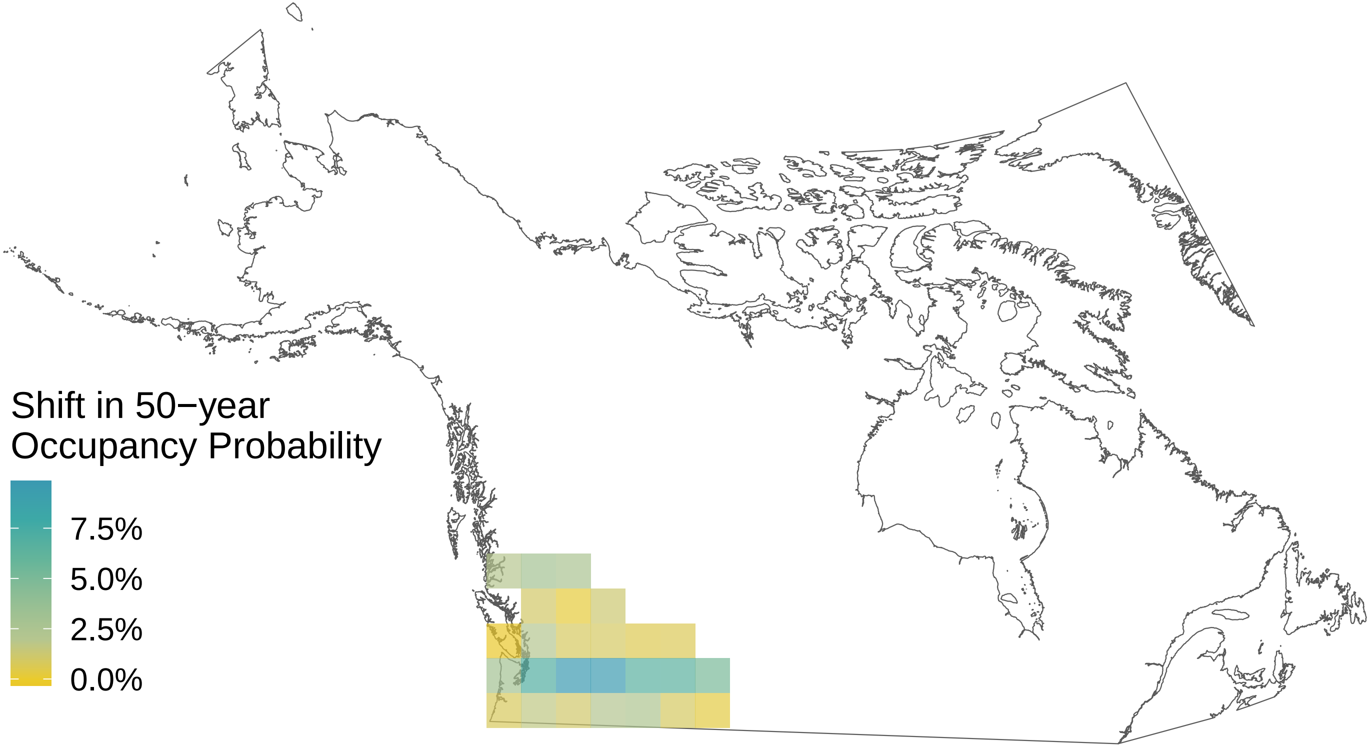

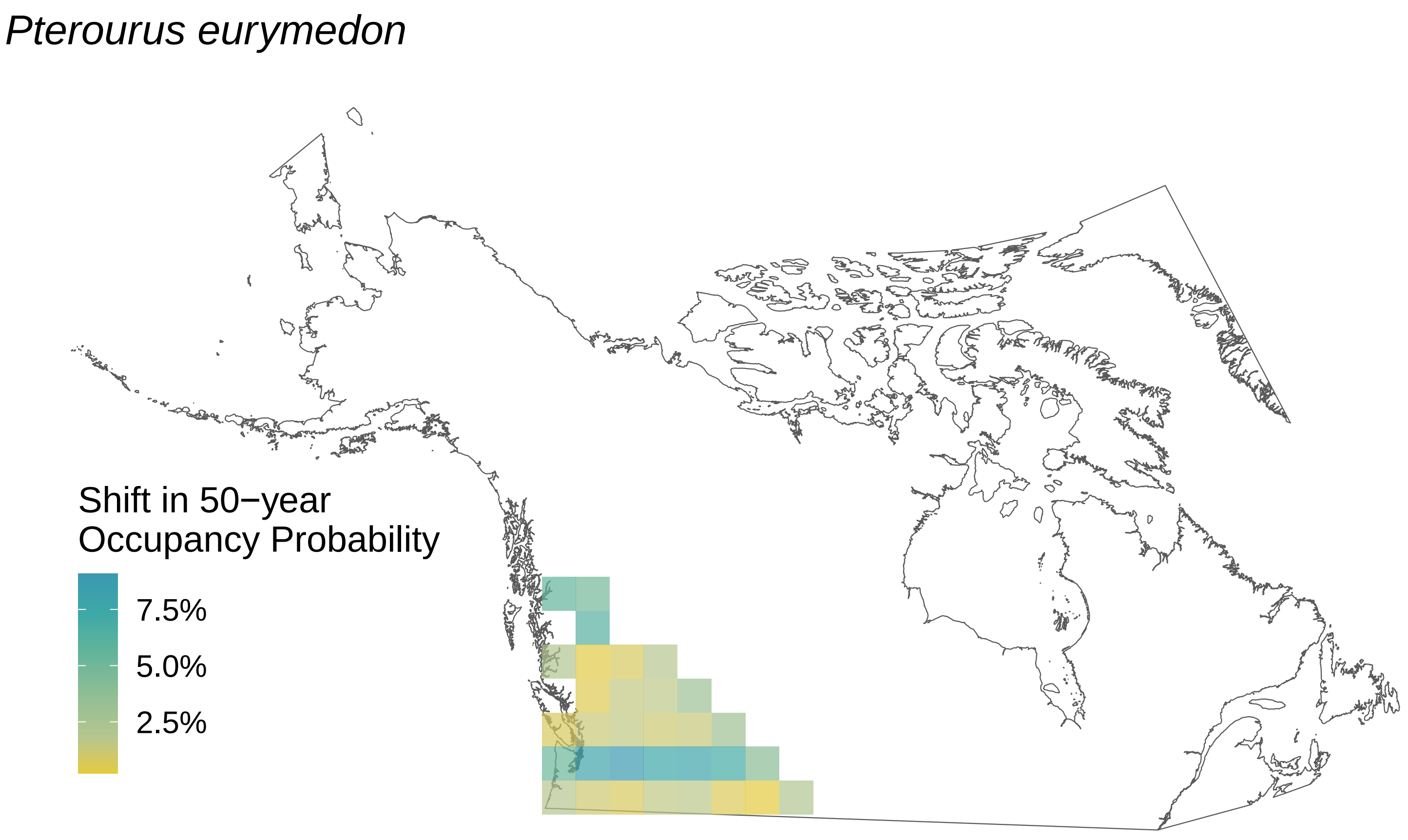

*Pterourus eurymedon*
Shift in 50−year
Occupancy Probability
7.5%
5.0%
2.5%

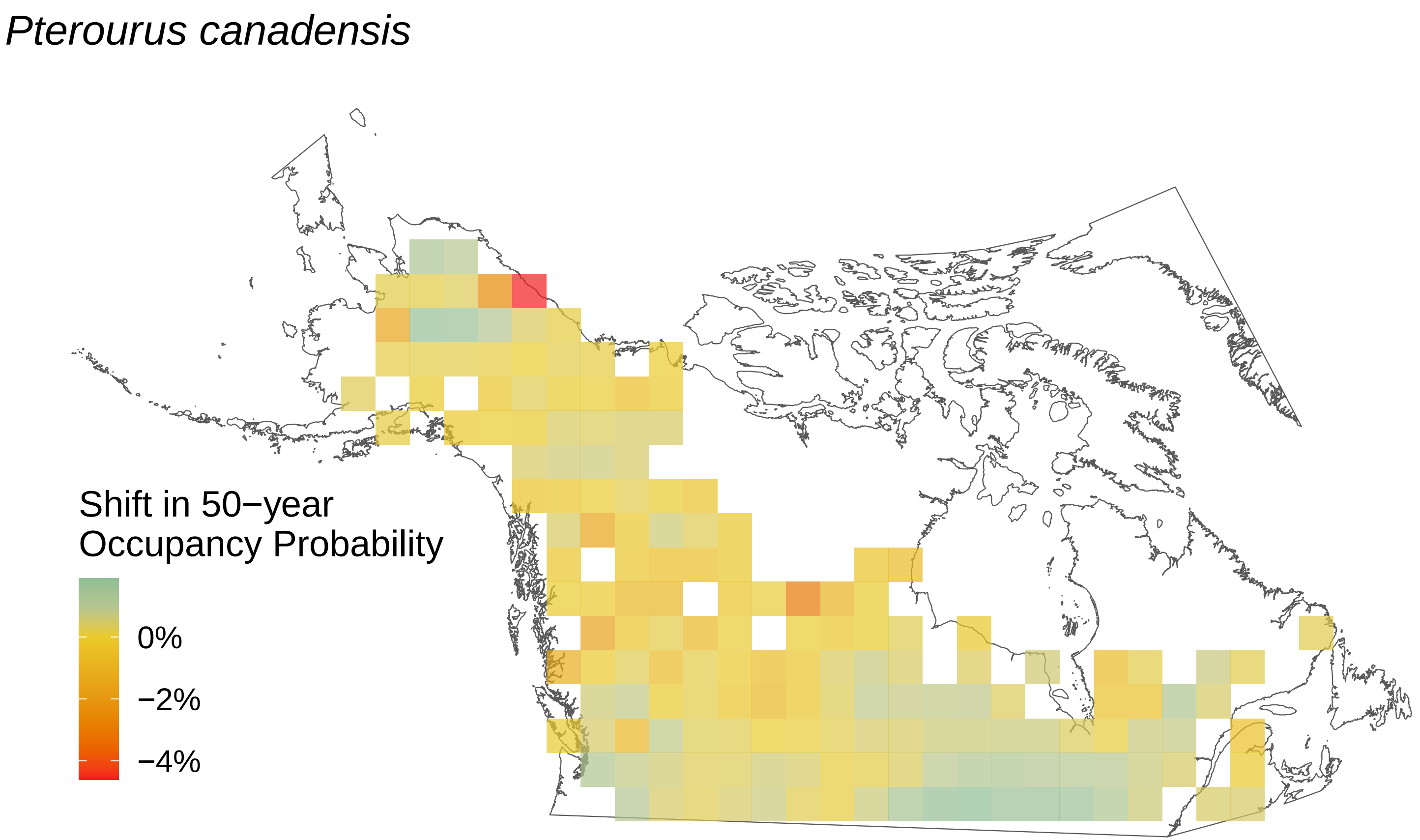

*Pterourus canadensis*
Shift in 50−year
Occupancy Probability
0%
−2%
−4%

*Pontia occidentalis*

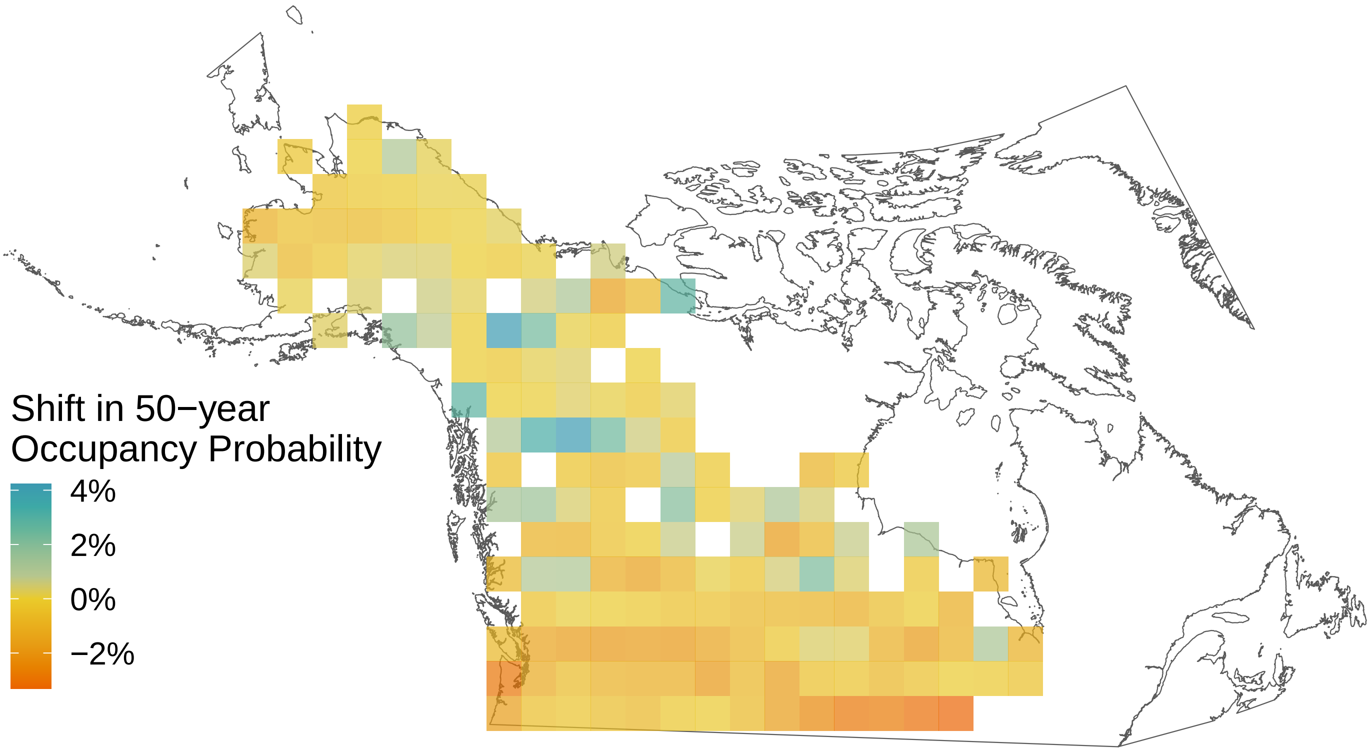

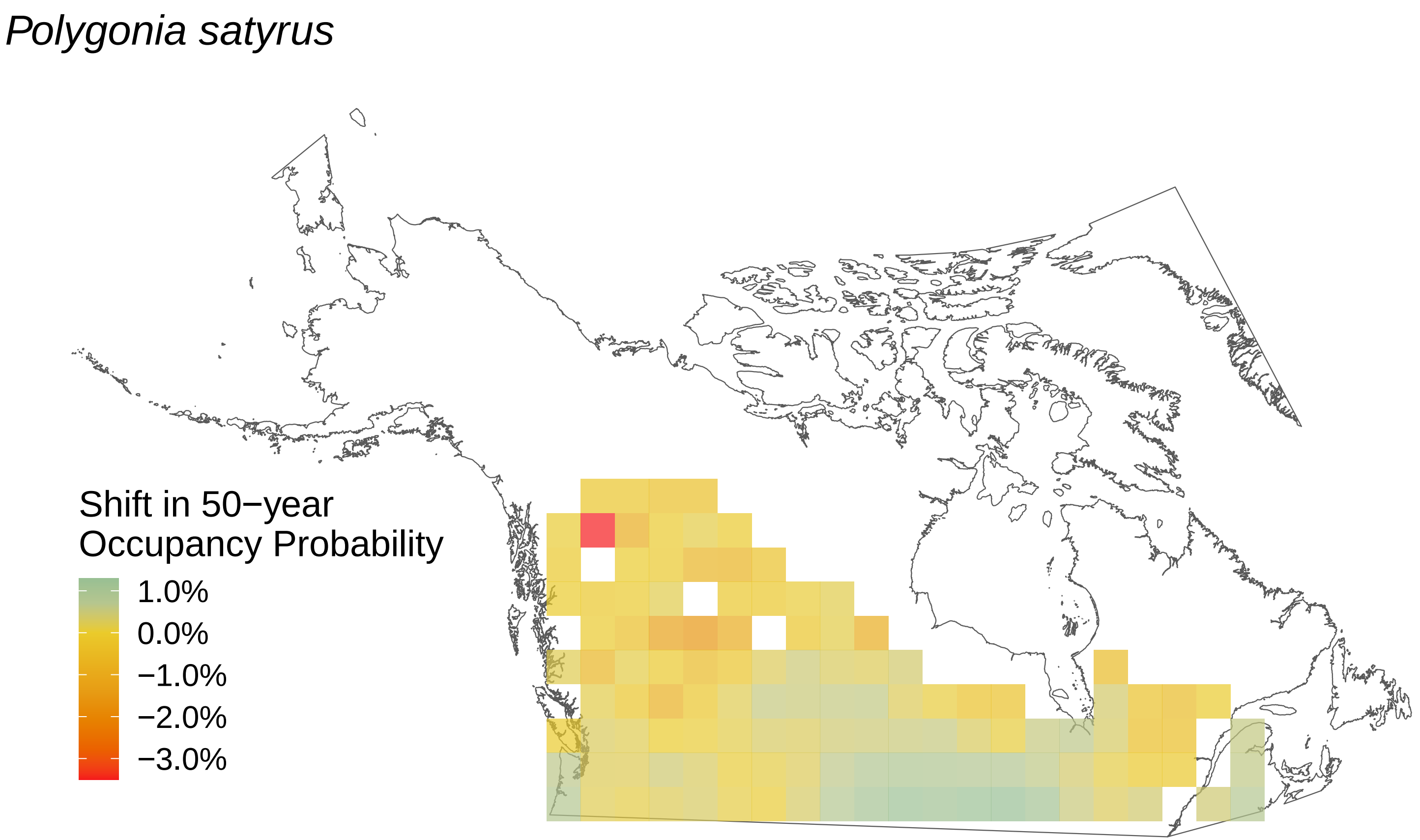

*Polygonia satyrus*
Shift in 50−year Occupancy Probability
1.0%
0.0%
−1.0%
−2.0%
−3.0%

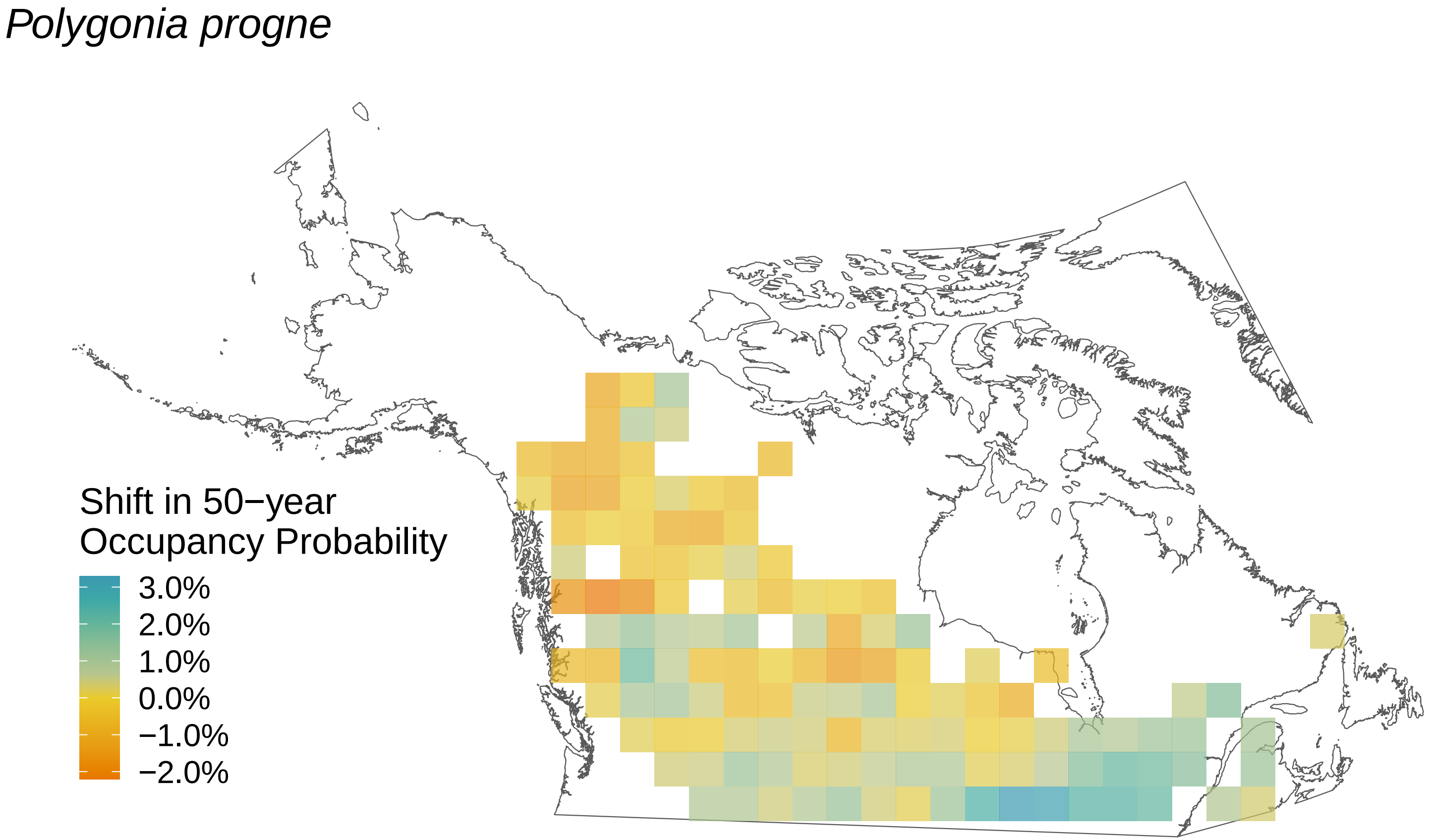

Polygonia progne
Shift in 50−year Occupancy Probability
3.0%
2.0%
1.0%
0.0%
−1.0%
−2.0%

*Polygonia interrogationis*

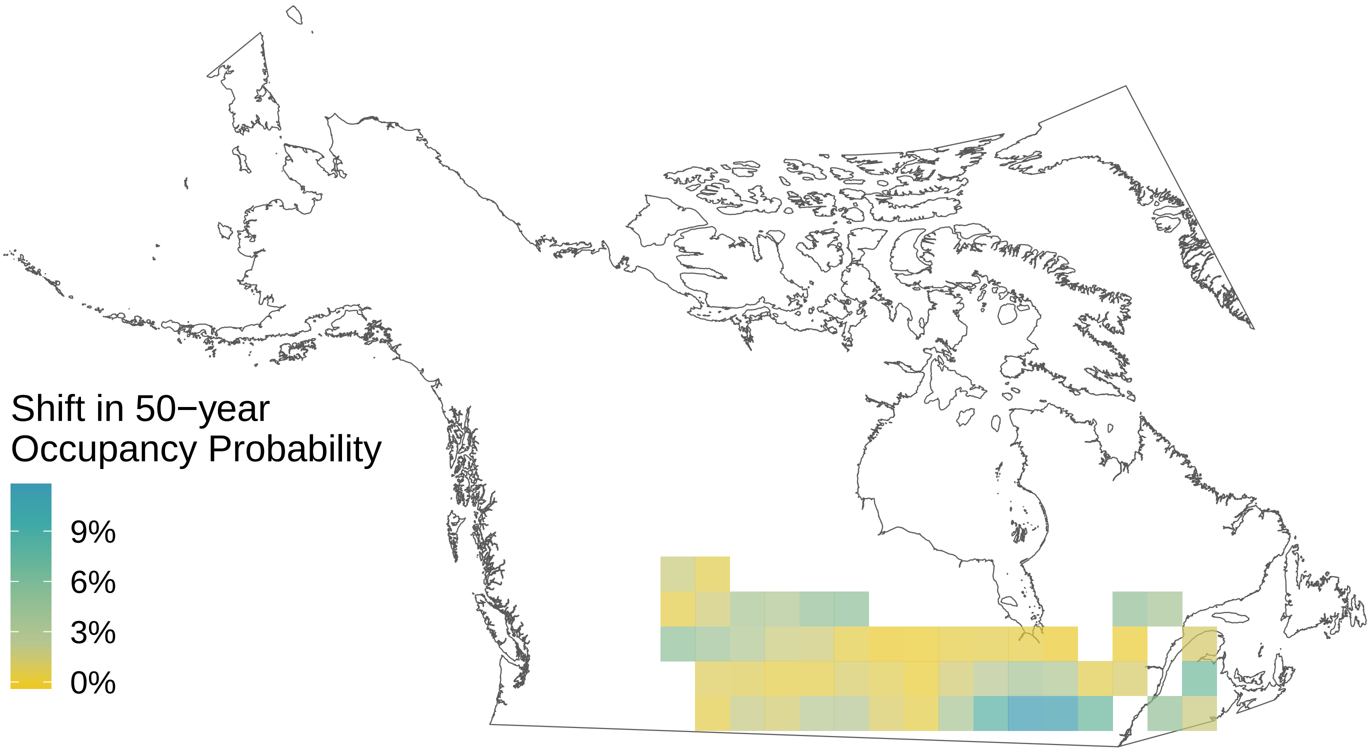

*Polygonia gracilis*

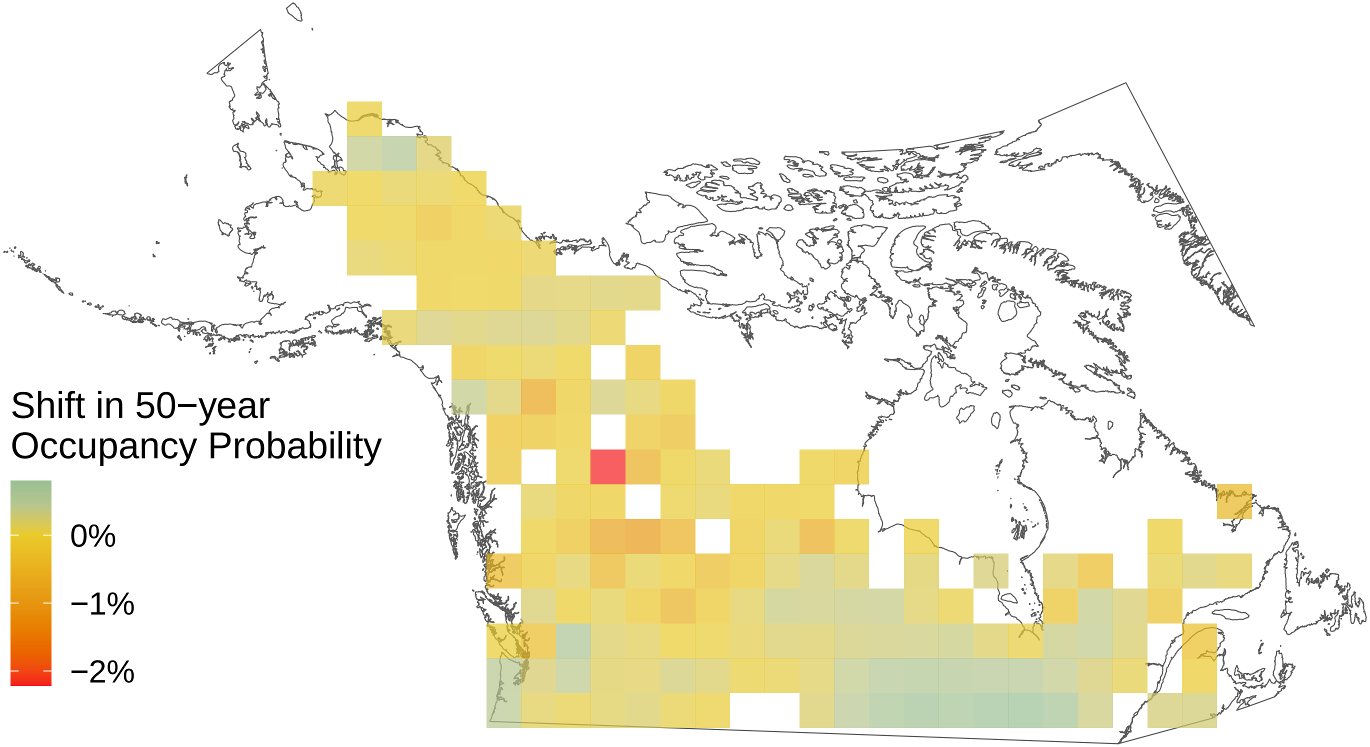

*Polygonia faunus*

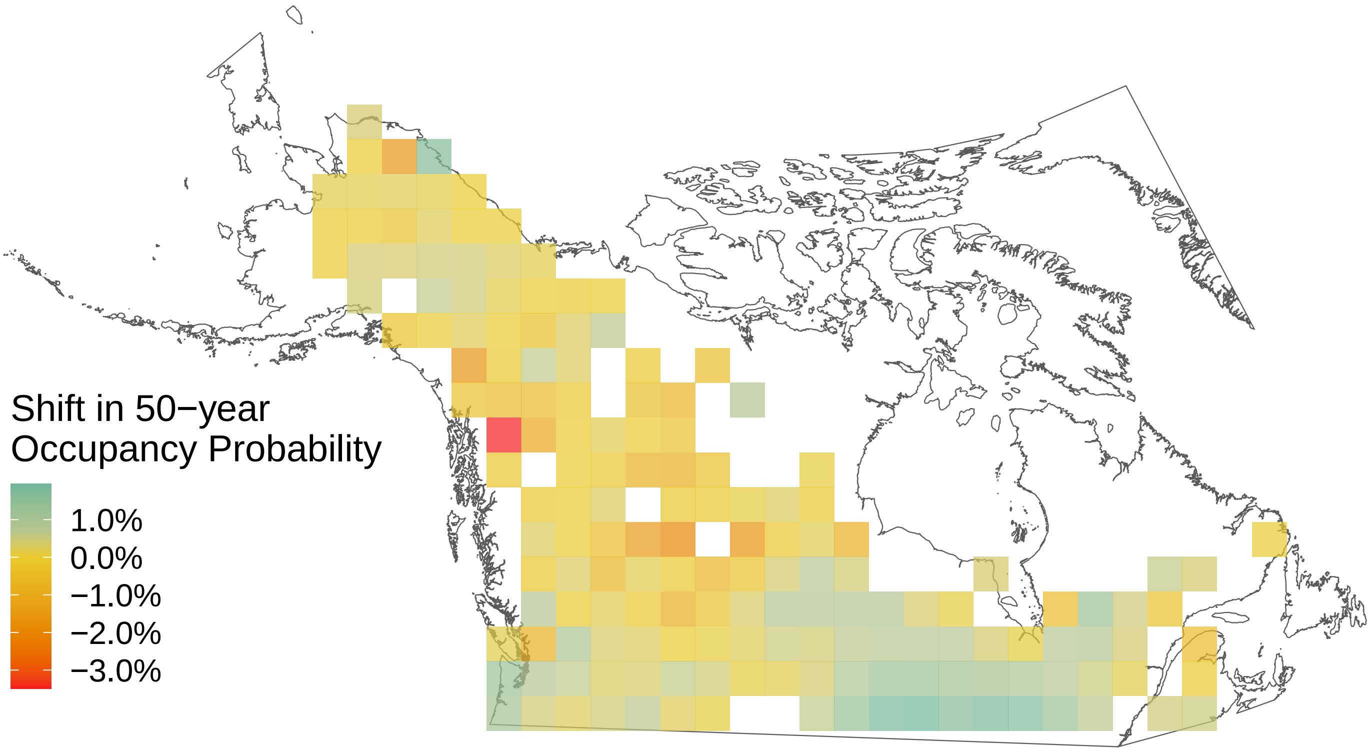

*Polites themistocles*

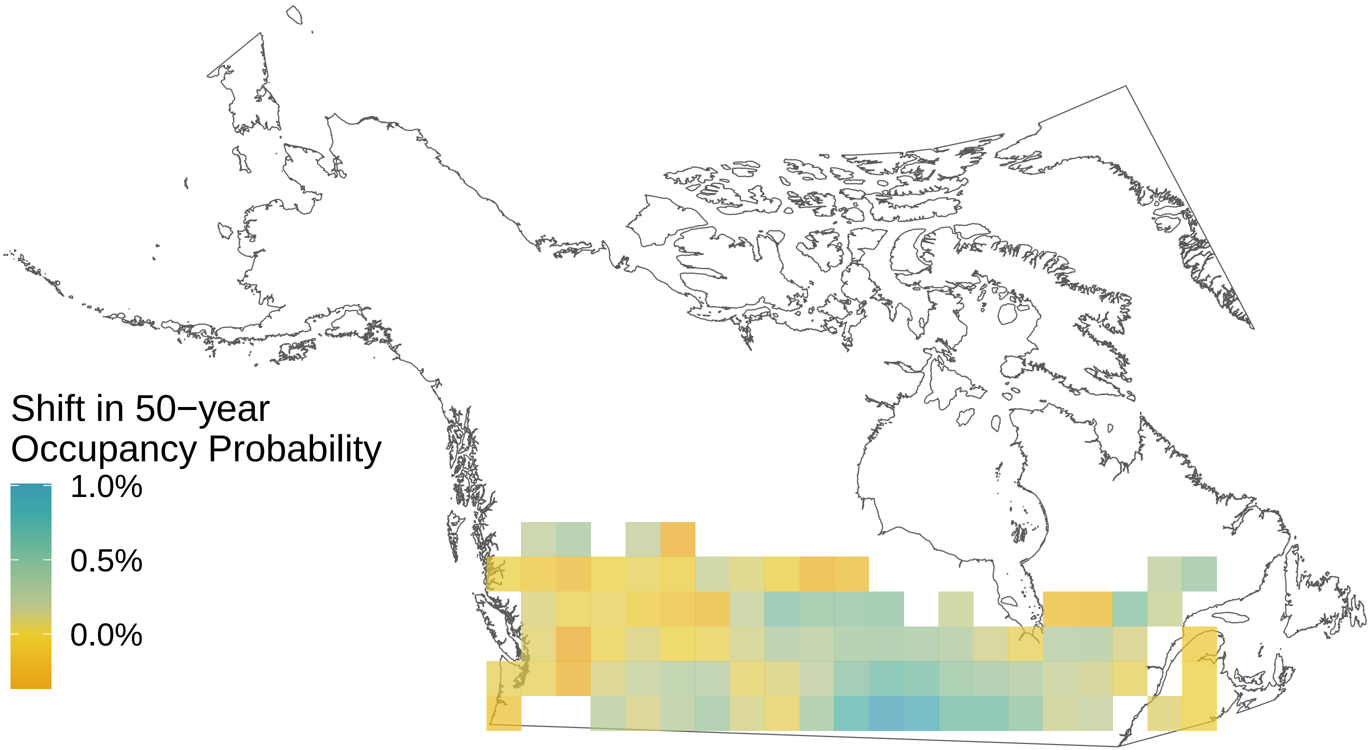

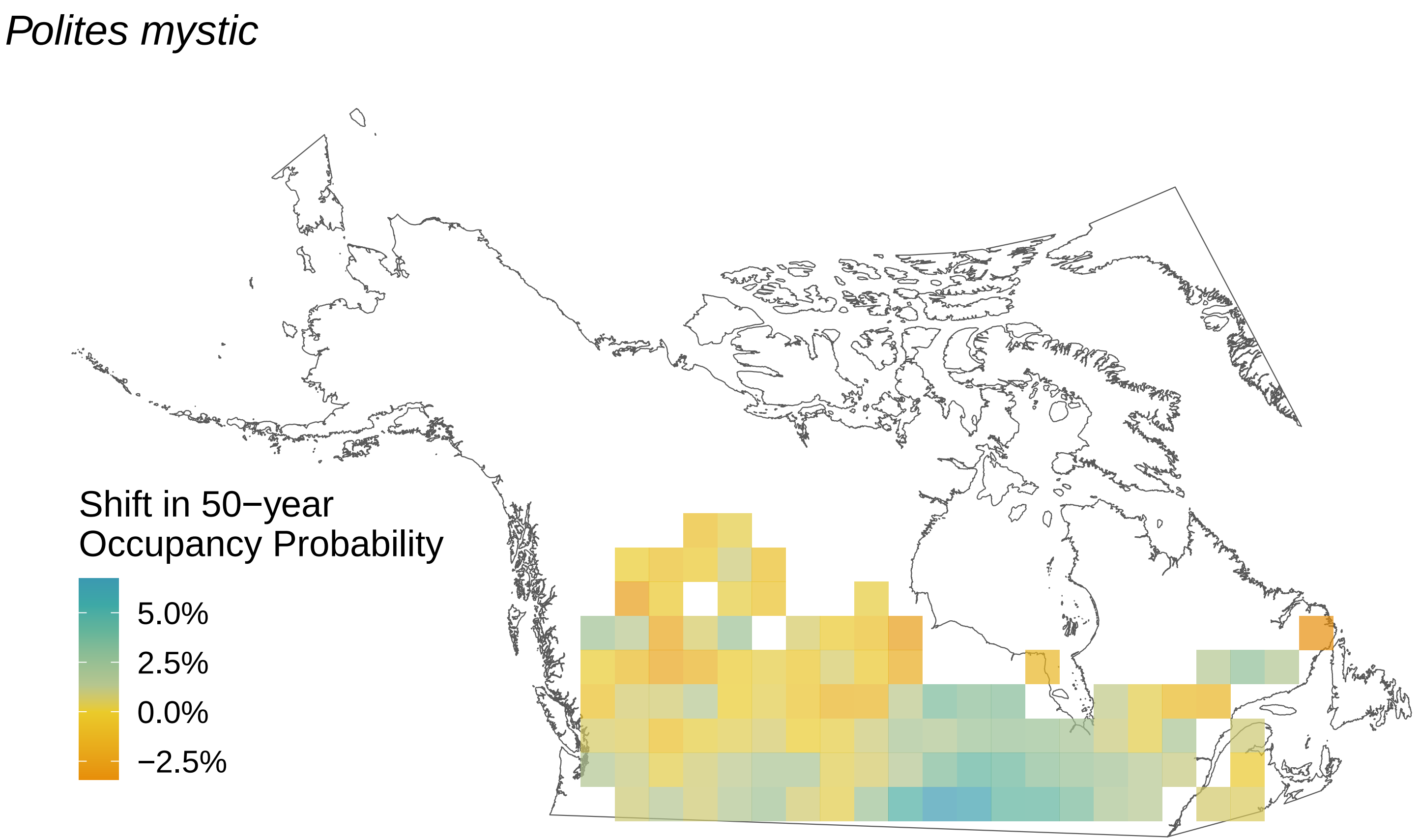
Polites mystic
Shift in 50−year Occupancy Probability
5.0%
2.5%
0.0%
−2.5%

*Poanes hobomok*

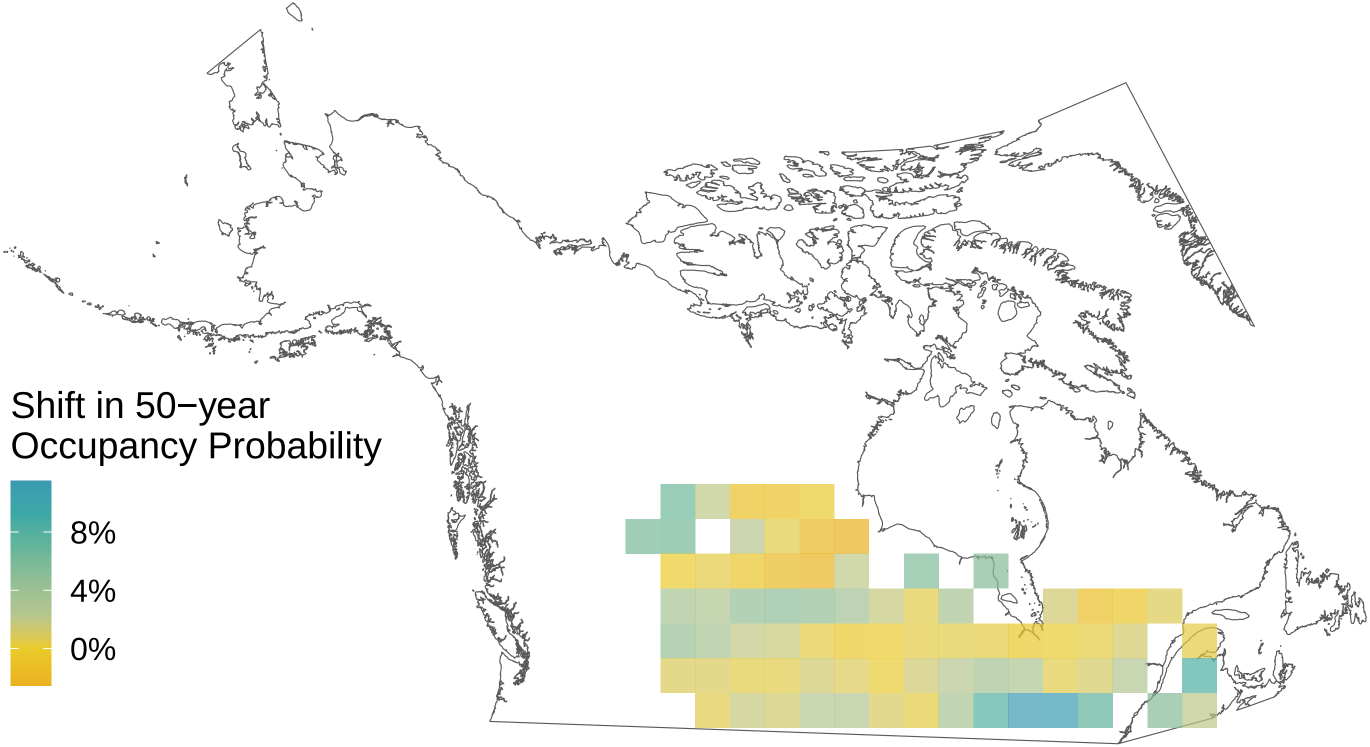

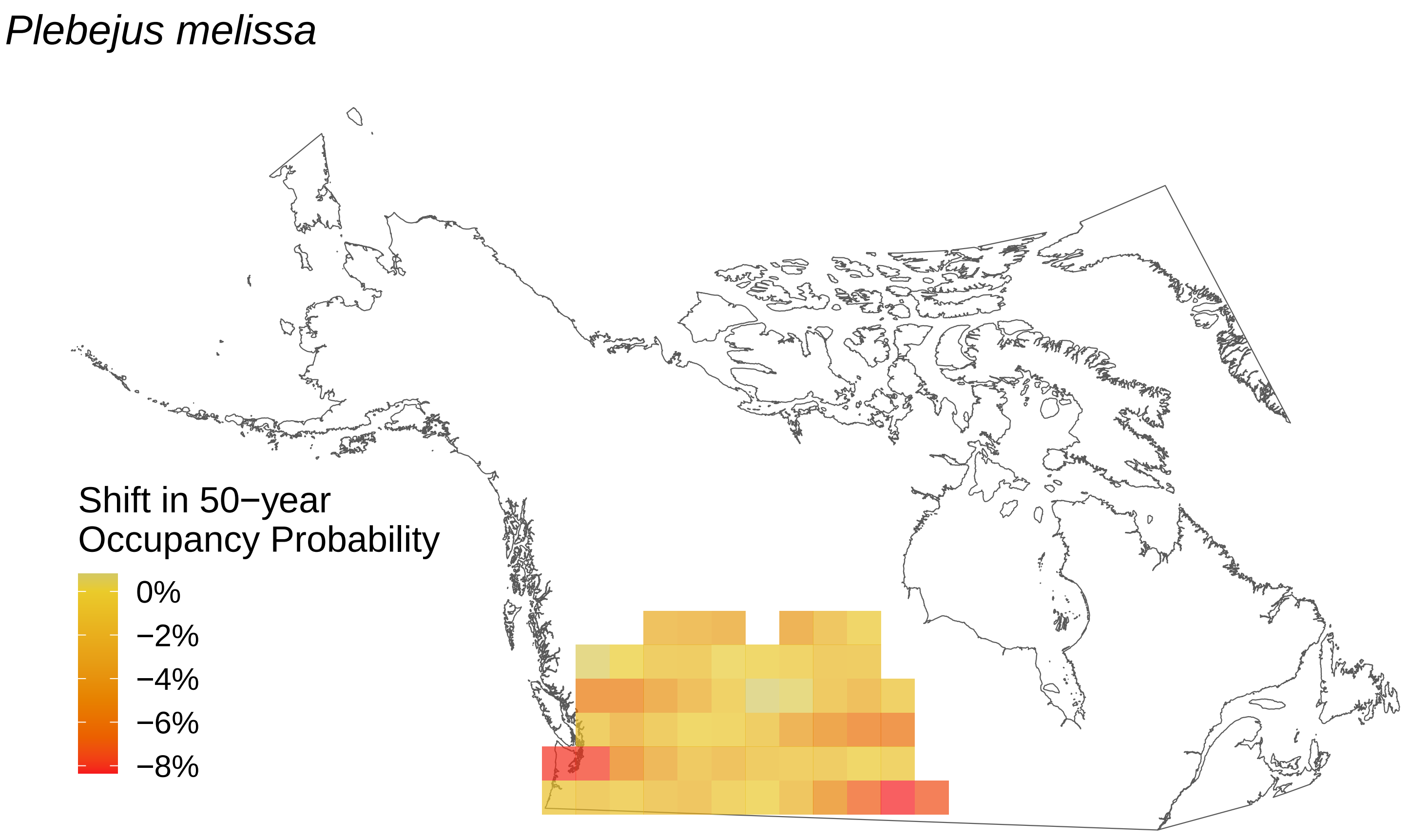

*Plebejus melissa*
Shift in 50−year
Occupancy Probability
0%
−2%
−4%
−6%
−8%

*Plebejus idas*

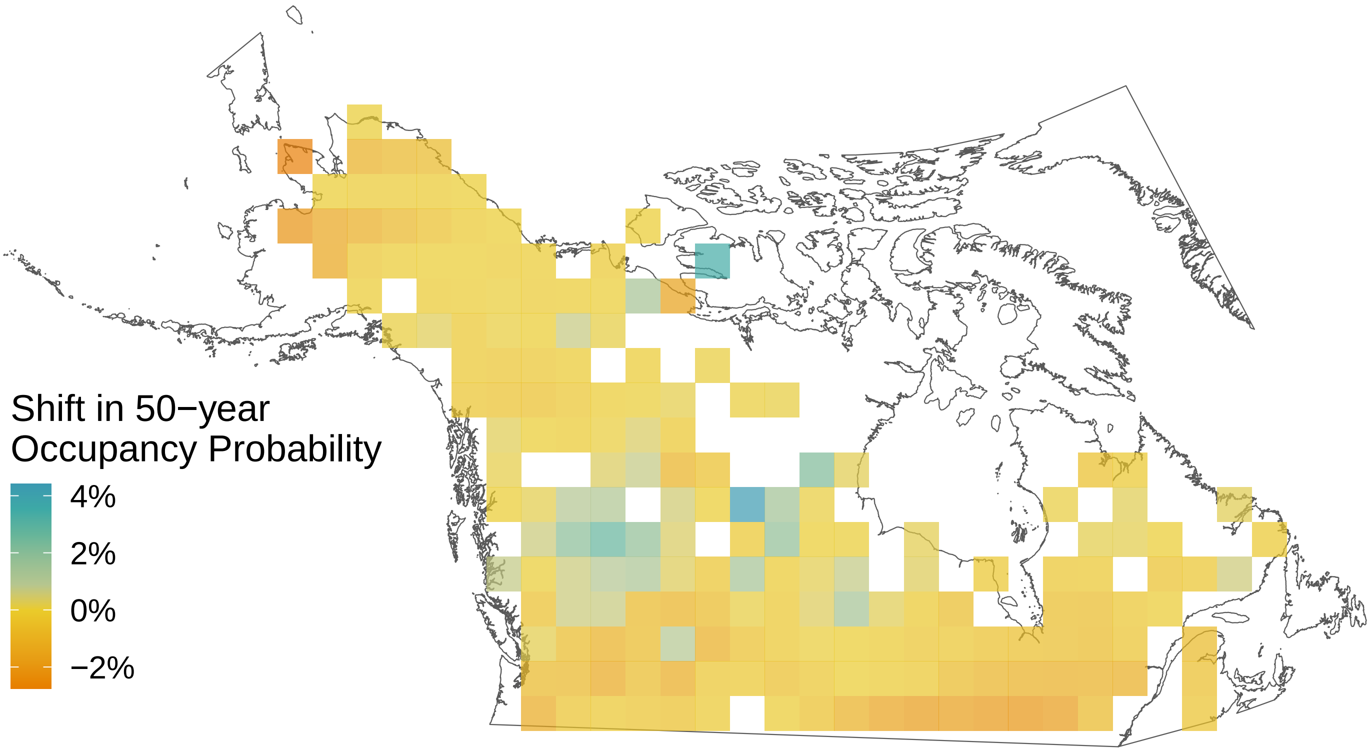

*Pieris rapae*

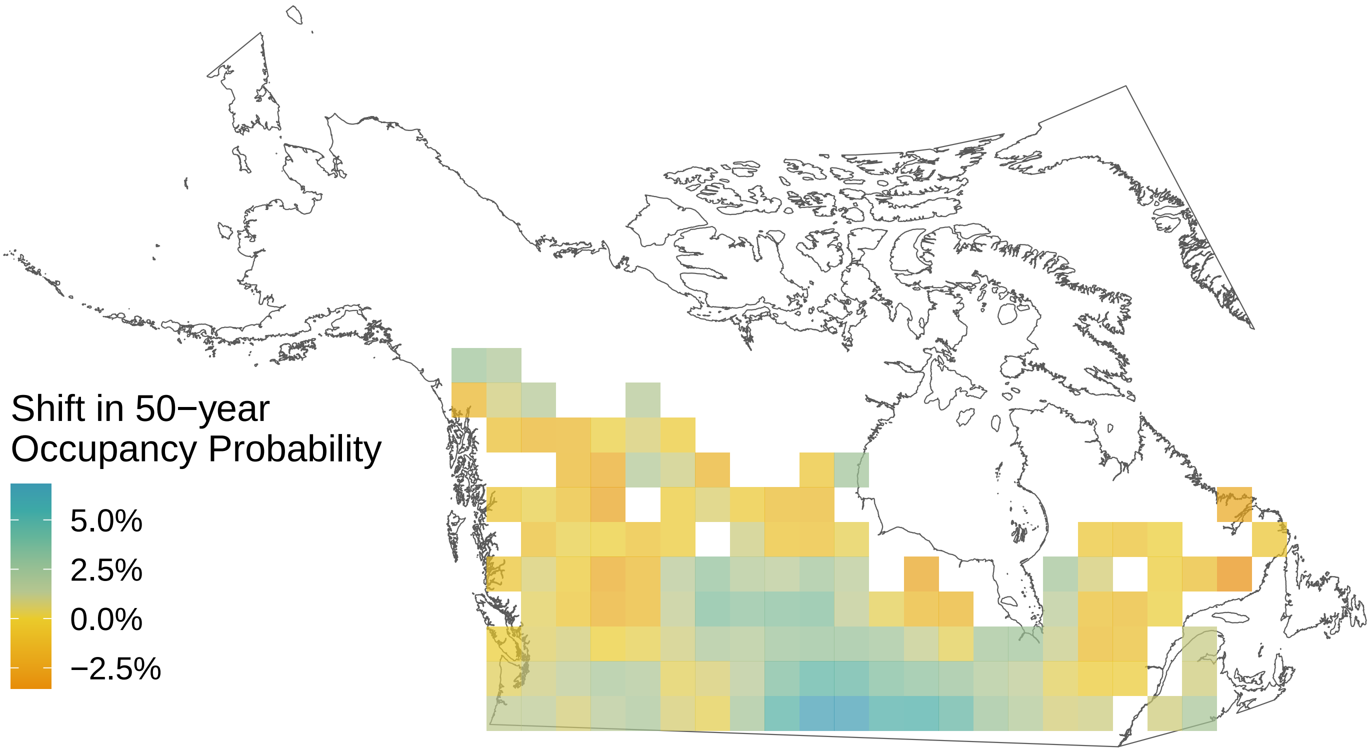

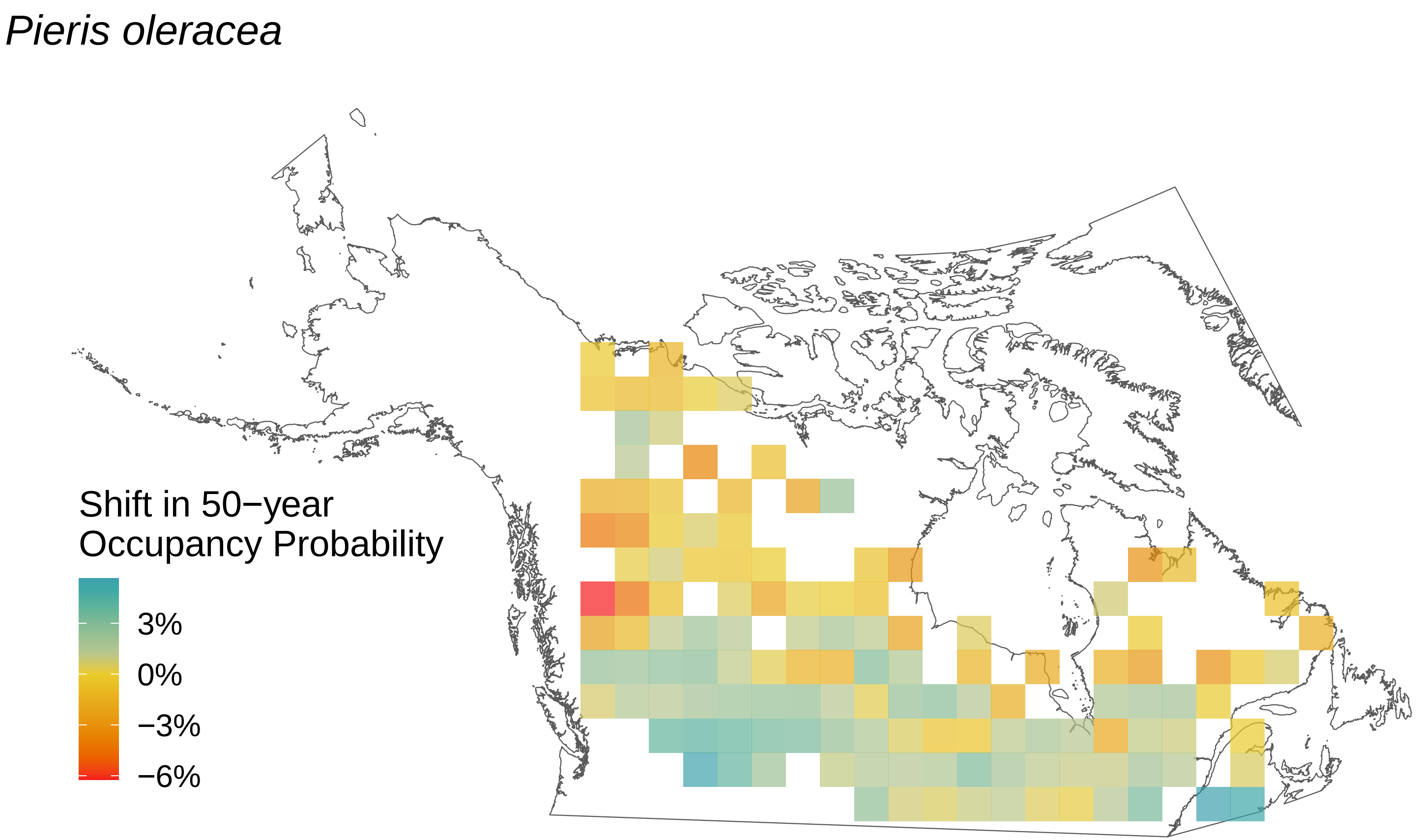
*Pieris oleracea*
Shift in 50−year
Occupancy Probability
3%
0%
−3%
−6%

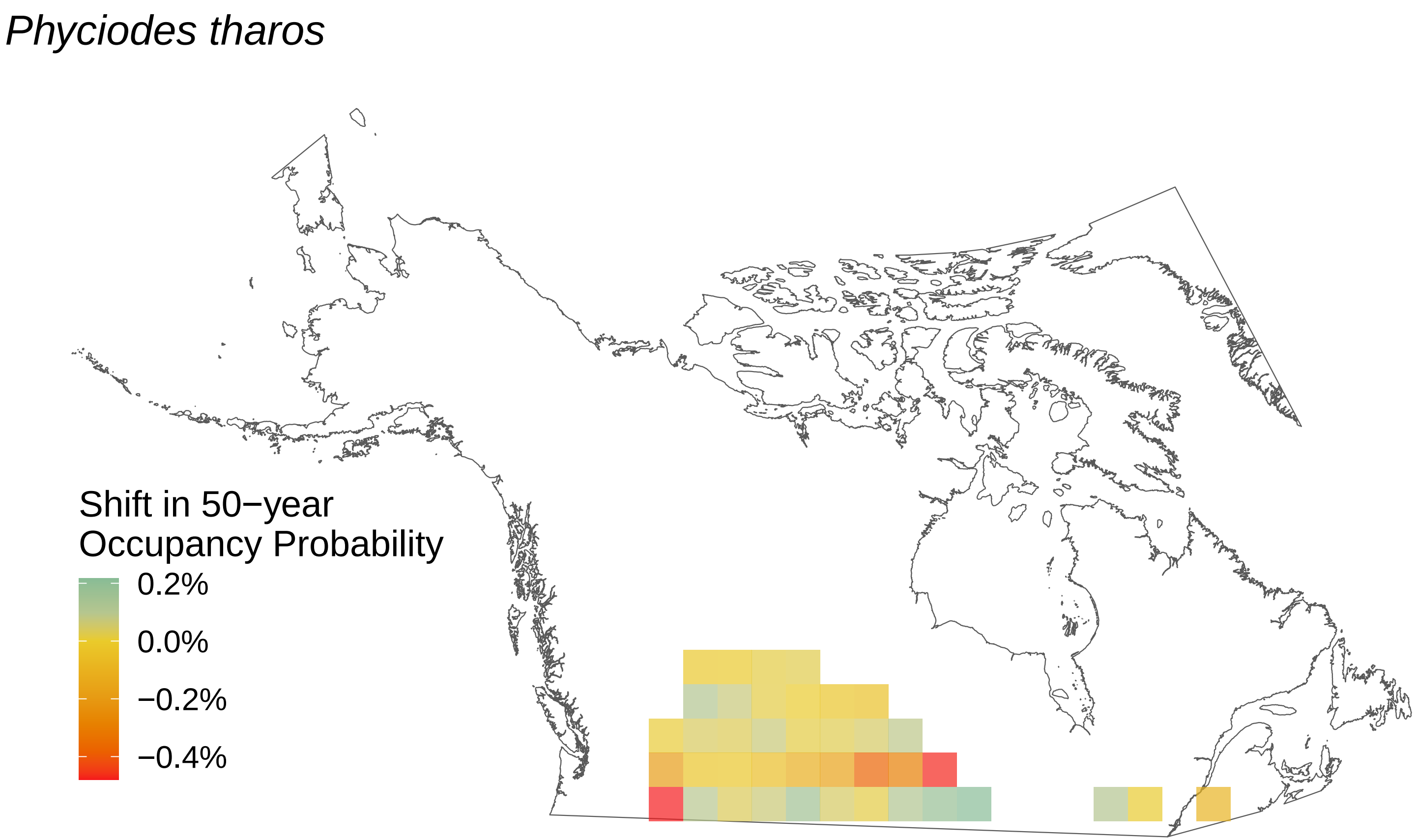
Phyciodes tharos
Shift in 50−year
Occupancy Probability
0.2%
0.0%
−0.2%
−0.4%

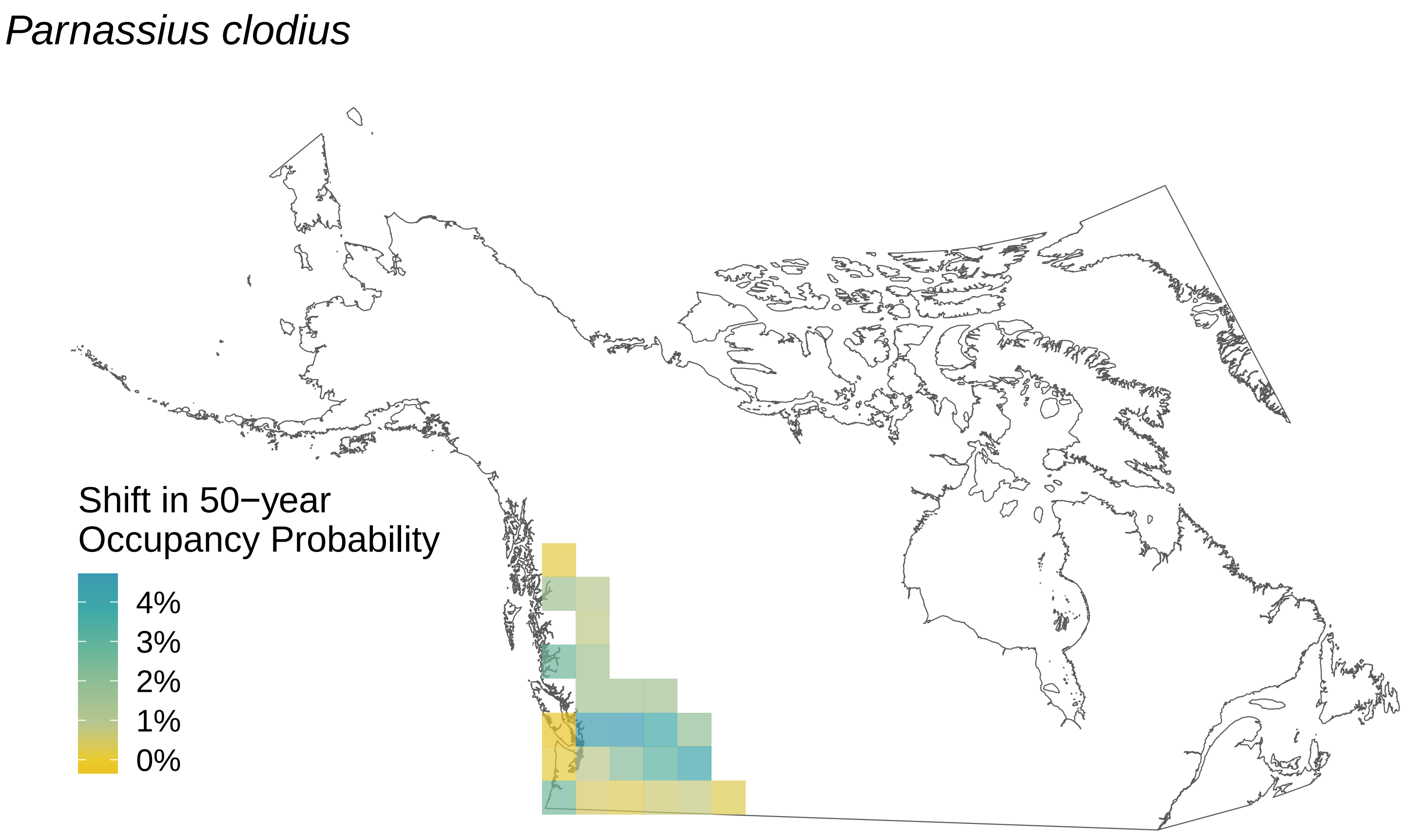
Parnassius clodius
Shift in 50−year
Occupancy Probability
4%
3%
2%
1%
0%

*Papilio zelicaon*

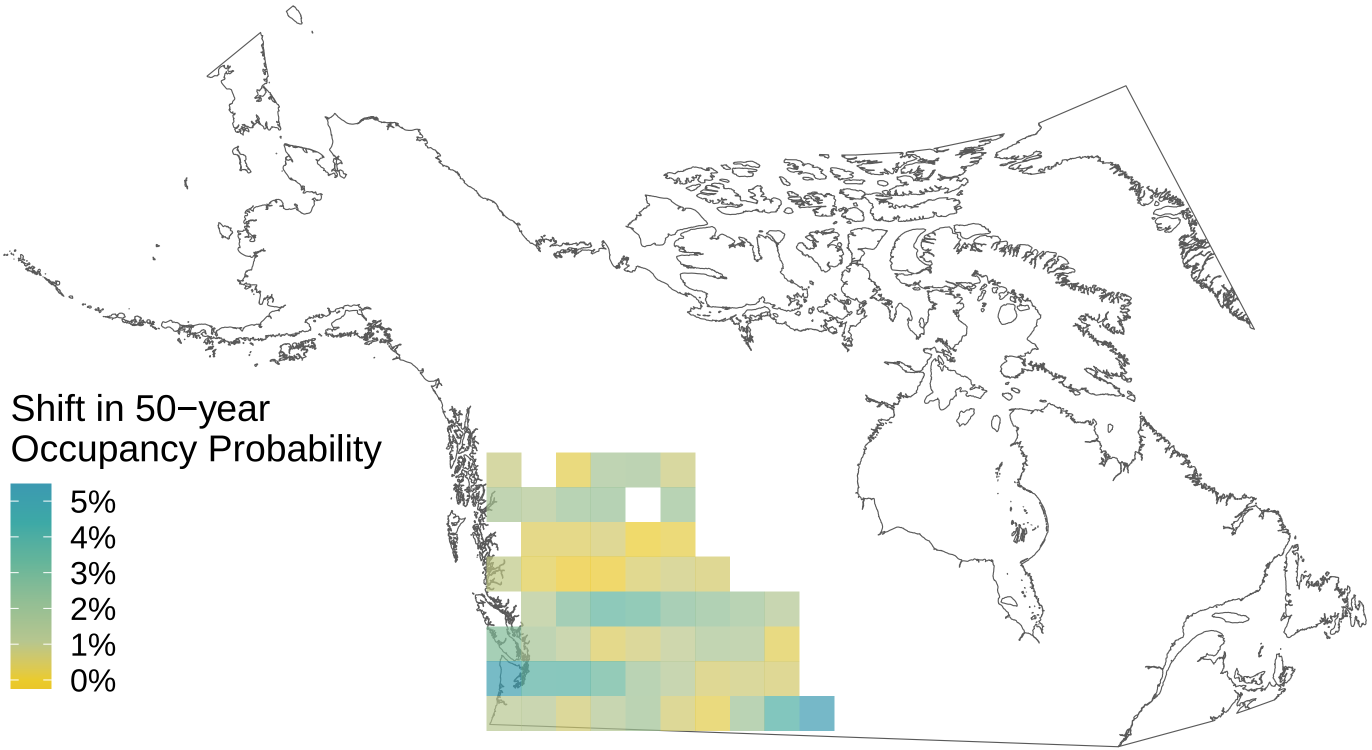

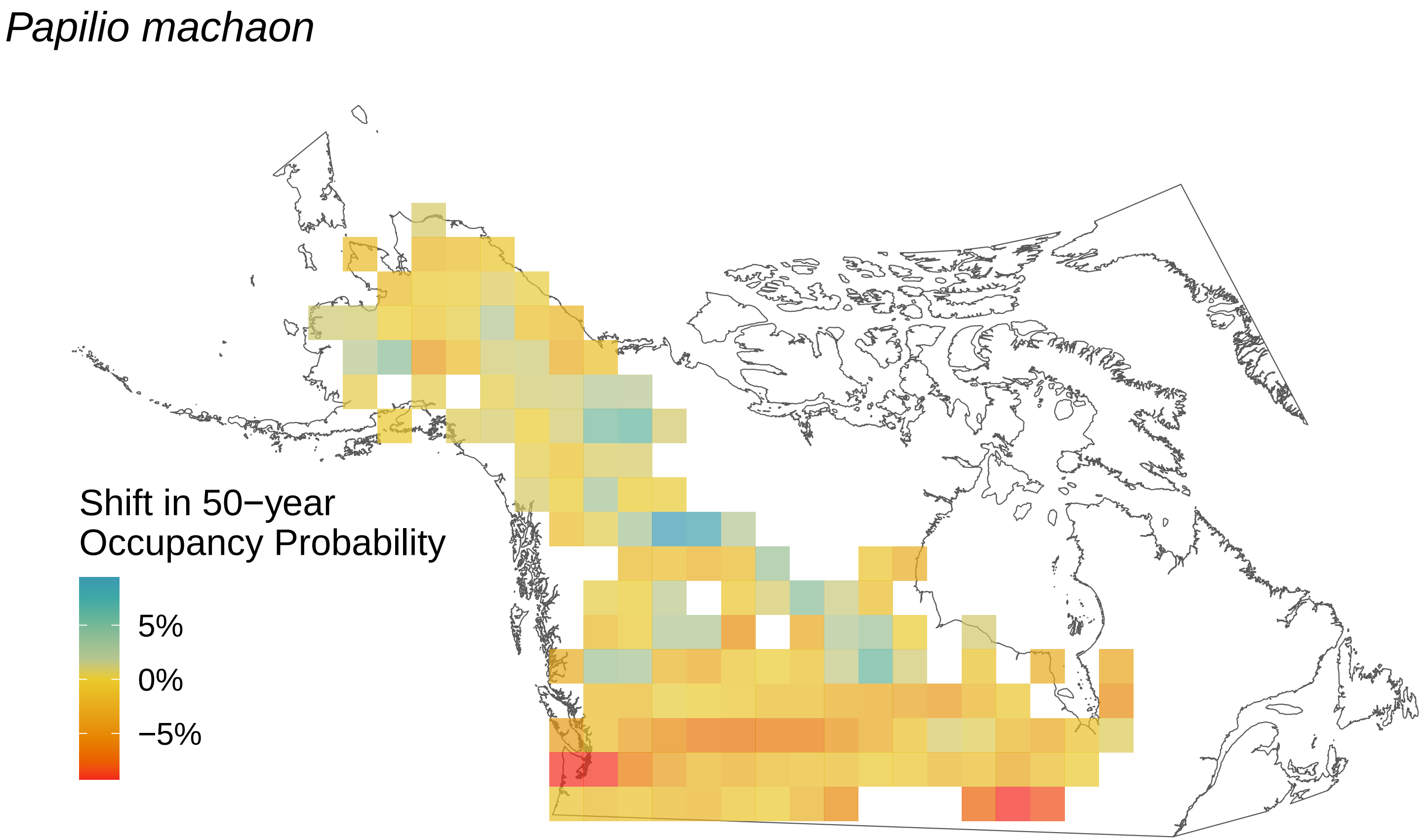

Papilio machaon
Shift in 50−year
Occupancy Probability
5%
0%
−5%

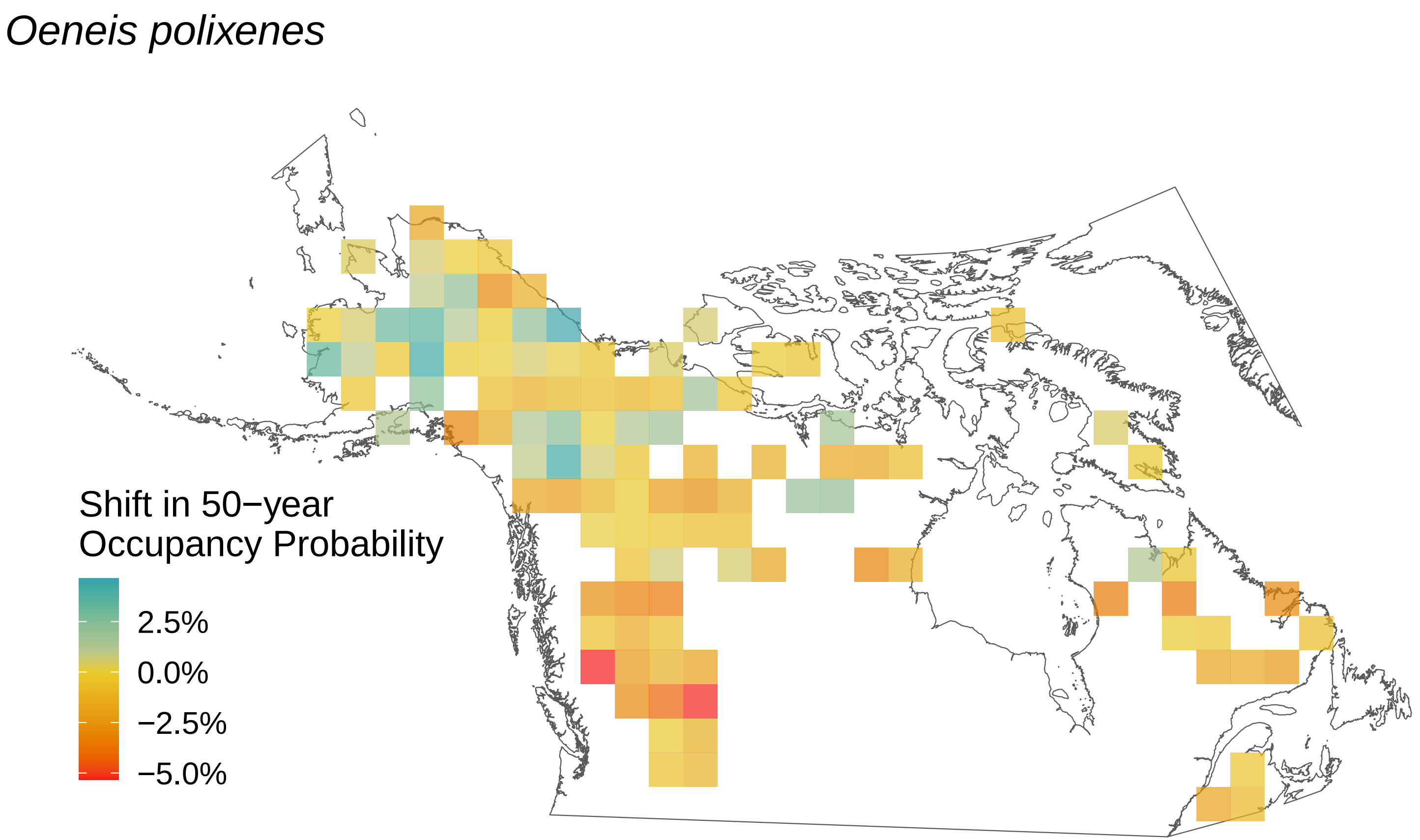
Oeneis polixenes
Shift in 50−year
Occupancy Probability
2.5%
0.0%
−2.5%
−5.0%

*Oeneis melissa*

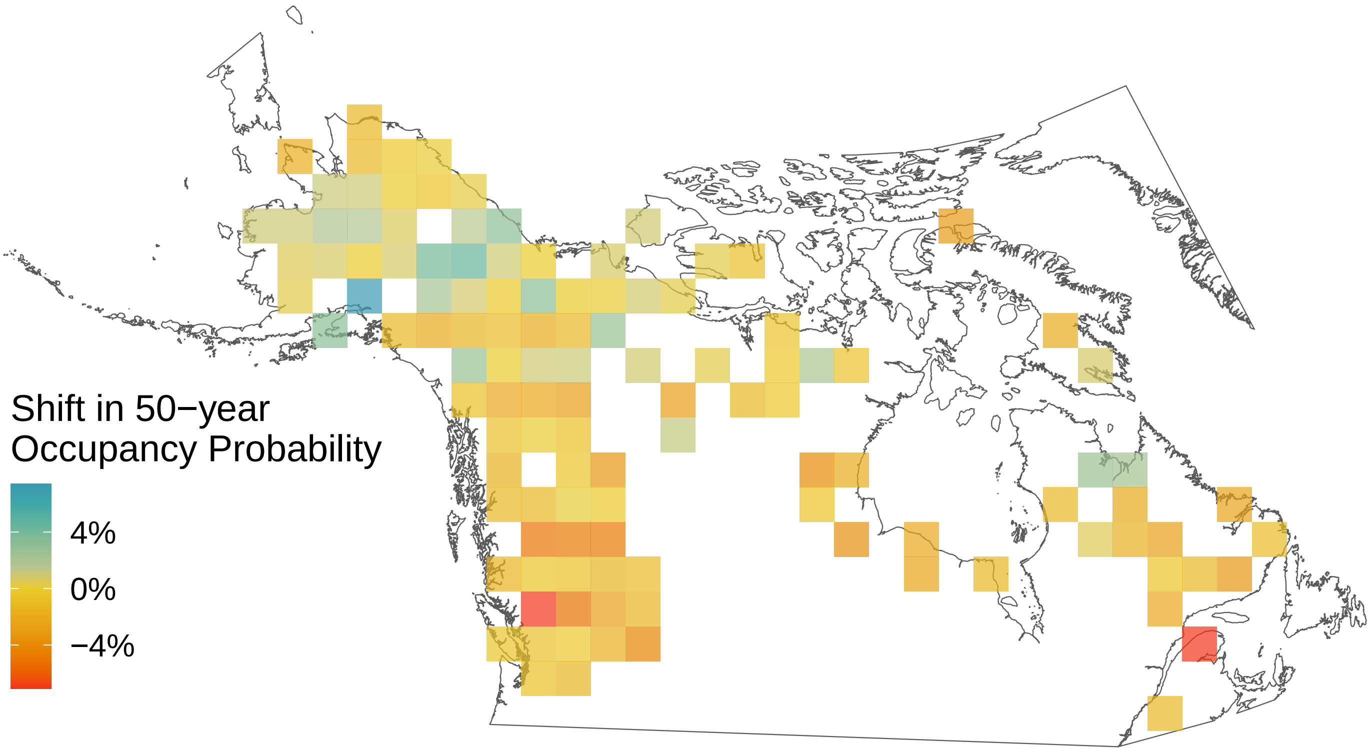

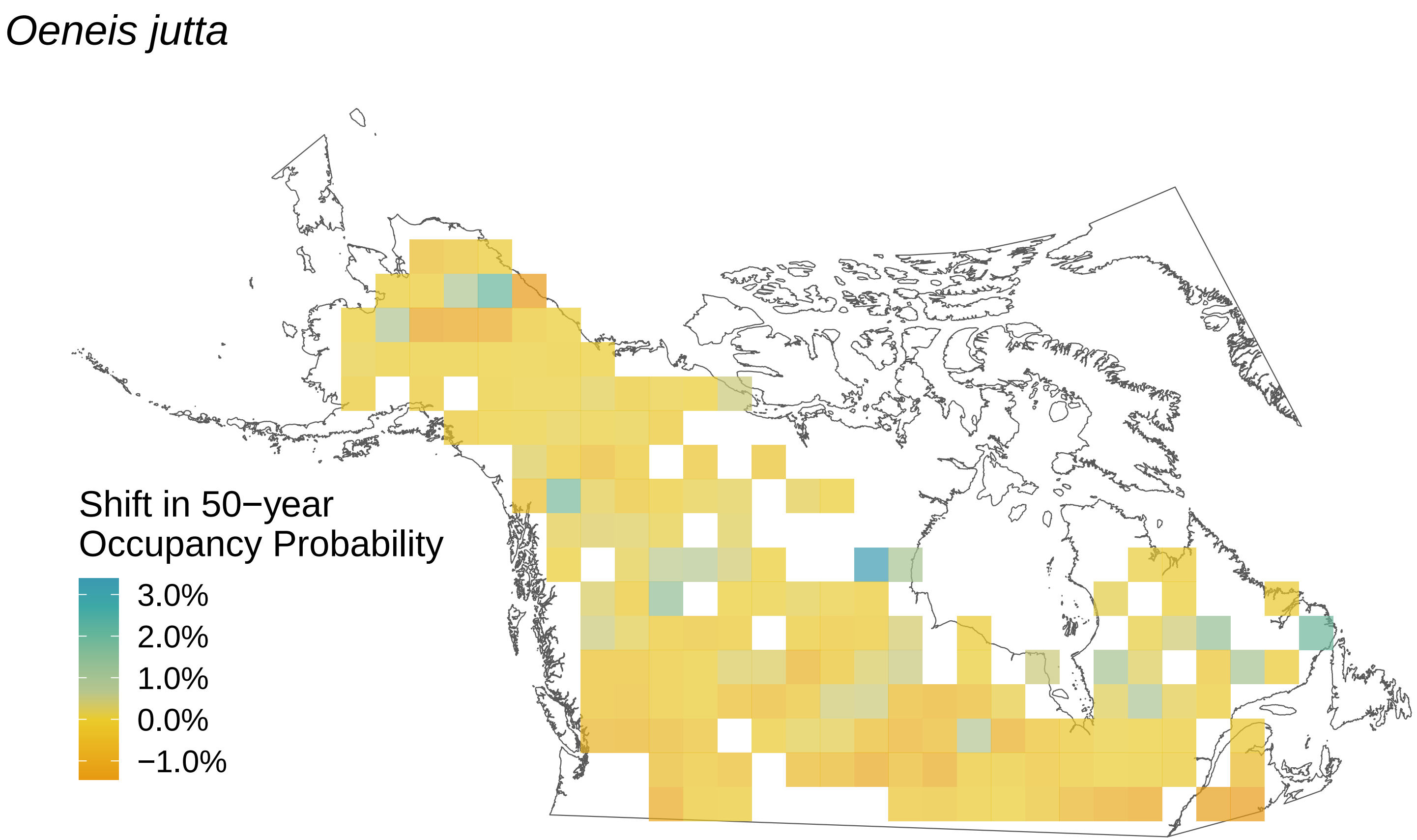

Oeneis jutta
Shift in 50−year
Occupancy Probability
3.0%
2.0%
1.0%
0.0%
−1.0%

*Oeneis chryxus*

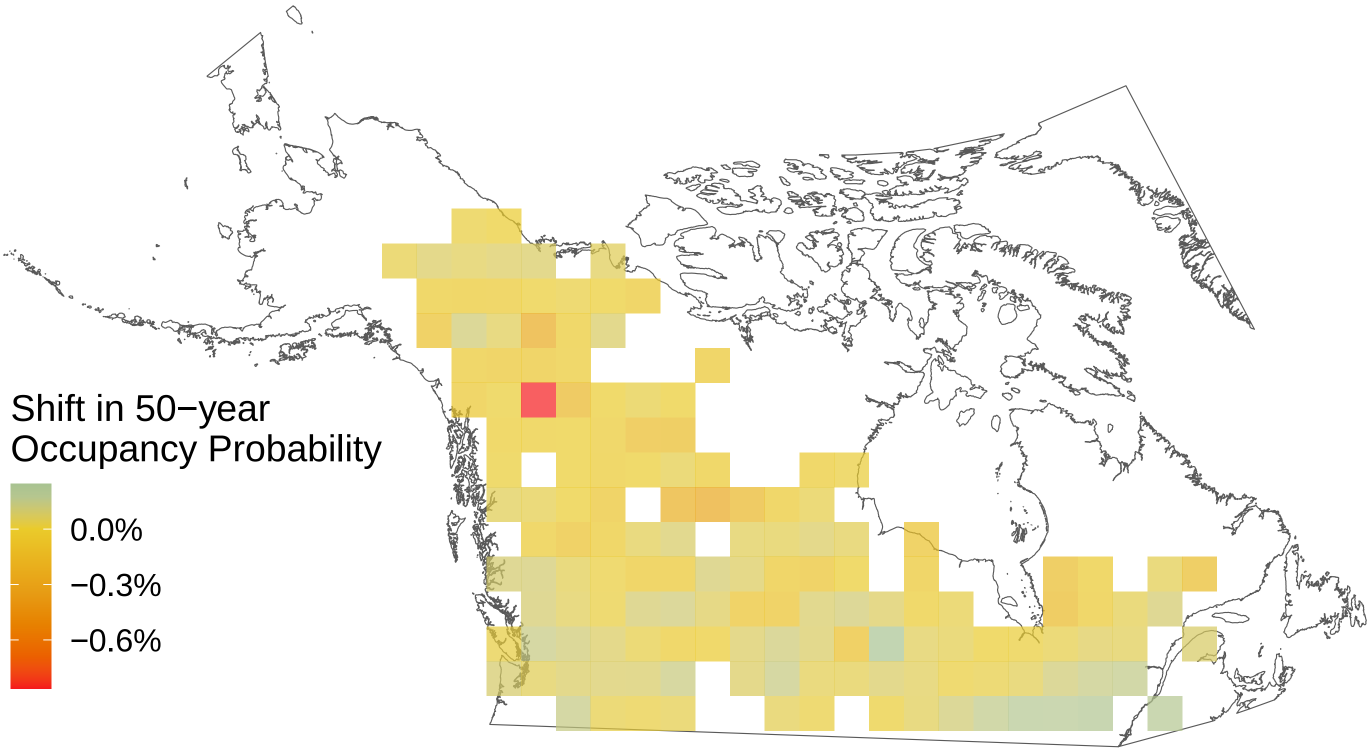

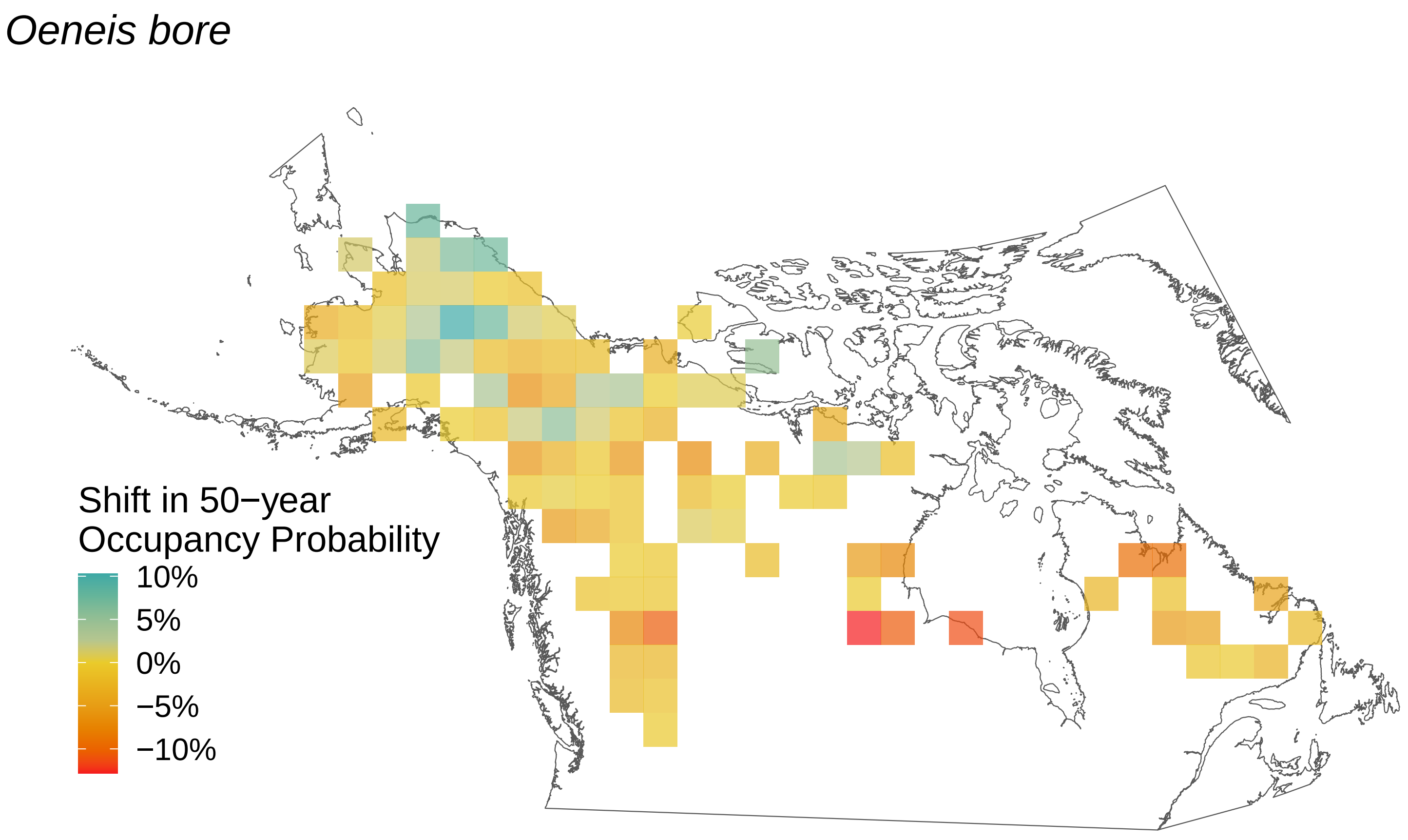
Oeneis bore
Shift in 50−year
Occupancy Probability
10%
5%
0%
−5%
−10%

### *Ochlodes sylvanoides*

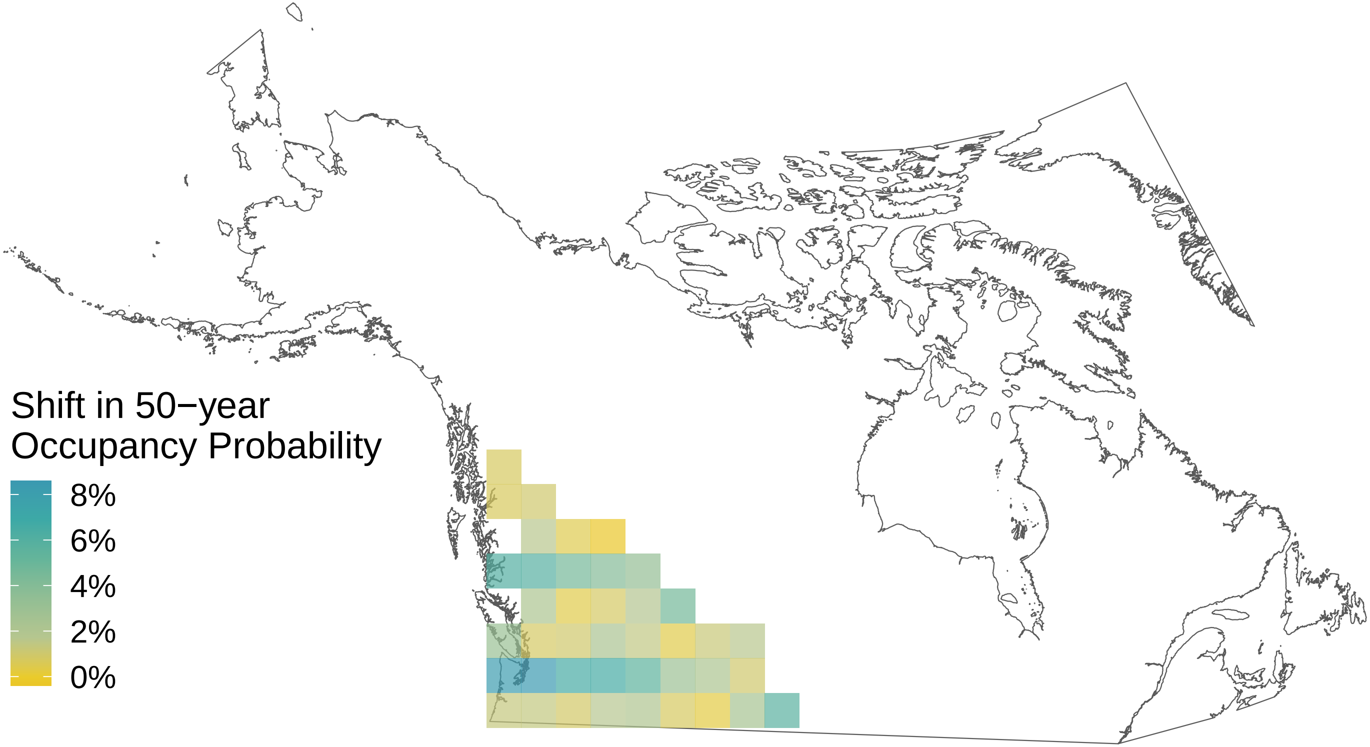

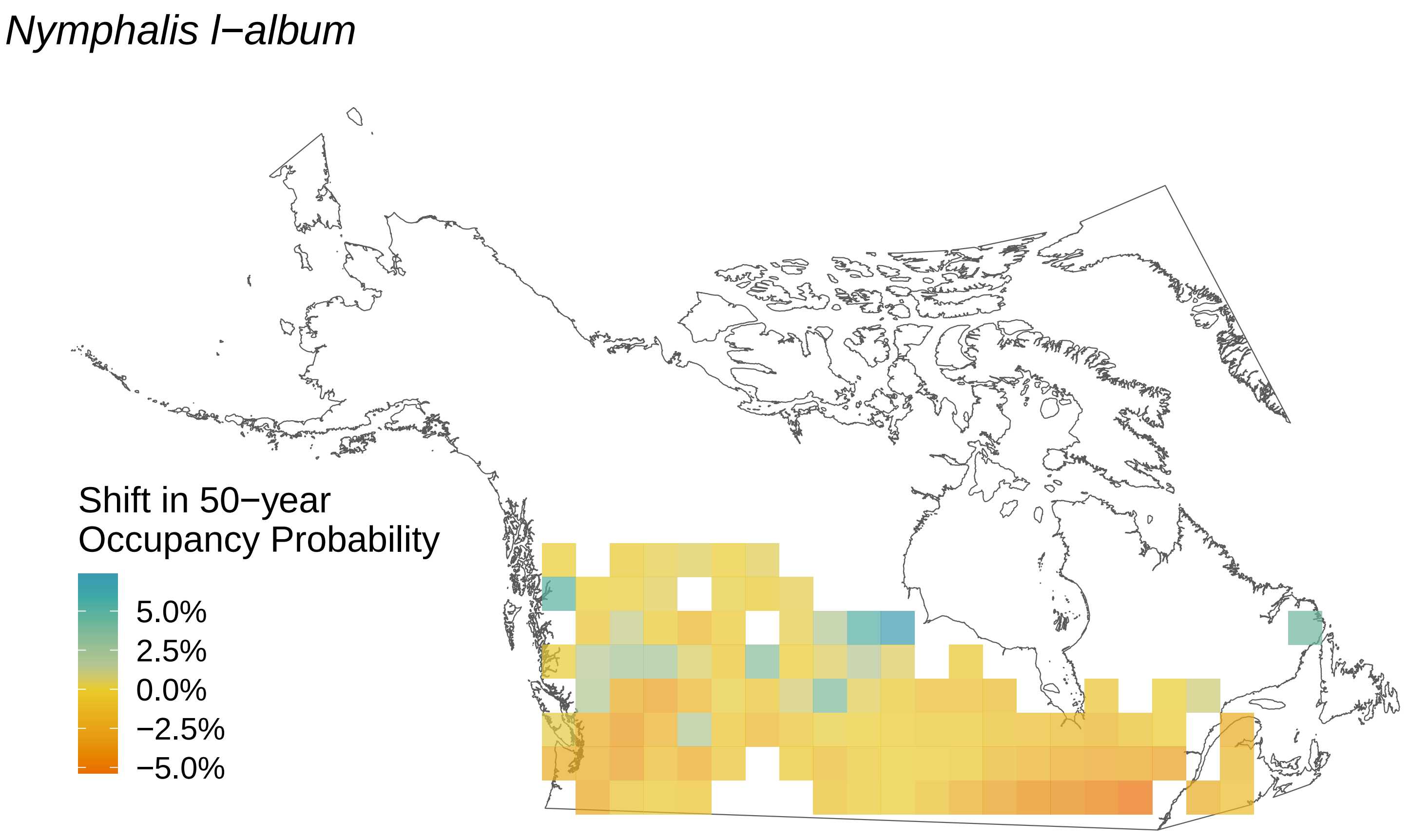
Nymphalis l−album
Shift in 50−year
Occupancy Probability
5.0%
2.5%
0.0%
−2.5%
−5.0%

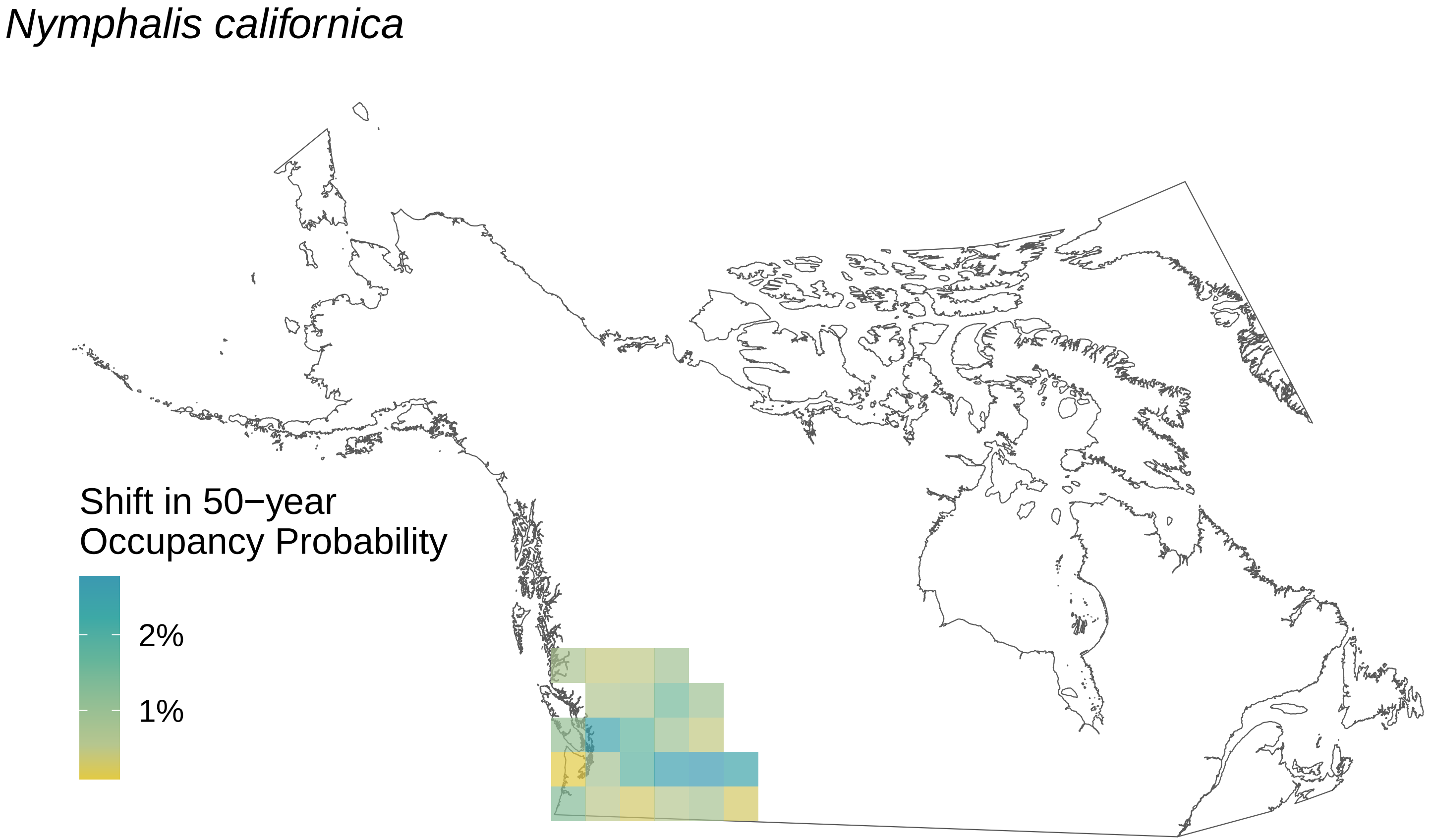

Nymphalis californica
Shift in 50−year
Occupancy Probability
2%
1%

*Nymphalis antiopa*

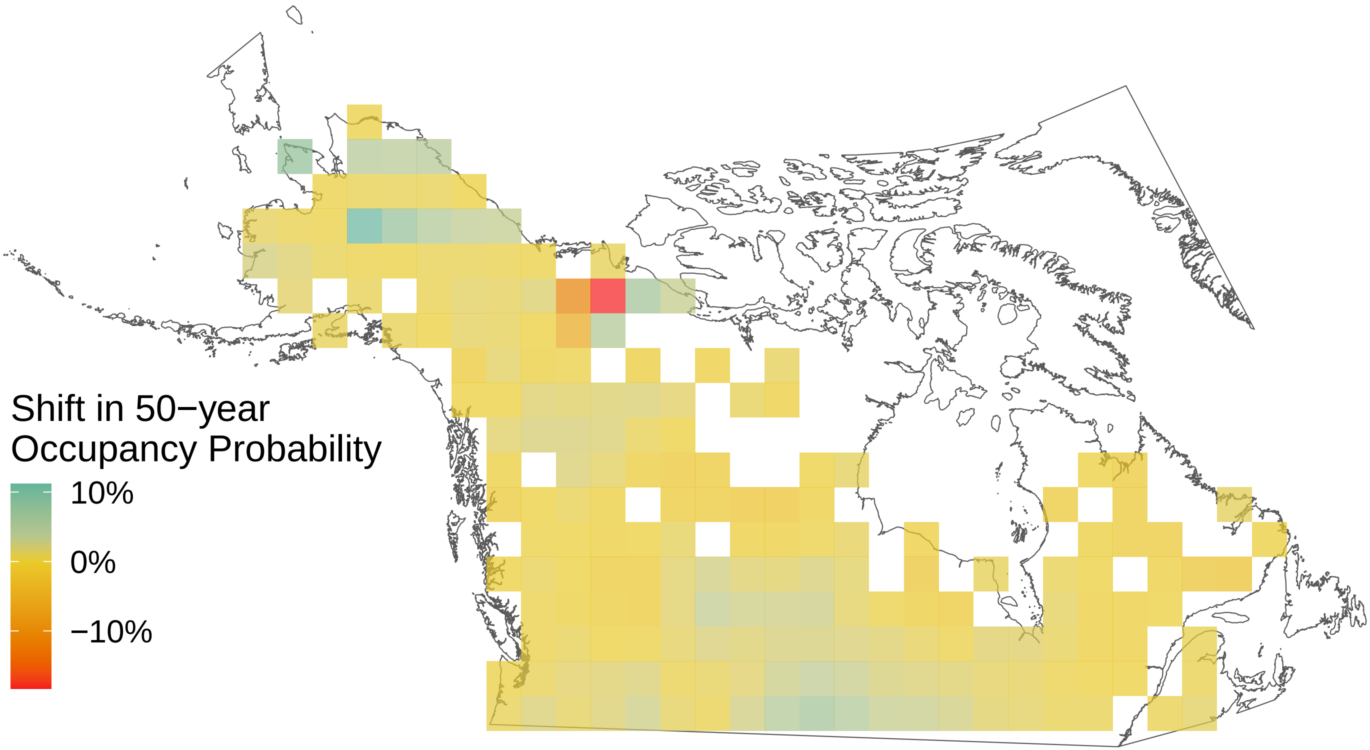

*Neophasia menapia*

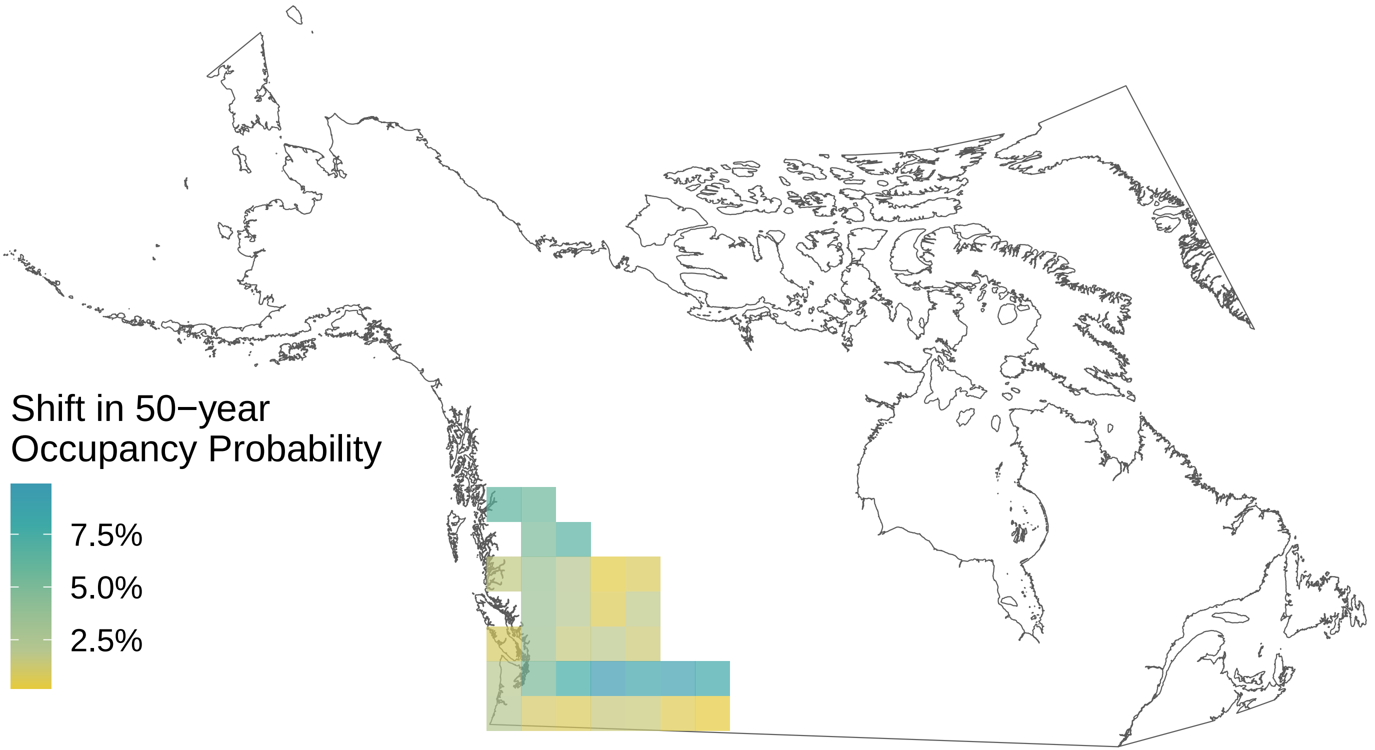

*Lycaena phlaeas*

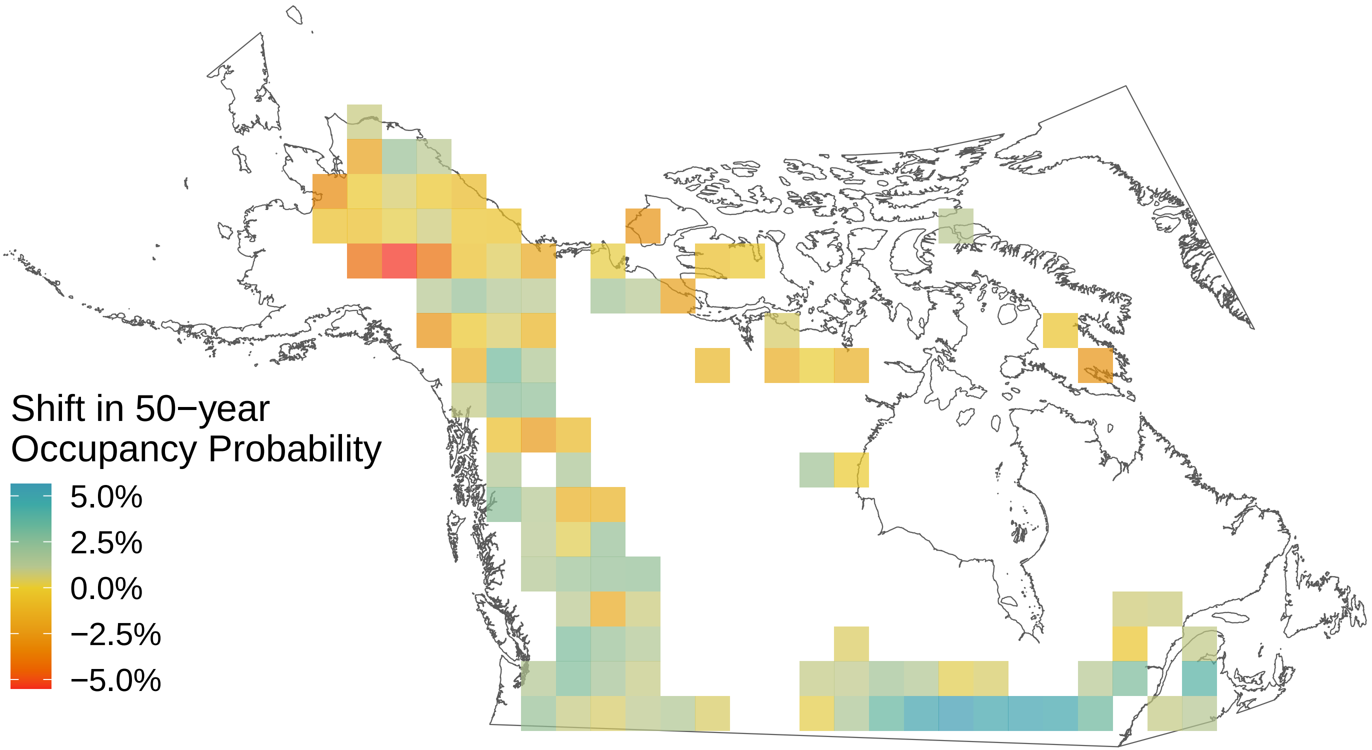

*Lycaena dorcas*

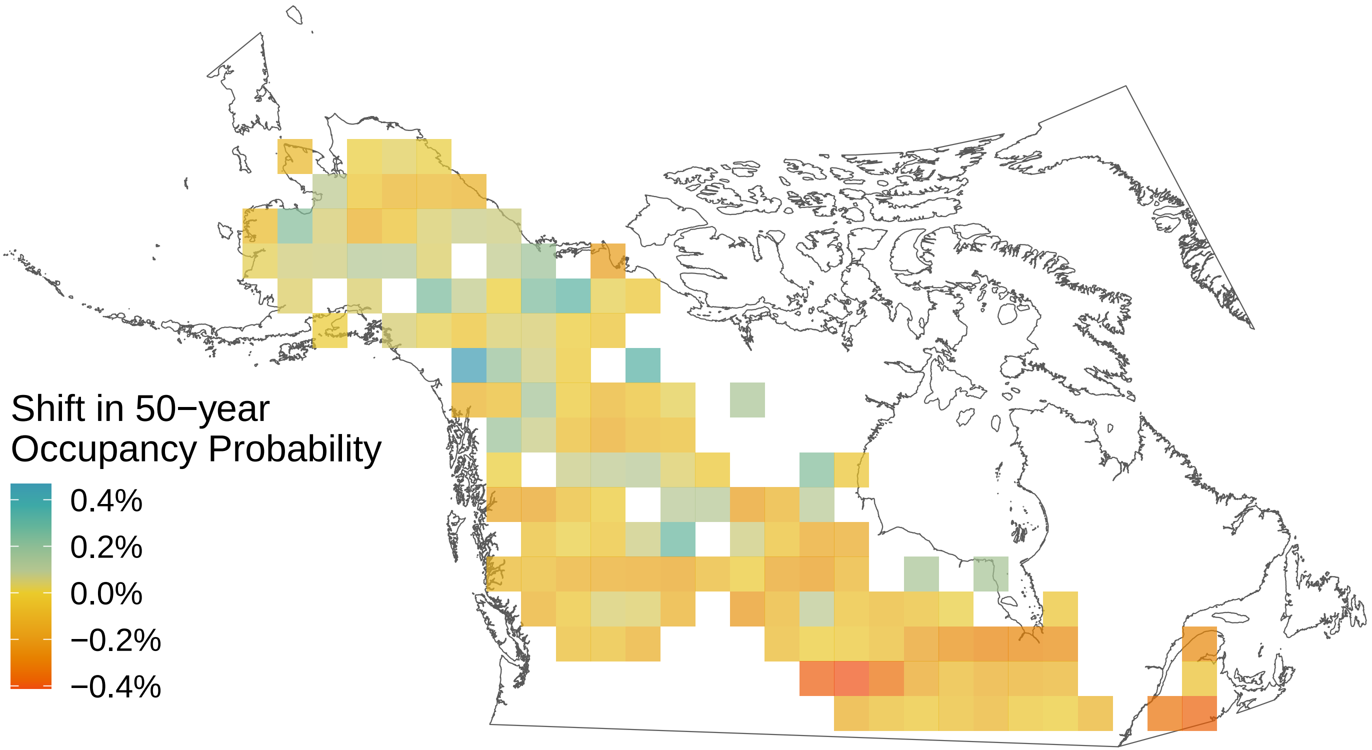

*Limenitis lorquini*

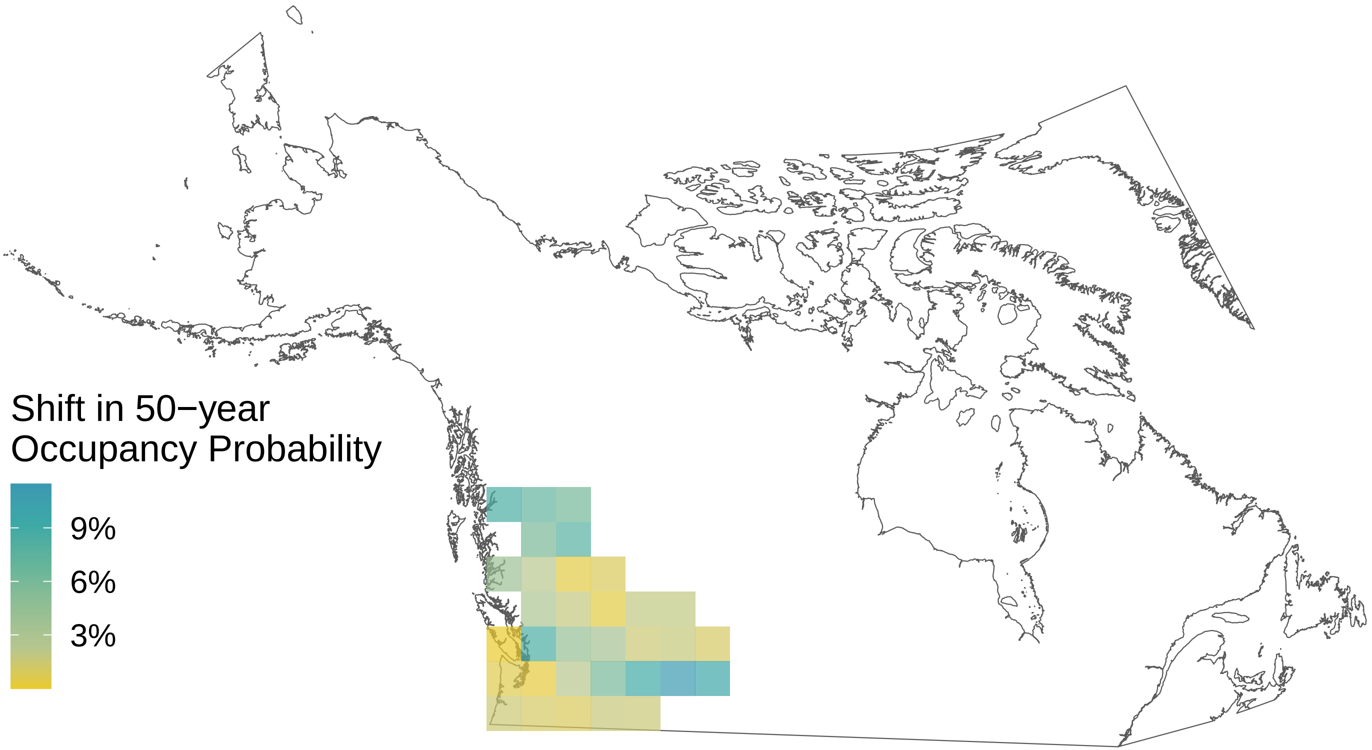

*Limenitis arthemis*

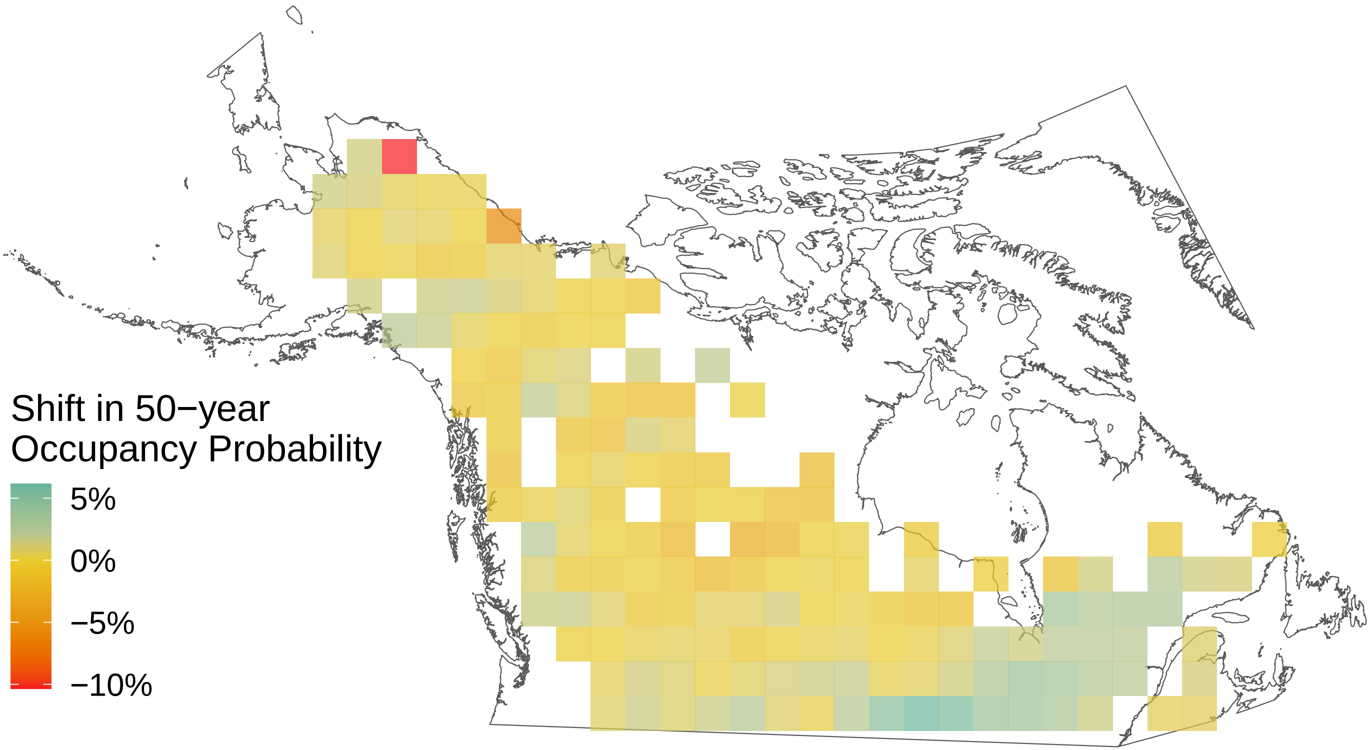

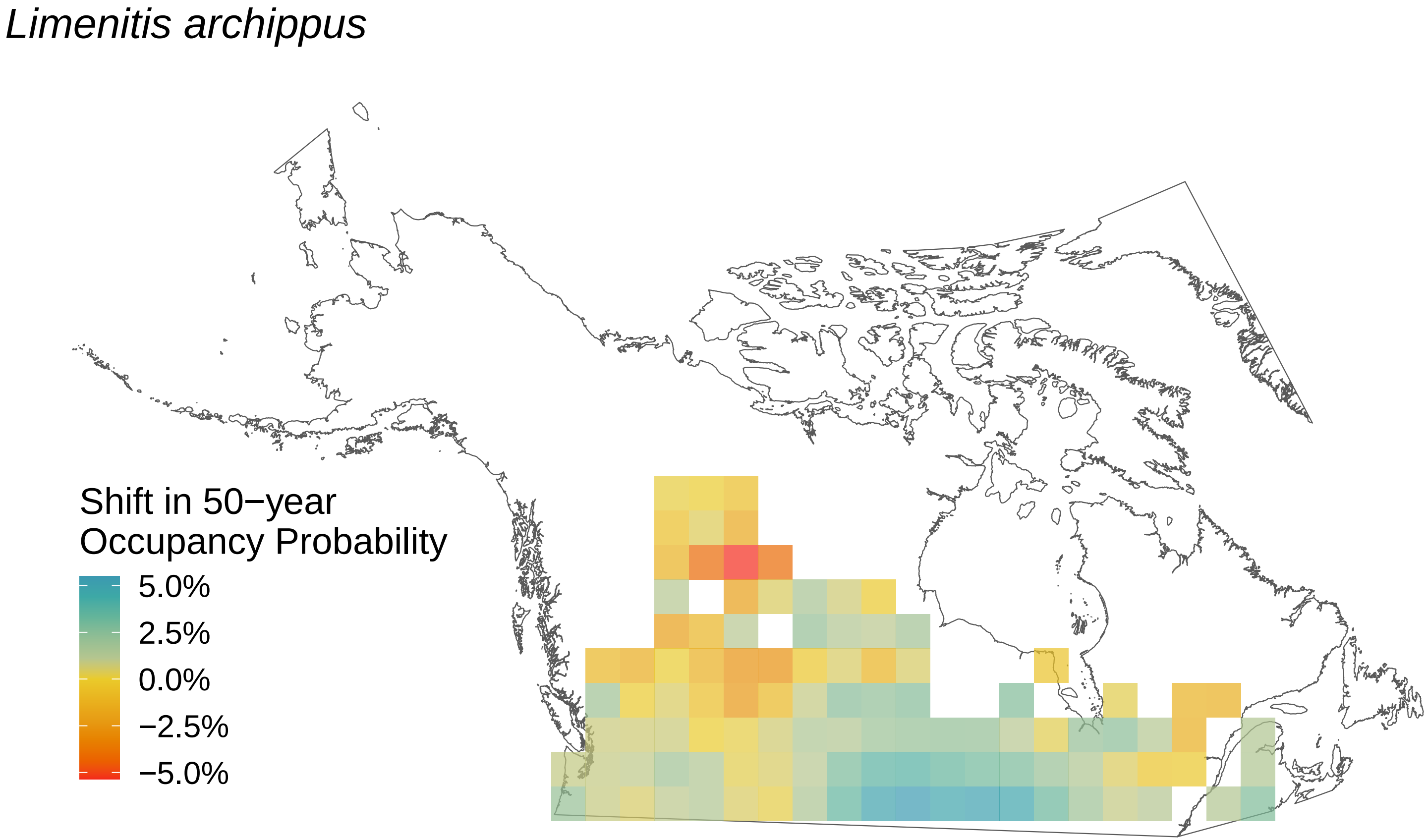
Limenitis archippus
Shift in 50−year
Occupancy Probability
5.0%
2.5%
0.0%
−2.5%
−5.0%

*Lethe eurydice*

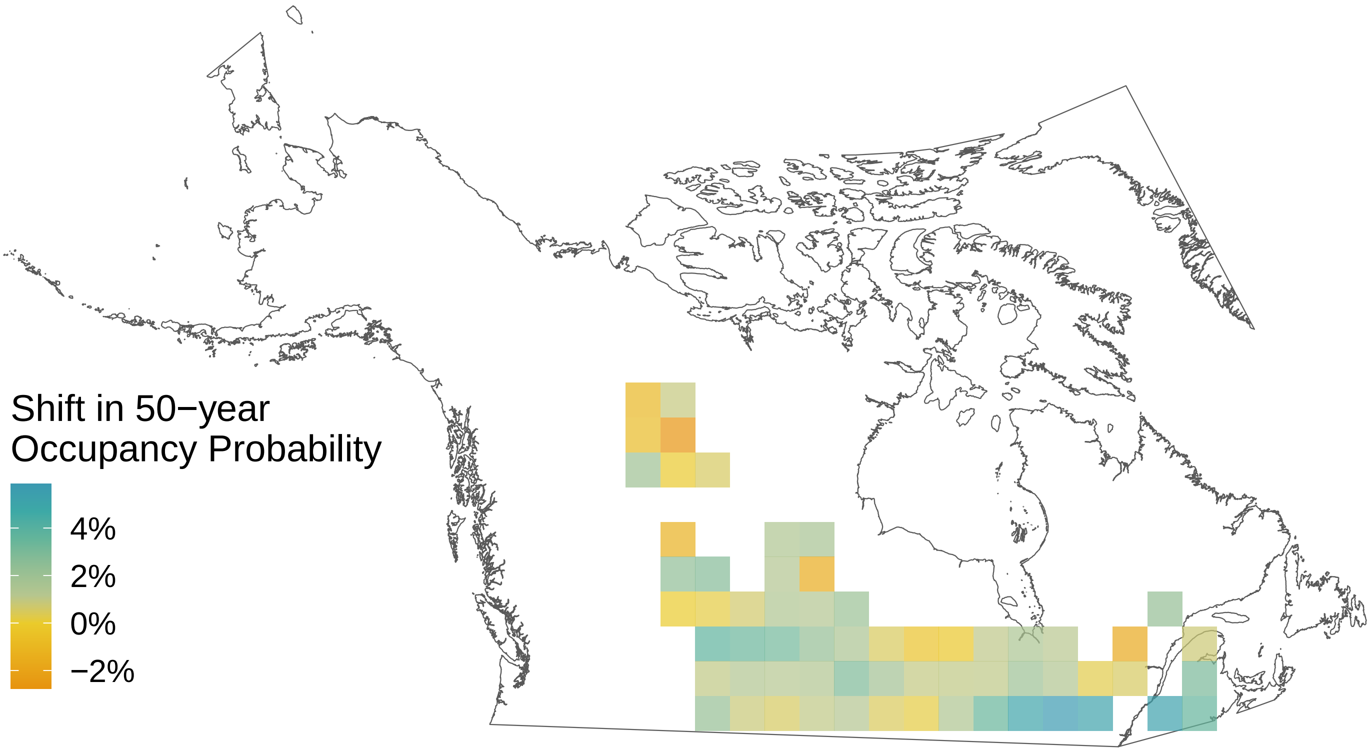

### *Icaricia saepiolus*

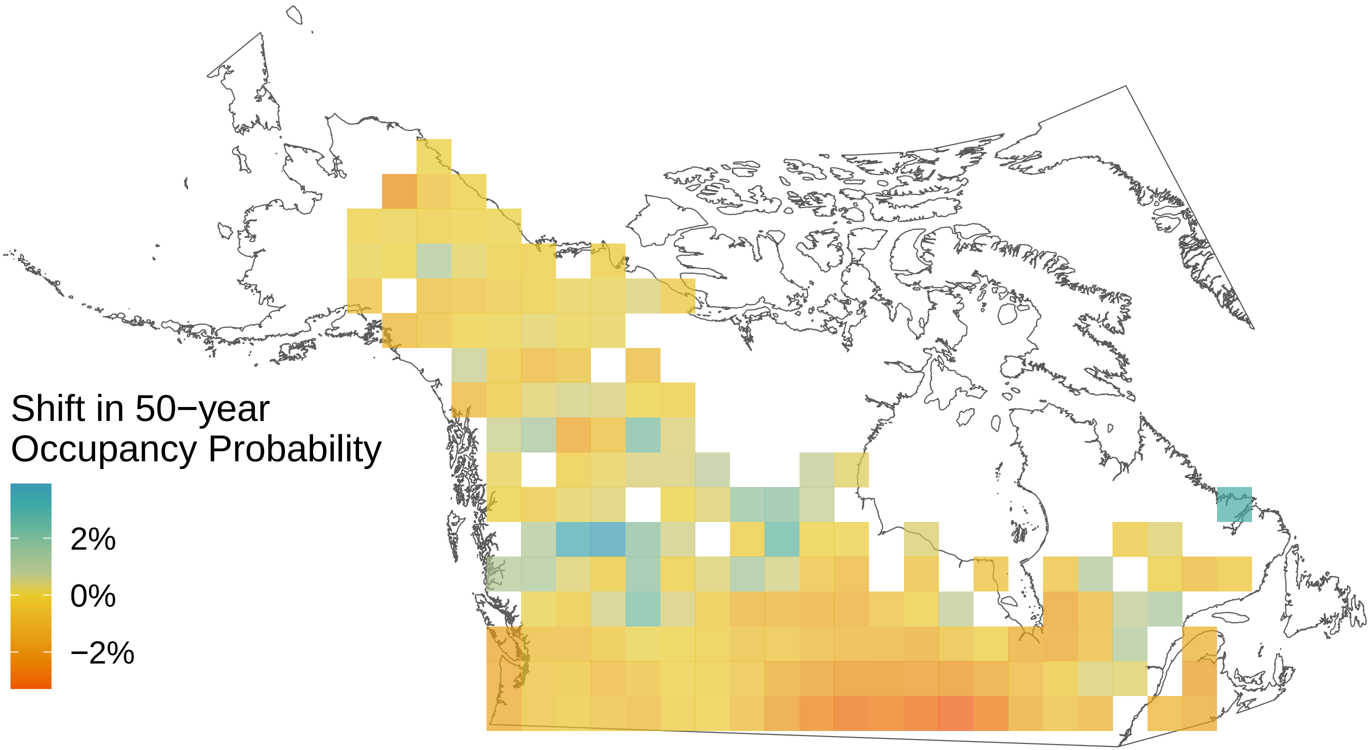

*Icaricia icarioides*

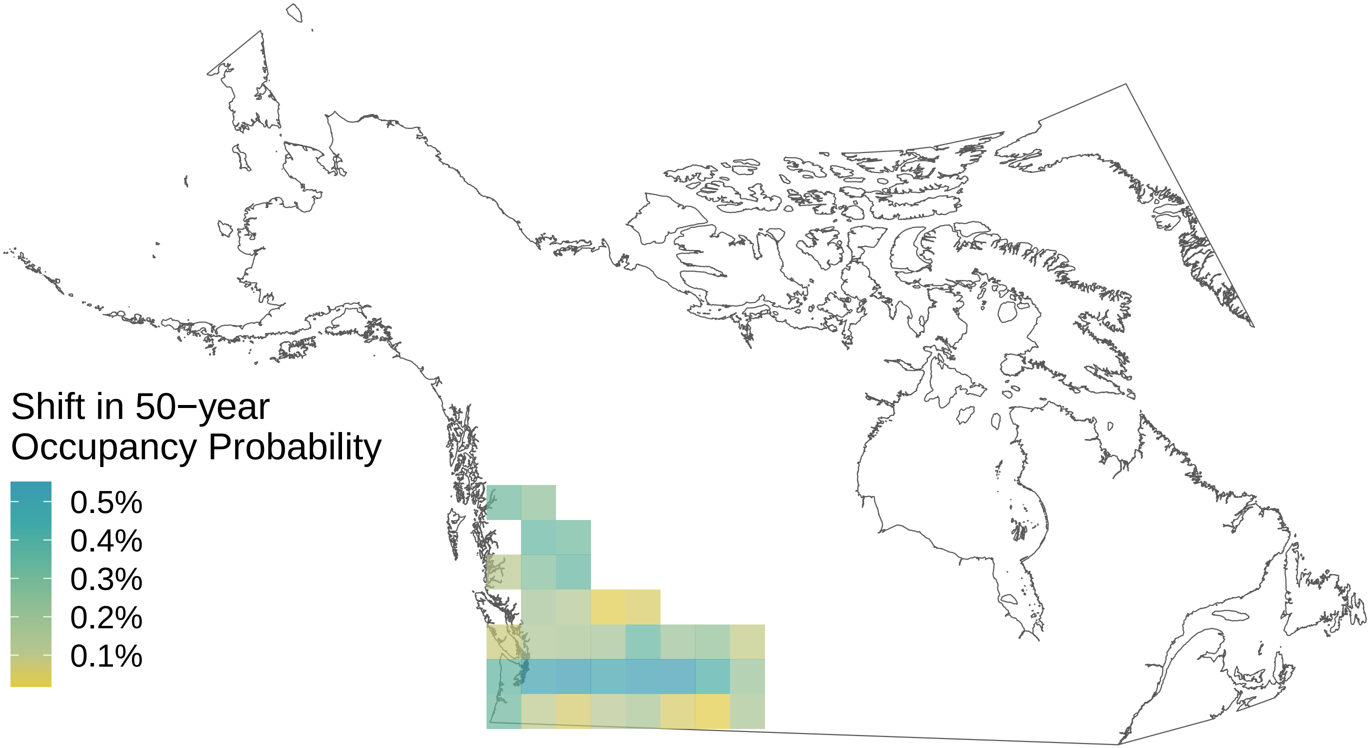

*Hesperia comma*

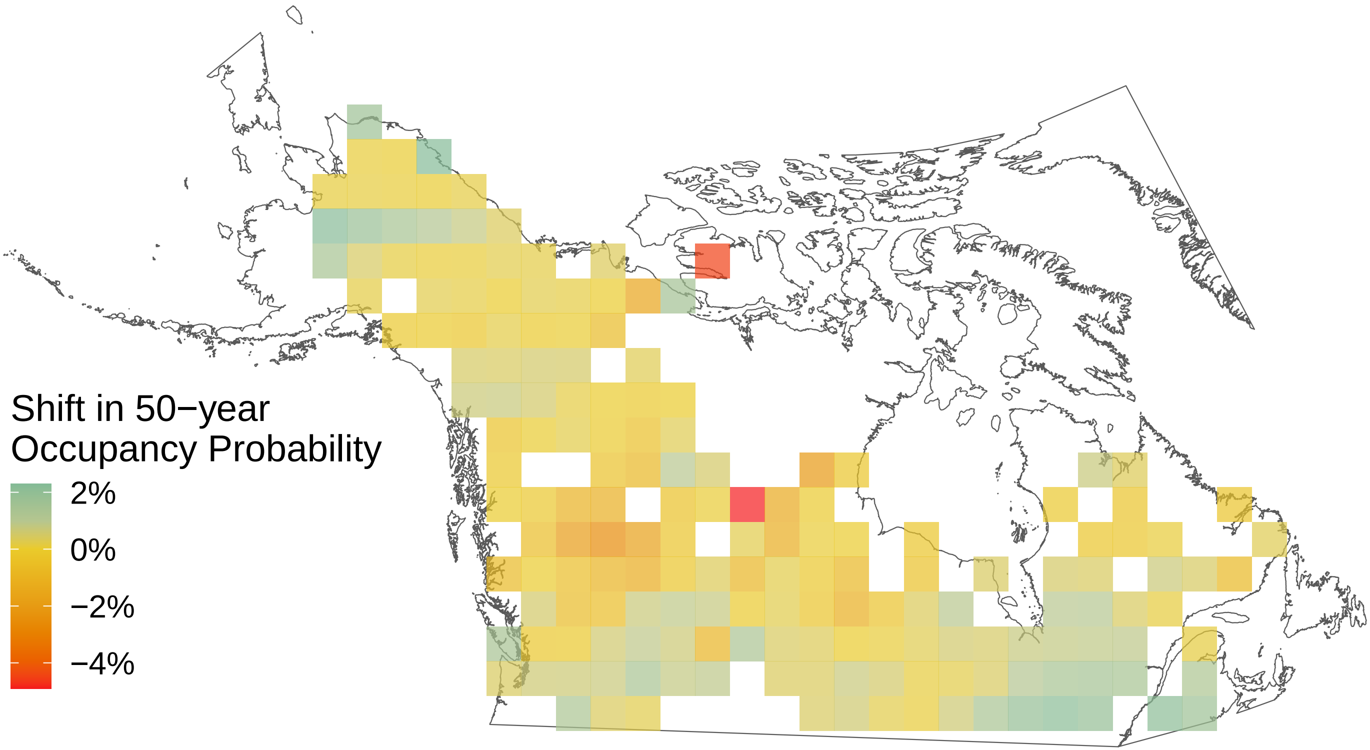

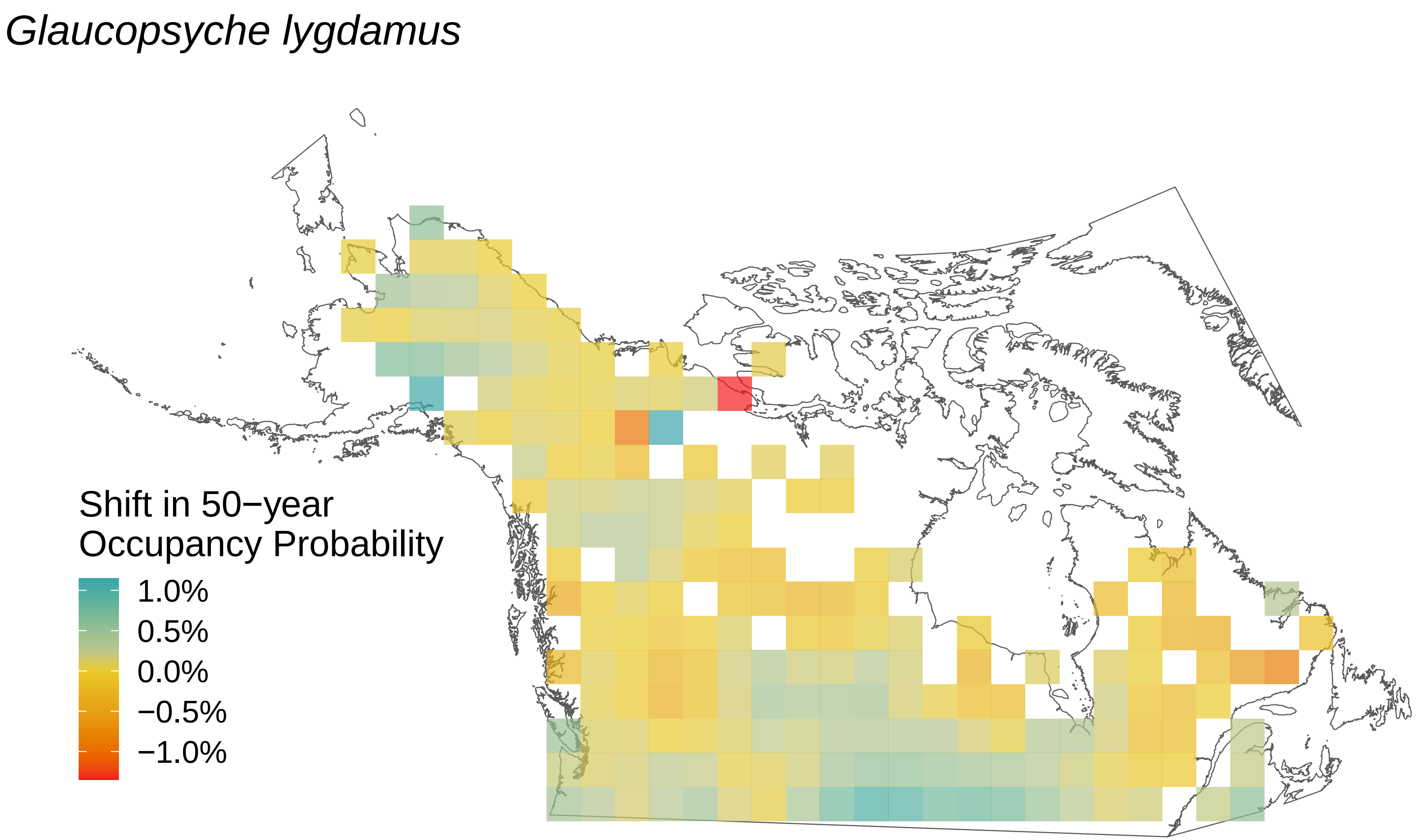
Glaucopsyche lygdamus
Shift in 50−year
Occupancy Probability
1.0%
0.5%
0.0%
−0.5%
−1.0%

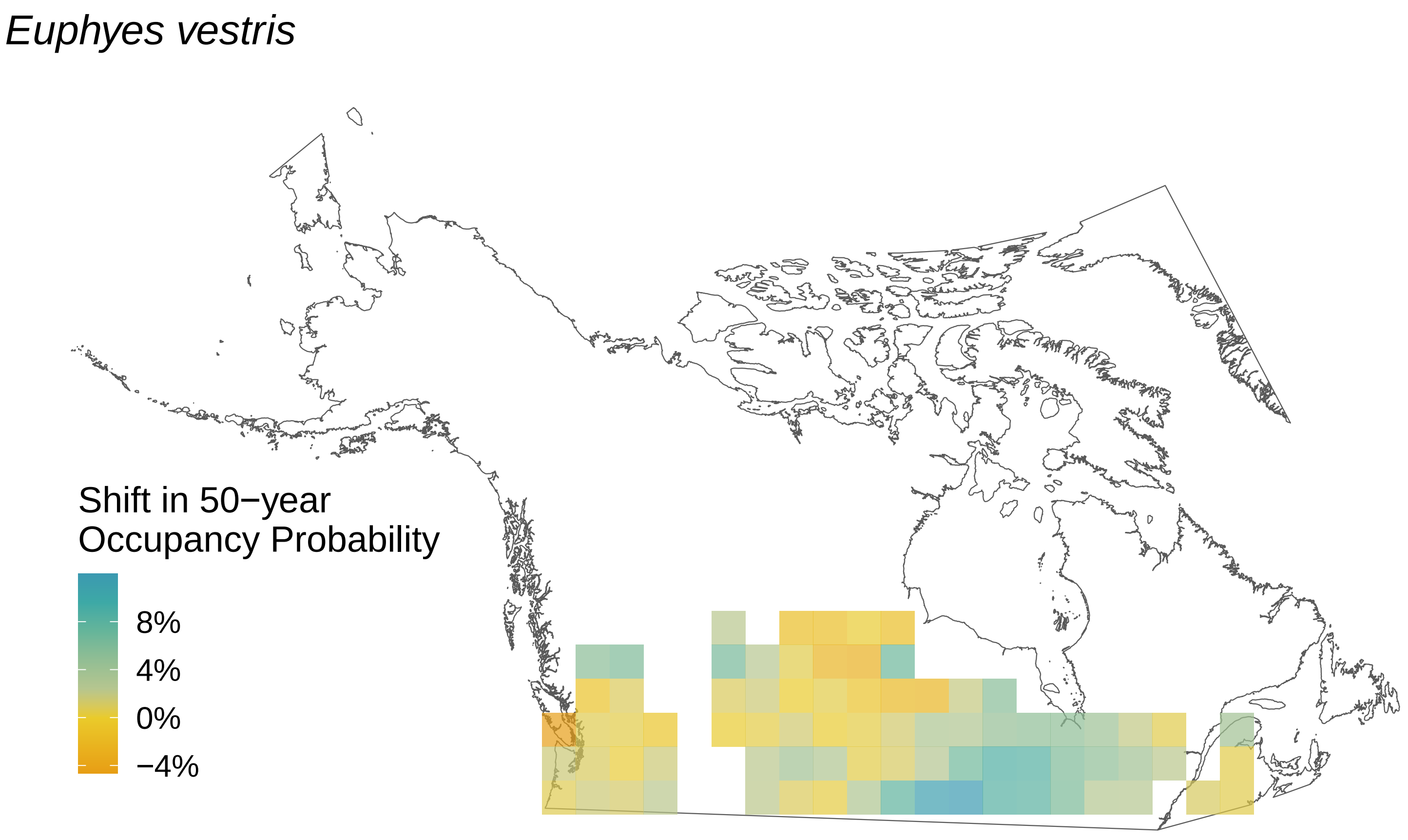
Euphyes vestris
Shift in 50−year Occupancy Probability
8%
4%
0%
−4%

*Euphydryas phaeton*

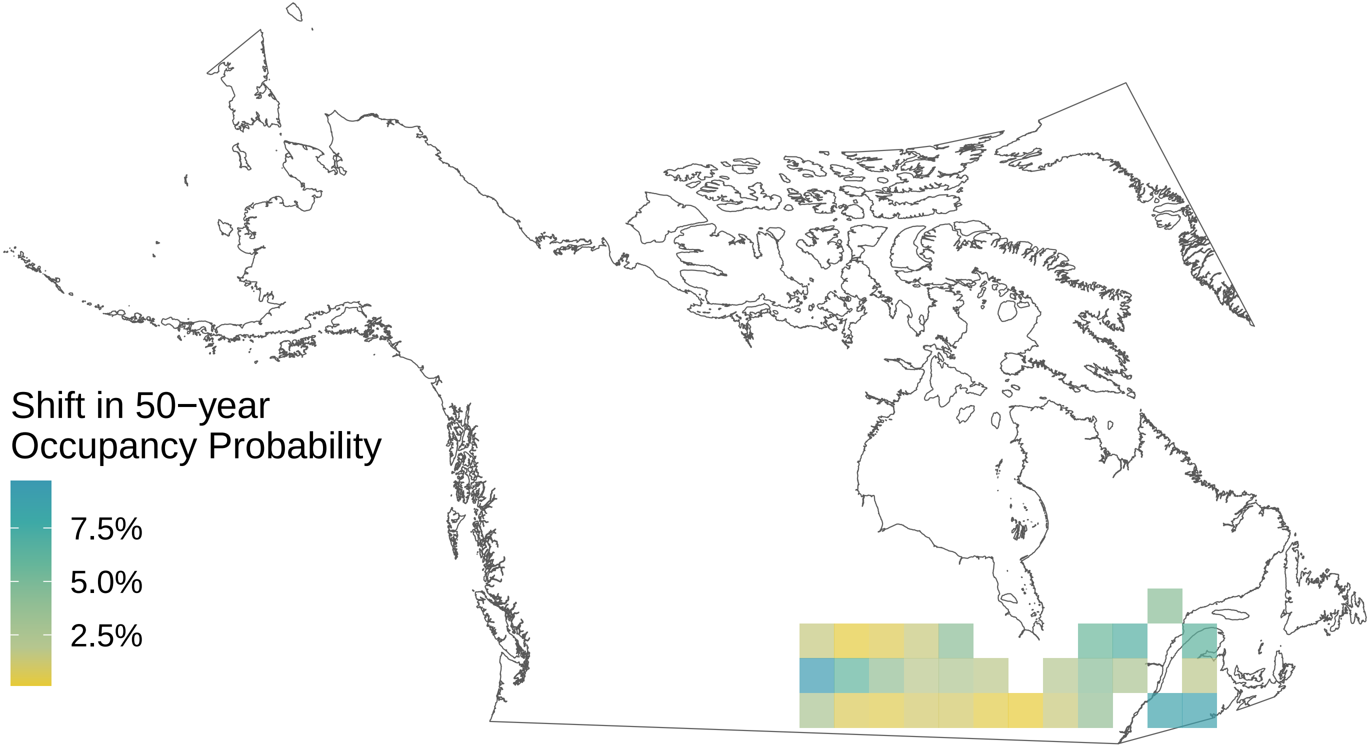

*Euphydryas anicia*

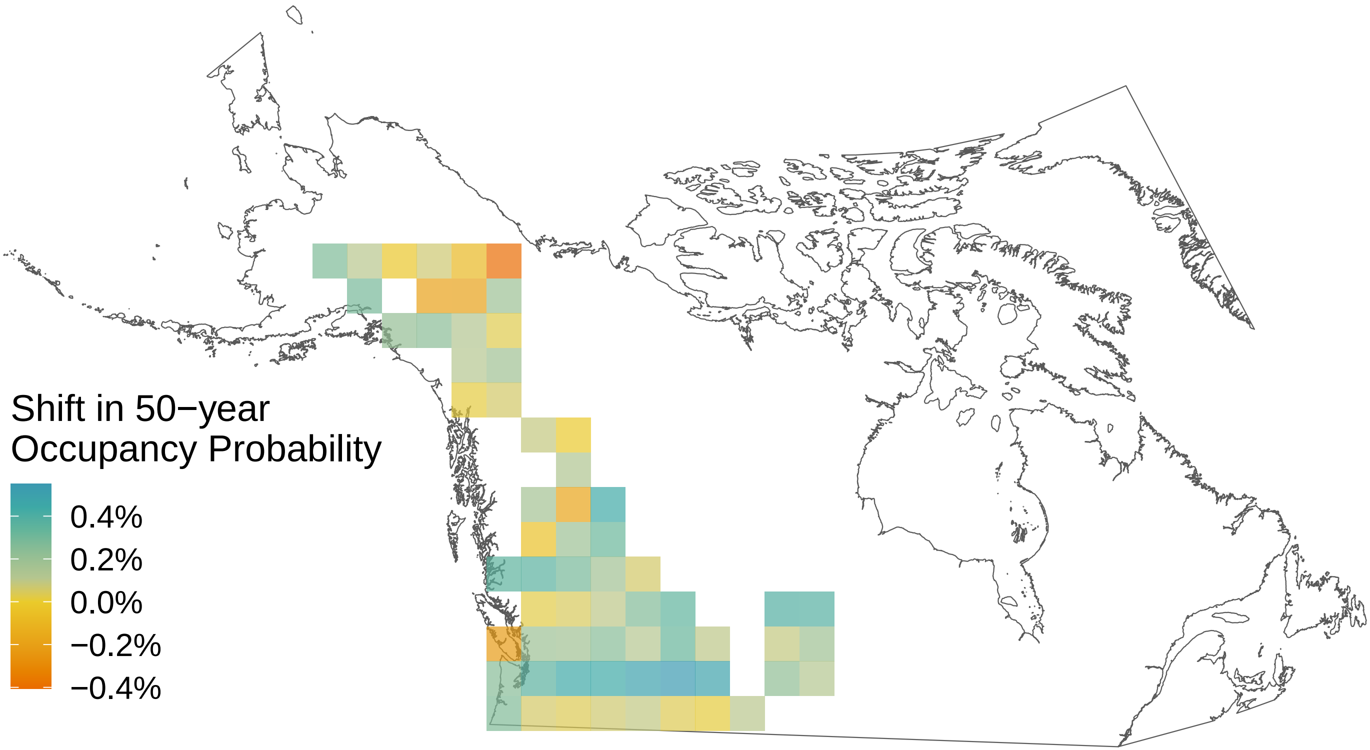

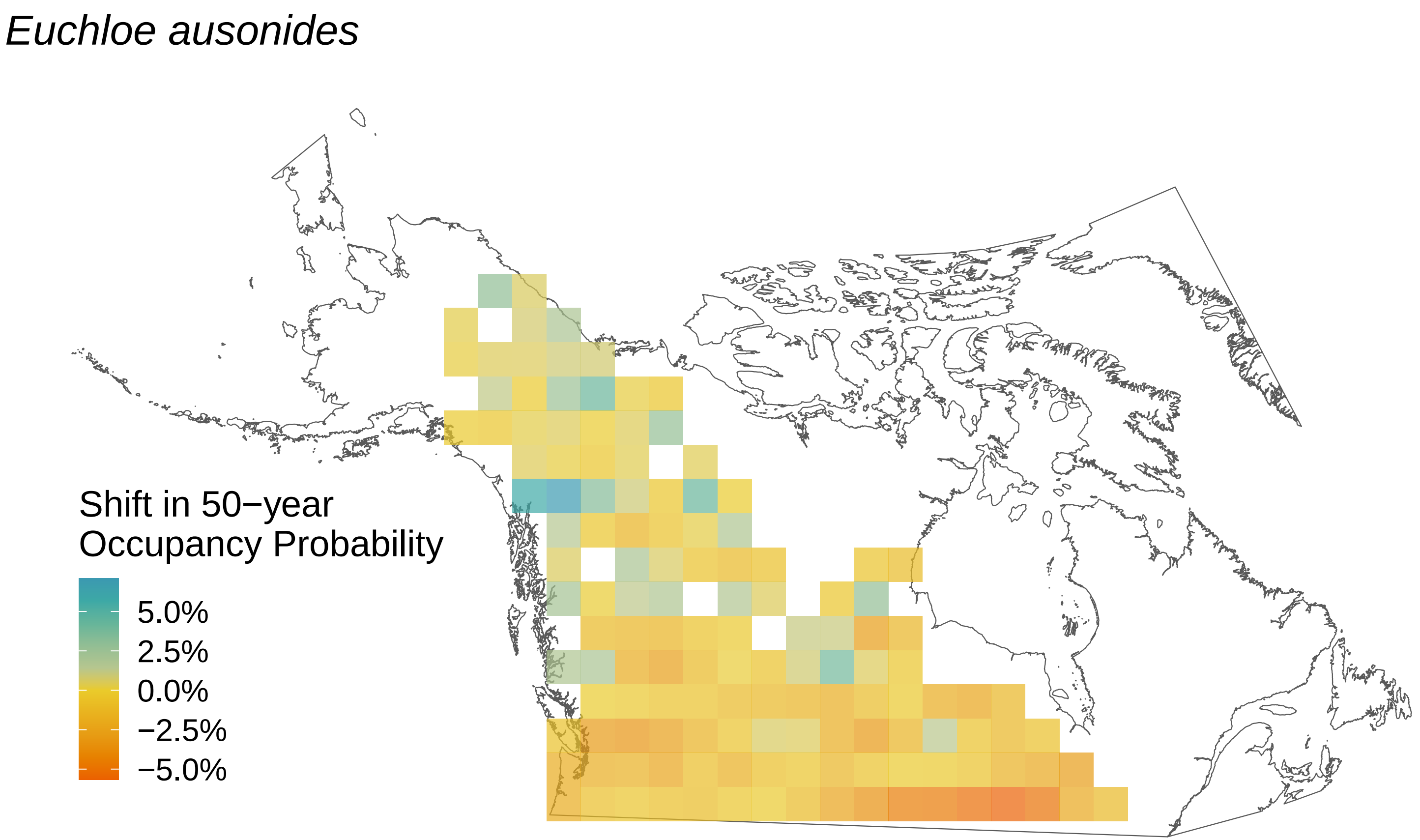
Euchloe ausonides
Shift in 50−year Occupancy Probability
5.0%
2.5%
0.0%
−2.5%
−5.0%

*Erynnis persius*

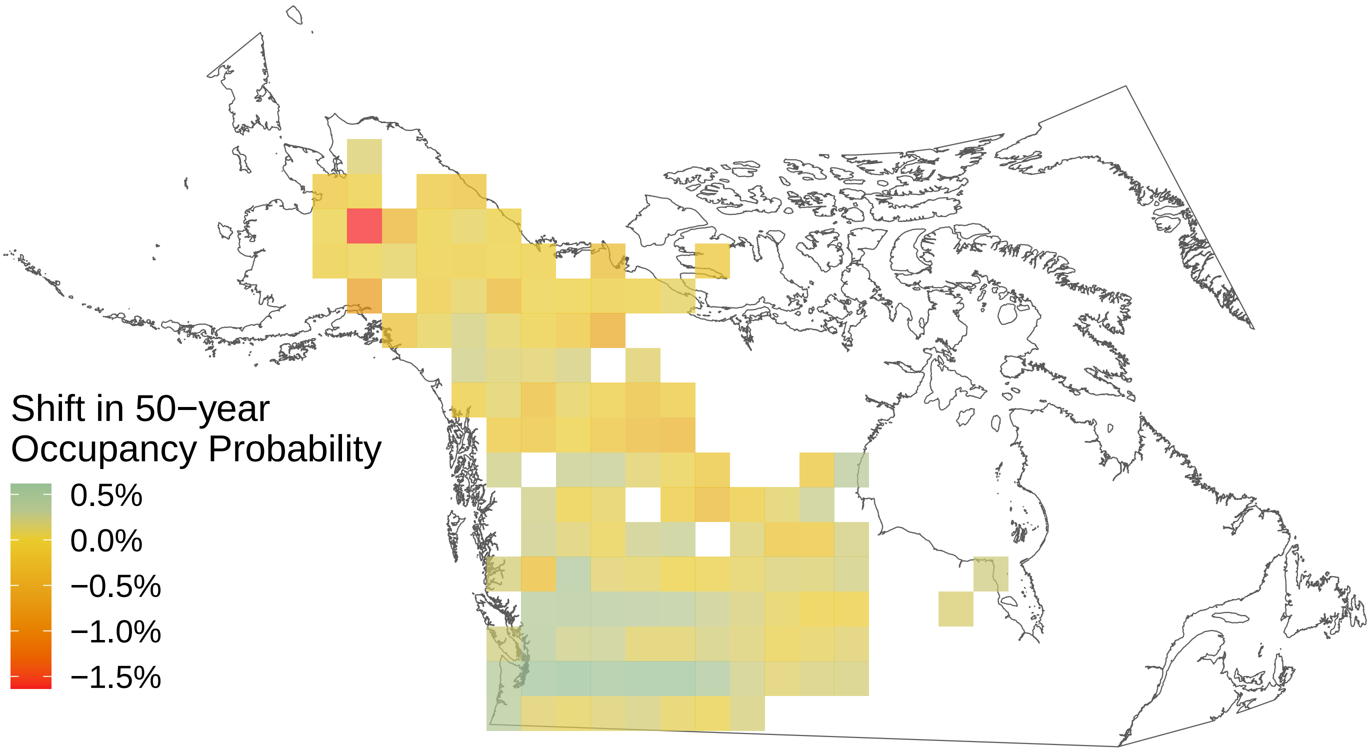

*Erynnis juvenalis*

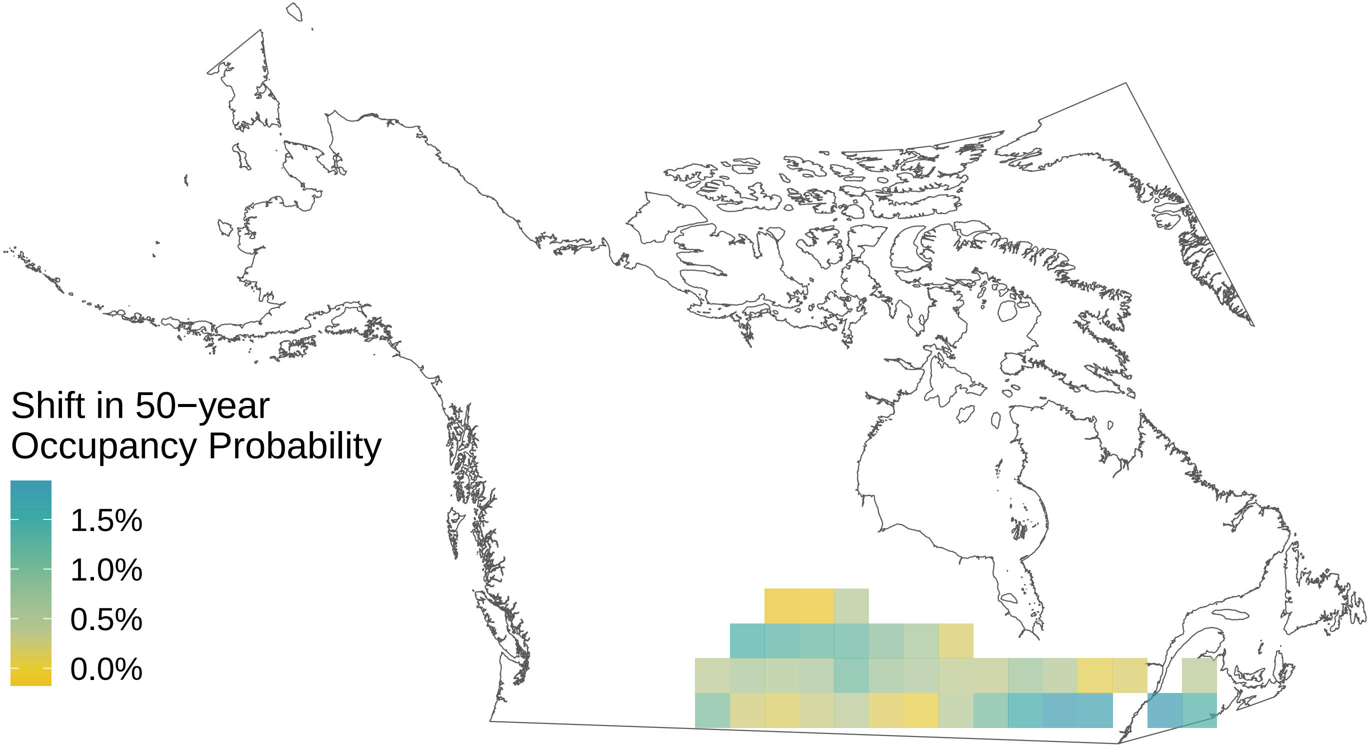

*Erynnis icelus*

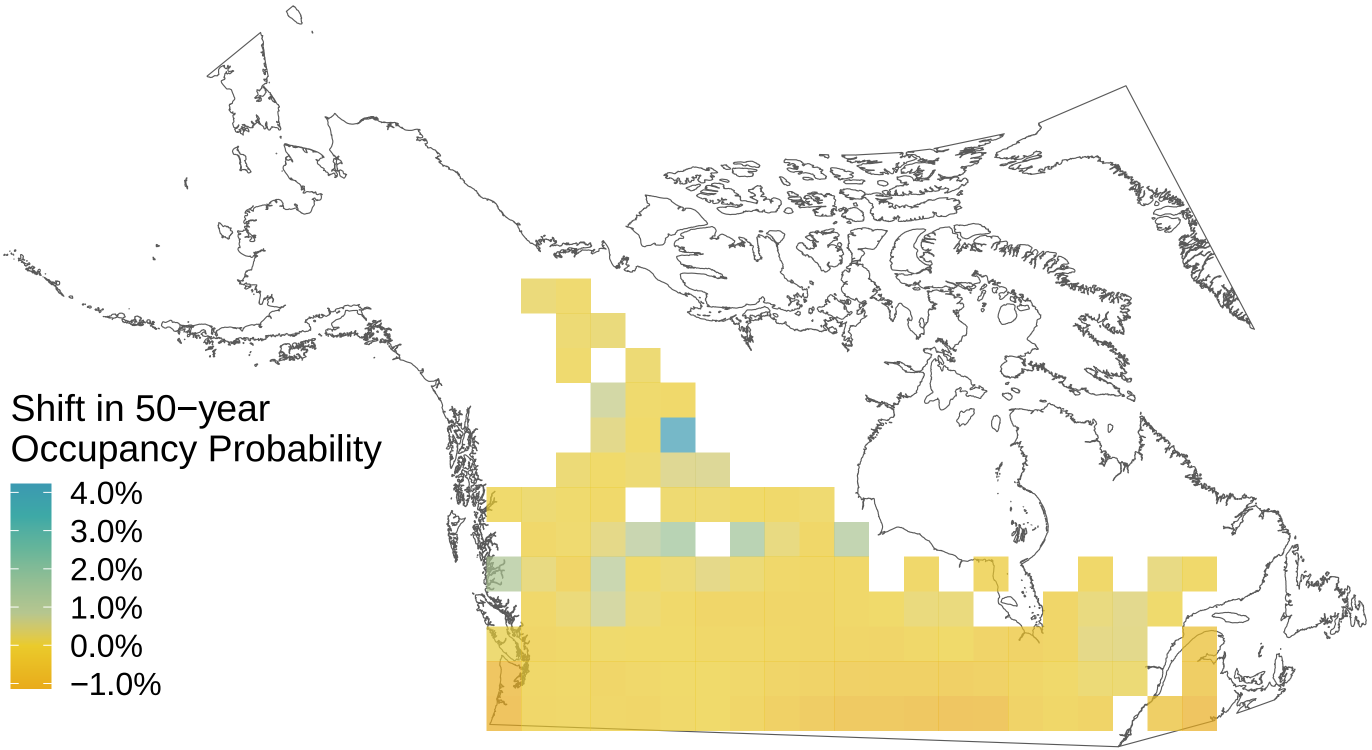

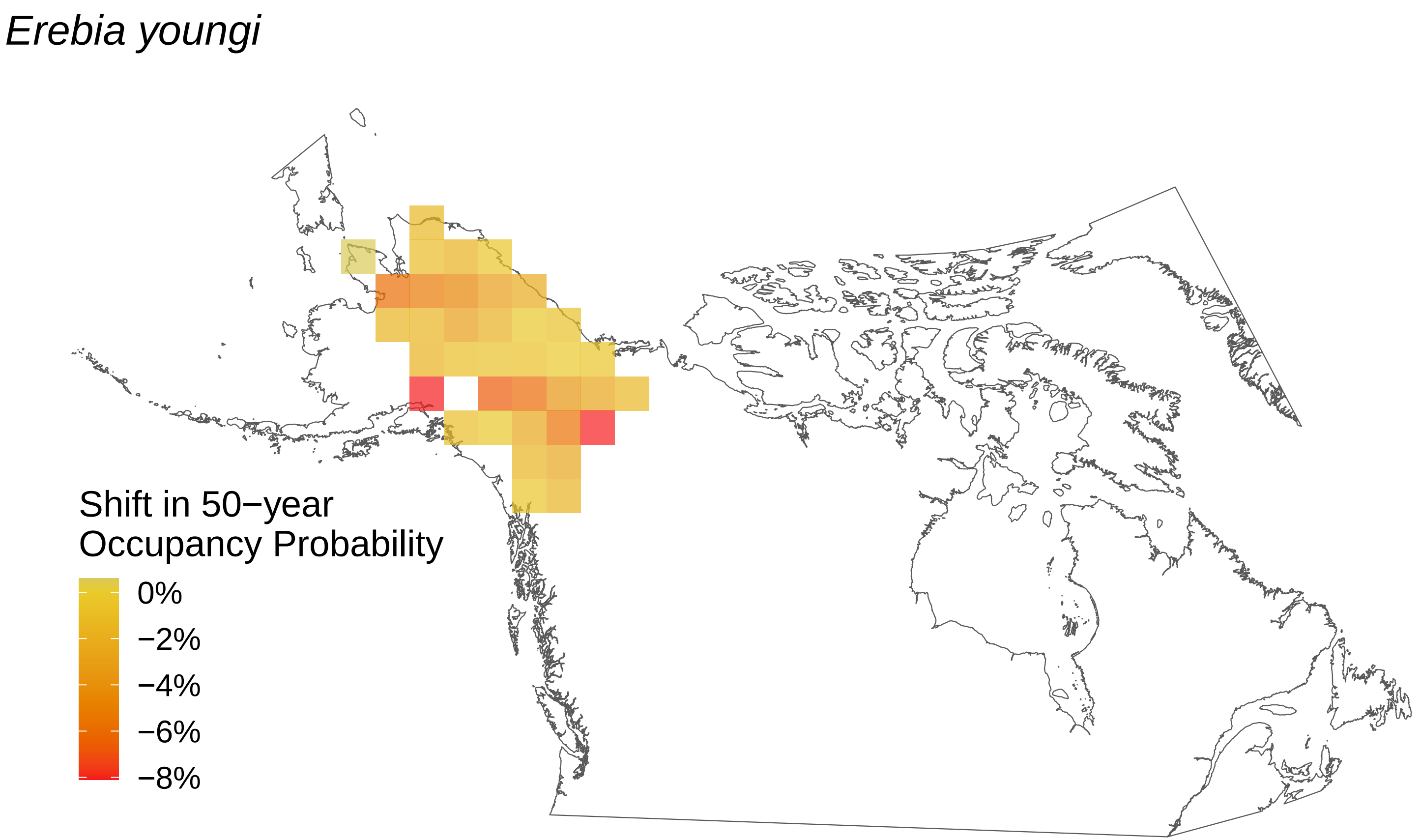
Erebia youngi
Shift in 50−year
Occupancy Probability
0%
−2%
−4%
−6%
−8%

*Erebia rossii*

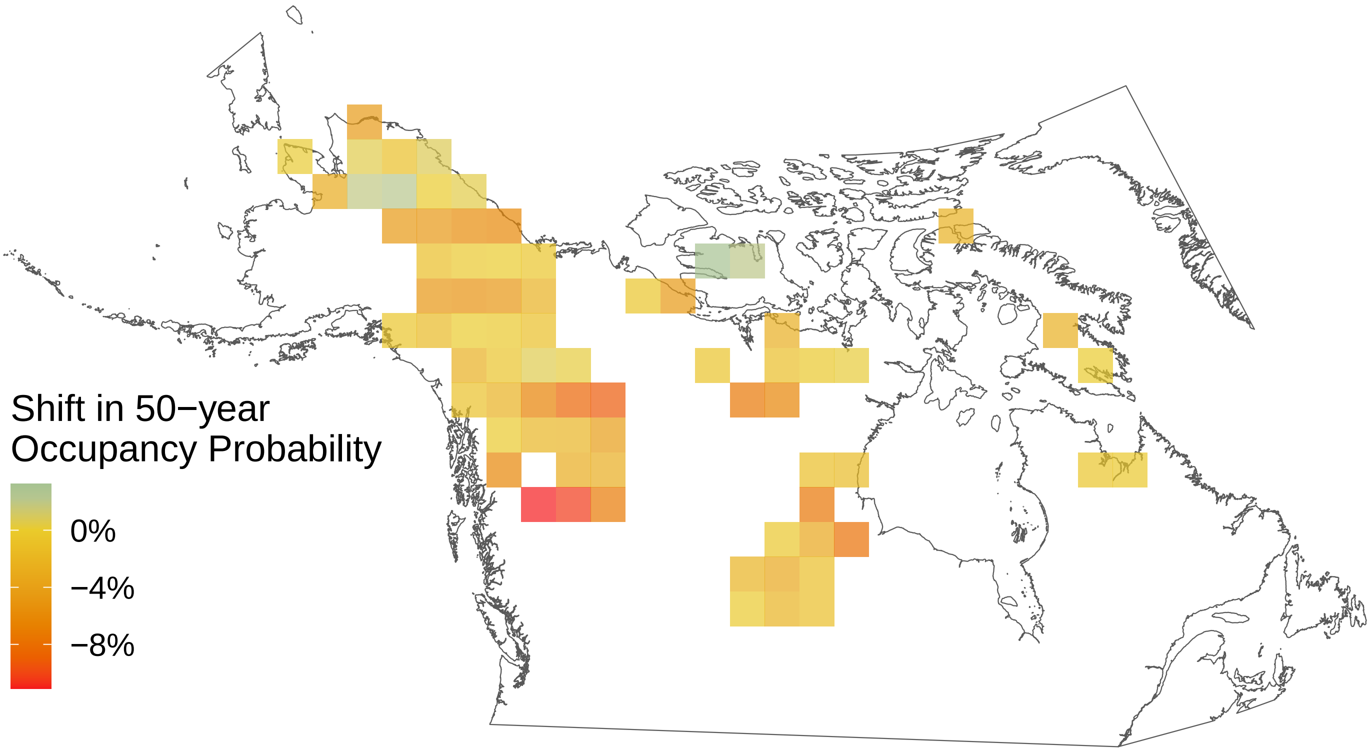

*Erebia fasciata*

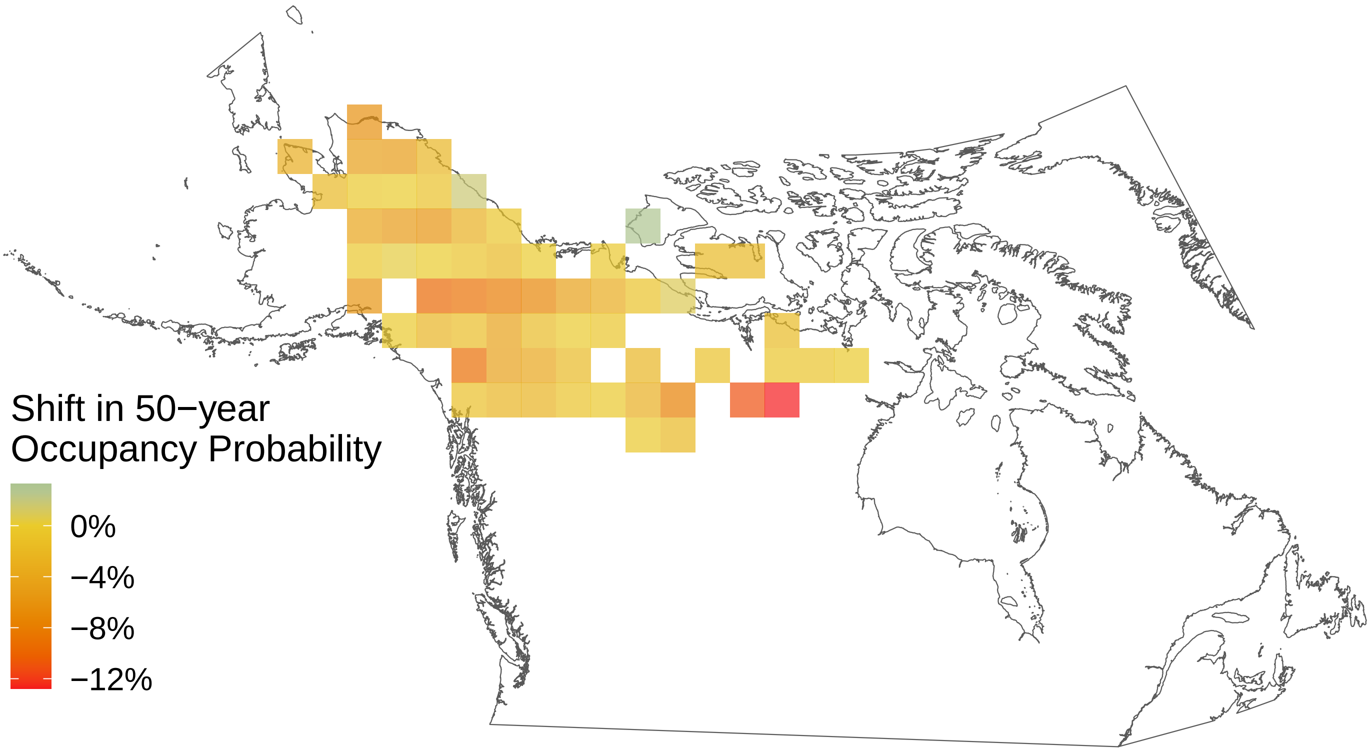

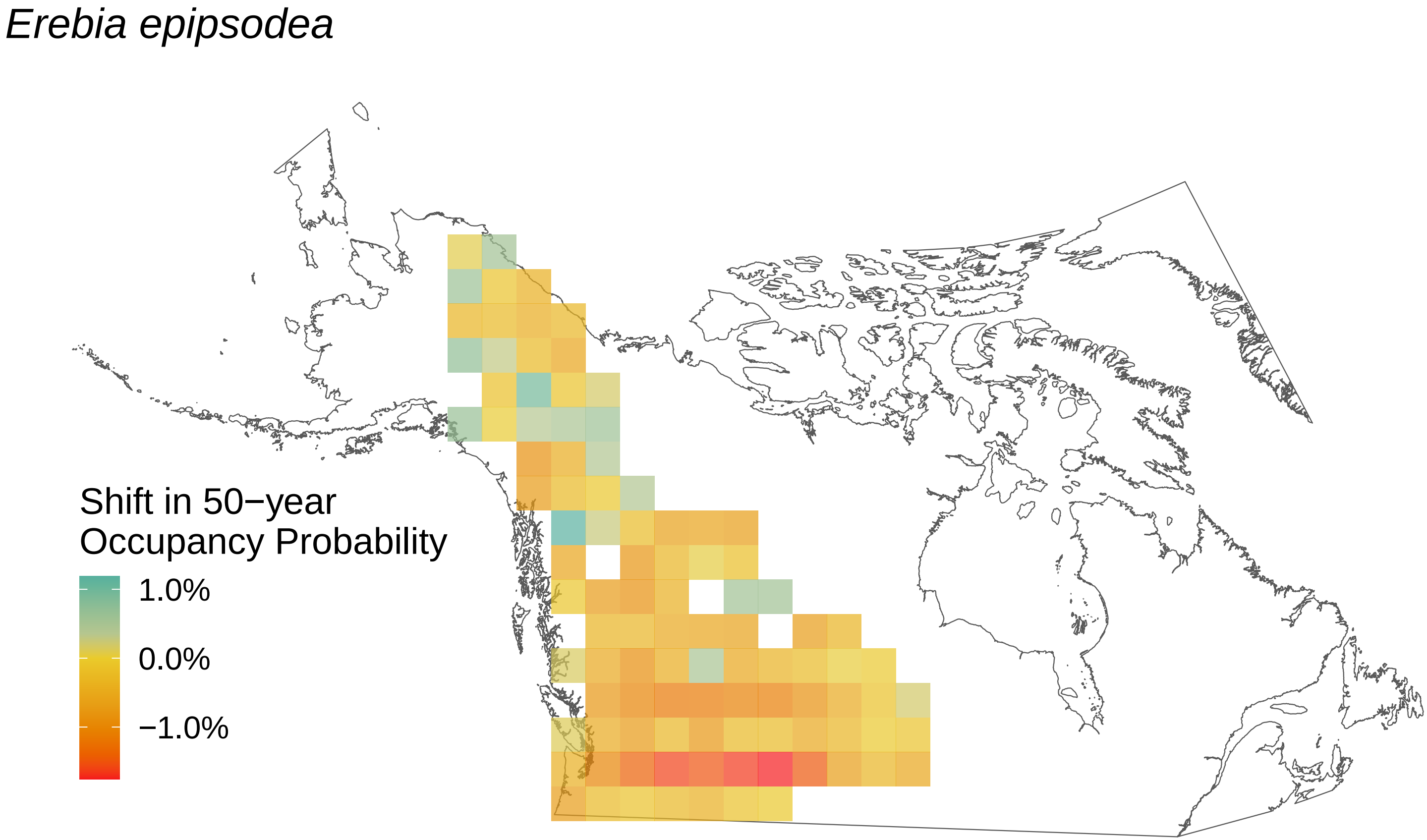
Erebia epipsodea
Shift in 50−year Occupancy Probability
1.0%
0.0%
−1.0%

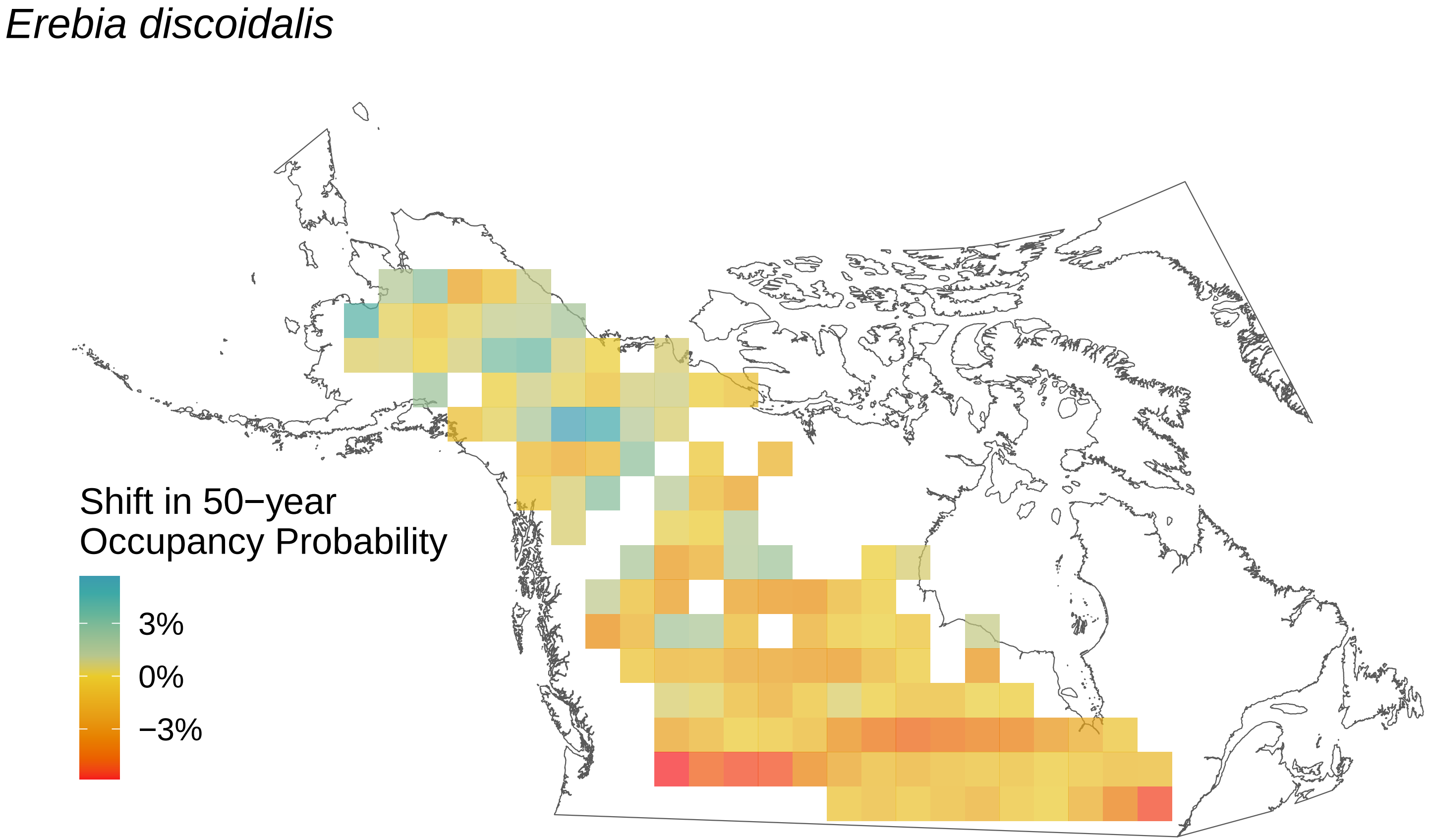
Erebia discoidalis
Shift in 50−year Occupancy Probability
3%
0%
−3%

*Epargyreus clarus*

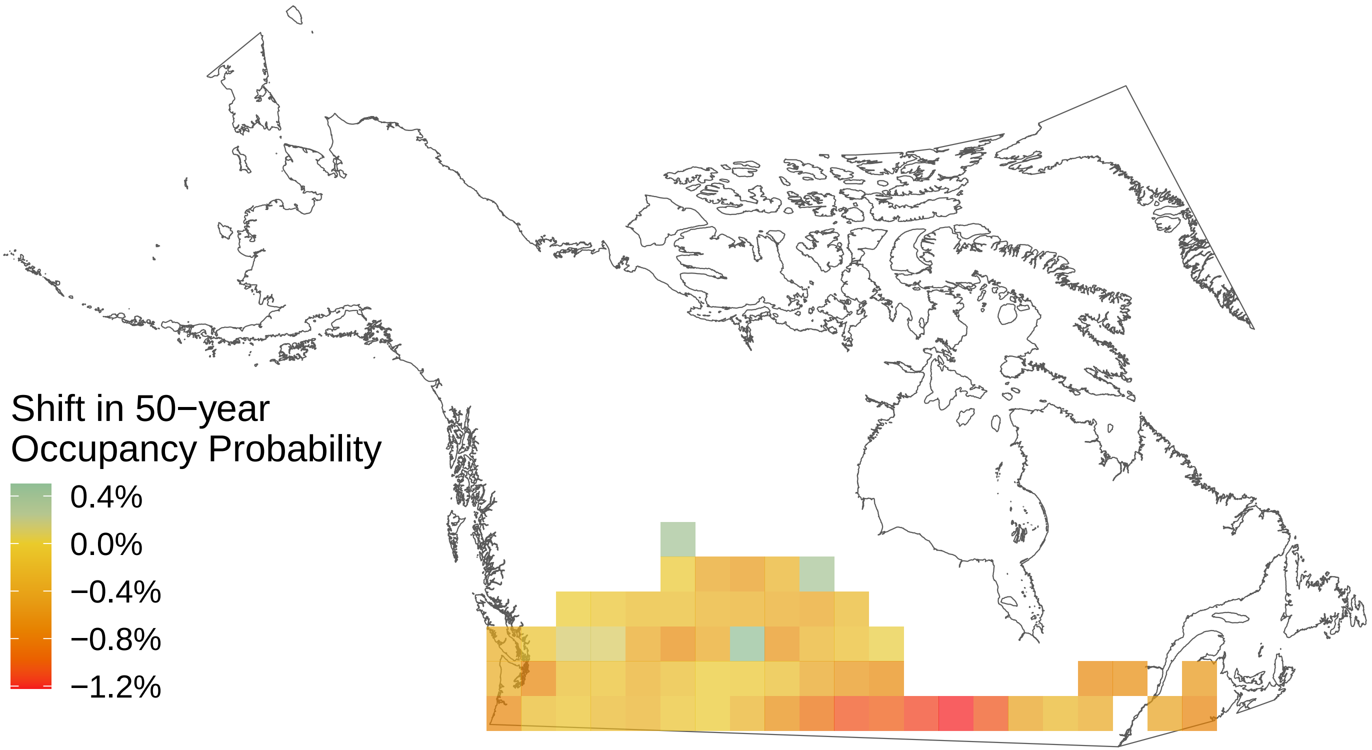

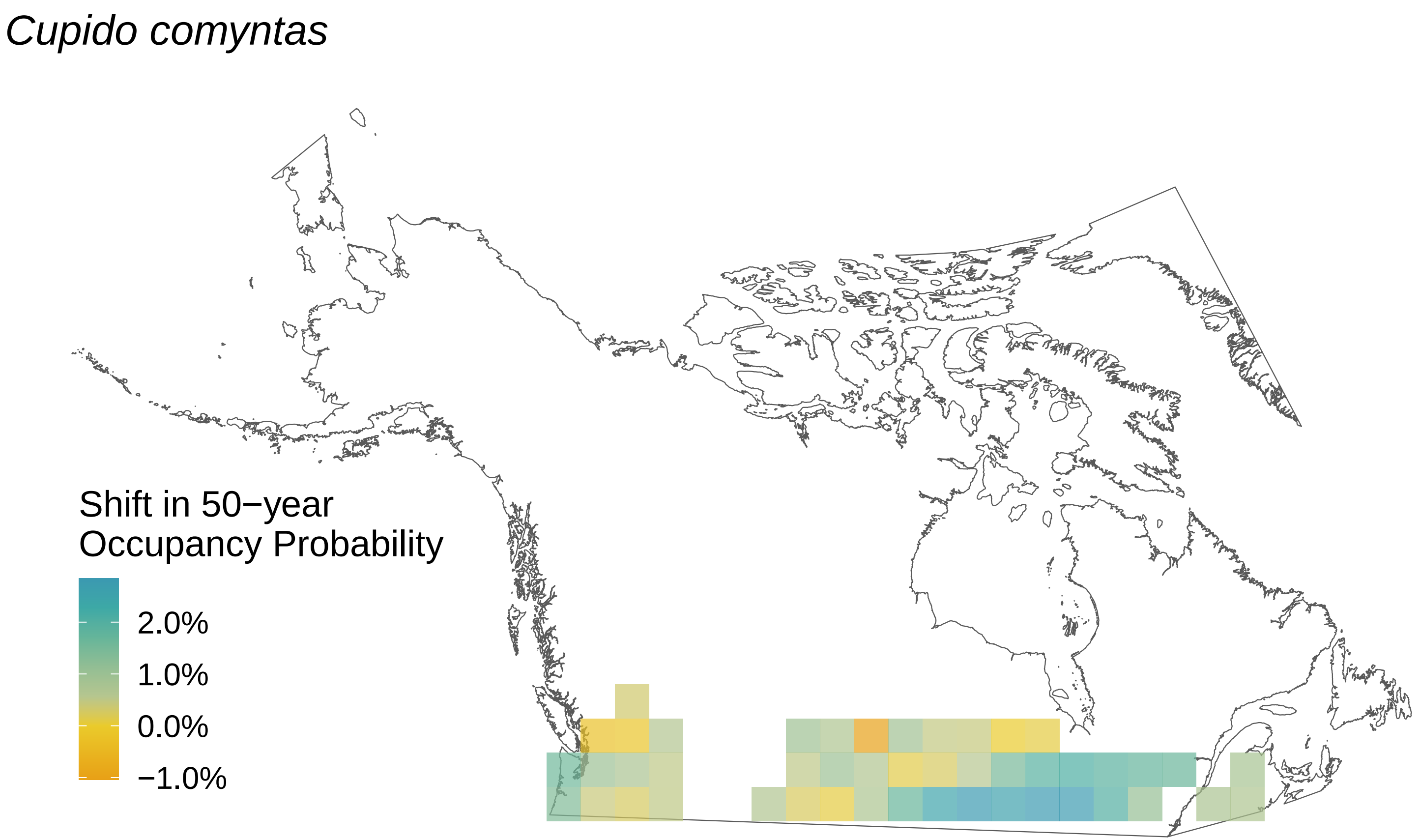
Cupido comyntas
Shift in 50−year
Occupancy Probability
2.0%
1.0%
0.0%
−1.0%

*Cupido amyntula*

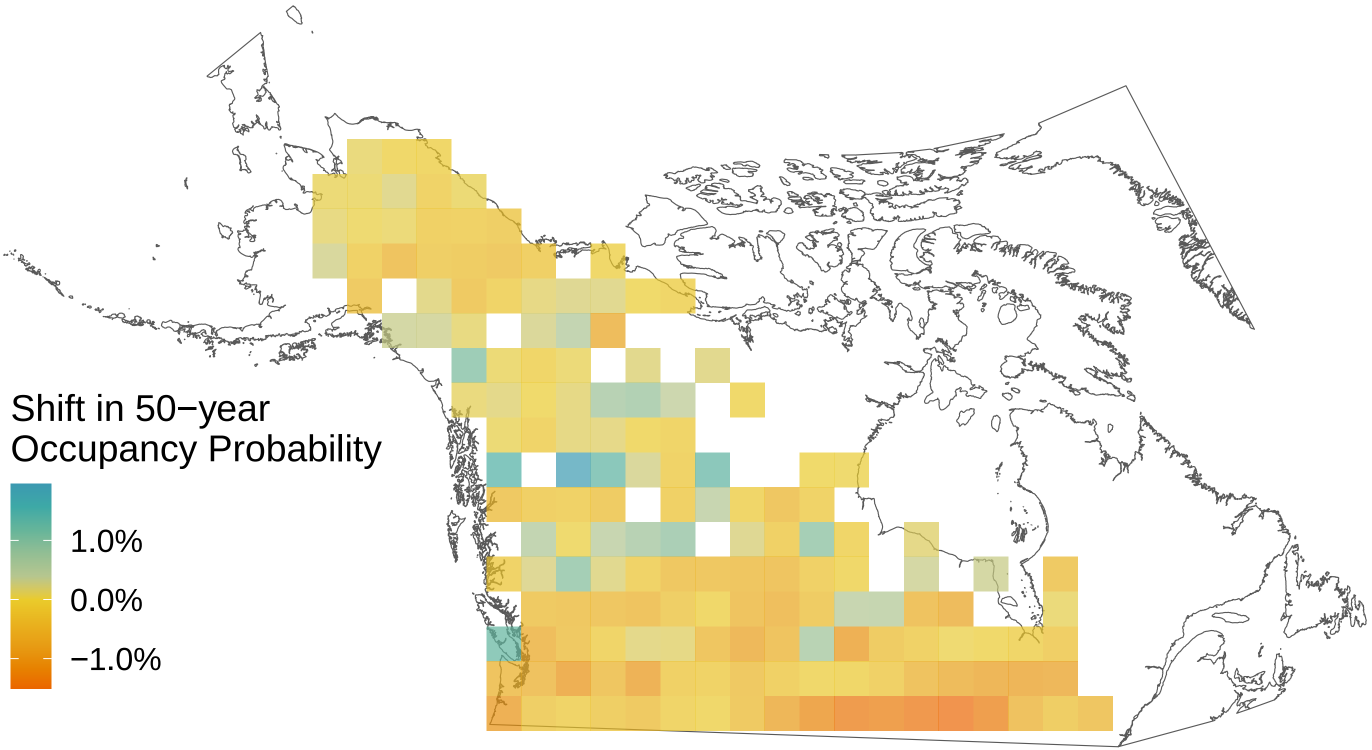

### *Colias philodice*

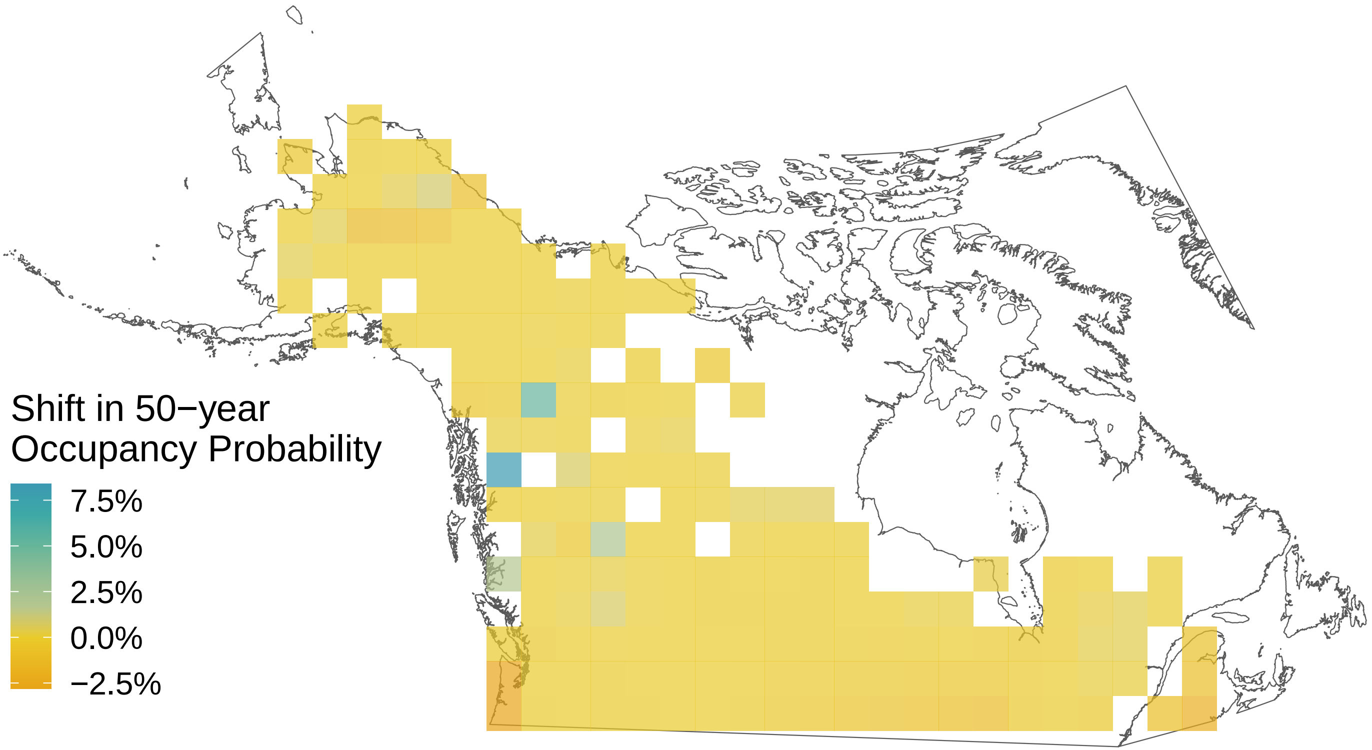

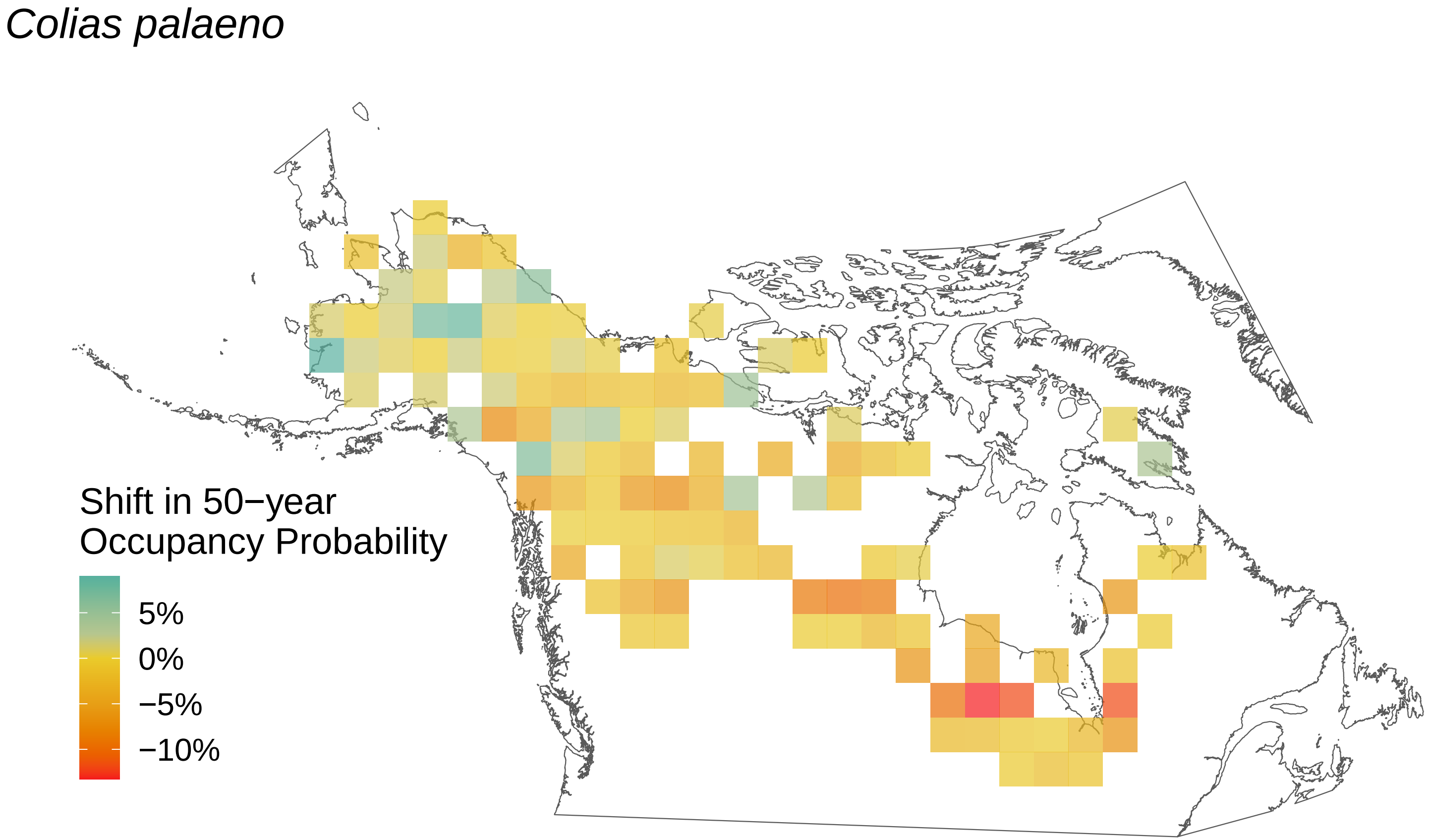
Colias palaeno
Shift in 50−year
Occupancy Probability
5%
0%
−5%
−10%

*Colias interior*

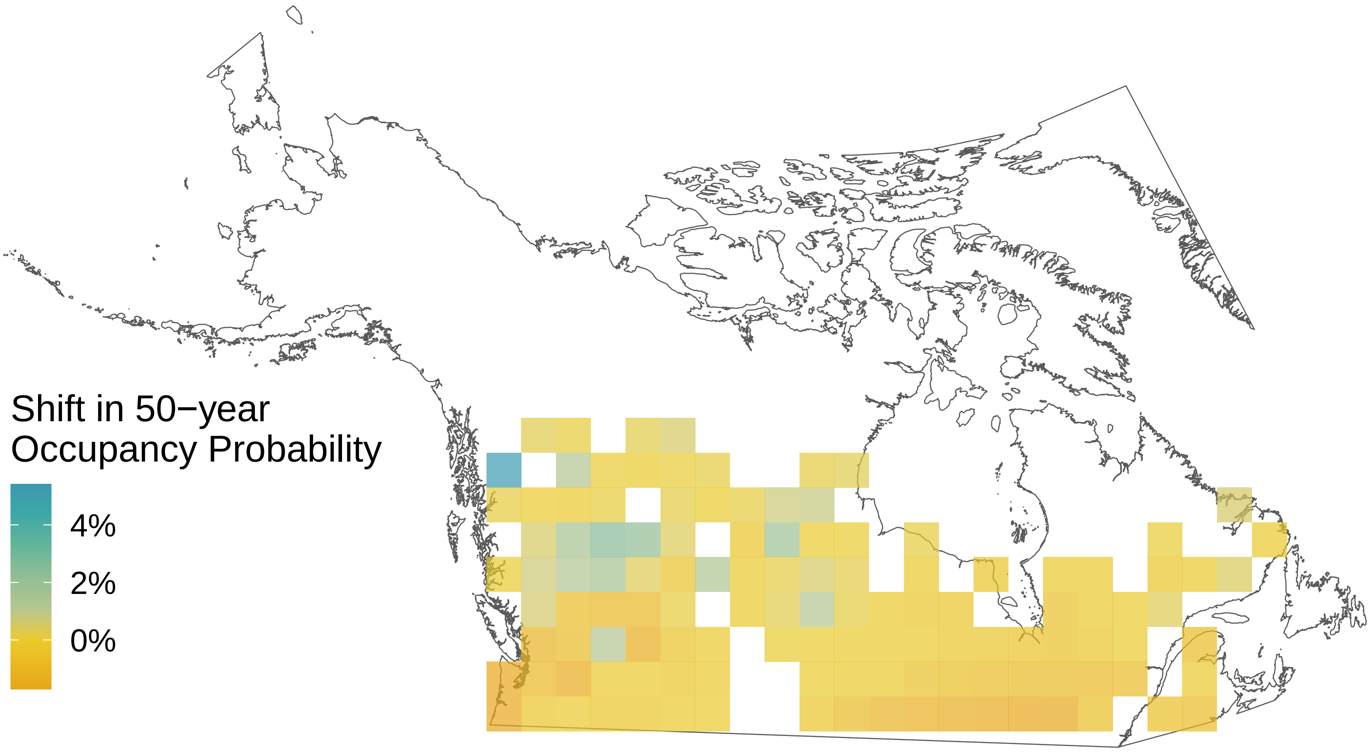

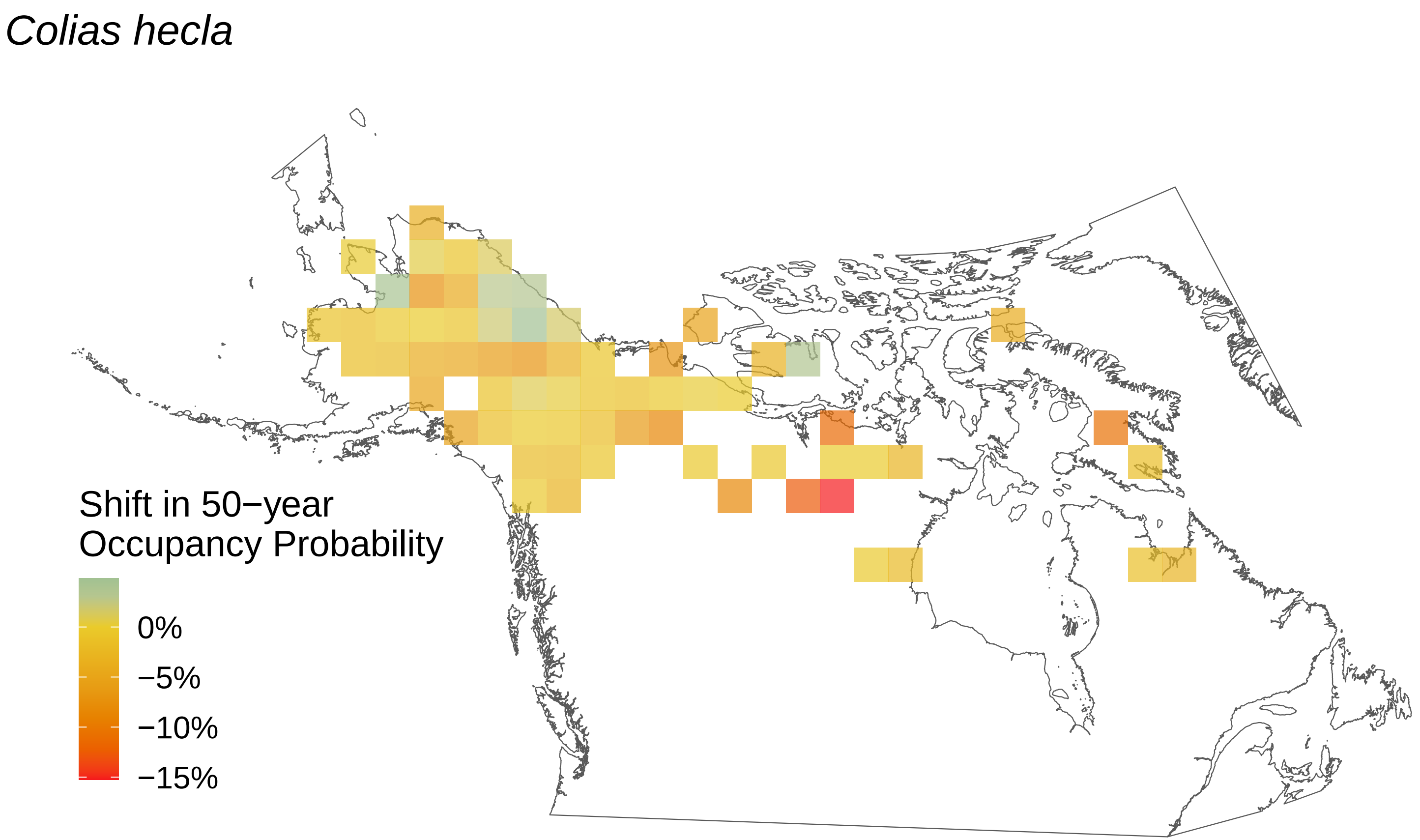

*Colias hecla*
Shift in 50−year
Occupancy Probability
0%
−5%
−10%
−15%

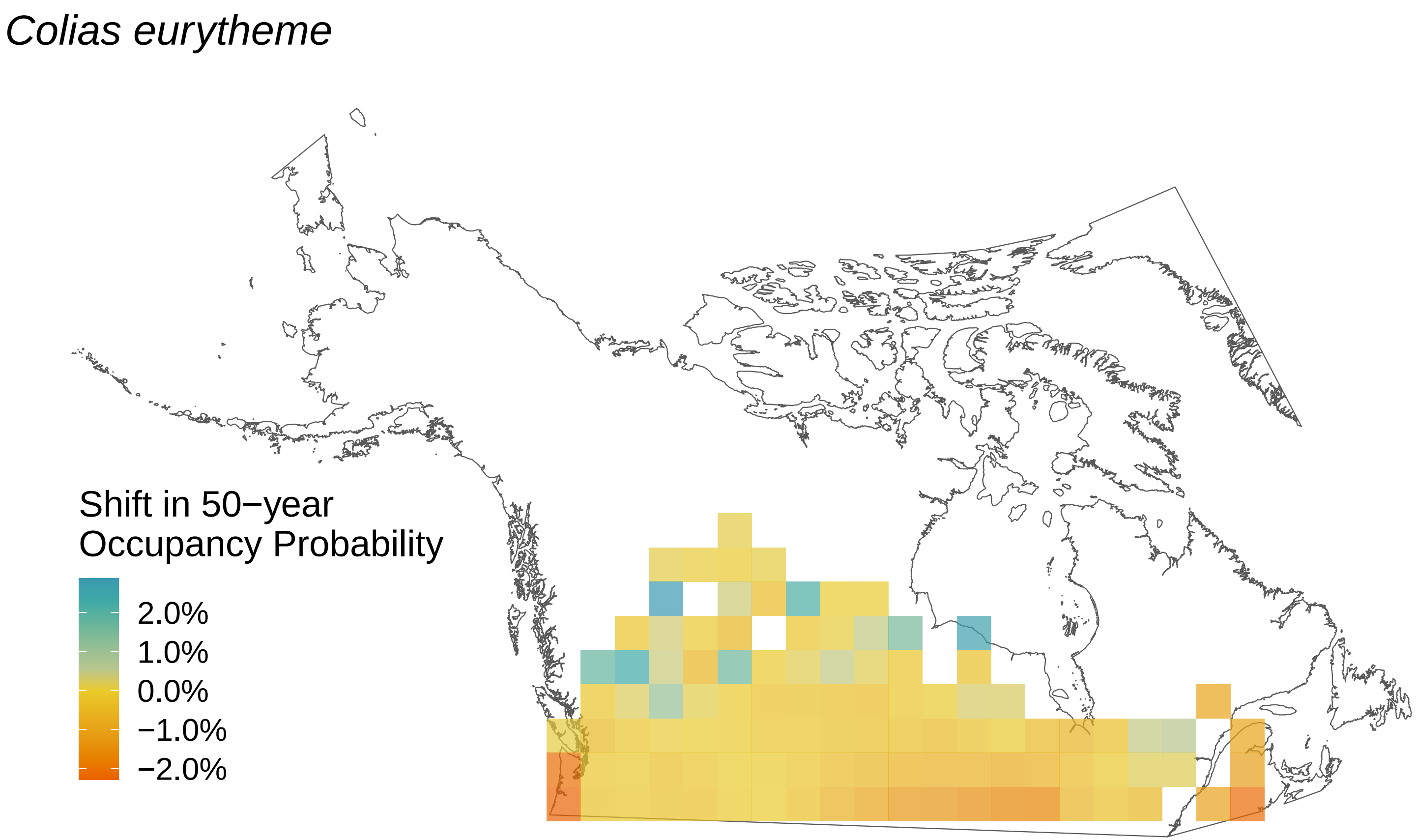
Colias eurytheme
Shift in 50−year
Occupancy Probability
2.0%
1.0%
0.0%
−1.0%
−2.0%

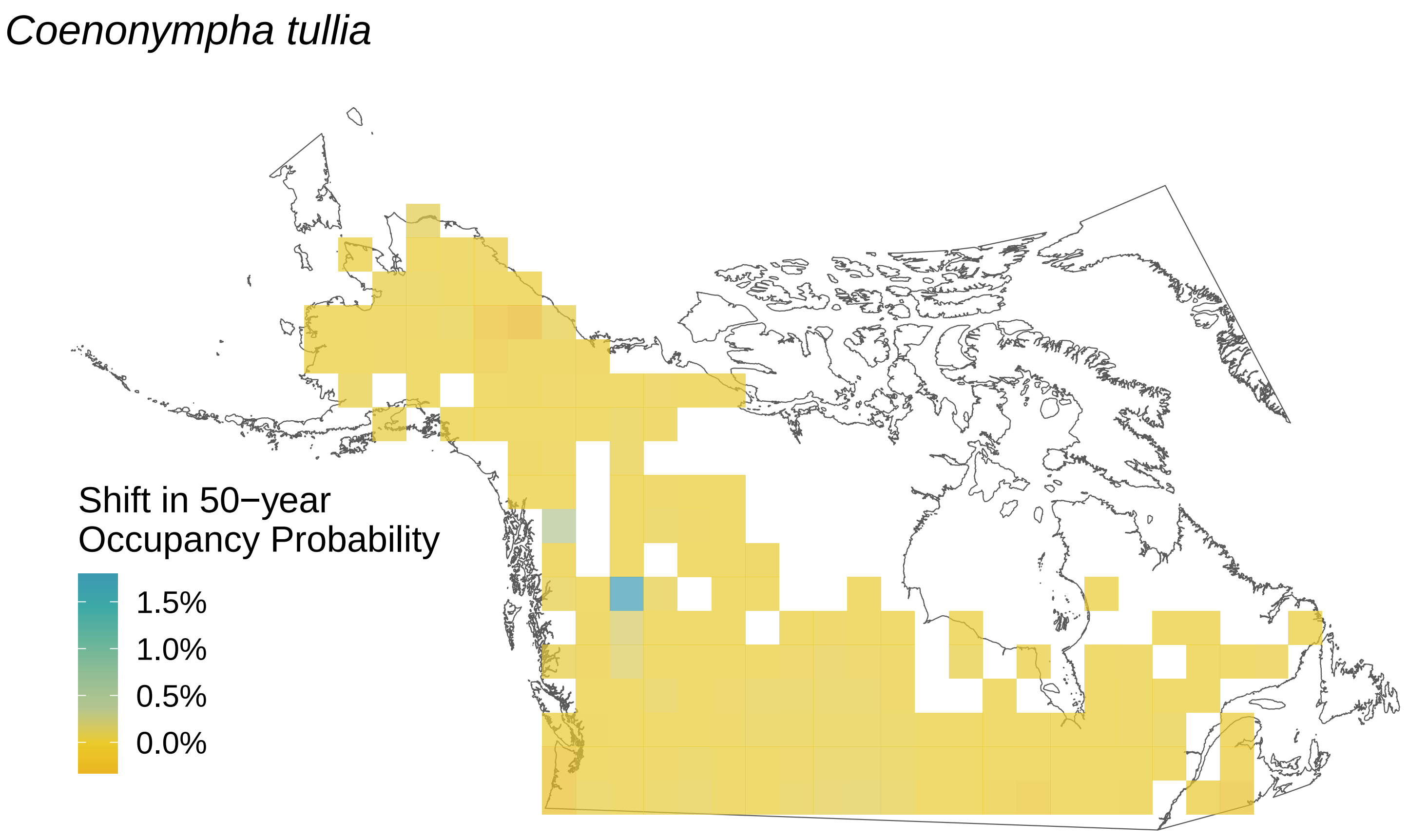

Coenonympha tullia
Shift in 50−year
Occupancy Probability
1.5%
1.0%
0.5%
0.0%

*Chlosyne harrisii*

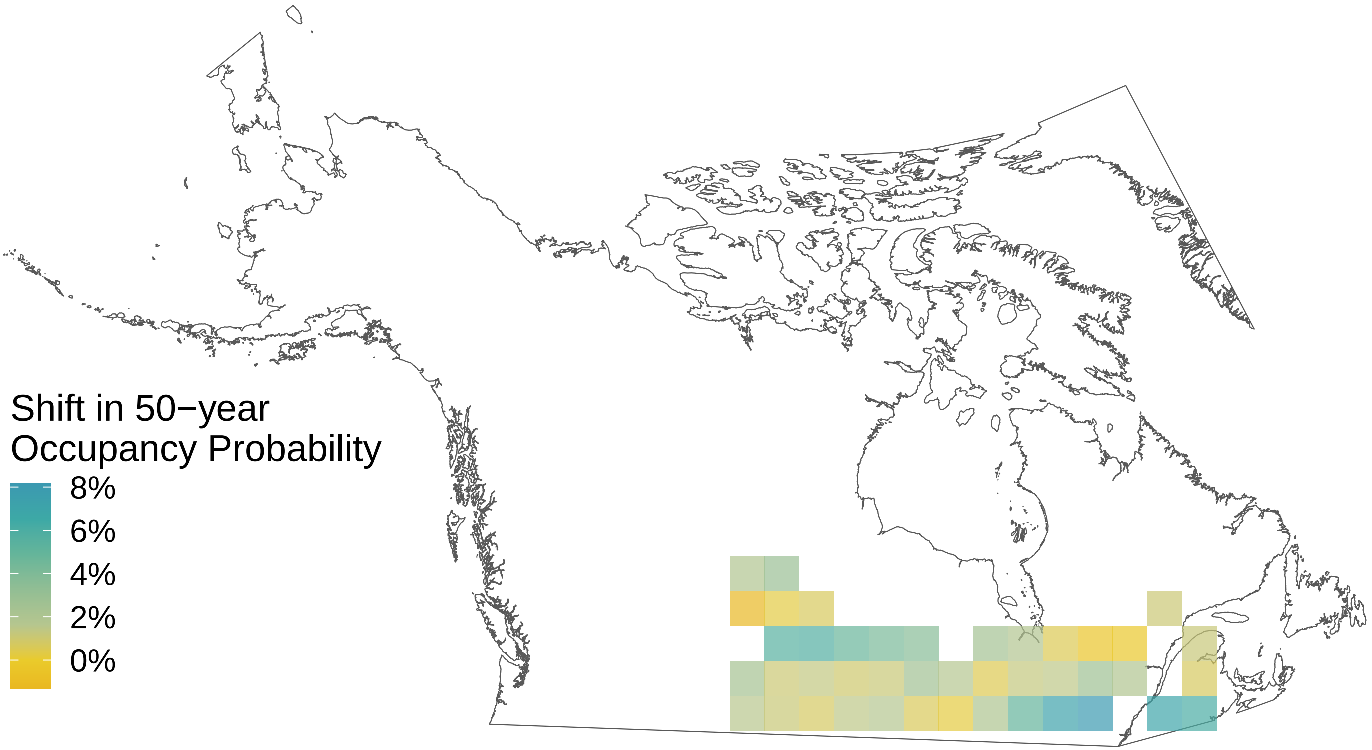

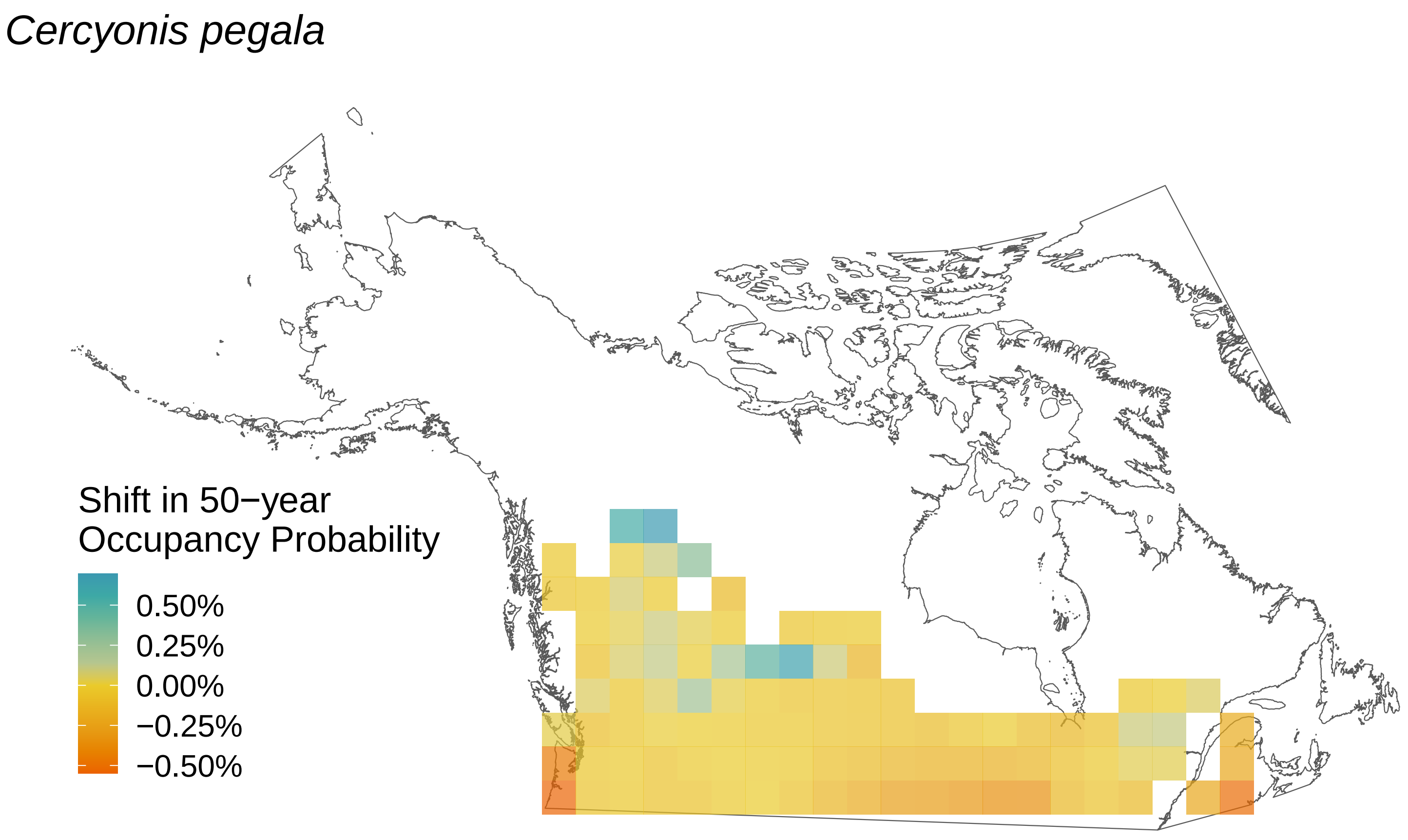
Cercyonis pegala
Shift in 50−year
Occupancy Probability
0.50%
0.25%
0.00%
−0.25%
−0.50%

*Carterocephalus palaemon*

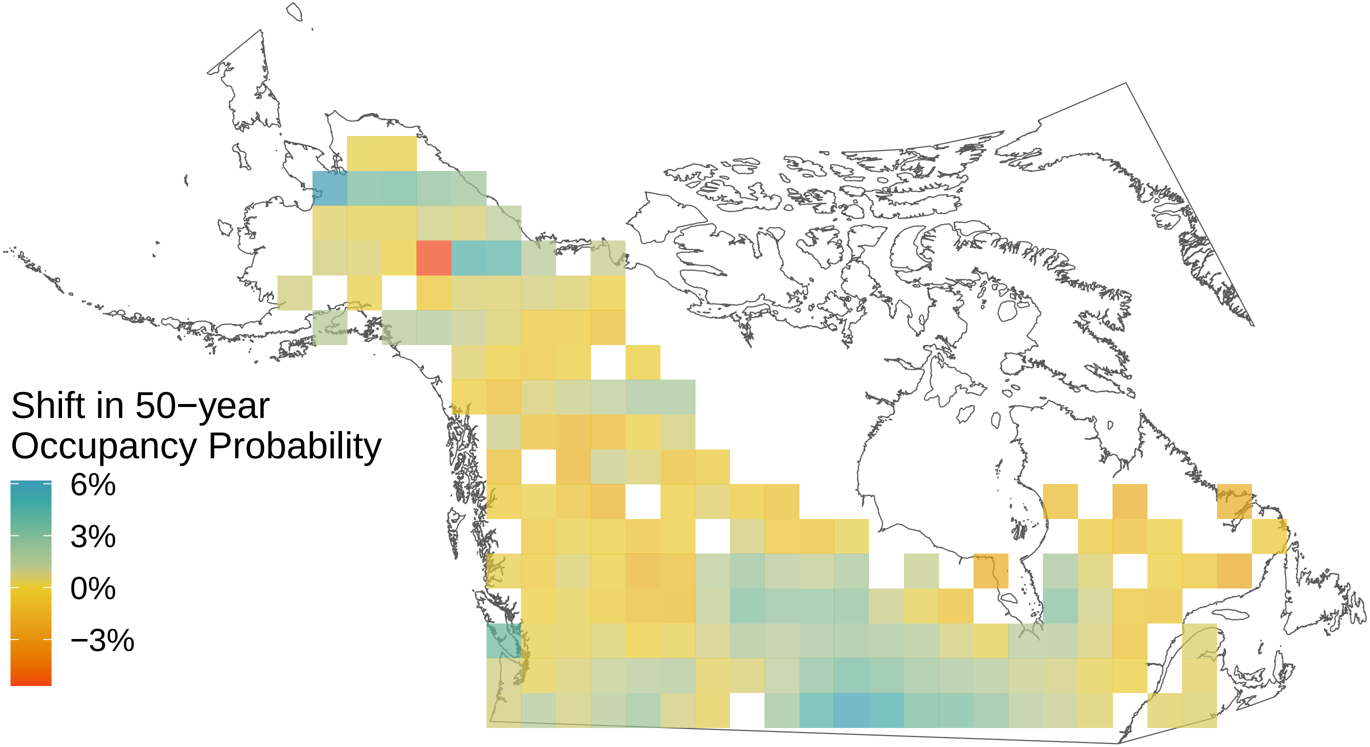

*Callophrys polios*

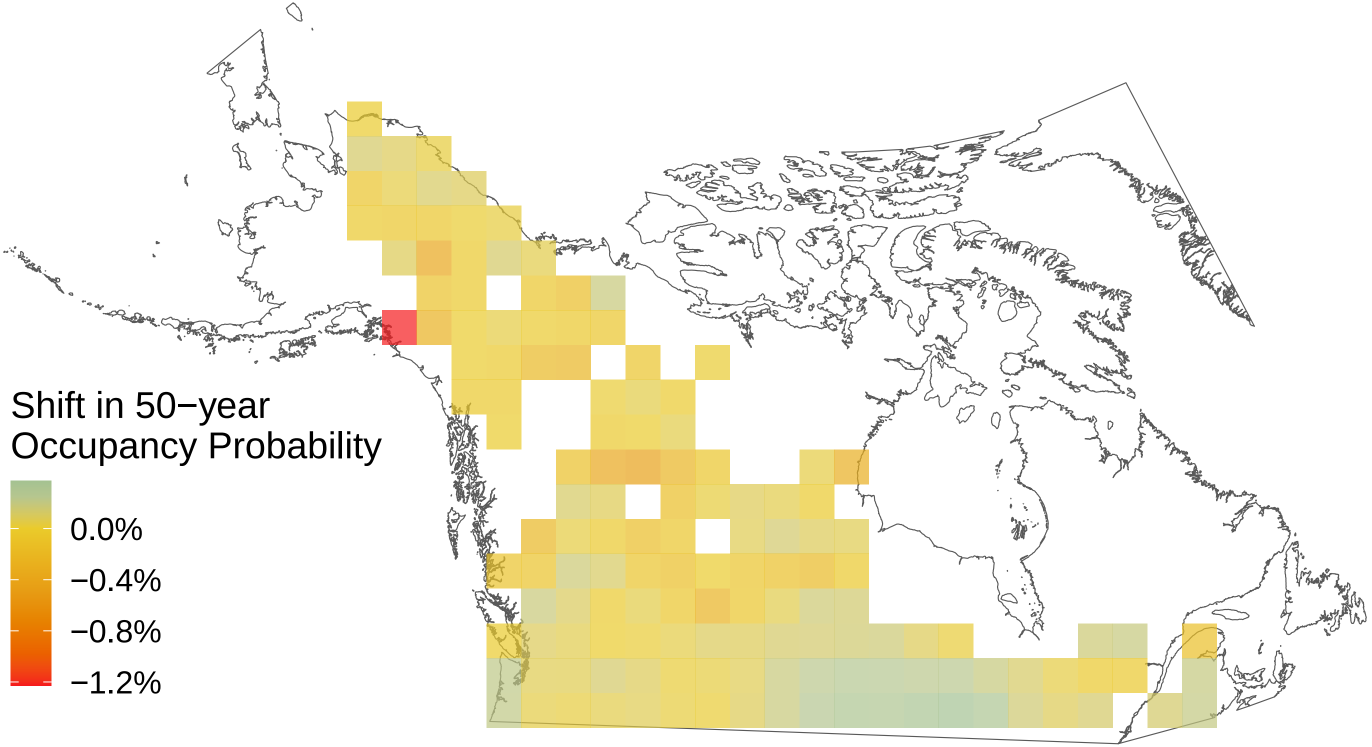

*Callophrys niphon*

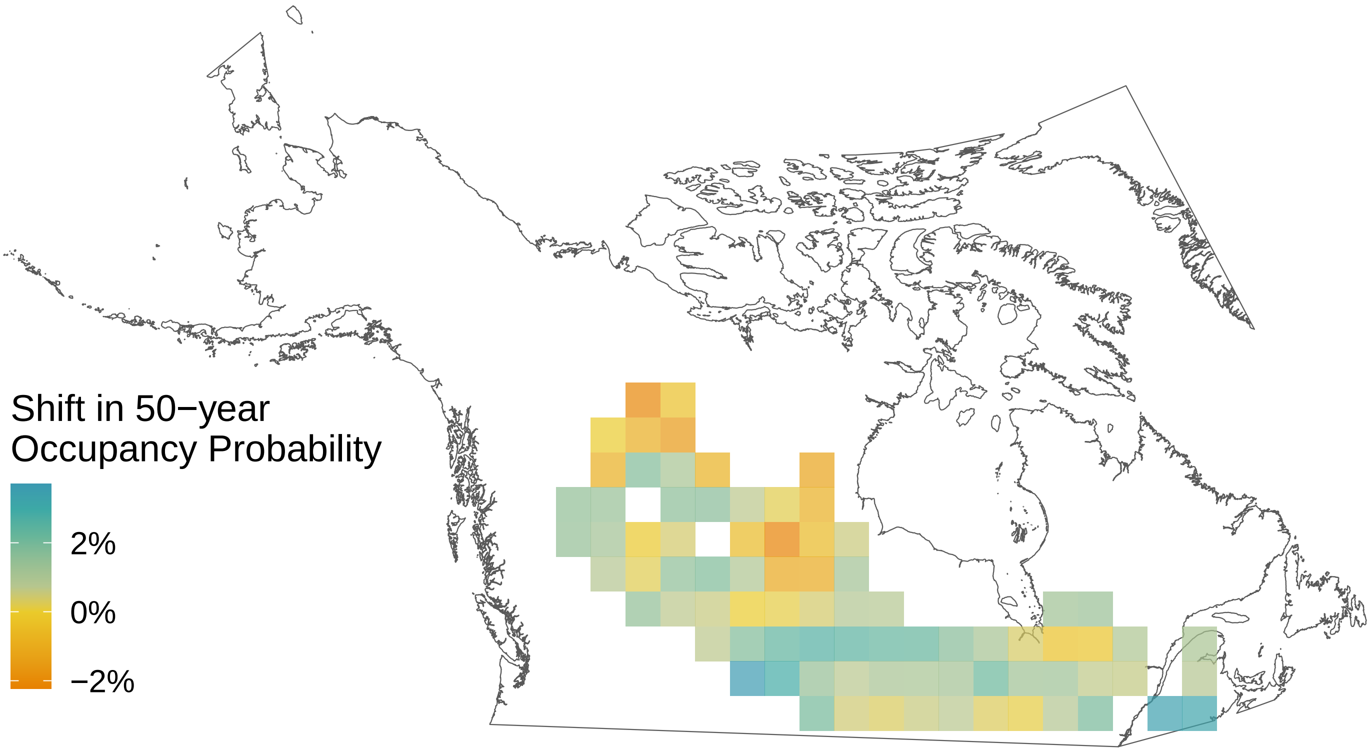

*Callophrys eryphon*

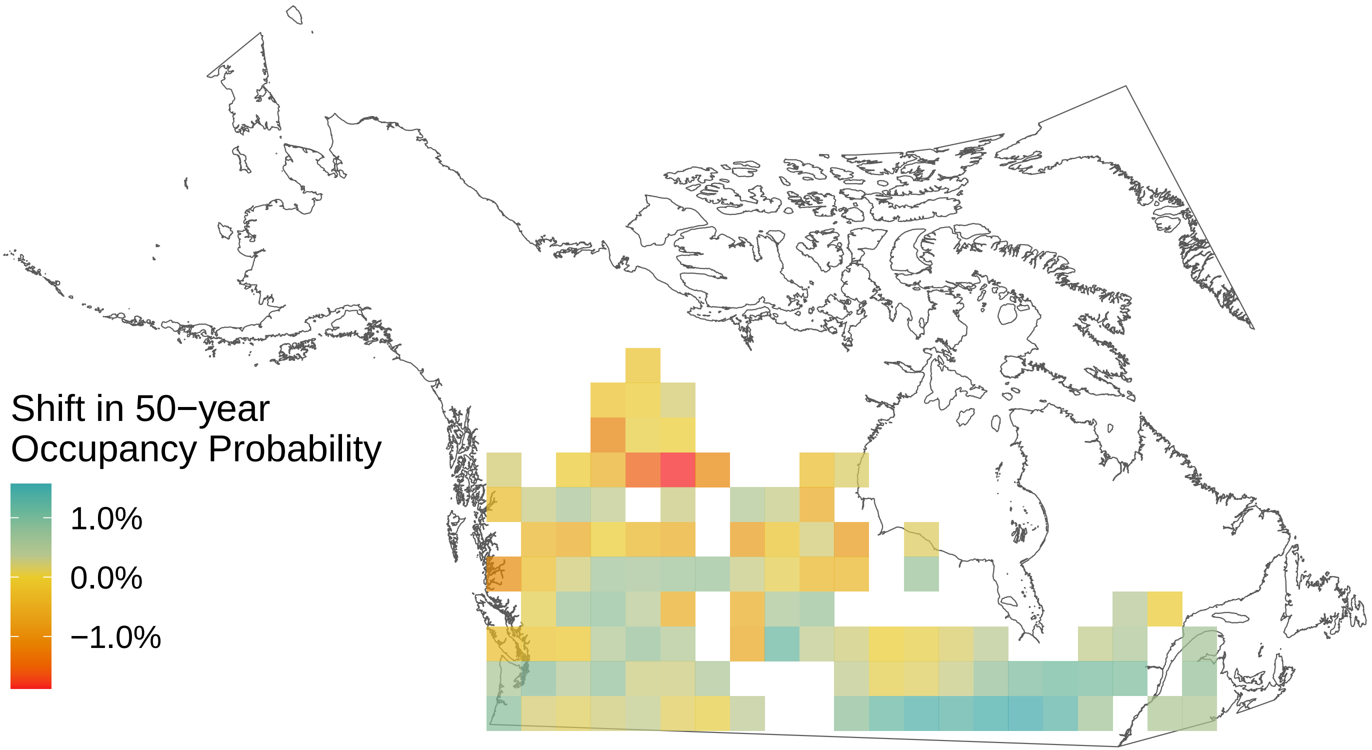

### *Callophrys augustinus*

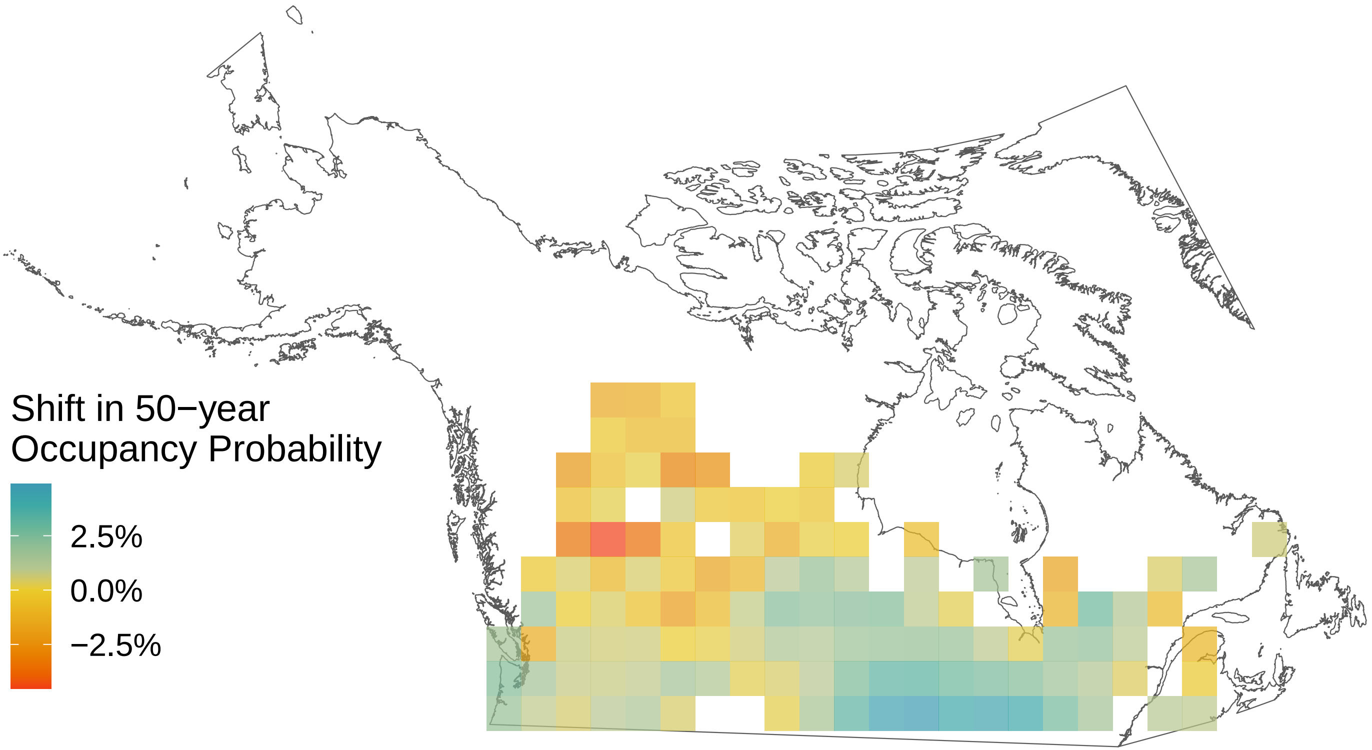

*Boloria selene*

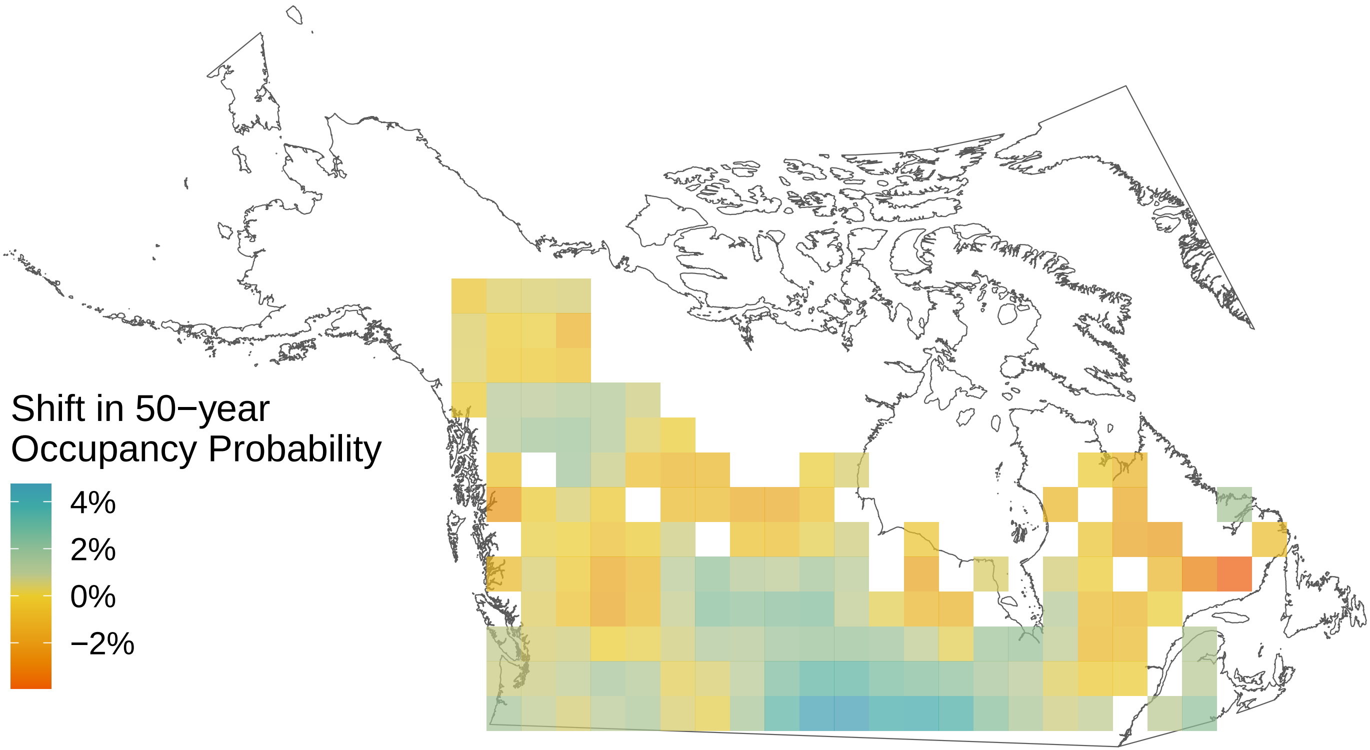

*Boloria polaris*

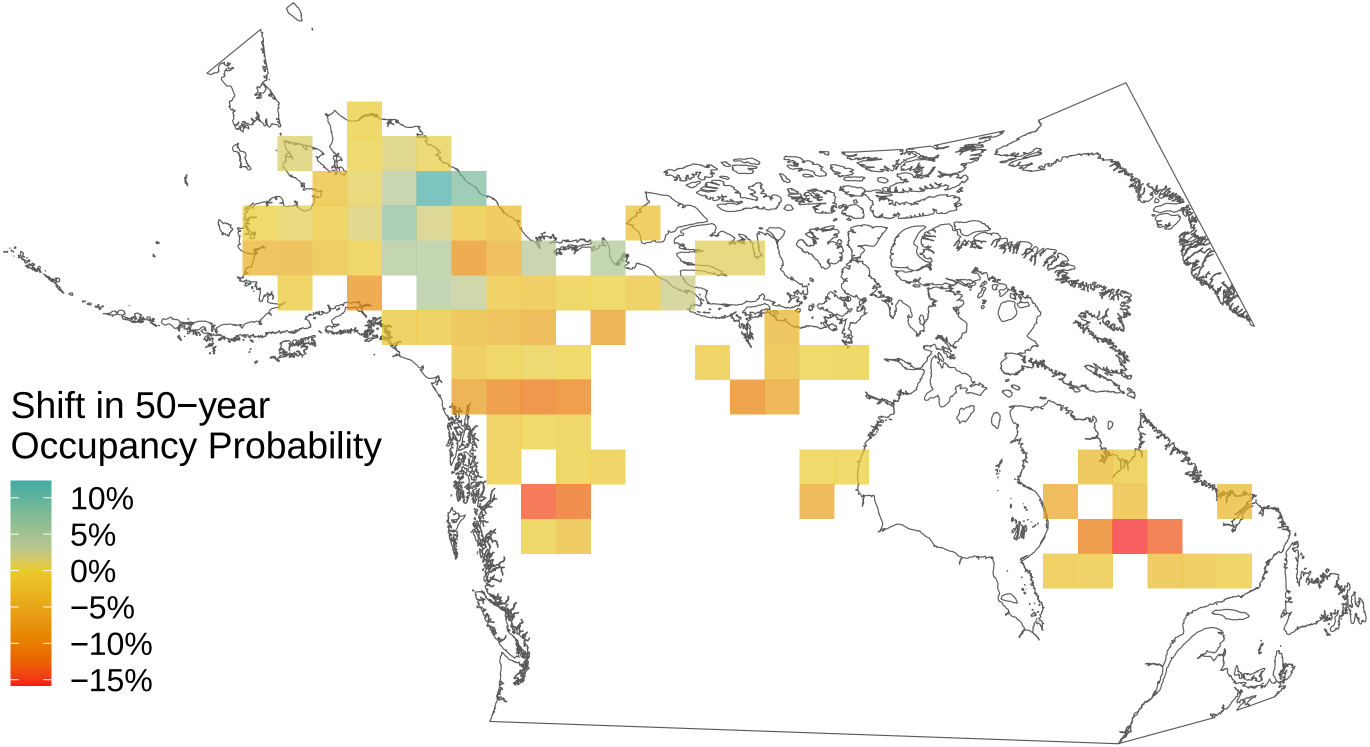

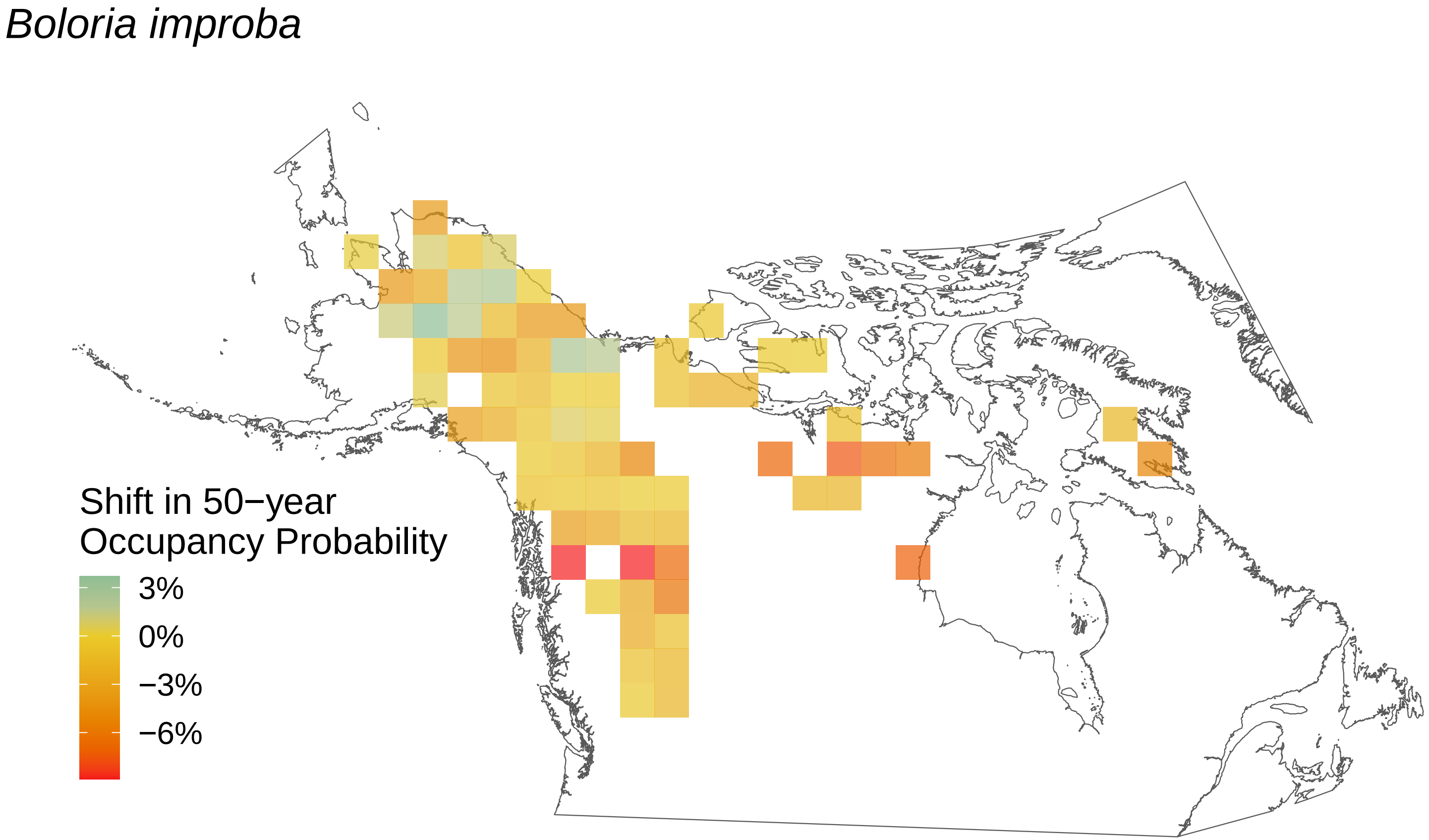

*Boloria improba*
Shift in 50−year
Occupancy Probability
3%
0%
−3%
−6%

*Boloria frigga*

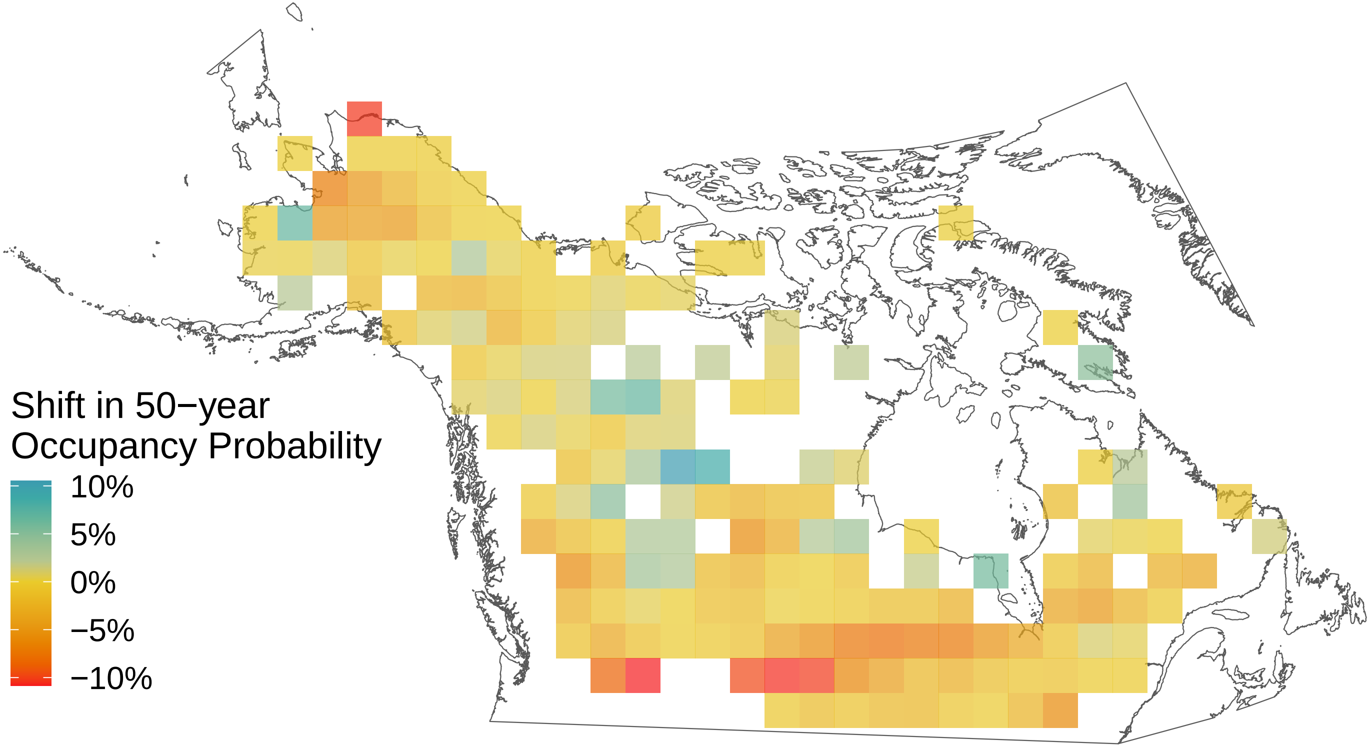

### *Boloria freija*

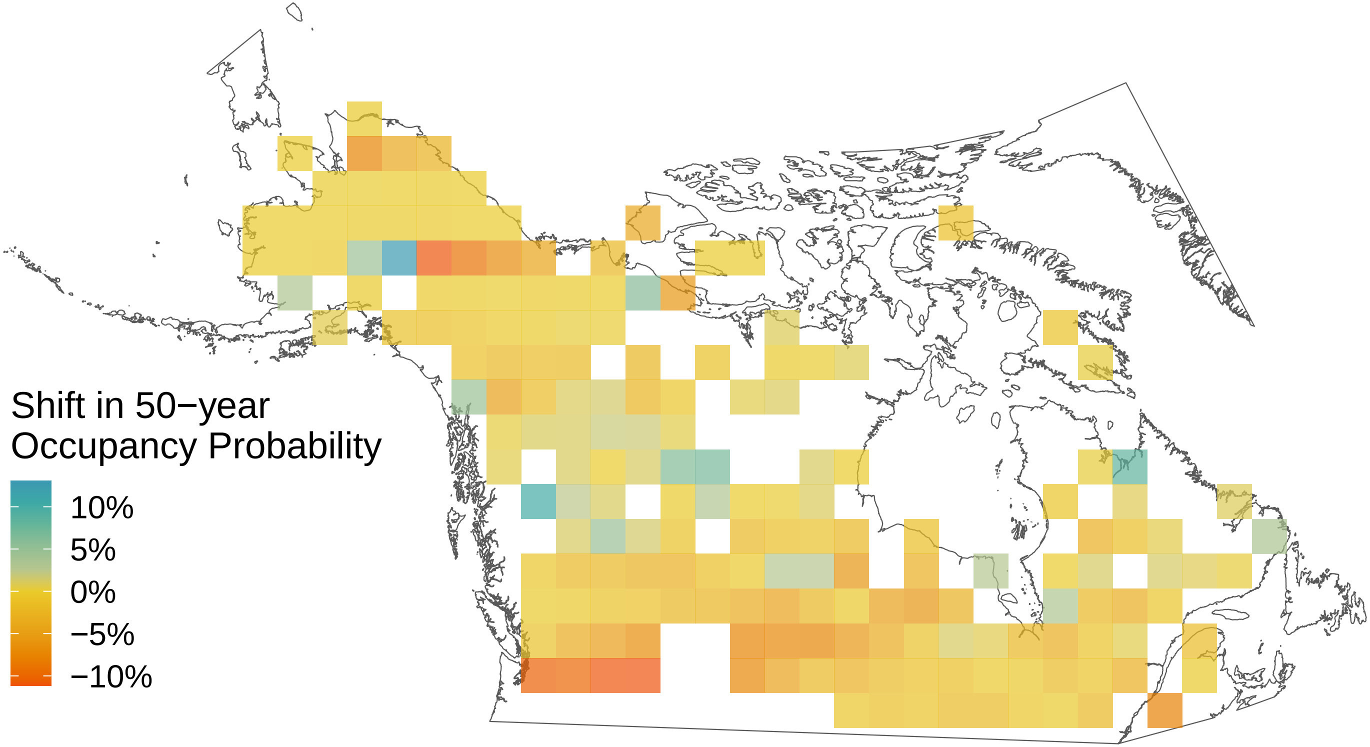

*Boloria eunomia*

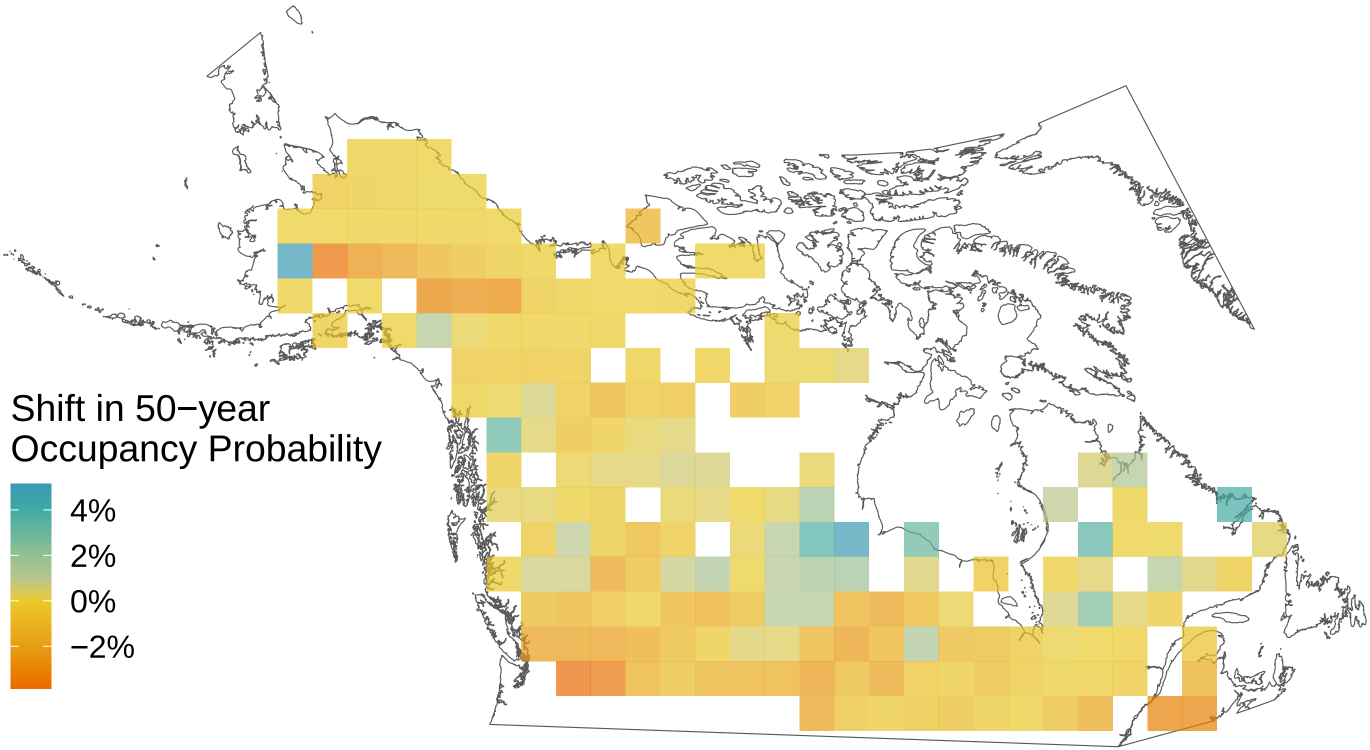

*Boloria chariclea*

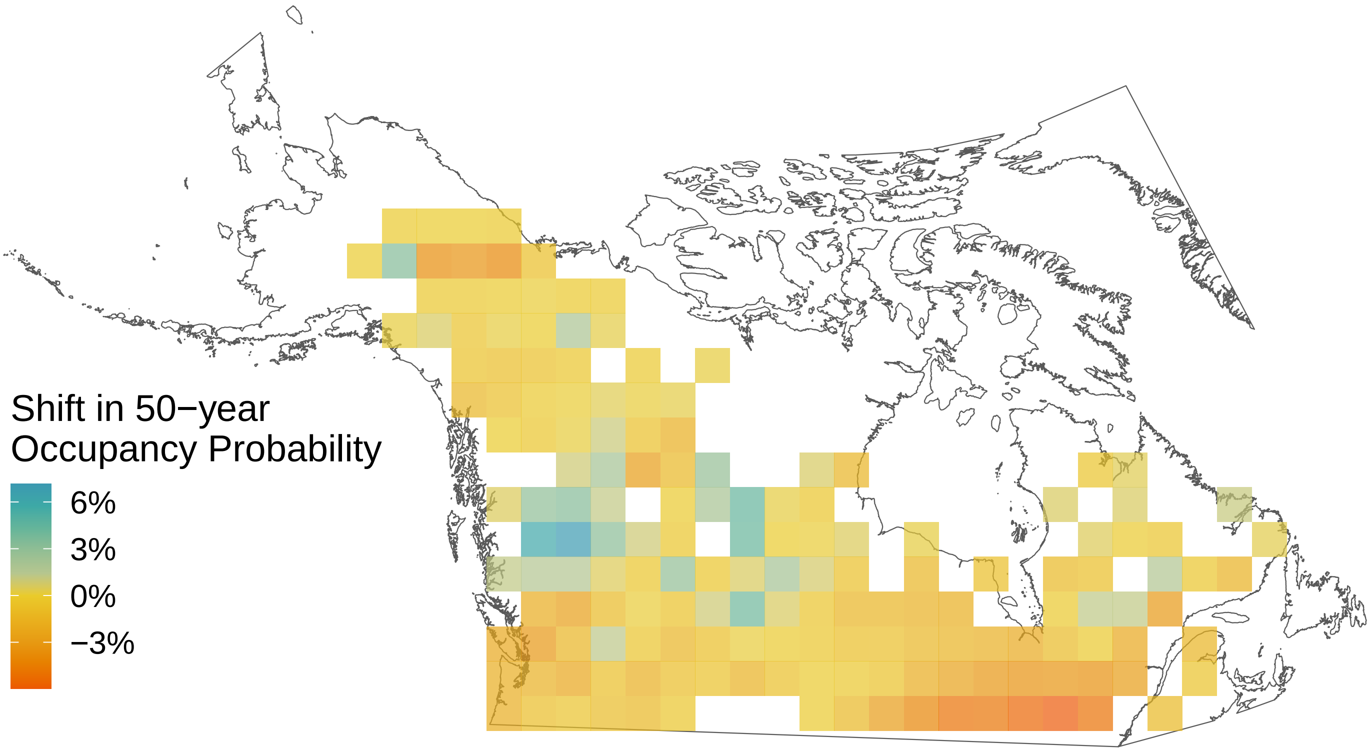

*Boloria bellona*

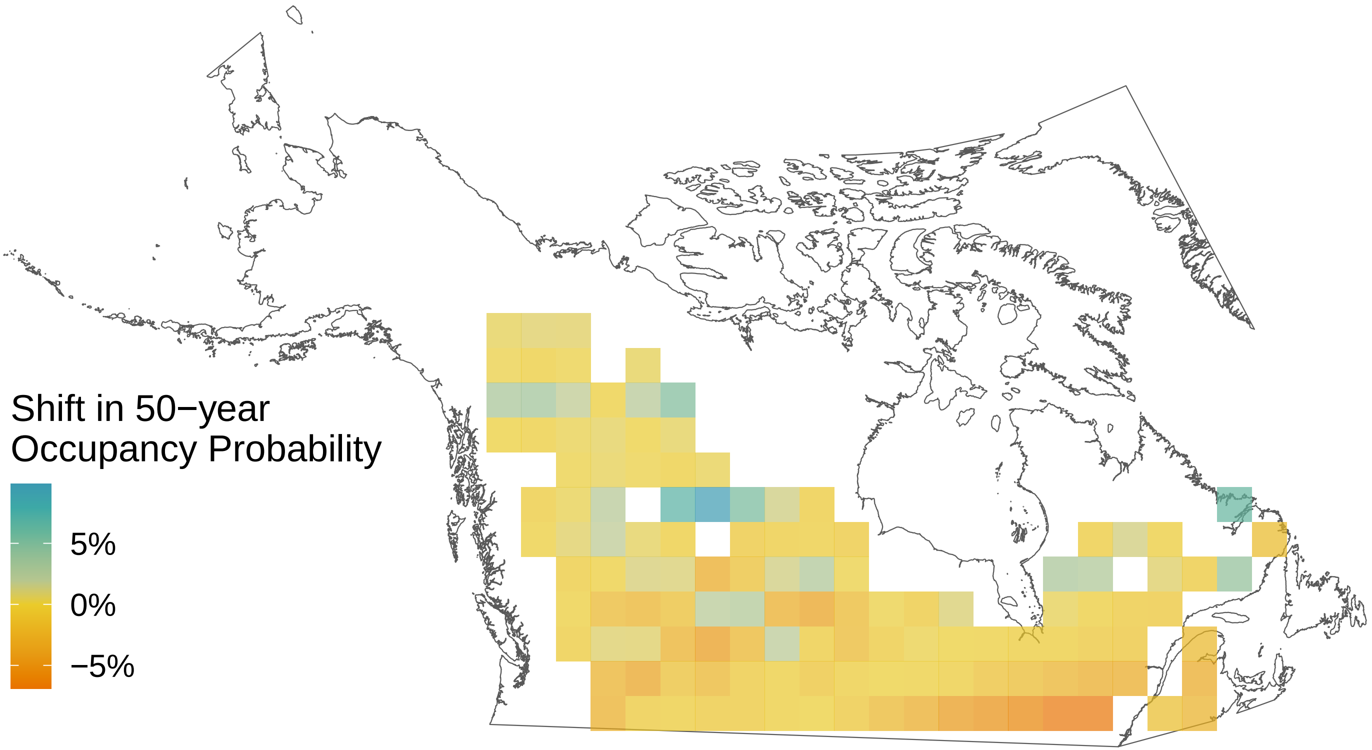

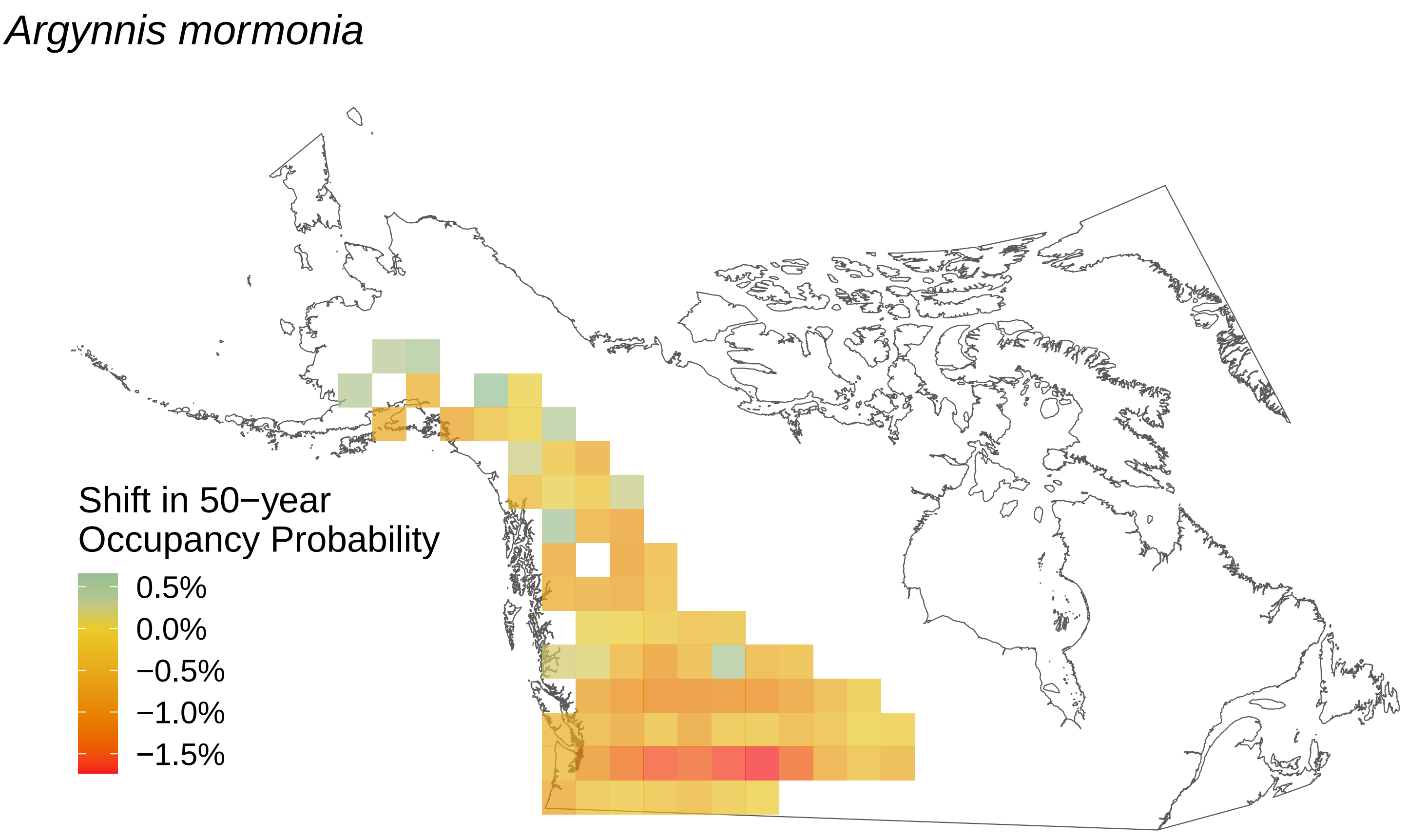
Argynnis mormonia
Shift in 50−year
Occupancy Probability
0.5%
0.0%
−0.5%
−1.0%
−1.5%

*Argynnis hydaspe*

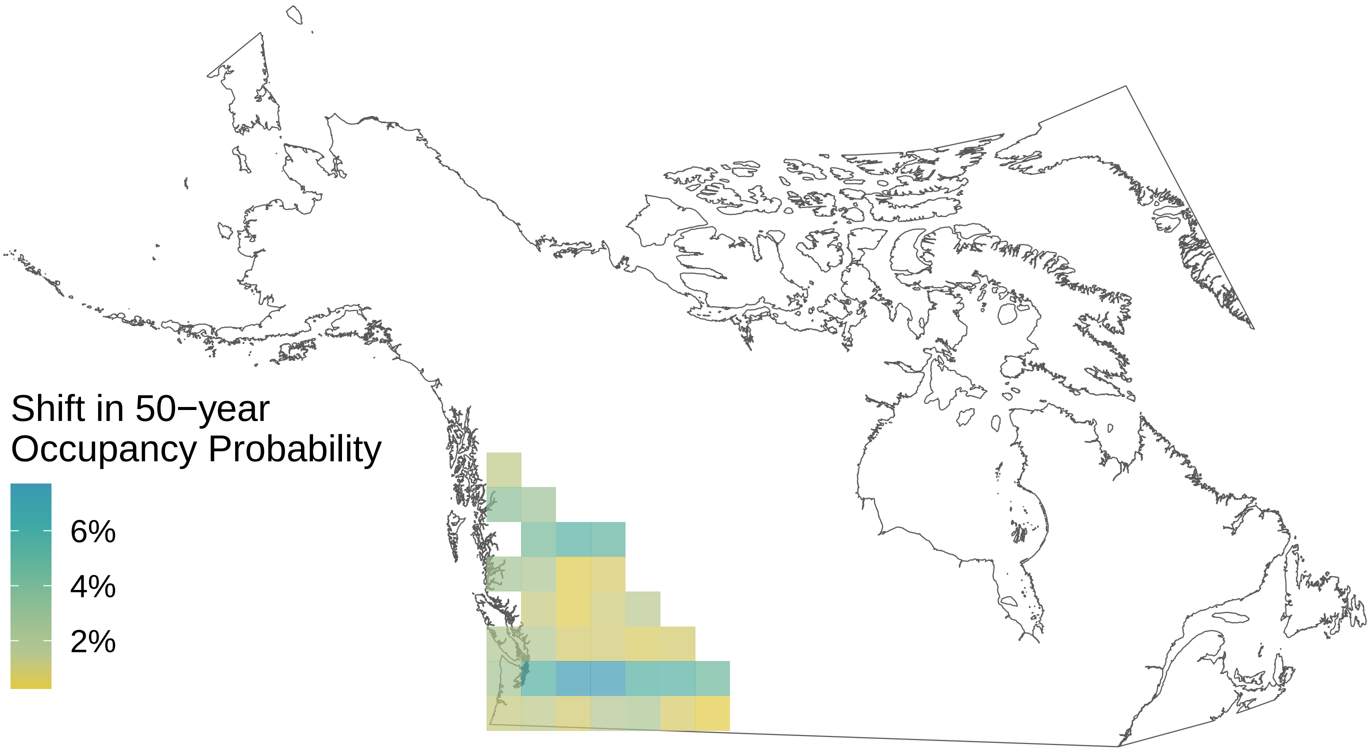

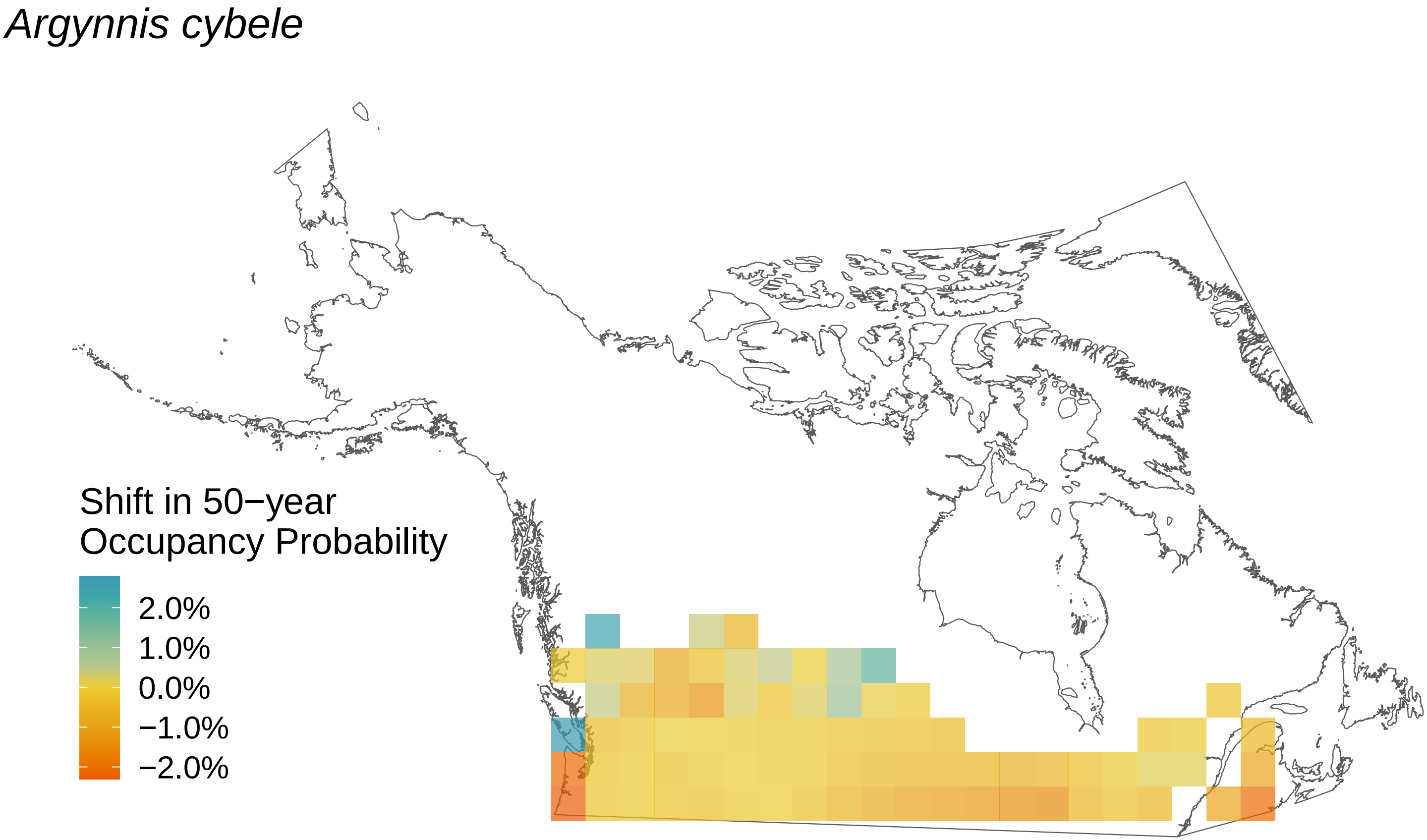
Argynnis cybele
Shift in 50−year
Occupancy Probability
2.0%
1.0%
0.0%
−1.0%
−2.0%

*Argynnis atlantis*

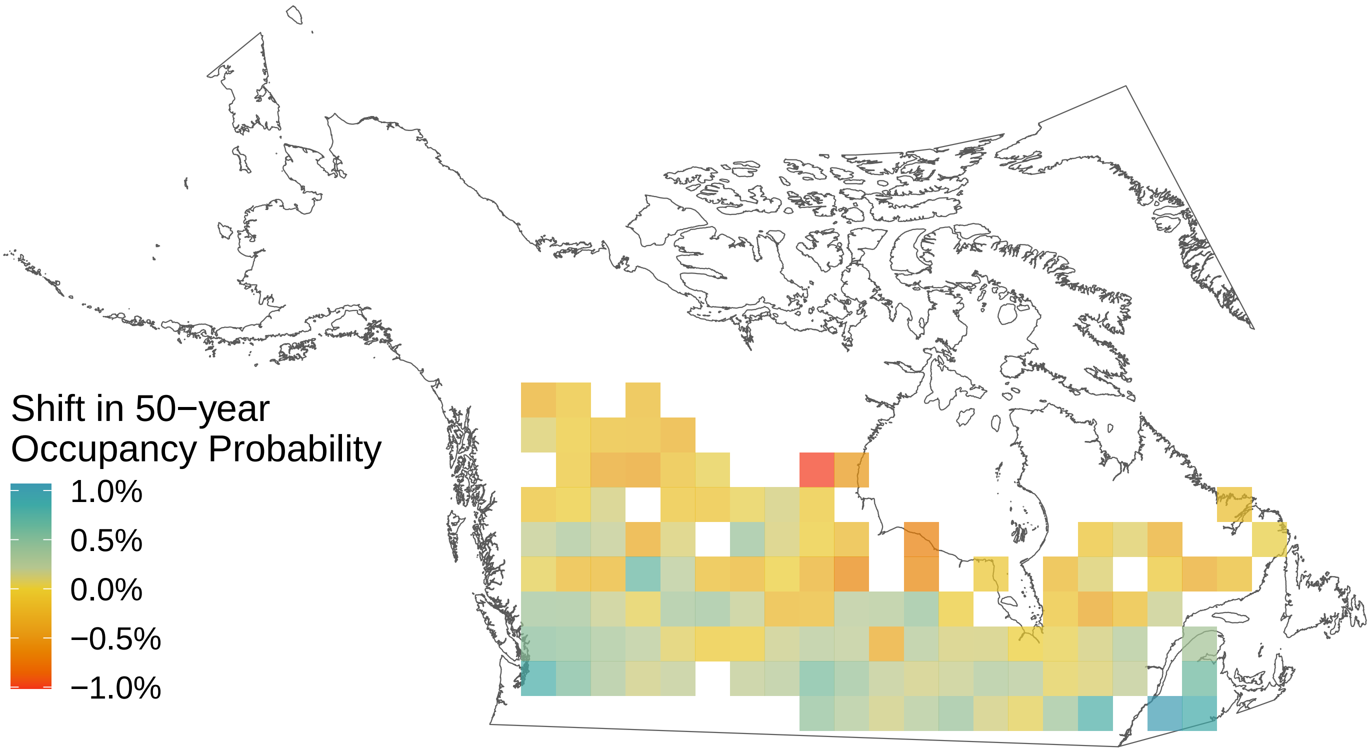

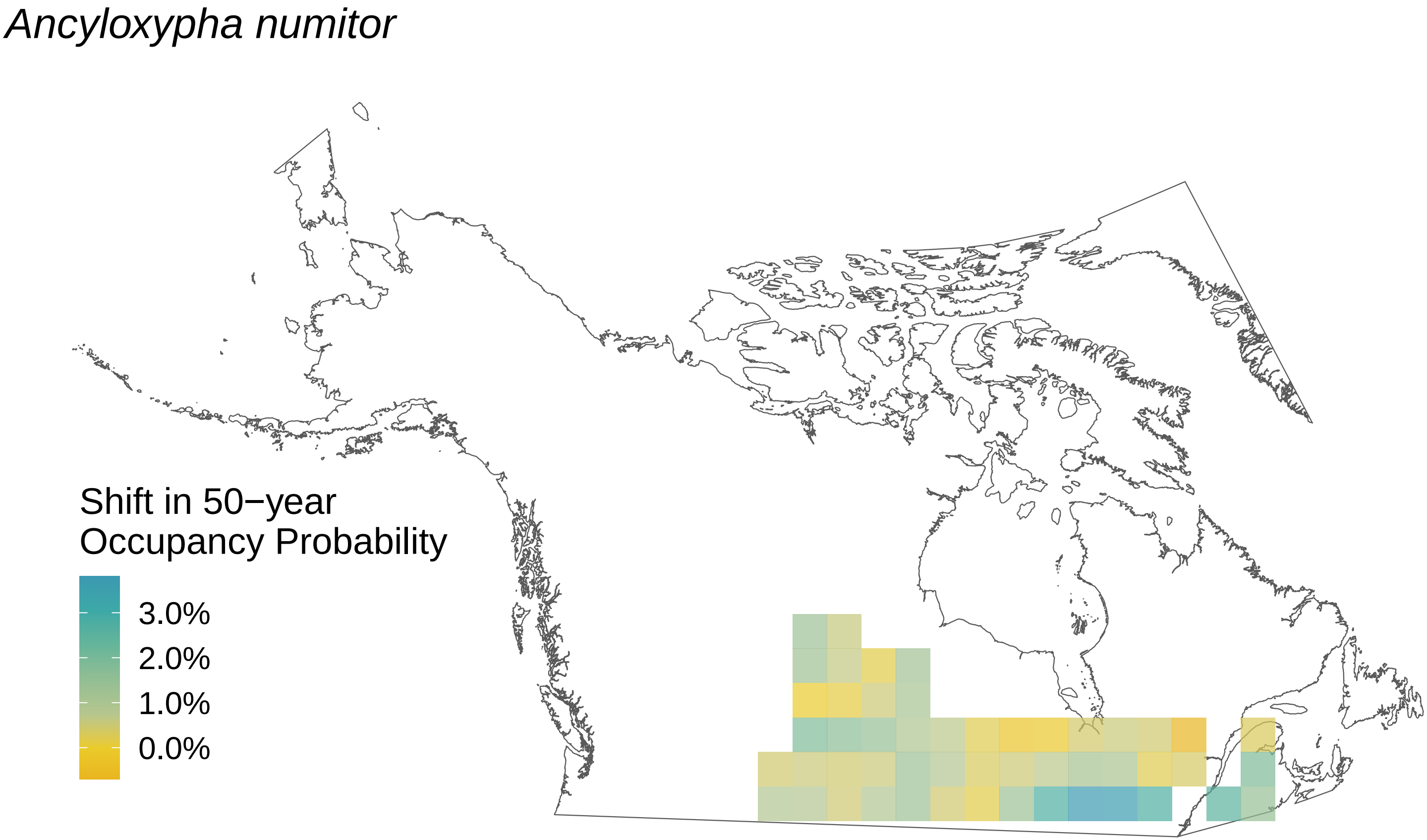

Ancyloxypha numitor
Shift in 50−year
Occupancy Probability
3.0%
2.0%
1.0%
0.0%

*Amblyscirtes vialis*

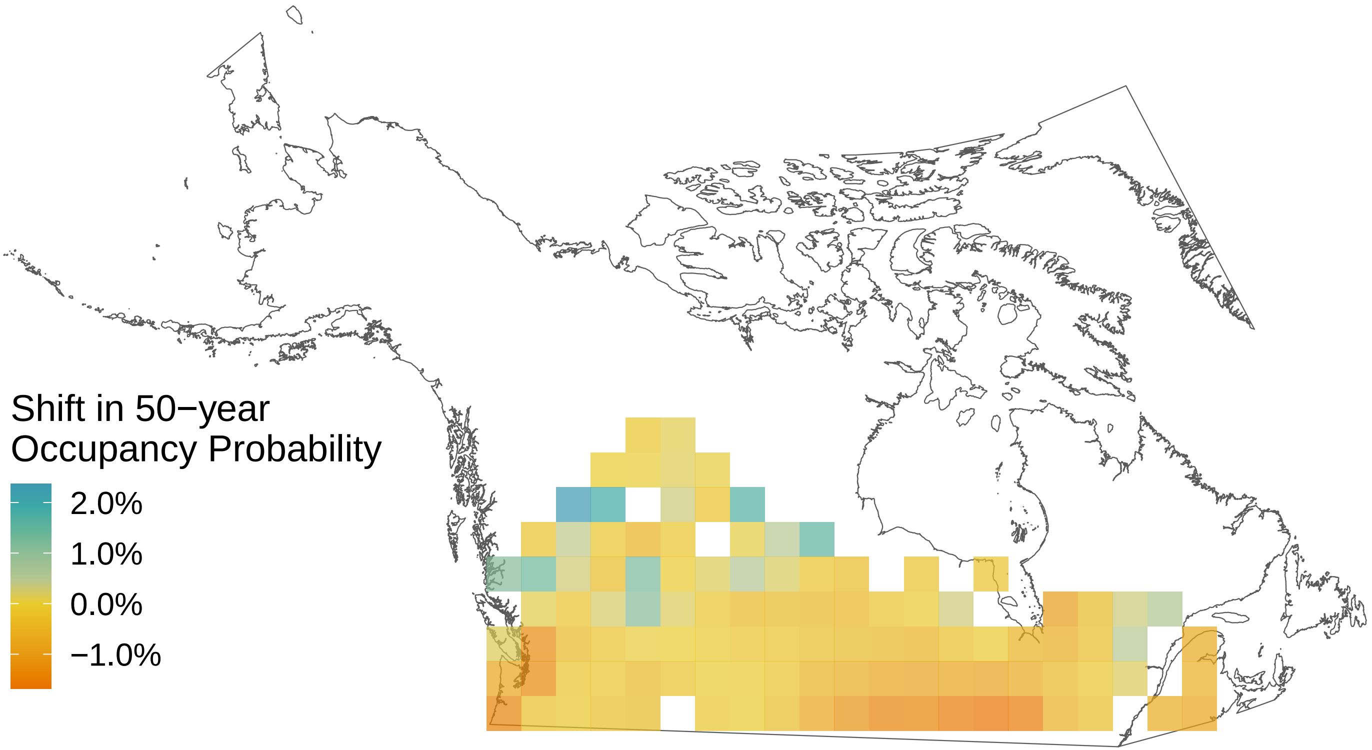

*Agriades glandon*

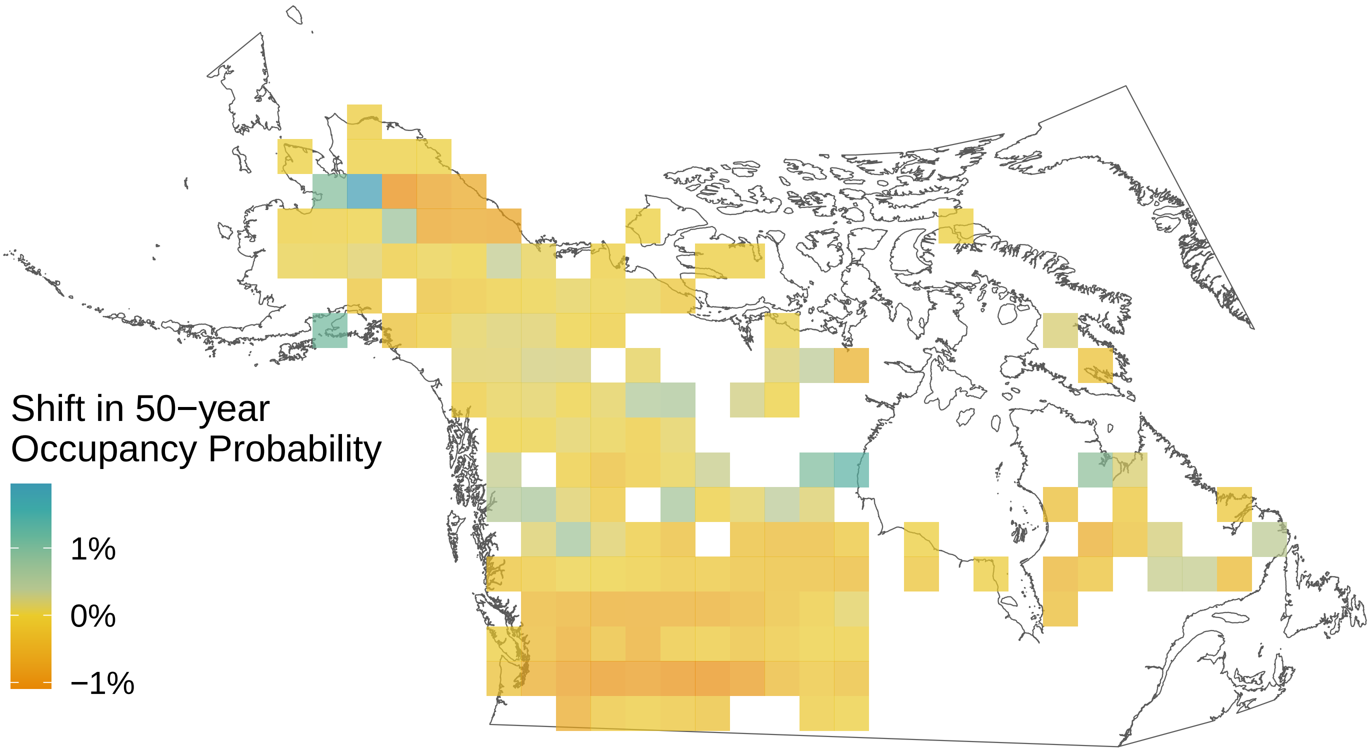

### *Aglais milberti*

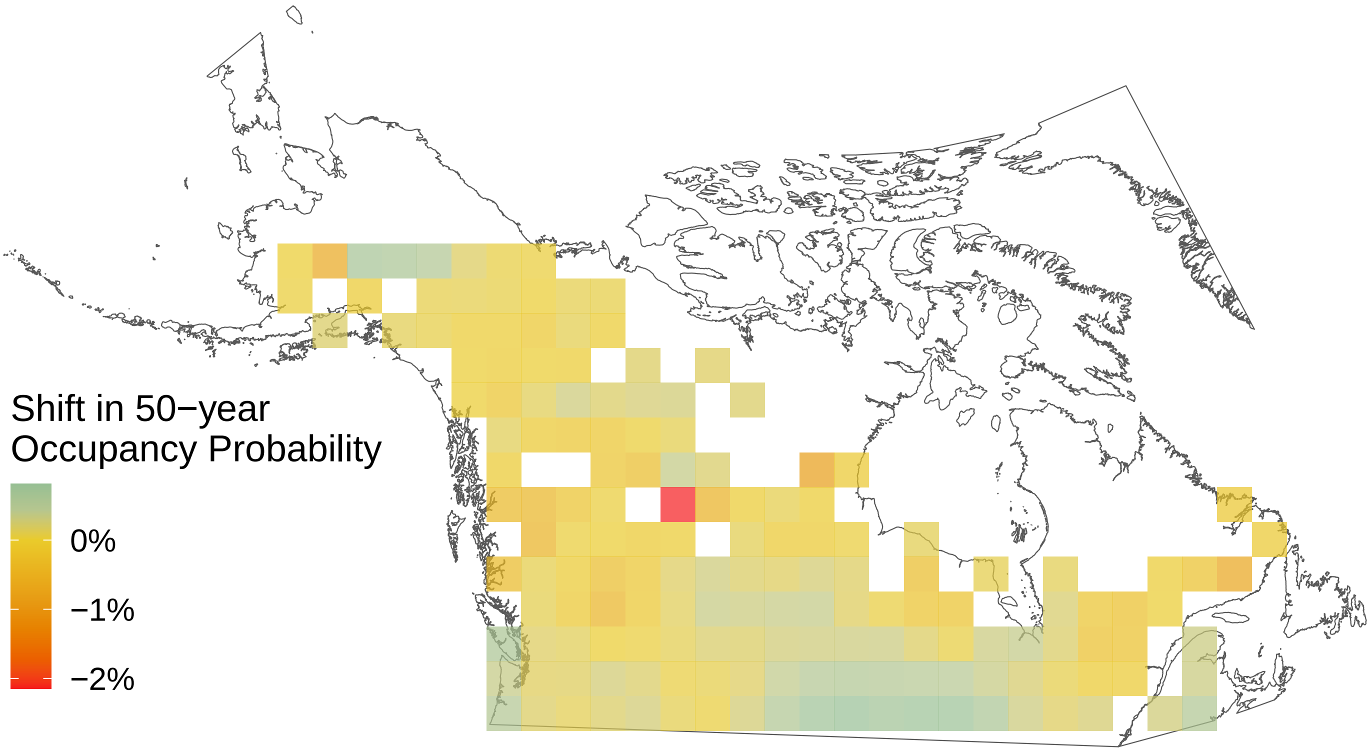
